# Fighting isn’t sexy in lekking Greater Sage-grouse (*Centrocercus urophasianus*)

**DOI:** 10.1101/2022.08.26.505294

**Authors:** Samuel S. Snow, Gail L. Patricelli, Carter T. Butts, Alan H. Krakauer, Anna C. Perry, Ryane Logsdon, Richard O. Prum

## Abstract

In lekking systems, females can observe both male courtship displays and fights. It has been theorized that male-male agonism may function as a display, giving females more information about mate quality. However, males in many species, such as Greater Sage-grouse, often fight when females are absent, and can even attack during copulation attempts in seeming conflict with females’ choices.

Traditional correlational approaches are inadequate to distinguish the underlying mechanisms of social interaction and can result in misleading associations between fighting and mating events. Using observations from a wild population, we posit a novel Relational Event Model that incorporates temporal dependencies of events among a network of individuals. We investigate how fighting among male sage-grouse predicts events such as future fights, copulation solicitations, and interrupted copulations.

Our analysis reveals that fighting’s primary function is not to impress females. Indeed, males are less likely to start and more likely to leave fights with females present, plausibly to avoid entanglement in conflict that reduces availability to mate. Moreover, being drawn into these latter viscous cycles of combat and retribution constitutes a significant risk associated with initiating attacks on other males. However, fighting serves other roles, e.g., to deter copulation interruptions and rebuff competitors. Our findings suggest that social systems that regulate conflict and promote females’ choice based on display are likely fundamental to the stable evolution of leks.

## Introduction

In lek-breeding systems, where males gather to display and females visit to evaluate and choose mates, females can often observe both male courtship displays and fights (Bradbury & Gibson, 1983; Avery, 1984; Wiley, 1991; Höglund & Alatalo, 1995). It has been theorized that male-male agonism may function as a display, giving females more information about mate quality (Kruijt & Hogan, 1967; Cox & Le Boeuf, 1977; Alatalo *et al*., 1991; Hovi *et al*., 1995; Candolin, 1999; Wong & Candolin, 2005; Hämäläinen *et al*., 2012). However, males in many lekking species, such as Greater Sage-grouse (*Centrocercus urophasianus*), often fight when females are absent, and can even attack during mating attempts in seeming conflict with females’ choices (Wiley, 1973; Foster, 1983; Trail, 1985; Trail & Koutnik, 1986; Gibson & Bradbury, 1987; Alatalo *et al*., 1996; Sæther *et al*., 1999; Nooker & Sandercock, 2008).

Whether male aggression is aligned or in conflict with female mate choice is crucial for our understanding of underlying evolutionary pressures and processes. Classic theory suggests that fighting demonstrates “good genes” fitness benefits for females (Cox & Le Boeuf, 1977; Andersson, 1994) and feeds into narratives of lek evolution wherein the lek system offers more opportunities for females to view contests (Beehler & Foster, 1988; Kokko, 1997). However, recent theoretical work has supported the notion that male aggression that hinders the efficacy of female mate choice can give rise to selection for female behaviors or morphologies that resist those effects, even when there is an apparent association between attractiveness and aggression in males (Snow *et al*., 2019). Selection for resistance behaviors can lead to drastically different expected outcomes in the trajectory of the evolution of mating interactions, as compared to models that assume male aggression and female mate choice are aligned (e.g., Rosenthal & Servedio, 1999; Lessells, 2006; Snow *et al*., 2019).

For example, if male aggression in the form of courtship disruption is in conflict with female mate choice, then fighting among males would be an obstacle to the evolution of a stable, functioning lek as an arena for female choice based on courtship display, rather than a force promoting the lek evolution. Given the extreme skew in mating success in lek breeding systems (Bradbury & Gibson, 1983), males would be incentivized to disrupt one-another’s copulations/courtship to the detriment of the group as a whole (Chapman, 1935; Foster, 1983; Prum, 2017) – a situation similar to the “Tragedy of the Commons” (Hardin, 1968). To overcome this “Tragedy of the Lek”, resist sexually coercive behavior, and allow for the persistence of lekking systems that we observe in nature, females would be expected to select for reduced aggression, and male behaviors that regulate aggression, instead of selecting for more aggressive males as previous theory has suggested, (Prum, 2017; Snow *et al*., 2019). Moreover, if fighting is in conflict with, as opposed to aligned with, female choice in some contexts, we may also expect to concurrently observe fighting behaviors serving a regulatory role in other contexts and/or the self-regulation of aggression.

One difficulty in teasing apart the various ideas about the role of aggression in mating systems (and in understanding social behaviors in general) is analytical. Mating success and histories of aggressive behavior are the result of episodic social interactions that unfold over time, and yet in evolutionary biology we usually aggregate these social events into counts (or, increasingly, metrics for behaviors embedded in a social network; e.g., Faust & Skvoretz, 2002; Croft *et al*., 2008) and then we correlate those behaviors or metrics of interest with outcomes like mating success (Alatalo *et al*., 1991; Höglund *et al*., 1997; Ryder *et al*., 2008, 2009; Hämäläinen *et al*., 2012). There has been a growing recognition that our ability to understand the processes underlying complex social dynamics within groups of individuals hinges on our understanding of how network structures and relationships change over time (e.g., Snijders, 2001; Koskinen & Snijders, 2007; Blonder *et al*., 2012; Hobson *et al*., 2013; Almquist & Butts, 2014; Pinter-Wollman *et al*., 2014; Krivitsky & Handcock, 2014; Dakin & Ryder, 2018; Dakin & Ryder, 2020; Ryder *et al*., 2020). However, previous static or dynamic approaches all rely on researchers to choose a temporal window over which to aggregate social events into a static network or networks. This temporal binning leads to a loss of information on the timing and sequence of events (Butts, 2008; Butts & Marcum, 2017), making it difficult or impossible to ask questions about mechanistic relationships among particular events. For example, eliminating information on sequence and timing of fighting and mating events would cause us to implicitly (and inappropriately) assume that many females are making mating decisions using information about fights that have not yet occurred.

To address this problem, we have for the first time implemented a Relational Event Model (REM) to study natural reproductive behavior (Butts, 2008; Marcum & Butts, 2015; Patison *et al*., 2015; Tranmer *et al*., 2015; Butts & Marcum, 2017). A REM is a statistical framework that explicitly models the temporal dependencies within sequences of social events among a network of individuals unfolding in continuous time. This allows us to directly ask questions about potential mechanistic relationships among events such as: (1) “Do particular histories or outcomes of fights in front of females precipitate copulation events?”, (2) “How does the presence of females influence male aggressive behavior?”, and (3) “Do past fights influence the likelihood of future fights, including the risk of being interrupted while copulating?” This last question allows us to gain insight into whether there is a regulatory role for fighting in this system. Because the framework leads to a generative model, it also allows us to probe the consequences of interaction mechanisms for the social system as a whole. In particular, we are able to examine the longer-term consequences of fighting for both copulation opportunities and future combat via indirect mechanisms, such as the initiation of persistent cycles of conflict that, once set in motion, become hard to escape. As we show, involvement in such cycles is a risk of initiating conflict.

In the REM, for every point in time, we model a *hazard* or conditional rate of occurrence for each type of behavioral event, such as “male A attacks male B” or a solicitation for copulation, that is possible at that moment (Fig. 1; Fig. 2). The hazard of any event is a function of the past observed event history, a set of effect statistics encoding various mechanisms that may affect the rates of occurrence of future events, and a set of parameters that govern the magnitude and direction of the effects (Eq. 1). We then estimate the values of the parameters associated with each effect statistic that are most probable given the observed interaction data under diffuse priors, employing the Bayesian Information Criterion (BIC) to choose the best model from among models built with different sets of possible effects.

**Figure 1.**
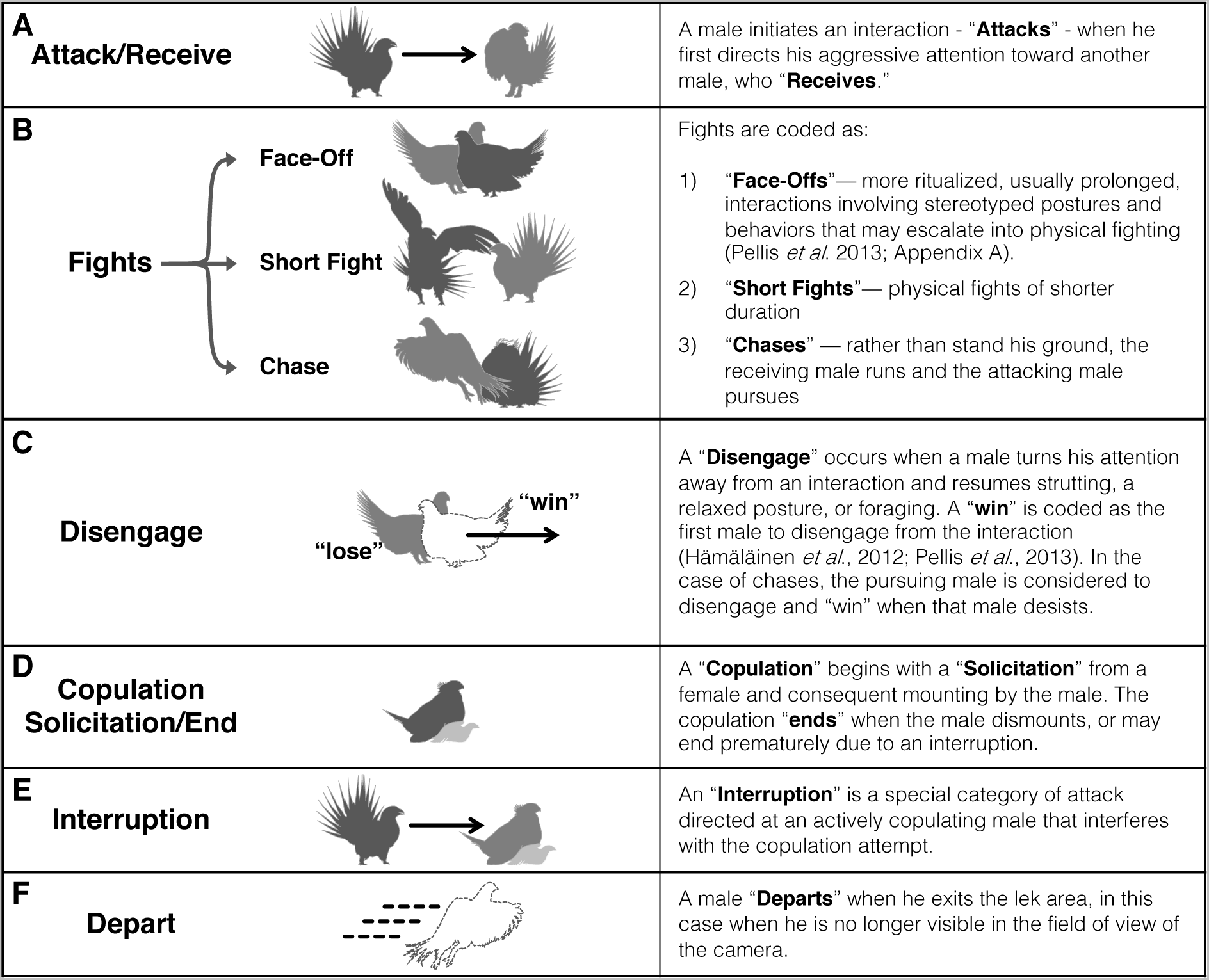
Ethogram of male behaviors. [Photo credit: A.H.K.]

**Figure 2.**
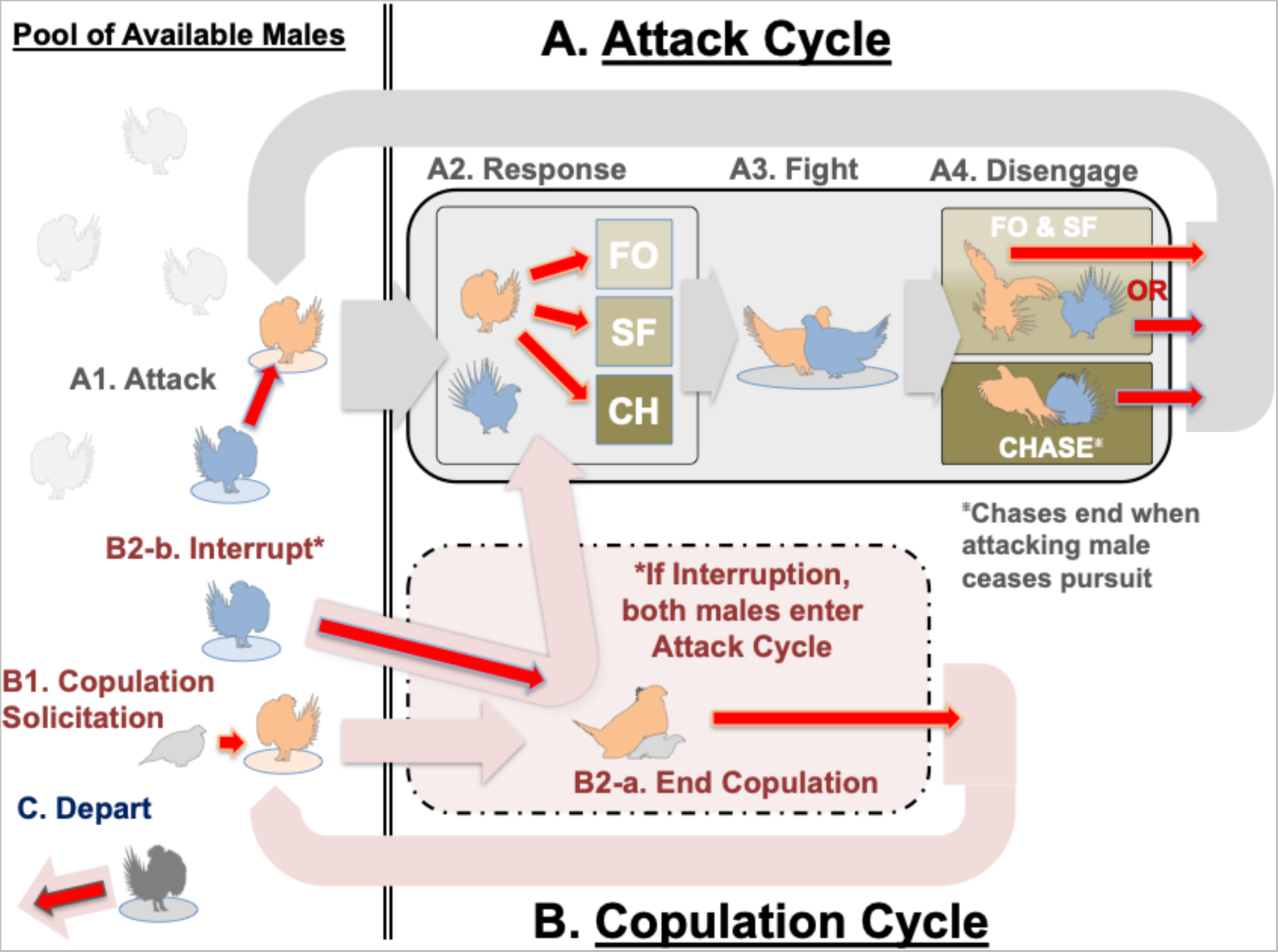
Event scheme and model constraints for the Relational Event Model (REM). Red arrows represent all eleven endogenous event types for which the model estimates hazards, as they are coded in the model. Wide arrows represent transitions of males out of and back into the pool of males that are available to begin an interaction/copulation/departure and transitions between stages of the behavioral cycles. Once a male is attacked (A1), he and the attacking male enter the Attack Cycle (A; solid box) and become unavailable to interact with any other males until they disengage and return to the pool (A4; wide gray arrow). Upon entering the Attack Cycle the receiver of the attack immediately responds (A2), dictating whether the interaction will be a Face-Off (FO), a Short Fight (SF), or a Chase (CH). These dictate the possibilities for the disengage event (A4): either male may disengage from a Face-Off or Short Fight, but only the attacking male may disengage from (and thereby win) a Chase. When a female solicits a copulation from a male (B1), that male enters a Copulation Cycle (B; dashed box). Males engaged in copulation are unavailable to attack other males (or depart, etc.) until the copulation ends and they return to the pool (B2-a; wide pink arrow). However, it is also possible for other males to attack the copulating male before it ends, and interrupt the copulation (B2-b), at which point both males enter an Attack Cycle (A2). Any male that is already in the pool of available males can depart the lek (C).

Using observations from a wild population of lekking sage-grouse in Wyoming, we investigated how aspects of fighting among male sage-grouse, and various other factors such as female presence, predict involvement in future fights, fight outcomes, solicitations for copulations (coupled with subsequent mounting by the male; hereafter referred to only as “solicitations,” but see *Event Types*), and interrupted copulations (see Table S1 for full list of possible model effect statistics with descriptions). In order to examine the idea of fighting as an attractive or informative element of male display, we fit two different base models: (1) the Aggression Model, which employs a single fixed effect for the solicitation hazard for all males, essentially forcing the model to use aspects of fighting to explain variation in copulation events for each male, and (2) the Differential Attractiveness Model, which uses individual-level fixed effects for solicitation hazards, allowing the model to attribute variation in solicitation hazard across males to some intrinsic property, such as their display attractiveness, in addition to (and independent of) the possible effects of their fighting behavior.

We find that fighting is not directly attractive to females. Rather, we show that fighting serves a complex and context-dependent role in this system, with some fighting, such as copulation interruptions and fighting with females present, being misaligned with female mate choice. We demonstrate clear incentives for males to interfere with one another through fighting, while at the same time we show that males regulate their own aggression in the presence of females and use fighting in a context-dependent manner to avoid future fights and copulation interruptions. These results suggest that the avoidance of fights and the overall regulation of aggression is likely necessary for a stable, functioning lek, with important implications for our understanding of lek evolution.

## Materials and Methods

### Field Data Collection

We conducted daily observations from March 11 to May 4, 2014 at the “Chugwater” lek, ∼25 km southeast of Hudson, in Fremont County, Wyoming. We video-recorded on-lek activity with one or two high-definition cameras (1080i; Sony HDR-HC1, Tokyo, and Canon Vixia HV40, New York) beginning at first light and continuing until all birds had departed the lek. Field assistants recorded videos from an observation blind on a hill ∼75 m from the lek, and made notes of individual males’ positions on the lek relative to a grid of stakes arrayed at 10 m intervals (Krakauer *et al*., 2009; Patricelli & Krakauer, 2010). This grid covered the center of activity on the lek, excluding peripheral males, who remained in the sagebrush surrounding the lek. Individual males were identified through spotting scopes via unique color band combinations and tail plumage patterns (Wiley, 1973; Perry *et al*., 2019). This combination of information enabled us to identify and track individual males on each video.

### Video Data Extraction

For this study, we extracted high temporal-density behavioral data from videos taken on consecutive days from March 19 to April 6, 2014, with the exception of March 27 due to poor weather, giving 18 video days of data. This sample coincides with the peak breeding period for this lek in 2014 and encompasses the longest stretch of days over which more than one female attended the lek, and the entire first-clutch breeding activity, including 72 of the 85 total copulations observed over the whole season. Often there is a second pulse of breeding later in the season when some broods fail and females re-mate before laying a second clutch (Wiley, 1973; G. Patricelli, Pers. Comm.). The first pulse likely represents females’ first choices.

Five research assistants were trained to extract data from videos using a standard training clip according to a standard protocol and vetted for precision and consistency. We extracted data describing the details and timing (to the nearest frame, or 1/30 s) of every direct interaction between every male on the lek as well as copulations with females for each day in the observation period. Because nearly every interaction we observed among males could be described as agonistic, we use the terms “fight” and “interaction” interchangeably. These behaviors, and how they were defined and coded, are described in Figure 1.

We also extracted a variety of other data from the videos. We recorded the timing of each male’s arrival on the lek, as well as the timing for females’ entrance and exit. Females’ entrance timing corresponds to the first female to appear, and their exit corresponds to the time of the last female to depart. We also defined female presence using a binary variable, where females are considered to be present when there are any females on the lek. We also recorded the instance of any disturbances to the lek that caused the birds to flush prematurely, like predators, airplanes, etc. We have four of these events in the dataset.

Finally, we recorded the position of every male relative to the 10 m stake array at the moment they either initiated or received an attack, as well as for all males present every five minutes independent of their activity. This allowed us to ascertain centroids of movement for each male on each day to control for spatial effects in our analyses.

We recognized three categories of males: males that have been positively identified by the field team and have persistent ID labels throughout the season, unknown “U males” that are consistently present on a particular observation day but are not observed attending on other days (either because they only attended for one day or the field team was unable to identify a known male on that day), and transient “X males,” usually juveniles/yearlings (Wiley, 1973), who typically arrive and then depart after a short time. Females can be distinguished from males by plumage, but they could not be individually identified.

Typically, a number of sage-grouse males will establish and occupy specific territories on the lek, returning to the same spot day after day to display (Wiley, 1973). For each day of video, we defined territory-holding males as those that had attended the lek for at least the previous two consecutive days.

Our video data from the 18-day observation period yielded 1506 unique interactions among up to 29 individual males, comprised of 14 identified males and 15 “U” males, plus interactions involving “X” males. On average, 8.56 (±2.01) males were present in the lek grid each day; females attended the lek for at least one span of time every day. We also observed 72 copulation events, 22 of which were disrupted by another male, a rate of interruption typical of Greater Sage-grouse (Wiley, 1973; Gibson & Bradbury, 1987).

### Relational Event Model (REM)

The REM directly models the unfolding of a sequence of events (in this case, observed social behaviors) through time. The goal of the REM is to identify the best-fitting model that elucidates underlying processes that could have generated a particular sequence, or set of sequences, of events.

This is done by the modeling a *hazard* for each kind of event that is possible at each moment in time. The hazard is a conditional rate of occurrence and can be thought of as an event’s propensity to occur. For example, an event that is impossible at a particular moment – e.g., female solicitation when females are absent – will have a hazard of 0, while the hazard for events that are more prevalent and quicker to occur will be larger. (Specifically, holding all other factors constant, and absent interruptions, the expected waiting time to an event with hazard *h* is 1/*h*.) An event hazard may be influenced by the past event history, the event type, and/or other covariates. To capture this, the hazard of each event is modeled as a multiplicative function of a series of statistics, each of which encodes the effect of some mechanism – be it sender/receiver identity, or some effect of a sub-sequence of historical events (e.g. recency of getting attacked) – that might enhance or inhibit the rate of occurrence (Butts & Marcum, 2017).

Adapting the notation from Butts and Marcum (2017) for the current study, we formalize the expression for an event hazard as follows: given a set of event types *C*, we can define a single event *a* as a list of elements containing the type of event *c* = *c*(*a*) ∈ *C* and the time the event occurred 1 = 1(*a*), such that *a*=(*c*,*t*). The history of past events from time 0 leading up to time *t* is denoted as *α_t_* = {*a_i_* : 1(*a*) :.: *t*}, and 𝔸 is the set of possible events at any moment. *X_a_* is a set of covariates that may be associated with a given event *a*, such as the categories of the males (identified, “U,” or “X”) and the individuals’ proximity in space. We can then express the hazard, ″α, of an event *a* at time *t*, given event history *α_t_* as:

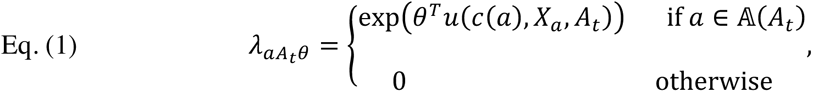

where *θ* is a vector of parameters governing the effects of *u*, a vector of statistics representing the effects of mechanisms that may influence the hazard of a given event. The *u* statistics may be functions of the event type and/or past events. Each unit change in *u_i_* multiplies the hazard of the event by exp(*θ_i_*). In this way, the *u* statistics are analogous to the effects in a conventional hazard model (Blossfeld & Rohwer, 2001), with *θ* being the model coefficients which we infer from data.

As is typical with REMs, we assume the state of the system – the set of possible events and their associated hazards – is “piecewise constant”, with changes only precipitated by the realization of new events. For example, a new event, such as an attack, might affect the propensity of other actions and/or limit the individuals available for interaction going forward. We update the state of the world to reflect this, and then the context is updated again when a new event occurs, such as a disengagement from the fight. This is reflected in Eq. 1: the hazard does not directly depend on *t* but rather on the temporally-ordered event history. Piecewise constancy is tantamount to the assumption that the waiting time between events is conditionally exponentially distributed, meaning that a given event with hazard *λ* at time *t* has a survival function *S*(*t*) = e^−*λ*^((t−t’) specifying its expected waiting time at time *t,* given a prior event at time *t*’. Thus, the probability of a particular event being the one to occur next (and at a particular time) is dictated by its hazard in proportion to the total hazard for all possible events. Exogenous events (introduced in Marcum & Butts, 2015) add flexibility within the piecewise constant model assumption, allowing for events to occur that are not endogenous to the system but nevertheless affect the system, or for bookkeeping devices such as “clock” events. We incorporate both here (see *Event Types* below).

The model thus specified, we test various hypotheses about the role of aggression in sage-grouse by selecting among possible models (sets of *u* effect statistics) and assessing goodness-of-fit to the observed sequence of events (see expression for model likelihood of an event history under the above-described process, incorporating exogenous effects, in Marcum & Butts, 2015). We conducted all REM analyses using the **relevent** package for R (Butts, 2015; R Core Team, 2019; code available upon request). We used Bayesian estimation of posterior modes to estimate model coefficients.

### Event Types

To effectively implement the REM, we had to create a modeling scheme that was faithful to the system and incorporated all the processes that might influence the behaviors of interest, in addition to the observed behavior events themselves. Although most of our event types map onto what would be considered dyadic sender/receiver “ties” in the conventional dynamic aggregated social network sense, our event type set includes several other types of events.

First, we have an event type for every pairwise permutation of males attacking/receiving, with the exception of males attacking themselves (Fig. 1A; Fig. 2A-1). To reduce the overall number of necessary event types, we defined an additional category of “response” event type wherein the receiver of an interaction nearly instantaneously responds to an attack by choosing the nature of the ensuing fight – they can choose to run away and begin a Chase, or stand their ground, beginning either a Short Fight or a Face-Off (Fig. 1B; Fig. 2A-2). There is also an event type encoding the disengagement from a fight for each male (Fig. 1C, Fig. 2A-4).

**Figure 3.**
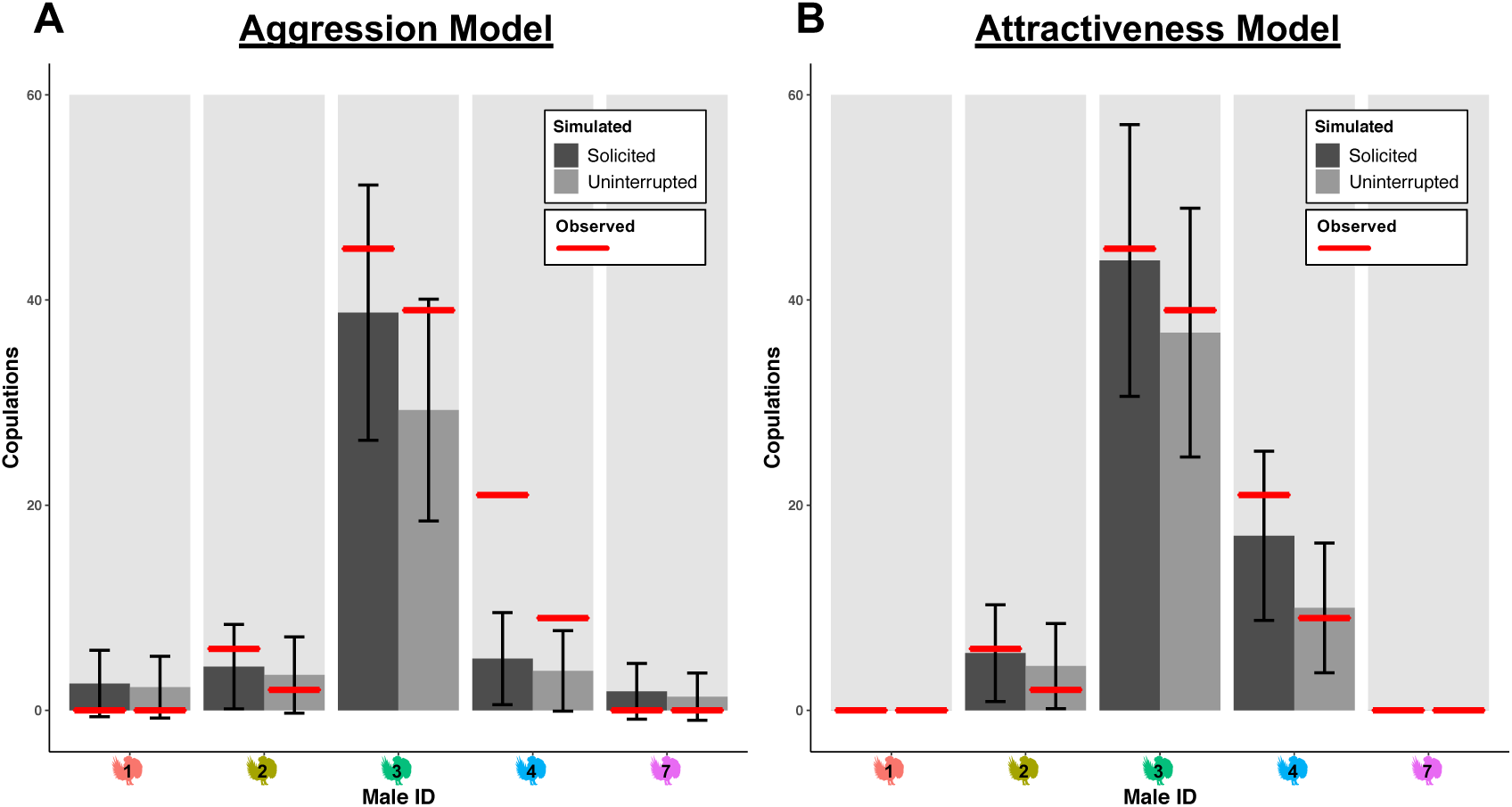
Comparison of simulation predictions of the numbers of solicitations for copulations and uninterrupted copulations for the Aggression Model (A) and the Differential Attractiveness Model (B). Plots show the mean total solicitations (dark bars) and mean total uninterrupted copulations (lighter bars) for each male across one hundred simulations for each day of the observation period. The simulated data were generated from the fitted coefficients of the alternative best-fitting Relational Event Models. The red lines are the true observed values. Error bars represent one standard deviation. The simulated numbers of solicitations and uninterrupted copulations from the Attractiveness model are much closer to the observed data than those from the Aggression Model. The data shown here are for the five individuals that attended the lek every day during the observation period

**Figure 4.**
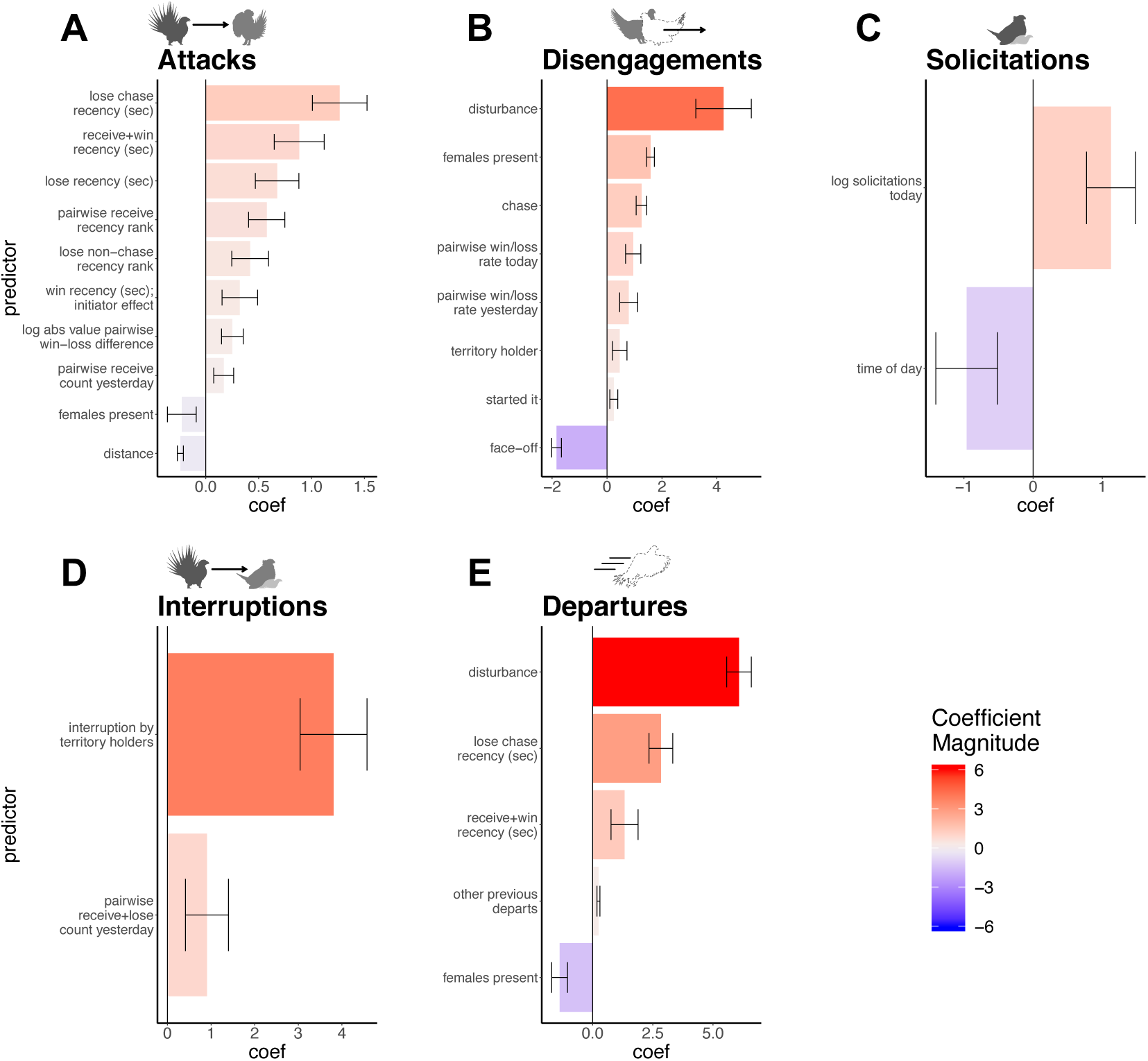
The values of the estimated coefficients for the set of effect statistics in the best-fitting Differential Attractiveness Model, arranged by category of event affected. Each unit change in the value of an effect statistic *i*, such as log(number of solicitations so far today), multiples the hazard (conditional rate of occurrence) of the event it affects by exp(coefi). Negative (bluer) values have a negative influence on the hazard of the events, positive (redder) values have a positive influence. Attacks (A) and Interruptions (D) refer to the hazard of a male receiving an attack or a copulation interruption, respectively, with the exception of statistics labelled “initiator effects.” Error bars represent two times the Bayesian posterior estimated standard errors, approximating 95% confidence intervals. See Table S3 for full model results including fixed effects.

Copulations are bounded by two events (Fig. 1D; Fig. 2B-1, 2B-2-a): (1) a solicitation/copulation start event – which includes a solicitation of copulation by a female and subsequent mounting by the male –and (2) an end copulation event, which occurs when the male dismounts. In some instances, a female may solicit a male to copulate, but the male does not mount (males are never observed to successfully copulate with a female without first being solicited). For the present analysis, we only include those events where a female solicited and then the male proceeded to mount. Thus, for the REM we defined “solicitation” events as beginning at moment the male mounts. Copulation interruptions (Fig. 1E; Fig. 2B-2-b) are defined in the REM as combination of other events: a solicitation/copulation followed by an attack directed at the copulating male prior to, and instead of, a successful copulation end.

We also included several “exogenous” event types – events with hazards that are dictated without respect to the events in the model. Thus, we assume that none of the events in the system can affect the occurrence of exogenous events. We treat events for the arrival of each male on the lek as exogenous (departures are treated as endogenous since they can be affected by events on the lek). Events for when females enter or exit the lek are treated as exogenous, dictating the presence of females, which may have an influence on the hazards of male behaviors (due to their size, crypsis, and high turnover during the day, females are modelled as a bulk actor such that females are considered present when there is at least one female on the lek). There are also exogenous “clock” events – a clock event occurs every fifteen minutes from the beginning of the observation period, allowing for effects of event history to change over time, and there is an event type that models disturbances from predators, humans, etc., that may cause the birds to prematurely flush from the lek. Finally, there is a special exogenous event type denoting the end of a daily observation period. In all, there are 1115 event types possible. Coded this way for the REM, the observed data contains a sequence of 5168 events.

### Support Constraints and Model Scheme

Not every event type is possible at every moment in time. Accordingly, we constrain the model such that it imposes zero hazards for impossible events (See Marcum & Butts 2015; Fig. 2). Constraints imposed on the model are as follows:

i. Only males currently present on the lek may interact.
ii. Solicitations are only possible if females are present on the lek.
iii. The “Attack Cycle”: A male may initiate or receive an attack only if he is available (present on the lek and not already fighting or copulating). Upon initiating an interaction, that pair of males is removed from the pool of available males (Fig. 2A-1). After a male receives an attack, the immediately following event is constrained to be a “response” event wherein the receiving male decides whether the interaction will be a Chase, a Short Fight, or a Face-Off (Fig. 2A-2). Once the attack cycle begins for a pair of males, other events may continue taking place, but the males in question are unavailable for other events involving them until one of them disengages (represented by the solid box around Fig. 2A-2 and 2A-3). Upon disengaging, both males involved return to the pool of available males (Fig. 2A-4). A male must disengage from an interaction and return to the pool of available males before becoming involved in a new one.
iv. The “Copulation Cycle”: A male that is actively copulating may be attacked by other males but cannot initiate an attack of his own until he has ended the copulation (represented by the dashed box around Fig. 2B-1 and 2B-2-a). An attack directed at a copulating male ends the copulation (obviates the copulation end event) and that pair of males then enters the attack cycle, where they are removed from the pool of available males (Fig. 2B-2-b).
v. X males: At a few points in the observation period, there is more than one unidentified male engaged in an interaction at once. For tractability, all X males act as a single bulk- actor which can potentially be involved in interactions with multiple partners simultaneously and is therefore not subject to the same constraints as the other types of males. X males are considered present and available for interaction if at least one is currently on the lek.
vi. Males can depart the lek only if they are already present and not otherwise engaged in an attack/copulation cycle (Fig. 2C).

### Fixed effects for the base intercept-only REM

We first selected the best-performing base intercept model with fixed effects for individual-level variability in intrinsic hazards of behaviors, as well as among-category differences in hazards between identified, “U,” and “X” males. We fit effects for attack and receive hazard, as well as the hazards of initiating different “response” events (Short Fights, Chases, and Face- Offs), disengagements, and departures (See Table S2 for detailed descriptions of the fixed effects included in all models).

Keeping all other fixed effects in common, we tested two different model formulations for fixed effects for solicitation hazard: (1) an “Aggression” model, in which we fit a single base solicitation hazard common to all males, forcing the model to attempt to use male-male fighting events to predict variation in solicitation; and (2) a “Differential Attractiveness” model, in which we fit separate fixed effects for solicitation hazard for each male that was ever solicited during the season. Males that never mated during the season were assigned the base hazard. This model allows for the possibility that individual males may possess intrinsically attractive attributes (such as display quality or effort) that in part dictate their hazard of being solicited for copulation, independent of their fighting behavior.

### Sufficient Statistics

We created a set of 301 possible effect statistics representing potential mechanisms by which histories of various fighting behaviors and other events – such as female presence, previous copulations, exogenous disturbances, and covariates such as distance between males – might influence the hazards of five different kinds of events: attacks, disengagements, solicitations, interruptions, and departures (for a full list of all statistics with descriptions, see Table S1). We developed the statistics based on the observable aspects of the behavior of individuals and the information about these behaviors plausibly available to each bird. For example, to avoid making assumptions about the social memory of individual sage-grouse, we tested statistics that were based on both pairwise interaction history as well as overall counts, and we tested these across multiple temporal scales, ranging from cumulatively across the entire observation period to on the order of seconds (Table S1). In keeping with this approach, statistics relating to solicitation hazard include only those events that occurred previously in the presence of females. We do not, therefore, directly test for higher-order effects of fighting or other activity on male display that are not mediated by female mate choice. For example, we do not examine whether fighting that occurs in the absence of females might cause fatigue, negatively affecting a male’s display effort.

Statistics specifically relating to copulation interruption only apply when a male on the lek has begun copulating. Therefore, these statistics can be thought of as testing for event-history-by-copulation interaction effects, in the statistical sense. Finally, on the first and on the ninth day—which is missing a true consecutive previous day due to poor weather—we simply set statistics that count events from the previous day to their base value, usually zero.

### Model Selection

For both alternative base sets of fixed model effects represented by the Aggression Model and the Differential Attractiveness Model, we began with the base model and used a forward selection algorithm based on BIC to choose the best fitting final model from among candidate sets of effects (different sets of effect statistics, *u*). We ran each forward selection process until the addition of effect statistics no longer improved BIC, and then we inspected the results to ascertain that no early statistics had become non-significant due to collinearity. Ultimately, the final models respectively identify subsets of the possible 301 effect statistics that, through their effects on event hazards governed by inferred parameter magnitudes, provide the most probable fit to the observed data.

### Simulations

We used the estimated coefficients from the final alternative REMs to simulate new event data for each day of the observation period. This approach allows us to compare how well different model formulations can approximate the actual grouse behaviors that we observed. These simulations also help us understand how inter-male aggression may indirectly impact males’ overall mating success. Although the REMs give insight into what fighting behaviors, or other event histories, influence a males’ propensity to be solicited, they fit hazards of solicitations conditional only on when they are possible in a given moment. This does not consider that aggressive behaviors might have a large effect on overall mating success by causing solicitations to be impossible for substantial stretches of time.

To generate simulated data after each new event, we can calculate new hazards for each predicted (and possible) next event using the REM coefficients and sufficient statistics, and then choose the next event and its timing based on those hazards and the model’s assumptions. Therefore, the next event’s timing is chosen randomly from an exponential distribution with the mean 1/(total hazard of all possible events), and the identity of the next event is chosen as a random categorical variable weighted by its hazard’s proportion of the total hazard (code available upon request). For each simulated day, we fed in the observed data from the previous day (the previous day being the longest relevant timescale identified in our models; see Results). We also used a scaffold of “scheduled” exogenous events based on the values observed on the day in question for: the first arrival times for each male present, the female entry and exit times, and the end of video recording. If a male departed the lek during the simulation, we used the average interval of absence for that male in the observed data as the mean waiting time of an exponential distribution to schedule their subsequent re-arrival. In the observed data, most males do not return once they have departed. However, X males often arrive and depart many times over the course of the day.

We calculated summary counts of events and summary network metrics for the simulated data using the **sna** package in R (Butts, 2016; R Core Team, 2019; code available upon request).

## Results

### Is fighting part of the display?

Our results indicate that fighting does not have a direct role in attracting females. While we find, like some previous studies (e.g., Alatalo *et al*., 1991; Höglund *et al*., 1997; Rintamäki *et al*., 2001; Hämäläinen *et al*., 2012), that solicitations (mating events) are positively correlated with fights in aggregate (Fig. S1), the REM does not support a temporal, mechanistic link between fighting and solicitations. Aggregation of fighting data over the entire time period eliminates information on the sequence and timing of events, and misleadingly implies a mechanistic relationship between fighting and solicitations that does not exist.

The best-fitting Differential Attractiveness Model is definitively preferred over the best-fitting Aggression Model by BIC (Tables S3 and S4; see Appendix A1 for analysis of model performance). In other words, the model which assumes that variation in solicitation is determined largely by male-male aggression does a substantially worse job in explaining the observed behavioral data than the model which allows for differential intrinsic male attractiveness to explain variation in solicitation hazards. Furthermore, the final set of effect statistics comprising the best-fitting Differential Attractiveness Model does not include any fighting-related effects on solicitation; effect statistics like perpetrating interruptions and winning fights were not necessary to explain solicitations for males to mate, regardless of the temporal scale of the event history. The number of copulations previously that day (indicative of day-level variation in display quality, or possibly mate choice copying; Gibson & Höglund, 1992) and time of day are sufficient to explain much of the variation in solicitation hazards once variation in intrinsic attractiveness is accounted for (Fig. 4C).

These findings are corroborated by our simulations. The Aggression Model, which is forced to use fighting effects to explain variation in solicitation hazards, performs much more poorly at predicting the distribution and number of copulations per male for the observation period (Fig. 3).

In addition to the superior performance in predicting solicitations, the Differential Attractiveness Model also reveals that males are less likely to start fights, more likely to end fights sooner, and less likely to depart the lek when females are present (Fig. 5). This strongly suggests that male sage-grouse have evolved to reduce aggression in favor of display in the presence of females, providing direct evidence against the hypothesis that females have evolved to use male aggression to inform their mate choices.

**Figure 5.**
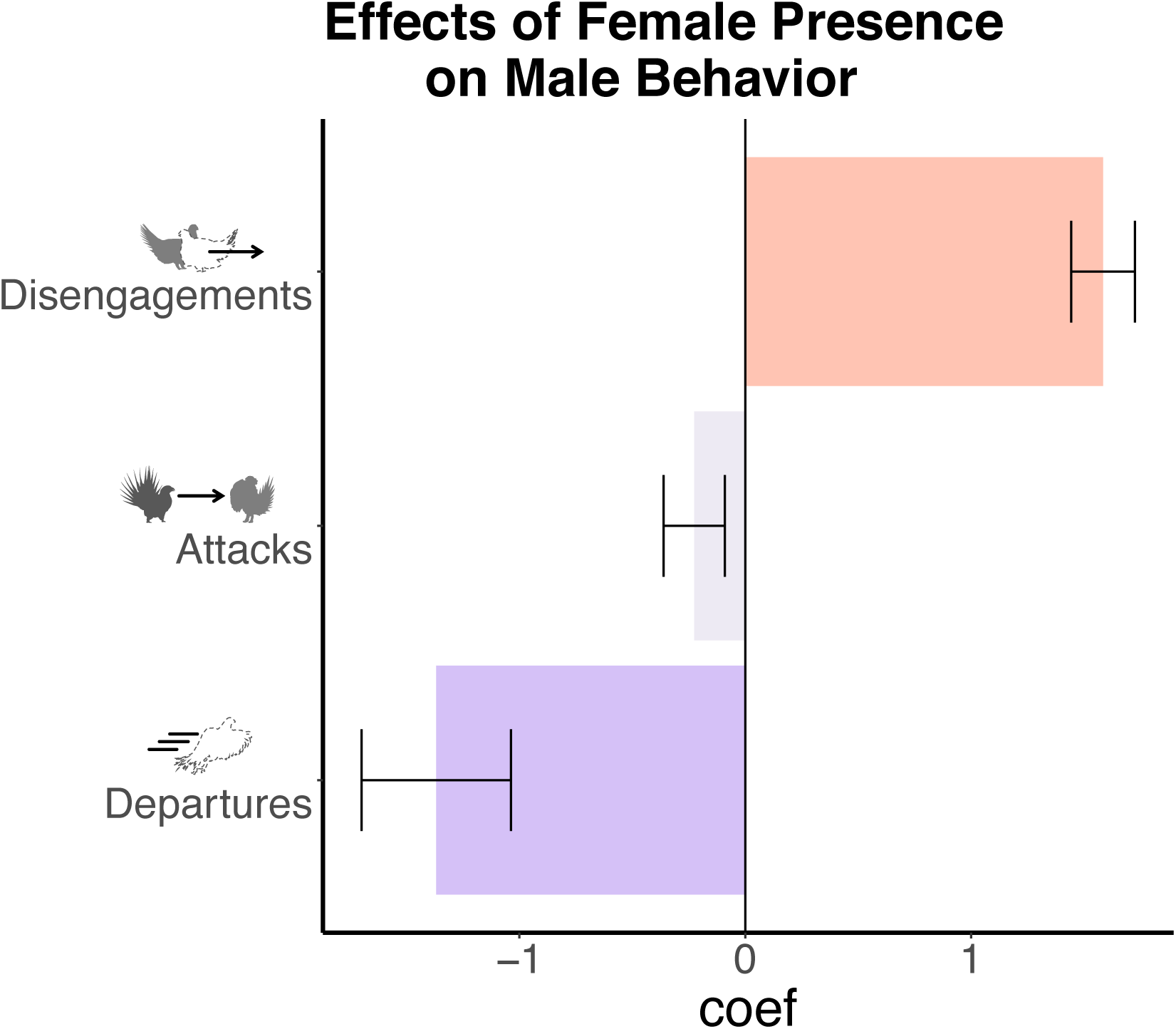
The effects (coefficient magnitudes) of female presence on the hazard of male Disengagements, Attacks, and Departures (y-axis).

### The indirect effects of fighting on mating success

The REM indicates that fighting begets more fighting, and males that fought amongst one another previously are likely to do so again. In general, being attacked and losing lead to a higher risk of being attacked in the future. The only category of interaction that does not greatly increase a male’s propensity to be attacked is “attacking + winning” (i.e., winning fights that it started; Figs. 4 and 6). One exception to this pattern is the “initiator effect” in the model, which shows that males that have just won a fight are more likely to go start another fight. However, when females are present, males are overall less likely to initiate attacks and the initiator effect is dampened such that “attack + win” interactions in particular, which do not have any other effects associated with them, do not appreciably increase a males’ hazard of initiating a new attack (Fig. 4A; Fig. 6).

**Figure 6.**
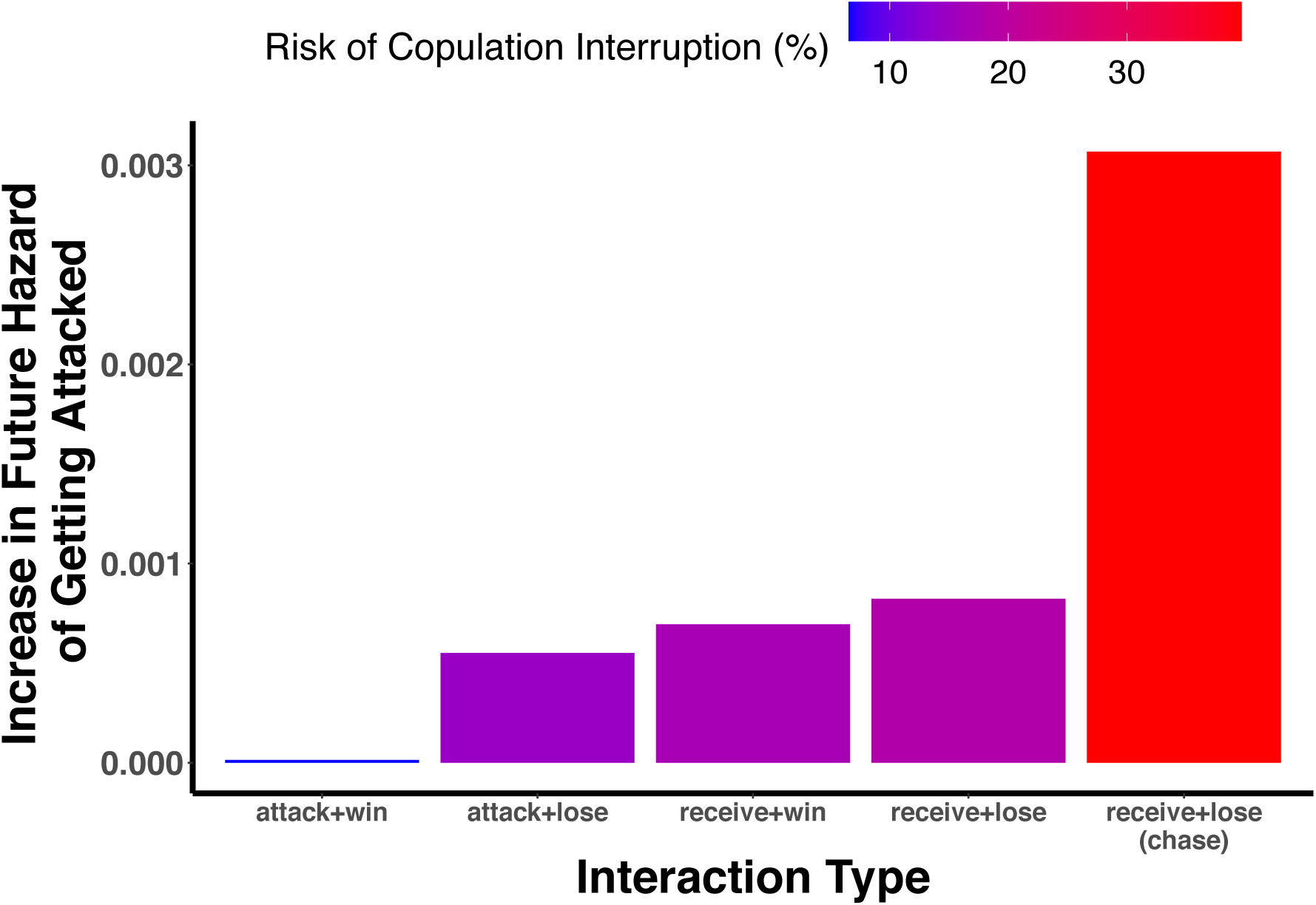
The effect of different types of interactions on an average male’s immediate future hazard of being attacked by another male (y-axis), and the corresponding probability of being interrupted if that male were to begin copulating (color scale). Calculations are based on the coefficients from the best-fitting Differential Attractiveness REM, for a simulated scenario involving six territory-holding males all with global-average attack rates and pairwise distances, with no prior history and with females present. Each single interaction is a combination of the initiation (the focal male attacks or receives; Fig. 1A) and the disengagement, which represents either a win or a loss for the focal male (Fig. 1C). All interactions are Short Fights (Fig. 1B) except for the rightmost bar representing the increase in attack hazard resulting from a Chase (Fig. 1B)

The absolute value of pairwise win-loss difference for the day also predicts the hazard of attacks received (Fig. 4A). This means males with drastically different records of fighting success are more likely to interact than pairs of males that are more evenly matched. This result appears to be substantially driven by the pattern of territorial males attacking and chasing juvenile/unidentifiable males.

Underlying the effects on the hazard of being attacked due to event histories, we also found significant individual variation in fixed effects for baseline hazard of receiving attacks (Tables S2 and S3), as well as a strong effect of pairwise centroid distance, with closer males being much more likely to interact (Fig. 4A).

In addition, there are several factors that influence the outcomes and durations of fights: pairwise win/loss rate today – and to a slightly lesser extent, yesterday – predict wins in future pairwise contests (Fig. 4B). Starting a fight is associated with winning (Fig. 4B). The remaining effects influencing disengagements (Fig. 4B) predict the overall duration of the fights. Fights end abruptly when there is a disturbance, and the presence of females is associated with much shorter fights overall. As expected, Chases tend to be short-lived, whereas Face-Offs may extend for a longer period than average before one male disengages (Fig. 1B; Fig. 4B).

The model also identifies effects that significantly influence the hazard of copulation interruptions on top of the background risk of being attacked (Fig. 4D). The effect statistic “interruption by territory holders” indicates that if one is going to be interrupted while copulating, it is likely going to be perpetrated by a territory-holding male. Additionally, it indicates that starting a copulation greatly increases the hazard of being attacked, suggesting that copulation is a major impetus for other males to attack when they otherwise would carry on displaying: the coefficient of 3.8 translates to an increase of an individual’s baseline hazard of being attacked by a factor exp(3.8)::45. Furthermore, interestingly, the overall interruption hazard as well as the identity of the interrupting male is highly influenced by the copulating male’s pairwise history of getting attacked by and losing to those other males (“receive + lose” count) on the previous day.

Finally, beyond disturbances flushing the birds from the lek, there are several social effects on lek departure by males. Males are more likely to remain on the lek in the presence of females, and are more likely to depart after recently losing a Chase (Fig. 4E). Indeed, Chase interactions often end with the losing male being chased off the lek entirely. Interestingly, “receive + win” recently also predicts departure from the lek, but this effect is entirely canceled by the effect of female presence (Fig. 4E). We may therefore interpret this as males departing the lek upon winning a challenge when females have already left for the day.

Our results make it clear that aggressively driving off competitors does not itself explain the observed variation in mating success. None of the males in our dataset that get chased off the lek are ever observed to mate (n=14), but there are many other males that also never mated and were never chased off (n=12). While fighting to drive off competitors (to establish and maintain a territory on the lek) likely plays an important role for mating success, we cannot speak to that here given that the lek was already formed at the beginning of our observation period.

Taken all together, the REM demonstrates that the role of fighting on the sage-grouse lek is dynamic and complex. While fighting does not directly induce females to choose mates, fighting affects mating success in two ways: through the time a male has available to mate (since a male cannot mate while he is fighting), and through managing the risk of copulation interruption.

Indeed, we observed considerable variation in the percentage of time that territory-holding males spent engaged in fights while females were present: 9.37% (±11.51) on average, with a single-day maximum of 64.1% (Fig. S2). It is clear, therefore, that mating success is in part related to a male’s tactical ability to avoid fighting or engage in fighting that does not consume time or increase his risk of interruption.

For example, on April 1, 2014 (Fig. 7), male 4 had the highest average inferred solicitation hazard of all males that day, driven by a streak of copulations in the morning. However, due to his history of receiving and losing fights the day before, a few key losses early in the day, and his overall aggressiveness, male 4 attained only one uninterrupted copulation out of ten mating attempts. Indeed, male 4 spent the most time fighting of any male on that day (15.3% of his 80 minutes in front of females) and the REM inferred an average moment-by-moment risk of interruption of more than 80%. In contrast, male 3 was attacked far less frequently on that day and therefore was less susceptible to interruption and to being drawn into time-consuming chains of aggressive interactions with other males, spending only 10% of his time fighting. Although male 3 had far fewer (four) solicitations that day than male 4, he was interrupted only twice, and so had the greatest number of uninterrupted copulations.

**Figure 7.**
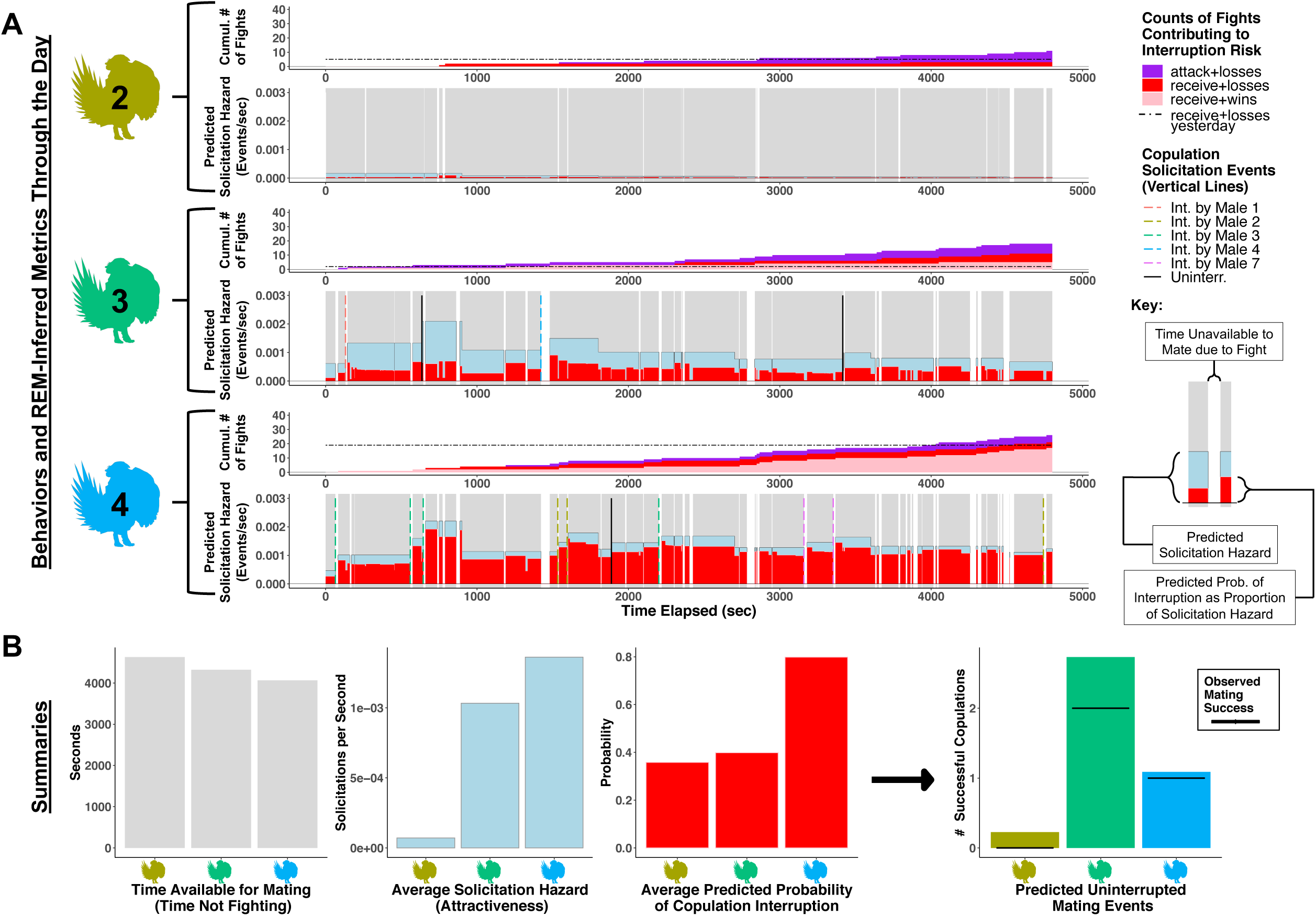
Example of the effects of fighting among sage-grouse males on availability to mate and copulation interruption risk from April 1, 2014. Data shown here is for the top three males in terms of mating success— ID’s 2, 3 and 4 — from the beginning of the day to the moment the last female exited the lek. Panel (A) shows counts of behaviors and Relation Event Model-inferred metrics through time. Moment-by-moment solicitation hazard (light blue) and copulation interruption risk (red) are calculated from the fitted coefficients of the best-fitting Differential Attractiveness Model. Interruption risk is the probability that, if a male were to begin copulating at that moment, another male would attack him before he naturally ended the copulation, based on the relative hazards of those events as predicted by the REM. In (A) the interruption risk (a percentage) is displayed as a proportion of the solicitation hazard for ease of comparison. The upper plots for each male show the cumulative counts of different kinds of fights that contribute the risk of being interrupted while copulating. These come in initiation- disengagement pairs: i.e., an attack+win is an interaction wherein the focal male initiated the attack and was also the winner. The horizontal dashed lines show the total number of receive+losses for each male from the previous day, a metric that is also important for interruption risk (Fig. 4D). Panel (B) shows summary bar-plots for total time available to mate, average solicitation hazard, and average probability of copulation interruption. The predicted total number of uninterrupted mating events is calculated as the Riemann sum over the rates of predicted uninterrupted copulations (the hazard of solicitation * the probability of interruption * the time interval available to mate between observed events) through time for each male.

## Discussion

Our analysis reveals that fighting behavior is unnecessary to predict copulation events in a Greater Sage-grouse lek, with individual attractiveness instead playing a central role. We recovered no evidence that fighting is directly attractive, informative, or functions in an additive way with display to contribute to the solicitation of copulation by females. In fact, males are less likely to start fights and more likely to end them sooner in the presence of females on the lek (Fig. 5). The importance of intrinsic male attractiveness is supported by extensive evidence that the elements of the famously elaborate display in Greater Sage-grouse, and those of other grouse species, are associated with mating success (Gibson & Bradbury, 1985; Gibson, 1996; Perry *et al*., 2019). However, our result that fighting is not attractive may strike as counter-intuitive, since fighting behavior in these systems is often also correlated with mating success in aggregate (e.g. Alatalo *et al*., 1991; Höglund *et al*., 1997; Rintamäki *et al*., 2001; Hämäläinen *et al*., 2012). Indeed, we recover a similar pattern in our aggregate data (Fig. S1). However, the causal relationship potentially implied by this correlation does not hold when we account for the sequence and timing of events in the Relational Event Model.

Our results demonstrate the Relation Event Modeling framework’s broad applicability to difficult behavioral questions in which the order and timing of events really matters for our ultimate interpretation of underlying processes: in this case, a traditional correlational study is inadequate to differentiate the causal relationships among fighting, display, and copulation events, yielding misleading results.

The main influences of fighting on mating success are indirect, affecting the opportunities for solicitations and the success of copulation attempts. The effects of fighting on time available to display and to mate, and particularly on risk of copulation interruption, had a large negative impact on overall mating success for the males in our dataset (Fig. 3; Fig. 7). Thus, it appears that the function of male aggression in lek behavior is really a question of how males might use fighting and social tolerance (non-fighting) to avoid the negative effects of fighting.

The best strategy for males seems to be to not start fights unless they are going to win. The REM demonstrates clearly how fights can take up a lot of time and build on one another, leading to self-sustaining webs of conflict that are difficult to diffuse (Figs. 6 and 7; Pellis *et al*., 2013; Balian & Bearman, 2018). We cannot say definitively why initiating fights would be good for an attractive male, but our results suggest the possibility that engaging other males’ time through fighting is a way for less-attractive males to attempt to increase their own relative probability of mating, thus flattening the skew in mating success. Given that an individual male cannot totally control whether they end up fighting, winning contests and having a history of winning contests is important for deterring interruptions in particular, since losing fights contributes greatly to the risk of interruption (Figs. 4D, 6 and 7).

The REM also indicated a role for individual males’ intrinsic hazard of being attacked (Tables S2 and S3). This could be associated with a covariate that might affect the risk of attack that other males can evaluate without needing to interact (such as body size or display behavior), or which other males could be aware of due to past history beyond the scope of our study, such as a competitor’s age. Alternatively, variation in individual-level hazard of being attacked could be a proxy for some aspect of territoriality, such as relative territory size given centrality, which has previously been associated with mating success in lekking grouse (e.g., Höglund *et al*., 1997; Rintamäki *et al*., 2001). However, our results suggest that females are not attracted to the territories or territory defense ability *per se*, but rather to the displays that successful territory defense facilitate, as some have previously speculated (Alatalo *et al*., 1996; Patricelli *et al*., 2011). Given that we see a strong effect of pairwise centroid distance on the risk of getting attacked (Fig. 4A), behavior such as attacking and winning may represent a constant underlying requirement for getting the space necessary for display and successful copulation. This is possibly why we do not observe a direct effect of these fighting behaviors on solicitation.

Overall, it is clear that male aggression has a complex relationship to female mate choice. Fighting is not directly attractive. Fighting and copulation interruption are forms of sexual coercion (Snow *et al*., 2019), distracting attractive males away from displaying, preventing females from evaluating those displays, and in some cases physically preventing females from fulfilling their mate choices. However, fighting is aligned with mating success insofar as winning fights contributes to social stability and protects female’s ability to choose based on display through deterring interruptions and possibly maintaining territory boundaries. In short, aggression is an essential part of successful male sage-grouse reproductive behavior, but to be used successfully, it must be deployed in a context-sensitive manner to reduce the risks of disruption to display and copulation. This is in line with other work on this group of birds showing that they can tactically allocate resources in other important domains such as display (Patricelli & Krakauer 2010; Perry *et al*., 2019).

### Implications for lek evolution

Understanding lek-breeding behavior has long been a challenge in evolutionary biology for a number of reasons (Taylor & Williams, 1982; Kirkpatrick & Ryan, 1991; Wiley, 1991; Höglund & Alatalo, 1995). One key has been to understand what forces cause males to cluster in space in the face of increasingly intense competition for mates (Emlen & Oring, 1977; Bradbury & Gibson, 1983; Queller, 1987; Beehler & Foster, 1988; Théry, 1992; Stillman *et al*., 1993; Kokko, 1997).

Our analysis shows lekking behavior to be a balancing act among several conflicting forces. When we consider that fighting is unlikely to be an aspect of display, it is clear that the regulation of aggression plays a crucial, yet underappreciated, role in how leks evolve in systems where females exercise choice.

The REM demonstrates that some males have clear incentives to start fights, both to waste other males’ time and interrupt competitors’ copulations, and that these behaviors interfere with the efficacy of female mate choice (Fig. 7). It is easy to see that all males, particularly less attractive males, acting in their own self-interest would lead to a “Tragedy of the Lek” wherein the lek would cease to function as an arena for females to freely make mating decisions. Clearly, achieving sexual success on the lek, and a stable lek overall, requires avoiding indiscriminately aggressive behavior in favor of a context-dependent strategy.

The REM also identifies specific behavioral mechanisms by which the “Tragedy of the Lek” may be avoided: (1) self-regulation of the initiation of fights, particularly when females are present, and prolonged fighting when females are absent (Fig. 5), and (2) reducing risk of copulation interruptions through adequate territory defense and dominating in pairwise fights over recent timescales. These all potentially represent evolved social behaviors that promote female choice (Patricelli *et al*., 2011), and are alternatives to the dissolution of leks in the face of courtship disruption (Foster, 1983; Westcott, 1997; Prum, 2017). Indeed, there is empirical evidence that disturbance to the established social structure of leks increases fighting and may decrease the overall mating success of lek members (Ballard, 1971; Ballard & Robel, 1974; Robel & Ballard, 1974; Westcott, 1997).

The selective regimes underlying the evolutionary transition from aggressive, solitary males to more socially-tolerant lekking males with more complex social lives warrants further theoretical and empirical study, particularly with respect to female behavioral responses to aggression. Our observation that sexual success is associated with the successful management of males’ own aggression and the aggressive attacks of rival males supports the hypothesis that stable lek behavior evolves through female choice for those males *and* those leks that reduce the costs of sexual coercion. This is consistent with the intriguing possibility that the existence of leks – i.e., avoiding the Tragedy of the Lek – is the result of a “remodeling” process (Prum, 2017), wherein females use mate choice to transform male social behavior in ways that lower sexual coercion and facilitate the efficacy of their mating preferences.

We recognize that the scope of this study is necessarily limited, because though we have high-density data, it derives from only a single lek in a single year. Sage grouse leks vary considerably in many ways: size, mating skew, level of aggression, sensitivity to disturbance, etc (Scott, 1942; Wiley, 1973; A. Krakauer, Pers. Comm.). Future studies should examine how the factors that influence behaviors of interest may vary across populations, geography, and time. Even more incisive would be to take a comparative approach to examine the variation in how aggression is regulated across related taxa.

## Supporting information

Supplemental_reproducible_code

## Acknowledgements

This work was supported by the NIH Predoctoral Training Program in Genetics under grant no. 5T32GM007499-38, Yale Institute for Biospheric Studies Doctoral Pilot Grant, Yale University EEB Chair’s Fund, and the French National Research Agency (ANR) under the Investments for the Future (Investissements d’Avenir) program (ANR-17-EUR-0010) to SSS; the W.R. Coe Fund to ROP; the National Science Foundation Graduate Research Fellowship under DGE-1148897 to ACP, DGE-1122492 to SSS, and DGE-1419118 to RL; National Science Foundation grant IOS-1258217 to GLP, AHK and Jennifer S Forbey (Boise State University) and USDA Hatch funds (Project #CA-D-EVE-2264-H) to GLP. Thank you to JS Forbey and the 2014 Sage Grouse Field Team, and thank you to the many helpful undergraduates at UC Davis and Team Grouse Yale who helped score behaviors from field videos. Fieldwork was approved by UC Davis IACUC Protocol# 18080 and Wyoming Game and Fish Permit #405. Thanks also to B. Evans and The Yale Center for Research Computing for support and access to high-performance computing resources.

## Appendix A1. Model Performance and Adequacy

Our best-fitting Differential Attractiveness Model (the best-fitting model overall) saw a 69.5% overall reduction in model deviance, with a 22.4% improvement over the base intercept model (see Tables S3 and S4 for full model results). This is on par with other well-performing REMs (Butts & Marcum, 2017). For each moment in time, the REM estimates a hazard for every event that could possibly occur next, with the event with the highest hazard being the one the model considers the most likely to occur (rank one), and events with successively smaller hazards being ranked lower in terms of likelihood to occur. We can look at another facet of model performance by looking at where the realized, observed event in each moment falls among the model’s rankings of hazards for the possible events at that moment. The model predicts the exact next event to occur (ranks the realized event as rank one) 54% of the time, with the mean rank of all observed events being 3.4 (Figs. S3 and S4). For attack events specifically, the mean conditional rank of a given observed attack event is 5.51 with a median of 3, out of a mean number of possible attack events in each instance of 46.25. Half of all observed attack events are ranked within the top three most likely events by the model, with 95% of all realized attack events ranked within the top 17 (again out of 46.25 possible attack events on average; Fig. S5). Analysis of deviance residuals for each event (an indicator of how “surprised” the model is; Butts & Marcum, 2017) shows that the model performs relatively similarly across different types of events, with the exception of “response” events, as we constrain them to always occur 0.001 seconds after the initiation of an interaction (Figs. S6 and S7).

Furthermore, our simulations show that the top models perform quite well at recapitulating the observed numbers of events, numbers of solicitations, proportions of copulations that were interrupted, distributions of fighting times and lek attendance, and individual-level and general network properties of the behaviors on each day, with the distributions produced by the simulations largely overlapping the observed data (Fig. 3; Figs. S8, S9, S10, and S11)

**Figure S1.**
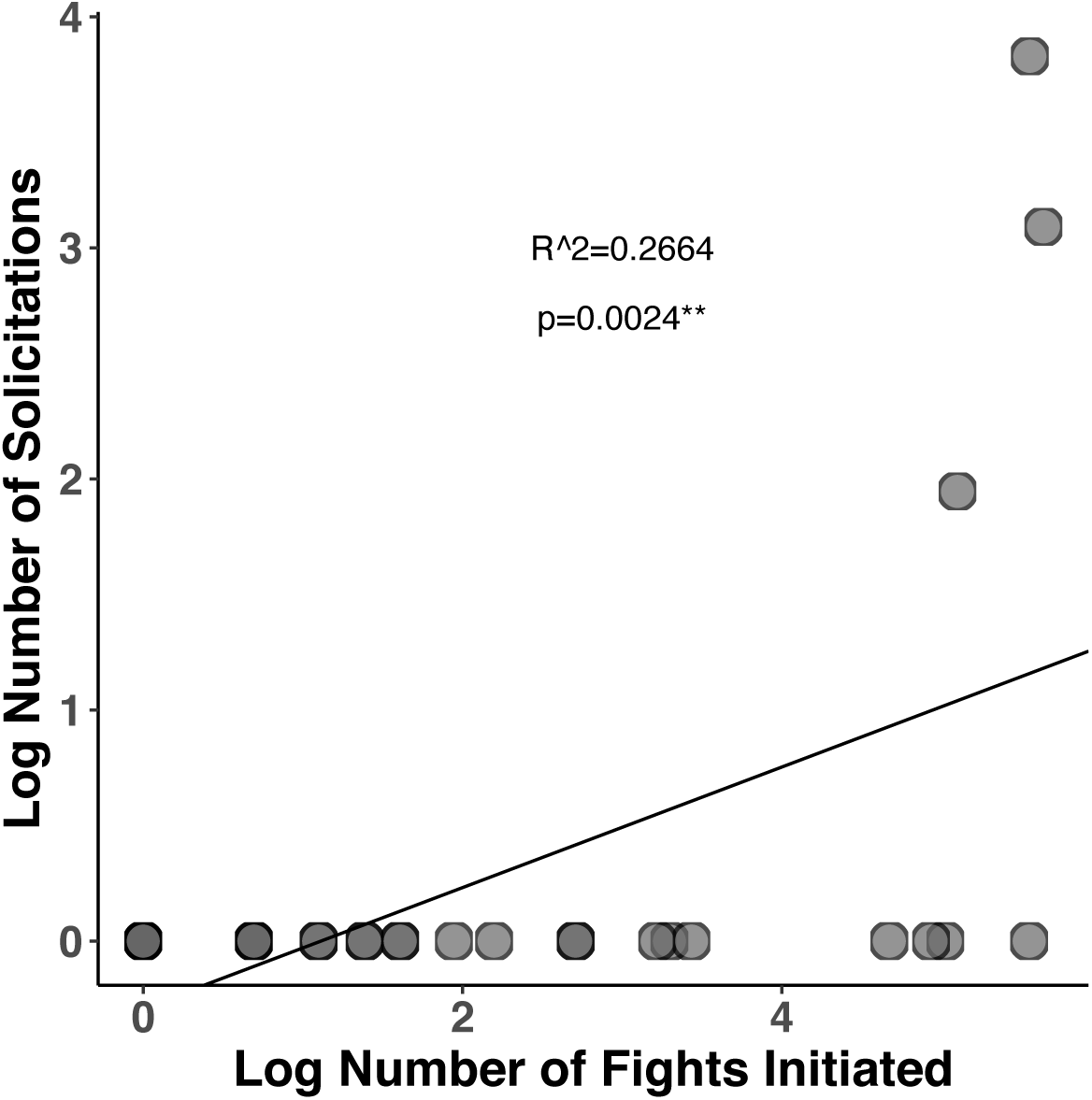
Misleading correlation between fighting and copulations in aggregate. Each point represents one male. Number of copulations and number of fights initiated are the totals for the entire dataset. The line is the line of best fit for a log- log linear model, with a statistically significant positive correlation (F=11.171,27, p=0.00245, R^2^=0.2664).

**Figure S2.**
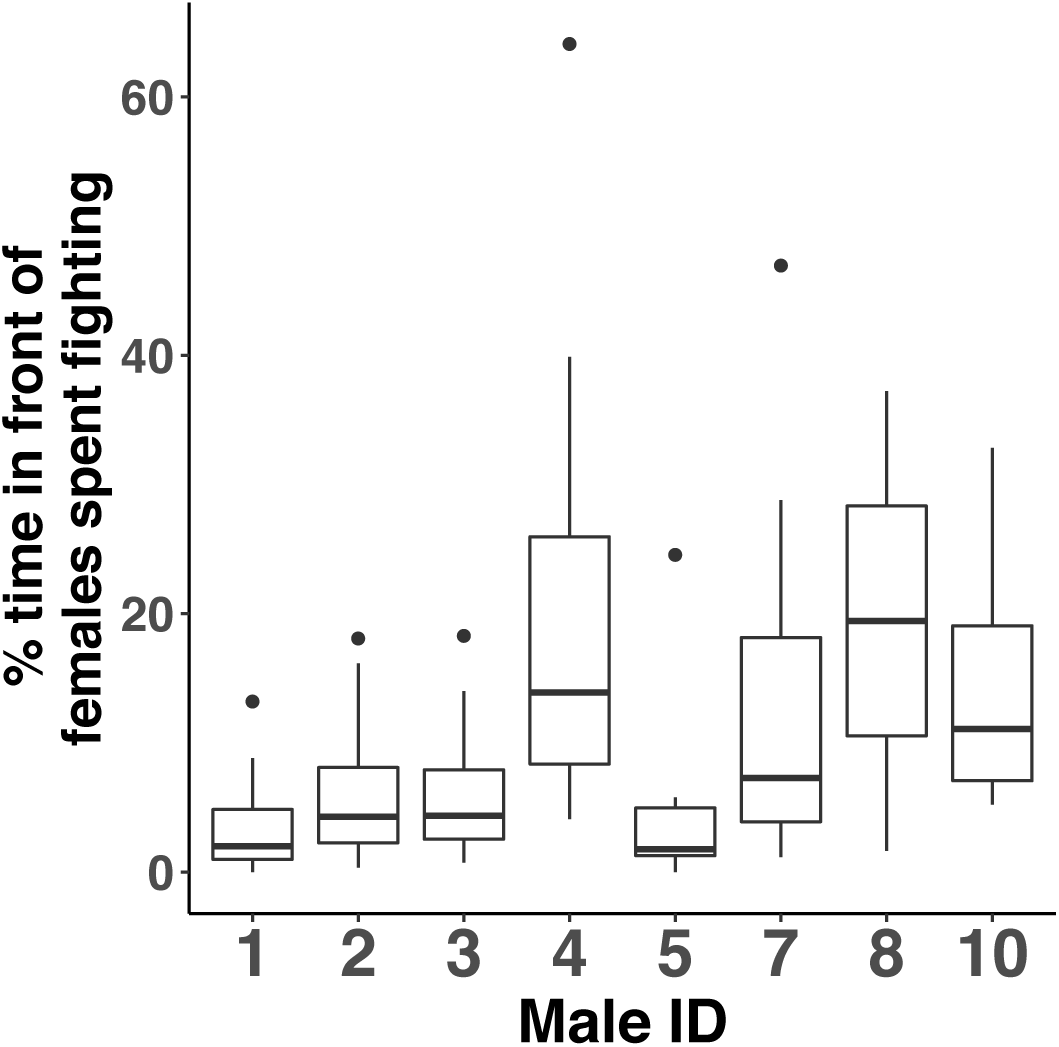
Summary of the percent of time territory-holding males spent fighting in front of females. Each data point underlying the boxplots is the amount of time a male spent fighting in front of females on a particular day divided by the total amount of time that he spent in attendance on the lek while females were also present. Male 12 was excluded because he was only considered a territory-holder for one day.

**Figure S3.**
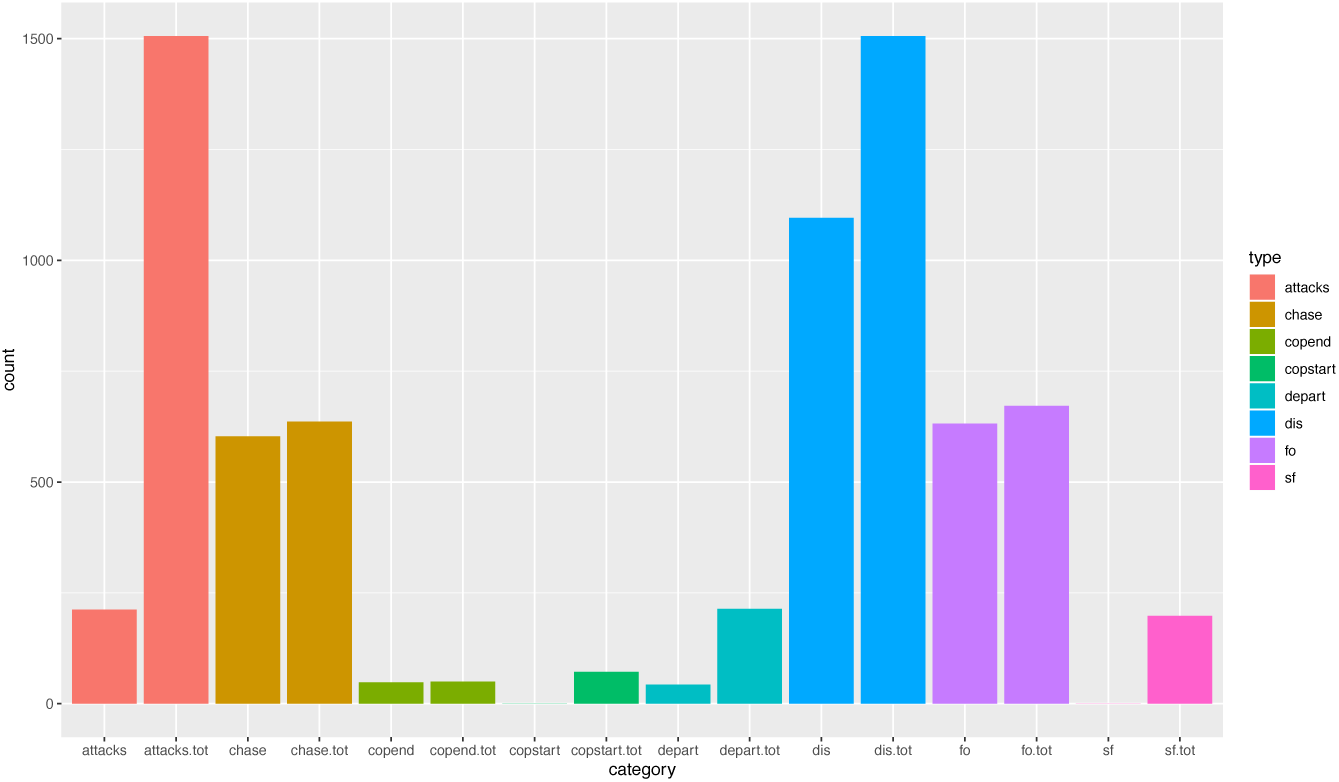
REM performance at predicting the exact next event to occur. Each color is a type of event: attacks, Chase response events, copulation ends, copulations starts, departures, disengagements, Face-Off (FO) response events, and Short Fight (SF) response event, respectively. Within each color, the left bar is the number of realized events that the Differential Attractiveness REM predicted perfectly (ranked them as the number one most likely event to occur next). The right bar is the total number of that type of event in the dataset, for reference.

**Figure S4.**
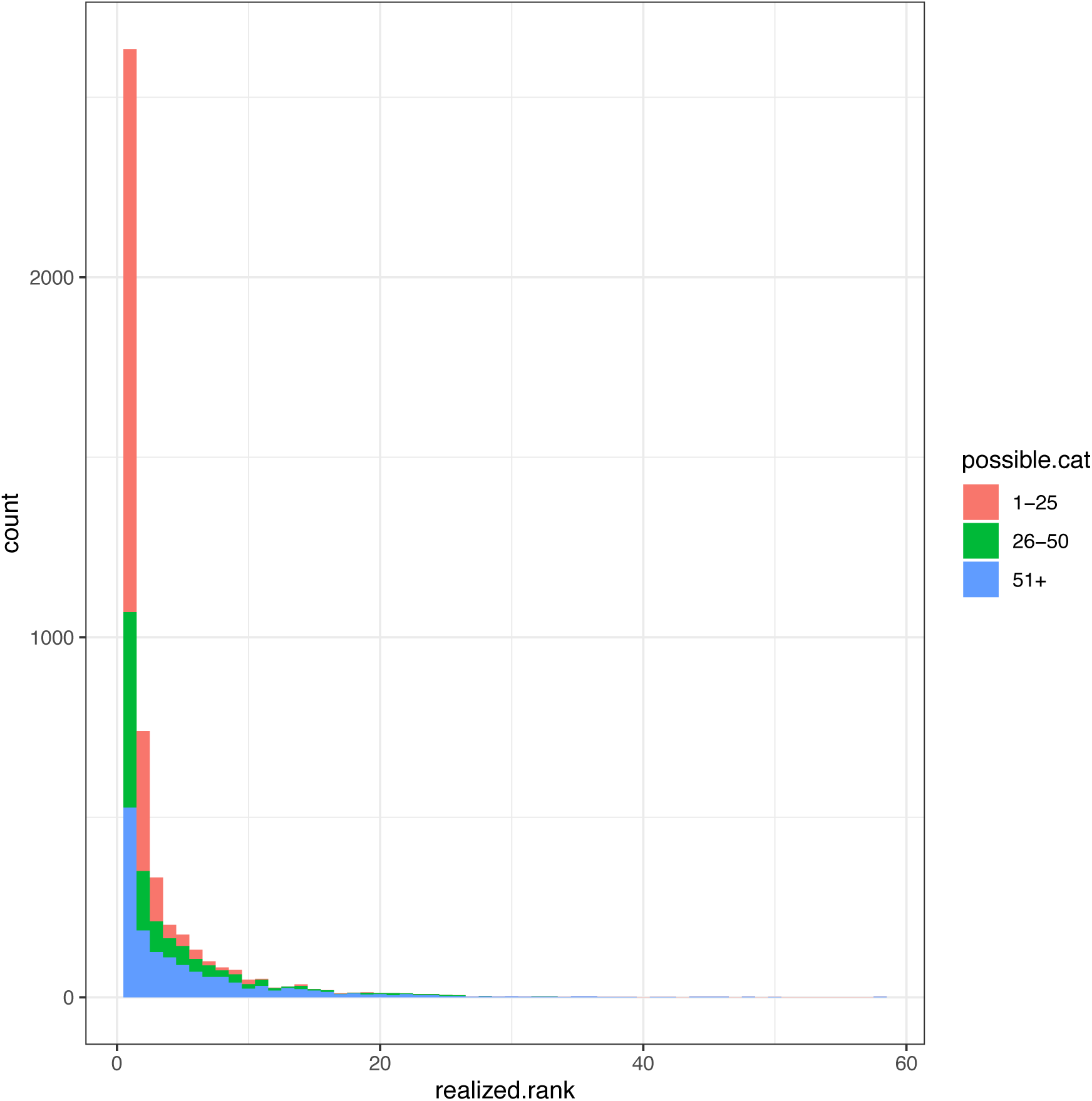
Frequency histogram of the ranking of each realized event by the Differential Attractiveness REM. A low-numbered rank means the model considered the actual observed event in the dataset to be very likely. A high-numbered rank indicated the model was relatively “surprised” by that particular event. Colors represent ranges for the total number of events that were possible at the time of each event.

**Figure S5.**
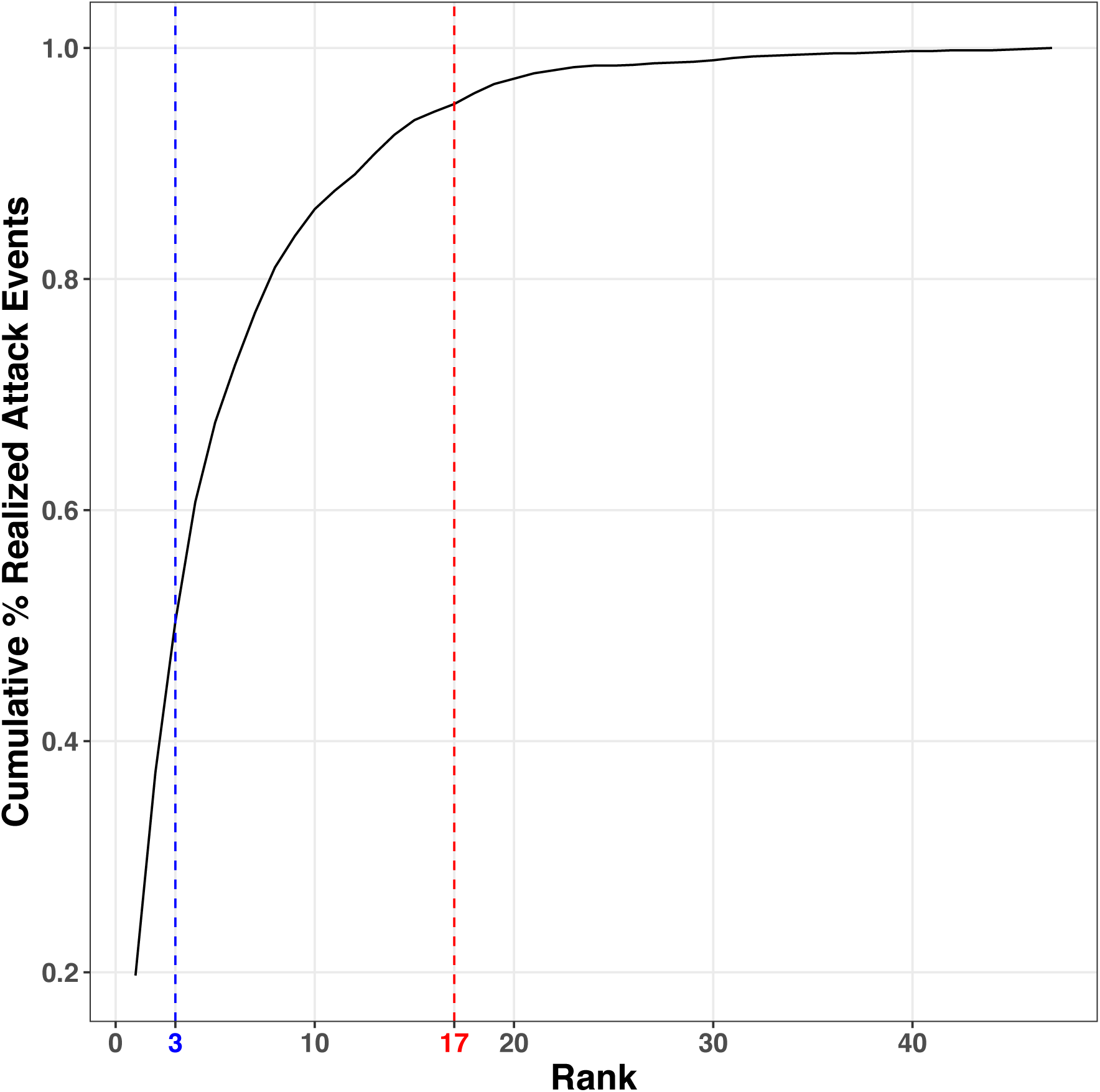
Recall plot for attack events. The black line represents the cumulative percentage of realized events in the dataset that the Differential Attractiveness REM predicted to be a particular rank or lower. The blue line at rank three shows that 50% of realized attack events are ranked by the model within the top three most likely to have occurred. The red line at rank seventeen shows that 95% of realized attack events are ranked by the model within the top seventeen most likely to have occurred.

**Figure S6.**
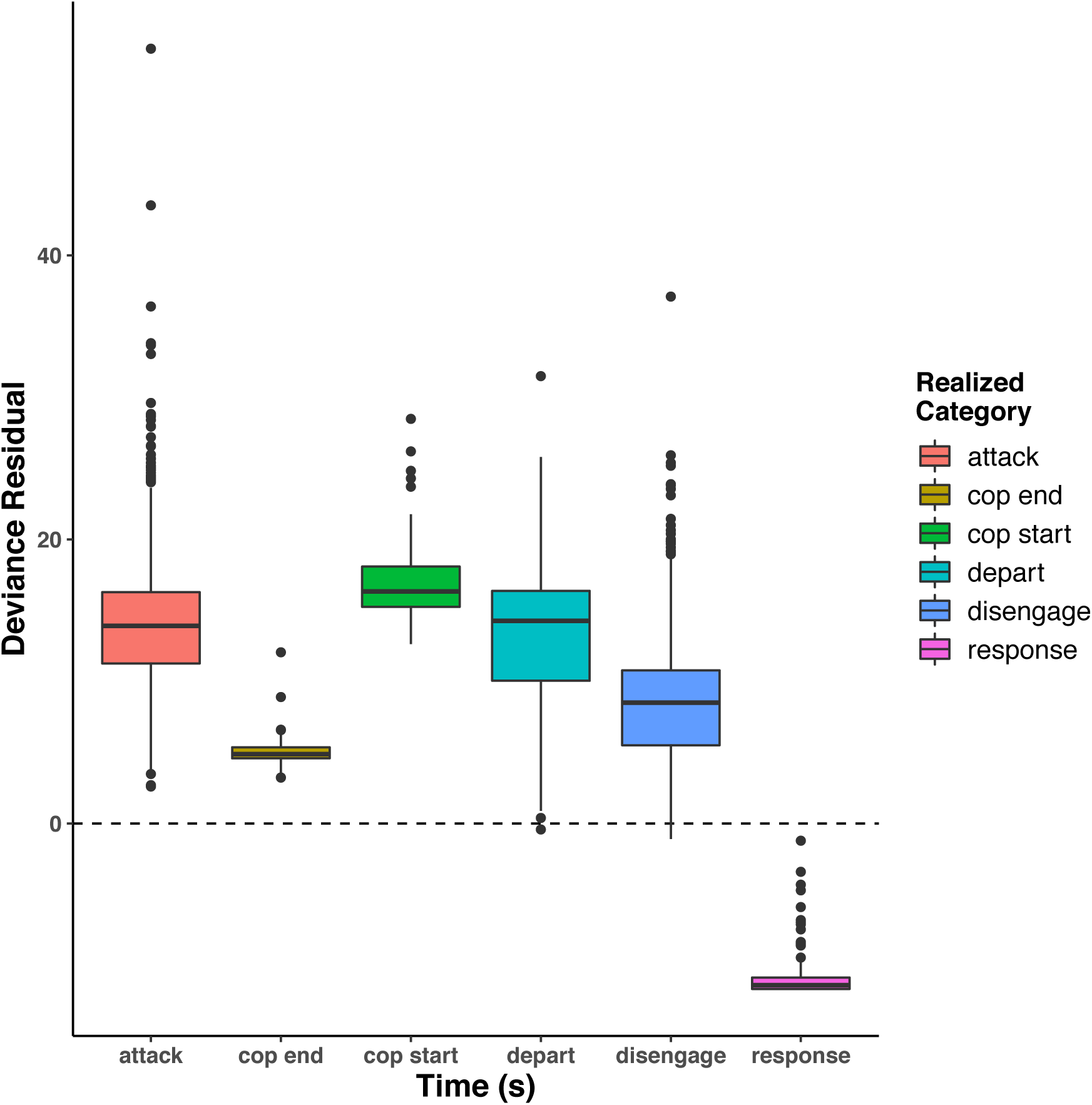
Residual deviance of the Differential Attractiveness REM by category of realized event. Smaller values indicate better event-specific model fit.

**Figure S7.**
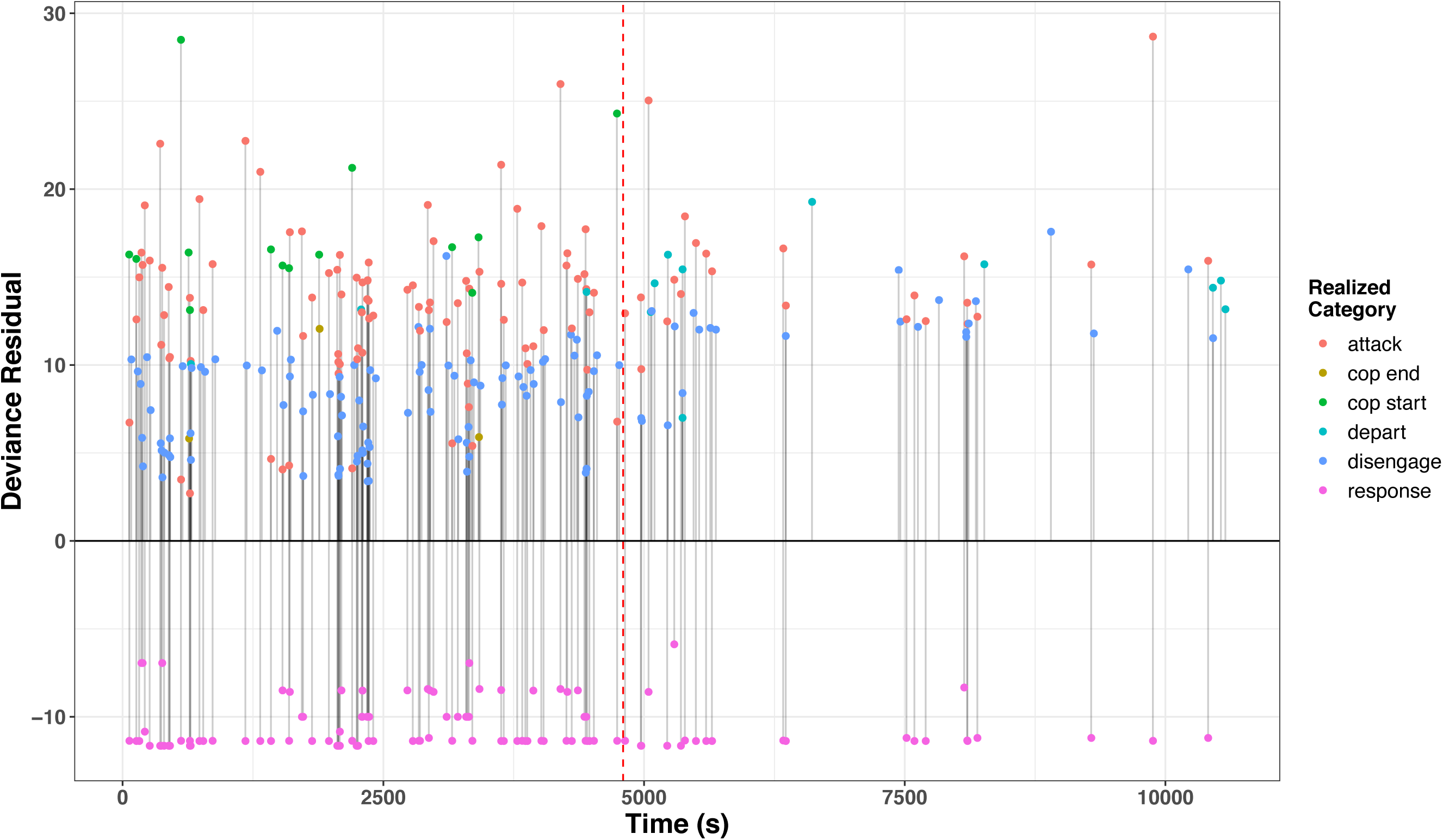
Example of deviance residuals of the Differential Attractiveness REM for every event through time for April 1, 2014. The vertical red dashed line marks the time when the last female departed from the lek.

**Figure S8.**
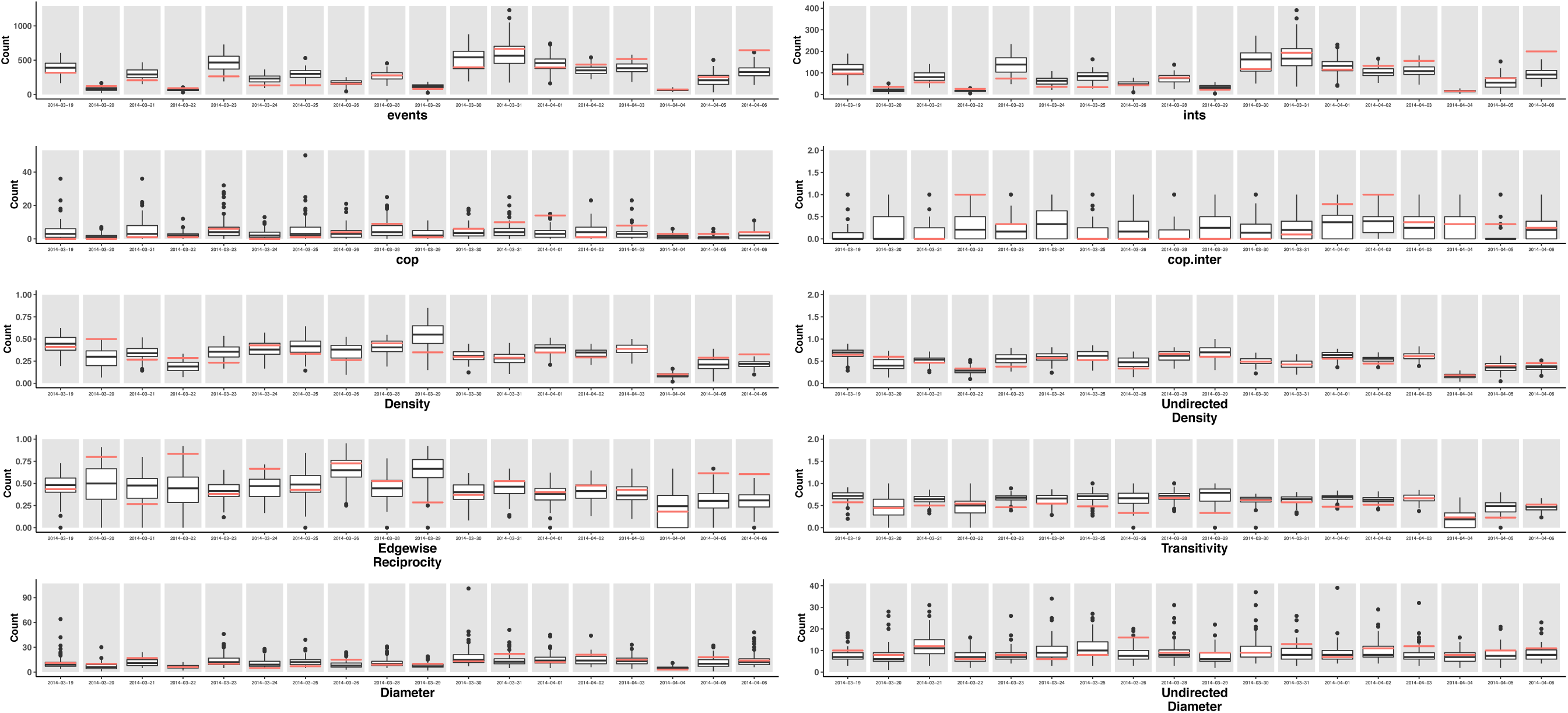
Summary of adequacy metrics for simulations generated from the fitted coefficients of the Differential Attractiveness REM for each of the eighteen days in the observation period. Red horizontal bars indicate the observed values for each metric on each day. Each boxplot represents the range of values for a particular metric across one hundred simulations for that particular day. “Events” is the total number of events, “ints” is the total number of interactions, “cop” is the total number of copulations, “cop.inter” is the proportion of copulations that were interrupted. The other panels are standard network metrics based on the network of interactions as ties, either directed or undirected, for that day. Directed ties go from attacker to receiver.

**Figure S9.**
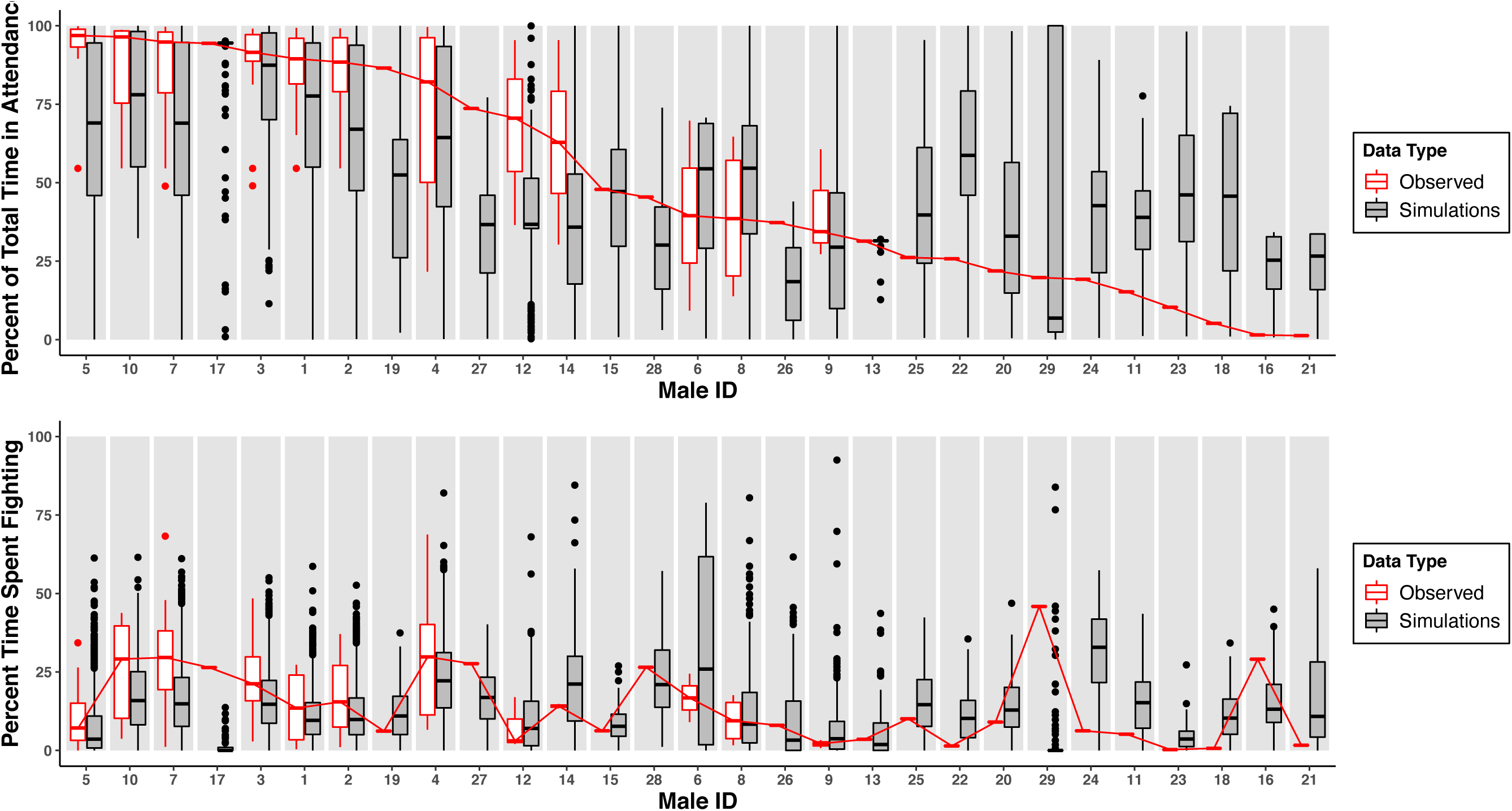
Observed (red) and simulated (black/grey) distributions of percent of time spent in attendance on the lek and percent of time spent fighting for each male across all days in the observation period. Simulated data is one hundred runs for each day generated from the fitted coefficients of the best-fitting Differential Attractiveness REM.

**Figure S10.**
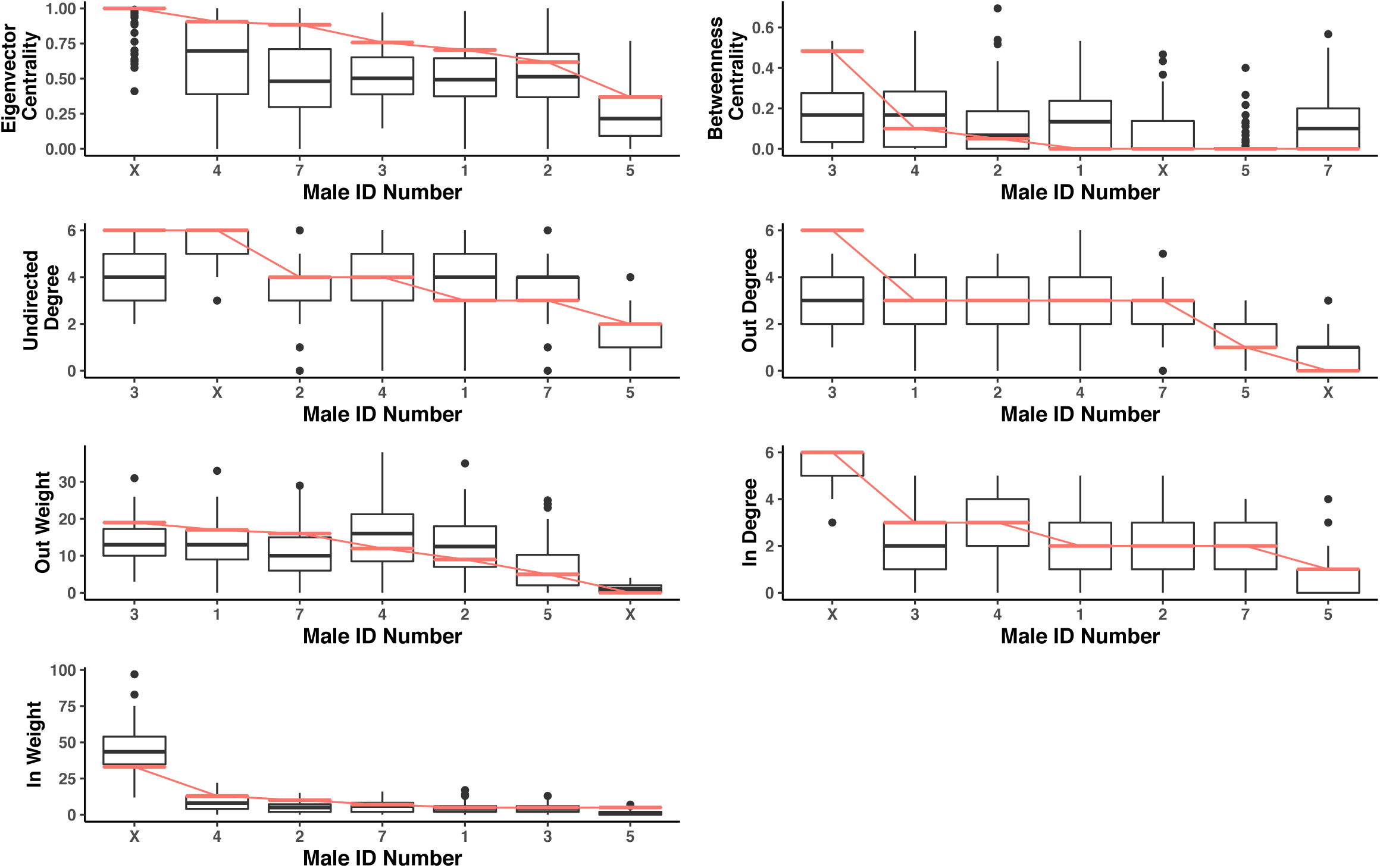
Example of observed (red) and simulated (black) distributions of node-level network metrics from day nine of eighteen in the observation period: March 28, 2014. Network nodes are each male present that day and ties are interactions; directed ties go from attacker to receiver. Simulated data is one hundred runs generated from the fitted coefficients of the best-fitting Differential Attractiveness REM.

**Figure S11.**
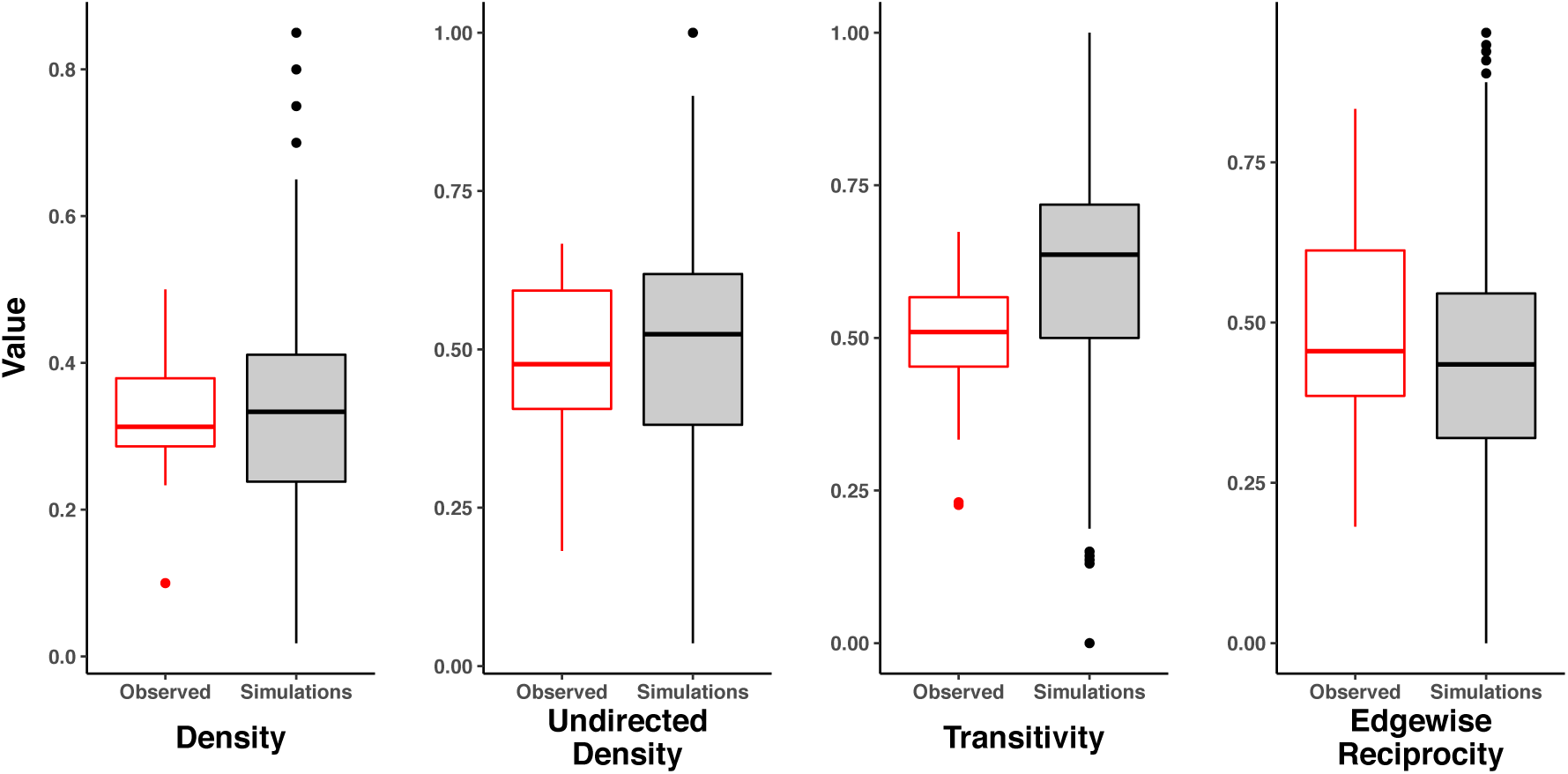
Observed (red) and simulated (black/grey) distributions for network-level metrics pooled across all eighteen observed days and across all eighteen hundred simulations, respectively.

**Table S1.**
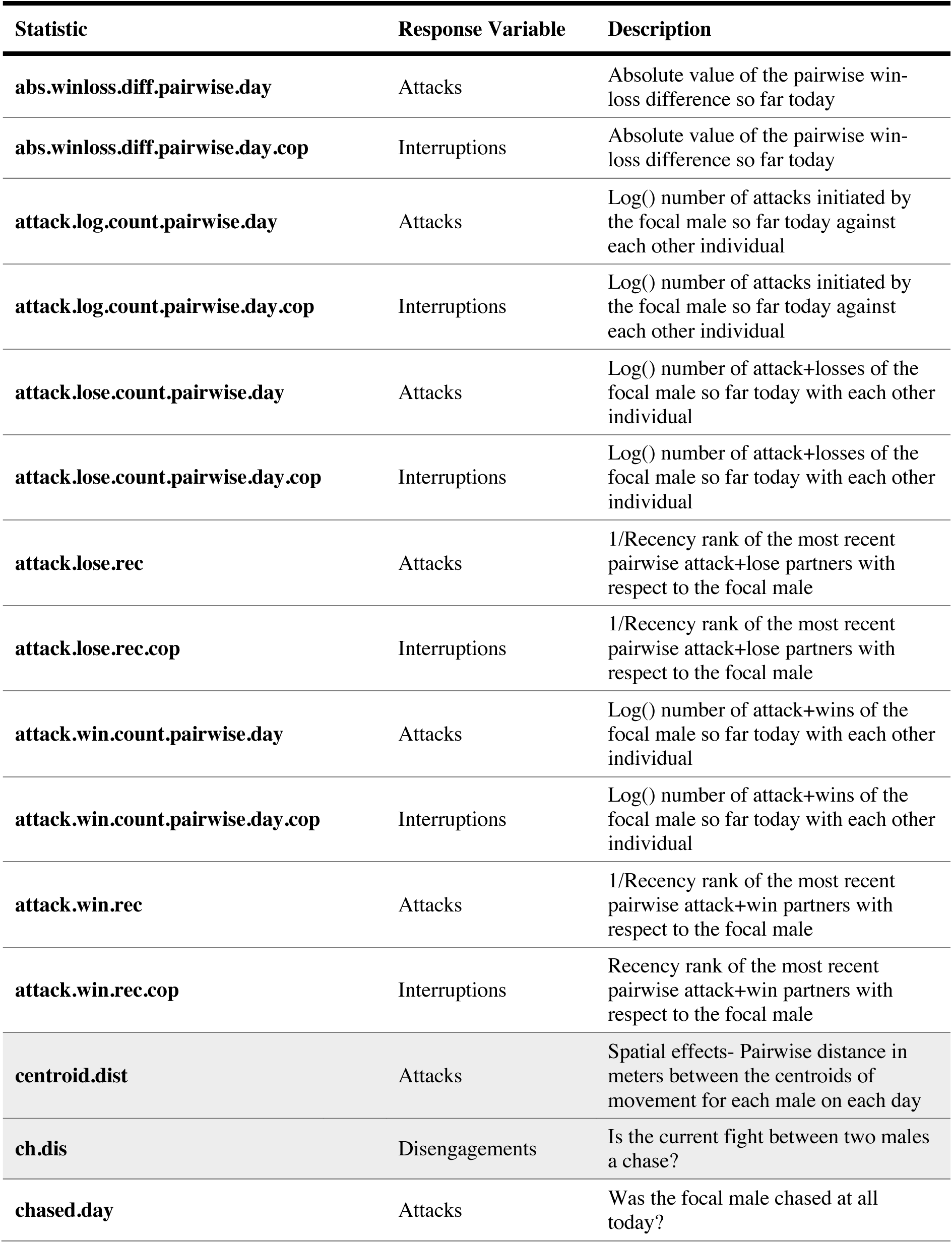
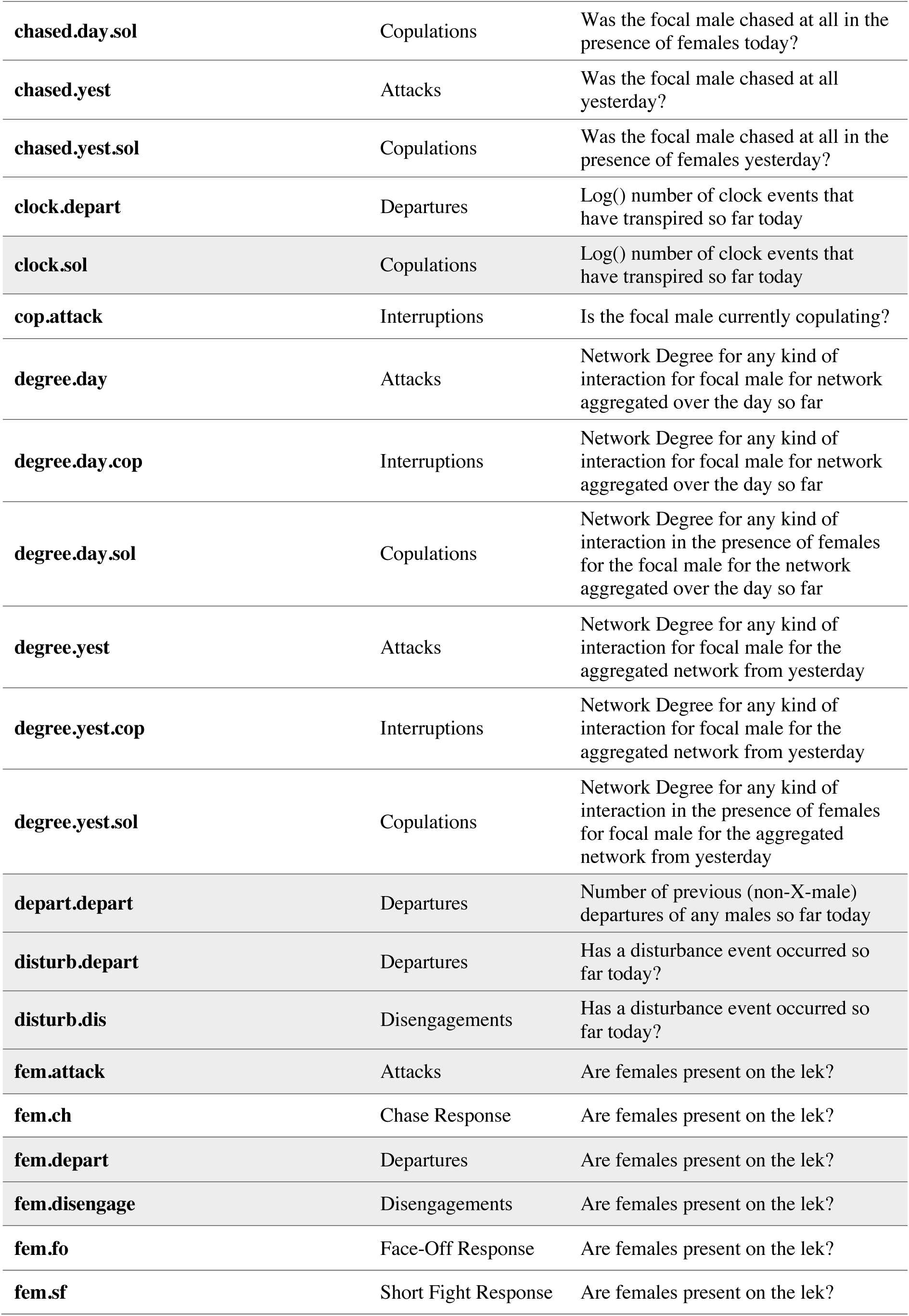
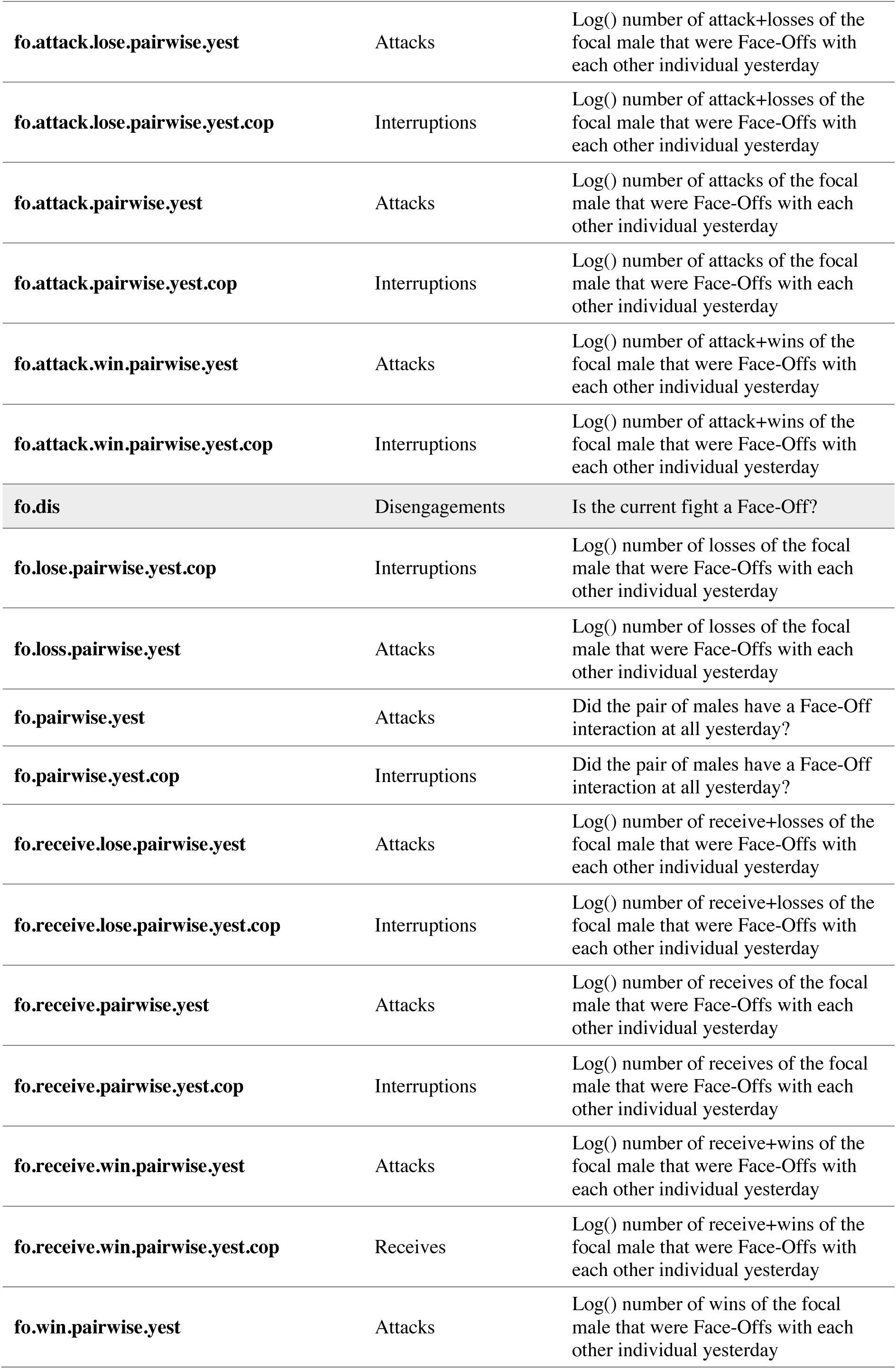
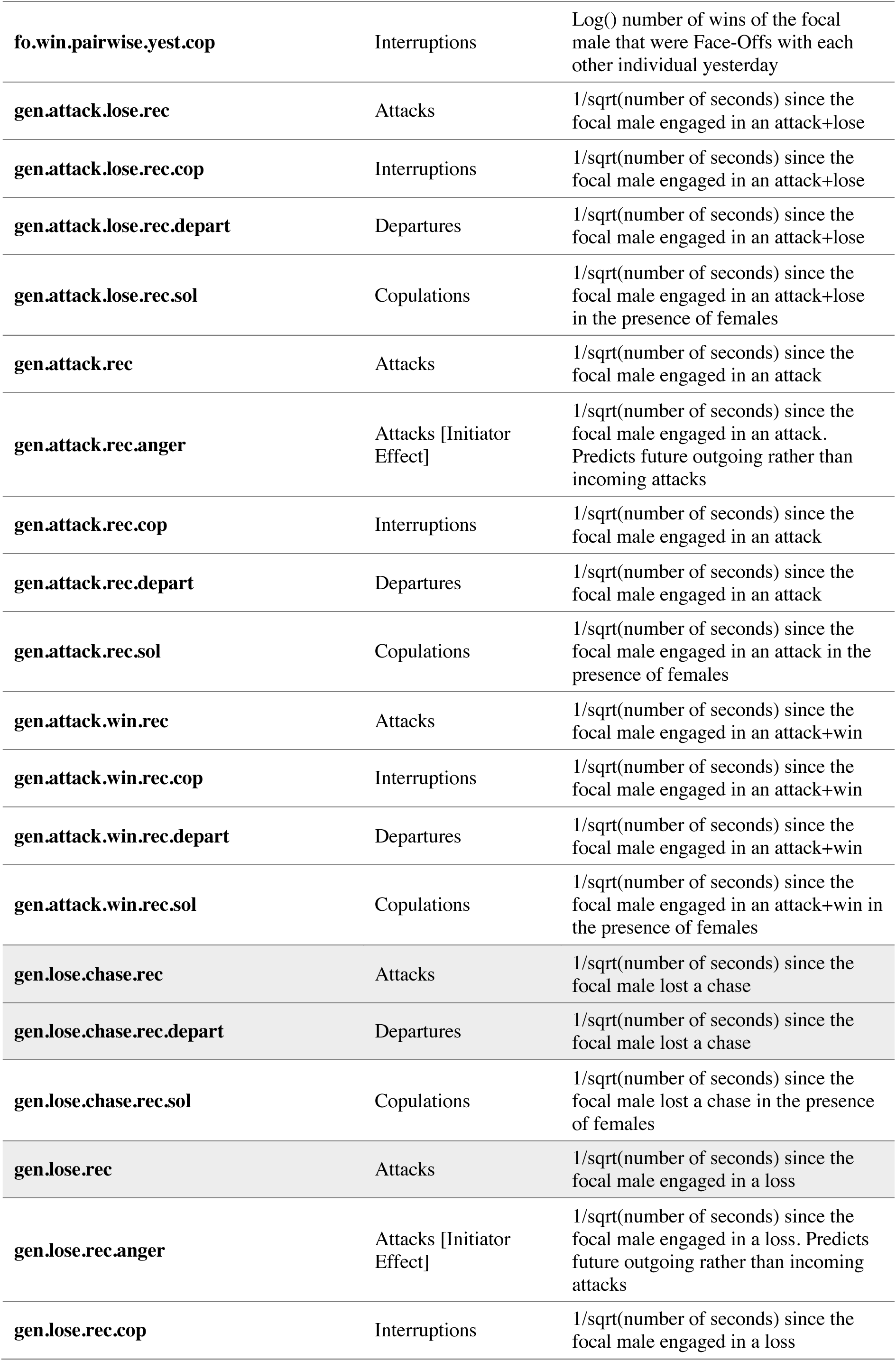
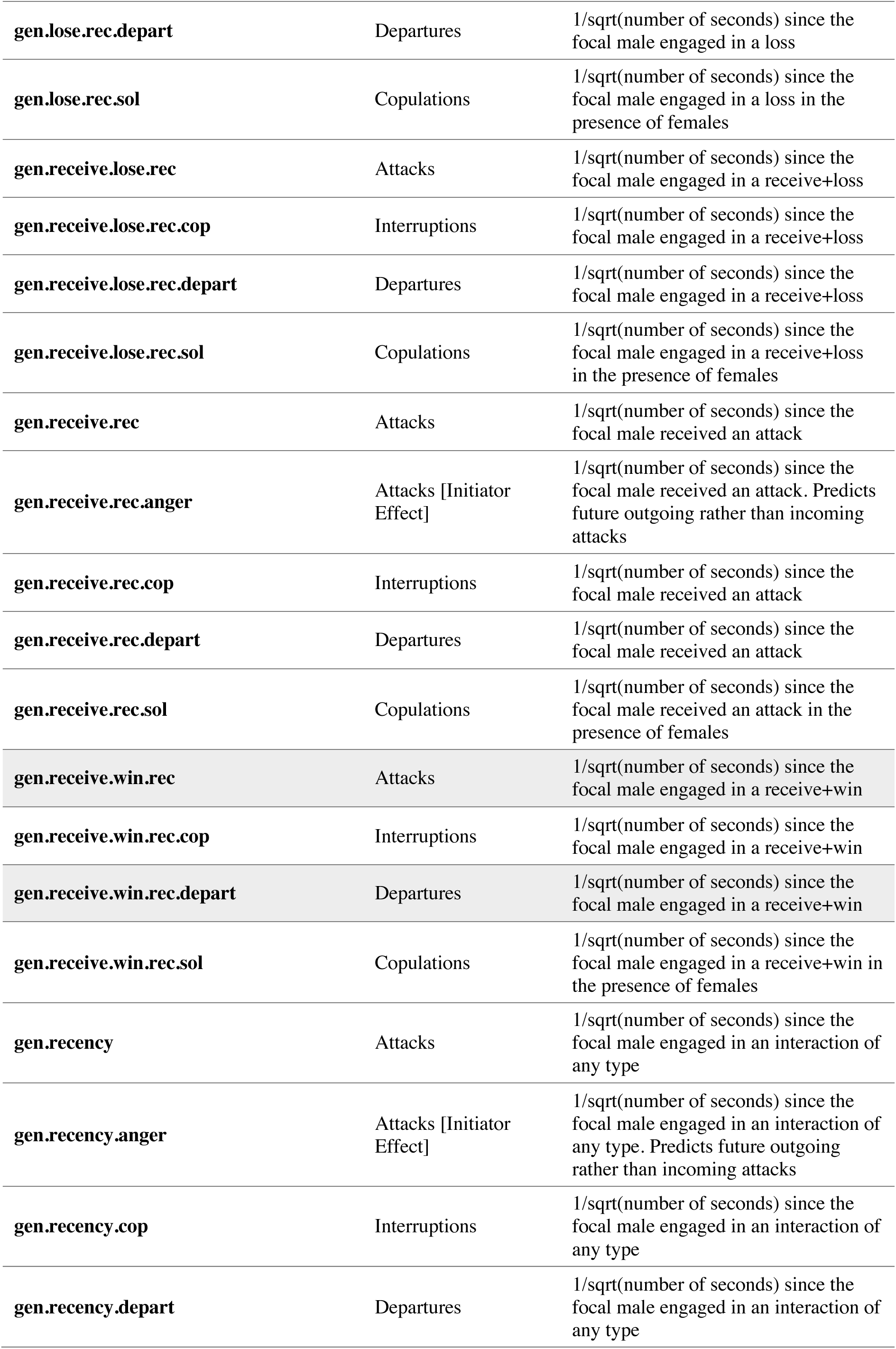
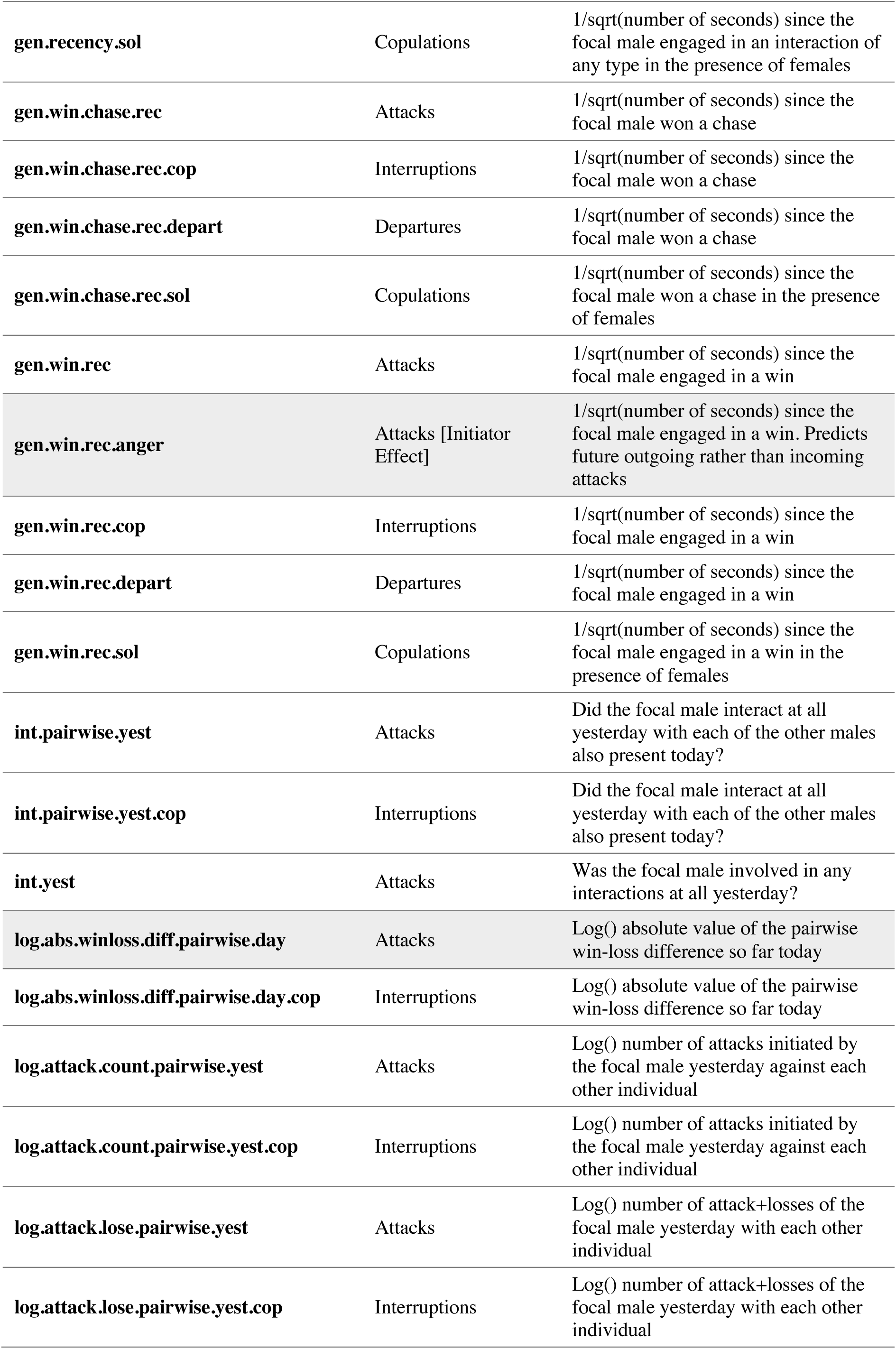
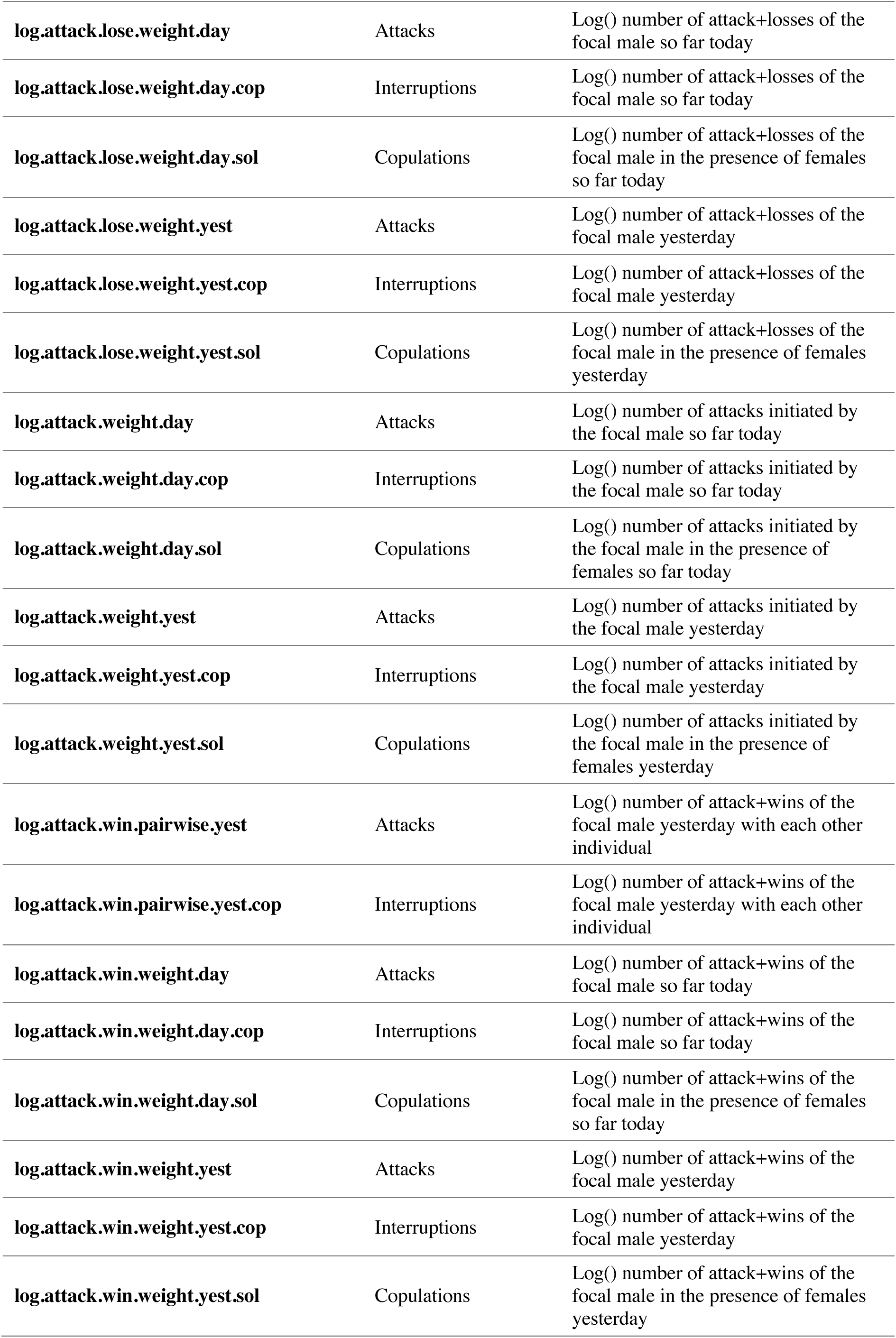
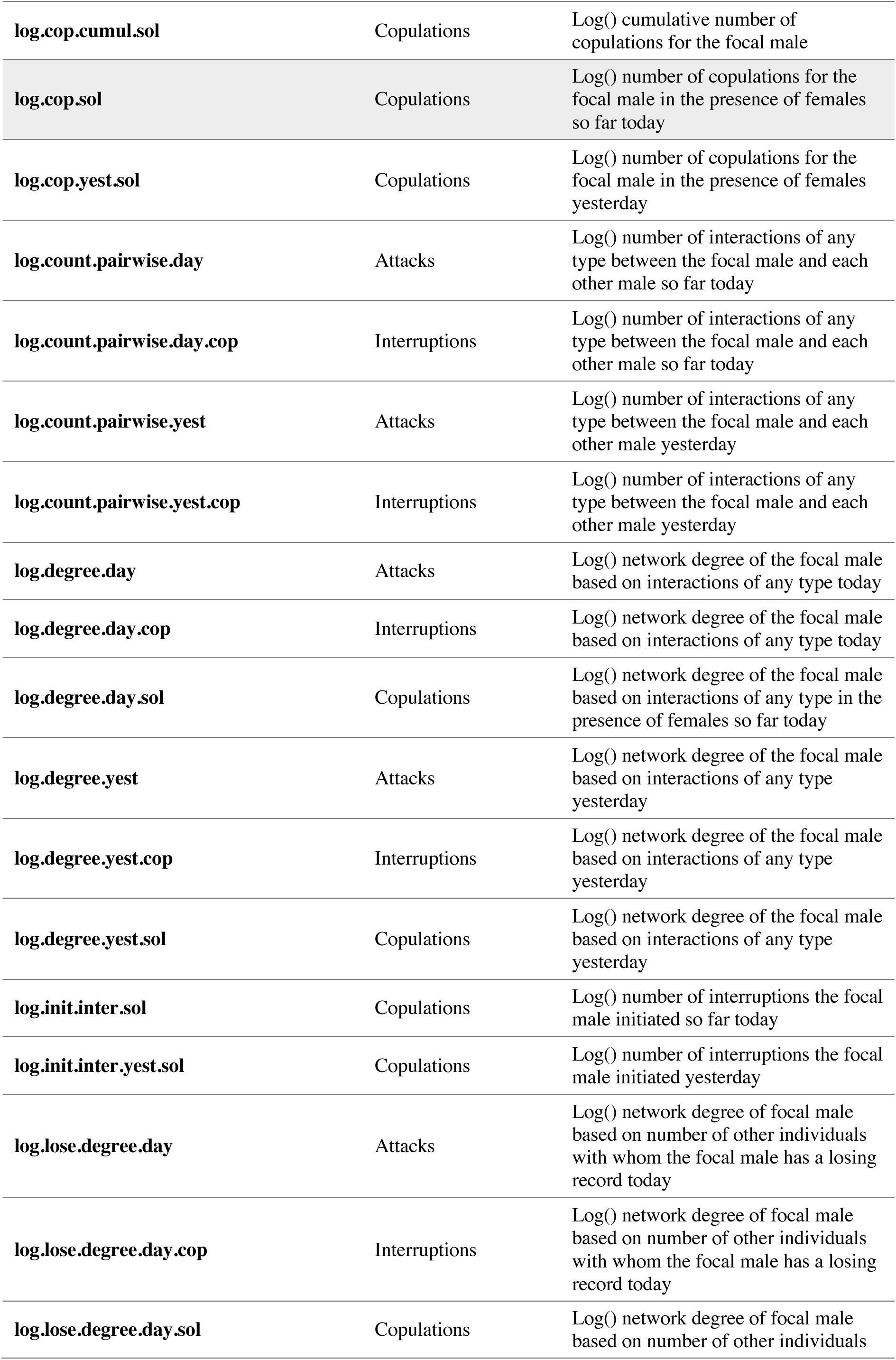
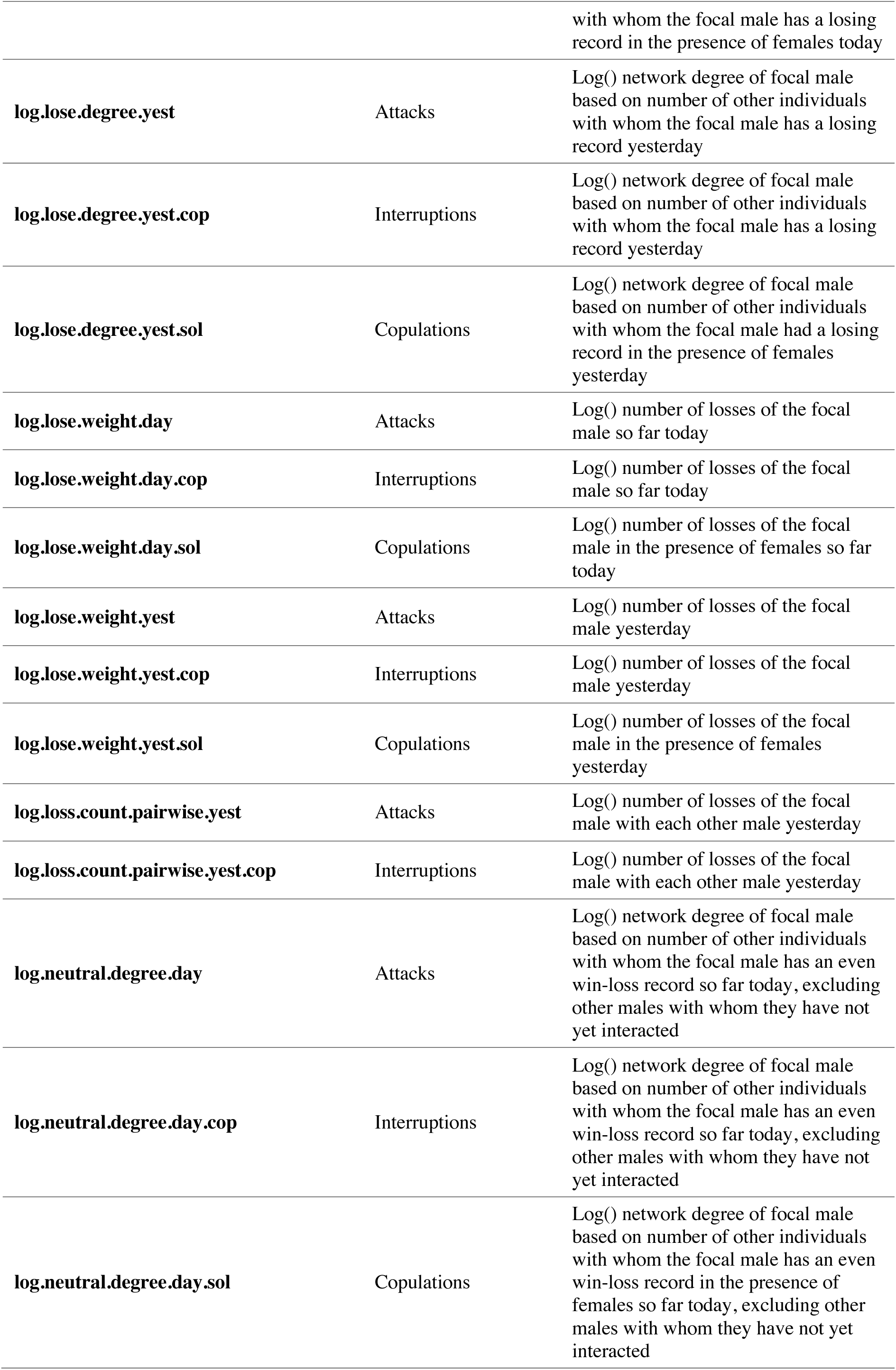
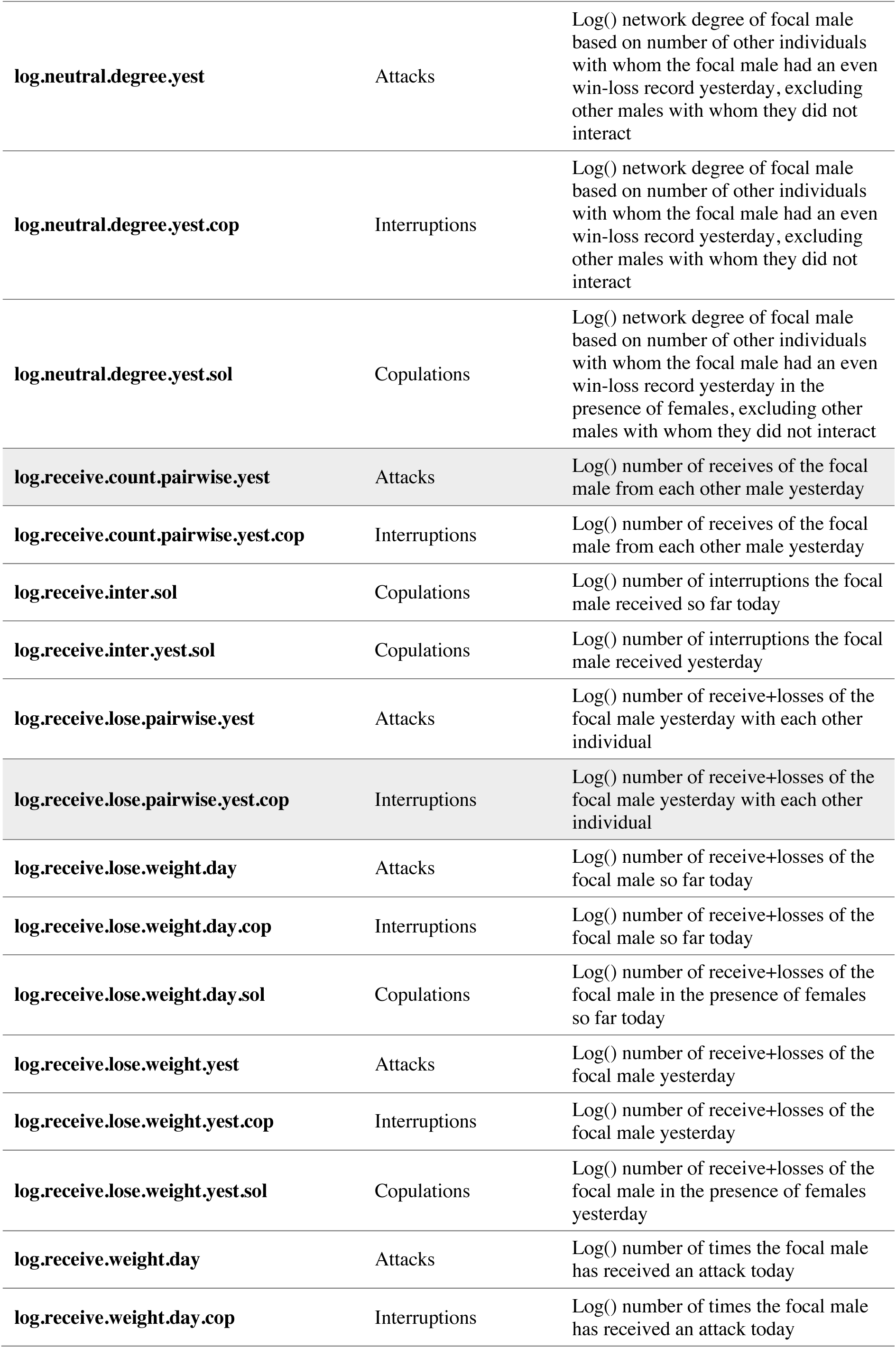
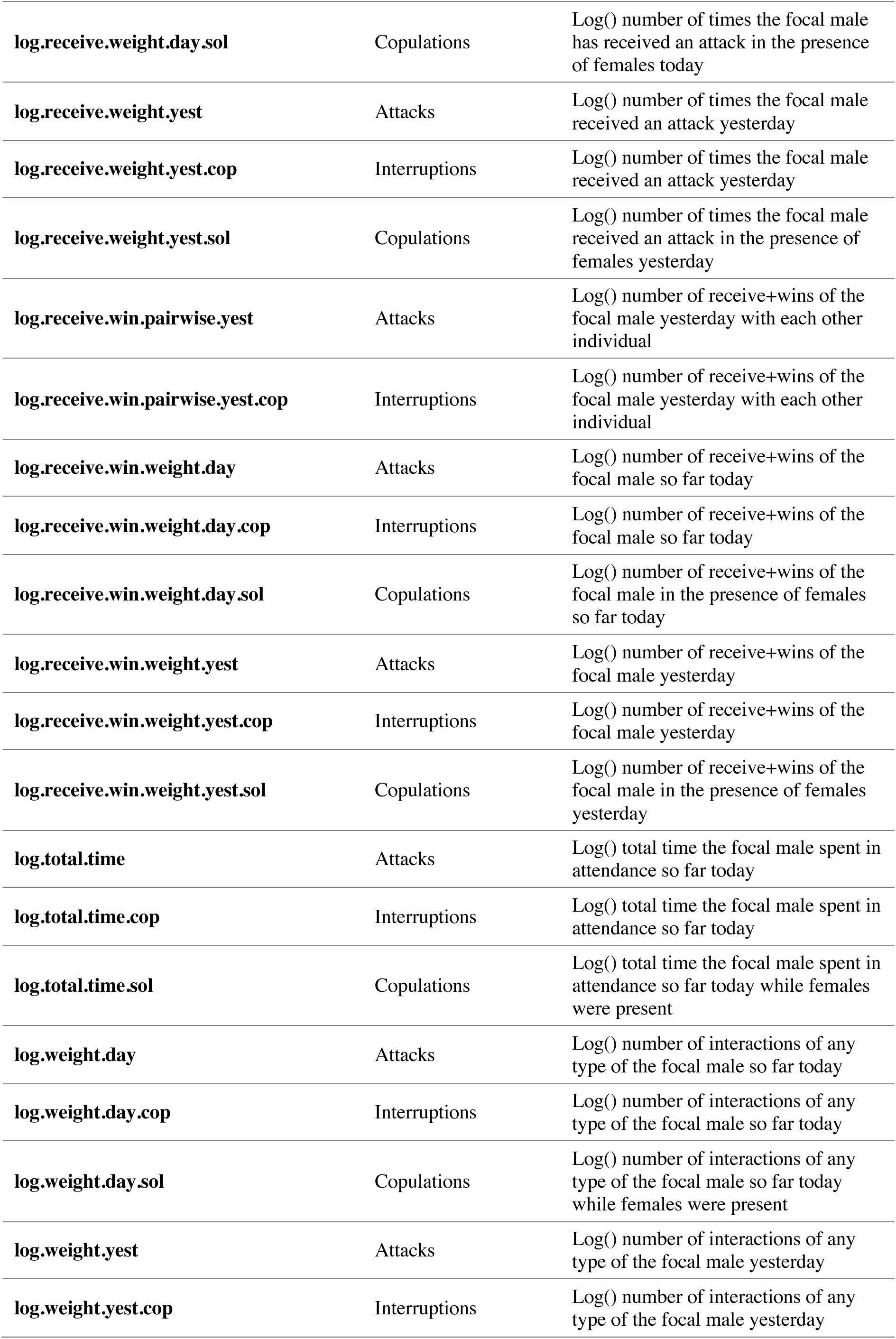
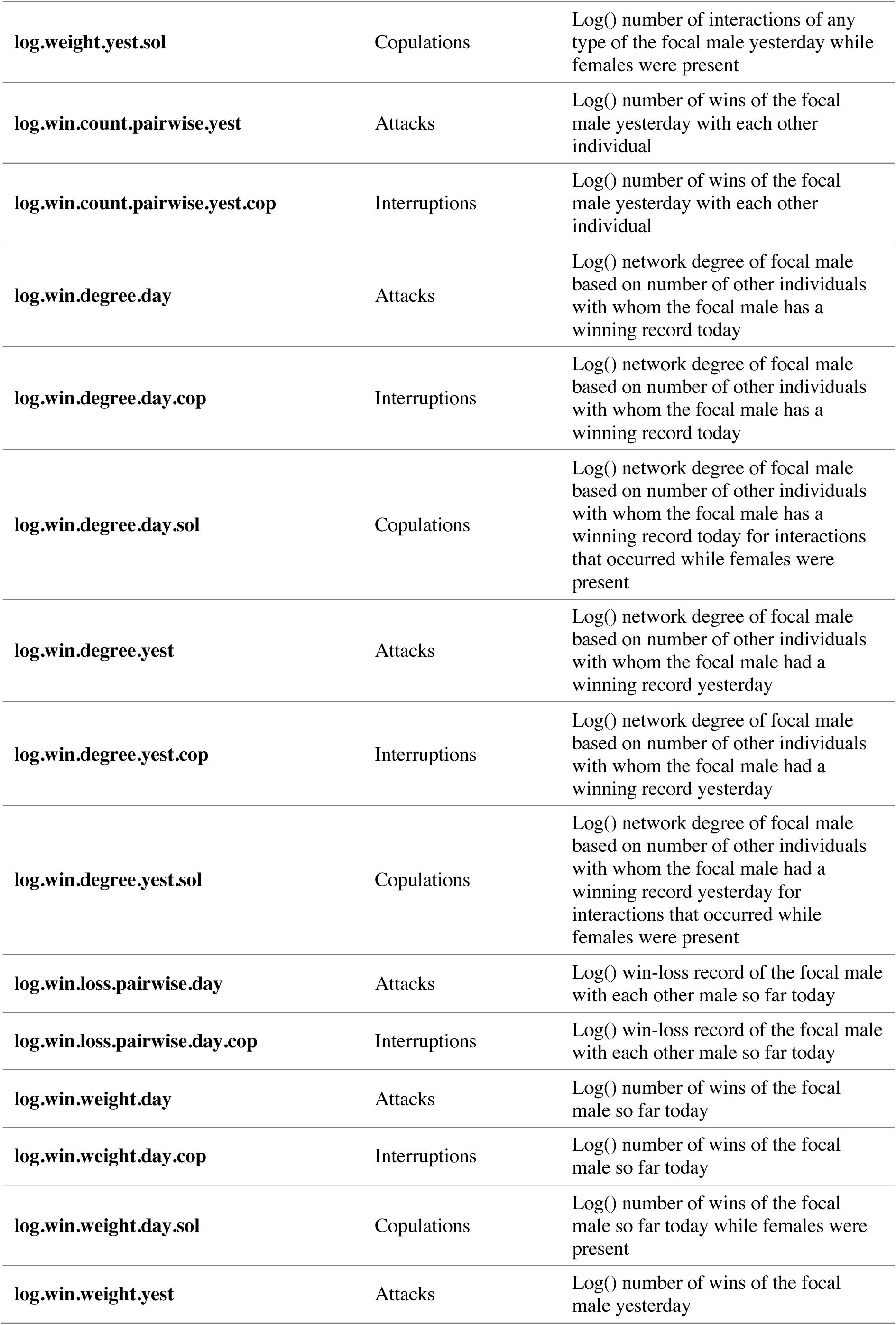
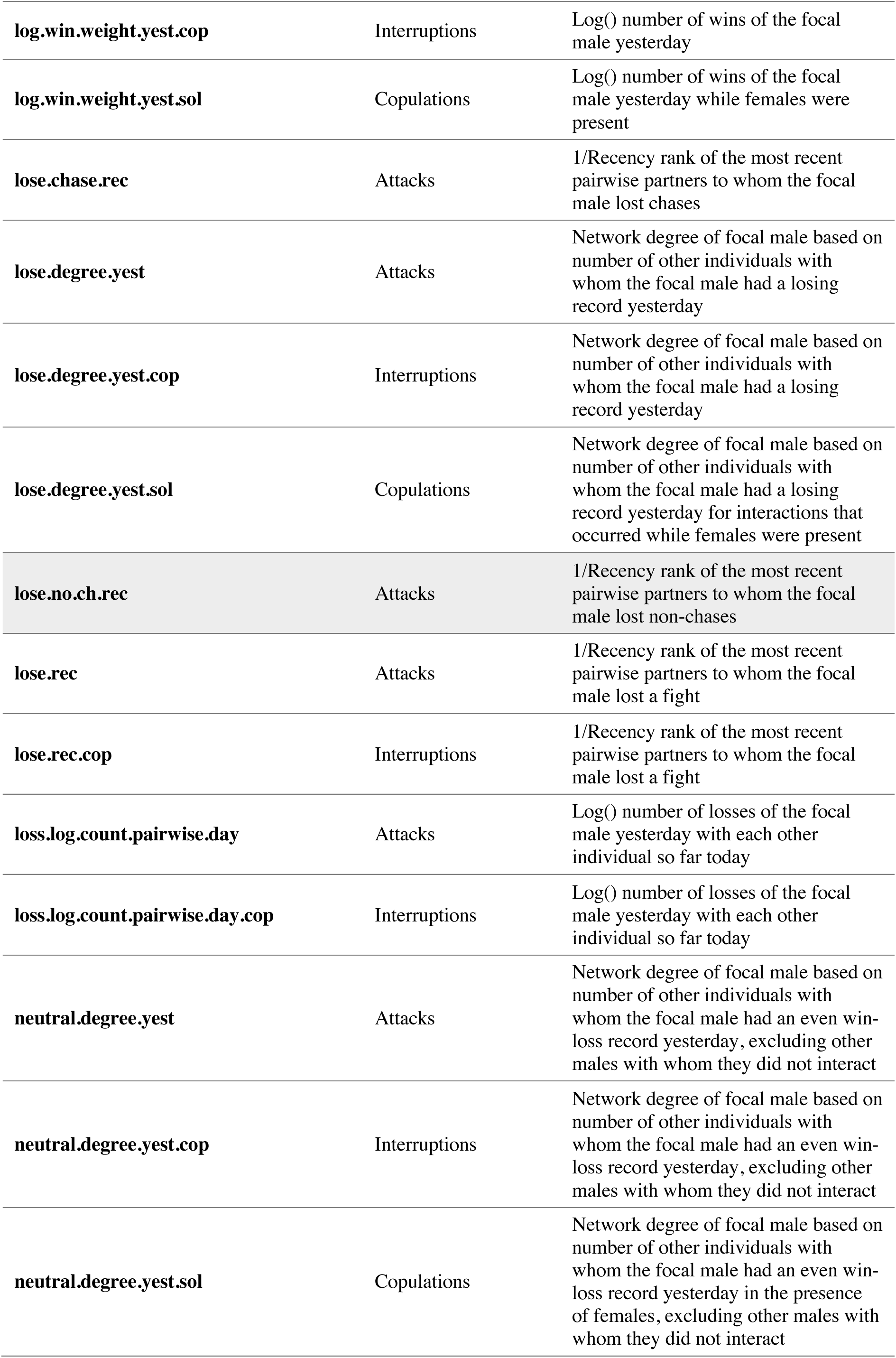
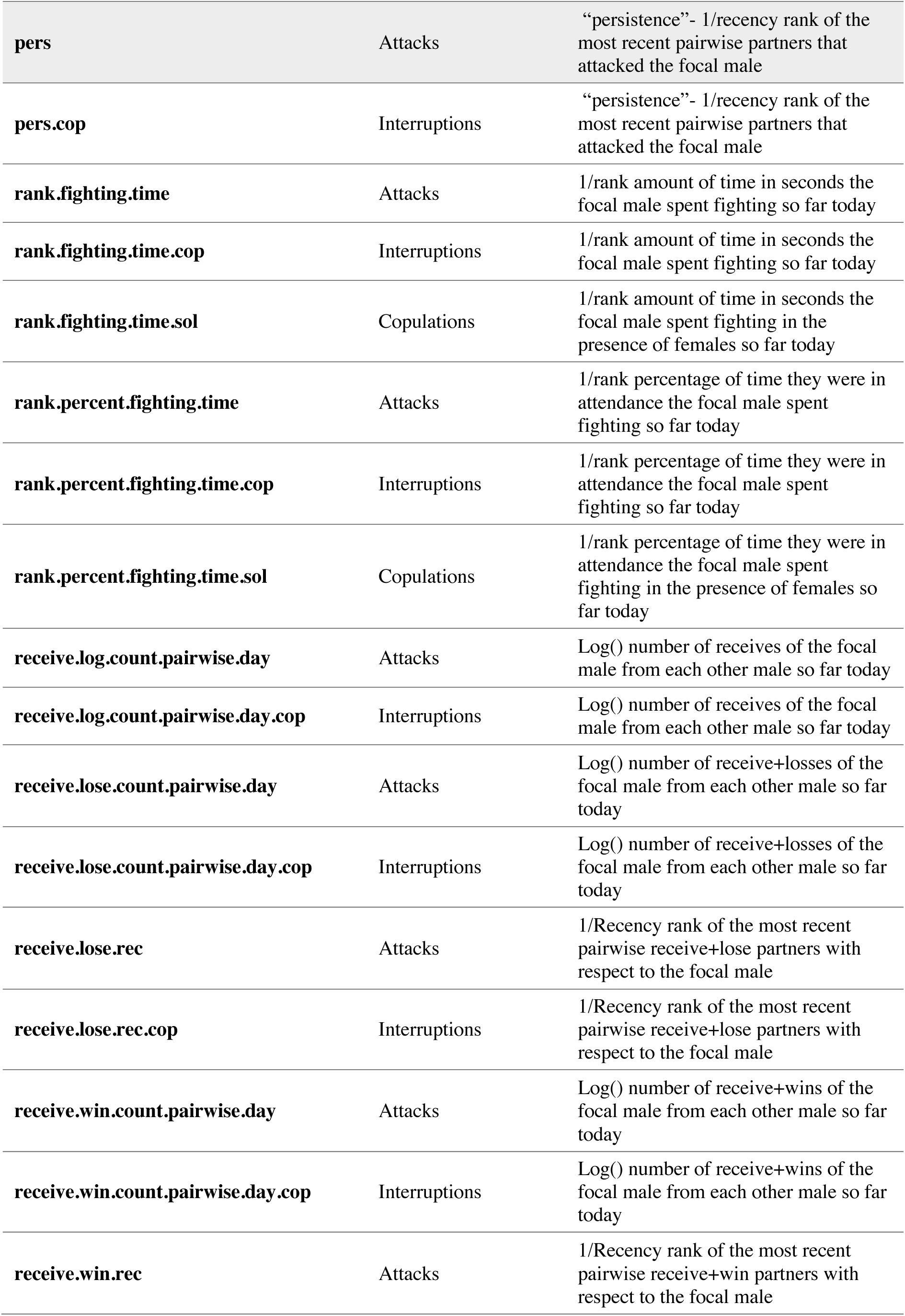
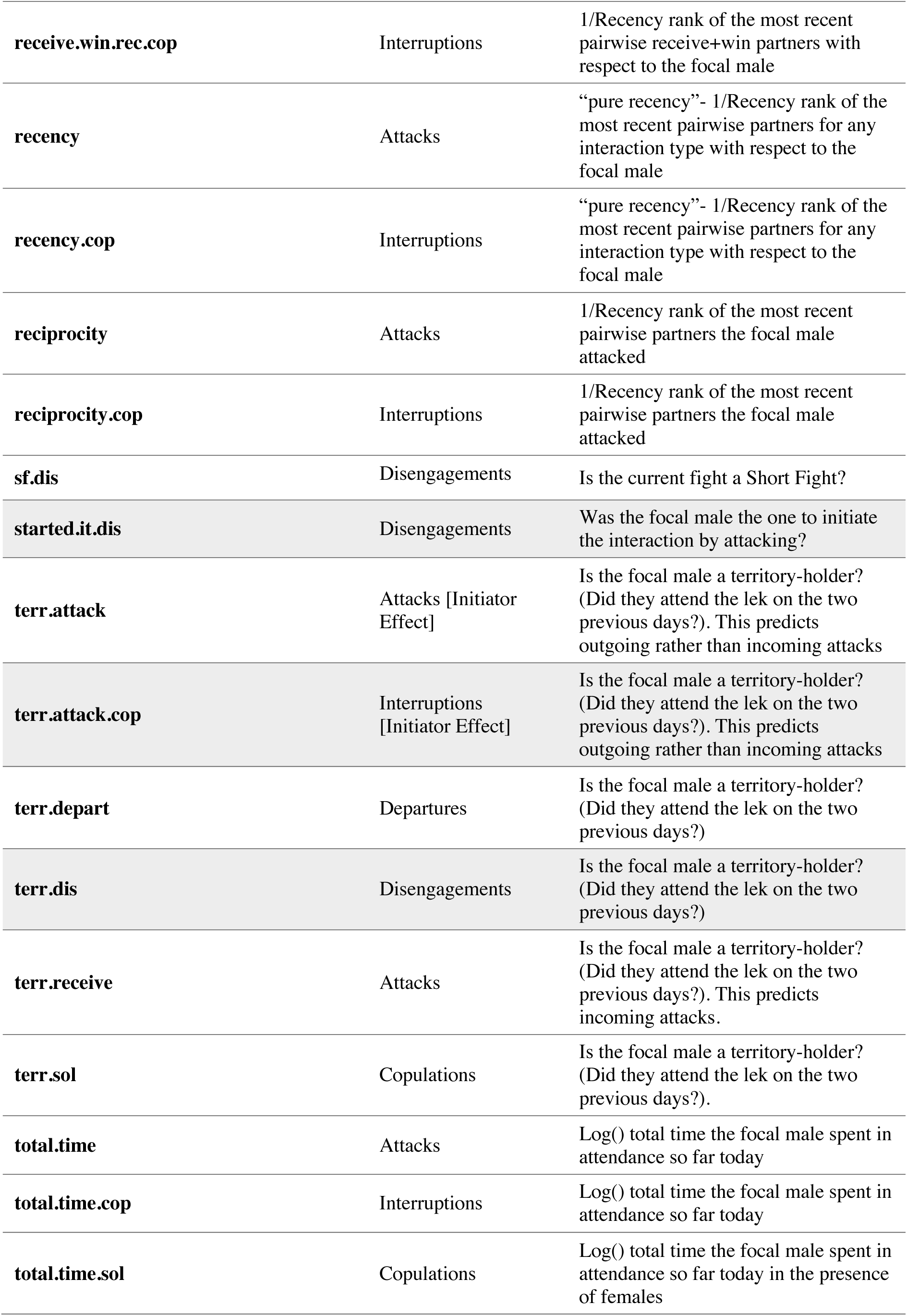
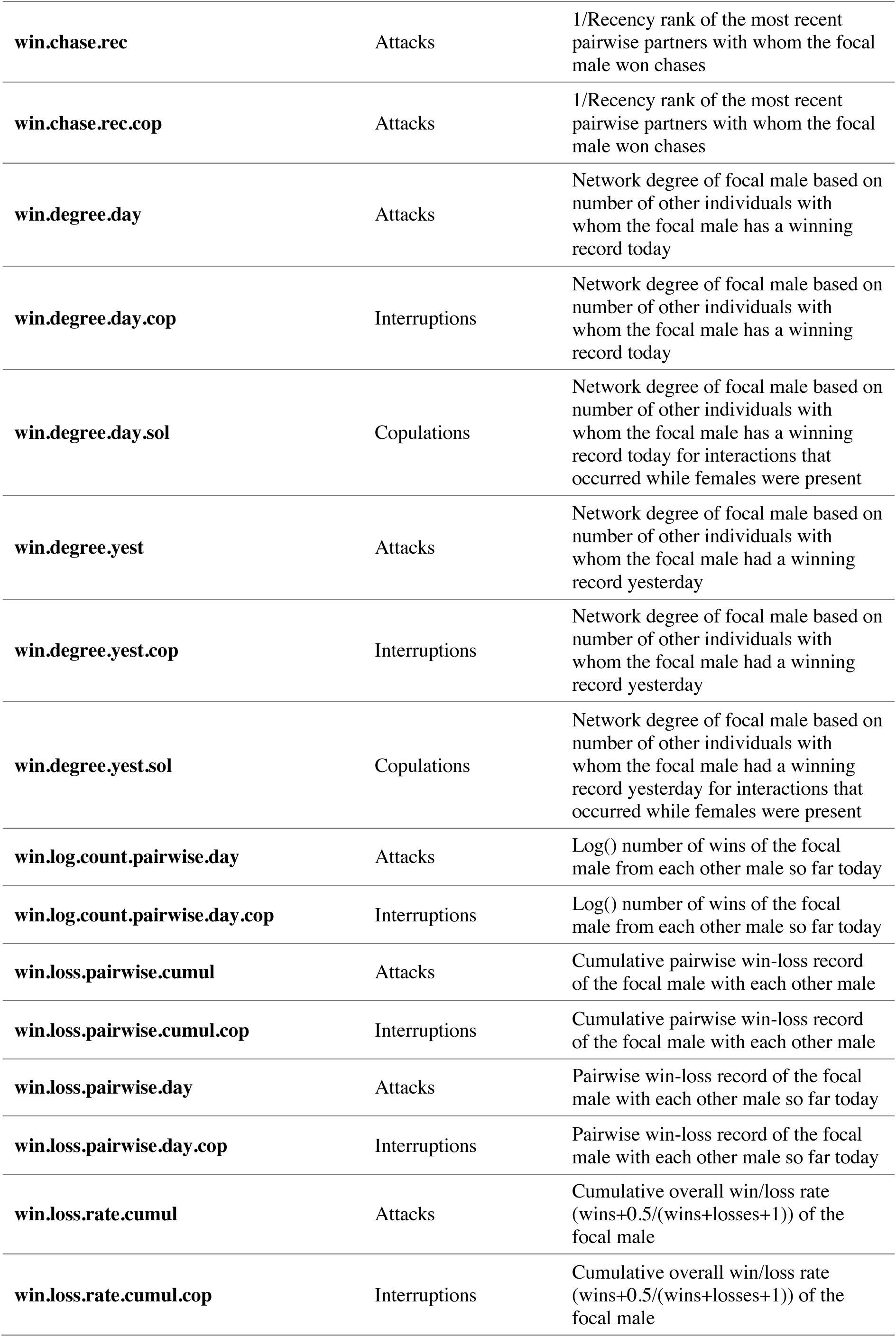
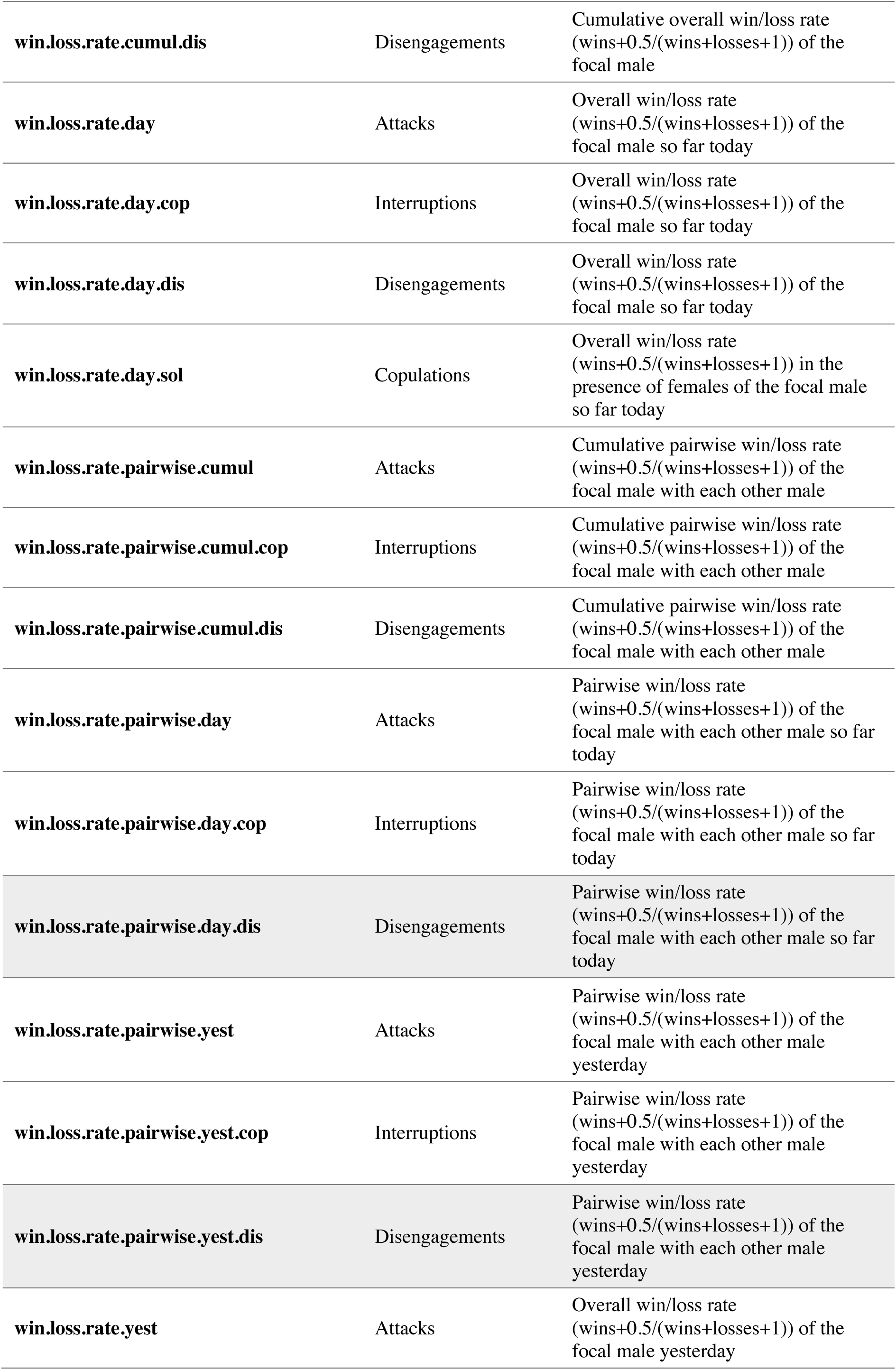
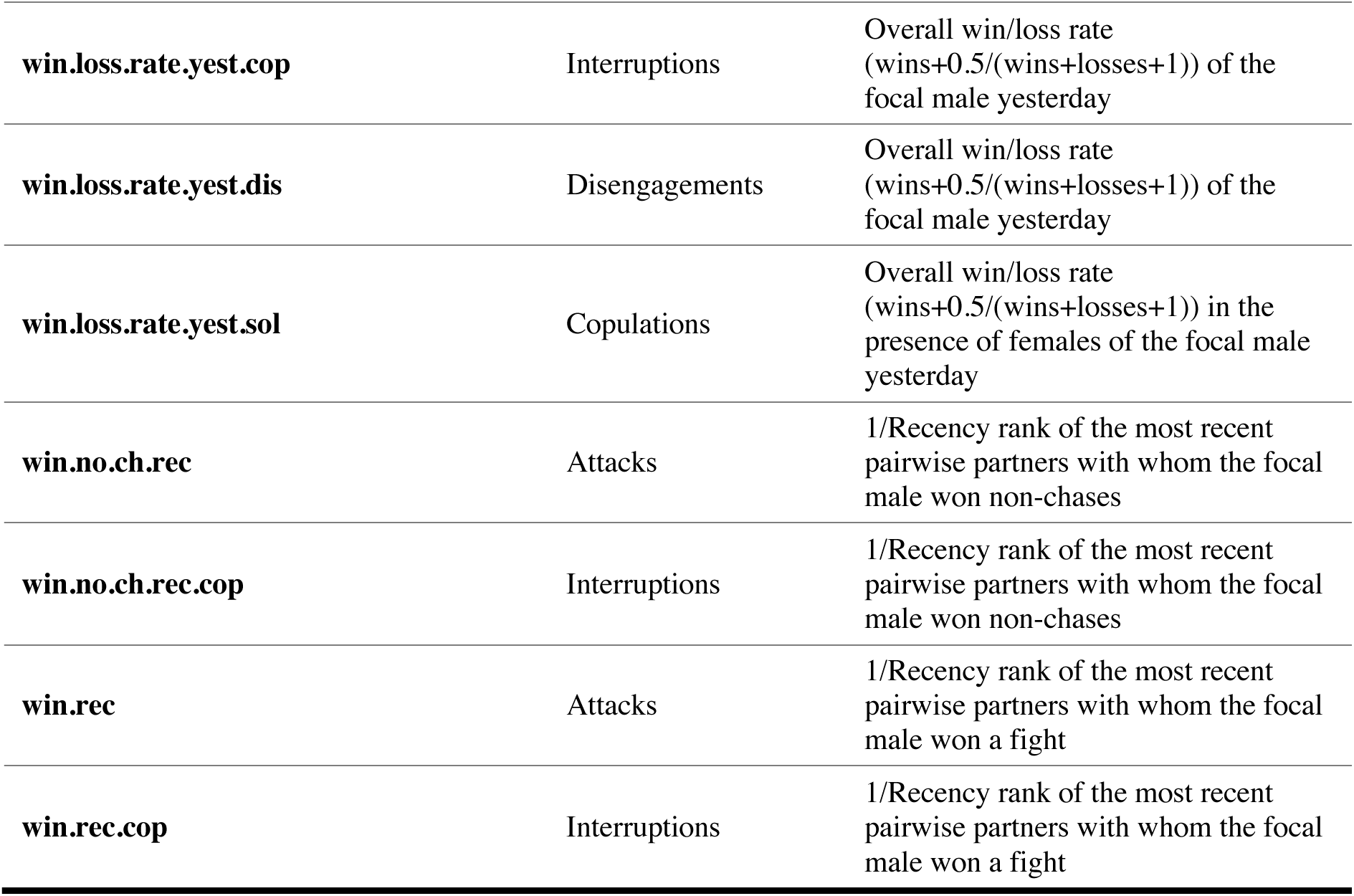
Table of full set of sufficient statistics tested in the Relational Event Model with descriptions. Shaded rows represent statistics that were significant effects in the best-fitting Differential Attractiveness Model. Unidentifiable “X males” were excluded from the calculations of any statistics dealing with pairwise histories, totals for the day, degree metrics, metrics from “yesterday,” and cumulative measures, since these males as they appear in the data cannot be guaranteed to be the same or different individuals through time.

**Table S2.**
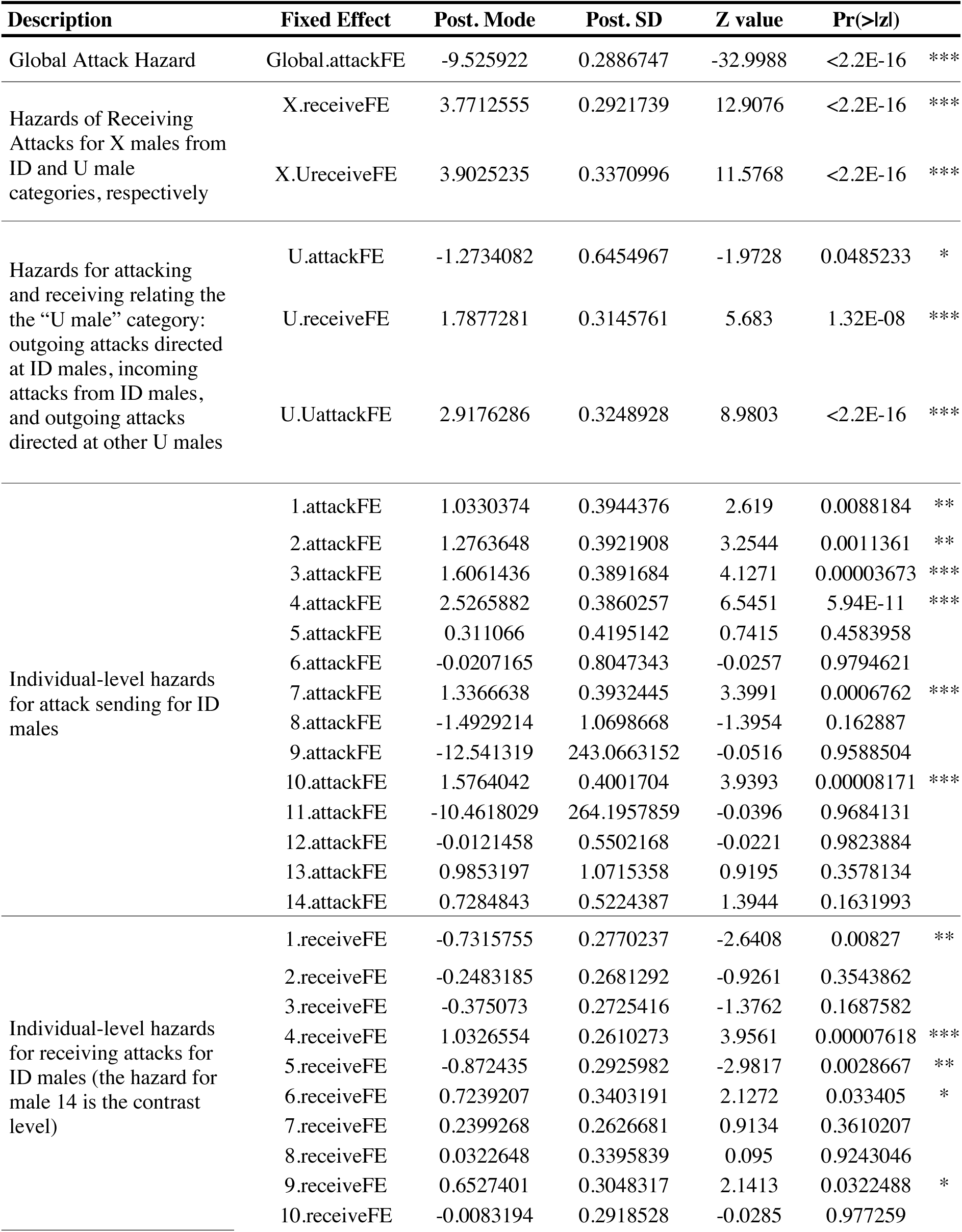
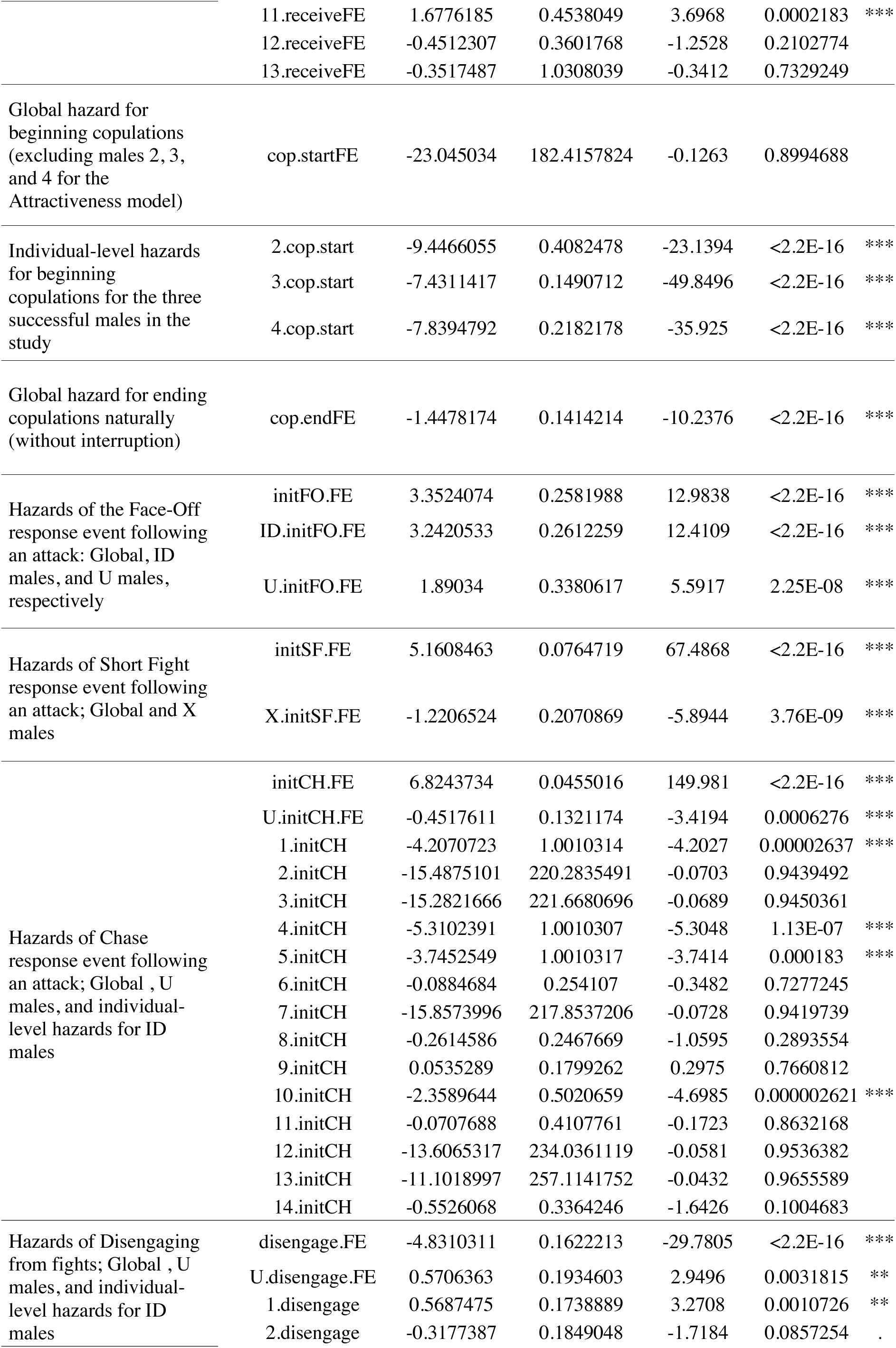
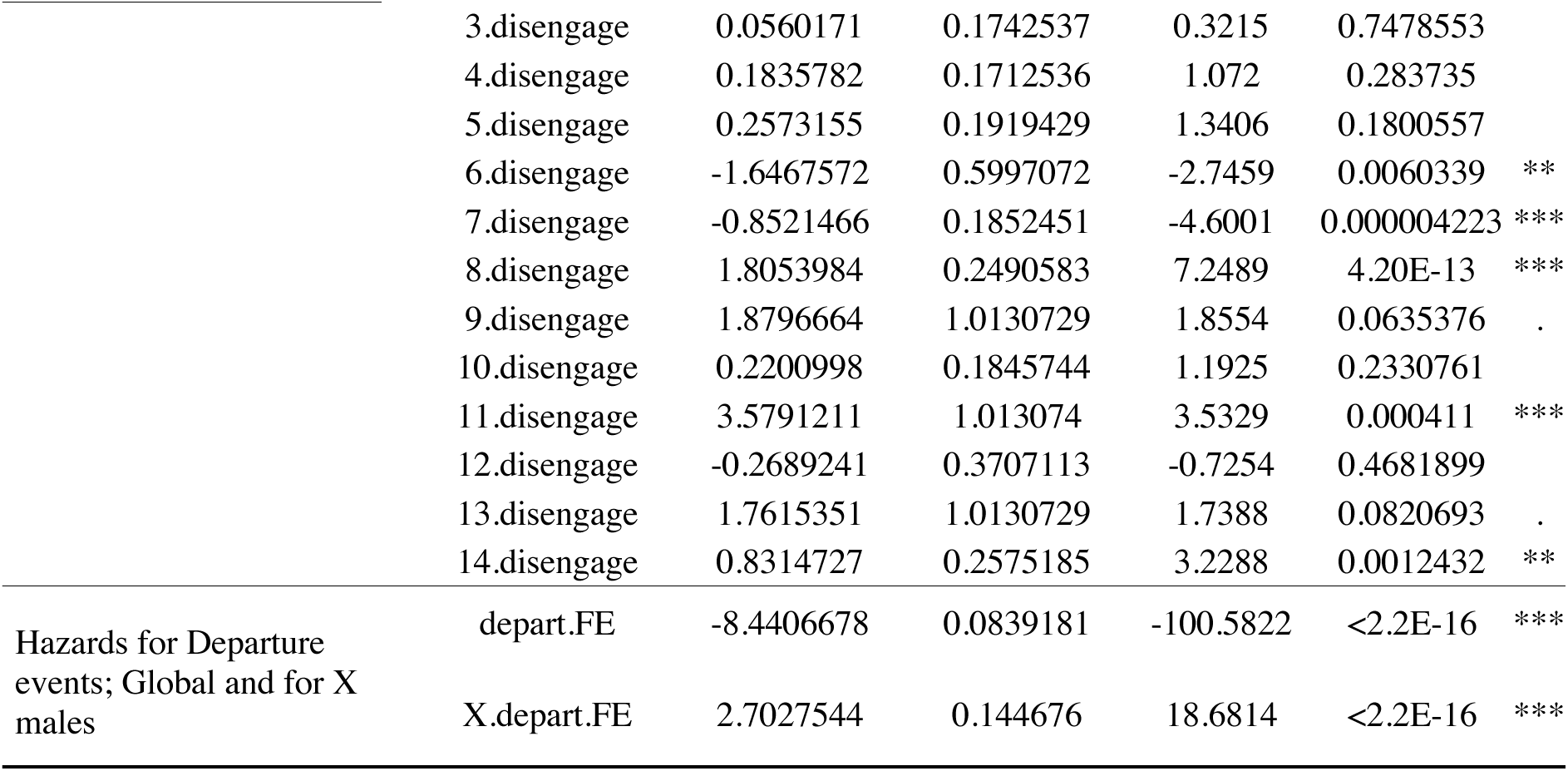
Fixed effects for the intercept-only REM. Differential Attractiveness Model fixed effects shown here; the Aggression Model is identical except that there is only a single global fixed effect for solicitations, and no individual-level effects. Null Deviance: 76628.21 on 4854 degrees of freedom. Residual Deviance: 30131.76 on 4777 degrees of freedom. Chi-square: 464496.45 on 77 degrees of freedom; asympototic p-value = 0. BIC: 30785.3. Log posterior: -92141.41. In the Bayesian framework, “p” values represent the probability that the sign of the estimated coefficient is incorrect. Significance codes: ‘***’ 0.001, ‘**’ 0.01, ‘*’ 0.05, ‘.’ 0.1.

**Table S3.** Differential Attractiveness REM full model results. BIC 24279.44. Statistics in bold represent the effect set for the best-fitting model. Non-bolded statistics are fixed effects. Null Deviance: 76628.21 on 4854 degrees of freedom. Residual Deviance: 23396.73 on 4750 deg. of freedom. Chi-square: 53231.48 on 104 degrees of freedom; asymptotic p-value=0. Log posterior: -115800.4. In the Bayesian framework, “p” values are the probability that the sign of the estimated coefficient is incorrect. Significance codes: ‘***’ 0.001, ‘**’ 0.01, ‘*’ 0.05, ‘.’ 0.1.

| **Statistic (Effect)** | **Post. Mode** | **Post. SD** | **Z value** | **Pr(>\|z\|)** |  |
| --- | --- | --- | --- | --- | --- |
| Global.attackFE | -9.5941184 | 0.294491 | -32.5786 | <2.2e-16 | *** |
| X.receiveFE | 2.6354545 | 0.30003 | 8.784 | <2.2e-16 | *** |
| X.UreceiveFE | 2.7456501 | 0.3432097 | 7.9999 | 1.33e-15 | *** |
| U.attackFE | 1.5330085 | 0.6627589 | 2.3131 | 0.0207187 | * |
| U.receiveFE | 4.1077 | 0.3478459 | 11.809 | <2.2e-16 | *** |
| U.UattackFE | 3.8164567 | 0.3443854 | 11.0819 | <2.2e-16 | *** |
| 1.attackFE | 3.5052453 | 0.4279641 | 8.1905 | 2.22e-16 | *** |
| 2.attackFE | 3.4534056 | 0.4244792 | 8.1356 | 4.44e-16 | *** |
| 3.attackFE | 3.6357434 | 0.4139669 | 8.7827 | <2.2e-16 | *** |
| 4.attackFE | 4.1186892 | 0.4260613 | 9.6669 | <2.2e-16 | *** |
| 5.attackFE | 3.3878531 | 0.4437625 | 7.6344 | 2.27e-14 | *** |
| 6.attackFE | 4.6055624 | 0.8319983 | 5.5355 | 3.10e-08 | *** |
| 7.attackFE | 3.483542 | 0.4250436 | 8.1957 | 2.22e-16 | *** |
| 8.attackFE | 1.9775035 | 1.0520656 | 1.8796 | 0.0601573 | . |
| 9.attackFE | -9.6275877 | 274.3683673 | -0.0351 | 0.972008 |  |
| 10.attackFE | 4.4199308 | 0.4505061 | 9.811 | <2.2e-16 | *** |
| 11.attackFE | -7.0703225 | 314.8496779 | -0.0225 | 0.9820841 |  |
| 12.attackFE | 3.6782825 | 0.6261727 | 5.8742 | 4.25e-09 | *** |
| 13.attackFE | 4.2658929 | 1.118182 | 3.815 | 0.0001362 | *** |
| 14.attackFE | 3.5711553 | 0.5594352 | 6.3835 | 1.73e-10 | *** |
| 1.receiveFE | -1.8819638 | 0.2891327 | -6.509 | 7.57e-11 | *** |
| 2.receiveFE | -1.9658524 | 0.2915699 | -6.7423 | 1.56e-11 | *** |
| 3.receiveFE | -1.8633946 | 0.2917904 | -6.3861 | 1.70e-10 | *** |
| 4.receiveFE | -1.1955048 | 0.2876318 | -4.1564 | 3.23e-05 | *** |
| 5.receiveFE | -1.0131437 | 0.3037432 | -3.3355 | 0.0008514 | *** |
| 6.receiveFE | 0.9828422 | 0.3493369 | 2.8135 | 0.0049013 | ** |
| 7.receiveFE | -1.6769817 | 0.286363 | -5.8561 | 4.74E-09 | *** |
| 8.receiveFE | 0.245951 | 0.3580668 | 0.6869 | 0.4921547 |  |
| 9.receiveFE | -0.1259594 | 0.3186162 | -0.3953 | 0.6925974 |  |
| 10.receiveFE | -0.0050913 | 0.3079875 | -0.0165 | 0.9868109 |  |
| 11.receiveFE | 0.8374651 | 0.482819 | 1.7345 | 0.0828237 | . |
| 12.receiveFE | -0.100656 | 0.4044671 | -0.2489 | 0.8034686 |  |
| 13.receiveFE | 0.4412992 | 1.1228393 | 0.393 | 0.6943041 |  |
| cop.startFE | -22.2491026 | 185.4741855 | -0.12 | 0.9045165 |  |
| 2.cop.start | -8.7536194 | 0.4347607 | -20.1343 | <2.2e-16 | *** |
| 3.cop.start | -7.4089176 | 0.2408981 | -30.7554 | <2.2e-16 | *** |
| 4.cop.start | -7.684103 | 0.2635543 | -29.1557 | <2.2e-16 | *** |
| cop.endFE | -1.4478174 | 0.1414214 | -10.2376 | <2.2e-16 | *** |
| initFO.FE | 3.3524074 | 0.2581988 | 12.9838 | <2.2e-16 | *** |
| ID.initFO.FE | 3.2420533 | 0.2612259 | 12.4109 | <2.2e-16 | *** |
| U.initFO.FE | 1.89034 | 0.3380617 | 5.5917 | 2.25e-08 | *** |
| initSF.FE | 5.1608463 | 0.0764719 | 67.4868 | <2.2e-16 | *** |
| X.initSF.FE | -1.2206524 | 0.2070869 | -5.8944 | 3.76e-09 | *** |
| initCH.FE | 6.8243734 | 0.0455016 | 149.981 | <2.2e-16 | *** |
| U.initCH.FE | -0.4517611 | 0.1321174 | -3.4194 | 0.0006276 | *** |
| 1.initCH | -4.2070723 | 1.0010314 | -4.2027 | 2.64e-05 | *** |
| 2.initCH | -15.4875101 | 220.2835491 | -0.0703 | 0.9439492 |  |
| 3.initCH | -15.2821666 | 221.6680696 | -0.0689 | 0.9450361 |  |
| 4.initCH | -5.3102391 | 1.0010307 | -5.3048 | 1.13e-07 | *** |
| 5.initCH | -3.7452549 | 1.0010317 | -3.7414 | 0.000183 | *** |
| 6.initCH | -0.0884684 | 0.254107 | -0.3482 | 0.7277245 |  |
| 7.initCH | -15.8573996 | 217.8537206 | -0.0728 | 0.9419739 |  |
| 8.initCH | -0.2614586 | 0.2467669 | -1.0595 | 0.2893554 |  |
| 9.initCH | 0.0535289 | 0.1799262 | 0.2975 | 0.7660812 |  |
| 10.initCH | -2.3589644 | 0.5020659 | -4.6985 | 2.62e-06 | *** |
| 11.initCH | -0.0707688 | 0.4107761 | -0.1723 | 0.8632168 |  |
| 12.initCH | -13.6065317 | 234.0361119 | -0.0581 | 0.9536382 |  |
| 13.initCH | -11.1018997 | 257.1141752 | -0.0432 | 0.9655589 |  |
| 14.initCH | -0.5526068 | 0.3364246 | -1.6426 | 0.1004683 |  |
| disengage.FE | -6.8895682 | 0.1989629 | -34.6274 | <2.2e-16 | *** |
| U.disengage.FE | 1.1775891 | 0.2104381 | 5.5959 | 2.20e-08 | *** |
| 1.disengage | 1.1302282 | 0.2414132 | 4.6817 | 2.85e-06 | *** |
| 2.disengage | 1.0585834 | 0.2450591 | 4.3197 | 1.56e-05 | *** |
| 3.disengage | 0.9989483 | 0.241377 | 4.1385 | 3.50e-05 | *** |
| 4.disengage | 1.1799068 | 0.234967 | 5.0216 | 5.13e-07 | *** |
| 5.disengage | 0.6272862 | 0.2266734 | 2.7674 | 0.0056513 | ** |
| 6.disengage | 1.1914852 | 0.6069975 | 1.9629 | 0.0496559 | * |
| 7.disengage | 0.7447842 | 0.243667 | 3.0566 | 0.0022389 | ** |
| 8.disengage | 1.236519 | 0.265623 | 4.6552 | 3.24e-06 | *** |
| 9.disengage | 3.2470071 | 1.0189348 | 3.1867 | 0.0014392 | ** |
| 10.disengage | 0.91412 | 0.2273969 | 4.0199 | 5.82e-05 | *** |
| 11.disengage | 3.1814565 | 1.0131276 | 3.1402 | 0.0016881 | ** |
| 12.disengage | 1.390207 | 0.3766766 | 3.6907 | 0.0002236 | *** |
| 13.disengage | 2.1678349 | 1.014493 | 2.1369 | 0.0326089 | * |
| 14.disengage | 1.3529772 | 0.2715032 | 4.9833 | 6.25e-07 | *** |
| depart.FE | -8.5393068 | 0.1487339 | -57.4133 | <2.2e-16 | *** |
| X.depart.FE | 2.0559312 | 0.2449126 | 8.3946 | <2.2e-16 | *** |
| fo.dis | -1.8407811 | 0.0882985 | -20.8473 | <2.2e-16 | *** |
| centroid.dist | -0.2391886 | 0.0143892 | -16.6228 | <2.2e-16 | *** |
| females.disengage | 1.5843288 | 0.070584 | 22.446 | <2.2e-16 | *** |
| gen.lose.rec | 0.6756445 | 0.1028391 | 6.5699 | 5.03e-11 | *** |
| disturb.depart | 6.077435 | 0.2543795 | 23.8912 | <2.2e-16 | *** |
| persistence | 0.577032 | 0.085633 | 6.7384 | 1.60e-11 | *** |
| ch.dis | 1.2550227 | 0.095248 | 13.1764 | <2.2e-16 | *** |
| terr.attack.cop | 3.8077728 | 0.3836473 | 9.9252 | <2.2e-16 | *** |
| depart.depart | 0.2447095 | 0.0299979 | 8.1575 | 4.44e-16 | *** |
| gen.lose.chase.rec.depart | 2.8355505 | 0.246584 | 11.4993 | <2.2e-16 | *** |
| gen.lose.chase.rec | 1.266837 | 0.1292308 | 9.8029 | <2.2e-16 | *** |
| win.loss.rate.pairwise.day.dis | 0.955583 | 0.139882 | 6.8314 | 8.41e-12 | *** |
| females.depart | -1.3681795 | 0.1652954 | -8.2772 | 2.22e-16 | *** |
| gen.receive.win.rec | 0.8838944 | 0.1182151 | 7.477 | 7.59e-14 | *** |
| log.cop.sol | 1.1270441 | 0.1768602 | 6.3725 | 1.86e-10 | *** |
| disturb.dis | 4.2437143 | 0.5050665 | 8.4023 | <2.2e-16 | *** |
| log.receive.count.pairwise.yest | 0.1703161 | 0.0470741 | 3.618 | 0.0002968 | *** |
| win.loss.rate.pairwise.yest.dis | 0.7881639 | 0.1648029 | 4.7825 | 1.73e-06 | *** |
| clock.sol | -0.9601822 | 0.2231803 | -4.3023 | 1.69e-05 | *** |
| gen.receive.win.rec.depart | 1.3301907 | 0.2809597 | 4.7345 | 2.20e-06 | *** |
| log.receive.lose.pairwise.yest.cop | 0.9059807 | 0.2466435 | 3.6732 | 0.0002395 | *** |
| gen.win.rec.anger | 0.3219296 | 0.0839812 | 3.8334 | 0.0001264 | *** |
| females.attack | -0.2259585 | 0.0679357 | -3.3261 | 0.0008808 | *** |
| terr.dis | 0.4611991 | 0.1335576 | 3.4532 | 0.000554 | *** |
| started.it.dis | 0.2471418 | 0.0715725 | 3.453 | 0.0005543 | *** |
| lose.noCH.rec | 0.4199225 | 0.0870307 | 4.825 | 1.40e-06 | *** |
| log.abs.winloss.diff.pairwise.day | 0.2519738 | 0.0512665 | 4.915 | 8.88e-07 | *** |

**Table S4.** Aggression Model REM full results. BIC 24285.54. Statistics in bold represent the effect set for the best-fitting model. Non-bolded statistics are fixed effects. Null Deviance: 76628.21 on 4854 degrees of freedom. Residual Deviance: 23419.81 on 4752 degrees of freedom. Chi-square: 53208.4 on 102 degrees of freedom; asymptotic p-value=0. Log posterior: -113810. In the Bayesian framework, “p” values are the probability that the sign of the estimated coefficient is incorrect. Significance codes: ‘***’ 0.001, ‘**’ 0.01, ‘*’ 0.05, ‘.’ 0.1.

| Statistic (Effect) | Post. Mode | Post. SD | Z value | Pr(>\|z\|) |  |
| --- | --- | --- | --- | --- | --- |
| Global.attackFE | -9.5941184 | 0.294491 | -32.5786 | <2.2e-16 | *** |
| X.receiveFE | 2.6354545 | 0.30003 | 8.784 | <2.2e-16 | *** |
| X.UreceiveFE | 2.7456501 | 0.3432097 | 7.9999 | 1.33E-15 | *** |
| U.attackFE | 1.5330085 | 0.6627589 | 2.3131 | 0.0207187 | * |
| U.receiveFE | 4.1077 | 0.3478459 | 11.809 | <2.2e-16 | *** |
| U.UattackFE | 3.8164567 | 0.3443854 | 11.0819 | <2.2e-16 | *** |
| 1.attackFE | 3.5052453 | 0.4279641 | 8.1905 | 2.22E-16 | *** |
| 2.attackFE | 3.4534056 | 0.4244792 | 8.1356 | 4.44E-16 | *** |
| 3.attackFE | 3.6357434 | 0.4139669 | 8.7827 | <2.2e-16 | *** |
| 4.attackFE | 4.1186892 | 0.4260613 | 9.6669 | <2.2e-16 | *** |
| 5.attackFE | 3.3878531 | 0.4437625 | 7.6344 | 2.27E-14 | *** |
| 6.attackFE | 4.6055624 | 0.8319983 | 5.5355 | 3.10E-08 | *** |
| 7.attackFE | 3.483542 | 0.4250436 | 8.1957 | 2.22E-16 | *** |
| 8.attackFE | 1.9775035 | 1.0520656 | 1.8796 | 0.0601573 | . |
| 9.attackFE | -9.6273195 | 274.3350443 | -0.0351 | 0.9720054 |  |
| 10.attackFE | 4.4199308 | 0.4505061 | 9.811 | <2.2e-16 | *** |
| 11.attackFE | -7.0703225 | 314.8496777 | -0.0225 | 0.9820841 |  |
| 12.attackFE | 3.6782825 | 0.6261727 | 5.8742 | 4.25E-09 | *** |
| 13.attackFE | 4.2658929 | 1.118182 | 3.815 | 0.0001362 | *** |
| 14.attackFE | 3.5711553 | 0.5594352 | 6.3835 | 1.73E-10 | *** |
| 1.receiveFE | -1.8819638 | 0.2891327 | -6.509 | 7.57E-11 | *** |
| 2.receiveFE | -1.9658524 | 0.2915699 | -6.7423 | 1.56E-11 | *** |
| 3.receiveFE | -1.8633946 | 0.2917904 | -6.3861 | 1.70E-10 | *** |
| 4.receiveFE | -1.1955048 | 0.2876318 | -4.1564 | 3.23E-05 | *** |
| 5.receiveFE | -1.0131437 | 0.3037432 | -3.3355 | 0.0008514 | *** |
| 6.receiveFE | 0.9828422 | 0.3493369 | 2.8135 | 0.0049013 | ** |
| 7.receiveFE | -1.6769817 | 0.286363 | -5.8561 | 4.74E-09 | *** |
| 8.receiveFE | 0.245951 | 0.3580668 | 0.6869 | 0.4921547 |  |
| 9.receiveFE | -0.1259594 | 0.3186162 | -0.3953 | 0.6925974 |  |
| 10.receiveFE | -0.0050913 | 0.3079875 | -0.0165 | 0.9868109 |  |
| 11.receiveFE | 0.8374651 | 0.482819 | 1.7345 | 0.0828237 | . |
| 12.receiveFE | -0.100656 | 0.4044671 | -0.2489 | 0.8034686 |  |
| 13.receiveFE | 0.4412992 | 1.1228393 | 0.393 | 0.6943041 |  |
| cop.startFE | -9.803251 | 0.2682293 | -36.548 | <2.2e-16 | *** |
| cop.endFE | -1.4478174 | 0.1414214 | -10.2376 | <2.2e-16 | *** |
| initFO.FE | 3.3524074 | 0.2581988 | 12.9838 | <2.2e-16 | *** |
| ID.initFO.FE | 3.2420533 | 0.2612259 | 12.4109 | <2.2e-16 | *** |
| U.initFO.FE | 1.89034 | 0.3380617 | 5.5917 | 2.25E-08 | *** |
| initSF.FE | 5.1608463 | 0.0764719 | 67.4868 | <2.2e-16 | *** |
| X.initSF.FE | -1.2206524 | 0.2070869 | -5.8944 | 3.76E-09 | *** |
| initCH.FE | 6.8243734 | 0.0455016 | 149.981 | <2.2e-16 | *** |
| U.initCH.FE | -0.4517611 | 0.1321174 | -3.4194 | 0.0006276 | *** |
| 1.initCH | -4.2070723 | 1.0010314 | -4.2027 | 2.64E-05 | *** |
| 2.initCH | -15.4875098 | 220.2835182 | -0.0703 | 0.9439492 |  |
| 3.initCH | -15.2821666 | 221.6680659 | -0.0689 | 0.945036 |  |
| 4.initCH | -5.3102391 | 1.0010307 | -5.3048 | 1.13E-07 | *** |
| 5.initCH | -3.7452549 | 1.0010317 | -3.7414 | 0.000183 | *** |
| 6.initCH | -0.0884684 | 0.254107 | -0.3482 | 0.7277245 |  |
| 7.initCH | -15.857393 | 217.8530504 | -0.0728 | 0.9419737 |  |
| 8.initCH | -0.2614586 | 0.2467669 | -1.0595 | 0.2893554 |  |
| 9.initCH | 0.0535289 | 0.1799262 | 0.2975 | 0.7660812 |  |
| 10.initCH | -2.3589644 | 0.5020659 | -4.6985 | 2.62E-06 | *** |
| 11.initCH | -0.0707688 | 0.4107761 | -0.1723 | 0.8632168 |  |
| 12.initCH | -13.6065317 | 234.0361119 | -0.0581 | 0.9536382 |  |
| 13.initCH | -11.1018997 | 257.1141752 | -0.0432 | 0.9655589 |  |
| 14.initCH | -0.5526068 | 0.3364246 | -1.6426 | 0.1004683 |  |
| disengage.FE | -6.8895682 | 0.1989629 | -34.6274 | <2.2e-16 | *** |
| U.disengage.FE | 1.1775891 | 0.2104381 | 5.5959 | 2.20E-08 | *** |
| 1.disengage | 1.1302282 | 0.2414132 | 4.6817 | 2.85E-06 | *** |
| 2.disengage | 1.0585834 | 0.2450591 | 4.3197 | 1.56E-05 | *** |
| 3.disengage | 0.9989483 | 0.241377 | 4.1385 | 3.50E-05 | *** |
| 4.disengage | 1.1799068 | 0.234967 | 5.0216 | 5.13E-07 | *** |
| 5.disengage | 0.6272862 | 0.2266734 | 2.7674 | 0.0056513 | ** |
| 6.disengage | 1.1914852 | 0.6069975 | 1.9629 | 0.0496559 | * |
| 7.disengage | 0.7447842 | 0.243667 | 3.0566 | 0.0022389 | ** |
| 8.disengage | 1.236519 | 0.265623 | 4.6552 | 3.24E-06 | *** |
| 9.disengage | 3.2470071 | 1.0189348 | 3.1867 | 0.0014392 | ** |
| 10.disengage | 0.91412 | 0.2273969 | 4.0199 | 5.82E-05 | *** |
| 11.disengage | 3.1814565 | 1.0131276 | 3.1402 | 0.0016881 | ** |
| 12.disengage | 1.390207 | 0.3766766 | 3.6907 | 0.0002236 | *** |
| 13.disengage | 2.1678349 | 1.014493 | 2.1369 | 0.0326089 | * |
| 14.disengage | 1.3529772 | 0.2715032 | 4.9833 | 6.25E-07 | *** |
| depart.FE | -8.5393068 | 0.1487339 | -57.4133 | <2.2e-16 | *** |
| X.depart.FE | 2.0559312 | 0.2449126 | 8.3946 | <2.2e-16 | *** |
| fo.dis | -1.8407811 | 0.0882985 | -20.8473 | <2.2e-16 | *** |
| centroid.dist | -0.2391886 | 0.0143892 | -16.6228 | <2.2e-16 | *** |
| females.disengage | 1.5843288 | 0.070584 | 22.446 | <2.2e-16 | *** |
| gen.lose.rec | 0.6756445 | 0.1028391 | 6.5699 | 5.03E-11 | *** |
| disturb.depart | 6.077435 | 0.2543795 | 23.8912 | <2.2e-16 | *** |
| persistence | 0.577032 | 0.085633 | 6.7384 | 1.60E-11 | *** |
| ch.dis | 1.2550227 | 0.095248 | 13.1764 | <2.2e-16 | *** |
| terr.attack.cop | 3.8077728 | 0.3836473 | 9.9252 | <2.2e-16 | *** |
| log.cop.cumul.sol | 0.8608124 | 0.0930913 | 9.247 | <2.2e-16 | *** |
| depart.depart | 0.2447095 | 0.0299979 | 8.1575 | 4.44E-16 | *** |
| gen.lose.chase.rec.depart | 2.8355505 | 0.246584 | 11.4993 | <2.2e-16 | *** |
| gen.lose.chase.rec | 1.266837 | 0.1292308 | 9.8029 | <2.2e-16 | *** |
| win.loss.rate.pairwise.day.dis | 0.955583 | 0.139882 | 6.8314 | 8.41E-12 | *** |
| females.depart | -1.3681795 | 0.1652954 | -8.2772 | 2.22E-16 | *** |
| gen.receive.win.rec | 0.8838944 | 0.1182151 | 7.477 | 7.59E-14 | *** |
| disturb.dis | 4.2437143 | 0.5050665 | 8.4023 | <2.2e-16 | *** |
| log.receive.count.pairwise.yest | 0.1703161 | 0.0470741 | 3.618 | 0.0002968 | *** |
| win.loss.rate.pairwise.yest.dis | 0.7881639 | 0.1648029 | 4.7825 | 1.73E-06 | *** |
| gen.receive.win.rec.depart | 1.3301907 | 0.2809597 | 4.7345 | 2.20E-06 | *** |
| log.receive.lose.pairwise.yest.cop | 0.9059807 | 0.2466435 | 3.6732 | 0.0002395 | *** |
| gen.win.rec.anger | 0.3219296 | 0.0839812 | 3.8334 | 0.0001264 | *** |
| females.attack | -0.2259585 | 0.0679357 | -3.3261 | 0.0008808 | *** |
| log.receive.inter.sol | 1.6301689 | 0.2724507 | 5.9834 | 2.19E-09 | *** |
| log.lose.weight.day.sol | -0.9711966 | 0.2096217 | -4.6331 | 3.60E-06 | *** |
| terr.dis | 0.4611991 | 0.1335576 | 3.4532 | 0.000554 | *** |
| started.it.dis | 0.2471418 | 0.0715725 | 3.453 | 0.0005543 | *** |
| lose.noCH.rec | 0.4199225 | 0.0870307 | 4.825 | 1.40E-06 | *** |
| log.abs.winloss.diff.pairwise.day | 0.2519738 | 0.0512665 | 4.915 | 8.88E-07 | *** |

## Notes

### Competing Interest Statement

The authors have declared no competing interest.

