## Supplemental_reproducible_code for "Fighting isn’t sexy in lekking Greater Sage-grouse (*Centrocercus urophasianus*)": relevent_analysis_base_udata_terr_adequacy.R

```
### Code for Analyzing Aggression Model Adequacy

rm(list=ls())

###make sure necessary packages are there and loaded

packages<-c("here","abind","relevent","lubridate","DescTools","gtools","e1071","ggplot2","magrittr","rcartocolor","ggpubr","cowplot")

package.check<-lapply(packages, FUN = function(x) {
    if (!require(x, character.only = TRUE, quietly=TRUE)) {
        install.packages(x, dependencies = TRUE)
        library(x, character.only = TRUE, quietly=TRUE)
    } 

})

#for project path management
#define the project root directory relative to the current script. This sets the working directory that gets called by the here() function. It is a relative path and will be robust to use on anyone's computer. In this case, I want the root directory to always be the enclosing folder for the whole sage grouse project

i_am("adequacy_checking/adequacy_aggression_model/relevent_analysis_base_udata_terr_adequacy.R")


##**If the per event adequacy data has already been produced, can skip the beginning portion of the code and go down to the Adequacy Analysis section

######


####If I need to re-run the model and reproduce the int.list:

#Build the intercept list, support array, event list, census array, etc

{
	
#make the base event list and intercept array


event.list<-read.table(file=here("data_processing","relevent_format_dataset_udata_wincol_disturbance.tsv"), header=T,  sep="\t")

event.list$Date<-as.factor(event.list$Date)
  
event.types<-read.table(file=here("data_processing","event_types_list_udata.csv"), header=F,  sep=",")
event.types<-as.character(event.types$V1)


#NOTE- the event.types vector now contains ENDOGENOUS event types as well as EXOGENOUS types: arrivals and clock events, and end.obs. Need to make sure everything is robust to this. 


### create eventlist for rem ###


event.list.dat<-split(event.list, event.list$Date)


#remove female context for now

basic.event.list<-event.list[-3]

#strip the "loser" from the disengage events so they are all the same event "type" and they match my previous code

basic.event.list$Events<-sapply(basic.event.list$Events, function(x) {
	
	event<-unlist(strsplit(as.character(x),".",fixed=T))
	if(event[2]=="disengage"){
		return(paste(event[1],event[2],sep="."))
	} else {return(as.character(x))}
}
)


#add factor levels for event types that don't actually happen (and put them in the same order as the event.type list) to maintain bookkeeping 

basic.event.list$Events<-as.character(basic.event.list$Events)

basic.event.list$Events<-factor(basic.event.list$Events, levels=as.character(event.types))

#turn event types into numeric values

basic.event.list$Events<-as.numeric(basic.event.list$Events)

#set the event type code for exogenous events to zero. I have there all at the trailing end of the vector

endogenous<-(length(event.types)-length(event.types[grep("arrive",event.types)])-5)

basic.event.list$Events[basic.event.list$Events>endogenous]<-0

rownames(basic.event.list)<-NULL
#need to divide the data up by grouping factor

evlist<-list()
evlist<-split(basic.event.list, basic.event.list$Date)

#strip the date column from the sub-dataframes so they're just two columns, event and time

evlist<-lapply(evlist, function(x) x<-x[-3])

#convert from dataframes to matrices

evlist<-lapply(evlist, function(x) x<-as.matrix(x))

## make event.types vector usable for intlist, etc. by excising exogenous event types

event.types<-event.types[1:endogenous]


#### create statlist ####

##"a three dimensional array, event number by event type by statistic, or a list thereof, containing the sufficient statistics for the model"

#start with the intercept sufficient stats

#seems like the statlist involves an array with an ojbect for every "group" according to grouping factor. In the example it's the individuals but for me it's the different DAYS. each day will have a matrix for each event in the sequence, and for the intercepts, each matrix will just be an identity matrix of event types... I think. give it a try. This, I think, should EXCLUDE rows for all the exogenous event types as well (except end.obs which is still in there). I know this isn't what Carter would prefer, but it's just adding dead events that we basically tell the model to ignore anyway. 


sequences<-table(event.list$Date)

#create list of dummy intercept names

idno<-length(event.types[grep("cop.start",event.types)])

dummy.ints.attack<-paste(1:14,".attackFE", sep="")
dummy.ints.receive<-paste(1:13, ".receiveFE", sep="")

IDintsCH<-paste(1:14,"initCH",sep=".")
IDdis<-paste(1:14,"disengage",sep=".")

int.names<-c("Global.attackFE", "X.receiveFE", "X.UreceiveFE", "U.attackFE", "U.receiveFE", "U.UattackFE", dummy.ints.attack, dummy.ints.receive, "cop.startFE","cop.endFE", "initFO.FE","ID.initFO.FE", "U.initFO.FE","initSF.FE","X.initSF.FE", "initCH.FE",  "U.initCH.FE", IDintsCH, "disengage.FE", "U.disengage.FE", IDdis, "depart.FE", "X.depart.FE")

int.list<-list()

for(i in 1:length(levels(event.list$Date))){

day.array<-array(0,dim=c(as.numeric(sequences[i]), length(event.types),length(int.names)))

#the third dimension is the stat slice.
dimnames(day.array)<-list(c(1:sequences[i]),as.character(event.types),as.character(int.names))

#add global attack effect, loops for "U" intercepts for attacks and receives, then one loop for typical intercepts

day.array[,grep("attack", event.types),1]<-1


for(h in 1:14){
day.array[,paste(h,".attack.",30, sep=""),2]<-1
}


for(h in 15:29){
day.array[,paste(h,".attack.",30, sep=""),3]<-1
}


for(h in 1:14){
	for(j in 15:29){	
day.array[,paste(j,".attack.", h, sep=""),4]<-1
}
}


for(h in 1:14){
	for(j in 15:29){
day.array[,paste(h,".attack.",j, sep=""),5]<-1
}
}

for(h in 15:29){
	for(j in 15:29){	
if(h!=j){
day.array[,paste(j,".attack.", h, sep=""),6]<-1
}
}
}

preloop<-6

#loop for dummy intercepts for attacks, loop for dummy intercepts for receives, add in cop.start and cop.end FEs, then one loop for typical intercepts

for(k in (preloop+1):(length(dummy.ints.attack)+preloop)){
	
	for(h in 1:14){
if(h!=(k-preloop)){
day.array[,paste((k-preloop),".attack.", h, sep=""),k]<-1
}
}
}

for(k in ((length(dummy.ints.attack)+preloop)+1):((length(dummy.ints.attack)+preloop)+length(dummy.ints.receive))){
	for(h in 1:14){
if(h!=(k-(length(dummy.ints.attack)+preloop))){
day.array[,paste(h,".attack.",(k-(length(dummy.ints.attack)+preloop)), sep=""),k]<-1
}
}
}

#cop.startFE
for(h in 1:idno){
day.array[,paste(h,".cop.start", sep=""),((length(dummy.ints.attack)+preloop)+length(dummy.ints.receive)+1)]<-1
}


#cop.endFE
for(h in 1:idno){
day.array[,paste(h,".cop.end", sep=""),((length(dummy.ints.attack)+preloop)+length(dummy.ints.receive)+2)]<-1
}

#initFO.FE
for(h in 1:idno){
day.array[,paste(h,".initFO", sep=""),((length(dummy.ints.attack)+preloop)+length(dummy.ints.receive)+3)]<-1
}

#ID.initFO.FE
for(h in 1:14){
day.array[,paste(h,".initFO", sep=""),((length(dummy.ints.attack)+preloop)+length(dummy.ints.receive)+4)]<-1
}

#U.initFO.FE
for(h in 15:29){
day.array[,paste(h,".initFO", sep=""),((length(dummy.ints.attack)+preloop)+length(dummy.ints.receive)+5)]<-1
}

#initSF.FE
for(h in 1:idno){
day.array[,paste(h,".initSF", sep=""),((length(dummy.ints.attack)+preloop)+length(dummy.ints.receive)+6)]<-1
}

#X.initSF.FE
day.array[,paste(30,".initSF", sep=""),((length(dummy.ints.attack)+preloop)+length(dummy.ints.receive)+7)]<-1


#initCH.FE
for(h in 1:idno){
day.array[,paste(h,".initCH", sep=""),((length(dummy.ints.attack)+preloop)+length(dummy.ints.receive)+8)]<-1
}


#U.initCH.FE
for(h in 15:29){
day.array[,paste(h,".initCH", sep=""),((length(dummy.ints.attack)+preloop)+length(dummy.ints.receive)+9)]<-1
}


#individual initCH effects for ID male
for(k in (((length(dummy.ints.attack)+preloop)+length(dummy.ints.receive)+10):((length(dummy.ints.attack)+preloop)+length(dummy.ints.receive)+10+(length(IDintsCH)-1)))){
	
	day.array[,int.names[k],k]<-1
	
}

preloop2<-((length(dummy.ints.attack)+preloop)+length(dummy.ints.receive)+10+(length(IDintsCH)-1))


#disengage.FE
for(h in 1:idno){
day.array[,paste(h,".disengage", sep=""),preloop2+1]<-1
}

#U.disengage.FE
for(h in 15:29){
day.array[,paste(h,".disengage", sep=""),preloop2+2]<-1
}

#individual disengage effect for ID males
for(k in (preloop2+3):(preloop2+3+length(IDdis)-1)){
	
	day.array[,int.names[k],k]<-1
	
}


preloop3<-(preloop2+3+length(IDdis)-1)

#depart.FE
for(h in 1:idno){
day.array[,paste(h,".depart", sep=""),preloop3+1]<-1
}

#X.depart.FE
day.array[,paste(30,".depart", sep=""),preloop3+2]<-1


#weird extra formatting to make it match the package example, seems like it becomes relevant *punintended!* if I want to fit day-specific effects

global<-list()

global[[1]]<-day.array

int.list[[i]]<-global

names(int.list[[i]])<-"global"

}


#### Create Support Matrix ####

#"should be a list withone element per history, each of which should be an event order by event type logical matrix. the ijth cell of this matrix should be TRUE if an event type of j was a possible next event given the preceding code i-1 events, and FALSE otherwise"

#need a way to ask, "what is each male doing right now" for each event in the sequence. That dictates the possibilities. 

#create a bookkeeping "census" matrix to keep track

census.list<-list()


for(j in 1:length(event.list.dat)){

census<-matrix("nothing", nrow=1,ncol=(idno), dimnames=list(1,c(1:(idno))))


##code that translates each new current event into what the males involved are doing at that time. The support matrix will be based on whatever the males were doing in the PREVIOUS timestep. This census matrix itself could be interesting to visualize. Each row of the census says what each male is doing going into the NEXT event.


for(i in 1:nrow(event.list.dat[[j]])){

#can ask if the second thing is something, and then move on to the first and third elements.

current<-census[nrow(census),]

#parse the event type. loop through test.seq rows
evname<-as.character(event.list.dat[[j]][i,1])
evsplit<-strsplit(evname, split=".", fixed=T)

#if it is an end obs, arrival, departure or clock event, it shouldn't change anything the males are doing in terms of interactions. Departures aren't exogenous, but they shouldn't change anything since they can only happen once a male is already doing nothing. 
if(evsplit[[1]][2]!="obs" & evsplit[[1]][2]!="depart" & evsplit[[1]][2]!="arrive" & evsplit[[1]][2]!="time" & evsplit[[1]][1]!="fem"){
#if the event was an attack, indicate the males involved and which was the receiver since this dictates who makes the initition decision
	if(evsplit[[1]][2]=="attack"){
		current[evsplit[[1]][1]]<-evname
		current[evsplit[[1]][3]]<-paste(evsplit[[1]][3], ".", "received", ".", evsplit[[1]][1], sep="")
			}
	#if it's an FO or SF, just indicate that that pair of males are involved in a fight now
		if(evsplit[[1]][2]=="initFO"||evsplit[[1]][2]=="initSF"){
			#need to refer back to the previous row of "census" to get the id of the other male
			
			current[evsplit[[1]][1]]<-paste(evsplit[[1]][1], ".", "fight", ".",strsplit(census[nrow(census),evsplit[[1]][1]], split=".",fixed=T)[[1]][3], sep="")
			current[[strsplit(census[nrow(census),evsplit[[1]][1]], split=".",fixed=T)[[1]][3]]]<-paste(strsplit(census[nrow(census),evsplit[[1]][1]], split=".",fixed=T)[[1]][3], ".", "fight", ".",evsplit[[1]][1], sep="")
			
			}
			
	#if it's a chase, indicate that the pair of males are involved in a fight, but there is an additional constraint that the individual DOING THE CHASING will be the disengager by definition
	if(evsplit[[1]][2]=="initCH"){
			
		current[evsplit[[1]][1]]<-paste(evsplit[[1]][1], ".", "fight", ".",strsplit(census[nrow(census),evsplit[[1]][1]], split=".",fixed=T)[[1]][3], ".chasedby",sep="")
			current[[strsplit(census[nrow(census),evsplit[[1]][1]], split=".",fixed=T)[[1]][3]]]<-paste(strsplit(census[nrow(census),evsplit[[1]][1]], split=".",fixed=T)[[1]][3], ".", "fight", ".",evsplit[[1]][1], ".chasing",sep="")

	}

#for disengaging, end the fight for the partner male as well
	if(evsplit[[1]][2]=="disengage"){
	current[evsplit[[1]][1]]<-"nothing"
	current[evsplit[[1]][3]]<-"nothing"	
	
}
 

#for copulations, end cops are only possible after start cops, but otherwise anything else is possible. EXCEPT the copulating male can't initiate an attack. If the next event involving that male ISN'T an end cop, it was interrupted

	if(evsplit[[1]][2]=="cop" && evsplit[[1]][3]=="start"){
		current[evsplit[[1]][1]]<-"copulating"
	}

	if(evsplit[[1]][2]=="cop" && evsplit[[1]][3]=="end"){
		current[evsplit[[1]][1]]<-"nothing"
	}

#if an X male is still involved in a fight at all, make sure he is still tagged as fighting (since sometimes more than one is interacting at once, but they can't depart until none are fighting)
	if((TRUE %in% grepl("fight.30",current) | TRUE %in% grepl("30.fight",current)) && !(grepl("receive", current[idno]) | grepl("attack", current[idno]))) {
		
		current[idno]<-"fight"

}

}

census<-rbind(census,current)


}


#don't want the first dummy row
census.list[[j]]<-census[-1,]

}


## Constraints to incorporate in the support matrix ##

#only males that are actually present can be involved in interactions on a given day and at a given time

#If "nothing" is happening, all "attacks", all start cops and all departs (given the male is actually present) are possible. Males cannot disengage or initiate fights without an attack happening first. End cop cannot happen without a previous start cop. end.obs is possible. (end.obs is not possible if any males are still fighting or copulating, so end.obs is ONLY possible when "nothing" is going on across the board)

#if the previous event was an "attack", the current event MUST be an initiation by the receiver, since we are assuming the attack is essentually instantaneous. 

#once initiatiated, subsequent events can only be a disengage from either male involved or any other event NOT invoving the interaction partners. Males actively fighting connot become engaged in a new fight until they disengage. 

#if the initiated fight was a chase only the male being chased can disengage, by definition

#Once a copulation starts, the only thing that male can do next is either end the copulation OR receive an attack. He cannot initiate anything. 

#copulations cannot occur when there are no females present

#a male can ONLY "depart" if "nothing" is happening

#Exceptions to these rules are for "X" males- there may be more than one unidentifiable male on the lek at a time. Right now we are dealing with this by having "X" males just be a mass actor category. 1) These males CAN do an "attack" before disengaging from an interaction. They can also be attacked. However, normal restrictions apply to the partner male. So if an X male is being chased, "disengage" becomes possible, but any "attack" (or cop.start, or depart) are still also possible 


### Build the actual support matrix ###

#for each event, there is a list of event types, with 1's for the types that are possible and 0's for types that are not. 


#code block to produce the support matrix

supplist<-list()


#bring in day-level male presence data

male.pres<-read.table(here("data_files","CHG_2014_male_presence_data_quest.csv"), sep=",", header=T)

#male IDs are already in ascending order, so I can convert them directly
colnames(male.pres)<-c("Date", 1:idno)

#bring in exact male presence data

pres.exact<-read.table(here("data_files","male_pres_exact_quest.csv"), sep=",",header=T)

	departlist<-list()
	
	for(j in 1:nrow(pres.exact)){
		
		arriverow<-pres.exact[j,startsWith(colnames(pres.exact),"arrive")]
		arriverow<-arriverow[!is.na(arriverow)]
		
		arrives<-t(sapply(arriverow, function(x) return(c(paste0(pres.exact$Num_ID[j],".arrive"),x,as.character(pres.exact$Date[j])))))
		
		departrow<-pres.exact[j,startsWith(colnames(pres.exact),"depart")]
		departrow<-departrow[!is.na(departrow)]
		
		departs<-t(sapply(departrow, function(x) return(c(paste0(pres.exact$Num_ID[j],".depart"),x,as.character(pres.exact$Date[j])))))
		
		allpres<-rbind(arrives,departs)
		
		departlist[[j]]<-allpres
		
	}

#collapse the departlist 

departevent<-abind(departlist, along=1)

departevent<-data.frame(departevent)
names(departevent)<-c("Events","Time", "Date")

departevent$Time<-as.numeric(as.character(departevent$Time))

library(abind)
arrstr<-abind(strsplit(as.character(departevent$Events),".",fixed=T),along=2)

departevent$Bird_ID<-arrstr[1,]
departevent$Type<-arrstr[2,]

departevent$Type<-as.factor(departevent$Type)
departevent$Bird_ID<-as.factor(departevent$Bird_ID)

# #split into day-specific data frames
pres.exact.dat<-split(departevent, departevent$Date)


#make a reference event list with data (most importantly female presence)
evlist.dat<-event.list.dat


for(j in 1:length(evlist)){

supplist[[j]]<-matrix(1, nrow=length(evlist[[j]][,1]), ncol=length(event.types))
colnames(supplist[[j]])<-event.types

#turn off all interactions involving males not present on that day
#make a list of all the event types that include the males that were entirely absent and remove them for all rows

supplist[[j]][,unique(unlist(lapply(grep(0,male.pres[j,2:(idno+1)]), function(x) c(event.types[startsWith(event.types, paste(x, ".", sep=""))], event.types[endsWith(event.types, paste(".", x, sep=""))]))))]<-0


#first row can be any "attack" or, technically, any "depart"
  #since now fem.enter is an actual event, cop.start is never possible as the very first thing
supplist[[j]][1,][-c(grep("attack", event.types), grep("depart", event.types))]<-0

#the exception is events involving males that will be present that day but haven't arrived yet


#basically I can make this code exactly the same as below- just want to ask who is actually there right now? 

arrsub<-subset(pres.exact.dat[[j]],Time<evlist[[j]][1,2] | Time==0 )
arrtab<-table(arrsub$Bird_ID,arrsub$Type)

arrpat<-apply(arrtab,1,function(x) sum(x))

#if the total number of previous arrives and departs is even, that male is currently absent
absent<-names(arrpat[(arrpat %% 2)==0])

if(length(absent)>0){
supplist[[j]][1,][unique(c(sapply(absent, function(x) c(event.types[startsWith(event.types, paste(x, ".", sep=""))], event.types[endsWith(event.types, paste(".", x, sep=""))]))))]<-0

} 


#loop through the rest of the events: 

for(i in 2:(length(evlist[[j]][,1])-1)){


#before anything else turn off events involving males that have either not yet arrived or have lef the lek early and are therefore no longer present that day or are otherwise not currently present


arrsub<-subset(pres.exact.dat[[j]],Time<evlist[[j]][i,2])
arrtab<-table(arrsub$Bird_ID,arrsub$Type)

arrpat<-apply(arrtab,1,function(x) sum(x))

#if the total number of previous arrives and departs is even, that male is currently absent
current.row.abs<-names(arrpat[(arrpat %% 2)==0])


if(length(current.row.abs)>0){
supplist[[j]][i,][unique(c(sapply(current.row.abs, function(x) c(event.types[startsWith(event.types, paste(x, ".", sep=""))], event.types[endsWith(event.types, paste(".", x, sep=""))]))))]<-0

} 


#determine if the previous event was an attack, since this dictates exactly what the next event can be: receiver initiates a type of fight
if((TRUE %in% grepl("received", census.list[[j]][i-1,]))==TRUE){

supplist[[j]][i,][-grep(TRUE, startsWith(event.types, paste(grep("received", census.list[[j]][i-1,]), ".", "init", sep="")))]<-0}


#(else) if the previous event wasn't an attack, the next event CAN'T be an initiation and the possibilities depend on what everyone is currently engaged in


if((TRUE %in% grepl("received", census.list[[j]][i-1,]))==FALSE){

supplist[[j]][i,][grep("init", event.types)]<-0

#if males are actively fighting, they can't attack or be attacked, or actually do anything else before disengaging (unless they are a ?? male, id 30)

if((TRUE %in% grepl("fight", census.list[[j]][i-1,]))==TRUE){
		
		for(k in grep("fight", census.list[[j]][i-1,])){
	
	if(k!=idno){
	
	supplist[[j]][i,][c(event.types[startsWith(event.types, paste(k, ".attack.", sep=""))], event.types[endsWith(event.types, paste("attack.", k, sep=""))], event.types[startsWith(event.types, paste(k, ".cop", sep=""))])]<-0
	
	}
	
}
	
	#ONLY those fighting can disengage. retain males that are fighting and remove the rest
	dis<-event.types[grep("disengage",event.types)]
	supplist[[j]][i,][dis[-grep("fight", census.list[[j]][i-1,])]]<-0
	
	#"chasing" and "chasedby" limit the possibilities further. Males getting chased DO NOT disengage by definition, again, with the exception of ?? males. This is taken care of by having ?? males just be "fight" -ing without any extra tags
	if(TRUE %in% grepl("chasedby", census.list[[j]][i-1,])){
		supplist[[j]][i,][dis[grep("chasedby", census.list[[j]][i-1,])]]<-0
	}
	
}

#if no one is fighting, no one can disengage, but all attack combinations are possible
if((TRUE %in% grepl("fight", census.list[[j]][i-1,]))==FALSE){
	
		supplist[[j]][i,][grep("disengage",event.types)]<-0
	
}


#actively copulating males: They cannot cop.start. ONLY they can cop.end. they CAN be attacked but they CANNOT attack anyone
if((TRUE %in% grepl("copulating", census.list[[j]][i-1,]))==TRUE){
	
	cop<-grep("copulating", census.list[[j]][i-1,]) 
	
	#stuff that needs to be turned off for only copulating males
	for(h in cop){

		supplist[[j]][i,][c(event.types[startsWith(event.types ,paste(h, ".attack.", sep=""))], event.types[startsWith(event.types, paste(h, ".cop.start", sep=""))])]<-0
				
	}
	
	#stuff that needs to be on for ONLY copulating males
	copendall<-event.types[grep("cop.end",event.types)]
	supplist[[j]][i,][copendall[-cop]]<-0
}

#if no one is copulating, all cop.starts are possible, but cop.ends are not

if((TRUE %in% grepl("copulating", census.list[[j]][i-1,]))==FALSE){
	
	copendall<-event.types[grep("cop.end",event.types)]
	supplist[[j]][i,][copendall]<-0
	
}


}

#if there are no females present, the next event could not have possibly been a cop.start

if(evlist.dat[[j]][i-1,]$Females==1){
	
	copstartall<-event.types[grep("cop.start",event.types)]
	supplist[[j]][i,][copstartall]<-0
	
}

#Finally, A male can only "depart" if he was already doing "nothing," so depart must be turned off for everyone doing not-nothing. (also he has to be present, but I've already taken care of that above)

noth<-as.character(which(census.list[[j]][i-1,]!="nothing"))
	
	if(length(noth)>0){
		
	supplist[[j]][i,][paste(noth, ".depart", sep="")]<-0

}

}
#end events loop here

#after done with all the middle events: onset of end.obs could depend on a lot of factors; not sure I want to fit it as part of the model. Just always say the last row must be end.obs (all endogenous events are no longer possible) 

supplist[[j]][length(evlist[[j]][,1]),]<-0

}
#end day loop here


###Make the win/loss census array

#i've adjusted the function to mark when a win or loss is the result of a chase or not
create.win.loss.census<-function(x){

#get disengagement events for focal male from evlist
win.loss.census<-x

#loop thru days and males

for(i in 1:length(win.loss.census)){

for(l in 1:idno){


dis.times<-which(startsWith(as.character(event.list.dat[[i]][,1]), paste(l,".disengage", sep="")))


if(length(dis.times)>0){

evsplit<-strsplit(as.character(event.list.dat[[i]][dis.times,1]),".",fixed=T)

#so I have a list of row numbers for disengagements and a list of interaction partners. This will cover wins for normal fights and losses for chases for the focal male. If I loop through all the males, all the wins and losses should get filled in.

#NOW DISENGAGES ALWAYS MEAN WINS, so I will still have chase information, but I don't need to do anything special with it here.

#need to make sure I am really precise since male 30 can be involved in multiple interactions, but they're usually all chases.


for(j in 1:length(dis.times)){
	
		
	if(grepl("chasing",win.loss.census[[i]][(dis.times[j]-1),l]) ==FALSE && grepl("chasedby",win.loss.census[[i]][(dis.times[j]-1),as.numeric(evsplit[[j]][3])]) ==FALSE){
	#apply the win in the focal male's column
	win.loss.census[[i]][dis.times[j],l]<-paste(l, "win",evsplit[[j]][3], sep=".")
	#apply the loss for the same interaction in the partner male's column
	win.loss.census[[i]][dis.times[j],as.numeric(evsplit[[j]][3])]<-paste(evsplit[[j]][3], "lose", l, sep=".")
	} else{
		#apply the won chase in the focal male's column
	win.loss.census[[i]][dis.times[j],l]<-paste(l, "win",evsplit[[j]][3],"chase", sep=".")
		#apply the loss for the same interaction in the partner male's column.
	win.loss.census[[i]][dis.times[j],as.numeric(evsplit[[j]][3])]<-paste(evsplit[[j]][3], "lose", l, "chase",sep=".")
		}
		
}

}

}

}

return(win.loss.census)
	
}

win.loss.census<-create.win.loss.census(census.list)


#other important stuff to load
### load in pairwise distance data
##bring in pairwise distances; I have a table for each day
##female exact presence data


file_list<-list.files(paste(here(),"/data_processing/udata_centroid_data/pairwise_dist",sep=""))

distlist<-list()

distlist<-lapply(file_list, function(x) read.table(paste(here(),"/data_processing/udata_centroid_data/pairwise_dist/",x,sep=""), sep=","))


#convert to martices

distlist.mat<-lapply(distlist, function(x) as.matrix(x, ))

#the names of the rows and columns are in the same order as the numerical ids passed to rem, so they can be used to index the matrix. Males that are absent on a given day have NA's for their centroid distances. I can change them to zeroes here so they don't produce errors, but the support array will take care of them anyway.

for(i in 1:length(distlist.mat)){
distlist.mat[[i]][is.na(distlist.mat[[i]])]<-0
}


fe_exact<-read.table(file=here("data_files","CHG_females_exact.tsv"), header=TRUE,sep="\t",stringsAsFactors=FALSE)

#reduce to only the necessary columns- this time I want time elapsed
fe_exact<-fe_exact[,c(1,3,5,7,9,11,13)]

#turn the first arrival NA's to 0s
fe_exact$first_arrival_elapsed[is.na(fe_exact$first_arrival_elapsed)]<-"00:00:00:00"

library(lubridate)
options(digits.secs=6)

#convert number of frames to partial seconds
for(h in 2:7){
fe_exact[,h]<-sapply(fe_exact[,h], function(x) {

fem.split<-unlist(strsplit(x,split=":", fixed=TRUE))
suppressWarnings(period_to_seconds(hms(paste(fem.split[1],fem.split[2],(as.numeric(fem.split[3])+as.numeric(fem.split[4])/30), sep=":"))))

}
)
}


}


##source the rest of my fuctions for predictor stats
#enable all the functions I have so far
source(here("rem_pre_analysis","rem.functions.R"), chdir = TRUE)

#make a list of the functions
fun.list<-ls()[grep("add.",ls())]
        

test.stat<-int.list


#bring in the function codes from the best result


prev.best<-read.table(paste(here(),"/cluster_optimization/Aggression_model/previous_best_remove_226_283_at_the_end",sep=""), sep=",")

prev.best.top<-as.character(prev.best[nrow(prev.best),2])
args.all=as.numeric(unlist(strsplit(prev.best.top, split="_", fixed=TRUE)))


arg.use<-c(1:length(args.all))

for(i in 1:length(arg.use)){

test.stat<-get(fun.list[args.all[i]])(test.stat)	

}


test.mod<-try(rem(evlist, test.stat, supplist=supplist, timing="interval", estimator="BPM"),silent=T)
summary(test.mod)


## Save the coefficients if I want

#setwd(paste(here(),"/adequacy_checking/adequacy_aggression_model",sep=""))

#mod_coef<-test.mod$coef
#write.table(test.mod$coef,file="rem_model_coef",sep=",")

#######
#OR bring in coefficients from the previously fitted REM 


mod_coef<-read.table(paste(here(),"/cluster_optimization/Aggression_model/rem_model_coef",sep=""), sep=",")
#######


#now we have the whole rem statlist object for the empirical data. At each timepoint, we can compare what the model would predict with the actual realized event. 


#dummy data output vector
output<-rep(0,(8+length(event.types)))


for(day in 1:length(test.stat)){


for(step in 1:nrow(evlist[[day]])){
#stat matrix predicting the next event!
stats<-test.stat[[day]][[1]][step,,]

#the coefficient vector gets multiplied by each row and the supplist vector gets multiplied by each column
statsup<-apply(stats,2,function(x) x*supplist[[day]][step,])

#for some reason the apply function across the row margin returns a transformed matrix, so I transform it back for sanity. Use the subset of coefficents for the predictors that are actually included for this timestep
statcoef<-t(apply(statsup,1,function(x) x*mod_coef[dimnames(stats)[[2]],]))

dimnames(statcoef)[[1]]<-event.types

#get the parameter-weighted sums of stats for each event type
log.hazards<-apply(statcoef,1,function(x) sum(x))

#restrict to only the events that are possible according to the supplist. These are still log hazards
possible.haz<-log.hazards[log.hazards!=0]

possible.num<-length(possible.haz)


#account for exogenous events in the dataset
if(possible.num>0){

#proportional hazards
prop.haz<-exp(possible.haz)/sum(exp(possible.haz))

tot.haz<-exp(possible.haz)

#also rank-order the proportions
rank.haz<-prop.haz[order(prop.haz,decreasing=T)]

#did the model get it right?

predicted<-names(rank.haz[1])

if(evlist[[day]][step,1]>0){

realized<-event.types[evlist[[day]][step,1]]		

#make sure there were no realized events that the model thought were impossible
if(!(realized %in% names(possible.haz))){
	
	realized<-"impossible"
	
} else {

##given that the realized event was possible and endogenous, calculate deviance residual

if(step==1 && evlist[[day]][step,2]!=0){
	
tdiff<-evlist[[day]][step,2]

dev.res<-(-2*(log(tot.haz[realized])-sum(tot.haz)*tdiff))
	
}


if(step>1){
	
tdiff<-evlist[[day]][step,2]-evlist[[day]][step-1,2]

dev.res<-(-2*(log(tot.haz[realized])-sum(tot.haz)*tdiff))

#NOTE-- this is equivalent to the log of the product of the hazard and all the survival functions of the events that didn't happen over the preceding period:
#-2*log(tot.haz[realized]*prod(exp(-tot.haz*tdiff)))
	
}

}

} else {

predicted<-"exogenous"	
realized<-"exogenous"	
	
}

if(predicted==realized){
	
	if(realized=="exogenous"){match<-"exogenous"} else {match<-"YES"}
	
} else {match<-"NO"}

#what rank predicted hazard was the realized event?

if(realized=="exogenous" | realized=="impossible"){

realized.rank<-1
dev.res<-0

} else {

realized.rank<-which(names(rank.haz)==realized)

}

#what was the distribution of hazards predicted? Essentially, was the mean greater than the median?
library(e1071)
skew<-skewness(prop.haz)

if(skew=="NaN"){skew<-0}

#save the proportional hazards for plotting
overall.haz<-rep(0,length(event.types))
names(overall.haz)<-event.types

#overall.haz[names(prop.haz)]<-prop.haz

overall.haz[names(tot.haz)]<-tot.haz

} else {
	
realized<-"exogenous"
predicted<-"exogenous"
match<-"exogenous"
realized.rank<-1
skew<-0
possible.num<-1
overall.haz<-rep(0,length(event.types))
names(overall.haz)<-event.types	
dev.res<-0
	
}


#append to a data frame

data<-c(realized,predicted,match,realized.rank,skew,possible.num,dev.res,day,overall.haz)
names(data)[1:8]<-c("realized","predicted","match","realized.rank","skew","possible.num","dev.res","day")

output<-rbind(output,data)

}
}
#remove dummy row after

output<-output[-1,]

output<-as.matrix(output)

str(output)

output.df<-data.frame(output,stringsAsFactors=F)

#just look at the summary columns for now

output.sum<-output.df[,c(1:8)]


setwd(paste(here(),"/adequacy_checking/adequacy_aggression_model",sep=""))

write.table(output.sum,file="per_event_adequacy_data_summary",sep=",", row.names=F)
write.table(output.df,file="per_event_adequacy_data_full",sep=",", row.names=F)


####### Adequacy Analyses #########

#If run already, can skip the first part of the code and bring in the saved data

setwd(paste(here(),"/adequacy_checking/adequacy_aggression_model",sep=""))

output.df<-read.table("per_event_adequacy_data_full", header=T,sep=",",stringsAsFactors=F)
output.sum<-read.table("per_event_adequacy_data_summary", header=T,sep=",",stringsAsFactors=F)


#how well did we match the right event?
table(output.sum$match)
# exogenous        NO       YES 
      # 314      2218      2636  

2636/(2218+2636)
#0.543 of the time we get it exactly right


#make a directory to hold new plots
if(!dir.exists("./main plots")){dir.create("./main plots")}
setwd("./main plots")

#what kind of events are the ones that we do really well on?

match.tab<-table(output.sum$realized[output.sum$match=="YES"])


attacks<-sum(match.tab[grep("attack",names(match.tab))])
fo<-sum(match.tab[grep("FO",names(match.tab))])
chase<-sum(match.tab[grep("CH",names(match.tab))])
sf<-sum(match.tab[grep("SF",names(match.tab))])
dis<-sum(match.tab[grep("disengage",names(match.tab))])
copstart<-sum(match.tab[grep("cop.start",names(match.tab))])
copend<-sum(match.tab[grep("cop.end",names(match.tab))])
depart<-sum(match.tab[grep("depart",names(match.tab))])

tot.tab<-table(output.sum$realized)

attacks.tot<-sum(tot.tab[grep("attack",names(tot.tab))])
fo.tot<-sum(tot.tab[grep("FO",names(tot.tab))])
chase.tot<-sum(tot.tab[grep("CH",names(tot.tab))])
sf.tot<-sum(tot.tab[grep("SF",names(tot.tab))])
dis.tot<-sum(tot.tab[grep("disengage",names(tot.tab))])
copstart.tot<-sum(tot.tab[grep("cop.start",names(tot.tab))])
copend.tot<-sum(tot.tab[grep("cop.end",names(tot.tab))])
depart.tot<-sum(tot.tab[grep("depart",names(tot.tab))])

plot.df<-data.frame(type=c("attacks","attacks","fo","fo","chase","chase","sf","sf","dis","dis","copstart","copstart","copend","copend","depart","depart"),count=c(attacks,attacks.tot,fo,fo.tot,chase,chase.tot,sf,sf.tot,dis,dis.tot,copstart,copstart.tot,copend,copend.tot,depart,depart.tot),category=c("attacks","attacks.tot","fo","fo.tot","chase","chase.tot","sf","sf.tot","dis","dis.tot","copstart","copstart.tot","copend","copend.tot","depart","depart.tot"))


library(ggplot2)

matches<-ggplot()+geom_bar(data=plot.df,aes(x=category,y=count,fill=type),stat="identity")

ggsave("matches.pdf",plot=matches,width=12,height=7,units="in")


## Looking at the rank hazard of the realized event is probably more informative

#mean rank predicted hazard for the realized event
library(magrittr)

output.sum$realized.rank<-as.numeric(output.sum$realized.rank)

#weed out the exogenous events

mean(output.sum$realized.rank[output.sum$match!="exogenous"])

#3.4

median(output.sum$realized.rank[output.sum$match!="exogenous"])
#1

#not too bad!
output.rank<-subset(output.sum,output.sum$match!="exogenous")

rank.hist<-ggplot()+geom_histogram(data=output.rank,aes(x=realized.rank),binwidth=1)+theme_bw()

ggsave("rank_hist.pdf",plot=rank.hist)

##partition by how many events were actually possible


output.rank$possible.cat<-sapply(as.numeric(output.rank$possible.num), function(x) {
	
	if(x<=25){return("1-25")}
	
	if(25<x & x<=50){return("26-50")}
	
	if(x>50){return("51+")}
	
})

rank_hist_cat<-ggplot()+geom_histogram(data=output.rank,aes(x=realized.rank,fill=possible.cat),binwidth=1)+theme_bw()

table(output.sum$realized.rank)


ggsave("rank_hist_cat.pdf",plot=rank_hist_cat)


##### Deviance Residuals #####

#How surprised was the model at a given moment? 

#take away exogenous events
output.dev<-subset(output.sum,output.sum$match!="exogenous")

#get the vector of corresponding event names
output.dev$realized.names<-event.list[,1][output.sum$match!="exogenous"]

output.dev$realized.names<-as.character(output.dev$realized.names)

#divide into categories

output.dev$realized.cat<-sapply(output.dev$realized.names, function(x){
	
	if(grepl("attack",x)){return("attack")}
	if(grepl("disengage",x)){return("disengage")}
	if(grepl("init",x)){return("response")}
	if(grepl("cop.start",x)){return("cop start")}
	if(grepl("cop.end",x)){return("cop end")}
	if(grepl("depart",x)){return("depart")}
})

table(output.dev$realized.cat)

#get the timing for the events

output.dev$realized.time<-event.list[,2][output.sum$match!="exogenous"]

output.dev$realized.time<-as.numeric(output.dev$realized.time)
output.dev$dev.res<-as.numeric(output.dev$dev.res)

#split by day

output.dev.dat<-split(output.dev,as.numeric(output.dev$day))

#add a vertical line for females exit

femline<-event.list.dat[[13]][grep("fem.exit",event.list.dat[[13]][,1]),]$Time


devtimeplot<-ggplot(data=output.dev.dat[[13]],aes(x=output.dev.dat[[13]]$realized.time,y=dev.res))+geom_segment(aes(x=output.dev.dat[[13]]$realized.time, y=dev.res, xend=output.dev.dat[[13]]$realized.time, yend=rep(0,length(dev.res))),alpha=0.2)+geom_abline(slope=0,intercept=0)+geom_point(aes(colour=realized.cat))+theme_bw()+theme(axis.title.x = element_text(size = 16, face = "bold"),axis.text.x=element_text(size=12,face="bold"),axis.text.y=element_text(size=12,face="bold"),axis.title.y=element_text(size=16,face="bold"),legend.text=element_text(size=12),legend.title=element_text(size=12,face="bold"))+labs(x="Time (s)", y="Deviance Residual",colour="Realized\nCategory")+geom_vline(xintercept=femline,colour="red",linetype=2)

ggsave("devtimeplot.pdf",plot=devtimeplot,width=12,height=7,units="in")

#consider highlighting the interrupted copulations here

## Overall Mean DR by Category ##

devmeanplot<-ggplot(data=output.dev)+geom_abline(slope=0,intercept=0,linetype=2)+geom_boxplot(aes(x=realized.cat,y=dev.res,fill=realized.cat))+theme_classic()+theme(axis.title.x = element_text(size = 14, face = "bold"),axis.text.x=element_text(size=10,face="bold"),axis.text.y=element_text(size=10,face="bold"),axis.title.y=element_text(size=14,face="bold"),legend.text=element_text(size=12),legend.title=element_text(size=12,face="bold"))+labs(x="Time (s)", y="Deviance Residual",x="Realized Category",fill="Realized\nCategory")

ggsave("devmeanplot.pdf",plot=devmeanplot)


##### Conditional Probability of Attack Events #####


#get a dataset of just the attack events 
output.cond<-output.df[grep("attack",event.list[,1]),]


#get the conditional probability of each realized event, given that it was an attack
cond.prob<-vector()
cond.rank<-vector()
cond.poss<-vector()

for(i in 1:nrow(output.cond)){
	
cond.haz<-as.numeric(output.cond[i,paste0("X",output.cond$realized[i])])


all.haz<-output.cond[i,grep("attack",dimnames(output.cond)[[2]])]
	
tot.cond.haz<-sum(as.numeric(all.haz))

cond.prob<-append(cond.prob,cond.haz/tot.cond.haz)

#get the conditional rank	
temp.rank<-which(names(all.haz[order(as.numeric(all.haz),decreasing=T)])==paste0("X",output.cond$realized[i]))

cond.rank<-append(cond.rank,temp.rank)

#number of possible attack events
temp.poss<-length(all.haz[as.numeric(all.haz)>0])

cond.poss<-append(cond.poss,temp.poss)
	
}

output.cond$cond.prob<-cond.prob
output.cond$cond.rank<-cond.rank
output.cond$cond.poss<-cond.poss

#Mean conditional probability of the realized attack event
mean(as.numeric(output.cond$cond.prob))
#0.12

#Mean/median conditional rank of the realized attack event
mean(as.numeric(output.cond$cond.rank))
#5.5

median(output.cond$cond.rank)
#3

#usually nailing it within the top 5. Consider putting this info in an inset box. 


#Mean/median number of possible attack events for each instance of a realized attack event
mean(as.numeric(output.cond$cond.poss))
#46.25365

median(as.numeric(output.cond$cond.poss))
#42

## Look at the distributions

cond.attack.plot<-ggplot()+geom_histogram(data=output.cond,aes(x=cond.rank),binwidth=1)+theme_bw()+labs(x="Attack Event Conditional Rank", y="Count")

ggsave("cond.attack.rank.plot.pdf",plot=cond.attack.plot)

#### "Recall Plot" ####

#more informative? asks the question: what % of my realized ranks were less than or equal to rank X?

#find the max rank possible
maxrank<-max(as.numeric(output.cond$cond.poss))
medrank<-median(as.numeric(output.cond$cond.poss))


#try with mean rank possible to get a better picture
meanrank<-ceiling(mean(as.numeric(output.cond$cond.poss)))


recall<-t(sapply(1:meanrank,function(x){
	
	c(x,length(output.cond$cond.rank[output.cond$cond.rank<=x])/length(output.cond$cond.rank))
		
}))


recall<-data.frame(recall)
colnames(recall)<-c("Rank","percent")


#lines for 50% and 95%
fifty<-length(recall[,2][recall[,2]<0.5])+1
ninefive<-length(recall[,2][recall[,2]<0.95])+1
rankscale<-c(seq(0,40,10),fifty,ninefive)

#get a list of colors for the axis
rankcol<-c(rep("grey30",length(rankscale)-2),"blue","red")
rankcol<-rankcol[order(rankscale)]

rankscale<-rankscale[order(rankscale)]

recallplot<-ggplot()+geom_line(data=recall,aes(x=Rank,y=percent))+geom_vline(xintercept=fifty, linetype=2,colour="blue")+geom_vline(xintercept=ninefive,linetype=2,colour="red")+theme_bw()+theme(axis.title.x = element_text(size = 16, face = "bold"),axis.text.x=element_text(size=12,face="bold",colour=rankcol),axis.text.y=element_text(size=12,face="bold"),axis.title.y=element_text(size=16,face="bold"),panel.grid.minor=element_blank())+labs(x="Rank", y="Cumulative % Realized Attack Events")+scale_x_continuous(breaks=rankscale)


ggsave("recall.plot.pdf",plot=recallplot)


##### Plot Overall hazards thru time #####


## create sums for total hazards for receive FEs. I want a sense of how much the overall amount an event type is favored is changing

#actually make the x axis TIME


for(i in 1:30){

datvect<-apply(output.df,1,function(x){
		sum(as.numeric(x[paste0("X",event.types[endsWith(event.types,paste0(".attack.",i))])]))
	
})

datname<-paste0(i,".receive.haz")

output.df$datvect<-datvect

colnames(output.df)[ncol(output.df)]<-datname

}

#split by day

haz.df<-output.df[,c(8,(ncol(output.df)-29):ncol(output.df))]

#add time column
haz.df<-cbind(event.list$Time,haz.df)
colnames(haz.df)[1]<-"Time"


haz.dat<-split(haz.df,as.numeric(haz.df$day))

#remove columns for males that were absent that day
haz.dat.pres<-lapply(haz.dat,function(z){
	z[,apply(z,2,function(x) sum(as.numeric(x))>0)]})

#Conditional Hazard: remove rows for which attacks weren't possible 
haz.dat.cond<-lapply(haz.dat.pres,function(z){
	z[apply(z,1,function(x) sum(as.numeric(x[3:length(x)]))>0),]})


#reformat for easier plotting

hazlist<-list()

hazlist<-lapply(haz.dat.cond,function(x){

	
	templist<-list()
	
	for(i in 3:ncol(x)){
		
		vect<-x[,i]
		logvect<-sapply(x[,i], function(y) if(y>0){log(y)} else {NA})
		sqvect<-sqrt(x[,i])
		names<-rep(colnames(x)[i],length(vect))
		rows<-x[,1]
		
		
	templist[[(i-1)]]<-data.frame(vect,logvect,sqvect,names,rows)	
		
	}
	
	tempdat<-data.frame(abind(templist,along=1),stringsAsFactors=F)
	
	tempdat$vect<-as.numeric(tempdat$vect)
	tempdat$logvect<-as.numeric(tempdat$logvect)
	tempdat$sqvect<-as.numeric(tempdat$sqvect)
	tempdat$rows<-as.numeric(tempdat$rows)
	
	return(tempdat)
	
	
})


#look at day 13

#try just a snapshot of before females leave
receiveplot.dat<-hazlist[[13]][hazlist[[13]]$rows<1750,]

#receive_haz_snap<-ggplot()+geom_step(data=receiveplot.dat,aes(x=rows,y=vect,colour=names),direction="vh")
receive_sqhaz_snap<-ggplot()+geom_step(data=receiveplot.dat,aes(x=rows,y=sqvect,colour=names),direction="vh")+labs(x="Time (sec)", y="Square Root of the Total Hazard\n of Getting Attacked")

#look again at the whole thing
#receive_haz<-ggplot()+geom_step(data=hazlist[[13]],aes(x=rows,y=vect,colour=names),direction="vh")
receive_sqhaz<-ggplot()+geom_step(data=hazlist[[13]],aes(x=rows,y=sqvect,colour=names),direction="vh")+labs(x="Time (sec)", y="Square Root of the Total Hazard\n of Getting Attacked")

#ggsave("overall_receive_haz_0401.pdf",plot=receive_haz,width=10,height=7,units="in")

ggsave("overall_receive_haz_0401_sqrt.pdf",plot=receive_sqhaz,width=10,height=7,units="in")
#ggsave("overall_receive_haz_0401_snapshot.pdf",plot=receive_haz_snap,width=10,height=7,units="in")
ggsave("overall_receive_haz_0401_snapshot_sqrt.pdf",plot=receive_sqhaz_snap,width=10,height=7,units="in")


#### Male-specific Receive Hazard Plots ###

###I might want to try to look at specific pairwise combinations for receive hazard- look at the receive hazards for just one male (male 3 vs male 4?) and see how they are regulating the hazard of being attacked


## Set a color pallette to use with the following plots for day 13

library(rcartocolor)


#emulate ggplot

gg_color_hue <- function(n) {
  hues = seq(15, 375, length = n + 1)
  hcl(h = hues, l = 65, c = 100)[1:n]
}

my_palette<-gg_color_hue(ncol(haz.dat.cond[[13]])-2)


#safe colorblind palettes	

#my_palette <- c("#88CCEE", "#CC6677", "#DDCC77", "#117733", "#332288", "#AA4499","#44AA99", "#999933", "#661100")

my_palette<-carto_pal(ncol(haz.dat.cond[[13]])-2,"Safe")

#flip the order
my_palette<-my_palette[length(my_palette):1]


#make the colors more transparent, better for plotting polygons that might overlap
append.alpha<-function(col,alpha=1){
	apply(sapply(col,col2rgb)/255,2,function(x)
	rgb(x[1],x[2],x[3], alpha=alpha))
}

#grab the darker blue color for male 3's border
male3<-carto_pal(12,"Safe")[11]

#make a data frame with only columns for attacking male 3

output.three<-cbind(output.df[,8],output.df[,endsWith(colnames(output.df),"attack.3")])

#add time column
output.three<-cbind(event.list$Time,output.three)
colnames(output.three)[c(1,2)]<-c("Time","Day")


		
#split by day

haz.three<-split(output.three,as.numeric(as.character(output.three$Day)))

#remove columns for males that were absent that day
haz.three.pres<-lapply(haz.three,function(z){
	z[,apply(z,2,function(x) sum(as.numeric(x))>0)]})


#Conditional Hazard: remove rows for which attacks weren't possible 
haz.three.cond<-lapply(haz.three.pres,function(z){
	z[apply(z,1,function(x) sum(as.numeric(x[3:length(x)]))>0),]})


##add column for the total hazard of getting attacked at that timepoint
haz.three.cond<-lapply(haz.three.cond, function(z){

z<-data.frame(z)

sumhaz<-apply(z[,c(3:ncol(z))],1,function(y){sum(as.numeric(y))})

z$sumhaz<-sumhaz

return(z)

})


### Plot of total hazard of getting attacked thru time for male 3 on day 13

#ggplot(data=haz.three.cond[[13]], aes(x=Time,y=sumhaz))+geom_step(direction="vh")
#nice!

#look at the square root hazard
#overall_3_haz<-ggplot()+geom_step(data=haz.three.cond[[13]], aes(x=Time,y=sqrt(sumhaz)),direction="vh",colour=male3)+theme_bw()+theme(axis.title.x = element_text(size = 16, face = "bold"),axis.text.x=element_text(size=12,face="bold"),axis.text.y=element_text(size=12,face="bold"),axis.title.y=element_text(size=16,face="bold"),panel.grid.minor=element_blank())+labs(x="Time (sec)", y="Square Root of the Total Hazard\n of Getting Attacked")

#look at the log hazard
overall_3_haz<-ggplot()+geom_step(data=haz.three.cond[[13]], aes(x=Time,y=log(sumhaz)),direction="vh",colour=male3)+theme_bw()+theme(axis.title.x = element_text(size = 16, face = "bold"),axis.text.x=element_text(size=12,face="bold"),axis.text.y=element_text(size=12,face="bold"),axis.title.y=element_text(size=16,face="bold"),panel.grid.minor=element_blank())+labs(x="Time (sec)", y="Log Total Hazard\n of Getting Attacked")
#version with colored fill under the line?

df<-haz.three.cond[[13]]

df$sqhaz<-log(df$sumhaz)

df1 <- rbind(
  df,
  transform(df[order(df$Time),],
    Time=Time + 1e-9,  # required to avoid crazy steps
    sqhaz=ave(sqhaz, FUN=function(z) c(tail(z, -1L),z[[length(z)]]))
) )


alcol<-append.alpha(my_palette[length(my_palette)],alpha=0.5)

#the area plot has to go in underneath the step line

overall_3_haz<-ggplot()+geom_ribbon(data=df1,aes(x=Time,ymin=-9.5,ymax=sqhaz),fill=alcol)+geom_step(data=haz.three.cond[[13]], aes(x=Time,y=log(sumhaz)),direction="vh",colour=male3)+theme_bw()+theme(axis.title.x = element_text(size = 16, face = "bold"),axis.text.x=element_text(size=12,face="bold"),axis.text.y=element_text(size=12,face="bold"),axis.title.y=element_text(size=16,face="bold"),panel.grid.minor=element_blank())+labs(x="Time (sec)", y="Log Total Hazard\n of Getting Attacked")


#Plots of receive hazard by other male ID 

hazlist<-list()

hazlist<-lapply(haz.three.cond,function(x){

	
	templist<-list()
	
	for(i in 3:(ncol(x)-1)){
		
		vect<-x[,i]
		percent<-as.numeric(x[,i])/x[,ncol(x)]
		names<-rep(colnames(x)[i],length(vect))
		rows<-x[,1]
		sumhaz<-x[,ncol(x)]
		
	templist[[(i-1)]]<-data.frame(vect,percent,names,rows,sumhaz)	
		
	}
	
	tempdat<-data.frame(abind(templist,along=1),stringsAsFactors=F)
	
	tempdat$vect<-as.numeric(tempdat$vect)
	tempdat$percent<-as.numeric(tempdat$percent)
	tempdat$rows<-as.numeric(tempdat$rows)
	tempdat$sumhaz<-as.numeric(tempdat$sumhaz)
	
	#create a % of sqrt(sum) column
	tempdat$persqsum<-tempdat$percent*sqrt(tempdat$sumhaz) 
	
	#reorder factors for names by initial % value
	tempdat$names<-factor(tempdat$names, levels=unique(tempdat$names)[order(tempdat[tempdat$rows==0,]$percent)])
	
	
	return(tempdat)
	
	
})


#set consistent names for male ID factors for colors
#names(my_palette)<-levels(hazlist[[13]]$names)


#ggplot()+geom_step(data=hazlist[[13]],aes(x=rows,y=vect,colour=names), direction="vh")+scale_colour_manual(values=my_palette)

#ggplot()+geom_step(data=hazlist[[13]],aes(x=rows,y=persqsum,colour=names),position="stack", direction="vh")+scale_colour_manual(values=my_palette)

#ggplot()+geom_step(data=hazlist[[13]],aes(x=rows,y=percent,colour=names),position="stack", direction="vh")+scale_colour_manual(values=my_palette)


##area plot version - possible to do stacked step+area?

df<-hazlist[[13]]


df2 <- rbind(
  df,
  transform(df[order(df$rows),],
    rows=rows + 1e-9,  # required to avoid crazy steps
    percent=ave(percent, names, FUN=function(z) c(tail(z, -1L),z[[length(z)]]))
) )

head(df2)

#make a vector of better factor level names
factmales<-unlist(sapply(strsplit(levels(df$names),split=".",fixed=T),function(x) {
	
	strsplit(x[1],split="X",fixed=T)[[1]][2]
	
}
))

#make a version for male 3 attack haz
factmales3<-factmales

names(factmales3)<-levels(df$names)

#add factor level for male number 3
factmales3<-c(factmales3,"3")

df2$names<-factor(factmales3[as.character(df2$names)],levels=factmales)

names(my_palette)<-factmales3


##percent hazard from each other male
percent.receive.haz<-ggplot()+geom_area(data=df2,aes(x=rows,y=percent,fill=names))+theme_classic()+theme(axis.title.x = element_blank(),axis.text.x=element_text(size=12,face="bold"),axis.text.y=element_text(size=12,face="bold"),axis.title.y=element_text(size=16,face="bold"), legend.text=element_text(size=12))+labs(y="Proportion of\nMale 3's Attack Hazard")+scale_fill_manual(name="Potential\nAttacker ID",labels=c(factmales),values=my_palette)


##as a portion of the sqrt(total hazard)? less informative?

df<-hazlist[[13]]


df3 <- rbind(
  df,
  transform(df[order(df$rows),],
    rows=rows + 1e-9,  # required to avoid crazy steps
    persqsum=ave(persqsum, names, FUN=function(z) c(tail(z, -1L),z[[length(z)]]))
) )

head(df3)

portion.receive.haz<-ggplot()+geom_area(data=df3,aes(x=rows,y=persqsum,fill=names,colour=names))


## put together in a combined plot?

library(ggpubr)
library(cowplot)

#add vertical lines for copulation events?

three.coptimes<-event.list.dat[[13]][startsWith(as.character(event.list.dat[[13]][,1]),"3.cop.start"),]$Time

percent.receive.haz.cop<-percent.receive.haz+geom_vline(xintercept=three.coptimes,linetype=2,colour="black", size=0.5)

overall_3_haz_cop<-overall_3_haz+geom_vline(xintercept=three.coptimes,linetype=2,colour="black", size=0.5)

#put the vert lines under the plot
overall_3_haz_cop$layers<-overall_3_haz_cop$layers[c(3,1,2)]

three_receive_haz_plot<-plot_grid(percent.receive.haz.cop,overall_3_haz_cop,nrow=2,ncol=1,align="v",axis="lr", rel_heights=c(1,1))

ggsave("three_receive_haz_plot.pdf",plot=three_receive_haz_plot,width=20,height=7,units="in")


## Layer on the same for Male 4?


#make a data frame with only columns for attacking male 3

output.four<-cbind(output.df[,8],output.df[,endsWith(colnames(output.df),"attack.4")])

#add time column
output.four<-cbind(event.list$Time,output.four)
colnames(output.four)[c(1,2)]<-c("Time","Day")


		
#split by day

haz.four<-split(output.four,as.numeric(as.character(output.four$Day)))

#remove columns for males that were absent that day
haz.four.pres<-lapply(haz.four,function(z){
	z[,apply(z,2,function(x) sum(as.numeric(x))>0)]})


#Conditional Hazard: remove rows for which attacks weren't possible 
haz.four.cond<-lapply(haz.four.pres,function(z){
	z[apply(z,1,function(x) sum(as.numeric(x[3:length(x)]))>0),]})


##add column for the total hazard of getting attacked at that timepoint
haz.four.cond<-lapply(haz.four.cond, function(z){

z<-data.frame(z)

sumhaz<-apply(z[,c(3:ncol(z))],1,function(y){sum(as.numeric(y))})

z$sumhaz<-sumhaz

return(z)

})


#use a subset of the data up until male 3 departs

haz.four.cond[[13]]<-haz.four.cond[[13]][haz.four.cond[[13]]$Time<=max(haz.three.cond[[13]]$Time),]

#4 is engaged in a FO during that interval, so add an extra row to bring the line out to the end

haz.four.cond[[13]]<-rbind(haz.four.cond[[13]],haz.four.cond[[13]][nrow(haz.four.cond[[13]]),])

haz.four.cond[[13]][nrow(haz.four.cond[[13]]),]$Time<-max(haz.three.cond[[13]]$Time)

### Plot of total hazard of getting attacked thru time for male 4 on day 13

ggplot(data=haz.four.cond[[13]], aes(x=Time,y=sumhaz))+geom_step(direction="vh")
#nice!

#look at the square root hazard
#overall_4_haz<-ggplot()+geom_step(data=haz.four.cond[[13]], aes(x=Time,y=sqrt(sumhaz)),direction="vh",colour=my_palette[length(my_palette)-1])+theme_bw()+theme(axis.title.x = element_text(size = 16, face = "bold"),axis.text.x=element_text(size=12,face="bold"),axis.text.y=element_text(size=12,face="bold"),axis.title.y=element_text(size=16,face="bold"),panel.grid.minor=element_blank())+labs(x="Time (sec)", y="Square Root of the Total Hazard\n of Getting Attacked")


#look at the log hazard
overall_4_haz<-ggplot()+geom_step(data=haz.four.cond[[13]], aes(x=Time,y=log(sumhaz)),direction="vh",colour=my_palette[length(my_palette)-1])+theme_bw()+theme(axis.title.x = element_text(size = 16, face = "bold"),axis.text.x=element_text(size=12,face="bold"),axis.text.y=element_text(size=12,face="bold"),axis.title.y=element_text(size=16,face="bold"),panel.grid.minor=element_blank())+labs(x="Time (sec)", y="Log Total Hazard\n of Getting Attacked")


#version with colored fill under the line?

df<-haz.four.cond[[13]]

df$sqhaz<-log(df$sumhaz)

df1.4 <- rbind(
  df,
  transform(df[order(df$Time),],
    Time=Time + 1e-9,  # required to avoid crazy steps
    sqhaz=ave(sqhaz, FUN=function(z) c(tail(z, -1L),z[[length(z)]]))
) )


alcol4<-append.alpha(my_palette[length(my_palette)-1],alpha=0.6)

#have to plot the area plot first

overall_4_haz<-ggplot()+geom_ribbon(data=df1.4,aes(x=Time,ymax=sqhaz,ymin=-9.5),fill=alcol4)+geom_step(data=haz.four.cond[[13]], aes(x=Time,y=log(sumhaz)),direction="vh",colour=my_palette[length(my_palette)-1])+theme_bw()+theme(axis.title.x = element_text(size = 16, face = "bold"),axis.text.x=element_text(size=12,face="bold"),axis.text.y=element_text(size=12,face="bold"),axis.title.y=element_text(size=16,face="bold"),panel.grid.minor=element_blank())+labs(x="Time (sec)", y="Log Total Hazard\n of Getting Attacked")


#layer this onto the plot for male 3?

#combine in a way that allows me to automatically produce a legend

id3<-rep("Male 3",nrow(haz.three.cond[[13]]))
id4<-rep("Male 4",nrow(haz.four.cond[[13]]))

overdat<-rbind(haz.three.cond[[13]][,c(1,ncol(haz.three.cond[[13]]))],haz.four.cond[[13]][,c(1,ncol(haz.four.cond[[13]]))])

overdat$male<-factor(c(id3,id4),levels=c("Male 4","Male 3"))

overcol<-c(my_palette[length(my_palette)-1],male3)
names(overcol)<-levels(overdat$male)

overall_3_4_combined<-ggplot()+geom_ribbon(data=df1.4,aes(x=Time,ymax=sqhaz,ymin=-9.5),fill=alcol4)+geom_ribbon(data=df1,aes(x=Time,ymin=-9.5,ymax=sqhaz),fill=alcol)+geom_step(data=overdat, aes(x=Time,y=log(sumhaz),colour=male),direction="vh")+scale_color_manual(values=overcol)+theme_bw()+theme(axis.title.x = element_text(size = 16, face = "bold"),axis.text.x=element_text(size=12,face="bold"),axis.text.y=element_text(size=12,face="bold"),axis.title.y=element_text(size=16,face="bold"),panel.grid.minor=element_blank(),legend.text=element_text(size=12),legend.title=element_text(face="bold"))+labs(x="Time (sec)", y="Log Total Hazard\n of Getting Attacked",colour="Male ID")+guides(colour=guide_legend(reverse=T))


#Plots of receive hazard by other male ID 

hazlist<-list()

hazlist<-lapply(haz.four.cond,function(x){

	
	templist<-list()
	
	for(i in 3:(ncol(x)-1)){
		
		vect<-x[,i]
		percent<-as.numeric(x[,i])/x[,ncol(x)]
		names<-rep(colnames(x)[i],length(vect))
		rows<-x[,1]
		sumhaz<-x[,ncol(x)]
		
	templist[[(i-1)]]<-data.frame(vect,percent,names,rows,sumhaz)	
		
	}
	
	tempdat<-data.frame(abind(templist,along=1),stringsAsFactors=F)
	
	tempdat$vect<-as.numeric(tempdat$vect)
	tempdat$percent<-as.numeric(tempdat$percent)
	tempdat$rows<-as.numeric(tempdat$rows)
	tempdat$sumhaz<-as.numeric(tempdat$sumhaz)
	
	#create a % of sqrt(sum) column
	tempdat$persqsum<-tempdat$percent*sqrt(tempdat$sumhaz) 
	
	#reorder factors for names by initial % value
	tempdat$names<-factor(tempdat$names, levels=c(unique(tempdat$names)[order(tempdat[tempdat$rows==0,]$percent)]))
	
	
	return(tempdat)
	
	
})


#set consistent names for male ID factors for colors
names(my_palette)<-levels(hazlist[[13]]$names)


#ggplot()+geom_step(data=hazlist[[13]],aes(x=rows,y=vect,colour=names), direction="vh")+scale_colour_manual(values=my_palette)

#ggplot()+geom_step(data=hazlist[[13]],aes(x=rows,y=persqsum,colour=names),position="stack", direction="vh")+scale_colour_manual(values=my_palette)

#ggplot()+geom_step(data=hazlist[[13]],aes(x=rows,y=percent,colour=names),position="stack", direction="vh")+scale_colour_manual(values=my_palette)


##area plot version - possible to do stacked step+area?

df<-hazlist[[13]]


df2.4 <- rbind(
  df,
  transform(df[order(df$rows),],
    rows=rows + 1e-9,  # required to avoid crazy steps
    percent=ave(percent, names, FUN=function(z) c(tail(z, -1L),z[[length(z)]]))
) )

head(df2.4)

#make a vector of better factor level names
factmales<-unlist(sapply(strsplit(levels(df$names),split=".",fixed=T),function(x) {
	
	strsplit(x[1],split="X",fixed=T)[[1]][2]
	
}
))

#make a version for male 4 attack haz
factmales4<-factmales

names(factmales4)<-levels(df$names)

#add factor level for male number 4
factmales4<-c(factmales4,"4")

#BUT use the same ordering as above


factmales4<-factmales4[match(factmales3,factmales4)]

df2.4$names<-factor(factmales4[as.character(df2.4$names)],levels=factmales4)

names(my_palette)<-factmales4


##percent hazard from each other male
percent.receive.haz.four<-ggplot()+geom_area(data=df2.4,aes(x=rows,y=percent,fill=names))+theme_classic()+theme(axis.title.x = element_blank(),axis.text.x=element_text(size=12,face="bold"),axis.text.y=element_text(size=12,face="bold"),axis.title.y=element_text(size=16,face="bold"), legend.text=element_text(size=12),legend.title=element_text(face="bold"))+labs(y="Proportion of\nMale 4's Attack Hazard")+scale_fill_manual(name="Potential\nAttacker ID",labels=c(factmales4),values=my_palette)


## Put it all together!


library(ggpubr)
library(cowplot)

#add vertical lines for copulation events?

three.coptimes<-event.list.dat[[13]][startsWith(as.character(event.list.dat[[13]][,1]),"3.cop.start"),]$Time

four.coptimes<-event.list.dat[[13]][startsWith(as.character(event.list.dat[[13]][,1]),"4.cop.start"),]$Time

#vert line for females exit?

femtime<-event.list.dat[[13]][startsWith(as.character(event.list.dat[[13]][,1]),"fem.exit"),]$Time

alcol.perc<-append.alpha(alcol,alpha=0.3)
alcol4.perc<-append.alpha(alcol4,alpha=0.3)

percent.receive.haz.cop<-percent.receive.haz+geom_vline(xintercept=three.coptimes,linetype=2,colour="black", size=0.5)+theme(panel.background=element_rect(fill=alcol.perc))

overall_3_4_combined_cop<-overall_3_4_combined+geom_vline(xintercept=three.coptimes,linetype=2,colour="black", size=0.5)+geom_vline(xintercept=four.coptimes,linetype=2,colour="dark red", size=0.5)+geom_vline(xintercept=femtime,linetype=1,colour="purple", size=0.4)

#layer the cop vertlines underneath
overall_3_4_combined_cop$layers<-overall_3_4_combined_cop$layers[c(4,5,1,2,3,6)]

percent.receive.haz.four.cop<-percent.receive.haz.four+geom_vline(xintercept=four.coptimes,linetype=2,colour="dark red", size=0.5)+theme(panel.background=element_rect(fill=alcol4.perc))

combined_receive_haz_plot<-plot_grid(percent.receive.haz.cop,overall_3_4_combined_cop,percent.receive.haz.four.cop,nrow=3,ncol=1,align="v",axis="lr", rel_heights=c(0.75,1,0.75))

ggsave("combined_receive_haz_plot.pdf",plot=combined_receive_haz_plot,width=20,height=10.5,units="in")
```
