## Supplemental_reproducible_code for "Fighting isn’t sexy in lekking Greater Sage-grouse (*Centrocercus urophasianus*)": availability_interruption_plots_terr_id_new.R

```
## Code for producing plots of individual males' hazards of being attacked and being solicited through time ##

rm(list=ls())

###make sure necessary packages are there and loaded

packages<-c("here","abind","relevent","lubridate","DescTools","gtools","e1071","ggplot2","magrittr","rcartocolor","ggpubr","cowplot","dplyr","tidyverse","grid","gtable","gridExtra")

package.check<-lapply(packages, FUN = function(x) {
    if (!require(x, character.only = TRUE, quietly=TRUE)) {
        install.packages(x, dependencies = TRUE)
        library(x, character.only = TRUE, quietly=TRUE)
    } 

})

#for project path management
#define the project root directory relative to the current script. This sets the working directory that gets called by the here() function. It is a relative path and will be robust to use on anyone's computer. In this case, I want the root directory to always be the enclosing folder for the whole sage grouse project


i_am("adequacy_checking/adequacy_attractiveness_model/availability_interruption_plots_terr_id_new.R")


#make the base event list and intercept array


event.list<-read.table(file=here("data_processing","relevent_format_dataset_udata_wincol_disturbance.tsv"), header=T,  sep="\t")

event.list$Date<-as.factor(event.list$Date)
  
event.types<-read.table(file=here("data_processing","event_types_list_udata.csv"), header=F,  sep=",")
event.types<-as.character(event.types$V1)


#NOTE- the event.types vector now contains ENDOGENOUS event types as well as EXOGENOUS types: arrivals and clock events, and end.obs. Need to make sure everything is robust to this. 


### create eventlist for rem ###


event.list.dat<-split(event.list, event.list$Date)


#remove female context for now

basic.event.list<-event.list[-3]

#strip the "loser" from the disengage events so they are all the same event "type" and they match my previous code

basic.event.list$Events<-sapply(basic.event.list$Events, function(x) {
	
	event<-unlist(strsplit(as.character(x),".",fixed=T))
	if(event[2]=="disengage"){
		return(paste(event[1],event[2],sep="."))
	} else {return(as.character(x))}
}
)


#add factor levels for event types that don't actually happen (and put them in the same order as the event.type list) to maintain bookkeeping 

basic.event.list$Events<-as.character(basic.event.list$Events)

basic.event.list$Events<-factor(basic.event.list$Events, levels=as.character(event.types))

#turn event types into numeric values

basic.event.list$Events<-as.numeric(basic.event.list$Events)

#set the event type code for exogenous events to zero. I have there all at the trailing end of the vector

endogenous<-(length(event.types)-length(event.types[grep("arrive",event.types)])-5)

basic.event.list$Events[basic.event.list$Events>endogenous]<-0

rownames(basic.event.list)<-NULL
#need to divide the data up by grouping factor

evlist<-list()
evlist<-split(basic.event.list, basic.event.list$Date)

#strip the date column from the sub-dataframes so they're just two columns, event and time

evlist<-lapply(evlist, function(x) x<-x[-3])

#convert from dataframes to matrices

evlist<-lapply(evlist, function(x) x<-as.matrix(x))

## make event.types vector usable for intlist, etc. by excising exogenous event types

event.types<-event.types[1:endogenous]


####### Bring in hazards from the REM model ######


setwd(paste(here(),"/adequacy_checking/adequacy_attractiveness_model",sep=""))

#bring in the saved data
output.df<-read.table("per_event_adequacy_data_full", header=T,sep=",",stringsAsFactors=F)
output.sum<-read.table("per_event_adequacy_data_summary", header=T,sep=",",stringsAsFactors=F)


### Availability Plot ###

#want intervals for male presence, and then for fighting/not fighting. Can get both of these numbers from the hazards 

#I want the hazard of getting attacked is greater than zero, Since males CAN be attacked during copulation but not during a fight.  BUT*** I need to also remove times when females are absent, since this doesn't cover that. 

#dim(output.df)


#split by day

#add time, Females, and Events columns
output.time<-cbind(event.list[,c(1,2,3)],output.df)
colnames(output.time)[c(1,2,3)]<-c("Events","Time","Females")


#remove rows for "init"s since the hazard for everything else is zero at that point and it messes up the intervals- makes it seem like the male is available to mate while he is just getting started fighting. 


output.time<-output.time[-grep("init",event.list$Events),]


library(dplyr)


avail.df<-split(output.time,as.numeric(as.character(output.time$day)))

#remove columns for males that were absent that day, and other columns we don't need right now
avail.df.pres<-lapply(avail.df,function(z){
	cbind(z[,c(1,2,3)],z[,c(12:ncol(z))][,apply(z[,c(12:ncol(z))],2,function(x) sum(as.numeric(x))>0)])})

library(abind)

avail.df.interval<-lapply(avail.df.pres,function(z){
	
	availlist<-list()
	
	for(i in 1:30){
	
	copcol<-paste0("X",i,".cop.start")
	cols<-colnames(z)[endsWith(colnames(z),paste0("attack.",i))]
	
	#make sure that male was there
	if(copcol %in% dimnames(z)[[2]]){
		
		newavail<-z[,cols]
		
		newavail<-apply(newavail,1,function(x) sum(x))
		
		intervail<-newavail

#if the hazard of being attacked is POSITIVE AND females are PRESENT, then they are available to mate in that interval

#maybe just label each timepoint as available or not and then pare down to the transitions using lag or lead

intervail[lead(newavail)>0 & z$Females==2]<-"avail"
intervail[lead(newavail)==0 | (lead(newavail)>0 & z$Females==1)]<-"not"

intervail[length(newavail)]<-"not"

#check
#cbind(cbind(cbind(intervail,newavail),z$Females),as.character(z$Events))

#then identify the actual beginnings and ends of the intervals
intervail.ends<-intervail

intervail.ends[intervail.ends==lag(intervail.ends)]<-"drop"

#since lag doesn't touch the first row
if(intervail.ends[1]=="not"){intervail.ends[1]<-"drop"}

#cbind(cbind(cbind(cbind(intervail.ends,intervail),newavail),z$Females),as.character(z$Events))
		
		lab<-rep(i,length(newavail))
		time<-z[,2]
				
	tempdat<-data.frame(avail=intervail,name=lab,time=time)
	
	#remove rows between interval termini
	tempdat<-tempdat[intervail.ends=="avail" | intervail.ends=="not",]	
	
	#turn into avail and not columns
	
	
	availlist[[i]]<-tempdat
		
	}
		
	}
	
	availday<-abind(availlist,along=1)
	
	availday<-data.frame(availday,stringsAsFactors=F)
	availday$time<-as.numeric(availday$time)
	
	return(availday)
})


#str(avail.df.interval[[13]])


availdat<-lapply(avail.df.interval, function(x){

mutdat<-x[x$avail=="avail",][,c(2,3)]

#match with interval ends
mutdat$not<-x[x$avail=="not",][,3]

colnames(mutdat)<-c("male","avail","not")
mutdat$male<-factor(mutdat$male,levels=c(1:30))

#restrict to territory-holders?
mutdat<-mutdat[(mutdat$male==1 | mutdat$male==2 | mutdat$male==3 | mutdat$male==4 | mutdat$male==7),]

return(mutdat)

})


library(ggplot2)
#availplot<-ggplot()+geom_segment(data=availdat[[13]],aes(x=avail,xend=not,y=0.0025,yend=0.0025),size=25,colour="grey70")+theme_classic()+coord_cartesian(ylim=c(0,0.005))+facet_wrap(.~male, nrow=5)+theme(strip.background=element_blank(),strip.text=element_blank())

#now layer on the .sol rates thru time- ridgeplot? violinplot?

#I want the rates for sols but the probabilities for interruptions, I think. That way I can calculate an "expected mating success": solrate*availability*(1-interruption_prob). Of course, one's history of copulations influences the future probability of copulations, so it's a little weird to take the average... this is just for visualizing what's going on. I could zoom in more and layer on all kinds of things that might matter - e.g. values of various coefficients that influence attack hazard

 

#exp(sum(coefs)) is the rate. So I want this number for each applicable timepoint. This is the raw hazard that I have here already, so that's good!


#Do I just want to remove rows that have cop.start rate==0? Then make a step function?


solrate.df<-list()

for(j in 1:length(avail.df.pres)){
	
	z<-avail.df.pres[[j]]
	
	availlist<-list()
	
	for(i in c(1,2,3,4,7)){
	
	copcol<-paste0("X",i,".cop.start")
	
	
	#make sure that male was there
	if(copcol %in% dimnames(z)[[2]]){
		
		newavail<-z[,copcol]
		
		#get the times associated with the FOLLOWING timestep, since the hazard in the current timestep describes the hazard of that event happening over the period leading up to the current time, but for the plot we want to see it starting at the beginning of the period over which it applies 
		
		rate<-lead(newavail)
		rate[length(rate)]<-rate[length(rate)-1]
		
		time<-z$Time
		
		#rate<-newavail[newavail>0]
		lab<-rep(i,length(rate))
		#time<-lag(z$Time)[newavail>0]
		
		#try to remove rows for which the male is copulating and that's why his hazard for cop.start is zero
			
	tempdat<-data.frame(rate=rate,male=lab,time=time)
		
		#want to locate rows where cop.start hazard is zero but that male is otherwise available
		
		intervals<-availdat[[j]][availdat[[j]]$male==i,c(2,3)]
		
		if(dim(intervals)[[1]]==1){intervals<-intervals[1,]} else {intervals<-as.matrix(intervals)}
		
		#test if any of the rows that have cop.start=0 occur in an available interval
		inside<-sapply(tempdat$time[which(tempdat$rate==0)],function(x) {
			
			if("TRUE" %in% (x %[)% intervals)){return(TRUE)} else {return(FALSE)}
			
			}
			)
		
		if(length(which(tempdat$rate==0)[inside])>0){
		
		#seems like I might not actually want to remove these, but just replace them with the previous non-zero rate. This way everything will line up with the interruption rates. 
		insiders<-which(tempdat$rate==0)[inside]
		
		for(m in insiders){
		
		#get the last non-zero rate before the missing rate 
		tempdat[m,1]<-tail(tempdat[1:(m-1),1][tempdat[1:(m-1),1]>0],1)
		
		}
		}
		
		
	availlist[[i]]<-tempdat
		
	}
		
	}
	
	availday<-abind(availlist,along=1)
	
	availday<-data.frame(availday,stringsAsFactors=F)
	availday$time<-as.numeric(availday$time)
	
	solrate.df[[j]]<-availday

}


###Just use one of the days for now

soldat<-solrate.df[[13]]
soldat$male<-factor(soldat$male,levels=c(1:30))

#for day 13, females depart at 4800, so pare down the data to that section for the plot

soldat<-soldat[!soldat$time>4800,]


#head(soldat)

#library(ggplot2)
#ggplot()+geom_step(data=soldat,aes(x=time,y=rate,colour=male),direction="hv")+theme_bw()

#ggplot()+geom_segment(data=availdat[[13]],aes(x=avail,xend=not,y=0,yend=0),size=55,colour="grey85")+geom_step(data=soldat,aes(x=time,y=rate,colour=male),direction="hv")+theme_classic()+coord_cartesian(ylim=c(0,0.005))+facet_wrap(.~male, nrow=5)+theme(strip.background=element_blank(),strip.text=element_blank())


#use code from other script to add shading under the step lines


#next up: write functions for calculating interruption risk

#############################################################
#############################################################


#need to build a whole intercept list. Basically I want the important copulation interruption functions to apply at all times using the coefficients from the REM. I need to just re-do the calculations for hazards using special functions that ignore the restriction that they only apply while a male is copulating. The idea is to know what the hazard of being iterrupted WOULD be IF the male were to begin copulating at that moment

#so I will again borrow from adequacy code


 {	


#### create statlist ####

##"a three dimensional array, event number by event type by statistic, or a list thereof, containing the sufficient statistics for the model"


sequences<-table(event.list$Date)

#create list of dummy intercept names

idno<-length(event.types[grep("cop.start",event.types)])

dummy.ints.attack<-paste(1:14,".attackFE", sep="")
dummy.ints.receive<-paste(1:13, ".receiveFE", sep="")

IDintsCH<-paste(1:14,"initCH",sep=".")
IDdis<-paste(1:14,"disengage",sep=".")

int.names<-c("Global.attackFE", "X.receiveFE", "X.UreceiveFE", "U.attackFE", "U.receiveFE", "U.UattackFE", dummy.ints.attack, dummy.ints.receive, "cop.startFE","2.cop.start","3.cop.start","4.cop.start","cop.endFE", "initFO.FE","ID.initFO.FE", "U.initFO.FE","initSF.FE","X.initSF.FE", "initCH.FE",  "U.initCH.FE", IDintsCH, "disengage.FE", "U.disengage.FE", IDdis, "depart.FE", "X.depart.FE")

int.list<-list()

for(i in 1:length(levels(event.list$Date))){

day.array<-array(0,dim=c(as.numeric(sequences[i]), length(event.types),length(int.names)))

#the third dimension is the stat slice.
dimnames(day.array)<-list(c(1:sequences[i]),as.character(event.types),as.character(int.names))

#add global attack effect, loops for "U" intercepts for attacks and receives, then one loop for typical intercepts

day.array[,grep("attack", event.types),1]<-1


for(h in 1:14){
day.array[,paste(h,".attack.",30, sep=""),2]<-1
}


for(h in 15:29){
day.array[,paste(h,".attack.",30, sep=""),3]<-1
}


for(h in 1:14){
	for(j in 15:29){	
day.array[,paste(j,".attack.", h, sep=""),4]<-1
}
}


for(h in 1:14){
	for(j in 15:29){
day.array[,paste(h,".attack.",j, sep=""),5]<-1
}
}

for(h in 15:29){
	for(j in 15:29){	
if(h!=j){
day.array[,paste(j,".attack.", h, sep=""),6]<-1
}
}
}

preloop<-6

#loop for dummy intercepts for attacks, loop for dummy intercepts for receives, add in cop.start and cop.end FEs, then one loop for typical intercepts

for(k in (preloop+1):(length(dummy.ints.attack)+preloop)){
	
	for(h in 1:14){
if(h!=(k-preloop)){
day.array[,paste((k-preloop),".attack.", h, sep=""),k]<-1
}
}
}

for(k in ((length(dummy.ints.attack)+preloop)+1):((length(dummy.ints.attack)+preloop)+length(dummy.ints.receive))){
	for(h in 1:14){
if(h!=(k-(length(dummy.ints.attack)+preloop))){
day.array[,paste(h,".attack.",(k-(length(dummy.ints.attack)+preloop)), sep=""),k]<-1
}
}
}

#cop.startFE
for(h in (1:idno)[-c(2,3,4)]){
day.array[,paste(h,".cop.start", sep=""),((length(dummy.ints.attack)+preloop)+length(dummy.ints.receive)+1)]<-1
}

#2.cop.start
day.array[,paste(2,".cop.start", sep=""),((length(dummy.ints.attack)+preloop)+length(dummy.ints.receive)+2)]<-1

#3.cop.start
day.array[,paste(3,".cop.start", sep=""),((length(dummy.ints.attack)+preloop)+length(dummy.ints.receive)+3)]<-1

#4.cop.start
day.array[,paste(4,".cop.start", sep=""),((length(dummy.ints.attack)+preloop)+length(dummy.ints.receive)+4)]<-1

#cop.endFE
for(h in 1:idno){
day.array[,paste(h,".cop.end", sep=""),((length(dummy.ints.attack)+preloop)+length(dummy.ints.receive)+5)]<-1
}

#initFO.FE
for(h in 1:idno){
day.array[,paste(h,".initFO", sep=""),((length(dummy.ints.attack)+preloop)+length(dummy.ints.receive)+6)]<-1
}

#ID.initFO.FE
for(h in 1:14){
day.array[,paste(h,".initFO", sep=""),((length(dummy.ints.attack)+preloop)+length(dummy.ints.receive)+7)]<-1
}

#U.initFO.FE
for(h in 15:29){
day.array[,paste(h,".initFO", sep=""),((length(dummy.ints.attack)+preloop)+length(dummy.ints.receive)+8)]<-1
}

#initSF.FE
for(h in 1:idno){
day.array[,paste(h,".initSF", sep=""),((length(dummy.ints.attack)+preloop)+length(dummy.ints.receive)+9)]<-1
}

#X.initSF.FE
day.array[,paste(30,".initSF", sep=""),((length(dummy.ints.attack)+preloop)+length(dummy.ints.receive)+10)]<-1


#initCH.FE
for(h in 1:idno){
day.array[,paste(h,".initCH", sep=""),((length(dummy.ints.attack)+preloop)+length(dummy.ints.receive)+11)]<-1
}


#U.initCH.FE
for(h in 15:29){
day.array[,paste(h,".initCH", sep=""),((length(dummy.ints.attack)+preloop)+length(dummy.ints.receive)+12)]<-1
}


#individual initCH effects for ID male
for(k in (((length(dummy.ints.attack)+preloop)+length(dummy.ints.receive)+13):((length(dummy.ints.attack)+preloop)+length(dummy.ints.receive)+13+(length(IDintsCH)-1)))){
	
	day.array[,int.names[k],k]<-1
	
}

preloop2<-((length(dummy.ints.attack)+preloop)+length(dummy.ints.receive)+13+(length(IDintsCH)-1))


#disengage.FE
for(h in 1:idno){
day.array[,paste(h,".disengage", sep=""),preloop2+1]<-1
}

#U.disengage.FE
for(h in 15:29){
day.array[,paste(h,".disengage", sep=""),preloop2+2]<-1
}

#individual disengage effect for ID males
for(k in (preloop2+3):(preloop2+3+length(IDdis)-1)){
	
	day.array[,int.names[k],k]<-1
	
}


preloop3<-(preloop2+3+length(IDdis)-1)

#depart.FE
for(h in 1:idno){
day.array[,paste(h,".depart", sep=""),preloop3+1]<-1
}

#X.depart.FE
day.array[,paste(30,".depart", sep=""),preloop3+2]<-1


#weird extra formatting to make it match the package example, seems like it becomes relevant *punintended!* if I want to fit day-specific effects

global<-list()

global[[1]]<-day.array

int.list[[i]]<-global

names(int.list[[i]])<-"global"

}


#### Create Support Matrix ####

#"should be a list withone element per history, each of which should be an event order by event type logical matrix. the ijth cell of this matrix should be TRUE if an event type of j was a possible next event given the preceding code i-1 events, and FALSE otherwise"

#need a way to ask, "what is each male doing right now" for each event in the sequence. That dictates the possibilities. 

#create a bookkeeping "census" matrix to keep track

census.list<-list()


for(j in 1:length(event.list.dat)){

census<-matrix("nothing", nrow=1,ncol=(idno), dimnames=list(1,c(1:(idno))))


##code that translates each new current event into what the males involved are doing at that time. The support matrix will be based on whatever the males were doing in the PREVIOUS timestep. This census matrix itself could be interesting to visualize. Each row of the census says what each male is doing going into the NEXT event.


for(i in 1:nrow(event.list.dat[[j]])){

#can ask if the second thing is something, and then move on to the first and third elements.

current<-census[nrow(census),]

#parse the event type. loop through test.seq rows
evname<-as.character(event.list.dat[[j]][i,1])
evsplit<-strsplit(evname, split=".", fixed=T)

#if it is an end obs, arrival, departure or clock event, it shouldn't change anything the males are doing in terms of interactions. Departures aren't exogenous, but they shouldn't change anything since they can only happen once a male is already doing nothing. 
if(evsplit[[1]][2]!="obs" & evsplit[[1]][2]!="depart" & evsplit[[1]][2]!="arrive" & evsplit[[1]][2]!="time" & evsplit[[1]][1]!="fem"){
#if the event was an attack, indicate the males involved and which was the receiver since this dictates who makes the initition decision
	if(evsplit[[1]][2]=="attack"){
		current[evsplit[[1]][1]]<-evname
		current[evsplit[[1]][3]]<-paste(evsplit[[1]][3], ".", "received", ".", evsplit[[1]][1], sep="")
			}
	#if it's an FO or SF, just indicate that that pair of males are involved in a fight now
		if(evsplit[[1]][2]=="initFO"||evsplit[[1]][2]=="initSF"){
			#need to refer back to the previous row of "census" to get the id of the other male
			
			current[evsplit[[1]][1]]<-paste(evsplit[[1]][1], ".", "fight", ".",strsplit(census[nrow(census),evsplit[[1]][1]], split=".",fixed=T)[[1]][3], sep="")
			current[[strsplit(census[nrow(census),evsplit[[1]][1]], split=".",fixed=T)[[1]][3]]]<-paste(strsplit(census[nrow(census),evsplit[[1]][1]], split=".",fixed=T)[[1]][3], ".", "fight", ".",evsplit[[1]][1], sep="")
			
			}
			
	#if it's a chase, indicate that the pair of males are involved in a fight, but there is an additional constraint that the individual DOING THE CHASING will be the disengager by definition
	if(evsplit[[1]][2]=="initCH"){
			
		current[evsplit[[1]][1]]<-paste(evsplit[[1]][1], ".", "fight", ".",strsplit(census[nrow(census),evsplit[[1]][1]], split=".",fixed=T)[[1]][3], ".chasedby",sep="")
			current[[strsplit(census[nrow(census),evsplit[[1]][1]], split=".",fixed=T)[[1]][3]]]<-paste(strsplit(census[nrow(census),evsplit[[1]][1]], split=".",fixed=T)[[1]][3], ".", "fight", ".",evsplit[[1]][1], ".chasing",sep="")

	}

#for disengaging, end the fight for the partner male as well
	if(evsplit[[1]][2]=="disengage"){
	current[evsplit[[1]][1]]<-"nothing"
	current[evsplit[[1]][3]]<-"nothing"	
	
}
 

#for copulations, end cops are only possible after start cops, but otherwise anything else is possible. EXCEPT the copulating male can't initiate an attack. If the next event involving that male ISN'T an end cop, it was interrupted

	if(evsplit[[1]][2]=="cop" && evsplit[[1]][3]=="start"){
		current[evsplit[[1]][1]]<-"copulating"
	}

	if(evsplit[[1]][2]=="cop" && evsplit[[1]][3]=="end"){
		current[evsplit[[1]][1]]<-"nothing"
	}

#if an X male is still involved in a fight at all, make sure he is still tagged as fighting (since sometimes more than one is interacting at once, but they can't depart until none are fighting)
	if((TRUE %in% grepl("fight.30",current) | TRUE %in% grepl("30.fight",current)) && !(grepl("receive", current[idno]) | grepl("attack", current[idno]))) {
		
		current[idno]<-"fight"

}

}

census<-rbind(census,current)


}


#don't want the first dummy row
census.list[[j]]<-census[-1,]

}


## Constraints to incorporate in the support matrix ##

#only males that are actually present can be involved in interactions on a given day and at a given time

#If "nothing" is happening, all "attacks", all start cops and all departs (given the male is actually present) are possible. Males cannot disengage or initiate fights without an attack happening first. End cop cannot happen without a previous start cop. end.obs is possible. (end.obs is not possible if any males are still fighting or copulating, so end.obs is ONLY possible when "nothing" is going on across the board)

#if the previous event was an "attack", the current event MUST be an initiation by the receiver, since we are assuming the attack is essentually instantaneous. 

#once initiatiated, subsequent events can only be a disengage from either male involved or any other event NOT invoving the interaction partners. Males actively fighting connot become engaged in a new fight until they disengage. 

#if the initiated fight was a chase only the male being chased can disengage, by definition

#Once a copulation starts, the only thing that male can do next is either end the copulation OR receive an attack. He cannot initiate anything. 

#copulations cannot occur when there are no females present

#a male can ONLY "depart" if "nothing" is happening

#Exceptions to these rules are for "X" males- there may be more than one unidentifiable male on the lek at a time. Right now we are dealing with this by having "X" males just be a mass actor category. 1) These males CAN do an "attack" before disengaging from an interaction. They can also be attacked. However, normal restrictions apply to the partner male. So if an X male is being chased, "disengage" becomes possible, but any "attack" (or cop.start, or depart) are still also possible 


### Build the actual support matrix ###

#for each event, there is a list of event types, with 1's for the types that are possible and 0's for types that are not. 


#code block to produce the support matrix

supplist<-list()


#bring in day-level male presence data

male.pres<-read.table(here("data_files","CHG_2014_male_presence_data_quest.csv"), sep=",", header=T)


#male IDs are already in ascending order, so I can convert them directly
colnames(male.pres)<-c("Date", 1:idno)

#bring in exact male presence data

pres.exact<-read.table(here("data_files","male_pres_exact_quest.csv"), sep=",",header=T)

	departlist<-list()
	
	for(j in 1:nrow(pres.exact)){
		
		arriverow<-pres.exact[j,startsWith(colnames(pres.exact),"arrive")]
		arriverow<-arriverow[!is.na(arriverow)]
		
		arrives<-t(sapply(arriverow, function(x) return(c(paste0(pres.exact$Num_ID[j],".arrive"),x,as.character(pres.exact$Date[j])))))
		
		departrow<-pres.exact[j,startsWith(colnames(pres.exact),"depart")]
		departrow<-departrow[!is.na(departrow)]
		
		departs<-t(sapply(departrow, function(x) return(c(paste0(pres.exact$Num_ID[j],".depart"),x,as.character(pres.exact$Date[j])))))
		
		allpres<-rbind(arrives,departs)
		
		departlist[[j]]<-allpres
		
	}

#collapse the departlist 

departevent<-abind(departlist, along=1)

departevent<-data.frame(departevent)
names(departevent)<-c("Events","Time", "Date")

departevent$Time<-as.numeric(as.character(departevent$Time))

library(abind)
arrstr<-abind(strsplit(as.character(departevent$Events),".",fixed=T),along=2)

departevent$Bird_ID<-arrstr[1,]
departevent$Type<-arrstr[2,]

departevent$Type<-as.factor(departevent$Type)
departevent$Bird_ID<-as.factor(departevent$Bird_ID)

# #split into day-specific data frames
pres.exact.dat<-split(departevent, departevent$Date)


#make a reference event list with data (most importantly female presence)
evlist.dat<-event.list.dat


for(j in 1:length(evlist)){

supplist[[j]]<-matrix(1, nrow=length(evlist[[j]][,1]), ncol=length(event.types))
colnames(supplist[[j]])<-event.types

#turn off all interactions involving males not present on that day
#make a list of all the event types that include the males that were entirely absent and remove them for all rows

supplist[[j]][,unique(unlist(lapply(grep(0,male.pres[j,2:(idno+1)]), function(x) c(event.types[startsWith(event.types, paste(x, ".", sep=""))], event.types[endsWith(event.types, paste(".", x, sep=""))]))))]<-0


#first row can be any "attack" or, technically, any "depart"
  #since now fem.enter is an actual event, cop.start is never possible as the very first thing
supplist[[j]][1,][-c(grep("attack", event.types), grep("depart", event.types))]<-0

#the exception is events involving males that will be present that day but haven't arrived yet


#basically I can make this code exactly the same as below- just want to ask who is actually there right now? 

arrsub<-subset(pres.exact.dat[[j]],Time<evlist[[j]][1,2] | Time==0 )
arrtab<-table(arrsub$Bird_ID,arrsub$Type)

arrpat<-apply(arrtab,1,function(x) sum(x))

#if the total number of previous arrives and departs is even, that male is currently absent
absent<-names(arrpat[(arrpat %% 2)==0])

if(length(absent)>0){
supplist[[j]][1,][unique(c(sapply(absent, function(x) c(event.types[startsWith(event.types, paste(x, ".", sep=""))], event.types[endsWith(event.types, paste(".", x, sep=""))]))))]<-0

} 


#loop through the rest of the events: 

for(i in 2:(length(evlist[[j]][,1])-1)){


#before anything else turn off events involving males that have either not yet arrived or have lef the lek early and are therefore no longer present that day or are otherwise not currently present


arrsub<-subset(pres.exact.dat[[j]],Time<evlist[[j]][i,2])
arrtab<-table(arrsub$Bird_ID,arrsub$Type)

arrpat<-apply(arrtab,1,function(x) sum(x))

#if the total number of previous arrives and departs is even, that male is currently absent
current.row.abs<-names(arrpat[(arrpat %% 2)==0])


if(length(current.row.abs)>0){
supplist[[j]][i,][unique(c(sapply(current.row.abs, function(x) c(event.types[startsWith(event.types, paste(x, ".", sep=""))], event.types[endsWith(event.types, paste(".", x, sep=""))]))))]<-0

} 


#determine if the previous event was an attack, since this dictates exactly what the next event can be: receiver initiates a type of fight
if((TRUE %in% grepl("received", census.list[[j]][i-1,]))==TRUE){

supplist[[j]][i,][-grep(TRUE, startsWith(event.types, paste(grep("received", census.list[[j]][i-1,]), ".", "init", sep="")))]<-0}


#(else) if the previous event wasn't an attack, the next event CAN'T be an initiation and the possibilities depend on what everyone is currently engaged in


if((TRUE %in% grepl("received", census.list[[j]][i-1,]))==FALSE){

supplist[[j]][i,][grep("init", event.types)]<-0

#if males are actively fighting, they can't attack or be attacked, or actually do anything else before disengaging (unless they are a ?? male, id 30)

if((TRUE %in% grepl("fight", census.list[[j]][i-1,]))==TRUE){
		
		for(k in grep("fight", census.list[[j]][i-1,])){
	
	if(k!=idno){
	
	supplist[[j]][i,][c(event.types[startsWith(event.types, paste(k, ".attack.", sep=""))], event.types[endsWith(event.types, paste("attack.", k, sep=""))], event.types[startsWith(event.types, paste(k, ".cop", sep=""))])]<-0
	
	}
	
}
	
	#ONLY those fighting can disengage. retain males that are fighting and remove the rest
	dis<-event.types[grep("disengage",event.types)]
	supplist[[j]][i,][dis[-grep("fight", census.list[[j]][i-1,])]]<-0
	
	#"chasing" and "chasedby" limit the possibilities further. Males getting chased DO NOT disengage by definition, again, with the exception of ?? males. This is taken care of by having ?? males just be "fight" -ing without any extra tags
	if(TRUE %in% grepl("chasedby", census.list[[j]][i-1,])){
		supplist[[j]][i,][dis[grep("chasedby", census.list[[j]][i-1,])]]<-0
	}
	
}

#if no one is fighting, no one can disengage, but all attack combinations are possible
if((TRUE %in% grepl("fight", census.list[[j]][i-1,]))==FALSE){
	
		supplist[[j]][i,][grep("disengage",event.types)]<-0
	
}


#actively copulating males: They cannot cop.start. ONLY they can cop.end. they CAN be attacked but they CANNOT attack anyone
if((TRUE %in% grepl("copulating", census.list[[j]][i-1,]))==TRUE){
	
	cop<-grep("copulating", census.list[[j]][i-1,]) 
	
	#stuff that needs to be turned off for only copulating males
	for(h in cop){

		supplist[[j]][i,][c(event.types[startsWith(event.types ,paste(h, ".attack.", sep=""))], event.types[startsWith(event.types, paste(h, ".cop.start", sep=""))])]<-0
				
	}
	
	#stuff that needs to be on for ONLY copulating males
	copendall<-event.types[grep("cop.end",event.types)]
	supplist[[j]][i,][copendall[-cop]]<-0
}

#if no one is copulating, all cop.starts are possible, but cop.ends are not

if((TRUE %in% grepl("copulating", census.list[[j]][i-1,]))==FALSE){
	
	copendall<-event.types[grep("cop.end",event.types)]
	supplist[[j]][i,][copendall]<-0
	
}


}

#if there are no females present, the next event could not have possibly been a cop.start

if(evlist.dat[[j]][i-1,]$Females==1){
	
	copstartall<-event.types[grep("cop.start",event.types)]
	supplist[[j]][i,][copstartall]<-0
	
}

#Finally, A male can only "depart" if he was already doing "nothing," so depart must be turned off for everyone doing not-nothing. (also he has to be present, but I've already taken care of that above)

noth<-as.character(which(census.list[[j]][i-1,]!="nothing"))
	
	if(length(noth)>0){
		
	supplist[[j]][i,][paste(noth, ".depart", sep="")]<-0

}

}
#end events loop here

#after done with all the middle events: onset of end.obs could depend on a lot of factors; not sure I want to fit it as part of the model. Just always say the last row must be end.obs (all endogenous events are no longer possible) 

supplist[[j]][length(evlist[[j]][,1]),]<-0

}
#end day loop here


###Make the win/loss census array

#i've adjusted the function to mark when a win or loss is the result of a chase or not
create.win.loss.census<-function(x){

#get disengagement events for focal male from evlist
win.loss.census<-x

#loop thru days and males

for(i in 1:length(win.loss.census)){

for(l in 1:idno){


dis.times<-which(startsWith(as.character(event.list.dat[[i]][,1]), paste(l,".disengage", sep="")))


if(length(dis.times)>0){

evsplit<-strsplit(as.character(event.list.dat[[i]][dis.times,1]),".",fixed=T)

#so I have a list of row numbers for disengagements and a list of interaction partners. This will cover wins for normal fights and losses for chases for the focal male. If I loop through all the males, all the wins and losses should get filled in.

#NOW DISENGAGES ALWAYS MEAN WINS, so I will still have chase information, but I don't need to do anything special with it here.

#need to make sure I am really precise since male 30 can be involved in multiple interactions, but they're usually all chases.


for(j in 1:length(dis.times)){
	
		
	if(grepl("chasing",win.loss.census[[i]][(dis.times[j]-1),l]) ==FALSE && grepl("chasedby",win.loss.census[[i]][(dis.times[j]-1),as.numeric(evsplit[[j]][3])]) ==FALSE){
	#apply the win in the focal male's column
	win.loss.census[[i]][dis.times[j],l]<-paste(l, "win",evsplit[[j]][3], sep=".")
	#apply the loss for the same interaction in the partner male's column
	win.loss.census[[i]][dis.times[j],as.numeric(evsplit[[j]][3])]<-paste(evsplit[[j]][3], "lose", l, sep=".")
	} else{
		#apply the won chase in the focal male's column
	win.loss.census[[i]][dis.times[j],l]<-paste(l, "win",evsplit[[j]][3],"chase", sep=".")
		#apply the loss for the same interaction in the partner male's column.
	win.loss.census[[i]][dis.times[j],as.numeric(evsplit[[j]][3])]<-paste(evsplit[[j]][3], "lose", l, "chase",sep=".")
		}
		
}

}

}

}

return(win.loss.census)
	
}

win.loss.census<-create.win.loss.census(census.list)


#other important stuff to load
### load in pairwise distance data
##bring in pairwise distances; I have a table for each day
##female exact presence data


file_list<-list.files(paste(here(),"/data_processing/udata_centroid_data/pairwise_dist",sep=""))

distlist<-list()

distlist<-lapply(file_list, function(x) read.table(paste(here(),"/data_processing/udata_centroid_data/pairwise_dist/",x,sep=""), sep=","))


#convert to martices

distlist.mat<-lapply(distlist, function(x) as.matrix(x, ))

#the names of the rows and columns are in the same order as the numerical ids passed to rem, so they can be used to index the matrix. Males that are absent on a given day have NA's for their centroid distances. I can change them to zeroes here so they don't produce errors, but the support array will take care of them anyway.

for(i in 1:length(distlist.mat)){
distlist.mat[[i]][is.na(distlist.mat[[i]])]<-0
}


fe_exact<-read.table(file=here("data_files","CHG_females_exact.tsv"), header=TRUE,sep="\t",stringsAsFactors=FALSE)

#reduce to only the necessary columns- this time I want time elapsed
fe_exact<-fe_exact[,c(1,3,5,7,9,11,13)]

#turn the first arrival NA's to 0s
fe_exact$first_arrival_elapsed[is.na(fe_exact$first_arrival_elapsed)]<-"00:00:00:00"

library(lubridate)
options(digits.secs=6)

#convert number of frames to partial seconds
for(h in 2:7){
fe_exact[,h]<-sapply(fe_exact[,h], function(x) {

fem.split<-unlist(strsplit(x,split=":", fixed=TRUE))
suppressWarnings(period_to_seconds(hms(paste(fem.split[1],fem.split[2],(as.numeric(fem.split[3])+as.numeric(fem.split[4])/30), sep=":"))))

}
)
}


}


##source the rest of my fuctions for predictor stats
#enable all the functions I have so far
source(here("rem_pre_analysis","rem.functions.R"), chdir = TRUE)

#make a list of the functions
fun.list<-ls()[grep("add.",ls())]
        

#replace the .cop functions with special ones that always apply


########
{
add.terr.attack.cop<-function(x){
	
rec.stat<-x

library(abind)

#loop through each day in the sequence

for(k in 1:length(rec.stat)){

#empty matrix of the right dimensions, event# x event type. In this case, the default is all zeroes. 
stat<-matrix(0, nrow=nrow(rec.stat[[k]][[1]][,,1]), ncol=length(event.types))
colnames(stat)<-event.types

#are you here today AND ALSO both of the past two days? (excluding X males)

#add attendance data from 3/17
extra<-rep(0,29)
names(extra)<-1:29
extra[c(1,2,3,4,5,7)]<-1

if(k==1 | k==2){
	
	if(k==1){
		
		terr.pres<-rbind(extra,extra)
		terr.pres<-rbind(terr.pres,male.pres[k,2:(ncol(male.pres)-1)])
		
		terr.holders<-which(apply(terr.pres,2,function(x) sum(x))==3)
		
	}
	
	if(k==2){
		
	
		terr.pres<-rbind(extra,male.pres[c(k-1,k),2:(ncol(male.pres)-1)])
		
		terr.holders<-which(apply(terr.pres,2,function(x) sum(x))==3)
		
	}
	
	
} else {
	
	terr.holders<-which(apply(male.pres[(k-2):k,2:(ncol(male.pres)-1)],2,function(x) sum(x))==3)
	
}

#this loops through each event in the sequence


if(length(terr.holders)>0){


#stat with 2 since the first event can't be a cop.start
for(j in 2:nrow(stat)){


#this gives the columns for which males were copulating, which corresponds to their id# if there are no males copulating, the loop will simply not run. ##FOR THE SPECIAL FUNCTION: REPLACE THIS WITH THE NUMBERS FOR ALL THE MALES
cop<-c(1:30) 


#for each event, loop through all currently copulating males
for(i in cop){
	
	#in case the copulating male IS a territory holder (he won't attack himself)
	
for(h in terr.holders){

if(h!=i){
#match with event types- 
stat[j,][paste0(h,".attack.",i)]<-1
}
}

}

}

}

rec.stat[[k]][[1]]<-abind(rec.stat[[k]][[1]], stat)

dimnames(rec.stat[[k]][[1]])[[3]][ncol(rec.stat[[k]][[1]][1,,])]<-"terr.attack.cop"

}

return(rec.stat)


}

	
	
#########	
	
	
add.log.receive.lose.pairwise.yest.cop<-function(x){

new.stat<-x

library(abind)


#loop through each day in the sequence, but applying 0 stat matrices to day 1 and 9 since there is no "yesterday" (March 27th data is missing due to field conditions)


for(k in 1:length(new.stat)){

#empty matrix of the right dimensions, event# x event type. In this case, the default is all zeroes. 
stat<-matrix(0, nrow=nrow(new.stat[[k]][[1]][,,1]), ncol=length(event.types))
colnames(stat)<-event.types

if(k!=1 && k!=9){
	
#this loops through each event in the sequence, in this case starting with the third event since this is the earliest possible win/loss event

for(j in 1:nrow(stat)){

#for each event, loop through all the males to get the recency ranks that will predict the next interaction- the support array will take care of males that weren't present that day.

#instead of looping through every focal male for this timestep, identify if any males were copulating in the previous timestep and loop through them. If no males are copulating, this row in stat remains all zeroes.

#this gives the columns for which males were copulating, which corresponds to their id# if there are no males copulating, the loop will simply not run. ##FOR THE SPECIAL FUNCTION: REPLACE THIS WITH THE NUMBERS FOR ALL THE MALES (except X males bacause "Pairwise" history doesn't apply to them)
cop<-c(1:29) 

		
for(i in cop){

	
#get the list of all the interactions for the focal male for yesterday

int.vector<-win.loss.census[[(k-1)]][,i]


#Remove the "fights" and "nothings" from the census list. This creates attack/receive and win/lose pairs in order of occurence
fight.times<-grep("fight",int.vector)
nothing.times<-grep("nothing",int.vector)

pairs<-int.vector[-c(fight.times, nothing.times)]


#need a way to do this that matches interaction partners but also is sensitive to the time-ordering 

receive.lose<-vector()

if(length(pairs)>0){

receive.lose<-unlist(sapply(1:ncol(census.list[[1]]), function(z){
	
	intset<-pairs[sapply(pairs, function(x) {
		
		strsplit(x,split=".",fixed=T)[[1]][3]==z
		
	})]
	
	names(intset)<-grep("TRUE",sapply(pairs, function(x) {
		
		strsplit(x,split=".",fixed=T)[[1]][3]==z
		
	}))
	
	receive<-grep("receive",intset)
	receivepluslose<-intset[receive+1][grep("lose", intset[receive+1])]
	
	return(receivepluslose)

}))
}

#if the male has not had any interactions yet, the hazard is equal across event types involving them, so set all to zero 
if(length(receive.lose)>0){

#strip the "chase" from the end of chase wins so I can include them
receive.lose<-sapply(receive.lose, function(x) if(grepl("chas",x)==TRUE){
temp<-strsplit(x, split=".", fixed=T)[[1]]	
new<-paste(temp[1],temp[2],temp[3], sep=".")
return(new)
}
else return(x)
)


#totals for each type- these will be the values for the stat
tots<-table(receive.lose)

#get just the interacting male id numbers
rankids<-sapply(names(tots), function(x) strsplit(x, split=".", fixed=T)[[1]][3])

#match with event types- this stat predicts INCOMING attacks based on total previous (yesterday's) attack+losses
#loop thru each element of rankids

for(l in 1:length(rankids)){

#incoming attacks directed at the focal male
stat[j,][paste(rankids[l],".attack.", i, sep="")]<-tots[l]
	}

		
}
	
}

}
}

#exclude attack events involving the X males 
stat[,(endsWith(event.types,paste(".attack.",ncol(census.list[[1]]), sep="")) | startsWith(event.types,paste(ncol(census.list[[1]]),".attack.", sep="")))]<-0

#take the log 
stat<-log(1+stat)

new.stat[[k]][[1]]<-abind(new.stat[[k]][[1]], stat)

dimnames(new.stat[[k]][[1]])[[3]][ncol(new.stat[[k]][[1]][1,,])]<-"log.receive.lose.pairwise.yest.cop"

}

return(new.stat)

}

	


	
########

add.receive.lose.count.pairwise.day.cop<-function(x){

new.stat<-x

library(abind)

#loop through each day sequence
for(k in 1:length(new.stat)){

#empty matrix of the right dimensions, event# x event type. In this case, the default is all zeroes. 
stat<-matrix(0, nrow=nrow(new.stat[[k]][[1]][,,1]), ncol=length(event.types))
colnames(stat)<-event.types


#this loops through each event in the sequence

for(j in 2:nrow(stat)){

#for each event, loop through all the males to get the recency ranks that will predict the next interaction- the support array will take care of males that weren't present that day.

#instead of looping through every focal male for this timestep, identify if any males were copulating in the previous timestep and loop through them. If no males are copulating, this row in stat remains all zeroes.

#this gives the columns for which males were copulating, which corresponds to their id# if there are no males copulating, the loop will simply not run. ##FOR THE SPECIAL FUNCTION: REPLACE THIS WITH THE NUMBERS FOR ALL THE MALES (except X males bacause "Pairwise" history doesn't apply to them)
cop<-c(1:29) 
		
for(i in cop){

	
#want to start the loop for the stat after the first interaction is complete, but I want to start the accounting of the past interactions from the beginning

int.vector<-win.loss.census[[k]][1:(j-1),i]


#Remove the "fights" and "nothings" from the census list. This creates attack/receive and win/lose pairs in order of occurence
fight.times<-grep("fight",int.vector)
nothing.times<-grep("nothing",int.vector)

pairs<-int.vector[-c(fight.times, nothing.times)]

#need a way to do this that matches interaction partners but also is sensitive to the time-ordering 

receive.losses<-vector()

if(length(pairs)>0){

receive.losses<-unlist(sapply(1:ncol(census.list[[1]]), function(z){
	
	intset<-pairs[sapply(pairs, function(x) {
		
		strsplit(x,split=".",fixed=T)[[1]][3]==z
		
	})]
	
	names(intset)<-grep("TRUE",sapply(pairs, function(x) {
		
		strsplit(x,split=".",fixed=T)[[1]][3]==z
		
	}))
	
	receives<-grep("receive",intset)
	receivepluslose<-intset[receives+1][grep("lose", intset[receives+1])]
	
	return(receivepluslose)
	
}))
}


#if the male has not had any interactions yet, the hazard is equal across event types involving them, so set all to zero 
if(length(receive.losses)>0){

#strip the "chase" from the end of chase wins so I can include them
receive.losses<-sapply(receive.losses, function(x) if(grepl("chas",x)==TRUE){
temp<-strsplit(x, split=".", fixed=T)[[1]]	
new<-paste(temp[1],temp[2],temp[3], sep=".")
return(new)
}
else return(x)
)


#get the number of interactions with each other male
counts<-table(receive.losses)	

#get just the interacting male id number
rankids<-sapply(names(counts), function(x) strsplit(x, split=".", fixed=T)[[1]][3])
	
#match with event types- this stat predicts new INCOMING attacks based on the number of previous OUTGOING attacks + WINS
#loop thru each element of rankids

for(l in 1:length(rankids)){

#incoming attacks directed at the focal male
stat[j,][paste(rankids[l],".attack.", i, sep="")]<-counts[l]
	}
		
}
	
}

}

#take the log 
stat<-log(1+stat)

new.stat[[k]][[1]]<-abind(new.stat[[k]][[1]], stat)

dimnames(new.stat[[k]][[1]])[[3]][ncol(new.stat[[k]][[1]][1,,])]<-"receive.lose.count.pairwise.day.cop"

}

return(new.stat)

}

}

#########


test.stat<-int.list


#bring in the function codes from the best result

prev.best<-read.table(paste(here(),"/cluster_optimization/Attractiveness_model/previous_best_remove_226_at_the_end",sep=""), sep=",")

prev.best.top<-as.character(prev.best[nrow(prev.best),2])
args.all=as.numeric(unlist(strsplit(prev.best.top, split="_", fixed=TRUE)))


arg.use<-c(1:length(args.all))

for(i in 1:length(arg.use)){

test.stat<-get(fun.list[args.all[i]])(test.stat)	

}


####


#next step is to bring in the original coefficients and then calculate attack hazards using the other bit of adequacy code. 

mod_coef<-read.table(paste(here(),"/cluster_optimization/Attractiveness_model/rem_model_coef",sep=""), sep=",")


#dummy data output vector
output<-rep(0,(1+length(event.types)))


for(day in 1:length(test.stat)){


for(step in 1:nrow(evlist[[day]])){
#stat matrix predicting the next event!
stats<-test.stat[[day]][[1]][step,,]

#the coefficient vector gets multiplied by each row and the supplist vector gets multiplied by each column
statsup<-apply(stats,2,function(x) x*supplist[[day]][step,])

#for some reason the apply function across the row margin returns a transformed matrix, so I transform it back for sanity. Use the subset of coefficents for the predictors that are acutally included for this timestep
statcoef<-t(apply(statsup,1,function(x) x*mod_coef[dimnames(stats)[[2]],]))

dimnames(statcoef)[[1]]<-event.types

#get the parameter-weighted sums of stats for each event type
log.hazards<-apply(statcoef,1,function(x) sum(x))

#restrict to only the events that are possible according to the supplist. These are still log hazards
possible.haz<-log.hazards[log.hazards!=0]

possible.num<-length(possible.haz)


#account for exogenous events in the dataset
if(possible.num>0){

tot.haz<-exp(possible.haz)

#save the proportional hazards for plotting
overall.haz<-rep(0,length(event.types))
names(overall.haz)<-event.types


overall.haz[names(tot.haz)]<-tot.haz

} else {
	
overall.haz<-rep(0,length(event.types))
names(overall.haz)<-event.types	
	
}


#append to a data frame

data<-c(day,overall.haz)
names(data)[1]<-"day"

output<-rbind(output,data)

}
}
#remove dummy row after

output<-output[-1,]

output<-as.matrix(output)

#str(output)

interout.df<-data.frame(output,stringsAsFactors=F)


##next extract the overall hazards of getting attacked for each male, which should now be re-adjusted so that it is as if they are copulating at each timestep. 


#split by day

#add time, Females, and Events columns
interout.time<-cbind(event.list[,c(1,2,3)],interout.df)
colnames(interout.time)[c(1,2,3)]<-c("Events","Time","Females")


#remove rows for "init"s since the hazard for everything else is zero at that point and it messes up the intervals


interout.time<-interout.time[-grep("init",event.list$Events),]


library(dplyr)


interday.df<-split(interout.time,as.numeric(as.character(interout.time$day)))

#remove columns for males that were absent that day
interday.df.pres<-lapply(interday.df,function(z){
	cbind(z[,c(1,2,3)],z[,c(5:ncol(z))][,apply(z[,c(5:ncol(z))],2,function(x) sum(as.numeric(x))>0)])})


#make data frames that sum up the total hazard of getting attacked for each male- very similar to what I did for the mating availability data


interrate.df<-lapply(interday.df.pres,function(z){
	
	interlist<-list()
	
	for(i in c(1,2,3,4,7)){
	
	copcol<-paste0("X",i,".cop.start")
	cols<-colnames(z)[endsWith(colnames(z),paste0("attack.",i))]
	
	#make sure that male was there
	if(copcol %in% dimnames(z)[[2]]){
		
		newavail<-z[,cols]
		
		newavail<-apply(newavail,1,function(x) sum(x))
		
		rate<-lead(newavail)
		rate[length(rate)]<-rate[length(rate)-1]
				
		lab<-rep(i,length(newavail))
		time<-z[,2]
		
		#use the hazards to get the PROBABILITY of a copulation that started at that moment being interrupted rather than ending naturally. (prob of an attack vs. a cop.end happening FIRST)
		
		#if the interruption rate is zero, that means the male is fighting, so keep it at zero because he wouldn't have been available to copulate anyway. 
		
		endrate<-exp(mod_coef["cop.endFE",])
		
		intprob<-rate/(rate+endrate)
				
	tempdat<-data.frame(interrate=rate,male=lab,time=time,intprob=intprob)
	
	#turn into avail and not columns
	
	
	interlist[[i]]<-tempdat
		
	}
		
	}
	
	availday<-abind(interlist,along=1)
	
	availday<-data.frame(availday,stringsAsFactors=F)
	availday$time<-as.numeric(availday$time)
	
	return(availday)
})


#######
##grab the example for day 13

interrate<-interrate.df[[13]]

interrate$male<-factor(interrate$male,levels=c(1:30))

#for day 13, females depart at 4800, so pare down the data to that section for the plot

interrate<-interrate[!interrate$time>4800,]


#head(interrate)

#bring over the solicitation data
interrate$solrate<-soldat$rate

#str(interrate)


#scale the interruption risk to the sol rate value? This would give a sense of the likeihood of a successful copulation at that moment. 

interrate$int.scaled<-interrate$intprob*interrate$solrate


#try plotting


#ggplot()+geom_segment(data=availdat[[13]],aes(x=avail,xend=not,y=0,yend=0),size=55,colour="grey85")+geom_step(data=interrate,aes(x=time,y=solrate,colour=male),direction="hv")+geom_step(data=interrate,aes(x=time,y=int.scaled),direction="hv",colour="red")+theme_classic()+coord_cartesian(ylim=c(0,0.003))+facet_wrap(.~male, nrow=5)+theme(strip.background=element_blank(),strip.text=element_blank())

#yay! now add in the shading?

#might have to arrange the plots differently if I want to do this- split up dataframes by male? Or can I just make sure they have the same factor levels?

df.sol<-interrate

dflist<-list()

for(i in c(1,2,3,4,7)){

df<-df.sol[df.sol$male==i,]

df.new<- rbind(
  df,
  transform(df[order(df$time),],
    time=time - 1e-9,  # required to avoid crazy steps
    solrate=ave(solrate, FUN=function(z) c(z[[1]],head(z, -1L))),
    int.scaled=ave(int.scaled, FUN=function(z) c(z[[1]],head(z, -1L)))
) )

dflist[[i]]<-df.new

}

df.sol<-dflist[[1]]

for(i in 2:length(dflist)){
	
	df.sol<-rbind(df.sol,dflist[[i]])
	
}


#put it all together


availdat.plot<-availdat[[13]]

#main<-ggplot()+geom_segment(data=availdat.plot,aes(x=avail,xend=not,y=0,yend=0),size=55,colour="grey85")+geom_step(data=interrate,aes(x=time,y=solrate,colour=male, alpha=0.9),direction="hv")+geom_ribbon(data=df.sol,aes(x=time,ymax=solrate,ymin=0,fill=male))+geom_step(data=interrate,aes(x=time,y=int.scaled),direction="hv",colour="pink")+geom_ribbon(data=df.sol,aes(x=time,ymax=int.scaled,ymin=0),fill="red")+theme_classic()+coord_cartesian(ylim=c(0,0.002))+facet_wrap(.~male, nrow=5)+theme(strip.background=element_blank(),strip.text=element_blank())

#cool! 

#add horizontal line at 0

#main<-main+geom_abline(slope=0,intercept=0,size=0.2)


#add copulations and ends/interruptions

day<-13

copevents<-event.list.dat[[day]][grep("cop.start|cop.end",event.list.dat[[day]]$Events),]

newcols<-t(sapply(copevents$Events,function(x){
	
	thing<-strsplit(as.character(x),split=".",fixed=T)

	return(c(thing[[1]][1],thing[[1]][3]))

}
))

copevents<-cbind(copevents,newcols)
colnames(copevents)[c(5,6)]<-c("male","type")

copevents$male<-factor(copevents$male,levels=c(1:30))


#interruptions are a little trickier


#want the row right after the last row they are copulating- grab any rows where they were attacked rather than copulation simply continuing or ending naturally
interlist<-apply(win.loss.census[[day]][1:(nrow(win.loss.census[[day]])-2),],2,function(x) (grep("copulating",x)+1)[grep("received",x[grep("copulating",x)+1])])


intevents<-event.list.dat[[day]][unlist(interlist),]

newcols<-t(sapply(intevents$Events,function(x){
	
	thing<-strsplit(as.character(x),split=".",fixed=T)

	return(c(thing[[1]][1],thing[[1]][3]))

}
))

intevents<-cbind(intevents,newcols)
colnames(intevents)[c(5,6)]<-c("perp","male")

intevents$male<-factor(intevents$male,levels=c(1:30))
intevents$perp<-factor(intevents$perp,levels=c(1:30))

#try including only cop.ends or interruptions for clarity. put them in the same dataframe so I can build I legend with the linetypes?

copends<-copevents[copevents$type=="end",]

#main<-ggplot()+geom_segment(data=availdat.plot,aes(x=avail,xend=not,y=0,yend=0),size=75,colour="grey85")+geom_step(data=interrate,aes(x=time,y=solrate,colour=male),direction="hv")+geom_ribbon(data=df.sol,aes(x=time,ymax=solrate,ymin=0,fill=male))+geom_step(data=interrate,aes(x=time,y=int.scaled),direction="hv",colour="pink")+geom_ribbon(data=df.sol,aes(x=time,ymax=int.scaled,ymin=0),fill="red")+theme_classic()+coord_cartesian(ylim=c(0,0.003))+geom_segment(data=copends,aes(x=Time,xend=Time,y=0,yend=0.003),size=0.8)+geom_segment(data=intevents,aes(x=Time,xend=Time,y=0,yend=0.003,colour=perp),size=0.8,linetype=5)+geom_abline(slope=0,intercept=0,size=0.2)+facet_wrap(.~male, nrow=5)+theme(strip.background=element_blank(),strip.text=element_blank())


##try restricting to just the top 3 males

#str(availdat.plot)
#str(interrate)
#str(df.sol)
#str(copends)
#str(intevents)

remove.males<-function(x){
	
	x<-filter(x,male=="2" | male=="3" | male=="4")
	return(x)
}

datlist<-list(availdat.plot,interrate,df.sol,copends,intevents)

datlist.short<-lapply(datlist,function(x) remove.males(x))


#define a color function so the colors don't get messed up
gg_color_hue<- function(n){
	hues=seq(15,375,length=n+1)
	hcl(h=hues,l=65,c=100)[1:n]
}

#set the colors for each male
mycolors<-gg_color_hue(5)
names(mycolors)<-levels(availdat.plot$male)[c(1:4,7)]

#main_small<-ggplot()+geom_segment(data=datlist.short[[1]],aes(x=avail,xend=not,y=0,yend=0),size=125,colour="grey85")+geom_step(data=datlist.short[[2]],aes(x=time,y=solrate,colour=male),direction="hv")+geom_ribbon(data=datlist.short[[3]],aes(x=time,ymax=solrate,ymin=0,fill=male))+scale_fill_manual(values=mycolors,guide=F)+geom_step(data=datlist.short[[2]],aes(x=time,y=int.scaled),direction="hv",colour="pink")+geom_ribbon(data=datlist.short[[3]],aes(x=time,ymax=int.scaled,ymin=0),fill="red")+theme_classic()+coord_cartesian(ylim=c(0,0.003))+geom_segment(data=datlist.short[[4]],aes(x=Time,xend=Time,y=0,yend=0.003,colour="black",linetype="black"),size=0.8)+geom_segment(data=datlist.short[[5]],aes(x=Time,xend=Time,y=0,yend=0.003,colour=perp,linetype=perp),size=0.8)+scale_colour_manual(name="Copulation\nEvents",values=c(mycolors,"black"),labels=c("Int. by 1","Int. by 2","Int. by 3","Int. by 4","Int. by 7","Uninterr."))+scale_linetype_manual(name="Copulation\nEvents",values=c(5,5,5,5,5,1),labels=c("Int. by 1","Int. by 2","Int. by 3","Int. by 4","Int. by 7","Uninterr."))+geom_abline(slope=0,intercept=0,size=0.2)+facet_wrap(.~male, nrow=5)+theme(strip.background=element_blank(),strip.text=element_blank(),axis.title.x=element_text(size=20,face="bold"),axis.title.y=element_text(size=20,face="bold"),axis.text.x=element_text(size=18,face="bold"),axis.text.y=element_text(size=18,face="bold"),legend.text=element_text(size=14),legend.title=element_text(size=16),legend.key.width=unit(1,"cm"),legend.position="right",legend.justification="bottom")+labs(x="Time Elapsed (sec)",y="Predicted Copulation Rate (Events/sec)")

newcolors<-c(mycolors,"black")
names(newcolors)[6]<-"black"

#main_small<-ggplot()+geom_segment(data=datlist.short[[1]],aes(x=avail,xend=not,y=0,yend=0),size=125,colour="grey85")+geom_step(data=datlist.short[[2]],aes(x=time,y=solrate,colour=male),direction="hv")+geom_ribbon(data=datlist.short[[3]],aes(x=time,ymax=solrate,ymin=0,fill=male))+scale_fill_manual(values=mycolors,guide=F)+geom_step(data=datlist.short[[2]],aes(x=time,y=int.scaled),direction="hv",colour="pink")+geom_ribbon(data=datlist.short[[3]],aes(x=time,ymax=int.scaled,ymin=0),fill="red")+theme_classic()+coord_cartesian(ylim=c(0,0.003))+geom_segment(data=datlist.short[[4]],aes(x=Time,xend=Time,y=0,yend=0.003,colour="black",linetype="Copulation\nEvents"),size=0.8)+geom_segment(data=datlist.short[[5]],aes(x=Time,xend=Time,y=0,yend=0.003,colour=perp,linetype=perp),size=0.8)+geom_abline(slope=0,intercept=0,size=0.2)+facet_wrap(.~male, nrow=5)+theme(strip.background=element_blank(),strip.text=element_blank(),axis.title.x=element_text(size=20,face="bold"),axis.title.y=element_text(size=20,face="bold"),axis.text.x=element_text(size=18,face="bold"),axis.text.y=element_text(size=18,face="bold"),legend.text=element_text(size=14),legend.title=element_text(size=16),legend.key.width=unit(1,"cm"),legend.position="right",legend.justification="bottom")+labs(x="Time Elapsed (sec)",y="Predicted Copulation Rate (Events/sec)")+scale_colour_manual(name="Copulation\nEvents",values=c(newcolors),labels=c("Int. by 1","Int. by 2","Int. by 3","Int. by 4","Int. by 7","Uninterr."))+scale_linetype_manual(name="Copulation\nEvents",values=c(5,5,5,5,5,1),labels=c("Int. by 1","Int. by 2","Int. by 3","Int. by 4","Int. by 7","Uninterr."))


#ggsave("plottest.pdf",main_small)

#for the smaller version:

availdat.plot<-remove.males(availdat.plot)
interrate<-remove.males(interrate)
df.sol<-remove.males(df.sol)
copends<-remove.males(copends)
intevents<-remove.males(intevents)


## Try to add in new metrics for stats- 

 {
# #I am imagining lines for the top few stats... I think I will simply focus on the contributions of different stats to the total hazard for each kind of event? This means I need to use the data from the statlist, not the saved hazards... I should be able to use the statlist with the altered hazards for interruptions by just leaving out those columns. 
# 
# 
# #This time, instead of using every column summed to get the hazard for an event, I'll break it down. 
# 
# library(tidyverse)
# library(dplyr)
# 
# #dummy data output vector
# output<-rep(0,(3+nrow(mod_coef)))
# names(output)<-c("male",rownames(mod_coef),"day","time")
# 
# for(day in 1:length(test.stat)){
#   
#   
#   for(step in 1:nrow(evlist[[day]])){
#     #stat matrix predicting the next event!
#     
#     #exclude "init" event rows
#     if(grepl("init",event.list.dat[[day]][step,1])==F){
#     
#     stats<-test.stat[[day]][[1]][step,,]
#     
#     #the coefficient vector gets multiplied by each row and the supplist vector gets multiplied by each column
#     statsup<-apply(stats,2,function(x) x*supplist[[day]][step,])
#     
#     #for some reason the apply function across the row margin returns a transformed matrix, so I transform it back for sanity. Use the subset of coefficents for the predictors that are acutally included for this timestep
#     statcoef<-t(apply(statsup,1,function(x) x*mod_coef[dimnames(stats)[[2]],]))
#     
#     statdat<-(data.frame(statcoef))
#     colnames(statdat)<-rownames(mod_coef)
#     
#     #now I have a matrix of event types by parameter-weighted contributions to the hazards. The sum of each row is the log hazard of the event type
#     
#     #I want to sum each column within each group of events that ocrrespond to the receive hazards for each male
#     
#     #restrict to only "attack" event types 
#     
#     statdat<-statdat %>%
#     mutate(event=event.types) %>%
#     filter(grepl("attack",event))
#     
#     statdat<-mutate(statdat,receiver=sapply(statdat$event,function(x)strsplit(x,split=".",fixed=T)[[1]][3]))
#   
#   statdat<-select(statdat,-event)
#   	statdat[,c(1:(ncol(statdat)-1))]<-apply(statdat[,c(1:(ncol(statdat)-1))],2,function(x) ifelse(x!=0,exp(x),0))
#   	
#   	
#   	statdat<-statdat %>%
#   	group_by(receiver) %>%
#     summarise(across(where(is.numeric),sum)) %>%
#     mutate(receiver=as.factor(receiver)) %>%
#     #restrict to just the top 3 males
#     filter(receiver=="2" | receiver=="3" | receiver=="4")%>%
#     rename(male=receiver)
#   
#    
# #ok so now I have a 3x105 data frame where each row represents a particular male's receive hazard, broken down into columns which are the sums of the exponentiated factors for each stat. These represent the multaplicitive contribution to the overall hazard of getting attacked, if everything else was equal
#   
#   #Now I need columns for date and for the timing and then I will rbind this with each other event in the series. 
#     
#     date<-rep(day,nrow(statdat))
#     time<-rep(event.list.dat[[day]][step,2],nrow(statdat))
#     
#     statdat<-statdat %>%
#     mutate(day=date)%>%
#     mutate(time=time)
#     
#     #append to a data frame
#     
#     output<-rbind(output,statdat)
#    } 
#    
#   }
# }
# 
# 
# 
# #remove dummy row after
# 
# output<-output[-1,]
# 
# #choose the top 5? predictors to show- the ones with the largest sums overall? largest average value?
# 
# stattots<-as.matrix(summarise(output,across(Global.attackFE:log.abs.winloss.diff.pairwise.day,mean)))
# 
# statord<-stattots[1,][order(stattots[1,],decreasing=T)]
# 
# #eliminate everything that's zero
# statord<-statord[statord>0]
# 
# #eliminateFEs, also cop stats since they're special in this statlist
# statord<-statord[!grepl("FE|cop",names(statord),perl=T)]
# 
# #eliminate others I'm not that interested in right now- females, centroid, and initiator effect
# statord<-statord[-c(which(names(statord)=="females.attack"),which(names(statord)=="centroid.dist"),which(names(statord)=="gen.win.rec.anger"))]
# 
# #keep the top 5
# stat.elim<-names(statord)[1:3]
# 
# #pare down to just these columns in the output dataframe
# 
# statlevels<-output[,c("male","time","day",stat.elim)]
# 
# #I will finally need to "pivot longer" the table to turn the stats I want into factors so then i can plot them all on the same graph
# 
# statlevels<-statlevels %>%
# 	pivot_longer(4:ncol(statlevels),names_to="stat",values_to="count")
# 
# #then split by day
# 
# statlev.day<-split(statlevels,statlevels$day)
# 
# 
# #######
# ##grab the example for day 13
# 
# statlev<-statlev.day[[13]]
# 
# #fix factor levels
# statlev$male<-factor(statlev$male,levels=c(1:30))
# 
# #for day 13, females depart at 4800, so pare down the data to that section for the plot
# 
# statlev<-statlev[!statlev$time>4800,]
# 
# #Awesome! try plotting:
# 
# ggplot()+geom_step(data=statlev,aes(x=time,y=count,colour=stat),direction="hv")+facet_grid(male~.)
# 
# ##hmmm... doesn't look very promising, since it's really messy to look at. Maybe not necessary after all... 
# 
# 
# 
# ##area plot version - possible to do stacked step+area? This code doesn't work since I have several males as categories as well... 
# 
# df<-split(statlev,statlev$male)
# 
# 
# 
# df2<-lapply(df,function(x){
# return(rbind(
#   x,
#   transform(x[order(x$time),],
#     time=time + 1e-9,  # required to avoid crazy steps
#     count=ave(count, stat, FUN=function(z) c(tail(z, -1L),z[[length(z)]]))
# ) ))
# 
# })
# 
# df3<-bind_rows(df2[[2]],df2[[3]],df2[[4]])
# 
# head(df2)
# 
# 
# ggplot()+geom_area(data=df3,aes(x=time,y=count,fill=stat))+facet_grid(male~.)


#issue still is that these are MULTIPLIERS so small differences yield big changes in hazard, especially at large magnitude

}

### Try something simpler- just look at counts of attacks, receives, losses, and wins, and then maybe receive+losses form yesterday, since it's important for cop inters

#this is ok since we already have a sense of the hazard of getting attacked with the risk of interruption, so we'll see if there's any discernible differences here


#do I have the winloss census available? yes. 

#Use code from my functions for counting the stats


#input will be the win.loss.census object

statcounts.fun<-function(x){
	
win.stat<-x

day.list<-list()

library(abind)

#loop through each day sequence
for(k in 1:length(win.stat)){

#need a dummy first event row

#malecols<-ncol(win.stat[[1]])  
#malenums<-c(1:malecols)
  
malenums<-c(2,3,4)
malecols<-length(malenums)  

baseline<-data.frame(male=as.character(malenums),attacks=rep(0,malecols),receives=rep(0,malecols),wins=rep(0,malecols),losses=rep(0,malecols),attackwins=rep(0,malecols),attacklosses=rep(0,malecols),receivewins=rep(0,malecols),receivelosses=rep(0,malecols),time=rep(0,malecols))

#this loops through each event in the sequence

for(j in 2:nrow(win.stat[[k]])){

#exclude "init" event rows here so I don't have to worry about it later
    if(grepl("init",event.list.dat[[k]][j,1])==F){

#for each event, loop through all the males to get the previous attack count that will predict the next interaction- the support array will take care of males that weren't present that day.
#for(i in 1:ncol(win.stat[[1]])){

#just go thru males 2 3 and 4 for now to save time and space      
for(i in c(2,3,4)){	

#list of all the attacks by the focal male
attacks<-grep("attack",win.stat[[k]][1:(j-1),i])
receives<-grep("received",win.stat[[k]][1:(j-1),i])
wins<-grep("win",win.stat[[k]][1:(j-1),i])
losses<-grep("lose",win.stat[[k]][1:(j-1),i])


#Remove the "fights" and "nothings" from the census list. This creates attack/receive and win/lose pairs in order of occurence
fight.times<-grep("fight",win.stat[[k]][1:(j-1),i])
nothing.times<-grep("nothing",win.stat[[k]][1:(j-1),i])

pairs<-win.stat[[k]][1:(j-1),i][-c(fight.times, nothing.times)]

#need a way to do this that matches interaction partners but also is sensitive to the time-ordering 

attack.losses<-vector()

if(length(pairs)>0){

attack.losses<-unlist(sapply(1:ncol(win.stat[[1]]), function(z){
	
	intset<-pairs[sapply(pairs, function(x) {
		
		strsplit(x,split=".",fixed=T)[[1]][3]==z
		
	})]
	
	names(intset)<-grep("TRUE",sapply(pairs, function(x) {
		
		strsplit(x,split=".",fixed=T)[[1]][3]==z
		
	}))
	
	attacks<-grep("attack",intset)
	attackpluslose<-intset[attacks+1][grep("lose", intset[attacks+1])]
	
	return(attackpluslose)

}))
}


attack.wins<-vector()

if(length(pairs)>0){

attack.wins<-unlist(sapply(1:ncol(win.stat[[1]]), function(z){
	
	intset<-pairs[sapply(pairs, function(x) {
		
		strsplit(x,split=".",fixed=T)[[1]][3]==z
		
	})]
	
	names(intset)<-grep("TRUE",sapply(pairs, function(x) {
		
		strsplit(x,split=".",fixed=T)[[1]][3]==z
		
	}))
	
	attacks<-grep("attack",intset)
	attackpluswin<-intset[attacks+1][grep("win", intset[attacks+1])]
	
	return(attackpluswin)

}))
}


receive.wins<-vector()

if(length(pairs)>0){

receive.wins<-unlist(sapply(1:ncol(win.stat[[1]]), function(z){
	
	intset<-pairs[sapply(pairs, function(x) {
		
		strsplit(x,split=".",fixed=T)[[1]][3]==z
		
	})]
	
	names(intset)<-grep("TRUE",sapply(pairs, function(x) {
		
		strsplit(x,split=".",fixed=T)[[1]][3]==z
		
	}))
	
	receives<-grep("receive",intset)
	receivepluswin<-intset[receives+1][grep("win", intset[receives+1])]
	
	return(receivepluswin)

}))
}


receive.losses<-vector()

if(length(pairs)>0){

receive.losses<-unlist(sapply(1:ncol(win.stat[[1]]), function(z){
	
	intset<-pairs[sapply(pairs, function(x) {
		
		strsplit(x,split=".",fixed=T)[[1]][3]==z
		
	})]
	
	names(intset)<-grep("TRUE",sapply(pairs, function(x) {
		
		strsplit(x,split=".",fixed=T)[[1]][3]==z
		
	}))
	
	receives<-grep("receive",intset)
	receivepluslose<-intset[receives+1][grep("lose", intset[receives+1])]
	
	return(receivepluslose)

}))
}


#put in the time of the event so I can index it later

#append to the list for the day
baseline<-rbind(baseline,c(i,length(attacks),length(receives),length(wins),length(losses),length(attack.wins),length(attack.losses),length(receive.wins),length(receive.losses),event.list.dat[[k]][j,2]))


}

}
}

#the j loop ends above, append the new object for the day to the daylist?

day.list[[k]]<-baseline

}

return(day.list)


}

statcounts<-statcounts.fun(win.loss.census)

stat.day<-statcounts[[13]]

#head(stat.day,35)

#restrict to tiems when females are present
stat.day<-stat.day[!stat.day$time>4800,]

#dim(stat.day)
#ok now I have a list of the times and counts for each of the three males. I can plot these as step functions or as dots? try both

#have to pivot longer to make the different counts into factors 

library(tidyverse)

stat.main<-stat.day %>%
  select(c(male:losses,time))%>%
  pivot_longer(attacks:losses,names_to="stat",values_to="count")

stat.interaction<-stat.day %>%
  select(male,c(attackwins:time))%>%
  pivot_longer(attackwins:receivelosses,names_to="stat",values_to="count")

ggplot()+geom_step(data=stat.interaction,aes(x=time,y=count,colour=stat))+facet_grid(male~.)

#the good thing about the interaction data is that the total is the total number of fights 

#try area plot version

df<-split(stat.interaction,stat.interaction$male)


df2<-lapply(df,function(x){
  return(rbind(
    x,
    transform(x[order(x$time),],
              time=time + 1e-9,  # required to avoid crazy steps
              count=ave(count, stat, FUN=function(z) c(tail(z, -1L),z[[length(z)]]))
    ) ))
  
})

df3<-bind_rows(df2[[1]],df2[[2]],df2[[3]])

#head(df3)

ggplot()+geom_area(data=df3,aes(x=time,y=count,fill=stat))+facet_grid(male~.)


#might be able to combine everything if I add a new factor for the datatset and shove it all together... 

df3<-df3%>%
  mutate(dattype=factor("statcount",levels=c("statcount","solplot")),ymin=0,ymax=40)

for(i in 1:5){

datlist.short[[i]]<-datlist.short[[i]]%>%
  mutate(dattype=factor("solplot",levels=c("statcount","solplot")),ymin=0,ymax=0.003)
 
}


#try putting it together

#need another new color scale to accomodate the different fills
newcolors<-c(mycolors,"black")
names(newcolors)[6]<-"black"

newcolors<-c(newcolors,"blue","purple","red","pink")
names(newcolors)[c(7,8,9,10)]<-levels(as.factor(df3$stat))

#add a line to make the y axis scales work

whiteline<-data.frame(male=2,height=0.003,time=0,dattype=factor("solplot",levels=c("statcount","solplot")))

#add horizontal lines for receive+losses yesterday?

day<-13

win.stat<-win.loss.census

rec.lose<-c(0,0)

for(i in c(2,3,4)){	

  #Remove the "fights" and "nothings" from the census list. This creates attack/receive and win/lose pairs in order of occurence
  fight.times<-grep("fight",win.stat[[day-1]][,i])
  nothing.times<-grep("nothing",win.stat[[day-1]][,i])
  
  pairs<-win.stat[[day-1]][,i][-c(fight.times, nothing.times)]

receive.losses<-vector()

if(length(pairs)>0){
  
  receive.losses<-unlist(sapply(1:ncol(win.stat[[1]]), function(z){
    
    intset<-pairs[sapply(pairs, function(x) {
      
      strsplit(x,split=".",fixed=T)[[1]][3]==z
      
    })]
    
    names(intset)<-grep("TRUE",sapply(pairs, function(x) {
      
      strsplit(x,split=".",fixed=T)[[1]][3]==z
      
    }))
    
    receives<-grep("receive",intset)
    receivepluslose<-intset[receives+1][grep("lose", intset[receives+1])]
    
    return(receivepluslose)
    
  }))
}

rec.lose<-rbind(rec.lose,c(i,length(receive.losses)))

}

rec.lose<-rec.lose[-1,]

rec.lose<-data.frame(male=rec.lose[,1],yint=rec.lose[,2],dattype=factor("statcount",levels=c("statcount","solplot")))


#main_small_counts<-ggplot()+geom_segment(data=datlist.short[[1]],aes(x=avail,xend=not,y=0,yend=0),size=125,colour="grey85")+geom_step(data=datlist.short[[2]],aes(x=time,y=solrate,colour=male),direction="hv")+geom_ribbon(data=datlist.short[[3]],aes(x=time,ymax=solrate,ymin=0,fill=male))+geom_step(data=datlist.short[[2]],aes(x=time,y=int.scaled),direction="hv",colour="pink")+geom_ribbon(data=datlist.short[[3]],aes(x=time,ymax=int.scaled,ymin=0),fill="red")+theme_classic()+geom_segment(data=datlist.short[[4]],aes(x=Time,xend=Time,y=0,yend=0.003,colour="black",linetype="Copulation\nEvents\n(Vertical Lines)"),size=0.8)+geom_segment(data=datlist.short[[5]],aes(x=Time,xend=Time,y=0,yend=0.003,colour=perp,linetype=perp),size=0.8)+geom_abline(slope=0,intercept=0,size=0.2)+theme(strip.background=element_blank(),strip.text=element_blank(),axis.title.x=element_text(size=20,face="bold"),axis.title.y=element_text(size=20,face="bold"),axis.text.x=element_text(size=16,face="bold"),axis.text.y=element_text(size=16,face="bold"),legend.text=element_text(size=16),legend.title=element_text(size=18),legend.key.width=unit(1,"cm"),legend.position="right",legend.justification="bottom")+labs(x="Time Elapsed (sec)",y=element_blank())+scale_colour_manual(name="Copulation\nEvents\n(Vertical Lines)",values=c(newcolors),labels=c("Int. by 1","Int. by 2","Int. by 3","Int. by 4","Int. by 7","Uninterr."))+scale_linetype_manual(name="Copulation\nEvents\n(Vertical Lines)",values=c(5,5,5,5,5,1),labels=c("Int. by 1","Int. by 2","Int. by 3","Int. by 4","Int. by 7","Uninterr."))+scale_fill_manual(name="Counts of\nBehaviors",values=newcolors,labels=c("2","3","4","attack+losses","attack+wins","receive+losses","receive+wins"))+geom_area(data=df3,aes(x=time,y=count,fill=stat))+geom_segment(data=whiteline,aes(x=time,xend=time,y=0,yend=height),colour="white")+geom_segment(data=rec.lose,aes(x=0,xend=4800,y=yint,yend=yint),colour="black",linetype=6)+facet_wrap(.~male+dattype, nrow=6,scales="free")+geom_blank(data=df3,aes(y=ymin))+geom_blank(data=df3,aes(y=ymax))+guides(colour=guide_legend(order=2),linetype=guide_legend(order=2),fill=guide_legend(order=1))

#try reorganizing the colors to make it easier to understand, since I'm going to be grouping the plots by male anyway using cartoons. Just change the solicitation rate to the same blue as the summary barplot

#main_small_counts<-ggplot()+geom_segment(data=datlist.short[[1]],aes(x=avail,xend=not,y=0,yend=0),size=125,colour="grey85")+geom_step(data=datlist.short[[2]],aes(x=time,y=solrate),colour="grey30",direction="hv")+geom_ribbon(data=datlist.short[[3]],aes(x=time,ymax=solrate,ymin=0),fill="light blue")+geom_step(data=datlist.short[[2]],aes(x=time,y=int.scaled),direction="hv",colour="pink")+geom_ribbon(data=datlist.short[[3]],aes(x=time,ymax=int.scaled,ymin=0),fill="red")+theme_classic()+geom_segment(data=datlist.short[[4]],aes(x=Time,xend=Time,y=0,yend=0.003,colour="black",linetype="Copulation\nEvents\n(Vertical Lines)"),size=0.8)+geom_segment(data=datlist.short[[5]],aes(x=Time,xend=Time,y=0,yend=0.003,colour=perp,linetype=perp),size=0.8)+geom_abline(slope=0,intercept=0,size=0.2)+theme(strip.background=element_blank(),strip.text=element_blank(),axis.title.x=element_text(size=20,face="bold"),axis.title.y=element_text(size=20,face="bold"),axis.text.x=element_text(size=16,face="bold"),axis.text.y=element_text(size=16,face="bold"),legend.text=element_text(size=16),legend.title=element_text(size=18),legend.key.width=unit(1,"cm"),legend.position="right",legend.justification="top")+labs(x="Time Elapsed (sec)",y=element_blank())+scale_colour_manual(name="Copulation\nEvents\n(Vertical Lines)",values=c(newcolors),labels=c("Int. by 1","Int. by 2","Int. by 3","Int. by 4","Int. by 7","Uninterr."))+scale_linetype_manual(name="Copulation\nEvents\n(Vertical Lines)",values=c(5,5,5,5,5,1),labels=c("Int. by 1","Int. by 2","Int. by 3","Int. by 4","Int. by 7","Uninterr."))+scale_fill_manual(name="Counts of\nBehaviors",values=newcolors,labels=c("attack+losses","attack+wins","receive+losses","receive+wins"))+geom_area(data=df3,aes(x=time,y=count,fill=stat))+geom_segment(data=whiteline,aes(x=time,xend=time,y=0,yend=height),colour="white")+geom_segment(data=rec.lose,aes(x=0,xend=4800,y=yint,yend=yint),colour="black",linetype=6)+facet_wrap(.~male+dattype, nrow=6,scales="free")+geom_blank(data=df3,aes(y=ymin))+geom_blank(data=df3,aes(y=ymax))+guides(colour=guide_legend(order=2),linetype=guide_legend(order=2),fill=guide_legend(order=1))


#also consider removing attack+wins? rename "counts of behaviors contributing to fight/interruption risk"

df4<-filter(df3,stat!="attackwins")

newcolors<-newcolors[-7]
names(newcolors)[7]<-"attacklosses"


main_small_counts<-ggplot()+geom_segment(data=datlist.short[[1]],aes(x=avail,xend=not,y=0,yend=0),size=125,colour="grey85")+geom_step(data=datlist.short[[2]],aes(x=time,y=solrate),colour="grey30",direction="hv")+geom_ribbon(data=datlist.short[[3]],aes(x=time,ymax=solrate,ymin=0),fill="light blue")+geom_step(data=datlist.short[[2]],aes(x=time,y=int.scaled),direction="hv",colour="pink")+geom_ribbon(data=datlist.short[[3]],aes(x=time,ymax=int.scaled,ymin=0),fill="red")+theme_classic()+geom_segment(data=datlist.short[[4]],aes(x=Time,xend=Time,y=0,yend=0.003,colour="black",linetype="Copulation\nSolicitation Events\n(Vertical Lines)"),size=0.8)+geom_segment(data=datlist.short[[5]],aes(x=Time,xend=Time,y=0,yend=0.003,colour=perp,linetype=perp),size=0.8)+geom_abline(slope=0,intercept=0,size=0.2)+theme(strip.background=element_blank(),strip.text=element_blank(),axis.title.x=element_text(size=20,face="bold"),axis.title.y=element_text(size=20,face="bold"),axis.text.x=element_text(size=16,face="bold"),axis.text.y=element_text(size=16,face="bold"),legend.text=element_text(size=20),legend.title=element_text(size=20,face="bold"),legend.key.width=unit(1,"cm"),legend.key.size=unit(0.25,"cm"),legend.position="right",legend.justification="top")+labs(x="Time Elapsed (sec)",y=element_blank())+scale_colour_manual(name="Copulation\nSolicitation Events\n(Vertical Lines)",values=c(newcolors),labels=c("Int. by Male 1","Int. by Male 2","Int. by Male 3","Int. by Male 4","Int. by Male 7","Uninterr."))+scale_linetype_manual(name="Copulation\nSolicitation Events\n(Vertical Lines)",values=c(5,5,5,5,5,1),labels=c("Int. by Male 1","Int. by Male 2","Int. by Male 3","Int. by Male 4","Int. by Male 7","Uninterr."))+scale_fill_manual(name="Counts of Fights\nContributing to\nInterruption Risk",values=newcolors,labels=c("attack+losses","receive+losses","receive+wins","receive+losses yesterday"))+geom_area(data=df4,aes(x=time,y=count,fill=stat))+geom_segment(data=whiteline,aes(x=time,xend=time,y=0,yend=height),colour="white")+geom_segment(data=rec.lose,aes(x=0,xend=4800,y=yint,yend=yint),colour="black",linetype=6)+facet_wrap(.~male+dattype, nrow=6,scales="free")+geom_blank(data=df3,aes(y=ymin))+geom_blank(data=df3,aes(y=ymax))+guides(colour=guide_legend(order=2),linetype=guide_legend(order=2),fill=guide_legend(order=1))

#don't worry about trying to add in the legend for receives yesterday, I'll just increase the spacing and add my own later

library(grid)
library(gtable)

gt<-ggplot_gtable(ggplot_build(main_small_counts))

#figure out which spacings I need to adjust; super cool function that lets you see the underlying layout of the plot!
#gtable_show_layout(gt)

#seems like I want to expand rows 13, 23, and 33 
gt$heights[c(13,23,33)]<-1.5*gt$heights[c(13,23,33)]

#and I can increase the spacing between plot groupings rows 16 and 26
gt$heights[c(5,6,7,15,16,17,25,26,27)]<-3*gt$heights[c(5,6,7,15,16,17,25,26,27)]

#pad out the plot for combining, and increase space between plots and legends
#1 on the left, 38 on the bottom and 11 on the right. 8 is the space between plots and guides
gt$widths[1]<-45*gt$widths[1]
gt$widths[11]<-6*gt$widths[11]
gt$widths[8]<-6*gt$widths[8]

gt$heights[38]<-6*gt$heights[38]

#finally, add a little space between the legends?
#get the index of the guidebox
lbox<-which(sapply(gt$grobs,paste)=="gtable[guide-box]")
#change the height of the spacing
gt$grobs[[lbox]]$heights[4]<-4*gt$grobs[[lbox]]$heights[4]


#grid.draw(gt)

#seems like the object is the gt thing and then that can be used in a new plot grid? 


######and calculate the total numbers to display in a table/chart on the side

availdat.plot$tot<-availdat.plot$not-availdat.plot$avail

head(availdat.plot)

#get total amount of time actually available for mating

coptable<-data.frame(male=unique(availdat.plot$male),total=unique(ave(availdat.plot$tot,availdat.plot$male, FUN=function(x) sum(x))))


#now calculate the average solrate

#the weighted average by the amount of time each rate obtains. The numerator of this also happens to be the riemann sum corresponding to the expected number of copulations. 

head(interrate)

mate.tot<-vector()
int.tot<-vector()
realized.tot<-vector()


for(i in 1:length(unique(interrate$male))){
	
	maleid<-unique(interrate$male)[i]
	
	tempdat<-interrate[interrate$male==maleid,]

	#get the legnth of each interval
	interval<-lead(tempdat$time)-tempdat$time
	
	#multiply by the solrate associated with each time interval
	interval.matenum<-interval*tempdat$solrate
	
	#get the sum for the "expected" total number of solicitations based on the model
	mate.total<-sum(interval.matenum,na.rm=T)
	
	mate.tot[i]<-mate.total


	#get the interruption probability associated with each time interval
	interval.intprob<-interval*tempdat$intprob
	
	#get the sum- this will have to be divided by the appropriate time available to get the average interruption risk 
	int.total<-sum(interval.intprob, na.rm=T) 

	int.tot[i]<-int.total
	
	
	
	#get the expected number of SUCCESSFUL COPULATIONS for the given time period
	interval.success<-interval*(tempdat$solrate-tempdat$int.scaled) 
	
	#get the sum for the expected total realized copulations expected
	success.total<-sum(interval.success, na.rm=T) 

	realized.tot[i]<-success.total

}

#add to the table
coptable<-cbind(coptable,mate.tot)
coptable<-cbind(coptable,int.tot)
coptable<-cbind(coptable,realized.tot)

#get the average solicitation rate- this is the more appropriate one to report since it does not also include the time component of availability to mate
coptable$sol.ave<-coptable$mate.tot/coptable$total

#get the average predicted interruption rate
coptable$int.ave<-coptable$int.tot/coptable$total


##add the actual observed realized mating success for the day


#all set!!

#move on to making the tables to display the info


timeplot<-ggplot()+geom_bar(data=coptable,aes(x=male,y=total),stat="identity",fill="grey85")+theme_classic()+theme(legend.position="none",axis.title.x=element_text(size=20,face="bold"),axis.title.y=element_text(size=18,face="bold"),axis.text.x=element_text(size=30,face="bold"),axis.text.y=element_text(size=16,face="bold"))+labs(x="Time Available for Mating\n (Time Not Fighting)", y="Seconds")

mateplot<-ggplot()+geom_bar(data=coptable,aes(x=male,y=sol.ave),fill="light blue",colour="grey60",stat="identity")+theme_classic()+theme(legend.position="none",axis.title.x=element_text(size=20,face="bold"),axis.title.y=element_text(size=18,face="bold"),axis.text.x=element_text(size=30,face="bold"),axis.text.y=element_text(size=16,face="bold"))+labs(x="Average Solicitation Hazard\n (Attractiveness)", y="Solicitations per Second")

intplot<-ggplot()+geom_bar(data=coptable,aes(x=male,y=int.ave),stat="identity",fill="red",colour="pink")+theme_classic()+theme(legend.position="none",axis.title.x=element_text(size=20,face="bold"),axis.title.y=element_text(size=18,face="bold"),axis.text.x=element_text(size=30,face="bold"),axis.text.y=element_text(size=16,face="bold"))+labs(x="Average Predicted Probability\nof Copulation Interruption", y="Probability")

##add in the actual observed number of uninterruptedmating events on top
#have to add in a legend manually later, I guess?
copends$Events<-as.factor(as.character(copends$Events))
copends$male<-factor(as.character(copends$male), levels=c("2","3","4"))

obscop<-apply(table(copends$Events,copends$male),2,function(x) sum(x))

obscop<-data.frame(male=names(obscop),obs=obscop)

realplot<-ggplot()+geom_bar(data=coptable,aes(x=male,y=realized.tot,fill=male),stat="identity")+scale_fill_manual(values=mycolors)+theme_classic()+theme(axis.title.x=element_text(size=20,face="bold"),axis.title.y=element_text(size=18,face="bold"),axis.text.x=element_text(size=30,face="bold"),axis.text.y=element_text(size=16,face="bold"),legend.text=element_blank(),legend.title=element_text(size=18,face="bold"), legend.key.width=unit(20,"mm"),legend.key.size=unit(1.5,"mm"),legend.background=element_rect(colour="black"))+labs(x="Predicted Uninterrupted\n Mating Events", y="#  Successful Copulations")+geom_boxplot(data=obscop,aes(x=male,y=obs,colour="Observed\nMating\nSuccess"))+scale_colour_manual(name="Observed\nMating\nSuccess",values="black")+guides(fill=F)


library(cowplot)

#add an arrow?
arrowplot<-ggplot()+geom_segment(aes(x=0,xend=0.5,y=1,yend=1),arrow=arrow(angle=50,type="closed",length=unit(0.5,"cm")),size=4)+xlim(c(0,0.6))+theme(legend.position = "none", panel.grid = element_blank(), axis.title = element_blank(), axis.text = element_blank(), axis.ticks = element_blank(), panel.background = element_blank())


library("gridExtra")

metrics<-grid.arrange(timeplot,mateplot,intplot,arrowplot,realplot,ncol=14,nrow=1,layout_matrix=matrix(c(1,1,1,2,2,2,3,3,3,4,5,5,5,5),nrow=1))


#figure out the particulars of labelling, etc. later if I decide I really want to use this one
#,annotate_figure(fig.lab="Overall")

plotall<-grid.arrange(gt,metrics,ncol=1,nrow=10,layout_matrix=matrix(c(1,1,1,1,1,1,1,2,2,2)))


setwd(paste(here(),"/adequacy_checking/adequacy_attractiveness_model",sep=""))

ggsave("matingplot_new_statlevs_newlabs.pdf",plot=plotall,height=17,width=23)

#increzse space between legends and plots to the right; increase space on the left as well. Pad out the bottom too.
#add receive+lose from yesterday as a horizontal line? 

#icrease the horizontal spacing a little more to make rom for stuff on the sides
#maybe change the ratio of pltos to 1.5 instead of 2 to emphasize the count a bit more
#increase size of legend text and titles


##### make new barplots for the overall fight counts for the three males in case I prefer to show it this way 


defcounts<-stat.interaction[stat.interaction$time==4800,]

defcounts<-defcounts[,-2]

rec.lose$stat<-"recloseyest"

def.rec<-rec.lose[,c(1,4,2)]
colnames(def.rec)[3]<-"count"

def.rec$male<-as.factor(def.rec$male)
def.all<-dplyr::bind_rows(defcounts,def.rec)

def.all$stat<-factor(def.all$stat,levels=c("attackwins","attacklosses","receivewins","receivelosses","recloseyest"))

#rename them something nicer
levels(def.all$stat)<-c("attack+win","attack+lose","receive+win","receive+lose","receive+lose\nyesterday  ")

defplot<-ggplot()+geom_bar(data=def.all,aes(x=stat,y=count),stat="identity")+facet_wrap(.~male,nrow=1,ncol=3)


setwd(paste(here(),"/adequacy_checking/adequacy_attractiveness_model",sep=""))

defcolors<-c("blue","purple","pink","red","grey")

trycolors<-gg_color_hue(5)[c(2,3,4)]

newplot<-ggplot()+geom_bar(data=def.all,aes(x=stat,y=count,fill=stat),stat="identity")+scale_fill_manual(values=defcolors)+facet_wrap(.~male,nrow=1,ncol=3)+theme_classic()+theme(strip.background=element_blank(),strip.text=element_blank(),axis.line.x=element_line(colour="black",size=1.5, linetype="solid"),axis.line.y=element_line(colour="black",size=1.5, linetype="solid"), axis.text.x=element_text(size=14,face="bold",angle=50,vjust=1,hjust=1), axis.text.y=element_text(size=18,face="bold"), axis.title.x=element_text(size=25,face="bold"), axis.title.y=element_text(size=25,face="bold"))+labs(x="Interaction Type",y="Total Count")+guides(fill=F)

#adjust the spacing between facets
library(grid)
library(gtable)

gt<-ggplot_gtable(ggplot_build(newplot))

#figure out which spacings I need to adjust; super cool function that lets you see the underlying layout of the plot!
#gtable_show_layout(gt)

gt$widths[7]<-7*gt$widths[7]
gt$widths[11]<-7*gt$widths[11]


ggsave("fight_count_fig.pdf",gt,height=5,width=10,units=c("in"))
```
