## Supplemental_reproducible_code for "Fighting isn’t sexy in lekking Greater Sage-grouse (*Centrocercus urophasianus*)": fight_effect_explore.R

```
######## Explore the effects of different events using the coefficients to produce hazards #######

rm(list=ls())

###make sure necessary packages are there and loaded

packages<-c("here","abind","relevent","lubridate","DescTools","gtools","ggplot2","tidyverse")

package.check<-lapply(packages, FUN = function(x) {
    if (!require(x, character.only = TRUE, quietly=TRUE)) {
        install.packages(x, dependencies = TRUE)
        library(x, character.only = TRUE, quietly=TRUE)
    } 

})

#for project path management
#define the project root directory relative to the current script. This sets the working directory that gets called by the here() function. It is a relative path and will be robust to use on anyone's computer. In this case, I want the root directory to always be the enclosing folder for the whole sage grouse project

i_am("adequacy_checking/adequacy_attractiveness_model/fight_effect_explore.R")

  
  ##source the rest of my fuctions for predictor stats- the standard functions should work fine in this case
  #enable all the functions I have so far
  source(here("rem_pre_analysis","rem.functions.R"), chdir = TRUE)
  
  #make a list of the functions
  fun.list<-ls()[grep("add.",ls())]

  tests<-data.frame(initiate=c("receive","receive","receive","attack","attack"),outcome=c("lose","lose","win","lose","win"),kind=c("CH","SF","SF","SF","SF"),stringsAsFactors=F)

  one.out<-c(1,1)  
  
  cop.out<-c(1,1)
  
 
  ###Build the intercept list, support array, event list, census array, etc; i'll turn this into a function I think since it has to happen five times

library(tidyverse)
  
for(r in 1:nrow(tests)){  
  
    
    #make the base event list and intercept array
    
   mod.day<-1 
    
      event.types<-read.table(file=here("data_processing","event_types_list_udata.csv"), header=F,  sep=",")
      event.types<-as.character(event.types$V1)
    

    #NOTE- the event.types vector now contains ENDOGENOUS event types as well as EXOGENOUS types: arrivals and clock events, and end.obs. Need to make sure everything is robust to this. 
    
    ## make event.types vector usable for intlist, etc. by excising exogenous event types
    
    
    endogenous<-(length(event.types)-length(event.types[grep("arrive",event.types)])-5)
    
   
     ##create my contrived event list: beginning of day 1, females arrive and then an interaction occurs between males 1 and 2
    
    #input<-c("receive","lose","SF")
    input<-as.character(tests[r,])
    
    if(input[1]=="receive"){attacker<-2; receiver<-1} else {attacker<-1; receiver<-2}
    
    if(input[2]=="lose"){winner<-2; loser<-1} else {winner<-1; loser<-2}
    
    fight.type<-input[3]
    
   
    #the final event before end.obs is just a placeholder
    
    testevents1<-c("fem.enter",paste0(attacker,".attack.",receiver),paste0(receiver,".init",fight.type),paste0(winner,".disengage"),"disturb.ance","disturb.ance","end.obs")
    
    #event history with a copulation to get interruption risk
    testevents2<-c("fem.enter",paste0(attacker,".attack.",receiver),paste0(receiver,".init",fight.type),paste0(winner,".disengage"),"1.cop.start","disturb.ance","end.obs")
    
    testtimes<-c(0,5,5.001,10,10.001,15,16)
    
    testev.list<-list(testevents1,testevents2)
    
     testout<-list()
    
for(w in 1:length(testev.list)){
    
    event.list<-data.frame(Events=testev.list[[w]],Time=testtimes,Females=2,Date="2014-03-19")
    
    event.list$Date<-as.factor(event.list$Date)
    
    event.list.dat<-split(event.list, event.list$Date)
    
    basic.event.list<-data.frame(Events=testev.list[[w]],Time=testtimes)
    
    basic.event.list$Events<-as.character(basic.event.list$Events)
    basic.event.list$Events<-factor(basic.event.list$Events, levels=as.character(event.types))
    
    #turn event types into numeric values
    
    basic.event.list$Events<-as.numeric(basic.event.list$Events)
    
    basic.event.list$Events[basic.event.list$Events>endogenous]<-0
    
    evlist<-list()

    #convert from dataframes to matrices
    
    evlist[[1]]<-as.matrix(basic.event.list)
    
  
    #remove the endogenous types from the event.types list
    
    event.types<-event.types[1:endogenous]
    
    
    ###other important objects
    
    ## format female exact time to use in simulation. Females will flip from present to absent at the average #time of first departure
fe_exact<-read.table(file=here("data_files","CHG_females_exact.tsv"), header=TRUE,sep="\t",stringsAsFactors=FALSE)

#reduce to only the necessary columns- this time I want time elapsed
fe_exact<-fe_exact[,c(1,3,5,7,9,11,13)]

#turn the first arrival NA's to 0s
fe_exact$first_arrival_elapsed[is.na(fe_exact$first_arrival_elapsed)]<-"00:00:00:00"

library(lubridate)
options(digits.secs=6)

#convert number of frames to partial seconds
for(h in 2:7){
fe_exact[,h]<-sapply(fe_exact[,h], function(x) {

fem.split<-unlist(strsplit(x,split=":", fixed=TRUE))
suppressWarnings(period_to_seconds(hms(paste(fem.split[1],fem.split[2],(as.numeric(fem.split[3])+as.numeric(fem.split[4])/30), sep=":"))))

}
)
}


#bring in day-level male presence data
male.pres<-read.table(here("data_files","CHG_2014_male_presence_data_quest.csv"), sep=",", header=T)
#male IDs are already in ascending order, so I can convert them directly
colnames(male.pres)<-c("Date", 1:30)

#alter to restrict to only six males (the territory holders for that day)
 male.pres[1,c(7,9:31)]<-0
 ##create pres.exact data

pres.test<-data.frame(Events=c("1.arrive","2.arrive","3.arrive","4.arrive","5.arrive","7.arrive"),Time=0,Date="2014-03-19",Bird_ID=c(1:5,7),Type="arrive")

 pres.test$Bird_ID<-factor(pres.test$Bird_ID,levels=as.character(1:30))
 pres.test$Type<-factor(pres.test$Type,levels=c("arrive","depart"))

pres.exact.dat<-list(pres.test)
  
  
  
### pairwise centroid distances

#this time, imagine they are all somehow equidistant- some sort of 3D space-lek? Or, rather, I'm focusing on male 1, so I can just say all the males are the same distance from him. I dont really have to worry about the rest. 
  
#the mean distance between any pair of males is:


file_list<-list.files(paste(here(),"/data_processing/udata_centroid_data/pairwise_dist",sep=""))

distlist<-list()

distlist<-lapply(file_list, function(x) read.table(paste(here(),"/data_processing/udata_centroid_data/pairwise_dist/",x,sep=""), sep=","))


#convert to martices

distlist.mat<-lapply(distlist, function(x) as.matrix(x, ))

#the names of the rows and columns are in the same order as the numerical ids passed to rem, so they can be used to index the matrix. Males that are absent on a given day have NA's for their centroid distances. I can change them to zeroes here so they don't produce errors, but the support array will take care of them anyway.

for(i in 1:length(distlist.mat)){
distlist.mat[[i]][is.na(distlist.mat[[i]])]<-0
}


#distance between every pair of territory-holding males for the whole season

meandist<-mean(unlist(lapply(distlist.mat,function(a){
b<-a[c(1,2,3,4,5,7),c(1,2,3,4,5,7)]
mean(b[upper.tri(b)][b[upper.tri(b)]>0])
}))) 

#6.539949

##change the data for the first day

distlist.mat[[1]][1,2:7]<-meandist
distlist.mat[[1]][2:7,1]<-meandist

##so now male 1 is equidistant from all his competitors


    #### create statlist ####
    
    ##"a three dimensional array, event number by event type by statistic, or a list thereof, containing the sufficient statistics for the model"
    
    #start with the intercept sufficient stats
    
    #seems like the statlist involves an array with an ojbect for every "group" according to grouping factor. In the example it's the individuals but for me it's the different DAYS. each day will have a matrix for each event in the sequence, and for the intercepts, each matrix will just be an identity matrix of event types... I think. give it a try. This, I think, should EXCLUDE rows for all the exogenous event types as well (except end.obs which is still in there). I know this isn't what Carter would prefer, but it's just adding dead events that we basically tell the model to ignore anyway. 
    
    
    sequences<-table(event.list$Date)
    
    #create list of dummy intercept names

idno<-length(event.types[grep("cop.start",event.types)])

dummy.ints.attack<-paste(1:14,".attackFE", sep="")
dummy.ints.receive<-paste(1:13, ".receiveFE", sep="")

IDintsCH<-paste(1:14,"initCH",sep=".")
IDdis<-paste(1:14,"disengage",sep=".")

int.names<-c("Global.attackFE", "X.receiveFE", "X.UreceiveFE", "U.attackFE", "U.receiveFE", "U.UattackFE", dummy.ints.attack, dummy.ints.receive, "cop.startFE","2.cop.start","3.cop.start","4.cop.start","cop.endFE", "initFO.FE","ID.initFO.FE", "U.initFO.FE","initSF.FE","X.initSF.FE", "initCH.FE",  "U.initCH.FE", IDintsCH, "disengage.FE", "U.disengage.FE", IDdis, "depart.FE", "X.depart.FE")

int.list<-list()

for(i in 1:length(levels(event.list$Date))){

day.array<-array(0,dim=c(as.numeric(sequences[i]), length(event.types),length(int.names)))

#the third dimension is the stat slice.
dimnames(day.array)<-list(c(1:sequences[i]),as.character(event.types),as.character(int.names))

#add global attack effect, loops for "U" intercepts for attacks and receives, then one loop for typical intercepts

day.array[,grep("attack", event.types),1]<-1


for(h in 1:14){
day.array[,paste(h,".attack.",30, sep=""),2]<-1
}


for(h in 15:29){
day.array[,paste(h,".attack.",30, sep=""),3]<-1
}


for(h in 1:14){
	for(j in 15:29){	
day.array[,paste(j,".attack.", h, sep=""),4]<-1
}
}


for(h in 1:14){
	for(j in 15:29){
day.array[,paste(h,".attack.",j, sep=""),5]<-1
}
}

for(h in 15:29){
	for(j in 15:29){	
if(h!=j){
day.array[,paste(j,".attack.", h, sep=""),6]<-1
}
}
}

preloop<-6

#loop for dummy intercepts for attacks, loop for dummy intercepts for receives, add in cop.start and cop.end FEs, then one loop for typical intercepts

for(k in (preloop+1):(length(dummy.ints.attack)+preloop)){
	
	for(h in 1:14){
if(h!=(k-preloop)){
day.array[,paste((k-preloop),".attack.", h, sep=""),k]<-1
}
}
}

for(k in ((length(dummy.ints.attack)+preloop)+1):((length(dummy.ints.attack)+preloop)+length(dummy.ints.receive))){
	for(h in 1:14){
if(h!=(k-(length(dummy.ints.attack)+preloop))){
day.array[,paste(h,".attack.",(k-(length(dummy.ints.attack)+preloop)), sep=""),k]<-1
}
}
}

#cop.startFE
for(h in (1:idno)[-c(2,3,4)]){
day.array[,paste(h,".cop.start", sep=""),((length(dummy.ints.attack)+preloop)+length(dummy.ints.receive)+1)]<-1
}

#2.cop.start
day.array[,paste(2,".cop.start", sep=""),((length(dummy.ints.attack)+preloop)+length(dummy.ints.receive)+2)]<-1

#3.cop.start
day.array[,paste(3,".cop.start", sep=""),((length(dummy.ints.attack)+preloop)+length(dummy.ints.receive)+3)]<-1

#4.cop.start
day.array[,paste(4,".cop.start", sep=""),((length(dummy.ints.attack)+preloop)+length(dummy.ints.receive)+4)]<-1

#cop.endFE
for(h in 1:idno){
day.array[,paste(h,".cop.end", sep=""),((length(dummy.ints.attack)+preloop)+length(dummy.ints.receive)+5)]<-1
}

#initFO.FE
for(h in 1:idno){
day.array[,paste(h,".initFO", sep=""),((length(dummy.ints.attack)+preloop)+length(dummy.ints.receive)+6)]<-1
}

#ID.initFO.FE
for(h in 1:14){
day.array[,paste(h,".initFO", sep=""),((length(dummy.ints.attack)+preloop)+length(dummy.ints.receive)+7)]<-1
}

#U.initFO.FE
for(h in 15:29){
day.array[,paste(h,".initFO", sep=""),((length(dummy.ints.attack)+preloop)+length(dummy.ints.receive)+8)]<-1
}

#initSF.FE
for(h in 1:idno){
day.array[,paste(h,".initSF", sep=""),((length(dummy.ints.attack)+preloop)+length(dummy.ints.receive)+9)]<-1
}

#X.initSF.FE
day.array[,paste(30,".initSF", sep=""),((length(dummy.ints.attack)+preloop)+length(dummy.ints.receive)+10)]<-1


#initCH.FE
for(h in 1:idno){
day.array[,paste(h,".initCH", sep=""),((length(dummy.ints.attack)+preloop)+length(dummy.ints.receive)+11)]<-1
}


#U.initCH.FE
for(h in 15:29){
day.array[,paste(h,".initCH", sep=""),((length(dummy.ints.attack)+preloop)+length(dummy.ints.receive)+12)]<-1
}


#individual initCH effects for ID male
for(k in (((length(dummy.ints.attack)+preloop)+length(dummy.ints.receive)+13):((length(dummy.ints.attack)+preloop)+length(dummy.ints.receive)+13+(length(IDintsCH)-1)))){
	
	day.array[,int.names[k],k]<-1
	
}

preloop2<-((length(dummy.ints.attack)+preloop)+length(dummy.ints.receive)+13+(length(IDintsCH)-1))


#disengage.FE
for(h in 1:idno){
day.array[,paste(h,".disengage", sep=""),preloop2+1]<-1
}

#U.disengage.FE
for(h in 15:29){
day.array[,paste(h,".disengage", sep=""),preloop2+2]<-1
}

#individual disengage effect for ID males
for(k in (preloop2+3):(preloop2+3+length(IDdis)-1)){
	
	day.array[,int.names[k],k]<-1
	
}


preloop3<-(preloop2+3+length(IDdis)-1)

#depart.FE
for(h in 1:idno){
day.array[,paste(h,".depart", sep=""),preloop3+1]<-1
}

#X.depart.FE
day.array[,paste(30,".depart", sep=""),preloop3+2]<-1


#weird extra formatting to make it match the package example, seems like it becomes relevant *punintended!* if I want to fit day-specific effects

global<-list()

global[[1]]<-day.array

int.list[[i]]<-global

names(int.list[[i]])<-"global"

}
    
    
    
    
    
    
    
    
    #### Create Support Matrix ####
    
    #"should be a list withone element per history, each of which should be an event order by event type logical matrix. the ijth cell of this matrix should be TRUE if an event type of j was a possible next event given the preceding code i-1 events, and FALSE otherwise"
    
    #need a way to ask, "what is each male doing right now" for each event in the sequence. That dictates the possibilities. 
    
    #create a bookkeeping "census" matrix to keep track
    
    
    
    census.list<-list()
    
    
    for(j in 1:length(event.list.dat)){
      
      
      if(j<mod.day){
        
        census<-matrix("nothing", nrow=1,ncol=(idno), dimnames=list(1,c(1:(idno))))
        
        
        ##code that translates each new current event into what the males involved are doing at that time. The support matrix will be based on whatever the males were doing in the PREVIOUS timestep. This census matrix itself could be interesting to visualize. Each row of the census says what each male is doing going into the NEXT event.
        
        
        for(i in 1:nrow(event.list.dat[[j]])){
          
          #can ask if the second thing is something, and then move on to the first and third elements.
          
          current<-census[nrow(census),]
          
          #parse the event type. loop through test.seq rows
          evname<-as.character(event.list.dat[[j]][i,1])
          evsplit<-strsplit(evname, split=".", fixed=T)
          
          #if it is an end obs, arrival, departure or clock event, it shouldn't change anything the males are doing in terms of interactions. Departures aren't exogenous, but they shouldn't change anything since they can only happen once a male is already doing nothing. 
          if(evsplit[[1]][2]!="obs" & evsplit[[1]][2]!="depart" & evsplit[[1]][2]!="arrive" & evsplit[[1]][2]!="time" & evsplit[[1]][1]!="fem"){
            #if the event was an attack, indicate the males involved and which was the receiver since this dictates who makes the initition decision
            if(evsplit[[1]][2]=="attack"){
              current[evsplit[[1]][1]]<-evname
              current[evsplit[[1]][3]]<-paste(evsplit[[1]][3], ".", "received", ".", evsplit[[1]][1], sep="")
            }
            #if it's an FO or SF, just indicate that that pair of males are involved in a fight now
            if(evsplit[[1]][2]=="initFO"||evsplit[[1]][2]=="initSF"){
              #need to refer back to the previous row of "census" to get the id of the other male
              
              current[evsplit[[1]][1]]<-paste(evsplit[[1]][1], ".", "fight", ".",strsplit(census[nrow(census),evsplit[[1]][1]], split=".",fixed=T)[[1]][3], sep="")
              current[[strsplit(census[nrow(census),evsplit[[1]][1]], split=".",fixed=T)[[1]][3]]]<-paste(strsplit(census[nrow(census),evsplit[[1]][1]], split=".",fixed=T)[[1]][3], ".", "fight", ".",evsplit[[1]][1], sep="")
              
            }
            
            #if it's a chase, indicate that the pair of males are involved in a fight, but there is an additional constraint that the individual DOING THE CHASING will be the disengager by definition
            if(evsplit[[1]][2]=="initCH"){
              
              current[evsplit[[1]][1]]<-paste(evsplit[[1]][1], ".", "fight", ".",strsplit(census[nrow(census),evsplit[[1]][1]], split=".",fixed=T)[[1]][3], ".chasedby",sep="")
              current[[strsplit(census[nrow(census),evsplit[[1]][1]], split=".",fixed=T)[[1]][3]]]<-paste(strsplit(census[nrow(census),evsplit[[1]][1]], split=".",fixed=T)[[1]][3], ".", "fight", ".",evsplit[[1]][1], ".chasing",sep="")
              
            }
            
            #for disengaging, end the fight for the partner male as well
            if(evsplit[[1]][2]=="disengage"){
              current[evsplit[[1]][1]]<-"nothing"
              current[evsplit[[1]][3]]<-"nothing"	
              
            }
            
            
            #for copulations, end cops are only possible after start cops, but otherwise anything else is possible. EXCEPT the copulating male can't initiate an attack. If the next event involving that male ISN'T an end cop, it was interrupted
            
            if(evsplit[[1]][2]=="cop" && evsplit[[1]][3]=="start"){
              current[evsplit[[1]][1]]<-"copulating"
            }
            
            if(evsplit[[1]][2]=="cop" && evsplit[[1]][3]=="end"){
              current[evsplit[[1]][1]]<-"nothing"
            }
            
            #if an X male is still involved in a fight at all, make sure he is still tagged as fighting (since sometimes more than one is interacting at once, but they can't depart until none are fighting)
            if((TRUE %in% grepl("fight.30",current) | TRUE %in% grepl("30.fight",current)) && !(grepl("receive", current[idno]) | grepl("attack", current[idno]))) {
              
              current[idno]<-"fight"
              
            }
            
          }
          
          census<-rbind(census,current)
          
          
        }
        
        
        #don't want the first dummy row
        census.list[[j]]<-census[-1,]
        
      } else {
        
        
        census<-matrix("nothing", nrow=1,ncol=(idno), dimnames=list(1,c(1:(idno))))
        
        
        ##code that translates each new current event into what the males involved are doing at that time. The support matrix will be based on whatever the males were doing in the PREVIOUS timestep. This census matrix itself could be interesting to visualize. Each row of the census says what each male is doing going into the NEXT event.
        
        
        for(i in 1:nrow(event.list.dat[[j]])){
          
          #can ask if the second thing is something, and then move on to the first and third elements.
          
          current<-census[nrow(census),]
          
          #parse the event type. loop through test.seq rows
          evname<-as.character(event.list.dat[[j]][i,1])
          evsplit<-strsplit(evname, split=".", fixed=T)
          
          #if it is an end obs, arrival, departure or clock event, it shouldn't change anything the males are doing in terms of interactions. Departures aren't exogenous, but they shouldn't change anything since they can only happen once a male is already doing nothing. 
          if(evsplit[[1]][2]!="obs" & evsplit[[1]][2]!="depart" & evsplit[[1]][2]!="arrive" & evsplit[[1]][2]!="time" & evsplit[[1]][1]!="fem"){
            #if the event was an attack, indicate the males involved and which was the receiver since this dictates who makes the initition decision
            if(evsplit[[1]][2]=="attack"){
              current[evsplit[[1]][1]]<-evname
              current[evsplit[[1]][3]]<-paste(evsplit[[1]][3], ".", "received", ".", evsplit[[1]][1], sep="")
            }
            #if it's an FO or SF, just indicate that that pair of males are involved in a fight now
            if(evsplit[[1]][2]=="initFO"||evsplit[[1]][2]=="initSF"){
              #need to refer back to the previous row of "census" to get the id of the other male
              
              current[evsplit[[1]][1]]<-paste(evsplit[[1]][1], ".", "fight", ".",strsplit(census[nrow(census),evsplit[[1]][1]], split=".",fixed=T)[[1]][3], sep="")
              current[[strsplit(census[nrow(census),evsplit[[1]][1]], split=".",fixed=T)[[1]][3]]]<-paste(strsplit(census[nrow(census),evsplit[[1]][1]], split=".",fixed=T)[[1]][3], ".", "fight", ".",evsplit[[1]][1], sep="")
              
            }
            
            #if it's a chase, indicate that the pair of males are involved in a fight, but there is an additional constraint that the individual DOING THE CHASING will be the disengager by definition
            if(evsplit[[1]][2]=="initCH"){
              
              current[evsplit[[1]][1]]<-paste(evsplit[[1]][1], ".", "fight", ".",strsplit(census[nrow(census),evsplit[[1]][1]], split=".",fixed=T)[[1]][3], ".chasedby",sep="")
              current[[strsplit(census[nrow(census),evsplit[[1]][1]], split=".",fixed=T)[[1]][3]]]<-paste(strsplit(census[nrow(census),evsplit[[1]][1]], split=".",fixed=T)[[1]][3], ".", "fight", ".",evsplit[[1]][1], ".chasing",sep="")
              
            }
            
            #for disengaging, need to search through the previous rows to find the most recent "fight" and identify the partner and end it
            if(evsplit[[1]][2]=="disengage"){
              current[evsplit[[1]][1]]<-"nothing"
              current[strsplit(census[max(grep("fight", census[,evsplit[[1]][1]])),evsplit[[1]][1]], split=".", fixed=T)[[1]][3]]<-"nothing"	
              
            }
            
            
            #for copulations, end cops are only possible after start cops, but otherwise anything else is possible. EXCEPT the copulating male can't initiate an attack. If the next event involving that male ISN'T an end cop, it was interrupted
            
            if(evsplit[[1]][2]=="cop" && evsplit[[1]][3]=="start"){
              current[evsplit[[1]][1]]<-"copulating"
            }
            
            if(evsplit[[1]][2]=="cop" && evsplit[[1]][3]=="end"){
              current[evsplit[[1]][1]]<-"nothing"
            }
            
            # #if an X male is still involved in a fight at all, make sure he is still tagged as fighting (since sometimes more than one is interacting at once, but they can't depart until none are fighting)
            # if((TRUE %in% grepl("fight.30",current) | TRUE %in% grepl("30.fight",current)) && !(grepl("receive", current[idno]) | grepl("attack", current[idno]))) {
            
            # current[idno]<-"fight"
            
            # }
            
          }
          
          census<-rbind(census,current)
          
          
        }
        
        #don't want the first dummy row
        census.list[[j]]<-census[-1,]
        
      }
      
    }
    
    
    #convert to win.loss census
    
    create.win.loss.census.sim<-function(x){
      
      #get disengagement events for focal male from evlist
      win.loss.census<-x
      
      #loop thru days and males
      
      for(i in 1:length(win.loss.census)){
        
        for(l in 1:idno){
          
          dis.times<-grep(TRUE,startsWith(as.character(event.list.dat[[i]][,1]), paste(l,".disengage", sep="")))
          
          
          if(length(dis.times)>0){
            
            #see who they were fighting with
            fights<-win.loss.census[[i]][dis.times-1,l]
            
            #so I have a list of row numbers for disengagements and a list of interaction partners. This will cover wins for normal fights and losses for chases for the focal male. If I loop through all the males, all the wins and losses should get filled in.
            
            #NOW DISENGAGES ALWAYS MEAN WINS, so I will still have chase information, but I don't need to do anything special with it here.
            
            for(j in 1:length(dis.times)){
              
              if(grepl("chasing",fights[j]) ==FALSE){
                #apply the win in the focal male's column
                win.loss.census[[i]][dis.times[j],l]<-paste(l, "win",strsplit(fights[j],".", fixed=T)[[1]][3], sep=".")
                #apply the loss for the same interaction in the partner male's column
                win.loss.census[[i]][dis.times[j],as.numeric(strsplit(fights[j],".", fixed=T)[[1]][3])]<-paste(strsplit(fights[j],".", fixed=T)[[1]][3], "lose", l, sep=".")
              } else{
                #apply the won chase in the focal male's column
                win.loss.census[[i]][dis.times[j],l]<-paste(l, "win",strsplit(fights[j],".", fixed=T)[[1]][3],"chase", sep=".")
                #apply the loss for the same interaction in the partner male's column.
                win.loss.census[[i]][dis.times[j],as.numeric(strsplit(fights[j],".", fixed=T)[[1]][3])]<-paste(strsplit(fights[j],".", fixed=T)[[1]][3], "lose", l, "chase",sep=".")
              }
              
            }
            
          }
          
        }
        
      }
      
      return(win.loss.census)
      
    }
    
    win.loss.census<-create.win.loss.census.sim(census.list)
    

    
    
    
    ## Constraints to incorporate in the support matrix ##
    
    #only males that are actually present can be involved in interactions on a given day and at a given time
    
    #If "nothing" is happening, all "attacks", all start cops and all departs (given the male is actually present) are possible. Males cannot disengage or initiate fights without an attack happening first. End cop cannot happen without a previous start cop. end.obs is possible. (end.obs is not possible if any males are still fighting or copulating, so end.obs is ONLY possible when "nothing" is going on across the board)
    
    #if the previous event was an "attack", the current event MUST be an initiation by the receiver, since we are assuming the attack is essentually instantaneous. 
    
    #once initiatiated, subsequent events can only be a disengage from either male involved or any other event NOT invoving the interaction partners. Males actively fighting connot become engaged in a new fight until they disengage. 
    
    #if the initiated fight was a chase only the male being chased can disengage, by definition
    
    #Once a copulation starts, the only thing that male can do next is either end the copulation OR receive an attack. He cannot initiate anything. 
    
    #copulations cannot occur when there are no females present
    
    #a male can ONLY "depart" if "nothing" is happening
    
    #Exceptions to these rules are for "X" males- there may be more than one unidentifiable male on the lek at a time. Right now we are dealing with this by having "X" males just be a mass actor category. 1) These males CAN do an "attack" before disengaging from an interaction. They can also be attacked. However, normal restrictions apply to the partner male. So if an X male is being chased, "disengage" becomes possible, but any "attack" (or cop.start, or depart) are still also possible 
    
    
    ### Build the actual support matrix ###
    
    #for each event, there is a list of event types, with 1's for the types that are possible and 0's for types that are not. 
    
    
    
    #code block to produce the support matrix
    
    supplist<-list()
    
    
      
    #make a reference event list with data (most importantly female presence)
    evlist.dat<-event.list.dat
    
    
    
    for(j in 1:length(evlist)){
      
      supplist[[j]]<-matrix(1, nrow=length(evlist[[j]][,1]), ncol=length(event.types))
      colnames(supplist[[j]])<-event.types
      
      #turn off all interactions involving males not present on that day
      #make a list of all the event types that include the males that were entirely absent and remove them for all rows
      
      supplist[[j]][,unique(unlist(lapply(grep(0,male.pres[j,2:(idno+1)]), function(x) c(event.types[startsWith(event.types, paste(x, ".", sep=""))], event.types[endsWith(event.types, paste(".", x, sep=""))]))))]<-0
      
      
      #first row can be any "attack" or, technically, any "depart"
      #since now fem.enter is an actual event, cop.start is never possible as the very first thing
      supplist[[j]][1,][-c(grep("attack", event.types), grep("depart", event.types))]<-0
      
      
      #the exception is events involving males that will be present that day but haven't arrived yet
      
      
      #basically I can make this code exactly the same as below- just want to ask who is actually there right now? 
      
      arrsub<-subset(pres.exact.dat[[j]],Time<evlist[[j]][1,2] | Time==0 )
      arrtab<-table(arrsub$Bird_ID,arrsub$Type)
      
      arrpat<-apply(arrtab,1,function(x) sum(x))
      
      #if the total number of previous arrives and departs is even, that male is currently absent
      absent<-names(arrpat[(arrpat %% 2)==0])
      
      if(length(absent)>0){
        supplist[[j]][1,][unique(c(sapply(absent, function(x) c(event.types[startsWith(event.types, paste(x, ".", sep=""))], event.types[endsWith(event.types, paste(".", x, sep=""))]))))]<-0
        
      } 
      
      
      
      #loop through the rest of the events: 
      
      for(i in 2:(length(evlist[[j]][,1])-1)){
        
        
        #before anything else turn off events involving males that have either not yet arrived or have lef the lek early and are therefore no longer present that day or are otherwise not currently present
        
        
        arrsub<-subset(pres.exact.dat[[j]],Time<evlist[[j]][i,2])
        arrtab<-table(arrsub$Bird_ID,arrsub$Type)
        
        arrpat<-apply(arrtab,1,function(x) sum(x))
        
        #if the total number of previous arrives and departs is even, that male is currently absent
        current.row.abs<-names(arrpat[(arrpat %% 2)==0])
        
        
        if(length(current.row.abs)>0){
          supplist[[j]][i,][unique(c(sapply(current.row.abs, function(x) c(event.types[startsWith(event.types, paste(x, ".", sep=""))], event.types[endsWith(event.types, paste(".", x, sep=""))]))))]<-0
          
        } 
        
        
        
        #determine if the previous event was an attack, since this dictates exactly what the next event can be: receiver initiates a type of fight
        if((TRUE %in% grepl("received", census.list[[j]][i-1,]))==TRUE){
          
          supplist[[j]][i,][-grep(TRUE, startsWith(event.types, paste(grep("received", census.list[[j]][i-1,]), ".", "init", sep="")))]<-0}
        
        
        #(else) if the previous event wasn't an attack, the next event CAN'T be an initiation and the possibilities depend on what everyone is currently engaged in
        
        
        if((TRUE %in% grepl("received", census.list[[j]][i-1,]))==FALSE){
          
          supplist[[j]][i,][grep("init", event.types)]<-0
          
          #if males are actively fighting, they can't attack or be attacked, or actually do anything else before disengaging (unless they are a ?? male, id 30)
          
          if((TRUE %in% grepl("fight", census.list[[j]][i-1,]))==TRUE){
            
            for(k in grep("fight", census.list[[j]][i-1,])){
              
              if(k!=idno){
                
                supplist[[j]][i,][c(event.types[startsWith(event.types, paste(k, ".attack.", sep=""))], event.types[endsWith(event.types, paste("attack.", k, sep=""))], event.types[startsWith(event.types, paste(k, ".cop", sep=""))])]<-0
                
              }
              
            }
            
            #ONLY those fighting can disengage. retain males that are fighting and remove the rest
            dis<-event.types[grep("disengage",event.types)]
            supplist[[j]][i,][dis[-grep("fight", census.list[[j]][i-1,])]]<-0
            
            #"chasing" and "chasedby" limit the possibilities further. Males getting chased DO NOT disengage by definition, again, with the exception of ?? males. This is taken care of by having ?? males just be "fight" -ing without any extra tags
            if(TRUE %in% grepl("chasedby", census.list[[j]][i-1,])){
              supplist[[j]][i,][dis[grep("chasedby", census.list[[j]][i-1,])]]<-0
            }
            
          }
          
          #if no one is fighting, no one can disengage, but all attack combinations are possible
          if((TRUE %in% grepl("fight", census.list[[j]][i-1,]))==FALSE){
            
            supplist[[j]][i,][grep("disengage",event.types)]<-0
            
          }
          
          
          #actively copulating males: They cannot cop.start. ONLY they can cop.end. they CAN be attacked but they CANNOT attack anyone
          if((TRUE %in% grepl("copulating", census.list[[j]][i-1,]))==TRUE){
            
            cop<-grep("copulating", census.list[[j]][i-1,]) 
            
            #stuff that needs to be turned off for only copulating males
            for(h in cop){
              
              supplist[[j]][i,][c(event.types[startsWith(event.types ,paste(h, ".attack.", sep=""))], event.types[startsWith(event.types, paste(h, ".cop.start", sep=""))])]<-0
              
            }
            
            #stuff that needs to be on for ONLY copulating males
            copendall<-event.types[grep("cop.end",event.types)]
            supplist[[j]][i,][copendall[-cop]]<-0
          }
          
          #if no one is copulating, all cop.starts are possible, but cop.ends are not
          
          if((TRUE %in% grepl("copulating", census.list[[j]][i-1,]))==FALSE){
            
            copendall<-event.types[grep("cop.end",event.types)]
            supplist[[j]][i,][copendall]<-0
            
          }
          
          
        }
        
        #if there are no females present, the next event could not have possibly been a cop.start
        
        if(evlist.dat[[j]][i-1,]$Females==1){
          
          copstartall<-event.types[grep("cop.start",event.types)]
          supplist[[j]][i,][copstartall]<-0
          
        }
        
        #Finally, A male can only "depart" if he was already doing "nothing," so depart must be turned off for everyone doing not-nothing. (also he has to be present, but I've already taken care of that above)
        
        noth<-as.character(which(census.list[[j]][i-1,]!="nothing"))
        
        if(length(noth)>0){
          
          supplist[[j]][i,][paste(noth, ".depart", sep="")]<-0
          
        }
        
      }
      #end events loop here
      
      #after done with all the middle events: onset of end.obs could depend on a lot of factors; not sure I want to fit it as part of the model. Just always say the last row must be end.obs (all endogenous events are no longer possible) 
      
      supplist[[j]][length(evlist[[j]][,1]),]<-0
      
    }
    #end day loop here
    


  test.stat<-int.list
  
  
  


  

  
  
  #bring in the function codes from the best result
  
  prev.best<-read.table(paste(here(),"/cluster_optimization/Attractiveness_model/previous_best_remove_226_at_the_end",sep=""), sep=",")
  
  prev.best.top<-as.character(prev.best[nrow(prev.best),2])
  args.all=as.numeric(unlist(strsplit(prev.best.top, split="_", fixed=TRUE)))
  
  arg.use<-c(1:length(args.all))
  
  for(i in 1:length(arg.use)){
    
    test.stat<-get(fun.list[args.all[i]])(test.stat)	
    
  }
  


#ok great!


#### Coefficients for calculating hazards

#need to bring in the coefficients from the model fit to the empirical data
  
coefs<-read.table(paste(here(),"/cluster_optimization/Attractiveness_model/rem_model_coef",sep=""), sep=",")$x

names(coefs)<-row.names(read.table(paste(here(),"/cluster_optimization/Attractiveness_model/rem_model_coef",sep=""), sep=","))

##First try: remove variability, see how it looks. Then maybe just use the original coefs?

#remove variability among the males

##fitted Global attack rate from the intercept model- this is the average attack rate across all males
#-7.763714

#I can try just putting this coefficient for global rate, and then removing the columns for the other coeffs?

newcoefs<-c(coefs["Global.attackFE"],coefs[which(names(coefs)=="cop.startFE"):length(coefs)])

newcoefs[1]<--7.763714


newint<-test.stat

#confirmed this way of adjusting the stats array for different sets of intercepts works fine
newint[[1]][[1]]<-test.stat[[1]][[1]][,,names(newcoefs)]

#dummy data output vector
output<-rep(0,length(event.types))

for(i in 1:nrow(evlist[[1]])){
	
   stats<-newint[[1]][[1]][i,,]
    
    #the coefficient vector gets multiplied by each row and the supplist vector gets multiplied by each column
 
    statsup<-apply(stats,2,function(x) x*supplist[[1]][i,])
    
    
    #for some reason the apply function across the row margin returns a transformed matrix, so I transform it back for sanity. Use the subset of coefficents for the predictors that are acutally included for this timestep
    statcoef<-t(apply(statsup,1,function(x) x*newcoefs[dimnames(stats)[[2]]]))
    
    dimnames(statcoef)[[1]]<-event.types
    
    #get the parameter-weighted sums of stats for each event type
    log.hazards<-apply(statcoef,1,function(x) sum(x))
 
	output<-rbind(output,log.hazards)
}
 
 
 #remove dummy row after

output<-output[-1,]
 
output<-as.matrix(output)

output.df<-data.frame(output,stringsAsFactors=F)
colnames(output.df)<-event.types

 #the hazard of male 1 getting attacked is going to the sum of the hazard of all the events representing possible attacks against him

one.rec<-vector()

for(i in 1:nrow(evlist[[1]])){ 
 step<-i
 
 rec.haz<-output.df[step,event.types[endsWith(event.types,"attack.1")]][output.df[step,event.types[endsWith(event.types,"attack.1")]]!=0]

total.receive<-sum(exp(rec.haz))

one.rec[step]<-total.receive
}

testout[[w]]<-one.rec

}


one.out<-rbind(one.out,testout[[1]][c(2,6)])
cop.out<-rbind(cop.out,testout[[2]][c(2,6)])

}

  #remove dummy row
  	one.out<-one.out[-1,]
	cop.out<-cop.out[-1,]

	mod<-data.frame(base=one.out[,1],attack=one.out[,2],inter=cop.out[,2])

data<-bind_cols(tests,mod)

type<-paste(data$initiate,data$outcome,sep="+")
type[1]<-"receive+lose\n(chase)"

data<-data %>%
mutate(type=type) %>%
mutate(diff=attack-base,perc.diff=(attack-base)/base) %>%
mutate(type=fct_reorder(type,diff))
  
  #calculate interruption risk
  
  endhaz<-exp(coefs["cop.endFE"])

data<-mutate(data,risk=100*inter/(inter+endhaz))

#   initiate outcome kind         base       attack      inter                  type         diff  perc.diff      risk
# 1  receive    lose   CH 0.0003546015 0.0034230047 0.15420633 receive+lose\n(chase) 3.068403e-03 8.65310159 39.612284
# 2  receive    lose   SF 0.0003546015 0.0011776544 0.05305332          receive+lose 8.230528e-04 2.32106383 18.412586
# 3  receive     win   SF 0.0003546015 0.0010501261 0.04730817           receive+win 6.955246e-04 1.96142577 16.752719
# 4   attack    lose   SF 0.0003546015 0.0009056811 0.04080093           attack+lose 5.510796e-04 1.55408118 14.789173
# 5   attack     win   SF 0.0003546015 0.0003681356 0.01658451            attack+win 1.353409e-05 0.03816704  6.589853

library(ggplot2)

#fix formatting later

newplot<-ggplot()+geom_bar(data=data,aes(x=type,y=diff,fill=risk),stat="identity")+scale_fill_gradient(name="Risk of Copulation Interruption (%)", low="blue",high="red")+theme_classic()+theme(axis.line.x=element_line(colour="black",size=1.5, linetype="solid"),axis.line.y=element_line(colour="black",size=1.5, linetype="solid"), axis.text.x=element_text(size=14,face="bold"), axis.text.y=element_text(size=18,face="bold"), axis.title.x=element_text(size=25,face="bold"), axis.title.y=element_text(size=25,face="bold"),legend.position="top",legend.title=element_text(size=20),legend.text=element_text(size=18),legend.key.size=unit(0.75,"cm"),legend.key.width=unit(1.5,"cm"))+labs(x="Interaction Type",y="Increase in Future Hazard \n of Getting Attacked")

setwd(paste(here(),"/adequacy_checking/adequacy_attractiveness_model",sep=""))

ggsave("fight_effect_fig2.pdf",newplot,height=7,width=10,units=c("in"))

#could consider doing a factorial thing to better show the effect of receive and lose independently

#ggplot(data)+geom_point(aes(x=1,y=1,size=perc.diff,colour=perc.diff))+facet_grid(initiate~outcome+kind)

# > one.rec[c(2,5)]
# [1] 0.0003546015 0.0011781518
```
