## Supplemental_reproducible_code for "Fighting isn’t sexy in lekking Greater Sage-grouse (*Centrocercus urophasianus*)": time_fighting.R

```
## Code for calculating male's time available to mate (not fighting), time fighting, and time present in front of females ##

rm(list=ls())

###make sure necessary packages are there and loaded

packages<-c("here","abind","relevent","lubridate","DescTools","gtools","e1071","ggplot2","magrittr","rcartocolor","ggpubr","cowplot","dplyr","tidyverse","grid","gtable","gridExtra")

package.check<-lapply(packages, FUN = function(x) {
  if (!require(x, character.only = TRUE, quietly=TRUE)) {
    install.packages(x, dependencies = TRUE)
    library(x, character.only = TRUE, quietly=TRUE)
  } 
  
})

#for project path management
#define the project root directory relative to the current script. This sets the working directory that gets called by the here() function. It is a relative path and will be robust to use on anyone's computer. In this case, I want the root directory to always be the enclosing folder for the whole sage grouse project


i_am("adequacy_checking/adequacy_attractiveness_model/time_fighting.R")


#make the base event list and intercept array


event.list<-read.table(file=here("data_processing","relevent_format_dataset_udata_wincol_disturbance.tsv"), header=T,  sep="\t")

event.list$Date<-as.factor(event.list$Date)

event.types<-read.table(file=here("data_processing","event_types_list_udata.csv"), header=F,  sep=",")
event.types<-as.character(event.types$V1)

basic.event.list<-event.list

basic.event.list$Events<-sapply(basic.event.list$Events, function(x) {
  
  event<-unlist(strsplit(as.character(x),".",fixed=T))
  if(event[2]=="disengage"){
    return(paste(event[1],event[2],sep="."))
  } else {return(as.character(x))}
}
)


#add factor levels for event types that don't actually happen (and put them in the same order as the event.type list) to maintain bookkeeping 

basic.event.list$Events<-as.character(basic.event.list$Events)

basic.event.list$Events<-factor(basic.event.list$Events, levels=as.character(event.types))

#NOTE- the event.types vector now contains ENDOGENOUS event types as well as EXOGENOUS types: arrivals and clock events, and end.obs. Need to make sure everything is robust to this. 


### create eventlist for rem ###


event.list.dat<-split(event.list, event.list$Date)


####### Bring in hazards from the REM model ######


setwd(paste(here(),"/adequacy_checking/adequacy_attractiveness_model",sep=""))

#bring in the saved data
output.df<-read.table("per_event_adequacy_data_full", header=T,sep=",",stringsAsFactors=F)
output.sum<-read.table("per_event_adequacy_data_summary", header=T,sep=",",stringsAsFactors=F)


### Availability Plot ###

#want intervals for male presence, and then for fighting/not fighting. Can get both of these numbers from the hazards 

#I want the hazard of getting attacked is greater than zero, Since males CAN be attacked during copulation but not during a fight.  BUT*** I need to also remove times when females are absent, since this doesn't cover that. 

#dim(output.df)


#split by day

#add time, Females, and Events columns
output.time<-cbind(event.list[,c(1,2,3)],output.df)
colnames(output.time)[c(1,2,3)]<-c("Events","Time","Females")


#remove rows for "init"s since the hazard for everything else is zero at that point and it messes up the intervals- makes it seem like the male is available to mate while he is just getting started fighting. 


output.time<-output.time[-grep("init",event.list$Events),]


library(dplyr)


avail.df<-split(output.time,as.numeric(as.character(output.time$day)))

#remove columns for males that were absent that day, and other columns we don't need right now
avail.df.pres<-lapply(avail.df,function(z){
  cbind(z[,c(1,2,3)],z[,c(12:ncol(z))][,apply(z[,c(12:ncol(z))],2,function(x) sum(as.numeric(x))>0)])})

library(abind)

avail.df.interval<-lapply(avail.df.pres,function(z){
  
  availlist<-list()
  
  for(i in 1:30){
    
    copcol<-paste0("X",i,".cop.start")
    cols<-colnames(z)[endsWith(colnames(z),paste0("attack.",i))]
    
    #make sure that male was there
    if(copcol %in% dimnames(z)[[2]]){
      
      newavail<-z[,cols]
      
      newavail<-apply(newavail,1,function(x) sum(x))
      
      intervail<-newavail
      
      #if the hazard of being attacked is POSITIVE AND females are PRESENT, then they are available to mate in that interval
      
      #maybe just label each timepoint as available or not and then pare down to the transitions using lag or lead
      
      intervail[lead(newavail)>0 & z$Females==2]<-"avail"
      intervail[lead(newavail)==0 | (lead(newavail)>0 & z$Females==1)]<-"not"
      
      intervail[length(newavail)]<-"not"
      
      #check
      #cbind(cbind(cbind(intervail,newavail),z$Females),as.character(z$Events))
      
      #then identify the actual beginnings and ends of the intervals
      intervail.ends<-intervail
      
      intervail.ends[intervail.ends==lag(intervail.ends)]<-"drop"
      
      #since lag doesn't touch the first row
      if(intervail.ends[1]=="not"){intervail.ends[1]<-"drop"}
      
      #cbind(cbind(cbind(cbind(intervail.ends,intervail),newavail),z$Females),as.character(z$Events))
      
      lab<-rep(i,length(newavail))
      time<-z[,2]
      
      tempdat<-data.frame(avail=intervail,name=lab,time=time)
      
      #remove rows between interval termini
      tempdat<-tempdat[intervail.ends=="avail" | intervail.ends=="not",]	
      
      #turn into avail and not columns
      
      
      availlist[[i]]<-tempdat
      
    }
    
  }
  
  availday<-abind(availlist,along=1)
  
  availday<-data.frame(availday,stringsAsFactors=F)
  availday$time<-as.numeric(availday$time)
  
  return(availday)
})


#str(avail.df.interval[[13]])


availdat<-lapply(avail.df.interval, function(x){
  
  mutdat<-x[x$avail=="avail",][,c(2,3)]
  
  #match with interval ends
  mutdat$not<-x[x$avail=="not",][,3]
  
  colnames(mutdat)<-c("male","avail","not")
  mutdat$male<-factor(mutdat$male,levels=c(1:30))
  
  #restrict to territory-holders?
 # mutdat<-mutdat[(mutdat$male==1 | mutdat$male==2 | mutdat$male==3 | mutdat$male==4 | mutdat$male==7),]
  
  return(mutdat)
  
})

######and calculate the total numbers to display in a table/chart on the side

availdat.plot<-lapply(availdat, function(x){
  x$tot<-x$not-x$avail
  return(x)})

#get total amount of time actually available for mating

availability<-lapply(availdat.plot, function(y){
z<-y %>%
  group_by(male) %>%
  summarise(timeavail=sum(tot))
return(z)
})


#this will gives the total time available to mate for each male each day. I can get means for these. 


#bring in day-level male presence data
male.pres<-read.table(file=here("data_files","CHG_2014_male_presence_data_quest.csv"), sep=",", header=T)
#male IDs are already in ascending order, so I can convert them directly
colnames(male.pres)<-c("Date", 1:30)


####### Can I adapt the availability code that uses the hazards at each timestep to use the census data instead in the same way to get fighting times instead?

#this code finds the amount of time each male spent fighting in front of females each day. Total time in attendance in front of females should be the sum of this data and the time available to mate. 


census.list<-list()
idno<-30

for(j in 1:length(event.list.dat)){
    
    census<-matrix("nothing", nrow=1,ncol=(idno), dimnames=list(1,c(1:(idno))))
    
    
    ##code that translates each new current event into what the males involved are doing at that time. The support matrix will be based on whatever the males were doing in the PREVIOUS timestep. This census matrix itself could be interesting to visualize. Each row of the census says what each male is doing going into the NEXT event.
    
    
    for(i in 1:nrow(event.list.dat[[j]])){
      
      #can ask if the second thing is something, and then move on to the first and third elements.
      
      current<-census[nrow(census),]
      
      #parse the event type. loop through test.seq rows
      evname<-as.character(event.list.dat[[j]][i,1])
      evsplit<-strsplit(evname, split=".", fixed=T)
      
      #if it is an end obs, arrival, departure or clock event, it shouldn't change anything the males are doing in terms of interactions. Departures aren't exogenous, but they shouldn't change anything since they can only happen once a male is already doing nothing. 
      if(evsplit[[1]][2]!="obs" & evsplit[[1]][2]!="depart" & evsplit[[1]][2]!="arrive" & evsplit[[1]][2]!="time" & evsplit[[1]][1]!="fem"){
        #if the event was an attack, indicate the males involved and which was the receiver since this dictates who makes the initition decision
        if(evsplit[[1]][2]=="attack"){
          current[evsplit[[1]][1]]<-evname
          current[evsplit[[1]][3]]<-paste(evsplit[[1]][3], ".", "received", ".", evsplit[[1]][1], sep="")
        }
        #if it's an FO or SF, just indicate that that pair of males are involved in a fight now
        if(evsplit[[1]][2]=="initFO"||evsplit[[1]][2]=="initSF"){
          #need to refer back to the previous row of "census" to get the id of the other male
          
          current[evsplit[[1]][1]]<-paste(evsplit[[1]][1], ".", "fight", ".",strsplit(census[nrow(census),evsplit[[1]][1]], split=".",fixed=T)[[1]][3], sep="")
          current[[strsplit(census[nrow(census),evsplit[[1]][1]], split=".",fixed=T)[[1]][3]]]<-paste(strsplit(census[nrow(census),evsplit[[1]][1]], split=".",fixed=T)[[1]][3], ".", "fight", ".",evsplit[[1]][1], sep="")
          
        }
        
        #if it's a chase, indicate that the pair of males are involved in a fight, but there is an additional constraint that the individual DOING THE CHASING will be the disengager by definition
        if(evsplit[[1]][2]=="initCH"){
          
          current[evsplit[[1]][1]]<-paste(evsplit[[1]][1], ".", "fight", ".",strsplit(census[nrow(census),evsplit[[1]][1]], split=".",fixed=T)[[1]][3], ".chasedby",sep="")
          current[[strsplit(census[nrow(census),evsplit[[1]][1]], split=".",fixed=T)[[1]][3]]]<-paste(strsplit(census[nrow(census),evsplit[[1]][1]], split=".",fixed=T)[[1]][3], ".", "fight", ".",evsplit[[1]][1], ".chasing",sep="")
          
        }
        
        #for disengaging, end the fight for the partner male as well
        if(evsplit[[1]][2]=="disengage"){
          current[evsplit[[1]][1]]<-"nothing"
          current[evsplit[[1]][3]]<-"nothing"	
          
        }
        
        
        #for copulations, end cops are only possible after start cops, but otherwise anything else is possible. EXCEPT the copulating male can't initiate an attack. If the next event involving that male ISN'T an end cop, it was interrupted
        
        if(evsplit[[1]][2]=="cop" && evsplit[[1]][3]=="start"){
          current[evsplit[[1]][1]]<-"copulating"
        }
        
        if(evsplit[[1]][2]=="cop" && evsplit[[1]][3]=="end"){
          current[evsplit[[1]][1]]<-"nothing"
        }
        
        #if an X male is still involved in a fight at all, make sure he is still tagged as fighting (since sometimes more than one is interacting at once, but they can't depart until none are fighting)
        if((TRUE %in% grepl("fight.30",current) | TRUE %in% grepl("30.fight",current)) && !(grepl("receive", current[idno]) | grepl("attack", current[idno]))) {
          
          current[idno]<-"fight"
          
        }
        
      }
      
      census<-rbind(census,current)
      
      
    }
    
    
    #don't want the first dummy row
    census.list[[j]]<-census[-1,]
    
  
}


#convert to win.loss census- again need two diferent functions for the different datasets b/c of constraints on X males 

create.win.loss.census.act<-function(x){
  
  #get disengagement events for focal male from evlist
  win.loss.census<-x
  
  #loop thru days and males
  
  for(i in 1:length(win.loss.census)){
    
    for(l in 1:idno){
      
      
      dis.times<-which(startsWith(as.character(event.list.dat[[i]][,1]), paste(l,".disengage", sep="")))
      
      
      if(length(dis.times)>0){
        
        evsplit<-strsplit(as.character(event.list.dat[[i]][dis.times,1]),".",fixed=T)
        
        #so we have a list of row numbers for disengagements and a list of interaction partners. This will cover wins for normal fights and losses for chases for the focal male. If we loop through all the males, all the wins and losses should get filled in.
        
        #NOW DISENGAGES ALWAYS MEAN WINS, so we will still have chase information, but we don't need to do anything special with it here.
        
        #need to make sure we are really precise since male 30 can be involved in multiple interactions, but they're usually all chases.
        
        
        for(j in 1:length(dis.times)){
          
          
          if(grepl("chasing",win.loss.census[[i]][(dis.times[j]-1),l]) ==FALSE && grepl("chasedby",win.loss.census[[i]][(dis.times[j]-1),as.numeric(evsplit[[j]][3])]) ==FALSE){
            #apply the win in the focal male's column
            win.loss.census[[i]][dis.times[j],l]<-paste(l, "win",evsplit[[j]][3], sep=".")
            #apply the loss for the same interaction in the partner male's column
            win.loss.census[[i]][dis.times[j],as.numeric(evsplit[[j]][3])]<-paste(evsplit[[j]][3], "lose", l, sep=".")
          } else{
            #apply the won chase in the focal male's column
            win.loss.census[[i]][dis.times[j],l]<-paste(l, "win",evsplit[[j]][3],"chase", sep=".")
            #apply the loss for the same interaction in the partner male's column.
            win.loss.census[[i]][dis.times[j],as.numeric(evsplit[[j]][3])]<-paste(evsplit[[j]][3], "lose", l, "chase",sep=".")
          }
          
        }
        
      }
      
    }
    
  }
  
  return(win.loss.census)
  
}

win.loss.census<-create.win.loss.census.act(census.list)

#win.loss.census[[1]]

## should be able to adapt the availability code to get the amount of time fighting in front of females as well. 


#add time, Females, and Events columns
census.time<-list()

for(i in 1:length(win.loss.census)){
  
  census.time[[i]]<-cbind(event.list.dat[[i]][,c(1,2,3)],win.loss.census[[i]])
  colnames(census.time[[i]])[c(1,2,3)]<-c("Events","Time","Females")
  
}


library(dplyr)


#remove columns for males that were absent that day, and other columns we don't need right now

census.time.pres<-sapply(c(1:length(census.time)),function(x){
  
  return(census.time[[x]][,c("Events","Time","Females",as.character(which(male.pres[x,]==1)-1))])
  
})

library(abind)

census.interval<-lapply(census.time.pres,function(z){
  
  availlist<-list()
  
  for(i in 1:30){
    
    #make sure that male was there
    if(as.character(i) %in% dimnames(z)[[2]]){
      
      newavail<-as.character(z[,as.character(i)])
      
      intervail<-newavail
      
      #if the male is attack, received, or fight, they are fighting. they become available again at the moment they win or lose. If females are not there, even if they are fighting, we don't count it here because they are fighting in front of females. 
      
      #maybe just label each timepoint as available or not and then pare down to the transitions using lag or lead
      
      intervail[grepl("fight|win|lose",lead(newavail)) & z$Females==2]<-"fight"
      intervail[!grepl("fight|win|lose",lead(newavail))  & z$Females==2 | z$Females==1]<-"not"
     
      #check
      #tail(data.frame(orig=newavail,new=intervail,time=z$Time,fem=z$Females),100)
      
     
      #then identify the actual beginnings and ends of the intervals
      intervail.ends<-intervail
      
      intervail.ends[intervail.ends==lag(intervail.ends)]<-"drop"
      
      #since lag doesn't touch the first row
      if(intervail.ends[1]=="not"){intervail.ends[1]<-"drop"}
      
     # data.frame(orig=newavail,new=intervail,drops=intervail.ends,time=z$Time,fem=z$Females)
      
      lab<-rep(i,length(newavail))
      time<-z[,2]
      
      tempdat<-data.frame(fight=intervail,name=lab,time=time)
      
      #remove rows between interval termini
      tempdat<-tempdat[intervail.ends=="fight" | intervail.ends=="not",]	
      
      #turn into avail and not columns
      
      
      availlist[[i]]<-tempdat
      
    }
    
  }
  
  availday<-abind(availlist,along=1)
  
  availday<-data.frame(availday,stringsAsFactors=F)
  availday$time<-as.numeric(availday$time)
  
  return(availday)
})


fightdat<-lapply(census.interval, function(x){
  
  mutdat<-x[x$fight=="fight",][,c(2,3)]
  
  #match with interval ends
  mutdat$not<-x[x$fight=="not",][,3]
  
  colnames(mutdat)<-c("male","fight","not")
  mutdat$male<-factor(mutdat$male,levels=c(1:30))
  
  #restrict to territory-holders?
  # mutdat<-mutdat[(mutdat$male==1 | mutdat$male==2 | mutdat$male==3 | mutdat$male==4 | mutdat$male==7),]
  
  return(mutdat)
  
})

######and calculate the total numbers to display in a table/chart on the side

fightdat.plot<-lapply(fightdat, function(x){
  x$tot<-x$not-x$fight
  return(x)})

#get total amount of time actually available for mating

fighting<-lapply(fightdat.plot, function(y){
  z<-y %>%
    group_by(male) %>%
    summarise(timefight=sum(tot))
  return(z)
})


##fuse together the datasets

totals<-list()
  
for(i in 1:length(availability)){
  
  totals[[i]]<-full_join(availability[[i]],fighting[[i]],by="male")
  
  }


###filter by territory-holding males on each day

#are you here today AND ALSO both of the past two days? (excluding X males)

#add attendance data from 3/17
extra<-rep(0,29)
names(extra)<-1:29
extra[c(1,2,3,4,5,7)]<-1

terr.list<-list()

for(k in 1:length(event.list.dat)){
  
  if(k==1 | k==2){
    
    if(k==1){
      
      terr.pres<-rbind(extra,extra)
      terr.pres<-rbind(terr.pres,male.pres[k,2:(ncol(male.pres)-1)])
      
      terr.holders<-which(apply(terr.pres,2,function(x) sum(x))==3)
      
    }
    
    if(k==2){
      
      
      terr.pres<-rbind(extra,male.pres[c(k-1,k),2:(ncol(male.pres)-1)])
      
      terr.holders<-which(apply(terr.pres,2,function(x) sum(x))==3)
      
    }
    
    
  } else {
    
    terr.holders<-which(apply(male.pres[(k-2):k,2:(ncol(male.pres)-1)],2,function(x) sum(x))==3)
    
  }
  
  terr.list[[k]]<-terr.holders
  
}


totals.terr<-list()

for(i in 1:length(totals)){
  
  totals.terr[[i]]<-totals[[i]][sapply(totals[[i]]$male, function(x){as.character(x)%in%terr.list[[i]]}),]
  
}

## change NA's to zeros for males that were present in fron tof females but didn't fight

totals.terr<-lapply(totals.terr,function(x){
  x$timefight[is.na(x$timefight)]<-0
  return(x)
})


##############################################################
###### Get fighting times for Day 13 example specifically ####

day13<-totals.terr[[13]]

day13<-mutate(day13,total=timeavail+timefight,perc.fight=timefight/(timeavail+timefight),availmin=timeavail/60,fightmin=timefight/60)

# male  timeavail timefight total perc.fight availmin fightmin
# <fct>     <dbl>     <dbl> <dbl>      <dbl>    <dbl>    <dbl>
# 1 1         4713.      86.7  4800     0.0181     78.6     1.45
# 2 2         4629.     171.   4800     0.0357     77.1     2.85
# 3 3         4322.     478.   4800     0.0997     72.0     7.97
# 4 4         4064.     736.   4800     0.153      67.7    12.3 
# 5 7         4613.     187.   4800     0.0390     76.9     3.12

##Male 3 spent 8 minutes fighting of his 80 mins in front of females (10%), compared to Male 4's 12.3 mins fighting (15.3%)

(day13$timeavail[day13$male=="3"]-day13$timeavail[day13$male=="4"])/day13$timeavail[day13$male=="3"]
# 0.05965497

(day13$timeavail[day13$male=="3"]-day13$timeavail[day13$male=="4"])/60
# 4.296683

##put another way, Male 4 was available to mate 6% less than male 3, a difference of 4.3 mins entirely attributable to fighting.

##############################################################

#### mush it into a big dataframe for the final steps

totaldat<-abind(totals.terr, along=1)

totaldatnew<-data.frame(totaldat,stringsAsFactors = F)

totaldatnew<-transmute(totaldatnew,male=as.factor(male),timeavail=as.numeric(timeavail),timefight=as.numeric(timefight))

totaldatnew<-mutate(totaldatnew,total=timeavail+timefight,perc.fight=timefight/(timeavail+timefight),availmin=timeavail/60,fightmin=timefight/60, totalmin=(timeavail+timefight)/60)

#get the number of instances for each male
table(totaldatnew$male)
#  1 10 12  2  3  4  5  7  8 
# 18  4  1 18 18 18  9 18  2 

#Have to drop male 12 since we only have one observation for him

totaldatpared<-totaldatnew[!totaldatnew$male=="12",]
#reset factor levels
totaldatpared$male<-as.factor(as.character(totaldatpared$male))

occurs<-table(totaldatpared$male)
#  1 10  2  3  4  5  7  8 
# 18  4 18 18 18  9 18  2 

max(totaldatpared$perc.fight)
#male 4 on day 10
#0.640997

totaldatpared[which(totaldatpared$perc.fight==max(totaldatpared$perc.fight)),]
# male timeavail timefight  total perc.fight availmin fightmin  totalmin
#   4  1045.632  1866.968 2912.6   0.640997  17.4272 31.11613    48.54333

min(totaldatpared$perc.fight)
#0

#maximum time spent fighting with females present was male 4 on day 10: 64% or 31.1 of his 48.4 mins in attendance in front of females


#get the mean and sd for percent time fighting for each male
fightsum<-totaldatpared %>%
  group_by(male)%>%
  summarise(mean.perc=mean(perc.fight),sd.perc=sd(perc.fight))

# male  mean.perc sd.perc
# <fct>     <dbl>   <dbl>
# 1 1        0.0345  0.0363
# 2 10       0.151   0.125 
# 3 2        0.0589  0.0517
# 4 3        0.0579  0.0482
# 5 4        0.191   0.154 
# 6 5        0.0479  0.0767
# 7 7        0.126   0.120 
# 8 8        0.194   0.252 

##calculate common mean and 
##propagate the SD appropriately for a mean of means

fightsum<-fightsum %>%
  mutate(occur=occurs) %>%
  mutate(sum.of.squares=occur*(sd.perc^2+mean.perc^2))

#common mean
sum(fightsum$occur*fightsum$mean.perc)/sum(fightsum$occur)
#[1] 0.09368897

#this is equivalent to just directly taking the mean overall
mean(totaldatpared$perc.fight)
#[1] 0.09368897

#common SD
sqrt((sum(fightsum$sum.of.squares)/sum(fightsum$occur))-mean(totaldatpared$perc.fight)^2)
#[1] 0.1151398

median(totaldatpared$perc.fight)
#[1] 0.05224468

##The overall mean percent fighting time in front of females for territory-holding males was 9.37% +- 0.12 SD, with a max of 64% and a minimum of 0 

#### Quick boxplot ####

library(ggplot2)

#reorder factor levels
totaldatpared$male.reord<-factor(totaldatpared$male, levels=levels(totaldatpared$male)[order(as.numeric(levels(totaldatpared$male)))])

percplot<-ggplot(data=totaldatpared)+geom_boxplot(aes(x=male.reord, y=100*perc.fight))+theme_classic()+theme(axis.title.x=element_text(size=20,face="bold"),axis.title.y=element_text(size=18,face="bold"),axis.text.x=element_text(size=22,face="bold"),axis.text.y=element_text(size=16,face="bold"))+labs(x="Male ID", y="% time in front of \nfemales spent fighting")

ggsave("time_fighting_fig.pdf",percplot,height=5,width=5,units=c("in"))


event.list$Events<-factor(event.list$Events, levels=event.types)

chased<-table(basic.event.list$Events)[grep("initCH",names(table(basic.event.list$Events)))]

# 1.initCH  2.initCH  3.initCH  4.initCH  5.initCH 
# 1         0         0         1         1 
# 6.initCH  7.initCH  8.initCH  9.initCH 10.initCH 
# 16         0        17        33         4 
# 11.initCH 12.initCH 13.initCH 14.initCH 15.initCH 
# 6         0         0         9         2 
# 16.initCH 17.initCH 18.initCH 19.initCH 20.initCH 
# 8         1         1         4        22 
# 21.initCH 22.initCH 23.initCH 24.initCH 25.initCH 
# 1         0         0        17         0 
# 26.initCH 27.initCH 28.initCH 29.initCH 30.initCH 
# 0         9         0         0       483 


#spot check the single-chase males 
chaseline<-which(startsWith(as.character(basic.event.list$Events),"21.initCH"))

basic.event.list[c((chaseline-10):(chaseline+30)),]

#18 DID get chased off, even though it was just once.
#so did 21

#omit males that were never chased off, and exclude X males

chased<-chased[chased>0 & (names(chased)!="30.initCH" &names(chased)!="1.initCH"&names(chased)!="4.initCH"&names(chased)!="5.initCH"&names(chased)!="17.initCH")]


# 6.initCH  8.initCH  9.initCH 10.initCH 
# 16        17        33         4 
# 11.initCH 14.initCH 15.initCH 16.initCH 
# 6         9         2         8 
# 18.initCH 19.initCH 20.initCH 21.initCH 
# 1         4        22         1 
# 24.initCH 27.initCH 
# 17         9 

#none of these males ever mated
names(chased)
# [1] "6.initCH"  "8.initCH"  "9.initCH" 
# [4] "10.initCH" "11.initCH" "14.initCH"
# [7] "15.initCH" "16.initCH" "18.initCH"
# [10] "19.initCH" "20.initCH" "21.initCH"
# [13] "24.initCH" "27.initCH"

length(chased)
#14

# 3 males that mated were never chased off
# 14 males were chased off AND never mated
# 29-(3+14)= 12 males that were never chased off AND never mated
```
