## Supplemental_reproducible_code for "Fighting isn’t sexy in lekking Greater Sage-grouse (*Centrocercus urophasianus*)": genremlist_forward_full_model_udata_terr.R

```
#load in the user-defined functions that add predictor stats
source('rem.functions.R', chdir = TRUE)

#make a list of the functions
fun.list<-ls()[grep("add.",ls())]

fun.num<-c(1:length(fun.list))

#bring in the index numbers of the functions remaining after winnowing down covariates and linear combinations

#fun.num<-read.table("remaining_function_list_eigen_BIC_nocol",sep=",")$V1


#since this is the first iteration, prev is Null but later it will be the vector of previous functions to always include
prev<-NULL

jobs<-sapply(fun.num, function(x) paste(c("module load R; Rscript rem_cluster_model_selection_stepwise_udata_terr.R", prev, x),collapse=" ", sep=" "))

write(as.matrix(jobs), file="joblist_1.txt")

#create a file to save the best results of each run, starting with a dummy result

write("10000000,none", file="previous_best")
```
