## Supplemental_reproducible_code for "Fighting isn’t sexy in lekking Greater Sage-grouse (*Centrocercus urophasianus*)": genremlist_next_forward_full_model_udata_terr.R

```
#load in the user-defined functions that add predictor stats
source('rem.functions.R', chdir = TRUE)

#make a list of the functions
fun.list<-ls()[grep("add.",ls())]

#get just the index numbers
fun.num<-c(1:length(fun.list))


#bring in the index numbers of the functions remaining after winnowing down covariates and linear combinations

#fun.num<-read.table("remaining_function_list_eigen_BIC_nocol",sep=",")$V1


#bring in the results from the current run
result.files<-list.files(path="./res_dump",pattern="result")

result.list<-lapply(result.files, function(x) read.table(paste0("./res_dump/",x), stringsAsFactors=F))

result.dat<-result.list[[1]]$V1

for(i in 2:length(result.list)){
	result.dat<-rbind(result.dat,result.list[[i]]$V1)
}

result.dat<-data.frame(result.dat,stringsAsFactors=F)
row.names(result.dat)<-NULL

#filter out "garbage" results- broken models where stats are linear combinations of one another

result.ok<-result.dat[result.dat$X4=="ok",]


#dump correlation matrices into a subdirectory
# if(!dir.exists("cor_dump")) dir.create("cor_dump")
# cor.files<-list.files(pattern="cors")
# dump<-sapply(cor.files, function(x) file.rename(x,paste0("./cor_dump/", x)))


#make sure there is at least one "ok" result this round

if(nrow(result.ok)>0){

AICC.list<-as.numeric(result.ok[,1])

BIC.list<-as.numeric(result.ok[,2])


##here I can choose whether I want to use AICC or BIC for my criterion
criterion.list<-BIC.list
#criterion.list<-AICC.list

current.best<-min(criterion.list)

#get the function list for the current best outcome

result.ok[grep(current.best,criterion.list),3]

current.best.func<-result.ok[grep(current.best,criterion.list),3]

#bring in the previous best result
previous.best<-read.table("previous_best", sep=",")


#if there are no more predictors to test, end the run

if(nrow(previous.best)==length(fun.num)){
	
	write("no more predictors to test",file="all_done.txt")
	
} else {

#if the current best improves the model, then append it to the list of models and create a new joblist. If not, end the run.
if(current.best < previous.best[nrow(previous.best),1]){
	
	write(paste(current.best,current.best.func, sep=","), file="previous_best", append=TRUE)
	
	#go back to simple forward stepwise protocol, see if I get the same answer
	
	prev<-unlist(strsplit(current.best.func,split="_"))
	
	#make robust to using subsets of the function list
	prev.index<-sapply(prev, function(x) grep("TRUE",as.numeric(x)==fun.num))
	
	fun.num.next<-fun.num[-c(as.numeric(prev.index))]
	
	
	#create a joblist that allows for the exclusion of previously accepted functions. Then bring in previous joblist and compare argument lists so I don't repeat anything

#forward selection joblist
	jobs<-sapply(fun.num.next, function(x) paste(c("module load R; Rscript rem_cluster_model_selection_stepwise_udata_terr.R", prev, x),collapse=" ", sep=" "))
	

#write new joblist for cluster
write(as.matrix(jobs), file="joblist_next.txt")
	
} else {write("no more model improvement",file="all_done.txt")}

}

#if no good models left, end the run
} else {write("no more ok models",file="all_done.txt")}
```
