## Supplemental_reproducible_code for "Fighting isn’t sexy in lekking Greater Sage-grouse (*Centrocercus urophasianus*)": genremlist_restart_full_model_udata_terr.R

```
#load in the user-defined functions that add predictor stats
source('rem.functions.R', chdir = TRUE)

#make a list of the functions
fun.list<-ls()[grep("add.",ls())]

fun.num<-c(1:length(fun.list))

#bring in the index numbers of the functions remaining after winnowing down covariates and linear combinations

#fun.num<-read.table("remaining_function_list_eigen_BIC_nocol",sep=",")$V1

#bring in the previous best result- here I put in the special version for resarting from an altered list
previous.best<-read.table("previous_best_no226_no283", sep=",")


current.best.func<-as.character(previous.best[nrow(previous.best),2])


	prev<-unlist(strsplit(current.best.func,split="_"))
	
	#make robust to using subsets of the function list
	prev.index<-sapply(prev, function(x) grep("TRUE",as.numeric(x)==fun.num))
	
	#removing function 226 and 283 (.sol function) from new joblists- see what happens
	fun.num.next<-fun.num[-c(as.numeric(prev.index),226,283)]
	
	
	#create a joblist that allows for the exclusion of previously accepted functions. Then bring in previous joblist and compare argument lists so I don't repeat anything

#forward selection joblist
jobs<-sapply(fun.num.next, function(x) paste(c("module load R; Rscript rem_cluster_model_selection_stepwise_udata_terr.R", prev, x),collapse=" ", sep=" "))

#if starting from an altered prev_best list (like if I removed a predictor and I want to see if the model adds more) then I tack on the list with the alterations to the joblist for comparison

jobs<-c(paste(c("module load R; Rscript rem_cluster_model_selection_stepwise_udata_terr_id.R", prev),collapse=" ", sep=" "),jobs)	
	
write(as.matrix(jobs), file="joblist_1.txt")
```
