## Supplemental_reproducible_code for "Fighting isn’t sexy in lekking Greater Sage-grouse (*Centrocercus urophasianus*)": rem_cluster_model_selection_stepwise_udata_terr.R

```
### Script for running REM in Parellel ###

#set up all the stuff I need to run the model, load in the files and the source code for the functions

#bring in the parameters I want to use, which will be a line from a list of all the combinations of predictors

#save the output as a line in a document

#GUI cluster acces: ood-farnam.hpc.yale.edu


##########################################################################

###make sure necessary packages are there and loaded

packages<-c("abind","relevent","lubridate","DescTools","gtools")

package.check<-lapply(packages, FUN = function(x) {
    if (!require(x, character.only = TRUE, quietly=TRUE)) {
 install.packages(x, dependencies = TRUE)
 library(x, character.only = TRUE, quietly=TRUE)
 } 

})


###Build the intercept list, support array, event list, census array, etc

{
	
#make the base event list and intercept array


event.list<-read.table(file="relevent_format_dataset_udata_wincol_disturbance.tsv", header=T,  sep="\t")

event.list$Date<-as.factor(event.list$Date)  
  
event.types<-read.table(file="event_types_list_udata.csv", header=F,  sep=",")
event.types<-as.character(event.types$V1)


#NOTE- the event.types vector now contains ENDOGENOUS event types as well as EXOGENOUS types: arrivals and clock events, and end.obs. Need to make sure everything is robust to this. 


### create eventlist for rem ###


event.list.dat<-split(event.list, event.list$Date)


#remove female context for now

basic.event.list<-event.list[-3]

#strip the "loser" from the disengage events so they are all the same event "type" and they match my previous code

basic.event.list$Events<-sapply(basic.event.list$Events, function(x) {
	
	event<-unlist(strsplit(as.character(x),".",fixed=T))
	if(event[2]=="disengage"){
		return(paste(event[1],event[2],sep="."))
	} else {return(as.character(x))}
}
)


#add factor levels for event types that don't actually happen (and put them in the same order as the event.type list) to maintain bookkeeping 

basic.event.list$Events<-as.character(basic.event.list$Events)

basic.event.list$Events<-factor(basic.event.list$Events, levels=as.character(event.types))

#turn event types into numeric values

basic.event.list$Events<-as.numeric(basic.event.list$Events)

#set the event type code for exogenous events to zero. I have there all at the trailing end of the vector

endogenous<-(length(event.types)-length(event.types[grep("arrive",event.types)])-5)

basic.event.list$Events[basic.event.list$Events>endogenous]<-0

rownames(basic.event.list)<-NULL
#need to divide the data up by grouping factor

evlist<-list()
evlist<-split(basic.event.list, basic.event.list$Date)

#strip the date column from the sub-dataframes so they're just two columns, event and time

evlist<-lapply(evlist, function(x) x<-x[-3])

#convert from dataframes to matrices

evlist<-lapply(evlist, function(x) x<-as.matrix(x))

## make event.types vector usable for intlist, etc. by excising exogenous event types

event.types<-event.types[1:endogenous]


#### create statlist ####

##"a three dimensional array, event number by event type by statistic, or a list thereof, containing the sufficient statistics for the model"

#start with the intercept sufficient stats

#seems like the statlist involves an array with an ojbect for every "group" according to grouping factor. In the example it's the individuals but for me it's the different DAYS. each day will have a matrix for each event in the sequence, and for the intercepts, each matrix will just be an identity matrix of event types... I think. give it a try. This, I think, should EXCLUDE rows for all the exogenous event types as well (except end.obs which is still in there). I know this isn't what Carter would prefer, but it's just adding dead events that we basically tell the model to ignore anyway. 


sequences<-table(event.list$Date)

#create list of dummy intercept names

idno<-length(event.types[grep("cop.start",event.types)])

dummy.ints.attack<-paste(1:14,".attackFE", sep="")
dummy.ints.receive<-paste(1:13, ".receiveFE", sep="")

IDintsCH<-paste(1:14,"initCH",sep=".")
IDdis<-paste(1:14,"disengage",sep=".")

int.names<-c("Global.attackFE", "X.receiveFE", "X.UreceiveFE", "U.attackFE", "U.receiveFE", "U.UattackFE", dummy.ints.attack, dummy.ints.receive, "cop.startFE","cop.endFE", "initFO.FE","ID.initFO.FE", "U.initFO.FE","initSF.FE","X.initSF.FE", "initCH.FE",  "U.initCH.FE", IDintsCH, "disengage.FE", "U.disengage.FE", IDdis, "depart.FE", "X.depart.FE")

int.list<-list()

for(i in 1:length(levels(event.list$Date))){

day.array<-array(0,dim=c(as.numeric(sequences[i]), length(event.types),length(int.names)))

#the third dimension is the stat slice.
dimnames(day.array)<-list(c(1:sequences[i]),as.character(event.types),as.character(int.names))

#add global attack effect, loops for "U" intercepts for attacks and receives, then one loop for typical intercepts

day.array[,grep("attack", event.types),1]<-1


for(h in 1:14){
day.array[,paste(h,".attack.",30, sep=""),2]<-1
}


for(h in 15:29){
day.array[,paste(h,".attack.",30, sep=""),3]<-1
}


for(h in 1:14){
	for(j in 15:29){	
day.array[,paste(j,".attack.", h, sep=""),4]<-1
}
}


for(h in 1:14){
	for(j in 15:29){
day.array[,paste(h,".attack.",j, sep=""),5]<-1
}
}

for(h in 15:29){
	for(j in 15:29){	
if(h!=j){
day.array[,paste(j,".attack.", h, sep=""),6]<-1
}
}
}

preloop<-6

#loop for dummy intercepts for attacks, loop for dummy intercepts for receives, add in cop.start and cop.end FEs, then one loop for typical intercepts

for(k in (preloop+1):(length(dummy.ints.attack)+preloop)){
	
	for(h in 1:14){
if(h!=(k-preloop)){
day.array[,paste((k-preloop),".attack.", h, sep=""),k]<-1
}
}
}

for(k in ((length(dummy.ints.attack)+preloop)+1):((length(dummy.ints.attack)+preloop)+length(dummy.ints.receive))){
	for(h in 1:14){
if(h!=(k-(length(dummy.ints.attack)+preloop))){
day.array[,paste(h,".attack.",(k-(length(dummy.ints.attack)+preloop)), sep=""),k]<-1
}
}
}

#cop.startFE
for(h in 1:idno){
day.array[,paste(h,".cop.start", sep=""),((length(dummy.ints.attack)+preloop)+length(dummy.ints.receive)+1)]<-1
}


#cop.endFE
for(h in 1:idno){
day.array[,paste(h,".cop.end", sep=""),((length(dummy.ints.attack)+preloop)+length(dummy.ints.receive)+2)]<-1
}

#initFO.FE
for(h in 1:idno){
day.array[,paste(h,".initFO", sep=""),((length(dummy.ints.attack)+preloop)+length(dummy.ints.receive)+3)]<-1
}

#ID.initFO.FE
for(h in 1:14){
day.array[,paste(h,".initFO", sep=""),((length(dummy.ints.attack)+preloop)+length(dummy.ints.receive)+4)]<-1
}

#U.initFO.FE
for(h in 15:29){
day.array[,paste(h,".initFO", sep=""),((length(dummy.ints.attack)+preloop)+length(dummy.ints.receive)+5)]<-1
}

#initSF.FE
for(h in 1:idno){
day.array[,paste(h,".initSF", sep=""),((length(dummy.ints.attack)+preloop)+length(dummy.ints.receive)+6)]<-1
}

#X.initSF.FE
day.array[,paste(30,".initSF", sep=""),((length(dummy.ints.attack)+preloop)+length(dummy.ints.receive)+7)]<-1


#initCH.FE
for(h in 1:idno){
day.array[,paste(h,".initCH", sep=""),((length(dummy.ints.attack)+preloop)+length(dummy.ints.receive)+8)]<-1
}


#U.initCH.FE
for(h in 15:29){
day.array[,paste(h,".initCH", sep=""),((length(dummy.ints.attack)+preloop)+length(dummy.ints.receive)+9)]<-1
}


#individual initCH effects for ID male
for(k in (((length(dummy.ints.attack)+preloop)+length(dummy.ints.receive)+10):((length(dummy.ints.attack)+preloop)+length(dummy.ints.receive)+10+(length(IDintsCH)-1)))){
	
	day.array[,int.names[k],k]<-1
	
}

preloop2<-((length(dummy.ints.attack)+preloop)+length(dummy.ints.receive)+10+(length(IDintsCH)-1))


#disengage.FE
for(h in 1:idno){
day.array[,paste(h,".disengage", sep=""),preloop2+1]<-1
}

#U.disengage.FE
for(h in 15:29){
day.array[,paste(h,".disengage", sep=""),preloop2+2]<-1
}

#individual disengage effect for ID males
for(k in (preloop2+3):(preloop2+3+length(IDdis)-1)){
	
	day.array[,int.names[k],k]<-1
	
}


preloop3<-(preloop2+3+length(IDdis)-1)

#depart.FE
for(h in 1:idno){
day.array[,paste(h,".depart", sep=""),preloop3+1]<-1
}

#X.depart.FE
day.array[,paste(30,".depart", sep=""),preloop3+2]<-1


#weird extra formatting to make it match the package example, seems like it becomes relevant *punintended!* if I want to fit day-specific effects

global<-list()

global[[1]]<-day.array

int.list[[i]]<-global

names(int.list[[i]])<-"global"

}


#### Create Support Matrix ####

#"should be a list withone element per history, each of which should be an event order by event type logical matrix. the ijth cell of this matrix should be TRUE if an event type of j was a possible next event given the preceding code i-1 events, and FALSE otherwise"

#need a way to ask, "what is each male doing right now" for each event in the sequence. That dictates the possibilities. 

#create a bookkeeping "census" matrix to keep track

census.list<-list()


for(j in 1:length(event.list.dat)){

census<-matrix("nothing", nrow=1,ncol=(idno), dimnames=list(1,c(1:(idno))))


##code that translates each new current event into what the males involved are doing at that time. The support matrix will be based on whatever the males were doing in the PREVIOUS timestep. This census matrix itself could be interesting to visualize. Each row of the census says what each male is doing going into the NEXT event.


for(i in 1:nrow(event.list.dat[[j]])){

#can ask if the second thing is something, and then move on to the first and third elements.

current<-census[nrow(census),]

#parse the event type. loop through test.seq rows
evname<-as.character(event.list.dat[[j]][i,1])
evsplit<-strsplit(evname, split=".", fixed=T)

#if it is an end obs, arrival, departure or clock event, it shouldn't change anything the males are doing in terms of interactions. Departures aren't exogenous, but they shouldn't change anything since they can only happen once a male is already doing nothing. 
if(evsplit[[1]][2]!="obs" & evsplit[[1]][2]!="depart" & evsplit[[1]][2]!="arrive" & evsplit[[1]][2]!="time" & evsplit[[1]][1]!="fem"){
#if the event was an attack, indicate the males involved and which was the receiver since this dictates who makes the initition decision
	if(evsplit[[1]][2]=="attack"){
		current[evsplit[[1]][1]]<-evname
		current[evsplit[[1]][3]]<-paste(evsplit[[1]][3], ".", "received", ".", evsplit[[1]][1], sep="")
			}
	#if it's an FO or SF, just indicate that that pair of males are involved in a fight now
		if(evsplit[[1]][2]=="initFO"||evsplit[[1]][2]=="initSF"){
			#need to refer back to the previous row of "census" to get the id of the other male
			
			current[evsplit[[1]][1]]<-paste(evsplit[[1]][1], ".", "fight", ".",strsplit(census[nrow(census),evsplit[[1]][1]], split=".",fixed=T)[[1]][3], sep="")
			current[[strsplit(census[nrow(census),evsplit[[1]][1]], split=".",fixed=T)[[1]][3]]]<-paste(strsplit(census[nrow(census),evsplit[[1]][1]], split=".",fixed=T)[[1]][3], ".", "fight", ".",evsplit[[1]][1], sep="")
			
			}
			
	#if it's a chase, indicate that the pair of males are involved in a fight, but there is an additional constraint that the individual DOING THE CHASING will be the disengager by definition
	if(evsplit[[1]][2]=="initCH"){
			
		current[evsplit[[1]][1]]<-paste(evsplit[[1]][1], ".", "fight", ".",strsplit(census[nrow(census),evsplit[[1]][1]], split=".",fixed=T)[[1]][3], ".chasedby",sep="")
			current[[strsplit(census[nrow(census),evsplit[[1]][1]], split=".",fixed=T)[[1]][3]]]<-paste(strsplit(census[nrow(census),evsplit[[1]][1]], split=".",fixed=T)[[1]][3], ".", "fight", ".",evsplit[[1]][1], ".chasing",sep="")

	}

#for disengaging, end the fight for the partner male as well
	if(evsplit[[1]][2]=="disengage"){
	current[evsplit[[1]][1]]<-"nothing"
	current[evsplit[[1]][3]]<-"nothing"	
	
}
 

#for copulations, end cops are only possible after start cops, but otherwise anything else is possible. EXCEPT the copulating male can't initiate an attack. If the next event involving that male ISN'T an end cop, it was interrupted

	if(evsplit[[1]][2]=="cop" && evsplit[[1]][3]=="start"){
		current[evsplit[[1]][1]]<-"copulating"
	}

	if(evsplit[[1]][2]=="cop" && evsplit[[1]][3]=="end"){
		current[evsplit[[1]][1]]<-"nothing"
	}

#if an X male is still involved in a fight at all, make sure he is still tagged as fighting (since sometimes more than one is interacting at once, but they can't depart until none are fighting)
	if((TRUE %in% grepl("fight.30",current) | TRUE %in% grepl("30.fight",current)) && !(grepl("receive", current[idno]) | grepl("attack", current[idno]))) {
		
		current[idno]<-"fight"

}

}

census<-rbind(census,current)


}


#don't want the first dummy row
census.list[[j]]<-census[-1,]

}


## Constraints to incorporate in the support matrix ##

#only males that are actually present can be involved in interactions on a given day and at a given time

#If "nothing" is happening, all "attacks", all start cops and all departs (given the male is actually present) are possible. Males cannot disengage or initiate fights without an attack happening first. End cop cannot happen without a previous start cop. end.obs is possible. (end.obs is not possible if any males are still fighting or copulating, so end.obs is ONLY possible when "nothing" is going on across the board)

#if the previous event was an "attack", the current event MUST be an initiation by the receiver, since we are assuming the attack is essentually instantaneous. 

#once initiatiated, subsequent events can only be a disengage from either male involved or any other event NOT invoving the interaction partners. Males actively fighting connot become engaged in a new fight until they disengage. 

#if the initiated fight was a chase only the male being chased can disengage, by definition

#Once a copulation starts, the only thing that male can do next is either end the copulation OR receive an attack. He cannot initiate anything. 

#copulations cannot occur when there are no females present

#a male can ONLY "depart" if "nothing" is happening

#Exceptions to these rules are for "X" males- there may be more than one unidentifiable male on the lek at a time. Right now we are dealing with this by having "X" males just be a mass actor category. 1) These males CAN do an "attack" before disengaging from an interaction. They can also be attacked. However, normal restrictions apply to the partner male. So if an X male is being chased, "disengage" becomes possible, but any "attack" (or cop.start, or depart) are still also possible 


### Build the actual support matrix ###

#for each event, there is a list of event types, with 1's for the types that are possible and 0's for types that are not. 


#code block to produce the support matrix

supplist<-list()


#bring in day-level male presence data

male.pres<-read.table("CHG_2014_male_presence_data_quest.csv", sep=",", header=T)

#male IDs are already in ascending order, so I can convert them directly
colnames(male.pres)<-c("Date", 1:idno)

#bring in exact male presence data

pres.exact<-read.table("male_pres_exact_quest.csv", sep=",",header=T)

	departlist<-list()
	
	for(j in 1:nrow(pres.exact)){
		
		arriverow<-pres.exact[j,startsWith(colnames(pres.exact),"arrive")]
		arriverow<-arriverow[!is.na(arriverow)]
		
		arrives<-t(sapply(arriverow, function(x) return(c(paste0(pres.exact$Num_ID[j],".arrive"),x,as.character(pres.exact$Date[j])))))
		
		departrow<-pres.exact[j,startsWith(colnames(pres.exact),"depart")]
		departrow<-departrow[!is.na(departrow)]
		
		departs<-t(sapply(departrow, function(x) return(c(paste0(pres.exact$Num_ID[j],".depart"),x,as.character(pres.exact$Date[j])))))
		
		allpres<-rbind(arrives,departs)
		
		departlist[[j]]<-allpres
		
	}

#collapse the departlist 

departevent<-abind(departlist, along=1)

departevent<-data.frame(departevent)
names(departevent)<-c("Events","Time", "Date")

departevent$Time<-as.numeric(as.character(departevent$Time))

library(abind)
arrstr<-abind(strsplit(as.character(departevent$Events),".",fixed=T),along=2)

departevent$Bird_ID<-arrstr[1,]
departevent$Type<-arrstr[2,]

departevent$Type<-as.factor(departevent$Type)
departevent$Bird_ID<-as.factor(departevent$Bird_ID)

# #split into day-specific data frames
pres.exact.dat<-split(departevent, departevent$Date)


#make a reference event list with data (most importantly female presence)
evlist.dat<-event.list.dat


for(j in 1:length(evlist)){

supplist[[j]]<-matrix(1, nrow=length(evlist[[j]][,1]), ncol=length(event.types))
colnames(supplist[[j]])<-event.types

#turn off all interactions involving males not present on that day
#make a list of all the event types that include the males that were entirely absent and remove them for all rows

supplist[[j]][,unique(unlist(lapply(grep(0,male.pres[j,2:(idno+1)]), function(x) c(event.types[startsWith(event.types, paste(x, ".", sep=""))], event.types[endsWith(event.types, paste(".", x, sep=""))]))))]<-0


#first row can be any "attack" or, technically, any "depart"
#since now fem.enter is an actual event, cop.start is never possible as the very first thing
supplist[[j]][1,][-c(grep("attack", event.types),grep("depart", event.types))]<-0


#the exception is events involving males that will be present that day but haven't arrived yet


#basically I can make this code exactly the same as below- just want to ask who is actually there right now? 

arrsub<-subset(pres.exact.dat[[j]],Time<evlist[[j]][1,2] | Time==0 )
arrtab<-table(arrsub$Bird_ID,arrsub$Type)

arrpat<-apply(arrtab,1,function(x) sum(x))

#if the total number of previous arrives and departs is even, that male is currently absent
absent<-names(arrpat[(arrpat %% 2)==0])

if(length(absent)>0){
supplist[[j]][1,][unique(c(sapply(absent, function(x) c(event.types[startsWith(event.types, paste(x, ".", sep=""))], event.types[endsWith(event.types, paste(".", x, sep=""))]))))]<-0

} 


#loop through the rest of the events: 

for(i in 2:(length(evlist[[j]][,1])-1)){


#before anything else turn off events involving males that have either not yet arrived or have lef the lek early and are therefore no longer present that day or are otherwise not currently present


arrsub<-subset(pres.exact.dat[[j]],Time<evlist[[j]][i,2])
arrtab<-table(arrsub$Bird_ID,arrsub$Type)

arrpat<-apply(arrtab,1,function(x) sum(x))

#if the total number of previous arrives and departs is even, that male is currently absent
current.row.abs<-names(arrpat[(arrpat %% 2)==0])


if(length(current.row.abs)>0){
supplist[[j]][i,][unique(c(sapply(current.row.abs, function(x) c(event.types[startsWith(event.types, paste(x, ".", sep=""))], event.types[endsWith(event.types, paste(".", x, sep=""))]))))]<-0

} 


#determine if the previous event was an attack, since this dictates exactly what the next event can be: receiver initiates a type of fight
if((TRUE %in% grepl("received", census.list[[j]][i-1,]))==TRUE){

supplist[[j]][i,][-grep(TRUE, startsWith(event.types, paste(grep("received", census.list[[j]][i-1,]), ".", "init", sep="")))]<-0}


#(else) if the previous event wasn't an attack, the next event CAN'T be an initiation and the possibilities depend on what everyone is currently engaged in


if((TRUE %in% grepl("received", census.list[[j]][i-1,]))==FALSE){

supplist[[j]][i,][grep("init", event.types)]<-0

#if males are actively fighting, they can't attack or be attacked, or actually do anything else before disengaging (unless they are a ?? male, id 30)

if((TRUE %in% grepl("fight", census.list[[j]][i-1,]))==TRUE){
		
		for(k in grep("fight", census.list[[j]][i-1,])){
	
	if(k!=idno){
	
	supplist[[j]][i,][c(event.types[startsWith(event.types, paste(k, ".attack.", sep=""))], event.types[endsWith(event.types, paste("attack.", k, sep=""))], event.types[startsWith(event.types, paste(k, ".cop", sep=""))])]<-0
	
	}
	
}
	
	#ONLY those fighting can disengage. retain males that are fighting and remove the rest
	dis<-event.types[grep("disengage",event.types)]
	supplist[[j]][i,][dis[-grep("fight", census.list[[j]][i-1,])]]<-0
	
	#"chasing" and "chasedby" limit the possibilities further. Males getting chased DO NOT disengage by definition, again, with the exception of ?? males. This is taken care of by having ?? males just be "fight" -ing without any extra tags
	if(TRUE %in% grepl("chasedby", census.list[[j]][i-1,])){
		supplist[[j]][i,][dis[grep("chasedby", census.list[[j]][i-1,])]]<-0
	}
	
}

#if no one is fighting, no one can disengage, but all attack combinations are possible
if((TRUE %in% grepl("fight", census.list[[j]][i-1,]))==FALSE){
	
		supplist[[j]][i,][grep("disengage",event.types)]<-0
	
}


#actively copulating males: They cannot cop.start. ONLY they can cop.end. they CAN be attacked but they CANNOT attack anyone
if((TRUE %in% grepl("copulating", census.list[[j]][i-1,]))==TRUE){
	
	cop<-grep("copulating", census.list[[j]][i-1,]) 
	
	#stuff that needs to be turned off for only copulating males
	for(h in cop){

		supplist[[j]][i,][c(event.types[startsWith(event.types ,paste(h, ".attack.", sep=""))], event.types[startsWith(event.types, paste(h, ".cop.start", sep=""))])]<-0
				
	}
	
	#stuff that needs to be on for ONLY copulating males
	copendall<-event.types[grep("cop.end",event.types)]
	supplist[[j]][i,][copendall[-cop]]<-0
}

#if no one is copulating, all cop.starts are possible, but cop.ends are not

if((TRUE %in% grepl("copulating", census.list[[j]][i-1,]))==FALSE){
	
	copendall<-event.types[grep("cop.end",event.types)]
	supplist[[j]][i,][copendall]<-0
	
}


}

#if there are no females present, the next event could not have possibly been a cop.start

if(evlist.dat[[j]][i-1,]$Females==1){
	
	copstartall<-event.types[grep("cop.start",event.types)]
	supplist[[j]][i,][copstartall]<-0
	
}

#Finally, A male can only "depart" if he was already doing "nothing," so depart must be turned off for everyone doing not-nothing. (also he has to be present, but I've already taken care of that above)

noth<-as.character(which(census.list[[j]][i-1,]!="nothing"))
	
	if(length(noth)>0){
		
	supplist[[j]][i,][paste(noth, ".depart", sep="")]<-0

}

}
#end events loop here

#after done with all the middle events: onset of end.obs could depend on a lot of factors; not sure I want to fit it as part of the model. Just always say the last row must be end.obs (all endogenous events are no longer possible) 

supplist[[j]][length(evlist[[j]][,1]),]<-0

}
#end day loop here


###Make the win/loss census array

#i've adjusted the function to mark when a win or loss is the result of a chase or not
create.win.loss.census<-function(x){

#get disengagement events for focal male from evlist
win.loss.census<-x

#loop thru days and males

for(i in 1:length(win.loss.census)){

for(l in 1:idno){


dis.times<-which(startsWith(as.character(event.list.dat[[i]][,1]), paste(l,".disengage", sep="")))


if(length(dis.times)>0){

evsplit<-strsplit(as.character(event.list.dat[[i]][dis.times,1]),".",fixed=T)

#so I have a list of row numbers for disengagements and a list of interaction partners. This will cover wins for normal fights and losses for chases for the focal male. If I loop through all the males, all the wins and losses should get filled in.

#NOW DISENGAGES ALWAYS MEAN WINS, so I will still have chase information, but I don't need to do anything special with it here.

#need to make sure I am really precise since male 30 can be involved in multiple interactions, but they're usually all chases.


for(j in 1:length(dis.times)){
	
		
	if(grepl("chasing",win.loss.census[[i]][(dis.times[j]-1),l]) ==FALSE && grepl("chasedby",win.loss.census[[i]][(dis.times[j]-1),as.numeric(evsplit[[j]][3])]) ==FALSE){
	#apply the win in the focal male's column
	win.loss.census[[i]][dis.times[j],l]<-paste(l, "win",evsplit[[j]][3], sep=".")
	#apply the loss for the same interaction in the partner male's column
	win.loss.census[[i]][dis.times[j],as.numeric(evsplit[[j]][3])]<-paste(evsplit[[j]][3], "lose", l, sep=".")
	} else{
		#apply the won chase in the focal male's column
	win.loss.census[[i]][dis.times[j],l]<-paste(l, "win",evsplit[[j]][3],"chase", sep=".")
		#apply the loss for the same interaction in the partner male's column.
	win.loss.census[[i]][dis.times[j],as.numeric(evsplit[[j]][3])]<-paste(evsplit[[j]][3], "lose", l, "chase",sep=".")
		}
		
}

}

}

}

return(win.loss.census)
	
}

win.loss.census<-create.win.loss.census(census.list)


#other important stuff to load
### load in pairwise distance data
##bring in pairwise distances; I have a table for each day
##female exact presence data


file_list<-list.files("./pairwise_dist")

distlist<-list()

distlist<-lapply(file_list, function(x) read.table(paste("./pairwise_dist/",x,sep=""), sep=","))


#convert to martices

distlist.mat<-lapply(distlist, function(x) as.matrix(x, ))

#the names of the rows and columns are in the same order as the numerical ids passed to rem, so they can be used to index the matrix. Males that are absent on a given day have NA's for their centroid distances. I can change them to zeroes here so they don't produce errors, but the support array will take care of them anyway.

for(i in 1:length(distlist.mat)){
distlist.mat[[i]][is.na(distlist.mat[[i]])]<-0
}


fe_exact<-read.table(file="CHG_females_exact.tsv", header=TRUE,sep="\t",stringsAsFactors=FALSE)

#reduce to only the necessary columns- this time I want time elapsed
fe_exact<-fe_exact[,c(1,3,5,7,9,11,13)]

#turn the first arrival NA's to 0s
fe_exact$first_arrival_elapsed[is.na(fe_exact$first_arrival_elapsed)]<-"00:00:00:00"

library(lubridate)
options(digits.secs=6)

#convert number of frames to partial seconds
for(h in 2:7){
fe_exact[,h]<-sapply(fe_exact[,h], function(x) {

fem.split<-unlist(strsplit(x,split=":", fixed=TRUE))
suppressWarnings(period_to_seconds(hms(paste(fem.split[1],fem.split[2],(as.numeric(fem.split[3])+as.numeric(fem.split[4])/30), sep=":"))))

}
)
}


}


##########################################################################
##########################################################################


#the following two lines grab the arguments from the command line and save them as a vector

#!/usr/bin/env Rscript
args=as.numeric(commandArgs(trailingOnly=TRUE))

#load in the user-defined functions that add predictor stats
source('rem.functions.R', chdir = TRUE)

fun.list<-lsf.str()[grep("add.",lsf.str())]


#add the new stats to the base intercept list

test.stat<-int.list

for(i in args){
test.stat<-get(fun.list[i])(test.stat)	
}


##run REM
test.mod<-try(rem(evlist, test.stat, supplist=supplist, timing="interval", estimator="BPM"),silent=T)


#if no problem, save output. Save impossible output if throws an error
if(inherits(test.mod,"try-error")==FALSE){

#this line gets saved to a text file
output<-c(test.mod$AICC,test.mod$BIC,paste(args, collapse="_"))
} else {output<-c(1000000,1000000,paste(args, collapse="_"))}


##screen the model for linear combos

##turn this off for now just to see

covmat<-summary(test.mod)$cov

if(length(args)>1){

covmat<-covmat[(nrow(covmat)-(length(args)-1)):nrow(covmat),(ncol(covmat)-(length(args)-1)):ncol(covmat)]

cormat<-cov2cor(covmat)

#save the correlation matrix too, just to see for now
#write.table(cormat,file=paste(c("cors", args), collapse="_"), sep=",") 

library(gtools)
comb<-combinations(length(args)+1,2)

testlist<-list()

for(k in 1:nrow(cormat)){

cors<-vector()

for(m in 1:nrow(comb)){

#add a logic gate that makes sure that the correlation cefficients are actually large enough to be meaningful, or else we will be removing variables for being too independent of one another
if(abs(cormat[k,][comb[m,1]])>0.1){	
cors<-append(cors,cormat[k,][comb[m,1]]+cormat[k,][comb[m,2]])
}

}

testlist[[k]]<-cors

}

#find if there are any instances of linear combinations, where the correlation coeffictients of two variables essentially cancel. This line will return TRUE if there are any hits and FALSE is everything seems fine

#based on spot-checking and testing, a threshold of .03 seems to work pretty well

if(TRUE %in% (lapply(testlist,function(x) TRUE %in% (x<0.03 & x>-0.03)))){
	
	#I changed this line here to say "ok" instead of "garbage," so the filtering is turned off
	output<-append(output,"ok")
} else {output<-append(output,"ok")}

} else {
	output<-append(output,"ok")}

#write out the file
if(!dir.exists("res_dump")) dir.create("res_dump")
write(output,file=paste0("./res_dump/",paste(c("result", args), collapse="_")))
```
