## Supplemental_reproducible_code for "Fighting isn’t sexy in lekking Greater Sage-grouse (*Centrocercus urophasianus*)": run_pipeline_stepwise.sh

```
#!/bin/bash
# -t 5:00
#SBATCH --mem=5G


# NOTES
# consider using sprintf("%03d", number) in R for formatting output. Will sort files numerically.

# \xC2\xA0 found in some R scripts. Is called non-breaking space, is wide UTF8 character which irritates R on cluster
for rfile in *.R;
 do
     sed -i 's/\xC2\xA0/ /g' $rfile
 done

module load R

if not [ -f joblist_1.txt ]
then
    Rscript genremlist_forward*
    jobfile='joblist_1.txt'
else
    Rscript genremlist_next_forward*
    jobfile='joblist_next.txt'
fi

if [ -f all_done.txt ]
then
    echo "simulations finished"
    exit 0
fi

module purge
module load dSQ/0.96
mkdir -p job_logfiles job_statusfiles

jid=$(dsq --jobfile $jobfile --mem=24G --time=2:00:00 -o job_logfiles/dsq-jobfile-%A_%a-%N.out --status-dir job_statusfiles --submit | awk '{print $NF}')
echo "submitted array $jid"

sbatch --dependency afterany:$jid --kill-on-invalid-dep=yes run_pipeline_stepwise.sh
```
