## Supplemental_reproducible_code for "Fighting isn’t sexy in lekking Greater Sage-grouse (*Centrocercus urophasianus*)": full_interaction_data_for_REM_import_code_udata_supp.R

```
#first, make sure R's memory is clean
rm(list=ls())

###make sure necessary packages are there and loaded

packages<-c("here","reshape2")

package.check<-lapply(packages, FUN = function(x) {
  if (!require(x, character.only = TRUE, quietly=TRUE)) {
    install.packages(x, dependencies = TRUE)
    library(x, character.only = TRUE, quietly=TRUE)
  } 
  
})


#for project path management

#define the project root directory relative to the current script. This sets the working directory that gets called bu the here() function. It is a relative path and will be robust to use on anyone's computer. In this case, I want the root directory to always be the enclosing folder for the whole sage grouse project
i_am("data_processing/full_interaction_data_for_REM_import_code_udata_supp.R")

file_list<-list.files(here("data_files","raw_interaction_data"))

#bring files in and create a dataframe called dataset
#first make a dataframe to append to

setwd(here("data_files","raw_interaction_data"))

dataset<-read.table(file_list[[1]],header=TRUE, sep="\t")

for(i in 2:length(file_list)){
	    temp_dataset <-read.table(file_list[[i]], header=TRUE, sep="\t")
    dataset<-rbind(dataset, temp_dataset)
    rm(temp_dataset)
}


#str(dataset)

full<-dataset

#save a copy of the full dataset without any modifications
#write.table(full, file="full_data_plus_cop.tsv", col.names=TRUE, row.names=F, sep="\t")


##replace NA's in the Face-off column with SF (short fight)

full$Face_Off[is.na(full$Face_Off)]<-"SF"


###Do operations on the full dataset including U's and ??'s 

quest<-full


#Make the Date and time into usable columns. I will make a formatted column just for time of day, and a factor for date

split<-colsplit(string=as.character(quest$Time_Stamp),pattern=" ",names=c("Date","Time"))

quest$Date<-split$Date
quest$Time<-split$Time

#turn time data into POSIXct format for the full dataset, use %OS to retain fractoinal seconds. The SRT only has out to three, and then python tacks three zeros on 
options(digits.secs=3)
quest$Time=as.POSIXct(quest$Time, format="%H:%M:%OS", tz=Sys.timezone())

#turn date into a factor (ordinal date info is stored in the datetime column)
quest$Date=as.factor(quest$Date)

#str(quest) #check

#for copulations that end in interruptions, the end stamp time is "inter" for now. 
#quest[quest$End_Caption_Stamp=="inter",]


######## Apply Females_exact data to make a new column representing whether there were any females on the grid at all at a particular time

fe_exact<-read.table(here("data_files","CHG_females_exact.tsv"), header=TRUE, sep="\t")

#str(fe_exact)

#reduce to only the necessary columns
fe_exact<-fe_exact[,c(1,2,4,6,8,10,12)]
#str(fe_exact)

#date I will leave as a factor, and then I will convert the other columns to time of day. This way I can apply my function by specific day.

for(i in c(2:7)){
	
	fe_exact[,i]<-as.POSIXct(fe_exact[,i], format="%H:%M:%S", tz=Sys.timezone())
	
	}

#Apply a function to the dataset indexing by the female exact times

for(i in c(1:length(quest$Date))){

quest$Females_Exact[i]<-ifelse(quest$Time[i]>=fe_exact[fe_exact$Date==quest$Date[i],]$third_depart,"N",
ifelse(quest$Time[i]<fe_exact[fe_exact$Date==quest$Date[i],]$third_depart && quest$Time[i]>=fe_exact[fe_exact$Date==quest$Date[i],]$third_arrival,"Y", 
ifelse(quest$Time[i]<fe_exact[fe_exact$Date==quest$Date[i],]$third_arrival && quest$Time[i]>=fe_exact[fe_exact$Date==quest$Date[i],]$second_depart,"N",
ifelse(quest$Time[i]<fe_exact[fe_exact$Date==quest$Date[i],]$second_depart && quest$Time[i]>=fe_exact[fe_exact$Date==quest$Date[i],]$second_arrival, "Y",
ifelse(quest$Time[i]<fe_exact[fe_exact$Date==quest$Date[i],]$second_arrival && quest$Time[i]>=fe_exact[fe_exact$Date==quest$Date[i],]$first_depart, "N",
ifelse(quest$Time[i]<fe_exact[fe_exact$Date==quest$Date[i],]$first_depart && quest$Time[i]>=fe_exact[fe_exact$Date==quest$Date[i],]$first_arrival, "Y",
ifelse(quest$Time[i]<fe_exact[fe_exact$Date==quest$Date[i],]$first_arrival, "N")))))))

}

#quest[,c(2,21)] #looks good!

#change to a factor
quest$Females_Exact<-as.factor(quest$Females_Exact)

#str(quest)


########## Add a column for distance between individuals at the beginning of an interaction

sqrt((c(quest$Initiator_Cartesian_X[1],quest$Initiator_Cartesian_Y[1])-c(quest$Partner_Cartesian_X[1],quest$Partner_Cartesian_Y[1]))[1]^2+(c(quest$Initiator_Cartesian_X[1],quest$Initiator_Cartesian_Y[1])-c(quest$Partner_Cartesian_X[1],quest$Partner_Cartesian_Y[1]))[2]^2)

disdif<-(cbind(quest$Initiator_Cartesian_X,quest$Initiator_Cartesian_Y)-cbind(quest$Partner_Cartesian_X,quest$Partner_Cartesian_Y))

quest$Dist=sqrt(disdif[,1]^2+disdif[,2]^2)

#the NA's in the data are from the copulations, where the partner coordinates are irrelevant. 


########### Re-code the U ids in quest so that they are usable in the REM ########

##IMPT##- we used the same U designations on multiple days, so we need to re-code the U male id's so that they are consistent. This will extend to the rest of the analyses as well. Can just append the date to the "U" to differentiate them while preserving the information

udata<-quest

udata$Initiator_ID<-as.character(udata$Initiator_ID)
udata$Partner_ID<-as.character(udata$Partner_ID)
udata$Disengager_ID<-as.character(udata$Disengager_ID)

for(i in 1:length(levels(udata$Date))){
	
	uday<-subset(udata,udata$Date==levels(udata$Date)[i])
	
	datenum<-strsplit(as.character(uday$Date[1]), split="-")[[1]][3]
	
	init.urows<-grep("U",uday$Initiator_ID)
	part.urows<-grep("U",uday$Partner_ID)
	dis.urows<-grep("U",uday$Disengager_ID)
	
	if(length(init.urows)>0){
	
	uday$Initiator_ID[init.urows]<-paste(uday$Initiator_ID[init.urows],datenum,sep=".")
}

	if(length(part.urows)>0){
	
	uday$Partner_ID[part.urows]<-paste(uday$Partner_ID[part.urows],datenum,sep=".")
}

	if(length(dis.urows)>0){
	
	uday$Disengager_ID[dis.urows]<-paste(uday$Disengager_ID[dis.urows],datenum,sep=".")
}

udata[udata$Date==levels(udata$Date)[i],]<-uday

}

#str(udata)

#union(unique(udata$Initiator_ID),unique(udata$Partner_ID))[order(union(unique(udata$Initiator_ID),unique(udata$Partner_ID)))]

 # [1] "??"    "315"   "340"   "351"   "702"   "705"   "709"   "710"   "711"   "712"  
# [11] "713"   "714"   "715"   "717"   "718"   "Fe"    "U.01"  "U.23"  "U1.22" "U1.23"
# [21] "U1.30" "U1.31" "U2.23" "U2.30" "U2.31" "U3.30" "U3.31" "U4.30" "U4.31" "U5.31"
# [31] "U6.31"


#so now we have 29 unique male IDs, and two mass-actors, Fe and ??

##### Re-code ?? males as well so as to preserve the order of things

udata$Initiator_ID[grep("??",udata$Initiator_ID, fixed=T)]<-"X"
udata$Partner_ID[grep("??",udata$Partner_ID, fixed=T)]<-"X"

#disengager is more complicated because there are NA's in there
udata$Disengager_ID[!is.na(udata$Disengager_ID)][udata$Disengager_ID[!is.na(udata$Disengager_ID)]=="??"]<-"X"

#str(udata)

########## SAVE DATASET ###########

setwd(here("data_processing"))

write.table(udata, file="full_data_plus_cop_quest.tsv", col.names=TRUE, row.names=F, sep="\t")
```
