## Supplemental_reproducible_code for "Fighting isn’t sexy in lekking Greater Sage-grouse (*Centrocercus urophasianus*)": relevent_formatting_dis_reverse_udata_wincolumn_disturbance.R

```
#first, make sure R's memory is clean
rm(list=ls())

###### Reformat data for use in the relevent package ######

###make sure necessary packages are there and loaded

packages<-c("here","lubridate","abind","dplyr")

package.check<-lapply(packages, FUN = function(x) {
  if (!require(x, character.only = TRUE, quietly=TRUE)) {
    install.packages(x, dependencies = TRUE)
    library(x, character.only = TRUE, quietly=TRUE)
  } 
  
})

#for project path management
#define the project root directory relative to the current script. This sets the working directory that gets called by the here() function. It is a relative path and will be robust to use on anyone's computer. In this case, I want the root directory to always be the enclosing folder for the whole sage grouse project
i_am("data_processing/relevent_formatting_dis_reverse_udata_wincolumn_disturbance.R")

data<-read.table(here("data_processing","full_data_plus_cop_quest.tsv"), header=TRUE, sep="\t")

#str(data)

###Identify and remove rows for which the disengager ID is ambiguous

#data[is.na(data$Disengager_ID),][-grep("C",data[is.na(data$Disengager_ID),"Reaction_Code"]),]
#row 1175 has an NA in the dataset, and 109 and 114 are problematic

data<-data[-c(109, 114, 1175),]

#1578 interactions


##### Male ID's as ordered numbers #####

ids<-union(unique(data$Initiator_ID), unique(data$Partner_ID))
#reorder

ids<-ids[order(ids)]
nums<-c(1:length(ids))

names(nums)<-ids
#Fe is number 15

#re-order to put Fe last. This is also how the ids are numbered in the pres.exact file as well

ids<-ids[c(1:14,16:31,15)]
names(nums)<-ids

#For reference
#nums
  # 315   340   351   702   705   709   710   711   712   713   714   715   717   718  U.01 
    # 1     2     3     4     5     6     7     8     9    10    11    12    13    14    15 
 
 # U.23 U1.22 U1.23 U1.30 U1.31 U2.23 U2.30 U2.31 U3.30 U3.31 U4.30 U4.31 U5.31 U6.31     X 
   # 16    17    18    19    20    21    22    23    24    25    26    27    28    29    30 
 
   # Fe 
   # 31 

data$Initiator_ID.num<-nums[as.character(data$Initiator_ID)]

data$Partner_ID.num<-nums[as.character(data$Partner_ID)]

data$Disengager_ID.num<-nums[as.character(data$Disengager_ID)]

#str(data)

###Time formatting

#library(lubridate)

options(digits.secs=6)
data$start.rem<-period_to_seconds(hms(as.character(data$Start_Caption_Stamp)))

#head(data$Start_Caption_Stamp)
#head(data$start.rem)

data$end.rem<-period_to_seconds(hms(as.character(data$End_Caption_Stamp)))
#NAs are produced for cops that end in interruptions, throwing warnings. That's fine for now. The entry for End_Caption_Stamp is still a character string.

#head(data$End_Caption_Stamp)
#head(data$end.rem)


##also should replace NA's in the Face-off column with SF (short fight)

data$Face_Off<-as.character(data$Face_Off)

data$Face_Off[is.na(data$Face_Off)]<-"SF"


#Split the dataframe up by day since the number of seconds is always reckoned from the beginning of each observation period, and we will also begin by analyzing each sequence separately.

datelist<-split(data,data$Date)


########## convert to event types for REM #############

#Go through each line and create a bunch of sets of rows, then bind them together in order at the end. Finally create a series of dataframes in a list.

#females_exact Y is 2, N is 1

##bring in pairwise distances; I have a table for each day


file_list<-list.files(here("data_processing","udata_centroid_data","pairwise_dist"))

distlist<-list()

setwd(here("data_processing","udata_centroid_data","pairwise_dist"))
distlist<-lapply(file_list, function(x) read.table(x, sep=","))

#change the column names to proper id names

for(i in 1:length(distlist)){
names(distlist[[i]])<-ids[-c(30,31)]
}
#note that this order doesn't necessarily have to match the order of ids in the interaction dataframe

#convert to martices

distlist.mat<-lapply(distlist, function(x) as.matrix(x, ))


##bring in end of observation times for each day

ends<-read.table(here("data_files","end_observation_times_CHG20140319_0406.csv"), sep=",")

ends<-as.matrix(ends)

ends<-rbind(ends, period_to_seconds(hms(ends[2,])))


#### create the eventlist ####


#make a loop through each day and then each row of the data
#for each set of events comprising an interaction we want to have females_exact, pairwise distance, and grouping factor (day)

#To reflect a change in the data collection protocol, here we implement a "reverse" coding for chase disengages, reflecting the fact that the disengager is now defined as the inititator who then leaves off chasing, rather than the receiver who is running away. This is more consistent with the idea that disengagements might represent "wins". SO, if the Face_Off column == "CH", we use the INITIATOR column for the disengager rather than the disengager column


#library(abind)

day.eventlist=list()

for(j in 1:length(datelist)){

eventlist<-list()


for(i in 1:nrow(datelist[[j]])){

if(datelist[[j]][i,]$Reaction_Code!="C"){
	entry<-matrix(nrow=3, ncol=4)
	entry[1,]<-c(paste(datelist[[j]][i,]$Initiator_ID.num,datelist[[j]][i,]$Partner_ID.num, sep=".attack."), datelist[[j]][i,]$start.rem, datelist[[j]][i,]$Females_Exact, as.character(datelist[[j]][i,]$Date))
	
	entry[2,]<-c(paste(datelist[[j]][i,]$Partner_ID.num, ".init", datelist[[j]][i,]$Face_Off, sep=""), datelist[[j]][i,]$start.rem+0.001, datelist[[j]][i,]$Females_Exact, as.character(datelist[[j]][i,]$Date))

if(as.character(datelist[[j]][i,]$Face_Off)!="CH"){
	
	
	if(datelist[[j]][i,]$Disengager_ID.num==datelist[[j]][i,]$Initiator_ID.num){
		
		entry[3,]<-c(paste(datelist[[j]][i,]$Disengager_ID.num, ".disengage.", datelist[[j]][i,]$Partner_ID.num, sep=""), datelist[[j]][i,]$end.rem, datelist[[j]][i,]$Females_Exact, as.character(datelist[[j]][i,]$Date))
	
	} else {
		
		entry[3,]<-c(paste(datelist[[j]][i,]$Disengager_ID.num, ".disengage.", datelist[[j]][i,]$Initiator_ID.num, sep=""), datelist[[j]][i,]$end.rem, datelist[[j]][i,]$Females_Exact, as.character(datelist[[j]][i,]$Date))
		
	}

} else {
	
	entry[3,]<-c(paste(datelist[[j]][i,]$Initiator_ID.num, ".disengage.", datelist[[j]][i,]$Partner_ID.num, sep=""), datelist[[j]][i,]$end.rem, datelist[[j]][i,]$Females_Exact, as.character(datelist[[j]][i,]$Date))
	
}
	
	eventlist[[i]]<-entry
}

if(datelist[[j]][i,]$Reaction_Code=="C"){
	
	if(datelist[[j]][i,]$End_Caption_Stamp=="inter"){
		entry<-matrix(nrow=1,ncol=4)
		entry[1,]<-c(paste(datelist[[j]][i,]$Initiator_ID.num, ".cop.start", sep=""), datelist[[j]][i,]$start.rem, datelist[[j]][i,]$Females_Exact, as.character(datelist[[j]][i,]$Date))
	}
	
	if(datelist[[j]][i,]$End_Caption_Stamp!="inter"){
	entry<-matrix(nrow=2,ncol=4)
	entry[1,]<-c(paste(datelist[[j]][i,]$Initiator_ID.num, ".cop.start", sep=""), datelist[[j]][i,]$start.rem, datelist[[j]][i,]$Females_Exact, as.character(datelist[[j]][i,]$Date))
	entry[2,]<-c(paste(datelist[[j]][i,]$Initiator_ID.num, ".cop.end", sep=""), datelist[[j]][i,]$end.rem, datelist[[j]][i,]$Females_Exact, as.character(datelist[[j]][i,]$Date))
}

eventlist[[i]]<-entry


}

}

eventlist[[nrow(datelist[[j]])+1]]<-c("end.obs", ends[3,j], "NA", as.character(datelist[[j]][1,]$Date))

eventlist.cat<-abind(eventlist, along=1)

day.eventlist[[j]]<-eventlist.cat

}

#str(day.eventlist)

rawevent<-abind(day.eventlist, along=1)

#order by time- end times may overlap with beginnings of other interactions, and all events need to be in order of occurrence for the rem package
rawevent<-rawevent[order(rawevent[,4], as.numeric(rawevent[,2])),]

rawevent.dat<-data.frame(rawevent)
names(rawevent.dat)<-c("Events","Time","Females", "Date")

#str(rawevent.dat)
#321 event types actually occur
#new total interaction rem events is 4658! 

#all the times from the video data are to the 1000th of a second

##checking

#this tells me whether rows are the same as the one after them, which should never happen in this dataset. 

#library(dplyr)

#which(dplyr::lag(rawevent.dat$Events)==rawevent.dat$Events)
#none!


##### departures themselves as events ######

#arrivals exogenous but departures endogenous 

pres.exact<-read.table(here("data_files","male_pres_exact_quest.csv"), sep=",",header=T)

	departlist<-list()
	
	for(j in 1:nrow(pres.exact)){
		
		arriverow<-pres.exact[j,startsWith(colnames(pres.exact),"arrive")]
		arriverow<-arriverow[!is.na(arriverow)]
		
		arrives<-t(sapply(arriverow, function(x) return(c(paste0(pres.exact$Num_ID[j],".arrive"),x,"NA",as.character(pres.exact$Date[j])))))
		
		departrow<-pres.exact[j,startsWith(colnames(pres.exact),"depart")]
		departrow<-departrow[!is.na(departrow)]
		
		departs<-t(sapply(departrow, function(x) return(c(paste0(pres.exact$Num_ID[j],".depart"),x,"NA",as.character(pres.exact$Date[j])))))
		
		allpres<-rbind(arrives,departs)
		
		departlist[[j]]<-allpres
		
	}

#collapse the departlist 

departevent<-abind(departlist, along=1)

#remove arrive at zero events since they are all simultaneous and might cause problems
departevent<-departevent[departevent[,2]!=0,]

departevent<-data.frame(departevent)
names(departevent)<-c("Events","Time","Females", "Date")

departevent$Time<-as.numeric(as.character(departevent$Time))
departevent$Females<-as.character(departevent$Females)


###Bring in Exact Female arrival and departure times

fe_exact<-read.table(here("data_files","CHG_females_exact.tsv"), header=TRUE, sep="\t")

#str(fe_exact)

#reduce to only the necessary columns- we want elapsed video time (as opposed to time of day) this time
fe_exact<-fe_exact[,c(1,3,5,7,9,11,13)]
#str(fe_exact)

#date we will leave as a factor, and then we will convert the other columns to elapsed time. This way we can apply the function by specific day.

#need to convert frames to fractional seconds

#library(lubridate)

options(digits.secs=6)

suppressWarnings(
for(i in c(2:7)){
	
	timesplit<-strsplit(as.character(fe_exact[,i]), split=":")
	
	frac<-sapply(timesplit, function(x) if(!is.na(x)) {paste(x[1],x[2],(as.numeric(x[3])+as.numeric(x[4])/30),sep=":")} else {if(i==2) "00:00:00.0" else "10:00:00.0"}
	)
	
fe_exact[,i]<-round(period_to_seconds(hms(as.character(frac))),digits=3)

	}
)


#need to max out the missing elapsed depart and arrival times like we did for the time of day 

	
#add to the whole event list and re-order again by date and time
	

#reformat

depart.mat<-as.matrix(departevent)
rawevent.mat<-as.matrix(rawevent.dat)

fullevent.mat<-rbind(depart.mat,rawevent.mat)

fullevent.mat<-fullevent.mat[order(fullevent.mat[,4], as.numeric(fullevent.mat[,2])),]


### Add exogenous clock events for time effects (every 15 mins / 900s)

#impt that they do not end up simultaneous with other events

#can have clock effects for solicitation rate and for rate of lek departure


clocklist<-list()

for(i in 1:ncol(ends)){
	
endat<-ends[c(1,3),i]	

clocktimes<-900*(1:floor(as.numeric(endat[2])/900))

clockevs<-t(sapply(clocktimes, function(x) return(c("clock.time",x,"NA",endat[1]))))

clocklist[[i]]<-clockevs
	
}

clockfull<-abind(clocklist, along=1)

clockfull<-data.frame(clockfull)
names(clockfull)<-c("Events","Time","Females", "Date")

clockfull$Time<-as.numeric(as.character(clockfull$Time))
clockfull$Females<-as.character(clockfull$Females)

#str(clockfull)


#again, re-integrate into the full event list and reorder

clockfull<-as.matrix(clockfull)

fullevent.matclock<-rbind(fullevent.mat,clockfull)

fullevent.matclock<-fullevent.matclock[order(fullevent.matclock[,4], as.numeric(fullevent.matclock[,2])),]

fullevent.dat<-data.frame(fullevent.matclock, stringsAsFactors=F)

#str(fullevent.dat)
#now I have 5122 rows!

#fullevent.dat[250:300,]


##incorporate exogenous events for Females arriving and departing

	femlist<-list()
	
	for(j in 1:nrow(fe_exact)){
		
		arriverow<-fe_exact[j,grepl("arrival",colnames(fe_exact))]
		#get rid of the dummy events that didn't actually happen
		arriverow<-arriverow[arriverow!=36000]
		
		arrives<-t(sapply(arriverow, function(x) return(c("fem.enter",x,"Y",as.character(fe_exact$Date[j])))))
		
		departrow<-fe_exact[j,grepl("depart",colnames(fe_exact))]
		departrow<-departrow[departrow!=36000]
		
		departs<-t(sapply(departrow, function(x) return(c("fem.exit",x,"N",as.character(fe_exact$Date[j])))))
		
		allpres<-rbind(arrives,departs)
		
		femlist[[j]]<-allpres
		
	}

#collapse the departlist 

femevent<-abind(femlist, along=1)

femevent<-data.frame(femevent,stringsAsFactors=F)
names(femevent)<-c("Events","Time","Females", "Date")


femevent$Time<-as.numeric(femevent$Time)


## integrate into dataset


femevent<-as.matrix(femevent)
fullevent.mat<-as.matrix(fullevent.dat)


fullevent.matfem<-rbind(fullevent.mat,femevent)

fullevent.matfem<-fullevent.matfem[order(fullevent.matfem[,4], as.numeric(fullevent.matfem[,2])),]

fullevent.dat<-data.frame(fullevent.matfem, stringsAsFactors=F)

#str(fullevent.dat)
#now 5164 events total


### incorporate exogenous events for disturbances such as predation events and planes


disturb<-read.table(here("data_files","CHG_2014_disturbance_events.csv"),sep=",",header=T)

#str(disturb)

disturb<-data.frame(Events=rep("disturb.ance",nrow(disturb)),Time=disturb$Time,Females=rep("NA",nrow(disturb)),Date=disturb$Date)

disturb<-disturb[!is.na(disturb$Time),]

disturb$Females<-as.character(disturb$Females)

#str(disturb)


#again, re-integrate into the full event list and reorder


disturb<-as.matrix(disturb)
fullevent.dist<-as.matrix(fullevent.dat)

fullevent.dist<-rbind(fullevent.dist,disturb)

fullevent.dist<-fullevent.dist[order(fullevent.dist[,4], as.numeric(fullevent.dist[,2])),]

fullevent.dist<-data.frame(fullevent.dist, stringsAsFactors=F, row.names=NULL)

#str(fullevent.dist)

#now 5168 events with the four disturbances included


###check and fix simultaneity

#this tells me which rows are the same as the one after them, which should never happen in this dataset.
#library(dplyr)

which(dplyr::lag(fullevent.dist$Time)==fullevent.dist$Time)
#none!

#now check to see if clock events or arrivals ever get in between fight attacks and receiver initiations, which would cause some problems with the support array later on

which(fullevent.dist$Events[grep("init",fullevent.dist$Events)-1]=="clock.time")
#again, luckily, never!

#lastly, check arrives and departs
"TRUE" %in% grepl("depart",fullevent.dist$Events[grep("init",fullevent.dist$Events)-1])

"TRUE" %in% grepl("arrive",fullevent.dist$Events[grep("init",fullevent.dist$Events)-1])

"TRUE" %in% grepl("enter",fullevent.dist$Events[grep("init",fullevent.dist$Events)-1])

"TRUE" %in% grepl("exit",fullevent.dist$Events[grep("init",fullevent.dist$Events)-1])

#ok, no issues here. So all functions should work ok. 


###There was an issue with the "female exact" times from the raw interaction data not giving the proper category once formatted for REM. This is due to females arriving/departing in the middle of an interaction, before the males disengage. We can apply a function to the whole dataset to apply female presence data to the new exogenous events (and endogenous male departure events) we added and to make sure to give the appropriate female presence category to every endogenous interaction event:

fullevent.dist$Time<-as.numeric(fullevent.dist$Time)

for(i in c(1:nrow(fullevent.dist))){

fullevent.dist$Females[i]<-ifelse(fullevent.dist$Time[i]>=fe_exact[fe_exact$Date==fullevent.dist$Date[i],]$third_depart_elapsed,"N",
ifelse(fullevent.dist$Time[i]<fe_exact[fe_exact$Date==fullevent.dist$Date[i],]$third_depart_elapsed && fullevent.dist$Time[i]>=fe_exact[fe_exact$Date==fullevent.dist$Date[i],]$third_arrival_elapsed,"Y", 
ifelse(fullevent.dist$Time[i]<fe_exact[fe_exact$Date==fullevent.dist$Date[i],]$third_arrival_elapsed && fullevent.dist$Time[i]>=fe_exact[fe_exact$Date==fullevent.dist$Date[i],]$second_depart_elapsed,"N",
ifelse(fullevent.dist$Time[i]<fe_exact[fe_exact$Date==fullevent.dist$Date[i],]$second_depart_elapsed && fullevent.dist$Time[i]>=fe_exact[fe_exact$Date==fullevent.dist$Date[i],]$second_arrival_elapsed, "Y",
ifelse(fullevent.dist$Time[i]<fe_exact[fe_exact$Date==fullevent.dist$Date[i],]$second_arrival_elapsed && fullevent.dist$Time[i]>=fe_exact[fe_exact$Date==fullevent.dist$Date[i],]$first_depart_elapsed, "N",
ifelse(fullevent.dist$Time[i]<fe_exact[fe_exact$Date==fullevent.dist$Date[i],]$first_depart_elapsed && fullevent.dist$Time[i]>=fe_exact[fe_exact$Date==fullevent.dist$Date[i],]$first_arrival_elapsed, "Y",
ifelse(fullevent.dist$Time[i]<fe_exact[fe_exact$Date==fullevent.dist$Date[i],]$first_arrival_elapsed, "N")))))))

}


#reformat Female column to numeric

fullevent.dist$Females<-as.numeric(as.factor(fullevent.dist$Females))

#format back into all characters for export to tsv

fullevent.dist$Females<-as.character(fullevent.dist$Females)
fullevent.dist$Time<-as.character(fullevent.dist$Time)


##export dataset

setwd(here("data_processing"))
write.table(fullevent.dist, file="relevent_format_dataset_udata_wincol_disturbance.tsv", sep='\t', row.names=F)


##### generate list of all the possible (endogenous) event types #####

#I'll use this list to create my support matrix and statlists

event.ids<-nums[-31]

attacks<-sapply(event.ids, function(x) paste(x, ".attack.", event.ids[-x], sep=""))

#flatten the matrix into a vector
attacks<-c(attacks[,1:30])

fo<-paste(event.ids, ".initFO", sep="")
ch<-paste(event.ids, ".initCH", sep="")
sf<-paste(event.ids, ".initSF", sep="")

dis<-paste(event.ids, ".disengage", sep="")

cop.s<-paste(event.ids, ".cop.start", sep="")
cop.e<-paste(event.ids, ".cop.end", sep="") 

depart<-paste(event.ids, ".depart", sep="")

arrive<-paste(event.ids, ".arrive", sep="")

event.types<-c(attacks,fo,ch,sf,dis,cop.s,cop.e, depart, arrive, "end.obs", "clock.time","fem.enter","fem.exit","disturb.ance")

length(event.types)
#now there are 1115 event types!!! but most of them are not actually possible most of the time

write.table(event.types, file="event_types_list_udata.csv", row.names=F, col.names=F)
```
