## Supplemental_reproducible_code for "Fighting isn’t sexy in lekking Greater Sage-grouse (*Centrocercus urophasianus*)": rem_udata_centroid_data.R

```
#Generate Pairwise Centroid Distance Matrices for REM Analysis

rm(list=ls())

###make sure necessary packages are there and loaded

packages<-c("here","reshape2")

package.check<-lapply(packages, FUN = function(x) {
  if (!require(x, character.only = TRUE, quietly=TRUE)) {
    install.packages(x, dependencies = TRUE)
    library(x, character.only = TRUE, quietly=TRUE)
  } 
  
})

#for project path management
#define the project root directory relative to the current script. This sets the working directory that gets called by the here() function. It is a relative path and will be robust to use on anyone's computer. In this case, I want the root directory to always be the enclosing folder for the whole sage grouse project
i_am("data_processing/rem_udata_centroid_data.R")


#set the working directory to the file with all the interaction data.
setwd(here("data_files","raw_position_data"))

#this is a way to bring in and concatenate all my interaction data files into one dataset, robust to the number of files in the folder
file_list<-list.files()

#bring files in and create a dataframe called dataset
#first make a daataframe to append to
dataset<-read.table(file_list[[1]],header=TRUE, sep="\t")

for(i in 2:length(file_list)){
	    temp_dataset <-read.table(file_list[[i]], header=TRUE, sep="\t")
    dataset<-rbind(dataset, temp_dataset)
    rm(temp_dataset)
}


#str(dataset)


###Make the Date and time into usable columns. I will make a formatted datetime column as well as one just for general time of day, and a factor for date

#turn time data into POSIXct format for the full datasetoptions(digits.secs=3)
dataset$datetime=as.POSIXct(as.character(dataset$Time_Stamp), format="%Y-%m-%d %H:%M:%OS", tz=Sys.timezone())

split<-colsplit(string=as.character(dataset$Time_Stamp),pattern=" ",names=c("Date","Time"))

dataset$Date<-split$Date
dataset$Time<-split$Time


dataset$Time=as.POSIXct(dataset$Time, format="%H:%M:%OS", tz=Sys.timezone())

#turn date into a factor (ordinal date info is stored in the datetime column)
dataset$Date=as.factor(dataset$Date)

str(dataset) #check

##IMPT##- we used the same U designations on multiple days, so we need to re-code the U male id's so that they are consistent. This will extend to the rest of the analyses as well. Can just append the date to the "U" to differentiate them while preserving the information

udata<-dataset

for(i in 1:length(levels(udata$Date))){
	
	uday<-subset(udata,udata$Date==levels(udata$Date)[i])
	
	datenum<-strsplit(as.character(uday$Date[1]), split="-")[[1]][3]
	urows<-grep("U",uday$Bird_ID)
	
	if(length(urows)>0){
	
	uday$Bird_ID[urows]<-paste(uday$Bird_ID[urows],datenum,sep=".")

udata[udata$Date==levels(udata$Date)[i],]<-uday
}
}

#table(udata$Bird_ID)

#ok

str(udata)


###### Integrate Interaction Data with Position Data##########

#bring in the data

inter<-read.table(here("data_processing","full_data_plus_cop_quest.tsv"), header=T, sep="\t")
#str(inter)

#Identify and remove rows for which the disengager ID is ambiguous

#inter[is.na(inter$Disengager_ID),][-grep("C",inter[is.na(inter$Disengager_ID),"Reaction_Code"]),]
#row 1175 has an NA in the dataset, and 109 and 114 are problematic

inter<-inter[-c(109, 114, 1175),]

#Leave in copulations and then remove centroids for Fe and X later?

#str(inter)
#1578 observations

#Have to integrate the data by treating the datasets separately at first, extracting for a particular male, and then fuse and get the centroid

##########################################

#### Get daily movement centroids and pairwise dist for REM analysis ####

#the centroid of a cluster of points is the mean of all the points in each direction

#split by date

dates_int<-split(inter,inter$Date)
#summary(dates_int)

dates_pos<-split(udata,udata$Date)
#summary(dates_pos)

#make a list of dataframes containing the x and y coordinates for the centoids for each male present

cent.list<-list()

for(i in 1:length(dates_int)){

males<-union(unique(dates_int[[i]]$Initiator_ID),unique(dates_int[[i]]$Partner_ID))

#remove X males
if("X" %in% males){males<-males[-grep("X",males)]}

#remove the Females if there were cops 
if("Fe" %in% males){males<-males[-grep("Fe",males)]}

males<-males[order(males)]

centroid<-data.frame(Bird_ID=males)


centroid$x_cent<-sapply(centroid$Bird_ID, function(x) {
	
	temp.init<-subset(dates_int[[i]],Initiator_ID==as.character(x))$Initiator_Cartesian_X
	
	temp.part<-subset(dates_int[[i]],Partner_ID==as.character(x))$Partner_Cartesian_X
	
	temp.pos<-subset(dates_pos[[i]],Bird_ID==as.character(x))$Cartesian_X
	
	mean(c(temp.init,temp.part,temp.pos))
	
	})

centroid$y_cent<-sapply(centroid$Bird_ID, function(x) {
	
	temp.init<-subset(dates_int[[i]],Initiator_ID==as.character(x))$Initiator_Cartesian_Y
	
	temp.part<-subset(dates_int[[i]],Partner_ID==as.character(x))$Partner_Cartesian_Y
	
	temp.pos<-subset(dates_pos[[i]],Bird_ID==as.character(x))$Cartesian_Y
	
	mean(c(temp.init,temp.part,temp.pos))
	
	})

cent.list[[i]]<-centroid

}

#create a matrix of pairwise distances for each day
#absent males get Na's? It doesn't matter since the support matrix will cause the model to always skip those entries.

males<-union(unique(inter$Initiator_ID),unique(inter$Partner_ID))

#remove X males
males<-males[-grep("X",males)]

#remove the Females
males<-males[-grep("Fe",males)]

names<-males[order(males)]

pair.dist<-matrix(nrow=length(names),ncol=length(names), dimnames=list(names,names))

#make a function for distance
pythag<-function(x1,y1,x2,y2){
	sqrt((x2-x1)^2+(y2-y1)^2)
}

pair.dist.list<-list()

for(k in 1:length(cent.list)){

pair.dist<-matrix(nrow=length(names),ncol=length(names), dimnames=list(names,names))

#loop through each column and row in the pair.dist matrix

for(i in 1:length(names)){
	for(j in 1:length(names)){
		
		#make sure the males in question are actually there that day, then apply the pythag function to that cell
		
		if(rownames(pair.dist)[i] %in% cent.list[[k]]$Bird_ID && rownames(pair.dist)[j] %in% cent.list[[k]]$Bird_ID) pair.dist[i,j]<-pythag(cent.list[[k]][grep(rownames(pair.dist)[i],cent.list[[k]]$Bird_ID),2],cent.list[[k]][grep(rownames(pair.dist)[i],cent.list[[k]]$Bird_ID),3], cent.list[[k]][grep(colnames(pair.dist)[j],cent.list[[k]]$Bird_ID),2], cent.list[[k]][grep(colnames(pair.dist)[j],cent.list[[k]]$Bird_ID),3])
		
	}
	
}

pair.dist.list[[k]]<-pair.dist

}

dir.create(here("data_processing","udata_centroid_data"),recursive=T)

dir.create(here("data_processing","udata_centroid_data","centroids"),recursive=T)
dir.create(here("data_processing","udata_centroid_data","pairwise_dist"),recursive=T)

#export data to save for later
setwd(here("data_processing","udata_centroid_data","centroids"))

dates<-names(dates_int)

#centroid data

for(i in 1:length(cent.list)){
	write.table(cent.list[[i]],file=paste("centroids_data_", dates[[i]], ".csv", sep=""), row.names=F, sep=",")
}

#pairwise distance matrices for each day

setwd(here("data_processing","udata_centroid_data","pairwise_dist"))

for(i in 1:length(pair.dist.list)){
	write.table(pair.dist.list[[i]],file=paste("pairwise_dist_data_", dates[[i]], ".csv", sep=""), row.names=T,sep=",")
}


##########################################
```
