## Supplemental_reproducible_code for "Fighting isn’t sexy in lekking Greater Sage-grouse (*Centrocercus urophasianus*)": genremlist_sim.R

```
## this will just say how many runs to do, and number them 

daynum<-1:18
runstart<-1
runend<-100

simjobslist<-list()

for(i in daynum){
	
	simjobslist[[i]]<-cbind(rep(i,length(runstart:runend)),runstart:runend)
	
}

jobs.list<-list()

for(i in daynum){
jobs.list[[i]]<-sapply(1:length(runstart:runend), function(x) paste(c("module load R/3.6.1-foss-2018b; Rscript rem_event_hist_simulation_cluster_udata_terr_actualday_newcensus.R", simjobslist[[i]][x,]),collapse=" ", sep=" "))
}


for(j in daynum){
write(as.matrix(jobs.list[[j]]), file=paste0("joblist_",j,".txt"))
}
```
