## Supplemental_reproducible_code for "Fighting isn’t sexy in lekking Greater Sage-grouse (*Centrocercus urophasianus*)": rem.functions.act.terr.sim.R

```
##Does the absolute pairwise distance between movement centroids predict attacks?

add.centroid.dist<-function(x){

abstat<-x

library(abind)

#loop through each day sequence
for(k in day){

#empty matrix of the right dimensions
stat<-matrix(nrow=step, ncol=length(event.types))
colnames(stat)<-event.types

#this loops through each event in the sequence
for(j in step){

#assign a value for the statistic for each potential next event type, given the current event in the sequence
for(i in 1:length(event.types)){
	
	if(grepl("attack",event.types[i])){
		ev<-unlist(strsplit(event.types[i], split=".", fixed=T))
		
		#exclude X males, here id 30
		if(!("30" %in% ev)){
		stat[j,i]<- distlist.mat[[k]][as.numeric(ev[1]),as.numeric(ev[3])]
		#leave the distance as absolute
		#if(stat[j,i]>0){stat[j,i]<-1/stat[j,i]}
		} 
		
		else {stat[j,i]<-0}
		
		}
		
		else {stat[j,i]<-0}

}
}

abstat[[k]][[1]]<-abind(abstat[[k]][[1]], stat)

dimnames(abstat[[k]][[1]])[[3]][ncol(abstat[[k]][[1]][1,,])]<-"centroid.dist"

}

return(abstat)

}


## Do females affect the hazard of attacks?

add.fem.attack<-function(x){

#get the data from the original event list


evlist.dat<-split(event.list, event.list$Date)

library(abind)

for(k in day){

#empty matrix of the right dimensions, event# x event type
stat<-matrix(0,nrow=step, ncol=length(event.types))
colnames(stat)<-event.types

#this loops through each event in the sequence
for(j in step){

#if females are present, certain event types were predicted to be affected somehow (have to deal with NA's since Females doesn't apply to end.obs)
if(evlist.dat[[k]]$Females[j]==2 && is.na(evlist.dat[[k]]$Females[j])==FALSE){

stat[j,][grep("attack",event.types)]<-1

}


}


x[[k]][[1]]<-abind(x[[k]][[1]], stat)

dimnames(x[[k]][[1]])[[3]][ncol(x[[k]][[1]][1,,])]<-"females.attack"

}

return(x)

}


## Do females affect the hazard of chases?

add.fem.ch<-function(x){

#get the data from the original event list


evlist.dat<-split(event.list, event.list$Date)

library(abind)

for(k in day){

#empty matrix of the right dimensions, event# x event type
stat<-matrix(0,nrow=step, ncol=length(event.types))
colnames(stat)<-event.types

#this loops through each event in the sequence
for(j in step){

#if females are present, certain event types were predicted to be affected somehow (have to deal with NA's since Females doesn't apply to end.obs)
if(evlist.dat[[k]]$Females[j]==2 && is.na(evlist.dat[[k]]$Females[j])==FALSE){

#assign a value for the statistic for each potential next event type, given the current event in the sequence

stat[j,][grep("initCH",event.types)]<-1

}


}


x[[k]][[1]]<-abind(x[[k]][[1]], stat)

dimnames(x[[k]][[1]])[[3]][ncol(x[[k]][[1]][1,,])]<-"females.initCH"

}

return(x)

}


## Do females affect the hazard of short fights?

add.fem.sf<-function(x){

#get the data from the original event list


evlist.dat<-split(event.list, event.list$Date)

library(abind)

for(k in day){

#empty matrix of the right dimensions, event# x event type
stat<-matrix(0,nrow=step, ncol=length(event.types))
colnames(stat)<-event.types

#this loops through each event in the sequence
for(j in step){

#if females are present, certain event types were predicted to be affected somehow (have to deal with NA's since Females doesn't apply to end.obs)
if(evlist.dat[[k]]$Females[j]==2 && is.na(evlist.dat[[k]]$Females[j])==FALSE){

#assign a value for the statistic for each potential next event type, given the current event in the sequence

stat[j,][grep("initSF",event.types)]<-1

}


}


x[[k]][[1]]<-abind(x[[k]][[1]], stat)

dimnames(x[[k]][[1]])[[3]][ncol(x[[k]][[1]][1,,])]<-"females.initSF"

}

return(x)

}


## Do females affect the hazard of Face-Offs?

add.fem.fo<-function(x){

#get the data from the original event list


evlist.dat<-split(event.list, event.list$Date)

library(abind)

for(k in day){

#empty matrix of the right dimensions, event# x event type
stat<-matrix(0,nrow=step, ncol=length(event.types))
colnames(stat)<-event.types

#this loops through each event in the sequence
for(j in step){

#if females are present, certain event types were predicted to be affected somehow (have to deal with NA's since Females doesn't apply to end.obs)
if(evlist.dat[[k]]$Females[j]==2 && is.na(evlist.dat[[k]]$Females[j])==FALSE){

#assign a value for the statistic for each potential next event type, given the current event in the sequence

stat[j,][grep("initFO",event.types)]<-1


}


}


x[[k]][[1]]<-abind(x[[k]][[1]], stat)

dimnames(x[[k]][[1]])[[3]][ncol(x[[k]][[1]][1,,])]<-"females.initFO"

}

return(x)

}


## Do females affect the hazard of disengages?

add.fem.disengage<-function(x){

#get the data from the original event list


evlist.dat<-split(event.list, event.list$Date)

library(abind)

for(k in day){

#empty matrix of the right dimensions, event# x event type
stat<-matrix(0,nrow=step, ncol=length(event.types))
colnames(stat)<-event.types

#this loops through each event in the sequence
for(j in step){

#if females are present, certain event types were predicted to be affected somehow (have to deal with NA's since Females doesn't apply to end.obs)
if(evlist.dat[[k]]$Females[j]==2 && is.na(evlist.dat[[k]]$Females[j])==FALSE){

#assign a value for the statistic for each potential next event type, given the current event in the sequence
	stat[j,][grep("disengage",event.types)]<-1
	
}

}


x[[k]][[1]]<-abind(x[[k]][[1]], stat)

dimnames(x[[k]][[1]])[[3]][ncol(x[[k]][[1]][1,,])]<-"females.disengage"

}

return(x)

}


## This statistic identifies when a copulation is occurring, and predicts that an attack on the copulating male is more likely

#the support array already takes care of what is actually possible at that particular point, so I just need to think about the types of events that might be affected

#go back to the "census" used for the support array- this tracks the behavioral state of each male through time 


add.cop.attack<-function(x){

cop.stat<-x

library(abind)

#loop through each day sequence
for(k in day){

#empty matrix of the right dimensions, event# x event type
stat<-matrix(nrow=step, ncol=length(event.types))
colnames(stat)<-event.types

#all event types are zero for this stat in the first timestep
stat[1,]<-0

#this loops through each event in the sequence, in this case starting with the second event since this stat requires a copulation to have started already in a previous timestep

for(j in step){

#if a male was actively copulating in the previous timestep according to the census (this includes the row containing the cop.start), we predict that male has an elevated hazard of being attacked in this timestep

if("copulating" %in% census.list[[k]][(j-1),]){
	
	cop<-grep("copulating", census.list[[k]][(j-1),]) 
	
	#turn on attacks directed at copulating males (robust to potential simultaneous copulations)
	
	for(h in cop){stat[j,][endsWith(event.types ,paste(".attack.", h, sep=""))]<-1}
	
#turn everything else off
	
	stat[j,][is.na(stat[j,])]<-0	
}

#if no one was copulating in the previous timestep, nothing special is predicted 

else { stat[j,]<-0 }

}

cop.stat[[k]][[1]]<-abind(cop.stat[[k]][[1]], stat)

dimnames(cop.stat[[k]][[1]])[[3]][ncol(cop.stat[[k]][[1]][1,,])]<-"cop.attack"

}

return(cop.stat)

}

############# Recency and Reciprocity ############

#many of the network-level measures like eigcent correlate highly with, and are likely largely explained by more proximate processes such as recency and reciprocity. This is likely to be true for this system, especially given that not all males are present every day. 

#I can think of reciprocity in terms of general interaction, attacks, or wins/losses. These all tell different stories and are relevant to identifying deterrence, "dominance," and punishment behaviors. 

##are you more likely to attack me if we have interacted in any way (attack AND Receive) recently?

#what if I change it from fight to attack+receive, since that's how I do it for the rest of them?

add.recency<-function(x){

rec.stat<-x

library(abind)

#loop through each day sequence
for(k in day){

#empty matrix of the right dimensions, event# x event type. In this case, the default is all zeroes. 
stat<-matrix(0, nrow=step, ncol=length(event.types))
colnames(stat)<-event.types


#this loops through each event in the sequence

for(j in step){

#for each event, loop through all the males to get the recency ranks that will predict the next interaction- the support array will take care of males that weren't present that day.
for(i in 1:(ncol(census.list[[1]]))){
	
#This is all the event steps during which fighting was happening for a particular male i, up to the current timestep, in order
#for looking at recency of attacks, change this from fight to attack. This predictor here is non-directional
fighting.times<-c(grep("attack",census.list[[k]][1:(j-1),i]),grep("receive",census.list[[k]][1:(j-1),i]))

#if the male has not had any interactions yet, the hazard is equal across event types involving them, so set all to zero 
if(length(fighting.times)>0){
	
#get the names of the interaction events put in descending order, most recent first

fighting.events.desc<-census.list[[k]][1:(j-1),i][fighting.times][order(1:length(fighting.times), decreasing=T)]

#Strip down to just the id of the interaction partners
fighting.events.desc.str<-sapply(strsplit(fighting.events.desc,".",fixed=T), function(x) x[3])

#get rid of duplicates and leave the most recent instance, leaving a list of fights ranked by recency
rankids<-unique(fighting.events.desc.str)	

#match with event types- for general interactions, this recency predicts the hazard of incoming AND outgoing attacks
#loop thru each element of rankids, the first being the most recent

for(l in 1:length(rankids)){

#incoming attacks directed at the focal male
stat[j,][paste(rankids[l],".attack.", i, sep="")]<-1/l

	}
		
}
	
}

}


rec.stat[[k]][[1]]<-abind(rec.stat[[k]][[1]], stat)

dimnames(rec.stat[[k]][[1]])[[3]][ncol(rec.stat[[k]][[1]][1,,])]<-"pure.recency"

}

return(rec.stat)

}


##Are you more likely to attack me if you attacked me recently?

add.pers<-function(x){

rec.stat<-x

library(abind)

#loop through each day sequence
for(k in day){

#empty matrix of the right dimensions, event# x event type. In this case, the default is all zeroes. 
stat<-matrix(0, nrow=step, ncol=length(event.types))
colnames(stat)<-event.types


#this loops through each event in the sequence

for(j in step){

#for each event, loop through all the males to get the recency ranks that will predict the next interaction- the support array will take care of males that weren't present that day.
for(i in 1:(ncol(census.list[[1]]))){
	
#This is all the event steps during which fighting was happening for a particular male i, up to the one previous timestep, in order. Remember, stat in this row is mean to predict the CURRENT event
#for looking at recency of attacks, change this from fight to received
fighting.times<-grep("received",census.list[[k]][1:(j-1),i])

#if the male has not had any interactions yet, the hazard is equal across event types involving them, so set all to zero 
if(length(fighting.times)>0){
	
#get the names of the interaction events put in descending order, most recent first

fighting.events.desc<-census.list[[k]][1:(j-1),i][fighting.times][order(1:length(fighting.times), decreasing=T)]

#get rid of duplicates and leave the most recent instance, leaving a list of fights ranked by recency
recent.ev<-unique(fighting.events.desc)	

#get just the interacting male id number
rankids<-sapply(recent.ev, function(x) strsplit(x, split=".", fixed=T)[[1]][3])
	
#match with event types- this stat predicts OUTGOING attacks based on previous INCOMING attacks
#loop thru each element of rankids, the first being the most recent

for(l in 1:length(rankids)){

#incoming attacks directed at the focal male
stat[j,][paste(rankids[l],".attack.", i, sep="")]<-1/l
	}
		
}
	
}

}


rec.stat[[k]][[1]]<-abind(rec.stat[[k]][[1]], stat)

dimnames(rec.stat[[k]][[1]])[[3]][ncol(rec.stat[[k]][[1]][1,,])]<-"persistence"

}

return(rec.stat)

}


##are you more likely to attack me if I attacked you recently? (as compared to out-of-the-blue)

add.reciprocity<-function(x){

rec.stat<-x

library(abind)

#loop through each day sequence
for(k in day){

#empty matrix of the right dimensions, event# x event type. In this case, the default is all zeroes. 
stat<-matrix(0, nrow=step, ncol=length(event.types))
colnames(stat)<-event.types


#this loops through each event in the sequence, in this case starting with the fourth event since this stat requires at least one previous fight to have taken place

for(j in step){

#for each event, loop through all the males to get the recency ranks that will predict the next interaction- the support array will take care of males that weren't present that day.
for(i in 1:ncol(census.list[[1]])){
	
#This is all the event steps during which fighting was happening for a particular male i, up to the the one previous timestep, in order
#in this case I am looking for whom the focal male attacked most recently
fighting.times<-grep("attack",census.list[[k]][1:(j-1),i])

#if the male has not had any interactions yet, the hazard is equal across event types involving them, so set all to zero 
if(length(fighting.times)>0){
	
#get the names of the interaction events put in descending order, most recent first

fighting.events.desc<-census.list[[k]][1:(j-1),i][fighting.times][order(1:length(fighting.times), decreasing=T)]

#get rid of duplicates and leave the most recent instance, leaving a list of fights ranked by recency
recent.ev<-unique(fighting.events.desc)	

#get just the interacting male id number
rankids<-sapply(recent.ev, function(x) strsplit(x, split=".", fixed=T)[[1]][3])
	
#match with event types- this stat predicts new OUTGOING attacks based on previous OUTGOING attacks
#loop thru each element of rankids, the first being the most recent

for(l in 1:length(rankids)){

#incoming attacks directed at the focal male
stat[j,][paste(rankids[l],".attack.", i, sep="")]<-1/l
	}
		
}
	
}

}


rec.stat[[k]][[1]]<-abind(rec.stat[[k]][[1]], stat)

dimnames(rec.stat[[k]][[1]])[[3]][ncol(rec.stat[[k]][[1]][1,,])]<-"reciprocity"

}

return(rec.stat)

}


#### Explore Recency Winner/Loser effects ####

##Are you more likely to attack me if I WON against you recently? (is winning protective?)

add.win.rec<-function(x){

new.stat<-x

library(abind)

#loop through each day sequence
for(k in day){

#empty matrix of the right dimensions, event# x event type. In this case, the default is all zeroes. 
stat<-matrix(0, nrow=step, ncol=length(event.types))
colnames(stat)<-event.types


#this loops through each event in the sequence

for(j in step){

#for each event, loop through all the males to get the recency ranks that will predict the next interaction- the support array will take care of males that weren't present that day.
for(i in 1:ncol(census.list[[1]])){
	
#This is all the event steps during which fighting was happening for a particular male i, up to the current timestep, in order
#in this case I am looking for whom the focal male won against most recently
win.times<-win.loss.census[[k]][1:(j-1),i][grep("win",win.loss.census[[k]][1:(j-1),i])]

#if the male has not had any interactions yet, the hazard is equal across event types involving them, so set all to zero 
if(length(win.times)>0){

#strip the "chase" from the end of chase wins so I can include them and lump them with normal wins
win.times<-sapply(win.times, function(x) if(grepl("chas",x)==TRUE){
temp<-strsplit(x, split=".", fixed=T)[[1]]	
new<-paste(temp[1],temp[2],temp[3], sep=".")
return(new)
}
else return(x)
)


#get the names of the interaction events put in descending order, most recent first

win.events.desc<-win.times[order(1:length(win.times), decreasing=T)]

#get rid of duplicates and leave the most recent instance, leaving a list of fights ranked by recency
recent.ev<-unique(win.events.desc)	

#get just the interacting male id number
rankids<-sapply(recent.ev, function(x) strsplit(x, split=".", fixed=T)[[1]][3])
	
#match with event types- this stat predicts new OUTGOING attacks based on previous WINS
#loop thru each element of rankids, the first being the most recent

for(l in 1:length(rankids)){

#incoming attacks directed at the focal male
stat[j,][paste(rankids[l],".attack.", i, sep="")]<-1/l
	}
		
}
	
}

}


new.stat[[k]][[1]]<-abind(new.stat[[k]][[1]], stat)

dimnames(new.stat[[k]][[1]])[[3]][ncol(new.stat[[k]][[1]][1,,])]<-"win.rec"

}

return(new.stat)

}


##Control for Chases: Are you more likely to attack me if I WON a CHASE against you recently? (equivalent to attacked you+chase)

add.win.chase.rec<-function(x){

new.stat<-x

library(abind)

#loop through each day sequence
for(k in day){

#empty matrix of the right dimensions, event# x event type. In this case, the default is all zeroes. 
stat<-matrix(0, nrow=step, ncol=length(event.types))
colnames(stat)<-event.types


#this loops through each event in the sequence

for(j in step){

#for each event, loop through all the males to get the recency ranks that will predict the next interaction- the support array will take care of males that weren't present that day.
for(i in 1:ncol(census.list[[1]])){
	
#This is all the event steps during which fighting was happening for a particular male i, up to the current timestep, in order
#in this case I am looking for whom the focal male attacked most recently
win.times<-grep("win",win.loss.census[[k]][1:(j-1),i])

#want the subset of these that were chases
win.chase.times<-win.loss.census[[k]][1:(j-1),i][win.times][endsWith(win.loss.census[[k]][1:(j-1),i][win.times],"chase")]


#if the male has not had any interactions yet, the hazard is equal across event types involving them, so set all to zero 
if(length(win.chase.times)>0){
	
#get the names of the interaction events put in descending order, most recent first
win.events.desc<-win.chase.times[order(1:length(win.chase.times), decreasing=T)]

#get rid of duplicates and leave the most recent instance, leaving a list of fights ranked by recency
recent.ev<-unique(win.events.desc)	

#get just the interacting male id number
rankids<-sapply(recent.ev, function(x) strsplit(x, split=".", fixed=T)[[1]][3])
	
#match with event types- this stat predicts new OUTGOING attacks based on previous OUTGOING attacks
#loop thru each element of rankids, the first being the most recent

for(l in 1:length(rankids)){

#incoming attacks directed at the focal male
stat[j,][paste(rankids[l],".attack.", i, sep="")]<-1/l
	}
		
}
	
}

}


new.stat[[k]][[1]]<-abind(new.stat[[k]][[1]], stat)

dimnames(new.stat[[k]][[1]])[[3]][ncol(new.stat[[k]][[1]][1,,])]<-"win.chase.rec"

}

return(new.stat)

}


##Are you more likely to attack me if I WON a non-chase against you recently? (is winning protective?)

add.win.no.ch.rec<-function(x){

new.stat<-x

library(abind)

#loop through each day sequence
for(k in day){

#empty matrix of the right dimensions, event# x event type. In this case, the default is all zeroes. 
stat<-matrix(0, nrow=step, ncol=length(event.types))
colnames(stat)<-event.types


#this loops through each event in the sequence

for(j in step){

#for each event, loop through all the males to get the recency ranks that will predict the next interaction- the support array will take care of males that weren't present that day.
for(i in 1:ncol(census.list[[1]])){
	
#This is all the event steps during which fighting was happening for a particular male i, up to the current timestep, in order
#in this case I am looking for whom the focal male won against most recently

int.vec<-win.loss.census[[k]][1:(j-1),i]

win.times<-grep("TRUE",(grepl("win",int.vec) & !grepl("chase",int.vec)))

#if the male has not had any interactions yet, the hazard is equal across event types involving them, so set all to zero 
if(length(win.times)>0){
	
#get the names of the interaction events put in descending order, most recent first

win.events.desc<-int.vec[win.times][order(1:length(win.times), decreasing=T)]

#get rid of duplicates and leave the most recent instance, leaving a list of fights ranked by recency
recent.ev<-unique(win.events.desc)	

#get just the interacting male id number
rankids<-sapply(recent.ev, function(x) strsplit(x, split=".", fixed=T)[[1]][3])
	
#match with event types- this stat predicts new OUTGOING attacks based on previous WINS
#loop thru each element of rankids, the first being the most recent

for(l in 1:length(rankids)){

#incoming attacks directed at the focal male
stat[j,][paste(rankids[l],".attack.", i, sep="")]<-1/l
	}
		
}
	
}

}


new.stat[[k]][[1]]<-abind(new.stat[[k]][[1]], stat)

dimnames(new.stat[[k]][[1]])[[3]][ncol(new.stat[[k]][[1]][1,,])]<-"win.noCH.rec"

}

return(new.stat)

}


##Are you more likely to attack me if I LOST against you recently?

add.lose.rec<-function(x){

new.stat<-x

library(abind)

#loop through each day sequence
for(k in day){

#empty matrix of the right dimensions, event# x event type. In this case, the default is all zeroes. 
stat<-matrix(0, nrow=step, ncol=length(event.types))
colnames(stat)<-event.types


#this loops through each event in the sequence

for(j in step){

#for each event, loop through all the males to get the recency ranks that will predict the next interaction- the support array will take care of males that weren't present that day.
for(i in 1:ncol(census.list[[1]])){
	
#This is all the event steps during which fighting was happening for a particular male i, up to the current timestep, in order
#in this case I am looking for whom the focal male lost to most recently
lose.times<-win.loss.census[[k]][1:(j-1),i][grep("lose",win.loss.census[[k]][1:(j-1),i])]

#if the male has not had any interactions yet, the hazard is equal across event types involving them, so set all to zero 
if(length(lose.times)>0){
	
#strip the "chase" from the end of chase wins so I can include them and lump them with normal wins
lose.times<-sapply(lose.times, function(x) if(grepl("chas",x)==TRUE){
temp<-strsplit(x, split=".", fixed=T)[[1]]	
new<-paste(temp[1],temp[2],temp[3], sep=".")
return(new)
}
else return(x)
)

#get the names of the interaction events put in descending order, most recent first

lose.events.desc<-lose.times[order(1:length(lose.times), decreasing=T)]

#get rid of duplicates and leave the most recent instance, leaving a list of losses ranked by recency
recent.ev<-unique(lose.events.desc)	

#get just the interacting male id number
rankids<-sapply(recent.ev, function(x) strsplit(x, split=".", fixed=T)[[1]][3])
	
#match with event types- this stat predicts new INCOMING attacks based on previous LOSSES
#loop thru each element of rankids, the first being the most recent

for(l in 1:length(rankids)){

#incoming attacks directed at the focal male
stat[j,][paste(rankids[l],".attack.", i, sep="")]<-1/l
	}
		
}
	
}

}


new.stat[[k]][[1]]<-abind(new.stat[[k]][[1]], stat)

dimnames(new.stat[[k]][[1]])[[3]][ncol(new.stat[[k]][[1]][1,,])]<-"lose.rec"

}

return(new.stat)

}


##Control for Chases: Are you more liekly to attack me if I LOST a CHASE against you recently? (equivalent to you attacked me+chase)


add.lose.chase.rec<-function(x){

new.stat<-x

library(abind)

#loop through each day sequence
for(k in day){

#empty matrix of the right dimensions, event# x event type. In this case, the default is all zeroes. 
stat<-matrix(0, nrow=step, ncol=length(event.types))
colnames(stat)<-event.types


#this loops through each event in the sequence

for(j in step){

#for each event, loop through all the males to get the recency ranks that will predict the next interaction- the support array will take care of males that weren't present that day.
for(i in 1:ncol(census.list[[1]])){
	
#This is all the event steps during which fighting was happening for a particular male i, up to the current timestep, in order
#in this case I am looking for whom the focal male attacked most recently
lose.times<-grep("lose",win.loss.census[[k]][1:(j-1),i])

#want the subset of these that were chases
lose.chase.times<-win.loss.census[[k]][1:(j-1),i][lose.times][endsWith(win.loss.census[[k]][1:(j-1),i][lose.times],"chase")]


#if the male has not had any interactions yet, the hazard is equal across event types involving them, so set all to zero 
if(length(lose.chase.times)>0){
	
#get the names of the interaction events put in descending order, most recent first
lose.events.desc<-lose.chase.times[order(1:length(lose.chase.times), decreasing=T)]

#get rid of duplicates and leave the most recent instance, leaving a list of fights ranked by recency
recent.ev<-unique(lose.events.desc)	

#get just the interacting male id number
rankids<-sapply(recent.ev, function(x) strsplit(x, split=".", fixed=T)[[1]][3])
	
#match with event types- this stat predicts new INCOMING attacks based on previous LOST CHASES
#loop thru each element of rankids, the first being the most recent

for(l in 1:length(rankids)){

#incoming attacks directed at the focal male
stat[j,][paste(rankids[l],".attack.", i, sep="")]<-1/l
	}
		
}
	
}

}


new.stat[[k]][[1]]<-abind(new.stat[[k]][[1]], stat)

dimnames(new.stat[[k]][[1]])[[3]][ncol(new.stat[[k]][[1]][1,,])]<-"lose.chase.rec"

}

return(new.stat)

}


##Are you more likely to attack me if I LOST a non-chase against you recently?

add.lose.no.ch.rec<-function(x){

new.stat<-x

library(abind)

#loop through each day sequence
for(k in day){

#empty matrix of the right dimensions, event# x event type. In this case, the default is all zeroes. 
stat<-matrix(0, nrow=step, ncol=length(event.types))
colnames(stat)<-event.types


#this loops through each event in the sequence

for(j in step){

#for each event, loop through all the males to get the recency ranks that will predict the next interaction- the support array will take care of males that weren't present that day.
for(i in 1:ncol(census.list[[1]])){
	
#This is all the event steps during which fighting was happening for a particular male i, up to the current timestep, in order
#in this case I am looking for whom the focal male won against most recently

int.vec<-win.loss.census[[k]][1:(j-1),i]

win.times<-grep("TRUE",(grepl("lose",int.vec) & !grepl("chase",int.vec)))

#if the male has not had any interactions yet, the hazard is equal across event types involving them, so set all to zero 
if(length(win.times)>0){
	
#get the names of the interaction events put in descending order, most recent first

win.events.desc<-int.vec[win.times][order(1:length(win.times), decreasing=T)]

#get rid of duplicates and leave the most recent instance, leaving a list of fights ranked by recency
recent.ev<-unique(win.events.desc)	

#get just the interacting male id number
rankids<-sapply(recent.ev, function(x) strsplit(x, split=".", fixed=T)[[1]][3])
	
#match with event types- this stat predicts new OUTGOING attacks based on previous LOSSES
#loop thru each element of rankids, the first being the most recent

for(l in 1:length(rankids)){

#incoming attacks directed at the focal male
stat[j,][paste(rankids[l],".attack.", i, sep="")]<-1/l
	}
		
}
	
}

}


new.stat[[k]][[1]]<-abind(new.stat[[k]][[1]], stat)

dimnames(new.stat[[k]][[1]])[[3]][ncol(new.stat[[k]][[1]][1,,])]<-"lose.noCH.rec"

}

return(new.stat)

}


## B->A | (A->B + AwinB)
######

add.attack.win.rec<-function(x){

new.stat<-x

library(abind)

#loop through each day sequence
for(k in day){

#empty matrix of the right dimensions, event# x event type. In this case, the default is all zeroes. 
stat<-matrix(0, nrow=step, ncol=length(event.types))
colnames(stat)<-event.types


#this loops through each event in the sequence

for(j in step){

#for each event, loop through all the males to get the recency ranks that will predict the next interaction- the support array will take care of males that weren't present that day.
for(i in 1:ncol(census.list[[1]])){
	
#want to start the loop for the stat after the first interaction is complete, but I want to start the accounting of the past interactions from the beginning

#Remove the "fights" and "nothings" from the census list. This creates attack/receive and win/lose pairs in order of occurence
fight.times<-grep("fight",win.loss.census[[k]][1:(j-1),i])
nothing.times<-grep("nothing",win.loss.census[[k]][1:(j-1),i])

pairs<-win.loss.census[[k]][1:(j-1),i][-c(fight.times, nothing.times)]

#Now I can go either way with it. Here I want all the attacks that are followed by wins

#need a way to do this that matches interaction partners but also is sensitive to the time-ordering 

attack.wins<-vector()

if(length(pairs)>0){

attack.wins<-unlist(sapply(1:ncol(census.list[[1]]), function(z){
	
	intset<-pairs[sapply(pairs, function(x) {
		
		strsplit(x,split=".",fixed=T)[[1]][3]==z
		
	})]
	
	names(intset)<-grep("TRUE",sapply(pairs, function(x) {
		
		strsplit(x,split=".",fixed=T)[[1]][3]==z
		
	}))
	
	attacks<-grep("attack",intset)
	attackpluswin<-intset[attacks+1][grep("win", intset[attacks+1])]
	
	return(attackpluswin)
	
}))
}


#if the male has not had any interactions yet, the hazard is equal across event types involving them, so set all to zero 
if(length(attack.wins)>0){

#strip the "chase" from the end of chase wins so I can include them and lump them with normal wins
attack.wins<-sapply(attack.wins, function(x) if(grepl("chas",x)==TRUE){
temp<-strsplit(x, split=".", fixed=T)[[1]]	
new<-paste(temp[1],temp[2],temp[3], sep=".")
return(new)
}
else return(x)
)

	
#get the names of the interaction events put in descending order, most recent first
attack.wins<-attack.wins[order(names(attack.wins), decreasing=T)]

#get rid of duplicates and leave the most recent instance, leaving a list of instances ranked by recency
recent.ev<-unique(attack.wins)	

#get just the interacting male id number
rankids<-sapply(recent.ev, function(x) strsplit(x, split=".", fixed=T)[[1]][3])
	
#match with event types- this stat predicts new INCOMING attacks based on previous OUTGOING attacks + WINS
#loop thru each element of rankids, the first being the most recent

for(l in 1:length(rankids)){

#incoming attacks directed at the focal male
stat[j,][paste(rankids[l],".attack.", i, sep="")]<-1/l
	}
		
}
	
}

}


new.stat[[k]][[1]]<-abind(new.stat[[k]][[1]], stat)

dimnames(new.stat[[k]][[1]])[[3]][ncol(new.stat[[k]][[1]][1,,])]<-"attack.win.rec"

}

return(new.stat)

}


## B->A | (A->B + AloseB)
######
add.attack.lose.rec<-function(x){

new.stat<-x

library(abind)

#loop through each day sequence
for(k in day){

#empty matrix of the right dimensions, event# x event type. In this case, the default is all zeroes. 
stat<-matrix(0, nrow=step, ncol=length(event.types))
colnames(stat)<-event.types


#this loops through each event in the sequence

for(j in step){

#for each event, loop through all the males to get the recency ranks that will predict the next interaction- the support array will take care of males that weren't present that day.
for(i in 1:ncol(census.list[[1]])){
	
#want to start the loop for the stat after the first interaction is complete, but I want to start the accounting of the past interactions from the beginning

#Remove the "fights" and "nothings" from the census list. This creates attack/receive and win/lose pairs in order of occurence
fight.times<-grep("fight",win.loss.census[[k]][1:(j-1),i])
nothing.times<-grep("nothing",win.loss.census[[k]][1:(j-1),i])

pairs<-win.loss.census[[k]][1:(j-1),i][-c(fight.times, nothing.times)]

#need a way to do this that matches interaction partners but also is sensitive to the time-ordering 

attack.losses<-vector()

if(length(pairs)>0){

attack.losses<-unlist(sapply(1:ncol(census.list[[1]]), function(z){
	
	intset<-pairs[sapply(pairs, function(x) {
		
		strsplit(x,split=".",fixed=T)[[1]][3]==z
		
	})]
	
	names(intset)<-grep("TRUE",sapply(pairs, function(x) {
		
		strsplit(x,split=".",fixed=T)[[1]][3]==z
		
	}))
	
	attacks<-grep("attack",intset)
	attackpluslose<-intset[attacks+1][grep("lose", intset[attacks+1])]
	
	return(attackpluslose)

}))
}


#if the male has not had any interactions yet, the hazard is equal across event types involving them, so set all to zero 
if(length(attack.losses)>0){
	
#strip the "chase" from the end of chase wins so I can include them and lump them with normal wins
attack.losses<-sapply(attack.losses, function(x) if(grepl("chas",x)==TRUE){
temp<-strsplit(x, split=".", fixed=T)[[1]]	
new<-paste(temp[1],temp[2],temp[3], sep=".")
return(new)
}
else return(x)
)

	
#get the names of the interaction events put in descending order, most recent first
attack.losses<-attack.losses[order(names(attack.losses), decreasing=T)]

#get rid of duplicates and leave the most recent instance, leaving a list of instances ranked by recency
recent.ev<-unique(attack.losses)	

#get just the interacting male id number
rankids<-sapply(recent.ev, function(x) strsplit(x, split=".", fixed=T)[[1]][3])
	
#match with event types- this stat predicts new INCOMING attacks based on previous OUTGOING attacks + LOSSES
#loop thru each element of rankids, the first being the most recent

for(l in 1:length(rankids)){

#incoming attacks directed at the focal male
stat[j,][paste(rankids[l],".attack.", i, sep="")]<-1/l
	}
		
}
	
}

}


new.stat[[k]][[1]]<-abind(new.stat[[k]][[1]], stat)

dimnames(new.stat[[k]][[1]])[[3]][ncol(new.stat[[k]][[1]][1,,])]<-"attack.lose.rec"

}

return(new.stat)

}


## B->A | (B->A + AwinB)
######


add.receive.win.rec<-function(x){

new.stat<-x

library(abind)

#loop through each day sequence
for(k in day){

#empty matrix of the right dimensions, event# x event type. In this case, the default is all zeroes. 
stat<-matrix(0, nrow=step, ncol=length(event.types))
colnames(stat)<-event.types


#this loops through each event in the sequence

for(j in step){

#for each event, loop through all the males to get the recency ranks that will predict the next interaction- the support array will take care of males that weren't present that day.
for(i in 1:ncol(census.list[[1]])){
	
#want to start the loop for the stat after the first interaction is complete, but I want to start the accounting of the past interactions from the beginning

#Remove the "fights" and "nothings" from the census list. This creates attack/receive and win/lose pairs in order of occurence
fight.times<-grep("fight",win.loss.census[[k]][1:(j-1),i])
nothing.times<-grep("nothing",win.loss.census[[k]][1:(j-1),i])

pairs<-win.loss.census[[k]][1:(j-1),i][-c(fight.times, nothing.times)]

#Now I can go either way with it. Here I want all the receives that are followed by wins

#need a way to do this that matches interaction partners but also is sensitive to the time-ordering 

receive.wins<-vector()

if(length(pairs)>0){

receive.wins<-unlist(sapply(1:ncol(census.list[[1]]), function(z){
	
	intset<-pairs[sapply(pairs, function(x) {
		
		strsplit(x,split=".",fixed=T)[[1]][3]==z
		
	})]
	
	names(intset)<-grep("TRUE",sapply(pairs, function(x) {
		
		strsplit(x,split=".",fixed=T)[[1]][3]==z
		
	}))
	
	receives<-grep("receive",intset)
	receivepluswin<-intset[receives+1][grep("win", intset[receives+1])]
	
	return(receivepluswin)

}))
}


#if the male has not had any interactions yet, the hazard is equal across event types involving them, so set all to zero 
if(length(receive.wins)>0){
	
#strip the "chase" from the end of chase wins so I can include them and lump them with normal wins
receive.wins<-sapply(receive.wins, function(x) if(grepl("chas",x)==TRUE){
temp<-strsplit(x, split=".", fixed=T)[[1]]	
new<-paste(temp[1],temp[2],temp[3], sep=".")
return(new)
}
else return(x)
)

#get the names of the interaction events put in descending order, most recent first
receive.wins<-receive.wins[order(names(receive.wins), decreasing=T)]

#get rid of duplicates and leave the most recent instance, leaving a list of instances ranked by recency
recent.ev<-unique(receive.wins)	

#get just the interacting male id number
rankids<-sapply(recent.ev, function(x) strsplit(x, split=".", fixed=T)[[1]][3])
	
#match with event types- this stat predicts new INCOMING attacks based on previous INCOMING attacks + WINS
#loop thru each element of rankids, the first being the most recent

for(l in 1:length(rankids)){

#incoming attacks directed at the focal male
stat[j,][paste(rankids[l],".attack.", i, sep="")]<-1/l
	}
		
}
	
}

}


new.stat[[k]][[1]]<-abind(new.stat[[k]][[1]], stat)

dimnames(new.stat[[k]][[1]])[[3]][ncol(new.stat[[k]][[1]][1,,])]<-"receive.win.rec"

}

return(new.stat)

}


## B->A | (B->A + AloseB)
######

add.receive.lose.rec<-function(x){

new.stat<-x

library(abind)

#loop through each day sequence
for(k in day){

#empty matrix of the right dimensions, event# x event type. In this case, the default is all zeroes. 
stat<-matrix(0, nrow=step, ncol=length(event.types))
colnames(stat)<-event.types


#this loops through each event in the sequence

for(j in step){

#for each event, loop through all the males to get the recency ranks that will predict the next interaction- the support array will take care of males that weren't present that day.
for(i in 1:ncol(census.list[[1]])){
	
#want to start the loop for the stat after the first interaction is complete, but I want to start the accounting of the past interactions from the beginning

#Remove the "fights" and "nothings" from the census list. This creates attack/receive and win/lose pairs in order of occurence
fight.times<-grep("fight",win.loss.census[[k]][1:(j-1),i])
nothing.times<-grep("nothing",win.loss.census[[k]][1:(j-1),i])

pairs<-win.loss.census[[k]][1:(j-1),i][-c(fight.times, nothing.times)]

#Now I can go either way with it. Here I want all the receives that are followed by losses

#need a way to do this that matches interaction partners but also is sensitive to the time-ordering 

receive.losses<-vector()

if(length(pairs)>0){

receive.losses<-unlist(sapply(1:ncol(census.list[[1]]), function(z){
	
	intset<-pairs[sapply(pairs, function(x) {
		
		strsplit(x,split=".",fixed=T)[[1]][3]==z
		
	})]
	
	names(intset)<-grep("TRUE",sapply(pairs, function(x) {
		
		strsplit(x,split=".",fixed=T)[[1]][3]==z
		
	}))
	
	receives<-grep("receive",intset)
	receivepluslose<-intset[receives+1][grep("lose", intset[receives+1])]
	
	return(receivepluslose)

}))
}

#if the male has not had any interactions yet, the hazard is equal across event types involving them, so set all to zero 
if(length(receive.losses)>0){

#strip the "chase" from the end of chase wins so I can include them and lump them with normal wins
receive.losses<-sapply(receive.losses, function(x) if(grepl("chas",x)==TRUE){
temp<-strsplit(x, split=".", fixed=T)[[1]]	
new<-paste(temp[1],temp[2],temp[3], sep=".")
return(new)
}
else return(x)
)

	
#get the names of the interaction events put in descending order, most recent first
receive.losses<-receive.losses[order(names(receive.losses), decreasing=T)]

#get rid of duplicates and leave the most recent instance, leaving a list of instances ranked by recency
recent.ev<-unique(receive.losses)	

#get just the interacting male id number
rankids<-sapply(recent.ev, function(x) strsplit(x, split=".", fixed=T)[[1]][3])
	
#match with event types- this stat predicts new INCOMING attacks based on previous INCOMING attacks + LOSSES
#loop thru each element of rankids, the first being the most recent

for(l in 1:length(rankids)){

#incoming attacks directed at the focal male
stat[j,][paste(rankids[l],".attack.", i, sep="")]<-1/l

	}
		
}
	
}

}


new.stat[[k]][[1]]<-abind(new.stat[[k]][[1]], stat)

dimnames(new.stat[[k]][[1]])[[3]][ncol(new.stat[[k]][[1]][1,,])]<-"receive.lose.rec"

}

return(new.stat)

}


######## General Effects of absolute recency #########

###Am I generally more likely to be attacked if I was recently ______ in terms of amount of time since the event? ####


###########################

##Am I generally more liekly to be attacked if I interacted at all recently (in seconds)?

add.gen.recency<-function(x){

rec.stat<-x

library(abind)

event.list.dat<-split(event.list,event.list$Date)

#loop through each day sequence
for(k in day){

#empty matrix of the right dimensions, event# x event type. In this case, the default is all zeroes. 
stat<-matrix(0, nrow=step, ncol=length(event.types))
colnames(stat)<-event.types


#this loops through each event in the sequence

for(j in step){

#for each event, loop through all the males- the support array will take care of males that weren't present that day.
for(i in 1:ncol(census.list[[1]])){
	
#This is all the event steps during which fighting was happening for a particular male i, up to the current timestep, in order
#for looking at recency of attacks, change this from fight to attack. This predictor here is non-directional
wins<-grep("win",win.loss.census[[k]][1:(j-1),i])
losses<-grep("lose",win.loss.census[[k]][1:(j-1),i])
end.times<-union(wins,losses)

#if the male has not had any interactions yet, the hazard is equal across event types involving them, so set all to zero 
if(length(end.times)>0){
	
#identify the most recent win or loss- this represents the end time of the most recent interaction for that male

most.rec<-max(end.times)

#get the time of the most recent end of interaction in seconds
most.rec.sec<-event.list.dat[[k]][most.rec,"Time"]

#get the difference with the current time
current.time<-event.list.dat[[k]][(j-1),"Time"]

time.diff<-(current.time-most.rec.sec)


#This will produce "infinities" in the stat array when the current event IS the disengage event. the support array should take care of this since males cannot be attacked before disengaging, but just in case, make it sqrt(timediff+1)


#match with event types- for general interactions, this recency predicts the hazard of incoming attacks

#I use a slightly more slowly decaying function here since the variance in recency can be quite large. Can explore other formulations later. 
#incoming attacks directed at the focal male
stat[j,][endsWith(event.types,paste(".attack.",i, sep=""))]<-1/sqrt(time.diff+1)

  		

}
	
}

}


rec.stat[[k]][[1]]<-abind(rec.stat[[k]][[1]], stat)

dimnames(rec.stat[[k]][[1]])[[3]][ncol(rec.stat[[k]][[1]][1,,])]<-"gen.recency"

}

return(rec.stat)

}


##Am I generally more liekly to be attacked if I attacked another male recently (in seconds)?

add.gen.attack.rec<-function(x){

rec.stat<-x

library(abind)

event.list.dat<-split(event.list,event.list$Date)

#loop through each day sequence
for(k in day){

#empty matrix of the right dimensions, event# x event type. In this case, the default is all zeroes. 
stat<-matrix(0, nrow=step, ncol=length(event.types))
colnames(stat)<-event.types


#this loops through each event in the sequence

for(j in step){

#for each event, loop through all the males- the support array will take care of males that weren't present that day.
for(i in 1:ncol(census.list[[1]])){


#Remove the "fights" and "nothings" from the census list. This creates attack/receive and win/lose pairs in order of occurence
fight.times<-grep("fight",win.loss.census[[k]][1:(j-1),i])
nothing.times<-grep("nothing",win.loss.census[[k]][1:(j-1),i])

pairs<-win.loss.census[[k]][1:(j-1),i][-c(fight.times, nothing.times)]
names(pairs)<-event.list.dat[[k]][1:(j-1),"Time"][-c(fight.times, nothing.times)]

#Now I can go either way with it. Here I want all the receives that are followed by losses

#need a way to do this that matches interaction partners but also is sensitive to the time-ordering 

ints<-vector()

#in this case I only care once a male has completed an interaction, so "pairs" has to be >=2
if(length(pairs)>=2 && (length(union(grep("receive",pairs),grep("attack",pairs)))==length(union(grep("win",pairs),grep("lose",pairs))))){

ints<-unlist(sapply(1:ncol(census.list[[1]]), function(z){
	
	intset<-pairs[sapply(pairs, function(x) {
		
		strsplit(x,split=".",fixed=T)[[1]][3]==z
		
	})]
	
	# names(intset)<-grep("TRUE",sapply(pairs, function(x) {
		
		# strsplit(x,split=".",fixed=T)[[1]][3]==z
		
	# }))
	
	attacks<-grep("attack",intset)
	attackplus<-intset[attacks+1]
	
	return(attackplus)

}))
}

if(length(ints)>0){
	
#get the time of the most recent end of interaction in seconds
most.rec.sec<-max(as.numeric(names(ints)))

#get the difference with the current time
current.time<-event.list.dat[[k]][(j-1),"Time"]

time.diff<-(current.time-most.rec.sec)

#This will produce "infinities" in the stat array when the current event IS the disengage event. the support array should take care of this since males cannot be attacked before disengaging, but just in case, make sure the timediff is greater than zero


#match with event types- for general interactions, this recency predicts the hazard of incoming attacks

#I use a slightly more slowly decaying function here since the variance in recency can be quite large. Can explore other formulations later. 
#incoming attacks directed at the focal male
stat[j,][endsWith(event.types,paste(".attack.",i, sep=""))]<-1/sqrt(time.diff+1)


}
			
}
	
}


rec.stat[[k]][[1]]<-abind(rec.stat[[k]][[1]], stat)

dimnames(rec.stat[[k]][[1]])[[3]][ncol(rec.stat[[k]][[1]][1,,])]<-"gen.attack.rec"

}

return(rec.stat)

}


##Am I generally more liekly to be attacked if another male attacked me recently (in seconds)?


add.gen.receive.rec<-function(x){

rec.stat<-x

library(abind)

event.list.dat<-split(event.list,event.list$Date)

#loop through each day sequence
for(k in day){

#empty matrix of the right dimensions, event# x event type. In this case, the default is all zeroes. 
stat<-matrix(0, nrow=step, ncol=length(event.types))
colnames(stat)<-event.types


#this loops through each event in the sequence

for(j in step){

#for each event, loop through all the males- the support array will take care of males that weren't present that day.
for(i in 1:ncol(census.list[[1]])){


#Remove the "fights" and "nothings" from the census list. This creates attack/receive and win/lose pairs in order of occurence
fight.times<-grep("fight",win.loss.census[[k]][1:(j-1),i])
nothing.times<-grep("nothing",win.loss.census[[k]][1:(j-1),i])

pairs<-win.loss.census[[k]][1:(j-1),i][-c(fight.times, nothing.times)]
names(pairs)<-event.list.dat[[k]][1:(j-1),"Time"][-c(fight.times, nothing.times)]

#Now I can go either way with it. Here I want all the receives that are followed by losses

#need a way to do this that matches interaction partners but also is sensitive to the time-ordering 

ints<-vector()

#in this case I only care once a male has completed an interaction, so "pairs" has to be >=2 and also an even number, with the same number of starts and ends, in the case of X males 


if(length(pairs)>=2 && (length(union(grep("receive",pairs),grep("attack",pairs)))==length(union(grep("win",pairs),grep("lose",pairs))))){

ints<-unlist(sapply(1:ncol(census.list[[1]]), function(z){
	
	intset<-pairs[sapply(pairs, function(x) {
		
		strsplit(x,split=".",fixed=T)[[1]][3]==z
		
	})]
	
	receives<-grep("receive",intset)
	receiveplus<-intset[receives+1]
	
	return(receiveplus)


}))
}

if(length(ints)>0){
	
#get the time of the most recent end of interaction in seconds
most.rec.sec<-max(as.numeric(names(ints)))

#get the difference with the current time
current.time<-event.list.dat[[k]][(j-1),"Time"]

time.diff<-(current.time-most.rec.sec)

#This will produce "infinities" in the stat array when the current event IS the disengage event. the support array should take care of this since males cannot be attacked before disengaging, but just in case, make sure the timediff is greater than zero


#match with event types- for general interactions, this recency predicts the hazard of incoming attacks

#I use a slightly more slowly decaying function here since the variance in recency can be quite large. Can explore other formulations later. 
#incoming attacks directed at the focal male
stat[j,][endsWith(event.types,paste(".attack.",i, sep=""))]<-1/sqrt(time.diff+1)


}
			
}
	
}


rec.stat[[k]][[1]]<-abind(rec.stat[[k]][[1]], stat)

dimnames(rec.stat[[k]][[1]])[[3]][ncol(rec.stat[[k]][[1]][1,,])]<-"gen.receive.rec"

}

return(rec.stat)

}


##Am I generally more liekly to be attacked if I "won" an interaction recently (in seconds)?


add.gen.win.rec<-function(x){

rec.stat<-x

library(abind)

event.list.dat<-split(event.list,event.list$Date)

#loop through each day sequence
for(k in day){

#empty matrix of the right dimensions, event# x event type. In this case, the default is all zeroes. 
stat<-matrix(0, nrow=step, ncol=length(event.types))
colnames(stat)<-event.types


#this loops through each event in the sequence

for(j in step){

#for each event, loop through all the males- the support array will take care of males that weren't present that day.
for(i in 1:ncol(census.list[[1]])){


#Remove the "fights" and "nothings" from the census list. This creates attack/receive and win/lose pairs in order of occurence
fight.times<-grep("fight",win.loss.census[[k]][1:(j-1),i])
nothing.times<-grep("nothing",win.loss.census[[k]][1:(j-1),i])

pairs<-win.loss.census[[k]][1:(j-1),i][-c(fight.times, nothing.times)]
names(pairs)<-event.list.dat[[k]][1:(j-1),"Time"][-c(fight.times, nothing.times)]

#Now I can go either way with it. Here I want all the receives that are followed by losses

#need a way to do this that matches interaction partners but also is sensitive to the time-ordering 

ints<-vector()

#in this case I only care once a male has completed an interaction, so "pairs" has to be >=2
if(length(pairs)>=2){

ints<-unlist(sapply(1:ncol(census.list[[1]]), function(z){
	
	intset<-pairs[sapply(pairs, function(x) {
		
		strsplit(x,split=".",fixed=T)[[1]][3]==z
		
	})]
	
	wins<-intset[grep("win",intset)]
	
	return(wins)

}))
}

if(length(ints)>0){
	
#get the time of the most recent end of interaction in seconds
most.rec.sec<-max(as.numeric(names(ints)))

#get the difference with the current time
current.time<-event.list.dat[[k]][(j-1),"Time"]

time.diff<-(current.time-most.rec.sec)

#This will produce "infinities" in the stat array when the current event IS the disengage event. the support array should take care of this since males cannot be attacked before disengaging, but just in case, make sure the timediff is greater than zero


#match with event types- for general interactions, this recency predicts the hazard of incoming attacks

#I use a slightly more slowly decaying function here since the variance in recency can be quite large. Can explore other formulations later. 
#incoming attacks directed at the focal male
stat[j,][endsWith(event.types,paste(".attack.",i, sep=""))]<-1/sqrt(time.diff+1)


}
			
}
	
}


rec.stat[[k]][[1]]<-abind(rec.stat[[k]][[1]], stat)

dimnames(rec.stat[[k]][[1]])[[3]][ncol(rec.stat[[k]][[1]][1,,])]<-"gen.win.rec"

}

return(rec.stat)

}


##Am I generally more liekly to be attacked if I "won" an chase recently (in seconds)?


add.gen.win.chase.rec<-function(x){

rec.stat<-x

library(abind)

event.list.dat<-split(event.list,event.list$Date)

#loop through each day sequence
for(k in day){

#empty matrix of the right dimensions, event# x event type. In this case, the default is all zeroes. 
stat<-matrix(0, nrow=step, ncol=length(event.types))
colnames(stat)<-event.types


#this loops through each event in the sequence

for(j in step){

#for each event, loop through all the males- the support array will take care of males that weren't present that day.
for(i in 1:ncol(census.list[[1]])){


#Remove the "fights" and "nothings" from the census list. This creates attack/receive and win/lose pairs in order of occurence
fight.times<-grep("fight",win.loss.census[[k]][1:(j-1),i])
nothing.times<-grep("nothing",win.loss.census[[k]][1:(j-1),i])

pairs<-win.loss.census[[k]][1:(j-1),i][-c(fight.times, nothing.times)]
names(pairs)<-event.list.dat[[k]][1:(j-1),"Time"][-c(fight.times, nothing.times)]

#Now I can go either way with it. Here I want all the receives that are followed by losses

#need a way to do this that matches interaction partners but also is sensitive to the time-ordering 

ints<-vector()

#in this case I only care once a male has completed an interaction, so "pairs" has to be >=2
if(length(pairs)>=2){

ints<-unlist(sapply(1:ncol(census.list[[1]]), function(z){
	
	intset<-pairs[sapply(pairs, function(x) {
		
		strsplit(x,split=".",fixed=T)[[1]][3]==z
		
	})]
	
	wins<-intset[intersect(grep("win",intset),grep("chase",intset))]
	
	return(wins)

}))
}

if(length(ints)>0){
	
#get the time of the most recent end of interaction in seconds
most.rec.sec<-max(as.numeric(names(ints)))

#get the difference with the current time
current.time<-event.list.dat[[k]][(j-1),"Time"]

time.diff<-(current.time-most.rec.sec)

#This will produce "infinities" in the stat array when the current event IS the disengage event. the support array should take care of this since males cannot be attacked before disengaging, but just in case, make sure the timediff is greater than zero


#match with event types- for general interactions, this recency predicts the hazard of incoming attacks

#I use a slightly more slowly decaying function here since the variance in recency can be quite large. Can explore other formulations later. 
#incoming attacks directed at the focal male
stat[j,][endsWith(event.types,paste(".attack.",i, sep=""))]<-1/sqrt(time.diff+1)


}
			
}
	
}


rec.stat[[k]][[1]]<-abind(rec.stat[[k]][[1]], stat)

dimnames(rec.stat[[k]][[1]])[[3]][ncol(rec.stat[[k]][[1]][1,,])]<-"gen.win.chase.rec"

}

return(rec.stat)

}


##Am I generally more liekly to be attacked if I "lost" an interaction recently (in seconds)?

add.gen.lose.rec<-function(x){

rec.stat<-x

library(abind)

event.list.dat<-split(event.list,event.list$Date)

#loop through each day sequence
for(k in day){

#empty matrix of the right dimensions, event# x event type. In this case, the default is all zeroes. 
stat<-matrix(0, nrow=step, ncol=length(event.types))
colnames(stat)<-event.types


#this loops through each event in the sequence

for(j in step){

#for each event, loop through all the males- the support array will take care of males that weren't present that day.
for(i in 1:ncol(census.list[[1]])){


#Remove the "fights" and "nothings" from the census list. This creates attack/receive and win/lose pairs in order of occurence
fight.times<-grep("fight",win.loss.census[[k]][1:(j-1),i])
nothing.times<-grep("nothing",win.loss.census[[k]][1:(j-1),i])

pairs<-win.loss.census[[k]][1:(j-1),i][-c(fight.times, nothing.times)]
names(pairs)<-event.list.dat[[k]][1:(j-1),"Time"][-c(fight.times, nothing.times)]

#Now I can go either way with it. Here I want all the receives that are followed by losses

#need a way to do this that matches interaction partners but also is sensitive to the time-ordering 

ints<-vector()

#in this case I only care once a male has completed an interaction, so "pairs" has to be >=2
if(length(pairs)>=2){

ints<-unlist(sapply(1:ncol(census.list[[1]]), function(z){
	
	intset<-pairs[sapply(pairs, function(x) {
		
		strsplit(x,split=".",fixed=T)[[1]][3]==z
		
	})]
	
	losses<-intset[grep("lose",intset)]
	
	return(losses)

}))
}

if(length(ints)>0){
	
#get the time of the most recent end of interaction in seconds
most.rec.sec<-max(as.numeric(names(ints)))

#get the difference with the current time
current.time<-event.list.dat[[k]][(j-1),"Time"]

time.diff<-(current.time-most.rec.sec)

#This will produce "infinities" in the stat array when the current event IS the disengage event. the support array should take care of this since males cannot be attacked before disengaging, but just in case, make sure the timediff is greater than zero


#match with event types- for general interactions, this recency predicts the hazard of incoming attacks

#I use a slightly more slowly decaying function here since the variance in recency can be quite large. Can explore other formulations later. 
#incoming attacks directed at the focal male
stat[j,][endsWith(event.types,paste(".attack.",i, sep=""))]<-1/sqrt(time.diff+1)


}
			
}
	
}


rec.stat[[k]][[1]]<-abind(rec.stat[[k]][[1]], stat)

dimnames(rec.stat[[k]][[1]])[[3]][ncol(rec.stat[[k]][[1]][1,,])]<-"gen.lose.rec"

}

return(rec.stat)

}


##Am I generally more liekly to be attacked if I "lost" an chase recently (in seconds)?


add.gen.lose.chase.rec<-function(x){

rec.stat<-x

library(abind)

event.list.dat<-split(event.list,event.list$Date)

#loop through each day sequence
for(k in day){

#empty matrix of the right dimensions, event# x event type. In this case, the default is all zeroes. 
stat<-matrix(0, nrow=step, ncol=length(event.types))
colnames(stat)<-event.types


#this loops through each event in the sequence

for(j in step){

#for each event, loop through all the males- the support array will take care of males that weren't present that day.
for(i in 1:ncol(census.list[[1]])){


#Remove the "fights" and "nothings" from the census list. This creates attack/receive and win/lose pairs in order of occurence
fight.times<-grep("fight",win.loss.census[[k]][1:(j-1),i])
nothing.times<-grep("nothing",win.loss.census[[k]][1:(j-1),i])

pairs<-win.loss.census[[k]][1:(j-1),i][-c(fight.times, nothing.times)]
names(pairs)<-event.list.dat[[k]][1:(j-1),"Time"][-c(fight.times, nothing.times)]

#Now I can go either way with it. Here I want all the receives that are followed by losses

#need a way to do this that matches interaction partners but also is sensitive to the time-ordering 

ints<-vector()

#in this case I only care once a male has completed an interaction, so "pairs" has to be >=2
if(length(pairs)>=2){

ints<-unlist(sapply(1:ncol(census.list[[1]]), function(z){
	
	intset<-pairs[sapply(pairs, function(x) {
		
		strsplit(x,split=".",fixed=T)[[1]][3]==z
		
	})]
	
	losses<-intset[intersect(grep("lose",intset),grep("chase",intset))]
	
	return(losses)

}))
}

if(length(ints)>0){
	
#get the time of the most recent end of interaction in seconds
most.rec.sec<-max(as.numeric(names(ints)))

#get the difference with the current time
current.time<-event.list.dat[[k]][(j-1),"Time"]

time.diff<-(current.time-most.rec.sec)

#This will produce "infinities" in the stat array when the current event IS the disengage event. the support array should take care of this since males cannot be attacked before disengaging, but just in case, make sure the timediff is greater than zero


#match with event types- for general interactions, this recency predicts the hazard of incoming attacks

#I use a slightly more slowly decaying function here since the variance in recency can be quite large. Can explore other formulations later. 
#incoming attacks directed at the focal male
stat[j,][endsWith(event.types,paste(".attack.",i, sep=""))]<-1/sqrt(time.diff+1)


}
			
}
	
}


rec.stat[[k]][[1]]<-abind(rec.stat[[k]][[1]], stat)

dimnames(rec.stat[[k]][[1]])[[3]][ncol(rec.stat[[k]][[1]][1,,])]<-"gen.lose.chase.rec"

}

return(rec.stat)

}


##Am I generally more likely to be attacked if I attack+won recently (in seconds)?


add.gen.attack.win.rec<-function(x){

rec.stat<-x

library(abind)

event.list.dat<-split(event.list,event.list$Date)

#loop through each day sequence
for(k in day){

#empty matrix of the right dimensions, event# x event type. In this case, the default is all zeroes. 
stat<-matrix(0, nrow=step, ncol=length(event.types))
colnames(stat)<-event.types


#this loops through each event in the sequence

for(j in step){

#for each event, loop through all the males- the support array will take care of males that weren't present that day.
for(i in 1:ncol(census.list[[1]])){


#Remove the "fights" and "nothings" from the census list. This creates attack/receive and win/lose pairs in order of occurence
fight.times<-grep("fight",win.loss.census[[k]][1:(j-1),i])
nothing.times<-grep("nothing",win.loss.census[[k]][1:(j-1),i])

pairs<-win.loss.census[[k]][1:(j-1),i][-c(fight.times, nothing.times)]
names(pairs)<-event.list.dat[[k]][1:(j-1),"Time"][-c(fight.times, nothing.times)]

#Now I can go either way with it. Here I want all the receives that are followed by losses

#need a way to do this that matches interaction partners but also is sensitive to the time-ordering 

ints<-vector()

#in this case I only care once a male has completed an interaction, so "pairs" has to be >=2
if(length(pairs)>=2){

ints<-unlist(sapply(1:ncol(census.list[[1]]), function(z){
	
	intset<-pairs[sapply(pairs, function(x) {
		
		strsplit(x,split=".",fixed=T)[[1]][3]==z
		
	})]
	
	
	attacks<-grep("attack",intset)
	attackpluswin<-intset[attacks+1][grep("win",intset[attacks+1])]
	
	return(attackpluswin)

}))
}

if(length(ints)>0){
	
#get the time of the most recent end of interaction in seconds
most.rec.sec<-max(as.numeric(names(ints)))

#get the difference with the current time
current.time<-event.list.dat[[k]][(j-1),"Time"]

time.diff<-(current.time-most.rec.sec)

#This will produce "infinities" in the stat array when the current event IS the disengage event. the support array should take care of this since males cannot be attacked before disengaging, but just in case, make sure the timediff is greater than zero


#match with event types- for general interactions, this recency predicts the hazard of incoming attacks

#I use a slightly more slowly decaying function here since the variance in recency can be quite large. Can explore other formulations later. 
#incoming attacks directed at the focal male
stat[j,][endsWith(event.types,paste(".attack.",i, sep=""))]<-1/sqrt(time.diff+1)


}
			
}
	
}


rec.stat[[k]][[1]]<-abind(rec.stat[[k]][[1]], stat)

dimnames(rec.stat[[k]][[1]])[[3]][ncol(rec.stat[[k]][[1]][1,,])]<-"gen.attack.win.rec"

}

return(rec.stat)

}


##Am I generally more likely to be attacked if I attack+lost recently (in seconds)?


add.gen.attack.lose.rec<-function(x){

rec.stat<-x

library(abind)

event.list.dat<-split(event.list,event.list$Date)

#loop through each day sequence
for(k in day){

#empty matrix of the right dimensions, event# x event type. In this case, the default is all zeroes. 
stat<-matrix(0, nrow=step, ncol=length(event.types))
colnames(stat)<-event.types


#this loops through each event in the sequence

for(j in step){

#for each event, loop through all the males- the support array will take care of males that weren't present that day.
for(i in 1:ncol(census.list[[1]])){


#Remove the "fights" and "nothings" from the census list. This creates attack/receive and win/lose pairs in order of occurence
fight.times<-grep("fight",win.loss.census[[k]][1:(j-1),i])
nothing.times<-grep("nothing",win.loss.census[[k]][1:(j-1),i])

pairs<-win.loss.census[[k]][1:(j-1),i][-c(fight.times, nothing.times)]
names(pairs)<-event.list.dat[[k]][1:(j-1),"Time"][-c(fight.times, nothing.times)]

#Now I can go either way with it. Here I want all the receives that are followed by losses

#need a way to do this that matches interaction partners but also is sensitive to the time-ordering 

ints<-vector()

#in this case I only care once a male has completed an interaction, so "pairs" has to be >=2
if(length(pairs)>=2){

ints<-unlist(sapply(1:ncol(census.list[[1]]), function(z){
	
	intset<-pairs[sapply(pairs, function(x) {
		
		strsplit(x,split=".",fixed=T)[[1]][3]==z
		
	})]
	
	
	attacks<-grep("attack",intset)
	attackpluswin<-intset[attacks+1][grep("lose",intset[attacks+1])]
	
	return(attackpluswin)

}))
}

if(length(ints)>0){
	
#get the time of the most recent end of interaction in seconds
most.rec.sec<-max(as.numeric(names(ints)))

#get the difference with the current time
current.time<-event.list.dat[[k]][(j-1),"Time"]

time.diff<-(current.time-most.rec.sec)

#This will produce "infinities" in the stat array when the current event IS the disengage event. the support array should take care of this since males cannot be attacked before disengaging, but just in case, make sure the timediff is greater than zero


#match with event types- for general interactions, this recency predicts the hazard of incoming attacks

#I use a slightly more slowly decaying function here since the variance in recency can be quite large. Can explore other formulations later. 
#incoming attacks directed at the focal male
stat[j,][endsWith(event.types,paste(".attack.",i, sep=""))]<-1/sqrt(time.diff+1)


}
			
}
	
}


rec.stat[[k]][[1]]<-abind(rec.stat[[k]][[1]], stat)

dimnames(rec.stat[[k]][[1]])[[3]][ncol(rec.stat[[k]][[1]][1,,])]<-"gen.attack.lose.rec"

}

return(rec.stat)

}


##Am I generally more likely to be attacked if I receive+won recently (in seconds)?


add.gen.receive.win.rec<-function(x){

rec.stat<-x

library(abind)

event.list.dat<-split(event.list,event.list$Date)

#loop through each day sequence
for(k in day){

#empty matrix of the right dimensions, event# x event type. In this case, the default is all zeroes. 
stat<-matrix(0, nrow=step, ncol=length(event.types))
colnames(stat)<-event.types


#this loops through each event in the sequence

for(j in step){

#for each event, loop through all the males- the support array will take care of males that weren't present that day.
for(i in 1:ncol(census.list[[1]])){


#Remove the "fights" and "nothings" from the census list. This creates attack/receive and win/lose pairs in order of occurence
fight.times<-grep("fight",win.loss.census[[k]][1:(j-1),i])
nothing.times<-grep("nothing",win.loss.census[[k]][1:(j-1),i])

pairs<-win.loss.census[[k]][1:(j-1),i][-c(fight.times, nothing.times)]
names(pairs)<-event.list.dat[[k]][1:(j-1),"Time"][-c(fight.times, nothing.times)]

#Now I can go either way with it. Here I want all the receives that are followed by losses

#need a way to do this that matches interaction partners but also is sensitive to the time-ordering 

ints<-vector()

#in this case I only care once a male has completed an interaction, so "pairs" has to be >=2
if(length(pairs)>=2){

ints<-unlist(sapply(1:ncol(census.list[[1]]), function(z){
	
	intset<-pairs[sapply(pairs, function(x) {
		
		strsplit(x,split=".",fixed=T)[[1]][3]==z
		
	})]
	
	
	receives<-grep("receive",intset)
	receivepluswin<-intset[receives+1][grep("win",intset[receives+1])]
	
	return(receivepluswin)

}))
}

if(length(ints)>0){
	
#get the time of the most recent end of interaction in seconds
most.rec.sec<-max(as.numeric(names(ints)))

#get the difference with the current time
current.time<-event.list.dat[[k]][(j-1),"Time"]

time.diff<-(current.time-most.rec.sec)

#This will produce "infinities" in the stat array when the current event IS the disengage event. the support array should take care of this since males cannot be attacked before disengaging, but just in case, make sure the timediff is greater than zero


#match with event types- for general interactions, this recency predicts the hazard of incoming attacks

#I use a slightly more slowly decaying function here since the variance in recency can be quite large. Can explore other formulations later. 
#incoming attacks directed at the focal male
stat[j,][endsWith(event.types,paste(".attack.",i, sep=""))]<-1/sqrt(time.diff+1)


}
			
}
	
}


rec.stat[[k]][[1]]<-abind(rec.stat[[k]][[1]], stat)

dimnames(rec.stat[[k]][[1]])[[3]][ncol(rec.stat[[k]][[1]][1,,])]<-"gen.receive.win.rec"

}

return(rec.stat)

}


##Am I generally more likely to be attacked if I receive+lost recently (in seconds)?


add.gen.receive.lose.rec<-function(x){

rec.stat<-x

library(abind)

event.list.dat<-split(event.list,event.list$Date)

#loop through each day sequence
for(k in day){

#empty matrix of the right dimensions, event# x event type. In this case, the default is all zeroes. 
stat<-matrix(0, nrow=step, ncol=length(event.types))
colnames(stat)<-event.types


#this loops through each event in the sequence

for(j in step){

#for each event, loop through all the males- the support array will take care of males that weren't present that day.
for(i in 1:ncol(census.list[[1]])){


#Remove the "fights" and "nothings" from the census list. This creates attack/receive and win/lose pairs in order of occurence
fight.times<-grep("fight",win.loss.census[[k]][1:(j-1),i])
nothing.times<-grep("nothing",win.loss.census[[k]][1:(j-1),i])

pairs<-win.loss.census[[k]][1:(j-1),i][-c(fight.times, nothing.times)]
names(pairs)<-event.list.dat[[k]][1:(j-1),"Time"][-c(fight.times, nothing.times)]

#Now I can go either way with it. Here I want all the receives that are followed by losses

#need a way to do this that matches interaction partners but also is sensitive to the time-ordering 

ints<-vector()

#in this case I only care once a male has completed an interaction, so "pairs" has to be >=2
if(length(pairs)>=2){

ints<-unlist(sapply(1:ncol(census.list[[1]]), function(z){
	
	intset<-pairs[sapply(pairs, function(x) {
		
		strsplit(x,split=".",fixed=T)[[1]][3]==z
		
	})]
	
	
	receives<-grep("receive",intset)
	receivepluslose<-intset[receives+1][grep("lose",intset[receives+1])]
	
	return(receivepluslose)

}))
}

if(length(ints)>0){
	
#get the time of the most recent end of interaction in seconds
most.rec.sec<-max(as.numeric(names(ints)))

#get the difference with the current time
current.time<-event.list.dat[[k]][(j-1),"Time"]

time.diff<-(current.time-most.rec.sec)

#This will produce "infinities" in the stat array when the current event IS the disengage event. the support array should take care of this since males cannot be attacked before disengaging, but just in case, make sure the timediff is greater than zero


#match with event types- for general interactions, this recency predicts the hazard of incoming attacks

#I use a slightly more slowly decaying function here since the variance in recency can be quite large. Can explore other formulations later. 
#incoming attacks directed at the focal male
stat[j,][endsWith(event.types,paste(".attack.",i, sep=""))]<-1/sqrt(time.diff+1)


}
			
}
	
}


rec.stat[[k]][[1]]<-abind(rec.stat[[k]][[1]], stat)

dimnames(rec.stat[[k]][[1]])[[3]][ncol(rec.stat[[k]][[1]][1,,])]<-"gen.receive.lose.rec"

}

return(rec.stat)

}


######## Effect of time spent on lek/fighting so far Today #########

#I feel like these rely heavily on the fact that the same male has been there all day, and the interpretation for X males is difficult. Therefore they are not included as FOCAL males in these stats. Fights other males have with them still count. 


##Am I generally more likely to be attacked if I have spent more time on the lek so far today?

add.total.time<-function(x){

rec.stat<-x

library(abind)

event.list.dat<-split(event.list,event.list$Date)

#loop through each day sequence
for(k in day){

#empty matrix of the right dimensions, event# x event type. In this case, the default is all zeroes. 
stat<-matrix(0, nrow=step, ncol=length(event.types))
colnames(stat)<-event.types


#this loops through each event in the sequence

for(j in step){

arrsub<-subset(pres.exact.dat[[k]],Time<event.list.dat[[k]][(j-1),"Time"])
arrsub<-arrsub[order(as.numeric(arrsub$Time)),]

#for each event, loop through all the males- the support array will take care of males that weren't present that day.
for(i in 1:(ncol(census.list[[1]])-1)){
	

arrind<-arrsub[as.character(arrsub$Bird_ID)==i,]

if(nrow(arrind)>0){

if((nrow(arrind) %% 2)==0){
	
	tot.time<-sum(sapply(1:(nrow(arrind)/2), function(x) return(arrind[(2*x),2]-arrind[(2*x-1),2])))
	
}

if(nrow(arrind)==1){
	
	tot.time<-(event.list.dat[[k]][(j-1),"Time"]-arrind[nrow(arrind),2])
}

if((nrow(arrind) %% 2)!=0 && nrow(arrind)!=1){
	
	tot.time<-sum(sapply(1:floor(nrow(arrind)/2), function(x) return(arrind[(2*x),2]-arrind[(2*x-1),2])))+(event.list.dat[[k]][(j-1),"Time"]-arrind[nrow(arrind),2])
	
}


#account for timesteps that occur before the male's arrival (just to keep it clean- the support array will take care of times when the focal male is not present)
if(length(tot.time)>0 && tot.time>0){

#match with event types- for general interactions, this recency predicts the hazard of incoming attacks

#I use a slightly more slowly decaying function here since the variance in recency can be quite large. Can explore other formulations later. 
#incoming attacks directed at the focal male
stat[j,][endsWith(event.types,paste(".attack.",i, sep=""))]<-tot.time

#exclude attack events involving the X males 
stat[j,][endsWith(event.types,paste(".attack.",ncol(census.list[[1]]), sep="")) | startsWith(event.types,paste(ncol(census.list[[1]]),".attack.", sep=""))]<-0

}
}			
}
	
}


rec.stat[[k]][[1]]<-abind(rec.stat[[k]][[1]], stat)

dimnames(rec.stat[[k]][[1]])[[3]][ncol(rec.stat[[k]][[1]][1,,])]<-"total.time"

}

return(rec.stat)

}


##Am I generally more likely to be attacked if I have spent log(1 + more time) on the lek so far today?


add.log.total.time<-function(x){

rec.stat<-x

library(abind)

event.list.dat<-split(event.list,event.list$Date)

#loop through each day sequence
for(k in day){

#empty matrix of the right dimensions, event# x event type. In this case, the default is all zeroes. 
stat<-matrix(0, nrow=step, ncol=length(event.types))
colnames(stat)<-event.types


#this loops through each event in the sequence

for(j in step){

arrsub<-subset(pres.exact.dat[[k]],Time<event.list.dat[[k]][(j-1),"Time"])
arrsub<-arrsub[order(as.numeric(arrsub$Time)),]

#for each event, loop through all the males- the support array will take care of males that weren't present that day.
for(i in 1:(ncol(census.list[[1]])-1)){
	

arrind<-arrsub[as.character(arrsub$Bird_ID)==i,]

if(nrow(arrind)>0){

if((nrow(arrind) %% 2)==0){
	
	tot.time<-sum(sapply(1:(nrow(arrind)/2), function(x) return(arrind[(2*x),2]-arrind[(2*x-1),2])))
	
}

if(nrow(arrind)==1){
	
	tot.time<-(event.list.dat[[k]][(j-1),"Time"]-arrind[nrow(arrind),2])
}

if((nrow(arrind) %% 2)!=0 && nrow(arrind)!=1){
	
	tot.time<-sum(sapply(1:floor(nrow(arrind)/2), function(x) return(arrind[(2*x),2]-arrind[(2*x-1),2])))+(event.list.dat[[k]][(j-1),"Time"]-arrind[nrow(arrind),2])
	
}


#account for timesteps that occur before the male's arrival (just to keep it clean- the support array will take care of times when the focal male is not present)
if(length(tot.time)>0 && tot.time>0){

#match with event types- for general interactions, this recency predicts the hazard of incoming attacks

#I use a slightly more slowly decaying function here since the variance in recency can be quite large. Can explore other formulations later. 
#incoming attacks directed at the focal male
stat[j,][endsWith(event.types,paste(".attack.",i, sep=""))]<-tot.time

#exclude attack events involving the X males 
stat[j,][endsWith(event.types,paste(".attack.",ncol(census.list[[1]]), sep="")) | startsWith(event.types,paste(ncol(census.list[[1]]),".attack.", sep=""))]<-0

}
}			
}
	
}

stat<-log(1+stat)

rec.stat[[k]][[1]]<-abind(rec.stat[[k]][[1]], stat)

dimnames(rec.stat[[k]][[1]])[[3]][ncol(rec.stat[[k]][[1]][1,,])]<-"log.total.time"

}

return(rec.stat)

}


##Am I generally more likely to be attacked if I have spent more time fighting so far today, compared to others?

add.rank.fighting.time<-function(x){

rec.stat<-x

library(abind)

event.list.dat<-split(event.list,event.list$Date)

#loop through each day sequence
for(k in day){

#empty matrix of the right dimensions, event# x event type. In this case, the default is all zeroes. 
stat<-matrix(0, nrow=step, ncol=length(event.types))
colnames(stat)<-event.types


#this loops through each event in the sequence

for(j in step){


#for each event, loop through all the males- the support array will take care of males that weren't present that day.
for(i in 1:(ncol(census.list[[1]])-1)){
		
#get pairs of starts and ends of interactions that preserve the row number of the original event sequence
	
wins<-grep("win",win.loss.census[[k]][1:(j-1),i])
losses<-grep("lose",win.loss.census[[k]][1:(j-1),i])
end.times<-union(wins,losses)
end.times<-end.times[order(end.times)]

attacks<-grep("attack",win.loss.census[[k]][1:(j-1),i])
receives<-grep("received",win.loss.census[[k]][1:(j-1),i])
start.times<-union(attacks,receives)
start.times<-start.times[order(start.times)]

pairs<-union(start.times,end.times)

#make sure the male has had any interactions so far and that any of these have already been completed. This will produce zero hazards for males currently engaged in interactions, which is fine since the support array makes them ineligible anyway
if(length(pairs)>=2){

pairs<-pairs[order(pairs)]

if(length(pairs) %% 2 != 0){pairs<-pairs[-length(pairs)]}


pairs.times<-event.list.dat[[k]][pairs,"Time"]

#sum up the time intervals spent fighting
intervals<-matrix(pairs.times,nrow=2)

fight.tot<-sum(intervals[2,]-intervals[1,])

#match with event types- for general interactions, this recency predicts the hazard of incoming attacks

#incoming attacks directed at the focal male
stat[j,][endsWith(event.types,paste(".attack.",i, sep=""))]<-fight.tot

#exclude attack events involving the X males 
stat[j,][endsWith(event.types,paste(".attack.",ncol(census.list[[1]]), sep="")) | startsWith(event.types,paste(ncol(census.list[[1]]),".attack.", sep=""))]<-0

}			
}


#scale by ranking due to the extreme skew in the data even in the upper half

if(max(stat[j,][grep("attack",event.types)])>0){
	
assign<-1:length(unique(stat[j,][grep("attack",event.types)]))

names(assign)<-unique(stat[j,][grep("attack",event.types)])[order(unique(stat[j,][grep("attack",event.types)]),decreasing=T)]

scaleassign<-assign[as.character(stat[j,][grep("attack",event.types)])]
scaleassign<-1/scaleassign

stat[j,][grep("attack",event.types)]<-scaleassign

}


	
}

rec.stat[[k]][[1]]<-abind(rec.stat[[k]][[1]], stat)

dimnames(rec.stat[[k]][[1]])[[3]][ncol(rec.stat[[k]][[1]][1,,])]<-"rank.fighting.time"

}

return(rec.stat)

}


##Am I generally more likely to be attacked if I have spent a larger percentage of my time on the lek fighting so far today? 


add.rank.percent.fighting.time<-function(x){

rec.stat<-x

library(abind)

event.list.dat<-split(event.list,event.list$Date)

#loop through each day sequence
for(k in day){

#empty matrix of the right dimensions, event# x event type. In this case, the default is all zeroes. 
stat<-matrix(0, nrow=step, ncol=length(event.types))
colnames(stat)<-event.types


#this loops through each event in the sequence

for(j in step){

arrsub<-subset(pres.exact.dat[[k]],Time<event.list.dat[[k]][(j-1),"Time"])
arrsub<-arrsub[order(as.numeric(arrsub$Time)),]

#for each event, loop through all the males- the support array will take care of males that weren't present that day.
for(i in 1:(ncol(census.list[[1]])-1)){
		
#get pairs of starts and ends of interactions that preserve the row number of the original event sequence
	
wins<-grep("win",win.loss.census[[k]][1:(j-1),i])
losses<-grep("lose",win.loss.census[[k]][1:(j-1),i])
end.times<-union(wins,losses)
end.times<-end.times[order(end.times)]

attacks<-grep("attack",win.loss.census[[k]][1:(j-1),i])
receives<-grep("received",win.loss.census[[k]][1:(j-1),i])
start.times<-union(attacks,receives)
start.times<-start.times[order(start.times)]

pairs<-union(start.times,end.times)

#make sure the male has had any interactions so far and that any of these have already been completed. This will produce zero hazards for males currently engaged in interactions, which is fine since the support array makes them ineligible anyway
if(length(pairs)>=2){

pairs<-pairs[order(pairs)]

if(length(pairs) %% 2 != 0){pairs<-pairs[-length(pairs)]}


pairs.times<-event.list.dat[[k]][pairs,"Time"]

#sum up the time intervals spent fighting
intervals<-matrix(pairs.times,nrow=2)

fight.tot<-sum(intervals[2,]-intervals[1,])


##get the total time the focal male was present on the lek so far today

arrind<-arrsub[as.character(arrsub$Bird_ID)==i,]

if(nrow(arrind)>0){

if((nrow(arrind) %% 2)==0){
	
	tot.time<-sum(sapply(1:(nrow(arrind)/2), function(x) return(arrind[(2*x),2]-arrind[(2*x-1),2])))
	
}

if(nrow(arrind)==1){
	
	tot.time<-(event.list.dat[[k]][(j-1),"Time"]-arrind[nrow(arrind),2])
}

if((nrow(arrind) %% 2)!=0 && nrow(arrind)!=1){
	
	tot.time<-sum(sapply(1:floor(nrow(arrind)/2), function(x) return(arrind[(2*x),2]-arrind[(2*x-1),2])))+(event.list.dat[[k]][(j-1),"Time"]-arrind[nrow(arrind),2])
	
}


if(length(tot.time)>0 && tot.time>0){
perc.fight<-100*(fight.tot/tot.time)
#match with event types- for general interactions, this recency predicts the hazard of incoming attacks

#incoming attacks directed at the focal male
stat[j,][endsWith(event.types,paste(".attack.",i, sep=""))]<-perc.fight

#exclude attack events involving the X males 
stat[j,][endsWith(event.types,paste(".attack.",ncol(census.list[[1]]), sep="")) | startsWith(event.types,paste(ncol(census.list[[1]]),".attack.", sep=""))]<-0

}
}

}			
}

#scale by ranking due to the extreme skew in the data even in the upper half

if(max(stat[j,][grep("attack",event.types)])>0){
	
assign<-1:length(unique(stat[j,][grep("attack",event.types)]))

names(assign)<-unique(stat[j,][grep("attack",event.types)])[order(unique(stat[j,][grep("attack",event.types)]),decreasing=T)]

scaleassign<-assign[as.character(stat[j,][grep("attack",event.types)])]
scaleassign<-1/scaleassign

stat[j,][grep("attack",event.types)]<-scaleassign

}

	
}

rec.stat[[k]][[1]]<-abind(rec.stat[[k]][[1]], stat)

dimnames(rec.stat[[k]][[1]])[[3]][ncol(rec.stat[[k]][[1]][1,,])]<-"rank.percent.fighting.time"

}

return(rec.stat)

}


###### Cumulative history of pairwise interaction for the day #

##test log NUMBER of attacks, receives, wins, losses


##Are you more or less likely to attack me if we interacted log(1+MORE TIMES) previously today?


add.log.count.pairwise.day<-function(x){
	
rec.stat<-x

library(abind)

#loop through each day sequence
for(k in day){

#empty matrix of the right dimensions, event# x event type. In this case, the default is all zeroes. 
stat<-matrix(0, nrow=step, ncol=length(event.types))
colnames(stat)<-event.types


#this loops through each event in the sequence

for(j in step){

#for each event, loop through all the males to get the previous attack count that will predict the next interaction- the support array will take care of males that weren't present that day.
for(i in 1:ncol(census.list[[1]])){
	
#list of all the attacks perpetrated by the focal male
attacks<-grep("attack",census.list[[k]][1:(j-1),i])

#list of all the attacks directed at the focal male
receives<-grep("received",census.list[[k]][1:(j-1),i])

#if the male has not had any interactions yet, the hazard is equal across event types involving them, so set all to zero 
if(length(attacks)>0){

#totals for each type- these will be the values for the stat
attack.tots<-table(census.list[[k]][1:(j-1),i][attacks])

#get just the interacting male id numbers
rankids<-sapply(names(attack.tots), function(x) strsplit(x, split=".", fixed=T)[[1]][3])

#match with event types- this stat predicts INCOMING attacks based on total previous OUTGOING + INCOMING attacks
#loop thru each element of rankids

for(l in 1:length(rankids)){

#incoming attacks directed at the focal male, the hazrd is proportional to the number of previous outgoing attacks
stat[j,][paste(rankids[l],".attack.", i, sep="")]<-attack.tots[l]
	}
}

#add receives if there are any
if(length(receives)>0){

#totals for each type- these will be the values for the stat
receive.tots<-table(census.list[[k]][1:(j-1),i][receives])

#get just the interacting male id numbers
rankids<-sapply(names(receive.tots), function(x) strsplit(x, split=".", fixed=T)[[1]][3])

#match with event types
#loop thru each element of rankids

for(l in 1:length(rankids)){

#incoming attacks directed at the focal male, the hazrd is proportional to the number of previous incoming attacks
stat[j,][paste(rankids[l],".attack.", i, sep="")]<-(stat[j,][paste(rankids[l],".attack.", i, sep="")]+receive.tots[l])
	}
		
}
	
}

#exclude attack events involving the X males 
stat[j,][endsWith(event.types,paste(".attack.",ncol(census.list[[1]]), sep="")) | startsWith(event.types,paste(ncol(census.list[[1]]),".attack.", sep=""))]<-0

}

stat<-log(1+stat)

rec.stat[[k]][[1]]<-abind(rec.stat[[k]][[1]], stat)

dimnames(rec.stat[[k]][[1]])[[3]][ncol(rec.stat[[k]][[1]][1,,])]<-"log.count.pairwise.day"

}

return(rec.stat)

	
}

remove<-function(x1,list){
	lapply(x1, function(x){

day<-x

day[[1]]<-day[[1]][,,-which(dimnames(day[[1]])[[3]] %in% list)]

global<-list()

global[[1]]<-day[[1]]

names(global)<-"global"

return(global)
	
}
)
}


##Are you more or less likely to attack me if I attacked you MORE TIMES in the past day?

add.attack.log.count.pairwise.day<-function(x){
	
rec.stat<-x

library(abind)

#loop through each day sequence
for(k in day){

#empty matrix of the right dimensions, event# x event type. In this case, the default is all zeroes. 
stat<-matrix(0, nrow=step, ncol=length(event.types))
colnames(stat)<-event.types


#this loops through each event in the sequence

for(j in step){

#for each event, loop through all the males to get the previous attack count that will predict the next interaction- the support array will take care of males that weren't present that day.
for(i in 1:ncol(census.list[[1]])){
	
#list of all the attacks perpetrated by the focal male
attacks<-grep("attack",census.list[[k]][1:(j-1),i])

#if the male has not had any interactions yet, the hazard is equal across event types involving them, so set all to zero 
if(length(attacks)>0){

#totals for each type- these will be the values for the stat
attack.tots<-table(census.list[[k]][1:(j-1),i][attacks])

#get just the interacting male id numbers
rankids<-sapply(names(attack.tots), function(x) strsplit(x, split=".", fixed=T)[[1]][3])

#match with event types- this stat predicts INCOMING attacks based on total previous OUTGOING attacks
#loop thru each element of rankids

for(l in 1:length(rankids)){

#incoming attacks directed at the focal male, the hazrd is proportional to the number of previous outgoing attacks
stat[j,][paste(rankids[l],".attack.", i, sep="")]<-attack.tots[l]
	}

		
}
	
}

#exclude attack events involving the X males 
stat[j,][endsWith(event.types,paste(".attack.",ncol(census.list[[1]]), sep="")) | startsWith(event.types,paste(ncol(census.list[[1]]),".attack.", sep=""))]<-0


}

stat<-log(1+stat)

rec.stat[[k]][[1]]<-abind(rec.stat[[k]][[1]], stat)

dimnames(rec.stat[[k]][[1]])[[3]][ncol(rec.stat[[k]][[1]][1,,])]<-"attack.log.count.pairwise.day"

}

return(rec.stat)

	
}


## Are you more or less likely to attack me if YOU attacked me MORE TIMES in the past day?

add.receive.log.count.pairwise.day<-function(x){
	
	rec.stat<-x

library(abind)

#loop through each day sequence
for(k in day){

#empty matrix of the right dimensions, event# x event type. In this case, the default is all zeroes. 
stat<-matrix(0, nrow=step, ncol=length(event.types))
colnames(stat)<-event.types


#this loops through each event in the sequence

for(j in step){

#for each event, loop through all the males to get the previous attack count that will predict the next interaction- the support array will take care of males that weren't present that day.
for(i in 1:ncol(census.list[[1]])){
	
#list of all the attacks directed at the focal male
receives<-grep("received",census.list[[k]][1:(j-1),i])

#if the male has not had any interactions yet, the hazard is equal across event types involving them, so set all to zero 
if(length(receives)>0){

#totals for each type- these will be the values for the stat
receive.tots<-table(census.list[[k]][1:(j-1),i][receives])

#get just the interacting male id numbers
rankids<-sapply(names(receive.tots), function(x) strsplit(x, split=".", fixed=T)[[1]][3])

#match with event types- this stat predicts INCOMING attacks based on total previous INCOMING attacks
#loop thru each element of rankids

for(l in 1:length(rankids)){

#incoming attacks directed at the focal male, the hazrd is proportional to the number of previous outgoing attacks
stat[j,][paste(rankids[l],".attack.", i, sep="")]<-receive.tots[l]
	}

		
}
	
}

#exclude attack events involving the X males 
stat[j,][endsWith(event.types,paste(".attack.",ncol(census.list[[1]]), sep="")) | startsWith(event.types,paste(ncol(census.list[[1]]),".attack.", sep=""))]<-0


}

stat<-log(1+stat)

rec.stat[[k]][[1]]<-abind(rec.stat[[k]][[1]], stat)

dimnames(rec.stat[[k]][[1]])[[3]][ncol(rec.stat[[k]][[1]][1,,])]<-"receive.log.count.pairwise.day"

}

return(rec.stat)

	
}


## Are you more or less likely to attack me if I WON against you MORE TIMES in the past day?

add.win.log.count.pairwise.day<-function(x){
	
rec.stat<-x

library(abind)

#loop through each day sequence
for(k in day){

#empty matrix of the right dimensions, event# x event type. In this case, the default is all zeroes. 
stat<-matrix(0, nrow=step, ncol=length(event.types))
colnames(stat)<-event.types


#this loops through each event in the sequence

for(j in step){

#for each event, loop through all the males to get the previous attack count that will predict the next interaction- the support array will take care of males that weren't present that day.
for(i in 1:ncol(census.list[[1]])){
	
#list of all the wins by the focal male
wins<-win.loss.census[[k]][1:(j-1),i][grep("win",win.loss.census[[k]][1:(j-1),i])]

#strip the "chase" from the end of chase wins so I can include them
wins<-sapply(wins, function(x) if(grepl("chas",x)==TRUE){
temp<-strsplit(x, split=".", fixed=T)[[1]]	
new<-paste(temp[1],temp[2],temp[3], sep=".")
return(new)
}
else return(x)
)

#if the male has not had any interactions yet, the hazard is equal across event types involving them, so set all to zero 
if(length(wins)>0){

#totals for each type- these will be the values for the stat
win.tots<-table(wins)

#get just the interacting male id numbers
rankids<-sapply(names(win.tots), function(x) strsplit(x, split=".", fixed=T)[[1]][3])

#match with event types- this stat predicts INCOMING attacks based on total previous WINS by the focal male
#loop thru each element of rankids

for(l in 1:length(rankids)){

#incoming attacks directed at the focal male, the hazrd is proportional to the number of previous outgoing attacks
stat[j,][paste(rankids[l],".attack.", i, sep="")]<-win.tots[l]
	}


		
}
	
}

#exclude attack events involving the X males 
stat[j,][endsWith(event.types,paste(".attack.",ncol(census.list[[1]]), sep="")) | startsWith(event.types,paste(ncol(census.list[[1]]),".attack.", sep=""))]<-0

}

stat<-log(1+stat)

rec.stat[[k]][[1]]<-abind(rec.stat[[k]][[1]], stat)

dimnames(rec.stat[[k]][[1]])[[3]][ncol(rec.stat[[k]][[1]][1,,])]<-"win.log.count.pairwise.day"

}

return(rec.stat)

	
}


## Are you more or less likely to attack me if I LOST to you MORE TIMES in the past day?

add.loss.log.count.pairwise.day<-function(x){
	
	rec.stat<-x

library(abind)

#loop through each day sequence
for(k in day){

#empty matrix of the right dimensions, event# x event type. In this case, the default is all zeroes. 
stat<-matrix(0, nrow=step, ncol=length(event.types))
colnames(stat)<-event.types


#this loops through each event in the sequence

for(j in step){

#for each event, loop through all the males to get the previous attack count that will predict the next interaction- the support array will take care of males that weren't present that day.
for(i in 1:ncol(census.list[[1]])){
	
#list of all the losses by the focal male
losses<-win.loss.census[[k]][1:(j-1),i][grep("lose",win.loss.census[[k]][1:(j-1),i])]

#strip the "chase" from the end of chase wins so I can include them
losses<-sapply(losses, function(x) if(grepl("chas",x)==TRUE){
temp<-strsplit(x, split=".", fixed=T)[[1]]	
new<-paste(temp[1],temp[2],temp[3], sep=".")
return(new)
}
else return(x)
)

#if the male has not had any interactions yet, the hazard is equal across event types involving them, so set all to zero 
if(length(losses)>0){

#totals for each type- these will be the values for the stat
lose.tots<-table(losses)

#get just the interacting male id numbers
rankids<-sapply(names(lose.tots), function(x) strsplit(x, split=".", fixed=T)[[1]][3])

#match with event types- this stat predicts INCOMING attacks based on total previous LOSSES by the focal male
#loop thru each element of rankids

for(l in 1:length(rankids)){

#incoming attacks directed at the focal male, the hazrd is proportional to the number of previous losses
stat[j,][paste(rankids[l],".attack.", i, sep="")]<-lose.tots[l]
	}


		
}
	
}

#exclude attack events involving the X males 
stat[j,][endsWith(event.types,paste(".attack.",ncol(census.list[[1]]), sep="")) | startsWith(event.types,paste(ncol(census.list[[1]]),".attack.", sep=""))]<-0

}

stat<-log(1+stat)

rec.stat[[k]][[1]]<-abind(rec.stat[[k]][[1]], stat)

dimnames(rec.stat[[k]][[1]])[[3]][ncol(rec.stat[[k]][[1]][1,,])]<-"loss.log.count.pairwise.day"

}

return(rec.stat)

}


#try accounting for "linkages" instead- same answer, or better? Right now I'm treating initiations and wins as independent, but they are always pairs, and they covary in particular, and apparently important ways that I am not capturing. Maybe the model is making up for the missing covariance term by forming these associations among the variables? so i want one for the total number and then four for the "linkages". Yeah, right now I'm double counting, since the total of all four is actually twice the total number of interactions. So it's proportional but not identical. Interesting. May want to rethink recency metrics too? (it's not the end of the world if my model is a little imperfect because I am using multiple metrics to approximate one better one- I'l still get a similar result at the end, it seems)


### log number of attack+wins   


add.attack.win.count.pairwise.day<-function(x){

new.stat<-x

library(abind)

#loop through each day sequence
for(k in day){

#empty matrix of the right dimensions, event# x event type. In this case, the default is all zeroes. 
stat<-matrix(0, nrow=step, ncol=length(event.types))
colnames(stat)<-event.types


#this loops through each event in the sequence

for(j in step){

#for each event, loop through all the males to get the recency ranks that will predict the next interaction- the support array will take care of males that weren't present that day.
for(i in 1:ncol(census.list[[1]])){
	
#want to start the loop for the stat after the first interaction is complete, but I want to start the accounting of the past interactions from the beginning

int.vector<-win.loss.census[[k]][1:(j-1),i]


#Remove the "fights" and "nothings" from the census list. This creates attack/receive and win/lose pairs in order of occurence
fight.times<-grep("fight",int.vector)
nothing.times<-grep("nothing",int.vector)

pairs<-int.vector[-c(fight.times, nothing.times)]

#need a way to do this that matches interaction partners but also is sensitive to the time-ordering 

attack.wins<-vector()

if(length(pairs)>0){

attack.wins<-unlist(sapply(1:ncol(census.list[[1]]), function(z){
	
	intset<-pairs[sapply(pairs, function(x) {
		
		strsplit(x,split=".",fixed=T)[[1]][3]==z
		
	})]
	
	names(intset)<-grep("TRUE",sapply(pairs, function(x) {
		
		strsplit(x,split=".",fixed=T)[[1]][3]==z
		
	}))
	
	attacks<-grep("attack",intset)
	attackpluswin<-intset[attacks+1][grep("win", intset[attacks+1])]
	
	return(attackpluswin)
	
}))
}


#if the male has not had any interactions yet, the hazard is equal across event types involving them, so set all to zero 
if(length(attack.wins)>0){

#strip the "chase" from the end of chase wins so I can include them
attack.wins<-sapply(attack.wins, function(x) if(grepl("chas",x)==TRUE){
temp<-strsplit(x, split=".", fixed=T)[[1]]	
new<-paste(temp[1],temp[2],temp[3], sep=".")
return(new)
}
else return(x)
)
	
#get the names of the interaction events put in descending order, most recent first
#attack.wins<-attack.wins[order(names(attack.wins), decreasing=T)]

#get the number of interactions with each other male
counts<-table(attack.wins)	

#get just the interacting male id number
rankids<-sapply(names(counts), function(x) strsplit(x, split=".", fixed=T)[[1]][3])
	
#match with event types- this stat predicts new INCOMING attacks based on the number of previous OUTGOING attacks + WINS
#loop thru each element of rankids

for(l in 1:length(rankids)){

#incoming attacks directed at the focal male
stat[j,][paste(rankids[l],".attack.", i, sep="")]<-counts[l]
	}
		
}
	
}

#exclude attack events involving the X males 
stat[j,][endsWith(event.types,paste(".attack.",ncol(census.list[[1]]), sep="")) | startsWith(event.types,paste(ncol(census.list[[1]]),".attack.", sep=""))]<-0

}

#take the log 
stat<-log(1+stat)

new.stat[[k]][[1]]<-abind(new.stat[[k]][[1]], stat)

dimnames(new.stat[[k]][[1]])[[3]][ncol(new.stat[[k]][[1]][1,,])]<-"attack.win.count.pairwise.day"

}

return(new.stat)

}


 ## log number of attack-losses  


add.attack.lose.count.pairwise.day<-function(x){

new.stat<-x

library(abind)

#loop through each day sequence
for(k in day){

#empty matrix of the right dimensions, event# x event type. In this case, the default is all zeroes. 
stat<-matrix(0, nrow=step, ncol=length(event.types))
colnames(stat)<-event.types


#this loops through each event in the sequence

for(j in step){

#for each event, loop through all the males to get the recency ranks that will predict the next interaction- the support array will take care of males that weren't present that day.
for(i in 1:ncol(census.list[[1]])){
	
#want to start the loop for the stat after the first interaction is complete, but I want to start the accounting of the past interactions from the beginning

int.vector<-win.loss.census[[k]][1:(j-1),i]


#Remove the "fights" and "nothings" from the census list. This creates attack/receive and win/lose pairs in order of occurence
fight.times<-grep("fight",int.vector)
nothing.times<-grep("nothing",int.vector)

pairs<-int.vector[-c(fight.times, nothing.times)]

#need a way to do this that matches interaction partners but also is sensitive to the time-ordering 

attack.losses<-vector()

if(length(pairs)>0){

attack.losses<-unlist(sapply(1:ncol(census.list[[1]]), function(z){
	
	intset<-pairs[sapply(pairs, function(x) {
		
		strsplit(x,split=".",fixed=T)[[1]][3]==z
		
	})]
	
	names(intset)<-grep("TRUE",sapply(pairs, function(x) {
		
		strsplit(x,split=".",fixed=T)[[1]][3]==z
		
	}))
	
	attacks<-grep("attack",intset)
	attackpluslose<-intset[attacks+1][grep("lose", intset[attacks+1])]
	
	return(attackpluslose)
	
}))
}


#if the male has not had any interactions yet, the hazard is equal across event types involving them, so set all to zero 
if(length(attack.losses)>0){

#strip the "chase" from the end of chase wins so I can include them
attack.losses<-sapply(attack.losses, function(x) if(grepl("chas",x)==TRUE){
temp<-strsplit(x, split=".", fixed=T)[[1]]	
new<-paste(temp[1],temp[2],temp[3], sep=".")
return(new)
}
else return(x)
)


#get the number of interactions with each other male
counts<-table(attack.losses)	

#get just the interacting male id number
rankids<-sapply(names(counts), function(x) strsplit(x, split=".", fixed=T)[[1]][3])
	
#match with event types- this stat predicts new INCOMING attacks based on the number of previous OUTGOING attacks + WINS
#loop thru each element of rankids

for(l in 1:length(rankids)){

#incoming attacks directed at the focal male
stat[j,][paste(rankids[l],".attack.", i, sep="")]<-counts[l]
	}
		
}
	
}

#exclude attack events involving the X males 
stat[j,][endsWith(event.types,paste(".attack.",ncol(census.list[[1]]), sep="")) | startsWith(event.types,paste(ncol(census.list[[1]]),".attack.", sep=""))]<-0

}

#take the log 
stat<-log(1+stat)

new.stat[[k]][[1]]<-abind(new.stat[[k]][[1]], stat)

dimnames(new.stat[[k]][[1]])[[3]][ncol(new.stat[[k]][[1]][1,,])]<-"attack.lose.count.pairwise.day"

}

return(new.stat)

}


## log number of Receive-wins?

add.receive.win.count.pairwise.day<-function(x){

new.stat<-x

library(abind)

#loop through each day sequence
for(k in day){

#empty matrix of the right dimensions, event# x event type. In this case, the default is all zeroes. 
stat<-matrix(0, nrow=step, ncol=length(event.types))
colnames(stat)<-event.types


#this loops through each event in the sequence

for(j in step){

#for each event, loop through all the males to get the recency ranks that will predict the next interaction- the support array will take care of males that weren't present that day.
for(i in 1:ncol(census.list[[1]])){
	
#want to start the loop for the stat after the first interaction is complete, but I want to start the accounting of the past interactions from the beginning

int.vector<-win.loss.census[[k]][1:(j-1),i]


#Remove the "fights" and "nothings" from the census list. This creates attack/receive and win/lose pairs in order of occurence
fight.times<-grep("fight",int.vector)
nothing.times<-grep("nothing",int.vector)

pairs<-int.vector[-c(fight.times, nothing.times)]

#need a way to do this that matches interaction partners but also is sensitive to the time-ordering 

receive.wins<-vector()

if(length(pairs)>0){

receive.wins<-unlist(sapply(1:ncol(census.list[[1]]), function(z){
	
	intset<-pairs[sapply(pairs, function(x) {
		
		strsplit(x,split=".",fixed=T)[[1]][3]==z
		
	})]
	
	names(intset)<-grep("TRUE",sapply(pairs, function(x) {
		
		strsplit(x,split=".",fixed=T)[[1]][3]==z
		
	}))
	
	receives<-grep("receive",intset)
	receivepluswin<-intset[receives+1][grep("win", intset[receives+1])]
	
	return(receivepluswin)
	
}))
}


#if the male has not had any interactions yet, the hazard is equal across event types involving them, so set all to zero 
if(length(receive.wins)>0){

#strip the "chase" from the end of chase wins so I can include them
receive.wins<-sapply(receive.wins, function(x) if(grepl("chas",x)==TRUE){
temp<-strsplit(x, split=".", fixed=T)[[1]]	
new<-paste(temp[1],temp[2],temp[3], sep=".")
return(new)
}
else return(x)
)


#get the number of interactions with each other male
counts<-table(receive.wins)	

#get just the interacting male id number
rankids<-sapply(names(counts), function(x) strsplit(x, split=".", fixed=T)[[1]][3])
	
#match with event types- this stat predicts new INCOMING attacks based on the number of previous OUTGOING attacks + WINS
#loop thru each element of rankids

for(l in 1:length(rankids)){

#incoming attacks directed at the focal male
stat[j,][paste(rankids[l],".attack.", i, sep="")]<-counts[l]
	}
		
}
	
}

#exclude attack events involving the X males 
stat[j,][endsWith(event.types,paste(".attack.",ncol(census.list[[1]]), sep="")) | startsWith(event.types,paste(ncol(census.list[[1]]),".attack.", sep=""))]<-0

}

#take the log 
stat<-log(1+stat)

new.stat[[k]][[1]]<-abind(new.stat[[k]][[1]], stat)

dimnames(new.stat[[k]][[1]])[[3]][ncol(new.stat[[k]][[1]][1,,])]<-"receive.win.count.pairwise.day"

}

return(new.stat)

}


## log number of receive-losses?


add.receive.lose.count.pairwise.day<-function(x){

new.stat<-x

library(abind)

#loop through each day sequence
for(k in day){

#empty matrix of the right dimensions, event# x event type. In this case, the default is all zeroes. 
stat<-matrix(0, nrow=step, ncol=length(event.types))
colnames(stat)<-event.types


#this loops through each event in the sequence

for(j in step){

#for each event, loop through all the males to get the recency ranks that will predict the next interaction- the support array will take care of males that weren't present that day.
for(i in 1:ncol(census.list[[1]])){
	
#want to start the loop for the stat after the first interaction is complete, but I want to start the accounting of the past interactions from the beginning

int.vector<-win.loss.census[[k]][1:(j-1),i]


#Remove the "fights" and "nothings" from the census list. This creates attack/receive and win/lose pairs in order of occurence
fight.times<-grep("fight",int.vector)
nothing.times<-grep("nothing",int.vector)

pairs<-int.vector[-c(fight.times, nothing.times)]

#need a way to do this that matches interaction partners but also is sensitive to the time-ordering 

receive.losses<-vector()

if(length(pairs)>0){

receive.losses<-unlist(sapply(1:ncol(census.list[[1]]), function(z){
	
	intset<-pairs[sapply(pairs, function(x) {
		
		strsplit(x,split=".",fixed=T)[[1]][3]==z
		
	})]
	
	names(intset)<-grep("TRUE",sapply(pairs, function(x) {
		
		strsplit(x,split=".",fixed=T)[[1]][3]==z
		
	}))
	
	receives<-grep("receive",intset)
	receivepluslose<-intset[receives+1][grep("lose", intset[receives+1])]
	
	return(receivepluslose)
	
}))
}


#if the male has not had any interactions yet, the hazard is equal across event types involving them, so set all to zero 
if(length(receive.losses)>0){

#strip the "chase" from the end of chase wins so I can include them
receive.losses<-sapply(receive.losses, function(x) if(grepl("chas",x)==TRUE){
temp<-strsplit(x, split=".", fixed=T)[[1]]	
new<-paste(temp[1],temp[2],temp[3], sep=".")
return(new)
}
else return(x)
)


#get the number of interactions with each other male
counts<-table(receive.losses)	

#get just the interacting male id number
rankids<-sapply(names(counts), function(x) strsplit(x, split=".", fixed=T)[[1]][3])
	
#match with event types- this stat predicts new INCOMING attacks based on the number of previous OUTGOING attacks + WINS
#loop thru each element of rankids

for(l in 1:length(rankids)){

#incoming attacks directed at the focal male
stat[j,][paste(rankids[l],".attack.", i, sep="")]<-counts[l]
	}
		
}
	
}

#exclude attack events involving the X males 
stat[j,][endsWith(event.types,paste(".attack.",ncol(census.list[[1]]), sep="")) | startsWith(event.types,paste(ncol(census.list[[1]]),".attack.", sep=""))]<-0

}

#take the log 
stat<-log(1+stat)

new.stat[[k]][[1]]<-abind(new.stat[[k]][[1]], stat)

dimnames(new.stat[[k]][[1]])[[3]][ncol(new.stat[[k]][[1]][1,,])]<-"receive.lose.count.pairwise.day"

}

return(new.stat)

}


### test the effects of pairwise win/loss record for the day

##Are you more likely to attack me if I have a positive win/loss record against you?


add.win.loss.pairwise.day<-function(x){
	
rec.stat<-x

library(abind)

#loop through each day sequence
for(k in day){

#empty matrix of the right dimensions, event# x event type. In this case, the default is all zeroes. 
stat<-matrix(0, nrow=step, ncol=length(event.types))
colnames(stat)<-event.types


#this loops through each event in the sequence

for(j in step){

#for each event, loop through all the males to get the previous attack count that will predict the next interaction- the support array will take care of males that weren't present that day.
for(i in 1:ncol(census.list[[1]])){
	
#start with simple +/-

#list of all the wins by the focal male
wins<-win.loss.census[[k]][1:(j-1),i][grep("win",win.loss.census[[k]][1:(j-1),i])]

#list of all the losses by the focal male
losses<-win.loss.census[[k]][1:(j-1),i][grep("lose",win.loss.census[[k]][1:(j-1),i])]

#if the male has not had any interactions yet, the hazard is equal across event types involving them, so set all to zero
#do wins first 
if(length(wins)>0){

#strip the "chase" from the end of chase wins so I can include them
wins<-sapply(wins, function(x) if(grepl("chas",x)==TRUE){
temp<-strsplit(x, split=".", fixed=T)[[1]]	
new<-paste(temp[1],temp[2],temp[3], sep=".")
return(new)
}
else return(x)
)


#totals for each type- these will be the values for the stat
win.tots<-table(wins)

#get just the interacting male id numbers
winids<-sapply(names(win.tots), function(x) strsplit(x, split=".", fixed=T)[[1]][3])

#match with event types- this stat predicts INCOMING attacks based on the difference of WINS and LOSSES of the FOCAL MALE

#loop thru each element of winids first

for(l in 1:length(winids)){

#incoming attacks directed at the focal male
stat[j,][paste(winids[l],".attack.", i, sep="")]<-win.tots[l]
	}
 
}

#now substract losses if there are any
if(length(losses)>0){

#strip the "chase" from the end of chase wins so I can include them
losses<-sapply(losses, function(x) if(grepl("chas",x)==TRUE){
temp<-strsplit(x, split=".", fixed=T)[[1]]	
new<-paste(temp[1],temp[2],temp[3], sep=".")
return(new)
}
else return(x)
)
	
	
lose.tots<-table(losses)
	
loseids<-sapply(names(lose.tots), function(x) strsplit(x, split=".", fixed=T)[[1]][3])

#loop thru each element of loseids, subtracting from whatever is there already

for(l in 1:length(loseids)){

#incoming attacks directed at the focal male
stat[j,][paste(loseids[l],".attack.", i, sep="")]<-stat[j,][paste(loseids[l],".attack.", i, sep="")]-lose.tots[l]
	}

}

}

#exclude attack events involving the X males 
stat[j,][endsWith(event.types,paste(".attack.",ncol(census.list[[1]]), sep="")) | startsWith(event.types,paste(ncol(census.list[[1]]),".attack.", sep=""))]<-0

}


rec.stat[[k]][[1]]<-abind(rec.stat[[k]][[1]], stat)

dimnames(rec.stat[[k]][[1]])[[3]][ncol(rec.stat[[k]][[1]][1,,])]<-"win.loss.pairwise.day"

}

return(rec.stat)

}


##Are you more likely to attack me if I have a positive log(1+win/loss) record against you?


add.log.win.loss.pairwise.day<-function(x){
	
rec.stat<-x

library(abind)

#loop through each day sequence
for(k in day){

#empty matrix of the right dimensions, event# x event type. In this case, the default is all zeroes. 
stat<-matrix(0, nrow=step, ncol=length(event.types))
colnames(stat)<-event.types


#this loops through each event in the sequence

for(j in step){

#for each event, loop through all the males to get the previous attack count that will predict the next interaction- the support array will take care of males that weren't present that day.
for(i in 1:ncol(census.list[[1]])){
	
#start with simple +/-

#list of all the wins by the focal male
wins<-win.loss.census[[k]][1:(j-1),i][grep("win",win.loss.census[[k]][1:(j-1),i])]

#list of all the losses by the focal male
losses<-win.loss.census[[k]][1:(j-1),i][grep("lose",win.loss.census[[k]][1:(j-1),i])]

#if the male has not had any interactions yet, the hazard is equal across event types involving them, so set all to zero
#do wins first 
if(length(wins)>0){

#strip the "chase" from the end of chase wins so I can include them
wins<-sapply(wins, function(x) if(grepl("chas",x)==TRUE){
temp<-strsplit(x, split=".", fixed=T)[[1]]	
new<-paste(temp[1],temp[2],temp[3], sep=".")
return(new)
}
else return(x)
)


#totals for each type- these will be the values for the stat
win.tots<-table(wins)

#get just the interacting male id numbers
winids<-sapply(names(win.tots), function(x) strsplit(x, split=".", fixed=T)[[1]][3])

#match with event types- this stat predicts INCOMING attacks based on the difference of WINS and LOSSES of the FOCAL MALE

#loop thru each element of winids first

for(l in 1:length(winids)){

#incoming attacks directed at the focal male
stat[j,][paste(winids[l],".attack.", i, sep="")]<-win.tots[l]
	}
 
}

#now substract losses if there are any
if(length(losses)>0){

#strip the "chase" from the end of chase wins so I can include them
losses<-sapply(losses, function(x) if(grepl("chas",x)==TRUE){
temp<-strsplit(x, split=".", fixed=T)[[1]]	
new<-paste(temp[1],temp[2],temp[3], sep=".")
return(new)
}
else return(x)
)
	
	
lose.tots<-table(losses)
	
loseids<-sapply(names(lose.tots), function(x) strsplit(x, split=".", fixed=T)[[1]][3])

#loop thru each element of loseids, subtracting from whatever is there already

for(l in 1:length(loseids)){

#incoming attacks directed at the focal male
stat[j,][paste(loseids[l],".attack.", i, sep="")]<-stat[j,][paste(loseids[l],".attack.", i, sep="")]-lose.tots[l]
	}

}

}

#exclude attack events involving the X males 
stat[j,][endsWith(event.types,paste(".attack.",ncol(census.list[[1]]), sep="")) | startsWith(event.types,paste(ncol(census.list[[1]]),".attack.", sep=""))]<-0

}

#take the log since these numbers can become quite large and we donot expect the hazard to increase exponentially forever with win-loss record. Have to do some fancy stuff to deal with the negative records

stat<-(log(1+abs(stat))/(stat))*abs(stat) 
stat[is.na(stat)]<-0

rec.stat[[k]][[1]]<-abind(rec.stat[[k]][[1]], stat)

dimnames(rec.stat[[k]][[1]])[[3]][ncol(rec.stat[[k]][[1]][1,,])]<-"log.win.loss.pairwise.day"

}

return(rec.stat)

}


## Are you more likely to attack me if we are more evenly matched (our win/loss record is closer to 0)?

#just take the absolute value of the win/loss record. Smaller values mean more evenly matched males


add.abs.winloss.diff.pairwise.day<-function(x){
	
rec.stat<-x

library(abind)

#loop through each day sequence
for(k in day){

#empty matrix of the right dimensions, event# x event type. In this case, the default is all zeroes. 
stat<-matrix(0, nrow=step, ncol=length(event.types))
colnames(stat)<-event.types


#this loops through each event in the sequence

for(j in step){

#for each event, loop through all the males to get the previous attack count that will predict the next interaction- the support array will take care of males that weren't present that day.
for(i in 1:ncol(census.list[[1]])){
	
#start with simple +/-

#list of all the wins by the focal male
wins<-win.loss.census[[k]][1:(j-1),i][grep("win",win.loss.census[[k]][1:(j-1),i])]

#list of all the losses by the focal male
losses<-win.loss.census[[k]][1:(j-1),i][grep("lose",win.loss.census[[k]][1:(j-1),i])]

#if the male has not had any interactions yet, the hazard is equal across event types involving them, so set all to zero
#do wins first 
if(length(wins)>0){

#strip the "chase" from the end of chase wins so I can include them
wins<-sapply(wins, function(x) if(grepl("chas",x)==TRUE){
temp<-strsplit(x, split=".", fixed=T)[[1]]	
new<-paste(temp[1],temp[2],temp[3], sep=".")
return(new)
}
else return(x)
)


#totals for each type- these will be the values for the stat
win.tots<-table(wins)

#get just the interacting male id numbers
winids<-sapply(names(win.tots), function(x) strsplit(x, split=".", fixed=T)[[1]][3])

#match with event types- this stat predicts INCOMING attacks based on the difference of WINS and LOSSES of the FOCAL MALE

#loop thru each element of winids first

for(l in 1:length(winids)){

#incoming attacks directed at the focal male
stat[j,][paste(winids[l],".attack.", i, sep="")]<-win.tots[l]
	}
 
}

#now substract losses if there are any
if(length(losses)>0){

#strip the "chase" from the end of chase wins so I can include them
losses<-sapply(losses, function(x) if(grepl("chas",x)==TRUE){
temp<-strsplit(x, split=".", fixed=T)[[1]]	
new<-paste(temp[1],temp[2],temp[3], sep=".")
return(new)
}
else return(x)
)
	
	
lose.tots<-table(losses)
	
loseids<-sapply(names(lose.tots), function(x) strsplit(x, split=".", fixed=T)[[1]][3])

#loop thru each element of loseids, subtracting from whatever is there already

for(l in 1:length(loseids)){

#incoming attacks directed at the focal male
stat[j,][paste(loseids[l],".attack.", i, sep="")]<-stat[j,][paste(loseids[l],".attack.", i, sep="")]-lose.tots[l]
	}

}

}

#exclude attack events involving the X males 
stat[j,][endsWith(event.types,paste(".attack.",ncol(census.list[[1]]), sep="")) | startsWith(event.types,paste(ncol(census.list[[1]]),".attack.", sep=""))]<-0

}

#take the absolute value of the stat matrix to turn all negative win/loss records positive so that larger numbers jsut mean farther from evenly matched

stat<-abs(stat)

rec.stat[[k]][[1]]<-abind(rec.stat[[k]][[1]], stat)

dimnames(rec.stat[[k]][[1]])[[3]][ncol(rec.stat[[k]][[1]][1,,])]<-"abs.winloss.diff.pairwise.day"

}

return(rec.stat)

}


## try log(evenly.matched)

add.log.abs.winloss.diff.pairwise.day<-function(x){
	
rec.stat<-x

library(abind)

#loop through each day sequence
for(k in day){

#empty matrix of the right dimensions, event# x event type. In this case, the default is all zeroes. 
stat<-matrix(0, nrow=step, ncol=length(event.types))
colnames(stat)<-event.types


#this loops through each event in the sequence

for(j in step){

#for each event, loop through all the males to get the previous attack count that will predict the next interaction- the support array will take care of males that weren't present that day.
for(i in 1:ncol(census.list[[1]])){
	
#start with simple +/-

#list of all the wins by the focal male
wins<-win.loss.census[[k]][1:(j-1),i][grep("win",win.loss.census[[k]][1:(j-1),i])]

#list of all the losses by the focal male
losses<-win.loss.census[[k]][1:(j-1),i][grep("lose",win.loss.census[[k]][1:(j-1),i])]

#if the male has not had any interactions yet, the hazard is equal across event types involving them, so set all to zero
#do wins first 
if(length(wins)>0){

#strip the "chase" from the end of chase wins so I can include them
wins<-sapply(wins, function(x) if(grepl("chas",x)==TRUE){
temp<-strsplit(x, split=".", fixed=T)[[1]]	
new<-paste(temp[1],temp[2],temp[3], sep=".")
return(new)
}
else return(x)
)


#totals for each type- these will be the values for the stat
win.tots<-table(wins)

#get just the interacting male id numbers
winids<-sapply(names(win.tots), function(x) strsplit(x, split=".", fixed=T)[[1]][3])

#match with event types- this stat predicts INCOMING attacks based on the difference of WINS and LOSSES of the FOCAL MALE

#loop thru each element of winids first

for(l in 1:length(winids)){

#incoming attacks directed at the focal male
stat[j,][paste(winids[l],".attack.", i, sep="")]<-win.tots[l]
	}
 
}

#now substract losses if there are any
if(length(losses)>0){

#strip the "chase" from the end of chase wins so I can include them
losses<-sapply(losses, function(x) if(grepl("chas",x)==TRUE){
temp<-strsplit(x, split=".", fixed=T)[[1]]	
new<-paste(temp[1],temp[2],temp[3], sep=".")
return(new)
}
else return(x)
)
	
	
lose.tots<-table(losses)
	
loseids<-sapply(names(lose.tots), function(x) strsplit(x, split=".", fixed=T)[[1]][3])

#loop thru each element of loseids, subtracting from whatever is there already

for(l in 1:length(loseids)){

#incoming attacks directed at the focal male
stat[j,][paste(loseids[l],".attack.", i, sep="")]<-stat[j,][paste(loseids[l],".attack.", i, sep="")]-lose.tots[l]
	}

}

}

#exclude attack events involving the X males 
stat[j,][endsWith(event.types,paste(".attack.",ncol(census.list[[1]]), sep="")) | startsWith(event.types,paste(ncol(census.list[[1]]),".attack.", sep=""))]<-0

}

#take the absolute value of the stat matrix to turn all negative win/loss records positive so that larger numbers jsut mean farther from evenly matched

stat<-abs(stat)

stat<-log(1+stat)

rec.stat[[k]][[1]]<-abind(rec.stat[[k]][[1]], stat)

dimnames(rec.stat[[k]][[1]])[[3]][ncol(rec.stat[[k]][[1]][1,,])]<-"log.abs.winloss.diff.pairwise.day"

}

return(rec.stat)

}


##Are you more likely to attack me if I have a greater win/loss rate against you?

add.win.loss.rate.pairwise.day<-function(x){
	
rec.stat<-x

library(abind)

#loop through each day sequence
for(k in day){

#empty matrix of the right dimensions, event# x event type. In this case, the default is all zeroes. 
stat<-matrix(0, nrow=step, ncol=length(event.types))
colnames(stat)<-event.types


#this loops through each event in the sequence

for(j in step){

#for each event, loop through all the males to get the previous attack count that will predict the next interaction- the support array will take care of males that weren't present that day.
for(i in 1:ncol(census.list[[1]])){
	
#start with simple +/-

#list of all the wins by the focal male
wins<-win.loss.census[[k]][1:(j-1),i][grep("win",win.loss.census[[k]][1:(j-1),i])]

#list of all the losses by the focal male
losses<-win.loss.census[[k]][1:(j-1),i][grep("lose",win.loss.census[[k]][1:(j-1),i])]

#if the male has not had any interactions yet, the hazard is equal across event types involving them, so set all to zero
#do wins first 
if(length(wins)>0){

#strip the "chase" from the end of chase wins so I can include them
wins<-sapply(wins, function(x) if(grepl("chas",x)==TRUE){
temp<-strsplit(x, split=".", fixed=T)[[1]]	
new<-paste(temp[1],temp[2],temp[3], sep=".")
return(new)
}
else return(x)
)


#totals for each type- these will be the values for the stat
win.tots<-table(wins)

#get just the interacting male id numbers
winids<-sapply(names(win.tots), function(x) strsplit(x, split=".", fixed=T)[[1]][3])

allwins<-rep(0,ncol(census.list[[1]]))
 
for(l in 1:length(winids)){
	
allwins[as.numeric(winids[l])]<-win.tots[l]
}

} else {
	allwins<-rep(0,ncol(census.list[[1]]))
	}


#now substract losses if there are any
if(length(losses)>0){

#strip the "chase" from the end of chase wins so I can include them
losses<-sapply(losses, function(x) if(grepl("chas",x)==TRUE){
temp<-strsplit(x, split=".", fixed=T)[[1]]	
new<-paste(temp[1],temp[2],temp[3], sep=".")
return(new)
}
else return(x)
)
	
	
lose.tots<-table(losses)
	
loseids<-sapply(names(lose.tots), function(x) strsplit(x, split=".", fixed=T)[[1]][3])

allloss<-rep(0,ncol(census.list[[1]]))
 
for(l in 1:length(loseids)){
	
allloss[as.numeric(loseids[l])]<-lose.tots[l]
}


} else {
	
	allloss<-rep(0,ncol(census.list[[1]]))
	
}


#now I can make a vector of all the win/loss rates with each other male in order

rates<-(allwins+0.5)/(allwins+allloss+1)

#get rid of the win rate for the focal male against itself
rates<-rates[-i]

#match with incoming attacks directed at the focal male

#incoming attacks directed at the focal male
stat[j,][event.types[endsWith(event.types,paste0(".attack.",i))]]<-rates

}

#exclude attack events involving the X males; here the baseline is 0.5 for 0 wins and 0 losses
stat[j,][endsWith(event.types,paste(".attack.",ncol(census.list[[1]]), sep="")) | startsWith(event.types,paste(ncol(census.list[[1]]),".attack.", sep=""))]<-0.5

}


rec.stat[[k]][[1]]<-abind(rec.stat[[k]][[1]], stat)

dimnames(rec.stat[[k]][[1]])[[3]][ncol(rec.stat[[k]][[1]][1,,])]<-"win.loss.rate.pairwise.day"

}

return(rec.stat)

}


############# General Effects, Day-specific ############

#cumulative day-specific stats that affect the hazard of being attacked in general rather than by specific individuals
#some of these correlate highly with a metric like eigenvector centrality: degree and edgeweight

## Am I more likely to be attacked if I have had a larger number of interaction partners today? (degree)

add.degree.day<-function(x){
	
rec.stat<-x

library(abind)

#loop through each day sequence
for(k in day){

#empty matrix of the right dimensions, event# x event type. In this case, the default is all zeroes. 
stat<-matrix(0, nrow=step, ncol=length(event.types))
colnames(stat)<-event.types


#this loops through each event in the sequence

for(j in step){

#for each event, loop through all the males to get the previous attack count that will predict the next interaction- the support array will take care of males that weren't present that day.
for(i in 1:ncol(census.list[[1]])){
	
#get number of unique interaction partners

#list of all the wins by the focal male
wins<-grep("win",win.loss.census[[k]][1:(j-1),i])

#list of all the losses by the focal male
losses<-grep("lose",win.loss.census[[k]][1:(j-1),i])

#this covers all the interactions that have happened so far for the focal male. Stick them all together
ints<-union(wins,losses)

#if the male has not had any interactions yet, the hazard is equal across event types involving them, so set all to zero
#do wins first 
if(length(ints)>0){

#totals for each type- these will be the values for the stat
int.names<-win.loss.census[[k]][1:(j-1),i][ints]

#get just the interacting male id numbers
intids<-sapply(int.names, function(x) strsplit(x, split=".", fixed=T)[[1]][3])

#collapse to unique parters
intids<-unique(intids)

#match with event types- this stat predicts general INCOMING attacks based on the total number of interaction partners so far that day for the focal male (degree)

stat[j,][endsWith(event.types,paste(".attack.",i, sep=""))]<-length(intids)
}


}

#exclude attack events involving the X males 
stat[j,][endsWith(event.types,paste(".attack.",ncol(census.list[[1]]), sep="")) | startsWith(event.types,paste(ncol(census.list[[1]]),".attack.", sep=""))]<-0

}


rec.stat[[k]][[1]]<-abind(rec.stat[[k]][[1]], stat)

dimnames(rec.stat[[k]][[1]])[[3]][ncol(rec.stat[[k]][[1]][1,,])]<-"degree.day"

}

return(rec.stat)

}


##try log(degree) just to see

add.log.degree.day<-function(x){
	
rec.stat<-x

library(abind)

#loop through each day sequence
for(k in day){

#empty matrix of the right dimensions, event# x event type. In this case, the default is all zeroes. 
stat<-matrix(0, nrow=step, ncol=length(event.types))
colnames(stat)<-event.types


#this loops through each event in the sequence

for(j in step){

#for each event, loop through all the males to get the previous attack count that will predict the next interaction- the support array will take care of males that weren't present that day.
for(i in 1:ncol(census.list[[1]])){
	
#get number of unique interaction partners

#list of all the wins by the focal male
wins<-grep("win",win.loss.census[[k]][1:(j-1),i])

#list of all the losses by the focal male
losses<-grep("lose",win.loss.census[[k]][1:(j-1),i])

#this covers all the interactions that have happened so far for the focal male. Stick them all together
ints<-union(wins,losses)

#if the male has not had any interactions yet, the hazard is equal across event types involving them, so set all to zero
#do wins first 
if(length(ints)>0){

#totals for each type- these will be the values for the stat
int.names<-win.loss.census[[k]][1:(j-1),i][ints]

#get just the interacting male id numbers
intids<-sapply(int.names, function(x) strsplit(x, split=".", fixed=T)[[1]][3])

#collapse to unique parters
intids<-unique(intids)

#match with event types- this stat predicts general INCOMING attacks based on the total number of interaction partners so far that day for the focal male (degree)

stat[j,][endsWith(event.types,paste(".attack.",i, sep=""))]<-length(intids)
}


}

#exclude attack events involving the X males 
stat[j,][endsWith(event.types,paste(".attack.",ncol(census.list[[1]]), sep="")) | startsWith(event.types,paste(ncol(census.list[[1]]),".attack.", sep=""))]<-0

}

stat<-log(1+stat)

rec.stat[[k]][[1]]<-abind(rec.stat[[k]][[1]], stat)

dimnames(rec.stat[[k]][[1]])[[3]][ncol(rec.stat[[k]][[1]][1,,])]<-"log.degree.day"

}

return(rec.stat)

}


## Am I more likely to be attacked if I have a larger number of other males that I have positive win records with? (win degree)

add.win.degree.day<-function(x){
	
rec.stat<-x

library(abind)

#loop through each day sequence
for(k in day){

#empty matrix of the right dimensions, event# x event type. In this case, the default is all zeroes. 
stat<-matrix(0, nrow=step, ncol=length(event.types))
colnames(stat)<-event.types


#this loops through each event in the sequence

for(j in step){

#for each event, loop through all the males to get the previous attack count that will predict the next interaction- the support array will take care of males that weren't present that day.
for(i in 1:ncol(census.list[[1]])){
	
#start with simple +/-

#list of all the wins by the focal male
wins<-grep("win",win.loss.census[[k]][1:(j-1),i])

#list of all the losses by the focal male
losses<-grep("lose",win.loss.census[[k]][1:(j-1),i])

win.record<-rep(0,ncol(census.list[[1]]))

#if the male has not had any interactions yet, the hazard is equal across event types involving them, so set all to zero
#do wins first 
if(length(wins)>0){

wins<-win.loss.census[[k]][1:(j-1),i][wins]

#strip the "chase" from the end of chase wins so I can include them
wins<-sapply(wins, function(x) if(grepl("chas",x)==TRUE){
temp<-strsplit(x, split=".", fixed=T)[[1]]	
new<-paste(temp[1],temp[2],temp[3], sep=".")
return(new)
}
else return(x)
)

#totals for each type
win.tots<-table(wins)

#get just the interacting male id numbers
winids<-sapply(names(win.tots), function(x) strsplit(x, split=".", fixed=T)[[1]][3])

win.record[as.numeric(winids)]<-win.tots
}


#now substract losses if there are any
if(length(losses)>0){

losses<-win.loss.census[[k]][1:(j-1),i][losses]

#strip the "chase" from the end of chase wins so I can include them
losses<-sapply(losses, function(x) if(grepl("chas",x)==TRUE){
temp<-strsplit(x, split=".", fixed=T)[[1]]	
new<-paste(temp[1],temp[2],temp[3], sep=".")
return(new)
}
else return(x)
)
	
	
lose.tots<-table(losses)
	
loseids<-sapply(names(lose.tots), function(x) strsplit(x, split=".", fixed=T)[[1]][3])

win.record[as.numeric(loseids)]<-win.record[as.numeric(loseids)]-lose.tots

} 
 
#test if the win records are positive
win.pos<-win.record>0 

#Get the total number of males with whom the focal male has positive win records 
if(is.na(table(win.pos)["TRUE"])){win.pos.num<-0} else {win.pos.num<-table(win.pos)["TRUE"]}

#match with event types- this stat predicts general INCOMING attacks based on the total number of interaction partners so far that day that the focal male has positive win records with (win degree)

stat[j,][endsWith(event.types,paste(".attack.",i, sep=""))]<-win.pos.num

}

#exclude attack events involving the X males 
stat[j,][endsWith(event.types,paste(".attack.",ncol(census.list[[1]]), sep="")) | startsWith(event.types,paste(ncol(census.list[[1]]),".attack.", sep=""))]<-0
}


rec.stat[[k]][[1]]<-abind(rec.stat[[k]][[1]], stat)

dimnames(rec.stat[[k]][[1]])[[3]][ncol(rec.stat[[k]][[1]][1,,])]<-"win.degree.day"

}

return(rec.stat)
	
	
}


##check log win degree


add.log.win.degree.day<-function(x){
	
rec.stat<-x

library(abind)

#loop through each day sequence
for(k in day){

#empty matrix of the right dimensions, event# x event type. In this case, the default is all zeroes. 
stat<-matrix(0, nrow=step, ncol=length(event.types))
colnames(stat)<-event.types


#this loops through each event in the sequence

for(j in step){

#for each event, loop through all the males to get the previous attack count that will predict the next interaction- the support array will take care of males that weren't present that day.
for(i in 1:ncol(census.list[[1]])){
	
#start with simple +/-

#list of all the wins by the focal male
wins<-grep("win",win.loss.census[[k]][1:(j-1),i])

#list of all the losses by the focal male
losses<-grep("lose",win.loss.census[[k]][1:(j-1),i])

win.record<-rep(0,ncol(census.list[[1]]))

#if the male has not had any interactions yet, the hazard is equal across event types involving them, so set all to zero
#do wins first 
if(length(wins)>0){

wins<-win.loss.census[[k]][1:(j-1),i][wins]

#strip the "chase" from the end of chase wins so I can include them
wins<-sapply(wins, function(x) if(grepl("chas",x)==TRUE){
temp<-strsplit(x, split=".", fixed=T)[[1]]	
new<-paste(temp[1],temp[2],temp[3], sep=".")
return(new)
}
else return(x)
)

#totals for each type
win.tots<-table(wins)

#get just the interacting male id numbers
winids<-sapply(names(win.tots), function(x) strsplit(x, split=".", fixed=T)[[1]][3])

win.record[as.numeric(winids)]<-win.tots
}


#now substract losses if there are any
if(length(losses)>0){

losses<-win.loss.census[[k]][1:(j-1),i][losses]

#strip the "chase" from the end of chase wins so I can include them
losses<-sapply(losses, function(x) if(grepl("chas",x)==TRUE){
temp<-strsplit(x, split=".", fixed=T)[[1]]	
new<-paste(temp[1],temp[2],temp[3], sep=".")
return(new)
}
else return(x)
)
	
	
lose.tots<-table(losses)
	
loseids<-sapply(names(lose.tots), function(x) strsplit(x, split=".", fixed=T)[[1]][3])

win.record[as.numeric(loseids)]<-win.record[as.numeric(loseids)]-lose.tots

} 
 
#test if the win records are positive
win.pos<-win.record>0 

#Get the total number of males with whom the focal male has positive win records 
if(is.na(table(win.pos)["TRUE"])){win.pos.num<-0} else {win.pos.num<-table(win.pos)["TRUE"]}

#match with event types- this stat predicts general INCOMING attacks based on the total number of interaction partners so far that day that the focal male has positive win records with (win degree)

stat[j,][endsWith(event.types,paste(".attack.",i, sep=""))]<-win.pos.num

}

#exclude attack events involving the X males 
stat[j,][endsWith(event.types,paste(".attack.",ncol(census.list[[1]]), sep="")) | startsWith(event.types,paste(ncol(census.list[[1]]),".attack.", sep=""))]<-0

}

stat<-log(1+stat)

rec.stat[[k]][[1]]<-abind(rec.stat[[k]][[1]], stat)

dimnames(rec.stat[[k]][[1]])[[3]][ncol(rec.stat[[k]][[1]][1,,])]<-"log.win.degree.day"

}

return(rec.stat)
	
	
}


##just out of curiosity, what does lose degree look like? (non-winning record)

add.log.lose.degree.day<-function(x){
	
rec.stat<-x

library(abind)

#loop through each day sequence
for(k in day){

#empty matrix of the right dimensions, event# x event type. In this case, the default is all zeroes. 
stat<-matrix(0, nrow=step, ncol=length(event.types))
colnames(stat)<-event.types


#this loops through each event in the sequence

for(j in step){

#for each event, loop through all the males to get the previous attack count that will predict the next interaction- the support array will take care of males that weren't present that day.
for(i in 1:ncol(census.list[[1]])){
	
#start with simple +/-

#list of all the wins by the focal male
wins<-grep("win",win.loss.census[[k]][1:(j-1),i])

#list of all the losses by the focal male
losses<-grep("lose",win.loss.census[[k]][1:(j-1),i])

win.record<-rep(0,ncol(census.list[[1]]))

#if the male has not had any interactions yet, the hazard is equal across event types involving them, so set all to zero
#do wins first 
if(length(wins)>0){

wins<-win.loss.census[[k]][1:(j-1),i][wins]

#strip the "chase" from the end of chase wins so I can include them
wins<-sapply(wins, function(x) if(grepl("chas",x)==TRUE){
temp<-strsplit(x, split=".", fixed=T)[[1]]	
new<-paste(temp[1],temp[2],temp[3], sep=".")
return(new)
}
else return(x)
)

#totals for each type
win.tots<-table(wins)

#get just the interacting male id numbers
winids<-sapply(names(win.tots), function(x) strsplit(x, split=".", fixed=T)[[1]][3])

win.record[as.numeric(winids)]<-win.tots
}


#now substract losses if there are any
if(length(losses)>0){

losses<-win.loss.census[[k]][1:(j-1),i][losses]

#strip the "chase" from the end of chase wins so I can include them
losses<-sapply(losses, function(x) if(grepl("chas",x)==TRUE){
temp<-strsplit(x, split=".", fixed=T)[[1]]	
new<-paste(temp[1],temp[2],temp[3], sep=".")
return(new)
}
else return(x)
)
	
	
lose.tots<-table(losses)
	
loseids<-sapply(names(lose.tots), function(x) strsplit(x, split=".", fixed=T)[[1]][3])

win.record[as.numeric(loseids)]<-win.record[as.numeric(loseids)]-lose.tots
 
 
for(g in 1:length(win.record)){
	
	if(g %in% as.numeric(loseids)){
	
	win.record[g]<-as.numeric(win.record[g])-lose.tots[loseids==g]
	}
	else{if(win.record[g]==0){win.record[g]<-"no.int"}}
	
}

}

if(length(losses)==0){

win.record[win.record==0]<-"no.int"

}
 
 
#test if the win records are less than or equal to zero
win.pos<-win.record<0 

#Get the total number of males with whom the focal male has positive win records 
if(is.na(table(win.pos)["TRUE"])){win.pos.num<-0}
else{win.pos.num<-table(win.pos)["TRUE"]}

#match with event types- this stat predicts general INCOMING attacks based on the total number of interaction partners so far that day that the focal male has positive win records with (win degree)

stat[j,][endsWith(event.types,paste(".attack.",i, sep=""))]<-win.pos.num

}

#exclude attack events involving the X males 
stat[j,][endsWith(event.types,paste(".attack.",ncol(census.list[[1]]), sep="")) | startsWith(event.types,paste(ncol(census.list[[1]]),".attack.", sep=""))]<-0

}

stat<-log(1+stat)

rec.stat[[k]][[1]]<-abind(rec.stat[[k]][[1]], stat)

dimnames(rec.stat[[k]][[1]])[[3]][ncol(rec.stat[[k]][[1]][1,,])]<-"log.lose.degree.day"

}

return(rec.stat)
	
	
}


##log number of other individuals with which you have an evenly matched record

add.log.neutral.degree.day<-function(x){
	
rec.stat<-x

library(abind)

#loop through each day sequence
for(k in day){

#empty matrix of the right dimensions, event# x event type. In this case, the default is all zeroes. 
stat<-matrix(0, nrow=step, ncol=length(event.types))
colnames(stat)<-event.types


#this loops through each event in the sequence

for(j in step){

#for each event, loop through all the males to get the previous attack count that will predict the next interaction- the support array will take care of males that weren't present that day.
for(i in 1:ncol(census.list[[1]])){
	
#start with simple +/-

#list of all the wins by the focal male
wins<-grep("win",win.loss.census[[k]][1:(j-1),i])

#list of all the losses by the focal male
losses<-grep("lose",win.loss.census[[k]][1:(j-1),i])

win.record<-rep(0,ncol(census.list[[1]]))

#if the male has not had any interactions yet, the hazard is equal across event types involving them, so set all to zero
#do wins first 
if(length(wins)>0){

wins<-win.loss.census[[k]][1:(j-1),i][wins]

#strip the "chase" from the end of chase wins so I can include them
wins<-sapply(wins, function(x) if(grepl("chas",x)==TRUE){
temp<-strsplit(x, split=".", fixed=T)[[1]]	
new<-paste(temp[1],temp[2],temp[3], sep=".")
return(new)
}
else return(x)
)

#totals for each type
win.tots<-table(wins)

#get just the interacting male id numbers
winids<-sapply(names(win.tots), function(x) strsplit(x, split=".", fixed=T)[[1]][3])

win.record[as.numeric(winids)]<-win.tots
}


#now substract losses if there are any
if(length(losses)>0){

losses<-win.loss.census[[k]][1:(j-1),i][losses]

#strip the "chase" from the end of chase wins so I can include them
losses<-sapply(losses, function(x) if(grepl("chas",x)==TRUE){
temp<-strsplit(x, split=".", fixed=T)[[1]]	
new<-paste(temp[1],temp[2],temp[3], sep=".")
return(new)
}
else return(x)
)
	
	
lose.tots<-table(losses)
	
loseids<-sapply(names(lose.tots), function(x) strsplit(x, split=".", fixed=T)[[1]][3])


for(g in 1:length(win.record)){
	
	if(g %in% as.numeric(loseids)){
	
	win.record[g]<-as.numeric(win.record[g])-lose.tots[loseids==g]
	}
	else{if(win.record[g]==0){win.record[g]<-"no.int"}}
	
}

}

if(length(losses)==0){

win.record[win.record==0]<-"no.int"

}

 
#test if the win records are equal to zero
win.pos<-win.record==0 

#Get the total number of males with whom the focal male has positive win records 
if(is.na(table(win.pos)["TRUE"])){win.pos.num<-0}
else{win.pos.num<-table(win.pos)["TRUE"]}

#match with event types- this stat predicts general INCOMING attacks based on the total number of interaction partners so far that day that the focal male has positive win records with (win degree)

stat[j,][endsWith(event.types,paste(".attack.",i, sep=""))]<-win.pos.num

}

#exclude attack events involving the X males 
stat[j,][endsWith(event.types,paste(".attack.",ncol(census.list[[1]]), sep="")) | startsWith(event.types,paste(ncol(census.list[[1]]),".attack.", sep=""))]<-0

}

stat<-log(1+stat)

rec.stat[[k]][[1]]<-abind(rec.stat[[k]][[1]], stat)

dimnames(rec.stat[[k]][[1]])[[3]][ncol(rec.stat[[k]][[1]][1,,])]<-"log.neutral.degree.day"

}

return(rec.stat)
	
	
}


### What about interaction weight?

## Am I more likely to be attacked in general if my total number of previous interactions is greater?


add.log.weight.day<-function(x){
	
rec.stat<-x

library(abind)

#loop through each day sequence
for(k in day){

#empty matrix of the right dimensions, event# x event type. In this case, the default is all zeroes. 
stat<-matrix(0, nrow=step, ncol=length(event.types))
colnames(stat)<-event.types


#this loops through each event in the sequence

for(j in step){

#for each event, loop through all the males to get the previous attack count that will predict the next interaction- the support array will take care of males that weren't present that day.
for(i in 1:ncol(census.list[[1]])){
	

#list of all the attacks by the focal male
attacks<-grep("attack",win.loss.census[[k]][1:(j-1),i])

receives<-grep("received",win.loss.census[[k]][1:(j-1),i])

all<-union(attacks,receives)

#if the male has not had any interactions yet, the hazard is equal across event types involving them, so set all to zero

if(length(all)>0){

#match with event types- this stat predicts general INCOMING attacks based on the total number of attacks so far that day by the focal male
stat[j,][endsWith(event.types,paste(".attack.",i, sep=""))]<-length(all)

}

}

#exclude attack events involving the X males 
stat[j,][endsWith(event.types,paste(".attack.",ncol(census.list[[1]]), sep="")) | startsWith(event.types,paste(ncol(census.list[[1]]),".attack.", sep=""))]<-0

}

stat<-log(1+stat)

rec.stat[[k]][[1]]<-abind(rec.stat[[k]][[1]], stat)

dimnames(rec.stat[[k]][[1]])[[3]][ncol(rec.stat[[k]][[1]][1,,])]<-"log.weight.day"

}

return(rec.stat)


}


## Am I more likely to be attacked in general if my total number of previous attacks initiated is greater?

add.log.attack.weight.day<-function(x){
	
rec.stat<-x

library(abind)

#loop through each day sequence
for(k in day){

#empty matrix of the right dimensions, event# x event type. In this case, the default is all zeroes. 
stat<-matrix(0, nrow=step, ncol=length(event.types))
colnames(stat)<-event.types


#this loops through each event in the sequence

for(j in step){

#for each event, loop through all the males to get the previous attack count that will predict the next interaction- the support array will take care of males that weren't present that day.
for(i in 1:ncol(census.list[[1]])){
	

#list of all the attacks by the focal male
attacks<-grep("attack",win.loss.census[[k]][1:(j-1),i])

#if the male has not had any interactions yet, the hazard is equal across event types involving them, so set all to zero

if(length(attacks)>0){

#match with event types- this stat predicts general INCOMING attacks based on the total number of attacks so far that day by the focal male
stat[j,][endsWith(event.types,paste(".attack.",i, sep=""))]<-length(attacks)

}

}

#exclude attack events involving the X males 
stat[j,][endsWith(event.types,paste(".attack.",ncol(census.list[[1]]), sep="")) | startsWith(event.types,paste(ncol(census.list[[1]]),".attack.", sep=""))]<-0

}

stat<-log(1+stat)

rec.stat[[k]][[1]]<-abind(rec.stat[[k]][[1]], stat)

dimnames(rec.stat[[k]][[1]])[[3]][ncol(rec.stat[[k]][[1]][1,,])]<-"log.attack.weight.day"

}

return(rec.stat)


}


## Am I more likely to be attacked in general if my total number of previous receives is greater?

add.log.receive.weight.day<-function(x){
	
rec.stat<-x

library(abind)

#loop through each day sequence
for(k in day){

#empty matrix of the right dimensions, event# x event type. In this case, the default is all zeroes. 
stat<-matrix(0, nrow=step, ncol=length(event.types))
colnames(stat)<-event.types


#this loops through each event in the sequence

for(j in step){

#for each event, loop through all the males to get the previous attack count that will predict the next interaction- the support array will take care of males that weren't present that day.
for(i in 1:ncol(census.list[[1]])){
	

#list of all the attacks by the focal male
receives<-grep("received",win.loss.census[[k]][1:(j-1),i])

#if the male has not had any interactions yet, the hazard is equal across event types involving them, so set all to zero

if(length(receives)>0){

#match with event types- this stat predicts general INCOMING attacks based on the total number of attacks so far that day by the focal male
stat[j,][endsWith(event.types,paste(".attack.",i, sep=""))]<-length(receives)

}

}

#exclude attack events involving the X males 
stat[j,][endsWith(event.types,paste(".attack.",ncol(census.list[[1]]), sep="")) | startsWith(event.types,paste(ncol(census.list[[1]]),".attack.", sep=""))]<-0

}

stat<-log(1+stat)

rec.stat[[k]][[1]]<-abind(rec.stat[[k]][[1]], stat)

dimnames(rec.stat[[k]][[1]])[[3]][ncol(rec.stat[[k]][[1]][1,,])]<-"log.receive.weight.day"

}

return(rec.stat)


}


## Am I more likely to be attacked in general if my total number of wins so far is greater?

add.log.win.weight.day<-function(x){
	
rec.stat<-x

library(abind)

#loop through each day sequence
for(k in day){

#empty matrix of the right dimensions, event# x event type. In this case, the default is all zeroes. 
stat<-matrix(0, nrow=step, ncol=length(event.types))
colnames(stat)<-event.types


#this loops through each event in the sequence

for(j in step){

#for each event, loop through all the males to get the previous attack count that will predict the next interaction- the support array will take care of males that weren't present that day.
for(i in 1:ncol(census.list[[1]])){
	

#list of all the attacks by the focal male
wins<-grep("win",win.loss.census[[k]][1:(j-1),i])

#if the male has not had any interactions yet, the hazard is equal across event types involving them, so set all to zero

if(length(wins)>0){

#match with event types- this stat predicts general INCOMING attacks based on the total number of attacks so far that day by the focal male
stat[j,][endsWith(event.types,paste(".attack.",i, sep=""))]<-length(wins)

}

}

#exclude attack events involving the X males 
stat[j,][endsWith(event.types,paste(".attack.",ncol(census.list[[1]]), sep="")) | startsWith(event.types,paste(ncol(census.list[[1]]),".attack.", sep=""))]<-0

}

stat<-log(1+stat)

rec.stat[[k]][[1]]<-abind(rec.stat[[k]][[1]], stat)

dimnames(rec.stat[[k]][[1]])[[3]][ncol(rec.stat[[k]][[1]][1,,])]<-"log.win.weight.day"

}

return(rec.stat)


}


## Am I more likely to be attacked in general if my total number of losses so far is greater?

add.log.lose.weight.day<-function(x){
	
rec.stat<-x

library(abind)

#loop through each day sequence
for(k in day){

#empty matrix of the right dimensions, event# x event type. In this case, the default is all zeroes. 
stat<-matrix(0, nrow=step, ncol=length(event.types))
colnames(stat)<-event.types


#this loops through each event in the sequence

for(j in step){

#for each event, loop through all the males to get the previous attack count that will predict the next interaction- the support array will take care of males that weren't present that day.
for(i in 1:ncol(census.list[[1]])){
	

#list of all the attacks by the focal male
losses<-grep("lose",win.loss.census[[k]][1:(j-1),i])

#if the male has not had any interactions yet, the hazard is equal across event types involving them, so set all to zero

if(length(losses)>0){

#match with event types- this stat predicts general INCOMING attacks based on the total number of attacks so far that day by the focal male
stat[j,][endsWith(event.types,paste(".attack.",i, sep=""))]<-length(losses)

}

}

#exclude attack events involving the X males 
stat[j,][endsWith(event.types,paste(".attack.",ncol(census.list[[1]]), sep="")) | startsWith(event.types,paste(ncol(census.list[[1]]),".attack.", sep=""))]<-0

}

stat<-log(1+stat)

rec.stat[[k]][[1]]<-abind(rec.stat[[k]][[1]], stat)

dimnames(rec.stat[[k]][[1]])[[3]][ncol(rec.stat[[k]][[1]][1,,])]<-"log.lose.weight.day"

}

return(rec.stat)


}


## Explore "linkages" again for weights ##

## Am I more liekly to be attacked in general is I have had more attack+losses?


add.log.attack.lose.weight.day<-function(x){

new.stat<-x

library(abind)

#loop through each day sequence
for(k in day){

#empty matrix of the right dimensions, event# x event type. In this case, the default is all zeroes. 
stat<-matrix(0, nrow=step, ncol=length(event.types))
colnames(stat)<-event.types


#this loops through each event in the sequence

for(j in step){

#for each event, loop through all the males to get the recency ranks that will predict the next interaction- the support array will take care of males that weren't present that day.
for(i in 1:ncol(census.list[[1]])){
	
#want to start the loop for the stat after the first interaction is complete, but I want to start the accounting of the past interactions from the beginning

#Remove the "fights" and "nothings" from the census list. This creates attack/receive and win/lose pairs in order of occurence
fight.times<-grep("fight",win.loss.census[[k]][1:(j-1),i])
nothing.times<-grep("nothing",win.loss.census[[k]][1:(j-1),i])

pairs<-win.loss.census[[k]][1:(j-1),i][-c(fight.times, nothing.times)]

#need a way to do this that matches interaction partners but also is sensitive to the time-ordering 

attack.losses<-vector()

if(length(pairs)>0){

attack.losses<-unlist(sapply(1:ncol(census.list[[1]]), function(z){
	
	intset<-pairs[sapply(pairs, function(x) {
		
		strsplit(x,split=".",fixed=T)[[1]][3]==z
		
	})]
	
	names(intset)<-grep("TRUE",sapply(pairs, function(x) {
		
		strsplit(x,split=".",fixed=T)[[1]][3]==z
		
	}))
	
	attacks<-grep("attack",intset)
	attackpluslose<-intset[attacks+1][grep("lose", intset[attacks+1])]
	
	return(attackpluslose)

}))
}


#if the male has not had any interactions yet, the hazard is equal across event types involving them, so set all to zero 
if(length(attack.losses)>0){

#incoming attacks directed at the focal male
stat[j,][endsWith(event.types,paste(".attack.", i, sep=""))]<-length(attack.losses)
	
		
}
	
}

#exclude attack events involving the X males 
stat[j,][endsWith(event.types,paste(".attack.",ncol(census.list[[1]]), sep="")) | startsWith(event.types,paste(ncol(census.list[[1]]),".attack.", sep=""))]<-0

}

#take the log 
stat<-log(1+stat)

new.stat[[k]][[1]]<-abind(new.stat[[k]][[1]], stat)

dimnames(new.stat[[k]][[1]])[[3]][ncol(new.stat[[k]][[1]][1,,])]<-"log.attack.lose.weight.day"

}

return(new.stat)

}


## Am I more liekly to be attacked in general is I have had more attack+wins?

add.log.attack.win.weight.day<-function(x){

new.stat<-x

library(abind)

#loop through each day sequence
for(k in day){

#empty matrix of the right dimensions, event# x event type. In this case, the default is all zeroes. 
stat<-matrix(0, nrow=step, ncol=length(event.types))
colnames(stat)<-event.types


#this loops through each event in the sequence

for(j in step){

#for each event, loop through all the males to get the recency ranks that will predict the next interaction- the support array will take care of males that weren't present that day.
for(i in 1:ncol(census.list[[1]])){
	
#want to start the loop for the stat after the first interaction is complete, but I want to start the accounting of the past interactions from the beginning

#Remove the "fights" and "nothings" from the census list. This creates attack/receive and win/lose pairs in order of occurence
fight.times<-grep("fight",win.loss.census[[k]][1:(j-1),i])
nothing.times<-grep("nothing",win.loss.census[[k]][1:(j-1),i])

pairs<-win.loss.census[[k]][1:(j-1),i][-c(fight.times, nothing.times)]

#need a way to do this that matches interaction partners but also is sensitive to the time-ordering 

attack.wins<-vector()

if(length(pairs)>0){

attack.wins<-unlist(sapply(1:ncol(census.list[[1]]), function(z){
	
	intset<-pairs[sapply(pairs, function(x) {
		
		strsplit(x,split=".",fixed=T)[[1]][3]==z
		
	})]
	
	names(intset)<-grep("TRUE",sapply(pairs, function(x) {
		
		strsplit(x,split=".",fixed=T)[[1]][3]==z
		
	}))
	
	attacks<-grep("attack",intset)
	attackpluswin<-intset[attacks+1][grep("win", intset[attacks+1])]
	
	return(attackpluswin)

}))
}


#if the male has not had any interactions yet, the hazard is equal across event types involving them, so set all to zero 
if(length(attack.wins)>0){

#incoming attacks directed at the focal male
stat[j,][endsWith(event.types,paste(".attack.", i, sep=""))]<-length(attack.wins)
	
		
}
	
}

#exclude attack events involving the X males 
stat[j,][endsWith(event.types,paste(".attack.",ncol(census.list[[1]]), sep="")) | startsWith(event.types,paste(ncol(census.list[[1]]),".attack.", sep=""))]<-0

}

#take the log 
stat<-log(1+stat)

new.stat[[k]][[1]]<-abind(new.stat[[k]][[1]], stat)

dimnames(new.stat[[k]][[1]])[[3]][ncol(new.stat[[k]][[1]][1,,])]<-"log.attack.win.weight.day"

}

return(new.stat)

}


## Am I more liekly to be attacked in general is I have had more receive+wins?


add.log.receive.win.weight.day<-function(x){

new.stat<-x

library(abind)

#loop through each day sequence
for(k in day){

#empty matrix of the right dimensions, event# x event type. In this case, the default is all zeroes. 
stat<-matrix(0, nrow=step, ncol=length(event.types))
colnames(stat)<-event.types


#this loops through each event in the sequence

for(j in step){

#for each event, loop through all the males to get the recency ranks that will predict the next interaction- the support array will take care of males that weren't present that day.
for(i in 1:ncol(census.list[[1]])){
	
#want to start the loop for the stat after the first interaction is complete, but I want to start the accounting of the past interactions from the beginning

#Remove the "fights" and "nothings" from the census list. This creates attack/receive and win/lose pairs in order of occurence
fight.times<-grep("fight",win.loss.census[[k]][1:(j-1),i])
nothing.times<-grep("nothing",win.loss.census[[k]][1:(j-1),i])

pairs<-win.loss.census[[k]][1:(j-1),i][-c(fight.times, nothing.times)]

#need a way to do this that matches interaction partners but also is sensitive to the time-ordering 

receive.wins<-vector()

if(length(pairs)>0){

receive.wins<-unlist(sapply(1:ncol(census.list[[1]]), function(z){
	
	intset<-pairs[sapply(pairs, function(x) {
		
		strsplit(x,split=".",fixed=T)[[1]][3]==z
		
	})]
	
	names(intset)<-grep("TRUE",sapply(pairs, function(x) {
		
		strsplit(x,split=".",fixed=T)[[1]][3]==z
		
	}))
	
	receives<-grep("receive",intset)
	receivepluswin<-intset[receives+1][grep("win", intset[receives+1])]
	
	return(receivepluswin)

}))
}


#if the male has not had any interactions yet, the hazard is equal across event types involving them, so set all to zero 
if(length(receive.wins)>0){

#incoming attacks directed at the focal male
stat[j,][endsWith(event.types,paste(".attack.", i, sep=""))]<-length(receive.wins)
	
		
}
	
}

#exclude attack events involving the X males 
stat[j,][endsWith(event.types,paste(".attack.",ncol(census.list[[1]]), sep="")) | startsWith(event.types,paste(ncol(census.list[[1]]),".attack.", sep=""))]<-0

}

#take the log 
stat<-log(1+stat)

new.stat[[k]][[1]]<-abind(new.stat[[k]][[1]], stat)

dimnames(new.stat[[k]][[1]])[[3]][ncol(new.stat[[k]][[1]][1,,])]<-"log.receive.win.weight.day"

}

return(new.stat)

}


## Am I more liekly to be attacked in general is I have had more receive+losses?


add.log.receive.lose.weight.day<-function(x){

new.stat<-x

library(abind)

#loop through each day sequence
for(k in day){

#empty matrix of the right dimensions, event# x event type. In this case, the default is all zeroes. 
stat<-matrix(0, nrow=step, ncol=length(event.types))
colnames(stat)<-event.types


#this loops through each event in the sequence

for(j in step){

#for each event, loop through all the males to get the recency ranks that will predict the next interaction- the support array will take care of males that weren't present that day.
for(i in 1:ncol(census.list[[1]])){
	
#want to start the loop for the stat after the first interaction is complete, but I want to start the accounting of the past interactions from the beginning

#Remove the "fights" and "nothings" from the census list. This creates attack/receive and win/lose pairs in order of occurence
fight.times<-grep("fight",win.loss.census[[k]][1:(j-1),i])
nothing.times<-grep("nothing",win.loss.census[[k]][1:(j-1),i])

pairs<-win.loss.census[[k]][1:(j-1),i][-c(fight.times, nothing.times)]

#need a way to do this that matches interaction partners but also is sensitive to the time-ordering 

receive.lose<-vector()

if(length(pairs)>0){

receive.lose<-unlist(sapply(1:ncol(census.list[[1]]), function(z){
	
	intset<-pairs[sapply(pairs, function(x) {
		
		strsplit(x,split=".",fixed=T)[[1]][3]==z
		
	})]
	
	names(intset)<-grep("TRUE",sapply(pairs, function(x) {
		
		strsplit(x,split=".",fixed=T)[[1]][3]==z
		
	}))
	
	receives<-grep("receive",intset)
	receivepluslose<-intset[receives+1][grep("lose", intset[receives+1])]
	
	return(receivepluslose)

}))
}


#if the male has not had any interactions yet, the hazard is equal across event types involving them, so set all to zero 
if(length(receive.lose)>0){

#incoming attacks directed at the focal male
stat[j,][endsWith(event.types,paste(".attack.", i, sep=""))]<-length(receive.lose)
	
		
}
	
}

#exclude attack events involving the X males 
stat[j,][endsWith(event.types,paste(".attack.",ncol(census.list[[1]]), sep="")) | startsWith(event.types,paste(ncol(census.list[[1]]),".attack.", sep=""))]<-0

}

#take the log 
stat<-log(1+stat)

new.stat[[k]][[1]]<-abind(new.stat[[k]][[1]], stat)

dimnames(new.stat[[k]][[1]])[[3]][ncol(new.stat[[k]][[1]][1,,])]<-"log.receive.lose.weight.day"

}

return(new.stat)

}


## Am I more likely to be attacked in general if my win/loss rate so far today is greater?

add.win.loss.rate.day<-function(x){
	
rec.stat<-x

library(abind)

#loop through each day sequence
for(k in day){

#empty matrix of the right dimensions, event# x event type. In this case, the default is all zeroes. 
stat<-matrix(0, nrow=step, ncol=length(event.types))
colnames(stat)<-event.types


#this loops through each event in the sequence

for(j in step){

#for each event, loop through all the males to get the previous attack count that will predict the next interaction- the support array will take care of males that weren't present that day.
for(i in 1:ncol(census.list[[1]])){
	

#list of all the wins by the focal male
wins<-length(grep("win",win.loss.census[[k]][1:(j-1),i]))

#all the losses
losses<-length(grep("lose",win.loss.census[[k]][1:(j-1),i]))

#parameterization robust to males with zero wins/losses
rate<-(wins+0.5)/(wins+losses+1)

#match with event types- this stat predicts general INCOMING attacks based on the win/loss rate so far that day by the focal male
stat[j,][endsWith(event.types,paste(".attack.",i, sep=""))]<-rate


}

#exclude attack events involving the X males 
stat[j,][endsWith(event.types,paste(".attack.",ncol(census.list[[1]]), sep="")) | startsWith(event.types,paste(ncol(census.list[[1]]),".attack.", sep=""))]<-0.5


}

rec.stat[[k]][[1]]<-abind(rec.stat[[k]][[1]], stat)

dimnames(rec.stat[[k]][[1]])[[3]][ncol(rec.stat[[k]][[1]][1,,])]<-"win.loss.rate.day"

}

return(rec.stat)


}


##last day-level test:

## Am I more likely to be attacked if I have been chased at all today? 

add.chased.day<-function(x){
	
rec.stat<-x

library(abind)

#loop through each day sequence
for(k in day){

#empty matrix of the right dimensions, event# x event type. In this case, the default is all zeroes. 
stat<-matrix(0, nrow=step, ncol=length(event.types))
colnames(stat)<-event.types


#this loops through each event in the sequence

for(j in step){

#for each event, loop through all the males to get the previous attack count that will predict the next interaction- the support array will take care of males that weren't present that day.
for(i in 1:ncol(census.list[[1]])){
	

#list of all the chases directed at the focal male
chases<-grep("chasedby",win.loss.census[[k]][1:(j-1),i])

#if the male has not had any interactions yet, the hazard is equal across event types involving them, so set all to zero

if(length(chases)>0){

#match with event types- this stat predicts general INCOMING attacks based on whether or not the focal male has been chased at all prevously that day
stat[j,][endsWith(event.types,paste(".attack.",i, sep=""))]<-1

}

}

#exclude attack events involving the X males 
stat[j,][endsWith(event.types,paste(".attack.",ncol(census.list[[1]]), sep="")) | startsWith(event.types,paste(ncol(census.list[[1]]),".attack.", sep=""))]<-0

}


rec.stat[[k]][[1]]<-abind(rec.stat[[k]][[1]], stat)

dimnames(rec.stat[[k]][[1]])[[3]][ncol(rec.stat[[k]][[1]][1,,])]<-"chased.day"

}

return(rec.stat)
	
	
}


################### Pairwise Effects From Yesterday ##################

## Are you more or less likely to attack me today if we interacted at all yesterday? 

add.int.pairwise.yest<-function(x){
	
rec.stat<-x

library(abind)


#loop through each day in the sequence, but applying 0 stat matrices to day 1 and 9 since there is no "yesterday" (March 27th data is missing due to field conditions)

for(k in day){

#empty matrix of the right dimensions, event# x event type. In this case, the default is all zeroes. 
stat<-matrix(0, nrow=step, ncol=length(event.types))
colnames(stat)<-event.types


if(k!=1 && mod.day!=1 && mod.day!=9){

#this loops through each event in the sequence

for(j in step){

#for each event, loop through all the males to get the previous attack count that will predict the next interaction- the support array will take care of males that weren't present that day.
for(i in 1:ncol(census.list[[1]])){
	
#list of all the attacks directed at the focal male YESTERDAY
receives<-grep("received",census.list[[(k-1)]][,i])

attacks<-grep("attack",census.list[[(k-1)]][,i])

ints<-union(receives, attacks)

#if the male has not had any yesterday, the hazard is equal across event types involving them, so set all to zero 
if(length(ints)>0){
	
int.names<-census.list[[(k-1)]][,i][ints]

#get just the interacting male id numbers
intids<-sapply(int.names, function(x) strsplit(x, split=".", fixed=T)[[1]][3])

#collapse to unique parters
intids<-unique(intids)


for(l in 1:length(intids)){

#incoming attacks directed at the focal male, the hazrd is proportional to the number of previous outgoing attacks
stat[j,][paste(intids[l],".attack.", i, sep="")]<-1
	}

#exclude attack events involving the X males 
stat[j,][endsWith(event.types,paste(".attack.",ncol(census.list[[1]]), sep="")) | startsWith(event.types,paste(ncol(census.list[[1]]),".attack.", sep=""))]<-0
		
}


}

}

}

rec.stat[[k]][[1]]<-abind(rec.stat[[k]][[1]], stat)

dimnames(rec.stat[[k]][[1]])[[3]][ncol(rec.stat[[k]][[1]][1,,])]<-"int.pairwise.yest"

}

return(rec.stat)
	
}


## Are you more or less likely to attack me today if we had more total interactions YESTERDAY?

add.log.count.pairwise.yest<-function(x){
	
rec.stat<-x

library(abind)

#loop through each day in the sequence, but applying 0 stat matrices to day 1 and 9 since there is no "yesterday" (March 27th data is missing due to field conditions)

for(k in day){

#empty matrix of the right dimensions, event# x event type. In this case, the default is all zeroes. 
stat<-matrix(0, nrow=step, ncol=length(event.types))
colnames(stat)<-event.types


if(k!=1 && mod.day!=1 && mod.day!=9){

#this loops through each event in the sequence

for(j in step){

#for each event, loop through all the males to get the previous attack count that will predict the next interaction- the support array will take care of males that weren't present that day.
for(i in 1:ncol(census.list[[1]])){
	
#list of all the attacks directed at the focal male YESTERDAY
receives<-grep("received",census.list[[(k-1)]][,i])

attacks<-grep("attack",census.list[[(k-1)]][,i])


#if the male has not had any interactions yet, the hazard is equal across event types involving them, so set all to zero 
if(length(attacks)>0){

#totals for each type- these will be the values for the stat
attack.tots<-table(census.list[[k-1]][,i][attacks])

#get just the interacting male id numbers
rankids<-sapply(names(attack.tots), function(x) strsplit(x, split=".", fixed=T)[[1]][3])

#match with event types- this stat predicts INCOMING attacks based on total previous (yesterday's) INCOMING attacks
#loop thru each element of rankids

for(l in 1:length(rankids)){

#incoming attacks directed at the focal male, the hazrd is proportional to the number of previous outgoing attacks
stat[j,][paste(rankids[l],".attack.", i, sep="")]<-attack.tots[l]
	}

		
}

if(length(receives)>0){

#totals for each type- these will be the values for the stat
receive.tots<-table(census.list[[k-1]][,i][receives])

#get just the interacting male id numbers
rankids<-sapply(names(receive.tots), function(x) strsplit(x, split=".", fixed=T)[[1]][3])

#match with event types- this stat predicts INCOMING attacks based on total previous (yesterday's) pairwise edgeweight
#loop thru each element of rankids

for(l in 1:length(rankids)){

#incoming attacks directed at the focal male, the hazrd is proportional to the number of previous outgoing attacks
stat[j,][paste(rankids[l],".attack.", i, sep="")]<-stat[j,][paste(rankids[l],".attack.", i, sep="")]+receive.tots[l]
	}
}	

}

}
}

stat[,(endsWith(event.types,paste(".attack.",ncol(census.list[[1]]), sep="")) | startsWith(event.types,paste(ncol(census.list[[1]]),".attack.", sep="")))]<-0

stat<-log(1+stat)

rec.stat[[k]][[1]]<-abind(rec.stat[[k]][[1]], stat)

dimnames(rec.stat[[k]][[1]])[[3]][ncol(rec.stat[[k]][[1]][1,,])]<-"log.count.pairwise.yest"

}

return(rec.stat)
	
}


## Are you more or less likely to attack me today if I attacked you more YESTERDAY?

add.log.attack.count.pairwise.yest<-function(x){
	
rec.stat<-x

library(abind)

#loop through each day in the sequence, but applying 0 stat matrices to day 1 and 9 since there is no "yesterday" (March 27th data is missing due to field conditions)

for(k in day){

#empty matrix of the right dimensions, event# x event type. In this case, the default is all zeroes. 
stat<-matrix(0, nrow=step, ncol=length(event.types))
colnames(stat)<-event.types


if(k!=1 && mod.day!=1 && mod.day!=9){

#this loops through each event in the sequence

for(j in step){

#for each event, loop through all the males to get the previous attack count that will predict the next interaction- the support array will take care of males that weren't present that day.
for(i in 1:ncol(census.list[[1]])){
	
#list of all the attacks initiated by the focal male YESTERDAY

attacks<-grep("attack",census.list[[(k-1)]][,i])


#if the male has not had any interactions yet, the hazard is equal across event types involving them, so set all to zero 
if(length(attacks)>0){

#totals for each type- these will be the values for the stat
attack.tots<-table(census.list[[k-1]][,i][attacks])

#get just the interacting male id numbers
rankids<-sapply(names(attack.tots), function(x) strsplit(x, split=".", fixed=T)[[1]][3])

#match with event types- this stat predicts INCOMING attacks based on total previous (yesterday's) INCOMING attacks
#loop thru each element of rankids

for(l in 1:length(rankids)){

#incoming attacks directed at the focal male, the hazrd is proportional to the number of previous outgoing attacks
stat[j,][paste(rankids[l],".attack.", i, sep="")]<-attack.tots[l]
	}

		
}

	
}

}
}

stat[,(endsWith(event.types,paste(".attack.",ncol(census.list[[1]]), sep="")) | startsWith(event.types,paste(ncol(census.list[[1]]),".attack.", sep="")))]<-0

stat<-log(1+stat)

rec.stat[[k]][[1]]<-abind(rec.stat[[k]][[1]], stat)

dimnames(rec.stat[[k]][[1]])[[3]][ncol(rec.stat[[k]][[1]][1,,])]<-"log.attack.count.pairwise.yest"

}

return(rec.stat)
	
}


## Are you more or less likely to attack me today if I was attacked by you more YESTERDAY?

add.log.receive.count.pairwise.yest<-function(x){
	
rec.stat<-x

library(abind)


#loop through each day in the sequence, but applying 0 stat matrices to day 1 and 9 since there is no "yesterday" (March 27th data is missing due to field conditions)

for(k in day){

#empty matrix of the right dimensions, event# x event type. In this case, the default is all zeroes. 
stat<-matrix(0, nrow=step, ncol=length(event.types))
colnames(stat)<-event.types


if(k!=1 && mod.day!=1 && mod.day!=9){


#this loops through each event in the sequence

for(j in step){

#for each event, loop through all the males to get the previous attack count that will predict the next interaction- the support array will take care of males that weren't present that day.
for(i in 1:ncol(census.list[[1]])){
	
#list of all the attacks directed at the focal male YESTERDAY
receives<-grep("received",census.list[[(k-1)]][,i])


#if the male has not had any interactions yet, the hazard is equal across event types involving them, so set all to zero 

if(length(receives)>0){

#totals for each type- these will be the values for the stat
receive.tots<-table(census.list[[k-1]][,i][receives])

#get just the interacting male id numbers
rankids<-sapply(names(receive.tots), function(x) strsplit(x, split=".", fixed=T)[[1]][3])

#match with event types- this stat predicts INCOMING attacks based on total previous (yesterday's) pairwise edgeweight
#loop thru each element of rankids

for(l in 1:length(rankids)){

#incoming attacks directed at the focal male, the hazrd is proportional to the number of previous outgoing attacks
stat[j,][paste(rankids[l],".attack.", i, sep="")]<-receive.tots[l]
	}
}	
}

}
}

stat[,(endsWith(event.types,paste(".attack.",ncol(census.list[[1]]), sep="")) | startsWith(event.types,paste(ncol(census.list[[1]]),".attack.", sep="")))]<-0

stat<-log(1+stat)

rec.stat[[k]][[1]]<-abind(rec.stat[[k]][[1]], stat)

dimnames(rec.stat[[k]][[1]])[[3]][ncol(rec.stat[[k]][[1]][1,,])]<-"log.receive.count.pairwise.yest"

}

return(rec.stat)
	
}


## Am I more likely to be attacked by you if I lost to you more times yesterday?

add.log.loss.count.pairwise.yest<-function(x){
	
rec.stat<-x

library(abind)

#loop through each day in the sequence, but applying 0 stat matrices to day 1 and 9 since there is no "yesterday" (March 27th data is missing due to field conditions)

for(k in day){

#empty matrix of the right dimensions, event# x event type. In this case, the default is all zeroes. 
stat<-matrix(0, nrow=step, ncol=length(event.types))
colnames(stat)<-event.types


if(k!=1 && mod.day!=1 && mod.day!=9){

#this loops through each event in the sequence

for(j in step){

#for each event, loop through all the males to get the previous attack count that will predict the next interaction- the support array will take care of males that weren't present that day.
for(i in 1:ncol(census.list[[1]])){
	
#list of all the instances the focal male lost YESTERDAY
losses<-win.loss.census[[(k-1)]][,i][grep("lose",win.loss.census[[(k-1)]][,i])]

#if the male has not had any interactions yet, the hazard is equal across event types involving them, so set all to zero 
if(length(losses)>0){

#strip the "chase" from the end of chase wins so I can include them
losses<-sapply(losses, function(x) if(grepl("chas",x)==TRUE){
temp<-strsplit(x, split=".", fixed=T)[[1]]	
new<-paste(temp[1],temp[2],temp[3], sep=".")
return(new)
}
else return(x)
)


#totals for each type- these will be the values for the stat
loss.tots<-table(losses)

#get just the interacting male id numbers
rankids<-sapply(names(loss.tots), function(x) strsplit(x, split=".", fixed=T)[[1]][3])

#match with event types- this stat predicts INCOMING attacks based on total previous (yesterday's) losses
#loop thru each element of rankids

for(l in 1:length(rankids)){

#incoming attacks directed at the focal male
stat[j,][paste(rankids[l],".attack.", i, sep="")]<-loss.tots[l]
	}

		
}
	
}

}
}

stat[,(endsWith(event.types,paste(".attack.",ncol(census.list[[1]]), sep="")) | startsWith(event.types,paste(ncol(census.list[[1]]),".attack.", sep="")))]<-0

stat<-log(1+stat)

rec.stat[[k]][[1]]<-abind(rec.stat[[k]][[1]], stat)

dimnames(rec.stat[[k]][[1]])[[3]][ncol(rec.stat[[k]][[1]][1,,])]<-"log.loss.count.pairwise.yest"

}

return(rec.stat)
	
}


## Am I more likely to be attacked by you if I won against you more times yesterday?

add.log.win.count.pairwise.yest<-function(x){
	
rec.stat<-x

library(abind)


#loop through each day in the sequence, but applying 0 stat matrices to day 1 and 9 since there is no "yesterday" (March 27th data is missing due to field conditions)

for(k in day){

#empty matrix of the right dimensions, event# x event type. In this case, the default is all zeroes. 
stat<-matrix(0, nrow=step, ncol=length(event.types))
colnames(stat)<-event.types


if(k!=1 && mod.day!=1 && mod.day!=9){

#this loops through each event in the sequence

for(j in step){

#for each event, loop through all the males to get the previous attack count that will predict the next interaction- the support array will take care of males that weren't present that day.
for(i in 1:ncol(census.list[[1]])){
	
#list of all the instances the focal male won YESTERDAY
losses<-win.loss.census[[(k-1)]][,i][grep("win",win.loss.census[[(k-1)]][,i])]

#if the male has not had any interactions yet, the hazard is equal across event types involving them, so set all to zero 
if(length(losses)>0){

#strip the "chase" from the end of chase wins so I can include them
losses<-sapply(losses, function(x) if(grepl("chas",x)==TRUE){
temp<-strsplit(x, split=".", fixed=T)[[1]]	
new<-paste(temp[1],temp[2],temp[3], sep=".")
return(new)
}
else return(x)
)


#totals for each type- these will be the values for the stat
loss.tots<-table(losses)

#get just the interacting male id numbers
rankids<-sapply(names(loss.tots), function(x) strsplit(x, split=".", fixed=T)[[1]][3])

#match with event types- this stat predicts INCOMING attacks based on total previous (yesterday's) losses
#loop thru each element of rankids

for(l in 1:length(rankids)){

#incoming attacks directed at the focal male
stat[j,][paste(rankids[l],".attack.", i, sep="")]<-loss.tots[l]
	}

		
}
	
}

}
}

stat[,(endsWith(event.types,paste(".attack.",ncol(census.list[[1]]), sep="")) | startsWith(event.types,paste(ncol(census.list[[1]]),".attack.", sep="")))]<-0

stat<-log(1+stat)

rec.stat[[k]][[1]]<-abind(rec.stat[[k]][[1]], stat)

dimnames(rec.stat[[k]][[1]])[[3]][ncol(rec.stat[[k]][[1]][1,,])]<-"log.win.count.pairwise.yest"

}

return(rec.stat)
	
}


## Are you more likely to attack me if I had more attack+lose with you yesterday?


add.log.attack.lose.pairwise.yest<-function(x){

new.stat<-x

library(abind)


#loop through each day in the sequence, but applying 0 stat matrices to day 1 and 9 since there is no "yesterday" (March 27th data is missing due to field conditions)


for(k in day){

#empty matrix of the right dimensions, event# x event type. In this case, the default is all zeroes. 
stat<-matrix(0, nrow=step, ncol=length(event.types))
colnames(stat)<-event.types

if(k!=1 && mod.day!=1 && mod.day!=9){
	
#this loops through each event in the sequence, in this case starting with the third event since this is the earliest possible win/loss event

for(j in step){

#for each event, loop through all the males to get the recency ranks that will predict the next interaction- the support array will take care of males that weren't present that day.
for(i in 1:ncol(census.list[[1]])){
	
#get the list of all the interactions for the focal male for yesterday

int.vector<-win.loss.census[[(k-1)]][,i]


#Remove the "fights" and "nothings" from the census list. This creates attack/receive and win/lose pairs in order of occurence
fight.times<-grep("fight",int.vector)
nothing.times<-grep("nothing",int.vector)

pairs<-int.vector[-c(fight.times, nothing.times)]


#need a way to do this that matches interaction partners but also is sensitive to the time-ordering 

attack.lose<-vector()

if(length(pairs)>0){

attack.lose<-unlist(sapply(1:ncol(census.list[[1]]), function(z){
	
	intset<-pairs[sapply(pairs, function(x) {
		
		strsplit(x,split=".",fixed=T)[[1]][3]==z
		
	})]
	
	names(intset)<-grep("TRUE",sapply(pairs, function(x) {
		
		strsplit(x,split=".",fixed=T)[[1]][3]==z
		
	}))
	
	attacks<-grep("attack",intset)
	attackpluslose<-intset[attacks+1][grep("lose", intset[attacks+1])]
	
	return(attackpluslose)

}))
}

#if the male has not had any interactions yet, the hazard is equal across event types involving them, so set all to zero 
if(length(attack.lose)>0){

#strip the "chase" from the end of chase wins so I can include them
attack.lose<-sapply(attack.lose, function(x) if(grepl("chas",x)==TRUE){
temp<-strsplit(x, split=".", fixed=T)[[1]]	
new<-paste(temp[1],temp[2],temp[3], sep=".")
return(new)
}
else return(x)
)


#totals for each type- these will be the values for the stat
tots<-table(attack.lose)

#get just the interacting male id numbers
rankids<-sapply(names(tots), function(x) strsplit(x, split=".", fixed=T)[[1]][3])

#match with event types- this stat predicts INCOMING attacks based on total previous (yesterday's) attack+losses
#loop thru each element of rankids

for(l in 1:length(rankids)){

#incoming attacks directed at the focal male
stat[j,][paste(rankids[l],".attack.", i, sep="")]<-tots[l]
	}

		
}
	
}

}
}

stat[,(endsWith(event.types,paste(".attack.",ncol(census.list[[1]]), sep="")) | startsWith(event.types,paste(ncol(census.list[[1]]),".attack.", sep="")))]<-0

#take the log 
stat<-log(1+stat)

new.stat[[k]][[1]]<-abind(new.stat[[k]][[1]], stat)

dimnames(new.stat[[k]][[1]])[[3]][ncol(new.stat[[k]][[1]][1,,])]<-"log.attack.lose.pairwise.yest"

}

return(new.stat)

}


## Are you more likely to attack me if I had more attack+wins with you yesterday?

add.log.attack.win.pairwise.yest<-function(x){

new.stat<-x

library(abind)


#loop through each day in the sequence, but applying 0 stat matrices to day 1 and 9 since there is no "yesterday" (March 27th data is missing due to field conditions)


for(k in day){

#empty matrix of the right dimensions, event# x event type. In this case, the default is all zeroes. 
stat<-matrix(0, nrow=step, ncol=length(event.types))
colnames(stat)<-event.types

if(k!=1 && mod.day!=1 && mod.day!=9){
	
#this loops through each event in the sequence, in this case starting with the third event since this is the earliest possible win/loss event

for(j in step){

#for each event, loop through all the males to get the recency ranks that will predict the next interaction- the support array will take care of males that weren't present that day.
for(i in 1:ncol(census.list[[1]])){
	
#get the list of all the interactions for the focal male for yesterday

int.vector<-win.loss.census[[(k-1)]][,i]


#Remove the "fights" and "nothings" from the census list. This creates attack/receive and win/lose pairs in order of occurence
fight.times<-grep("fight",int.vector)
nothing.times<-grep("nothing",int.vector)

pairs<-int.vector[-c(fight.times, nothing.times)]


#need a way to do this that matches interaction partners but also is sensitive to the time-ordering 

attack.win<-vector()

if(length(pairs)>0){

attack.win<-unlist(sapply(1:ncol(census.list[[1]]), function(z){
	
	intset<-pairs[sapply(pairs, function(x) {
		
		strsplit(x,split=".",fixed=T)[[1]][3]==z
		
	})]
	
	names(intset)<-grep("TRUE",sapply(pairs, function(x) {
		
		strsplit(x,split=".",fixed=T)[[1]][3]==z
		
	}))
	
	attacks<-grep("attack",intset)
	attackpluswin<-intset[attacks+1][grep("win", intset[attacks+1])]
	
	return(attackpluswin)

}))
}

#if the male has not had any interactions yet, the hazard is equal across event types involving them, so set all to zero 
if(length(attack.win)>0){

#strip the "chase" from the end of chase wins so I can include them
attack.win<-sapply(attack.win, function(x) if(grepl("chas",x)==TRUE){
temp<-strsplit(x, split=".", fixed=T)[[1]]	
new<-paste(temp[1],temp[2],temp[3], sep=".")
return(new)
}
else return(x)
)


#totals for each type- these will be the values for the stat
tots<-table(attack.win)

#get just the interacting male id numbers
rankids<-sapply(names(tots), function(x) strsplit(x, split=".", fixed=T)[[1]][3])

#match with event types- this stat predicts INCOMING attacks based on total previous (yesterday's) attack+losses
#loop thru each element of rankids

for(l in 1:length(rankids)){

#incoming attacks directed at the focal male
stat[j,][paste(rankids[l],".attack.", i, sep="")]<-tots[l]
	}

		
}
	
}

}
}

stat[,(endsWith(event.types,paste(".attack.",ncol(census.list[[1]]), sep="")) | startsWith(event.types,paste(ncol(census.list[[1]]),".attack.", sep="")))]<-0

#take the log 
stat<-log(1+stat)

new.stat[[k]][[1]]<-abind(new.stat[[k]][[1]], stat)

dimnames(new.stat[[k]][[1]])[[3]][ncol(new.stat[[k]][[1]][1,,])]<-"log.attack.win.pairwise.yest"

}

return(new.stat)

}


## Are you more likely to attack me if I had more receive+wins with you yesterday?


add.log.receive.win.pairwise.yest<-function(x){

new.stat<-x

library(abind)


#loop through each day in the sequence, but applying 0 stat matrices to day 1 and 9 since there is no "yesterday" (March 27th data is missing due to field conditions)


for(k in day){

#empty matrix of the right dimensions, event# x event type. In this case, the default is all zeroes. 
stat<-matrix(0, nrow=step, ncol=length(event.types))
colnames(stat)<-event.types

if(k!=1 && mod.day!=1 && mod.day!=9){
	
#this loops through each event in the sequence, in this case starting with the third event since this is the earliest possible win/loss event

for(j in step){

#for each event, loop through all the males to get the recency ranks that will predict the next interaction- the support array will take care of males that weren't present that day.
for(i in 1:ncol(census.list[[1]])){
	
#get the list of all the interactions for the focal male for yesterday

int.vector<-win.loss.census[[(k-1)]][,i]


#Remove the "fights" and "nothings" from the census list. This creates attack/receive and win/lose pairs in order of occurence
fight.times<-grep("fight",int.vector)
nothing.times<-grep("nothing",int.vector)

pairs<-int.vector[-c(fight.times, nothing.times)]


#need a way to do this that matches interaction partners but also is sensitive to the time-ordering 

receive.win<-vector()

if(length(pairs)>0){

receive.win<-unlist(sapply(1:ncol(census.list[[1]]), function(z){
	
	intset<-pairs[sapply(pairs, function(x) {
		
		strsplit(x,split=".",fixed=T)[[1]][3]==z
		
	})]
	
	names(intset)<-grep("TRUE",sapply(pairs, function(x) {
		
		strsplit(x,split=".",fixed=T)[[1]][3]==z
		
	}))
	
	receive<-grep("receive",intset)
	receivepluswin<-intset[receive+1][grep("win", intset[receive+1])]
	
	return(receivepluswin)

}))
}

#if the male has not had any interactions yet, the hazard is equal across event types involving them, so set all to zero 
if(length(receive.win)>0){

#strip the "chase" from the end of chase wins so I can include them
receive.win<-sapply(receive.win, function(x) if(grepl("chas",x)==TRUE){
temp<-strsplit(x, split=".", fixed=T)[[1]]	
new<-paste(temp[1],temp[2],temp[3], sep=".")
return(new)
}
else return(x)
)


#totals for each type- these will be the values for the stat
tots<-table(receive.win)

#get just the interacting male id numbers
rankids<-sapply(names(tots), function(x) strsplit(x, split=".", fixed=T)[[1]][3])

#match with event types- this stat predicts INCOMING attacks based on total previous (yesterday's) attack+losses
#loop thru each element of rankids

for(l in 1:length(rankids)){

#incoming attacks directed at the focal male
stat[j,][paste(rankids[l],".attack.", i, sep="")]<-tots[l]
	}

		
}
	
}

}
}

stat[,(endsWith(event.types,paste(".attack.",ncol(census.list[[1]]), sep="")) | startsWith(event.types,paste(ncol(census.list[[1]]),".attack.", sep="")))]<-0

#take the log 
stat<-log(1+stat)

new.stat[[k]][[1]]<-abind(new.stat[[k]][[1]], stat)

dimnames(new.stat[[k]][[1]])[[3]][ncol(new.stat[[k]][[1]][1,,])]<-"log.receive.win.pairwise.yest"

}

return(new.stat)

}


## Are you more likely to attack me if I had more receive+losses with you yesterday?


add.log.receive.lose.pairwise.yest<-function(x){

new.stat<-x

library(abind)


#loop through each day in the sequence, but applying 0 stat matrices to day 1 and 9 since there is no "yesterday" (March 27th data is missing due to field conditions)


for(k in day){

#empty matrix of the right dimensions, event# x event type. In this case, the default is all zeroes. 
stat<-matrix(0, nrow=step, ncol=length(event.types))
colnames(stat)<-event.types

if(k!=1 && mod.day!=1 && mod.day!=9){
	
#this loops through each event in the sequence, in this case starting with the third event since this is the earliest possible win/loss event

for(j in step){

#for each event, loop through all the males to get the recency ranks that will predict the next interaction- the support array will take care of males that weren't present that day.
for(i in 1:ncol(census.list[[1]])){
	
#get the list of all the interactions for the focal male for yesterday

int.vector<-win.loss.census[[(k-1)]][,i]


#Remove the "fights" and "nothings" from the census list. This creates attack/receive and win/lose pairs in order of occurence
fight.times<-grep("fight",int.vector)
nothing.times<-grep("nothing",int.vector)

pairs<-int.vector[-c(fight.times, nothing.times)]


#need a way to do this that matches interaction partners but also is sensitive to the time-ordering 

receive.lose<-vector()

if(length(pairs)>0){

receive.lose<-unlist(sapply(1:ncol(census.list[[1]]), function(z){
	
	intset<-pairs[sapply(pairs, function(x) {
		
		strsplit(x,split=".",fixed=T)[[1]][3]==z
		
	})]
	
	names(intset)<-grep("TRUE",sapply(pairs, function(x) {
		
		strsplit(x,split=".",fixed=T)[[1]][3]==z
		
	}))
	
	receive<-grep("receive",intset)
	receivepluslose<-intset[receive+1][grep("lose", intset[receive+1])]
	
	return(receivepluslose)

}))
}

#if the male has not had any interactions yet, the hazard is equal across event types involving them, so set all to zero 
if(length(receive.lose)>0){

#strip the "chase" from the end of chase wins so I can include them
receive.lose<-sapply(receive.lose, function(x) if(grepl("chas",x)==TRUE){
temp<-strsplit(x, split=".", fixed=T)[[1]]	
new<-paste(temp[1],temp[2],temp[3], sep=".")
return(new)
}
else return(x)
)


#totals for each type- these will be the values for the stat
tots<-table(receive.lose)

#get just the interacting male id numbers
rankids<-sapply(names(tots), function(x) strsplit(x, split=".", fixed=T)[[1]][3])

#match with event types- this stat predicts INCOMING attacks based on total previous (yesterday's) attack+losses
#loop thru each element of rankids

for(l in 1:length(rankids)){

#incoming attacks directed at the focal male
stat[j,][paste(rankids[l],".attack.", i, sep="")]<-tots[l]
	}

		
}
	
}

}
}

stat[,(endsWith(event.types,paste(".attack.",ncol(census.list[[1]]), sep="")) | startsWith(event.types,paste(ncol(census.list[[1]]),".attack.", sep="")))]<-0

#take the log 
stat<-log(1+stat)

new.stat[[k]][[1]]<-abind(new.stat[[k]][[1]], stat)

dimnames(new.stat[[k]][[1]])[[3]][ncol(new.stat[[k]][[1]][1,,])]<-"log.receive.lose.pairwise.yest"

}

return(new.stat)

}


##Are you more likely to attack me if I had a greater win/loss rate against you yesterday?


add.win.loss.rate.pairwise.yest<-function(x){
	
rec.stat<-x

library(abind)

#loop through each day sequence
for(k in day){

#empty matrix of the right dimensions, event# x event type. In this case, the default is all zeroes. 
stat<-matrix(0, nrow=step, ncol=length(event.types))
colnames(stat)<-event.types

if(k!=1 && mod.day!=1 && mod.day!=9){

#this loops through each event in the sequence

for(j in step){

#for each event, loop through all the males to get the previous attack count that will predict the next interaction- the support array will take care of males that weren't present that day.
for(i in 1:ncol(census.list[[1]])){
	
#start with simple +/-

#list of all the instances the focal male won YESTERDAY
wins<-win.loss.census[[(k-1)]][,i][grep("win",win.loss.census[[(k-1)]][,i])]

#list of all the instances the focal male lost YESTERDAY
losses<-win.loss.census[[(k-1)]][,i][grep("lose",win.loss.census[[(k-1)]][,i])]

#if the male has not had any interactions yet, the hazard is equal across event types involving them, so set all to zero
#do wins first 
if(length(wins)>0){

#strip the "chase" from the end of chase wins so I can include them
wins<-sapply(wins, function(x) if(grepl("chas",x)==TRUE){
temp<-strsplit(x, split=".", fixed=T)[[1]]	
new<-paste(temp[1],temp[2],temp[3], sep=".")
return(new)
}
else return(x)
)


#totals for each type- these will be the values for the stat
win.tots<-table(wins)

#get just the interacting male id numbers
winids<-sapply(names(win.tots), function(x) strsplit(x, split=".", fixed=T)[[1]][3])

allwins<-rep(0,ncol(census.list[[1]]))
 
for(l in 1:length(winids)){
	
allwins[as.numeric(winids[l])]<-win.tots[l]
}

} else {
	allwins<-rep(0,ncol(census.list[[1]]))
	}


#now substract losses if there are any
if(length(losses)>0){

#strip the "chase" from the end of chase wins so I can include them
losses<-sapply(losses, function(x) if(grepl("chas",x)==TRUE){
temp<-strsplit(x, split=".", fixed=T)[[1]]	
new<-paste(temp[1],temp[2],temp[3], sep=".")
return(new)
}
else return(x)
)
	
	
lose.tots<-table(losses)
	
loseids<-sapply(names(lose.tots), function(x) strsplit(x, split=".", fixed=T)[[1]][3])

allloss<-rep(0,ncol(census.list[[1]]))
 
for(l in 1:length(loseids)){
	
allloss[as.numeric(loseids[l])]<-lose.tots[l]
}


} else {
	
	allloss<-rep(0,ncol(census.list[[1]]))
	
}


#now I can make a vector of all the win/loss rates with each other male in order

rates<-(allwins+0.5)/(allwins+allloss+1)

#get rid of the win rate for the focal male against itself
rates<-rates[-i]

#match with incoming attacks directed at the focal male

#incoming attacks directed at the focal male
stat[j,][event.types[endsWith(event.types,paste0(".attack.",i))]]<-rates

}

}
} else {
	
	#if it is day 1 or 9, everyone gets the baseline stat value of 0.5
	stat[,grep("attack",event.types)]<-0.5
	
}

#exclude X males
stat[,(endsWith(event.types,paste(".attack.",ncol(census.list[[1]]), sep="")) | startsWith(event.types,paste(ncol(census.list[[1]]),".attack.", sep="")))]<-0.5

rec.stat[[k]][[1]]<-abind(rec.stat[[k]][[1]], stat)

dimnames(rec.stat[[k]][[1]])[[3]][ncol(rec.stat[[k]][[1]][1,,])]<-"win.loss.rate.pairwise.yest"

}

return(rec.stat)

}


########## General Effects From Yesterday ###########


### Explore yesterday effects of edgeweight

## Am I more/less likely to be attacked if I was involved in any interactions yesterday?

add.int.yest<-function(x){
	
rec.stat<-x

library(abind)

#loop through each day in the sequence, but applying 0 stat matrices to day 1 and 9 since there is no "yesterday" (March 27th data is missing due to field conditions)

for(k in day){

#empty matrix of the right dimensions, event# x event type. In this case, the default is all zeroes. 
stat<-matrix(0, nrow=step, ncol=length(event.types))
colnames(stat)<-event.types


if(k!=1 && mod.day!=1 && mod.day!=9){

#this loops through each event in the sequence

for(j in step){

#for each event, loop through all the males to get the previous attack count that will predict the next interaction- the support array will take care of males that weren't present that day.
for(i in 1:ncol(census.list[[1]])){
	

#list of all the attacks by the focal male
attacks<-grep("attack",win.loss.census[[k-1]][,i])

receives<-grep("received",win.loss.census[[k-1]][,i])

ints<-union(attacks, receives)

#if the male has not had any interactions yet, the hazard is equal across event types involving them, so set all to zero

if(length(ints)>0){

#match with event types- this stat predicts general INCOMING attacks based on if the focal male interacted yesterday
stat[j,][endsWith(event.types,paste(".attack.",i, sep=""))]<-1

}

}

}
}

#exclude X males
stat[,(endsWith(event.types,paste(".attack.",ncol(census.list[[1]]), sep="")) | startsWith(event.types,paste(ncol(census.list[[1]]),".attack.", sep="")))]<-0

rec.stat[[k]][[1]]<-abind(rec.stat[[k]][[1]], stat)

dimnames(rec.stat[[k]][[1]])[[3]][ncol(rec.stat[[k]][[1]][1,,])]<-"int.yest"

}

return(rec.stat)


}


## Am I more/less likely to be attacked if I was involved in a large number of total interactions yesterday?

add.log.weight.yest<-function(x){
	
rec.stat<-x

library(abind)

#loop through each day in the sequence, but applying 0 stat matrices to day 1 and 9 since there is no "yesterday" (March 27th data is missing due to field conditions)

for(k in day){

#empty matrix of the right dimensions, event# x event type. In this case, the default is all zeroes. 
stat<-matrix(0, nrow=step, ncol=length(event.types))
colnames(stat)<-event.types


if(k!=1 && mod.day!=1 && mod.day!=9){

#this loops through each event in the sequence

for(j in step){

#for each event, loop through all the males to get the previous attack count that will predict the next interaction- the support array will take care of males that weren't present that day.
for(i in 1:ncol(census.list[[1]])){
	

#list of all the attacks by the focal male
attacks<-grep("attack",win.loss.census[[k-1]][,i])

receives<-grep("received",win.loss.census[[k-1]][,i])

ints<-union(attacks, receives)

#if the male has not had any interactions yet, the hazard is equal across event types involving them, so set all to zero

if(length(ints)>0){

#match with event types- this stat predicts general INCOMING attacks based on the total number of interactions had yesterday by the focal male
stat[j,][endsWith(event.types,paste(".attack.",i, sep=""))]<-length(ints)

}

}

}
}


#exclude X males
stat[,(endsWith(event.types,paste(".attack.",ncol(census.list[[1]]), sep="")) | startsWith(event.types,paste(ncol(census.list[[1]]),".attack.", sep="")))]<-0


stat<-log(1+stat)

rec.stat[[k]][[1]]<-abind(rec.stat[[k]][[1]], stat)

dimnames(rec.stat[[k]][[1]])[[3]][ncol(rec.stat[[k]][[1]][1,,])]<-"log.weight.yest"

}

return(rec.stat)


}


## Am I more/less likely to be attacked if I inititated a lot of attacks yesterday?

add.log.attack.weight.yest<-function(x){
	
rec.stat<-x

library(abind)


#loop through each day in the sequence, but applying 0 stat matrices to day 1 and 9 since there is no "yesterday" (March 27th data is missing due to field conditions)

for(k in day){

#empty matrix of the right dimensions, event# x event type. In this case, the default is all zeroes. 
stat<-matrix(0, nrow=step, ncol=length(event.types))
colnames(stat)<-event.types


if(k!=1 && mod.day!=1 && mod.day!=9){

#this loops through each event in the sequence

for(j in step){

#for each event, loop through all the males to get the previous attack count that will predict the next interaction- the support array will take care of males that weren't present that day.
for(i in 1:ncol(census.list[[1]])){
	

#list of all the attacks by the focal male
attacks<-grep("attack",win.loss.census[[k-1]][,i])


#if the male has not had any interactions yet, the hazard is equal across event types involving them, so set all to zero

if(length(attacks)>0){

#match with event types- this stat predicts general INCOMING attacks based on the total number of interactions had yesterday by the focal male
stat[j,][endsWith(event.types,paste(".attack.",i, sep=""))]<-length(attacks)

}

}

}
}

#exclude X males
stat[,(endsWith(event.types,paste(".attack.",ncol(census.list[[1]]), sep="")) | startsWith(event.types,paste(ncol(census.list[[1]]),".attack.", sep="")))]<-0


stat<-log(1+stat)

rec.stat[[k]][[1]]<-abind(rec.stat[[k]][[1]], stat)

dimnames(rec.stat[[k]][[1]])[[3]][ncol(rec.stat[[k]][[1]][1,,])]<-"log.attack.weight.yest"

}

return(rec.stat)


}


## Am I more/less likely to be attacked if I was attacked more times yesterday?

add.log.receive.weight.yest<-function(x){
	
rec.stat<-x

library(abind)

#loop through each day in the sequence, but applying 0 stat matrices to day 1 and 9 since there is no "yesterday" (March 27th data is missing due to field conditions)

for(k in day){

#empty matrix of the right dimensions, event# x event type. In this case, the default is all zeroes. 
stat<-matrix(0, nrow=step, ncol=length(event.types))
colnames(stat)<-event.types


if(k!=1 && mod.day!=1 && mod.day!=9){

#this loops through each event in the sequence

for(j in step){

#for each event, loop through all the males to get the previous attack count that will predict the next interaction- the support array will take care of males that weren't present that day.
for(i in 1:ncol(census.list[[1]])){
	

#list of all the attacks by the focal male


receives<-grep("received",win.loss.census[[k-1]][,i])


#if the male has not had any interactions yet, the hazard is equal across event types involving them, so set all to zero

if(length(receives)>0){

#match with event types- this stat predicts general INCOMING attacks based on the total number of interactions had yesterday by the focal male
stat[j,][endsWith(event.types,paste(".attack.",i, sep=""))]<-length(receives)

}

}

}

}


#exclude X males
stat[,(endsWith(event.types,paste(".attack.",ncol(census.list[[1]]), sep="")) | startsWith(event.types,paste(ncol(census.list[[1]]),".attack.", sep="")))]<-0


stat<-log(1+stat)

rec.stat[[k]][[1]]<-abind(rec.stat[[k]][[1]], stat)

dimnames(rec.stat[[k]][[1]])[[3]][ncol(rec.stat[[k]][[1]][1,,])]<-"log.receive.weight.yest"

}

return(rec.stat)


}


## Am I more likely to be attacked in general if my total number of wins yesterday is greater?


add.log.win.weight.yest<-function(x){
	
rec.stat<-x

library(abind)


#loop through each day in the sequence, but applying 0 stat matrices to day 1 and 9 since there is no "yesterday" (March 27th data is missing due to field conditions)

for(k in day){

#empty matrix of the right dimensions, event# x event type. In this case, the default is all zeroes. 
stat<-matrix(0, nrow=step, ncol=length(event.types))
colnames(stat)<-event.types


if(k!=1 && mod.day!=1 && mod.day!=9){

#this loops through each event in the sequence, in this case starting with the fourth event since this stat requires at least one previous win/loss to have taken place

for(j in step){

#for each event, loop through all the males to get the previous attack count that will predict the next interaction- the support array will take care of males that weren't present that day.
for(i in 1:ncol(census.list[[1]])){
	

#list of all the attacks by the focal male
wins<-grep("win",win.loss.census[[(k-1)]][,i])

#if the male has not had any interactions yet, the hazard is equal across event types involving them, so set all to zero

if(length(wins)>0){

#match with event types- this stat predicts general INCOMING attacks based on the total number of attacks so far that day by the focal male
stat[j,][endsWith(event.types,paste(".attack.",i, sep=""))]<-length(wins)

}

}

}
}

#exclude X males
stat[,(endsWith(event.types,paste(".attack.",ncol(census.list[[1]]), sep="")) | startsWith(event.types,paste(ncol(census.list[[1]]),".attack.", sep="")))]<-0


stat<-log(1+stat)

rec.stat[[k]][[1]]<-abind(rec.stat[[k]][[1]], stat)

dimnames(rec.stat[[k]][[1]])[[3]][ncol(rec.stat[[k]][[1]][1,,])]<-"log.win.weight.yest"

}

return(rec.stat)


}


## Am I more likely to be attacked in general if my total number of losses yesterday is greater?

add.log.lose.weight.yest<-function(x){
	
rec.stat<-x

library(abind)


#loop through each day in the sequence, but applying 0 stat matrices to day 1 and 9 since there is no "yesterday" (March 27th data is missing due to field conditions)

for(k in day){

#empty matrix of the right dimensions, event# x event type. In this case, the default is all zeroes. 
stat<-matrix(0, nrow=step, ncol=length(event.types))
colnames(stat)<-event.types


if(k!=1 && mod.day!=1 && mod.day!=9){


#this loops through each event in the sequence, in this case starting with the fourth event since this stat requires at least one previous win/loss to have taken place

for(j in step){

#for each event, loop through all the males to get the previous attack count that will predict the next interaction- the support array will take care of males that weren't present that day.
for(i in 1:ncol(census.list[[1]])){
	

#list of all the attacks by the focal male
wins<-grep("lose",win.loss.census[[(k-1)]][,i])

#if the male has not had any interactions yet, the hazard is equal across event types involving them, so set all to zero

if(length(wins)>0){

#match with event types- this stat predicts general INCOMING attacks based on the total number of attacks so far that day by the focal male
stat[j,][endsWith(event.types,paste(".attack.",i, sep=""))]<-length(wins)

}

}

}
}


#exclude X males
stat[,(endsWith(event.types,paste(".attack.",ncol(census.list[[1]]), sep="")) | startsWith(event.types,paste(ncol(census.list[[1]]),".attack.", sep="")))]<-0


stat<-log(1+stat)

rec.stat[[k]][[1]]<-abind(rec.stat[[k]][[1]], stat)

dimnames(rec.stat[[k]][[1]])[[3]][ncol(rec.stat[[k]][[1]][1,,])]<-"log.lose.weight.yest"

}

return(rec.stat)


}


## Am I more likely to be attacked in general if my total number of receive+losses yesterday is greater?


add.log.receive.lose.weight.yest<-function(x){

new.stat<-x

library(abind)

#loop through each day in the sequence, but applying 0 stat matrices to day 1 and 9 since there is no "yesterday" (March 27th data is missing due to field conditions)

for(k in day){

#empty matrix of the right dimensions, event# x event type. In this case, the default is all zeroes. 
stat<-matrix(0, nrow=step, ncol=length(event.types))
colnames(stat)<-event.types


if(k!=1 && mod.day!=1 && mod.day!=9){

#this loops through each event in the sequence, in this case starting with the third event since this is the earliest possible win/loss event

for(j in step){

#for each event, loop through all the males to get the recency ranks that will predict the next interaction- the support array will take care of males that weren't present that day.
for(i in 1:ncol(census.list[[1]])){
	
#want to start the loop for the stat after the first interaction is complete, but I want to start the accounting of the past interactions from the beginning

int.vector<-win.loss.census[[(k-1)]][,i]

#Remove the "fights" and "nothings" from the census list. This creates attack/receive and win/lose pairs in order of occurence
fight.times<-grep("fight",int.vector)
nothing.times<-grep("nothing",int.vector)

pairs<-int.vector[-c(fight.times, nothing.times)]

#need a way to do this that matches interaction partners but also is sensitive to the time-ordering 

receive.lose<-vector()

if(length(pairs)>0){

receive.lose<-unlist(sapply(1:ncol(census.list[[1]]), function(z){
	
	intset<-pairs[sapply(pairs, function(x) {
		
		strsplit(x,split=".",fixed=T)[[1]][3]==z
		
	})]
	
	names(intset)<-grep("TRUE",sapply(pairs, function(x) {
		
		strsplit(x,split=".",fixed=T)[[1]][3]==z
		
	}))
	
	receive<-grep("receive",intset)
	receivepluslose<-intset[receive+1][grep("lose", intset[receive+1])]
	
	return(receivepluslose)

}))
}


#if the male has not had any interactions yet, the hazard is equal across event types involving them, so set all to zero 
if(length(receive.lose)>0){

#incoming attacks directed at the focal male
stat[j,][endsWith(event.types,paste(".attack.", i, sep=""))]<-length(receive.lose)
	
		
}
	
}

}
}


#exclude X males
stat[,(endsWith(event.types,paste(".attack.",ncol(census.list[[1]]), sep="")) | startsWith(event.types,paste(ncol(census.list[[1]]),".attack.", sep="")))]<-0


#take the log 
stat<-log(1+stat)

new.stat[[k]][[1]]<-abind(new.stat[[k]][[1]], stat)

dimnames(new.stat[[k]][[1]])[[3]][ncol(new.stat[[k]][[1]][1,,])]<-"log.receive.lose.weight.yest"

}

return(new.stat)

}


## Am I more likely to be attacked in general if my total number of receive+wins yesterday is greater?


add.log.receive.win.weight.yest<-function(x){

new.stat<-x

library(abind)

#loop through each day in the sequence, but applying 0 stat matrices to day 1 and 9 since there is no "yesterday" (March 27th data is missing due to field conditions)

for(k in day){

#empty matrix of the right dimensions, event# x event type. In this case, the default is all zeroes. 
stat<-matrix(0, nrow=step, ncol=length(event.types))
colnames(stat)<-event.types


if(k!=1 && mod.day!=1 && mod.day!=9){

#this loops through each event in the sequence, in this case starting with the third event since this is the earliest possible win/loss event

for(j in step){

#for each event, loop through all the males to get the recency ranks that will predict the next interaction- the support array will take care of males that weren't present that day.
for(i in 1:ncol(census.list[[1]])){
	
#want to start the loop for the stat after the first interaction is complete, but I want to start the accounting of the past interactions from the beginning

int.vector<-win.loss.census[[(k-1)]][,i]

#Remove the "fights" and "nothings" from the census list. This creates attack/receive and win/lose pairs in order of occurence
fight.times<-grep("fight",int.vector)
nothing.times<-grep("nothing",int.vector)

pairs<-int.vector[-c(fight.times, nothing.times)]

#need a way to do this that matches interaction partners but also is sensitive to the time-ordering 

receive.win<-vector()

if(length(pairs)>0){

receive.win<-unlist(sapply(1:ncol(census.list[[1]]), function(z){
	
	intset<-pairs[sapply(pairs, function(x) {
		
		strsplit(x,split=".",fixed=T)[[1]][3]==z
		
	})]
	
	names(intset)<-grep("TRUE",sapply(pairs, function(x) {
		
		strsplit(x,split=".",fixed=T)[[1]][3]==z
		
	}))
	
	receive<-grep("receive",intset)
	receivepluswin<-intset[receive+1][grep("win", intset[receive+1])]
	
	return(receivepluswin)

}))
}


#if the male has not had any interactions yet, the hazard is equal across event types involving them, so set all to zero 
if(length(receive.win)>0){

#incoming attacks directed at the focal male
stat[j,][endsWith(event.types,paste(".attack.", i, sep=""))]<-length(receive.win)
	
		
}
	
}

}
}


#exclude X males
stat[,(endsWith(event.types,paste(".attack.",ncol(census.list[[1]]), sep="")) | startsWith(event.types,paste(ncol(census.list[[1]]),".attack.", sep="")))]<-0


#take the log 
stat<-log(1+stat)

new.stat[[k]][[1]]<-abind(new.stat[[k]][[1]], stat)

dimnames(new.stat[[k]][[1]])[[3]][ncol(new.stat[[k]][[1]][1,,])]<-"log.receive.win.weight.yest"

}

return(new.stat)

}


## Am I more likely to be attacked in general if my total number of attack+losses yesterday is greater?


add.log.attack.lose.weight.yest<-function(x){

new.stat<-x

library(abind)

#loop through each day in the sequence, but applying 0 stat matrices to day 1 and 9 since there is no "yesterday" (March 27th data is missing due to field conditions)

for(k in day){

#empty matrix of the right dimensions, event# x event type. In this case, the default is all zeroes. 
stat<-matrix(0, nrow=step, ncol=length(event.types))
colnames(stat)<-event.types


if(k!=1 && mod.day!=1 && mod.day!=9){

#this loops through each event in the sequence, in this case starting with the third event since this is the earliest possible win/loss event

for(j in step){

#for each event, loop through all the males to get the recency ranks that will predict the next interaction- the support array will take care of males that weren't present that day.
for(i in 1:ncol(census.list[[1]])){
	
#want to start the loop for the stat after the first interaction is complete, but I want to start the accounting of the past interactions from the beginning

int.vector<-win.loss.census[[(k-1)]][,i]

#Remove the "fights" and "nothings" from the census list. This creates attack/receive and win/lose pairs in order of occurence
fight.times<-grep("fight",int.vector)
nothing.times<-grep("nothing",int.vector)

pairs<-int.vector[-c(fight.times, nothing.times)]

#need a way to do this that matches interaction partners but also is sensitive to the time-ordering 

attack.lose<-vector()

if(length(pairs)>0){

attack.lose<-unlist(sapply(1:ncol(census.list[[1]]), function(z){
	
	intset<-pairs[sapply(pairs, function(x) {
		
		strsplit(x,split=".",fixed=T)[[1]][3]==z
		
	})]
	
	names(intset)<-grep("TRUE",sapply(pairs, function(x) {
		
		strsplit(x,split=".",fixed=T)[[1]][3]==z
		
	}))
	
	attack<-grep("attack",intset)
	attackpluslose<-intset[attack+1][grep("lose", intset[attack+1])]
	
	return(attackpluslose)

}))
}


#if the male has not had any interactions yet, the hazard is equal across event types involving them, so set all to zero 
if(length(attack.lose)>0){

#incoming attacks directed at the focal male
stat[j,][endsWith(event.types,paste(".attack.", i, sep=""))]<-length(attack.lose)
	
		
}
	
}

}
}

#exclude X males
stat[,(endsWith(event.types,paste(".attack.",ncol(census.list[[1]]), sep="")) | startsWith(event.types,paste(ncol(census.list[[1]]),".attack.", sep="")))]<-0


#take the log 
stat<-log(1+stat)

new.stat[[k]][[1]]<-abind(new.stat[[k]][[1]], stat)

dimnames(new.stat[[k]][[1]])[[3]][ncol(new.stat[[k]][[1]][1,,])]<-"log.attack.lose.weight.yest"

}

return(new.stat)

}


## Am I more likely to be attacked in general if my total number of attack+wins yesterday is greater?


add.log.attack.win.weight.yest<-function(x){

new.stat<-x

library(abind)

#loop through each day in the sequence, but applying 0 stat matrices to day 1 and 9 since there is no "yesterday" (March 27th data is missing due to field conditions)

for(k in day){

#empty matrix of the right dimensions, event# x event type. In this case, the default is all zeroes. 
stat<-matrix(0, nrow=step, ncol=length(event.types))
colnames(stat)<-event.types


if(k!=1 && mod.day!=1 && mod.day!=9){

#this loops through each event in the sequence, in this case starting with the third event since this is the earliest possible win/loss event

for(j in step){

#for each event, loop through all the males to get the recency ranks that will predict the next interaction- the support array will take care of males that weren't present that day.
for(i in 1:ncol(census.list[[1]])){
	
#want to start the loop for the stat after the first interaction is complete, but I want to start the accounting of the past interactions from the beginning

int.vector<-win.loss.census[[(k-1)]][,i]

#Remove the "fights" and "nothings" from the census list. This creates attack/receive and win/lose pairs in order of occurence
fight.times<-grep("fight",int.vector)
nothing.times<-grep("nothing",int.vector)

pairs<-int.vector[-c(fight.times, nothing.times)]

#need a way to do this that matches interaction partners but also is sensitive to the time-ordering 

attack.win<-vector()

if(length(pairs)>0){

attack.win<-unlist(sapply(1:ncol(census.list[[1]]), function(z){
	
	intset<-pairs[sapply(pairs, function(x) {
		
		strsplit(x,split=".",fixed=T)[[1]][3]==z
		
	})]
	
	names(intset)<-grep("TRUE",sapply(pairs, function(x) {
		
		strsplit(x,split=".",fixed=T)[[1]][3]==z
		
	}))
	
	attack<-grep("attack",intset)
	attackpluswin<-intset[attack+1][grep("win", intset[attack+1])]
	
	return(attackpluswin)

}))
}


#if the male has not had any interactions yet, the hazard is equal across event types involving them, so set all to zero 
if(length(attack.win)>0){

#incoming attacks directed at the focal male
stat[j,][endsWith(event.types,paste(".attack.", i, sep=""))]<-length(attack.win)
	
		
}
	
}

}
}

#exclude X males
stat[,(endsWith(event.types,paste(".attack.",ncol(census.list[[1]]), sep="")) | startsWith(event.types,paste(ncol(census.list[[1]]),".attack.", sep="")))]<-0


#take the log 
stat<-log(1+stat)

new.stat[[k]][[1]]<-abind(new.stat[[k]][[1]], stat)

dimnames(new.stat[[k]][[1]])[[3]][ncol(new.stat[[k]][[1]][1,,])]<-"log.attack.win.weight.yest"

}

return(new.stat)

}


## Am I more likely to be attacked in general if my win/loss rate yesterday was greater?

add.win.loss.rate.yest<-function(x){
	
rec.stat<-x

library(abind)

#loop through each day sequence
for(k in day){

#empty matrix of the right dimensions, event# x event type. In this case, the default is all zeroes. 
stat<-matrix(0, nrow=step, ncol=length(event.types))
colnames(stat)<-event.types


if(k!=1 && mod.day!=1 && mod.day!=9){

#this loops through each event in the sequence

for(j in step){

#for each event, loop through all the males to get the previous attack count that will predict the next interaction- the support array will take care of males that weren't present that day.
for(i in 1:ncol(census.list[[1]])){
	


#list of all the instances the focal male won YESTERDAY
wins<-length(win.loss.census[[(k-1)]][,i][grep("win",win.loss.census[[(k-1)]][,i])])


#list of all the instances the focal male lost YESTERDAY
losses<-length(win.loss.census[[(k-1)]][,i][grep("lose",win.loss.census[[(k-1)]][,i])])

#parameterization robust to males with zero wins/losses
rate<-(wins+0.5)/(wins+losses+1)

#match with event types- this stat predicts general INCOMING attacks based on the win/loss rate so far that day by the focal male
stat[j,][endsWith(event.types,paste(".attack.",i, sep=""))]<-rate


}
}
} else {
	
	#if it is day 1 or 9, everyone gets the baseline stat value of 0.5
	stat[,grep("attack",event.types)]<-0.5
	
}

#exclude X males
stat[,(endsWith(event.types,paste(".attack.",ncol(census.list[[1]]), sep="")) | startsWith(event.types,paste(ncol(census.list[[1]]),".attack.", sep="")))]<-0.5


rec.stat[[k]][[1]]<-abind(rec.stat[[k]][[1]], stat)

dimnames(rec.stat[[k]][[1]])[[3]][ncol(rec.stat[[k]][[1]][1,,])]<-"win.loss.rate.yest"

}

return(rec.stat)


}


### Look at degree metrics for yesterday ###

## Am I more likely to be attacked today if I interacted with log(1+more different other males) yesterday?


add.log.degree.yest<-function(x){
	
rec.stat<-x

library(abind)


#loop through each day in the sequence, but applying 0 stat matrices to day 1 and 9 since there is no "yesterday" (March 27th data is missing due to field conditions)

for(k in day){

#empty matrix of the right dimensions, event# x event type. In this case, the default is all zeroes. 
stat<-matrix(0, nrow=step, ncol=length(event.types))
colnames(stat)<-event.types


if(k!=1 && mod.day!=1 && mod.day!=9){


#this loops through each event in the sequence, in this case starting with the fourth event since this stat requires at least one previous win/loss to have taken place

for(j in step){

#for each event, loop through all the males to get the previous attack count that will predict the next interaction- the support array will take care of males that weren't present that day.
for(i in 1:ncol(census.list[[1]])){
	
#get number of unique interaction partners

#list of all the wins by the focal male
wins<-grep("win",win.loss.census[[(k-1)]][,i])

#list of all the losses by the focal male
losses<-grep("lose",win.loss.census[[(k-1)]][,i])

#this covers all the interactions that happened yesterday for the focal male. Stick them all together
ints<-union(wins,losses)

#if the male has not had any interactions yet, the hazard is equal across event types involving them, so set all to zero
#do wins first 
if(length(ints)>0){

#totals for each type- these will be the values for the stat
int.names<-win.loss.census[[(k-1)]][,i][ints]

#get just the interacting male id numbers
intids<-sapply(int.names, function(x) strsplit(x, split=".", fixed=T)[[1]][3])

#collapse to unique parters
intids<-unique(intids)

#match with event types- this stat predicts general INCOMING attacks based on the total number of interaction partners so far that day for the focal male (degree)

stat[j,][endsWith(event.types,paste(".attack.",i, sep=""))]<-length(intids)
}


}

}
}

#exclude X males
stat[,(endsWith(event.types,paste(".attack.",ncol(census.list[[1]]), sep="")) | startsWith(event.types,paste(ncol(census.list[[1]]),".attack.", sep="")))]<-0


stat<-log(1+stat)

rec.stat[[k]][[1]]<-abind(rec.stat[[k]][[1]], stat)

dimnames(rec.stat[[k]][[1]])[[3]][ncol(rec.stat[[k]][[1]][1,,])]<-"log.degree.yest"

}

return(rec.stat)

}


## Am I more likely to be attacked today if I interacted with more different other males yesterday?

add.degree.yest<-function(x){
	
rec.stat<-x

library(abind)


#loop through each day in the sequence, but applying 0 stat matrices to day 1 and 9 since there is no "yesterday" (March 27th data is missing due to field conditions)

for(k in day){

#empty matrix of the right dimensions, event# x event type. In this case, the default is all zeroes. 
stat<-matrix(0, nrow=step, ncol=length(event.types))
colnames(stat)<-event.types

if(k!=1 && mod.day!=1 && mod.day!=9){
	
#this loops through each event in the sequence, in this case starting with the fourth event since this stat requires at least one previous win/loss to have taken place

for(j in step){

#for each event, loop through all the males to get the previous attack count that will predict the next interaction- the support array will take care of males that weren't present that day.
for(i in 1:ncol(census.list[[1]])){
	
#get number of unique interaction partners

#list of all the wins by the focal male
wins<-grep("win",win.loss.census[[(k-1)]][,i])

#list of all the losses by the focal male
losses<-grep("lose",win.loss.census[[(k-1)]][,i])

#this covers all the interactions that happened yesterday for the focal male. Stick them all together
ints<-union(wins,losses)

#if the male has not had any interactions yet, the hazard is equal across event types involving them, so set all to zero
#do wins first 
if(length(ints)>0){

#totals for each type- these will be the values for the stat
int.names<-win.loss.census[[(k-1)]][,i][ints]

#get just the interacting male id numbers
intids<-sapply(int.names, function(x) strsplit(x, split=".", fixed=T)[[1]][3])

#collapse to unique parters
intids<-unique(intids)

#match with event types- this stat predicts general INCOMING attacks based on the total number of interaction partners so far that day for the focal male (degree)

stat[j,][endsWith(event.types,paste(".attack.",i, sep=""))]<-length(intids)
}


}

}

}

#exclude X males
stat[,(endsWith(event.types,paste(".attack.",ncol(census.list[[1]]), sep="")) | startsWith(event.types,paste(ncol(census.list[[1]]),".attack.", sep="")))]<-0


rec.stat[[k]][[1]]<-abind(rec.stat[[k]][[1]], stat)

dimnames(rec.stat[[k]][[1]])[[3]][ncol(rec.stat[[k]][[1]][1,,])]<-"degree.yest"

}

return(rec.stat)

}


## Am I more likely to be attacked today if I have a winning record with more different other males yesterday?


add.win.degree.yest<-function(x){
	
rec.stat<-x

library(abind)

#loop through each day in the sequence, but applying 0 stat matrices to day 1 and 9 since there is no "yesterday" (March 27th data is missing due to field conditions)

for(k in day){

#empty matrix of the right dimensions, event# x event type. In this case, the default is all zeroes. 
stat<-matrix(0, nrow=step, ncol=length(event.types))
colnames(stat)<-event.types


if(k!=1 && mod.day!=1 && mod.day!=9){
	
#this loops through each event in the sequence
for(j in step){

#for each event, loop through all the males to get the previous attack count that will predict the next interaction- the support array will take care of males that weren't present that day.
for(i in 1:ncol(census.list[[1]])){
	
#start with simple +/-

#list of all the wins by the focal male
wins<-grep("win",win.loss.census[[(k-1)]][,i])

#list of all the losses by the focal male
losses<-grep("lose",win.loss.census[[(k-1)]][,i])

win.record<-rep(0,ncol(census.list[[1]]))

#if the male has not had any interactions yet, the hazard is equal across event types involving them, so set all to zero
#do wins first 
if(length(wins)>0){

wins<-win.loss.census[[(k-1)]][,i][wins]

#strip the "chase" from the end of chase wins so I can include them
wins<-sapply(wins, function(x) if(grepl("chas",x)==TRUE){
temp<-strsplit(x, split=".", fixed=T)[[1]]	
new<-paste(temp[1],temp[2],temp[3], sep=".")
return(new)
}
else return(x)
)

#totals for each type
win.tots<-table(wins)

#get just the interacting male id numbers
winids<-sapply(names(win.tots), function(x) strsplit(x, split=".", fixed=T)[[1]][3])

win.record[as.numeric(winids)]<-win.tots
}


#now substract losses if there are any
if(length(losses)>0){

losses<-win.loss.census[[(k-1)]][,i][losses]

#strip the "chase" from the end of chase wins so I can include them
losses<-sapply(losses, function(x) if(grepl("chas",x)==TRUE){
temp<-strsplit(x, split=".", fixed=T)[[1]]	
new<-paste(temp[1],temp[2],temp[3], sep=".")
return(new)
}
else return(x)
)
	
	
lose.tots<-table(losses)
	
loseids<-sapply(names(lose.tots), function(x) strsplit(x, split=".", fixed=T)[[1]][3])

win.record[as.numeric(loseids)]<-win.record[as.numeric(loseids)]-lose.tots

} 
 
#test if the win records are positive
win.pos<-win.record>0 

#Get the total number of males with whom the focal male has positive win records 
if(is.na(table(win.pos)["TRUE"])){win.pos.num<-0}
else{win.pos.num<-table(win.pos)["TRUE"]}

#match with event types- this stat predicts general INCOMING attacks based on the total number of interaction partners the focal male has positive win records with (win degree) from yesterday

stat[j,][endsWith(event.types,paste(".attack.",i, sep=""))]<-win.pos.num

}

}
}

#exclude X males
stat[,(endsWith(event.types,paste(".attack.",ncol(census.list[[1]]), sep="")) | startsWith(event.types,paste(ncol(census.list[[1]]),".attack.", sep="")))]<-0


rec.stat[[k]][[1]]<-abind(rec.stat[[k]][[1]], stat)

dimnames(rec.stat[[k]][[1]])[[3]][ncol(rec.stat[[k]][[1]][1,,])]<-"win.degree.yest"

}

return(rec.stat)
	
	
}


## Am I more likely to be attacked today if I have a winning record with log(1+more males) yesterday?

add.log.win.degree.yest<-function(x){
	
rec.stat<-x

library(abind)

#loop through each day in the sequence, but applying 0 stat matrices to day 1 and 9 since there is no "yesterday" (March 27th data is missing due to field conditions)

for(k in day){

#empty matrix of the right dimensions, event# x event type. In this case, the default is all zeroes. 
stat<-matrix(0, nrow=step, ncol=length(event.types))
colnames(stat)<-event.types


if(k!=1 && mod.day!=1 && mod.day!=9){

#this loops through each event in the sequence
for(j in step){

#for each event, loop through all the males to get the previous attack count that will predict the next interaction- the support array will take care of males that weren't present that day.
for(i in 1:ncol(census.list[[1]])){
	
#start with simple +/-

#list of all the wins by the focal male
wins<-grep("win",win.loss.census[[(k-1)]][,i])

#list of all the losses by the focal male
losses<-grep("lose",win.loss.census[[(k-1)]][,i])

win.record<-rep(0,ncol(census.list[[1]]))

#if the male has not had any interactions yet, the hazard is equal across event types involving them, so set all to zero
#do wins first 
if(length(wins)>0){

wins<-win.loss.census[[(k-1)]][,i][wins]

#strip the "chase" from the end of chase wins so I can include them
wins<-sapply(wins, function(x) if(grepl("chas",x)==TRUE){
temp<-strsplit(x, split=".", fixed=T)[[1]]	
new<-paste(temp[1],temp[2],temp[3], sep=".")
return(new)
}
else return(x)
)

#totals for each type
win.tots<-table(wins)

#get just the interacting male id numbers
winids<-sapply(names(win.tots), function(x) strsplit(x, split=".", fixed=T)[[1]][3])

win.record[as.numeric(winids)]<-win.tots
}


#now substract losses if there are any
if(length(losses)>0){

losses<-win.loss.census[[(k-1)]][,i][losses]

#strip the "chase" from the end of chase wins so I can include them
losses<-sapply(losses, function(x) if(grepl("chas",x)==TRUE){
temp<-strsplit(x, split=".", fixed=T)[[1]]	
new<-paste(temp[1],temp[2],temp[3], sep=".")
return(new)
}
else return(x)
)
	
	
lose.tots<-table(losses)
	
loseids<-sapply(names(lose.tots), function(x) strsplit(x, split=".", fixed=T)[[1]][3])

win.record[as.numeric(loseids)]<-win.record[as.numeric(loseids)]-lose.tots

} 
 
#test if the win records are positive
win.pos<-win.record>0 

#Get the total number of males with whom the focal male has positive win records 
if(is.na(table(win.pos)["TRUE"])){win.pos.num<-0}
else{win.pos.num<-table(win.pos)["TRUE"]}

#match with event types- this stat predicts general INCOMING attacks based on the total number of interaction partners the focal male has positive win records with (win degree) from yesterday

stat[j,][endsWith(event.types,paste(".attack.",i, sep=""))]<-win.pos.num

}

}
}

#exclude X males
stat[,(endsWith(event.types,paste(".attack.",ncol(census.list[[1]]), sep="")) | startsWith(event.types,paste(ncol(census.list[[1]]),".attack.", sep="")))]<-0


stat<-log(1+stat)

rec.stat[[k]][[1]]<-abind(rec.stat[[k]][[1]], stat)

dimnames(rec.stat[[k]][[1]])[[3]][ncol(rec.stat[[k]][[1]][1,,])]<-"log.win.degree.yest"

}

return(rec.stat)
	
	
}


## Am I more likely to be attacked today if I have a LOSING record with more males yesterday?


add.lose.degree.yest<-function(x){
	
rec.stat<-x

library(abind)

#loop through each day in the sequence, but applying 0 stat matrices to day 1 and 9 since there is no "yesterday" (March 27th data is missing due to field conditions)

for(k in day){

#empty matrix of the right dimensions, event# x event type. In this case, the default is all zeroes. 
stat<-matrix(0, nrow=step, ncol=length(event.types))
colnames(stat)<-event.types


if(k!=1 && mod.day!=1 && mod.day!=9){

#this loops through each event in the sequence
for(j in step){

#for each event, loop through all the males to get the previous attack count that will predict the next interaction- the support array will take care of males that weren't present that day.
for(i in 1:ncol(census.list[[1]])){
	
#start with simple +/-

#list of all the wins by the focal male
wins<-grep("win",win.loss.census[[(k-1)]][,i])

#list of all the losses by the focal male
losses<-grep("lose",win.loss.census[[(k-1)]][,i])

win.record<-rep(0,ncol(census.list[[1]]))

#if the male has not had any interactions yet, the hazard is equal across event types involving them, so set all to zero
#do wins first 
if(length(wins)>0){

wins<-win.loss.census[[(k-1)]][,i][wins]

#strip the "chase" from the end of chase wins so I can include them
wins<-sapply(wins, function(x) if(grepl("chas",x)==TRUE){
temp<-strsplit(x, split=".", fixed=T)[[1]]	
new<-paste(temp[1],temp[2],temp[3], sep=".")
return(new)
}
else return(x)
)

#totals for each type
win.tots<-table(wins)

#get just the interacting male id numbers
winids<-sapply(names(win.tots), function(x) strsplit(x, split=".", fixed=T)[[1]][3])

win.record[as.numeric(winids)]<-win.tots
}


#now substract losses if there are any
if(length(losses)>0){

losses<-win.loss.census[[(k-1)]][,i][losses]

#strip the "chase" from the end of chase wins so I can include them
losses<-sapply(losses, function(x) if(grepl("chas",x)==TRUE){
temp<-strsplit(x, split=".", fixed=T)[[1]]	
new<-paste(temp[1],temp[2],temp[3], sep=".")
return(new)
}
else return(x)
)
	
	
lose.tots<-table(losses)
	
loseids<-sapply(names(lose.tots), function(x) strsplit(x, split=".", fixed=T)[[1]][3])

win.record[as.numeric(loseids)]<-win.record[as.numeric(loseids)]-lose.tots

 
for(g in 1:length(win.record)){
	
	if(g %in% as.numeric(loseids)){
	
	win.record[g]<-as.numeric(win.record[g])-lose.tots[loseids==g]
	}
	else{if(win.record[g]==0){win.record[g]<-"no.int"}}
	
}

}

if(length(losses)==0){

win.record[win.record==0]<-"no.int"

}
 
 
#test if the win records are less than zero
win.pos<-win.record<0 

#Get the total number of males with whom the focal male has positive win records 
if(is.na(table(win.pos)["TRUE"])){win.pos.num<-0}
else{win.pos.num<-table(win.pos)["TRUE"]}

#match with event types- this stat predicts general INCOMING attacks based on the total number of interaction partners the focal male has positive win records with (win degree) from yesterday

stat[j,][endsWith(event.types,paste(".attack.",i, sep=""))]<-win.pos.num

}

}
}

#exclude X males
stat[,(endsWith(event.types,paste(".attack.",ncol(census.list[[1]]), sep="")) | startsWith(event.types,paste(ncol(census.list[[1]]),".attack.", sep="")))]<-0


rec.stat[[k]][[1]]<-abind(rec.stat[[k]][[1]], stat)

dimnames(rec.stat[[k]][[1]])[[3]][ncol(rec.stat[[k]][[1]][1,,])]<-"lose.degree.yest"

}

return(rec.stat)
	
	
}


## Am I more likely to be attacked today if I have a LOSING record with log(1+more) males yesterday?


add.log.lose.degree.yest<-function(x){
	
rec.stat<-x

library(abind)

#loop through each day in the sequence, but applying 0 stat matrices to day 1 and 9 since there is no "yesterday" (March 27th data is missing due to field conditions)

for(k in day){

#empty matrix of the right dimensions, event# x event type. In this case, the default is all zeroes. 
stat<-matrix(0, nrow=step, ncol=length(event.types))
colnames(stat)<-event.types


if(k!=1 && mod.day!=1 && mod.day!=9){

#this loops through each event in the sequence
for(j in step){

#for each event, loop through all the males to get the previous attack count that will predict the next interaction- the support array will take care of males that weren't present that day.
for(i in 1:ncol(census.list[[1]])){
	
#start with simple +/-

#list of all the wins by the focal male
wins<-grep("win",win.loss.census[[(k-1)]][,i])

#list of all the losses by the focal male
losses<-grep("lose",win.loss.census[[(k-1)]][,i])

win.record<-rep(0,ncol(census.list[[1]]))

#if the male has not had any interactions yet, the hazard is equal across event types involving them, so set all to zero
#do wins first 
if(length(wins)>0){

wins<-win.loss.census[[(k-1)]][,i][wins]

#strip the "chase" from the end of chase wins so I can include them
wins<-sapply(wins, function(x) if(grepl("chas",x)==TRUE){
temp<-strsplit(x, split=".", fixed=T)[[1]]	
new<-paste(temp[1],temp[2],temp[3], sep=".")
return(new)
}
else return(x)
)

#totals for each type
win.tots<-table(wins)

#get just the interacting male id numbers
winids<-sapply(names(win.tots), function(x) strsplit(x, split=".", fixed=T)[[1]][3])

win.record[as.numeric(winids)]<-win.tots
}


#now substract losses if there are any
if(length(losses)>0){

losses<-win.loss.census[[(k-1)]][,i][losses]

#strip the "chase" from the end of chase wins so I can include them
losses<-sapply(losses, function(x) if(grepl("chas",x)==TRUE){
temp<-strsplit(x, split=".", fixed=T)[[1]]	
new<-paste(temp[1],temp[2],temp[3], sep=".")
return(new)
}
else return(x)
)
	
	
lose.tots<-table(losses)
	
loseids<-sapply(names(lose.tots), function(x) strsplit(x, split=".", fixed=T)[[1]][3])

win.record[as.numeric(loseids)]<-win.record[as.numeric(loseids)]-lose.tots

 
for(g in 1:length(win.record)){
	
	if(g %in% as.numeric(loseids)){
	
	win.record[g]<-as.numeric(win.record[g])-lose.tots[loseids==g]
	}
	else{if(win.record[g]==0){win.record[g]<-"no.int"}}
	
}

}

if(length(losses)==0){

win.record[win.record==0]<-"no.int"

}
 
 
#test if the win records are less than zero
win.pos<-win.record<0 

#Get the total number of males with whom the focal male has positive win records 
if(is.na(table(win.pos)["TRUE"])){win.pos.num<-0}
else{win.pos.num<-table(win.pos)["TRUE"]}

#match with event types- this stat predicts general INCOMING attacks based on the total number of interaction partners the focal male has positive win records with (win degree) from yesterday

stat[j,][endsWith(event.types,paste(".attack.",i, sep=""))]<-win.pos.num

}

}
}

#exclude X males
stat[,(endsWith(event.types,paste(".attack.",ncol(census.list[[1]]), sep="")) | startsWith(event.types,paste(ncol(census.list[[1]]),".attack.", sep="")))]<-0


stat<-log(1+stat)

rec.stat[[k]][[1]]<-abind(rec.stat[[k]][[1]], stat)

dimnames(rec.stat[[k]][[1]])[[3]][ncol(rec.stat[[k]][[1]][1,,])]<-"log.lose.degree.yest"

}

return(rec.stat)
	
	
}


## test "neutral" degree, excluding those males that have a zero win-loss record because they didn't interact


add.neutral.degree.yest<-function(x){
	
rec.stat<-x

library(abind)

#loop through each day in the sequence, but applying 0 stat matrices to day 1 and 9 since there is no "yesterday" (March 27th data is missing due to field conditions)

for(k in day){

#empty matrix of the right dimensions, event# x event type. In this case, the default is all zeroes. 
stat<-matrix(0, nrow=step, ncol=length(event.types))
colnames(stat)<-event.types


if(k!=1 && mod.day!=1 && mod.day!=9){

#this loops through each event in the sequence
for(j in step){

#for each event, loop through all the males to get the previous attack count that will predict the next interaction- the support array will take care of males that weren't present that day.
for(i in 1:ncol(census.list[[1]])){
	
#start with simple +/-

#list of all the wins by the focal male
wins<-grep("win",win.loss.census[[(k-1)]][,i])

#list of all the losses by the focal male
losses<-grep("lose",win.loss.census[[(k-1)]][,i])

win.record<-rep(0,ncol(census.list[[1]]))

#if the male has not had any interactions yet, the hazard is equal across event types involving them, so set all to zero
#do wins first 
if(length(wins)>0){

wins<-win.loss.census[[(k-1)]][,i][wins]

#strip the "chase" from the end of chase wins so I can include them
wins<-sapply(wins, function(x) if(grepl("chas",x)==TRUE){
temp<-strsplit(x, split=".", fixed=T)[[1]]	
new<-paste(temp[1],temp[2],temp[3], sep=".")
return(new)
}
else return(x)
)

#totals for each type
win.tots<-table(wins)

#get just the interacting male id numbers
winids<-sapply(names(win.tots), function(x) strsplit(x, split=".", fixed=T)[[1]][3])

win.record[as.numeric(winids)]<-win.tots
}


#now substract losses if there are any
if(length(losses)>0){

losses<-win.loss.census[[(k-1)]][,i][losses]

#strip the "chase" from the end of chase wins so I can include them
losses<-sapply(losses, function(x) if(grepl("chas",x)==TRUE){
temp<-strsplit(x, split=".", fixed=T)[[1]]	
new<-paste(temp[1],temp[2],temp[3], sep=".")
return(new)
}
else return(x)
)
	
	
lose.tots<-table(losses)
	
loseids<-sapply(names(lose.tots), function(x) strsplit(x, split=".", fixed=T)[[1]][3])

for(g in 1:length(win.record)){
	
	if(g %in% as.numeric(loseids)){
	
	win.record[g]<-as.numeric(win.record[g])-lose.tots[loseids==g]
	}
	else{if(win.record[g]==0){win.record[g]<-"no.int"}}
	
}

}

if(length(losses)==0){

win.record[win.record==0]<-"no.int"

}
 
#test if the win records are strictly negative
win.pos<-win.record==0 

#Get the total number of males with whom the focal male has positive win records 
if(is.na(table(win.pos)["TRUE"])){win.pos.num<-0}
else{win.pos.num<-table(win.pos)["TRUE"]}

#match with event types- this stat predicts general INCOMING attacks based on the total number of interaction partners the focal male has positive win records with (win degree) from yesterday

stat[j,][endsWith(event.types,paste(".attack.",i, sep=""))]<-win.pos.num

}

}
}

#exclude X males
stat[,(endsWith(event.types,paste(".attack.",ncol(census.list[[1]]), sep="")) | startsWith(event.types,paste(ncol(census.list[[1]]),".attack.", sep="")))]<-0


rec.stat[[k]][[1]]<-abind(rec.stat[[k]][[1]], stat)

dimnames(rec.stat[[k]][[1]])[[3]][ncol(rec.stat[[k]][[1]][1,,])]<-"neutral.degree.yest"

}

return(rec.stat)
	
	
}


## log "neutral" degree, excluding those males that have a zero win-loss record because they didn't interact

add.log.neutral.degree.yest<-function(x){
	
rec.stat<-x

library(abind)


#loop through each day in the sequence, but applying 0 stat matrices to day 1 and 9 since there is no "yesterday" (March 27th data is missing due to field conditions)

for(k in day){

#empty matrix of the right dimensions, event# x event type. In this case, the default is all zeroes. 
stat<-matrix(0, nrow=step, ncol=length(event.types))
colnames(stat)<-event.types


if(k!=1 && mod.day!=1 && mod.day!=9){


#this loops through each event in the sequence
for(j in step){

#for each event, loop through all the males to get the previous attack count that will predict the next interaction- the support array will take care of males that weren't present that day.
for(i in 1:ncol(census.list[[1]])){
	
#start with simple +/-

#list of all the wins by the focal male
wins<-grep("win",win.loss.census[[(k-1)]][,i])

#list of all the losses by the focal male
losses<-grep("lose",win.loss.census[[(k-1)]][,i])

win.record<-rep(0,ncol(census.list[[1]]))

#if the male has not had any interactions yet, the hazard is equal across event types involving them, so set all to zero
#do wins first 
if(length(wins)>0){

wins<-win.loss.census[[(k-1)]][,i][wins]

#strip the "chase" from the end of chase wins so I can include them
wins<-sapply(wins, function(x) if(grepl("chas",x)==TRUE){
temp<-strsplit(x, split=".", fixed=T)[[1]]	
new<-paste(temp[1],temp[2],temp[3], sep=".")
return(new)
}
else return(x)
)

#totals for each type
win.tots<-table(wins)

#get just the interacting male id numbers
winids<-sapply(names(win.tots), function(x) strsplit(x, split=".", fixed=T)[[1]][3])

win.record[as.numeric(winids)]<-win.tots
}


#now substract losses if there are any
if(length(losses)>0){

losses<-win.loss.census[[(k-1)]][,i][losses]

#strip the "chase" from the end of chase wins so I can include them
losses<-sapply(losses, function(x) if(grepl("chas",x)==TRUE){
temp<-strsplit(x, split=".", fixed=T)[[1]]	
new<-paste(temp[1],temp[2],temp[3], sep=".")
return(new)
}
else return(x)
)
	
	
lose.tots<-table(losses)
	
loseids<-sapply(names(lose.tots), function(x) strsplit(x, split=".", fixed=T)[[1]][3])

for(g in 1:length(win.record)){
	
	if(g %in% as.numeric(loseids)){
	
	win.record[g]<-as.numeric(win.record[g])-lose.tots[loseids==g]
	}
	else{if(win.record[g]==0){win.record[g]<-"no.int"}}
	
}

}

if(length(losses)==0){

win.record[win.record==0]<-"no.int"

}
 
#test if the win records are strictly negative
win.pos<-win.record==0 

#Get the total number of males with whom the focal male has positive win records 
if(is.na(table(win.pos)["TRUE"])){win.pos.num<-0}
else{win.pos.num<-table(win.pos)["TRUE"]}

#match with event types- this stat predicts general INCOMING attacks based on the total number of interaction partners the focal male has positive win records with (win degree) from yesterday

stat[j,][endsWith(event.types,paste(".attack.",i, sep=""))]<-win.pos.num

}

}
}

#exclude X males
stat[,(endsWith(event.types,paste(".attack.",ncol(census.list[[1]]), sep="")) | startsWith(event.types,paste(ncol(census.list[[1]]),".attack.", sep="")))]<-0


stat<-log(1+stat)

rec.stat[[k]][[1]]<-abind(rec.stat[[k]][[1]], stat)

dimnames(rec.stat[[k]][[1]])[[3]][ncol(rec.stat[[k]][[1]][1,,])]<-"log.neutral.degree.yest"

}

return(rec.stat)
	
	
}


## Am I more/less likely to be attacked if I was chased yesterday?

add.chased.yest<-function(x){
	
rec.stat<-x

library(abind)


#loop through each day in the sequence, but applying 0 stat matrices to day 1 and 9 since there is no "yesterday" (March 27th data is missing due to field conditions)


for(k in day){

#empty matrix of the right dimensions, event# x event type. In this case, the default is all zeroes. 
stat<-matrix(0, nrow=step, ncol=length(event.types))
colnames(stat)<-event.types


if(k!=1 && mod.day!=1 && mod.day!=9){


#this loops through each event in the sequence, in this case starting with the third event since this stat requires at least one previous interaction to have taken place

for(j in step){

#for each event, loop through all the males to get the previous attack count that will predict the next interaction- the support array will take care of males that weren't present that day.
for(i in 1:ncol(census.list[[1]])){
	

#list of all the chases directed at the focal male
chases<-grep("chasedby",win.loss.census[[(k-1)]][,i])

#if the male has not had any interactions yet, the hazard is equal across event types involving them, so set all to zero

if(length(chases)>0){

#match with event types- this stat predicts general INCOMING attacks based on whether or not the focal male has been chased at all prevously that day
stat[j,][endsWith(event.types,paste(".attack.",i, sep=""))]<-1

}

}

}

}

#exclude X males
stat[,(endsWith(event.types,paste(".attack.",ncol(census.list[[1]]), sep="")) | startsWith(event.types,paste(ncol(census.list[[1]]),".attack.", sep="")))]<-0


rec.stat[[k]][[1]]<-abind(rec.stat[[k]][[1]], stat)

dimnames(rec.stat[[k]][[1]])[[3]][ncol(rec.stat[[k]][[1]][1,,])]<-"chased.yest"

}

return(rec.stat)
	
	
}


####################


### Effects of Faceoffs Yesterday with Females Absent ###


### Am I less likely to be attacked by you if I was involved in a FO with you yesterday after the females left?


add.fo.pairwise.yest<-function(x){
	
rec.stat<-x

evlist.dat<-split(event.list, event.list$Date)

library(abind)


#loop through each day in the sequence, but applying 0 stat matrices to day 1 and 9 since there is no "yesterday" (March 27th data is missing due to field conditions)

for(k in day){

#empty matrix of the right dimensions, event# x event type. In this case, the default is all zeroes. 
stat<-matrix(0, nrow=step, ncol=length(event.types))
colnames(stat)<-event.types


if(k!=1 && mod.day!=1 && mod.day!=9){

#this loops through each event in the sequence

for(j in step){

#for each event, loop through all the males to get the previous attack count that will predict the next interaction- the support array will take care of males that weren't present that day.
for(i in 1:ncol(census.list[[1]])){
	
attacks<-grep("attack",win.loss.census[[(k-1)]][,i])

#include these since I am just looking at whether the pair of males was involved in a FO. If I want to look at the effect of the focal male attacking and then the receiver initiating a FO, I would just use the attacks
receives<-grep("received",win.loss.census[[(k-1)]][,i])

ints<-union(attacks,receives)

if(length(ints)>0){

#associate interactions with initiations (FO, SF, CH) 

ints.names<-data.frame(win.loss.census[[(k-1)]][ints,i])

ints.names<-cbind(ints.names,evlist.dat[[(k-1)]][ints+1,])

#were the interactions FO's?
fos<-ints.names[grep("initFO", ints.names$Events),]


#did they happen in the absence of females? + reduce downto associated interaction to get the partner id
fo.fem<-fos[grep("1",fos$Females),1]

if(length(fo.fem)>0){
	
	#just want y/n instead of totals right now
	fo.fem<-unique(fo.fem)
	
	#get just the interacting male id numbers
rankids<-sapply(as.character(fo.fem), function(x) strsplit(x, split=".", fixed=T)[[1]][3])

#match with event types- this stat predicts INCOMING attacks based on whether the focal male was involved in a FO with the other male yesterday after the females were gone

#loop thru each element of rankids

for(l in 1:length(rankids)){

#incoming attacks directed at the focal male
stat[j,][paste(rankids[l],".attack.", i, sep="")]<-1

}

}

}
	
}

}
}

#exclude X males
stat[,(endsWith(event.types,paste(".attack.",ncol(census.list[[1]]), sep="")) | startsWith(event.types,paste(ncol(census.list[[1]]),".attack.", sep="")))]<-0


rec.stat[[k]][[1]]<-abind(rec.stat[[k]][[1]], stat)

dimnames(rec.stat[[k]][[1]])[[3]][ncol(rec.stat[[k]][[1]][1,,])]<-"FO.pairwise.yest"

}

return(rec.stat)
	
}


## Am I more likely to be attacked by you if I attacked and precipitated a FO with you yesterday when females were absent?

add.fo.attack.pairwise.yest<-function(x){
	
rec.stat<-x

evlist.dat<-split(event.list, event.list$Date)

library(abind)


#loop through each day in the sequence, but applying 0 stat matrices to day 1 and 9 since there is no "yesterday" (March 27th data is missing due to field conditions)

for(k in day){

#empty matrix of the right dimensions, event# x event type. In this case, the default is all zeroes. 
stat<-matrix(0, nrow=step, ncol=length(event.types))
colnames(stat)<-event.types

if(k!=1 && mod.day!=1 && mod.day!=9){

#this loops through each event in the sequence

for(j in step){

#for each event, loop through all the males to get the previous attack count that will predict the next interaction- the support array will take care of males that weren't present that day.
for(i in 1:ncol(census.list[[1]])){

#attacks from yesterday	
attacks<-grep("attack",win.loss.census[[(k-1)]][,i])

ints<-attacks

if(length(ints)>0){

#associate interactions with initiations (FO, SF, CH) 

ints.names<-data.frame(win.loss.census[[(k-1)]][ints,i])

ints.names<-cbind(ints.names,evlist.dat[[(k-1)]][ints+1,])

#were the interactions FO's?
fos<-ints.names[grep("initFO", ints.names$Events),]


#did they happen in the absence of females? + reduce downto associated interaction to get the partner id
fo.fem<-fos[grep("1",fos$Females),1]

if(length(fo.fem)>0){
	
	#just want y/n instead of totals right now
	fo.fem<-unique(fo.fem)
	
	#get just the interacting male id numbers
rankids<-sapply(as.character(fo.fem), function(x) strsplit(x, split=".", fixed=T)[[1]][3])

#match with event types- this stat predicts INCOMING attacks based on whether the focal male STARTED a FO with the other male yesterday after the females were gone

#loop thru each element of rankids

for(l in 1:length(rankids)){

#incoming attacks directed at the focal male
stat[j,][paste(rankids[l],".attack.", i, sep="")]<-1

}

}

}
	
}

}

}

#exclude X males
stat[,(endsWith(event.types,paste(".attack.",ncol(census.list[[1]]), sep="")) | startsWith(event.types,paste(ncol(census.list[[1]]),".attack.", sep="")))]<-0


rec.stat[[k]][[1]]<-abind(rec.stat[[k]][[1]], stat)

dimnames(rec.stat[[k]][[1]])[[3]][ncol(rec.stat[[k]][[1]][1,,])]<-"FO.attack.pairwise.yest"

}

return(rec.stat)
	
}


## Am I more likely to be attacked by you if you attacked me and started a FO with me yesterday while females were absent?

add.fo.receive.pairwise.yest<-function(x){
	
rec.stat<-x

evlist.dat<-split(event.list, event.list$Date)

library(abind)

#loop through each day in the sequence, but applying 0 stat matrices to day 1 and 9 since there is no "yesterday" (March 27th data is missing due to field conditions)

for(k in day){

#empty matrix of the right dimensions, event# x event type. In this case, the default is all zeroes. 
stat<-matrix(0, nrow=step, ncol=length(event.types))
colnames(stat)<-event.types


if(k!=1 && mod.day!=1 && mod.day!=9){

#this loops through each event in the sequence

for(j in step){

#for each event, loop through all the males to get the previous attack count that will predict the next interaction- the support array will take care of males that weren't present that day.
for(i in 1:ncol(census.list[[1]])){

#receives from yesterday	
receives<-grep("received",win.loss.census[[(k-1)]][,i])

ints<-receives

if(length(ints)>0){

#associate interactions with initiations (FO, SF, CH) 

ints.names<-data.frame(win.loss.census[[(k-1)]][ints,i])

ints.names<-cbind(ints.names,evlist.dat[[(k-1)]][ints+1,])

#were the interactions FO's?
fos<-ints.names[grep("initFO", ints.names$Events),]


#did they happen in the absence of females? + reduce downto associated interaction to get the partner id
fo.fem<-fos[grep("1",fos$Females),1]

if(length(fo.fem)>0){
	
	#just want y/n instead of totals right now
	fo.fem<-unique(fo.fem)
	
	#get just the interacting male id numbers
rankids<-sapply(as.character(fo.fem), function(x) strsplit(x, split=".", fixed=T)[[1]][3])

#match with event types- this stat predicts INCOMING attacks based on whether the focal male STARTED a FO with the other male yesterday after the females were gone

#loop thru each element of rankids

for(l in 1:length(rankids)){

#incoming attacks directed at the focal male
stat[j,][paste(rankids[l],".attack.", i, sep="")]<-1

}

}

}
	
}

}
}

#exclude X males
stat[,(endsWith(event.types,paste(".attack.",ncol(census.list[[1]]), sep="")) | startsWith(event.types,paste(ncol(census.list[[1]]),".attack.", sep="")))]<-0


rec.stat[[k]][[1]]<-abind(rec.stat[[k]][[1]], stat)

dimnames(rec.stat[[k]][[1]])[[3]][ncol(rec.stat[[k]][[1]][1,,])]<-"FO.receive.pairwise.yest"

}

return(rec.stat)
	
}


## Am I more likely to be attacked by you today if I WON a FO with you while females were absent yesterday?

add.fo.win.pairwise.yest<-function(x){
	
rec.stat<-x

evlist.dat<-split(event.list, event.list$Date)

#add row numbers as a column to census so I can match the the attacks/receives associated with wins/losses with FO's in the event list
win.loss.census.rownum<-list()

for(n in 1:length(win.loss.census)){
win.loss.census.rownum[[n]]<-cbind(win.loss.census[[n]],c(1:nrow(win.loss.census[[n]])))
}


library(abind)

#loop through each day in the sequence, but applying 0 stat matrices to day 1 and 9 since there is no "yesterday" (March 27th data is missing due to field conditions)

for(k in day){

#empty matrix of the right dimensions, event# x event type. In this case, the default is all zeroes. 
stat<-matrix(0, nrow=step, ncol=length(event.types))
colnames(stat)<-event.types


if(k!=1 && mod.day!=1 && mod.day!=9){

#this loops through each event in the sequence

for(j in step){

#for each event, loop through all the males to get the previous attack count that will predict the next interaction- the support array will take care of males that weren't present that day.
for(i in 1:ncol(census.list[[1]])){

#Remove the "fights" and "nothings" from the census list. This creates attack/receive and win/lose pairs in order of occurence
fight.times<-grep("fight",win.loss.census.rownum[[(k-1)]][,i])
nothing.times<-grep("nothing",win.loss.census.rownum[[(k-1)]][,i])

pairs<-win.loss.census.rownum[[(k-1)]][,c(i,ncol(win.loss.census.rownum[[1]]))][-c(fight.times, nothing.times),]
	
pairs.ord<-vector()

if(length(pairs)>0){

pairs.ord<-unlist(sapply(1:ncol(census.list[[1]]), function(z){
	
	intset<-pairs[,1][sapply(pairs[,1], function(x) {
		
		strsplit(x,split=".",fixed=T)[[1]][3]==z
		
	})]
	
	names(intset)<-pairs[,2][grep("TRUE",sapply(pairs, function(x) {
		
		strsplit(x,split=".",fixed=T)[[1]][3]==z
		
	}))]
	

	winstarts<-intset[(grep("win", intset)-1)]

	return(winstarts)

}))


#match with the event list and step forward to identify FO's


ints.names<-evlist.dat[[(k-1)]][(as.numeric(names(pairs.ord))+1),]

#did the FOs happen in the presence of females? + pare down to just the event type to get the partner id

fo.fem<-pairs.ord[intersect(grep("initFO", ints.names$Events),grep("1",ints.names$Females))]

if(length(fo.fem)>0){
	
	#just want y/n instead of totals right now
	fo.fem<-unique(fo.fem)
	
	#get just the interacting male id numbers
rankids<-sapply(as.character(fo.fem), function(x) strsplit(x, split=".", fixed=T)[[1]][3])

#match with event types- this stat predicts INCOMING attacks based on whether the focal male WON a FO with the other male yesterday after the females were gone

#loop thru each element of rankids

for(l in 1:length(rankids)){

#incoming attacks directed at the focal male
stat[j,][paste(rankids[l],".attack.", i, sep="")]<-1

}
}
}
}	
}

}

#exclude X males
stat[,(endsWith(event.types,paste(".attack.",ncol(census.list[[1]]), sep="")) | startsWith(event.types,paste(ncol(census.list[[1]]),".attack.", sep="")))]<-0


rec.stat[[k]][[1]]<-abind(rec.stat[[k]][[1]], stat)

dimnames(rec.stat[[k]][[1]])[[3]][ncol(rec.stat[[k]][[1]][1,,])]<-"FO.win.pairwise.yest"

}

return(rec.stat)
	
}


## Am I more likely to be attacked by you today if I LOST a FO with you while females were absent yesterday?

add.fo.loss.pairwise.yest<-function(x){
	
rec.stat<-x

evlist.dat<-split(event.list, event.list$Date)

#add row numbers as a column to census so I can match the the attacks/receives associated with wins/losses with FO's in the event list
win.loss.census.rownum<-list()

for(n in 1:length(win.loss.census)){
win.loss.census.rownum[[n]]<-cbind(win.loss.census[[n]],c(1:nrow(win.loss.census[[n]])))
}


library(abind)

#loop through each day in the sequence, but applying 0 stat matrices to day 1 and 9 since there is no "yesterday" (March 27th data is missing due to field conditions)

for(k in day){

#empty matrix of the right dimensions, event# x event type. In this case, the default is all zeroes. 
stat<-matrix(0, nrow=step, ncol=length(event.types))
colnames(stat)<-event.types


if(k!=1 && mod.day!=1 && mod.day!=9){

#this loops through each event in the sequence

for(j in step){

#for each event, loop through all the males to get the previous attack count that will predict the next interaction- the support array will take care of males that weren't present that day.
for(i in 1:ncol(census.list[[1]])){

#Remove the "fights" and "nothings" from the census list. This creates attack/receive and win/lose pairs in order of occurence
fight.times<-grep("fight",win.loss.census.rownum[[(k-1)]][,i])
nothing.times<-grep("nothing",win.loss.census.rownum[[(k-1)]][,i])

pairs<-win.loss.census.rownum[[(k-1)]][,c(i,ncol(win.loss.census.rownum[[1]]))][-c(fight.times, nothing.times),]
	
pairs.ord<-vector()

if(length(pairs)>0){

pairs.ord<-unlist(sapply(1:ncol(census.list[[1]]), function(z){
	
	intset<-pairs[,1][sapply(pairs[,1], function(x) {
		
		strsplit(x,split=".",fixed=T)[[1]][3]==z
		
	})]
	
	names(intset)<-pairs[,2][grep("TRUE",sapply(pairs, function(x) {
		
		strsplit(x,split=".",fixed=T)[[1]][3]==z
		
	}))]
	

	winstarts<-intset[(grep("lose", intset)-1)]

	return(winstarts)

}))


#match with the event list and step forward to identify FO's


ints.names<-evlist.dat[[(k-1)]][(as.numeric(names(pairs.ord))+1),]

#did the FOs happen in the presence of females? + pare down to just the event type to get the partner id

fo.fem<-pairs.ord[intersect(grep("initFO", ints.names$Events),grep("1",ints.names$Females))]

if(length(fo.fem)>0){
	
	#just want y/n instead of totals right now
	fo.fem<-unique(fo.fem)
	
	#get just the interacting male id numbers
rankids<-sapply(as.character(fo.fem), function(x) strsplit(x, split=".", fixed=T)[[1]][3])

#match with event types- this stat predicts INCOMING attacks based on whether the focal male WON a FO with the other male yesterday after the females were gone

#loop thru each element of rankids

for(l in 1:length(rankids)){

#incoming attacks directed at the focal male
stat[j,][paste(rankids[l],".attack.", i, sep="")]<-1

}
}
}
}	
}

}

#exclude X males
stat[,(endsWith(event.types,paste(".attack.",ncol(census.list[[1]]), sep="")) | startsWith(event.types,paste(ncol(census.list[[1]]),".attack.", sep="")))]<-0


rec.stat[[k]][[1]]<-abind(rec.stat[[k]][[1]], stat)

dimnames(rec.stat[[k]][[1]])[[3]][ncol(rec.stat[[k]][[1]][1,,])]<-"FO.lose.pairwise.yest"

}

return(rec.stat)
	
}


## Am I more likely to be attacked by you if i received a FO from you AND WON  YESTERDAY?


add.fo.receive.win.pairwise.yest<-function(x){
	
rec.stat<-x

evlist.dat<-split(event.list, event.list$Date)

#add row numbers as a column to census so I can match the the attacks/receives associated with wins/losses with FO's in the event list
win.loss.census.rownum<-list()

for(n in 1:length(win.loss.census)){
win.loss.census.rownum[[n]]<-cbind(win.loss.census[[n]],c(1:nrow(win.loss.census[[n]])))
}


library(abind)

#loop through each day in the sequence, but applying 0 stat matrices to day 1 and 9 since there is no "yesterday" (March 27th data is missing due to field conditions)

for(k in day){

#empty matrix of the right dimensions, event# x event type. In this case, the default is all zeroes. 
stat<-matrix(0, nrow=step, ncol=length(event.types))
colnames(stat)<-event.types


if(k!=1 && mod.day!=1 && mod.day!=9){

#this loops through each event in the sequence

for(j in step){

#for each event, loop through all the males to get the previous attack count that will predict the next interaction- the support array will take care of males that weren't present that day.
for(i in 1:ncol(census.list[[1]])){

#Remove the "fights" and "nothings" from the census list. This creates attack/receive and win/lose pairs in order of occurence
fight.times<-grep("fight",win.loss.census.rownum[[(k-1)]][,i])
nothing.times<-grep("nothing",win.loss.census.rownum[[(k-1)]][,i])

pairs<-win.loss.census.rownum[[(k-1)]][,c(i,ncol(win.loss.census.rownum[[1]]))][-c(fight.times, nothing.times),]
	
pairs.ord<-vector()

if(length(pairs)>0){

pairs.ord<-unlist(sapply(1:ncol(census.list[[1]]), function(z){
	
	intset<-pairs[,1][sapply(pairs[,1], function(x) {
		
		strsplit(x,split=".",fixed=T)[[1]][3]==z
		
	})]
	
	names(intset)<-pairs[,2][grep("TRUE",sapply(pairs, function(x) {
		
		strsplit(x,split=".",fixed=T)[[1]][3]==z
		
	}))]
	
	receives<-grep("receive",intset)
	#this time I am retaining the initiation event
	receivepluswin<-intset[receives][grep("win", intset[receives+1])]

	return(receivepluswin)

}))


#match with the event list and step forward to identify FO's


ints.names<-evlist.dat[[(k-1)]][(as.numeric(names(pairs.ord))+1),]

#did the FOs happen in the presence of females? + pare down to just the event type to get the partner id

fo.fem<-pairs.ord[intersect(grep("initFO", ints.names$Events),grep("1",ints.names$Females))]

if(length(fo.fem)>0){
	
	#just want y/n instead of totals right now
	fo.fem<-unique(fo.fem)
	
	#get just the interacting male id numbers
rankids<-sapply(as.character(fo.fem), function(x) strsplit(x, split=".", fixed=T)[[1]][3])

#match with event types- this stat predicts INCOMING attacks based on whether the focal male WON a FO with the other male yesterday after the females were gone

#loop thru each element of rankids

for(l in 1:length(rankids)){

#incoming attacks directed at the focal male
stat[j,][paste(rankids[l],".attack.", i, sep="")]<-1

}
}
}
}	
}

}

#exclude X males
stat[,(endsWith(event.types,paste(".attack.",ncol(census.list[[1]]), sep="")) | startsWith(event.types,paste(ncol(census.list[[1]]),".attack.", sep="")))]<-0


rec.stat[[k]][[1]]<-abind(rec.stat[[k]][[1]], stat)

dimnames(rec.stat[[k]][[1]])[[3]][ncol(rec.stat[[k]][[1]][1,,])]<-"FO.receive.win.pairwise.yest"

}

return(rec.stat)
	
}


## Am I more likely to be attacked by you if I precipitated an FO with you AND WON YESTERDAY?


add.fo.attack.win.pairwise.yest<-function(x){
	
rec.stat<-x

evlist.dat<-split(event.list, event.list$Date)

#add row numbers as a column to census so I can match the the attacks/receives associated with wins/losses with FO's in the event list
win.loss.census.rownum<-list()

for(n in 1:length(win.loss.census)){
win.loss.census.rownum[[n]]<-cbind(win.loss.census[[n]],c(1:nrow(win.loss.census[[n]])))
}


library(abind)

#loop through each day in the sequence, but applying 0 stat matrices to day 1 and 9 since there is no "yesterday" (March 27th data is missing due to field conditions)

for(k in day){

#empty matrix of the right dimensions, event# x event type. In this case, the default is all zeroes. 
stat<-matrix(0, nrow=step, ncol=length(event.types))
colnames(stat)<-event.types


if(k!=1 && mod.day!=1 && mod.day!=9){

#this loops through each event in the sequence

for(j in step){

#for each event, loop through all the males to get the previous attack count that will predict the next interaction- the support array will take care of males that weren't present that day.
for(i in 1:ncol(census.list[[1]])){

#Remove the "fights" and "nothings" from the census list. This creates attack/receive and win/lose pairs in order of occurence
fight.times<-grep("fight",win.loss.census.rownum[[(k-1)]][,i])
nothing.times<-grep("nothing",win.loss.census.rownum[[(k-1)]][,i])

pairs<-win.loss.census.rownum[[(k-1)]][,c(i,ncol(win.loss.census.rownum[[1]]))][-c(fight.times, nothing.times),]
	
pairs.ord<-vector()

if(length(pairs)>0){

pairs.ord<-unlist(sapply(1:ncol(census.list[[1]]), function(z){
	
	intset<-pairs[,1][sapply(pairs[,1], function(x) {
		
		strsplit(x,split=".",fixed=T)[[1]][3]==z
		
	})]
	
	names(intset)<-pairs[,2][grep("TRUE",sapply(pairs, function(x) {
		
		strsplit(x,split=".",fixed=T)[[1]][3]==z
		
	}))]
	
	attacks<-grep("attack",intset)
	#this time I am retaining the initiation event
	attackpluswin<-intset[attacks][grep("win", intset[attacks+1])]

	return(attackpluswin)

}))


#match with the event list and step forward to identify FO's


ints.names<-evlist.dat[[(k-1)]][(as.numeric(names(pairs.ord))+1),]

#did the FOs happen in the presence of females? + pare down to just the event type to get the partner id

fo.fem<-pairs.ord[intersect(grep("initFO", ints.names$Events),grep("1",ints.names$Females))]

if(length(fo.fem)>0){
	
	#just want y/n instead of totals right now
	fo.fem<-unique(fo.fem)
	
	#get just the interacting male id numbers
rankids<-sapply(as.character(fo.fem), function(x) strsplit(x, split=".", fixed=T)[[1]][3])

#match with event types- this stat predicts INCOMING attacks based on whether the focal male WON a FO with the other male yesterday after the females were gone

#loop thru each element of rankids

for(l in 1:length(rankids)){

#incoming attacks directed at the focal male
stat[j,][paste(rankids[l],".attack.", i, sep="")]<-1

}
}
}
}	
}

}

#exclude X males
stat[,(endsWith(event.types,paste(".attack.",ncol(census.list[[1]]), sep="")) | startsWith(event.types,paste(ncol(census.list[[1]]),".attack.", sep="")))]<-0


rec.stat[[k]][[1]]<-abind(rec.stat[[k]][[1]], stat)

dimnames(rec.stat[[k]][[1]])[[3]][ncol(rec.stat[[k]][[1]][1,,])]<-"FO.attack.win.pairwise.yest"

}

return(rec.stat)
	
}


## Am I more likely to be attacked by you if I received a FO from you AND LOST YESTERDAY?

add.fo.receive.lose.pairwise.yest<-function(x){
	
rec.stat<-x

evlist.dat<-split(event.list, event.list$Date)

#add row numbers as a column to census so I can match the the attacks/receives associated with wins/losses with FO's in the event list
win.loss.census.rownum<-list()

for(n in 1:length(win.loss.census)){
win.loss.census.rownum[[n]]<-cbind(win.loss.census[[n]],c(1:nrow(win.loss.census[[n]])))
}


library(abind)

#loop through each day in the sequence, but applying 0 stat matrices to day 1 and 9 since there is no "yesterday" (March 27th data is missing due to field conditions)

for(k in day){

#empty matrix of the right dimensions, event# x event type. In this case, the default is all zeroes. 
stat<-matrix(0, nrow=step, ncol=length(event.types))
colnames(stat)<-event.types


if(k!=1 && mod.day!=1 && mod.day!=9){

#this loops through each event in the sequence

for(j in step){

#for each event, loop through all the males to get the previous attack count that will predict the next interaction- the support array will take care of males that weren't present that day.
for(i in 1:ncol(census.list[[1]])){

#Remove the "fights" and "nothings" from the census list. This creates attack/receive and win/lose pairs in order of occurence
fight.times<-grep("fight",win.loss.census.rownum[[(k-1)]][,i])
nothing.times<-grep("nothing",win.loss.census.rownum[[(k-1)]][,i])

pairs<-win.loss.census.rownum[[(k-1)]][,c(i,ncol(win.loss.census.rownum[[1]]))][-c(fight.times, nothing.times),]
	
pairs.ord<-vector()

if(length(pairs)>0){

pairs.ord<-unlist(sapply(1:ncol(census.list[[1]]), function(z){
	
	intset<-pairs[,1][sapply(pairs[,1], function(x) {
		
		strsplit(x,split=".",fixed=T)[[1]][3]==z
		
	})]
	
	names(intset)<-pairs[,2][grep("TRUE",sapply(pairs, function(x) {
		
		strsplit(x,split=".",fixed=T)[[1]][3]==z
		
	}))]
	
	receive<-grep("receive",intset)
	#this time I am retaining the initiation event
	receivepluslose<-intset[receive][grep("lose", intset[receive+1])]

	return(receivepluslose)

}))


#match with the event list and step forward to identify FO's


ints.names<-evlist.dat[[(k-1)]][(as.numeric(names(pairs.ord))+1),]

#did the FOs happen in the presence of females? + pare down to just the event type to get the partner id

fo.fem<-pairs.ord[intersect(grep("initFO", ints.names$Events),grep("1",ints.names$Females))]

if(length(fo.fem)>0){
	
	#just want y/n instead of totals right now
	fo.fem<-unique(fo.fem)
	
	#get just the interacting male id numbers
rankids<-sapply(as.character(fo.fem), function(x) strsplit(x, split=".", fixed=T)[[1]][3])

#match with event types- this stat predicts INCOMING attacks based on whether the focal male WON a FO with the other male yesterday after the females were gone

#loop thru each element of rankids

for(l in 1:length(rankids)){

#incoming attacks directed at the focal male
stat[j,][paste(rankids[l],".attack.", i, sep="")]<-1

}
}
}
}	
}

}

#exclude X males
stat[,(endsWith(event.types,paste(".attack.",ncol(census.list[[1]]), sep="")) | startsWith(event.types,paste(ncol(census.list[[1]]),".attack.", sep="")))]<-0


rec.stat[[k]][[1]]<-abind(rec.stat[[k]][[1]], stat)

dimnames(rec.stat[[k]][[1]])[[3]][ncol(rec.stat[[k]][[1]][1,,])]<-"FO.receive.lose.pairwise.yest"

}

return(rec.stat)
	
}


#### Am I more likely to be attacked by you if I started a FO with you AND LOST YESTERDAY?


add.fo.attack.lose.pairwise.yest<-function(x){
	
rec.stat<-x

evlist.dat<-split(event.list, event.list$Date)

#add row numbers as a column to census so I can match the the attacks/receives associated with wins/losses with FO's in the event list
win.loss.census.rownum<-list()

for(n in 1:length(win.loss.census)){
win.loss.census.rownum[[n]]<-cbind(win.loss.census[[n]],c(1:nrow(win.loss.census[[n]])))
}


library(abind)

#loop through each day in the sequence, but applying 0 stat matrices to day 1 and 9 since there is no "yesterday" (March 27th data is missing due to field conditions)

for(k in day){

#empty matrix of the right dimensions, event# x event type. In this case, the default is all zeroes. 
stat<-matrix(0, nrow=step, ncol=length(event.types))
colnames(stat)<-event.types


if(k!=1 && mod.day!=1 && mod.day!=9){

#this loops through each event in the sequence

for(j in step){

#for each event, loop through all the males to get the previous attack count that will predict the next interaction- the support array will take care of males that weren't present that day.
for(i in 1:ncol(census.list[[1]])){

#Remove the "fights" and "nothings" from the census list. This creates attack/receive and win/lose pairs in order of occurence
fight.times<-grep("fight",win.loss.census.rownum[[(k-1)]][,i])
nothing.times<-grep("nothing",win.loss.census.rownum[[(k-1)]][,i])

pairs<-win.loss.census.rownum[[(k-1)]][,c(i,ncol(win.loss.census.rownum[[1]]))][-c(fight.times, nothing.times),]
	
pairs.ord<-vector()

if(length(pairs)>0){

pairs.ord<-unlist(sapply(1:ncol(census.list[[1]]), function(z){
	
	intset<-pairs[,1][sapply(pairs[,1], function(x) {
		
		strsplit(x,split=".",fixed=T)[[1]][3]==z
		
	})]
	
	names(intset)<-pairs[,2][grep("TRUE",sapply(pairs, function(x) {
		
		strsplit(x,split=".",fixed=T)[[1]][3]==z
		
	}))]
	
	attacks<-grep("attack",intset)
	#this time I am retaining the initiation event
	attackpluslose<-intset[attacks][grep("lose", intset[attacks+1])]

	return(attackpluslose)

}))


#match with the event list and step forward to identify FO's


ints.names<-evlist.dat[[(k-1)]][(as.numeric(names(pairs.ord))+1),]

#did the FOs happen in the presence of females? + pare down to just the event type to get the partner id

fo.fem<-pairs.ord[intersect(grep("initFO", ints.names$Events),grep("1",ints.names$Females))]

if(length(fo.fem)>0){
	
	#just want y/n instead of totals right now
	fo.fem<-unique(fo.fem)
	
	#get just the interacting male id numbers
rankids<-sapply(as.character(fo.fem), function(x) strsplit(x, split=".", fixed=T)[[1]][3])

#match with event types- this stat predicts INCOMING attacks based on whether the focal male WON a FO with the other male yesterday after the females were gone

#loop thru each element of rankids

for(l in 1:length(rankids)){

#incoming attacks directed at the focal male
stat[j,][paste(rankids[l],".attack.", i, sep="")]<-1

}
}
}
}	
}

}

#exclude X males
stat[,(endsWith(event.types,paste(".attack.",ncol(census.list[[1]]), sep="")) | startsWith(event.types,paste(ncol(census.list[[1]]),".attack.", sep="")))]<-0


rec.stat[[k]][[1]]<-abind(rec.stat[[k]][[1]], stat)

dimnames(rec.stat[[k]][[1]])[[3]][ncol(rec.stat[[k]][[1]][1,,])]<-"FO.attack.lose.pairwise.yest"

}

return(rec.stat)
	
}


######################################################

# Cumulative Predictors
######################################################

## Am I less likely to be attacked by you if my cumulative win-loss record is greater? 

add.win.loss.pairwise.cumul<-function(x){
	
rec.stat<-x

library(abind)


#loop through each day sequence
for(k in day){

#empty matrix of the right dimensions, event# x event type. In this case, the default is all zeroes. 
stat<-matrix(0, nrow=step, ncol=length(event.types))
colnames(stat)<-event.types


#this loops through each event in the sequence

for(j in step){

		
for(i in 1:ncol(census.list[[1]])){
	
#start with +/-

#list of all the wins by the focal male up to now
wins<-win.loss.census[[k]][1:(j-1),i][grep("win",win.loss.census[[k]][1:(j-1),i])]
if(k>1){
for(m in 1:(k-1)){
	wins<-append(wins, win.loss.census[[k-m]][,i][grep("win",win.loss.census[[k-m]][,i])])	
}
}

#list of all the losses by the focal male up to now
losses<-win.loss.census[[k]][1:(j-1),i][grep("lose",win.loss.census[[k]][1:(j-1),i])]
if(k>1){
for(m in 1:(k-1)){
	losses<-append(losses, win.loss.census[[k-m]][,i][grep("lose",win.loss.census[[k-m]][,i])])	
}
}

#if the male has not had any interactions yet, the hazard is equal across event types involving them, so set all to zero
#do wins first 
if(length(wins)>0){

#strip the "chase" from the end of chase wins so I can include them
wins<-sapply(wins, function(x) if(grepl("chas",x)==TRUE){
temp<-strsplit(x, split=".", fixed=T)[[1]]	
new<-paste(temp[1],temp[2],temp[3], sep=".")
return(new)
}
else return(x)
)

#totals for each type- these will be the values for the stat
win.tots<-table(wins)

#get just the interacting male id numbers
winids<-sapply(names(win.tots), function(x) strsplit(x, split=".", fixed=T)[[1]][3])

#match with event types- this stat predicts INCOMING attacks based on the difference of WINS and LOSSES of the FOCAL MALE

#loop thru each element of winids first

for(l in 1:length(winids)){

#incoming attacks directed at the focal male
stat[j,][paste(winids[l],".attack.", i, sep="")]<-win.tots[l]
	}
 
}

#now substract losses if there are any
if(length(losses)>0){

#strip the "chase" from the end of chase wins so I can include them
losses<-sapply(losses, function(x) if(grepl("chas",x)==TRUE){
temp<-strsplit(x, split=".", fixed=T)[[1]]	
new<-paste(temp[1],temp[2],temp[3], sep=".")
return(new)
}
else return(x)
)
	
lose.tots<-table(losses)
	
loseids<-sapply(names(lose.tots), function(x) strsplit(x, split=".", fixed=T)[[1]][3])

#loop thru each element of loseids, subtracting from whatever is there already

for(l in 1:length(loseids)){

#incoming attacks directed at the focal male
stat[j,][paste(loseids[l],".attack.", i, sep="")]<-stat[j,][paste(loseids[l],".attack.", i, sep="")]-lose.tots[l]
	}

}
}


}

#take the log since these numbers can become quite large and we donot expect the hazard to increase exponentially forever with win-loss record. Have to do some fancy stuff to deal with the negative records

stat<-(log(1+abs(stat))/(stat))*abs(stat) 
stat[is.na(stat)]<-0

#exclude X males
stat[,(endsWith(event.types,paste(".attack.",ncol(census.list[[1]]), sep="")) | startsWith(event.types,paste(ncol(census.list[[1]]),".attack.", sep="")))]<-0


rec.stat[[k]][[1]]<-abind(rec.stat[[k]][[1]], stat)

dimnames(rec.stat[[k]][[1]])[[3]][ncol(rec.stat[[k]][[1]][1,,])]<-"win.loss.pairwise.cumul"

}

return(rec.stat)

}


##############################################################################

#### Predictors of getting attacked in the context of copulation ####

# Am I more likely to be attacked by you WHILE I AM COPULATING if _______ #

##############################################################################


############# Recency and Reciprocity ############

#many of the network-level measures like eigcent correlate highly with, and are likely largely explained by more proximate processes such as recency and reciprocity. This is likely to be true for this system, especially given that not all males are present every day. 

#I can think of reciprocity in terms of general interaction, attacks, or wins/losses. These all tell different stories and are relevant to identifying deterrence, "dominance," and punishment behaviors. 

##are you more likely to attack me if we have interacted in any way (attack AND Receive) recently?

add.recency.cop<-function(x){

rec.stat<-x

library(abind)

#loop through each day sequence
for(k in day){

#empty matrix of the right dimensions, event# x event type. In this case, the default is all zeroes. 
stat<-matrix(0, nrow=step, ncol=length(event.types))
colnames(stat)<-event.types


#this loops through each event in the sequence

for(j in step){

#for each event, loop through all the males to get the recency ranks that will predict the next interaction- the support array will take care of males that weren't present that day.


#instead of looping through every focal male for this timestep, identify if any males were copulating in the previous timestep and loop through them. If no males are copulating, this row in stat remains all zeroes.

#this gives the columns for which males were copulating, which corresponds to their id# if there are no males copulating, the loop will simply not run. 
cop<-grep("copulating", win.loss.census[[k]][(j-1),]) 
		
for(i in cop){
	
#This is all the event steps during which fighting was happening for a particular male i, up to the current timestep, in order
#for looking at recency of attacks, change this from fight to attack. This predictor here is non-directional
fighting.times<-c(grep("attack",census.list[[k]][1:(j-1),i]),grep("receive",census.list[[k]][1:(j-1),i]))

#if the male has not had any interactions yet, the hazard is equal across event types involving them, so set all to zero 
if(length(fighting.times)>0){
	
#get the names of the interaction events put in descending order, most recent first

fighting.events.desc<-census.list[[k]][1:(j-1),i][fighting.times][order(1:length(fighting.times), decreasing=T)]

#Strip down to just the id of the interaction partners
fighting.events.desc.str<-sapply(strsplit(fighting.events.desc,".",fixed=T), function(x) x[3])

#get rid of duplicates and leave the most recent instance, leaving a list of fights ranked by recency
rankids<-unique(fighting.events.desc.str)	

#match with event types- for general interactions, this recency predicts the hazard of incoming AND outgoing attacks
#loop thru each element of rankids, the first being the most recent

for(l in 1:length(rankids)){

#incoming attacks directed at the focal male
stat[j,][paste(rankids[l],".attack.", i, sep="")]<-1/l

	}
		
}
	
}

}


rec.stat[[k]][[1]]<-abind(rec.stat[[k]][[1]], stat)

dimnames(rec.stat[[k]][[1]])[[3]][ncol(rec.stat[[k]][[1]][1,,])]<-"pure.recency.cop"

}

return(rec.stat)

}


##Are you more likely to attack me if you attacked me recently?

add.pers.cop<-function(x){

rec.stat<-x

library(abind)

#loop through each day sequence
for(k in day){

#empty matrix of the right dimensions, event# x event type. In this case, the default is all zeroes. 
stat<-matrix(0, nrow=step, ncol=length(event.types))
colnames(stat)<-event.types


#this loops through each event in the sequence

for(j in step){

#for each event, loop through all the males to get the recency ranks that will predict the next interaction- the support array will take care of males that weren't present that day.

#instead of looping through every focal male for this timestep, identify if any males were copulating in the previous timestep and loop through them. If no males are copulating, this row in stat remains all zeroes.

#this gives the columns for which males were copulating, which corresponds to their id# if there are no males copulating, the loop will simply not run. 
cop<-grep("copulating", win.loss.census[[k]][(j-1),]) 
		
for(i in cop){
	
#This is all the event steps during which fighting was happening for a particular male i, up to the one previous timestep, in order. Remember, stat in this row is mean to predict the CURRENT event
#for looking at recency of attacks, change this from fight to received
fighting.times<-grep("received",census.list[[k]][1:(j-1),i])

#if the male has not had any interactions yet, the hazard is equal across event types involving them, so set all to zero 
if(length(fighting.times)>0){
	
#get the names of the interaction events put in descending order, most recent first

fighting.events.desc<-census.list[[k]][1:(j-1),i][fighting.times][order(1:length(fighting.times), decreasing=T)]

#get rid of duplicates and leave the most recent instance, leaving a list of fights ranked by recency
recent.ev<-unique(fighting.events.desc)	

#get just the interacting male id number
rankids<-sapply(recent.ev, function(x) strsplit(x, split=".", fixed=T)[[1]][3])
	
#match with event types- this stat predicts OUTGOING attacks based on previous INCOMING attacks
#loop thru each element of rankids, the first being the most recent

for(l in 1:length(rankids)){

#incoming attacks directed at the focal male
stat[j,][paste(rankids[l],".attack.", i, sep="")]<-1/l
	}
		
}
	
}

}


rec.stat[[k]][[1]]<-abind(rec.stat[[k]][[1]], stat)

dimnames(rec.stat[[k]][[1]])[[3]][ncol(rec.stat[[k]][[1]][1,,])]<-"persistence.cop"

}

return(rec.stat)

}


##are you more likely to attack me if I attacked you recently? (as compared to out-of-the-blue)

add.reciprocity.cop<-function(x){

rec.stat<-x

library(abind)

#loop through each day sequence
for(k in day){

#empty matrix of the right dimensions, event# x event type. In this case, the default is all zeroes. 
stat<-matrix(0, nrow=step, ncol=length(event.types))
colnames(stat)<-event.types


#this loops through each event in the sequence, in this case starting with the fourth event since this stat requires at least one previous fight to have taken place

for(j in step){

#for each event, loop through all the males to get the recency ranks that will predict the next interaction- the support array will take care of males that weren't present that day.

#instead of looping through every focal male for this timestep, identify if any males were copulating in the previous timestep and loop through them. If no males are copulating, this row in stat remains all zeroes.

#this gives the columns for which males were copulating, which corresponds to their id# if there are no males copulating, the loop will simply not run. 
cop<-grep("copulating", win.loss.census[[k]][(j-1),]) 
		
for(i in cop){
	
#This is all the event steps during which fighting was happening for a particular male i, up to the the one previous timestep, in order
#in this case I am looking for whom the focal male attacked most recently
fighting.times<-grep("attack",census.list[[k]][1:(j-1),i])

#if the male has not had any interactions yet, the hazard is equal across event types involving them, so set all to zero 
if(length(fighting.times)>0){
	
#get the names of the interaction events put in descending order, most recent first

fighting.events.desc<-census.list[[k]][1:(j-1),i][fighting.times][order(1:length(fighting.times), decreasing=T)]

#get rid of duplicates and leave the most recent instance, leaving a list of fights ranked by recency
recent.ev<-unique(fighting.events.desc)	

#get just the interacting male id number
rankids<-sapply(recent.ev, function(x) strsplit(x, split=".", fixed=T)[[1]][3])
	
#match with event types- this stat predicts new OUTGOING attacks based on previous OUTGOING attacks
#loop thru each element of rankids, the first being the most recent

for(l in 1:length(rankids)){

#incoming attacks directed at the focal male
stat[j,][paste(rankids[l],".attack.", i, sep="")]<-1/l
	}
		
}
	
}

}


rec.stat[[k]][[1]]<-abind(rec.stat[[k]][[1]], stat)

dimnames(rec.stat[[k]][[1]])[[3]][ncol(rec.stat[[k]][[1]][1,,])]<-"reciprocity.cop"

}

return(rec.stat)

}


#### Explore Recency Winner/Loser effects ####

##Are you more likely to attack me if I WON against you recently? (is winning protective?)

add.win.rec.cop<-function(x){

new.stat<-x

library(abind)

#loop through each day sequence
for(k in day){

#empty matrix of the right dimensions, event# x event type. In this case, the default is all zeroes. 
stat<-matrix(0, nrow=step, ncol=length(event.types))
colnames(stat)<-event.types


#this loops through each event in the sequence

for(j in step){

#for each event, loop through all the males to get the recency ranks that will predict the next interaction- the support array will take care of males that weren't present that day.

#instead of looping through every focal male for this timestep, identify if any males were copulating in the previous timestep and loop through them. If no males are copulating, this row in stat remains all zeroes.

#this gives the columns for which males were copulating, which corresponds to their id# if there are no males copulating, the loop will simply not run. 
cop<-grep("copulating", win.loss.census[[k]][(j-1),]) 
		
for(i in cop){

	
#This is all the event steps during which fighting was happening for a particular male i, up to the current timestep, in order
#in this case I am looking for whom the focal male won against most recently
win.times<-win.loss.census[[k]][1:(j-1),i][grep("win",win.loss.census[[k]][1:(j-1),i])]

#if the male has not had any interactions yet, the hazard is equal across event types involving them, so set all to zero 
if(length(win.times)>0){

#strip the "chase" from the end of chase wins so I can include them and lump them with normal wins
win.times<-sapply(win.times, function(x) if(grepl("chas",x)==TRUE){
temp<-strsplit(x, split=".", fixed=T)[[1]]	
new<-paste(temp[1],temp[2],temp[3], sep=".")
return(new)
}
else return(x)
)


#get the names of the interaction events put in descending order, most recent first

win.events.desc<-win.times[order(1:length(win.times), decreasing=T)]

#get rid of duplicates and leave the most recent instance, leaving a list of fights ranked by recency
recent.ev<-unique(win.events.desc)	

#get just the interacting male id number
rankids<-sapply(recent.ev, function(x) strsplit(x, split=".", fixed=T)[[1]][3])
	
#match with event types- this stat predicts new OUTGOING attacks based on previous WINS
#loop thru each element of rankids, the first being the most recent

for(l in 1:length(rankids)){

#incoming attacks directed at the focal male
stat[j,][paste(rankids[l],".attack.", i, sep="")]<-1/l
	}
		
}
	
}

}


new.stat[[k]][[1]]<-abind(new.stat[[k]][[1]], stat)

dimnames(new.stat[[k]][[1]])[[3]][ncol(new.stat[[k]][[1]][1,,])]<-"win.rec.cop"

}

return(new.stat)

}


##Control for Chases: Are you more likely to attack me if I WON a CHASE against you recently? (equivalent to attacked you+chase)

add.win.chase.rec.cop<-function(x){

new.stat<-x

library(abind)

#loop through each day sequence
for(k in day){

#empty matrix of the right dimensions, event# x event type. In this case, the default is all zeroes. 
stat<-matrix(0, nrow=step, ncol=length(event.types))
colnames(stat)<-event.types


#this loops through each event in the sequence

for(j in step){

#for each event, loop through all the males to get the recency ranks that will predict the next interaction- the support array will take care of males that weren't present that day.

#instead of looping through every focal male for this timestep, identify if any males were copulating in the previous timestep and loop through them. If no males are copulating, this row in stat remains all zeroes.

#this gives the columns for which males were copulating, which corresponds to their id# if there are no males copulating, the loop will simply not run. 
cop<-grep("copulating", win.loss.census[[k]][(j-1),]) 
		
for(i in cop){
	
#This is all the event steps during which fighting was happening for a particular male i, up to the current timestep, in order
#in this case I am looking for whom the focal male attacked most recently
win.times<-grep("win",win.loss.census[[k]][1:(j-1),i])

#want the subset of these that were chases
win.chase.times<-win.loss.census[[k]][1:(j-1),i][win.times][endsWith(win.loss.census[[k]][1:(j-1),i][win.times],"chase")]


#if the male has not had any interactions yet, the hazard is equal across event types involving them, so set all to zero 
if(length(win.chase.times)>0){
	
#get the names of the interaction events put in descending order, most recent first
win.events.desc<-win.chase.times[order(1:length(win.chase.times), decreasing=T)]

#get rid of duplicates and leave the most recent instance, leaving a list of fights ranked by recency
recent.ev<-unique(win.events.desc)	

#get just the interacting male id number
rankids<-sapply(recent.ev, function(x) strsplit(x, split=".", fixed=T)[[1]][3])
	
#match with event types- this stat predicts new OUTGOING attacks based on previous OUTGOING attacks
#loop thru each element of rankids, the first being the most recent

for(l in 1:length(rankids)){

#incoming attacks directed at the focal male
stat[j,][paste(rankids[l],".attack.", i, sep="")]<-1/l
	}
		
}
	
}

}


new.stat[[k]][[1]]<-abind(new.stat[[k]][[1]], stat)

dimnames(new.stat[[k]][[1]])[[3]][ncol(new.stat[[k]][[1]][1,,])]<-"win.chase.rec.cop"

}

return(new.stat)

}


##Are you more likely to attack me if I WON a non-chase against you recently? (is winning protective?)

add.win.no.ch.rec.cop<-function(x){

new.stat<-x

library(abind)

#loop through each day sequence
for(k in day){

#empty matrix of the right dimensions, event# x event type. In this case, the default is all zeroes. 
stat<-matrix(0, nrow=step, ncol=length(event.types))
colnames(stat)<-event.types


#this loops through each event in the sequence

for(j in step){

#for each event, loop through all the males to get the recency ranks that will predict the next interaction- the support array will take care of males that weren't present that day.

#instead of looping through every focal male for this timestep, identify if any males were copulating in the previous timestep and loop through them. If no males are copulating, this row in stat remains all zeroes.

#this gives the columns for which males were copulating, which corresponds to their id# if there are no males copulating, the loop will simply not run. 
cop<-grep("copulating", win.loss.census[[k]][(j-1),]) 
		
for(i in cop){
	
#This is all the event steps during which fighting was happening for a particular male i, up to the current timestep, in order
#in this case I am looking for whom the focal male won against most recently

int.vec<-win.loss.census[[k]][1:(j-1),i]

win.times<-grep("TRUE",(grepl("win",int.vec) & !grepl("chase",int.vec)))

#if the male has not had any interactions yet, the hazard is equal across event types involving them, so set all to zero 
if(length(win.times)>0){
	
#get the names of the interaction events put in descending order, most recent first

win.events.desc<-int.vec[win.times][order(1:length(win.times), decreasing=T)]

#get rid of duplicates and leave the most recent instance, leaving a list of fights ranked by recency
recent.ev<-unique(win.events.desc)	

#get just the interacting male id number
rankids<-sapply(recent.ev, function(x) strsplit(x, split=".", fixed=T)[[1]][3])
	
#match with event types- this stat predicts new OUTGOING attacks based on previous WINS
#loop thru each element of rankids, the first being the most recent

for(l in 1:length(rankids)){

#incoming attacks directed at the focal male
stat[j,][paste(rankids[l],".attack.", i, sep="")]<-1/l
	}
		
}
	
}

}


new.stat[[k]][[1]]<-abind(new.stat[[k]][[1]], stat)

dimnames(new.stat[[k]][[1]])[[3]][ncol(new.stat[[k]][[1]][1,,])]<-"win.noCH.rec.cop"

}

return(new.stat)

}


##Are you more likely to attack me if I LOST against you recently?


add.lose.rec.cop<-function(x){

new.stat<-x

library(abind)

#loop through each day sequence
for(k in day){

#empty matrix of the right dimensions, event# x event type. In this case, the default is all zeroes. 
stat<-matrix(0, nrow=step, ncol=length(event.types))
colnames(stat)<-event.types


#this loops through each event in the sequence

for(j in step){

#for each event, loop through all the males to get the recency ranks that will predict the next interaction- the support array will take care of males that weren't present that day.

#instead of looping through every focal male for this timestep, identify if any males were copulating in the previous timestep and loop through them. If no males are copulating, this row in stat remains all zeroes.

#this gives the columns for which males were copulating, which corresponds to their id# if there are no males copulating, the loop will simply not run. 
cop<-grep("copulating", win.loss.census[[k]][(j-1),]) 
		
for(i in cop){

	
#This is all the event steps during which fighting was happening for a particular male i, up to the current timestep, in order
#in this case I am looking for whom the focal male lost to most recently
lose.times<-win.loss.census[[k]][1:(j-1),i][grep("lose",win.loss.census[[k]][1:(j-1),i])]

#if the male has not had any interactions yet, the hazard is equal across event types involving them, so set all to zero 
if(length(lose.times)>0){
	
#strip the "chase" from the end of chase wins so I can include them and lump them with normal wins
lose.times<-sapply(lose.times, function(x) if(grepl("chas",x)==TRUE){
temp<-strsplit(x, split=".", fixed=T)[[1]]	
new<-paste(temp[1],temp[2],temp[3], sep=".")
return(new)
}
else return(x)
)

#get the names of the interaction events put in descending order, most recent first

lose.events.desc<-lose.times[order(1:length(lose.times), decreasing=T)]

#get rid of duplicates and leave the most recent instance, leaving a list of losses ranked by recency
recent.ev<-unique(lose.events.desc)	

#get just the interacting male id number
rankids<-sapply(recent.ev, function(x) strsplit(x, split=".", fixed=T)[[1]][3])
	
#match with event types- this stat predicts new INCOMING attacks based on previous LOSSES
#loop thru each element of rankids, the first being the most recent

for(l in 1:length(rankids)){

#incoming attacks directed at the focal male
stat[j,][paste(rankids[l],".attack.", i, sep="")]<-1/l
	}
		
}
	
}

}


new.stat[[k]][[1]]<-abind(new.stat[[k]][[1]], stat)

dimnames(new.stat[[k]][[1]])[[3]][ncol(new.stat[[k]][[1]][1,,])]<-"lose.rec.cop"

}

return(new.stat)

}


## B->A | (A->B + AwinB)
######

add.attack.win.rec.cop<-function(x){

new.stat<-x

library(abind)

#loop through each day sequence
for(k in day){

#empty matrix of the right dimensions, event# x event type. In this case, the default is all zeroes. 
stat<-matrix(0, nrow=step, ncol=length(event.types))
colnames(stat)<-event.types


#this loops through each event in the sequence

for(j in step){

#for each event, loop through all the males to get the recency ranks that will predict the next interaction- the support array will take care of males that weren't present that day.

#instead of looping through every focal male for this timestep, identify if any males were copulating in the previous timestep and loop through them. If no males are copulating, this row in stat remains all zeroes.

#this gives the columns for which males were copulating, which corresponds to their id# if there are no males copulating, the loop will simply not run. 
cop<-grep("copulating", win.loss.census[[k]][(j-1),]) 
		
for(i in cop){
	
#want to start the loop for the stat after the first interaction is complete, but I want to start the accounting of the past interactions from the beginning

#Remove the "fights" and "nothings" from the census list. This creates attack/receive and win/lose pairs in order of occurence
fight.times<-grep("fight",win.loss.census[[k]][1:(j-1),i])
nothing.times<-grep("nothing",win.loss.census[[k]][1:(j-1),i])

pairs<-win.loss.census[[k]][1:(j-1),i][-c(fight.times, nothing.times)]

#Now I can go either way with it. Here I want all the attacks that are followed by wins

#need a way to do this that matches interaction partners but also is sensitive to the time-ordering 

attack.wins<-vector()

if(length(pairs)>0){

attack.wins<-unlist(sapply(1:ncol(census.list[[1]]), function(z){
	
	intset<-pairs[sapply(pairs, function(x) {
		
		strsplit(x,split=".",fixed=T)[[1]][3]==z
		
	})]
	
	names(intset)<-grep("TRUE",sapply(pairs, function(x) {
		
		strsplit(x,split=".",fixed=T)[[1]][3]==z
		
	}))
	
	attacks<-grep("attack",intset)
	attackpluswin<-intset[attacks+1][grep("win", intset[attacks+1])]
	
	return(attackpluswin)
	
}))
}


#if the male has not had any interactions yet, the hazard is equal across event types involving them, so set all to zero 
if(length(attack.wins)>0){

#strip the "chase" from the end of chase wins so I can include them and lump them with normal wins
attack.wins<-sapply(attack.wins, function(x) if(grepl("chas",x)==TRUE){
temp<-strsplit(x, split=".", fixed=T)[[1]]	
new<-paste(temp[1],temp[2],temp[3], sep=".")
return(new)
}
else return(x)
)

	
#get the names of the interaction events put in descending order, most recent first
attack.wins<-attack.wins[order(names(attack.wins), decreasing=T)]

#get rid of duplicates and leave the most recent instance, leaving a list of instances ranked by recency
recent.ev<-unique(attack.wins)	

#get just the interacting male id number
rankids<-sapply(recent.ev, function(x) strsplit(x, split=".", fixed=T)[[1]][3])
	
#match with event types- this stat predicts new INCOMING attacks based on previous OUTGOING attacks + WINS
#loop thru each element of rankids, the first being the most recent

for(l in 1:length(rankids)){

#incoming attacks directed at the focal male
stat[j,][paste(rankids[l],".attack.", i, sep="")]<-1/l
	}
		
}
	
}

}


new.stat[[k]][[1]]<-abind(new.stat[[k]][[1]], stat)

dimnames(new.stat[[k]][[1]])[[3]][ncol(new.stat[[k]][[1]][1,,])]<-"attack.win.rec.cop"

}

return(new.stat)

}


## B->A | (A->B + AloseB)
######


add.attack.lose.rec.cop<-function(x){

new.stat<-x

library(abind)

#loop through each day sequence
for(k in day){

#empty matrix of the right dimensions, event# x event type. In this case, the default is all zeroes. 
stat<-matrix(0, nrow=step, ncol=length(event.types))
colnames(stat)<-event.types


#this loops through each event in the sequence

for(j in step){

#for each event, loop through all the males to get the recency ranks that will predict the next interaction- the support array will take care of males that weren't present that day.

#instead of looping through every focal male for this timestep, identify if any males were copulating in the previous timestep and loop through them. If no males are copulating, this row in stat remains all zeroes.

#this gives the columns for which males were copulating, which corresponds to their id# if there are no males copulating, the loop will simply not run. 
cop<-grep("copulating", win.loss.census[[k]][(j-1),]) 
		
for(i in cop){
	
#want to start the loop for the stat after the first interaction is complete, but I want to start the accounting of the past interactions from the beginning

#Remove the "fights" and "nothings" from the census list. This creates attack/receive and win/lose pairs in order of occurence
fight.times<-grep("fight",win.loss.census[[k]][1:(j-1),i])
nothing.times<-grep("nothing",win.loss.census[[k]][1:(j-1),i])

pairs<-win.loss.census[[k]][1:(j-1),i][-c(fight.times, nothing.times)]

#need a way to do this that matches interaction partners but also is sensitive to the time-ordering 

attack.losses<-vector()

if(length(pairs)>0){

attack.losses<-unlist(sapply(1:ncol(census.list[[1]]), function(z){
	
	intset<-pairs[sapply(pairs, function(x) {
		
		strsplit(x,split=".",fixed=T)[[1]][3]==z
		
	})]
	
	names(intset)<-grep("TRUE",sapply(pairs, function(x) {
		
		strsplit(x,split=".",fixed=T)[[1]][3]==z
		
	}))
	
	attacks<-grep("attack",intset)
	attackpluslose<-intset[attacks+1][grep("lose", intset[attacks+1])]
	
	return(attackpluslose)

}))
}


#if the male has not had any interactions yet, the hazard is equal across event types involving them, so set all to zero 
if(length(attack.losses)>0){
	
#strip the "chase" from the end of chase wins so I can include them and lump them with normal wins
attack.losses<-sapply(attack.losses, function(x) if(grepl("chas",x)==TRUE){
temp<-strsplit(x, split=".", fixed=T)[[1]]	
new<-paste(temp[1],temp[2],temp[3], sep=".")
return(new)
}
else return(x)
)

	
#get the names of the interaction events put in descending order, most recent first
attack.losses<-attack.losses[order(names(attack.losses), decreasing=T)]

#get rid of duplicates and leave the most recent instance, leaving a list of instances ranked by recency
recent.ev<-unique(attack.losses)	

#get just the interacting male id number
rankids<-sapply(recent.ev, function(x) strsplit(x, split=".", fixed=T)[[1]][3])
	
#match with event types- this stat predicts new INCOMING attacks based on previous OUTGOING attacks + LOSSES
#loop thru each element of rankids, the first being the most recent

for(l in 1:length(rankids)){

#incoming attacks directed at the focal male
stat[j,][paste(rankids[l],".attack.", i, sep="")]<-1/l
	}
		
}
	
}

}


new.stat[[k]][[1]]<-abind(new.stat[[k]][[1]], stat)

dimnames(new.stat[[k]][[1]])[[3]][ncol(new.stat[[k]][[1]][1,,])]<-"attack.lose.rec.cop"

}

return(new.stat)

}


## B->A | (B->A + AwinB)
######

add.receive.win.rec.cop<-function(x){

new.stat<-x

library(abind)

#loop through each day sequence
for(k in day){

#empty matrix of the right dimensions, event# x event type. In this case, the default is all zeroes. 
stat<-matrix(0, nrow=step, ncol=length(event.types))
colnames(stat)<-event.types


#this loops through each event in the sequence

for(j in step){

#for each event, loop through all the males to get the recency ranks that will predict the next interaction- the support array will take care of males that weren't present that day.

#instead of looping through every focal male for this timestep, identify if any males were copulating in the previous timestep and loop through them. If no males are copulating, this row in stat remains all zeroes.

#this gives the columns for which males were copulating, which corresponds to their id# if there are no males copulating, the loop will simply not run. 
cop<-grep("copulating", win.loss.census[[k]][(j-1),]) 
		
for(i in cop){

	
#want to start the loop for the stat after the first interaction is complete, but I want to start the accounting of the past interactions from the beginning

#Remove the "fights" and "nothings" from the census list. This creates attack/receive and win/lose pairs in order of occurence
fight.times<-grep("fight",win.loss.census[[k]][1:(j-1),i])
nothing.times<-grep("nothing",win.loss.census[[k]][1:(j-1),i])

pairs<-win.loss.census[[k]][1:(j-1),i][-c(fight.times, nothing.times)]

#Now I can go either way with it. Here I want all the receives that are followed by wins

#need a way to do this that matches interaction partners but also is sensitive to the time-ordering 

receive.wins<-vector()

if(length(pairs)>0){

receive.wins<-unlist(sapply(1:ncol(census.list[[1]]), function(z){
	
	intset<-pairs[sapply(pairs, function(x) {
		
		strsplit(x,split=".",fixed=T)[[1]][3]==z
		
	})]
	
	names(intset)<-grep("TRUE",sapply(pairs, function(x) {
		
		strsplit(x,split=".",fixed=T)[[1]][3]==z
		
	}))
	
	receives<-grep("receive",intset)
	receivepluswin<-intset[receives+1][grep("win", intset[receives+1])]
	
	return(receivepluswin)

}))
}


#if the male has not had any interactions yet, the hazard is equal across event types involving them, so set all to zero 
if(length(receive.wins)>0){
	
#strip the "chase" from the end of chase wins so I can include them and lump them with normal wins
receive.wins<-sapply(receive.wins, function(x) if(grepl("chas",x)==TRUE){
temp<-strsplit(x, split=".", fixed=T)[[1]]	
new<-paste(temp[1],temp[2],temp[3], sep=".")
return(new)
}
else return(x)
)

#get the names of the interaction events put in descending order, most recent first
receive.wins<-receive.wins[order(names(receive.wins), decreasing=T)]

#get rid of duplicates and leave the most recent instance, leaving a list of instances ranked by recency
recent.ev<-unique(receive.wins)	

#get just the interacting male id number
rankids<-sapply(recent.ev, function(x) strsplit(x, split=".", fixed=T)[[1]][3])
	
#match with event types- this stat predicts new INCOMING attacks based on previous INCOMING attacks + WINS
#loop thru each element of rankids, the first being the most recent

for(l in 1:length(rankids)){

#incoming attacks directed at the focal male
stat[j,][paste(rankids[l],".attack.", i, sep="")]<-1/l
	}
		
}
	
}

}


new.stat[[k]][[1]]<-abind(new.stat[[k]][[1]], stat)

dimnames(new.stat[[k]][[1]])[[3]][ncol(new.stat[[k]][[1]][1,,])]<-"receive.win.rec.cop"

}

return(new.stat)

}


## B->A | (B->A + AloseB)
######

add.receive.lose.rec.cop<-function(x){

new.stat<-x

library(abind)

#loop through each day sequence
for(k in day){

#empty matrix of the right dimensions, event# x event type. In this case, the default is all zeroes. 
stat<-matrix(0, nrow=step, ncol=length(event.types))
colnames(stat)<-event.types


#this loops through each event in the sequence

for(j in step){

#for each event, loop through all the males to get the recency ranks that will predict the next interaction- the support array will take care of males that weren't present that day.

#instead of looping through every focal male for this timestep, identify if any males were copulating in the previous timestep and loop through them. If no males are copulating, this row in stat remains all zeroes.

#this gives the columns for which males were copulating, which corresponds to their id# if there are no males copulating, the loop will simply not run. 
cop<-grep("copulating", win.loss.census[[k]][(j-1),]) 
		
for(i in cop){

	
#want to start the loop for the stat after the first interaction is complete, but I want to start the accounting of the past interactions from the beginning

#Remove the "fights" and "nothings" from the census list. This creates attack/receive and win/lose pairs in order of occurence
fight.times<-grep("fight",win.loss.census[[k]][1:(j-1),i])
nothing.times<-grep("nothing",win.loss.census[[k]][1:(j-1),i])

pairs<-win.loss.census[[k]][1:(j-1),i][-c(fight.times, nothing.times)]

#Now I can go either way with it. Here I want all the receives that are followed by losses

#need a way to do this that matches interaction partners but also is sensitive to the time-ordering 

receive.losses<-vector()

if(length(pairs)>0){

receive.losses<-unlist(sapply(1:ncol(census.list[[1]]), function(z){
	
	intset<-pairs[sapply(pairs, function(x) {
		
		strsplit(x,split=".",fixed=T)[[1]][3]==z
		
	})]
	
	names(intset)<-grep("TRUE",sapply(pairs, function(x) {
		
		strsplit(x,split=".",fixed=T)[[1]][3]==z
		
	}))
	
	receives<-grep("receive",intset)
	receivepluslose<-intset[receives+1][grep("lose", intset[receives+1])]
	
	return(receivepluslose)

}))
}

#if the male has not had any interactions yet, the hazard is equal across event types involving them, so set all to zero 
if(length(receive.losses)>0){

#strip the "chase" from the end of chase wins so I can include them and lump them with normal wins
receive.losses<-sapply(receive.losses, function(x) if(grepl("chas",x)==TRUE){
temp<-strsplit(x, split=".", fixed=T)[[1]]	
new<-paste(temp[1],temp[2],temp[3], sep=".")
return(new)
}
else return(x)
)

	
#get the names of the interaction events put in descending order, most recent first
receive.losses<-receive.losses[order(names(receive.losses), decreasing=T)]

#get rid of duplicates and leave the most recent instance, leaving a list of instances ranked by recency
recent.ev<-unique(receive.losses)	

#get just the interacting male id number
rankids<-sapply(recent.ev, function(x) strsplit(x, split=".", fixed=T)[[1]][3])
	
#match with event types- this stat predicts new INCOMING attacks based on previous INCOMING attacks + LOSSES
#loop thru each element of rankids, the first being the most recent

for(l in 1:length(rankids)){

#incoming attacks directed at the focal male
stat[j,][paste(rankids[l],".attack.", i, sep="")]<-1/l

	}
		
}
	
}

}


new.stat[[k]][[1]]<-abind(new.stat[[k]][[1]], stat)

dimnames(new.stat[[k]][[1]])[[3]][ncol(new.stat[[k]][[1]][1,,])]<-"receive.lose.rec.cop"

}

return(new.stat)

}


######## General Effects of absolute recency #########

###Am I generally more likely to be attacked if I was recently ______ in terms of amount of time since the event? ####


###########################

##Am I generally more liekly to be attacked if I interacted at all recently (in seconds)?

add.gen.recency.cop<-function(x){

rec.stat<-x

library(abind)

event.list.dat<-split(event.list,event.list$Date)

#loop through each day sequence
for(k in day){

#empty matrix of the right dimensions, event# x event type. In this case, the default is all zeroes. 
stat<-matrix(0, nrow=step, ncol=length(event.types))
colnames(stat)<-event.types


#this loops through each event in the sequence

for(j in step){

#for each event, loop through all the males- the support array will take care of males that weren't present that day.

#instead of looping through every focal male for this timestep, identify if any males were copulating in the previous timestep and loop through them. If no males are copulating, this row in stat remains all zeroes.

#this gives the columns for which males were copulating, which corresponds to their id# if there are no males copulating, the loop will simply not run. 
cop<-grep("copulating", win.loss.census[[k]][(j-1),]) 
		
for(i in cop){
	
#This is all the event steps during which fighting was happening for a particular male i, up to the current timestep, in order
#for looking at recency of attacks, change this from fight to attack. This predictor here is non-directional
wins<-grep("win",win.loss.census[[k]][1:(j-1),i])
losses<-grep("lose",win.loss.census[[k]][1:(j-1),i])
end.times<-union(wins,losses)

#if the male has not had any interactions yet, the hazard is equal across event types involving them, so set all to zero 
if(length(end.times)>0){
	
#identify the most recent win or loss- this represents the end time of the most recent interaction for that male

most.rec<-max(end.times)

#get the time of the most recent end of interaction in seconds
most.rec.sec<-event.list.dat[[k]][most.rec,"Time"]

#get the difference with the current time
current.time<-event.list.dat[[k]][(j-1),"Time"]

time.diff<-(current.time-most.rec.sec)


#This will produce "infinities" in the stat array when the current event IS the disengage event. the support array should take care of this since males cannot be attacked before disengaging, but just in case, make it sqrt(timediff+1)


#match with event types- for general interactions, this recency predicts the hazard of incoming attacks

#I use a slightly more slowly decaying function here since the variance in recency can be quite large. Can explore other formulations later. 
#incoming attacks directed at the focal male
stat[j,][endsWith(event.types,paste(".attack.",i, sep=""))]<-1/sqrt(time.diff+1)

  		

}
	
}

}


rec.stat[[k]][[1]]<-abind(rec.stat[[k]][[1]], stat)

dimnames(rec.stat[[k]][[1]])[[3]][ncol(rec.stat[[k]][[1]][1,,])]<-"gen.recency.cop"

}

return(rec.stat)

}


##Am I generally more liekly to be attacked if I attacked another male recently (in seconds)?

add.gen.attack.rec.cop<-function(x){

rec.stat<-x

library(abind)

event.list.dat<-split(event.list,event.list$Date)

#loop through each day sequence
for(k in day){

#empty matrix of the right dimensions, event# x event type. In this case, the default is all zeroes. 
stat<-matrix(0, nrow=step, ncol=length(event.types))
colnames(stat)<-event.types


#this loops through each event in the sequence

for(j in step){

#for each event, loop through all the males- the support array will take care of males that weren't present that day.

#instead of looping through every focal male for this timestep, identify if any males were copulating in the previous timestep and loop through them. If no males are copulating, this row in stat remains all zeroes.

#this gives the columns for which males were copulating, which corresponds to their id# if there are no males copulating, the loop will simply not run. 
cop<-grep("copulating", win.loss.census[[k]][(j-1),]) 
		
for(i in cop){


#Remove the "fights" and "nothings" from the census list. This creates attack/receive and win/lose pairs in order of occurence
fight.times<-grep("fight",win.loss.census[[k]][1:(j-1),i])
nothing.times<-grep("nothing",win.loss.census[[k]][1:(j-1),i])

pairs<-win.loss.census[[k]][1:(j-1),i][-c(fight.times, nothing.times)]
names(pairs)<-event.list.dat[[k]][1:(j-1),"Time"][-c(fight.times, nothing.times)]

#Now I can go either way with it. Here I want all the receives that are followed by losses

#need a way to do this that matches interaction partners but also is sensitive to the time-ordering 

ints<-vector()

#in this case I only care once a male has completed an interaction, so "pairs" has to be >=2
if(length(pairs)>=2){

ints<-unlist(sapply(1:ncol(census.list[[1]]), function(z){
	
	intset<-pairs[sapply(pairs, function(x) {
		
		strsplit(x,split=".",fixed=T)[[1]][3]==z
		
	})]
	
	# names(intset)<-grep("TRUE",sapply(pairs, function(x) {
		
		# strsplit(x,split=".",fixed=T)[[1]][3]==z
		
	# }))
	
	attacks<-grep("attack",intset)
	attackplus<-intset[attacks+1]
	
	return(attackplus)

}))
}

if(length(ints)>0){
	
#get the time of the most recent end of interaction in seconds
most.rec.sec<-max(as.numeric(names(ints)))

#get the difference with the current time
current.time<-event.list.dat[[k]][(j-1),"Time"]

time.diff<-(current.time-most.rec.sec)

#This will produce "infinities" in the stat array when the current event IS the disengage event. the support array should take care of this since males cannot be attacked before disengaging, but just in case, make sure the timediff is greater than zero


#match with event types- for general interactions, this recency predicts the hazard of incoming attacks

#I use a slightly more slowly decaying function here since the variance in recency can be quite large. Can explore other formulations later. 
#incoming attacks directed at the focal male
stat[j,][endsWith(event.types,paste(".attack.",i, sep=""))]<-1/sqrt(time.diff+1)


}
			
}
	
}


rec.stat[[k]][[1]]<-abind(rec.stat[[k]][[1]], stat)

dimnames(rec.stat[[k]][[1]])[[3]][ncol(rec.stat[[k]][[1]][1,,])]<-"gen.attack.rec.cop"

}

return(rec.stat)

}


##Am I generally more liekly to be attacked if another male attacked me recently (in seconds)?

add.gen.receive.rec.cop<-function(x){

rec.stat<-x

library(abind)

event.list.dat<-split(event.list,event.list$Date)

#loop through each day sequence
for(k in day){

#empty matrix of the right dimensions, event# x event type. In this case, the default is all zeroes. 
stat<-matrix(0, nrow=step, ncol=length(event.types))
colnames(stat)<-event.types


#this loops through each event in the sequence

for(j in step){

#for each event, loop through all the males- the support array will take care of males that weren't present that day.

#instead of looping through every focal male for this timestep, identify if any males were copulating in the previous timestep and loop through them. If no males are copulating, this row in stat remains all zeroes.

#this gives the columns for which males were copulating, which corresponds to their id# if there are no males copulating, the loop will simply not run. 
cop<-grep("copulating", win.loss.census[[k]][(j-1),]) 
		
for(i in cop){


#Remove the "fights" and "nothings" from the census list. This creates attack/receive and win/lose pairs in order of occurence
fight.times<-grep("fight",win.loss.census[[k]][1:(j-1),i])
nothing.times<-grep("nothing",win.loss.census[[k]][1:(j-1),i])

pairs<-win.loss.census[[k]][1:(j-1),i][-c(fight.times, nothing.times)]
names(pairs)<-event.list.dat[[k]][1:(j-1),"Time"][-c(fight.times, nothing.times)]

#Now I can go either way with it. Here I want all the receives that are followed by losses

#need a way to do this that matches interaction partners but also is sensitive to the time-ordering 

ints<-vector()

#in this case I only care once a male has completed an interaction, so "pairs" has to be >=2
if(length(pairs)>=2){

ints<-unlist(sapply(1:ncol(census.list[[1]]), function(z){
	
	intset<-pairs[sapply(pairs, function(x) {
		
		strsplit(x,split=".",fixed=T)[[1]][3]==z
		
	})]
	
	receives<-grep("receive",intset)
	receiveplus<-intset[receives+1]
	
	return(receiveplus)

}))
}

if(length(ints)>0){
	
#get the time of the most recent end of interaction in seconds
most.rec.sec<-max(as.numeric(names(ints)))

#get the difference with the current time
current.time<-event.list.dat[[k]][(j-1),"Time"]

time.diff<-(current.time-most.rec.sec)

#This will produce "infinities" in the stat array when the current event IS the disengage event. the support array should take care of this since males cannot be attacked before disengaging, but just in case, make sure the timediff is greater than zero


#match with event types- for general interactions, this recency predicts the hazard of incoming attacks

#I use a slightly more slowly decaying function here since the variance in recency can be quite large. Can explore other formulations later. 
#incoming attacks directed at the focal male
stat[j,][endsWith(event.types,paste(".attack.",i, sep=""))]<-1/sqrt(time.diff+1)


}
			
}
	
}


rec.stat[[k]][[1]]<-abind(rec.stat[[k]][[1]], stat)

dimnames(rec.stat[[k]][[1]])[[3]][ncol(rec.stat[[k]][[1]][1,,])]<-"gen.receive.rec.cop"

}

return(rec.stat)

}


##Am I generally more liekly to be attacked if I "won" an interaction recently (in seconds)?

add.gen.win.rec.cop<-function(x){

rec.stat<-x

library(abind)

event.list.dat<-split(event.list,event.list$Date)

#loop through each day sequence
for(k in day){

#empty matrix of the right dimensions, event# x event type. In this case, the default is all zeroes. 
stat<-matrix(0, nrow=step, ncol=length(event.types))
colnames(stat)<-event.types


#this loops through each event in the sequence

for(j in step){

#for each event, loop through all the males- the support array will take care of males that weren't present that day.

#instead of looping through every focal male for this timestep, identify if any males were copulating in the previous timestep and loop through them. If no males are copulating, this row in stat remains all zeroes.

#this gives the columns for which males were copulating, which corresponds to their id# if there are no males copulating, the loop will simply not run. 
cop<-grep("copulating", win.loss.census[[k]][(j-1),]) 
		
for(i in cop){


#Remove the "fights" and "nothings" from the census list. This creates attack/receive and win/lose pairs in order of occurence
fight.times<-grep("fight",win.loss.census[[k]][1:(j-1),i])
nothing.times<-grep("nothing",win.loss.census[[k]][1:(j-1),i])

pairs<-win.loss.census[[k]][1:(j-1),i][-c(fight.times, nothing.times)]
names(pairs)<-event.list.dat[[k]][1:(j-1),"Time"][-c(fight.times, nothing.times)]

#Now I can go either way with it. Here I want all the receives that are followed by losses

#need a way to do this that matches interaction partners but also is sensitive to the time-ordering 

ints<-vector()

#in this case I only care once a male has completed an interaction, so "pairs" has to be >=2
if(length(pairs)>=2){

ints<-unlist(sapply(1:ncol(census.list[[1]]), function(z){
	
	intset<-pairs[sapply(pairs, function(x) {
		
		strsplit(x,split=".",fixed=T)[[1]][3]==z
		
	})]
	
	wins<-intset[grep("win",intset)]
	
	return(wins)

}))
}

if(length(ints)>0){
	
#get the time of the most recent end of interaction in seconds
most.rec.sec<-max(as.numeric(names(ints)))

#get the difference with the current time
current.time<-event.list.dat[[k]][(j-1),"Time"]

time.diff<-(current.time-most.rec.sec)

#This will produce "infinities" in the stat array when the current event IS the disengage event. the support array should take care of this since males cannot be attacked before disengaging, but just in case, make sure the timediff is greater than zero


#match with event types- for general interactions, this recency predicts the hazard of incoming attacks

#I use a slightly more slowly decaying function here since the variance in recency can be quite large. Can explore other formulations later. 
#incoming attacks directed at the focal male
stat[j,][endsWith(event.types,paste(".attack.",i, sep=""))]<-1/sqrt(time.diff+1)


}
			
}
	
}


rec.stat[[k]][[1]]<-abind(rec.stat[[k]][[1]], stat)

dimnames(rec.stat[[k]][[1]])[[3]][ncol(rec.stat[[k]][[1]][1,,])]<-"gen.win.rec.cop"

}

return(rec.stat)

}


##Am I generally more liekly to be attacked if I "won" an chase recently (in seconds)?

add.gen.win.chase.rec.cop<-function(x){

rec.stat<-x

library(abind)

event.list.dat<-split(event.list,event.list$Date)

#loop through each day sequence
for(k in day){

#empty matrix of the right dimensions, event# x event type. In this case, the default is all zeroes. 
stat<-matrix(0, nrow=step, ncol=length(event.types))
colnames(stat)<-event.types


#this loops through each event in the sequence

for(j in step){

#for each event, loop through all the males- the support array will take care of males that weren't present that day.

#instead of looping through every focal male for this timestep, identify if any males were copulating in the previous timestep and loop through them. If no males are copulating, this row in stat remains all zeroes.

#this gives the columns for which males were copulating, which corresponds to their id# if there are no males copulating, the loop will simply not run. 
cop<-grep("copulating", win.loss.census[[k]][(j-1),]) 
		
for(i in cop){


#Remove the "fights" and "nothings" from the census list. This creates attack/receive and win/lose pairs in order of occurence
fight.times<-grep("fight",win.loss.census[[k]][1:(j-1),i])
nothing.times<-grep("nothing",win.loss.census[[k]][1:(j-1),i])

pairs<-win.loss.census[[k]][1:(j-1),i][-c(fight.times, nothing.times)]
names(pairs)<-event.list.dat[[k]][1:(j-1),"Time"][-c(fight.times, nothing.times)]

#Now I can go either way with it. Here I want all the receives that are followed by losses

#need a way to do this that matches interaction partners but also is sensitive to the time-ordering 

ints<-vector()

#in this case I only care once a male has completed an interaction, so "pairs" has to be >=2
if(length(pairs)>=2){

ints<-unlist(sapply(1:ncol(census.list[[1]]), function(z){
	
	intset<-pairs[sapply(pairs, function(x) {
		
		strsplit(x,split=".",fixed=T)[[1]][3]==z
		
	})]
	
	wins<-intset[intersect(grep("win",intset),grep("chase",intset))]
	
	return(wins)

}))
}

if(length(ints)>0){
	
#get the time of the most recent end of interaction in seconds
most.rec.sec<-max(as.numeric(names(ints)))

#get the difference with the current time
current.time<-event.list.dat[[k]][(j-1),"Time"]

time.diff<-(current.time-most.rec.sec)

#This will produce "infinities" in the stat array when the current event IS the disengage event. the support array should take care of this since males cannot be attacked before disengaging, but just in case, make sure the timediff is greater than zero


#match with event types- for general interactions, this recency predicts the hazard of incoming attacks

#I use a slightly more slowly decaying function here since the variance in recency can be quite large. Can explore other formulations later. 
#incoming attacks directed at the focal male
stat[j,][endsWith(event.types,paste(".attack.",i, sep=""))]<-1/sqrt(time.diff+1)


}
			
}
	
}


rec.stat[[k]][[1]]<-abind(rec.stat[[k]][[1]], stat)

dimnames(rec.stat[[k]][[1]])[[3]][ncol(rec.stat[[k]][[1]][1,,])]<-"gen.win.chase.rec.cop"

}

return(rec.stat)

}


##Am I generally more liekly to be attacked if I "lost" an interaction recently (in seconds)?

add.gen.lose.rec.cop<-function(x){

rec.stat<-x

library(abind)

event.list.dat<-split(event.list,event.list$Date)

#loop through each day sequence
for(k in day){

#empty matrix of the right dimensions, event# x event type. In this case, the default is all zeroes. 
stat<-matrix(0, nrow=step, ncol=length(event.types))
colnames(stat)<-event.types


#this loops through each event in the sequence

for(j in step){

#for each event, loop through all the males- the support array will take care of males that weren't present that day.

#instead of looping through every focal male for this timestep, identify if any males were copulating in the previous timestep and loop through them. If no males are copulating, this row in stat remains all zeroes.

#this gives the columns for which males were copulating, which corresponds to their id# if there are no males copulating, the loop will simply not run. 
cop<-grep("copulating", win.loss.census[[k]][(j-1),]) 
		
for(i in cop){


#Remove the "fights" and "nothings" from the census list. This creates attack/receive and win/lose pairs in order of occurence
fight.times<-grep("fight",win.loss.census[[k]][1:(j-1),i])
nothing.times<-grep("nothing",win.loss.census[[k]][1:(j-1),i])

pairs<-win.loss.census[[k]][1:(j-1),i][-c(fight.times, nothing.times)]
names(pairs)<-event.list.dat[[k]][1:(j-1),"Time"][-c(fight.times, nothing.times)]

#Now I can go either way with it. Here I want all the receives that are followed by losses

#need a way to do this that matches interaction partners but also is sensitive to the time-ordering 

ints<-vector()

#in this case I only care once a male has completed an interaction, so "pairs" has to be >=2
if(length(pairs)>=2){

ints<-unlist(sapply(1:ncol(census.list[[1]]), function(z){
	
	intset<-pairs[sapply(pairs, function(x) {
		
		strsplit(x,split=".",fixed=T)[[1]][3]==z
		
	})]
	
	losses<-intset[grep("lose",intset)]
	
	return(losses)

}))
}

if(length(ints)>0){
	
#get the time of the most recent end of interaction in seconds
most.rec.sec<-max(as.numeric(names(ints)))

#get the difference with the current time
current.time<-event.list.dat[[k]][(j-1),"Time"]

time.diff<-(current.time-most.rec.sec)

#This will produce "infinities" in the stat array when the current event IS the disengage event. the support array should take care of this since males cannot be attacked before disengaging, but just in case, make sure the timediff is greater than zero


#match with event types- for general interactions, this recency predicts the hazard of incoming attacks

#I use a slightly more slowly decaying function here since the variance in recency can be quite large. Can explore other formulations later. 
#incoming attacks directed at the focal male
stat[j,][endsWith(event.types,paste(".attack.",i, sep=""))]<-1/sqrt(time.diff+1)


}
			
}
	
}


rec.stat[[k]][[1]]<-abind(rec.stat[[k]][[1]], stat)

dimnames(rec.stat[[k]][[1]])[[3]][ncol(rec.stat[[k]][[1]][1,,])]<-"gen.lose.rec.cop"

}

return(rec.stat)

}


##Am I generally more likely to be attacked if I attack+won recently (in seconds)?

add.gen.attack.win.rec.cop<-function(x){

rec.stat<-x

library(abind)

event.list.dat<-split(event.list,event.list$Date)

#loop through each day sequence
for(k in day){

#empty matrix of the right dimensions, event# x event type. In this case, the default is all zeroes. 
stat<-matrix(0, nrow=step, ncol=length(event.types))
colnames(stat)<-event.types


#this loops through each event in the sequence

for(j in step){

#for each event, loop through all the males- the support array will take care of males that weren't present that day.

#instead of looping through every focal male for this timestep, identify if any males were copulating in the previous timestep and loop through them. If no males are copulating, this row in stat remains all zeroes.

#this gives the columns for which males were copulating, which corresponds to their id# if there are no males copulating, the loop will simply not run. 
cop<-grep("copulating", win.loss.census[[k]][(j-1),]) 
		
for(i in cop){


#Remove the "fights" and "nothings" from the census list. This creates attack/receive and win/lose pairs in order of occurence
fight.times<-grep("fight",win.loss.census[[k]][1:(j-1),i])
nothing.times<-grep("nothing",win.loss.census[[k]][1:(j-1),i])

pairs<-win.loss.census[[k]][1:(j-1),i][-c(fight.times, nothing.times)]
names(pairs)<-event.list.dat[[k]][1:(j-1),"Time"][-c(fight.times, nothing.times)]

#Now I can go either way with it. Here I want all the receives that are followed by losses

#need a way to do this that matches interaction partners but also is sensitive to the time-ordering 

ints<-vector()

#in this case I only care once a male has completed an interaction, so "pairs" has to be >=2
if(length(pairs)>=2){

ints<-unlist(sapply(1:ncol(census.list[[1]]), function(z){
	
	intset<-pairs[sapply(pairs, function(x) {
		
		strsplit(x,split=".",fixed=T)[[1]][3]==z
		
	})]
	
	
	attacks<-grep("attack",intset)
	attackpluswin<-intset[attacks+1][grep("win",intset[attacks+1])]
	
	return(attackpluswin)

}))
}

if(length(ints)>0){
	
#get the time of the most recent end of interaction in seconds
most.rec.sec<-max(as.numeric(names(ints)))

#get the difference with the current time
current.time<-event.list.dat[[k]][(j-1),"Time"]

time.diff<-(current.time-most.rec.sec)

#This will produce "infinities" in the stat array when the current event IS the disengage event. the support array should take care of this since males cannot be attacked before disengaging, but just in case, make sure the timediff is greater than zero


#match with event types- for general interactions, this recency predicts the hazard of incoming attacks

#I use a slightly more slowly decaying function here since the variance in recency can be quite large. Can explore other formulations later. 
#incoming attacks directed at the focal male
stat[j,][endsWith(event.types,paste(".attack.",i, sep=""))]<-1/sqrt(time.diff+1)


}
			
}
	
}


rec.stat[[k]][[1]]<-abind(rec.stat[[k]][[1]], stat)

dimnames(rec.stat[[k]][[1]])[[3]][ncol(rec.stat[[k]][[1]][1,,])]<-"gen.attack.win.rec.cop"

}

return(rec.stat)

}


##Am I generally more likely to be attacked if I attack+lost recently (in seconds)?

add.gen.attack.lose.rec.cop<-function(x){

rec.stat<-x

library(abind)

event.list.dat<-split(event.list,event.list$Date)

#loop through each day sequence
for(k in day){

#empty matrix of the right dimensions, event# x event type. In this case, the default is all zeroes. 
stat<-matrix(0, nrow=step, ncol=length(event.types))
colnames(stat)<-event.types


#this loops through each event in the sequence

for(j in step){

#for each event, loop through all the males- the support array will take care of males that weren't present that day.

#instead of looping through every focal male for this timestep, identify if any males were copulating in the previous timestep and loop through them. If no males are copulating, this row in stat remains all zeroes.

#this gives the columns for which males were copulating, which corresponds to their id# if there are no males copulating, the loop will simply not run. 
cop<-grep("copulating", win.loss.census[[k]][(j-1),]) 
		
for(i in cop){


#Remove the "fights" and "nothings" from the census list. This creates attack/receive and win/lose pairs in order of occurence
fight.times<-grep("fight",win.loss.census[[k]][1:(j-1),i])
nothing.times<-grep("nothing",win.loss.census[[k]][1:(j-1),i])

pairs<-win.loss.census[[k]][1:(j-1),i][-c(fight.times, nothing.times)]
names(pairs)<-event.list.dat[[k]][1:(j-1),"Time"][-c(fight.times, nothing.times)]

#Now I can go either way with it. Here I want all the receives that are followed by losses

#need a way to do this that matches interaction partners but also is sensitive to the time-ordering 

ints<-vector()

#in this case I only care once a male has completed an interaction, so "pairs" has to be >=2
if(length(pairs)>=2){

ints<-unlist(sapply(1:ncol(census.list[[1]]), function(z){
	
	intset<-pairs[sapply(pairs, function(x) {
		
		strsplit(x,split=".",fixed=T)[[1]][3]==z
		
	})]
	
	
	attacks<-grep("attack",intset)
	attackpluswin<-intset[attacks+1][grep("lose",intset[attacks+1])]
	
	return(attackpluswin)

}))
}

if(length(ints)>0){
	
#get the time of the most recent end of interaction in seconds
most.rec.sec<-max(as.numeric(names(ints)))

#get the difference with the current time
current.time<-event.list.dat[[k]][(j-1),"Time"]

time.diff<-(current.time-most.rec.sec)

#This will produce "infinities" in the stat array when the current event IS the disengage event. the support array should take care of this since males cannot be attacked before disengaging, but just in case, make sure the timediff is greater than zero


#match with event types- for general interactions, this recency predicts the hazard of incoming attacks

#I use a slightly more slowly decaying function here since the variance in recency can be quite large. Can explore other formulations later. 
#incoming attacks directed at the focal male
stat[j,][endsWith(event.types,paste(".attack.",i, sep=""))]<-1/sqrt(time.diff+1)


}
			
}
	
}


rec.stat[[k]][[1]]<-abind(rec.stat[[k]][[1]], stat)

dimnames(rec.stat[[k]][[1]])[[3]][ncol(rec.stat[[k]][[1]][1,,])]<-"gen.attack.lose.rec.cop"

}

return(rec.stat)

}


##Am I generally more likely to be attacked if I receive+won recently (in seconds)?

add.gen.receive.win.rec.cop<-function(x){

rec.stat<-x

library(abind)

event.list.dat<-split(event.list,event.list$Date)

#loop through each day sequence
for(k in day){

#empty matrix of the right dimensions, event# x event type. In this case, the default is all zeroes. 
stat<-matrix(0, nrow=step, ncol=length(event.types))
colnames(stat)<-event.types


#this loops through each event in the sequence

for(j in step){

#for each event, loop through all the males- the support array will take care of males that weren't present that day.

#instead of looping through every focal male for this timestep, identify if any males were copulating in the previous timestep and loop through them. If no males are copulating, this row in stat remains all zeroes.

#this gives the columns for which males were copulating, which corresponds to their id# if there are no males copulating, the loop will simply not run. 
cop<-grep("copulating", win.loss.census[[k]][(j-1),]) 
		
for(i in cop){


#Remove the "fights" and "nothings" from the census list. This creates attack/receive and win/lose pairs in order of occurence
fight.times<-grep("fight",win.loss.census[[k]][1:(j-1),i])
nothing.times<-grep("nothing",win.loss.census[[k]][1:(j-1),i])

pairs<-win.loss.census[[k]][1:(j-1),i][-c(fight.times, nothing.times)]
names(pairs)<-event.list.dat[[k]][1:(j-1),"Time"][-c(fight.times, nothing.times)]

#Now I can go either way with it. Here I want all the receives that are followed by losses

#need a way to do this that matches interaction partners but also is sensitive to the time-ordering 

ints<-vector()

#in this case I only care once a male has completed an interaction, so "pairs" has to be >=2
if(length(pairs)>=2){

ints<-unlist(sapply(1:ncol(census.list[[1]]), function(z){
	
	intset<-pairs[sapply(pairs, function(x) {
		
		strsplit(x,split=".",fixed=T)[[1]][3]==z
		
	})]
	
	
	receives<-grep("receive",intset)
	receivepluswin<-intset[receives+1][grep("win",intset[receives+1])]
	
	return(receivepluswin)

}))
}

if(length(ints)>0){
	
#get the time of the most recent end of interaction in seconds
most.rec.sec<-max(as.numeric(names(ints)))

#get the difference with the current time
current.time<-event.list.dat[[k]][(j-1),"Time"]

time.diff<-(current.time-most.rec.sec)

#This will produce "infinities" in the stat array when the current event IS the disengage event. the support array should take care of this since males cannot be attacked before disengaging, but just in case, make sure the timediff is greater than zero


#match with event types- for general interactions, this recency predicts the hazard of incoming attacks

#I use a slightly more slowly decaying function here since the variance in recency can be quite large. Can explore other formulations later. 
#incoming attacks directed at the focal male
stat[j,][endsWith(event.types,paste(".attack.",i, sep=""))]<-1/sqrt(time.diff+1)


}
			
}
	
}


rec.stat[[k]][[1]]<-abind(rec.stat[[k]][[1]], stat)

dimnames(rec.stat[[k]][[1]])[[3]][ncol(rec.stat[[k]][[1]][1,,])]<-"gen.receive.win.rec.cop"

}

return(rec.stat)

}


##Am I generally more likely to be attacked if I receive+lost recently (in seconds)?

add.gen.receive.lose.rec.cop<-function(x){

rec.stat<-x

library(abind)

event.list.dat<-split(event.list,event.list$Date)

#loop through each day sequence
for(k in day){

#empty matrix of the right dimensions, event# x event type. In this case, the default is all zeroes. 
stat<-matrix(0, nrow=step, ncol=length(event.types))
colnames(stat)<-event.types


#this loops through each event in the sequence

for(j in step){

#for each event, loop through all the males- the support array will take care of males that weren't present that day.

#instead of looping through every focal male for this timestep, identify if any males were copulating in the previous timestep and loop through them. If no males are copulating, this row in stat remains all zeroes.

#this gives the columns for which males were copulating, which corresponds to their id# if there are no males copulating, the loop will simply not run. 
cop<-grep("copulating", win.loss.census[[k]][(j-1),]) 
		
for(i in cop){


#Remove the "fights" and "nothings" from the census list. This creates attack/receive and win/lose pairs in order of occurence
fight.times<-grep("fight",win.loss.census[[k]][1:(j-1),i])
nothing.times<-grep("nothing",win.loss.census[[k]][1:(j-1),i])

pairs<-win.loss.census[[k]][1:(j-1),i][-c(fight.times, nothing.times)]
names(pairs)<-event.list.dat[[k]][1:(j-1),"Time"][-c(fight.times, nothing.times)]

#Now I can go either way with it. Here I want all the receives that are followed by losses

#need a way to do this that matches interaction partners but also is sensitive to the time-ordering 

ints<-vector()

#in this case I only care once a male has completed an interaction, so "pairs" has to be >=2
if(length(pairs)>=2){

ints<-unlist(sapply(1:ncol(census.list[[1]]), function(z){
	
	intset<-pairs[sapply(pairs, function(x) {
		
		strsplit(x,split=".",fixed=T)[[1]][3]==z
		
	})]
	
	
	receives<-grep("receive",intset)
	receivepluslose<-intset[receives+1][grep("lose",intset[receives+1])]
	
	return(receivepluslose)

}))
}

if(length(ints)>0){
	
#get the time of the most recent end of interaction in seconds
most.rec.sec<-max(as.numeric(names(ints)))

#get the difference with the current time
current.time<-event.list.dat[[k]][(j-1),"Time"]

time.diff<-(current.time-most.rec.sec)

#This will produce "infinities" in the stat array when the current event IS the disengage event. the support array should take care of this since males cannot be attacked before disengaging, but just in case, make sure the timediff is greater than zero


#match with event types- for general interactions, this recency predicts the hazard of incoming attacks

#I use a slightly more slowly decaying function here since the variance in recency can be quite large. Can explore other formulations later. 
#incoming attacks directed at the focal male
stat[j,][endsWith(event.types,paste(".attack.",i, sep=""))]<-1/sqrt(time.diff+1)


}
			
}
	
}


rec.stat[[k]][[1]]<-abind(rec.stat[[k]][[1]], stat)

dimnames(rec.stat[[k]][[1]])[[3]][ncol(rec.stat[[k]][[1]][1,,])]<-"gen.receive.lose.rec.cop"

}

return(rec.stat)

}


######## Effect of time spent on lek/fighting so far Today #########


##Am I generally more likely to be attacked if I have spent more time on the lek so far today?

add.total.time.cop<-function(x){

rec.stat<-x

library(abind)

event.list.dat<-split(event.list,event.list$Date)

#loop through each day sequence
for(k in day){

#empty matrix of the right dimensions, event# x event type. In this case, the default is all zeroes. 
stat<-matrix(0, nrow=step, ncol=length(event.types))
colnames(stat)<-event.types


#this loops through each event in the sequence

for(j in step){

arrsub<-subset(pres.exact.dat[[k]],Time<event.list.dat[[k]][(j-1),"Time"])
arrsub<-arrsub[order(as.numeric(arrsub$Time)),]

#for each event, loop through all the males- the support array will take care of males that weren't present that day.

#instead of looping through every focal male for this timestep, identify if any males were copulating in the previous timestep and loop through them. If no males are copulating, this row in stat remains all zeroes.

#this gives the columns for which males were copulating, which corresponds to their id# if there are no males copulating, the loop will simply not run. 
cop<-grep("copulating", win.loss.census[[k]][(j-1),]) 
		
for(i in cop){

	

arrind<-arrsub[as.character(arrsub$Bird_ID)==i,]

if(nrow(arrind)>0){

if((nrow(arrind) %% 2)==0){
	
	tot.time<-sum(sapply(1:(nrow(arrind)/2), function(x) return(arrind[(2*x),2]-arrind[(2*x-1),2])))
	
}

if(nrow(arrind)==1){
	
	tot.time<-(event.list.dat[[k]][(j-1),"Time"]-arrind[nrow(arrind),2])
}

if((nrow(arrind) %% 2)!=0 && nrow(arrind)!=1){
	
	tot.time<-sum(sapply(1:floor(nrow(arrind)/2), function(x) return(arrind[(2*x),2]-arrind[(2*x-1),2])))+(event.list.dat[[k]][(j-1),"Time"]-arrind[nrow(arrind),2])
	
}

}

#account for timesteps that occur before the male's arrival (just to keep it clean- the support array will take care of times when the focal male is not present)
if(length(tot.time)>0 && tot.time>0){

#match with event types- for general interactions, this recency predicts the hazard of incoming attacks

#I use a slightly more slowly decaying function here since the variance in recency can be quite large. Can explore other formulations later. 
#incoming attacks directed at the focal male
stat[j,][endsWith(event.types,paste(".attack.",i, sep=""))]<-tot.time

}			
}
	
}

#exclude X males
stat[,(endsWith(event.types,paste(".attack.",ncol(census.list[[1]]), sep="")) | startsWith(event.types,paste(ncol(census.list[[1]]),".attack.", sep="")))]<-0


rec.stat[[k]][[1]]<-abind(rec.stat[[k]][[1]], stat)

dimnames(rec.stat[[k]][[1]])[[3]][ncol(rec.stat[[k]][[1]][1,,])]<-"total.time.cop"

}

return(rec.stat)

}


##Am I generally more likely to be attacked if I have spent log(1 + more time) on the lek so far today?

add.log.total.time.cop<-function(x){

rec.stat<-x

library(abind)

event.list.dat<-split(event.list,event.list$Date)

#loop through each day sequence
for(k in day){

#empty matrix of the right dimensions, event# x event type. In this case, the default is all zeroes. 
stat<-matrix(0, nrow=step, ncol=length(event.types))
colnames(stat)<-event.types


#this loops through each event in the sequence

for(j in step){

arrsub<-subset(pres.exact.dat[[k]],Time<event.list.dat[[k]][(j-1),"Time"])
arrsub<-arrsub[order(as.numeric(arrsub$Time)),]

#for each event, loop through all the males- the support array will take care of males that weren't present that day.

#instead of looping through every focal male for this timestep, identify if any males were copulating in the previous timestep and loop through them. If no males are copulating, this row in stat remains all zeroes.

#this gives the columns for which males were copulating, which corresponds to their id# if there are no males copulating, the loop will simply not run. 
cop<-grep("copulating", win.loss.census[[k]][(j-1),]) 
		
for(i in cop){


arrind<-arrsub[as.character(arrsub$Bird_ID)==i,]

if(nrow(arrind)>0){

if((nrow(arrind) %% 2)==0){
	
	tot.time<-sum(sapply(1:(nrow(arrind)/2), function(x) return(arrind[(2*x),2]-arrind[(2*x-1),2])))
	
}

if(nrow(arrind)==1){
	
	tot.time<-(event.list.dat[[k]][(j-1),"Time"]-arrind[nrow(arrind),2])
}

if((nrow(arrind) %% 2)!=0 && nrow(arrind)!=1){
	
	tot.time<-sum(sapply(1:floor(nrow(arrind)/2), function(x) return(arrind[(2*x),2]-arrind[(2*x-1),2])))+(event.list.dat[[k]][(j-1),"Time"]-arrind[nrow(arrind),2])
	
}

}

#account for timesteps that occur before the male's arrival (just to keep it clean- the support array will take care of times when the focal male is not present)
if(length(tot.time)>0 && tot.time>0){

#match with event types- for general interactions, this recency predicts the hazard of incoming attacks

#I use a slightly more slowly decaying function here since the variance in recency can be quite large. Can explore other formulations later. 
#incoming attacks directed at the focal male
stat[j,][endsWith(event.types,paste(".attack.",i, sep=""))]<-tot.time

}			
}
	
}

#exclude X males
stat[,(endsWith(event.types,paste(".attack.",ncol(census.list[[1]]), sep="")) | startsWith(event.types,paste(ncol(census.list[[1]]),".attack.", sep="")))]<-0


stat<-log(1+stat)

rec.stat[[k]][[1]]<-abind(rec.stat[[k]][[1]], stat)

dimnames(rec.stat[[k]][[1]])[[3]][ncol(rec.stat[[k]][[1]][1,,])]<-"log.total.time.cop"

}

return(rec.stat)

}


##Am I generally more likely to be interrupted if I have spent more time fighting so far today? 

add.rank.fighting.time.cop<-function(x){

rec.stat<-x

library(abind)

event.list.dat<-split(event.list,event.list$Date)

#loop through each day sequence
for(k in day){

#empty matrix of the right dimensions, event# x event type. In this case, the default is all zeroes. 
stat<-matrix(0, nrow=step, ncol=length(event.types))
colnames(stat)<-event.types


#this loops through each event in the sequence

for(j in step){


#for each event, loop through all the males- the support array will take care of males that weren't present that day.

#instead of looping through every focal male for this timestep, identify if any males were copulating in the previous timestep and loop through them. If no males are copulating, this row in stat remains all zeroes.

#this gives the columns for which males were copulating, which corresponds to their id# if there are no males copulating, the loop will simply not run. 
cop<-grep("copulating", win.loss.census[[k]][(j-1),]) 
		
for(i in cop){
	
		
#get pairs of starts and ends of interactions that preserve the row number of the original event sequence
	
wins<-grep("win",win.loss.census[[k]][1:(j-1),i])
losses<-grep("lose",win.loss.census[[k]][1:(j-1),i])
end.times<-union(wins,losses)
end.times<-end.times[order(end.times)]

attacks<-grep("attack",win.loss.census[[k]][1:(j-1),i])
receives<-grep("received",win.loss.census[[k]][1:(j-1),i])
start.times<-union(attacks,receives)
start.times<-start.times[order(start.times)]

pairs<-union(start.times,end.times)

#make sure the male has had any interactions so far and that any of these have already been completed. This will produce zero hazards for males currently engaged in interactions, which is fine since the support array makes them ineligible anyway
if(length(pairs)>=2){

pairs<-pairs[order(pairs)]

if(length(pairs) %% 2 != 0){pairs<-pairs[-length(pairs)]}


pairs.times<-event.list.dat[[k]][pairs,"Time"]

#sum up the time intervals spent fighting
intervals<-matrix(pairs.times,nrow=2)

fight.tot<-sum(intervals[2,]-intervals[1,])

#match with event types- for general interactions, this recency predicts the hazard of incoming attacks

#incoming attacks directed at the focal male
stat[j,][endsWith(event.types,paste(".attack.",i, sep=""))]<-fight.tot

#exclude attack events involving the X males 
stat[j,][endsWith(event.types,paste(".attack.",ncol(census.list[[1]]), sep="")) | startsWith(event.types,paste(ncol(census.list[[1]]),".attack.", sep=""))]<-0


}			
}


#scale by ranking due to the extreme skew in the data even in the upper half

if(max(stat[j,][grep("attack",event.types)])>0){
	
assign<-1:length(unique(stat[j,][grep("attack",event.types)]))

names(assign)<-unique(stat[j,][grep("attack",event.types)])[order(unique(stat[j,][grep("attack",event.types)]),decreasing=T)]

scaleassign<-assign[as.character(stat[j,][grep("attack",event.types)])]
scaleassign<-1/scaleassign

stat[j,][grep("attack",event.types)]<-scaleassign

}


	
}

rec.stat[[k]][[1]]<-abind(rec.stat[[k]][[1]], stat)

dimnames(rec.stat[[k]][[1]])[[3]][ncol(rec.stat[[k]][[1]][1,,])]<-"rank.fighting.time.cop"

}

return(rec.stat)

}


##Am I generally more likely to be interrupted if I have spent a larger % of time fighting so far today?  

add.rank.percent.fighting.time.cop<-function(x){

rec.stat<-x

library(abind)

event.list.dat<-split(event.list,event.list$Date)

#loop through each day sequence
for(k in day){

#empty matrix of the right dimensions, event# x event type. In this case, the default is all zeroes. 
stat<-matrix(0, nrow=step, ncol=length(event.types))
colnames(stat)<-event.types


#this loops through each event in the sequence

for(j in step){

arrsub<-subset(pres.exact.dat[[k]],Time<event.list.dat[[k]][(j-1),"Time"])
arrsub<-arrsub[order(as.numeric(arrsub$Time)),]

#for each event, loop through all the males- the support array will take care of males that weren't present that day.

#instead of looping through every focal male for this timestep, identify if any males were copulating in the previous timestep and loop through them. If no males are copulating, this row in stat remains all zeroes.

#this gives the columns for which males were copulating, which corresponds to their id# if there are no males copulating, the loop will simply not run. 
cop<-grep("copulating", win.loss.census[[k]][(j-1),]) 
		
for(i in cop){

		
#get pairs of starts and ends of interactions that preserve the row number of the original event sequence
	
wins<-grep("win",win.loss.census[[k]][1:(j-1),i])
losses<-grep("lose",win.loss.census[[k]][1:(j-1),i])
end.times<-union(wins,losses)
end.times<-end.times[order(end.times)]

attacks<-grep("attack",win.loss.census[[k]][1:(j-1),i])
receives<-grep("received",win.loss.census[[k]][1:(j-1),i])
start.times<-union(attacks,receives)
start.times<-start.times[order(start.times)]

pairs<-union(start.times,end.times)

#make sure the male has had any interactions so far and that any of these have already been completed. This will produce zero hazards for males currently engaged in interactions, which is fine since the support array makes them ineligible anyway
if(length(pairs)>=2){

pairs<-pairs[order(pairs)]

if(length(pairs) %% 2 != 0){pairs<-pairs[-length(pairs)]}


pairs.times<-event.list.dat[[k]][pairs,"Time"]

#sum up the time intervals spent fighting
intervals<-matrix(pairs.times,nrow=2)

fight.tot<-sum(intervals[2,]-intervals[1,])


##get the total time the focal male was present on the lek so far today

arrind<-arrsub[as.character(arrsub$Bird_ID)==i,]

if(nrow(arrind)>0){

if((nrow(arrind) %% 2)==0){
	
	tot.time<-sum(sapply(1:(nrow(arrind)/2), function(x) return(arrind[(2*x),2]-arrind[(2*x-1),2])))
	
}

if(nrow(arrind)==1){
	
	tot.time<-(event.list.dat[[k]][(j-1),"Time"]-arrind[nrow(arrind),2])
}

if((nrow(arrind) %% 2)!=0 && nrow(arrind)!=1){
	
	tot.time<-sum(sapply(1:floor(nrow(arrind)/2), function(x) return(arrind[(2*x),2]-arrind[(2*x-1),2])))+(event.list.dat[[k]][(j-1),"Time"]-arrind[nrow(arrind),2])
	
}

}


if(length(tot.time)>0 && tot.time>0){
perc.fight<-100*(fight.tot/tot.time)
#match with event types- for general interactions, this recency predicts the hazard of incoming attacks

#incoming attacks directed at the focal male
stat[j,][endsWith(event.types,paste(".attack.",i, sep=""))]<-perc.fight

#exclude attack events involving the X males 
stat[j,][endsWith(event.types,paste(".attack.",ncol(census.list[[1]]), sep="")) | startsWith(event.types,paste(ncol(census.list[[1]]),".attack.", sep=""))]<-0
}

}			
}

#scale by ranking due to the extreme skew in the data even in the upper half

if(max(stat[j,][grep("attack",event.types)])>0){
	
assign<-1:length(unique(stat[j,][grep("attack",event.types)]))

names(assign)<-unique(stat[j,][grep("attack",event.types)])[order(unique(stat[j,][grep("attack",event.types)]),decreasing=T)]

scaleassign<-assign[as.character(stat[j,][grep("attack",event.types)])]
scaleassign<-1/scaleassign

stat[j,][grep("attack",event.types)]<-scaleassign

}

	
}

rec.stat[[k]][[1]]<-abind(rec.stat[[k]][[1]], stat)

dimnames(rec.stat[[k]][[1]])[[3]][ncol(rec.stat[[k]][[1]][1,,])]<-"rank.percent.fighting.time.cop"

}

return(rec.stat)

}


###### Cumulative history of pairwise interaction for the day #

##test log NUMBER of attacks, receives, wins, losses


##Are you more or less likely to attack me if we interacted log(1+MORE TIMES) previously today?


add.log.count.pairwise.day.cop<-function(x){
	
rec.stat<-x

library(abind)

#loop through each day sequence
for(k in day){

#empty matrix of the right dimensions, event# x event type. In this case, the default is all zeroes. 
stat<-matrix(0, nrow=step, ncol=length(event.types))
colnames(stat)<-event.types


#this loops through each event in the sequence

for(j in step){

#for each event, loop through all the males to get the previous attack count that will predict the next interaction- the support array will take care of males that weren't present that day.

#instead of looping through every focal male for this timestep, identify if any males were copulating in the previous timestep and loop through them. If no males are copulating, this row in stat remains all zeroes.

#this gives the columns for which males were copulating, which corresponds to their id# if there are no males copulating, the loop will simply not run. 
cop<-grep("copulating", win.loss.census[[k]][(j-1),]) 
		
for(i in cop){
	
#list of all the attacks perpetrated by the focal male
attacks<-grep("attack",census.list[[k]][1:(j-1),i])

#list of all the attacks directed at the focal male
receives<-grep("received",census.list[[k]][1:(j-1),i])

#if the male has not had any interactions yet, the hazard is equal across event types involving them, so set all to zero 
if(length(attacks)>0){

#totals for each type- these will be the values for the stat
attack.tots<-table(census.list[[k]][1:(j-1),i][attacks])

#get just the interacting male id numbers
rankids<-sapply(names(attack.tots), function(x) strsplit(x, split=".", fixed=T)[[1]][3])

#match with event types- this stat predicts INCOMING attacks based on total previous OUTGOING + INCOMING attacks
#loop thru each element of rankids

for(l in 1:length(rankids)){

#incoming attacks directed at the focal male, the hazrd is proportional to the number of previous outgoing attacks
stat[j,][paste(rankids[l],".attack.", i, sep="")]<-attack.tots[l]
	}
}

#add receives if there are any
if(length(receives)>0){

#totals for each type- these will be the values for the stat
receive.tots<-table(census.list[[k]][1:(j-1),i][receives])

#get just the interacting male id numbers
rankids<-sapply(names(receive.tots), function(x) strsplit(x, split=".", fixed=T)[[1]][3])

#match with event types
#loop thru each element of rankids

for(l in 1:length(rankids)){

#incoming attacks directed at the focal male, the hazrd is proportional to the number of previous incoming attacks
stat[j,][paste(rankids[l],".attack.", i, sep="")]<-(stat[j,][paste(rankids[l],".attack.", i, sep="")]+receive.tots[l])
	}
		
}
	
}

#exclude attack events involving the X males 
stat[j,][endsWith(event.types,paste(".attack.",ncol(census.list[[1]]), sep="")) | startsWith(event.types,paste(ncol(census.list[[1]]),".attack.", sep=""))]<-0


}

stat<-log(1+stat)

rec.stat[[k]][[1]]<-abind(rec.stat[[k]][[1]], stat)

dimnames(rec.stat[[k]][[1]])[[3]][ncol(rec.stat[[k]][[1]][1,,])]<-"log.count.pairwise.day.cop"

}

return(rec.stat)

	
}


##Are you more or less likely to attack me if I attacked you MORE TIMES in the past day?

add.attack.log.count.pairwise.day.cop<-function(x){
	
rec.stat<-x

library(abind)

#loop through each day sequence
for(k in day){

#empty matrix of the right dimensions, event# x event type. In this case, the default is all zeroes. 
stat<-matrix(0, nrow=step, ncol=length(event.types))
colnames(stat)<-event.types


#this loops through each event in the sequence

for(j in step){

#for each event, loop through all the males to get the previous attack count that will predict the next interaction- the support array will take care of males that weren't present that day.

#instead of looping through every focal male for this timestep, identify if any males were copulating in the previous timestep and loop through them. If no males are copulating, this row in stat remains all zeroes.

#this gives the columns for which males were copulating, which corresponds to their id# if there are no males copulating, the loop will simply not run. 
cop<-grep("copulating", win.loss.census[[k]][(j-1),]) 
		
for(i in cop){
	
#list of all the attacks perpetrated by the focal male
attacks<-grep("attack",census.list[[k]][1:(j-1),i])

#if the male has not had any interactions yet, the hazard is equal across event types involving them, so set all to zero 
if(length(attacks)>0){

#totals for each type- these will be the values for the stat
attack.tots<-table(census.list[[k]][1:(j-1),i][attacks])

#get just the interacting male id numbers
rankids<-sapply(names(attack.tots), function(x) strsplit(x, split=".", fixed=T)[[1]][3])

#match with event types- this stat predicts INCOMING attacks based on total previous OUTGOING attacks
#loop thru each element of rankids

for(l in 1:length(rankids)){

#incoming attacks directed at the focal male, the hazrd is proportional to the number of previous outgoing attacks
stat[j,][paste(rankids[l],".attack.", i, sep="")]<-attack.tots[l]
	}

		
}
	
}

#exclude attack events involving the X males 
stat[j,][endsWith(event.types,paste(".attack.",ncol(census.list[[1]]), sep="")) | startsWith(event.types,paste(ncol(census.list[[1]]),".attack.", sep=""))]<-0


}

stat<-log(1+stat)

rec.stat[[k]][[1]]<-abind(rec.stat[[k]][[1]], stat)

dimnames(rec.stat[[k]][[1]])[[3]][ncol(rec.stat[[k]][[1]][1,,])]<-"attack.log.count.pairwise.day.cop"

}

return(rec.stat)

	
}


## Are you more or less likely to attack me if YOU attacked me MORE TIMES in the past day?

add.receive.log.count.pairwise.day.cop<-function(x){
	
	rec.stat<-x

library(abind)

#loop through each day sequence
for(k in day){

#empty matrix of the right dimensions, event# x event type. In this case, the default is all zeroes. 
stat<-matrix(0, nrow=step, ncol=length(event.types))
colnames(stat)<-event.types


#this loops through each event in the sequence

for(j in step){

#for each event, loop through all the males to get the previous attack count that will predict the next interaction- the support array will take care of males that weren't present that day.

#instead of looping through every focal male for this timestep, identify if any males were copulating in the previous timestep and loop through them. If no males are copulating, this row in stat remains all zeroes.

#this gives the columns for which males were copulating, which corresponds to their id# if there are no males copulating, the loop will simply not run. 
cop<-grep("copulating", win.loss.census[[k]][(j-1),]) 
		
for(i in cop){
	
#list of all the attacks directed at the focal male
receives<-grep("received",census.list[[k]][1:(j-1),i])

#if the male has not had any interactions yet, the hazard is equal across event types involving them, so set all to zero 
if(length(receives)>0){

#totals for each type- these will be the values for the stat
receive.tots<-table(census.list[[k]][1:(j-1),i][receives])

#get just the interacting male id numbers
rankids<-sapply(names(receive.tots), function(x) strsplit(x, split=".", fixed=T)[[1]][3])

#match with event types- this stat predicts INCOMING attacks based on total previous INCOMING attacks
#loop thru each element of rankids

for(l in 1:length(rankids)){

#incoming attacks directed at the focal male, the hazrd is proportional to the number of previous outgoing attacks
stat[j,][paste(rankids[l],".attack.", i, sep="")]<-receive.tots[l]
	}

		
}
	
}

#exclude attack events involving the X males 
stat[j,][endsWith(event.types,paste(".attack.",ncol(census.list[[1]]), sep="")) | startsWith(event.types,paste(ncol(census.list[[1]]),".attack.", sep=""))]<-0


}

stat<-log(1+stat)

rec.stat[[k]][[1]]<-abind(rec.stat[[k]][[1]], stat)

dimnames(rec.stat[[k]][[1]])[[3]][ncol(rec.stat[[k]][[1]][1,,])]<-"receive.log.count.pairwise.day.cop"

}

return(rec.stat)

	
}


## Are you more or less likely to attack me if I WON against you MORE TIMES in the past day?

add.win.log.count.pairwise.day.cop<-function(x){
	
rec.stat<-x

library(abind)

#loop through each day sequence
for(k in day){

#empty matrix of the right dimensions, event# x event type. In this case, the default is all zeroes. 
stat<-matrix(0, nrow=step, ncol=length(event.types))
colnames(stat)<-event.types


#this loops through each event in the sequence

for(j in step){

#for each event, loop through all the males to get the previous attack count that will predict the next interaction- the support array will take care of males that weren't present that day.

#instead of looping through every focal male for this timestep, identify if any males were copulating in the previous timestep and loop through them. If no males are copulating, this row in stat remains all zeroes.

#this gives the columns for which males were copulating, which corresponds to their id# if there are no males copulating, the loop will simply not run. 
cop<-grep("copulating", win.loss.census[[k]][(j-1),]) 
		
for(i in cop){
	
#list of all the wins by the focal male
wins<-win.loss.census[[k]][1:(j-1),i][grep("win",win.loss.census[[k]][1:(j-1),i])]

#strip the "chase" from the end of chase wins so I can include them
wins<-sapply(wins, function(x) if(grepl("chas",x)==TRUE){
temp<-strsplit(x, split=".", fixed=T)[[1]]	
new<-paste(temp[1],temp[2],temp[3], sep=".")
return(new)
}
else return(x)
)

#if the male has not had any interactions yet, the hazard is equal across event types involving them, so set all to zero 
if(length(wins)>0){

#totals for each type- these will be the values for the stat
win.tots<-table(wins)

#get just the interacting male id numbers
rankids<-sapply(names(win.tots), function(x) strsplit(x, split=".", fixed=T)[[1]][3])

#match with event types- this stat predicts INCOMING attacks based on total previous WINS by the focal male
#loop thru each element of rankids

for(l in 1:length(rankids)){

#incoming attacks directed at the focal male, the hazrd is proportional to the number of previous outgoing attacks
stat[j,][paste(rankids[l],".attack.", i, sep="")]<-win.tots[l]
	}


		
}
	
}

#exclude attack events involving the X males 
stat[j,][endsWith(event.types,paste(".attack.",ncol(census.list[[1]]), sep="")) | startsWith(event.types,paste(ncol(census.list[[1]]),".attack.", sep=""))]<-0


}

stat<-log(1+stat)

rec.stat[[k]][[1]]<-abind(rec.stat[[k]][[1]], stat)

dimnames(rec.stat[[k]][[1]])[[3]][ncol(rec.stat[[k]][[1]][1,,])]<-"win.log.count.pairwise.day.cop"

}

return(rec.stat)

	
}


## Are you more or less likely to attack me if I LOST to you MORE TIMES in the past day?

add.loss.log.count.pairwise.day.cop<-function(x){
	
	rec.stat<-x

library(abind)

#loop through each day sequence
for(k in day){

#empty matrix of the right dimensions, event# x event type. In this case, the default is all zeroes. 
stat<-matrix(0, nrow=step, ncol=length(event.types))
colnames(stat)<-event.types


#this loops through each event in the sequence

for(j in step){

#for each event, loop through all the males to get the previous attack count that will predict the next interaction- the support array will take care of males that weren't present that day.

#instead of looping through every focal male for this timestep, identify if any males were copulating in the previous timestep and loop through them. If no males are copulating, this row in stat remains all zeroes.

#this gives the columns for which males were copulating, which corresponds to their id# if there are no males copulating, the loop will simply not run. 
cop<-grep("copulating", win.loss.census[[k]][(j-1),]) 
		
for(i in cop){
	
#list of all the losses by the focal male
losses<-win.loss.census[[k]][1:(j-1),i][grep("lose",win.loss.census[[k]][1:(j-1),i])]

#strip the "chase" from the end of chase wins so I can include them
losses<-sapply(losses, function(x) if(grepl("chas",x)==TRUE){
temp<-strsplit(x, split=".", fixed=T)[[1]]	
new<-paste(temp[1],temp[2],temp[3], sep=".")
return(new)
}
else return(x)
)

#if the male has not had any interactions yet, the hazard is equal across event types involving them, so set all to zero 
if(length(losses)>0){

#totals for each type- these will be the values for the stat
lose.tots<-table(losses)

#get just the interacting male id numbers
rankids<-sapply(names(lose.tots), function(x) strsplit(x, split=".", fixed=T)[[1]][3])

#match with event types- this stat predicts INCOMING attacks based on total previous LOSSES by the focal male
#loop thru each element of rankids

for(l in 1:length(rankids)){

#incoming attacks directed at the focal male, the hazrd is proportional to the number of previous losses
stat[j,][paste(rankids[l],".attack.", i, sep="")]<-lose.tots[l]
	}


		
}
	
}

#exclude attack events involving the X males 
stat[j,][endsWith(event.types,paste(".attack.",ncol(census.list[[1]]), sep="")) | startsWith(event.types,paste(ncol(census.list[[1]]),".attack.", sep=""))]<-0


}

stat<-log(1+stat)

rec.stat[[k]][[1]]<-abind(rec.stat[[k]][[1]], stat)

dimnames(rec.stat[[k]][[1]])[[3]][ncol(rec.stat[[k]][[1]][1,,])]<-"loss.log.count.pairwise.day.cop"

}

return(rec.stat)

}


#try accounting for "linkages" instead- same answer, or better? Right now I'm treating initiations and wins as independent, but they are always pairs, and they covary in particular, and apparently important ways that I am not capturing. Maybe the model is making up for the missing covariance term by forming these associations among the variables? so i want one for the total number and then four for the "linkages". Yeah, right now I'm double counting, since the total of all four is actually twice the total number of interactions. So it's proportional but not identical. Interesting. May want to rethink recency metrics too? (it's not the end of the world if my model is a little imperfect because I am using multiple metrics to approximate one better one- I'l still get a similar result at the end, it seems)

### log number of attack+wins   


add.attack.win.count.pairwise.day.cop<-function(x){

new.stat<-x

library(abind)

#loop through each day sequence
for(k in day){

#empty matrix of the right dimensions, event# x event type. In this case, the default is all zeroes. 
stat<-matrix(0, nrow=step, ncol=length(event.types))
colnames(stat)<-event.types


#this loops through each event in the sequence

for(j in step){

#for each event, loop through all the males to get the recency ranks that will predict the next interaction- the support array will take care of males that weren't present that day.

#instead of looping through every focal male for this timestep, identify if any males were copulating in the previous timestep and loop through them. If no males are copulating, this row in stat remains all zeroes.

#this gives the columns for which males were copulating, which corresponds to their id# if there are no males copulating, the loop will simply not run. 
cop<-grep("copulating", win.loss.census[[k]][(j-1),]) 
		
for(i in cop){

	
#want to start the loop for the stat after the first interaction is complete, but I want to start the accounting of the past interactions from the beginning

int.vector<-win.loss.census[[k]][1:(j-1),i]


#Remove the "fights" and "nothings" from the census list. This creates attack/receive and win/lose pairs in order of occurence
fight.times<-grep("fight",int.vector)
nothing.times<-grep("nothing",int.vector)

pairs<-int.vector[-c(fight.times, nothing.times)]

#need a way to do this that matches interaction partners but also is sensitive to the time-ordering 

attack.wins<-vector()

if(length(pairs)>0){

attack.wins<-unlist(sapply(1:ncol(census.list[[1]]), function(z){
	
	intset<-pairs[sapply(pairs, function(x) {
		
		strsplit(x,split=".",fixed=T)[[1]][3]==z
		
	})]
	
	names(intset)<-grep("TRUE",sapply(pairs, function(x) {
		
		strsplit(x,split=".",fixed=T)[[1]][3]==z
		
	}))
	
	attacks<-grep("attack",intset)
	attackpluswin<-intset[attacks+1][grep("win", intset[attacks+1])]
	
	return(attackpluswin)
	
}))
}


#if the male has not had any interactions yet, the hazard is equal across event types involving them, so set all to zero 
if(length(attack.wins)>0){

#strip the "chase" from the end of chase wins so I can include them
attack.wins<-sapply(attack.wins, function(x) if(grepl("chas",x)==TRUE){
temp<-strsplit(x, split=".", fixed=T)[[1]]	
new<-paste(temp[1],temp[2],temp[3], sep=".")
return(new)
}
else return(x)
)
	
#get the names of the interaction events put in descending order, most recent first
#attack.wins<-attack.wins[order(names(attack.wins), decreasing=T)]

#get the number of interactions with each other male
counts<-table(attack.wins)	

#get just the interacting male id number
rankids<-sapply(names(counts), function(x) strsplit(x, split=".", fixed=T)[[1]][3])
	
#match with event types- this stat predicts new INCOMING attacks based on the number of previous OUTGOING attacks + WINS
#loop thru each element of rankids

for(l in 1:length(rankids)){

#incoming attacks directed at the focal male
stat[j,][paste(rankids[l],".attack.", i, sep="")]<-counts[l]
	}
		
}
	
}

#exclude attack events involving the X males 
stat[j,][endsWith(event.types,paste(".attack.",ncol(census.list[[1]]), sep="")) | startsWith(event.types,paste(ncol(census.list[[1]]),".attack.", sep=""))]<-0


}

#take the log 
stat<-log(1+stat)

new.stat[[k]][[1]]<-abind(new.stat[[k]][[1]], stat)

dimnames(new.stat[[k]][[1]])[[3]][ncol(new.stat[[k]][[1]][1,,])]<-"attack.win.count.pairwise.day.cop"

}

return(new.stat)

}


 ## log number of attack-losses  

add.attack.lose.count.pairwise.day.cop<-function(x){

new.stat<-x

library(abind)

#loop through each day sequence
for(k in day){

#empty matrix of the right dimensions, event# x event type. In this case, the default is all zeroes. 
stat<-matrix(0, nrow=step, ncol=length(event.types))
colnames(stat)<-event.types


#this loops through each event in the sequence

for(j in step){

#for each event, loop through all the males to get the recency ranks that will predict the next interaction- the support array will take care of males that weren't present that day.

#instead of looping through every focal male for this timestep, identify if any males were copulating in the previous timestep and loop through them. If no males are copulating, this row in stat remains all zeroes.

#this gives the columns for which males were copulating, which corresponds to their id# if there are no males copulating, the loop will simply not run. 
cop<-grep("copulating", win.loss.census[[k]][(j-1),]) 
		
for(i in cop){

	
#want to start the loop for the stat after the first interaction is complete, but I want to start the accounting of the past interactions from the beginning

int.vector<-win.loss.census[[k]][1:(j-1),i]


#Remove the "fights" and "nothings" from the census list. This creates attack/receive and win/lose pairs in order of occurence
fight.times<-grep("fight",int.vector)
nothing.times<-grep("nothing",int.vector)

pairs<-int.vector[-c(fight.times, nothing.times)]

#need a way to do this that matches interaction partners but also is sensitive to the time-ordering 

attack.losses<-vector()

if(length(pairs)>0){

attack.losses<-unlist(sapply(1:ncol(census.list[[1]]), function(z){
	
	intset<-pairs[sapply(pairs, function(x) {
		
		strsplit(x,split=".",fixed=T)[[1]][3]==z
		
	})]
	
	names(intset)<-grep("TRUE",sapply(pairs, function(x) {
		
		strsplit(x,split=".",fixed=T)[[1]][3]==z
		
	}))
	
	attacks<-grep("attack",intset)
	attackpluslose<-intset[attacks+1][grep("lose", intset[attacks+1])]
	
	return(attackpluslose)
	
}))
}


#if the male has not had any interactions yet, the hazard is equal across event types involving them, so set all to zero 
if(length(attack.losses)>0){

#strip the "chase" from the end of chase wins so I can include them
attack.losses<-sapply(attack.losses, function(x) if(grepl("chas",x)==TRUE){
temp<-strsplit(x, split=".", fixed=T)[[1]]	
new<-paste(temp[1],temp[2],temp[3], sep=".")
return(new)
}
else return(x)
)


#get the number of interactions with each other male
counts<-table(attack.losses)	

#get just the interacting male id number
rankids<-sapply(names(counts), function(x) strsplit(x, split=".", fixed=T)[[1]][3])
	
#match with event types- this stat predicts new INCOMING attacks based on the number of previous OUTGOING attacks + WINS
#loop thru each element of rankids

for(l in 1:length(rankids)){

#incoming attacks directed at the focal male
stat[j,][paste(rankids[l],".attack.", i, sep="")]<-counts[l]
	}
		
}
	
}

#exclude attack events involving the X males 
stat[j,][endsWith(event.types,paste(".attack.",ncol(census.list[[1]]), sep="")) | startsWith(event.types,paste(ncol(census.list[[1]]),".attack.", sep=""))]<-0


}

#take the log 
stat<-log(1+stat)

new.stat[[k]][[1]]<-abind(new.stat[[k]][[1]], stat)

dimnames(new.stat[[k]][[1]])[[3]][ncol(new.stat[[k]][[1]][1,,])]<-"attack.lose.count.pairwise.day.cop"

}

return(new.stat)

}


## log number of Receive-wins?

add.receive.win.count.pairwise.day.cop<-function(x){

new.stat<-x

library(abind)

#loop through each day sequence
for(k in day){

#empty matrix of the right dimensions, event# x event type. In this case, the default is all zeroes. 
stat<-matrix(0, nrow=step, ncol=length(event.types))
colnames(stat)<-event.types


#this loops through each event in the sequence

for(j in step){

#for each event, loop through all the males to get the recency ranks that will predict the next interaction- the support array will take care of males that weren't present that day.

#instead of looping through every focal male for this timestep, identify if any males were copulating in the previous timestep and loop through them. If no males are copulating, this row in stat remains all zeroes.

#this gives the columns for which males were copulating, which corresponds to their id# if there are no males copulating, the loop will simply not run. 
cop<-grep("copulating", win.loss.census[[k]][(j-1),]) 
		
for(i in cop){

	
#want to start the loop for the stat after the first interaction is complete, but I want to start the accounting of the past interactions from the beginning

int.vector<-win.loss.census[[k]][1:(j-1),i]


#Remove the "fights" and "nothings" from the census list. This creates attack/receive and win/lose pairs in order of occurence
fight.times<-grep("fight",int.vector)
nothing.times<-grep("nothing",int.vector)

pairs<-int.vector[-c(fight.times, nothing.times)]

#need a way to do this that matches interaction partners but also is sensitive to the time-ordering 

receive.wins<-vector()

if(length(pairs)>0){

receive.wins<-unlist(sapply(1:ncol(census.list[[1]]), function(z){
	
	intset<-pairs[sapply(pairs, function(x) {
		
		strsplit(x,split=".",fixed=T)[[1]][3]==z
		
	})]
	
	names(intset)<-grep("TRUE",sapply(pairs, function(x) {
		
		strsplit(x,split=".",fixed=T)[[1]][3]==z
		
	}))
	
	receives<-grep("receive",intset)
	receivepluswin<-intset[receives+1][grep("win", intset[receives+1])]
	
	return(receivepluswin)
	
}))
}


#if the male has not had any interactions yet, the hazard is equal across event types involving them, so set all to zero 
if(length(receive.wins)>0){

#strip the "chase" from the end of chase wins so I can include them
receive.wins<-sapply(receive.wins, function(x) if(grepl("chas",x)==TRUE){
temp<-strsplit(x, split=".", fixed=T)[[1]]	
new<-paste(temp[1],temp[2],temp[3], sep=".")
return(new)
}
else return(x)
)


#get the number of interactions with each other male
counts<-table(receive.wins)	

#get just the interacting male id number
rankids<-sapply(names(counts), function(x) strsplit(x, split=".", fixed=T)[[1]][3])
	
#match with event types- this stat predicts new INCOMING attacks based on the number of previous OUTGOING attacks + WINS
#loop thru each element of rankids

for(l in 1:length(rankids)){

#incoming attacks directed at the focal male
stat[j,][paste(rankids[l],".attack.", i, sep="")]<-counts[l]
	}
		
}
	
}

#exclude attack events involving the X males 
stat[j,][endsWith(event.types,paste(".attack.",ncol(census.list[[1]]), sep="")) | startsWith(event.types,paste(ncol(census.list[[1]]),".attack.", sep=""))]<-0


}

#take the log 
stat<-log(1+stat)

new.stat[[k]][[1]]<-abind(new.stat[[k]][[1]], stat)

dimnames(new.stat[[k]][[1]])[[3]][ncol(new.stat[[k]][[1]][1,,])]<-"receive.win.count.pairwise.day.cop"

}

return(new.stat)

}


## log number of receive-losses?

add.receive.lose.count.pairwise.day.cop<-function(x){

new.stat<-x

library(abind)

#loop through each day sequence
for(k in day){

#empty matrix of the right dimensions, event# x event type. In this case, the default is all zeroes. 
stat<-matrix(0, nrow=step, ncol=length(event.types))
colnames(stat)<-event.types


#this loops through each event in the sequence

for(j in step){

#for each event, loop through all the males to get the recency ranks that will predict the next interaction- the support array will take care of males that weren't present that day.

#instead of looping through every focal male for this timestep, identify if any males were copulating in the previous timestep and loop through them. If no males are copulating, this row in stat remains all zeroes.

#this gives the columns for which males were copulating, which corresponds to their id# if there are no males copulating, the loop will simply not run. 
cop<-grep("copulating", win.loss.census[[k]][(j-1),]) 
		
for(i in cop){

	
#want to start the loop for the stat after the first interaction is complete, but I want to start the accounting of the past interactions from the beginning

int.vector<-win.loss.census[[k]][1:(j-1),i]


#Remove the "fights" and "nothings" from the census list. This creates attack/receive and win/lose pairs in order of occurence
fight.times<-grep("fight",int.vector)
nothing.times<-grep("nothing",int.vector)

pairs<-int.vector[-c(fight.times, nothing.times)]

#need a way to do this that matches interaction partners but also is sensitive to the time-ordering 

receive.losses<-vector()

if(length(pairs)>0){

receive.losses<-unlist(sapply(1:ncol(census.list[[1]]), function(z){
	
	intset<-pairs[sapply(pairs, function(x) {
		
		strsplit(x,split=".",fixed=T)[[1]][3]==z
		
	})]
	
	names(intset)<-grep("TRUE",sapply(pairs, function(x) {
		
		strsplit(x,split=".",fixed=T)[[1]][3]==z
		
	}))
	
	receives<-grep("receive",intset)
	receivepluslose<-intset[receives+1][grep("lose", intset[receives+1])]
	
	return(receivepluslose)
	
}))
}


#if the male has not had any interactions yet, the hazard is equal across event types involving them, so set all to zero 
if(length(receive.losses)>0){

#strip the "chase" from the end of chase wins so I can include them
receive.losses<-sapply(receive.losses, function(x) if(grepl("chas",x)==TRUE){
temp<-strsplit(x, split=".", fixed=T)[[1]]	
new<-paste(temp[1],temp[2],temp[3], sep=".")
return(new)
}
else return(x)
)


#get the number of interactions with each other male
counts<-table(receive.losses)	

#get just the interacting male id number
rankids<-sapply(names(counts), function(x) strsplit(x, split=".", fixed=T)[[1]][3])
	
#match with event types- this stat predicts new INCOMING attacks based on the number of previous OUTGOING attacks + WINS
#loop thru each element of rankids

for(l in 1:length(rankids)){

#incoming attacks directed at the focal male
stat[j,][paste(rankids[l],".attack.", i, sep="")]<-counts[l]
	}
		
}
	
}

#exclude attack events involving the X males 
stat[j,][endsWith(event.types,paste(".attack.",ncol(census.list[[1]]), sep="")) | startsWith(event.types,paste(ncol(census.list[[1]]),".attack.", sep=""))]<-0


}

#take the log 
stat<-log(1+stat)

new.stat[[k]][[1]]<-abind(new.stat[[k]][[1]], stat)

dimnames(new.stat[[k]][[1]])[[3]][ncol(new.stat[[k]][[1]][1,,])]<-"receive.lose.count.pairwise.day.cop"

}

return(new.stat)

}


### test the effects of pairwise win/loss record for the day

##Are you more likely to attack me if I have a positive win/loss record against you?


add.win.loss.pairwise.day.cop<-function(x){
	
rec.stat<-x

library(abind)

#loop through each day sequence
for(k in day){

#empty matrix of the right dimensions, event# x event type. In this case, the default is all zeroes. 
stat<-matrix(0, nrow=step, ncol=length(event.types))
colnames(stat)<-event.types


#this loops through each event in the sequence

for(j in step){

#for each event, loop through all the males to get the previous attack count that will predict the next interaction- the support array will take care of males that weren't present that day.

#instead of looping through every focal male for this timestep, identify if any males were copulating in the previous timestep and loop through them. If no males are copulating, this row in stat remains all zeroes.

#this gives the columns for which males were copulating, which corresponds to their id# if there are no males copulating, the loop will simply not run. 
cop<-grep("copulating", win.loss.census[[k]][(j-1),]) 
		
for(i in cop){
	
#start with simple +/-

#list of all the wins by the focal male
wins<-win.loss.census[[k]][1:(j-1),i][grep("win",win.loss.census[[k]][1:(j-1),i])]

#list of all the losses by the focal male
losses<-win.loss.census[[k]][1:(j-1),i][grep("lose",win.loss.census[[k]][1:(j-1),i])]

#if the male has not had any interactions yet, the hazard is equal across event types involving them, so set all to zero
#do wins first 
if(length(wins)>0){

#strip the "chase" from the end of chase wins so I can include them
wins<-sapply(wins, function(x) if(grepl("chas",x)==TRUE){
temp<-strsplit(x, split=".", fixed=T)[[1]]	
new<-paste(temp[1],temp[2],temp[3], sep=".")
return(new)
}
else return(x)
)


#totals for each type- these will be the values for the stat
win.tots<-table(wins)

#get just the interacting male id numbers
winids<-sapply(names(win.tots), function(x) strsplit(x, split=".", fixed=T)[[1]][3])

#match with event types- this stat predicts INCOMING attacks based on the difference of WINS and LOSSES of the FOCAL MALE

#loop thru each element of winids first

for(l in 1:length(winids)){

#incoming attacks directed at the focal male
stat[j,][paste(winids[l],".attack.", i, sep="")]<-win.tots[l]
	}
 
}

#now substract losses if there are any
if(length(losses)>0){

#strip the "chase" from the end of chase wins so I can include them
losses<-sapply(losses, function(x) if(grepl("chas",x)==TRUE){
temp<-strsplit(x, split=".", fixed=T)[[1]]	
new<-paste(temp[1],temp[2],temp[3], sep=".")
return(new)
}
else return(x)
)
	
	
lose.tots<-table(losses)
	
loseids<-sapply(names(lose.tots), function(x) strsplit(x, split=".", fixed=T)[[1]][3])

#loop thru each element of loseids, subtracting from whatever is there already

for(l in 1:length(loseids)){

#incoming attacks directed at the focal male
stat[j,][paste(loseids[l],".attack.", i, sep="")]<-stat[j,][paste(loseids[l],".attack.", i, sep="")]-lose.tots[l]
	}

}

}

#exclude attack events involving the X males 
stat[j,][endsWith(event.types,paste(".attack.",ncol(census.list[[1]]), sep="")) | startsWith(event.types,paste(ncol(census.list[[1]]),".attack.", sep=""))]<-0


}


rec.stat[[k]][[1]]<-abind(rec.stat[[k]][[1]], stat)

dimnames(rec.stat[[k]][[1]])[[3]][ncol(rec.stat[[k]][[1]][1,,])]<-"win.loss.pairwise.day.cop"

}

return(rec.stat)

}


##Are you more likely to attack me if I have a positive log(1+win/loss) record against you?


add.log.win.loss.pairwise.day.cop<-function(x){
	
rec.stat<-x

library(abind)

#loop through each day sequence
for(k in day){

#empty matrix of the right dimensions, event# x event type. In this case, the default is all zeroes. 
stat<-matrix(0, nrow=step, ncol=length(event.types))
colnames(stat)<-event.types


#this loops through each event in the sequence

for(j in step){

#for each event, loop through all the males to get the previous attack count that will predict the next interaction- the support array will take care of males that weren't present that day.

#instead of looping through every focal male for this timestep, identify if any males were copulating in the previous timestep and loop through them. If no males are copulating, this row in stat remains all zeroes.

#this gives the columns for which males were copulating, which corresponds to their id# if there are no males copulating, the loop will simply not run. 
cop<-grep("copulating", win.loss.census[[k]][(j-1),]) 
		
for(i in cop){
	
#start with simple +/-

#list of all the wins by the focal male
wins<-win.loss.census[[k]][1:(j-1),i][grep("win",win.loss.census[[k]][1:(j-1),i])]

#list of all the losses by the focal male
losses<-win.loss.census[[k]][1:(j-1),i][grep("lose",win.loss.census[[k]][1:(j-1),i])]

#if the male has not had any interactions yet, the hazard is equal across event types involving them, so set all to zero
#do wins first 
if(length(wins)>0){

#strip the "chase" from the end of chase wins so I can include them
wins<-sapply(wins, function(x) if(grepl("chas",x)==TRUE){
temp<-strsplit(x, split=".", fixed=T)[[1]]	
new<-paste(temp[1],temp[2],temp[3], sep=".")
return(new)
}
else return(x)
)


#totals for each type- these will be the values for the stat
win.tots<-table(wins)

#get just the interacting male id numbers
winids<-sapply(names(win.tots), function(x) strsplit(x, split=".", fixed=T)[[1]][3])

#match with event types- this stat predicts INCOMING attacks based on the difference of WINS and LOSSES of the FOCAL MALE

#loop thru each element of winids first

for(l in 1:length(winids)){

#incoming attacks directed at the focal male
stat[j,][paste(winids[l],".attack.", i, sep="")]<-win.tots[l]
	}
 
}

#now substract losses if there are any
if(length(losses)>0){

#strip the "chase" from the end of chase wins so I can include them
losses<-sapply(losses, function(x) if(grepl("chas",x)==TRUE){
temp<-strsplit(x, split=".", fixed=T)[[1]]	
new<-paste(temp[1],temp[2],temp[3], sep=".")
return(new)
}
else return(x)
)
	
	
lose.tots<-table(losses)
	
loseids<-sapply(names(lose.tots), function(x) strsplit(x, split=".", fixed=T)[[1]][3])

#loop thru each element of loseids, subtracting from whatever is there already

for(l in 1:length(loseids)){

#incoming attacks directed at the focal male
stat[j,][paste(loseids[l],".attack.", i, sep="")]<-stat[j,][paste(loseids[l],".attack.", i, sep="")]-lose.tots[l]
	}

}

}

#exclude attack events involving the X males 
stat[j,][endsWith(event.types,paste(".attack.",ncol(census.list[[1]]), sep="")) | startsWith(event.types,paste(ncol(census.list[[1]]),".attack.", sep=""))]<-0


}

#take the log since these numbers can become quite large and we donot expect the hazard to increase exponentially forever with win-loss record. Have to do some fancy stuff to deal with the negative records

stat<-(log(1+abs(stat))/(stat))*abs(stat) 
stat[is.na(stat)]<-0

rec.stat[[k]][[1]]<-abind(rec.stat[[k]][[1]], stat)

dimnames(rec.stat[[k]][[1]])[[3]][ncol(rec.stat[[k]][[1]][1,,])]<-"log.win.loss.pairwise.day.cop"

}

return(rec.stat)

}


## Are you more likely to attack me if we are more evenly matched (our win/loss record is closer to 0)?

#just take the absolute value of the win/loss record. Smaller values mean more evenly matched males


add.abs.winloss.diff.pairwise.day.cop<-function(x){
	
rec.stat<-x

library(abind)

#loop through each day sequence
for(k in day){

#empty matrix of the right dimensions, event# x event type. In this case, the default is all zeroes. 
stat<-matrix(0, nrow=step, ncol=length(event.types))
colnames(stat)<-event.types


#this loops through each event in the sequence

for(j in step){

#for each event, loop through all the males to get the previous attack count that will predict the next interaction- the support array will take care of males that weren't present that day.

#instead of looping through every focal male for this timestep, identify if any males were copulating in the previous timestep and loop through them. If no males are copulating, this row in stat remains all zeroes.

#this gives the columns for which males were copulating, which corresponds to their id# if there are no males copulating, the loop will simply not run. 
cop<-grep("copulating", win.loss.census[[k]][(j-1),]) 
		
for(i in cop){
	
#start with simple +/-

#list of all the wins by the focal male
wins<-win.loss.census[[k]][1:(j-1),i][grep("win",win.loss.census[[k]][1:(j-1),i])]

#list of all the losses by the focal male
losses<-win.loss.census[[k]][1:(j-1),i][grep("lose",win.loss.census[[k]][1:(j-1),i])]

#if the male has not had any interactions yet, the hazard is equal across event types involving them, so set all to zero
#do wins first 
if(length(wins)>0){

#strip the "chase" from the end of chase wins so I can include them
wins<-sapply(wins, function(x) if(grepl("chas",x)==TRUE){
temp<-strsplit(x, split=".", fixed=T)[[1]]	
new<-paste(temp[1],temp[2],temp[3], sep=".")
return(new)
}
else return(x)
)


#totals for each type- these will be the values for the stat
win.tots<-table(wins)

#get just the interacting male id numbers
winids<-sapply(names(win.tots), function(x) strsplit(x, split=".", fixed=T)[[1]][3])

#match with event types- this stat predicts INCOMING attacks based on the difference of WINS and LOSSES of the FOCAL MALE

#loop thru each element of winids first

for(l in 1:length(winids)){

#incoming attacks directed at the focal male
stat[j,][paste(winids[l],".attack.", i, sep="")]<-win.tots[l]
	}
 
}

#now substract losses if there are any
if(length(losses)>0){

#strip the "chase" from the end of chase wins so I can include them
losses<-sapply(losses, function(x) if(grepl("chas",x)==TRUE){
temp<-strsplit(x, split=".", fixed=T)[[1]]	
new<-paste(temp[1],temp[2],temp[3], sep=".")
return(new)
}
else return(x)
)
	
	
lose.tots<-table(losses)
	
loseids<-sapply(names(lose.tots), function(x) strsplit(x, split=".", fixed=T)[[1]][3])

#loop thru each element of loseids, subtracting from whatever is there already

for(l in 1:length(loseids)){

#incoming attacks directed at the focal male
stat[j,][paste(loseids[l],".attack.", i, sep="")]<-stat[j,][paste(loseids[l],".attack.", i, sep="")]-lose.tots[l]
	}

}

}

#exclude attack events involving the X males 
stat[j,][endsWith(event.types,paste(".attack.",ncol(census.list[[1]]), sep="")) | startsWith(event.types,paste(ncol(census.list[[1]]),".attack.", sep=""))]<-0


}

#take the absolute value of the stat matrix to turn all negative win/loss records positive so that larger numbers jsut mean farther from evenly matched

stat<-abs(stat)

rec.stat[[k]][[1]]<-abind(rec.stat[[k]][[1]], stat)

dimnames(rec.stat[[k]][[1]])[[3]][ncol(rec.stat[[k]][[1]][1,,])]<-"abs.winloss.diff.pairwise.day.cop"

}

return(rec.stat)

}


## try log(evenly.matched)

add.log.abs.winloss.diff.pairwise.day.cop<-function(x){
	
rec.stat<-x

library(abind)

#loop through each day sequence
for(k in day){

#empty matrix of the right dimensions, event# x event type. In this case, the default is all zeroes. 
stat<-matrix(0, nrow=step, ncol=length(event.types))
colnames(stat)<-event.types


#this loops through each event in the sequence

for(j in step){

#for each event, loop through all the males to get the previous attack count that will predict the next interaction- the support array will take care of males that weren't present that day.

#instead of looping through every focal male for this timestep, identify if any males were copulating in the previous timestep and loop through them. If no males are copulating, this row in stat remains all zeroes.

#this gives the columns for which males were copulating, which corresponds to their id# if there are no males copulating, the loop will simply not run. 
cop<-grep("copulating", win.loss.census[[k]][(j-1),]) 
		
for(i in cop){
	
#start with simple +/-

#list of all the wins by the focal male
wins<-win.loss.census[[k]][1:(j-1),i][grep("win",win.loss.census[[k]][1:(j-1),i])]

#list of all the losses by the focal male
losses<-win.loss.census[[k]][1:(j-1),i][grep("lose",win.loss.census[[k]][1:(j-1),i])]

#if the male has not had any interactions yet, the hazard is equal across event types involving them, so set all to zero
#do wins first 
if(length(wins)>0){

#strip the "chase" from the end of chase wins so I can include them
wins<-sapply(wins, function(x) if(grepl("chas",x)==TRUE){
temp<-strsplit(x, split=".", fixed=T)[[1]]	
new<-paste(temp[1],temp[2],temp[3], sep=".")
return(new)
}
else return(x)
)


#totals for each type- these will be the values for the stat
win.tots<-table(wins)

#get just the interacting male id numbers
winids<-sapply(names(win.tots), function(x) strsplit(x, split=".", fixed=T)[[1]][3])

#match with event types- this stat predicts INCOMING attacks based on the difference of WINS and LOSSES of the FOCAL MALE

#loop thru each element of winids first

for(l in 1:length(winids)){

#incoming attacks directed at the focal male
stat[j,][paste(winids[l],".attack.", i, sep="")]<-win.tots[l]
	}
 
}

#now substract losses if there are any
if(length(losses)>0){

#strip the "chase" from the end of chase wins so I can include them
losses<-sapply(losses, function(x) if(grepl("chas",x)==TRUE){
temp<-strsplit(x, split=".", fixed=T)[[1]]	
new<-paste(temp[1],temp[2],temp[3], sep=".")
return(new)
}
else return(x)
)
	
	
lose.tots<-table(losses)
	
loseids<-sapply(names(lose.tots), function(x) strsplit(x, split=".", fixed=T)[[1]][3])

#loop thru each element of loseids, subtracting from whatever is there already

for(l in 1:length(loseids)){

#incoming attacks directed at the focal male
stat[j,][paste(loseids[l],".attack.", i, sep="")]<-stat[j,][paste(loseids[l],".attack.", i, sep="")]-lose.tots[l]
	}

}

}

#exclude attack events involving the X males 
stat[j,][endsWith(event.types,paste(".attack.",ncol(census.list[[1]]), sep="")) | startsWith(event.types,paste(ncol(census.list[[1]]),".attack.", sep=""))]<-0


}

#take the absolute value of the stat matrix to turn all negative win/loss records positive so that larger numbers jsut mean farther from evenly matched

stat<-abs(stat)

stat<-log(1+stat)

rec.stat[[k]][[1]]<-abind(rec.stat[[k]][[1]], stat)

dimnames(rec.stat[[k]][[1]])[[3]][ncol(rec.stat[[k]][[1]][1,,])]<-"log.abs.winloss.diff.pairwise.day.cop"

}

return(rec.stat)

}


############# General Effects, Day-specific ############

#cumulative day-specific stats that affect the hazard of being attacked in general rather than by specific individuals
#some of these correlate highly with a metric like eigenvector centrality: degree and edgeweight

## Am I more likely to be attacked if I have had a larger number of interaction partners today? (degree)

add.degree.day.cop<-function(x){
	
rec.stat<-x

library(abind)

#loop through each day sequence
for(k in day){

#empty matrix of the right dimensions, event# x event type. In this case, the default is all zeroes. 
stat<-matrix(0, nrow=step, ncol=length(event.types))
colnames(stat)<-event.types


#this loops through each event in the sequence

for(j in step){

#for each event, loop through all the males to get the previous attack count that will predict the next interaction- the support array will take care of males that weren't present that day.

#instead of looping through every focal male for this timestep, identify if any males were copulating in the previous timestep and loop through them. If no males are copulating, this row in stat remains all zeroes.

#this gives the columns for which males were copulating, which corresponds to their id# if there are no males copulating, the loop will simply not run. 
cop<-grep("copulating", win.loss.census[[k]][(j-1),]) 
		
for(i in cop){
	
#get number of unique interaction partners

#list of all the wins by the focal male
wins<-grep("win",win.loss.census[[k]][1:(j-1),i])

#list of all the losses by the focal male
losses<-grep("lose",win.loss.census[[k]][1:(j-1),i])

#this covers all the interactions that have happened so far for the focal male. Stick them all together
ints<-union(wins,losses)

#if the male has not had any interactions yet, the hazard is equal across event types involving them, so set all to zero
#do wins first 
if(length(ints)>0){

#totals for each type- these will be the values for the stat
int.names<-win.loss.census[[k]][1:(j-1),i][ints]

#get just the interacting male id numbers
intids<-sapply(int.names, function(x) strsplit(x, split=".", fixed=T)[[1]][3])

#collapse to unique parters
intids<-unique(intids)

#match with event types- this stat predicts general INCOMING attacks based on the total number of interaction partners so far that day for the focal male (degree)

stat[j,][endsWith(event.types,paste(".attack.",i, sep=""))]<-length(intids)
}


}

#exclude attack events involving the X males 
stat[j,][endsWith(event.types,paste(".attack.",ncol(census.list[[1]]), sep="")) | startsWith(event.types,paste(ncol(census.list[[1]]),".attack.", sep=""))]<-0


}


rec.stat[[k]][[1]]<-abind(rec.stat[[k]][[1]], stat)

dimnames(rec.stat[[k]][[1]])[[3]][ncol(rec.stat[[k]][[1]][1,,])]<-"degree.day.cop"

}

return(rec.stat)

}


##try log(degree) just to see

add.log.degree.day.cop<-function(x){
	
rec.stat<-x

library(abind)

#loop through each day sequence
for(k in day){

#empty matrix of the right dimensions, event# x event type. In this case, the default is all zeroes. 
stat<-matrix(0, nrow=step, ncol=length(event.types))
colnames(stat)<-event.types


#this loops through each event in the sequence

for(j in step){

#for each event, loop through all the males to get the previous attack count that will predict the next interaction- the support array will take care of males that weren't present that day.

#instead of looping through every focal male for this timestep, identify if any males were copulating in the previous timestep and loop through them. If no males are copulating, this row in stat remains all zeroes.

#this gives the columns for which males were copulating, which corresponds to their id# if there are no males copulating, the loop will simply not run. 
cop<-grep("copulating", win.loss.census[[k]][(j-1),]) 
		
for(i in cop){
	
#get number of unique interaction partners

#list of all the wins by the focal male
wins<-grep("win",win.loss.census[[k]][1:(j-1),i])

#list of all the losses by the focal male
losses<-grep("lose",win.loss.census[[k]][1:(j-1),i])

#this covers all the interactions that have happened so far for the focal male. Stick them all together
ints<-union(wins,losses)

#if the male has not had any interactions yet, the hazard is equal across event types involving them, so set all to zero
#do wins first 
if(length(ints)>0){

#totals for each type- these will be the values for the stat
int.names<-win.loss.census[[k]][1:(j-1),i][ints]

#get just the interacting male id numbers
intids<-sapply(int.names, function(x) strsplit(x, split=".", fixed=T)[[1]][3])

#collapse to unique parters
intids<-unique(intids)

#match with event types- this stat predicts general INCOMING attacks based on the total number of interaction partners so far that day for the focal male (degree)

stat[j,][endsWith(event.types,paste(".attack.",i, sep=""))]<-length(intids)
}


}

#exclude attack events involving the X males 
stat[j,][endsWith(event.types,paste(".attack.",ncol(census.list[[1]]), sep="")) | startsWith(event.types,paste(ncol(census.list[[1]]),".attack.", sep=""))]<-0


}

stat<-log(1+stat)

rec.stat[[k]][[1]]<-abind(rec.stat[[k]][[1]], stat)

dimnames(rec.stat[[k]][[1]])[[3]][ncol(rec.stat[[k]][[1]][1,,])]<-"log.degree.day.cop"

}

return(rec.stat)

}


## Am I more likely to be attacked if I have a larger number of other males that I have positive win records with? (win degree)

add.win.degree.day.cop<-function(x){
	
rec.stat<-x

library(abind)

#loop through each day sequence
for(k in day){

#empty matrix of the right dimensions, event# x event type. In this case, the default is all zeroes. 
stat<-matrix(0, nrow=step, ncol=length(event.types))
colnames(stat)<-event.types


#this loops through each event in the sequence

for(j in step){

#for each event, loop through all the males to get the previous attack count that will predict the next interaction- the support array will take care of males that weren't present that day.

#instead of looping through every focal male for this timestep, identify if any males were copulating in the previous timestep and loop through them. If no males are copulating, this row in stat remains all zeroes.

#this gives the columns for which males were copulating, which corresponds to their id# if there are no males copulating, the loop will simply not run. 
cop<-grep("copulating", win.loss.census[[k]][(j-1),]) 
		
for(i in cop){

	
#start with simple +/-

#list of all the wins by the focal male
wins<-grep("win",win.loss.census[[k]][1:(j-1),i])

#list of all the losses by the focal male
losses<-grep("lose",win.loss.census[[k]][1:(j-1),i])

win.record<-rep(0,ncol(census.list[[1]]))

#if the male has not had any interactions yet, the hazard is equal across event types involving them, so set all to zero
#do wins first 
if(length(wins)>0){

wins<-win.loss.census[[k]][1:(j-1),i][wins]

#strip the "chase" from the end of chase wins so I can include them
wins<-sapply(wins, function(x) if(grepl("chas",x)==TRUE){
temp<-strsplit(x, split=".", fixed=T)[[1]]	
new<-paste(temp[1],temp[2],temp[3], sep=".")
return(new)
}
else return(x)
)

#totals for each type
win.tots<-table(wins)

#get just the interacting male id numbers
winids<-sapply(names(win.tots), function(x) strsplit(x, split=".", fixed=T)[[1]][3])

win.record[as.numeric(winids)]<-win.tots
}


#now substract losses if there are any
if(length(losses)>0){

losses<-win.loss.census[[k]][1:(j-1),i][losses]

#strip the "chase" from the end of chase wins so I can include them
losses<-sapply(losses, function(x) if(grepl("chas",x)==TRUE){
temp<-strsplit(x, split=".", fixed=T)[[1]]	
new<-paste(temp[1],temp[2],temp[3], sep=".")
return(new)
}
else return(x)
)
	
	
lose.tots<-table(losses)
	
loseids<-sapply(names(lose.tots), function(x) strsplit(x, split=".", fixed=T)[[1]][3])

win.record[as.numeric(loseids)]<-win.record[as.numeric(loseids)]-lose.tots

} 
 
#test if the win records are positive
win.pos<-win.record>0 

#Get the total number of males with whom the focal male has positive win records 
if(is.na(table(win.pos)["TRUE"])){win.pos.num<-0} else {win.pos.num<-table(win.pos)["TRUE"]}

#match with event types- this stat predicts general INCOMING attacks based on the total number of interaction partners so far that day that the focal male has positive win records with (win degree)

stat[j,][endsWith(event.types,paste(".attack.",i, sep=""))]<-win.pos.num

}

#exclude attack events involving the X males 
stat[j,][endsWith(event.types,paste(".attack.",ncol(census.list[[1]]), sep="")) | startsWith(event.types,paste(ncol(census.list[[1]]),".attack.", sep=""))]<-0


}


rec.stat[[k]][[1]]<-abind(rec.stat[[k]][[1]], stat)

dimnames(rec.stat[[k]][[1]])[[3]][ncol(rec.stat[[k]][[1]][1,,])]<-"win.degree.day.cop"

}

return(rec.stat)
	
	
}


##check log win degree

add.log.win.degree.day.cop<-function(x){
	
rec.stat<-x

library(abind)

#loop through each day sequence
for(k in day){

#empty matrix of the right dimensions, event# x event type. In this case, the default is all zeroes. 
stat<-matrix(0, nrow=step, ncol=length(event.types))
colnames(stat)<-event.types


#this loops through each event in the sequence

for(j in step){

#for each event, loop through all the males to get the previous attack count that will predict the next interaction- the support array will take care of males that weren't present that day.

#instead of looping through every focal male for this timestep, identify if any males were copulating in the previous timestep and loop through them. If no males are copulating, this row in stat remains all zeroes.

#this gives the columns for which males were copulating, which corresponds to their id# if there are no males copulating, the loop will simply not run. 
cop<-grep("copulating", win.loss.census[[k]][(j-1),]) 
		
for(i in cop){

	
#start with simple +/-

#list of all the wins by the focal male
wins<-grep("win",win.loss.census[[k]][1:(j-1),i])

#list of all the losses by the focal male
losses<-grep("lose",win.loss.census[[k]][1:(j-1),i])

win.record<-rep(0,ncol(census.list[[1]]))

#if the male has not had any interactions yet, the hazard is equal across event types involving them, so set all to zero
#do wins first 
if(length(wins)>0){

wins<-win.loss.census[[k]][1:(j-1),i][wins]

#strip the "chase" from the end of chase wins so I can include them
wins<-sapply(wins, function(x) if(grepl("chas",x)==TRUE){
temp<-strsplit(x, split=".", fixed=T)[[1]]	
new<-paste(temp[1],temp[2],temp[3], sep=".")
return(new)
}
else return(x)
)

#totals for each type
win.tots<-table(wins)

#get just the interacting male id numbers
winids<-sapply(names(win.tots), function(x) strsplit(x, split=".", fixed=T)[[1]][3])

win.record[as.numeric(winids)]<-win.tots
}


#now substract losses if there are any
if(length(losses)>0){

losses<-win.loss.census[[k]][1:(j-1),i][losses]
```
