## Supplemental_reproducible_code for "Fighting isn’t sexy in lekking Greater Sage-grouse (*Centrocercus urophasianus*)": rem_event_hist_simulation_cluster_udata_terr_actualday_newcensus.R

```
	
rm(list=ls())

###make sure necessary packages are there and loaded

packages<-c("abind","relevent","lubridate","DescTools","gtools")

package.check<-lapply(packages, FUN = function(x) {
	if (!require(x, character.only = TRUE, quietly=TRUE)) {
		install.packages(x, dependencies = TRUE)
		library(x, character.only = TRUE, quietly=TRUE)
	} 

})


#!/usr/bin/env Rscript
args=as.numeric(commandArgs(trailingOnly=TRUE))

#args<-c(18,1)

## Choose the day to model

mod.day<-args[1]

if(mod.day>1){
yester.day<-(mod.day-1)
} else {yester.day<-mod.day}

#day 9 issue is taken care of inside the stat functions 

#code to run to ensure a copulation in the demo sim

#make a dummy list to start
# event.list.split<-list(1,data.frame(Events="none"))
# while(!(TRUE %in% grepl("cop.start",event.list.split[[2]]$Events))){

#run to reset everything
{
#make the base event list and intercept array


event.list<-read.table(file="relevent_format_dataset_udata_wincol_disturbance.tsv", header=T,  sep="\t")

event.list$Date<-as.factor(event.list$Date)  

event.types<-read.table(file="event_types_list_udata.csv", header=F,  sep=",")
event.types<-as.character(event.types$V1)


#NOTE- the event.types vector now contains ENDOGENOUS event types as well as EXOGENOUS types: arrivals and clock events, and end.obs. Need to make sure everything is robust to this. 
### create eventlist for rem ###


event.list.dat<-split(event.list, event.list$Date)


#remove female context for now

basic.event.list<-event.list[-3]

#strip the "loser" from the disengage events so they are all the same event "type" and they match my previous code

basic.event.list$Events<-sapply(basic.event.list$Events, function(x) {
	
	event<-unlist(strsplit(as.character(x),".",fixed=T))
	if(event[2]=="disengage"){
		return(paste(event[1],event[2],sep="."))
	} else {return(as.character(x))}
}
)


#add factor levels for event types that don't actually happen (and put them in the same order as the event.type list) to maintain bookkeeping 

basic.event.list$Events<-as.character(basic.event.list$Events)

basic.event.list$Events<-factor(basic.event.list$Events, levels=as.character(event.types))

#turn event types into numeric values

basic.event.list$Events<-as.numeric(basic.event.list$Events)

#set the event type code for exogenous events to zero. I have there all at the trailing end of the vector

endogenous<-(length(event.types)-length(event.types[grep("arrive",event.types)])-5)

basic.event.list$Events[basic.event.list$Events>endogenous]<-0

rownames(basic.event.list)<-NULL
#need to divide the data up by grouping factor

evlist<-list()
evlist<-split(basic.event.list, basic.event.list$Date)

#strip the date column from the sub-dataframes so they're just two columns, event and time

evlist<-lapply(evlist, function(x) x<-x[-3])

#convert from dataframes to matrices

evlist<-lapply(evlist, function(x) x<-as.matrix(x))

## make event.types vector usable for intlist, etc. by excising exogenous event types

event.types<-event.types[1:endogenous]

#grab the max time for the day we are simulating
tday<-event.list.dat[[mod.day]][nrow(event.list.dat[[mod.day]]),2]


#extract cop.start data to use for the cumulative effect of cops on sol

#want all the cops by male from the beginning thru "yester.day"

#if the mod.day is day 1, then the function will ignore coplist. It just needs to have some value to avoid errors.

coplist<-list()

for(i in 1:yester.day){

coplist[[i]]<-as.character(event.list.dat[[i]][,"Events"][grep(".cop.start", event.list.dat[[i]][,"Events"])])

}

coplist<-unlist(coplist)


##########

#other important stuff to load
## format female exact time to use in simulation. Females will flip from present to absent at the average #time of first departure
fe_exact<-read.table("CHG_females_exact.tsv", header=TRUE,sep="\t",stringsAsFactors=FALSE)

#reduce to only the necessary columns- this time I want time elapsed
fe_exact<-fe_exact[,c(1,3,5,7,9,11,13)]

#turn the first arrival NA's to 0s
fe_exact$first_arrival_elapsed[is.na(fe_exact$first_arrival_elapsed)]<-"00:00:00:00"

library(lubridate)
options(digits.secs=6)

#convert number of frames to partial seconds
for(h in 2:7){
fe_exact[,h]<-sapply(fe_exact[,h], function(x) {

fem.split<-unlist(strsplit(x,split=":", fixed=TRUE))
suppressWarnings(period_to_seconds(hms(paste(fem.split[1],fem.split[2],(as.numeric(fem.split[3])+as.numeric(fem.split[4])/30), sep=":"))))

}
)
}

#mean(fe_exact$first_depart_elapsed)
#[1] 4377.154

#simulation version of fe_exact that will still work inside my stat functions

fe_exact<-fe_exact[c(yester.day,mod.day),]
#fe_sim[,3]<-4377.154

#get the corresponding female exogenous events
fem.vect<-event.list.dat[[mod.day]][startsWith(as.character(event.list.dat[[mod.day]]$Events),"fem"),][,c(1,2)]

#bring in day-level male presence data
male.pres<-read.table("CHG_2014_male_presence_data_quest.csv", sep=",", header=T)
#male IDs are already in ascending order, so I can convert them directly
colnames(male.pres)<-c("Date", 1:30)


#get territory-holder data for the terr effects functions so they don't have to look back two days inside the functions

#are you here today AND ALSO both of the past two days? (excluding X males)

#add attendance data from 3/17
extra<-rep(0,29)
names(extra)<-1:29
extra[c(1,2,3,4,5,7)]<-1

terr.list<-list()

for(k in 1:length(event.list.dat)){

if(k==1 | k==2){
	
	if(k==1){
		
		terr.pres<-rbind(extra,extra)
		terr.pres<-rbind(terr.pres,male.pres[k,2:(ncol(male.pres)-1)])
		
		terr.holders<-which(apply(terr.pres,2,function(x) sum(x))==3)
		
	}
	
	if(k==2){
		
	
		terr.pres<-rbind(extra,male.pres[c(k-1,k),2:(ncol(male.pres)-1)])
		
		terr.holders<-which(apply(terr.pres,2,function(x) sum(x))==3)
		
	}
	
	
} else {
	
	terr.holders<-which(apply(male.pres[(k-2):k,2:(ncol(male.pres)-1)],2,function(x) sum(x))==3)
	
}

terr.list[[k]]<-terr.holders

}


#pare down to the two days in question
male.pres<-male.pres[c(yester.day,mod.day),]

terr.list<-terr.list[c(yester.day,mod.day)]

#now that I have depart events as endogenous features, and I don't have the whole eventlist at the outset, I should be producing a new pres.exact every step

#make the base pres.exact object- in this case I will have the arrivals at the actual times, with the exception of subsequent X male arriveals  


pres.exact<-read.table("male_pres_exact_quest.csv", sep=",",header=T)

	departlist<-list()
	
	for(j in 1:nrow(pres.exact)){
		
		arriverow<-pres.exact[j,startsWith(colnames(pres.exact),"arrive")]
		arriverow<-arriverow[!is.na(arriverow)]
		
		arrives<-t(sapply(arriverow, function(x) return(c(paste0(pres.exact$Num_ID[j],".arrive"),x,as.character(pres.exact$Date[j])))))
		
		departrow<-pres.exact[j,startsWith(colnames(pres.exact),"depart")]
		departrow<-departrow[!is.na(departrow)]
		
		departs<-t(sapply(departrow, function(x) return(c(paste0(pres.exact$Num_ID[j],".depart"),x,as.character(pres.exact$Date[j])))))
		
		allpres<-rbind(arrives,departs)
		
		departlist[[j]]<-allpres
		
	}

#collapse the departlist 

departevent<-abind(departlist, along=1)

departevent<-data.frame(departevent)
names(departevent)<-c("Events","Time", "Date")

departevent$Time<-as.numeric(as.character(departevent$Time))

library(abind)
arrstr<-abind(strsplit(as.character(departevent$Events),".",fixed=T),along=2)

departevent$Bird_ID<-arrstr[1,]
departevent$Type<-arrstr[2,]

departevent$Type<-as.factor(departevent$Type)
departevent$Bird_ID<-as.factor(departevent$Bird_ID)

# #split into day-specific data frames
pres.exact.dat<-split(departevent, departevent$Date)

pres.exact.dat<-lapply(pres.exact.dat, function(x) {
	
	x<-x[order(x$Time),]
	
	return(x)
})


#are there any males that leave and then return? Get the amount of time between the departure and return so I can model it in case they depart in the simulation. 

#mean waiting time between departures and new arrivals by ID across the whole dataset


meanarr<-list()

for(m in 1:30){

things<-pres.exact[grep(m,pres.exact[,2]),-c(1:5)]

meanlist<-list()

for(i in 1:nrow(pres.exact[grep(m,pres.exact[,2]),-c(1:5)])){
	
	meanlist[[i]]<-list()
	
	for(j in 1:((length(things[i,][!is.na(things[i,])])/2)-1)){
			
			if(j!=0){
					
			meanlist[[i]][j]<-things[i,j*2+1]-things[i,(j*2)]
			
		}	
			
		}
	
}

meanarr[[m]]<-mean(unlist(meanlist)[!is.na(unlist(meanlist))])

names(meanarr[[m]])<-paste0(m,".arr")

}

meanarr<-unlist(meanarr)

#use these average waiting times for when these males leave the lek. 


###pare down to the days in question. Again, issues of actual day 1 or 9 for yesterday are taken care of inside the stat functions

pres.exact.dat<-list(pres.exact.dat[[yester.day]],pres.exact.dat[[mod.day]])

dates<-c(as.character(pres.exact.dat[[1]][1,3]),as.character(pres.exact.dat[[2]][1,3]))


#remove departs because I am modeling them endogenously
#I will have to make rules to make sure they don't happen impossibly

pres.exact.dat[[2]]<-pres.exact.dat[[2]][!endsWith(as.character(pres.exact.dat[[2]]$Events),".depart"),]

#retain only the first instace of each male's arrival
pres.exact.dat[[2]]<-pres.exact.dat[[2]][match(unique(pres.exact.dat[[2]]$Events),pres.exact.dat[[2]]$Events),]


#vector of scheduled arrivals after the beginning
arr.vect<-pres.exact.dat[[2]][pres.exact.dat[[2]]$Time>0,c(1,2)]

#since the arrivals that happen after t=0 get incorporated into the exo.vect, and then later into the pres.exact.dat matrix as it grows, I actually just want only arrivals at t=0 for this.

pres.exact.dat[[2]]<-pres.exact.dat[[2]][pres.exact.dat[[2]]$Time==0,]


### pairwise centroid distances

file_list<-list.files("./pairwise_dist")

distlist<-list()

distlist<-lapply(file_list, function(x) read.table(paste("./pairwise_dist/",x,sep=""), sep=","))


#convert to martices

distlist.mat<-lapply(distlist, function(x) as.matrix(x, ))

#the names of the rows and columns are in the same order as the numerical ids passed to rem, so they can be used to index the matrix. Males that are absent on a given day have NA's for their centroid distances. I can change them to zeroes here so they don't produce errors, but the support array will take care of them anyway.

for(i in 1:length(distlist.mat)){
distlist.mat[[i]][is.na(distlist.mat[[i]])]<-0
}

#for the sim, pare down to the two days in question

dist.one<-distlist.mat[[yester.day]]
dist.two<-distlist.mat[[mod.day]]

#replace the distlist.mat object with the special sim version so it will work with my functions

distlist.mat<-list(dist.one,dist.two)


#####

#coefficients from the fitted REM 

mod_coef<-read.table("rem_model_coef", sep=",")


####

#source the rest of my fuctions for predictor stats
#enable all the functions I have so far
source('rem.functions.act.terr.sim.R', chdir = TRUE)

#make a list of the functions
fun.list<-ls()[grep("add.",ls())]
        
#bring in the function codes from the best result

prev.best<-read.table("previous_best_remove_226_283_at_the_end", sep=",")

prev.best.top<-as.character(prev.best[nrow(prev.best),2])
args.all=as.numeric(unlist(strsplit(prev.best.top, split="_", fixed=TRUE)))

#args.all<-c(256,257,258,259,260,261,30,29)

arg.use<-c(1:length(args.all))


#############


### create eventlist for rem ###

##THIS is basically the object I will be growing in the sim. 

#for simulating the first day, which doesn't have a yesterday, it's a little different. I simply need to rename the dates column so that the dataframe can be split

#create the seed for the evlist object

tempev<-list()
tempev[[1]]<-evlist[[yester.day]]
tempev[[2]]<-matrix(c(1,1),nrow=1,ncol=2)
colnames(tempev[[1]])<-c("Events","Time")
colnames(tempev[[2]])<-c("Events","Time")

#not sure this matters but I'm afraid of screwing everything up
#names(tempev)<-dates

evlist<-tempev


#vector to index date names
if(mod.day==1){dates[1]<-paste0("1a-",dates[1])}
days<-dates

#create the seed for the event.list object

eventtemp<-event.list.dat[[yester.day]]

##if I want to start with simday 1:

# firstevent<-matrix(nrow=1,ncol=4)
# firstevent[,3]<-1
# firstevent[,4]<-days[2]
# colnames(firstevent)<-c("Events","Time","Females","Date")

# event.list<-rbind(eventtemp,firstevent)

event.list<-eventtemp
event.list$Events<-as.character(event.list$Events)
event.list$Females<-as.numeric(event.list$Females)
event.list$Time<-as.numeric(event.list$Time)
event.list$Date<-rep(days[1],nrow(event.list))

#dummy vector for saving hazards
overall.haz.dat<-rep(0,length(event.types))

#set up a data frame of all the currently scheduled exogenous events

exo.vect<-rbind(arr.vect,fem.vect)

#add "clock" events

clocknum<-floor(tday/900)

clock.vect<-data.frame(Events=rep("clock.time",clocknum),Time=900*(1:clocknum),stringsAsFactors=F)

exo.vect<-rbind(exo.vect,clock.vect)


## bring in the actual exogenous disturbance events that casued the birds to flush for that day if there were any

disturb<-read.table("CHG_2014_disturbance_events.csv", header=T,sep=",") 

if(!is.na(disturb[mod.day,]$Time)){
	
	exo.vect<-rbind(exo.vect,c("disturb.ance",as.character(disturb[mod.day,]$Time)))
	
}


exo.vect$Time<-as.numeric(exo.vect$Time)
exo.vect<-exo.vect[order(exo.vect$Time),]
exo.vect$Events<-as.character(exo.vect$Events)

#if I want to bring in a specific event history up to a certain point and then do an intervention, I can just seed with these objects instead of the ones I created above. I would then just set the initial "step" to nrow evlist[[day]]


}


##Daily Scenario

step<-1
day<-2
Females<-1
t<-0
tmax<-tday
#tmax<-8021.944
#tmax<-500

####

#schedule exogenous events- this is taken care of by the exo.vect object. Other return-arrives are governed by the meanarr vector

########### Start the simulation loop here #######

#library(tictoc)
#tic()
while(day<2 | t<tmax){


#code for simulating multiple days:
#if t is past tmax, flip the parameters over to start day 2
# if(t>=tmax && day==1){
	
	# t<-0
	# day<-2
	# step<-1
	# Females<-2
	# x.arr<-rexp(1,1/1720.291)
	# #x.arr<-120
# }


#append the new row to event.list here with the female presence info, and dummy data for the rest
if(!(day==1 & step==1)){
	
	event.list<-rbind(event.list, c("NA","NA",Females,days[day]))
	event.list$Females<-as.numeric(event.list$Females)
	suppressWarnings(event.list$Time<-as.numeric(event.list$Time))
	
	if((TRUE %in% grepl("depart",event.list[nrow(event.list)-1,1])) | (TRUE %in% grepl("arrive",event.list[nrow(event.list)-1,1]))){
		
		ar.row<-event.list[nrow(event.list)-1,]
		newline<-unlist(c(ar.row[1],ar.row[2],ar.row[4],strsplit(as.character(ar.row[1]),".",fixed=T)[[1]][1],strsplit(as.character(ar.row[1]),".",fixed=T)[[1]][2]))
		
		names(newline)<-c("Events","Time","Date","Bird_ID","Type")
		
		pres.exact.dat[[day]]<-rbind(pres.exact.dat[[day]],newline)
		
		pres.exact.dat[[day]]$Time<-as.numeric(pres.exact.dat[[day]]$Time)
		
	}
	
	
	}


##set new .arr time if particular males have departed. Append to the exo.vect?

if(step>1){
if(endsWith(as.character(event.list[nrow(event.list)-1,1]),".depart") && !(day==2 & step==1)){
	
	#match the departed male with an arrive hazard, if it exists. 
	
	depev<-strsplit(as.character(event.list[nrow(event.list)-1,1]),".",fixed=T)[[1]][1]
	
	if(!is.na(meanarr[paste0(depev,".arr")])){
		
		#Schedule a return based on the mean waiting time for that male
		
		tarr<-t+rexp(1,1/meanarr[paste0(depev,".arr")])
		
		#append the new scheduled exogenous event to the exo.vect
	
	exo.vect<-rbind(exo.vect,c(paste0(depev,".arrive"),tarr))
	exo.vect$Time<-as.numeric(exo.vect$Time)
	exo.vect<-exo.vect[order(exo.vect$Time),]
		
	}
	
	
}
}


event.list.dat<-split(event.list,event.list$Date)


#### create new intercept statlist ####

##"a three dimensional array, event number by event type by statistic, or a list thereof, containing the sufficient statistics for the model"

#start with the intercept sufficient stats

#for it to be compatible with the stat functions, I might have to go ahead and re-create it every time in the usual way and then just look at the latest stat layer. One excpetion is that it needs to be one event layer longer so that it can be used to predict the next future event


if(day==1){
	sequences<-step
}

if(day==2){
sequences<-c(nrow(evlist[[1]]),step)	
}

#create list of dummy intercept names

idno<-length(event.types[grep("cop.start",event.types)])

dummy.ints.attack<-paste(1:14,".attackFE", sep="")
dummy.ints.receive<-paste(1:13, ".receiveFE", sep="")

IDintsCH<-paste(1:14,"initCH",sep=".")
IDdis<-paste(1:14,"disengage",sep=".")

int.names<-c("Global.attackFE", "X.receiveFE", "X.UreceiveFE", "U.attackFE", "U.receiveFE", "U.UattackFE", dummy.ints.attack, dummy.ints.receive, "cop.startFE","cop.endFE", "initFO.FE","ID.initFO.FE", "U.initFO.FE","initSF.FE","X.initSF.FE", "initCH.FE",  "U.initCH.FE", IDintsCH, "disengage.FE", "U.disengage.FE", IDdis, "depart.FE", "X.depart.FE")

int.list<-list()


for(i in c(1:day)){

day.array<-array(0,dim=c(as.numeric(sequences[i]), length(event.types),length(int.names)))

#the third dimension is the stat slice.
dimnames(day.array)<-list(c(1:sequences[i]),as.character(event.types),as.character(int.names))

#add global attack effect, loops for "U" intercepts for attacks and receives, then one loop for typical intercepts

day.array[,grep("attack", event.types),1]<-1


for(h in 1:14){
day.array[,paste(h,".attack.",30, sep=""),2]<-1
}


for(h in 15:29){
day.array[,paste(h,".attack.",30, sep=""),3]<-1
}


for(h in 1:14){
	for(j in 15:29){	
day.array[,paste(j,".attack.", h, sep=""),4]<-1
}
}


for(h in 1:14){
	for(j in 15:29){
day.array[,paste(h,".attack.",j, sep=""),5]<-1
}
}

for(h in 15:29){
	for(j in 15:29){	
if(h!=j){
day.array[,paste(j,".attack.", h, sep=""),6]<-1
}
}
}

preloop<-6

#loop for dummy intercepts for attacks, loop for dummy intercepts for receives, add in cop.start and cop.end FEs, then one loop for typical intercepts

for(k in (preloop+1):(length(dummy.ints.attack)+preloop)){
	
	for(h in 1:14){
if(h!=(k-preloop)){
day.array[,paste((k-preloop),".attack.", h, sep=""),k]<-1
}
}
}

for(k in ((length(dummy.ints.attack)+preloop)+1):((length(dummy.ints.attack)+preloop)+length(dummy.ints.receive))){
	for(h in 1:14){
if(h!=(k-(length(dummy.ints.attack)+preloop))){
day.array[,paste(h,".attack.",(k-(length(dummy.ints.attack)+preloop)), sep=""),k]<-1
}
}
}

#cop.startFE
for(h in 1:idno){
day.array[,paste(h,".cop.start", sep=""),((length(dummy.ints.attack)+preloop)+length(dummy.ints.receive)+1)]<-1
}


#cop.endFE
for(h in 1:idno){
day.array[,paste(h,".cop.end", sep=""),((length(dummy.ints.attack)+preloop)+length(dummy.ints.receive)+2)]<-1
}

#initFO.FE
for(h in 1:idno){
day.array[,paste(h,".initFO", sep=""),((length(dummy.ints.attack)+preloop)+length(dummy.ints.receive)+3)]<-1
}

#ID.initFO.FE
for(h in 1:14){
day.array[,paste(h,".initFO", sep=""),((length(dummy.ints.attack)+preloop)+length(dummy.ints.receive)+4)]<-1
}

#U.initFO.FE
for(h in 15:29){
day.array[,paste(h,".initFO", sep=""),((length(dummy.ints.attack)+preloop)+length(dummy.ints.receive)+5)]<-1
}

#initSF.FE
for(h in 1:idno){
day.array[,paste(h,".initSF", sep=""),((length(dummy.ints.attack)+preloop)+length(dummy.ints.receive)+6)]<-1
}

#X.initSF.FE
day.array[,paste(30,".initSF", sep=""),((length(dummy.ints.attack)+preloop)+length(dummy.ints.receive)+7)]<-1


#initCH.FE
for(h in 1:idno){
day.array[,paste(h,".initCH", sep=""),((length(dummy.ints.attack)+preloop)+length(dummy.ints.receive)+8)]<-1
}


#U.initCH.FE
for(h in 15:29){
day.array[,paste(h,".initCH", sep=""),((length(dummy.ints.attack)+preloop)+length(dummy.ints.receive)+9)]<-1
}


#individual initCH effects for ID male
for(k in (((length(dummy.ints.attack)+preloop)+length(dummy.ints.receive)+10):((length(dummy.ints.attack)+preloop)+length(dummy.ints.receive)+10+(length(IDintsCH)-1)))){
	
	day.array[,int.names[k],k]<-1
	
}

preloop2<-((length(dummy.ints.attack)+preloop)+length(dummy.ints.receive)+10+(length(IDintsCH)-1))


#disengage.FE
for(h in 1:idno){
day.array[,paste(h,".disengage", sep=""),preloop2+1]<-1
}

#U.disengage.FE
for(h in 15:29){
day.array[,paste(h,".disengage", sep=""),preloop2+2]<-1
}

#individual disengage effect for ID males
for(k in (preloop2+3):(preloop2+3+length(IDdis)-1)){
	
	day.array[,int.names[k],k]<-1
	
}


preloop3<-(preloop2+3+length(IDdis)-1)

#depart.FE
for(h in 1:idno){
day.array[,paste(h,".depart", sep=""),preloop3+1]<-1
}

#X.depart.FE
day.array[,paste(30,".depart", sep=""),preloop3+2]<-1


#weird extra formatting to make it match the package example, seems like it becomes relevant *punintended!* if I want to fit day-specific effects

global<-list()

global[[1]]<-day.array

int.list[[i]]<-global

names(int.list[[i]])<-"global"

}


#the support matrix and associated objects need to be built every timestep. So actually it will be a vector for each timestep telling me whats possible.  

#### Create Support Matrix ####

#"should be a list withone element per history, each of which should be an event order by event type logical matrix. the ijth cell of this matrix should be TRUE if an event type of j was a possible next event given the preceding code i-1 events, and FALSE otherwise"

#this is the tool that lets me see the list of all event that are possible at the next timestep for caluclating the conditional hazard
#looks like I can keep this code the way it is

#need a way to ask, "what is each male doing right now" for each event in the sequence. That dictates the possibilities. 

#create a bookkeeping "census" matrix to keep track
{
census.list<-list()

for(j in 1:day){

census<-matrix("nothing", nrow=1,ncol=(idno), dimnames=list(1,c(1:(idno))))


##code that translates each new current event into what the males involved are doing at that time. The support matrix will be based on whatever the males were doing in the PREVIOUS timestep. This census matrix itself could be interesting to visualize. Each row of the census says what each male is doing going into the NEXT event.
if(step>1|(step==1 && day==2 && j==1)){

#census list gets built differently for the observed data, which is always day 1 here
if(j==1){
	
  for(i in 1:length(evlist[[j]][,1])){
          
          #can ask if the second thing is something, and then move on to the first and third elements.
          
          current<-census[nrow(census),]
          
          #parse the event type. loop through test.seq rows
          evname<-as.character(event.list.dat[[j]][i,1])
          evsplit<-strsplit(evname, split=".", fixed=T)
          
          #if it is an end obs, arrival, departure or clock event, it shouldn't change anything the males are doing in terms of interactions. Departures aren't exogenous, but they shouldn't change anything since they can only happen once a male is already doing nothing. 
          if(evsplit[[1]][2]!="obs" & evsplit[[1]][2]!="depart" & evsplit[[1]][2]!="arrive" & evsplit[[1]][2]!="time" & evsplit[[1]][1]!="fem"){
            #if the event was an attack, indicate the males involved and which was the receiver since this dictates who makes the initition decision
            if(evsplit[[1]][2]=="attack"){
              current[evsplit[[1]][1]]<-evname
              current[evsplit[[1]][3]]<-paste(evsplit[[1]][3], ".", "received", ".", evsplit[[1]][1], sep="")
            }
            #if it's an FO or SF, just indicate that that pair of males are involved in a fight now
            if(evsplit[[1]][2]=="initFO"||evsplit[[1]][2]=="initSF"){
              #need to refer back to the previous row of "census" to get the id of the other male
              
              current[evsplit[[1]][1]]<-paste(evsplit[[1]][1], ".", "fight", ".",strsplit(census[nrow(census),evsplit[[1]][1]], split=".",fixed=T)[[1]][3], sep="")
              current[[strsplit(census[nrow(census),evsplit[[1]][1]], split=".",fixed=T)[[1]][3]]]<-paste(strsplit(census[nrow(census),evsplit[[1]][1]], split=".",fixed=T)[[1]][3], ".", "fight", ".",evsplit[[1]][1], sep="")
              
            }
            
            #if it's a chase, indicate that the pair of males are involved in a fight, but there is an additional constraint that the individual DOING THE CHASING will be the disengager by definition
            if(evsplit[[1]][2]=="initCH"){
              
              current[evsplit[[1]][1]]<-paste(evsplit[[1]][1], ".", "fight", ".",strsplit(census[nrow(census),evsplit[[1]][1]], split=".",fixed=T)[[1]][3], ".chasedby",sep="")
              current[[strsplit(census[nrow(census),evsplit[[1]][1]], split=".",fixed=T)[[1]][3]]]<-paste(strsplit(census[nrow(census),evsplit[[1]][1]], split=".",fixed=T)[[1]][3], ".", "fight", ".",evsplit[[1]][1], ".chasing",sep="")
              
            }
            
            #for disengaging, end the fight for the partner male as well
            if(evsplit[[1]][2]=="disengage"){
              current[evsplit[[1]][1]]<-"nothing"
              current[evsplit[[1]][3]]<-"nothing"	
              
            }
            
            
            #for copulations, end cops are only possible after start cops, but otherwise anything else is possible. EXCEPT the copulating male can't initiate an attack. If the next event involving that male ISN'T an end cop, it was interrupted
            
            if(evsplit[[1]][2]=="cop" && evsplit[[1]][3]=="start"){
              current[evsplit[[1]][1]]<-"copulating"
            }
            
            if(evsplit[[1]][2]=="cop" && evsplit[[1]][3]=="end"){
              current[evsplit[[1]][1]]<-"nothing"
            }
            
            #if an X male is still involved in a fight at all, make sure he is still tagged as fighting (since sometimes more than one is interacting at once, but they can't depart until none are fighting)
            if((TRUE %in% grepl("fight.30",current) | TRUE %in% grepl("30.fight",current)) && !(grepl("receive", current[idno]) | grepl("attack", current[idno]))) {
              
              current[idno]<-"fight"
              
            }
            
          }
          
          census<-rbind(census,current)
          
          
        }	
	
} else {
	
for(i in 1:length(evlist[[j]][,1])){

#can ask if the second thing is something, and then move on to the first and third elements.

current<-census[nrow(census),]

#parse the event type. loop through test.seq rows
evname<-as.character(event.list.dat[[j]][i,1])
evsplit<-strsplit(evname, split=".", fixed=T)

#if it is an end obs, arrival, departure or clock event, it shouldn't change anything the males are doing in terms of interactions. Departures aren't exogenous, but they shouldn't change anything since they can only happen once a male is already doing nothing. 
if(evsplit[[1]][2]!="obs" & evsplit[[1]][2]!="depart" & evsplit[[1]][2]!="arrive" & evsplit[[1]][2]!="time" & evsplit[[1]][1]!="fem"){
#if the event was an attack, indicate the males involved and which was the receiver since this dictates who makes the initition decision
	if(evsplit[[1]][2]=="attack"){
		current[evsplit[[1]][1]]<-evname
		current[evsplit[[1]][3]]<-paste(evsplit[[1]][3], ".", "received", ".", evsplit[[1]][1], sep="")
			}
	#if it's an FO or SF, just indicate that that pair of males are involved in a fight now
		if(evsplit[[1]][2]=="initFO"||evsplit[[1]][2]=="initSF"){
			#need to refer back to the previous row of "census" to get the id of the other male
			
			current[evsplit[[1]][1]]<-paste(evsplit[[1]][1], ".", "fight", ".",strsplit(census[nrow(census),evsplit[[1]][1]], split=".",fixed=T)[[1]][3], sep="")
			current[[strsplit(census[nrow(census),evsplit[[1]][1]], split=".",fixed=T)[[1]][3]]]<-paste(strsplit(census[nrow(census),evsplit[[1]][1]], split=".",fixed=T)[[1]][3], ".", "fight", ".",evsplit[[1]][1], sep="")
			
			}
			
	#if it's a chase, indicate that the pair of males are involved in a fight, but there is an additional constraint that the individual DOING THE CHASING will be the disengager by definition
	if(evsplit[[1]][2]=="initCH"){
			
		current[evsplit[[1]][1]]<-paste(evsplit[[1]][1], ".", "fight", ".",strsplit(census[nrow(census),evsplit[[1]][1]], split=".",fixed=T)[[1]][3], ".chasedby",sep="")
			current[[strsplit(census[nrow(census),evsplit[[1]][1]], split=".",fixed=T)[[1]][3]]]<-paste(strsplit(census[nrow(census),evsplit[[1]][1]], split=".",fixed=T)[[1]][3], ".", "fight", ".",evsplit[[1]][1], ".chasing",sep="")

	}

#for disengaging, need to search through the previous rows to find the most recent "fight" and identify the partner and end it
	if(evsplit[[1]][2]=="disengage"){
	current[evsplit[[1]][1]]<-"nothing"
	current[strsplit(census[max(grep("fight", census[,evsplit[[1]][1]])),evsplit[[1]][1]], split=".", fixed=T)[[1]][3]]<-"nothing"	
	
}
 

#for copulations, end cops are only possible after start cops, but otherwise anything else is possible. EXCEPT the copulating male can't initiate an attack. If the next event involving that male ISN'T an end cop, it was interrupted

	if(evsplit[[1]][2]=="cop" && evsplit[[1]][3]=="start"){
		current[evsplit[[1]][1]]<-"copulating"
	}

	if(evsplit[[1]][2]=="cop" && evsplit[[1]][3]=="end"){
		current[evsplit[[1]][1]]<-"nothing"
	}

# #if an X male is still involved in a fight at all, make sure he is still tagged as fighting (since sometimes more than one is interacting at once, but they can't depart until none are fighting)
	# if((TRUE %in% grepl("fight.30",current) | TRUE %in% grepl("30.fight",current)) && !(grepl("receive", current[idno]) | grepl("attack", current[idno]))) {
		
		# current[idno]<-"fight"

# }

}

census<-rbind(census,current)


}

}


#don't want the first dummy row
census.list[[j]]<-census[-1,]


#if it's the second step, then the census list for that day still has only one row, so it needs to be formatted properly.

if((step==2 && day==1)|(step==2 && j==2)){
	
	census.list[[j]]<-t(as.matrix(census.list[[j]]))
}


} 

#for these cases that step=1 on day one or day two
if((step==1 && day==1 && j==1)|(step==1 && day==2 && j==2)){
	
	
	census.list[[j]]<-census
	#census.list[[j]]<-t(as.matrix(census.list[[j]]))
	
	}

}
}


###Make the win/loss census array


    #convert to win.loss census- again need two diferent functions for the different datasets b/c of constraints on X males 
    
    create.win.loss.census.act<-function(x){
      
      #get disengagement events for focal male from evlist
      win.loss.census<-x
      
      #loop thru days and males
      
      for(i in 1:length(win.loss.census)){
        
        for(l in 1:idno){
          
          
          dis.times<-which(startsWith(as.character(event.list.dat[[i]][,1]), paste(l,".disengage", sep="")))
          
          
          if(length(dis.times)>0){
            
            evsplit<-strsplit(as.character(event.list.dat[[i]][dis.times,1]),".",fixed=T)
            
            #so I have a list of row numbers for disengagements and a list of interaction partners. This will cover wins for normal fights and losses for chases for the focal male. If I loop through all the males, all the wins and losses should get filled in.
            
            #NOW DISENGAGES ALWAYS MEAN WINS, so I will still have chase information, but I don't need to do anything special with it here.
            
            #need to make sure I am really precise since male 30 can be involved in multiple interactions, but they're usually all chases.
            
            
            for(j in 1:length(dis.times)){
              
              
              if(grepl("chasing",win.loss.census[[i]][(dis.times[j]-1),l]) ==FALSE && grepl("chasedby",win.loss.census[[i]][(dis.times[j]-1),as.numeric(evsplit[[j]][3])]) ==FALSE){
                #apply the win in the focal male's column
                win.loss.census[[i]][dis.times[j],l]<-paste(l, "win",evsplit[[j]][3], sep=".")
                #apply the loss for the same interaction in the partner male's column
                win.loss.census[[i]][dis.times[j],as.numeric(evsplit[[j]][3])]<-paste(evsplit[[j]][3], "lose", l, sep=".")
              } else{
                #apply the won chase in the focal male's column
                win.loss.census[[i]][dis.times[j],l]<-paste(l, "win",evsplit[[j]][3],"chase", sep=".")
                #apply the loss for the same interaction in the partner male's column.
                win.loss.census[[i]][dis.times[j],as.numeric(evsplit[[j]][3])]<-paste(evsplit[[j]][3], "lose", l, "chase",sep=".")
              }
              
            }
            
          }
          
        }
        
      }
      
      return(win.loss.census)
      
    }


#i've adjusted the function to mark when a win or loss is the result of a chase or not
create.win.loss.census.sim<-function(x){

#get disengagement events for focal male from evlist
win.loss.census<-x

#loop thru days and males

for(i in 1:length(win.loss.census)){

for(l in 1:idno){

dis.times<-which(startsWith(as.character(event.list.dat[[(i+1)]][,1]), paste(l,".disengage", sep="")))

if(length(dis.times)>0){

#see who they were fighting with
fights<-win.loss.census[[i]][dis.times-1,l]

#so I have a list of row numbers for disengagements and a list of interaction partners. This will cover wins for normal fights and losses for chases for the focal male. If I loop through all the males, all the wins and losses should get filled in.

#NOW DISENGAGES ALWAYS MEAN WINS, so I will still have chase information, but I don't need to do anything special with it here.

for(j in 1:length(dis.times)){
	
	if(grepl("chasing",fights[j]) ==FALSE){
	#apply the win in the focal male's column
	win.loss.census[[i]][dis.times[j],l]<-paste(l, "win",strsplit(fights[j],".", fixed=T)[[1]][3], sep=".")
	#apply the loss for the same interaction in the partner male's column
	win.loss.census[[i]][dis.times[j],as.numeric(strsplit(fights[j],".", fixed=T)[[1]][3])]<-paste(strsplit(fights[j],".", fixed=T)[[1]][3], "lose", l, sep=".")
	} else{
		#apply the won chase in the focal male's column
	win.loss.census[[i]][dis.times[j],l]<-paste(l, "win",strsplit(fights[j],".", fixed=T)[[1]][3],"chase", sep=".")
		#apply the loss for the same interaction in the partner male's column.
	win.loss.census[[i]][dis.times[j],as.numeric(strsplit(fights[j],".", fixed=T)[[1]][3])]<-paste(strsplit(fights[j],".", fixed=T)[[1]][3], "lose", l, "chase",sep=".")
		}
		
}

}

}

}

return(win.loss.census)
	
}


#put it together

win.loss.census.act<-create.win.loss.census.act(census.list[1])

win.loss.census.sim<-create.win.loss.census.sim(census.list[c(2:length(census.list))])
    
    
win.loss.census<-append(win.loss.census.act,win.loss.census.sim)


## Constraints to incorporate in the support matrix ##

#only males that are actually present can be involved in interactions on a given day and at a given time

#If "nothing" is happening, all "attacks", all start cops and all departs (given the male is actually present) are possible. Males cannot disengage or initiate fights without an attack happening first. End cop cannot happen without a previous start cop. end.obs is possible. (end.obs is not possible if any males are still fighting or copulating, so end.obs is ONLY possible when "nothing" is going on across the board)

#if the previous event was an "attack", the current event MUST be an initiation by the receiver, since we are assuming the attack is essentually instantaneous. 

#once initiatiated, subsequent events can only be a disengage from either male involved or any other event NOT invoving the interaction partners. Males actively fighting connot become engaged in a new fight until they disengage. 

#if the initiated fight was a chase only the male being chased can disengage, by definition

#Once a copulation starts, the only thing that male can do next is either end the copulation OR receive an attack. He cannot initiate anything. 

#copulations cannot occur when there are no females present

#a male can ONLY "depart" if "nothing" is happening

#Exceptions to these rules are for "X" males- there may be more than one unidentifiable male on the lek at a time. Right now we are dealing with this by having "X" males just be a mass actor category. 1) These males CAN do an "attack" before disengaging from an interaction. They can also be attacked. However, normal restrictions apply to the partner male. So if an X male is being chased, "disengage" becomes possible, but any "attack" (or cop.start, or depart) are still also possible 


### Build the actual support matrix ###


#for each event, there is a list of event types, with 1's for the types that are possible and 0's for types that are not. 

#str(evlist)

#code block to produce the support matrix


#make a reference event list with data (most importantly female presence)

evlist.dat<-event.list.dat

supplist<-list()

#for(j in 1:length(evlist)){

supplist<-matrix(1, nrow=1, ncol=length(event.types))
colnames(supplist)<-event.types

#turn off all interactions involving males not present on that day
#make a list of all the event types that include the males that were entirely absent and remove them for all rows

supplist[,unique(unlist(lapply(grep(0,male.pres[day,2:(idno+1)]), function(x) c(event.types[startsWith(event.types, paste(x, ".", sep=""))], event.types[endsWith(event.types, paste(".", x, sep=""))]))))]<-0


#for the first event of the day:

if(step==1){
  #first row can be any "attack" or, technically, any "depart"
  #since now fem.enter is an actual event, cop.start is never possible as the very first thing
supplist[1,][-c(grep("attack", event.types), grep("depart", event.types))]<-0

#the exception is the events involving males who will be there at some point but haven't arrived yet

arrsub<-subset(pres.exact.dat[[day]],Time==0)
arrtab<-table(arrsub$Bird_ID,arrsub$Type)

arrpat<-apply(arrtab,1,function(x) sum(x))

#if the total number of previous arrives and departs is even, that male is currently absent
current.row.abs<-names(arrpat[(arrpat %% 2)==0])


if(length(current.row.abs)>0){
supplist[1,][unique(c(sapply(current.row.abs, function(x) c(event.types[startsWith(event.types, paste(x, ".", sep=""))], event.types[endsWith(event.types, paste(".", x, sep=""))]))))]<-0

}

#If females have not arrived yet, then cop.starts are also not possible
if(Females==1){
	
	copstartall<-event.types[grep("cop.start",event.types)]
	supplist[1,][copstartall]<-0
	
}

 
}

#for all subsequent events:

if(step>1){ 

for(i in step){

#before anything else turn off events involving males that have either not yet arrived or have lef the lek early and are therefore no longer present that day or are otherwise not currently present


arrsub<-subset(pres.exact.dat[[day]],Time<=event.list.dat[[day]][(i-1),2])
arrtab<-table(arrsub$Bird_ID,arrsub$Type)

arrpat<-apply(arrtab,1,function(x) sum(x))

#if the total number of previous arrives and departs is even, that male is currently absent
current.row.abs<-names(arrpat[(arrpat %% 2)==0])


if(length(current.row.abs)>0){
supplist[1,][unique(c(sapply(current.row.abs, function(x) c(event.types[startsWith(event.types, paste(x, ".", sep=""))], event.types[endsWith(event.types, paste(".", x, sep=""))]))))]<-0

} 


#determine if the previous event was an attack, since this dictates exactly what the next event can be: receiver initiates a type of fight
if((TRUE %in% grepl("received", census.list[[day]][i-1,]))==TRUE){

supplist[1,][-grep(TRUE, startsWith(event.types, paste(grep("received", census.list[[day]][i-1,]), ".", "init", sep="")))]<-0}


#(else) if the previous event wasn't an attack, the next event CAN'T be an initiation and the possibilities depend on what everyone is currently engaged in


if((TRUE %in% grepl("received", census.list[[day]][i-1,]))==FALSE){

supplist[1,][grep("init", event.types)]<-0

#if males are actively fighting, they can't attack or be attacked, or actually do anything else before disengaging

if((TRUE %in% grepl("fight", census.list[[day]][i-1,]))==TRUE){
		
		for(k in grep("fight", census.list[[day]][i-1,])){
	
	
	
	supplist[1,][c(event.types[startsWith(event.types, paste(k, ".attack.", sep=""))], event.types[endsWith(event.types, paste("attack.", k, sep=""))], event.types[startsWith(event.types, paste(k, ".cop", sep=""))])]<-0
	
}
	
	#ONLY those fighting can disengage. retain males that are fighting and remove the rest
	dis<-event.types[grep("disengage",event.types)]
	supplist[1,][dis[-grep("fight", census.list[[day]][i-1,])]]<-0
	
	#"chasing" and "chasedby" limit the possibilities further. Males getting chased DO NOT disengage by definition, again, with the exception of ?? males. This is taken care of by having ?? males just be "fight" -ing without any extra tags
	if(TRUE %in% grepl("chasedby", census.list[[day]][i-1,])){
		supplist[1,][dis[grep("chasedby", census.list[[day]][i-1,])]]<-0
	}
	
}

#if no one is fighting, no one can disengage, but all attack combinations are possible
if((TRUE %in% grepl("fight", census.list[[day]][i-1,]))==FALSE){
	
		supplist[1,][grep("disengage",event.types)]<-0
	
}


#actively copulating males: They cannot cop.start. ONLY they can cop.end. they CAN be attacked but they CANNOT attack anyone
if((TRUE %in% grepl("copulating", census.list[[day]][i-1,]))==TRUE){
	
	cop<-grep("copulating", census.list[[day]][i-1,]) 
	
	#stuff that needs to be turned off for only copulating males
	for(h in cop){

		supplist[1,][c(event.types[startsWith(event.types ,paste(h, ".attack.", sep=""))], event.types[startsWith(event.types, paste(h, ".cop.start", sep=""))])]<-0
				
	}
	
	#stuff that needs to be on for ONLY copulating males
	copendall<-event.types[grep("cop.end",event.types)]
	supplist[1,][copendall[-cop]]<-0
}

#if no one is copulating, all cop.starts are possible, but cop.ends are not

if((TRUE %in% grepl("copulating", census.list[[day]][i-1,]))==FALSE){
	
	copendall<-event.types[grep("cop.end",event.types)]
	supplist[1,][copendall]<-0
	
}


}

#finally, if there are no females present, the next event could not have possibly been a cop.start

if(Females==1){
	
	copstartall<-event.types[grep("cop.start",event.types)]
	supplist[1,][copstartall]<-0
	
}

#Finally, A male can only "depart" if he was already doing "nothing," so depart must be turned off for everyone doing not-nothing. (also he has to be present, but I've already taken care of that above)

noth<-as.character(which(census.list[[day]][i-1,]!="nothing"))
	
	if(length(noth)>0){
		
	supplist[1,][paste(noth, ".depart", sep="")]<-0

}


}
#end events loop here


}


########

#add the new stats to the base intercept list

test.stat<-int.list

#reset arg.use each loop, or I'll end up not using any of my functions!
arg.use<-c(1:length(args.all))

#rememebr that for the very first event on day 2, I have to leave out the non-yest stats

#if it's the first event on day 1, only centroid distance and females apply
if(step==1 & day==1){
	
	arg.use<-grep("fem|centroid",fun.list[args.all])

}

#if it's the first event of day 2, females, centroid , cumul.sol and yest (except .yest.cop and .yest.dis) apply, but none others
if(step==1 & day==2){
	
	arg.use<-grep("fem|centroid|cumul.sol|yest.*(?<!.cop|.dis)$",fun.list[args.all], perl=T)
}


#this makes sure there are placeholders for functions that don't apply in the step=1 cases

for(i in 1:length(args.all)){


if(i %in% arg.use){
	
	test.stat<-get(fun.list[args.all[i]])(test.stat)	
	
} else {

for(k in 1:day){

if(day>1 && k<2){
#empty matrix of the right dimensions
stat<-matrix(0,nrow=nrow(test.stat[[k]][[1]][,,1]), ncol=length(event.types))
colnames(stat)<-event.types
} 
else {
stat<-matrix(0,nrow=step, ncol=length(event.types))
colnames(stat)<-event.types
}

test.stat[[k]][[1]]<-abind(test.stat[[k]][[1]], stat)
dimnames(test.stat[[k]][[1]])[[3]][ncol(test.stat[[k]][[1]][1,,])]<-rownames(mod_coef)[length(mod_coef[,1])-length(args.all)+i]

}

}
}

#for checking:
#test.stat[[2]][[1]][step-1,"4.cop.start",arg.use+74]


#next step is to calculate the hazard for the next timestep, choose the delta t and the next event, then make the loop

#stat matrix predicting the next event!
stats<-test.stat[[day]][[1]][step,,]

#the coefficient vector gets multiplied by each row and the supplist vector gets multiplied by each column
statsup<-apply(stats,2,function(x) x*supplist)

#for some reason the apply function across the row margin returns a transformed matrix, so I transform it back for sanity. Use the subset of coefficents for the predictors that are acutally included for this timestep
statcoef<-t(apply(statsup,1,function(x) x*as.numeric(mod_coef[,1])[rownames(mod_coef) == dimnames(stats)[[2]]]))

dimnames(statcoef)[[1]]<-event.types

#get the parameter-weighted sums of stats for each event type
log.hazards<-apply(statcoef,1,function(x) sum(x))

#restrict to only the events that are possible according to the supplist
possible.haz<-log.hazards[log.hazards!=0]

#get the total hazard for any event to happen next
tot.haz<-sum(exp(possible.haz))

if(tot.haz>0){

delta.t<-rexp(1,tot.haz)

##next is to choose the next event as a categorical random variable

#proportions of the total hazards
prop.haz<-exp(possible.haz)/sum(exp(possible.haz))

#sample the next event as a categorical random variable
nextev<-sample(names(prop.haz),1,prob=prop.haz)

#find the event number in the event list
nextev.num<-grep(paste0("^",nextev,"$"),event.types)

#set a provisional new t, barring any exogenous events

tcurr<-t

#provisionally set a new t
t<-t+delta.t


## Is there an exogenous event scheduled to happen within the interval tcurr - t? Special case for t=0
if(step==1 && tcurr==0){

sched<-exo.vect[(tcurr<=exo.vect$Time & exo.vect$Time<t),]	
	
} else {
	
sched<-exo.vect[(tcurr<exo.vect$Time & exo.vect$Time<t),]

}


if(length(sched$Time)>0){
	
	#even if there is more than one, just take the first one (the one happening soonest)
	sched.ev<-sched[1,]
	
	if(sched.ev$Events=="fem.enter"){
		
	nextev<-"fem.enter"
	nextev.num<-0
	
	#update females for eventlist
	event.list[nrow(event.list),3]<-2
	Females<-2
	t<-sched.ev$Time

		
	}
	
	if(sched.ev$Events=="fem.exit"){
		
	nextev<-"fem.exit"
	nextev.num<-0
	
	#update females for eventlist
	event.list[nrow(event.list),3]<-1
	Females<-1
	t<-sched.ev$Time
		
	}
	
	if(sched.ev$Events=="clock.time"){
		
	nextev<-"clock.time"
	nextev.num<-0
	
	#just update time
	t<-sched.ev$Time
		
	}
	
	if(sched.ev$Events=="disturb.ance"){
		
	nextev<-"disturb.ance"
	nextev.num<-0
	
	#just update time
	t<-sched.ev$Time
		
	}
	
	if(endsWith(sched.ev$Events,".arrive")){
		
	nextev<-sched.ev$Events
	nextev.num<-0
	
	#just update time
	t<-sched.ev$Time
		
	}
	
}


#end the sim if we run out of time
if(t>tmax){
	
	nextev<-"end.obs"
	nextev.num<-0
	
	#update time
	t<-tmax	
	
}

#if none if these above are true, it wont do anything and it will retain the natural event assignment

} 

if(tot.haz==0){
	
	#if tot.haz==0, then there are no males on the lek and we end the sim
	
	nextev<-"end.obs"
	nextev.num<-0
	
	#update time
	t<-tmax		
	
	#signal that this one ended with all males departing
	print("no males on lek")
	
}


if(step==1){evlist[[day]][1,]<-c(nextev.num,t)} else {evlist[[day]]<-rbind(evlist[[day]], c(nextev.num,t))}

#remember to also make event.list object to hold females present info in a format that the functions can use
event.list[nrow(event.list),c(1,2)]<-c(nextev,t)

#save the proportional hazards for plotting
overall.haz<-rep(0,length(event.types))
names(overall.haz)<-event.types

overall.haz[names(possible.haz)]<-exp(possible.haz)

overall.haz.dat<-rbind(overall.haz.dat,overall.haz)

step<-step+1

##back to the beginning of the loop here
}
#toc()
###################################
#1657.887 sec elapsed

event.list.split<-split(event.list,event.list$Date)

#event.list.split[[2]]

#this curly is for the cop while loop that I will comment out
#}

#write code to export data

#for parallelization, have it use a command line argument to number the run, and use that to label the output folder
if(!dir.exists("sim_results_actualday")) dir.create("sim_results_actualday")

#folder for the simulated day
datfile<-paste0("./sim_results_actualday/sim_results_",days[2])
if(!dir.exists(datfile)) dir.create(datfile)

#folder for the specific sim run for that day
outfile<-paste0(datfile,"/","sim_results_",days[2],"_",args[2])
dir.create(outfile)

write.table(evlist[[1]],file=paste0(outfile,"/","evlist_day1"),sep=",",row.names=F)

write.table(evlist[[2]],file=paste0(outfile,"/","evlist_day2"),sep=",",row.names=F)

write.table(event.list,file=paste0(outfile,"/","full_event_list"),sep=",",row.names=F)

write.table(event.list.split[[1]],file=paste0(outfile,"/","full_event_list_day1"),sep=",",row.names=F)

write.table(event.list.split[[2]],file=paste0(outfile,"/","full_event_list_day2"),sep=",",row.names=F)

write.table(overall.haz.dat,file=paste0(outfile,"/","hazards"),sep=",",row.names=F)


write.table(pres.exact.dat[[1]],file=paste0(outfile,"/","pres_exact_day_1"),sep=",",row.names=F)
  
write.table(pres.exact.dat[[2]],file=paste0(outfile,"/","pres_exact_day_2"),sep=",",row.names=F)
```
