## Supplemental_reproducible_code for "Fighting isn’t sexy in lekking Greater Sage-grouse (*Centrocercus urophasianus*)": run_pipeline_sim_act.sh

```
#!/bin/bash
# -t 5:00

# NOTES
# consider using sprintf("%03d", number) in R for formatting output. Will sort files numerically.

# \xC2\xA0 found in some R scripts. Is called non-breaking space, is wide UTF8 character which irritates R on cluster
for rfile in *.R;
 do
     sed -i 's/\xC2\xA0/ /g' $rfile
 done

module load R

if not [ -f joblist_1.txt ]
then
    Rscript genremlist_sim.R
    jobfile="ls joblist*";
else
    echo "simulations finished"
    exit 0
fi

module purge
module load dSQ/0.96
mkdir -p job_logfiles job_statusfiles

for jobby in $jobfile
 do
 	jid=$(dsq --jobfile $jobby --mem=10G -o job_logfiles/dsq-jobfile-%A_%a-%N.out --status-dir job_statusfiles --submit | awk '{print $NF}')
 	echo "submitted array $jid"
 done
 
 
sbatch --dependency afterany:$jid --kill-on-invalid-dep=yes run_pipeline_sim_act.sh
```
