## Supplemental_reproducible_code for "Fighting isn’t sexy in lekking Greater Sage-grouse (*Centrocercus urophasianus*)": sim_output_analysis_cluster_new_3.R

```
### Code for Visualization of Simulation Results

rm(list=ls())

###make sure necessary packages are there and loaded

packages<-c("here","abind","ggplot2","cowplot","gridExtra","tidyverse","forcats","igraph","sna")

package.check<-lapply(packages, FUN = function(x) {
    if (!require(x, character.only = TRUE, quietly=TRUE)) {
        install.packages(x, dependencies = TRUE)
        library(x, character.only = TRUE, quietly=TRUE)
    } 

})

#for project path management
#define the project root directory relative to the current script. This sets the working directory that gets called by the here() function. It is a relative path and will be robust to use on anyone's computer. In this case, we want the root directory to always be the enclosing folder for the whole sage grouse project

i_am("event_history_simulation/Aggression_model/sim_output_analysis_cluster_new_2.R")

setwd(here("event_history_simulation","Aggression_model"))


#Bring in the empirical data for comparison

event.list<-read.table(file="relevent_format_dataset_udata_wincol_disturbance.tsv", header=T,  sep="\t")

event.types<-read.table(file="event_types_list_udata.csv", header=F,  sep=",")
event.types<-as.character(event.types$V1)


#NOTE- the event.types vector now contains ENDOGENOUS event types as well as EXOGENOUS types: arrivals and clock events, and end.obs. Need to make sure everything is robust to this. 
### create eventlist for rem ###


event.list.dat<-split(event.list, event.list$Date)

dates<-as.character(unique(event.list$Date))


####### Per-Day Visualizations ######### 


####
#Make a dataset with all the sims

sim.all.dat<-list()

for(j in 1:length(event.list.dat)){

mod.day<-j

daydata<-list()

daydata[[1]]<-event.list.dat[[mod.day]]

dayfolders<-list.files(paste0("./sim_results_actualday/sim_results_",dates[[mod.day]]), include.dirs=T,full.names=T)

simlist<-list()

if(length(dayfolders)>0){

for(i in 1:length(dayfolders)){

simdata<-list.files(dayfolders[i],full.names=T)

simevents<-read.table(simdata[endsWith(simdata,"full_event_list")], sep=",",header=T, stringsAsFactors=F)
#str(simevents)

simevents<-split(simevents,simevents$Date)


#get event list for the simulated day based on an actual yesterday

simday<-simevents[[2]]


simlist[[i]]<-simday

}
}

daydata[[2]]<-simlist

sim.all.dat[[j]]<-daydata

}


#### Make summary plots for each day 


if(!dir.exists("./sim_actualday_plots")) dir.create("./sim_actualday_plots")


#make a list to hold overall data

overallcop<-list()

overallfight<-list()

overallnet<-list()

overallcounts<-list()


for(b in 1:length(sim.all.dat)){
	#make sure I acutally have sim data for the day
	if(length(sim.all.dat[[b]][[2]])>0){

actday<-sim.all.dat[[b]][[1]]

simlist<-sim.all.dat[[b]][[2]]


#### Summary stats ####

#Start with counts of stuff

counts.df<-data.frame()


##event number

evs<-matrix(data=c("events","act",length(actday[,1])),nrow=1,ncol=3)


evsim<-lapply(simlist,function(x) {
	
	c("events","sim",length(x[,1]))
	
})

library(abind)

evsim<-t(abind(evsim,along=2))

evdat<-data.frame(rbind(evs,evsim))

colnames(evdat)<-c("Category","Data","Count")

counts.df<-rbind(counts.df,evdat)


##interactions


ints<-matrix(data=c("ints","act",length(grep("attack", actday[,1]))),nrow=1,ncol=3)


intsim<-lapply(simlist,function(x) {
	
	c("ints","sim",length(grep("attack", x[,1])))
	
})

library(abind)

intsim<-t(abind(intsim,along=2))

intdat<-data.frame(rbind(ints,intsim))

colnames(intdat)<-c("Category","Data","Count")

counts.df<-rbind(counts.df,intdat)


##cop number

cops<-matrix(data=c("cop","act",length(grep(".cop.start", actday[,1]))),nrow=1,ncol=3)


copsim<-lapply(simlist,function(x) {
	
	c("cop","sim",length(grep(".cop.start", x[,1])))
	
})

library(abind)

copsim<-t(abind(copsim,along=2))

copdat<-data.frame(rbind(cops,copsim))

colnames(copdat)<-c("Category","Data","Count")

counts.df<-rbind(counts.df,copdat)


## Overall Proportion of cops interrupted

copints<-matrix(data=c("cop.inter","act",1-(length(grep(".cop.end", actday[,1]))/length(grep(".cop.start", actday[,1])))),nrow=1,ncol=3)


copintsim<-lapply(simlist,function(x) {
	
	c("cop.inter","sim",1-(length(grep(".cop.end", x[,1]))/length(grep(".cop.start", x[,1]))))
	
})

library(abind)

copintsim<-t(abind(copintsim,along=2))

copintdat<-data.frame(rbind(copints,copintsim))

colnames(copintdat)<-c("Category","Data","Count")

counts.df<-rbind(counts.df,copintdat)


## Make the plots of overall counts for a day

#str(counts.df)
counts.df$Count<-as.numeric(as.character(counts.df$Count))
counts.df$Category<-as.factor(counts.df$Category)

counts.split<-split(counts.df,counts.df$Category)

library(ggplot2)
library(gridExtra)
library(cowplot)

plotlist<-list()

for(i in 1:length(counts.split)){

pic<-ggplot()+geom_boxplot(data=subset(counts.split[[i]],counts.split[[i]]$Data=="sim"),aes(y=Count))+geom_boxplot(data=subset(counts.split[[i]],counts.split[[i]]$Data=="act"),aes(y=Count,colour="red"))+theme_classic()+theme(legend.position="none",axis.title.x = element_text(size = 14, face = "bold"), axis.ticks.x=element_blank(),axis.text.x=element_blank(),axis.text.y=element_text(face="bold"),axis.title.y=element_text(face="bold"))+labs(x=levels(counts.split[[i]]$Category)[i])

if(i>1){pic<-pic+theme(axis.title.y=element_blank())}

plotlist[[i]]<-pic

}

countsplot<-plot_grid(plotlist[[1]],plotlist[[2]],plotlist[[3]],plotlist[[4]],nrow=1,ncol=length(plotlist),align="hv",axis="lb")

datfile<-paste0("./sim_actualday_plots/day_level_plots_",dates[b])
if(!dir.exists(datfile)) dir.create(datfile)

suppressWarnings(ggsave(paste0(datfile,"/countsplot_",dates[b]),plot=countsplot,device="pdf",width=6,height=4,units="in"))

counts.df$Category<-as.character(counts.df$Category)
overallcounts[[b]]<-counts.df


######## Freq. Distributions of metrics across nodes on one day ########

daylist<-list()
daylist[[1]]<-actday
daylist<-append(daylist,simlist)


male.pres<-read.table("CHG_2014_male_presence_data_quest.csv", sep=",", header=T)
#male IDs are already in ascending order, so I can convert them directly
colnames(male.pres)<-c("Date", 1:30)

#pare down to the day in question
male.pres<-male.pres[b,]


#######
## Distribution of mating success
#######


copdataday<-list()

copdist<-factor(as.character(actday[,1][grep(".cop.start", actday[,1])]),levels=event.types[grep(".cop.start",event.types)])

coptab<-table(copdist)

#this is the one I can save for the overall plot
coptab.df<-data.frame(coptab)
coptab.df[male.pres[,2:ncol(male.pres)]==0,2]<-NA

copdataday[[1]]<-coptab.df

#rank-order for day-level plot 

coptab.plot<-coptab.df
coptab.plot$copdist<-factor(coptab.plot$copdist,levels=event.types[grep(".cop.start",event.types)][order(table(copdist),decreasing=T)])

coptab.plot[male.pres[,2:ncol(male.pres)]==0,2]<-NA


#go thru the rest of the days 

coplist<-list()

for(i in 2:length(daylist)){

	
copdf<-factor(as.character(daylist[[i]][,1][grep(".cop.start", daylist[[i]][,1])]),levels=event.types[grep(".cop.start",event.types)])

if(length(copdf)>0){


copdf<-table(copdf)
copdf<-data.frame(copdf)

coplist[[i]]<-copdf

} else {coplist[[i]]<-data.frame("copdf"=event.types[grep(".cop.start",event.types)],"Freq"=rep(0,30))}


}

copdist.sim<-coplist[[1]]

for(i in 2:length(coplist)){
	
	copdist.sim<-rbind(copdist.sim,coplist[[i]])
}

copdist.sim[male.pres[,2:ncol(male.pres)]==0,2]<-NA


#rank-order sim factor levels to match the actual
copdist.sim$copdf<-factor(copdist.sim$copdf,levels=levels(coptab.plot$copdist))

xlabs<-sapply(strsplit(levels(coptab.plot$copdist),split=".",fixed=T),function(x) x[1])

#revert the label for X males so it's clear in the figure
xlabs[grep("30",xlabs)]<-"X"

copdist.plot<-ggplot()+geom_boxplot(data=copdist.sim, aes(x=copdf, y=Freq))+geom_boxplot(data=coptab.plot, aes(x=copdist, y=Freq,colour="red"))+geom_line(data=coptab.plot,aes(x=as.numeric(copdist),y=Freq,colour="red"))+theme_classic()+theme(legend.position="none",axis.title.x = element_text(size = 14, face="bold"),axis.text.x=element_text(face="bold"),axis.text.y=element_text(face="bold"),axis.title.y=element_text(face="bold"))+scale_x_discrete(labels=xlabs)+scale_y_continuous(breaks=function(x) pretty(x))+labs(x="Distribution of Solicitations by Male ID",y="Count")


suppressWarnings(ggsave(paste0(datfile,"/cop_distribution_plot_",dates[b]), plot=copdist.plot, device="pdf",width=6,height=4,units="in"))


#get the means and sds for the day and save them
#these are not in rank-order

meancops<-ave(copdist.sim$Freq,copdist.sim$copdf)[1:30]

#for a poisson-distrubuted variable, the variance=mean, so the sd=sqrt(variance)

sdcops<-sqrt(meancops)

#sdcops<-ave(copdist.sim$Freq,copdist.sim$copdf, FUN=function(x) sd(x))[1:30]

simcoptab<-data.frame(copdist=coptab.df$copdist,meancops=meancops,sdcops=sdcops)

copdataday[[2]]<-simcoptab


#######
## Dist. of Copulation Interruption RATES
#######


intdist<-c(1,1,1)

for(m in 1:30){
	
	copnum<-length(actday[,1][startsWith(as.character(actday[,1]),paste0(m,".cop.start"))])

if(copnum>0){
	intrate<-
	(1-(length(actday[,1][startsWith(as.character(actday[,1]),paste0(m,".cop.end"))])/copnum))	
	
	intnum<-copnum-length(actday[,1][startsWith(as.character(actday[,1]),paste0(m,".cop.end"))])
	
	} else {
		intrate<-NA
		intnum<-0
		}
	

	
intdist<-rbind(intdist,c(m,intrate,intnum))	
	
}

#remove dummy row
intdist<-intdist[-1,]

#reorder by rank for plot

intdist.plot<-intdist[order(intdist[,2],decreasing=T),]

intdist.plot<-data.frame(intdist.plot)
colnames(intdist.plot)<-c("Male","rate","count")
intdist.plot$Male<-factor(intdist.plot$Male, levels=as.character(intdist.plot$Male))


#go thru the rest of the sim days 

interlist<-list()

for(i in 2:length(daylist)){

simdist<-c(1,1,1)

for(m in 1:30){
	
	copnum<-length(daylist[[i]][,1][startsWith(as.character(daylist[[i]][,1]),paste0(m,".cop.start"))])

if(copnum>0){
	intrate<-
	(1-(length(daylist[[i]][,1][startsWith(as.character(daylist[[i]][,1]),paste0(m,".cop.end"))])/copnum))	
	intnum<-copnum-length(daylist[[i]][,1][startsWith(as.character(daylist[[i]][,1]),paste0(m,".cop.end"))])

	} else {
		intrate<-NA
		intnum<-0
		}
	
simdist<-rbind(simdist,c(m,intrate,intnum))	
	
}

#remove dummy row
simdist<-simdist[-1,]

simdist<-data.frame(simdist)
colnames(simdist)<-c("Male","rate","count")

#reorder by actual rank for plot
simdist$Male<-factor(simdist$Male, levels=as.character(intdist.plot$Male))

interlist[[i-1]]<-simdist	


}

interlist.sim<-interlist[[1]]

for(i in 2:length(interlist)){
	
	interlist.sim<-rbind(interlist.sim,interlist[[i]])
}

xlabs<-levels(interlist.sim$Male)

#revert the label for X males so it's clear in the figure
xlabs[grep("30",xlabs)]<-"X"

interdist.plot<-ggplot()+geom_boxplot(data=interlist.sim, aes(x=Male, y=rate))+geom_boxplot(data=intdist.plot, aes(x=Male, y=rate,colour="red"))+geom_line(data=intdist.plot,aes(x=as.numeric(1:30),y=rate,colour="red"))+theme_classic()+theme(legend.position="none",axis.title.x = element_text(size = 14, face="bold"),axis.text.x=element_text(face="bold"),axis.text.y=element_text(face="bold"),axis.title.y=element_text(face="bold"))+scale_x_discrete(labels=xlabs)+scale_y_continuous(breaks=function(x) pretty(x))+labs(x="Distribution of Copulation Interruption Rates by Male ID")


suppressWarnings(ggsave(paste0(datfile,"/inter_distribution_plot_",dates[b]), plot=interdist.plot, device="pdf",width=6,height=4,units="in"))


#non rankordered data frames for the overall plot
intdist<-data.frame(intdist)
colnames(intdist)<-c("Male","rate","count")

intdist$Male<-factor(intdist$Male)

copdataday[[3]]<-intdist


#get the means and sds for the day and save them
#these are not in rank-order

meanints<-ave(interlist.sim$count,interlist.sim$Male, FUN=function(x) mean(x, na.rm=T))[1:30]

sdints<-ave(interlist.sim$count,interlist.sim$Male, FUN=function(x) sd(x, na.rm=T))[1:30]

siminttab<-data.frame(Male=intdist$Male,meanints=meanints,sdints=sdints)

copdataday[[4]]<-siminttab


#######
## Distribution of uninterrupted copulations
#######

#UNinterrupted copulations are simply the number of cop.ends! 


#first the observed data

enddist<-factor(as.character(actday[,1][grep(".cop.end", actday[,1])]),levels=event.types[grep(".cop.end",event.types)])

endtab<-table(enddist)

#this is the one I can save for the overall plot
endtab.df<-data.frame(endtab)
endtab.df[male.pres[,2:ncol(male.pres)]==0,2]<-NA

copdataday[[5]]<-endtab.df

#rank-order for day-level plot 

endtab.plot<-endtab.df
endtab.plot$enddist<-factor(endtab.plot$enddist,levels=event.types[grep(".cop.end",event.types)][order(table(enddist),decreasing=T)])

endtab.plot[male.pres[,2:ncol(male.pres)]==0,2]<-NA


#go thru the rest of the days 

endlist<-list()

for(i in 2:length(daylist)){

	
enddf<-factor(as.character(daylist[[i]][,1][grep(".cop.end", daylist[[i]][,1])]),levels=event.types[grep(".cop.end",event.types)])

if(length(enddf)>0){


enddf<-table(enddf)
enddf<-data.frame(enddf)

endlist[[i]]<-enddf

} else {endlist[[i]]<-data.frame("enddf"=event.types[grep(".cop.end",event.types)],"Freq"=rep(0,30))}


}

enddist.sim<-endlist[[1]]

for(i in 2:length(endlist)){
	
	enddist.sim<-rbind(enddist.sim,endlist[[i]])
}

enddist.sim[male.pres[,2:ncol(male.pres)]==0,2]<-NA


#rank-order sim factor levels to match the actual
enddist.sim$enddf<-factor(enddist.sim$enddf,levels=levels(endtab.plot$enddist))

xlabs<-sapply(strsplit(levels(endtab.plot$enddist),split=".",fixed=T),function(x) x[1])

#revert the label for X males so it's clear in the figure
xlabs[grep("30",xlabs)]<-"X"

enddist.plot<-ggplot()+geom_boxplot(data=enddist.sim, aes(x=enddf, y=Freq))+geom_boxplot(data=endtab.plot, aes(x=enddist, y=Freq,colour="red"))+geom_line(data=endtab.plot,aes(x=as.numeric(enddist),y=Freq,colour="red"))+theme_classic()+theme(legend.position="none",axis.title.x = element_text(size = 14, face="bold"),axis.text.x=element_text(face="bold"),axis.text.y=element_text(face="bold"),axis.title.y=element_text(face="bold"))+scale_x_discrete(labels=xlabs)+scale_y_continuous(breaks=function(x) pretty(x))+labs(x="Distribution of Uninterrupted\nCopulations by Male ID",y="Count")


suppressWarnings(ggsave(paste0(datfile,"/end_distribution_plot_",dates[b]), plot=enddist.plot, device="pdf",width=6,height=4,units="in"))


#get the means and sds for the day and save them
#these are not in rank-order

meanends<-ave(enddist.sim$Freq,enddist.sim$enddf)[1:30]

#for a poisson distribution, which these events are (I think!) the variance=mean and the sd=sqrt(variance)

sdends<-sqrt(meanends)

#sdends<-ave(enddist.sim$Freq,enddist.sim$enddf, FUN=function(x) sd(x))[1:30]

simendtab<-data.frame(enddist=endtab.df$enddist,meanends=meanends,sdends=sdends)

copdataday[[6]]<-simendtab


##I'll need to do a version of this that averages across all the days

#save to the full dataset object
overallcop[[b]]<-copdataday


########
## Dist. of Fighting times
########

#need to get the win.loss.census for the simulated day

daylist<-list()
daylist[[1]]<-actday
daylist<-append(daylist,simlist)

idno<-30

#I actually need to grab different code for these census lists for the actual data vs the simulated data


census.list<-list()


for(j in 1:length(daylist)){


if(j==1){

census<-matrix("nothing", nrow=1,ncol=(idno), dimnames=list(1,c(1:(idno))))


##code that translates each new current event into what the males involved are doing at that time. The support matrix will be based on whatever the males were doing in the PREVIOUS timestep. This census matrix itself could be interesting to visualize. Each row of the census says what each male is doing going into the NEXT event.


for(i in 1:nrow(daylist[[j]])){

#can ask if the second thing is something, and then move on to the first and third elements.

current<-census[nrow(census),]

#parse the event type. loop through test.seq rows
evname<-as.character(daylist[[j]][i,1])
evsplit<-strsplit(evname, split=".", fixed=T)

#if it is an end obs, arrival, departure or clock event, it shouldn't change anything the males are doing in terms of interactions. Departures aren't exogenous, but they shouldn't change anything since they can only happen once a male is already doing nothing. 
if(evsplit[[1]][2]!="obs" & evsplit[[1]][2]!="depart" & evsplit[[1]][2]!="arrive" & evsplit[[1]][2]!="time" & evsplit[[1]][1]!="fem"){
#if the event was an attack, indicate the males involved and which was the receiver since this dictates who makes the initition decision
	if(evsplit[[1]][2]=="attack"){
		current[evsplit[[1]][1]]<-evname
		current[evsplit[[1]][3]]<-paste(evsplit[[1]][3], ".", "received", ".", evsplit[[1]][1], sep="")
			}
	#if it's an FO or SF, just indicate that that pair of males are involved in a fight now
		if(evsplit[[1]][2]=="initFO"||evsplit[[1]][2]=="initSF"){
			#need to refer back to the previous row of "census" to get the id of the other male
			
			current[evsplit[[1]][1]]<-paste(evsplit[[1]][1], ".", "fight", ".",strsplit(census[nrow(census),evsplit[[1]][1]], split=".",fixed=T)[[1]][3], sep="")
			current[[strsplit(census[nrow(census),evsplit[[1]][1]], split=".",fixed=T)[[1]][3]]]<-paste(strsplit(census[nrow(census),evsplit[[1]][1]], split=".",fixed=T)[[1]][3], ".", "fight", ".",evsplit[[1]][1], sep="")
			
			}
			
	#if it's a chase, indicate that the pair of males are involved in a fight, but there is an additional constraint that the individual DOING THE CHASING will be the disengager by definition
	if(evsplit[[1]][2]=="initCH"){
			
		current[evsplit[[1]][1]]<-paste(evsplit[[1]][1], ".", "fight", ".",strsplit(census[nrow(census),evsplit[[1]][1]], split=".",fixed=T)[[1]][3], ".chasedby",sep="")
			current[[strsplit(census[nrow(census),evsplit[[1]][1]], split=".",fixed=T)[[1]][3]]]<-paste(strsplit(census[nrow(census),evsplit[[1]][1]], split=".",fixed=T)[[1]][3], ".", "fight", ".",evsplit[[1]][1], ".chasing",sep="")

	}

#for disengaging, end the fight for the partner male as well
	if(evsplit[[1]][2]=="disengage"){
	current[evsplit[[1]][1]]<-"nothing"
	current[evsplit[[1]][3]]<-"nothing"	
	
}
 

#for copulations, end cops are only possible after start cops, but otherwise anything else is possible. EXCEPT the copulating male can't initiate an attack. If the next event involving that male ISN'T an end cop, it was interrupted

	if(evsplit[[1]][2]=="cop" && evsplit[[1]][3]=="start"){
		current[evsplit[[1]][1]]<-"copulating"
	}

	if(evsplit[[1]][2]=="cop" && evsplit[[1]][3]=="end"){
		current[evsplit[[1]][1]]<-"nothing"
	}

#if an X male is still involved in a fight at all, make sure he is still tagged as fighting (since sometimes more than one is interacting at once, but they can't depart until none are fighting)
	if((TRUE %in% grepl("fight.30",current) | TRUE %in% grepl("30.fight",current)) && !(grepl("receive", current[idno]) | grepl("attack", current[idno]))) {
		
		current[idno]<-"fight"

}

}

census<-rbind(census,current)


}


#don't want the first dummy row
census.list[[j]]<-census[-1,]

} else {
	
	
census<-matrix("nothing", nrow=1,ncol=(idno), dimnames=list(1,c(1:(idno))))


##code that translates each new current event into what the males involved are doing at that time. The support matrix will be based on whatever the males were doing in the PREVIOUS timestep. This census matrix itself could be interesting to visualize. Each row of the census says what each male is doing going into the NEXT event.


for(i in 1:nrow(daylist[[j]])){

#can ask if the second thing is something, and then move on to the first and third elements.

current<-census[nrow(census),]

#parse the event type. loop through test.seq rows
evname<-as.character(daylist[[j]][i,1])
evsplit<-strsplit(evname, split=".", fixed=T)

#if it is an end obs, arrival, departure or clock event, it shouldn't change anything the males are doing in terms of interactions. Departures aren't exogenous, but they shouldn't change anything since they can only happen once a male is already doing nothing. 
if(evsplit[[1]][2]!="obs" & evsplit[[1]][2]!="depart" & evsplit[[1]][2]!="arrive" & evsplit[[1]][2]!="time" & evsplit[[1]][1]!="fem"){
#if the event was an attack, indicate the males involved and which was the receiver since this dictates who makes the initition decision
	if(evsplit[[1]][2]=="attack"){
		current[evsplit[[1]][1]]<-evname
		current[evsplit[[1]][3]]<-paste(evsplit[[1]][3], ".", "received", ".", evsplit[[1]][1], sep="")
			}
	#if it's an FO or SF, just indicate that that pair of males are involved in a fight now
		if(evsplit[[1]][2]=="initFO"||evsplit[[1]][2]=="initSF"){
			#need to refer back to the previous row of "census" to get the id of the other male
			
			current[evsplit[[1]][1]]<-paste(evsplit[[1]][1], ".", "fight", ".",strsplit(census[nrow(census),evsplit[[1]][1]], split=".",fixed=T)[[1]][3], sep="")
			current[[strsplit(census[nrow(census),evsplit[[1]][1]], split=".",fixed=T)[[1]][3]]]<-paste(strsplit(census[nrow(census),evsplit[[1]][1]], split=".",fixed=T)[[1]][3], ".", "fight", ".",evsplit[[1]][1], sep="")
			
			}
			
	#if it's a chase, indicate that the pair of males are involved in a fight, but there is an additional constraint that the individual DOING THE CHASING will be the disengager by definition
	if(evsplit[[1]][2]=="initCH"){
			
		current[evsplit[[1]][1]]<-paste(evsplit[[1]][1], ".", "fight", ".",strsplit(census[nrow(census),evsplit[[1]][1]], split=".",fixed=T)[[1]][3], ".chasedby",sep="")
			current[[strsplit(census[nrow(census),evsplit[[1]][1]], split=".",fixed=T)[[1]][3]]]<-paste(strsplit(census[nrow(census),evsplit[[1]][1]], split=".",fixed=T)[[1]][3], ".", "fight", ".",evsplit[[1]][1], ".chasing",sep="")

	}

#for disengaging, need to search through the previous rows to find the most recent "fight" and identify the partner and end it
	if(evsplit[[1]][2]=="disengage"){
	current[evsplit[[1]][1]]<-"nothing"
	current[strsplit(census[max(grep("fight", census[,evsplit[[1]][1]])),evsplit[[1]][1]], split=".", fixed=T)[[1]][3]]<-"nothing"	
	
}
 

#for copulations, end cops are only possible after start cops, but otherwise anything else is possible. EXCEPT the copulating male can't initiate an attack. If the next event involving that male ISN'T an end cop, it was interrupted

	if(evsplit[[1]][2]=="cop" && evsplit[[1]][3]=="start"){
		current[evsplit[[1]][1]]<-"copulating"
	}

	if(evsplit[[1]][2]=="cop" && evsplit[[1]][3]=="end"){
		current[evsplit[[1]][1]]<-"nothing"
	}

# #if an X male is still involved in a fight at all, make sure he is still tagged as fighting (since sometimes more than one is interacting at once, but they can't depart until none are fighting)
	# if((TRUE %in% grepl("fight.30",current) | TRUE %in% grepl("30.fight",current)) && !(grepl("receive", current[idno]) | grepl("attack", current[idno]))) {
		
		# current[idno]<-"fight"

# }

}

census<-rbind(census,current)


}

#don't want the first dummy row
census.list[[j]]<-census[-1,]

	}

}


#convert to win.loss census- again need two diferent functions for the different datasets b/c of constraints on X males 

create.win.loss.census.act<-function(x){

#get disengagement events for focal male from evlist
win.loss.census<-x

#loop thru days and males

for(i in 1:length(win.loss.census)){

for(l in 1:idno){


dis.times<-which(startsWith(as.character(daylist[[i]][,1]), paste(l,".disengage", sep="")))


if(length(dis.times)>0){

evsplit<-strsplit(as.character(daylist[[i]][dis.times,1]),".",fixed=T)

#so we have a list of row numbers for disengagements and a list of interaction partners. This will cover wins for normal fights and losses for chases for the focal male. If we loop through all the males, all the wins and losses should get filled in.

#NOW DISENGAGES ALWAYS MEAN WINS, so we will still have chase information, but we don't need to do anything special with it here.

#need to make sure we are really precise since male 30 can be involved in multiple interactions, but they're usually all chases.


for(j in 1:length(dis.times)){
	
		
	if(grepl("chasing",win.loss.census[[i]][(dis.times[j]-1),l]) ==FALSE && grepl("chasedby",win.loss.census[[i]][(dis.times[j]-1),as.numeric(evsplit[[j]][3])]) ==FALSE){
	#apply the win in the focal male's column
	win.loss.census[[i]][dis.times[j],l]<-paste(l, "win",evsplit[[j]][3], sep=".")
	#apply the loss for the same interaction in the partner male's column
	win.loss.census[[i]][dis.times[j],as.numeric(evsplit[[j]][3])]<-paste(evsplit[[j]][3], "lose", l, sep=".")
	} else{
		#apply the won chase in the focal male's column
	win.loss.census[[i]][dis.times[j],l]<-paste(l, "win",evsplit[[j]][3],"chase", sep=".")
		#apply the loss for the same interaction in the partner male's column.
	win.loss.census[[i]][dis.times[j],as.numeric(evsplit[[j]][3])]<-paste(evsplit[[j]][3], "lose", l, "chase",sep=".")
		}
		
}

}

}

}

return(win.loss.census)
	
}

win.loss.census<-create.win.loss.census.act(census.list[1])


create.win.loss.census.sim<-function(x){

#get disengagement events for focal male from evlist
win.loss.census<-x

#loop thru days and males

for(i in 1:length(win.loss.census)){

for(l in 1:idno){

dis.times<-grep(TRUE,startsWith(as.character(daylist[[i+1]][,1]), paste(l,".disengage", sep="")))


if(length(dis.times)>0){

#see who they were fighting with
fights<-win.loss.census[[i]][dis.times-1,l]

#so we have a list of row numbers for disengagements and a list of interaction partners. This will cover wins for normal fights and losses for chases for the focal male. If we loop through all the males, all the wins and losses should get filled in.

#NOW DISENGAGES ALWAYS MEAN WINS, so we will still have chase information, but we don't need to do anything special with it here.

for(j in 1:length(dis.times)){
	
	if(grepl("chasing",fights[j]) ==FALSE){
	#apply the win in the focal male's column
	win.loss.census[[i]][dis.times[j],l]<-paste(l, "win",strsplit(fights[j],".", fixed=T)[[1]][3], sep=".")
	#apply the loss for the same interaction in the partner male's column
	win.loss.census[[i]][dis.times[j],as.numeric(strsplit(fights[j],".", fixed=T)[[1]][3])]<-paste(strsplit(fights[j],".", fixed=T)[[1]][3], "lose", l, sep=".")
	} else{
		#apply the won chase in the focal male's column
	win.loss.census[[i]][dis.times[j],l]<-paste(l, "win",strsplit(fights[j],".", fixed=T)[[1]][3],"chase", sep=".")
		#apply the loss for the same interaction in the partner male's column.
	win.loss.census[[i]][dis.times[j],as.numeric(strsplit(fights[j],".", fixed=T)[[1]][3])]<-paste(strsplit(fights[j],".", fixed=T)[[1]][3], "lose", l, "chase",sep=".")
		}
		
}

}

}

}

return(win.loss.census)
	
}

win.loss.census.sim<-create.win.loss.census.sim(census.list[2:length(census.list)])


win.loss.census<-append(win.loss.census,win.loss.census.sim)


## Now get fighting and attendance times

dist.fighting.time<-function(x){

rec.stat<-x

fightlist<-list()

#loop through each day sequence
for(k in 1:length(rec.stat)){

#this loops through each event in the sequence

step<-nrow(rec.stat[[k]])

for(j in step){

#maybe make this an empty matrix instead
fightvect<-matrix(0,nrow=4,ncol=(ncol(census.list[[1]])-1))

#for each event, loop through all the males- the support array will take care of males that weren't present that day.
for(i in 1:(ncol(census.list[[1]])-1)){
	
#get pairs of starts and ends of interactions that preserve the row number of the original event sequence
	
wins<-grep("win",win.loss.census[[k]][1:(j-1),i])
losses<-grep("lose",win.loss.census[[k]][1:(j-1),i])
end.times<-union(wins,losses)
end.times<-end.times[order(end.times)]

attacks<-grep("attack",win.loss.census[[k]][1:(j-1),i])
receives<-grep("received",win.loss.census[[k]][1:(j-1),i])
start.times<-union(attacks,receives)
start.times<-start.times[order(start.times)]

pairs<-union(start.times,end.times)


if(length(pairs)>=2){

pairs<-pairs[order(pairs)]

#in this case, I still want to know what the cumulative fighting time was for a male, even if he was in the middle of an interaction when the simulation ended. 
if((length(pairs) %% 2)==1){pairs<-pairs[-length(pairs)]}

pairs.times<-rec.stat[[k]][pairs,"Time"]

#sum up the time intervals spent fighting
intervals<-matrix(pairs.times,nrow=2)

fight.tot<-sum(intervals[2,]-intervals[1,])

fightvect[1,i]<-fight.tot

} else {fight.tot<-0}


##get the total time the focal male was present on the lek so far today

arrsub<-rec.stat[[k]][grep("arrive|depart",rec.stat[[k]][,1]),]

arrsub$Events<-as.character(arrsub$Events)

arrind<-arrsub[startsWith(arrsub$Events,paste0(i,".")),]

#if the male was there that day according to the presence data but has no departure, then they were there the whole time. Create a special object in this case. 

if(!nrow(arrind)>0 && male.pres[,i+1]==1){arrind[1,]<-c(paste0(i,".arrive"),0,1,dates[b])}

if(nrow(arrind)>0){

#if the first entry is a depart, they were there at time 0, so add one in
if(endsWith(arrind[1,1],"depart")){arrind<-rbind(c(paste0(i,".arrive"),0,1,dates[b]),arrind)}

arrind$Time<-as.numeric(arrind$Time)


if((nrow(arrind) %% 2)==0){
	
	tot.time<-sum(sapply(1:(nrow(arrind)/2), function(x) return(arrind[(2*x),2]-arrind[(2*x-1),2])))
	
}

if(nrow(arrind)==1){
	
	tot.time<-(rec.stat[[k]][(j),"Time"]-arrind[nrow(arrind),2])
}

if((nrow(arrind) %% 2)!=0 && nrow(arrind)!=1){
	
	tot.time<-sum(sapply(1:floor(nrow(arrind)/2), function(x) return(arrind[(2*x),2]-arrind[(2*x-1),2])))+(rec.stat[[k]][(j),"Time"]-arrind[nrow(arrind),2])
	
}


if(length(tot.time)>0 && tot.time>0){
perc.fight<-100*(fight.tot/tot.time)

fightvect[2,i]<-tot.time
fightvect[3,i]<-perc.fight

#get % of total day in attendance
tmax<-rec.stat[[k]][(j),"Time"]
perc.attend<-100*(tot.time/tmax)

fightvect[4,i]<-perc.attend

}

}


}

}

fightlist[[k]]<-fightvect

}

return(fightlist)

}


fight.list<-dist.fighting.time(daylist)


## Finally, can plot these

#use male.pres data to replace zeroes with NAs

for(i in 1:length(fight.list)){
	
	fight.list[[i]][,male.pres[,2:(ncol(male.pres)-1)]==0]<-NA
	
}


day1<-data.frame(t(fight.list[[1]]))

day1$Male<-as.factor(c(1:29))

colnames(day1)<-c("Fighting.Time","Total.Time","Perc.Fighting","Perc.Attend","Male")

for(i in 2:length(fight.list)){
	
	fight.list[[i]]<-data.frame(t(fight.list[[i]]))
	fight.list[[i]]$Male<-c(1:29)
}

restfight<-data.frame(abind(fight.list[2:length(fight.list)],along=1))

restfight[,1]<-as.numeric(as.character(restfight[,1]))
restfight[,2]<-as.numeric(as.character(restfight[,2]))
restfight[,3]<-as.numeric(as.character(restfight[,3]))
restfight$Male<-as.factor(as.numeric(as.character(restfight$Male)))

colnames(restfight)<-c("Fighting.Time","Total.Time","Perc.Fighting","Perc.Attend","Male")

#save for later
day1save<-day1
day1save$Date<-rep(dates[b],nrow(day1save))

restfightsave<-restfight
restfightsave$Date<-rep(dates[b],nrow(restfightsave))


fightframe<-list()

fightframe[[1]]<-day1save
fightframe[[2]]<-restfightsave

overallfight[[b]]<-fightframe

## try rank-ordering but retain the ID numbers

library(tidyverse)
library(forcats)

day1<-day1 %>%
mutate(Male=fct_reorder(Male,desc(Fighting.Time), .fun='median'))

restfight$Male<-factor(restfight$Male,levels(day1$Male))


#just for the plots, remove males that weren't there 
rankord<-levels(day1$Male)
rankord<-rankord[male.pres[,1+as.numeric(rankord)]==1]

day1plot<-day1[!is.na(day1$Total.Time),]
day1plot$Male<-factor(day1plot$Male,rankord)

restfightplot<-restfight[!is.na(restfight$Total.Time),]
restfightplot$Male<-factor(restfightplot$Male,rankord)

#emulate ggplot colors

gg_color_hue <- function(n) {
  hues = seq(15, 375, length = n + 1)
  hcl(h = hues, l = 65, c = 100)[1:n]
}

actfight<-ggplot()+geom_boxplot(data=restfightplot,aes(x=Male,y=Fighting.Time,colour="black"))+geom_boxplot(data=day1plot,aes(x=Male,y=Fighting.Time,colour="red"))+theme_classic()+theme(axis.title.x = element_text(size = 14, face = "bold"),axis.text.x=element_text(face="bold"),axis.text.y=element_text(face="bold"),axis.title.y=element_text(face="bold"),legend.title=element_text(face="bold"))+geom_line(data=day1plot,aes(x=as.numeric(Male),y=Fighting.Time,colour="red"))+scale_colour_manual(values=c("black",gg_color_hue(1)),name="Data Type",labels=c("Simulations","Observed"))+labs(x="Male ID", y="Total Time Spent Fighting")

# acttot<-ggplot()+geom_boxplot(data=restfightplot,aes(x=Male,y=Total.Time))+geom_boxplot(data=day1plot,aes(x=Male,y=Total.Time,colour="red"))+theme_classic()+theme(legend.position="none",axis.title.x = element_text(size = 14, face = "bold"),axis.text.x=element_text(face="bold"),axis.text.y=element_text(face="bold"),axis.title.y=element_text(face="bold"))+geom_line(data=day1plot,aes(x=as.numeric(Male),y=Total.Time,colour="red"))

# actperc<-ggplot()+geom_boxplot(data=restfightplot,aes(x=Male,y=Perc.Fighting))+geom_boxplot(data=day1plot,aes(x=Male,y=Perc.Fighting,colour="red"))+theme_classic()+theme(legend.position="none",axis.title.x = element_text(size = 14, face = "bold"),axis.text.x=element_text(face="bold"),axis.text.y=element_text(face="bold"),axis.title.y=element_text(face="bold"))+geom_line(data=day1plot,aes(x=as.numeric(Male),y=Perc.Fighting,colour="red"))

# actattend<-ggplot()+geom_boxplot(data=restfightplot,aes(x=Male,y=Perc.Attend))+geom_boxplot(data=day1plot,aes(x=Male,y=Perc.Attend,colour="red"))+theme_classic()+theme(legend.position="none",axis.title.x = element_text(size = 14, face = "bold"),axis.text.x=element_text(face="bold"),axis.text.y=element_text(face="bold"),axis.title.y=element_text(face="bold"))+geom_line(data=day1plot,aes(x=as.numeric(Male),y=Perc.Attend,colour="red"))


# fightplot<-plot_grid(actfight,actperc,acttot,actattend,nrow=2,ncol=2,align="vh",axis="lb")


# Rearrange- Plot totals and %'s together with male ID as a factor? 

percfight<-day1plot[,c(3,5)]
percfight$type<-rep("percfight",nrow(day1plot))
colnames(percfight)[1]<-"percent"

percatt<-day1plot[,c(4,5)]
percatt$type<-rep("percatt",nrow(day1plot)) 
colnames(percatt)[1]<-"percent"

percdat<-rbind(percfight,percatt)

#for the sims

percfight<-restfightplot[,c(3,5)]
percfight$type<-rep("percfight",nrow(restfightplot))
colnames(percfight)[1]<-"percent"

percatt<-restfightplot[,c(4,5)]
percatt$type<-rep("percatt",nrow(restfightplot)) 
colnames(percatt)[1]<-"percent"

percsim<-rbind(percfight,percatt)

#bars in the background to group the boxplots
barback<-cbind(as.character(day1plot$Male),rep(100,nrow(day1plot)))
barback<-data.frame(barback)
barback$X1<-factor(barback$X1,levels(day1plot$Male))
barback$X2<-as.numeric(as.character(barback$X2))


#center the lines within the "boxes" for the different data types

percdat$midbox<-sapply(c(1:nrow(percdat)),function(x){
	
	if(percdat$type[x]=="percfight"){return(as.numeric(percdat$Male[x])+(11/60))} else {return(as.numeric(percdat$Male[x])-(11/60))}
	
	
})


percplot<-ggplot()+geom_bar(data=barback,stat="identity",aes(x=X1,y=X2),fill="grey90")+geom_boxplot(data=percsim,aes(x=Male,y=percent,fill=type))+scale_fill_manual(values=c("white","grey"),name="Simulations", labels=c("Percent of Total Time\nin Attendance on Lek\n ","Percent of Time\non the Lek Spent Fighting\n"))+geom_boxplot(data=percdat,aes(x=Male,y=percent,colour=type))+scale_colour_manual(values=c("red","red"),name="acttype",guide=FALSE)+geom_line(data=percdat,aes(x=midbox,y=percent,linetype=type),colour="red")+scale_linetype_discrete(name="Observed",labels=c("Percent of Total Time\nin Attendance on Lek\n ","Percent of Time\non the Lek Spent Fighting\n"))+theme_classic()+theme(axis.title.x = element_text(size = 14, face = "bold"),axis.text.x=element_text(face="bold"),axis.text.y=element_text(face="bold"),axis.title.y=element_text(face="bold"),legend.title=element_text(face="bold"),legend.position="right")+guides(acttype=FALSE)+labs(x="Male ID", y="Percent Time")


fightsplots<-plot_grid(actfight,percplot,nrow=2,ncol=1,align="v",axis="b")

ggsave(paste0(datfile,"/fighting_time_dist_",dates[b]),plot=fightsplots,device="pdf",height=7,width=7,units="in")


## Try rank-ordering but without retaining IDs

# # dayrank<-day1

# dayrank<-dayrank[order(dayrank$Time, decreasing=T),]

# dayrank$Male<-c(1:29)

# dayrank$Male<-as.factor(as.numeric(as.character(dayrank$Male)))


# for(i in 2:length(fight.list)){
	
	# fight.list[[i]]<-data.frame(fight.list[[i]])
	# fight.list[[i]][,1]<-fight.list[[i]][,1][order(fight.list[[i]][,1], decreasing=T)]
	# fight.list[[i]]$Male<-c(1:29)
# }

# restrank<-data.frame(abind(fight.list[2:length(fight.list)],along=1))

# restrank[,1]<-as.numeric(as.character(restrank[,1]))
# restrank$Male<-as.factor(as.numeric(as.character(restrank$Male)))

# colnames(restrank)<-c("Time","Male")


# rankfight<-ggplot()+geom_boxplot(data=restrank,aes(x=Male,y=Time))+geom_boxplot(data=dayrank,aes(x=Male,y=Time,colour="red"))+theme_classic()+theme(legend.position="none",axis.title.x = element_text(size = 14, face = "bold"),axis.text.x=element_text(face="bold"),axis.text.y=element_text(face="bold"),axis.title.y=element_text(face="bold"))+geom_line(data=dayrank,aes(x=as.numeric(Male),y=Time,colour="red"))+labs(x="Rank Order", y="Total Fighting Time")

# ggsave("rank_fighting_time_dist",plot=rankfight,device="pdf")


######## Dist. of network metrics #########


#turn rem-format data back into network data 

#focus on attacks/receives for now. BUT since we did it for the fighting times, we actually have a list of the accurate win.loss.census matrices, with the first one being the actual day, and the rest being sims of that day


#make sure to grab everyone who was there in case the sim ends up with certain males not interacting at all. 

male.pres<-read.table("CHG_2014_male_presence_data_quest.csv", sep=",", header=T)
#male IDs are already in ascending order, so I can convert them directly
colnames(male.pres)<-c("Date", 1:30)

#pare down to the day in question
male.pres<-male.pres[b,]

nodes<-as.character((which(male.pres==1)-1))


##create networks for the actual day and each sim day

metnodelist<-list()
metnetlist<-list()

for(j in 1:length(daylist)){


netevents<-table(grep("attack",daylist[[j]][,1],value=T))

#check for days with no interactions
if(dim(netevents)!=0){

#get the ties for the network, in this case attacks
edgelist<-t(abind(strsplit(names(netevents),".",fixed=T),along=2))[,c(1,3)]

#in the case of a one- or no-attack day:
if(length(netevents)<=1){
edgelist<-t(edgelist)
}

edgelist<-cbind(edgelist,netevents)

colnames(edgelist)[3]<-"weight"


#reset empty adjacency matrix
adj<-matrix(0, nrow=length(nodes),ncol=length(nodes),dimnames=list(nodes,nodes))

#apply numbers in appropriate places in the adjacency matrix

for(i in 1:nrow(edgelist)){
	
	adj[edgelist[i,1],edgelist[i,2]]<-edgelist[i,3]
	
}


library(igraph)

#make compatible network objects for the different packages
#the igraph object handles the weighted edge attributes

edgelist.df<-data.frame(edgelist)
edgelist.df$weight<-as.numeric(as.character(edgelist.df$weight))

i.net<-igraph::graph.data.frame(edgelist.df, vertices=cbind(nodes,rep(0,length(nodes))),directed=T)

# summary(i.net)
# summary(sna.net)

#plot(i.net)
#plot(sna.net)

#check that they're equivalent
#get.adjacency(i.net,sparse=F,attr="weight")


#undirected version of weighted network for clustering analysis
undirnet<-igraph::as.undirected(i.net, mode="collapse", edge.attr.comb= "sum")
#plot(undirnet)


######### Caluculate Individual Network Metrics ###########


metlist<-nodes

#eigenvector centrality: highly connected, giving weight to being connected to other highly connected individuals. Got rid of Directed eigenvector centrality.
eigens_tot<-igraph::evcent(i.net)$vector
metlist<-cbind(metlist,eigens_tot)

#betweenness centrality- are you on the way to lots of pairs of nodes?
between<-igraph::betweenness(i.net, normalized=T)
metlist<-cbind(metlist,between)


#undirected degree distribution
deg<-igraph::degree(undirnet)
metlist<-cbind(metlist,deg)


#out-degree centrality:
outdegree<-igraph::degree(i.net, mode="out")
metlist<-cbind(metlist,outdegree)


#out-strength
outweight<-igraph::strength(i.net,mode="out")
metlist<-cbind(metlist,outweight)


#in-degree centrality
indegree<-igraph::degree(i.net, mode="in")
metlist<-cbind(metlist,indegree)

#in-strength
inweight<-igraph::strength(i.net,mode="in")
metlist<-cbind(metlist,inweight)


metnodelist[[j]]<-metlist


###### Calculate Network-Level Metrics ######

netlevel<-vector()

netlevel<-append(netlevel,igraph::graph.density(i.net))

netlevel<-append(netlevel,igraph::graph.density(undirnet))

netlevel<-append(netlevel,igraph::diameter(i.net))

netlevel<-append(netlevel,igraph::diameter(undirnet))

netlevel<-append(netlevel,igraph::transitivity(i.net, type="global"))

#use sna for edgewise reciprocity
library(sna)
snadj<-apply(adj,2,function(x) as.numeric(x))
sna.net<-network(snadj,directed=T,ignore.eval=F,names.eval="weight")

netlevel<-append(netlevel,sna::grecip(sna.net, measure="edgewise"))

names(netlevel)<-c("Density","Undir.Density","Diameter","Undir.Diameter","Transitivity", "Edgewise\nReciprocity")

metnetlist[[j]]<-netlevel

}
}


## In general, I think I want these to be rank-ordered rather than associated with specific males. 


#need a different ordering for each metric

library(tidyverse)

plotnames<-c("Eigenvector\n Centrality","Betweenness\n Centrality","Undirected\n Degree", "Out Degree", "Out Weight", "In Degree", "In Weight")

#first make the dataframes for the plots

metplotlist<-list()

for(k in 2:ncol(metnodelist[[1]])){


day1<-data.frame(metnodelist[[1]])[,c(1,k)]
colnames(day1)<-c("Male","Metric")

day1[,2]<-as.numeric(as.character(day1[,2]))

day1<-day1 %>%
mutate(Male=fct_reorder(Male,desc(Metric), .fun='median'))

colnames(day1)[2]<-colnames(metnodelist[[1]])[k]


temp<-list()

for(i in 2:length(metnodelist)){
	
	if(length(metnodelist[[i]])>0){
	
	temp[[i]]<-data.frame(metnodelist[[i]])[,c(1,k)]
	
	}
	
}

restmet<-data.frame(abind(temp[2:length(temp)],along=1))

restmet[,2]<-as.numeric(as.character(restmet[,2]))
colnames(restmet)<-c("Male",colnames(metnodelist[[1]])[k])

restmet$Male<-factor(restmet$Male,levels(day1$Male))


metplotlist[[k-1]]<-list(day1,restmet)


}

netmetsnodes<-function(j){

xlabs<-levels(metplotlist[[j]][[2]]$Male)

#revert the label for X males so it's clear in the figure
xlabs[grep("30",xlabs)]<-"X"
  
ggplot()+geom_boxplot(data=metplotlist[[j]][[2]],aes(x=Male,y=metplotlist[[j]][[2]][,2]))+geom_boxplot(data=metplotlist[[j]][[1]],aes(x=Male,y=metplotlist[[j]][[1]][,2],colour="red"))+theme_classic()+theme(legend.position="none",axis.title.x = element_text(size = 12, face = "bold"),axis.text.x=element_text(face="bold"),axis.text.y=element_text(face="bold"),axis.title.y=element_text(face="bold"))+geom_line(data=metplotlist[[j]][[1]],aes(x=as.numeric(Male),y=metplotlist[[j]][[1]][,2],colour="red"))+scale_x_discrete(labels=xlabs)+labs(x="Male ID Number",y=plotnames[j])

}


library(gridExtra)
library(cowplot)

nodemetsplot<-plot_grid(netmetsnodes(1),netmetsnodes(2),netmetsnodes(3),netmetsnodes(4),netmetsnodes(5),netmetsnodes(6),netmetsnodes(7),nrow=4,ncol=2, align="hv",axis="lb")

ggsave(paste0(datfile,"/nodemetsplot_",dates[b]),plot=nodemetsplot,device="pdf",width=11,height=7,units="in")


##Note: look into plotrange padding for ggplot to make the axees line up in the grid


## Plot the network-level measures in a form similar to the overall counts

#metnetlist

#re-organize a little bit

metnetdf<-data.frame("Category"=1,"Data"=1,"Count"=1)
 
for(i in 1:length(metnetlist)){
	
if(i==1){
	
	for(j in 1:length(metnetlist[[i]])){
		
	metnetdf<-rbind(metnetdf,c(names(metnetlist[[i]][j]),"act",metnetlist[[i]][j]))
	
	}
	
} else {
	if(length(metnetlist[[i]]>0)){
		
	for(j in 1:length(metnetlist[[i]])){
		
	metnetdf<-rbind(metnetdf,c(names(metnetlist[[i]][j]),"sim",metnetlist[[i]][j]))
	
	}
	
}	

}	
		
}

#remove dummy row
metnetdf<-metnetdf[-1,]

metnetdf$Category<-as.factor(metnetdf$Category)
metnetdf$Data<-as.factor(metnetdf$Data)
metnetdf$Count<-as.numeric(metnetdf$Count)


## Make the plots of overall counts for a day

levels(metnetdf$Category)<-c("Density","Diameter","Edgewise\nReciprocity","Transitivity","Undirected\nDensity","Undirected\nDiameter")

mets.split<-split(metnetdf,metnetdf$Category)

library(ggplot2)
library(gridExtra)

metlist<-list()

for(i in 1:length(mets.split)){

pic<-ggplot()+geom_boxplot(data=subset(mets.split[[i]],mets.split[[i]]$Data=="sim"),aes(y=Count))+geom_boxplot(data=subset(mets.split[[i]],mets.split[[i]]$Data=="act"),aes(y=Count,colour="red"))+theme_classic()+theme(legend.position="none",axis.title.x = element_text(size = 12, face = "bold"), axis.ticks.x=element_blank(),axis.text.x=element_blank(),axis.text.y=element_text(size=10,face="bold"),axis.title.y=element_text(face="bold"))+labs(x=levels(mets.split[[i]]$Category)[i])

if(i>1){pic<-pic+theme(axis.title.y=element_blank())}

metlist[[i]]<-pic

}

metsplot<-plot_grid(metlist[[1]],metlist[[2]],metlist[[3]],metlist[[4]],metlist[[5]],metlist[[6]],nrow=1,ncol=length(metlist), align="hv",axis="lb")

ggsave(paste0(datfile,"/metsplot_",dates[b]),plot=metsplot,device="pdf",width=9,height=5,units="in")

#save data for later
overallnet[[b]]<-mets.split

metnetdf$Category<-as.character(metnetdf$Category)

overallcounts[[b]]<-rbind(overallcounts[[b]],metnetdf)

overallcounts[[b]]$Date<-rep(dates[b],nrow(overallcounts[[b]]))
	
}	


}


############ FUll-SEQUENCE Visualizations ###############

#use the series of sim "tomorrows" based on actual "yesterdays."


#str(overallcop)

#str(overallfight)

#str(overallnet)


## Distributions by Male ##

####overall copulation dist

#------># this is the sum of the means for each day? Must to propagate that error appropriately. 

#standard deviation for a sum of means is the square root of the sum of squared sds: sqrt(sum(sd_i^2)). I can use 2x this as my error bar 

#so I need to save the mean and sd for number of cops and # of interruptions for each male for each day, and then I can add them up. Then getting the overall rates is simple.


overcopact<-overallcop[[1]][[1]]

for(i in 2:length(overallcop)){
	
	overcopact<-rbind(overcopact,overallcop[[i]][[1]])
	
}

coptots<-ave(overcopact$Freq,overcopact$copdist, FUN=function(x) sum(x,na.rm=T))[1:30]

coptots.df<-data.frame(Male=c(1:30),coptots=coptots)
coptots.df$Male<-factor(coptots.df$Male,levels=as.character(c(1:30)))


overcopsim<-overallcop[[1]][[2]]

for(i in 2:length(overallcop)){
	
	overcopsim<-rbind(overcopsim,overallcop[[i]][[2]])
	
}

coptotsim<-ave(overcopsim$meancops,overcopact$copdist, FUN=function(x) sum(x,na.rm=T))[1:30]

#aggregate the sds appropriately
copsds<-ave(overcopsim$sdcops,overcopact$copdist, FUN=function(x) sqrt(sum(x^2,na.rm=T)))[1:30]

coptotsim.df<-data.frame(Male=c(1:30),coptots=coptotsim,copsds=copsds)
coptotsim.df$Male<-factor(coptotsim.df$Male,levels=as.character(c(1:30)))


#bars in the background to group the boxplots
barback.cop<-cbind(as.character(coptots.df$Male),rep(max(coptots.df$coptots)+2*max(coptotsim.df$copsds)+1,nrow(coptots.df)))
barback.cop<-data.frame(barback.cop)
barback.cop$X1<-factor(barback.cop$X1,levels(coptots.df$Male))
barback.cop$X2<-as.numeric(as.character(barback.cop$X2))

xlabs<-levels(coptotsim.df$Male)

#revert the label for X males so it's clear in the figure
xlabs[grep("30",xlabs)]<-"X"

totalcops.plot<-ggplot()+geom_bar(data=barback.cop,aes(x=X1,y=X2),stat="identity",fill="grey90")+geom_bar(data=coptotsim.df,aes(x=Male,y=coptots),stat="identity")+geom_boxplot(data=coptots.df,aes(x=Male,y=coptots),colour="red")+geom_errorbar(data=coptotsim.df,aes(x=Male,ymin=coptots-2*copsds,ymax=coptots+2*copsds),width=0.6)+theme_classic()+theme(legend.position="none",axis.title.x = element_text(size = 14, face="bold"),axis.text.x=element_text(face="bold"),axis.text.y=element_text(face="bold"),axis.title.y=element_text(face="bold"))+scale_y_continuous(breaks=function(x) pretty(x))+scale_x_discrete(labels=xlabs)+labs(x="Distribution of Total Solicitations by Male ID", y="Total Solicitations")


#+geom_line(data=coptots.df,aes(x=as.numeric(Male),y=coptots),colour="red")

ggsave(paste0("./sim_actualday_plots","/totalcops.plot.pdf"),plot=totalcops.plot,width=9,height=7,units="in")


### overall number of uninterrupted copulations ####


overendact<-overallcop[[1]][[5]]

for(i in 2:length(overallcop)){
	
	overendact<-rbind(overendact,overallcop[[i]][[5]])
	
}

endtots<-ave(overendact$Freq,overendact$enddist, FUN=function(x) sum(x,na.rm=T))[1:30]

endtots.df<-data.frame(Male=c(1:30),endtots=endtots)
endtots.df$Male<-factor(endtots.df$Male,levels=as.character(c(1:30)))


overendsim<-overallcop[[1]][[6]]

for(i in 2:length(overallcop)){
	
	overendsim<-rbind(overendsim,overallcop[[i]][[6]])
	
}

endtotsim<-ave(overendsim$meanends,overendact$enddist, FUN=function(x) sum(x,na.rm=T))[1:30]

#aggregate the sds appropriately
endsds<-ave(overendsim$sdends,overendact$enddist, FUN=function(x) sqrt(sum(x^2,na.rm=T)))[1:30]

endtotsim.df<-data.frame(Male=c(1:30),endtots=endtotsim,endsds=endsds)
endtotsim.df$Male<-factor(endtotsim.df$Male,levels=as.character(c(1:30)))


#bars in the background to group the boxplots

barback.end<-cbind(as.character(endtots.df$Male),rep(max(endtots.df$endtots)+2*max(endtotsim.df$endsds)+1,nrow(endtots.df)))
barback.end<-data.frame(barback.end)
barback.end$X1<-factor(barback.end$X1,levels(endtots.df$Male))
barback.end$X2<-as.numeric(as.character(barback.end$X2))


xlabs<-levels(endtotsim.df$Male)

#revert the label for X males so it's clear in the figure
xlabs[grep("30",xlabs)]<-"X"

totalends.plot<-ggplot()+geom_bar(data=barback.end,aes(x=X1,y=X2),stat="identity",fill="grey90")+geom_bar(data=endtotsim.df,aes(x=Male,y=endtots),stat="identity")+geom_boxplot(data=endtots.df,aes(x=Male,y=endtots),colour="red")+geom_errorbar(data=endtotsim.df,aes(x=Male,ymin=endtots-2*endsds,ymax=endtots+2*endsds),width=0.6)+theme_classic()+theme(legend.position="none",axis.title.x = element_text(size = 14, face="bold"),axis.text.x=element_text(face="bold"),axis.text.y=element_text(face="bold"),axis.title.y=element_text(face="bold"))+scale_y_continuous(breaks=function(x) pretty(x))+labs(x="Distribution of Uninterrupted Copulations by Male ID", y="Total Solicitations")


#+geom_line(data=coptots.df,aes(x=as.numeric(Male),y=coptots),colour="red")

ggsave(paste0("./sim_actualday_plots","/totalends.plot.pdf"),plot=totalends.plot,width=9,height=7,units="in")


###combine plots for total vs. uninterrupted matings

endtotsim.df$success<-rep("Uninterrupted",30)
colnames(endtotsim.df)<-c("Male","tots","sds","success")

coptotsim.df$success<-rep("Solicited",30)
colnames(coptotsim.df)<-c("Male","tots","sds","success")

combocop<-rbind(coptotsim.df,endtotsim.df)


#get the observed data too
endtots.df$success<-rep("Uninterrupted",30)
colnames(endtots.df)<-c("Male","tots","success")

coptots.df$success<-rep("Solicited",30)
colnames(coptots.df)<-c("Male","tots","success")

comboact<-rbind(coptots.df,endtots.df)


#I can do kind of a work-around where I re-order the observed data and set the x-values for the specific columns I want. This way I can get the other color. 

comboact<-comboact[order(comboact$Male),] 

comboact$plotx<-as.numeric(comboact$Male)

#even rows
comboact$plotx[(1:length(comboact$plotx) %% 2)==0]<-comboact$plotx[(1:length(comboact$plotx) %% 2)==0]+0.2125

#odd rows
comboact$plotx[(1:length(comboact$plotx) %% 2)==1]<-comboact$plotx[(1:length(comboact$plotx) %% 2)==1]-0.2125

xlabs<-levels(combocop$Male)

#revert the label for X males so it's clear in the figure
xlabs[grep("30",xlabs)]<-"X"

matebase<-ggplot()+geom_bar(data=barback.cop,aes(x=X1,y=X2),stat="identity",fill="grey90")+geom_bar(data=combocop,aes(x=Male,y=tots,fill=success),stat="identity", position="dodge")+geom_errorbar(data=combocop,aes(x=Male,ymin=tots-2*sds,ymax=tots+2*sds,fill=success),width=0.3,position=position_dodge(0.9))+geom_boxplot(data=comboact,aes(x=Male,y=tots,colour=success),position=position_dodge(0.9))+scale_colour_manual(name="Observed",values=c("red","red"))+scale_fill_manual(name="Simulated",values=c("grey30","grey60"))+scale_x_discrete(labels=xlabs)+theme_classic()+theme(axis.title.x = element_text(size = 14, face = "bold"),axis.text.x=element_text(face="bold"),axis.text.y=element_text(face="bold"),axis.title.y=element_text(size=14,face="bold"),legend.title=element_text(face="bold"),legend.position="right",legend.text=element_text(size=12),legend.background=element_rect(colour="black"))+labs(x="Male ID", y="Copulations")+theme(legend.position=c(0.875,0.75))+guides(colour=F)

#maybe just do the rest in illustrator

ggsave(paste0("./sim_actualday_plots","/matebase.pdf"),plot=matebase,width=9,height=7,units="in")


## restrict to just the main 5 males?

smallcop<-combocop[combocop$Male %in% c(1,2,3,4,7),] 

smallact<-comboact[comboact$Male %in% c(1,2,3,4,7),] 

smallbar<-barback.cop[c(1,2,3,4,7),]

#to make the height of the grey bars consistent with the non-id sim version
smallbar[,2]<-60


matesmall<-ggplot()+geom_bar(data=smallbar,aes(x=X1,y=X2),stat="identity",fill="grey90")+geom_bar(data=smallcop,aes(x=Male,y=tots,fill=success),stat="identity", position="dodge")+ylim(c(-1,60))+geom_errorbar(data=smallcop,aes(x=Male,ymin=tots-2*sds,ymax=tots+2*sds,fill=success),width=0.3,position=position_dodge(0.9),size=1)+geom_boxplot(data=smallact,aes(x=Male,y=tots,colour=success),position=position_dodge(0.9),size=1)+scale_colour_manual(name="Observed",values=c("red","red"))+scale_fill_manual(name="Simulated",values=c("grey30","grey60"))+theme_classic()+theme(axis.title.x = element_text(size = 14, face = "bold"),axis.text.x=element_text(size=18,face="bold"),axis.text.y=element_text(face="bold"),axis.title.y=element_text(size=14,face="bold"),legend.title=element_text(face="bold"),legend.position="right",legend.text=element_text(size=12),legend.background=element_rect(colour="black"))+labs(x="Male ID", y="Copulations")+theme(legend.position=c(0.825,0.865))+guides(colour=F)

ggsave(paste0("./sim_actualday_plots","/matesmall.pdf"),plot=matesmall,width=7,height=7,units="in")


####overall interruption rate distribution


# Take the rate of interruptions for each day and then take the mean across days. So for simulations, the rate will be total# ints / total# cops

#for the actual data

overintact<-overallcop[[1]][[3]]

for(i in 2:length(overallcop)){
	
	overintact<-rbind(overintact,overallcop[[i]][[3]])
	
}

inttots<-ave(overintact$rate,overintact$Male, FUN=function(x) mean(x,na.rm=T))[1:30]

##consider only using days when they actually mate for the sample size in the SE calculation
intSEs<-ave(overintact$rate,overintact$Male, FUN=function(x) sd(x,na.rm=T))[1:30]

inttots.df<-data.frame(Male=c(1:30),ratio=inttots, intSE=intSEs)
inttots.df$Male<-factor(inttots.df$Male,levels=as.character(c(1:30)))

#inttots.df$ratio<-inttots.df$inttots/coptots.df$coptots

#find the number of days that the male actually copulated (has a rate associated with interruptions)
intdays<-apply(table(overintact$rate,overintact$Male),2,function(x) sum(x))

inttots.df$intSE.corr<-inttots.df$intSE/sqrt(intdays)

#inttots.df$intSE.corr<-inttots.df$intSE

#for the sims


#go to the raw data


alldatint<-lapply(sim.all.dat, function(y){

fullcoplist<-lapply(y[[2]], function(x){
	
	x$Events<-factor(x$Events, levels=event.types)
	
	table(x$Events)[endsWith(names(table(x$Events)),"cop.start")]
	
	})

#remove the first element since that is the observed data
fullcoplist<-fullcoplist[-1]

fullcop<-fullcoplist[[1]]

for(i in 2:length(fullcoplist)){
fullcop<-fullcop+fullcoplist[[i]]
}

return(fullcop)

})


## now do the same for cop.ends

alldatend<-lapply(sim.all.dat, function(y){
  
  fullcoplist<-lapply(y[[2]], function(x){
  	
  	x$Events<-factor(x$Events, levels=event.types)
	
	table(x$Events)[endsWith(names(table(x$Events)),"cop.end")]
	
	})
  
  #remove the first eement since that is the observed data
  fullcoplist<-fullcoplist[-1]
  
  
fullcop<-fullcoplist[[1]]

for(i in 2:length(fullcoplist)){
fullcop<-fullcop+fullcoplist[[i]]
}

return(fullcop)
  
})


#Divide to get the rate of successful cops

intratio<-list()

for(i in 1:length(alldatint)){

ratio<-(1-alldatend[[i]]/alldatint[[i]])

intratio[[i]]<-data.frame(ratio)
colnames(intratio[[i]])<-c("Male","ratio")

}

intsim.df<-intratio[[1]]

for(i in 2:length(intratio)){
	
	intsim.df<-rbind(intsim.df,intratio[[i]])
	
}

#get the means and SE's to match the observed data

intsimtots<-ave(intsim.df$ratio,intsim.df$Male, FUN=function(x) mean(x,na.rm=T))[1:30]

#intsimSEs<-ave(intsim.df$ratio,intsim.df$Male, FUN=function(x) sd(x,na.rm=T))[1:30]
intsimSEs<-ave(intsim.df$ratio,intsim.df$Male, FUN=function(x) sd(x,na.rm=T)/sqrt(18))[1:30]

intsim.sum.df<-data.frame(Male=c(1:30),ratio=intsimtots, intSE=intsimSEs)
intsim.sum.df$Male<-factor(intsim.sum.df$Male,levels=as.character(c(1:30)))


# overintsim<-overallcop[[1]][[4]]
# 
# for(i in 2:length(overallcop)){
# 	
# 	overintsim<-rbind(overintsim,overallcop[[i]][[4]])
# 	
# }
# 
# inttotsim<-ave(overintsim$meanints,overintsim$Male, FUN=function(x) sum(x,na.rm=T))[1:30]
# 
# #aggregate the sds appropriately for the sum of the mean interruptions
# intsds<-ave(overintsim$sdints,overintsim$Male, FUN=function(x) sqrt(sum(x^2,na.rm=T)))[1:30]
# 
# intsim.df<-data.frame(Male=c(1:30),inttots=inttotsim,intsds=intsds)
# intsim.df$Male<-factor(intsim.df$Male,levels=as.character(c(1:30)))
# 
# intsim.df$ratio<-intsim.df$inttots/coptotsim.df$coptots
# 
# #propagate the error for the sds of the ratio of means
# intsim.df$sds<-sqrt((intsim.df$intsds/coptotsim.df$coptots)^2+((intsim.df$inttots*coptotsim.df$copsds)/coptotsim.df$coptots^2)^2)

#add a dataset for the observed points to show the spread

intbox<-overintact[,c(1,2)]
colnames(intbox)<-c("Male","ratio")
intbox$Data<-rep("Observed",nrow(intbox))


#bars in the background to group the boxplots
barback<-cbind(as.character(inttots.df$Male),rep(1,nrow(inttots.df)))
barback<-data.frame(barback)
barback$X1<-factor(barback$X1,levels(inttots.df$Male))
barback$X2<-as.numeric(as.character(barback$X2))

xlabs<-levels(intsim.sum.df$Male)

#revert the label for X males so it's clear in the figure
xlabs[grep("30",xlabs)]<-"X"


totalints.plot<-ggplot()+geom_bar(data=barback,aes(x=X1,y=X2),stat="identity",fill="grey90")+geom_bar(data=intsim.sum.df,aes(x=Male,y=ratio,fill="Simulated"),stat="identity")+geom_errorbar(data=intsim.sum.df,aes(x=Male,ymin=ratio-intSE,ymax=ratio+intSE),width=0.3)+theme_classic()+geom_boxplot(data=inttots.df,aes(x=Male,y=ratio,colour="Observed"))+geom_errorbar(data=inttots.df,aes(x=Male,ymin=ratio-intSE.corr,ymax=ratio+intSE.corr),width=0.3, colour="red")+geom_jitter(data=intbox,aes(x=Male,y=ratio,colour="Observed"),height=0,width=0.3)+scale_fill_manual(name="Simulated",values=c("grey40"))+scale_colour_manual(name="Observed",values=c("red"))+theme(legend.title=element_blank(),legend.position=c(0.875,0.75),legend.text=element_text(size=12),legend.background=element_rect(colour="black"),axis.title.x = element_text(size = 14, face="bold"),axis.text.x=element_text(face="bold"),axis.text.y=element_text(face="bold"),axis.title.y=element_text(size=14,face="bold"))+scale_x_discrete(labels=xlabs)+scale_y_continuous(breaks=function(x) pretty(x))+labs(x="Male ID", y="Mean Interruption Rate")+guides(fill=guide_legend(order=1),colour=guide_legend(order=2))


###version that is restricted to just the five main males

smallintsim<-intsim.sum.df[c(1,2,3,4,7),]

smallintact<-inttots.df[c(1,2,3,4,7),]

barback.int.small<-barback[c(1,2,3,4,7),]

intbox.small<-intbox[intbox$Male %in% c(1,2,3,4,7),]


totalints.small<-ggplot()+geom_bar(data=barback.int.small,aes(x=X1,y=X2),stat="identity",fill="grey90")+geom_bar(data=smallintsim,aes(x=Male,y=ratio,fill="Simulated"),stat="identity")+geom_errorbar(data=smallintsim,aes(x=Male,ymin=ratio-intSE,ymax=ratio+intSE),width=0.3,size=1)+theme_classic()+geom_boxplot(data=smallintact,aes(x=Male,y=ratio,colour="Observed"),size=1)+geom_errorbar(data=smallintact,aes(x=Male,ymin=ratio-intSE.corr,ymax=ratio+intSE.corr),width=0.3, colour="red",size=1)+geom_jitter(data=intbox.small,aes(x=Male,y=ratio,colour="Observed"),height=0,width=0.3,size=2.5)+scale_fill_manual(name="Simulated",values=c("grey40"))+scale_colour_manual(name="Observed",values=c("red"))+theme(legend.title=element_blank(),legend.position=c(0.875,0.75),legend.text=element_text(size=12),legend.background=element_rect(colour="black"),axis.title.x = element_text(size = 14, face="bold"),axis.text.x=element_text(size=18, face="bold"),axis.text.y=element_text(face="bold"),axis.title.y=element_text(size=14,face="bold"))+scale_y_continuous(breaks=function(x) pretty(x))+labs(x="Male ID", y="Mean Interruption Rate")+guides(fill=guide_legend(order=1),colour=guide_legend(order=2))+theme(legend.position=c(0.825,0.85))


#add in errorbars for the observed means? just to show the variance, basically 
#+geom_errorbar(data=inttots.df,aes(x=Male,ymin=ratio-intSE.corr,ymax=ratio+intSE.corr),width=0.6, colour="red")


#what about a boxplot approach? This gives a better sense of the data spread and sampling


# # 
##intsim.df$Male<-sapply(intsim.df$Male, function(x) strsplit(as.character(x),".",fixed=T)[[1]][1])
##intsim.df$Male<-factor(intsim.df$Male,levels=as.character(c(1:30)))


# intbox<-overintact[,c(1,2)]
# colnames(intbox)<-c("Male","ratio")
# intbox$Data<-rep("Observed",nrow(intbox))

# intsim.df$Data<-rep("Simulations",nrow(intsim.df))

# intbox<-rbind(intbox,intsim.df)


# totalints.plot<-ggplot()+geom_bar(data=barback,aes(x=X1,y=X2),stat="identity",fill="grey90")+geom_boxplot(data=intbox,aes(x=Male,y=ratio,colour=Data,fill=Data),position=position_dodge(preserve="single"))+scale_colour_manual(values=c("red","black"),labels=c("Observed","Simulations"))+scale_fill_manual(values=c("white","grey"),labels=c("Observed","Simulations"))+theme_classic()+theme(axis.title.x = element_text(size = 14, face = "bold"),axis.text.x=element_text(face="bold"),axis.text.y=element_text(face="bold"),axis.title.y=element_text(size=14,face="bold"),legend.title=element_text(face="bold"),legend.position="right",legend.text=element_text(size=12),legend.background=element_rect(colour="black"))+guides(name="Data Type", labels=c("Observed","Simulation"))+labs(x="Distribution of Interruption Rate by Male ID", y="Interruption Rate",fill="Data Type", colour="Data Type")+theme(legend.position=c(0.875,0.75))


ggsave(paste0("./sim_actualday_plots","/totalints.plot.pdf"),plot=totalints.plot,width=9,height=7,units="in")

ggsave(paste0("./sim_actualday_plots","/totalints.small.pdf"),plot=totalints.small,width=7,height=7,units="in")


#should put the cops and the ints together so you can eyeball the expected success rate

copint_combined<-plot_grid(matebase,totalints.plot,ncol=1,nrow=2,align="hv")


ggsave(paste0("./sim_actualday_plots","/copint_combined.pdf"),plot=copint_combined,width=9,height=7,units="in")


#combined small plots:

copint_small<-plot_grid(matesmall,totalints.small,ncol=1,nrow=2,align="hv")

ggsave(paste0("./sim_actualday_plots","/copint_small.pdf"),plot=copint_small,width=7,height=14,units="in")


#### ave % time fighting for each male: just pool everything (since all my days will have the same number of sims)


fightact<-overallfight[[1]][[1]]

for(i in 2:length(overallfight)){
	
	fightact<-rbind(fightact,overallfight[[i]][[1]])
	
}

fightact$data<-rep("act",nrow(fightact))


fightsim<-overallfight[[1]][[2]]

for(i in 2:length(overallfight)){
	
	fightsim<-rbind(fightsim,overallfight[[i]][[2]])
	
}

fightsim$data<-rep("sim",nrow(fightsim))

#reorder by rank magnitude for percent attendance

library(tidyverse)
library(forcats)

fightact<-fightact %>%
mutate(Male=fct_reorder(Male,desc(Perc.Attend), .fun=function(x) median(x,na.rm=T)))

fightsim$Male<-factor(fightsim$Male,levels(fightact$Male))


fightall<-rbind(fightact,fightsim)

#bars in the background to group the boxplots
barback<-cbind(levels(fightall$Male),rep(100,length(levels(fightall$Male))))
barback<-data.frame(barback)
barback$X1<-factor(barback$X1,levels(fightall$Male))
barback$X2<-as.numeric(as.character(barback$X2))

#draw a line thru them
meds<-ave(fightact$Perc.Fighting,fightact$Male,FUN=function(x) median(x, na.rm=T))[1:29]

meds<-data.frame(medians=meds,Male=c(1:29))
meds$Male<-factor(meds$Male,levels=levels(fightact$Male))

overallfightplot<-ggplot()+geom_bar(data=barback,aes(x=X1,y=X2),stat="identity",fill="grey90")+geom_boxplot(data=fightall,aes(x=Male,y=Perc.Fighting,colour=data,fill=data))+scale_colour_manual(values=c("red","black"),labels=c("Observed","Simulations"))+scale_fill_manual(values=c("white","grey"),labels=c("Observed","Simulations"))+theme_classic()+theme(axis.title.x = element_text(size = 14, face = "bold"),axis.text.x=element_text(face="bold"),axis.text.y=element_text(face="bold"),axis.title.y=element_text(size=14,face="bold"),legend.title=element_text(face="bold"),legend.position="right",legend.text=element_text(size=12),legend.background=element_rect(colour="black"))+guides(name="Data Type", labels=c("Observed","Simulation"))+labs(x="Male ID", y="Percent Time Spent Fighting",fill="Data Type", colour="Data Type")+geom_line(data=meds,aes(x=as.numeric(Male)-(11/60),y=medians),colour="red")


ggsave(paste0("./sim_actualday_plots","/overallfightplot.pdf"),plot=overallfightplot,width=11,height=7,units="in")


#### ave % of total time in attendance

#draw a line thru them
meds.att<-ave(fightact$Perc.Attend,fightact$Male,FUN=function(x) median(x, na.rm=T))[1:29]

meds.att<-data.frame(medians=meds.att,Male=c(1:29))
meds.att$Male<-factor(meds.att$Male,levels=levels(fightact$Male))

overallattendplot<-ggplot()+geom_bar(data=barback,aes(x=X1,y=X2),stat="identity",fill="grey90")+geom_boxplot(data=fightall,aes(x=Male,y=Perc.Attend,colour=data,fill=data))+scale_colour_manual(values=c("red","black"),labels=c("Observed","Simulations"))+scale_fill_manual(values=c("white","grey"),labels=c("Observed","Simulations"))+theme_classic()+theme(axis.title.x = element_text(size = 14, face = "bold"),axis.text.x=element_text(face="bold"),axis.text.y=element_text(face="bold"),axis.title.y=element_text(size=14,face="bold"),legend.title=element_text(face="bold"),legend.position="right",legend.text=element_text(size=12),legend.background=element_rect(colour="black"))+guides(name="Data Type", labels=c("Observed","Simulation"))+labs(x="Male ID", y="Percent of Total Time in Attendance",fill="Data Type", colour="Data Type")+geom_line(data=meds.att,aes(x=as.numeric(Male)-(11/60),y=medians),colour="red")


attendfight_combined<-plot_grid(overallattendplot,overallfightplot,nrow=2,ncol=1,align="hv")


ggsave(paste0("./sim_actualday_plots","/attendfight_combined.pdf"),plot=attendfight_combined,width=13,height=7,units="in")


## Overall Average Counts ##


#str(overallnet)


#average density

density<-overallnet[[1]][[1]]

for(i in 2:length(overallnet)){
	
	density<-rbind(density,overallnet[[i]][[1]])
	
}

levels(density$Data)<-c("Observed","Simulations")

density.plot<-ggplot()+geom_boxplot(data=density,aes(x=Data,y=Count,colour=Data,fill=Data))+scale_colour_manual(values=c("red","black"))+scale_fill_manual(values=c("white","grey80"))+theme_classic()+theme(axis.title.x = element_text(size = 14, face = "bold"),axis.text.x=element_text(face="bold"),axis.text.y=element_text(face="bold"),axis.title.y=element_text(size=14,face="bold"),legend.position="none")+labs(x="Density", y="Value")


#average undir.density


undir<-overallnet[[1]][[5]]

for(i in 2:length(overallnet)){
	
	undir<-rbind(undir,overallnet[[i]][[5]])
	
}

levels(undir$Data)<-c("Observed","Simulations")

undir.plot<-ggplot()+geom_boxplot(data=undir,aes(x=Data,y=Count,colour=Data,fill=Data))+scale_colour_manual(values=c("red","black"))+scale_fill_manual(values=c("white","grey80"))+theme_classic()+theme(axis.title.x = element_text(size = 14, face = "bold"),axis.text.x=element_text(face="bold"),axis.text.y=element_text(face="bold"),axis.title.y=element_blank(),legend.position="none")+labs(x="Undirected\nDensity", y="Value")


#average transitivity

trans<-overallnet[[1]][[4]]

for(i in 2:length(overallnet)){
	
	trans<-rbind(trans,overallnet[[i]][[4]])
	
}

levels(trans$Data)<-c("Observed","Simulations")

trans.plot<-ggplot()+geom_boxplot(data=trans,aes(x=Data,y=Count,colour=Data,fill=Data))+scale_colour_manual(values=c("red","black"))+scale_fill_manual(values=c("white","grey80"))+theme_classic()+theme(axis.title.x = element_text(size = 14, face = "bold"),axis.text.x=element_text(face="bold"),axis.text.y=element_text(face="bold"),axis.title.y=element_blank(),legend.position="none")+labs(x="Transitivity", y="Value")


#average reciprocity


recip<-overallnet[[1]][[3]]

for(i in 2:length(overallnet)){
	
	recip<-rbind(recip,overallnet[[i]][[3]])
	
}

levels(recip$Data)<-c("Observed","Simulations")

recip.plot<-ggplot()+geom_boxplot(data=recip,aes(x=Data,y=Count,colour=Data,fill=Data))+scale_colour_manual(values=c("red","black"))+scale_fill_manual(values=c("white","grey80"))+theme_classic()+theme(axis.title.x = element_text(size = 14, face = "bold"),axis.text.x=element_text(face="bold"),axis.text.y=element_text(face="bold"),axis.title.y=element_blank(),legend.position="none")+labs(x="Edgewise\nReciprocity", y="Value")


overallnet_plot<-plot_grid(density.plot,undir.plot,trans.plot,recip.plot,nrow=1,ncol=4,align="hv")

ggsave(paste0("./sim_actualday_plots/","overallnet_plot.pdf"),plot=overallnet_plot,width=10,height=5,units="in")


#### Look at counts by Day rather than global averages to compare

overallcounts.plot<-overallcounts[[1]]

for(i in 2:length(overallcounts)){
  
  overallcounts.plot<-rbind(overallcounts.plot,overallcounts[[i]])
  
}

overallcounts.plot$Date<-as.factor(overallcounts.plot$Date)
overallcounts.plot$Category<-as.factor(overallcounts.plot$Category)
overallcounts.plot$Category<-factor(overallcounts.plot$Category,levels=levels(overallcounts.plot$Category)[c(6,7,1,2,3,9,5,8,4,10)])

levels(overallcounts.plot$Data)<-c("Observed","Simulations")

overallcounts.plot.split<-split(overallcounts.plot,overallcounts.plot$Category)

countbacks<-list()

for(i in 1:length(overallcounts.plot.split)){

#bars in the background to group the boxplots
barback<-cbind(levels(overallcounts.plot.split[[i]]$Date),rep(ceiling(1.05*max(overallcounts.plot.split[[i]]$Count,na.rm=T)),length(levels(overallcounts.plot.split[[i]]$Date))))
barback<-data.frame(barback)
barback$X1<-factor(barback$X1,levels(overallcounts.plot.split[[i]]$Date))
barback$X2<-as.numeric(as.character(barback$X2))

countbacks[[i]]<-barback

}


plotlist<-list()

for(i in 1:length(overallcounts.plot.split)){
  
  pic<-ggplot()+geom_boxplot(data=subset(overallcounts.plot.split[[i]],overallcounts.plot.split[[i]]$Data=="sim"),aes(x=Date,y=Count))+geom_boxplot(data=subset(overallcounts.plot.split[[i]],overallcounts.plot.split[[i]]$Data=="act"),aes(x=Date,y=Count,colour="red"))+theme_classic()+theme(legend.position="none",axis.title.x = element_text(size = 14, face = "bold"),axis.text.x=element_text(size=5,face="bold"),axis.text.y=element_text(face="bold"),axis.title.y=element_text(face="bold"))+labs(x=levels(overallcounts.plot.split[[i]]$Category)[i])+geom_bar(data=countbacks[[i]],aes(x=X1,y=X2),stat="identity",fill="grey90")
  
  pic$layers<-pic$layers[c(3,1,2)]
  
  #if(i<length(overallcounts.plot.split)){pic<-pic+theme(axis.title.x=element_blank())}
  
  plotlist[[i]]<-pic
  
}

countsplot<-plot_grid(plotlist[[1]],plotlist[[2]],plotlist[[3]],plotlist[[4]],plotlist[[5]],plotlist[[6]],plotlist[[7]],plotlist[[8]],plotlist[[9]],plotlist[[10]],nrow=length(plotlist)/2,ncol=2,align="hv",axis="lb")

ggsave(paste0("./sim_actualday_plots/","overallcount_plot.pdf"),plot=countsplot,width=24,height=11,units="in")
```
