## Supplemental_reproducible_code for "Fighting isn’t sexy in lekking Greater Sage-grouse (*Centrocercus urophasianus*)": funlist_formatting.R

```
rm(list=ls())

###make sure necessary packages are there and loaded

packages<-c("here")

package.check<-lapply(packages, FUN = function(x) {
    if (!require(x, character.only = TRUE, quietly=TRUE)) {
        install.packages(x, dependencies = TRUE)
        library(x, character.only = TRUE, quietly=TRUE)
    } 

})

i_am("other_supporting_materials/funlist_formatting.R")

#load in the user-defined functions that add predictor stats
source(here("rem_pre_analysis","rem.functions.R"), chdir = TRUE)

##make sure no extraneous functions are floating around in R's brain, that will really mess things up
fun.list<-lsf.str()[grep("add.",lsf.str())]

#strip the "add." of the fronts of the function names

stripped<-sapply(fun.list, function(x) gsub("^[^.]*.","",x))

#add sf.dis to the list 
stripped<-c(stripped,"sf.dis")

names(stripped)<-NULL

setwd(here("other_supporting_materials"))
write.table(stripped, file="funlist_text",sep=",",row.names=F)
```
