## Supplemental_reproducible_code for "Fighting isn’t sexy in lekking Greater Sage-grouse (*Centrocercus urophasianus*)": supp_corplot_fight_by_mating.R

```
#first, make sure R's memory is clean
rm(list=ls())

###make sure necessary packages are there and loaded

packages<-c("here","abind","relevent","lubridate","DescTools","gtools","ggplot2")

package.check<-lapply(packages, FUN = function(x) {
 if (!require(x, character.only = TRUE, quietly=TRUE)) {
 install.packages(x, dependencies = TRUE)
 library(x, character.only = TRUE, quietly=TRUE)
 } 

})


#for project path management
#define the project root directory relative to the current script. This sets the working directory that gets called by the here() function. It is a relative path and will be robust to use on anyone's computer. In this case, I want the root directory to always be the enclosing folder for the whole sage grouse project
i_am("other_supporting_materials/supp_corplot_fight_by_mating.R")

###Build the intercept list, support array, event list, census array, etc

{
	
#make the base event list and intercept array

event.list<-read.table(file=here("data_processing","relevent_format_dataset_udata_wincol_disturbance.tsv"), header=T,  sep="\t")

event.list$Date<-as.factor(event.list$Date)  

event.types<-read.table(file=here("data_processing","event_types_list_udata.csv"), header=F,  sep=",")
event.types<-as.character(event.types$V1)


#NOTE- the event.types vector now contains ENDOGENOUS event types as well as EXOGENOUS types: arrivals and clock events, and end.obs. Need to make sure everything is robust to this. 


### create eventlist for rem ###


event.list.dat<-split(event.list, event.list$Date)


#remove female context for now

basic.event.list<-event.list[-3]

#strip the "loser" from the disengage events so they are all the same event "type" and they match my previous code

basic.event.list$Events<-sapply(basic.event.list$Events, function(x) {
	
	event<-unlist(strsplit(as.character(x),".",fixed=T))
	if(event[2]=="disengage"){
		return(paste(event[1],event[2],sep="."))
	} else {return(as.character(x))}
}
)


#add factor levels for event types that don't actually happen (and put them in the same order as the event.type list) to maintain bookkeeping 

basic.event.list$Events<-as.character(basic.event.list$Events)

basic.event.list$Events<-factor(basic.event.list$Events, levels=as.character(event.types))

#turn event types into numeric values

basic.event.list$Events<-as.numeric(basic.event.list$Events)

#set the event type code for exogenous events to zero. I have there all at the trailing end of the vector

endogenous<-(length(event.types)-length(event.types[grep("arrive",event.types)])-5)

basic.event.list$Events[basic.event.list$Events>endogenous]<-0

rownames(basic.event.list)<-NULL
#need to divide the data up by grouping factor

evlist<-list()
evlist<-split(basic.event.list, basic.event.list$Date)

#strip the date column from the sub-dataframes so they're just two columns, event and time

evlist<-lapply(evlist, function(x) x<-x[-3])

#convert from dataframes to matrices

evlist<-lapply(evlist, function(x) x<-as.matrix(x))

## make event.types vector usable for intlist, etc. by excising exogenous event types

event.types<-event.types[1:endogenous]

}


#get a data frame with each row is a male

#count attacks and count wins


evtab<-table(event.list$Events)

copcounts<-evtab[grep("cop.start",names(evtab))]

attacks<-evtab[grep("attack",names(evtab))]

attack.vec<-vector()

for(i in 1:29){

attack.vec[i]<-sum(attacks[startsWith(names(attacks),paste0(i,"."))])

}

cop.vec<-vector()


for(i in 1:29){

cop.vec[i]<-sum(copcounts[startsWith(names(copcounts),paste0(i,"."))])

}

example<-data.frame(male=as.character(1:29),attack=attack.vec,cop=cop.vec)


#poisson regression

lintest<-summary(glm(cop~attack,data=example,family="poisson"))


# # Call:
# glm(formula = cop ~ attack, family = "poisson", data = example)

# Deviance Residuals: 
    # Min       1Q   Median       3Q      Max  
# -5.9021  -0.3688  -0.2870  -0.2768   5.3809  

# Coefficients:
             # Estimate Std. Error z value Pr(>|z|)    
# (Intercept) -3.261740   0.701374  -4.650 3.31e-06 ***
# attack       0.023903   0.002702   8.846  < 2e-16 ***
# ---
# Signif. codes:  
# 0 ‘***’ 0.001 ‘**’ 0.01 ‘*’ 0.05 ‘.’ 0.1 ‘ ’ 1

# (Dispersion parameter for poisson family taken to be 1)

    # Null deviance: 361.021  on 28  degrees of freedom
# Residual deviance:  80.927  on 27  degrees of freedom
# AIC: 99.123

# Number of Fisher Scoring iterations: 5


#actually, apparently log-log is better
loglintest<-summary(glm(log(1+cop)~log(1+attack),data=example))


# Call:
# glm(formula = log(1 + cop) ~ log(1 + attack), data = example)

# Deviance Residuals: 
     # Min        1Q    Median        3Q       Max  
# -1.15753  -0.41625  -0.07138   0.29034   2.67010  

# Coefficients:
                # Estimate Std. Error t value Pr(>|t|)
# (Intercept)     -0.29034    0.23379  -1.242  0.22497
# log(1 + attack)  0.26092    0.07808   3.342  0.00245
                  
# (Intercept)       
# log(1 + attack) **
# ---
# Signif. codes:  
# 0 ‘***’ 0.001 ‘**’ 0.01 ‘*’ 0.05 ‘.’ 0.1 ‘ ’ 1

# (Dispersion parameter for gaussian family taken to be 0.6625775)

    # Null deviance: 25.289  on 28  degrees of freedom
# Residual deviance: 17.890  on 27  degrees of freedom
# AIC: 74.289

# Number of Fisher Scoring iterations: 2

#can I get an R-squared?

lmsum<-summary(lm(log(1+cop)~log(1+attack),data=example))


# Call:
# lm(formula = log(1 + cop) ~ log(1 + attack), data = example)

# Residuals:
     # Min       1Q   Median       3Q      Max 
# -1.15753 -0.41625 -0.07138  0.29034  2.67010 

# Coefficients:
                # Estimate Std. Error t value Pr(>|t|)   
# (Intercept)     -0.29034    0.23379  -1.242  0.22497   
# log(1 + attack)  0.26092    0.07808   3.342  0.00245 **
# ---
# Signif. codes:  0 ‘***’ 0.001 ‘**’ 0.01 ‘*’ 0.05 ‘.’ 0.1 ‘ ’ 1

# Residual standard error: 0.814 on 27 degrees of freedom
# Multiple R-squared:  0.2926,	Adjusted R-squared:  0.2664 
# F-statistic: 11.17 on 1 and 27 DF,  p-value: 0.002447


library(ggplot2)

basic<-ggplot(example)+geom_point(aes(x=log(1+attack),y=log(1+cop)),shape=21,size=5,fill="grey40",stroke=1,alpha=0.7)+geom_abline(intercept=loglintest$coeff[1],slope=loglintest$coeff[2])

full<-basic+theme_classic()+labs(x="Log Number of Fights Initiated", y="Log Number of Solicitations")+theme(axis.title.x=element_text(size=16,face="bold"),axis.text.x=element_text(size=14,face="bold"),axis.title.y=element_text(size=16,face="bold"),axis.text.y=element_text(size=14,face="bold"))+geom_text(aes(x=3,y=3,label=paste0("R^2=",round(lmsum$adj.r.squared,4))))+geom_text(aes(x=3,y=2.7,label=paste0("p=",round(lmsum$coefficients[2,4],4),"**")))

setwd(here("other_supporting_materials"))

ggsave("supp_corplot_fight_by_mating.pdf",full,height=5,width=5) 
```
