## Supplemental_reproducible_code for "Fighting isn’t sexy in lekking Greater Sage-grouse (*Centrocercus urophasianus*)": README_sage_grouse_REM.docx

Steps to reproduce sage grouse REM results

1. Data Processing
   1. Run the script: **full_interaction_data_for_REM_import_code_udata_supp.R**
      - This reproduces the consolidated data file: **full_data_plus_cop_quest.tsv**
   2. Run the script: **rem_udata_centroid_data.R**
      - This reproduces the folder: **udata_centroid_data** and its subfolders
        - contains spatial data for the positions of the sage grouse: centroids of movement for each male on each day, and the pairwise distance between centroids for all males on each day
   3. Run the script: **relevent_formatting_dis_reverse_udata_wincolumn_disturbance.**R
      - This reproduces the formatted data file: **relevent_format_dataset_udata_wincol_disturbance.tsv** as well as the file **event_types_list_udata.csv**
        - The data file is the interaction network data formatted for use in the REM analysis. The event types list is a list of all the unique types of events that can happen in the scheme of the REM
2. REM pre-analysis
   1. Script **relevent_analysis_intercepts_udata.R** shows the process by which we established the base intercept-only model for the REM. Optimized models all use this intercept model as its baseline
   2. The two text files contain the final intercept-only model output for the aggression and attractiveness models, respectively.
   3. Script **rem.functions.R** contains code for each of the functions that can add a particular sufficient statistic to the REM model. The final models are composed of the basic intercepts and optimized subsets of these statistics
3. Cluster Optimization: These folders contain all the materials and scripts used for running our model selection algorithm on a high-performance computing cluster from a single directory, subsequent analysis, and the creation of summary figures
   1. Aggression Model
      - CHG_2014_male_presence_data_quest.csv, CHG_females_exact.tsv, event_types_list_udata.csv, male_pres_exact_quest.csv, relevent_format_dataset_udata_wincol_disturbance.tsv, rem.functions.R, and pairwise_dist are all data files identical to their counterparts in the data_files and data_processing folders
      - **run_pipeline_stepwise.sh** and **run_pipeline_stepwise_restart.sh** are shell scripts that orchestrate the parallel runs of the REM model on the cluster. The former is for starting fresh, and the latter is for restarting in the middle of a run
      - **genremlist*** scripts generate the sets of joblists that get sent to the cluster. **genremlist_next*** is triggered by the shell script after each round of model selection; it assesses which combination of statistics so far has performed the best and generates new sets to test. **genremlist_restart*** creates the next joblist after a run on the cluster is paused and needs to be restarted at the point where it left off.
      - The script **rem_cluster_model_selection_stepwise_udata_terr.R** encodes a single run of the REM model on the cluster for a given set of sufficient statistics.
      - Cluster run outcomes:
        - **previous_best_initial_run** shows the set of statistics from the initial run of model selection, including some non-significant statistics
        - **previous_best_re-run_without_226_283** shows the result from removing the non-significant statistics from the pool and re-running the algorithm from the beginning
        - **previous_best_remove_226_283_at_the_end** shows the result from removing the non-significant statistics from the final results of the initial run, and then restarting the algorithm from that point. It converges immediately and with the best BIC; this represents the final statistic set reflected in the final aggression model results, and this file is the one referred to in the analysis script.

- The script **best_result_full_model_forward_udata_terrsol_analysis.R** goes through the final model selection process and the best-fit statistic set result in detail and produces the figures contained in the **summary_figs** folder, as well as the output files **rem_model_coef** and **rem_model_sd**, giving the values for the coefficients for the best-fitting model and the standard deviations for the coefficients, respectively.
- For reference, **aggression_model_output_full.txt** contains the full model output for the best-fitting aggression model
2. Attractiveness Model
   - The data files, shell scripts, and “genremlist” scripts, are equivalent to their counterparts found in the aggression model folder
   - The script **rem_cluster_model_selection_stepwise_udata_terr_id.R** encodes a single run of the REM model on the cluster for a given set of sufficient statistics, using the base intercept model that accounts for individual-level attractiveness effects.
   - Cluster run outcomes:
     - **previous_best_remove_226_at_the_end** shows the set of statistics from the initial run of model selection in the second-to-last row, and the result of removing the non-significant statistic and then restarting the algorithm from that point in the last row. The algorithm converges immediately, and so the result is the same as what one gets by simply removing the single non-significant predictor. This is the final set of statistics reflected in the final attractiveness model results, and this file is the one referred to in the analysis script.
     - **previous_best_re_run_without_226** shows the result from removing the non-significant statistic from the pool and re-running the algorithm from the beginning.
   - Analogous to the script in the Aggression Model folder, the script **best_result_full_model_forward_udata_terrsol_id_analysis.R** goes through the final model selection process and the best-fit statistic set result in detail and produces the figures contained in the **summary_figs** folder, as well as the output files **rem_model_coef** and **rem_model_sd**, giving the values for the coefficients for the best-fitting model and the standard deviations for the coefficients, respectively.
     - The plot **allplot_terr_id_model_grid** corresponds to the final **Figure 4** in the main text
     - The plot **femplot** corresponds to the final **Figure 5** in the main text
   - For reference, **attractiveness_model_output_full.txt** contains the full model output for the best-fitting attractiveness model

4. Adequacy Checking: These folders contain the scripts (and associated products) used for examining the adequacy of the optimized REM models for describing the observed data.
   1. Aggression Model
      - Run the script **relevent_analysis_base_udata_terr_adequacy.R** to reproduce the basic adequacy analysis as well as:
        - CSV files **per_event_adequacy_data_full** and **per_event_adequacy_data_summary**; these contain the lists of hazards the REM model calculates for each event in the observed dataset and some summary statistics for the calculated hazards, respectively.
        - Folder **main plots**; contains summary plots describing Aggression Model adequacy, including event matching, event rank histograms, recall, and deviance residuals.
   2. Attractiveness Model

- Run the script **relevent_analysis_base_udata_terr_adequacy.R** to reproduce the basic adequacy analysis as well as:
  - CSV files **per_event_adequacy_data_full** and **per_event_adequacy_data_summary**; as above, these contain the lists of hazards the REM model calculates for each event in the observed dataset and some summary statistics for the calculated hazards, respectively.
  - Folder **main plots**; contains summary plots describing Aggression Model adequacy, including event matching, event rank histograms, recall, and deviance residuals.
    - **matches.pdf** corresponds to **Figure S3**
    - **rank_hist_cat.pdf** corresponds to **Figure S4**
    - **recall.plot.pdf** corresponds to **Figure S5**
    - **devmeanplot.pdf** corresponds to **Figure S6**
    - **devtimeplot.pdf** corresponds to **Figure S7**
- Run the script **availability_interruption_plots_terr_id_new.R** to reproduce:
  - **matingplot_new_statlevs_newlabs.pdf** which corresponds to **Figure 7** in the main text
  - summary plot **fight_count_fig.pdf**
- Run the script **fight_effect_explore.R** to reproduce:
  - **fight_effect_fig.pdf** which corresponds to **Figure 6** in the main text
- Run the script **time_fighting.R** to reproduce:
  - **time_fighting_fig.pdf** which corresponds to **Figure S2**

5. Event History Simulation: These folders contain all the materials and scripts used for producing simulated event history datasets based on the best-fit REM models on a high-performance computing cluster from a single directory, subsequent analysis, and the creation of summary figures.
   1. Aggression Model
      - CHG_2014_disturbance_events.csv, CHG_2014_male_presence_data_quest.csv, CHG_females_exact.tsv, event_types_list_udata.csv, male_pres_exact_quest.csv, relevent_format_dataset_udata_wincol_disturbance.tsv, and pairwise_dist are all data files identical to their counterparts in the data_files and data_processing folders
      - **rem.functions.act.terr.sim.R** contains code for each of the functions that can add a particular sufficient statistic to the REM model. They correspond to the functions in the rem.functions.R file in the rem_pre_analysis folder, but in this case they are coded in order to be robust to a growing, simulated event history, rather than an existing observed dataset.
      - **previous_best_remove_226_283_at_the_end and rem_model_coef** are identical to the files representing the final statistic set for the best-fitting Aggression Model results, and the corresponding coefficients, respectively, found in cluster_optimization 🡪 Aggression_model.
      - **run_pipeline_sim_act.sh** is a shell script that organizes the parallel simulations on the cluster
      - **genremlist_sim.R** script is called by the shell script and generates the joblist that gets sent to the cluster
      - **rem_event_hist_simulation_cluster_udata_terr_actualday_newcensus.R** encodes a single simulation run, in this case, based on the best-fit Aggression REM.
      - The folder **sim_results_actualday** contains all the simulated data produced, in this case based on the best-fit Aggression Model: 100 simulations for each of the 18 days in the empirical dataset.
      - The script **sim_output_analysis_cluster_new_3.R** brings together the simulation data and produces summary analyses and figures, contained in the folder **sim_actualday_plots**:
        - Each **day_level*** subfolder contains sets of plots summarizing the simulations for each simulated day in the dataset
        - Other plots summarize the characteristics of the simulated “season” of data overall, and compares them to the observed data, including overall counts of different types of events and overall network characteristics of the event histories
          - **matesmall.pdf** corresponds to **Figure 3, panel A** in the main text
   2. Attractiveness Model
      - The data files, shell script, “genremlist” script, and function list are equivalent to their counterparts found in event_history_simulation 🡪 Aggression_model
      - **previous_best_remove_226_at_the_end and rem_model_coef** are identical to the files representing the final statistic set for the best-fitting Attractiveness Model results, and the corresponding coefficients, respectively, found in cluster_optimization 🡪 Attractiveness_model.
      - **rem_event_hist_simulation_cluster_udata_terr_id_actualday_newcensus.R** encodes a single simulation run, in this case, based on the best-fit Attractiveness REM.
      - The folder **sim_results_actualday** contains all the simulated data produced, in this case based on the best-fit Attractiveness Model: 100 simulations for each of the 18 days in the empirical dataset.
      - As above, the script **sim_output_analysis_cluster_new_3.R** brings together the simulation data and produces summary analyses and figures, contained in the folder **sim_actualday_plots**:
        - As above, each **day_level*** subfolder contains sets of plots summarizing the simulations for each simulated day in the dataset
          - **nodemetsplot_2014-03-28.pdf** corresponds to **Figure S8**
        - Other plots summarize the characteristics of the simulated “season” of data overall, and compares them to the observed data, including overall counts of different types of events and overall network characteristics of the event histories
          - **overallcount_plot.pdf** corresponds to **Figure S8**
          - **attendfight_combined.pdf** corresponds to **Figure S9**
          - **overallnet_plot.pdf** corresponds to **Figure S11**
          - **matesmall.pdf** corresponds to **Figure 3, panel B** in the main text
6. Other supporting materials
   1. Run script **funlist_formatting.R** to reproduce **funlist_text**, which is the basis for **Table S1**, the full annotated function list.
   2. Run script **supp_corplot_fight_by_mating.R**:
      - Reproduces **supp_corplot_fight_by_mating.pdf** which corresponds to **Figure S1**
