## Supplemental_reproducible_code for "Fighting isn’t sexy in lekking Greater Sage-grouse (*Centrocercus urophasianus*)": relevent_analysis_intercepts_udata.R

```
#This code goes through our process of selecting the basic intercept model (analogous to choosing fixed effects) for the REM analysis. The model deemed most appropriate is used in the subsequent analysis.


#########################################

#first, make sure R's memory is clean
rm(list=ls())

###### Reformat data for use in the relevent package ######

###make sure necessary packages are there and loaded

packages<-c("here","abind","relevent","tictoc")

package.check<-lapply(packages, FUN = function(x) {
  if (!require(x, character.only = TRUE, quietly=TRUE)) {
    install.packages(x, dependencies = TRUE)
    library(x, character.only = TRUE, quietly=TRUE)
  } 
  
})

#for project path management
#define the project root directory relative to the current script. This sets the working directory that gets called by the here() function. It is a relative path and will be robust to use on anyone's computer. In this case, I want the root directory to always be the enclosing folder for the whole sage grouse project
i_am("rem_pre_analysis/relevent_analysis_intercepts_udata.R")


#make the base event list and intercept array


event.list<-read.table(file=here("data_processing","relevent_format_dataset_udata_wincol_disturbance.tsv"), header=T,  sep="\t")

event.list$Date<-as.factor(event.list$Date)  
  
event.types<-read.table(file=here("data_processing","event_types_list_udata.csv"), header=F,  sep=",")
event.types<-as.character(event.types$V1)


#NOTE- the event.types vector now contains ENDOGENOUS event types as well as EXOGENOUS types: arrivals and clock events, and end.obs. Need to make sure everything is robust to this. 

### create eventlist for rem ###


event.list.dat<-split(event.list, event.list$Date)


#remove female context column for now

basic.event.list<-event.list[-3]

#strip the "loser" from the disengage events so they are all the same event "type" and they match my previous code

basic.event.list$Events<-sapply(basic.event.list$Events, function(x) {
  
  event<-unlist(strsplit(as.character(x),".",fixed=T))
  if(event[2]=="disengage"){
    return(paste(event[1],event[2],sep="."))
  } else {return(as.character(x))}
}
)


#add factor levels for event types that don't actually happen (and put them in the same order as the event.type list) to maintain bookkeeping 

basic.event.list$Events<-as.character(basic.event.list$Events)

basic.event.list$Events<-factor(basic.event.list$Events, levels=as.character(event.types))

#turn event types into numeric values

basic.event.list$Events<-as.numeric(basic.event.list$Events)

#set the event type code for exogenous events to zero. They are all at the trailing end of the vector

endogenous<-(length(event.types)-length(event.types[grep("arrive",event.types)])-5)

basic.event.list$Events[basic.event.list$Events>endogenous]<-0

rownames(basic.event.list)<-NULL
#need to divide the data up by grouping factor

evlist<-list()
evlist<-split(basic.event.list, basic.event.list$Date)

#strip the date column from the sub-dataframes so they're just two columns, event and time

evlist<-lapply(evlist, function(x) x<-x[-3])

#convert from dataframes to matrices

evlist<-lapply(evlist, function(x) x<-as.matrix(x))

## make event.types vector usable for intlist, etc. by excising exogenous event types

event.types<-event.types[1:endogenous]


#### create statlist ####

##"a three dimensional array, event number by event type by statistic, or a list thereof, containing the sufficient statistics for the model"

#start with the intercept sufficient stats


sequences<-table(event.list$Date)

#create list of dummy intercept names. Leave male 30.receives as the baseline- this is the receive rate for X males- expect everyone else's receive rate to be less

idno<-length(event.types[grep("cop.start",event.types)])

dummy.ints.attack<-paste(1:idno,".attackFE", sep="")
dummy.ints.receive<-paste(1:(idno-1), ".receiveFE", sep="")


int.names<-c(dummy.ints.attack,dummy.ints.receive,event.types[-grep("attack", event.types)])

int.list<-list()

for(i in 1:length(levels(event.list$Date))){

day.array<-array(0,dim=c(as.numeric(sequences[i]), length(event.types),length(int.names)))

#the third dimension is the stat slice.
dimnames(day.array)<-list(c(1:sequences[i]),as.character(event.types),as.character(int.names))

#loop for dummy intercepts for attacks, loop for dummy intercepts for receives, then one loop for typical intercepts

for(k in 1:length(dummy.ints.attack)){
	
day.array[,startsWith(names(day.array[1,,k]), paste(k,".attack", sep="")),k]<-1

}

for(k in (length(dummy.ints.attack)+1):(length(dummy.ints.attack)+length(dummy.ints.receive))){
	
day.array[,endsWith(names(day.array[1,,k]), paste("attack.",(k-idno), sep="")),k]<-1

}

for(k in (length(dummy.ints.attack)+length(dummy.ints.receive)+1):length(int.names)){
	
	day.array[,int.names[k],k]<-1
	
}

#weird extra formatting to make it match the package example, seems like it becomes relevant *punintended!* if I want to fit day-specific effects

global<-list()

global[[1]]<-day.array

int.list[[i]]<-global

names(int.list[[i]])<-"global"

}


### give it a shot without the support matrix ###

##NOTE- this fails to converge (at least within ~3 hours) in the udata case- this is'nt surprising since there are tons of event types that only happen on particular days or not at all, and the model would have a hard time estimate a rate for them. 

#library(relevent)

#fit.rem<-rem(evlist, int.list, timing="interval", estimator="BPM")
#summary(fit.rem)


#### Create Support Matrix ####

#"should be a list withone element per history, each of which should be an event order by event type logical matrix. the ijth cell of this matrix should be TRUE if an event type of j was a possible next event given the preceding code i-1 events, and FALSE otherwise"

#need a way to ask, "what is each male doing right now" for each event in the sequence. That dictates the possibilities. 

#create a bookkeeping "census" matrix to keep track

census.list<-list()


for(j in 1:length(event.list.dat)){
  
  census<-matrix("nothing", nrow=1,ncol=(idno), dimnames=list(1,c(1:(idno))))
  
  
  ##code that translates each new current event into what the males involved are doing at that time. The support matrix will be based on whatever the males were doing in the PREVIOUS timestep. This census matrix itself could be interesting to visualize. Each row of the census says what each male is doing going into the NEXT event.
  
  
  for(i in 1:nrow(event.list.dat[[j]])){
    
    #can ask if the second thing is something, and then move on to the first and third elements.
    
    current<-census[nrow(census),]
    
    #parse the event type. loop through test.seq rows
    evname<-as.character(event.list.dat[[j]][i,1])
    evsplit<-strsplit(evname, split=".", fixed=T)
    
    #if it is an end obs, arrival, departure or clock event, it shouldn't change anything the males are doing in terms of interactions. Departures aren't exogenous, but they shouldn't change anything since they can only happen once a male is already doing nothing. 
    if(evsplit[[1]][2]!="obs" & evsplit[[1]][2]!="depart" & evsplit[[1]][2]!="arrive" & evsplit[[1]][2]!="time" & evsplit[[1]][1]!="fem"){
      #if the event was an attack, indicate the males involved and which was the receiver since this dictates who makes the initition decision
      if(evsplit[[1]][2]=="attack"){
        current[evsplit[[1]][1]]<-evname
        current[evsplit[[1]][3]]<-paste(evsplit[[1]][3], ".", "received", ".", evsplit[[1]][1], sep="")
      }
      #if it's an FO or SF, just indicate that that pair of males are involved in a fight now
      if(evsplit[[1]][2]=="initFO"||evsplit[[1]][2]=="initSF"){
        #need to refer back to the previous row of "census" to get the id of the other male
        
        current[evsplit[[1]][1]]<-paste(evsplit[[1]][1], ".", "fight", ".",strsplit(census[nrow(census),evsplit[[1]][1]], split=".",fixed=T)[[1]][3], sep="")
        current[[strsplit(census[nrow(census),evsplit[[1]][1]], split=".",fixed=T)[[1]][3]]]<-paste(strsplit(census[nrow(census),evsplit[[1]][1]], split=".",fixed=T)[[1]][3], ".", "fight", ".",evsplit[[1]][1], sep="")
        
      }
      
      #if it's a chase, indicate that the pair of males are involved in a fight, but there is an additional constraint that the individual DOING THE CHASING will be the disengager by definition
      if(evsplit[[1]][2]=="initCH"){
        
        current[evsplit[[1]][1]]<-paste(evsplit[[1]][1], ".", "fight", ".",strsplit(census[nrow(census),evsplit[[1]][1]], split=".",fixed=T)[[1]][3], ".chasedby",sep="")
        current[[strsplit(census[nrow(census),evsplit[[1]][1]], split=".",fixed=T)[[1]][3]]]<-paste(strsplit(census[nrow(census),evsplit[[1]][1]], split=".",fixed=T)[[1]][3], ".", "fight", ".",evsplit[[1]][1], ".chasing",sep="")
        
      }
      
      #for disengaging, end the fight for the partner male as well
      if(evsplit[[1]][2]=="disengage"){
        current[evsplit[[1]][1]]<-"nothing"
        current[evsplit[[1]][3]]<-"nothing"	
        
      }
      
      
      #for copulations, end cops are only possible after start cops, but otherwise anything else is possible. EXCEPT the copulating male can't initiate an attack. If the next event involving that male ISN'T an end cop, it was interrupted
      
      if(evsplit[[1]][2]=="cop" && evsplit[[1]][3]=="start"){
        current[evsplit[[1]][1]]<-"copulating"
      }
      
      if(evsplit[[1]][2]=="cop" && evsplit[[1]][3]=="end"){
        current[evsplit[[1]][1]]<-"nothing"
      }
      
      #if an X male is still involved in a fight at all, make sure he is still tagged as fighting (since sometimes more than one is interacting at once, but they can't depart until none are fighting)
      if((TRUE %in% grepl("fight.30",current) | TRUE %in% grepl("30.fight",current)) && !(grepl("receive", current[idno]) | grepl("attack", current[idno]))) {
        
        current[idno]<-"fight"
        
      }
      
    }
    
    census<-rbind(census,current)
    
    
  }
  
  
  #don't want the first dummy row
  census.list[[j]]<-census[-1,]
  
}


## Constraints to incorporate in the support matrix ##

#only males that are actually present can be involved in interactions on a given day and at a given time

#If "nothing" is happening, all "attacks", all start cops and all departs (given the male is actually present) are possible. Males cannot disengage or initiate fights without an attack happening first. End cop cannot happen without a previous start cop. end.obs is possible. (end.obs is not possible if any males are still fighting or copulating, so end.obs is ONLY possible when "nothing" is going on across the board)

#if the previous event was an "attack", the current event MUST be an initiation by the receiver, since we are assuming the attack is essentually instantaneous. 

#once initiatiated, subsequent events can only be a disengage from either male involved or any other event NOT invoving the interaction partners. Males actively fighting connot become engaged in a new fight until they disengage. 

#if the initiated fight was a chase only the male being chased can disengage, by definition

#Once a copulation starts, the only thing that male can do next is either end the copulation OR receive an attack. He cannot initiate anything. 

#copulations cannot occur when there are no females present

#a male can ONLY "depart" if "nothing" is happening

#Exceptions to these rules are for "X" males- there may be more than one unidentifiable male on the lek at a time. Right now we are dealing with this by having "X" males just be a mass actor category. 1) These males CAN do an "attack" before disengaging from an interaction. They can also be attacked. However, normal restrictions apply to the partner male. So if an X male is being chased, "disengage" becomes possible, but any "attack" (or cop.start, or depart) are still also possible 


### Build the actual support matrix ###

#for each event, there is a list of event types, with 1's for the types that are possible and 0's for types that are not. 


#code block to produce the support matrix

supplist<-list()


#bring in day-level male presence data

male.pres<-read.table(here("data_files","CHG_2014_male_presence_data_quest.csv"), sep=",", header=T)

#male IDs are already in ascending order, so I can convert them directly
colnames(male.pres)<-c("Date", 1:idno)

#bring in exact male presence data

pres.exact<-read.table(here("data_files","male_pres_exact_quest.csv"), sep=",",header=T)

departlist<-list()

for(j in 1:nrow(pres.exact)){
  
  arriverow<-pres.exact[j,startsWith(colnames(pres.exact),"arrive")]
  arriverow<-arriverow[!is.na(arriverow)]
  
  arrives<-t(sapply(arriverow, function(x) return(c(paste0(pres.exact$Num_ID[j],".arrive"),x,as.character(pres.exact$Date[j])))))
  
  departrow<-pres.exact[j,startsWith(colnames(pres.exact),"depart")]
  departrow<-departrow[!is.na(departrow)]
  
  departs<-t(sapply(departrow, function(x) return(c(paste0(pres.exact$Num_ID[j],".depart"),x,as.character(pres.exact$Date[j])))))
  
  allpres<-rbind(arrives,departs)
  
  departlist[[j]]<-allpres
  
}

#collapse the departlist 

departevent<-abind(departlist, along=1)

departevent<-data.frame(departevent)
names(departevent)<-c("Events","Time", "Date")

departevent$Time<-as.numeric(as.character(departevent$Time))

#library(abind)
arrstr<-abind(strsplit(as.character(departevent$Events),".",fixed=T),along=2)

departevent$Bird_ID<-arrstr[1,]
departevent$Type<-arrstr[2,]

departevent$Type<-as.factor(departevent$Type)
departevent$Bird_ID<-as.factor(departevent$Bird_ID)

# #split into day-specific data frames
pres.exact.dat<-split(departevent, departevent$Date)


#make a reference event list with data (most importantly female presence)
evlist.dat<-event.list.dat


for(j in 1:length(evlist)){
  
  supplist[[j]]<-matrix(1, nrow=length(evlist[[j]][,1]), ncol=length(event.types))
  colnames(supplist[[j]])<-event.types
  
  #turn off all interactions involving males not present on that day
  #make a list of all the event types that include the males that were entirely absent and remove them for all rows
  
  supplist[[j]][,unique(unlist(lapply(grep(0,male.pres[j,2:(idno+1)]), function(x) c(event.types[startsWith(event.types, paste(x, ".", sep=""))], event.types[endsWith(event.types, paste(".", x, sep=""))]))))]<-0
  
  
  #first row can be any "attack" or, technically, any "depart"
  #since now fem.enter is an actual event, cop.start is never possible as the very first thing
  supplist[[j]][1,][-c(grep("attack", event.types),grep("depart", event.types))]<-0
  
  
  #the exception is events involving males that will be present that day but haven't arrived yet
  
  
  #basically I can make this code exactly the same as below- just want to ask who is actually there right now? 
  
  arrsub<-subset(pres.exact.dat[[j]],Time<evlist[[j]][1,2] | Time==0 )
  arrtab<-table(arrsub$Bird_ID,arrsub$Type)
  
  arrpat<-apply(arrtab,1,function(x) sum(x))
  
  #if the total number of previous arrives and departs is even, that male is currently absent
  absent<-names(arrpat[(arrpat %% 2)==0])
  
  if(length(absent)>0){
    supplist[[j]][1,][unique(c(sapply(absent, function(x) c(event.types[startsWith(event.types, paste(x, ".", sep=""))], event.types[endsWith(event.types, paste(".", x, sep=""))]))))]<-0
    
  } 
  
  
  
  #loop through the rest of the events: 
  
  for(i in 2:(length(evlist[[j]][,1])-1)){
    
    
    #before anything else turn off events involving males that have either not yet arrived or have lef the lek early and are therefore no longer present that day or are otherwise not currently present
    
    
    arrsub<-subset(pres.exact.dat[[j]],Time<evlist[[j]][i,2])
    arrtab<-table(arrsub$Bird_ID,arrsub$Type)
    
    arrpat<-apply(arrtab,1,function(x) sum(x))
    
    #if the total number of previous arrives and departs is even, that male is currently absent
    current.row.abs<-names(arrpat[(arrpat %% 2)==0])
    
    
    if(length(current.row.abs)>0){
      supplist[[j]][i,][unique(c(sapply(current.row.abs, function(x) c(event.types[startsWith(event.types, paste(x, ".", sep=""))], event.types[endsWith(event.types, paste(".", x, sep=""))]))))]<-0
      
    } 
    
    
    
    #determine if the previous event was an attack, since this dictates exactly what the next event can be: receiver initiates a type of fight
    if((TRUE %in% grepl("received", census.list[[j]][i-1,]))==TRUE){
      
      supplist[[j]][i,][-grep(TRUE, startsWith(event.types, paste(grep("received", census.list[[j]][i-1,]), ".", "init", sep="")))]<-0}
    
    
    #(else) if the previous event wasn't an attack, the next event CAN'T be an initiation and the possibilities depend on what everyone is currently engaged in
    
    
    if((TRUE %in% grepl("received", census.list[[j]][i-1,]))==FALSE){
      
      supplist[[j]][i,][grep("init", event.types)]<-0
      
      #if males are actively fighting, they can't attack or be attacked, or actually do anything else before disengaging (unless they are a ?? male, id 30)
      
      if((TRUE %in% grepl("fight", census.list[[j]][i-1,]))==TRUE){
        
        for(k in grep("fight", census.list[[j]][i-1,])){
          
          if(k!=idno){
            
            supplist[[j]][i,][c(event.types[startsWith(event.types, paste(k, ".attack.", sep=""))], event.types[endsWith(event.types, paste("attack.", k, sep=""))], event.types[startsWith(event.types, paste(k, ".cop", sep=""))])]<-0
            
          }
          
        }
        
        #ONLY those fighting can disengage. retain males that are fighting and remove the rest
        dis<-event.types[grep("disengage",event.types)]
        supplist[[j]][i,][dis[-grep("fight", census.list[[j]][i-1,])]]<-0
        
        #"chasing" and "chasedby" limit the possibilities further. Males getting chased DO NOT disengage by definition, again, with the exception of ?? males. This is taken care of by having ?? males just be "fight" -ing without any extra tags
        if(TRUE %in% grepl("chasedby", census.list[[j]][i-1,])){
          supplist[[j]][i,][dis[grep("chasedby", census.list[[j]][i-1,])]]<-0
        }
        
      }
      
      #if no one is fighting, no one can disengage, but all attack combinations are possible
      if((TRUE %in% grepl("fight", census.list[[j]][i-1,]))==FALSE){
        
        supplist[[j]][i,][grep("disengage",event.types)]<-0
        
      }
      
      
      #actively copulating males: They cannot cop.start. ONLY they can cop.end. they CAN be attacked but they CANNOT attack anyone
      if((TRUE %in% grepl("copulating", census.list[[j]][i-1,]))==TRUE){
        
        cop<-grep("copulating", census.list[[j]][i-1,]) 
        
        #stuff that needs to be turned off for only copulating males
        for(h in cop){
          
          supplist[[j]][i,][c(event.types[startsWith(event.types ,paste(h, ".attack.", sep=""))], event.types[startsWith(event.types, paste(h, ".cop.start", sep=""))])]<-0
          
        }
        
        #stuff that needs to be on for ONLY copulating males
        copendall<-event.types[grep("cop.end",event.types)]
        supplist[[j]][i,][copendall[-cop]]<-0
      }
      
      #if no one is copulating, all cop.starts are possible, but cop.ends are not
      
      if((TRUE %in% grepl("copulating", census.list[[j]][i-1,]))==FALSE){
        
        copendall<-event.types[grep("cop.end",event.types)]
        supplist[[j]][i,][copendall]<-0
        
      }
      
      
    }
    
    #if there are no females present, the next event could not have possibly been a cop.start
    
    if(evlist.dat[[j]][i-1,]$Females==1){
      
      copstartall<-event.types[grep("cop.start",event.types)]
      supplist[[j]][i,][copstartall]<-0
      
    }
    
    #Finally, A male can only "depart" if he was already doing "nothing," so depart must be turned off for everyone doing not-nothing. (also he has to be present, but I've already taken care of that above)
    
    noth<-as.character(which(census.list[[j]][i-1,]!="nothing"))
    
    if(length(noth)>0){
      
      supplist[[j]][i,][paste(noth, ".depart", sep="")]<-0
      
    }
    
  }
  #end events loop here
  
  #after done with all the middle events: onset of end.obs could depend on a lot of factors; not sure I want to fit it as part of the model. Just always say the last row must be end.obs (all endogenous events are no longer possible) 
  
  supplist[[j]][length(evlist[[j]][,1]),]<-0
  
}
#end day loop here


##### Run Rem with intercept statlist + supplort list #####

#library(relevent)
#library(tictoc)

tic()
intfit.sup<-rem(evlist, int.list, supplist=supplist, timing="interval", estimator="BPM")
toc()

options(max.print=2000)
summary(intfit.sup)
#looks good! the model performs MUCH better since it eliminates the possibility of impossible events. Also this runs super fast compared to before.

#results here
{
# Egocentric Relational Event Model (Interval Likelihood)
# 
#               Post.Mode    Post.SD  Z value  Pr(>|z|)    
# 1.attackFE    -5.691936   0.072902 -78.0767 < 2.2e-16 ***
# 2.attackFE    -6.061305   0.086983 -69.6834 < 2.2e-16 ***
# 3.attackFE    -5.594034   0.073058 -76.5696 < 2.2e-16 ***
# 4.attackFE    -5.139261   0.069087 -74.3877 < 2.2e-16 ***
# 5.attackFE    -6.076087   0.104436 -58.1798 < 2.2e-16 ***
# 6.attackFE    -7.745222   0.707845 -10.9420 < 2.2e-16 ***
# 7.attackFE    -5.999958   0.089411 -67.1055 < 2.2e-16 ***
# 8.attackFE    -5.851973   0.199312 -29.3609 < 2.2e-16 ***
# 9.attackFE    -9.216134   1.000670  -9.2100 < 2.2e-16 ***
# 10.attackFE   -5.303609   0.091194 -58.1574 < 2.2e-16 ***
# 11.attackFE  -18.232062 203.963285  -0.0894 0.9287728    
# 12.attackFE   -7.307190   0.355752 -20.5402 < 2.2e-16 ***
# 13.attackFE   -6.465665   1.001167  -6.4581 1.060e-10 ***
# 14.attackFE   -5.795849   0.207437 -27.9402 < 2.2e-16 ***
# 15.attackFE   -7.160362   0.579494 -12.3562 < 2.2e-16 ***
# 16.attackFE  -15.243124 221.934731  -0.0687 0.9452420    
# 17.attackFE   -5.841598   0.709511  -8.2333 2.220e-16 ***
# 18.attackFE  -17.267794 209.276230  -0.0825 0.9342396    
# 19.attackFE   -5.614105   0.278309 -20.1722 < 2.2e-16 ***
# 20.attackFE   -7.546369   1.001074  -7.5383 4.752e-14 ***
# 21.attackFE  -16.792494 212.052811  -0.0792 0.9368814    
# 22.attackFE   -6.760439   0.583388 -11.5882 < 2.2e-16 ***
# 23.attackFE  -17.770218 206.456883  -0.0861 0.9314089    
# 24.attackFE  -17.999513 205.207489  -0.0877 0.9301042    
# 25.attackFE   -4.825562   0.193550 -24.9318 < 2.2e-16 ***
# 26.attackFE   -5.326821   0.514602 -10.3513 < 2.2e-16 ***
# 27.attackFE   -6.942945   0.502202 -13.8250 < 2.2e-16 ***
# 28.attackFE   -5.401014   0.274847 -19.6510 < 2.2e-16 ***
# 29.attackFE   -4.877860   0.414753 -11.7609 < 2.2e-16 ***
# 30.attackFE   -7.274863   0.294002 -24.7442 < 2.2e-16 ***
# 1.receiveFE   -3.035067   0.125627 -24.1593 < 2.2e-16 ***
# 2.receiveFE   -2.649488   0.104687 -25.3087 < 2.2e-16 ***
# 3.receiveFE   -2.755423   0.114850 -23.9914 < 2.2e-16 ***
# 4.receiveFE   -1.507390   0.081842 -18.4182 < 2.2e-16 ***
# 5.receiveFE   -3.097196   0.154354 -20.0656 < 2.2e-16 ***
# 6.receiveFE   -1.641032   0.234168  -7.0079 2.419e-12 ***
# 7.receiveFE   -2.129663   0.089759 -23.7265 < 2.2e-16 ***
# 8.receiveFE   -2.042537   0.209122  -9.7672 < 2.2e-16 ***
# 9.receiveFE   -1.719290   0.177471  -9.6877 < 2.2e-16 ***
# 10.receiveFE  -2.322298   0.154550 -15.0262 < 2.2e-16 ***
# 11.receiveFE  -0.881359   0.381251  -2.3118 0.0207913 *  
# 12.receiveFE  -2.854943   0.262473 -10.8771 < 2.2e-16 ***
# 13.receiveFE  -2.780729   1.001152  -2.7775 0.0054774 ** 
# 14.receiveFE  -2.258105   0.246840  -9.1481 < 2.2e-16 ***
# 15.receiveFE  -2.408252   0.336498  -7.1568 8.258e-13 ***
# 16.receiveFE   1.642929   0.356611   4.6071 4.084e-06 ***
# 17.receiveFE  -2.537486   0.708617  -3.5809 0.0003424 ***
# 18.receiveFE  -2.182725   1.001104  -2.1803 0.0292340 *  
# 19.receiveFE  -2.030680   0.300046  -6.7679 1.307e-11 ***
# 20.receiveFE  -0.421994   0.195102  -2.1629 0.0305457 *  
# 21.receiveFE  -0.610787   1.001075  -0.6101 0.5417752    
# 22.receiveFE -14.616240 226.344456  -0.0646 0.9485122    
# 23.receiveFE  -3.165828   1.002293  -3.1586 0.0015854 ** 
# 24.receiveFE  -0.535976   0.246548  -2.1739 0.0297111 *  
# 25.receiveFE  -3.100881   0.709013  -4.3735 1.223e-05 ***
# 26.receiveFE  -2.595370   0.714355  -3.6332 0.0002800 ***
# 27.receiveFE  -1.537261   0.229662  -6.6936 2.178e-11 ***
# 28.receiveFE  -2.934238   0.579966  -5.0593 4.207e-07 ***
# 29.receiveFE  -2.283943   1.002518  -2.2782 0.0227142 *  
# 1.initFO       6.744430   0.127000  53.1057 < 2.2e-16 ***
# 2.initFO       6.700116   0.104828  63.9150 < 2.2e-16 ***
# 3.initFO       6.738679   0.114708  58.7464 < 2.2e-16 ***
# 4.initFO       6.684612   0.075378  88.6814 < 2.2e-16 ***
# 5.initFO       6.792686   0.156174  43.4944 < 2.2e-16 ***
# 6.initFO       5.061926   0.577351   8.7675 < 2.2e-16 ***
# 7.initFO       6.715748   0.085436  78.6058 < 2.2e-16 ***
# 8.initFO       5.115994   0.500000  10.2320 < 2.2e-16 ***
# 9.initFO       3.381391   1.000001   3.3814 0.0007212 ***
# 10.initFO      6.662633   0.166667  39.9758 < 2.2e-16 ***
# 11.initFO     -6.724740 321.814902  -0.0209 0.9833284    
# 12.initFO      5.809142   0.447214  12.9896 < 2.2e-16 ***
# 13.initFO     -5.062715 363.256388  -0.0139 0.9888802    
# 14.initFO      5.460835   0.500000  10.9217 < 2.2e-16 ***
# 15.initFO      6.096823   0.500000  12.1936 < 2.2e-16 ***
# 16.initFO     -6.841114 319.417929  -0.0214 0.9829127    
# 17.initFO     -5.646696 346.931976  -0.0163 0.9870141    
# 18.initFO     -5.062715 363.256388  -0.0139 0.9888802    
# 19.initFO      4.422843   1.000002   4.4228 9.741e-06 ***
# 20.initFO      3.506554   1.000002   3.5065 0.0004540 ***
# 21.initFO     -5.062715 363.256388  -0.0139 0.9888802    
# 22.initFO      0.000000 894.427191   0.0000 1.0000000    
# 23.initFO     -5.062715 363.256388  -0.0139 0.9888802    
# 24.initFO     -7.553047 305.835850  -0.0247 0.9802971    
# 25.initFO      6.214600   1.000003   6.2146 5.146e-10 ***
# 26.initFO      6.907751   0.707108   9.7690 < 2.2e-16 ***
# 27.initFO      6.060457   0.333333  18.1814 < 2.2e-16 ***
# 28.initFO      6.502286   0.707108   9.1956 < 2.2e-16 ***
# 29.initFO      6.907747   1.000004   6.9077 4.925e-12 ***
# 30.initFO      3.352407   0.258199  12.9838 < 2.2e-16 ***
# 1.initCH       2.617293   1.000001   2.6173 0.0088631 ** 
# 2.initCH      -9.185505 280.258756  -0.0328 0.9738539    
# 3.initCH      -8.988496 283.008974  -0.0318 0.9746631    
# 4.initCH       1.514126   1.000000   1.5141 0.1299940    
# 5.initCH       3.079110   1.000001   3.0791 0.0020762 ** 
# 6.initCH       6.735904   0.250000  26.9436 < 2.2e-16 ***
# 7.initCH      -9.541022 275.492334  -0.0346 0.9723727    
# 8.initCH       6.562914   0.242536  27.0596 < 2.2e-16 ***
# 9.initCH       6.877902   0.174078  39.5105 < 2.2e-16 ***
# 10.initCH      4.465407   0.500000   8.9308 < 2.2e-16 ***
# 11.initCH      6.753603   0.408249  16.5429 < 2.2e-16 ***
# 12.initCH     -7.392244 308.751880  -0.0239 0.9808986    
# 13.initCH     -5.062715 363.256388  -0.0139 0.9888802    
# 14.initCH      6.271766   0.333333  18.8153 < 2.2e-16 ***
# 15.initCH      5.403675   0.707108   7.6419 2.132e-14 ***
# 16.initCH      6.907754   0.353554  19.5381 < 2.2e-16 ***
# 17.initCH      6.214600   1.000003   6.2146 5.146e-10 ***
# 18.initCH      6.907747   1.000004   6.9077 4.925e-12 ***
# 19.initCH      5.809141   0.500000  11.6183 < 2.2e-16 ***
# 20.initCH      6.597600   0.213201  30.9455 < 2.2e-16 ***
# 21.initCH      6.907747   1.000004   6.9077 4.925e-12 ***
# 22.initCH      0.000000 894.427191   0.0000 1.0000000    
# 23.initCH     -5.062715 363.256388  -0.0139 0.9888802    
# 24.initCH      6.850596   0.242536  28.2457 < 2.2e-16 ***
# 25.initCH     -5.646696 346.931976  -0.0163 0.9870141    
# 26.initCH     -5.646696 346.931976  -0.0163 0.9870141    
# 27.initCH      6.060457   0.333333  18.1814 < 2.2e-16 ***
# 28.initCH     -5.992692 338.240255  -0.0177 0.9858644    
# 29.initCH     -5.062715 363.256388  -0.0139 0.9888802    
# 30.initCH      6.824374   0.045502 149.9810 < 2.2e-16 ***
# 1.initSF       4.919880   0.316228  15.5580 < 2.2e-16 ***
# 2.initSF       5.233779   0.218218  23.9842 < 2.2e-16 ***
# 3.initSF       5.047002   0.267261  18.8842 < 2.2e-16 ***
# 4.initSF       5.275328   0.152499  34.5926 < 2.2e-16 ***
# 5.initSF       4.465407   0.500000   8.9308 < 2.2e-16 ***
# 6.initSF      -7.600810 304.985488  -0.0249 0.9801173    
# 7.initSF       5.163063   0.185695  27.8039 < 2.2e-16 ***
# 8.initSF       4.828312   0.577351   8.3629 < 2.2e-16 ***
# 9.initSF      -8.117025 296.225732  -0.0274 0.9781395    
# 10.initSF      4.870872   0.408248  11.9311 < 2.2e-16 ***
# 11.initSF      4.961839   1.000002   4.9618 6.983e-07 ***
# 12.initSF      6.502289   0.316228  20.5620 < 2.2e-16 ***
# 13.initSF      6.907747   1.000004   6.9077 4.925e-12 ***
# 14.initSF      5.460835   0.500000  10.9217 < 2.2e-16 ***
# 15.initSF      5.809141   0.577351  10.0617 < 2.2e-16 ***
# 16.initSF     -6.841114 319.417929  -0.0214 0.9829127    
# 17.initSF      6.214600   1.000003   6.2146 5.146e-10 ***
# 18.initSF     -5.062715 363.256388  -0.0139 0.9888802    
# 19.initSF      6.368758   0.377965  16.8501 < 2.2e-16 ***
# 20.initSF      5.452467   0.377965  14.4259 < 2.2e-16 ***
# 21.initSF     -5.062715 363.256388  -0.0139 0.9888802    
# 22.initSF      0.000000 894.427191   0.0000 1.0000000    
# 23.initSF      6.907747   1.000004   6.9077 4.925e-12 ***
# 24.initSF      4.017378   1.000002   4.0174 5.885e-05 ***
# 25.initSF      6.214600   1.000003   6.2146 5.146e-10 ***
# 26.initSF     -5.646696 346.931976  -0.0163 0.9870141    
# 27.initSF      4.961843   0.577351   8.5942 < 2.2e-16 ***
# 28.initSF      5.809136   1.000003   5.8091 6.280e-09 ***
# 29.initSF     -5.062715 363.256388  -0.0139 0.9888802    
# 30.initSF      3.940194   0.192450  20.4738 < 2.2e-16 ***
# 1.disengage   -4.262284   0.062622 -68.0632 < 2.2e-16 ***
# 2.disengage   -5.148770   0.088736 -58.0237 < 2.2e-16 ***
# 3.disengage   -4.775014   0.063628 -75.0452 < 2.2e-16 ***
# 4.disengage   -4.647453   0.054882 -84.6806 < 2.2e-16 ***
# 5.disengage   -4.573716   0.102598 -44.5791 < 2.2e-16 ***
# 6.disengage   -6.477786   0.577349 -11.2199 < 2.2e-16 ***
# 7.disengage   -5.683178   0.089443 -63.5399 < 2.2e-16 ***
# 8.disengage   -3.025632   0.188982 -16.0101 < 2.2e-16 ***
# 9.disengage   -2.951359   0.999998  -2.9514 0.0031637 ** 
# 10.disengage  -4.610931   0.088045 -52.3701 < 2.2e-16 ***
# 11.disengage  -1.251904   0.999999  -1.2519 0.2106042    
# 12.disengage  -5.099954   0.333333 -15.2999 < 2.2e-16 ***
# 13.disengage  -3.069490   0.999997  -3.0695 0.0021442 ** 
# 14.disengage  -3.999558   0.200000 -19.9978 < 2.2e-16 ***
# 15.disengage  -3.711562   0.377964  -9.8199 < 2.2e-16 ***
# 16.disengage   0.000000 894.427191   0.0000 1.0000000    
# 17.disengage  -5.585480   0.707105  -7.8991 2.887e-15 ***
# 18.disengage   0.000000 894.427191   0.0000 1.0000000    
# 19.disengage  -3.306133   0.250000 -13.2245 < 2.2e-16 ***
# 20.disengage  -2.450509   0.353553  -6.9311 4.176e-12 ***
# 21.disengage   0.000000 894.427191   0.0000 1.0000000    
# 22.disengage  -3.514373   0.999997  -3.5144 0.0004408 ***
# 23.disengage -12.189754 246.283852  -0.0495 0.9605250    
# 24.disengage  -1.891904   0.999998  -1.8919 0.0585033 .  
# 25.disengage  -2.434066   0.200000 -12.1703 < 2.2e-16 ***
# 26.disengage  -3.804916   0.408248  -9.3201 < 2.2e-16 ***
# 27.disengage  -5.368294   0.316228 -16.9760 < 2.2e-16 ***
# 28.disengage  -5.092138   0.353553 -14.4027 < 2.2e-16 ***
# 29.disengage  -5.097787   0.408248 -12.4870 < 2.2e-16 ***
# 30.disengage  -4.831031   0.162221 -29.7805 < 2.2e-16 ***
# 1.cop.start  -21.775230 187.429984  -0.1162 0.9075115    
# 2.cop.start   -9.446606   0.408248 -23.1394 < 2.2e-16 ***
# 3.cop.start   -7.431142   0.149071 -49.8496 < 2.2e-16 ***
# 4.cop.start   -7.839479   0.218218 -35.9250 < 2.2e-16 ***
# 5.cop.start  -21.334710 189.269729  -0.1127 0.9102516    
# 6.cop.start  -19.273179 198.658073  -0.0970 0.9227130    
# 7.cop.start  -21.663933 187.889841  -0.1153 0.9082063    
# 8.cop.start  -19.504105 197.536450  -0.0987 0.9213473    
# 9.cop.start  -19.225865 198.890250  -0.0967 0.9229919    
# 10.cop.start -20.397433 193.369880  -0.1055 0.9159917    
# 11.cop.start -16.297433 215.065251  -0.0758 0.9395949    
# 12.cop.start -19.261927 198.713216  -0.0969 0.9227794    
# 13.cop.start -17.124683 210.100682  -0.0815 0.9350387    
# 14.cop.start -19.045919 199.780762  -0.0953 0.9240495    
# 15.cop.start -19.059755 199.711865  -0.0954 0.9239683    
# 16.cop.start -15.410565 220.799296  -0.0698 0.9443573    
# 17.cop.start -17.846074 206.041010  -0.0866 0.9309782    
# 18.cop.start -16.894762 211.445896  -0.0799 0.9363159    
# 19.cop.start -19.197853 199.028089  -0.0965 0.9231568    
# 20.cop.start -18.175836 204.262033  -0.0890 0.9290955    
# 21.cop.start -15.559441 219.804656  -0.0708 0.9435668    
# 22.cop.start -18.414696 203.001851  -0.0907 0.9277215    
# 23.cop.start -17.543193 207.716617  -0.0845 0.9326928    
# 24.cop.start -18.088681 204.727722  -0.0884 0.9295947    
# 25.cop.start -18.516001 202.474390  -0.0914 0.9271361    
# 26.cop.start -17.327815 208.933327  -0.0829 0.9339035    
# 27.cop.start -18.831473 200.857796  -0.0938 0.9253036    
# 28.cop.start -18.717694 201.436365  -0.0929 0.9259662    
# 29.cop.start   0.000000 894.427191   0.0000 1.0000000    
# 30.cop.start -20.543324 192.714163  -0.1066 0.9151063    
# 1.cop.end      0.000000 894.427191   0.0000 1.0000000    
# 2.cop.end     -1.651538   0.707106  -2.3356 0.0195106 *  
# 3.cop.end     -1.354380   0.160128  -8.4581 < 2.2e-16 ***
# 4.cop.end     -1.737753   0.333333  -5.2133 1.856e-07 ***
# 5.cop.end      0.000000 894.427191   0.0000 1.0000000    
# 6.cop.end      0.000000 894.427191   0.0000 1.0000000    
# 7.cop.end      0.000000 894.427191   0.0000 1.0000000    
# 8.cop.end      0.000000 894.427191   0.0000 1.0000000    
# 9.cop.end      0.000000 894.427191   0.0000 1.0000000    
# 10.cop.end     0.000000 894.427191   0.0000 1.0000000    
# 11.cop.end     0.000000 894.427191   0.0000 1.0000000    
# 12.cop.end     0.000000 894.427191   0.0000 1.0000000    
# 13.cop.end     0.000000 894.427191   0.0000 1.0000000    
# 14.cop.end     0.000000 894.427191   0.0000 1.0000000    
# 15.cop.end     0.000000 894.427191   0.0000 1.0000000    
# 16.cop.end     0.000000 894.427191   0.0000 1.0000000    
# 17.cop.end     0.000000 894.427191   0.0000 1.0000000    
# 18.cop.end     0.000000 894.427191   0.0000 1.0000000    
# 19.cop.end     0.000000 894.427191   0.0000 1.0000000    
# 20.cop.end     0.000000 894.427191   0.0000 1.0000000    
# 21.cop.end     0.000000 894.427191   0.0000 1.0000000    
# 22.cop.end     0.000000 894.427191   0.0000 1.0000000    
# 23.cop.end     0.000000 894.427191   0.0000 1.0000000    
# 24.cop.end     0.000000 894.427191   0.0000 1.0000000    
# 25.cop.end     0.000000 894.427191   0.0000 1.0000000    
# 26.cop.end     0.000000 894.427191   0.0000 1.0000000    
# 27.cop.end     0.000000 894.427191   0.0000 1.0000000    
# 28.cop.end     0.000000 894.427191   0.0000 1.0000000    
# 29.cop.end     0.000000 894.427191   0.0000 1.0000000    
# 30.cop.end     0.000000 894.427191   0.0000 1.0000000    
# 1.depart      -8.701475   0.235702 -36.9172 < 2.2e-16 ***
# 2.depart      -8.645592   0.235702 -36.6802 < 2.2e-16 ***
# 3.depart      -8.603133   0.235702 -36.5000 < 2.2e-16 ***
# 4.depart      -8.213108   0.235702 -34.8453 < 2.2e-16 ***
# 5.depart      -8.797853   0.288675 -30.4767 < 2.2e-16 ***
# 6.depart      -8.107312   0.707105 -11.4655 < 2.2e-16 ***
# 7.depart      -8.466583   0.235702 -35.9207 < 2.2e-16 ***
# 8.depart      -7.624461   0.499999 -15.2489 < 2.2e-16 ***
# 9.depart      -8.213118   0.577349 -14.2256 < 2.2e-16 ***
# 10.depart     -8.336652   0.377964 -22.0567 < 2.2e-16 ***
# 11.depart     -6.136106   0.577349 -10.6281 < 2.2e-16 ***
# 12.depart     -8.430566   0.577349 -14.6022 < 2.2e-16 ***
# 13.depart     -6.374262   0.999995  -6.3743 1.838e-10 ***
# 14.depart     -8.432862   0.707105 -11.9259 < 2.2e-16 ***
# 15.depart     -8.472487   0.999994  -8.4725 < 2.2e-16 ***
# 16.depart     -4.553187   0.999997  -4.5532 5.284e-06 ***
# 17.depart     -7.332594   0.999995  -7.3326 2.256e-13 ***
# 18.depart     -6.129319   0.999996  -6.1293 8.824e-10 ***
# 19.depart     -8.897214   0.999994  -8.8973 < 2.2e-16 ***
# 20.depart     -7.676151   0.999995  -7.6762 1.643e-14 ***
# 21.depart     -4.711675   0.999996  -4.7117 2.457e-06 ***
# 22.depart     -7.735384   0.999995  -7.7354 1.021e-14 ***
# 23.depart     -7.017221   0.999995  -7.0173 2.263e-12 ***
# 24.depart     -7.391510   0.999995  -7.3915 1.452e-13 ***
# 25.depart     -7.842174   0.999994  -7.8422 4.441e-15 ***
# 26.depart     -8.035618   0.999994  -8.0357 8.882e-16 ***
# 27.depart     -8.659685   0.999994  -8.6597 < 2.2e-16 ***
# 28.depart     -8.191807   0.999994  -8.1919 2.220e-16 ***
# 29.depart     -7.054816   0.999995  -7.0549 1.728e-12 ***
# 30.depart     -5.737913   0.117851 -48.6878 < 2.2e-16 ***
# ---
# Signif. codes:  0 ‘***’ 0.001 ‘**’ 0.01 ‘*’ 0.05 ‘.’ 0.1 ‘ ’ 1
# Null deviance: 76628.21 on 4854 degrees of freedom
# Residual deviance: 29745.98 on 4585 degrees of freedom
# 	Chi-square: 46882.23 on 269 degrees of freedom, asymptotic p-value 0 
# AIC: 30283.98 AICC: 30315.67 BIC: 32029.13 
# Log posterior: -284136.8 
# Prior parameters: mu=0 sigma=1000 nu=4 
}


#########################################################################

###Testing different intercept models###

## try int.list with one rate for all receivers, see if that model does better. This is equivalent to fitting a global attack rate


{


#### create statlist ####

##"a three dimensional array, event number by event type by statistic, or a list thereof, containing the sufficient statistics for the model"

#start with the intercept sufficient stats

sequences<-table(event.list$Date)

#create list of dummy intercept names. Leave male 30.receives as the baseline- this is the receive rate for ?? males- I expect everyone else's receive rate to be less

idno<-length(event.types[grep("cop.start",event.types)])

dummy.ints.attack<-paste(1:(idno-1),".attackFE", sep="")
#dummy.ints.receive<-paste(1:(idno-1), ".receiveFE", sep="")


int.names<-c("Global.attackFE", dummy.ints.attack,event.types[-grep("attack", event.types)])

int.list<-list()

for(i in 1:length(levels(event.list$Date))){

day.array<-array(0,dim=c(as.numeric(sequences[i]), length(event.types),length(int.names)))

#the third dimension is the stat slice.
dimnames(day.array)<-list(c(1:sequences[i]),as.character(event.types),as.character(int.names))

#add global attack effect, loop for dummy intercepts for attacks, then one loop for typical intercepts

day.array[,grep("attack", event.types),1]<-1

for(k in 2:(length(dummy.ints.attack)+1)){
	
day.array[,startsWith(names(day.array[1,,k]), paste(k-1,".attack", sep="")),k]<-1

}


for(k in (length(dummy.ints.attack)+2):length(int.names)){
	
	day.array[,int.names[k],k]<-1
	
}

#weird extra formatting to make it match the package example, seems like it becomes relevant *punintended!* if I want to fit day-specific effects

global<-list()

global[[1]]<-day.array

int.list[[i]]<-global

names(int.list[[i]])<-"global"

}


}

#int.list[[1]][[1]][1,,2]
#int.list[[1]][[1]][1,,3]

#library(tictoc)

#library(relevent)

tic()
intfit.sup<-rem(evlist, int.list, supplist=supplist, timing="interval", estimator="BPM")
toc()

summary(intfit.sup)

#quite a bit worse than fitting individual FEs for receiving. 

# Egocentric Relational Event Model (Interval Likelihood)
# 
#                  Post.Mode    Post.SD  Z value  Pr(>|z|)    
# Global.attackFE  -9.525925   0.288675 -32.9988 < 2.2e-16 ***
# 1.attackFE        1.939537   0.295363   6.5666 5.147e-11 ***
# 2.attackFE        1.517461   0.299112   5.0732 3.912e-07 ***
# 3.attackFE        1.990941   0.295338   6.7412 1.570e-11 ***
# 4.attackFE        2.357111   0.294796   7.9957 1.332e-15 ***
# 5.attackFE        1.495783   0.304577   4.9110 9.061e-07 ***
# 6.attackFE       -0.196523   0.763762  -0.2573 0.7969402    
# 7.attackFE        1.555175   0.299926   5.1852 2.158e-07 ***
# 8.attackFE        1.821930   0.348991   5.2206 1.784e-07 ***
# 9.attackFE       -1.507956   1.040831  -1.4488 0.1473935    
# 10.attackFE       2.419982   0.300964   8.0408 8.882e-16 ***
# 11.attackFE     -10.608213 262.524409  -0.0404 0.9677675    
# 12.attackFE       0.294406   0.456435   0.6450 0.5189199    
# 13.attackFE       0.955885   1.040833   0.9184 0.3584177    
# 14.attackFE       1.803741   0.353553   5.1018 3.365e-07 ***
# 15.attackFE       0.321364   0.645497   0.4979 0.6185856    
# 16.attackFE      -8.269266 293.783224  -0.0281 0.9775445    
# 17.attackFE       1.252211   0.763763   1.6395 0.1011032    
# 18.attackFE      -9.804771 272.109438  -0.0360 0.9712565    
# 19.attackFE       1.836411   0.393398   4.6681 3.040e-06 ***
# 20.attackFE       0.112428   1.040832   0.1080 0.9139816    
# 21.attackFE      -8.401427 291.711085  -0.0288 0.9770237    
# 22.attackFE       1.114675   0.645497   1.7268 0.0841949 .  
# 23.attackFE     -10.473254 264.063814  -0.0397 0.9683627    
# 24.attackFE     -10.903385 259.249258  -0.0421 0.9664528    
# 25.attackFE       3.247766   0.341565   9.5085 < 2.2e-16 ***
# 26.attackFE       1.897968   0.577350   3.2874 0.0010113 ** 
# 27.attackFE       0.704884   0.577350   1.2209 0.2221256    
# 28.attackFE       2.303485   0.393398   5.8554 4.760e-09 ***
# 29.attackFE       2.969206   0.500000   5.9384 2.878e-09 ***
# 1.initFO          6.744430   0.127000  53.1057 < 2.2e-16 ***
# 2.initFO          6.700116   0.104828  63.9150 < 2.2e-16 ***
# 3.initFO          6.738679   0.114708  58.7464 < 2.2e-16 ***
# 4.initFO          6.684612   0.075378  88.6814 < 2.2e-16 ***
# 5.initFO          6.792686   0.156174  43.4944 < 2.2e-16 ***
# 6.initFO          5.061926   0.577351   8.7675 < 2.2e-16 ***
# 7.initFO          6.715748   0.085436  78.6058 < 2.2e-16 ***
# 8.initFO          5.115994   0.500000  10.2320 < 2.2e-16 ***
# 9.initFO          3.381391   1.000001   3.3814 0.0007212 ***
# 10.initFO         6.662633   0.166667  39.9758 < 2.2e-16 ***
# 11.initFO        -6.724740 321.814902  -0.0209 0.9833284    
# 12.initFO         5.809142   0.447214  12.9896 < 2.2e-16 ***
# 13.initFO        -5.062715 363.256388  -0.0139 0.9888802    
# 14.initFO         5.460835   0.500000  10.9217 < 2.2e-16 ***
# 15.initFO         6.096823   0.500000  12.1936 < 2.2e-16 ***
# 16.initFO        -6.841114 319.417929  -0.0214 0.9829127    
# 17.initFO        -5.646696 346.931976  -0.0163 0.9870141    
# 18.initFO        -5.062715 363.256388  -0.0139 0.9888802    
# 19.initFO         4.422843   1.000002   4.4228 9.741e-06 ***
# 20.initFO         3.506554   1.000002   3.5065 0.0004540 ***
# 21.initFO        -5.062715 363.256388  -0.0139 0.9888802    
# 22.initFO         0.000000 894.427191   0.0000 1.0000000    
# 23.initFO        -5.062715 363.256388  -0.0139 0.9888802    
# 24.initFO        -7.553047 305.835850  -0.0247 0.9802971    
# 25.initFO         6.214600   1.000003   6.2146 5.146e-10 ***
# 26.initFO         6.907751   0.707108   9.7690 < 2.2e-16 ***
# 27.initFO         6.060457   0.333333  18.1814 < 2.2e-16 ***
# 28.initFO         6.502286   0.707108   9.1956 < 2.2e-16 ***
# 29.initFO         6.907747   1.000004   6.9077 4.925e-12 ***
# 30.initFO         3.352407   0.258199  12.9838 < 2.2e-16 ***
# 1.initCH          2.617293   1.000001   2.6173 0.0088631 ** 
# 2.initCH         -9.185505 280.258756  -0.0328 0.9738539    
# 3.initCH         -8.988496 283.008974  -0.0318 0.9746631    
# 4.initCH          1.514126   1.000000   1.5141 0.1299940    
# 5.initCH          3.079110   1.000001   3.0791 0.0020762 ** 
# 6.initCH          6.735904   0.250000  26.9436 < 2.2e-16 ***
# 7.initCH         -9.541022 275.492334  -0.0346 0.9723727    
# 8.initCH          6.562914   0.242536  27.0596 < 2.2e-16 ***
# 9.initCH          6.877902   0.174078  39.5105 < 2.2e-16 ***
# 10.initCH         4.465407   0.500000   8.9308 < 2.2e-16 ***
# 11.initCH         6.753603   0.408249  16.5429 < 2.2e-16 ***
# 12.initCH        -7.392244 308.751880  -0.0239 0.9808986    
# 13.initCH        -5.062715 363.256388  -0.0139 0.9888802    
# 14.initCH         6.271766   0.333333  18.8153 < 2.2e-16 ***
# 15.initCH         5.403675   0.707108   7.6419 2.132e-14 ***
# 16.initCH         6.907754   0.353554  19.5381 < 2.2e-16 ***
# 17.initCH         6.214600   1.000003   6.2146 5.146e-10 ***
# 18.initCH         6.907747   1.000004   6.9077 4.925e-12 ***
# 19.initCH         5.809141   0.500000  11.6183 < 2.2e-16 ***
# 20.initCH         6.597600   0.213201  30.9455 < 2.2e-16 ***
# 21.initCH         6.907747   1.000004   6.9077 4.925e-12 ***
# 22.initCH         0.000000 894.427191   0.0000 1.0000000    
# 23.initCH        -5.062715 363.256388  -0.0139 0.9888802    
# 24.initCH         6.850596   0.242536  28.2457 < 2.2e-16 ***
# 25.initCH        -5.646696 346.931976  -0.0163 0.9870141    
# 26.initCH        -5.646696 346.931976  -0.0163 0.9870141    
# 27.initCH         6.060457   0.333333  18.1814 < 2.2e-16 ***
# 28.initCH        -5.992692 338.240255  -0.0177 0.9858644    
# 29.initCH        -5.062715 363.256388  -0.0139 0.9888802    
# 30.initCH         6.824374   0.045502 149.9810 < 2.2e-16 ***
# 1.initSF          4.919880   0.316228  15.5580 < 2.2e-16 ***
# 2.initSF          5.233779   0.218218  23.9842 < 2.2e-16 ***
# 3.initSF          5.047002   0.267261  18.8842 < 2.2e-16 ***
# 4.initSF          5.275328   0.152499  34.5926 < 2.2e-16 ***
# 5.initSF          4.465407   0.500000   8.9308 < 2.2e-16 ***
# 6.initSF         -7.600810 304.985488  -0.0249 0.9801173    
# 7.initSF          5.163063   0.185695  27.8039 < 2.2e-16 ***
# 8.initSF          4.828312   0.577351   8.3629 < 2.2e-16 ***
# 9.initSF         -8.117025 296.225732  -0.0274 0.9781395    
# 10.initSF         4.870872   0.408248  11.9311 < 2.2e-16 ***
# 11.initSF         4.961839   1.000002   4.9618 6.983e-07 ***
# 12.initSF         6.502289   0.316228  20.5620 < 2.2e-16 ***
# 13.initSF         6.907747   1.000004   6.9077 4.925e-12 ***
# 14.initSF         5.460835   0.500000  10.9217 < 2.2e-16 ***
# 15.initSF         5.809141   0.577351  10.0617 < 2.2e-16 ***
# 16.initSF        -6.841114 319.417929  -0.0214 0.9829127    
# 17.initSF         6.214600   1.000003   6.2146 5.146e-10 ***
# 18.initSF        -5.062715 363.256388  -0.0139 0.9888802    
# 19.initSF         6.368758   0.377965  16.8501 < 2.2e-16 ***
# 20.initSF         5.452467   0.377965  14.4259 < 2.2e-16 ***
# 21.initSF        -5.062715 363.256388  -0.0139 0.9888802    
# 22.initSF         0.000000 894.427191   0.0000 1.0000000    
# 23.initSF         6.907747   1.000004   6.9077 4.925e-12 ***
# 24.initSF         4.017378   1.000002   4.0174 5.885e-05 ***
# 25.initSF         6.214600   1.000003   6.2146 5.146e-10 ***
# 26.initSF        -5.646696 346.931976  -0.0163 0.9870141    
# 27.initSF         4.961843   0.577351   8.5942 < 2.2e-16 ***
# 28.initSF         5.809136   1.000003   5.8091 6.280e-09 ***
# 29.initSF        -5.062715 363.256388  -0.0139 0.9888802    
# 30.initSF         3.940194   0.192450  20.4738 < 2.2e-16 ***
# 1.disengage      -4.262284   0.062622 -68.0632 < 2.2e-16 ***
# 2.disengage      -5.148770   0.088736 -58.0237 < 2.2e-16 ***
# 3.disengage      -4.775014   0.063628 -75.0452 < 2.2e-16 ***
# 4.disengage      -4.647453   0.054882 -84.6806 < 2.2e-16 ***
# 5.disengage      -4.573716   0.102598 -44.5791 < 2.2e-16 ***
# 6.disengage      -6.477786   0.577349 -11.2199 < 2.2e-16 ***
# 7.disengage      -5.683178   0.089443 -63.5399 < 2.2e-16 ***
# 8.disengage      -3.025632   0.188982 -16.0101 < 2.2e-16 ***
# 9.disengage      -2.951359   0.999998  -2.9514 0.0031637 ** 
# 10.disengage     -4.610931   0.088045 -52.3701 < 2.2e-16 ***
# 11.disengage     -1.251904   0.999999  -1.2519 0.2106042    
# 12.disengage     -5.099954   0.333333 -15.2999 < 2.2e-16 ***
# 13.disengage     -3.069490   0.999997  -3.0695 0.0021442 ** 
# 14.disengage     -3.999558   0.200000 -19.9978 < 2.2e-16 ***
# 15.disengage     -3.711562   0.377964  -9.8199 < 2.2e-16 ***
# 16.disengage      0.000000 894.427191   0.0000 1.0000000    
# 17.disengage     -5.585480   0.707105  -7.8991 2.887e-15 ***
# 18.disengage      0.000000 894.427191   0.0000 1.0000000    
# 19.disengage     -3.306133   0.250000 -13.2245 < 2.2e-16 ***
# 20.disengage     -2.450509   0.353553  -6.9311 4.176e-12 ***
# 21.disengage      0.000000 894.427191   0.0000 1.0000000    
# 22.disengage     -3.514373   0.999997  -3.5144 0.0004408 ***
# 23.disengage    -12.189754 246.283852  -0.0495 0.9605250    
# 24.disengage     -1.891904   0.999998  -1.8919 0.0585033 .  
# 25.disengage     -2.434066   0.200000 -12.1703 < 2.2e-16 ***
# 26.disengage     -3.804916   0.408248  -9.3201 < 2.2e-16 ***
# 27.disengage     -5.368294   0.316228 -16.9760 < 2.2e-16 ***
# 28.disengage     -5.092138   0.353553 -14.4027 < 2.2e-16 ***
# 29.disengage     -5.097787   0.408248 -12.4870 < 2.2e-16 ***
# 30.disengage     -4.831031   0.162221 -29.7805 < 2.2e-16 ***
# 1.cop.start     -21.775226 187.429636  -0.1162 0.9075114    
# 2.cop.start      -9.446606   0.408248 -23.1394 < 2.2e-16 ***
# 3.cop.start      -7.431142   0.149071 -49.8496 < 2.2e-16 ***
# 4.cop.start      -7.839479   0.218218 -35.9250 < 2.2e-16 ***
# 5.cop.start     -21.334706 189.269432  -0.1127 0.9102515    
# 6.cop.start     -19.273179 198.658064  -0.0970 0.9227130    
# 7.cop.start     -21.663929 187.889476  -0.1153 0.9082062    
# 8.cop.start     -19.504105 197.536437  -0.0987 0.9213473    
# 9.cop.start     -19.225865 198.890242  -0.0967 0.9229919    
# 10.cop.start    -20.397432 193.369811  -0.1055 0.9159917    
# 11.cop.start    -16.297433 215.065250  -0.0758 0.9395949    
# 12.cop.start    -19.261927 198.713207  -0.0969 0.9227794    
# 13.cop.start    -17.124683 210.100682  -0.0815 0.9350387    
# 14.cop.start    -19.045919 199.780757  -0.0953 0.9240495    
# 15.cop.start    -19.059755 199.711859  -0.0954 0.9239683    
# 16.cop.start    -15.410565 220.799295  -0.0698 0.9443573    
# 17.cop.start    -17.846074 206.041009  -0.0866 0.9309782    
# 18.cop.start    -16.894762 211.445896  -0.0799 0.9363159    
# 19.cop.start    -19.197853 199.028082  -0.0965 0.9231568    
# 20.cop.start    -18.175836 204.262031  -0.0890 0.9290955    
# 21.cop.start    -15.559441 219.804656  -0.0708 0.9435668    
# 22.cop.start    -18.414696 203.001849  -0.0907 0.9277215    
# 23.cop.start    -17.543193 207.716617  -0.0845 0.9326928    
# 24.cop.start    -18.088681 204.727721  -0.0884 0.9295947    
# 25.cop.start    -18.516001 202.474387  -0.0914 0.9271361    
# 26.cop.start    -17.327815 208.933327  -0.0829 0.9339035    
# 27.cop.start    -18.831473 200.857793  -0.0938 0.9253036    
# 28.cop.start    -18.717694 201.436362  -0.0929 0.9259662    
# 29.cop.start      0.000000 894.427191   0.0000 1.0000000    
# 30.cop.start    -20.543323 192.714073  -0.1066 0.9151063    
# 1.cop.end         0.000000 894.427191   0.0000 1.0000000    
# 2.cop.end        -1.651538   0.707106  -2.3356 0.0195106 *  
# 3.cop.end        -1.354380   0.160128  -8.4581 < 2.2e-16 ***
# 4.cop.end        -1.737753   0.333333  -5.2133 1.856e-07 ***
# 5.cop.end         0.000000 894.427191   0.0000 1.0000000    
# 6.cop.end         0.000000 894.427191   0.0000 1.0000000    
# 7.cop.end         0.000000 894.427191   0.0000 1.0000000    
# 8.cop.end         0.000000 894.427191   0.0000 1.0000000    
# 9.cop.end         0.000000 894.427191   0.0000 1.0000000    
# 10.cop.end        0.000000 894.427191   0.0000 1.0000000    
# 11.cop.end        0.000000 894.427191   0.0000 1.0000000    
# 12.cop.end        0.000000 894.427191   0.0000 1.0000000    
# 13.cop.end        0.000000 894.427191   0.0000 1.0000000    
# 14.cop.end        0.000000 894.427191   0.0000 1.0000000    
# 15.cop.end        0.000000 894.427191   0.0000 1.0000000    
# 16.cop.end        0.000000 894.427191   0.0000 1.0000000    
# 17.cop.end        0.000000 894.427191   0.0000 1.0000000    
# 18.cop.end        0.000000 894.427191   0.0000 1.0000000    
# 19.cop.end        0.000000 894.427191   0.0000 1.0000000    
# 20.cop.end        0.000000 894.427191   0.0000 1.0000000    
# 21.cop.end        0.000000 894.427191   0.0000 1.0000000    
# 22.cop.end        0.000000 894.427191   0.0000 1.0000000    
# 23.cop.end        0.000000 894.427191   0.0000 1.0000000    
# 24.cop.end        0.000000 894.427191   0.0000 1.0000000    
# 25.cop.end        0.000000 894.427191   0.0000 1.0000000    
# 26.cop.end        0.000000 894.427191   0.0000 1.0000000    
# 27.cop.end        0.000000 894.427191   0.0000 1.0000000    
# 28.cop.end        0.000000 894.427191   0.0000 1.0000000    
# 29.cop.end        0.000000 894.427191   0.0000 1.0000000    
# 30.cop.end        0.000000 894.427191   0.0000 1.0000000    
# 1.depart         -8.701475   0.235702 -36.9172 < 2.2e-16 ***
# 2.depart         -8.645592   0.235702 -36.6802 < 2.2e-16 ***
# 3.depart         -8.603133   0.235702 -36.5000 < 2.2e-16 ***
# 4.depart         -8.213108   0.235702 -34.8453 < 2.2e-16 ***
# 5.depart         -8.797853   0.288675 -30.4767 < 2.2e-16 ***
# 6.depart         -8.107312   0.707105 -11.4655 < 2.2e-16 ***
# 7.depart         -8.466583   0.235702 -35.9207 < 2.2e-16 ***
# 8.depart         -7.624461   0.499999 -15.2489 < 2.2e-16 ***
# 9.depart         -8.213118   0.577349 -14.2256 < 2.2e-16 ***
# 10.depart        -8.336652   0.377964 -22.0567 < 2.2e-16 ***
# 11.depart        -6.136106   0.577349 -10.6281 < 2.2e-16 ***
# 12.depart        -8.430566   0.577349 -14.6022 < 2.2e-16 ***
# 13.depart        -6.374262   0.999995  -6.3743 1.838e-10 ***
# 14.depart        -8.432862   0.707105 -11.9259 < 2.2e-16 ***
# 15.depart        -8.472487   0.999994  -8.4725 < 2.2e-16 ***
# 16.depart        -4.553187   0.999997  -4.5532 5.284e-06 ***
# 17.depart        -7.332594   0.999995  -7.3326 2.256e-13 ***
# 18.depart        -6.129319   0.999996  -6.1293 8.824e-10 ***
# 19.depart        -8.897214   0.999994  -8.8973 < 2.2e-16 ***
# 20.depart        -7.676151   0.999995  -7.6762 1.643e-14 ***
# 21.depart        -4.711675   0.999996  -4.7117 2.457e-06 ***
# 22.depart        -7.735384   0.999995  -7.7354 1.021e-14 ***
# 23.depart        -7.017221   0.999995  -7.0173 2.263e-12 ***
# 24.depart        -7.391510   0.999995  -7.3915 1.452e-13 ***
# 25.depart        -7.842174   0.999994  -7.8422 4.441e-15 ***
# 26.depart        -8.035618   0.999994  -8.0357 8.882e-16 ***
# 27.depart        -8.659685   0.999994  -8.6597 < 2.2e-16 ***
# 28.depart        -8.191807   0.999994  -8.1919 2.220e-16 ***
# 29.depart        -7.054816   0.999995  -7.0549 1.728e-12 ***
# 30.depart        -5.737913   0.117851 -48.6878 < 2.2e-16 ***
# ---
# Signif. codes:  0 ‘***’ 0.001 ‘**’ 0.01 ‘*’ 0.05 ‘.’ 0.1 ‘ ’ 1
# Null deviance: 76628.21 on 4854 degrees of freedom
# Residual deviance: 31465.49 on 4614 degrees of freedom
# 	Chi-square: 45162.72 on 240 degrees of freedom, asymptotic p-value 0 
# AIC: 31945.49 AICC: 31970.57 BIC: 33502.51 
# Log posterior: -255968.2 
# Prior parameters: mu=0 sigma=1000 nu=4 

##########


## Try fitting an Attack FE and a Receive FE that lumps all the U's 

#right now, male nums 1-14 are ID'd males, and then 15-29 are U's, and 30 is ??'s 

sequences<-table(event.list$Date)

#create list of dummy intercept names. Leave male 30.receives as the baseline- this is the receive rate for ?? males- I expect everyone else's receive rate to be less?

idno<-length(event.types[grep("cop.start",event.types)])


dummy.ints.attack<-paste(c(1:14,"U",idno),".attackFE", sep="")
dummy.ints.receive<-paste(c(1:14,"U"), ".receiveFE", sep="")


int.names<-c(dummy.ints.attack,dummy.ints.receive,event.types[-grep("attack", event.types)])

int.list<-list()

for(i in 1:length(levels(event.list$Date))){

day.array<-array(0,dim=c(as.numeric(sequences[i]), length(event.types),length(int.names)))

#the third dimension is the stat slice.
dimnames(day.array)<-list(c(1:sequences[i]),as.character(event.types),as.character(int.names))

#loop for dummy intercepts for attacks, loop for dummy intercepts for receives, then one loop for typical intercepts

for(k in 1:length(dummy.ints.attack)){
	
	if(k<(length(dummy.ints.attack)-2)){
day.array[,startsWith(names(day.array[1,,k]), paste(k,".attack", sep="")),k]<-1
} 

if(k==(length(dummy.ints.attack)-1)){
	
	for(h in 15:29){
	day.array[,startsWith(names(day.array[1,,k]), paste(h,".attack", sep="")),k]<-1
}

}

if(k==length(dummy.ints.attack)){
	day.array[,startsWith(names(day.array[1,,k]), paste(30,".attack", sep="")),k]<-1
}

}

for(k in (length(dummy.ints.attack)+1):(length(dummy.ints.attack)+length(dummy.ints.receive))){

if(k<(length(dummy.ints.attack)+length(dummy.ints.receive)-1)){
day.array[,endsWith(names(day.array[1,,k]), paste("attack.",(k-length(dummy.ints.attack)), sep="")),k]<-1
}
	
if(k==(length(dummy.ints.attack)+length(dummy.ints.receive))){
	
	for(h in 15:29){
day.array[,endsWith(names(day.array[1,,k]), paste("attack.",h, sep="")),k]<-1
}

}

# if(k==(length(dummy.ints.attack)+length(dummy.ints.receive))){
# day.array[,endsWith(names(day.array[1,,k]), paste("attack.",30, sep="")),k]<-1
# }

}

for(k in (length(dummy.ints.attack)+length(dummy.ints.receive)+1):length(int.names)){
	
	day.array[,int.names[k],k]<-1
	
}

#weird extra formatting to make it match the package example, seems like it becomes relevant *punintended!* if I want to fit day-specific effects

global<-list()

global[[1]]<-day.array

int.list[[i]]<-global

names(int.list[[i]])<-"global"

}


#library(tictoc)

#library(relevent)

tic()
intfit.sup<-rem(evlist, int.list, supplist=supplist, timing="interval", estimator="BPM")
toc()

summary(intfit.sup)

# Egocentric Relational Event Model (Interval Likelihood)
# 
#                 Post.Mode      Post.SD  Z value  Pr(>|z|)    
# 1.attackFE    -5.0972e+00   6.3706e-02 -80.0114 < 2.2e-16 ***
# 2.attackFE    -5.5110e+00   7.9180e-02 -69.6007 < 2.2e-16 ***
# 3.attackFE    -5.0395e+00   6.3517e-02 -79.3403 < 2.2e-16 ***
# 4.attackFE    -4.7170e+00   6.0535e-02 -77.9219 < 2.2e-16 ***
# 5.attackFE    -5.3758e+00   9.7739e-02 -55.0013 < 2.2e-16 ***
# 6.attackFE    -7.1244e+00   7.0714e-01 -10.0750 < 2.2e-16 ***
# 7.attackFE    -5.4850e+00   8.2050e-02 -66.8490 < 2.2e-16 ***
# 8.attackFE    -5.4381e+00   1.9635e-01 -27.6955 < 2.2e-16 ***
# 9.attackFE    -8.5898e+00   1.0000e+00  -8.5896 < 2.2e-16 ***
# 10.attackFE   -5.0365e+00   8.5512e-02 -58.8979 < 2.2e-16 ***
# 11.attackFE   -1.8084e+01   2.0475e+02  -0.0883 0.9296228    
# 12.attackFE   -7.4130e+00   3.5359e-01 -20.9650 < 2.2e-16 ***
# 13.attackFE   -5.4092e+00   1.0001e+00  -5.4089 6.342e-08 ***
# 14.attackFE    6.8091e-15   8.9443e+02   0.0000 1.0000000    
# U.attackFE    -5.2159e+00   1.1395e-01 -45.7754 < 2.2e-16 ***
# 30.attackFE   -6.3349e+00   2.8970e-01 -21.8671 < 2.2e-16 ***
# 1.receiveFE   -4.7675e+00   1.1740e-01 -40.6083 < 2.2e-16 ***
# 2.receiveFE   -4.4877e+00   9.4838e-02 -47.3198 < 2.2e-16 ***
# 3.receiveFE   -4.6843e+00   1.0569e-01 -44.3229 < 2.2e-16 ***
# 4.receiveFE   -3.5508e+00   6.7785e-02 -52.3829 < 2.2e-16 ***
# 5.receiveFE   -4.5004e+00   1.4798e-01 -30.4114 < 2.2e-16 ***
# 6.receiveFE   -2.2085e+00   2.3169e-01  -9.5322 < 2.2e-16 ***
# 7.receiveFE   -3.9979e+00   7.8011e-02 -51.2479 < 2.2e-16 ***
# 8.receiveFE   -4.2965e+00   2.0419e-01 -21.0419 < 2.2e-16 ***
# 9.receiveFE   -2.2404e+00   1.7399e-01 -12.8764 < 2.2e-16 ***
# 10.receiveFE  -4.9741e+00   1.4749e-01 -33.7260 < 2.2e-16 ***
# 11.receiveFE  -3.1618e+00   3.7803e-01  -8.3638 < 2.2e-16 ***
# 12.receiveFE  -6.3038e+00   2.5820e-01 -24.4140 < 2.2e-16 ***
# 13.receiveFE  -3.2557e+00   1.0005e+00  -3.2539 0.0011381 ** 
# 14.receiveFE -2.0701e-171   8.9443e+02   0.0000 1.0000000    
# U.receiveFE   -2.2339e+00   1.0065e-01 -22.1937 < 2.2e-16 ***
# 1.initFO       6.7444e+00   1.2700e-01  53.1057 < 2.2e-16 ***
# 2.initFO       6.7001e+00   1.0483e-01  63.9150 < 2.2e-16 ***
# 3.initFO       6.7387e+00   1.1471e-01  58.7464 < 2.2e-16 ***
# 4.initFO       6.6846e+00   7.5378e-02  88.6814 < 2.2e-16 ***
# 5.initFO       6.7927e+00   1.5617e-01  43.4944 < 2.2e-16 ***
# 6.initFO       5.0619e+00   5.7735e-01   8.7675 < 2.2e-16 ***
# 7.initFO       6.7157e+00   8.5436e-02  78.6058 < 2.2e-16 ***
# 8.initFO       5.1160e+00   5.0000e-01  10.2320 < 2.2e-16 ***
# 9.initFO       3.3814e+00   1.0000e+00   3.3814 0.0007212 ***
# 10.initFO      6.6626e+00   1.6667e-01  39.9758 < 2.2e-16 ***
# 11.initFO     -6.7247e+00   3.2181e+02  -0.0209 0.9833284    
# 12.initFO      5.8091e+00   4.4721e-01  12.9896 < 2.2e-16 ***
# 13.initFO     -5.0627e+00   3.6326e+02  -0.0139 0.9888802    
# 14.initFO      5.4608e+00   5.0000e-01  10.9217 < 2.2e-16 ***
# 15.initFO      6.0968e+00   5.0000e-01  12.1936 < 2.2e-16 ***
# 16.initFO     -6.8411e+00   3.1942e+02  -0.0214 0.9829127    
# 17.initFO     -5.6467e+00   3.4693e+02  -0.0163 0.9870141    
# 18.initFO     -5.0627e+00   3.6326e+02  -0.0139 0.9888802    
# 19.initFO      4.4228e+00   1.0000e+00   4.4228 9.741e-06 ***
# 20.initFO      3.5066e+00   1.0000e+00   3.5065 0.0004540 ***
# 21.initFO     -5.0627e+00   3.6326e+02  -0.0139 0.9888802    
# 22.initFO      0.0000e+00   8.9443e+02   0.0000 1.0000000    
# 23.initFO     -5.0627e+00   3.6326e+02  -0.0139 0.9888802    
# 24.initFO     -7.5530e+00   3.0584e+02  -0.0247 0.9802971    
# 25.initFO      6.2146e+00   1.0000e+00   6.2146 5.146e-10 ***
# 26.initFO      6.9078e+00   7.0711e-01   9.7690 < 2.2e-16 ***
# 27.initFO      6.0605e+00   3.3333e-01  18.1814 < 2.2e-16 ***
# 28.initFO      6.5023e+00   7.0711e-01   9.1956 < 2.2e-16 ***
# 29.initFO      6.9077e+00   1.0000e+00   6.9077 4.925e-12 ***
# 30.initFO      3.3524e+00   2.5820e-01  12.9838 < 2.2e-16 ***
# 1.initCH       2.6173e+00   1.0000e+00   2.6173 0.0088631 ** 
# 2.initCH      -9.1855e+00   2.8026e+02  -0.0328 0.9738539    
# 3.initCH      -8.9885e+00   2.8301e+02  -0.0318 0.9746631    
# 4.initCH       1.5141e+00   1.0000e+00   1.5141 0.1299940    
# 5.initCH       3.0791e+00   1.0000e+00   3.0791 0.0020762 ** 
# 6.initCH       6.7359e+00   2.5000e-01  26.9436 < 2.2e-16 ***
# 7.initCH      -9.5410e+00   2.7549e+02  -0.0346 0.9723727    
# 8.initCH       6.5629e+00   2.4254e-01  27.0596 < 2.2e-16 ***
# 9.initCH       6.8779e+00   1.7408e-01  39.5105 < 2.2e-16 ***
# 10.initCH      4.4654e+00   5.0000e-01   8.9308 < 2.2e-16 ***
# 11.initCH      6.7536e+00   4.0825e-01  16.5429 < 2.2e-16 ***
# 12.initCH     -7.3922e+00   3.0875e+02  -0.0239 0.9808986    
# 13.initCH     -5.0627e+00   3.6326e+02  -0.0139 0.9888802    
# 14.initCH      6.2718e+00   3.3333e-01  18.8153 < 2.2e-16 ***
# 15.initCH      5.4037e+00   7.0711e-01   7.6419 2.132e-14 ***
# 16.initCH      6.9078e+00   3.5355e-01  19.5381 < 2.2e-16 ***
# 17.initCH      6.2146e+00   1.0000e+00   6.2146 5.146e-10 ***
# 18.initCH      6.9077e+00   1.0000e+00   6.9077 4.925e-12 ***
# 19.initCH      5.8091e+00   5.0000e-01  11.6183 < 2.2e-16 ***
# 20.initCH      6.5976e+00   2.1320e-01  30.9455 < 2.2e-16 ***
# 21.initCH      6.9077e+00   1.0000e+00   6.9077 4.925e-12 ***
# 22.initCH      0.0000e+00   8.9443e+02   0.0000 1.0000000    
# 23.initCH     -5.0627e+00   3.6326e+02  -0.0139 0.9888802    
# 24.initCH      6.8506e+00   2.4254e-01  28.2457 < 2.2e-16 ***
# 25.initCH     -5.6467e+00   3.4693e+02  -0.0163 0.9870141    
# 26.initCH     -5.6467e+00   3.4693e+02  -0.0163 0.9870141    
# 27.initCH      6.0605e+00   3.3333e-01  18.1814 < 2.2e-16 ***
# 28.initCH     -5.9927e+00   3.3824e+02  -0.0177 0.9858644    
# 29.initCH     -5.0627e+00   3.6326e+02  -0.0139 0.9888802    
# 30.initCH      6.8244e+00   4.5502e-02 149.9810 < 2.2e-16 ***
# 1.initSF       4.9199e+00   3.1623e-01  15.5580 < 2.2e-16 ***
# 2.initSF       5.2338e+00   2.1822e-01  23.9842 < 2.2e-16 ***
# 3.initSF       5.0470e+00   2.6726e-01  18.8842 < 2.2e-16 ***
# 4.initSF       5.2753e+00   1.5250e-01  34.5926 < 2.2e-16 ***
# 5.initSF       4.4654e+00   5.0000e-01   8.9308 < 2.2e-16 ***
# 6.initSF      -7.6008e+00   3.0499e+02  -0.0249 0.9801173    
# 7.initSF       5.1631e+00   1.8570e-01  27.8039 < 2.2e-16 ***
# 8.initSF       4.8283e+00   5.7735e-01   8.3629 < 2.2e-16 ***
# 9.initSF      -8.1170e+00   2.9623e+02  -0.0274 0.9781395    
# 10.initSF      4.8709e+00   4.0825e-01  11.9311 < 2.2e-16 ***
# 11.initSF      4.9618e+00   1.0000e+00   4.9618 6.983e-07 ***
# 12.initSF      6.5023e+00   3.1623e-01  20.5620 < 2.2e-16 ***
# 13.initSF      6.9077e+00   1.0000e+00   6.9077 4.925e-12 ***
# 14.initSF      5.4608e+00   5.0000e-01  10.9217 < 2.2e-16 ***
# 15.initSF      5.8091e+00   5.7735e-01  10.0617 < 2.2e-16 ***
# 16.initSF     -6.8411e+00   3.1942e+02  -0.0214 0.9829127    
# 17.initSF      6.2146e+00   1.0000e+00   6.2146 5.146e-10 ***
# 18.initSF     -5.0627e+00   3.6326e+02  -0.0139 0.9888802    
# 19.initSF      6.3688e+00   3.7796e-01  16.8501 < 2.2e-16 ***
# 20.initSF      5.4525e+00   3.7796e-01  14.4259 < 2.2e-16 ***
# 21.initSF     -5.0627e+00   3.6326e+02  -0.0139 0.9888802    
# 22.initSF      0.0000e+00   8.9443e+02   0.0000 1.0000000    
# 23.initSF      6.9077e+00   1.0000e+00   6.9077 4.925e-12 ***
# 24.initSF      4.0174e+00   1.0000e+00   4.0174 5.885e-05 ***
# 25.initSF      6.2146e+00   1.0000e+00   6.2146 5.146e-10 ***
# 26.initSF     -5.6467e+00   3.4693e+02  -0.0163 0.9870141    
# 27.initSF      4.9618e+00   5.7735e-01   8.5942 < 2.2e-16 ***
# 28.initSF      5.8091e+00   1.0000e+00   5.8091 6.280e-09 ***
# 29.initSF     -5.0627e+00   3.6326e+02  -0.0139 0.9888802    
# 30.initSF      3.9402e+00   1.9245e-01  20.4738 < 2.2e-16 ***
# 1.disengage   -4.2623e+00   6.2622e-02 -68.0632 < 2.2e-16 ***
# 2.disengage   -5.1488e+00   8.8736e-02 -58.0237 < 2.2e-16 ***
# 3.disengage   -4.7750e+00   6.3628e-02 -75.0452 < 2.2e-16 ***
# 4.disengage   -4.6475e+00   5.4882e-02 -84.6806 < 2.2e-16 ***
# 5.disengage   -4.5737e+00   1.0260e-01 -44.5791 < 2.2e-16 ***
# 6.disengage   -6.4778e+00   5.7735e-01 -11.2199 < 2.2e-16 ***
# 7.disengage   -5.6832e+00   8.9443e-02 -63.5399 < 2.2e-16 ***
# 8.disengage   -3.0256e+00   1.8898e-01 -16.0101 < 2.2e-16 ***
# 9.disengage   -2.9514e+00   1.0000e+00  -2.9514 0.0031637 ** 
# 10.disengage  -4.6109e+00   8.8045e-02 -52.3701 < 2.2e-16 ***
# 11.disengage  -1.2519e+00   1.0000e+00  -1.2519 0.2106042    
# 12.disengage  -5.1000e+00   3.3333e-01 -15.2999 < 2.2e-16 ***
# 13.disengage  -3.0695e+00   1.0000e+00  -3.0695 0.0021442 ** 
# 14.disengage  -3.9996e+00   2.0000e-01 -19.9978 < 2.2e-16 ***
# 15.disengage  -3.7116e+00   3.7796e-01  -9.8199 < 2.2e-16 ***
# 16.disengage   0.0000e+00   8.9443e+02   0.0000 1.0000000    
# 17.disengage  -5.5855e+00   7.0711e-01  -7.8991 2.887e-15 ***
# 18.disengage   0.0000e+00   8.9443e+02   0.0000 1.0000000    
# 19.disengage  -3.3061e+00   2.5000e-01 -13.2245 < 2.2e-16 ***
# 20.disengage  -2.4505e+00   3.5355e-01  -6.9311 4.176e-12 ***
# 21.disengage   0.0000e+00   8.9443e+02   0.0000 1.0000000    
# 22.disengage  -3.5144e+00   1.0000e+00  -3.5144 0.0004408 ***
# 23.disengage  -1.2190e+01   2.4628e+02  -0.0495 0.9605250    
# 24.disengage  -1.8919e+00   1.0000e+00  -1.8919 0.0585033 .  
# 25.disengage  -2.4341e+00   2.0000e-01 -12.1703 < 2.2e-16 ***
# 26.disengage  -3.8049e+00   4.0825e-01  -9.3201 < 2.2e-16 ***
# 27.disengage  -5.3683e+00   3.1623e-01 -16.9760 < 2.2e-16 ***
# 28.disengage  -5.0921e+00   3.5355e-01 -14.4027 < 2.2e-16 ***
# 29.disengage  -5.0978e+00   4.0825e-01 -12.4870 < 2.2e-16 ***
# 30.disengage  -4.8310e+00   1.6222e-01 -29.7805 < 2.2e-16 ***
# 1.cop.start   -2.1775e+01   1.8743e+02  -0.1162 0.9075119    
# 2.cop.start   -9.4466e+00   4.0825e-01 -23.1394 < 2.2e-16 ***
# 3.cop.start   -7.4311e+00   1.4907e-01 -49.8496 < 2.2e-16 ***
# 4.cop.start   -7.8395e+00   2.1822e-01 -35.9250 < 2.2e-16 ***
# 5.cop.start   -2.1335e+01   1.8927e+02  -0.1127 0.9102517    
# 6.cop.start   -1.9273e+01   1.9866e+02  -0.0970 0.9227130    
# 7.cop.start   -2.1664e+01   1.8789e+02  -0.1153 0.9082066    
# 8.cop.start   -1.9504e+01   1.9754e+02  -0.0987 0.9213473    
# 9.cop.start   -1.9226e+01   1.9889e+02  -0.0967 0.9229919    
# 10.cop.start  -2.0397e+01   1.9337e+02  -0.1055 0.9159918    
# 11.cop.start  -1.6297e+01   2.1507e+02  -0.0758 0.9395949    
# 12.cop.start  -1.9262e+01   1.9871e+02  -0.0969 0.9227794    
# 13.cop.start  -1.7125e+01   2.1010e+02  -0.0815 0.9350387    
# 14.cop.start  -1.9046e+01   1.9978e+02  -0.0953 0.9240495    
# 15.cop.start  -1.9060e+01   1.9971e+02  -0.0954 0.9239683    
# 16.cop.start  -1.5411e+01   2.2080e+02  -0.0698 0.9443573    
# 17.cop.start  -1.7846e+01   2.0604e+02  -0.0866 0.9309782    
# 18.cop.start  -1.6895e+01   2.1145e+02  -0.0799 0.9363159    
# 19.cop.start  -1.9198e+01   1.9903e+02  -0.0965 0.9231568    
# 20.cop.start  -1.8176e+01   2.0426e+02  -0.0890 0.9290955    
# 21.cop.start  -1.5559e+01   2.1980e+02  -0.0708 0.9435668    
# 22.cop.start  -1.8415e+01   2.0300e+02  -0.0907 0.9277215    
# 23.cop.start  -1.7543e+01   2.0772e+02  -0.0845 0.9326928    
# 24.cop.start  -1.8089e+01   2.0473e+02  -0.0884 0.9295947    
# 25.cop.start  -1.8516e+01   2.0247e+02  -0.0914 0.9271361    
# 26.cop.start  -1.7328e+01   2.0893e+02  -0.0829 0.9339035    
# 27.cop.start  -1.8831e+01   2.0086e+02  -0.0938 0.9253036    
# 28.cop.start  -1.8718e+01   2.0144e+02  -0.0929 0.9259662    
# 29.cop.start   0.0000e+00   8.9443e+02   0.0000 1.0000000    
# 30.cop.start  -2.0543e+01   1.9271e+02  -0.1066 0.9151063    
# 1.cop.end      0.0000e+00   8.9443e+02   0.0000 1.0000000    
# 2.cop.end     -1.6515e+00   7.0711e-01  -2.3356 0.0195106 *  
# 3.cop.end     -1.3544e+00   1.6013e-01  -8.4581 < 2.2e-16 ***
# 4.cop.end     -1.7378e+00   3.3333e-01  -5.2133 1.856e-07 ***
# 5.cop.end      0.0000e+00   8.9443e+02   0.0000 1.0000000    
# 6.cop.end      0.0000e+00   8.9443e+02   0.0000 1.0000000    
# 7.cop.end      0.0000e+00   8.9443e+02   0.0000 1.0000000    
# 8.cop.end      0.0000e+00   8.9443e+02   0.0000 1.0000000    
# 9.cop.end      0.0000e+00   8.9443e+02   0.0000 1.0000000    
# 10.cop.end     0.0000e+00   8.9443e+02   0.0000 1.0000000    
# 11.cop.end     0.0000e+00   8.9443e+02   0.0000 1.0000000    
# 12.cop.end     0.0000e+00   8.9443e+02   0.0000 1.0000000    
# 13.cop.end     0.0000e+00   8.9443e+02   0.0000 1.0000000    
# 14.cop.end     0.0000e+00   8.9443e+02   0.0000 1.0000000    
# 15.cop.end     0.0000e+00   8.9443e+02   0.0000 1.0000000    
# 16.cop.end     0.0000e+00   8.9443e+02   0.0000 1.0000000    
# 17.cop.end     0.0000e+00   8.9443e+02   0.0000 1.0000000    
# 18.cop.end     0.0000e+00   8.9443e+02   0.0000 1.0000000    
# 19.cop.end     0.0000e+00   8.9443e+02   0.0000 1.0000000    
# 20.cop.end     0.0000e+00   8.9443e+02   0.0000 1.0000000    
# 21.cop.end     0.0000e+00   8.9443e+02   0.0000 1.0000000    
# 22.cop.end     0.0000e+00   8.9443e+02   0.0000 1.0000000    
# 23.cop.end     0.0000e+00   8.9443e+02   0.0000 1.0000000    
# 24.cop.end     0.0000e+00   8.9443e+02   0.0000 1.0000000    
# 25.cop.end     0.0000e+00   8.9443e+02   0.0000 1.0000000    
# 26.cop.end     0.0000e+00   8.9443e+02   0.0000 1.0000000    
# 27.cop.end     0.0000e+00   8.9443e+02   0.0000 1.0000000    
# 28.cop.end     0.0000e+00   8.9443e+02   0.0000 1.0000000    
# 29.cop.end     0.0000e+00   8.9443e+02   0.0000 1.0000000    
# 30.cop.end     0.0000e+00   8.9443e+02   0.0000 1.0000000    
# 1.depart      -8.7015e+00   2.3570e-01 -36.9172 < 2.2e-16 ***
# 2.depart      -8.6456e+00   2.3570e-01 -36.6802 < 2.2e-16 ***
# 3.depart      -8.6031e+00   2.3570e-01 -36.5000 < 2.2e-16 ***
# 4.depart      -8.2131e+00   2.3570e-01 -34.8453 < 2.2e-16 ***
# 5.depart      -8.7979e+00   2.8867e-01 -30.4767 < 2.2e-16 ***
# 6.depart      -8.1073e+00   7.0710e-01 -11.4655 < 2.2e-16 ***
# 7.depart      -8.4666e+00   2.3570e-01 -35.9207 < 2.2e-16 ***
# 8.depart      -7.6245e+00   5.0000e-01 -15.2489 < 2.2e-16 ***
# 9.depart      -8.2131e+00   5.7735e-01 -14.2256 < 2.2e-16 ***
# 10.depart     -8.3367e+00   3.7796e-01 -22.0567 < 2.2e-16 ***
# 11.depart     -6.1361e+00   5.7735e-01 -10.6281 < 2.2e-16 ***
# 12.depart     -8.4306e+00   5.7735e-01 -14.6022 < 2.2e-16 ***
# 13.depart     -6.3743e+00   1.0000e+00  -6.3743 1.838e-10 ***
# 14.depart     -8.4329e+00   7.0710e-01 -11.9259 < 2.2e-16 ***
# 15.depart     -8.4725e+00   9.9999e-01  -8.4725 < 2.2e-16 ***
# 16.depart     -4.5532e+00   1.0000e+00  -4.5532 5.284e-06 ***
# 17.depart     -7.3326e+00   9.9999e-01  -7.3326 2.256e-13 ***
# 18.depart     -6.1293e+00   1.0000e+00  -6.1293 8.824e-10 ***
# 19.depart     -8.8972e+00   9.9999e-01  -8.8973 < 2.2e-16 ***
# 20.depart     -7.6762e+00   9.9999e-01  -7.6762 1.643e-14 ***
# 21.depart     -4.7117e+00   1.0000e+00  -4.7117 2.457e-06 ***
# 22.depart     -7.7354e+00   9.9999e-01  -7.7354 1.021e-14 ***
# 23.depart     -7.0172e+00   9.9999e-01  -7.0173 2.263e-12 ***
# 24.depart     -7.3915e+00   9.9999e-01  -7.3915 1.452e-13 ***
# 25.depart     -7.8422e+00   9.9999e-01  -7.8422 4.441e-15 ***
# 26.depart     -8.0356e+00   9.9999e-01  -8.0357 8.882e-16 ***
# 27.depart     -8.6597e+00   9.9999e-01  -8.6597 < 2.2e-16 ***
# 28.depart     -8.1918e+00   9.9999e-01  -8.1919 2.220e-16 ***
# 29.depart     -7.0548e+00   9.9999e-01  -7.0549 1.728e-12 ***
# 30.depart     -5.7379e+00   1.1785e-01 -48.6878 < 2.2e-16 ***
# ---
# Signif. codes:  0 ‘***’ 0.001 ‘**’ 0.01 ‘*’ 0.05 ‘.’ 0.1 ‘ ’ 1
# Null deviance: 76628.21 on 4854 degrees of freedom
# Residual deviance: 38794.2 on 4613 degrees of freedom
# 	Chi-square: 37834.01 on 241 degrees of freedom, asymptotic p-value 0 
# AIC: 39276.2 AICC: 39301.49 BIC: 40839.7 
# Log posterior: -260633.5 
# Prior parameters: mu=0 sigma=1000 nu=4 

#WAY WORSE!! 


## compare to an Attack FE and a Receive FE that lumps all the (U's + ??'s) 


#right now, male nums 1-14 are ID'd males, and then 15-29 are U's, and 30 is ??'s 

sequences<-table(event.list$Date)

#create list of dummy intercept names. This time make male 14 the baseline since I am lumping everyone else

idno<-length(event.types[grep("cop.start",event.types)])


dummy.ints.attack<-paste(c(1:14,"U"),".attackFE", sep="")
dummy.ints.receive<-paste(c(1:13,"U"), ".receiveFE", sep="")


int.names<-c(dummy.ints.attack,dummy.ints.receive,event.types[-grep("attack", event.types)])

int.list<-list()

for(i in 1:length(levels(event.list$Date))){

day.array<-array(0,dim=c(as.numeric(sequences[i]), length(event.types),length(int.names)))

#the third dimension is the stat slice.
dimnames(day.array)<-list(c(1:sequences[i]),as.character(event.types),as.character(int.names))

#loop for dummy intercepts for attacks, loop for dummy intercepts for receives, then one loop for typical intercepts

for(k in 1:length(dummy.ints.attack)){
	
	if(k<(length(dummy.ints.attack)-1)){
day.array[,startsWith(names(day.array[1,,k]), paste(k,".attack", sep="")),k]<-1
} 

if(k==(length(dummy.ints.attack))){
	
	for(h in 15:idno){
	day.array[,startsWith(names(day.array[1,,k]), paste(h,".attack", sep="")),k]<-1
}

}


}

for(k in (length(dummy.ints.attack)+1):(length(dummy.ints.attack)+length(dummy.ints.receive))){

if(k<(length(dummy.ints.attack)+length(dummy.ints.receive)-1)){
day.array[,endsWith(names(day.array[1,,k]), paste("attack.",(k-length(dummy.ints.attack)), sep="")),k]<-1
}
	
if(k==(length(dummy.ints.attack)+length(dummy.ints.receive))){
	
	for(h in 15:idno){
day.array[,endsWith(names(day.array[1,,k]), paste("attack.",h, sep="")),k]<-1
}

}


}

for(k in (length(dummy.ints.attack)+length(dummy.ints.receive)+1):length(int.names)){
	
	day.array[,int.names[k],k]<-1
	
}

#weird extra formatting to make it match the package example, seems like it becomes relevant *punintended!* if I want to fit day-specific effects

global<-list()

global[[1]]<-day.array

int.list[[i]]<-global

names(int.list[[i]])<-"global"

}


library(tictoc)

library(relevent)

tic()
intfit.sup<-rem(evlist, int.list, supplist=supplist, timing="interval", estimator="BPM")
toc()

summary(intfit.sup)

# Egocentric Relational Event Model (Interval Likelihood)
# 
#                Post.Mode     Post.SD  Z value  Pr(>|z|)    
# 1.attackFE   -4.1688e+00  6.5474e-02 -63.6719 < 2.2e-16 ***
# 2.attackFE   -4.6470e+00  8.0267e-02 -57.8945 < 2.2e-16 ***
# 3.attackFE   -4.1918e+00  6.4784e-02 -64.7045 < 2.2e-16 ***
# 4.attackFE   -3.8014e+00  6.1940e-02 -61.3713 < 2.2e-16 ***
# 5.attackFE   -4.3052e+00  9.9421e-02 -43.3030 < 2.2e-16 ***
# 6.attackFE   -5.4473e+00  7.0791e-01  -7.6949 1.421e-14 ***
# 7.attackFE   -4.5891e+00  8.3136e-02 -55.1997 < 2.2e-16 ***
# 8.attackFE   -4.7224e+00  1.9643e-01 -24.0418 < 2.2e-16 ***
# 9.attackFE   -7.0066e+00  1.0006e+00  -7.0027 2.510e-12 ***
# 10.attackFE  -4.2364e+00  8.5900e-02 -49.3184 < 2.2e-16 ***
# 11.attackFE  -1.6591e+01  2.1326e+02  -0.0778 0.9379881    
# 12.attackFE  -7.0815e+00  3.5358e-01 -20.0281 < 2.2e-16 ***
# 13.attackFE  -4.0807e+00  1.0007e+00  -4.0779 4.544e-05 ***
# 14.attackFE  -2.8558e-15  8.9443e+02   0.0000 1.0000000    
# U.attackFE   -4.7586e+00  1.0616e-01 -44.8239 < 2.2e-16 ***
# 1.receiveFE  -5.1212e+00  1.1830e-01 -43.2907 < 2.2e-16 ***
# 2.receiveFE  -4.8200e+00  9.5815e-02 -50.3059 < 2.2e-16 ***
# 3.receiveFE  -4.9938e+00  1.0652e-01 -46.8810 < 2.2e-16 ***
# 4.receiveFE  -3.8319e+00  6.8904e-02 -55.6122 < 2.2e-16 ***
# 5.receiveFE  -4.9738e+00  1.4895e-01 -33.3929 < 2.2e-16 ***
# 6.receiveFE  -3.1531e+00  2.3213e-01 -13.5838 < 2.2e-16 ***
# 7.receiveFE  -4.3228e+00  7.9148e-02 -54.6163 < 2.2e-16 ***
# 8.receiveFE  -4.5168e+00  2.0444e-01 -22.0938 < 2.2e-16 ***
# 9.receiveFE  -3.1330e+00  1.7435e-01 -17.9700 < 2.2e-16 ***
# 10.receiveFE -5.1372e+00  1.4768e-01 -34.7874 < 2.2e-16 ***
# 11.receiveFE -3.4259e+00  3.7825e-01  -9.0573 < 2.2e-16 ***
# 12.receiveFE -6.3722e+00  2.5822e-01 -24.6769 < 2.2e-16 ***
# 13.receiveFE  2.5223e-27  8.9443e+02   0.0000 1.0000000    
# U.receiveFE  -2.4819e+00  4.3892e-02 -56.5454 < 2.2e-16 ***
# 1.initFO      6.7444e+00  1.2700e-01  53.1057 < 2.2e-16 ***
# 2.initFO      6.7001e+00  1.0483e-01  63.9150 < 2.2e-16 ***
# 3.initFO      6.7387e+00  1.1471e-01  58.7464 < 2.2e-16 ***
# 4.initFO      6.6846e+00  7.5378e-02  88.6814 < 2.2e-16 ***
# 5.initFO      6.7927e+00  1.5617e-01  43.4944 < 2.2e-16 ***
# 6.initFO      5.0619e+00  5.7735e-01   8.7675 < 2.2e-16 ***
# 7.initFO      6.7157e+00  8.5436e-02  78.6058 < 2.2e-16 ***
# 8.initFO      5.1160e+00  5.0000e-01  10.2320 < 2.2e-16 ***
# 9.initFO      3.3814e+00  1.0000e+00   3.3814 0.0007212 ***
# 10.initFO     6.6626e+00  1.6667e-01  39.9758 < 2.2e-16 ***
# 11.initFO    -6.7247e+00  3.2181e+02  -0.0209 0.9833284    
# 12.initFO     5.8091e+00  4.4721e-01  12.9896 < 2.2e-16 ***
# 13.initFO    -5.0627e+00  3.6326e+02  -0.0139 0.9888802    
# 14.initFO     5.4608e+00  5.0000e-01  10.9217 < 2.2e-16 ***
# 15.initFO     6.0968e+00  5.0000e-01  12.1936 < 2.2e-16 ***
# 16.initFO    -6.8411e+00  3.1942e+02  -0.0214 0.9829127    
# 17.initFO    -5.6467e+00  3.4693e+02  -0.0163 0.9870141    
# 18.initFO    -5.0627e+00  3.6326e+02  -0.0139 0.9888802    
# 19.initFO     4.4228e+00  1.0000e+00   4.4228 9.741e-06 ***
# 20.initFO     3.5066e+00  1.0000e+00   3.5065 0.0004540 ***
# 21.initFO    -5.0627e+00  3.6326e+02  -0.0139 0.9888802    
# 22.initFO     0.0000e+00  8.9443e+02   0.0000 1.0000000    
# 23.initFO    -5.0627e+00  3.6326e+02  -0.0139 0.9888802    
# 24.initFO    -7.5530e+00  3.0584e+02  -0.0247 0.9802971    
# 25.initFO     6.2146e+00  1.0000e+00   6.2146 5.146e-10 ***
# 26.initFO     6.9078e+00  7.0711e-01   9.7690 < 2.2e-16 ***
# 27.initFO     6.0605e+00  3.3333e-01  18.1814 < 2.2e-16 ***
# 28.initFO     6.5023e+00  7.0711e-01   9.1956 < 2.2e-16 ***
# 29.initFO     6.9077e+00  1.0000e+00   6.9077 4.925e-12 ***
# 30.initFO     3.3524e+00  2.5820e-01  12.9838 < 2.2e-16 ***
# 1.initCH      2.6173e+00  1.0000e+00   2.6173 0.0088631 ** 
# 2.initCH     -9.1855e+00  2.8026e+02  -0.0328 0.9738539    
# 3.initCH     -8.9885e+00  2.8301e+02  -0.0318 0.9746631    
# 4.initCH      1.5141e+00  1.0000e+00   1.5141 0.1299940    
# 5.initCH      3.0791e+00  1.0000e+00   3.0791 0.0020762 ** 
# 6.initCH      6.7359e+00  2.5000e-01  26.9436 < 2.2e-16 ***
# 7.initCH     -9.5410e+00  2.7549e+02  -0.0346 0.9723727    
# 8.initCH      6.5629e+00  2.4254e-01  27.0596 < 2.2e-16 ***
# 9.initCH      6.8779e+00  1.7408e-01  39.5105 < 2.2e-16 ***
# 10.initCH     4.4654e+00  5.0000e-01   8.9308 < 2.2e-16 ***
# 11.initCH     6.7536e+00  4.0825e-01  16.5429 < 2.2e-16 ***
# 12.initCH    -7.3922e+00  3.0875e+02  -0.0239 0.9808986    
# 13.initCH    -5.0627e+00  3.6326e+02  -0.0139 0.9888802    
# 14.initCH     6.2718e+00  3.3333e-01  18.8153 < 2.2e-16 ***
# 15.initCH     5.4037e+00  7.0711e-01   7.6419 2.132e-14 ***
# 16.initCH     6.9078e+00  3.5355e-01  19.5381 < 2.2e-16 ***
# 17.initCH     6.2146e+00  1.0000e+00   6.2146 5.146e-10 ***
# 18.initCH     6.9077e+00  1.0000e+00   6.9077 4.925e-12 ***
# 19.initCH     5.8091e+00  5.0000e-01  11.6183 < 2.2e-16 ***
# 20.initCH     6.5976e+00  2.1320e-01  30.9455 < 2.2e-16 ***
# 21.initCH     6.9077e+00  1.0000e+00   6.9077 4.925e-12 ***
# 22.initCH     0.0000e+00  8.9443e+02   0.0000 1.0000000    
# 23.initCH    -5.0627e+00  3.6326e+02  -0.0139 0.9888802    
# 24.initCH     6.8506e+00  2.4254e-01  28.2457 < 2.2e-16 ***
# 25.initCH    -5.6467e+00  3.4693e+02  -0.0163 0.9870141    
# 26.initCH    -5.6467e+00  3.4693e+02  -0.0163 0.9870141    
# 27.initCH     6.0605e+00  3.3333e-01  18.1814 < 2.2e-16 ***
# 28.initCH    -5.9927e+00  3.3824e+02  -0.0177 0.9858644    
# 29.initCH    -5.0627e+00  3.6326e+02  -0.0139 0.9888802    
# 30.initCH     6.8244e+00  4.5502e-02 149.9810 < 2.2e-16 ***
# 1.initSF      4.9199e+00  3.1623e-01  15.5580 < 2.2e-16 ***
# 2.initSF      5.2338e+00  2.1822e-01  23.9842 < 2.2e-16 ***
# 3.initSF      5.0470e+00  2.6726e-01  18.8842 < 2.2e-16 ***
# 4.initSF      5.2753e+00  1.5250e-01  34.5926 < 2.2e-16 ***
# 5.initSF      4.4654e+00  5.0000e-01   8.9308 < 2.2e-16 ***
# 6.initSF     -7.6008e+00  3.0499e+02  -0.0249 0.9801173    
# 7.initSF      5.1631e+00  1.8570e-01  27.8039 < 2.2e-16 ***
# 8.initSF      4.8283e+00  5.7735e-01   8.3629 < 2.2e-16 ***
# 9.initSF     -8.1170e+00  2.9623e+02  -0.0274 0.9781395    
# 10.initSF     4.8709e+00  4.0825e-01  11.9311 < 2.2e-16 ***
# 11.initSF     4.9618e+00  1.0000e+00   4.9618 6.983e-07 ***
# 12.initSF     6.5023e+00  3.1623e-01  20.5620 < 2.2e-16 ***
# 13.initSF     6.9077e+00  1.0000e+00   6.9077 4.925e-12 ***
# 14.initSF     5.4608e+00  5.0000e-01  10.9217 < 2.2e-16 ***
# 15.initSF     5.8091e+00  5.7735e-01  10.0617 < 2.2e-16 ***
# 16.initSF    -6.8411e+00  3.1942e+02  -0.0214 0.9829127    
# 17.initSF     6.2146e+00  1.0000e+00   6.2146 5.146e-10 ***
# 18.initSF    -5.0627e+00  3.6326e+02  -0.0139 0.9888802    
# 19.initSF     6.3688e+00  3.7796e-01  16.8501 < 2.2e-16 ***
# 20.initSF     5.4525e+00  3.7796e-01  14.4259 < 2.2e-16 ***
# 21.initSF    -5.0627e+00  3.6326e+02  -0.0139 0.9888802    
# 22.initSF     0.0000e+00  8.9443e+02   0.0000 1.0000000    
# 23.initSF     6.9077e+00  1.0000e+00   6.9077 4.925e-12 ***
# 24.initSF     4.0174e+00  1.0000e+00   4.0174 5.885e-05 ***
# 25.initSF     6.2146e+00  1.0000e+00   6.2146 5.146e-10 ***
# 26.initSF    -5.6467e+00  3.4693e+02  -0.0163 0.9870141    
# 27.initSF     4.9618e+00  5.7735e-01   8.5942 < 2.2e-16 ***
# 28.initSF     5.8091e+00  1.0000e+00   5.8091 6.280e-09 ***
# 29.initSF    -5.0627e+00  3.6326e+02  -0.0139 0.9888802    
# 30.initSF     3.9402e+00  1.9245e-01  20.4738 < 2.2e-16 ***
# 1.disengage  -4.2623e+00  6.2622e-02 -68.0632 < 2.2e-16 ***
# 2.disengage  -5.1488e+00  8.8736e-02 -58.0237 < 2.2e-16 ***
# 3.disengage  -4.7750e+00  6.3628e-02 -75.0452 < 2.2e-16 ***
# 4.disengage  -4.6475e+00  5.4882e-02 -84.6806 < 2.2e-16 ***
# 5.disengage  -4.5737e+00  1.0260e-01 -44.5791 < 2.2e-16 ***
# 6.disengage  -6.4778e+00  5.7735e-01 -11.2199 < 2.2e-16 ***
# 7.disengage  -5.6832e+00  8.9443e-02 -63.5399 < 2.2e-16 ***
# 8.disengage  -3.0256e+00  1.8898e-01 -16.0101 < 2.2e-16 ***
# 9.disengage  -2.9514e+00  1.0000e+00  -2.9514 0.0031637 ** 
# 10.disengage -4.6109e+00  8.8045e-02 -52.3701 < 2.2e-16 ***
# 11.disengage -1.2519e+00  1.0000e+00  -1.2519 0.2106042    
# 12.disengage -5.1000e+00  3.3333e-01 -15.2999 < 2.2e-16 ***
# 13.disengage -3.0695e+00  1.0000e+00  -3.0695 0.0021442 ** 
# 14.disengage -3.9996e+00  2.0000e-01 -19.9978 < 2.2e-16 ***
# 15.disengage -3.7116e+00  3.7796e-01  -9.8199 < 2.2e-16 ***
# 16.disengage  0.0000e+00  8.9443e+02   0.0000 1.0000000    
# 17.disengage -5.5855e+00  7.0711e-01  -7.8991 2.887e-15 ***
# 18.disengage  0.0000e+00  8.9443e+02   0.0000 1.0000000    
# 19.disengage -3.3061e+00  2.5000e-01 -13.2245 < 2.2e-16 ***
# 20.disengage -2.4505e+00  3.5355e-01  -6.9311 4.176e-12 ***
# 21.disengage  0.0000e+00  8.9443e+02   0.0000 1.0000000    
# 22.disengage -3.5144e+00  1.0000e+00  -3.5144 0.0004408 ***
# 23.disengage -1.2190e+01  2.4628e+02  -0.0495 0.9605250    
# 24.disengage -1.8919e+00  1.0000e+00  -1.8919 0.0585033 .  
# 25.disengage -2.4341e+00  2.0000e-01 -12.1703 < 2.2e-16 ***
# 26.disengage -3.8049e+00  4.0825e-01  -9.3201 < 2.2e-16 ***
# 27.disengage -5.3683e+00  3.1623e-01 -16.9760 < 2.2e-16 ***
# 28.disengage -5.0921e+00  3.5355e-01 -14.4027 < 2.2e-16 ***
# 29.disengage -5.0978e+00  4.0825e-01 -12.4870 < 2.2e-16 ***
# 30.disengage -4.8310e+00  1.6222e-01 -29.7805 < 2.2e-16 ***
# 1.cop.start  -2.1775e+01  1.8743e+02  -0.1162 0.9075119    
# 2.cop.start  -9.4466e+00  4.0825e-01 -23.1394 < 2.2e-16 ***
# 3.cop.start  -7.4311e+00  1.4907e-01 -49.8496 < 2.2e-16 ***
# 4.cop.start  -7.8395e+00  2.1822e-01 -35.9250 < 2.2e-16 ***
# 5.cop.start  -2.1335e+01  1.8927e+02  -0.1127 0.9102517    
# 6.cop.start  -1.9273e+01  1.9866e+02  -0.0970 0.9227130    
# 7.cop.start  -2.1664e+01  1.8789e+02  -0.1153 0.9082066    
# 8.cop.start  -1.9504e+01  1.9754e+02  -0.0987 0.9213473    
# 9.cop.start  -1.9226e+01  1.9889e+02  -0.0967 0.9229919    
# 10.cop.start -2.0397e+01  1.9337e+02  -0.1055 0.9159918    
# 11.cop.start -1.6297e+01  2.1507e+02  -0.0758 0.9395949    
# 12.cop.start -1.9262e+01  1.9871e+02  -0.0969 0.9227794    
# 13.cop.start -1.7125e+01  2.1010e+02  -0.0815 0.9350387    
# 14.cop.start -1.9046e+01  1.9978e+02  -0.0953 0.9240495    
# 15.cop.start -1.9060e+01  1.9971e+02  -0.0954 0.9239683    
# 16.cop.start -1.5411e+01  2.2080e+02  -0.0698 0.9443573    
# 17.cop.start -1.7846e+01  2.0604e+02  -0.0866 0.9309782    
# 18.cop.start -1.6895e+01  2.1145e+02  -0.0799 0.9363159    
# 19.cop.start -1.9198e+01  1.9903e+02  -0.0965 0.9231568    
# 20.cop.start -1.8176e+01  2.0426e+02  -0.0890 0.9290955    
# 21.cop.start -1.5559e+01  2.1980e+02  -0.0708 0.9435668    
# 22.cop.start -1.8415e+01  2.0300e+02  -0.0907 0.9277215    
# 23.cop.start -1.7543e+01  2.0772e+02  -0.0845 0.9326928    
# 24.cop.start -1.8089e+01  2.0473e+02  -0.0884 0.9295947    
# 25.cop.start -1.8516e+01  2.0247e+02  -0.0914 0.9271361    
# 26.cop.start -1.7328e+01  2.0893e+02  -0.0829 0.9339035    
# 27.cop.start -1.8831e+01  2.0086e+02  -0.0938 0.9253036    
# 28.cop.start -1.8718e+01  2.0144e+02  -0.0929 0.9259662    
# 29.cop.start  0.0000e+00  8.9443e+02   0.0000 1.0000000    
# 30.cop.start -2.0543e+01  1.9271e+02  -0.1066 0.9151063    
# 1.cop.end     0.0000e+00  8.9443e+02   0.0000 1.0000000    
# 2.cop.end    -1.6515e+00  7.0711e-01  -2.3356 0.0195106 *  
# 3.cop.end    -1.3544e+00  1.6013e-01  -8.4581 < 2.2e-16 ***
# 4.cop.end    -1.7378e+00  3.3333e-01  -5.2133 1.856e-07 ***
# 5.cop.end     0.0000e+00  8.9443e+02   0.0000 1.0000000    
# 6.cop.end     0.0000e+00  8.9443e+02   0.0000 1.0000000    
# 7.cop.end     0.0000e+00  8.9443e+02   0.0000 1.0000000    
# 8.cop.end     0.0000e+00  8.9443e+02   0.0000 1.0000000    
# 9.cop.end     0.0000e+00  8.9443e+02   0.0000 1.0000000    
# 10.cop.end    0.0000e+00  8.9443e+02   0.0000 1.0000000    
# 11.cop.end    0.0000e+00  8.9443e+02   0.0000 1.0000000    
# 12.cop.end    0.0000e+00  8.9443e+02   0.0000 1.0000000    
# 13.cop.end    0.0000e+00  8.9443e+02   0.0000 1.0000000    
# 14.cop.end    0.0000e+00  8.9443e+02   0.0000 1.0000000    
# 15.cop.end    0.0000e+00  8.9443e+02   0.0000 1.0000000    
# 16.cop.end    0.0000e+00  8.9443e+02   0.0000 1.0000000    
# 17.cop.end    0.0000e+00  8.9443e+02   0.0000 1.0000000    
# 18.cop.end    0.0000e+00  8.9443e+02   0.0000 1.0000000    
# 19.cop.end    0.0000e+00  8.9443e+02   0.0000 1.0000000    
# 20.cop.end    0.0000e+00  8.9443e+02   0.0000 1.0000000    
# 21.cop.end    0.0000e+00  8.9443e+02   0.0000 1.0000000    
# 22.cop.end    0.0000e+00  8.9443e+02   0.0000 1.0000000    
# 23.cop.end    0.0000e+00  8.9443e+02   0.0000 1.0000000    
# 24.cop.end    0.0000e+00  8.9443e+02   0.0000 1.0000000    
# 25.cop.end    0.0000e+00  8.9443e+02   0.0000 1.0000000    
# 26.cop.end    0.0000e+00  8.9443e+02   0.0000 1.0000000    
# 27.cop.end    0.0000e+00  8.9443e+02   0.0000 1.0000000    
# 28.cop.end    0.0000e+00  8.9443e+02   0.0000 1.0000000    
# 29.cop.end    0.0000e+00  8.9443e+02   0.0000 1.0000000    
# 30.cop.end    0.0000e+00  8.9443e+02   0.0000 1.0000000    
# 1.depart     -8.7015e+00  2.3570e-01 -36.9172 < 2.2e-16 ***
# 2.depart     -8.6456e+00  2.3570e-01 -36.6802 < 2.2e-16 ***
# 3.depart     -8.6031e+00  2.3570e-01 -36.5000 < 2.2e-16 ***
# 4.depart     -8.2131e+00  2.3570e-01 -34.8453 < 2.2e-16 ***
# 5.depart     -8.7979e+00  2.8867e-01 -30.4767 < 2.2e-16 ***
# 6.depart     -8.1073e+00  7.0710e-01 -11.4655 < 2.2e-16 ***
# 7.depart     -8.4666e+00  2.3570e-01 -35.9207 < 2.2e-16 ***
# 8.depart     -7.6245e+00  5.0000e-01 -15.2489 < 2.2e-16 ***
# 9.depart     -8.2131e+00  5.7735e-01 -14.2256 < 2.2e-16 ***
# 10.depart    -8.3367e+00  3.7796e-01 -22.0567 < 2.2e-16 ***
# 11.depart    -6.1361e+00  5.7735e-01 -10.6281 < 2.2e-16 ***
# 12.depart    -8.4306e+00  5.7735e-01 -14.6022 < 2.2e-16 ***
# 13.depart    -6.3743e+00  1.0000e+00  -6.3743 1.838e-10 ***
# 14.depart    -8.4329e+00  7.0710e-01 -11.9259 < 2.2e-16 ***
# 15.depart    -8.4725e+00  9.9999e-01  -8.4725 < 2.2e-16 ***
# 16.depart    -4.5532e+00  1.0000e+00  -4.5532 5.284e-06 ***
# 17.depart    -7.3326e+00  9.9999e-01  -7.3326 2.256e-13 ***
# 18.depart    -6.1293e+00  1.0000e+00  -6.1293 8.824e-10 ***
# 19.depart    -8.8972e+00  9.9999e-01  -8.8973 < 2.2e-16 ***
# 20.depart    -7.6762e+00  9.9999e-01  -7.6762 1.643e-14 ***
# 21.depart    -4.7117e+00  1.0000e+00  -4.7117 2.457e-06 ***
# 22.depart    -7.7354e+00  9.9999e-01  -7.7354 1.021e-14 ***
# 23.depart    -7.0172e+00  9.9999e-01  -7.0173 2.263e-12 ***
# 24.depart    -7.3915e+00  9.9999e-01  -7.3915 1.452e-13 ***
# 25.depart    -7.8422e+00  9.9999e-01  -7.8422 4.441e-15 ***
# 26.depart    -8.0356e+00  9.9999e-01  -8.0357 8.882e-16 ***
# 27.depart    -8.6597e+00  9.9999e-01  -8.6597 < 2.2e-16 ***
# 28.depart    -8.1918e+00  9.9999e-01  -8.1919 2.220e-16 ***
# 29.depart    -7.0548e+00  9.9999e-01  -7.0549 1.728e-12 ***
# 30.depart    -5.7379e+00  1.1785e-01 -48.6878 < 2.2e-16 ***
# ---
# Signif. codes:  0 ‘***’ 0.001 ‘**’ 0.01 ‘*’ 0.05 ‘.’ 0.1 ‘ ’ 1
# Null deviance: 76628.21 on 4854 degrees of freedom
# Residual deviance: 33406.07 on 4615 degrees of freedom
# 	Chi-square: 43222.14 on 239 degrees of freedom, asymptotic p-value 0 
# AIC: 33884.07 AICC: 33908.94 BIC: 35434.6 
# Log posterior: -255937.5 
# Prior parameters: mu=0 sigma=1000 nu=4 

#not bad, but worse than fitting a global receive rate or just individual rates


## Since there are so many males now with not a lot of data associated with them, what about just a single global attack rate? 


sequences<-table(event.list$Date)

#create list of dummy intercept names. Leave male 30.receives as the baseline- this is the receive rate for ?? males- I expect everyone else's receive rate to be less

idno<-length(event.types[grep("cop.start",event.types)])

#dummy.ints.attack<-paste(1:(idno-1),".attackFE", sep="")
#dummy.ints.receive<-paste(1:(idno-1), ".receiveFE", sep="")


int.names<-c("Global.attackFE", event.types[-grep("attack", event.types)])

int.list<-list()

for(i in 1:length(levels(event.list$Date))){

day.array<-array(0,dim=c(as.numeric(sequences[i]), length(event.types),length(int.names)))

#the third dimension is the stat slice.
dimnames(day.array)<-list(c(1:sequences[i]),as.character(event.types),as.character(int.names))

#add global attack effect, loop for dummy intercepts for attacks, then one loop for typical intercepts

day.array[,grep("attack", event.types),1]<-1

# for(k in 2:(length(dummy.ints.attack)+1)){
	
# day.array[,startsWith(names(day.array[1,,k]), paste(k-1,".attack", sep="")),k]<-1

# }


for(k in 2:length(int.names)){
	
	day.array[,int.names[k],k]<-1
	
}

#weird extra formatting to make it match the package example, seems like it becomes relevant *punintended!* if I want to fit day-specific effects

global<-list()

global[[1]]<-day.array

int.list[[i]]<-global

names(int.list[[i]])<-"global"

}

tic()
intfit.sup<-rem(evlist, int.list, supplist=supplist, timing="interval", estimator="BPM")
toc()

summary(intfit.sup)

# Egocentric Relational Event Model (Interval Likelihood)
# 
#                  Post.Mode    Post.SD   Z value  Pr(>|z|)    
# Global.attackFE  -7.729663   0.025768 -299.9667 < 2.2e-16 ***
# 1.initFO          6.744430   0.127000   53.1057 < 2.2e-16 ***
# 2.initFO          6.700116   0.104828   63.9150 < 2.2e-16 ***
# 3.initFO          6.738679   0.114708   58.7464 < 2.2e-16 ***
# 4.initFO          6.684612   0.075378   88.6814 < 2.2e-16 ***
# 5.initFO          6.792686   0.156174   43.4944 < 2.2e-16 ***
# 6.initFO          5.061926   0.577351    8.7675 < 2.2e-16 ***
# 7.initFO          6.715748   0.085436   78.6058 < 2.2e-16 ***
# 8.initFO          5.115994   0.500000   10.2320 < 2.2e-16 ***
# 9.initFO          3.381391   1.000001    3.3814 0.0007212 ***
# 10.initFO         6.662633   0.166667   39.9758 < 2.2e-16 ***
# 11.initFO        -6.724740 321.814902   -0.0209 0.9833284    
# 12.initFO         5.809142   0.447214   12.9896 < 2.2e-16 ***
# 13.initFO        -5.062715 363.256388   -0.0139 0.9888802    
# 14.initFO         5.460835   0.500000   10.9217 < 2.2e-16 ***
# 15.initFO         6.096823   0.500000   12.1936 < 2.2e-16 ***
# 16.initFO        -6.841114 319.417929   -0.0214 0.9829127    
# 17.initFO        -5.646696 346.931976   -0.0163 0.9870141    
# 18.initFO        -5.062715 363.256388   -0.0139 0.9888802    
# 19.initFO         4.422843   1.000002    4.4228 9.741e-06 ***
# 20.initFO         3.506554   1.000002    3.5065 0.0004540 ***
# 21.initFO        -5.062715 363.256388   -0.0139 0.9888802    
# 22.initFO         0.000000 894.427191    0.0000 1.0000000    
# 23.initFO        -5.062715 363.256388   -0.0139 0.9888802    
# 24.initFO        -7.553047 305.835850   -0.0247 0.9802971    
# 25.initFO         6.214600   1.000003    6.2146 5.146e-10 ***
# 26.initFO         6.907751   0.707108    9.7690 < 2.2e-16 ***
# 27.initFO         6.060457   0.333333   18.1814 < 2.2e-16 ***
# 28.initFO         6.502286   0.707108    9.1956 < 2.2e-16 ***
# 29.initFO         6.907747   1.000004    6.9077 4.925e-12 ***
# 30.initFO         3.352407   0.258199   12.9838 < 2.2e-16 ***
# 1.initCH          2.617293   1.000001    2.6173 0.0088631 ** 
# 2.initCH         -9.185505 280.258756   -0.0328 0.9738539    
# 3.initCH         -8.988496 283.008974   -0.0318 0.9746631    
# 4.initCH          1.514126   1.000000    1.5141 0.1299940    
# 5.initCH          3.079110   1.000001    3.0791 0.0020762 ** 
# 6.initCH          6.735904   0.250000   26.9436 < 2.2e-16 ***
# 7.initCH         -9.541022 275.492334   -0.0346 0.9723727    
# 8.initCH          6.562914   0.242536   27.0596 < 2.2e-16 ***
# 9.initCH          6.877902   0.174078   39.5105 < 2.2e-16 ***
# 10.initCH         4.465407   0.500000    8.9308 < 2.2e-16 ***
# 11.initCH         6.753603   0.408249   16.5429 < 2.2e-16 ***
# 12.initCH        -7.392244 308.751880   -0.0239 0.9808986    
# 13.initCH        -5.062715 363.256388   -0.0139 0.9888802    
# 14.initCH         6.271766   0.333333   18.8153 < 2.2e-16 ***
# 15.initCH         5.403675   0.707108    7.6419 2.132e-14 ***
# 16.initCH         6.907754   0.353554   19.5381 < 2.2e-16 ***
# 17.initCH         6.214600   1.000003    6.2146 5.146e-10 ***
# 18.initCH         6.907747   1.000004    6.9077 4.925e-12 ***
# 19.initCH         5.809141   0.500000   11.6183 < 2.2e-16 ***
# 20.initCH         6.597600   0.213201   30.9455 < 2.2e-16 ***
# 21.initCH         6.907747   1.000004    6.9077 4.925e-12 ***
# 22.initCH         0.000000 894.427191    0.0000 1.0000000    
# 23.initCH        -5.062715 363.256388   -0.0139 0.9888802    
# 24.initCH         6.850596   0.242536   28.2457 < 2.2e-16 ***
# 25.initCH        -5.646696 346.931976   -0.0163 0.9870141    
# 26.initCH        -5.646696 346.931976   -0.0163 0.9870141    
# 27.initCH         6.060457   0.333333   18.1814 < 2.2e-16 ***
# 28.initCH        -5.992692 338.240255   -0.0177 0.9858644    
# 29.initCH        -5.062715 363.256388   -0.0139 0.9888802    
# 30.initCH         6.824374   0.045502  149.9810 < 2.2e-16 ***
# 1.initSF          4.919880   0.316228   15.5580 < 2.2e-16 ***
# 2.initSF          5.233779   0.218218   23.9842 < 2.2e-16 ***
# 3.initSF          5.047002   0.267261   18.8842 < 2.2e-16 ***
# 4.initSF          5.275328   0.152499   34.5926 < 2.2e-16 ***
# 5.initSF          4.465407   0.500000    8.9308 < 2.2e-16 ***
# 6.initSF         -7.600810 304.985488   -0.0249 0.9801173    
# 7.initSF          5.163063   0.185695   27.8039 < 2.2e-16 ***
# 8.initSF          4.828312   0.577351    8.3629 < 2.2e-16 ***
# 9.initSF         -8.117025 296.225732   -0.0274 0.9781395    
# 10.initSF         4.870872   0.408248   11.9311 < 2.2e-16 ***
# 11.initSF         4.961839   1.000002    4.9618 6.983e-07 ***
# 12.initSF         6.502289   0.316228   20.5620 < 2.2e-16 ***
# 13.initSF         6.907747   1.000004    6.9077 4.925e-12 ***
# 14.initSF         5.460835   0.500000   10.9217 < 2.2e-16 ***
# 15.initSF         5.809141   0.577351   10.0617 < 2.2e-16 ***
# 16.initSF        -6.841114 319.417929   -0.0214 0.9829127    
# 17.initSF         6.214600   1.000003    6.2146 5.146e-10 ***
# 18.initSF        -5.062715 363.256388   -0.0139 0.9888802    
# 19.initSF         6.368758   0.377965   16.8501 < 2.2e-16 ***
# 20.initSF         5.452467   0.377965   14.4259 < 2.2e-16 ***
# 21.initSF        -5.062715 363.256388   -0.0139 0.9888802    
# 22.initSF         0.000000 894.427191    0.0000 1.0000000    
# 23.initSF         6.907747   1.000004    6.9077 4.925e-12 ***
# 24.initSF         4.017378   1.000002    4.0174 5.885e-05 ***
# 25.initSF         6.214600   1.000003    6.2146 5.146e-10 ***
# 26.initSF        -5.646696 346.931976   -0.0163 0.9870141    
# 27.initSF         4.961843   0.577351    8.5942 < 2.2e-16 ***
# 28.initSF         5.809136   1.000003    5.8091 6.280e-09 ***
# 29.initSF        -5.062715 363.256388   -0.0139 0.9888802    
# 30.initSF         3.940194   0.192450   20.4738 < 2.2e-16 ***
# 1.disengage      -4.262284   0.062622  -68.0632 < 2.2e-16 ***
# 2.disengage      -5.148770   0.088736  -58.0237 < 2.2e-16 ***
# 3.disengage      -4.775014   0.063628  -75.0452 < 2.2e-16 ***
# 4.disengage      -4.647453   0.054882  -84.6806 < 2.2e-16 ***
# 5.disengage      -4.573716   0.102598  -44.5791 < 2.2e-16 ***
# 6.disengage      -6.477786   0.577349  -11.2199 < 2.2e-16 ***
# 7.disengage      -5.683178   0.089443  -63.5399 < 2.2e-16 ***
# 8.disengage      -3.025632   0.188982  -16.0101 < 2.2e-16 ***
# 9.disengage      -2.951359   0.999998   -2.9514 0.0031637 ** 
# 10.disengage     -4.610931   0.088045  -52.3701 < 2.2e-16 ***
# 11.disengage     -1.251904   0.999999   -1.2519 0.2106042    
# 12.disengage     -5.099954   0.333333  -15.2999 < 2.2e-16 ***
# 13.disengage     -3.069490   0.999997   -3.0695 0.0021442 ** 
# 14.disengage     -3.999558   0.200000  -19.9978 < 2.2e-16 ***
# 15.disengage     -3.711562   0.377964   -9.8199 < 2.2e-16 ***
# 16.disengage      0.000000 894.427191    0.0000 1.0000000    
# 17.disengage     -5.585480   0.707105   -7.8991 2.887e-15 ***
# 18.disengage      0.000000 894.427191    0.0000 1.0000000    
# 19.disengage     -3.306133   0.250000  -13.2245 < 2.2e-16 ***
# 20.disengage     -2.450509   0.353553   -6.9311 4.176e-12 ***
# 21.disengage      0.000000 894.427191    0.0000 1.0000000    
# 22.disengage     -3.514373   0.999997   -3.5144 0.0004408 ***
# 23.disengage    -12.189754 246.283852   -0.0495 0.9605250    
# 24.disengage     -1.891904   0.999998   -1.8919 0.0585033 .  
# 25.disengage     -2.434066   0.200000  -12.1703 < 2.2e-16 ***
# 26.disengage     -3.804916   0.408248   -9.3201 < 2.2e-16 ***
# 27.disengage     -5.368294   0.316228  -16.9760 < 2.2e-16 ***
# 28.disengage     -5.092138   0.353553  -14.4027 < 2.2e-16 ***
# 29.disengage     -5.097787   0.408248  -12.4870 < 2.2e-16 ***
# 30.disengage     -4.831031   0.162221  -29.7805 < 2.2e-16 ***
# 1.cop.start     -21.775227 187.429658   -0.1162 0.9075114    
# 2.cop.start      -9.446606   0.408248  -23.1394 < 2.2e-16 ***
# 3.cop.start      -7.431142   0.149071  -49.8496 < 2.2e-16 ***
# 4.cop.start      -7.839479   0.218218  -35.9250 < 2.2e-16 ***
# 5.cop.start     -21.334707 189.269456   -0.1127 0.9102515    
# 6.cop.start     -19.273179 198.658066   -0.0970 0.9227130    
# 7.cop.start     -21.663930 187.889501   -0.1153 0.9082062    
# 8.cop.start     -19.504105 197.536439   -0.0987 0.9213473    
# 9.cop.start     -19.225865 198.890244   -0.0967 0.9229919    
# 10.cop.start    -20.397432 193.369819   -0.1055 0.9159917    
# 11.cop.start    -16.297433 215.065251   -0.0758 0.9395949    
# 12.cop.start    -19.261927 198.713209   -0.0969 0.9227794    
# 13.cop.start    -17.124683 210.100682   -0.0815 0.9350387    
# 14.cop.start    -19.045919 199.780758   -0.0953 0.9240495    
# 15.cop.start    -19.059755 199.711860   -0.0954 0.9239683    
# 16.cop.start    -15.410565 220.799295   -0.0698 0.9443573    
# 17.cop.start    -17.846074 206.041009   -0.0866 0.9309782    
# 18.cop.start    -16.894762 211.445896   -0.0799 0.9363159    
# 19.cop.start    -19.197853 199.028083   -0.0965 0.9231568    
# 20.cop.start    -18.175836 204.262032   -0.0890 0.9290955    
# 21.cop.start    -15.559441 219.804656   -0.0708 0.9435668    
# 22.cop.start    -18.414696 203.001849   -0.0907 0.9277215    
# 23.cop.start    -17.543193 207.716617   -0.0845 0.9326928    
# 24.cop.start    -18.088681 204.727721   -0.0884 0.9295947    
# 25.cop.start    -18.516001 202.474388   -0.0914 0.9271361    
# 26.cop.start    -17.327815 208.933327   -0.0829 0.9339035    
# 27.cop.start    -18.831473 200.857793   -0.0938 0.9253036    
# 28.cop.start    -18.717694 201.436363   -0.0929 0.9259662    
# 29.cop.start      0.000000 894.427191    0.0000 1.0000000    
# 30.cop.start    -20.543323 192.714083   -0.1066 0.9151063    
# 1.cop.end         0.000000 894.427191    0.0000 1.0000000    
# 2.cop.end        -1.651538   0.707106   -2.3356 0.0195106 *  
# 3.cop.end        -1.354380   0.160128   -8.4581 < 2.2e-16 ***
# 4.cop.end        -1.737753   0.333333   -5.2133 1.856e-07 ***
# 5.cop.end         0.000000 894.427191    0.0000 1.0000000    
# 6.cop.end         0.000000 894.427191    0.0000 1.0000000    
# 7.cop.end         0.000000 894.427191    0.0000 1.0000000    
# 8.cop.end         0.000000 894.427191    0.0000 1.0000000    
# 9.cop.end         0.000000 894.427191    0.0000 1.0000000    
# 10.cop.end        0.000000 894.427191    0.0000 1.0000000    
# 11.cop.end        0.000000 894.427191    0.0000 1.0000000    
# 12.cop.end        0.000000 894.427191    0.0000 1.0000000    
# 13.cop.end        0.000000 894.427191    0.0000 1.0000000    
# 14.cop.end        0.000000 894.427191    0.0000 1.0000000    
# 15.cop.end        0.000000 894.427191    0.0000 1.0000000    
# 16.cop.end        0.000000 894.427191    0.0000 1.0000000    
# 17.cop.end        0.000000 894.427191    0.0000 1.0000000    
# 18.cop.end        0.000000 894.427191    0.0000 1.0000000    
# 19.cop.end        0.000000 894.427191    0.0000 1.0000000    
# 20.cop.end        0.000000 894.427191    0.0000 1.0000000    
# 21.cop.end        0.000000 894.427191    0.0000 1.0000000    
# 22.cop.end        0.000000 894.427191    0.0000 1.0000000    
# 23.cop.end        0.000000 894.427191    0.0000 1.0000000    
# 24.cop.end        0.000000 894.427191    0.0000 1.0000000    
# 25.cop.end        0.000000 894.427191    0.0000 1.0000000    
# 26.cop.end        0.000000 894.427191    0.0000 1.0000000    
# 27.cop.end        0.000000 894.427191    0.0000 1.0000000    
# 28.cop.end        0.000000 894.427191    0.0000 1.0000000    
# 29.cop.end        0.000000 894.427191    0.0000 1.0000000    
# 30.cop.end        0.000000 894.427191    0.0000 1.0000000    
# 1.depart         -8.701475   0.235702  -36.9172 < 2.2e-16 ***
# 2.depart         -8.645592   0.235702  -36.6802 < 2.2e-16 ***
# 3.depart         -8.603133   0.235702  -36.5000 < 2.2e-16 ***
# 4.depart         -8.213108   0.235702  -34.8453 < 2.2e-16 ***
# 5.depart         -8.797853   0.288675  -30.4767 < 2.2e-16 ***
# 6.depart         -8.107312   0.707105  -11.4655 < 2.2e-16 ***
# 7.depart         -8.466583   0.235702  -35.9207 < 2.2e-16 ***
# 8.depart         -7.624461   0.499999  -15.2489 < 2.2e-16 ***
# 9.depart         -8.213118   0.577349  -14.2256 < 2.2e-16 ***
# 10.depart        -8.336652   0.377964  -22.0567 < 2.2e-16 ***
# 11.depart        -6.136106   0.577349  -10.6281 < 2.2e-16 ***
# 12.depart        -8.430566   0.577349  -14.6022 < 2.2e-16 ***
# 13.depart        -6.374262   0.999995   -6.3743 1.838e-10 ***
# 14.depart        -8.432862   0.707105  -11.9259 < 2.2e-16 ***
# 15.depart        -8.472487   0.999994   -8.4725 < 2.2e-16 ***
# 16.depart        -4.553187   0.999997   -4.5532 5.284e-06 ***
# 17.depart        -7.332594   0.999995   -7.3326 2.256e-13 ***
# 18.depart        -6.129319   0.999996   -6.1293 8.824e-10 ***
# 19.depart        -8.897214   0.999994   -8.8973 < 2.2e-16 ***
# 20.depart        -7.676151   0.999995   -7.6762 1.643e-14 ***
# 21.depart        -4.711675   0.999996   -4.7117 2.457e-06 ***
# 22.depart        -7.735384   0.999995   -7.7354 1.021e-14 ***
# 23.depart        -7.017221   0.999995   -7.0173 2.263e-12 ***
# 24.depart        -7.391510   0.999995   -7.3915 1.452e-13 ***
# 25.depart        -7.842174   0.999994   -7.8422 4.441e-15 ***
# 26.depart        -8.035618   0.999994   -8.0357 8.882e-16 ***
# 27.depart        -8.659685   0.999994   -8.6597 < 2.2e-16 ***
# 28.depart        -8.191807   0.999994   -8.1919 2.220e-16 ***
# 29.depart        -7.054816   0.999995   -7.0549 1.728e-12 ***
# 30.depart        -5.737913   0.117851  -48.6878 < 2.2e-16 ***
# ---
# Signif. codes:  0 ‘***’ 0.001 ‘**’ 0.01 ‘*’ 0.05 ‘.’ 0.1 ‘ ’ 1
# Null deviance: 76628.21 on 4854 degrees of freedom
# Residual deviance: 31906.13 on 4643 degrees of freedom
# 	Chi-square: 44722.09 on 211 degrees of freedom, asymptotic p-value 0 
# AIC: 32328.13 AICC: 32347.4 BIC: 33697 
# Log posterior: -227160 
# Prior parameters: mu=0 sigma=1000 nu=4 


## How about single global attack + ??FE for attacks and receives?


sequences<-table(event.list$Date)

#create list of dummy intercept names. Leave male 30.receives as the baseline- this is the receive rate for ?? males- I expect everyone else's receive rate to be less

idno<-length(event.types[grep("cop.start",event.types)])

#dummy.ints.attack<-paste(1:(idno-1),".attackFE", sep="")
#dummy.ints.receive<-paste(1:(idno-1), ".receiveFE", sep="")


int.names<-c("Global.attackFE", "X.attackFE", "X.receiveFE", event.types[-grep("attack", event.types)])

int.list<-list()

for(i in 1:length(levels(event.list$Date))){

day.array<-array(0,dim=c(as.numeric(sequences[i]), length(event.types),length(int.names)))

#the third dimension is the stat slice.
dimnames(day.array)<-list(c(1:sequences[i]),as.character(event.types),as.character(int.names))

#add global attack effect, set dummy "U" intercepts for attacks and receives, then one loop for typical intercepts

day.array[,grep("attack", event.types),1]<-1

day.array[,startsWith(names(day.array[1,,2]), paste(30,".attack", sep="")),2]<-1

day.array[,endsWith(names(day.array[1,,3]), paste("attack.",30, sep="")),3]<-1


for(k in 4:length(int.names)){
	
	day.array[,int.names[k],k]<-1
	
}

#weird extra formatting to make it match the package example, seems like it becomes relevant *punintended!* if I want to fit day-specific effects

global<-list()

global[[1]]<-day.array

int.list[[i]]<-global

names(int.list[[i]])<-"global"

}

tic()
intfit.sup<-rem(evlist, int.list, supplist=supplist, timing="interval", estimator="BPM")
toc()

summary(intfit.sup)


# Egocentric Relational Event Model (Interval Likelihood)
# 
#                  Post.Mode    Post.SD   Z value  Pr(>|z|)    
# Global.attackFE  -8.069659   0.032125 -251.1985 < 2.2e-16 ***
# X.attackFE       -1.456264   0.290457   -5.0137 5.339e-07 ***
# X.receiveFE       2.322755   0.054192   42.8617 < 2.2e-16 ***
# 1.initFO          6.744430   0.127000   53.1057 < 2.2e-16 ***
# 2.initFO          6.700116   0.104828   63.9150 < 2.2e-16 ***
# 3.initFO          6.738679   0.114708   58.7464 < 2.2e-16 ***
# 4.initFO          6.684612   0.075378   88.6814 < 2.2e-16 ***
# 5.initFO          6.792686   0.156174   43.4944 < 2.2e-16 ***
# 6.initFO          5.061926   0.577351    8.7675 < 2.2e-16 ***
# 7.initFO          6.715748   0.085436   78.6058 < 2.2e-16 ***
# 8.initFO          5.115994   0.500000   10.2320 < 2.2e-16 ***
# 9.initFO          3.381391   1.000001    3.3814 0.0007212 ***
# 10.initFO         6.662633   0.166667   39.9758 < 2.2e-16 ***
# 11.initFO        -6.724740 321.814902   -0.0209 0.9833284    
# 12.initFO         5.809142   0.447214   12.9896 < 2.2e-16 ***
# 13.initFO        -5.062715 363.256388   -0.0139 0.9888802    
# 14.initFO         5.460835   0.500000   10.9217 < 2.2e-16 ***
# 15.initFO         6.096823   0.500000   12.1936 < 2.2e-16 ***
# 16.initFO        -6.841114 319.417929   -0.0214 0.9829127    
# 17.initFO        -5.646696 346.931976   -0.0163 0.9870141    
# 18.initFO        -5.062715 363.256388   -0.0139 0.9888802    
# 19.initFO         4.422843   1.000002    4.4228 9.741e-06 ***
# 20.initFO         3.506554   1.000002    3.5065 0.0004540 ***
# 21.initFO        -5.062715 363.256388   -0.0139 0.9888802    
# 22.initFO         0.000000 894.427191    0.0000 1.0000000    
# 23.initFO        -5.062715 363.256388   -0.0139 0.9888802    
# 24.initFO        -7.553047 305.835850   -0.0247 0.9802971    
# 25.initFO         6.214600   1.000003    6.2146 5.146e-10 ***
# 26.initFO         6.907751   0.707108    9.7690 < 2.2e-16 ***
# 27.initFO         6.060457   0.333333   18.1814 < 2.2e-16 ***
# 28.initFO         6.502286   0.707108    9.1956 < 2.2e-16 ***
# 29.initFO         6.907747   1.000004    6.9077 4.925e-12 ***
# 30.initFO         3.352407   0.258199   12.9838 < 2.2e-16 ***
# 1.initCH          2.617293   1.000001    2.6173 0.0088631 ** 
# 2.initCH         -9.185505 280.258756   -0.0328 0.9738539    
# 3.initCH         -8.988496 283.008974   -0.0318 0.9746631    
# 4.initCH          1.514126   1.000000    1.5141 0.1299940    
# 5.initCH          3.079110   1.000001    3.0791 0.0020762 ** 
# 6.initCH          6.735904   0.250000   26.9436 < 2.2e-16 ***
# 7.initCH         -9.541022 275.492334   -0.0346 0.9723727    
# 8.initCH          6.562914   0.242536   27.0596 < 2.2e-16 ***
# 9.initCH          6.877902   0.174078   39.5105 < 2.2e-16 ***
# 10.initCH         4.465407   0.500000    8.9308 < 2.2e-16 ***
# 11.initCH         6.753603   0.408249   16.5429 < 2.2e-16 ***
# 12.initCH        -7.392244 308.751880   -0.0239 0.9808986    
# 13.initCH        -5.062715 363.256388   -0.0139 0.9888802    
# 14.initCH         6.271766   0.333333   18.8153 < 2.2e-16 ***
# 15.initCH         5.403675   0.707108    7.6419 2.132e-14 ***
# 16.initCH         6.907754   0.353554   19.5381 < 2.2e-16 ***
# 17.initCH         6.214600   1.000003    6.2146 5.146e-10 ***
# 18.initCH         6.907747   1.000004    6.9077 4.925e-12 ***
# 19.initCH         5.809141   0.500000   11.6183 < 2.2e-16 ***
# 20.initCH         6.597600   0.213201   30.9455 < 2.2e-16 ***
# 21.initCH         6.907747   1.000004    6.9077 4.925e-12 ***
# 22.initCH         0.000000 894.427191    0.0000 1.0000000    
# 23.initCH        -5.062715 363.256388   -0.0139 0.9888802    
# 24.initCH         6.850596   0.242536   28.2457 < 2.2e-16 ***
# 25.initCH        -5.646696 346.931976   -0.0163 0.9870141    
# 26.initCH        -5.646696 346.931976   -0.0163 0.9870141    
# 27.initCH         6.060457   0.333333   18.1814 < 2.2e-16 ***
# 28.initCH        -5.992692 338.240255   -0.0177 0.9858644    
# 29.initCH        -5.062715 363.256388   -0.0139 0.9888802    
# 30.initCH         6.824374   0.045502  149.9810 < 2.2e-16 ***
# 1.initSF          4.919880   0.316228   15.5580 < 2.2e-16 ***
# 2.initSF          5.233779   0.218218   23.9842 < 2.2e-16 ***
# 3.initSF          5.047002   0.267261   18.8842 < 2.2e-16 ***
# 4.initSF          5.275328   0.152499   34.5926 < 2.2e-16 ***
# 5.initSF          4.465407   0.500000    8.9308 < 2.2e-16 ***
# 6.initSF         -7.600810 304.985488   -0.0249 0.9801173    
# 7.initSF          5.163063   0.185695   27.8039 < 2.2e-16 ***
# 8.initSF          4.828312   0.577351    8.3629 < 2.2e-16 ***
# 9.initSF         -8.117025 296.225732   -0.0274 0.9781395    
# 10.initSF         4.870872   0.408248   11.9311 < 2.2e-16 ***
# 11.initSF         4.961839   1.000002    4.9618 6.983e-07 ***
# 12.initSF         6.502289   0.316228   20.5620 < 2.2e-16 ***
# 13.initSF         6.907747   1.000004    6.9077 4.925e-12 ***
# 14.initSF         5.460835   0.500000   10.9217 < 2.2e-16 ***
# 15.initSF         5.809141   0.577351   10.0617 < 2.2e-16 ***
# 16.initSF        -6.841114 319.417929   -0.0214 0.9829127    
# 17.initSF         6.214600   1.000003    6.2146 5.146e-10 ***
# 18.initSF        -5.062715 363.256388   -0.0139 0.9888802    
# 19.initSF         6.368758   0.377965   16.8501 < 2.2e-16 ***
# 20.initSF         5.452467   0.377965   14.4259 < 2.2e-16 ***
# 21.initSF        -5.062715 363.256388   -0.0139 0.9888802    
# 22.initSF         0.000000 894.427191    0.0000 1.0000000    
# 23.initSF         6.907747   1.000004    6.9077 4.925e-12 ***
# 24.initSF         4.017378   1.000002    4.0174 5.885e-05 ***
# 25.initSF         6.214600   1.000003    6.2146 5.146e-10 ***
# 26.initSF        -5.646696 346.931976   -0.0163 0.9870141    
# 27.initSF         4.961843   0.577351    8.5942 < 2.2e-16 ***
# 28.initSF         5.809136   1.000003    5.8091 6.280e-09 ***
# 29.initSF        -5.062715 363.256388   -0.0139 0.9888802    
# 30.initSF         3.940194   0.192450   20.4738 < 2.2e-16 ***
# 1.disengage      -4.262284   0.062622  -68.0632 < 2.2e-16 ***
# 2.disengage      -5.148770   0.088736  -58.0237 < 2.2e-16 ***
# 3.disengage      -4.775014   0.063628  -75.0452 < 2.2e-16 ***
# 4.disengage      -4.647453   0.054882  -84.6806 < 2.2e-16 ***
# 5.disengage      -4.573716   0.102598  -44.5791 < 2.2e-16 ***
# 6.disengage      -6.477786   0.577349  -11.2199 < 2.2e-16 ***
# 7.disengage      -5.683178   0.089443  -63.5399 < 2.2e-16 ***
# 8.disengage      -3.025632   0.188982  -16.0101 < 2.2e-16 ***
# 9.disengage      -2.951359   0.999998   -2.9514 0.0031637 ** 
# 10.disengage     -4.610931   0.088045  -52.3701 < 2.2e-16 ***
# 11.disengage     -1.251904   0.999999   -1.2519 0.2106042    
# 12.disengage     -5.099954   0.333333  -15.2999 < 2.2e-16 ***
# 13.disengage     -3.069490   0.999997   -3.0695 0.0021442 ** 
# 14.disengage     -3.999558   0.200000  -19.9978 < 2.2e-16 ***
# 15.disengage     -3.711562   0.377964   -9.8199 < 2.2e-16 ***
# 16.disengage      0.000000 894.427191    0.0000 1.0000000    
# 17.disengage     -5.585480   0.707105   -7.8991 2.887e-15 ***
# 18.disengage      0.000000 894.427191    0.0000 1.0000000    
# 19.disengage     -3.306133   0.250000  -13.2245 < 2.2e-16 ***
# 20.disengage     -2.450509   0.353553   -6.9311 4.176e-12 ***
# 21.disengage      0.000000 894.427191    0.0000 1.0000000    
# 22.disengage     -3.514373   0.999997   -3.5144 0.0004408 ***
# 23.disengage    -12.189754 246.283852   -0.0495 0.9605250    
# 24.disengage     -1.891904   0.999998   -1.8919 0.0585033 .  
# 25.disengage     -2.434066   0.200000  -12.1703 < 2.2e-16 ***
# 26.disengage     -3.804916   0.408248   -9.3201 < 2.2e-16 ***
# 27.disengage     -5.368294   0.316228  -16.9760 < 2.2e-16 ***
# 28.disengage     -5.092138   0.353553  -14.4027 < 2.2e-16 ***
# 29.disengage     -5.097787   0.408248  -12.4870 < 2.2e-16 ***
# 30.disengage     -4.831031   0.162221  -29.7805 < 2.2e-16 ***
# 1.cop.start     -21.775226 187.429632   -0.1162 0.9075114    
# 2.cop.start      -9.446606   0.408248  -23.1394 < 2.2e-16 ***
# 3.cop.start      -7.431142   0.149071  -49.8496 < 2.2e-16 ***
# 4.cop.start      -7.839479   0.218218  -35.9250 < 2.2e-16 ***
# 5.cop.start     -21.334707 189.269443   -0.1127 0.9102515    
# 6.cop.start     -19.273179 198.658065   -0.0970 0.9227130    
# 7.cop.start     -21.663929 187.889479   -0.1153 0.9082062    
# 8.cop.start     -19.504105 197.536439   -0.0987 0.9213473    
# 9.cop.start     -19.225865 198.890243   -0.0967 0.9229919    
# 10.cop.start    -20.397432 193.369817   -0.1055 0.9159917    
# 11.cop.start    -16.297433 215.065251   -0.0758 0.9395949    
# 12.cop.start    -19.261927 198.713208   -0.0969 0.9227794    
# 13.cop.start    -17.124683 210.100682   -0.0815 0.9350387    
# 14.cop.start    -19.045919 199.780758   -0.0953 0.9240495    
# 15.cop.start    -19.059755 199.711860   -0.0954 0.9239683    
# 16.cop.start    -15.410565 220.799295   -0.0698 0.9443573    
# 17.cop.start    -17.846074 206.041009   -0.0866 0.9309782    
# 18.cop.start    -16.894762 211.445896   -0.0799 0.9363159    
# 19.cop.start    -19.197853 199.028083   -0.0965 0.9231568    
# 20.cop.start    -18.175836 204.262032   -0.0890 0.9290955    
# 21.cop.start    -15.559441 219.804656   -0.0708 0.9435668    
# 22.cop.start    -18.414696 203.001849   -0.0907 0.9277215    
# 23.cop.start    -17.543193 207.716617   -0.0845 0.9326928    
# 24.cop.start    -18.088681 204.727721   -0.0884 0.9295947    
# 25.cop.start    -18.516001 202.474388   -0.0914 0.9271361    
# 26.cop.start    -17.327815 208.933327   -0.0829 0.9339035    
# 27.cop.start    -18.831473 200.857793   -0.0938 0.9253036    
# 28.cop.start    -18.717694 201.436362   -0.0929 0.9259662    
# 29.cop.start      0.000000 894.427191    0.0000 1.0000000    
# 30.cop.start    -20.543323 192.714080   -0.1066 0.9151063    
# 1.cop.end         0.000000 894.427191    0.0000 1.0000000    
# 2.cop.end        -1.651538   0.707106   -2.3356 0.0195106 *  
# 3.cop.end        -1.354380   0.160128   -8.4581 < 2.2e-16 ***
# 4.cop.end        -1.737753   0.333333   -5.2133 1.856e-07 ***
# 5.cop.end         0.000000 894.427191    0.0000 1.0000000    
# 6.cop.end         0.000000 894.427191    0.0000 1.0000000    
# 7.cop.end         0.000000 894.427191    0.0000 1.0000000    
# 8.cop.end         0.000000 894.427191    0.0000 1.0000000    
# 9.cop.end         0.000000 894.427191    0.0000 1.0000000    
# 10.cop.end        0.000000 894.427191    0.0000 1.0000000    
# 11.cop.end        0.000000 894.427191    0.0000 1.0000000    
# 12.cop.end        0.000000 894.427191    0.0000 1.0000000    
# 13.cop.end        0.000000 894.427191    0.0000 1.0000000    
# 14.cop.end        0.000000 894.427191    0.0000 1.0000000    
# 15.cop.end        0.000000 894.427191    0.0000 1.0000000    
# 16.cop.end        0.000000 894.427191    0.0000 1.0000000    
# 17.cop.end        0.000000 894.427191    0.0000 1.0000000    
# 18.cop.end        0.000000 894.427191    0.0000 1.0000000    
# 19.cop.end        0.000000 894.427191    0.0000 1.0000000    
# 20.cop.end        0.000000 894.427191    0.0000 1.0000000    
# 21.cop.end        0.000000 894.427191    0.0000 1.0000000    
# 22.cop.end        0.000000 894.427191    0.0000 1.0000000    
# 23.cop.end        0.000000 894.427191    0.0000 1.0000000    
# 24.cop.end        0.000000 894.427191    0.0000 1.0000000    
# 25.cop.end        0.000000 894.427191    0.0000 1.0000000    
# 26.cop.end        0.000000 894.427191    0.0000 1.0000000    
# 27.cop.end        0.000000 894.427191    0.0000 1.0000000    
# 28.cop.end        0.000000 894.427191    0.0000 1.0000000    
# 29.cop.end        0.000000 894.427191    0.0000 1.0000000    
# 30.cop.end        0.000000 894.427191    0.0000 1.0000000    
# 1.depart         -8.701475   0.235702  -36.9172 < 2.2e-16 ***
# 2.depart         -8.645592   0.235702  -36.6802 < 2.2e-16 ***
# 3.depart         -8.603133   0.235702  -36.5000 < 2.2e-16 ***
# 4.depart         -8.213108   0.235702  -34.8453 < 2.2e-16 ***
# 5.depart         -8.797853   0.288675  -30.4767 < 2.2e-16 ***
# 6.depart         -8.107312   0.707105  -11.4655 < 2.2e-16 ***
# 7.depart         -8.466583   0.235702  -35.9207 < 2.2e-16 ***
# 8.depart         -7.624461   0.499999  -15.2489 < 2.2e-16 ***
# 9.depart         -8.213118   0.577349  -14.2256 < 2.2e-16 ***
# 10.depart        -8.336652   0.377964  -22.0567 < 2.2e-16 ***
# 11.depart        -6.136106   0.577349  -10.6281 < 2.2e-16 ***
# 12.depart        -8.430566   0.577349  -14.6022 < 2.2e-16 ***
# 13.depart        -6.374262   0.999995   -6.3743 1.838e-10 ***
# 14.depart        -8.432862   0.707105  -11.9259 < 2.2e-16 ***
# 15.depart        -8.472487   0.999994   -8.4725 < 2.2e-16 ***
# 16.depart        -4.553187   0.999997   -4.5532 5.284e-06 ***
# 17.depart        -7.332594   0.999995   -7.3326 2.256e-13 ***
# 18.depart        -6.129319   0.999996   -6.1293 8.824e-10 ***
# 19.depart        -8.897214   0.999994   -8.8973 < 2.2e-16 ***
# 20.depart        -7.676151   0.999995   -7.6762 1.643e-14 ***
# 21.depart        -4.711675   0.999996   -4.7117 2.457e-06 ***
# 22.depart        -7.735384   0.999995   -7.7354 1.021e-14 ***
# 23.depart        -7.017221   0.999995   -7.0173 2.263e-12 ***
# 24.depart        -7.391510   0.999995   -7.3915 1.452e-13 ***
# 25.depart        -7.842174   0.999994   -7.8422 4.441e-15 ***
# 26.depart        -8.035618   0.999994   -8.0357 8.882e-16 ***
# 27.depart        -8.659685   0.999994   -8.6597 < 2.2e-16 ***
# 28.depart        -8.191807   0.999994   -8.1919 2.220e-16 ***
# 29.depart        -7.054816   0.999995   -7.0549 1.728e-12 ***
# 30.depart        -5.737913   0.117851  -48.6878 < 2.2e-16 ***
# ---
# Signif. codes:  0 ‘***’ 0.001 ‘**’ 0.01 ‘*’ 0.05 ‘.’ 0.1 ‘ ’ 1
# Null deviance: 76628.21 on 4854 degrees of freedom
# Residual deviance: 30526.25 on 4641 degrees of freedom
# 	Chi-square: 46101.96 on 213 degrees of freedom, asymptotic p-value 0 
# AIC: 30952.25 AICC: 30971.9 BIC: 32334.1 
# Log posterior: -228472.1 
# Prior parameters: mu=0 sigma=1000 nu=4 

#getting close, but still slightly worse than the individual effects 


### Try partitioning the rates into attacks/receive vs non-transients and vs. other transients


sequences<-table(event.list$Date)

#create list of dummy intercept names. Leave male 30.receives as the baseline- this is the receive rate for ?? males- I expect everyone else's receive rate to be less

idno<-length(event.types[grep("cop.start",event.types)])

#dummy.ints.attack<-paste(1:(idno-1),".attackFE", sep="")
#dummy.ints.receive<-paste(1:(idno-1), ".receiveFE", sep="")


int.names<-c("Global.attackFE", "X.attackFE", "X.receiveFE", "X.UattackFE", "X.UreceiveFE", event.types[-grep("attack", event.types)])

int.list<-list()

for(i in 1:length(levels(event.list$Date))){

day.array<-array(0,dim=c(as.numeric(sequences[i]), length(event.types),length(int.names)))

#the third dimension is the stat slice.
dimnames(day.array)<-list(c(1:sequences[i]),as.character(event.types),as.character(int.names))

#add global attack effect, loops for "U" intercepts for attacks and receives, then one loop for typical intercepts

day.array[,grep("attack", event.types),1]<-1

for(h in 1:14){
day.array[,paste(30,".attack.", h, sep=""),2]<-1
}

for(h in 1:14){
day.array[,paste(h,".attack.",30, sep=""),3]<-1
}

for(h in 15:29){
day.array[,paste(30,".attack.", h, sep=""),4]<-1
}

for(h in 15:29){
day.array[,paste(h,".attack.",30, sep=""),5]<-1
}


for(k in 6:length(int.names)){
	
	day.array[,int.names[k],k]<-1
	
}

#weird extra formatting to make it match the package example, seems like it becomes relevant *punintended!* if I want to fit day-specific effects

global<-list()

global[[1]]<-day.array

int.list[[i]]<-global

names(int.list[[i]])<-"global"

}

tic()
intfit.sup<-rem(evlist, int.list, supplist=supplist, timing="interval", estimator="BPM")
toc()

summary(intfit.sup)


# Egocentric Relational Event Model (Interval Likelihood)
# 
#                  Post.Mode    Post.SD   Z value  Pr(>|z|)    
# Global.attackFE  -8.069659   0.032125 -251.1985 < 2.2e-16 ***
# X.attackFE       -1.581459   0.317855   -4.9754 6.511e-07 ***
# X.receiveFE       2.314992   0.055358   41.8185 < 2.2e-16 ***
# X.UattackFE      -0.357100   0.707836   -0.5045 0.6139136    
# X.UreceiveFE      2.446260   0.177017   13.8194 < 2.2e-16 ***
# 1.initFO          6.744430   0.127000   53.1057 < 2.2e-16 ***
# 2.initFO          6.700116   0.104828   63.9150 < 2.2e-16 ***
# 3.initFO          6.738679   0.114708   58.7464 < 2.2e-16 ***
# 4.initFO          6.684612   0.075378   88.6814 < 2.2e-16 ***
# 5.initFO          6.792686   0.156174   43.4944 < 2.2e-16 ***
# 6.initFO          5.061926   0.577351    8.7675 < 2.2e-16 ***
# 7.initFO          6.715748   0.085436   78.6058 < 2.2e-16 ***
# 8.initFO          5.115994   0.500000   10.2320 < 2.2e-16 ***
# 9.initFO          3.381391   1.000001    3.3814 0.0007212 ***
# 10.initFO         6.662633   0.166667   39.9758 < 2.2e-16 ***
# 11.initFO        -6.724740 321.814902   -0.0209 0.9833284    
# 12.initFO         5.809142   0.447214   12.9896 < 2.2e-16 ***
# 13.initFO        -5.062715 363.256388   -0.0139 0.9888802    
# 14.initFO         5.460835   0.500000   10.9217 < 2.2e-16 ***
# 15.initFO         6.096823   0.500000   12.1936 < 2.2e-16 ***
# 16.initFO        -6.841114 319.417929   -0.0214 0.9829127    
# 17.initFO        -5.646696 346.931976   -0.0163 0.9870141    
# 18.initFO        -5.062715 363.256388   -0.0139 0.9888802    
# 19.initFO         4.422843   1.000002    4.4228 9.741e-06 ***
# 20.initFO         3.506554   1.000002    3.5065 0.0004540 ***
# 21.initFO        -5.062715 363.256388   -0.0139 0.9888802    
# 22.initFO         0.000000 894.427191    0.0000 1.0000000    
# 23.initFO        -5.062715 363.256388   -0.0139 0.9888802    
# 24.initFO        -7.553047 305.835850   -0.0247 0.9802971    
# 25.initFO         6.214600   1.000003    6.2146 5.146e-10 ***
# 26.initFO         6.907751   0.707108    9.7690 < 2.2e-16 ***
# 27.initFO         6.060457   0.333333   18.1814 < 2.2e-16 ***
# 28.initFO         6.502286   0.707108    9.1956 < 2.2e-16 ***
# 29.initFO         6.907747   1.000004    6.9077 4.925e-12 ***
# 30.initFO         3.352407   0.258199   12.9838 < 2.2e-16 ***
# 1.initCH          2.617293   1.000001    2.6173 0.0088631 ** 
# 2.initCH         -9.185505 280.258756   -0.0328 0.9738539    
# 3.initCH         -8.988496 283.008974   -0.0318 0.9746631    
# 4.initCH          1.514126   1.000000    1.5141 0.1299940    
# 5.initCH          3.079110   1.000001    3.0791 0.0020762 ** 
# 6.initCH          6.735904   0.250000   26.9436 < 2.2e-16 ***
# 7.initCH         -9.541022 275.492334   -0.0346 0.9723727    
# 8.initCH          6.562914   0.242536   27.0596 < 2.2e-16 ***
# 9.initCH          6.877902   0.174078   39.5105 < 2.2e-16 ***
# 10.initCH         4.465407   0.500000    8.9308 < 2.2e-16 ***
# 11.initCH         6.753603   0.408249   16.5429 < 2.2e-16 ***
# 12.initCH        -7.392244 308.751880   -0.0239 0.9808986    
# 13.initCH        -5.062715 363.256388   -0.0139 0.9888802    
# 14.initCH         6.271766   0.333333   18.8153 < 2.2e-16 ***
# 15.initCH         5.403675   0.707108    7.6419 2.132e-14 ***
# 16.initCH         6.907754   0.353554   19.5381 < 2.2e-16 ***
# 17.initCH         6.214600   1.000003    6.2146 5.146e-10 ***
# 18.initCH         6.907747   1.000004    6.9077 4.925e-12 ***
# 19.initCH         5.809141   0.500000   11.6183 < 2.2e-16 ***
# 20.initCH         6.597600   0.213201   30.9455 < 2.2e-16 ***
# 21.initCH         6.907747   1.000004    6.9077 4.925e-12 ***
# 22.initCH         0.000000 894.427191    0.0000 1.0000000    
# 23.initCH        -5.062715 363.256388   -0.0139 0.9888802    
# 24.initCH         6.850596   0.242536   28.2457 < 2.2e-16 ***
# 25.initCH        -5.646696 346.931976   -0.0163 0.9870141    
# 26.initCH        -5.646696 346.931976   -0.0163 0.9870141    
# 27.initCH         6.060457   0.333333   18.1814 < 2.2e-16 ***
# 28.initCH        -5.992692 338.240255   -0.0177 0.9858644    
# 29.initCH        -5.062715 363.256388   -0.0139 0.9888802    
# 30.initCH         6.824374   0.045502  149.9810 < 2.2e-16 ***
# 1.initSF          4.919880   0.316228   15.5580 < 2.2e-16 ***
# 2.initSF          5.233779   0.218218   23.9842 < 2.2e-16 ***
# 3.initSF          5.047002   0.267261   18.8842 < 2.2e-16 ***
# 4.initSF          5.275328   0.152499   34.5926 < 2.2e-16 ***
# 5.initSF          4.465407   0.500000    8.9308 < 2.2e-16 ***
# 6.initSF         -7.600810 304.985488   -0.0249 0.9801173    
# 7.initSF          5.163063   0.185695   27.8039 < 2.2e-16 ***
# 8.initSF          4.828312   0.577351    8.3629 < 2.2e-16 ***
# 9.initSF         -8.117025 296.225732   -0.0274 0.9781395    
# 10.initSF         4.870872   0.408248   11.9311 < 2.2e-16 ***
# 11.initSF         4.961839   1.000002    4.9618 6.983e-07 ***
# 12.initSF         6.502289   0.316228   20.5620 < 2.2e-16 ***
# 13.initSF         6.907747   1.000004    6.9077 4.925e-12 ***
# 14.initSF         5.460835   0.500000   10.9217 < 2.2e-16 ***
# 15.initSF         5.809141   0.577351   10.0617 < 2.2e-16 ***
# 16.initSF        -6.841114 319.417929   -0.0214 0.9829127    
# 17.initSF         6.214600   1.000003    6.2146 5.146e-10 ***
# 18.initSF        -5.062715 363.256388   -0.0139 0.9888802    
# 19.initSF         6.368758   0.377965   16.8501 < 2.2e-16 ***
# 20.initSF         5.452467   0.377965   14.4259 < 2.2e-16 ***
# 21.initSF        -5.062715 363.256388   -0.0139 0.9888802    
# 22.initSF         0.000000 894.427191    0.0000 1.0000000    
# 23.initSF         6.907747   1.000004    6.9077 4.925e-12 ***
# 24.initSF         4.017378   1.000002    4.0174 5.885e-05 ***
# 25.initSF         6.214600   1.000003    6.2146 5.146e-10 ***
# 26.initSF        -5.646696 346.931976   -0.0163 0.9870141    
# 27.initSF         4.961843   0.577351    8.5942 < 2.2e-16 ***
# 28.initSF         5.809136   1.000003    5.8091 6.280e-09 ***
# 29.initSF        -5.062715 363.256388   -0.0139 0.9888802    
# 30.initSF         3.940194   0.192450   20.4738 < 2.2e-16 ***
# 1.disengage      -4.262284   0.062622  -68.0632 < 2.2e-16 ***
# 2.disengage      -5.148770   0.088736  -58.0237 < 2.2e-16 ***
# 3.disengage      -4.775014   0.063628  -75.0452 < 2.2e-16 ***
# 4.disengage      -4.647453   0.054882  -84.6806 < 2.2e-16 ***
# 5.disengage      -4.573716   0.102598  -44.5791 < 2.2e-16 ***
# 6.disengage      -6.477786   0.577349  -11.2199 < 2.2e-16 ***
# 7.disengage      -5.683178   0.089443  -63.5399 < 2.2e-16 ***
# 8.disengage      -3.025632   0.188982  -16.0101 < 2.2e-16 ***
# 9.disengage      -2.951359   0.999998   -2.9514 0.0031637 ** 
# 10.disengage     -4.610931   0.088045  -52.3701 < 2.2e-16 ***
# 11.disengage     -1.251904   0.999999   -1.2519 0.2106042    
# 12.disengage     -5.099954   0.333333  -15.2999 < 2.2e-16 ***
# 13.disengage     -3.069490   0.999997   -3.0695 0.0021442 ** 
# 14.disengage     -3.999558   0.200000  -19.9978 < 2.2e-16 ***
# 15.disengage     -3.711562   0.377964   -9.8199 < 2.2e-16 ***
# 16.disengage      0.000000 894.427191    0.0000 1.0000000    
# 17.disengage     -5.585480   0.707105   -7.8991 2.887e-15 ***
# 18.disengage      0.000000 894.427191    0.0000 1.0000000    
# 19.disengage     -3.306133   0.250000  -13.2245 < 2.2e-16 ***
# 20.disengage     -2.450509   0.353553   -6.9311 4.176e-12 ***
# 21.disengage      0.000000 894.427191    0.0000 1.0000000    
# 22.disengage     -3.514373   0.999997   -3.5144 0.0004408 ***
# 23.disengage    -12.189754 246.283852   -0.0495 0.9605250    
# 24.disengage     -1.891904   0.999998   -1.8919 0.0585033 .  
# 25.disengage     -2.434066   0.200000  -12.1703 < 2.2e-16 ***
# 26.disengage     -3.804916   0.408248   -9.3201 < 2.2e-16 ***
# 27.disengage     -5.368294   0.316228  -16.9760 < 2.2e-16 ***
# 28.disengage     -5.092138   0.353553  -14.4027 < 2.2e-16 ***
# 29.disengage     -5.097787   0.408248  -12.4870 < 2.2e-16 ***
# 30.disengage     -4.831031   0.162221  -29.7805 < 2.2e-16 ***
# 1.cop.start     -21.775226 187.429617   -0.1162 0.9075114    
# 2.cop.start      -9.446606   0.408248  -23.1394 < 2.2e-16 ***
# 3.cop.start      -7.431142   0.149071  -49.8496 < 2.2e-16 ***
# 4.cop.start      -7.839479   0.218218  -35.9250 < 2.2e-16 ***
# 5.cop.start     -21.334706 189.269436   -0.1127 0.9102515    
# 6.cop.start     -19.273179 198.658065   -0.0970 0.9227130    
# 7.cop.start     -21.663929 187.889467   -0.1153 0.9082062    
# 8.cop.start     -19.504105 197.536438   -0.0987 0.9213473    
# 9.cop.start     -19.225865 198.890243   -0.0967 0.9229919    
# 10.cop.start    -20.397432 193.369815   -0.1055 0.9159917    
# 11.cop.start    -16.297433 215.065251   -0.0758 0.9395949    
# 12.cop.start    -19.261927 198.713208   -0.0969 0.9227794    
# 13.cop.start    -17.124683 210.100682   -0.0815 0.9350387    
# 14.cop.start    -19.045919 199.780757   -0.0953 0.9240495    
# 15.cop.start    -19.059755 199.711860   -0.0954 0.9239683    
# 16.cop.start    -15.410565 220.799295   -0.0698 0.9443573    
# 17.cop.start    -17.846074 206.041009   -0.0866 0.9309782    
# 18.cop.start    -16.894762 211.445896   -0.0799 0.9363159    
# 19.cop.start    -19.197853 199.028082   -0.0965 0.9231568    
# 20.cop.start    -18.175836 204.262032   -0.0890 0.9290955    
# 21.cop.start    -15.559441 219.804656   -0.0708 0.9435668    
# 22.cop.start    -18.414696 203.001849   -0.0907 0.9277215    
# 23.cop.start    -17.543193 207.716617   -0.0845 0.9326928    
# 24.cop.start    -18.088681 204.727721   -0.0884 0.9295947    
# 25.cop.start    -18.516001 202.474388   -0.0914 0.9271361    
# 26.cop.start    -17.327815 208.933327   -0.0829 0.9339035    
# 27.cop.start    -18.831473 200.857793   -0.0938 0.9253036    
# 28.cop.start    -18.717694 201.436362   -0.0929 0.9259662    
# 29.cop.start      0.000000 894.427191    0.0000 1.0000000    
# 30.cop.start    -20.543323 192.714078   -0.1066 0.9151063    
# 1.cop.end         0.000000 894.427191    0.0000 1.0000000    
# 2.cop.end        -1.651538   0.707106   -2.3356 0.0195106 *  
# 3.cop.end        -1.354380   0.160128   -8.4581 < 2.2e-16 ***
# 4.cop.end        -1.737753   0.333333   -5.2133 1.856e-07 ***
# 5.cop.end         0.000000 894.427191    0.0000 1.0000000    
# 6.cop.end         0.000000 894.427191    0.0000 1.0000000    
# 7.cop.end         0.000000 894.427191    0.0000 1.0000000    
# 8.cop.end         0.000000 894.427191    0.0000 1.0000000    
# 9.cop.end         0.000000 894.427191    0.0000 1.0000000    
# 10.cop.end        0.000000 894.427191    0.0000 1.0000000    
# 11.cop.end        0.000000 894.427191    0.0000 1.0000000    
# 12.cop.end        0.000000 894.427191    0.0000 1.0000000    
# 13.cop.end        0.000000 894.427191    0.0000 1.0000000    
# 14.cop.end        0.000000 894.427191    0.0000 1.0000000    
# 15.cop.end        0.000000 894.427191    0.0000 1.0000000    
# 16.cop.end        0.000000 894.427191    0.0000 1.0000000    
# 17.cop.end        0.000000 894.427191    0.0000 1.0000000    
# 18.cop.end        0.000000 894.427191    0.0000 1.0000000    
# 19.cop.end        0.000000 894.427191    0.0000 1.0000000    
# 20.cop.end        0.000000 894.427191    0.0000 1.0000000    
# 21.cop.end        0.000000 894.427191    0.0000 1.0000000    
# 22.cop.end        0.000000 894.427191    0.0000 1.0000000    
# 23.cop.end        0.000000 894.427191    0.0000 1.0000000    
# 24.cop.end        0.000000 894.427191    0.0000 1.0000000    
# 25.cop.end        0.000000 894.427191    0.0000 1.0000000    
# 26.cop.end        0.000000 894.427191    0.0000 1.0000000    
# 27.cop.end        0.000000 894.427191    0.0000 1.0000000    
# 28.cop.end        0.000000 894.427191    0.0000 1.0000000    
# 29.cop.end        0.000000 894.427191    0.0000 1.0000000    
# 30.cop.end        0.000000 894.427191    0.0000 1.0000000    
# 1.depart         -8.701475   0.235702  -36.9172 < 2.2e-16 ***
# 2.depart         -8.645592   0.235702  -36.6802 < 2.2e-16 ***
# 3.depart         -8.603133   0.235702  -36.5000 < 2.2e-16 ***
# 4.depart         -8.213108   0.235702  -34.8453 < 2.2e-16 ***
# 5.depart         -8.797853   0.288675  -30.4767 < 2.2e-16 ***
# 6.depart         -8.107312   0.707105  -11.4655 < 2.2e-16 ***
# 7.depart         -8.466583   0.235702  -35.9207 < 2.2e-16 ***
# 8.depart         -7.624461   0.499999  -15.2489 < 2.2e-16 ***
# 9.depart         -8.213118   0.577349  -14.2256 < 2.2e-16 ***
# 10.depart        -8.336652   0.377964  -22.0567 < 2.2e-16 ***
# 11.depart        -6.136106   0.577349  -10.6281 < 2.2e-16 ***
# 12.depart        -8.430566   0.577349  -14.6022 < 2.2e-16 ***
# 13.depart        -6.374262   0.999995   -6.3743 1.838e-10 ***
# 14.depart        -8.432862   0.707105  -11.9259 < 2.2e-16 ***
# 15.depart        -8.472487   0.999994   -8.4725 < 2.2e-16 ***
# 16.depart        -4.553187   0.999997   -4.5532 5.284e-06 ***
# 17.depart        -7.332594   0.999995   -7.3326 2.256e-13 ***
# 18.depart        -6.129319   0.999996   -6.1293 8.824e-10 ***
# 19.depart        -8.897214   0.999994   -8.8973 < 2.2e-16 ***
# 20.depart        -7.676151   0.999995   -7.6762 1.643e-14 ***
# 21.depart        -4.711675   0.999996   -4.7117 2.457e-06 ***
# 22.depart        -7.735384   0.999995   -7.7354 1.021e-14 ***
# 23.depart        -7.017221   0.999995   -7.0173 2.263e-12 ***
# 24.depart        -7.391510   0.999995   -7.3915 1.452e-13 ***
# 25.depart        -7.842174   0.999994   -7.8422 4.441e-15 ***
# 26.depart        -8.035618   0.999994   -8.0357 8.882e-16 ***
# 27.depart        -8.659685   0.999994   -8.6597 < 2.2e-16 ***
# 28.depart        -8.191807   0.999994   -8.1919 2.220e-16 ***
# 29.depart        -7.054816   0.999995   -7.0549 1.728e-12 ***
# 30.depart        -5.737913   0.117851  -48.6878 < 2.2e-16 ***
# ---
# Signif. codes:  0 ‘***’ 0.001 ‘**’ 0.01 ‘*’ 0.05 ‘.’ 0.1 ‘ ’ 1
# Null deviance: 76628.21 on 4854 degrees of freedom
# Residual deviance: 30523.84 on 4639 degrees of freedom
# 	Chi-square: 46104.37 on 215 degrees of freedom, asymptotic p-value 0 
# AIC: 30953.84 AICC: 30973.87 BIC: 32348.67 
# Log posterior: -230472.8 
# Prior parameters: mu=0 sigma=1000 nu=4 


#seems like X males receive attacks from U's slightly more, but they don't attack them at a different rate
#soo... what if I remove that term?


##Model that partitions out only Ureceives from X interactions 

sequences<-table(event.list$Date)

#create list of dummy intercept names. Leave male 30.receives as the baseline- this is the receive rate for ?? males- I expect everyone else's receive rate to be less

idno<-length(event.types[grep("cop.start",event.types)])

#dummy.ints.attack<-paste(1:(idno-1),".attackFE", sep="")
#dummy.ints.receive<-paste(1:(idno-1), ".receiveFE", sep="")


int.names<-c("Global.attackFE", "X.attackFE", "X.receiveFE", "X.UreceiveFE", event.types[-grep("attack", event.types)])

int.list<-list()

for(i in 1:length(levels(event.list$Date))){

day.array<-array(0,dim=c(as.numeric(sequences[i]), length(event.types),length(int.names)))

#the third dimension is the stat slice.
dimnames(day.array)<-list(c(1:sequences[i]),as.character(event.types),as.character(int.names))

#add global attack effect, loops for "U" intercepts for attacks and receives, then one loop for typical intercepts

day.array[,grep("attack", event.types),1]<-1

for(h in 1:14){
day.array[,paste(30,".attack.", h, sep=""),2]<-1
}

for(h in 1:14){
day.array[,paste(h,".attack.",30, sep=""),3]<-1
}


for(h in 15:29){
day.array[,paste(h,".attack.",30, sep=""),4]<-1
}


for(k in 5:length(int.names)){
	
	day.array[,int.names[k],k]<-1
	
}

#weird extra formatting to make it match the package example, seems like it becomes relevant *punintended!* if I want to fit day-specific effects

global<-list()

global[[1]]<-day.array

int.list[[i]]<-global

names(int.list[[i]])<-"global"

}

tic()
intfit.sup<-rem(evlist, int.list, supplist=supplist, timing="interval", estimator="BPM")
toc()

summary(intfit.sup)


# Egocentric Relational Event Model (Interval Likelihood)
# 
#                  Post.Mode    Post.SD   Z value  Pr(>|z|)    
# Global.attackFE  -8.070543   0.032092 -251.4852 < 2.2e-16 ***
# X.attackFE       -1.580576   0.317852   -4.9727 6.603e-07 ***
# X.receiveFE       2.315876   0.055339   41.8490 < 2.2e-16 ***
# X.UreceiveFE      2.447144   0.177011   13.8248 < 2.2e-16 ***
# 1.initFO          6.744430   0.127000   53.1057 < 2.2e-16 ***
# 2.initFO          6.700116   0.104828   63.9150 < 2.2e-16 ***
# 3.initFO          6.738679   0.114708   58.7464 < 2.2e-16 ***
# 4.initFO          6.684612   0.075378   88.6814 < 2.2e-16 ***
# 5.initFO          6.792686   0.156174   43.4944 < 2.2e-16 ***
# 6.initFO          5.061926   0.577351    8.7675 < 2.2e-16 ***
# 7.initFO          6.715748   0.085436   78.6058 < 2.2e-16 ***
# 8.initFO          5.115994   0.500000   10.2320 < 2.2e-16 ***
# 9.initFO          3.381391   1.000001    3.3814 0.0007212 ***
# 10.initFO         6.662633   0.166667   39.9758 < 2.2e-16 ***
# 11.initFO        -6.724740 321.814902   -0.0209 0.9833284    
# 12.initFO         5.809142   0.447214   12.9896 < 2.2e-16 ***
# 13.initFO        -5.062715 363.256388   -0.0139 0.9888802    
# 14.initFO         5.460835   0.500000   10.9217 < 2.2e-16 ***
# 15.initFO         6.096823   0.500000   12.1936 < 2.2e-16 ***
# 16.initFO        -6.841114 319.417929   -0.0214 0.9829127    
# 17.initFO        -5.646696 346.931976   -0.0163 0.9870141    
# 18.initFO        -5.062715 363.256388   -0.0139 0.9888802    
# 19.initFO         4.422843   1.000002    4.4228 9.741e-06 ***
# 20.initFO         3.506554   1.000002    3.5065 0.0004540 ***
# 21.initFO        -5.062715 363.256388   -0.0139 0.9888802    
# 22.initFO         0.000000 894.427191    0.0000 1.0000000    
# 23.initFO        -5.062715 363.256388   -0.0139 0.9888802    
# 24.initFO        -7.553047 305.835850   -0.0247 0.9802971    
# 25.initFO         6.214600   1.000003    6.2146 5.146e-10 ***
# 26.initFO         6.907751   0.707108    9.7690 < 2.2e-16 ***
# 27.initFO         6.060457   0.333333   18.1814 < 2.2e-16 ***
# 28.initFO         6.502286   0.707108    9.1956 < 2.2e-16 ***
# 29.initFO         6.907747   1.000004    6.9077 4.925e-12 ***
# 30.initFO         3.352407   0.258199   12.9838 < 2.2e-16 ***
# 1.initCH          2.617293   1.000001    2.6173 0.0088631 ** 
# 2.initCH         -9.185505 280.258756   -0.0328 0.9738539    
# 3.initCH         -8.988496 283.008974   -0.0318 0.9746631    
# 4.initCH          1.514126   1.000000    1.5141 0.1299940    
# 5.initCH          3.079110   1.000001    3.0791 0.0020762 ** 
# 6.initCH          6.735904   0.250000   26.9436 < 2.2e-16 ***
# 7.initCH         -9.541022 275.492334   -0.0346 0.9723727    
# 8.initCH          6.562914   0.242536   27.0596 < 2.2e-16 ***
# 9.initCH          6.877902   0.174078   39.5105 < 2.2e-16 ***
# 10.initCH         4.465407   0.500000    8.9308 < 2.2e-16 ***
# 11.initCH         6.753603   0.408249   16.5429 < 2.2e-16 ***
# 12.initCH        -7.392244 308.751880   -0.0239 0.9808986    
# 13.initCH        -5.062715 363.256388   -0.0139 0.9888802    
# 14.initCH         6.271766   0.333333   18.8153 < 2.2e-16 ***
# 15.initCH         5.403675   0.707108    7.6419 2.132e-14 ***
# 16.initCH         6.907754   0.353554   19.5381 < 2.2e-16 ***
# 17.initCH         6.214600   1.000003    6.2146 5.146e-10 ***
# 18.initCH         6.907747   1.000004    6.9077 4.925e-12 ***
# 19.initCH         5.809141   0.500000   11.6183 < 2.2e-16 ***
# 20.initCH         6.597600   0.213201   30.9455 < 2.2e-16 ***
# 21.initCH         6.907747   1.000004    6.9077 4.925e-12 ***
# 22.initCH         0.000000 894.427191    0.0000 1.0000000    
# 23.initCH        -5.062715 363.256388   -0.0139 0.9888802    
# 24.initCH         6.850596   0.242536   28.2457 < 2.2e-16 ***
# 25.initCH        -5.646696 346.931976   -0.0163 0.9870141    
# 26.initCH        -5.646696 346.931976   -0.0163 0.9870141    
# 27.initCH         6.060457   0.333333   18.1814 < 2.2e-16 ***
# 28.initCH        -5.992692 338.240255   -0.0177 0.9858644    
# 29.initCH        -5.062715 363.256388   -0.0139 0.9888802    
# 30.initCH         6.824374   0.045502  149.9810 < 2.2e-16 ***
# 1.initSF          4.919880   0.316228   15.5580 < 2.2e-16 ***
# 2.initSF          5.233779   0.218218   23.9842 < 2.2e-16 ***
# 3.initSF          5.047002   0.267261   18.8842 < 2.2e-16 ***
# 4.initSF          5.275328   0.152499   34.5926 < 2.2e-16 ***
# 5.initSF          4.465407   0.500000    8.9308 < 2.2e-16 ***
# 6.initSF         -7.600810 304.985488   -0.0249 0.9801173    
# 7.initSF          5.163063   0.185695   27.8039 < 2.2e-16 ***
# 8.initSF          4.828312   0.577351    8.3629 < 2.2e-16 ***
# 9.initSF         -8.117025 296.225732   -0.0274 0.9781395    
# 10.initSF         4.870872   0.408248   11.9311 < 2.2e-16 ***
# 11.initSF         4.961839   1.000002    4.9618 6.983e-07 ***
# 12.initSF         6.502289   0.316228   20.5620 < 2.2e-16 ***
# 13.initSF         6.907747   1.000004    6.9077 4.925e-12 ***
# 14.initSF         5.460835   0.500000   10.9217 < 2.2e-16 ***
# 15.initSF         5.809141   0.577351   10.0617 < 2.2e-16 ***
# 16.initSF        -6.841114 319.417929   -0.0214 0.9829127    
# 17.initSF         6.214600   1.000003    6.2146 5.146e-10 ***
# 18.initSF        -5.062715 363.256388   -0.0139 0.9888802    
# 19.initSF         6.368758   0.377965   16.8501 < 2.2e-16 ***
# 20.initSF         5.452467   0.377965   14.4259 < 2.2e-16 ***
# 21.initSF        -5.062715 363.256388   -0.0139 0.9888802    
# 22.initSF         0.000000 894.427191    0.0000 1.0000000    
# 23.initSF         6.907747   1.000004    6.9077 4.925e-12 ***
# 24.initSF         4.017378   1.000002    4.0174 5.885e-05 ***
# 25.initSF         6.214600   1.000003    6.2146 5.146e-10 ***
# 26.initSF        -5.646696 346.931976   -0.0163 0.9870141    
# 27.initSF         4.961843   0.577351    8.5942 < 2.2e-16 ***
# 28.initSF         5.809136   1.000003    5.8091 6.280e-09 ***
# 29.initSF        -5.062715 363.256388   -0.0139 0.9888802    
# 30.initSF         3.940194   0.192450   20.4738 < 2.2e-16 ***
# 1.disengage      -4.262284   0.062622  -68.0632 < 2.2e-16 ***
# 2.disengage      -5.148770   0.088736  -58.0237 < 2.2e-16 ***
# 3.disengage      -4.775014   0.063628  -75.0452 < 2.2e-16 ***
# 4.disengage      -4.647453   0.054882  -84.6806 < 2.2e-16 ***
# 5.disengage      -4.573716   0.102598  -44.5791 < 2.2e-16 ***
# 6.disengage      -6.477786   0.577349  -11.2199 < 2.2e-16 ***
# 7.disengage      -5.683178   0.089443  -63.5399 < 2.2e-16 ***
# 8.disengage      -3.025632   0.188982  -16.0101 < 2.2e-16 ***
# 9.disengage      -2.951359   0.999998   -2.9514 0.0031637 ** 
# 10.disengage     -4.610931   0.088045  -52.3701 < 2.2e-16 ***
# 11.disengage     -1.251904   0.999999   -1.2519 0.2106042    
# 12.disengage     -5.099954   0.333333  -15.2999 < 2.2e-16 ***
# 13.disengage     -3.069490   0.999997   -3.0695 0.0021442 ** 
# 14.disengage     -3.999558   0.200000  -19.9978 < 2.2e-16 ***
# 15.disengage     -3.711562   0.377964   -9.8199 < 2.2e-16 ***
# 16.disengage      0.000000 894.427191    0.0000 1.0000000    
# 17.disengage     -5.585480   0.707105   -7.8991 2.887e-15 ***
# 18.disengage      0.000000 894.427191    0.0000 1.0000000    
# 19.disengage     -3.306133   0.250000  -13.2245 < 2.2e-16 ***
# 20.disengage     -2.450509   0.353553   -6.9311 4.176e-12 ***
# 21.disengage      0.000000 894.427191    0.0000 1.0000000    
# 22.disengage     -3.514373   0.999997   -3.5144 0.0004408 ***
# 23.disengage    -12.189754 246.283852   -0.0495 0.9605250    
# 24.disengage     -1.891904   0.999998   -1.8919 0.0585033 .  
# 25.disengage     -2.434066   0.200000  -12.1703 < 2.2e-16 ***
# 26.disengage     -3.804916   0.408248   -9.3201 < 2.2e-16 ***
# 27.disengage     -5.368294   0.316228  -16.9760 < 2.2e-16 ***
# 28.disengage     -5.092138   0.353553  -14.4027 < 2.2e-16 ***
# 29.disengage     -5.097787   0.408248  -12.4870 < 2.2e-16 ***
# 30.disengage     -4.831031   0.162221  -29.7805 < 2.2e-16 ***
# 1.cop.start     -21.775226 187.429617   -0.1162 0.9075114    
# 2.cop.start      -9.446606   0.408248  -23.1394 < 2.2e-16 ***
# 3.cop.start      -7.431142   0.149071  -49.8496 < 2.2e-16 ***
# 4.cop.start      -7.839479   0.218218  -35.9250 < 2.2e-16 ***
# 5.cop.start     -21.334706 189.269436   -0.1127 0.9102515    
# 6.cop.start     -19.273179 198.658065   -0.0970 0.9227130    
# 7.cop.start     -21.663929 187.889467   -0.1153 0.9082062    
# 8.cop.start     -19.504105 197.536438   -0.0987 0.9213473    
# 9.cop.start     -19.225865 198.890243   -0.0967 0.9229919    
# 10.cop.start    -20.397432 193.369815   -0.1055 0.9159917    
# 11.cop.start    -16.297433 215.065251   -0.0758 0.9395949    
# 12.cop.start    -19.261927 198.713208   -0.0969 0.9227794    
# 13.cop.start    -17.124683 210.100682   -0.0815 0.9350387    
# 14.cop.start    -19.045919 199.780757   -0.0953 0.9240495    
# 15.cop.start    -19.059755 199.711860   -0.0954 0.9239683    
# 16.cop.start    -15.410565 220.799295   -0.0698 0.9443573    
# 17.cop.start    -17.846074 206.041009   -0.0866 0.9309782    
# 18.cop.start    -16.894762 211.445896   -0.0799 0.9363159    
# 19.cop.start    -19.197853 199.028082   -0.0965 0.9231568    
# 20.cop.start    -18.175836 204.262032   -0.0890 0.9290955    
# 21.cop.start    -15.559441 219.804656   -0.0708 0.9435668    
# 22.cop.start    -18.414696 203.001849   -0.0907 0.9277215    
# 23.cop.start    -17.543193 207.716617   -0.0845 0.9326928    
# 24.cop.start    -18.088681 204.727721   -0.0884 0.9295947    
# 25.cop.start    -18.516001 202.474388   -0.0914 0.9271361    
# 26.cop.start    -17.327815 208.933327   -0.0829 0.9339035    
# 27.cop.start    -18.831473 200.857793   -0.0938 0.9253036    
# 28.cop.start    -18.717694 201.436362   -0.0929 0.9259662    
# 29.cop.start      0.000000 894.427191    0.0000 1.0000000    
# 30.cop.start    -20.543323 192.714078   -0.1066 0.9151063    
# 1.cop.end         0.000000 894.427191    0.0000 1.0000000    
# 2.cop.end        -1.651538   0.707106   -2.3356 0.0195106 *  
# 3.cop.end        -1.354380   0.160128   -8.4581 < 2.2e-16 ***
# 4.cop.end        -1.737753   0.333333   -5.2133 1.856e-07 ***
# 5.cop.end         0.000000 894.427191    0.0000 1.0000000    
# 6.cop.end         0.000000 894.427191    0.0000 1.0000000    
# 7.cop.end         0.000000 894.427191    0.0000 1.0000000    
# 8.cop.end         0.000000 894.427191    0.0000 1.0000000    
# 9.cop.end         0.000000 894.427191    0.0000 1.0000000    
# 10.cop.end        0.000000 894.427191    0.0000 1.0000000    
# 11.cop.end        0.000000 894.427191    0.0000 1.0000000    
# 12.cop.end        0.000000 894.427191    0.0000 1.0000000    
# 13.cop.end        0.000000 894.427191    0.0000 1.0000000    
# 14.cop.end        0.000000 894.427191    0.0000 1.0000000    
# 15.cop.end        0.000000 894.427191    0.0000 1.0000000    
# 16.cop.end        0.000000 894.427191    0.0000 1.0000000    
# 17.cop.end        0.000000 894.427191    0.0000 1.0000000    
# 18.cop.end        0.000000 894.427191    0.0000 1.0000000    
# 19.cop.end        0.000000 894.427191    0.0000 1.0000000    
# 20.cop.end        0.000000 894.427191    0.0000 1.0000000    
# 21.cop.end        0.000000 894.427191    0.0000 1.0000000    
# 22.cop.end        0.000000 894.427191    0.0000 1.0000000    
# 23.cop.end        0.000000 894.427191    0.0000 1.0000000    
# 24.cop.end        0.000000 894.427191    0.0000 1.0000000    
# 25.cop.end        0.000000 894.427191    0.0000 1.0000000    
# 26.cop.end        0.000000 894.427191    0.0000 1.0000000    
# 27.cop.end        0.000000 894.427191    0.0000 1.0000000    
# 28.cop.end        0.000000 894.427191    0.0000 1.0000000    
# 29.cop.end        0.000000 894.427191    0.0000 1.0000000    
# 30.cop.end        0.000000 894.427191    0.0000 1.0000000    
# 1.depart         -8.701475   0.235702  -36.9172 < 2.2e-16 ***
# 2.depart         -8.645592   0.235702  -36.6802 < 2.2e-16 ***
# 3.depart         -8.603133   0.235702  -36.5000 < 2.2e-16 ***
# 4.depart         -8.213108   0.235702  -34.8453 < 2.2e-16 ***
# 5.depart         -8.797853   0.288675  -30.4767 < 2.2e-16 ***
# 6.depart         -8.107312   0.707105  -11.4655 < 2.2e-16 ***
# 7.depart         -8.466583   0.235702  -35.9207 < 2.2e-16 ***
# 8.depart         -7.624461   0.499999  -15.2489 < 2.2e-16 ***
# 9.depart         -8.213118   0.577349  -14.2256 < 2.2e-16 ***
# 10.depart        -8.336652   0.377964  -22.0567 < 2.2e-16 ***
# 11.depart        -6.136106   0.577349  -10.6281 < 2.2e-16 ***
# 12.depart        -8.430566   0.577349  -14.6022 < 2.2e-16 ***
# 13.depart        -6.374262   0.999995   -6.3743 1.838e-10 ***
# 14.depart        -8.432862   0.707105  -11.9259 < 2.2e-16 ***
# 15.depart        -8.472487   0.999994   -8.4725 < 2.2e-16 ***
# 16.depart        -4.553187   0.999997   -4.5532 5.284e-06 ***
# 17.depart        -7.332594   0.999995   -7.3326 2.256e-13 ***
# 18.depart        -6.129319   0.999996   -6.1293 8.824e-10 ***
# 19.depart        -8.897214   0.999994   -8.8973 < 2.2e-16 ***
# 20.depart        -7.676151   0.999995   -7.6762 1.643e-14 ***
# 21.depart        -4.711675   0.999996   -4.7117 2.457e-06 ***
# 22.depart        -7.735384   0.999995   -7.7354 1.021e-14 ***
# 23.depart        -7.017221   0.999995   -7.0173 2.263e-12 ***
# 24.depart        -7.391510   0.999995   -7.3915 1.452e-13 ***
# 25.depart        -7.842174   0.999994   -7.8422 4.441e-15 ***
# 26.depart        -8.035618   0.999994   -8.0357 8.882e-16 ***
# 27.depart        -8.659685   0.999994   -8.6597 < 2.2e-16 ***
# 28.depart        -8.191807   0.999994   -8.1919 2.220e-16 ***
# 29.depart        -7.054816   0.999995   -7.0549 1.728e-12 ***
# 30.depart        -5.737913   0.117851  -48.6878 < 2.2e-16 ***
# ---
# Signif. codes:  0 ‘***’ 0.001 ‘**’ 0.01 ‘*’ 0.05 ‘.’ 0.1 ‘ ’ 1
# Null deviance: 76628.21 on 4854 degrees of freedom
# Residual deviance: 30524.13 on 4640 degrees of freedom
# 	Chi-square: 46104.08 on 214 degrees of freedom, asymptotic p-value 0 
# AIC: 30952.13 AICC: 30971.97 BIC: 32340.47 
# Log posterior: -229472 
# Prior parameters: mu=0 sigma=1000 nu=4 


#a little better but still not as good as the baseline


### Try partitioning out the interactions among the U males now

#these guys almost NEVER interact because mostly they are not present simultaneously on the same day

sequences<-table(event.list$Date)

#create list of dummy intercept names. Leave male 30.receives as the baseline- this is the receive rate for ?? males- I expect everyone else's receive rate to be less

idno<-length(event.types[grep("cop.start",event.types)])

#dummy.ints.attack<-paste(1:(idno-1),".attackFE", sep="")
#dummy.ints.receive<-paste(1:(idno-1), ".receiveFE", sep="")


int.names<-c("Global.attackFE", "X.attackFE", "X.receiveFE", "X.UreceiveFE", "U.attackFE", "U.receiveFE", "U.UattackFE", event.types[-grep("attack", event.types)])

int.list<-list()

for(i in 1:length(levels(event.list$Date))){

day.array<-array(0,dim=c(as.numeric(sequences[i]), length(event.types),length(int.names)))

#the third dimension is the stat slice.
dimnames(day.array)<-list(c(1:sequences[i]),as.character(event.types),as.character(int.names))

#add global attack effect, loops for "U" intercepts for attacks and receives, then one loop for typical intercepts

day.array[,grep("attack", event.types),1]<-1

for(h in 1:14){
day.array[,paste(30,".attack.", h, sep=""),2]<-1
}

for(h in 1:14){
day.array[,paste(h,".attack.",30, sep=""),3]<-1
}


for(h in 15:29){
day.array[,paste(h,".attack.",30, sep=""),4]<-1
}


for(h in 1:14){
	for(j in 15:29){	
day.array[,paste(j,".attack.", h, sep=""),5]<-1
}
}


for(h in 1:14){
	for(j in 15:29){
day.array[,paste(h,".attack.",j, sep=""),6]<-1
}
}

for(h in 15:29){
	for(j in 15:29){	
if(h!=j){
day.array[,paste(j,".attack.", h, sep=""),7]<-1
}
}
}


for(k in 8:length(int.names)){
	
	day.array[,int.names[k],k]<-1
	
}

#weird extra formatting to make it match the package example, seems like it becomes relevant *punintended!* if I want to fit day-specific effects

global<-list()

global[[1]]<-day.array

int.list[[i]]<-global

names(int.list[[i]])<-"global"

}

tic()
intfit.sup<-rem(evlist, int.list, supplist=supplist, timing="interval", estimator="BPM")
toc()

summary(intfit.sup)

# Egocentric Relational Event Model (Interval Likelihood)
# 
#                  Post.Mode    Post.SD   Z value  Pr(>|z|)    
# Global.attackFE  -8.081802   0.034120 -236.8671 < 2.2e-16 ***
# X.attackFE       -1.569316   0.318063   -4.9340 8.057e-07 ***
# X.receiveFE       2.327136   0.056539   41.1598 < 2.2e-16 ***
# X.UreceiveFE      2.458404   0.177390   13.8588 < 2.2e-16 ***
# U.attackFE       -2.717527   0.578357   -4.6987 2.618e-06 ***
# U.receiveFE       0.343608   0.129573    2.6519 0.0080051 ** 
# U.UattackFE       1.473509   0.152926    9.6354 < 2.2e-16 ***
# 1.initFO          6.744430   0.127000   53.1057 < 2.2e-16 ***
# 2.initFO          6.700116   0.104828   63.9150 < 2.2e-16 ***
# 3.initFO          6.738679   0.114708   58.7464 < 2.2e-16 ***
# 4.initFO          6.684612   0.075378   88.6814 < 2.2e-16 ***
# 5.initFO          6.792686   0.156174   43.4944 < 2.2e-16 ***
# 6.initFO          5.061926   0.577351    8.7675 < 2.2e-16 ***
# 7.initFO          6.715748   0.085436   78.6058 < 2.2e-16 ***
# 8.initFO          5.115994   0.500000   10.2320 < 2.2e-16 ***
# 9.initFO          3.381391   1.000001    3.3814 0.0007212 ***
# 10.initFO         6.662633   0.166667   39.9758 < 2.2e-16 ***
# 11.initFO        -6.724740 321.814902   -0.0209 0.9833284    
# 12.initFO         5.809142   0.447214   12.9896 < 2.2e-16 ***
# 13.initFO        -5.062715 363.256388   -0.0139 0.9888802    
# 14.initFO         5.460835   0.500000   10.9217 < 2.2e-16 ***
# 15.initFO         6.096823   0.500000   12.1936 < 2.2e-16 ***
# 16.initFO        -6.841114 319.417929   -0.0214 0.9829127    
# 17.initFO        -5.646696 346.931976   -0.0163 0.9870141    
# 18.initFO        -5.062715 363.256388   -0.0139 0.9888802    
# 19.initFO         4.422843   1.000002    4.4228 9.741e-06 ***
# 20.initFO         3.506554   1.000002    3.5065 0.0004540 ***
# 21.initFO        -5.062715 363.256388   -0.0139 0.9888802    
# 22.initFO         0.000000 894.427191    0.0000 1.0000000    
# 23.initFO        -5.062715 363.256388   -0.0139 0.9888802    
# 24.initFO        -7.553047 305.835850   -0.0247 0.9802971    
# 25.initFO         6.214600   1.000003    6.2146 5.146e-10 ***
# 26.initFO         6.907751   0.707108    9.7690 < 2.2e-16 ***
# 27.initFO         6.060457   0.333333   18.1814 < 2.2e-16 ***
# 28.initFO         6.502286   0.707108    9.1956 < 2.2e-16 ***
# 29.initFO         6.907747   1.000004    6.9077 4.925e-12 ***
# 30.initFO         3.352407   0.258199   12.9838 < 2.2e-16 ***
# 1.initCH          2.617293   1.000001    2.6173 0.0088631 ** 
# 2.initCH         -9.185505 280.258756   -0.0328 0.9738539    
# 3.initCH         -8.988496 283.008974   -0.0318 0.9746631    
# 4.initCH          1.514126   1.000000    1.5141 0.1299940    
# 5.initCH          3.079110   1.000001    3.0791 0.0020762 ** 
# 6.initCH          6.735904   0.250000   26.9436 < 2.2e-16 ***
# 7.initCH         -9.541022 275.492334   -0.0346 0.9723727    
# 8.initCH          6.562914   0.242536   27.0596 < 2.2e-16 ***
# 9.initCH          6.877902   0.174078   39.5105 < 2.2e-16 ***
# 10.initCH         4.465407   0.500000    8.9308 < 2.2e-16 ***
# 11.initCH         6.753603   0.408249   16.5429 < 2.2e-16 ***
# 12.initCH        -7.392244 308.751880   -0.0239 0.9808986    
# 13.initCH        -5.062715 363.256388   -0.0139 0.9888802    
# 14.initCH         6.271766   0.333333   18.8153 < 2.2e-16 ***
# 15.initCH         5.403675   0.707108    7.6419 2.132e-14 ***
# 16.initCH         6.907754   0.353554   19.5381 < 2.2e-16 ***
# 17.initCH         6.214600   1.000003    6.2146 5.146e-10 ***
# 18.initCH         6.907747   1.000004    6.9077 4.925e-12 ***
# 19.initCH         5.809141   0.500000   11.6183 < 2.2e-16 ***
# 20.initCH         6.597600   0.213201   30.9455 < 2.2e-16 ***
# 21.initCH         6.907747   1.000004    6.9077 4.925e-12 ***
# 22.initCH         0.000000 894.427191    0.0000 1.0000000    
# 23.initCH        -5.062715 363.256388   -0.0139 0.9888802    
# 24.initCH         6.850596   0.242536   28.2457 < 2.2e-16 ***
# 25.initCH        -5.646696 346.931976   -0.0163 0.9870141    
# 26.initCH        -5.646696 346.931976   -0.0163 0.9870141    
# 27.initCH         6.060457   0.333333   18.1814 < 2.2e-16 ***
# 28.initCH        -5.992692 338.240255   -0.0177 0.9858644    
# 29.initCH        -5.062715 363.256388   -0.0139 0.9888802    
# 30.initCH         6.824374   0.045502  149.9810 < 2.2e-16 ***
# 1.initSF          4.919880   0.316228   15.5580 < 2.2e-16 ***
# 2.initSF          5.233779   0.218218   23.9842 < 2.2e-16 ***
# 3.initSF          5.047002   0.267261   18.8842 < 2.2e-16 ***
# 4.initSF          5.275328   0.152499   34.5926 < 2.2e-16 ***
# 5.initSF          4.465407   0.500000    8.9308 < 2.2e-16 ***
# 6.initSF         -7.600810 304.985488   -0.0249 0.9801173    
# 7.initSF          5.163063   0.185695   27.8039 < 2.2e-16 ***
# 8.initSF          4.828312   0.577351    8.3629 < 2.2e-16 ***
# 9.initSF         -8.117025 296.225732   -0.0274 0.9781395    
# 10.initSF         4.870872   0.408248   11.9311 < 2.2e-16 ***
# 11.initSF         4.961839   1.000002    4.9618 6.983e-07 ***
# 12.initSF         6.502289   0.316228   20.5620 < 2.2e-16 ***
# 13.initSF         6.907747   1.000004    6.9077 4.925e-12 ***
# 14.initSF         5.460835   0.500000   10.9217 < 2.2e-16 ***
# 15.initSF         5.809141   0.577351   10.0617 < 2.2e-16 ***
# 16.initSF        -6.841114 319.417929   -0.0214 0.9829127    
# 17.initSF         6.214600   1.000003    6.2146 5.146e-10 ***
# 18.initSF        -5.062715 363.256388   -0.0139 0.9888802    
# 19.initSF         6.368758   0.377965   16.8501 < 2.2e-16 ***
# 20.initSF         5.452467   0.377965   14.4259 < 2.2e-16 ***
# 21.initSF        -5.062715 363.256388   -0.0139 0.9888802    
# 22.initSF         0.000000 894.427191    0.0000 1.0000000    
# 23.initSF         6.907747   1.000004    6.9077 4.925e-12 ***
# 24.initSF         4.017378   1.000002    4.0174 5.885e-05 ***
# 25.initSF         6.214600   1.000003    6.2146 5.146e-10 ***
# 26.initSF        -5.646696 346.931976   -0.0163 0.9870141    
# 27.initSF         4.961843   0.577351    8.5942 < 2.2e-16 ***
# 28.initSF         5.809136   1.000003    5.8091 6.280e-09 ***
# 29.initSF        -5.062715 363.256388   -0.0139 0.9888802    
# 30.initSF         3.940194   0.192450   20.4738 < 2.2e-16 ***
# 1.disengage      -4.262284   0.062622  -68.0632 < 2.2e-16 ***
# 2.disengage      -5.148770   0.088736  -58.0237 < 2.2e-16 ***
# 3.disengage      -4.775014   0.063628  -75.0452 < 2.2e-16 ***
# 4.disengage      -4.647453   0.054882  -84.6806 < 2.2e-16 ***
# 5.disengage      -4.573716   0.102598  -44.5791 < 2.2e-16 ***
# 6.disengage      -6.477786   0.577349  -11.2199 < 2.2e-16 ***
# 7.disengage      -5.683178   0.089443  -63.5399 < 2.2e-16 ***
# 8.disengage      -3.025632   0.188982  -16.0101 < 2.2e-16 ***
# 9.disengage      -2.951359   0.999998   -2.9514 0.0031637 ** 
# 10.disengage     -4.610931   0.088045  -52.3701 < 2.2e-16 ***
# 11.disengage     -1.251904   0.999999   -1.2519 0.2106042    
# 12.disengage     -5.099954   0.333333  -15.2999 < 2.2e-16 ***
# 13.disengage     -3.069490   0.999997   -3.0695 0.0021442 ** 
# 14.disengage     -3.999558   0.200000  -19.9978 < 2.2e-16 ***
# 15.disengage     -3.711562   0.377964   -9.8199 < 2.2e-16 ***
# 16.disengage      0.000000 894.427191    0.0000 1.0000000    
# 17.disengage     -5.585480   0.707105   -7.8991 2.887e-15 ***
# 18.disengage      0.000000 894.427191    0.0000 1.0000000    
# 19.disengage     -3.306133   0.250000  -13.2245 < 2.2e-16 ***
# 20.disengage     -2.450509   0.353553   -6.9311 4.176e-12 ***
# 21.disengage      0.000000 894.427191    0.0000 1.0000000    
# 22.disengage     -3.514373   0.999997   -3.5144 0.0004408 ***
# 23.disengage    -12.189754 246.283852   -0.0495 0.9605250    
# 24.disengage     -1.891904   0.999998   -1.8919 0.0585033 .  
# 25.disengage     -2.434066   0.200000  -12.1703 < 2.2e-16 ***
# 26.disengage     -3.804916   0.408248   -9.3201 < 2.2e-16 ***
# 27.disengage     -5.368294   0.316228  -16.9760 < 2.2e-16 ***
# 28.disengage     -5.092138   0.353553  -14.4027 < 2.2e-16 ***
# 29.disengage     -5.097787   0.408248  -12.4870 < 2.2e-16 ***
# 30.disengage     -4.831031   0.162221  -29.7805 < 2.2e-16 ***
# 1.cop.start     -21.775226 187.429600   -0.1162 0.9075113    
# 2.cop.start      -9.446606   0.408248  -23.1394 < 2.2e-16 ***
# 3.cop.start      -7.431142   0.149071  -49.8496 < 2.2e-16 ***
# 4.cop.start      -7.839479   0.218218  -35.9250 < 2.2e-16 ***
# 5.cop.start     -21.334706 189.269428   -0.1127 0.9102515    
# 6.cop.start     -19.273179 198.658065   -0.0970 0.9227130    
# 7.cop.start     -21.663929 187.889453   -0.1153 0.9082062    
# 8.cop.start     -19.504105 197.536438   -0.0987 0.9213473    
# 9.cop.start     -19.225865 198.890243   -0.0967 0.9229919    
# 10.cop.start    -20.397432 193.369813   -0.1055 0.9159917    
# 11.cop.start    -16.297433 215.065251   -0.0758 0.9395949    
# 12.cop.start    -19.261927 198.713208   -0.0969 0.9227794    
# 13.cop.start    -17.124683 210.100682   -0.0815 0.9350387    
# 14.cop.start    -19.045919 199.780757   -0.0953 0.9240495    
# 15.cop.start    -19.059755 199.711859   -0.0954 0.9239683    
# 16.cop.start    -15.410565 220.799295   -0.0698 0.9443573    
# 17.cop.start    -17.846074 206.041009   -0.0866 0.9309782    
# 18.cop.start    -16.894762 211.445896   -0.0799 0.9363159    
# 19.cop.start    -19.197853 199.028082   -0.0965 0.9231568    
# 20.cop.start    -18.175836 204.262032   -0.0890 0.9290955    
# 21.cop.start    -15.559441 219.804656   -0.0708 0.9435668    
# 22.cop.start    -18.414696 203.001849   -0.0907 0.9277215    
# 23.cop.start    -17.543193 207.716617   -0.0845 0.9326928    
# 24.cop.start    -18.088681 204.727721   -0.0884 0.9295947    
# 25.cop.start    -18.516001 202.474388   -0.0914 0.9271361    
# 26.cop.start    -17.327815 208.933327   -0.0829 0.9339035    
# 27.cop.start    -18.831473 200.857793   -0.0938 0.9253036    
# 28.cop.start    -18.717694 201.436362   -0.0929 0.9259662    
# 29.cop.start      0.000000 894.427191    0.0000 1.0000000    
# 30.cop.start    -20.543323 192.714076   -0.1066 0.9151063    
# 1.cop.end         0.000000 894.427191    0.0000 1.0000000    
# 2.cop.end        -1.651538   0.707106   -2.3356 0.0195106 *  
# 3.cop.end        -1.354380   0.160128   -8.4581 < 2.2e-16 ***
# 4.cop.end        -1.737753   0.333333   -5.2133 1.856e-07 ***
# 5.cop.end         0.000000 894.427191    0.0000 1.0000000    
# 6.cop.end         0.000000 894.427191    0.0000 1.0000000    
# 7.cop.end         0.000000 894.427191    0.0000 1.0000000    
# 8.cop.end         0.000000 894.427191    0.0000 1.0000000    
# 9.cop.end         0.000000 894.427191    0.0000 1.0000000    
# 10.cop.end        0.000000 894.427191    0.0000 1.0000000    
# 11.cop.end        0.000000 894.427191    0.0000 1.0000000    
# 12.cop.end        0.000000 894.427191    0.0000 1.0000000    
# 13.cop.end        0.000000 894.427191    0.0000 1.0000000    
# 14.cop.end        0.000000 894.427191    0.0000 1.0000000    
# 15.cop.end        0.000000 894.427191    0.0000 1.0000000    
# 16.cop.end        0.000000 894.427191    0.0000 1.0000000    
# 17.cop.end        0.000000 894.427191    0.0000 1.0000000    
# 18.cop.end        0.000000 894.427191    0.0000 1.0000000    
# 19.cop.end        0.000000 894.427191    0.0000 1.0000000    
# 20.cop.end        0.000000 894.427191    0.0000 1.0000000    
# 21.cop.end        0.000000 894.427191    0.0000 1.0000000    
# 22.cop.end        0.000000 894.427191    0.0000 1.0000000    
# 23.cop.end        0.000000 894.427191    0.0000 1.0000000    
# 24.cop.end        0.000000 894.427191    0.0000 1.0000000    
# 25.cop.end        0.000000 894.427191    0.0000 1.0000000    
# 26.cop.end        0.000000 894.427191    0.0000 1.0000000    
# 27.cop.end        0.000000 894.427191    0.0000 1.0000000    
# 28.cop.end        0.000000 894.427191    0.0000 1.0000000    
# 29.cop.end        0.000000 894.427191    0.0000 1.0000000    
# 30.cop.end        0.000000 894.427191    0.0000 1.0000000    
# 1.depart         -8.701475   0.235702  -36.9172 < 2.2e-16 ***
# 2.depart         -8.645592   0.235702  -36.6802 < 2.2e-16 ***
# 3.depart         -8.603133   0.235702  -36.5000 < 2.2e-16 ***
# 4.depart         -8.213108   0.235702  -34.8453 < 2.2e-16 ***
# 5.depart         -8.797853   0.288675  -30.4767 < 2.2e-16 ***
# 6.depart         -8.107312   0.707105  -11.4655 < 2.2e-16 ***
# 7.depart         -8.466583   0.235702  -35.9207 < 2.2e-16 ***
# 8.depart         -7.624461   0.499999  -15.2489 < 2.2e-16 ***
# 9.depart         -8.213118   0.577349  -14.2256 < 2.2e-16 ***
# 10.depart        -8.336652   0.377964  -22.0567 < 2.2e-16 ***
# 11.depart        -6.136106   0.577349  -10.6281 < 2.2e-16 ***
# 12.depart        -8.430566   0.577349  -14.6022 < 2.2e-16 ***
# 13.depart        -6.374262   0.999995   -6.3743 1.838e-10 ***
# 14.depart        -8.432862   0.707105  -11.9259 < 2.2e-16 ***
# 15.depart        -8.472487   0.999994   -8.4725 < 2.2e-16 ***
# 16.depart        -4.553187   0.999997   -4.5532 5.284e-06 ***
# 17.depart        -7.332594   0.999995   -7.3326 2.256e-13 ***
# 18.depart        -6.129319   0.999996   -6.1293 8.824e-10 ***
# 19.depart        -8.897214   0.999994   -8.8973 < 2.2e-16 ***
# 20.depart        -7.676151   0.999995   -7.6762 1.643e-14 ***
# 21.depart        -4.711675   0.999996   -4.7117 2.457e-06 ***
# 22.depart        -7.735384   0.999995   -7.7354 1.021e-14 ***
# 23.depart        -7.017221   0.999995   -7.0173 2.263e-12 ***
# 24.depart        -7.391510   0.999995   -7.3915 1.452e-13 ***
# 25.depart        -7.842174   0.999994   -7.8422 4.441e-15 ***
# 26.depart        -8.035618   0.999994   -8.0357 8.882e-16 ***
# 27.depart        -8.659685   0.999994   -8.6597 < 2.2e-16 ***
# 28.depart        -8.191807   0.999994   -8.1919 2.220e-16 ***
# 29.depart        -7.054816   0.999995   -7.0549 1.728e-12 ***
# 30.depart        -5.737913   0.117851  -48.6878 < 2.2e-16 ***
# ---
# Signif. codes:  0 ‘***’ 0.001 ‘**’ 0.01 ‘*’ 0.05 ‘.’ 0.1 ‘ ’ 1
# Null deviance: 76628.21 on 4854 degrees of freedom
# Residual deviance: 30385.71 on 4637 degrees of freedom
# 	Chi-square: 46242.51 on 217 degrees of freedom, asymptotic p-value 0 
# AIC: 30819.71 AICC: 30840.11 BIC: 32227.51 
# Log posterior: -232405.7 
# Prior parameters: mu=0 sigma=1000 nu=4 

#So it seems like U males are indeed less likely to attack id males than average, more likely to attack each other than average, and then even more likely to attack X males. Also slightly more likely to be attacked by id male than average. 

#does it make sense to try lumping X and U? seems like no, we're getting a lot of model improvement this way

#looks like since I am leaving out X.Uattack, I can use that as my contrast term? yeah 

#but, maybe I still need to leave out a contrast term for the within-ID attack/receives? since otherwise they will be totally double-covered?


#### Last thing to try: Partition out ID males as individual rates within thier category, since I know that is favored over global 


sequences<-table(event.list$Date)

#create list of dummy intercept names. Leave male 30.receives as the baseline- this is the receive rate for ?? males- I expect everyone else's receive rate to be less

idno<-length(event.types[grep("cop.start",event.types)])

dummy.ints.attack<-paste(1:14,".attackFE", sep="")
dummy.ints.receive<-paste(1:13, ".receiveFE", sep="")


int.names<-c("Global.attackFE", "X.attackFE", "X.receiveFE", "X.UreceiveFE", "U.attackFE", "U.receiveFE", "U.UattackFE", dummy.ints.attack, dummy.ints.receive, event.types[-grep("attack", event.types)])

int.list<-list()

for(i in 1:length(levels(event.list$Date))){

day.array<-array(0,dim=c(as.numeric(sequences[i]), length(event.types),length(int.names)))

#the third dimension is the stat slice.
dimnames(day.array)<-list(c(1:sequences[i]),as.character(event.types),as.character(int.names))

#add global attack effect, loops for "U" intercepts for attacks and receives, then one loop for typical intercepts

day.array[,grep("attack", event.types),1]<-1

for(h in 1:14){
day.array[,paste(30,".attack.", h, sep=""),2]<-1
}

for(h in 1:14){
day.array[,paste(h,".attack.",30, sep=""),3]<-1
}


for(h in 15:29){
day.array[,paste(h,".attack.",30, sep=""),4]<-1
}


for(h in 1:14){
	for(j in 15:29){	
day.array[,paste(j,".attack.", h, sep=""),5]<-1
}
}


for(h in 1:14){
	for(j in 15:29){
day.array[,paste(h,".attack.",j, sep=""),6]<-1
}
}

for(h in 15:29){
	for(j in 15:29){	
if(h!=j){
day.array[,paste(j,".attack.", h, sep=""),7]<-1
}
}
}


#loop for dummy intercepts for attacks, loop for dummy intercepts for receives, then one loop for typical intercepts

for(k in 8:(length(dummy.ints.attack)+7)){
	
	for(h in 1:14){
if(h!=(k-7)){
day.array[,paste((k-7),".attack.", h, sep=""),k]<-1
}
}
}

for(k in ((length(dummy.ints.attack)+7)+1):((length(dummy.ints.attack)+7)+length(dummy.ints.receive))){
	for(h in 1:14){
if(h!=(k-(length(dummy.ints.attack)+7))){
day.array[,paste(h,".attack.",(k-(length(dummy.ints.attack)+7)), sep=""),k]<-1
}
}
}


for(k in (((length(dummy.ints.attack)+7)+length(dummy.ints.receive))+1):length(int.names)){
	
	day.array[,int.names[k],k]<-1
	
}

#weird extra formatting to make it match the package example, seems like it becomes relevant *punintended!* if I want to fit day-specific effects

global<-list()

global[[1]]<-day.array

int.list[[i]]<-global

names(int.list[[i]])<-"global"

}

tic()
intfit.sup<-rem(evlist, int.list, supplist=supplist, timing="interval", estimator="BPM")
toc()

summary(intfit.sup)

# Egocentric Relational Event Model (Interval Likelihood)
# 
#                   Post.Mode     Post.SD  Z value  Pr(>|z|)    
# Global.attackFE  -8.4267684   0.7071056 -11.9173 < 2.2e-16 ***
# X.attackFE       -1.2243498   0.7745955  -1.5806 0.1139624    
# X.receiveFE       2.6721020   0.7085414   3.7713 0.0001624 ***
# X.UreceiveFE      2.8033700   0.7282179   3.8496 0.0001183 ***
# U.attackFE       -2.3725613   0.9128695  -2.5990 0.0093492 ** 
# U.receiveFE       0.6885746   0.7180692   0.9589 0.3375965    
# U.UattackFE       1.8184751   0.7226483   2.5164 0.0118559 *  
# 1.attackFE       -0.0661150   0.7564690  -0.0874 0.9303540    
# 2.attackFE        0.1772124   0.7552999   0.2346 0.8144996    
# 3.attackFE        0.5069912   0.7537349   0.6726 0.5011772    
# 4.attackFE        1.4274358   0.7521171   1.8979 0.0577105 .  
# 5.attackFE       -0.7880864   0.7698418  -1.0237 0.3059774    
# 6.attackFE       -1.1198682   1.0316301  -1.0855 0.2776858    
# 7.attackFE        0.2375114   0.7558475   0.3142 0.7533449    
# 8.attackFE       -2.5920725   1.2495106  -2.0745 0.0380357 *  
# 9.attackFE      -13.5622188 234.3929314  -0.0579 0.9538593    
# 10.attackFE       0.4772518   0.7594739   0.6284 0.5297433    
# 11.attackFE     -11.4690404 253.3018603  -0.0453 0.9638856    
# 12.attackFE      -1.1112981   0.8481761  -1.3102 0.1901211    
# 13.attackFE      -0.1138313   1.2509400  -0.0910 0.9274953    
# 14.attackFE      -0.3706679   0.8304254  -0.4464 0.6553379    
# 1.receiveFE      -0.7315767   0.2770235  -2.6408 0.0082699 ** 
# 2.receiveFE      -0.2483197   0.2681291  -0.9261 0.3543836    
# 3.receiveFE      -0.3750742   0.2725414  -1.3762 0.1687566    
# 4.receiveFE       1.0326542   0.2610272   3.9561 7.618e-05 ***
# 5.receiveFE      -0.8724362   0.2925981  -2.9817 0.0028666 ** 
# 6.receiveFE       0.7239195   0.3403190   2.1272 0.0334052 *  
# 7.receiveFE       0.2399256   0.2626679   0.9134 0.3610229    
# 8.receiveFE       0.0322636   0.3395838   0.0950 0.9243074    
# 9.receiveFE       0.6527389   0.3048316   2.1413 0.0322491 *  
# 10.receiveFE     -0.0083206   0.2918527  -0.0285 0.9772557    
# 11.receiveFE      1.6776173   0.4538048   3.6968 0.0002183 ***
# 12.receiveFE     -0.4512319   0.3601767  -1.2528 0.2102760    
# 13.receiveFE     -0.3517499   1.0308038  -0.3412 0.7329241    
# 1.initFO          6.7444301   0.1270001  53.1057 < 2.2e-16 ***
# 2.initFO          6.7001158   0.1048285  63.9150 < 2.2e-16 ***
# 3.initFO          6.7386788   0.1147079  58.7464 < 2.2e-16 ***
# 4.initFO          6.6846117   0.0753778  88.6814 < 2.2e-16 ***
# 5.initFO          6.7926857   0.1561738  43.4944 < 2.2e-16 ***
# 6.initFO          5.0619265   0.5773508   8.7675 < 2.2e-16 ***
# 7.initFO          6.7157484   0.0854358  78.6058 < 2.2e-16 ***
# 8.initFO          5.1159942   0.5000003  10.2320 < 2.2e-16 ***
# 9.initFO          3.3813905   1.0000015   3.3814 0.0007212 ***
# 10.initFO         6.6626326   0.1666667  39.9758 < 2.2e-16 ***
# 11.initFO        -6.7247399 321.8149019  -0.0209 0.9833284    
# 12.initFO         5.8091415   0.4472139  12.9896 < 2.2e-16 ***
# 13.initFO        -5.0627152 363.2563881  -0.0139 0.9888802    
# 14.initFO         5.4608346   0.5000003  10.9217 < 2.2e-16 ***
# 15.initFO         6.0968232   0.5000004  12.1936 < 2.2e-16 ***
# 16.initFO        -6.8411143 319.4179285  -0.0214 0.9829127    
# 17.initFO        -5.6466962 346.9319758  -0.0163 0.9870141    
# 18.initFO        -5.0627152 363.2563881  -0.0139 0.9888802    
# 19.initFO         4.4228431   1.0000021   4.4228 9.741e-06 ***
# 20.initFO         3.5065535   1.0000016   3.5065 0.0004540 ***
# 21.initFO        -5.0627152 363.2563881  -0.0139 0.9888802    
# 22.initFO         0.0000000 894.4271910   0.0000 1.0000000    
# 23.initFO        -5.0627152 363.2563881  -0.0139 0.9888802    
# 24.initFO        -7.5530467 305.8358503  -0.0247 0.9802971    
# 25.initFO         6.2146003   1.0000033   6.2146 5.146e-10 ***
# 26.initFO         6.9077510   0.7071081   9.7690 < 2.2e-16 ***
# 27.initFO         6.0604566   0.3333335  18.1814 < 2.2e-16 ***
# 28.initFO         6.5022861   0.7071080   9.1956 < 2.2e-16 ***
# 29.initFO         6.9077466   1.0000037   6.9077 4.925e-12 ***
# 30.initFO         3.3524069   0.2581989  12.9838 < 2.2e-16 ***
# 1.initCH          2.6172926   1.0000010   2.6173 0.0088631 ** 
# 2.initCH         -9.1855050 280.2587561  -0.0328 0.9738539    
# 3.initCH         -8.9884961 283.0089737  -0.0318 0.9746631    
# 4.initCH          1.5141258   1.0000003   1.5141 0.1299940    
# 5.initCH          3.0791100   1.0000013   3.0791 0.0020762 ** 
# 6.initCH          6.7359045   0.2500001  26.9436 < 2.2e-16 ***
# 7.initCH         -9.5410217 275.4923345  -0.0346 0.9723727    
# 8.initCH          6.5629143   0.2425357  27.0596 < 2.2e-16 ***
# 9.initCH          6.8779021   0.1740777  39.5105 < 2.2e-16 ***
# 10.initCH         4.4654068   0.5000003   8.9308 < 2.2e-16 ***
# 11.initCH         6.7536032   0.4082485  16.5429 < 2.2e-16 ***
# 12.initCH        -7.3922442 308.7518796  -0.0239 0.9808986    
# 13.initCH        -5.0627152 363.2563881  -0.0139 0.9888802    
# 14.initCH         6.2717656   0.3333335  18.8153 < 2.2e-16 ***
# 15.initCH         5.4036745   0.7071078   7.6419 2.132e-14 ***
# 16.initCH         6.9077542   0.3535536  19.5381 < 2.2e-16 ***
# 17.initCH         6.2146003   1.0000033   6.2146 5.146e-10 ***
# 18.initCH         6.9077466   1.0000037   6.9077 4.925e-12 ***
# 19.initCH         5.8091412   0.5000004  11.6183 < 2.2e-16 ***
# 20.initCH         6.5976000   0.2132008  30.9455 < 2.2e-16 ***
# 21.initCH         6.9077466   1.0000037   6.9077 4.925e-12 ***
# 22.initCH         0.0000000 894.4271910   0.0000 1.0000000    
# 23.initCH        -5.0627152 363.2563881  -0.0139 0.9888802    
# 24.initCH         6.8505964   0.2425357  28.2457 < 2.2e-16 ***
# 25.initCH        -5.6466962 346.9319758  -0.0163 0.9870141    
# 26.initCH        -5.6466962 346.9319758  -0.0163 0.9870141    
# 27.initCH         6.0604566   0.3333335  18.1814 < 2.2e-16 ***
# 28.initCH        -5.9926922 338.2402555  -0.0177 0.9858644    
# 29.initCH        -5.0627152 363.2563881  -0.0139 0.9888802    
# 30.initCH         6.8243737   0.0455016 149.9810 < 2.2e-16 ***
# 1.initSF          4.9198803   0.3162278  15.5580 < 2.2e-16 ***
# 2.initSF          5.2337785   0.2182179  23.9842 < 2.2e-16 ***
# 3.initSF          5.0470025   0.2672613  18.8842 < 2.2e-16 ***
# 4.initSF          5.2753277   0.1524986  34.5926 < 2.2e-16 ***
# 5.initSF          4.4654068   0.5000003   8.9308 < 2.2e-16 ***
# 6.initSF         -7.6008103 304.9854879  -0.0249 0.9801173    
# 7.initSF          5.1630631   0.1856954  27.8039 < 2.2e-16 ***
# 8.initSF          4.8283117   0.5773507   8.3629 < 2.2e-16 ***
# 9.initSF         -8.1170250 296.2257322  -0.0274 0.9781395    
# 10.initSF         4.8708723   0.4082485  11.9311 < 2.2e-16 ***
# 11.initSF         4.9618389   1.0000025   4.9618 6.983e-07 ***
# 12.initSF         6.5022894   0.3162279  20.5620 < 2.2e-16 ***
# 13.initSF         6.9077466   1.0000037   6.9077 4.925e-12 ***
# 14.initSF         5.4608346   0.5000003  10.9217 < 2.2e-16 ***
# 15.initSF         5.8091406   0.5773508  10.0617 < 2.2e-16 ***
# 16.initSF        -6.8411143 319.4179285  -0.0214 0.9829127    
# 17.initSF         6.2146003   1.0000033   6.2146 5.146e-10 ***
# 18.initSF        -5.0627152 363.2563881  -0.0139 0.9888802    
# 19.initSF         6.3687576   0.3779647  16.8501 < 2.2e-16 ***
# 20.initSF         5.4524671   0.3779646  14.4259 < 2.2e-16 ***
# 21.initSF        -5.0627152 363.2563881  -0.0139 0.9888802    
# 22.initSF         0.0000000 894.4271910   0.0000 1.0000000    
# 23.initSF         6.9077466   1.0000037   6.9077 4.925e-12 ***
# 24.initSF         4.0173785   1.0000019   4.0174 5.885e-05 ***
# 25.initSF         6.2146003   1.0000033   6.2146 5.146e-10 ***
# 26.initSF        -5.6466962 346.9319758  -0.0163 0.9870141    
# 27.initSF         4.9618431   0.5773507   8.5942 < 2.2e-16 ***
# 28.initSF         5.8091357   1.0000030   5.8091 6.280e-09 ***
# 29.initSF        -5.0627152 363.2563881  -0.0139 0.9888802    
# 30.initSF         3.9401937   0.1924501  20.4738 < 2.2e-16 ***
# 1.disengage      -4.2622836   0.0626224 -68.0632 < 2.2e-16 ***
# 2.disengage      -5.1487697   0.0887356 -58.0237 < 2.2e-16 ***
# 3.disengage      -4.7750140   0.0636285 -75.0452 < 2.2e-16 ***
# 4.disengage      -4.6474529   0.0548821 -84.6806 < 2.2e-16 ***
# 5.disengage      -4.5737155   0.1025978 -44.5791 < 2.2e-16 ***
# 6.disengage      -6.4777863   0.5773494 -11.2199 < 2.2e-16 ***
# 7.disengage      -5.6831776   0.0894427 -63.5399 < 2.2e-16 ***
# 8.disengage      -3.0256324   0.1889822 -16.0101 < 2.2e-16 ***
# 9.disengage      -2.9513586   0.9999975  -2.9514 0.0031637 ** 
# 10.disengage     -4.6109312   0.0880451 -52.3701 < 2.2e-16 ***
# 11.disengage     -1.2519039   0.9999986  -1.2519 0.2106042    
# 12.disengage     -5.0999545   0.3333332 -15.2999 < 2.2e-16 ***
# 13.disengage     -3.0694899   0.9999975  -3.0695 0.0021442 ** 
# 14.disengage     -3.9995581   0.2000000 -19.9978 < 2.2e-16 ***
# 15.disengage     -3.7115624   0.3779643  -9.8199 < 2.2e-16 ***
# 16.disengage      0.0000000 894.4271910   0.0000 1.0000000    
# 17.disengage     -5.5854796   0.7071053  -7.8991 2.887e-15 ***
# 18.disengage      0.0000000 894.4271910   0.0000 1.0000000    
# 19.disengage     -3.3061330   0.2500000 -13.2245 < 2.2e-16 ***
# 20.disengage     -2.4505089   0.3535533  -6.9311 4.176e-12 ***
# 21.disengage      0.0000000 894.4271910   0.0000 1.0000000    
# 22.disengage     -3.5143729   0.9999972  -3.5144 0.0004408 ***
# 23.disengage    -12.1897538 246.2838524  -0.0495 0.9605250    
# 24.disengage     -1.8919041   0.9999982  -1.8919 0.0585033 .  
# 25.disengage     -2.4340658   0.2000000 -12.1703 < 2.2e-16 ***
# 26.disengage     -3.8049157   0.4082481  -9.3201 < 2.2e-16 ***
# 27.disengage     -5.3682937   0.3162276 -16.9760 < 2.2e-16 ***
# 28.disengage     -5.0921384   0.3535532 -14.4027 < 2.2e-16 ***
# 29.disengage     -5.0977870   0.4082480 -12.4870 < 2.2e-16 ***
# 30.disengage     -4.8310314   0.1622214 -29.7805 < 2.2e-16 ***
# 1.cop.start     -21.7752272 187.4297190  -0.1162 0.9075114    
# 2.cop.start      -9.4466055   0.4082478 -23.1394 < 2.2e-16 ***
# 3.cop.start      -7.4311417   0.1490712 -49.8496 < 2.2e-16 ***
# 4.cop.start      -7.8394792   0.2182178 -35.9250 < 2.2e-16 ***
# 5.cop.start     -21.3347068 189.2694639  -0.1127 0.9102515    
# 6.cop.start     -19.2731790 198.6580647  -0.0970 0.9227130    
# 7.cop.start     -21.6639301 187.8895411  -0.1153 0.9082062    
# 8.cop.start     -19.5041052 197.5364378  -0.0987 0.9213473    
# 9.cop.start     -19.2258646 198.8902427  -0.0967 0.9229919    
# 10.cop.start    -20.3974324 193.3698157  -0.1055 0.9159917    
# 11.cop.start    -16.2974335 215.0652505  -0.0758 0.9395949    
# 12.cop.start    -19.2619267 198.7132076  -0.0969 0.9227794    
# 13.cop.start    -17.1246828 210.1006818  -0.0815 0.9350387    
# 14.cop.start    -19.0459187 199.7807571  -0.0953 0.9240495    
# 15.cop.start    -19.0597550 199.7118593  -0.0954 0.9239683    
# 16.cop.start    -15.4105655 220.7992955  -0.0698 0.9443573    
# 17.cop.start    -17.8460741 206.0410091  -0.0866 0.9309782    
# 18.cop.start    -16.8947616 211.4458963  -0.0799 0.9363159    
# 19.cop.start    -19.1978533 199.0280820  -0.0965 0.9231568    
# 20.cop.start    -18.1758355 204.2620316  -0.0890 0.9290955    
# 21.cop.start    -15.5594410 219.8046557  -0.0708 0.9435668    
# 22.cop.start    -18.4146963 203.0018489  -0.0907 0.9277215    
# 23.cop.start    -17.5431931 207.7166170  -0.0845 0.9326928    
# 24.cop.start    -18.0886812 204.7277209  -0.0884 0.9295947    
# 25.cop.start    -18.5160008 202.4743875  -0.0914 0.9271361    
# 26.cop.start    -17.3278154 208.9333269  -0.0829 0.9339035    
# 27.cop.start    -18.8314734 200.8577929  -0.0938 0.9253036    
# 28.cop.start    -18.7176943 201.4363622  -0.0929 0.9259662    
# 29.cop.start      0.0000000 894.4271910   0.0000 1.0000000    
# 30.cop.start    -20.5433232 192.7140797  -0.1066 0.9151063    
# 1.cop.end         0.0000000 894.4271910   0.0000 1.0000000    
# 2.cop.end        -1.6515381   0.7071062  -2.3356 0.0195106 *  
# 3.cop.end        -1.3543802   0.1601281  -8.4581 < 2.2e-16 ***
# 4.cop.end        -1.7377527   0.3333333  -5.2133 1.856e-07 ***
# 5.cop.end         0.0000000 894.4271910   0.0000 1.0000000    
# 6.cop.end         0.0000000 894.4271910   0.0000 1.0000000    
# 7.cop.end         0.0000000 894.4271910   0.0000 1.0000000    
# 8.cop.end         0.0000000 894.4271910   0.0000 1.0000000    
# 9.cop.end         0.0000000 894.4271910   0.0000 1.0000000    
# 10.cop.end        0.0000000 894.4271910   0.0000 1.0000000    
# 11.cop.end        0.0000000 894.4271910   0.0000 1.0000000    
# 12.cop.end        0.0000000 894.4271910   0.0000 1.0000000    
# 13.cop.end        0.0000000 894.4271910   0.0000 1.0000000    
# 14.cop.end        0.0000000 894.4271910   0.0000 1.0000000    
# 15.cop.end        0.0000000 894.4271910   0.0000 1.0000000    
# 16.cop.end        0.0000000 894.4271910   0.0000 1.0000000    
# 17.cop.end        0.0000000 894.4271910   0.0000 1.0000000    
# 18.cop.end        0.0000000 894.4271910   0.0000 1.0000000    
# 19.cop.end        0.0000000 894.4271910   0.0000 1.0000000    
# 20.cop.end        0.0000000 894.4271910   0.0000 1.0000000    
# 21.cop.end        0.0000000 894.4271910   0.0000 1.0000000    
# 22.cop.end        0.0000000 894.4271910   0.0000 1.0000000    
# 23.cop.end        0.0000000 894.4271910   0.0000 1.0000000    
# 24.cop.end        0.0000000 894.4271910   0.0000 1.0000000    
# 25.cop.end        0.0000000 894.4271910   0.0000 1.0000000    
# 26.cop.end        0.0000000 894.4271910   0.0000 1.0000000    
# 27.cop.end        0.0000000 894.4271910   0.0000 1.0000000    
# 28.cop.end        0.0000000 894.4271910   0.0000 1.0000000    
# 29.cop.end        0.0000000 894.4271910   0.0000 1.0000000    
# 30.cop.end        0.0000000 894.4271910   0.0000 1.0000000    
# 1.depart         -8.7014746   0.2357022 -36.9172 < 2.2e-16 ***
# 2.depart         -8.6455924   0.2357022 -36.6802 < 2.2e-16 ***
# 3.depart         -8.6031331   0.2357022 -36.5000 < 2.2e-16 ***
# 4.depart         -8.2131081   0.2357022 -34.8453 < 2.2e-16 ***
# 5.depart         -8.7978526   0.2886750 -30.4767 < 2.2e-16 ***
# 6.depart         -8.1073118   0.7071048 -11.4655 < 2.2e-16 ***
# 7.depart         -8.4665832   0.2357022 -35.9207 < 2.2e-16 ***
# 8.depart         -7.6244610   0.4999993 -15.2489 < 2.2e-16 ***
# 9.depart         -8.2131177   0.5773492 -14.2256 < 2.2e-16 ***
# 10.depart        -8.3366519   0.3779642 -22.0567 < 2.2e-16 ***
# 11.depart        -6.1361062   0.5773494 -10.6281 < 2.2e-16 ***
# 12.depart        -8.4305661   0.5773491 -14.6022 < 2.2e-16 ***
# 13.depart        -6.3742619   0.9999954  -6.3743 1.838e-10 ***
# 14.depart        -8.4328617   0.7071047 -11.9259 < 2.2e-16 ***
# 15.depart        -8.4724875   0.9999941  -8.4725 < 2.2e-16 ***
# 16.depart        -4.5531868   0.9999965  -4.5532 5.284e-06 ***
# 17.depart        -7.3325942   0.9999948  -7.3326 2.256e-13 ***
# 18.depart        -6.1293192   0.9999955  -6.1293 8.824e-10 ***
# 19.depart        -8.8972145   0.9999938  -8.8973 < 2.2e-16 ***
# 20.depart        -7.6761506   0.9999946  -7.6762 1.643e-14 ***
# 21.depart        -4.7116752   0.9999964  -4.7117 2.457e-06 ***
# 22.depart        -7.7353839   0.9999945  -7.7354 1.021e-14 ***
# 23.depart        -7.0172213   0.9999950  -7.0173 2.263e-12 ***
# 24.depart        -7.3915096   0.9999948  -7.3915 1.452e-13 ***
# 25.depart        -7.8421735   0.9999945  -7.8422 4.441e-15 ***
# 26.depart        -8.0356182   0.9999944  -8.0357 8.882e-16 ***
# 27.depart        -8.6596852   0.9999940  -8.6597 < 2.2e-16 ***
# 28.depart        -8.1918068   0.9999943  -8.1919 2.220e-16 ***
# 29.depart        -7.0548164   0.9999950  -7.0549 1.728e-12 ***
# 30.depart        -5.7379132   0.1178511 -48.6878 < 2.2e-16 ***
# ---
# Signif. codes:  0 ‘***’ 0.001 ‘**’ 0.01 ‘*’ 0.05 ‘.’ 0.1 ‘ ’ 1
# Null deviance: 76628.21 on 4854 degrees of freedom
# Residual deviance: 29756.44 on 4610 degrees of freedom
# 	Chi-square: 46871.77 on 244 degrees of freedom, asymptotic p-value 0 
# AIC: 30244.44 AICC: 30270.38 BIC: 31827.4 
# Log posterior: -259117.6 
# Prior parameters: mu=0 sigma=1000 nu=4 

#interestingly, the individual rates seems largely uninformative? They also seem colinear with U.receiveFE and X.attackFE? X attackFE goes away I think because it becomes indistinguishable from the global base rate. U males it seems are not any more/less likely to be attacked than ID males


### Try to refine a bit more- lump ID.attack but keep individual receive rates 

sequences<-table(event.list$Date)

#create list of dummy intercept names. Leave male 30.receives as the baseline- this is the receive rate for ?? males- I expect everyone else's receive rate to be less

idno<-length(event.types[grep("cop.start",event.types)])

#dummy.ints.attack<-paste(1:14,".attackFE", sep="")
dummy.ints.receive<-paste(1:13, ".receiveFE", sep="")


int.names<-c("Global.attackFE", "X.attackFE", "X.receiveFE", "X.UreceiveFE", "U.attackFE", "U.receiveFE", "U.UattackFE", "ID.attackFE", dummy.ints.receive, event.types[-grep("attack", event.types)])

int.list<-list()

for(i in 1:length(levels(event.list$Date))){

day.array<-array(0,dim=c(as.numeric(sequences[i]), length(event.types),length(int.names)))

#the third dimension is the stat slice.
dimnames(day.array)<-list(c(1:sequences[i]),as.character(event.types),as.character(int.names))

#add global attack effect, loops for "U" intercepts for attacks and receives, then one loop for typical intercepts

day.array[,grep("attack", event.types),1]<-1

for(h in 1:14){
day.array[,paste(30,".attack.", h, sep=""),2]<-1
}

for(h in 1:14){
day.array[,paste(h,".attack.",30, sep=""),3]<-1
}


for(h in 15:29){
day.array[,paste(h,".attack.",30, sep=""),4]<-1
}


for(h in 1:14){
	for(j in 15:29){	
day.array[,paste(j,".attack.", h, sep=""),5]<-1
}
}


for(h in 1:14){
	for(j in 15:29){
day.array[,paste(h,".attack.",j, sep=""),6]<-1
}
}

for(h in 15:29){
	for(j in 15:29){	
if(h!=j){
day.array[,paste(j,".attack.", h, sep=""),7]<-1
}
}
}

for(h in 1:14){
	for(j in 1:14){	
if(h!=j){
day.array[,paste(j,".attack.", h, sep=""),8]<-1
}
}
}


#loop for dummy intercepts for receives, then one loop for typical intercepts

for(k in 9:(length(dummy.ints.receive)+8)){
	
	for(h in 1:14){
if(h!=(k-8)){
day.array[,paste(h, ".attack.", (k-8), sep=""),k]<-1
}
}
}


for(k in ((length(dummy.ints.receive)+8)+1):length(int.names)){
	
	day.array[,int.names[k],k]<-1
	
}

#weird extra formatting to make it match the package example, seems like it becomes relevant *punintended!* if I want to fit day-specific effects

global<-list()

global[[1]]<-day.array

int.list[[i]]<-global

names(int.list[[i]])<-"global"

}

tic()
intfit.sup<-rem(evlist, int.list, supplist=supplist, timing="interval", estimator="BPM")
toc()

summary(intfit.sup)

# Egocentric Relational Event Model (Interval Likelihood)
# 
#                  Post.Mode    Post.SD  Z value  Pr(>|z|)    
# Global.attackFE  -8.426750   0.707102 -11.9173 < 2.2e-16 ***
# X.attackFE       -1.224368   0.774592  -1.5807 0.1139556    
# X.receiveFE       2.672084   0.708538   3.7713 0.0001624 ***
# X.UreceiveFE      2.803352   0.728215   3.8496 0.0001183 ***
# U.attackFE       -2.372579   0.912867  -2.5990 0.0093484 ** 
# U.receiveFE       0.688557   0.718066   0.9589 0.3376067    
# U.UattackFE       1.818457   0.722645   2.5164 0.0118564 *  
# ID.attackFE       0.358794   0.749995   0.4784 0.6323690    
# 1.receiveFE      -0.662839   0.276385  -2.3982 0.0164739 *  
# 2.receiveFE      -0.199693   0.267261  -0.7472 0.4549534    
# 3.receiveFE      -0.412247   0.271544  -1.5182 0.1289736    
# 4.receiveFE       0.763206   0.258971   2.9471 0.0032080 ** 
# 5.receiveFE      -0.786770   0.291937  -2.6950 0.0070389 ** 
# 6.receiveFE       0.661084   0.339310   1.9483 0.0513771 .  
# 7.receiveFE       0.297103   0.261771   1.1350 0.2563871    
# 8.receiveFE       0.114674   0.339310   0.3380 0.7353912    
# 9.receiveFE       0.649688   0.304636   2.1327 0.0329516 *  
# 10.receiveFE     -0.052736   0.291071  -0.1812 0.8562268    
# 11.receiveFE      1.305717   0.453163   2.8813 0.0039599 ** 
# 12.receiveFE     -0.481256   0.359398  -1.3391 0.1805501    
# 13.receiveFE     -0.469192   1.030775  -0.4552 0.6489773    
# 1.initFO          6.744430   0.127000  53.1057 < 2.2e-16 ***
# 2.initFO          6.700116   0.104828  63.9150 < 2.2e-16 ***
# 3.initFO          6.738679   0.114708  58.7464 < 2.2e-16 ***
# 4.initFO          6.684612   0.075378  88.6814 < 2.2e-16 ***
# 5.initFO          6.792686   0.156174  43.4944 < 2.2e-16 ***
# 6.initFO          5.061926   0.577351   8.7675 < 2.2e-16 ***
# 7.initFO          6.715748   0.085436  78.6058 < 2.2e-16 ***
# 8.initFO          5.115994   0.500000  10.2320 < 2.2e-16 ***
# 9.initFO          3.381391   1.000001   3.3814 0.0007212 ***
# 10.initFO         6.662633   0.166667  39.9758 < 2.2e-16 ***
# 11.initFO        -6.724740 321.814902  -0.0209 0.9833284    
# 12.initFO         5.809142   0.447214  12.9896 < 2.2e-16 ***
# 13.initFO        -5.062715 363.256388  -0.0139 0.9888802    
# 14.initFO         5.460835   0.500000  10.9217 < 2.2e-16 ***
# 15.initFO         6.096823   0.500000  12.1936 < 2.2e-16 ***
# 16.initFO        -6.841114 319.417929  -0.0214 0.9829127    
# 17.initFO        -5.646696 346.931976  -0.0163 0.9870141    
# 18.initFO        -5.062715 363.256388  -0.0139 0.9888802    
# 19.initFO         4.422843   1.000002   4.4228 9.741e-06 ***
# 20.initFO         3.506554   1.000002   3.5065 0.0004540 ***
# 21.initFO        -5.062715 363.256388  -0.0139 0.9888802    
# 22.initFO         0.000000 894.427191   0.0000 1.0000000    
# 23.initFO        -5.062715 363.256388  -0.0139 0.9888802    
# 24.initFO        -7.553047 305.835850  -0.0247 0.9802971    
# 25.initFO         6.214600   1.000003   6.2146 5.146e-10 ***
# 26.initFO         6.907751   0.707108   9.7690 < 2.2e-16 ***
# 27.initFO         6.060457   0.333333  18.1814 < 2.2e-16 ***
# 28.initFO         6.502286   0.707108   9.1956 < 2.2e-16 ***
# 29.initFO         6.907747   1.000004   6.9077 4.925e-12 ***
# 30.initFO         3.352407   0.258199  12.9838 < 2.2e-16 ***
# 1.initCH          2.617293   1.000001   2.6173 0.0088631 ** 
# 2.initCH         -9.185505 280.258756  -0.0328 0.9738539    
# 3.initCH         -8.988496 283.008974  -0.0318 0.9746631    
# 4.initCH          1.514126   1.000000   1.5141 0.1299940    
# 5.initCH          3.079110   1.000001   3.0791 0.0020762 ** 
# 6.initCH          6.735904   0.250000  26.9436 < 2.2e-16 ***
# 7.initCH         -9.541022 275.492334  -0.0346 0.9723727    
# 8.initCH          6.562914   0.242536  27.0596 < 2.2e-16 ***
# 9.initCH          6.877902   0.174078  39.5105 < 2.2e-16 ***
# 10.initCH         4.465407   0.500000   8.9308 < 2.2e-16 ***
# 11.initCH         6.753603   0.408249  16.5429 < 2.2e-16 ***
# 12.initCH        -7.392244 308.751880  -0.0239 0.9808986    
# 13.initCH        -5.062715 363.256388  -0.0139 0.9888802    
# 14.initCH         6.271766   0.333333  18.8153 < 2.2e-16 ***
# 15.initCH         5.403675   0.707108   7.6419 2.132e-14 ***
# 16.initCH         6.907754   0.353554  19.5381 < 2.2e-16 ***
# 17.initCH         6.214600   1.000003   6.2146 5.146e-10 ***
# 18.initCH         6.907747   1.000004   6.9077 4.925e-12 ***
# 19.initCH         5.809141   0.500000  11.6183 < 2.2e-16 ***
# 20.initCH         6.597600   0.213201  30.9455 < 2.2e-16 ***
# 21.initCH         6.907747   1.000004   6.9077 4.925e-12 ***
# 22.initCH         0.000000 894.427191   0.0000 1.0000000    
# 23.initCH        -5.062715 363.256388  -0.0139 0.9888802    
# 24.initCH         6.850596   0.242536  28.2457 < 2.2e-16 ***
# 25.initCH        -5.646696 346.931976  -0.0163 0.9870141    
# 26.initCH        -5.646696 346.931976  -0.0163 0.9870141    
# 27.initCH         6.060457   0.333333  18.1814 < 2.2e-16 ***
# 28.initCH        -5.992692 338.240255  -0.0177 0.9858644    
# 29.initCH        -5.062715 363.256388  -0.0139 0.9888802    
# 30.initCH         6.824374   0.045502 149.9810 < 2.2e-16 ***
# 1.initSF          4.919880   0.316228  15.5580 < 2.2e-16 ***
# 2.initSF          5.233779   0.218218  23.9842 < 2.2e-16 ***
# 3.initSF          5.047002   0.267261  18.8842 < 2.2e-16 ***
# 4.initSF          5.275328   0.152499  34.5926 < 2.2e-16 ***
# 5.initSF          4.465407   0.500000   8.9308 < 2.2e-16 ***
# 6.initSF         -7.600810 304.985488  -0.0249 0.9801173    
# 7.initSF          5.163063   0.185695  27.8039 < 2.2e-16 ***
# 8.initSF          4.828312   0.577351   8.3629 < 2.2e-16 ***
# 9.initSF         -8.117025 296.225732  -0.0274 0.9781395    
# 10.initSF         4.870872   0.408248  11.9311 < 2.2e-16 ***
# 11.initSF         4.961839   1.000002   4.9618 6.983e-07 ***
# 12.initSF         6.502289   0.316228  20.5620 < 2.2e-16 ***
# 13.initSF         6.907747   1.000004   6.9077 4.925e-12 ***
# 14.initSF         5.460835   0.500000  10.9217 < 2.2e-16 ***
# 15.initSF         5.809141   0.577351  10.0617 < 2.2e-16 ***
# 16.initSF        -6.841114 319.417929  -0.0214 0.9829127    
# 17.initSF         6.214600   1.000003   6.2146 5.146e-10 ***
# 18.initSF        -5.062715 363.256388  -0.0139 0.9888802    
# 19.initSF         6.368758   0.377965  16.8501 < 2.2e-16 ***
# 20.initSF         5.452467   0.377965  14.4259 < 2.2e-16 ***
# 21.initSF        -5.062715 363.256388  -0.0139 0.9888802    
# 22.initSF         0.000000 894.427191   0.0000 1.0000000    
# 23.initSF         6.907747   1.000004   6.9077 4.925e-12 ***
# 24.initSF         4.017378   1.000002   4.0174 5.885e-05 ***
# 25.initSF         6.214600   1.000003   6.2146 5.146e-10 ***
# 26.initSF        -5.646696 346.931976  -0.0163 0.9870141    
# 27.initSF         4.961843   0.577351   8.5942 < 2.2e-16 ***
# 28.initSF         5.809136   1.000003   5.8091 6.280e-09 ***
# 29.initSF        -5.062715 363.256388  -0.0139 0.9888802    
# 30.initSF         3.940194   0.192450  20.4738 < 2.2e-16 ***
# 1.disengage      -4.262284   0.062622 -68.0632 < 2.2e-16 ***
# 2.disengage      -5.148770   0.088736 -58.0237 < 2.2e-16 ***
# 3.disengage      -4.775014   0.063628 -75.0452 < 2.2e-16 ***
# 4.disengage      -4.647453   0.054882 -84.6806 < 2.2e-16 ***
# 5.disengage      -4.573716   0.102598 -44.5791 < 2.2e-16 ***
# 6.disengage      -6.477786   0.577349 -11.2199 < 2.2e-16 ***
# 7.disengage      -5.683178   0.089443 -63.5399 < 2.2e-16 ***
# 8.disengage      -3.025632   0.188982 -16.0101 < 2.2e-16 ***
# 9.disengage      -2.951359   0.999998  -2.9514 0.0031637 ** 
# 10.disengage     -4.610931   0.088045 -52.3701 < 2.2e-16 ***
# 11.disengage     -1.251904   0.999999  -1.2519 0.2106042    
# 12.disengage     -5.099954   0.333333 -15.2999 < 2.2e-16 ***
# 13.disengage     -3.069490   0.999997  -3.0695 0.0021442 ** 
# 14.disengage     -3.999558   0.200000 -19.9978 < 2.2e-16 ***
# 15.disengage     -3.711562   0.377964  -9.8199 < 2.2e-16 ***
# 16.disengage      0.000000 894.427191   0.0000 1.0000000    
# 17.disengage     -5.585480   0.707105  -7.8991 2.887e-15 ***
# 18.disengage      0.000000 894.427191   0.0000 1.0000000    
# 19.disengage     -3.306133   0.250000 -13.2245 < 2.2e-16 ***
# 20.disengage     -2.450509   0.353553  -6.9311 4.176e-12 ***
# 21.disengage      0.000000 894.427191   0.0000 1.0000000    
# 22.disengage     -3.514373   0.999997  -3.5144 0.0004408 ***
# 23.disengage    -12.189754 246.283852  -0.0495 0.9605250    
# 24.disengage     -1.891904   0.999998  -1.8919 0.0585033 .  
# 25.disengage     -2.434066   0.200000 -12.1703 < 2.2e-16 ***
# 26.disengage     -3.804916   0.408248  -9.3201 < 2.2e-16 ***
# 27.disengage     -5.368294   0.316228 -16.9760 < 2.2e-16 ***
# 28.disengage     -5.092138   0.353553 -14.4027 < 2.2e-16 ***
# 29.disengage     -5.097787   0.408248 -12.4870 < 2.2e-16 ***
# 30.disengage     -4.831031   0.162221 -29.7805 < 2.2e-16 ***
# 1.cop.start     -21.775230 187.429988  -0.1162 0.9075115    
# 2.cop.start      -9.446606   0.408248 -23.1394 < 2.2e-16 ***
# 3.cop.start      -7.431142   0.149071 -49.8496 < 2.2e-16 ***
# 4.cop.start      -7.839479   0.218218 -35.9250 < 2.2e-16 ***
# 5.cop.start     -21.334708 189.269565  -0.1127 0.9102515    
# 6.cop.start     -19.273179 198.658066  -0.0970 0.9227130    
# 7.cop.start     -21.663932 187.889751  -0.1153 0.9082063    
# 8.cop.start     -19.504105 197.536440  -0.0987 0.9213473    
# 9.cop.start     -19.225865 198.890244  -0.0967 0.9229919    
# 10.cop.start    -20.397433 193.369830  -0.1055 0.9159917    
# 11.cop.start    -16.297433 215.065251  -0.0758 0.9395949    
# 12.cop.start    -19.261927 198.713209  -0.0969 0.9227794    
# 13.cop.start    -17.124683 210.100682  -0.0815 0.9350387    
# 14.cop.start    -19.045919 199.780758  -0.0953 0.9240495    
# 15.cop.start    -19.059755 199.711860  -0.0954 0.9239683    
# 16.cop.start    -15.410565 220.799295  -0.0698 0.9443573    
# 17.cop.start    -17.846074 206.041009  -0.0866 0.9309782    
# 18.cop.start    -16.894762 211.445896  -0.0799 0.9363159    
# 19.cop.start    -19.197853 199.028083  -0.0965 0.9231568    
# 20.cop.start    -18.175836 204.262032  -0.0890 0.9290955    
# 21.cop.start    -15.559441 219.804656  -0.0708 0.9435668    
# 22.cop.start    -18.414696 203.001849  -0.0907 0.9277215    
# 23.cop.start    -17.543193 207.716617  -0.0845 0.9326928    
# 24.cop.start    -18.088681 204.727721  -0.0884 0.9295947    
# 25.cop.start    -18.516001 202.474388  -0.0914 0.9271361    
# 26.cop.start    -17.327815 208.933327  -0.0829 0.9339035    
# 27.cop.start    -18.831473 200.857794  -0.0938 0.9253036    
# 28.cop.start    -18.717694 201.436363  -0.0929 0.9259662    
# 29.cop.start      0.000000 894.427191   0.0000 1.0000000    
# 30.cop.start    -20.543323 192.714099  -0.1066 0.9151063    
# 1.cop.end         0.000000 894.427191   0.0000 1.0000000    
# 2.cop.end        -1.651538   0.707106  -2.3356 0.0195106 *  
# 3.cop.end        -1.354380   0.160128  -8.4581 < 2.2e-16 ***
# 4.cop.end        -1.737753   0.333333  -5.2133 1.856e-07 ***
# 5.cop.end         0.000000 894.427191   0.0000 1.0000000    
# 6.cop.end         0.000000 894.427191   0.0000 1.0000000    
# 7.cop.end         0.000000 894.427191   0.0000 1.0000000    
# 8.cop.end         0.000000 894.427191   0.0000 1.0000000    
# 9.cop.end         0.000000 894.427191   0.0000 1.0000000    
# 10.cop.end        0.000000 894.427191   0.0000 1.0000000    
# 11.cop.end        0.000000 894.427191   0.0000 1.0000000    
# 12.cop.end        0.000000 894.427191   0.0000 1.0000000    
# 13.cop.end        0.000000 894.427191   0.0000 1.0000000    
# 14.cop.end        0.000000 894.427191   0.0000 1.0000000    
# 15.cop.end        0.000000 894.427191   0.0000 1.0000000    
# 16.cop.end        0.000000 894.427191   0.0000 1.0000000    
# 17.cop.end        0.000000 894.427191   0.0000 1.0000000    
# 18.cop.end        0.000000 894.427191   0.0000 1.0000000    
# 19.cop.end        0.000000 894.427191   0.0000 1.0000000    
# 20.cop.end        0.000000 894.427191   0.0000 1.0000000    
# 21.cop.end        0.000000 894.427191   0.0000 1.0000000    
# 22.cop.end        0.000000 894.427191   0.0000 1.0000000    
# 23.cop.end        0.000000 894.427191   0.0000 1.0000000    
# 24.cop.end        0.000000 894.427191   0.0000 1.0000000    
# 25.cop.end        0.000000 894.427191   0.0000 1.0000000    
# 26.cop.end        0.000000 894.427191   0.0000 1.0000000    
# 27.cop.end        0.000000 894.427191   0.0000 1.0000000    
# 28.cop.end        0.000000 894.427191   0.0000 1.0000000    
# 29.cop.end        0.000000 894.427191   0.0000 1.0000000    
# 30.cop.end        0.000000 894.427191   0.0000 1.0000000    
# 1.depart         -8.701475   0.235702 -36.9172 < 2.2e-16 ***
# 2.depart         -8.645592   0.235702 -36.6802 < 2.2e-16 ***
# 3.depart         -8.603133   0.235702 -36.5000 < 2.2e-16 ***
# 4.depart         -8.213108   0.235702 -34.8453 < 2.2e-16 ***
# 5.depart         -8.797853   0.288675 -30.4767 < 2.2e-16 ***
# 6.depart         -8.107312   0.707105 -11.4655 < 2.2e-16 ***
# 7.depart         -8.466583   0.235702 -35.9207 < 2.2e-16 ***
# 8.depart         -7.624461   0.499999 -15.2489 < 2.2e-16 ***
# 9.depart         -8.213118   0.577349 -14.2256 < 2.2e-16 ***
# 10.depart        -8.336652   0.377964 -22.0567 < 2.2e-16 ***
# 11.depart        -6.136106   0.577349 -10.6281 < 2.2e-16 ***
# 12.depart        -8.430566   0.577349 -14.6022 < 2.2e-16 ***
# 13.depart        -6.374262   0.999995  -6.3743 1.838e-10 ***
# 14.depart        -8.432862   0.707105 -11.9259 < 2.2e-16 ***
# 15.depart        -8.472487   0.999994  -8.4725 < 2.2e-16 ***
# 16.depart        -4.553187   0.999997  -4.5532 5.284e-06 ***
# 17.depart        -7.332594   0.999995  -7.3326 2.256e-13 ***
# 18.depart        -6.129319   0.999996  -6.1293 8.824e-10 ***
# 19.depart        -8.897214   0.999994  -8.8973 < 2.2e-16 ***
# 20.depart        -7.676151   0.999995  -7.6762 1.643e-14 ***
# 21.depart        -4.711675   0.999996  -4.7117 2.457e-06 ***
# 22.depart        -7.735384   0.999995  -7.7354 1.021e-14 ***
# 23.depart        -7.017221   0.999995  -7.0173 2.263e-12 ***
# 24.depart        -7.391510   0.999995  -7.3915 1.452e-13 ***
# 25.depart        -7.842174   0.999994  -7.8422 4.441e-15 ***
# 26.depart        -8.035618   0.999994  -8.0357 8.882e-16 ***
# 27.depart        -8.659685   0.999994  -8.6597 < 2.2e-16 ***
# 28.depart        -8.191807   0.999994  -8.1919 2.220e-16 ***
# 29.depart        -7.054816   0.999995  -7.0549 1.728e-12 ***
# 30.depart        -5.737913   0.117851 -48.6878 < 2.2e-16 ***
# ---
# Signif. codes:  0 ‘***’ 0.001 ‘**’ 0.01 ‘*’ 0.05 ‘.’ 0.1 ‘ ’ 1
# Null deviance: 76628.21 on 4854 degrees of freedom
# Residual deviance: 30142.71 on 4623 degrees of freedom
# 	Chi-square: 46485.5 on 231 degrees of freedom, asymptotic p-value 0 
# AIC: 30604.71 AICC: 30627.9 BIC: 32103.34 
# Log posterior: -246297.9 
# Prior parameters: mu=0 sigma=1000 nu=4


#dang thats a lot worse! Looks like individual rates for ID males are still preferred, which is not surprising since it was before, and now I am accounting for the other rates


###Try cleaning up the non-significant FE's still in there- X.attack  

#This should basically cause Global Rate to represent the base attack rate for X males 


sequences<-table(event.list$Date)

#create list of dummy intercept names

idno<-length(event.types[grep("cop.start",event.types)])

dummy.ints.attack<-paste(1:14,".attackFE", sep="")
dummy.ints.receive<-paste(1:13, ".receiveFE", sep="")


int.names<-c("Global.attackFE", "X.receiveFE", "X.UreceiveFE", "U.attackFE", "U.receiveFE", "U.UattackFE", dummy.ints.attack, dummy.ints.receive, event.types[-grep("attack", event.types)])

int.list<-list()

for(i in 1:length(levels(event.list$Date))){

day.array<-array(0,dim=c(as.numeric(sequences[i]), length(event.types),length(int.names)))

#the third dimension is the stat slice.
dimnames(day.array)<-list(c(1:sequences[i]),as.character(event.types),as.character(int.names))

#add global attack effect, loops for "U" intercepts for attacks and receives, then one loop for typical intercepts

day.array[,grep("attack", event.types),1]<-1


for(h in 1:14){
day.array[,paste(h,".attack.",30, sep=""),2]<-1
}


for(h in 15:29){
day.array[,paste(h,".attack.",30, sep=""),3]<-1
}


for(h in 1:14){
	for(j in 15:29){	
day.array[,paste(j,".attack.", h, sep=""),4]<-1
}
}


for(h in 1:14){
	for(j in 15:29){
day.array[,paste(h,".attack.",j, sep=""),5]<-1
}
}

for(h in 15:29){
	for(j in 15:29){	
if(h!=j){
day.array[,paste(j,".attack.", h, sep=""),6]<-1
}
}
}

preloop<-6

#loop for dummy intercepts for attacks, loop for dummy intercepts for receives, then one loop for typical intercepts

for(k in (preloop+1):(length(dummy.ints.attack)+preloop)){
	
	for(h in 1:14){
if(h!=(k-preloop)){
day.array[,paste((k-preloop),".attack.", h, sep=""),k]<-1
}
}
}

for(k in ((length(dummy.ints.attack)+preloop)+1):((length(dummy.ints.attack)+preloop)+length(dummy.ints.receive))){
	for(h in 1:14){
if(h!=(k-(length(dummy.ints.attack)+preloop))){
day.array[,paste(h,".attack.",(k-(length(dummy.ints.attack)+preloop)), sep=""),k]<-1
}
}
}


for(k in (((length(dummy.ints.attack)+preloop)+length(dummy.ints.receive))+1):length(int.names)){
	
	day.array[,int.names[k],k]<-1
	
}

#weird extra formatting to make it match the package example, seems like it becomes relevant *punintended!* if I want to fit day-specific effects

global<-list()

global[[1]]<-day.array

int.list[[i]]<-global

names(int.list[[i]])<-"global"

}

tic()
intfit.sup<-rem(evlist, int.list, supplist=supplist, timing="interval", estimator="BPM")
toc()
alarm()

summary(intfit.sup)

# Egocentric Relational Event Model (Interval Likelihood)
# 
#                   Post.Mode     Post.SD  Z value  Pr(>|z|)    
# Global.attackFE  -9.5259220   0.2886747 -32.9988 < 2.2e-16 ***
# X.receiveFE       3.7712555   0.2921739  12.9076 < 2.2e-16 ***
# X.UreceiveFE      3.9025235   0.3370996  11.5768 < 2.2e-16 ***
# U.attackFE       -1.2734082   0.6454967  -1.9728 0.0485233 *  
# U.receiveFE       1.7877281   0.3145761   5.6830 1.324e-08 ***
# U.UattackFE       2.9176286   0.3248928   8.9803 < 2.2e-16 ***
# 1.attackFE        1.0330374   0.3944376   2.6190 0.0088184 ** 
# 2.attackFE        1.2763648   0.3921908   3.2544 0.0011361 ** 
# 3.attackFE        1.6061436   0.3891684   4.1271 3.673e-05 ***
# 4.attackFE        2.5265882   0.3860257   6.5451 5.944e-11 ***
# 5.attackFE        0.3110660   0.4195142   0.7415 0.4583958    
# 6.attackFE       -0.0207165   0.8047343  -0.0257 0.9794621    
# 7.attackFE        1.3366638   0.3932445   3.3991 0.0006762 ***
# 8.attackFE       -1.4929214   1.0698668  -1.3954 0.1628870    
# 9.attackFE      -12.5413190 243.0663151  -0.0516 0.9588504    
# 10.attackFE       1.5764042   0.4001704   3.9393 8.171e-05 ***
# 11.attackFE     -10.4618029 264.1957859  -0.0396 0.9684131    
# 12.attackFE      -0.0121458   0.5502168  -0.0221 0.9823884    
# 13.attackFE       0.9853197   1.0715358   0.9195 0.3578134    
# 14.attackFE       0.7284843   0.5224387   1.3944 0.1631993    
# 1.receiveFE      -0.7315755   0.2770237  -2.6408 0.0082700 ** 
# 2.receiveFE      -0.2483185   0.2681292  -0.9261 0.3543862    
# 3.receiveFE      -0.3750730   0.2725416  -1.3762 0.1687582    
# 4.receiveFE       1.0326554   0.2610273   3.9561 7.618e-05 ***
# 5.receiveFE      -0.8724350   0.2925982  -2.9817 0.0028667 ** 
# 6.receiveFE       0.7239207   0.3403191   2.1272 0.0334050 *  
# 7.receiveFE       0.2399268   0.2626681   0.9134 0.3610207    
# 8.receiveFE       0.0322648   0.3395839   0.0950 0.9243046    
# 9.receiveFE       0.6527401   0.3048317   2.1413 0.0322488 *  
# 10.receiveFE     -0.0083194   0.2918528  -0.0285 0.9772590    
# 11.receiveFE      1.6776185   0.4538049   3.6968 0.0002183 ***
# 12.receiveFE     -0.4512307   0.3601768  -1.2528 0.2102774    
# 13.receiveFE     -0.3517487   1.0308039  -0.3412 0.7329249    
# 1.initFO          6.7444301   0.1270001  53.1057 < 2.2e-16 ***
# 2.initFO          6.7001158   0.1048285  63.9150 < 2.2e-16 ***
# 3.initFO          6.7386788   0.1147079  58.7464 < 2.2e-16 ***
# 4.initFO          6.6846117   0.0753778  88.6814 < 2.2e-16 ***
# 5.initFO          6.7926857   0.1561738  43.4944 < 2.2e-16 ***
# 6.initFO          5.0619265   0.5773508   8.7675 < 2.2e-16 ***
# 7.initFO          6.7157484   0.0854358  78.6058 < 2.2e-16 ***
# 8.initFO          5.1159942   0.5000003  10.2320 < 2.2e-16 ***
# 9.initFO          3.3813905   1.0000015   3.3814 0.0007212 ***
# 10.initFO         6.6626326   0.1666667  39.9758 < 2.2e-16 ***
# 11.initFO        -6.7247399 321.8149019  -0.0209 0.9833284    
# 12.initFO         5.8091415   0.4472139  12.9896 < 2.2e-16 ***
# 13.initFO        -5.0627152 363.2563881  -0.0139 0.9888802    
# 14.initFO         5.4608346   0.5000003  10.9217 < 2.2e-16 ***
# 15.initFO         6.0968232   0.5000004  12.1936 < 2.2e-16 ***
# 16.initFO        -6.8411143 319.4179285  -0.0214 0.9829127    
# 17.initFO        -5.6466962 346.9319758  -0.0163 0.9870141    
# 18.initFO        -5.0627152 363.2563881  -0.0139 0.9888802    
# 19.initFO         4.4228431   1.0000021   4.4228 9.741e-06 ***
# 20.initFO         3.5065535   1.0000016   3.5065 0.0004540 ***
# 21.initFO        -5.0627152 363.2563881  -0.0139 0.9888802    
# 22.initFO         0.0000000 894.4271910   0.0000 1.0000000    
# 23.initFO        -5.0627152 363.2563881  -0.0139 0.9888802    
# 24.initFO        -7.5530467 305.8358503  -0.0247 0.9802971    
# 25.initFO         6.2146003   1.0000033   6.2146 5.146e-10 ***
# 26.initFO         6.9077510   0.7071081   9.7690 < 2.2e-16 ***
# 27.initFO         6.0604566   0.3333335  18.1814 < 2.2e-16 ***
# 28.initFO         6.5022861   0.7071080   9.1956 < 2.2e-16 ***
# 29.initFO         6.9077466   1.0000037   6.9077 4.925e-12 ***
# 30.initFO         3.3524069   0.2581989  12.9838 < 2.2e-16 ***
# 1.initCH          2.6172926   1.0000010   2.6173 0.0088631 ** 
# 2.initCH         -9.1855050 280.2587561  -0.0328 0.9738539    
# 3.initCH         -8.9884961 283.0089737  -0.0318 0.9746631    
# 4.initCH          1.5141258   1.0000003   1.5141 0.1299940    
# 5.initCH          3.0791100   1.0000013   3.0791 0.0020762 ** 
# 6.initCH          6.7359045   0.2500001  26.9436 < 2.2e-16 ***
# 7.initCH         -9.5410217 275.4923345  -0.0346 0.9723727    
# 8.initCH          6.5629143   0.2425357  27.0596 < 2.2e-16 ***
# 9.initCH          6.8779021   0.1740777  39.5105 < 2.2e-16 ***
# 10.initCH         4.4654068   0.5000003   8.9308 < 2.2e-16 ***
# 11.initCH         6.7536032   0.4082485  16.5429 < 2.2e-16 ***
# 12.initCH        -7.3922442 308.7518796  -0.0239 0.9808986    
# 13.initCH        -5.0627152 363.2563881  -0.0139 0.9888802    
# 14.initCH         6.2717656   0.3333335  18.8153 < 2.2e-16 ***
# 15.initCH         5.4036745   0.7071078   7.6419 2.132e-14 ***
# 16.initCH         6.9077542   0.3535536  19.5381 < 2.2e-16 ***
# 17.initCH         6.2146003   1.0000033   6.2146 5.146e-10 ***
# 18.initCH         6.9077466   1.0000037   6.9077 4.925e-12 ***
# 19.initCH         5.8091412   0.5000004  11.6183 < 2.2e-16 ***
# 20.initCH         6.5976000   0.2132008  30.9455 < 2.2e-16 ***
# 21.initCH         6.9077466   1.0000037   6.9077 4.925e-12 ***
# 22.initCH         0.0000000 894.4271910   0.0000 1.0000000    
# 23.initCH        -5.0627152 363.2563881  -0.0139 0.9888802    
# 24.initCH         6.8505964   0.2425357  28.2457 < 2.2e-16 ***
# 25.initCH        -5.6466962 346.9319758  -0.0163 0.9870141    
# 26.initCH        -5.6466962 346.9319758  -0.0163 0.9870141    
# 27.initCH         6.0604566   0.3333335  18.1814 < 2.2e-16 ***
# 28.initCH        -5.9926922 338.2402555  -0.0177 0.9858644    
# 29.initCH        -5.0627152 363.2563881  -0.0139 0.9888802    
# 30.initCH         6.8243737   0.0455016 149.9810 < 2.2e-16 ***
# 1.initSF          4.9198803   0.3162278  15.5580 < 2.2e-16 ***
# 2.initSF          5.2337785   0.2182179  23.9842 < 2.2e-16 ***
# 3.initSF          5.0470025   0.2672613  18.8842 < 2.2e-16 ***
# 4.initSF          5.2753277   0.1524986  34.5926 < 2.2e-16 ***
# 5.initSF          4.4654068   0.5000003   8.9308 < 2.2e-16 ***
# 6.initSF         -7.6008103 304.9854879  -0.0249 0.9801173    
# 7.initSF          5.1630631   0.1856954  27.8039 < 2.2e-16 ***
# 8.initSF          4.8283117   0.5773507   8.3629 < 2.2e-16 ***
# 9.initSF         -8.1170250 296.2257322  -0.0274 0.9781395    
# 10.initSF         4.8708723   0.4082485  11.9311 < 2.2e-16 ***
# 11.initSF         4.9618389   1.0000025   4.9618 6.983e-07 ***
# 12.initSF         6.5022894   0.3162279  20.5620 < 2.2e-16 ***
# 13.initSF         6.9077466   1.0000037   6.9077 4.925e-12 ***
# 14.initSF         5.4608346   0.5000003  10.9217 < 2.2e-16 ***
# 15.initSF         5.8091406   0.5773508  10.0617 < 2.2e-16 ***
# 16.initSF        -6.8411143 319.4179285  -0.0214 0.9829127    
# 17.initSF         6.2146003   1.0000033   6.2146 5.146e-10 ***
# 18.initSF        -5.0627152 363.2563881  -0.0139 0.9888802    
# 19.initSF         6.3687576   0.3779647  16.8501 < 2.2e-16 ***
# 20.initSF         5.4524671   0.3779646  14.4259 < 2.2e-16 ***
# 21.initSF        -5.0627152 363.2563881  -0.0139 0.9888802    
# 22.initSF         0.0000000 894.4271910   0.0000 1.0000000    
# 23.initSF         6.9077466   1.0000037   6.9077 4.925e-12 ***
# 24.initSF         4.0173785   1.0000019   4.0174 5.885e-05 ***
# 25.initSF         6.2146003   1.0000033   6.2146 5.146e-10 ***
# 26.initSF        -5.6466962 346.9319758  -0.0163 0.9870141    
# 27.initSF         4.9618431   0.5773507   8.5942 < 2.2e-16 ***
# 28.initSF         5.8091357   1.0000030   5.8091 6.280e-09 ***
# 29.initSF        -5.0627152 363.2563881  -0.0139 0.9888802    
# 30.initSF         3.9401937   0.1924501  20.4738 < 2.2e-16 ***
# 1.disengage      -4.2622836   0.0626224 -68.0632 < 2.2e-16 ***
# 2.disengage      -5.1487697   0.0887356 -58.0237 < 2.2e-16 ***
# 3.disengage      -4.7750140   0.0636285 -75.0452 < 2.2e-16 ***
# 4.disengage      -4.6474529   0.0548821 -84.6806 < 2.2e-16 ***
# 5.disengage      -4.5737155   0.1025978 -44.5791 < 2.2e-16 ***
# 6.disengage      -6.4777863   0.5773494 -11.2199 < 2.2e-16 ***
# 7.disengage      -5.6831776   0.0894427 -63.5399 < 2.2e-16 ***
# 8.disengage      -3.0256324   0.1889822 -16.0101 < 2.2e-16 ***
# 9.disengage      -2.9513586   0.9999975  -2.9514 0.0031637 ** 
# 10.disengage     -4.6109312   0.0880451 -52.3701 < 2.2e-16 ***
# 11.disengage     -1.2519039   0.9999986  -1.2519 0.2106042    
# 12.disengage     -5.0999545   0.3333332 -15.2999 < 2.2e-16 ***
# 13.disengage     -3.0694899   0.9999975  -3.0695 0.0021442 ** 
# 14.disengage     -3.9995581   0.2000000 -19.9978 < 2.2e-16 ***
# 15.disengage     -3.7115624   0.3779643  -9.8199 < 2.2e-16 ***
# 16.disengage      0.0000000 894.4271910   0.0000 1.0000000    
# 17.disengage     -5.5854796   0.7071053  -7.8991 2.887e-15 ***
# 18.disengage      0.0000000 894.4271910   0.0000 1.0000000    
# 19.disengage     -3.3061330   0.2500000 -13.2245 < 2.2e-16 ***
# 20.disengage     -2.4505089   0.3535533  -6.9311 4.176e-12 ***
# 21.disengage      0.0000000 894.4271910   0.0000 1.0000000    
# 22.disengage     -3.5143729   0.9999972  -3.5144 0.0004408 ***
# 23.disengage    -12.1897538 246.2838524  -0.0495 0.9605250    
# 24.disengage     -1.8919041   0.9999982  -1.8919 0.0585033 .  
# 25.disengage     -2.4340658   0.2000000 -12.1703 < 2.2e-16 ***
# 26.disengage     -3.8049157   0.4082481  -9.3201 < 2.2e-16 ***
# 27.disengage     -5.3682937   0.3162276 -16.9760 < 2.2e-16 ***
# 28.disengage     -5.0921384   0.3535532 -14.4027 < 2.2e-16 ***
# 29.disengage     -5.0977870   0.4082480 -12.4870 < 2.2e-16 ***
# 30.disengage     -4.8310314   0.1622214 -29.7805 < 2.2e-16 ***
# 1.cop.start     -21.7752266 187.4296704  -0.1162 0.9075114    
# 2.cop.start      -9.4466055   0.4082478 -23.1394 < 2.2e-16 ***
# 3.cop.start      -7.4311417   0.1490712 -49.8496 < 2.2e-16 ***
# 4.cop.start      -7.8394792   0.2182178 -35.9250 < 2.2e-16 ***
# 5.cop.start     -21.3347065 189.2694410  -0.1127 0.9102515    
# 6.cop.start     -19.2731790 198.6580640  -0.0970 0.9227130    
# 7.cop.start     -21.6639297 187.8895009  -0.1153 0.9082062    
# 8.cop.start     -19.5041051 197.5364367  -0.0987 0.9213473    
# 9.cop.start     -19.2258646 198.8902420  -0.0967 0.9229919    
# 10.cop.start    -20.3974323 193.3698111  -0.1055 0.9159917    
# 11.cop.start    -16.2974335 215.0652505  -0.0758 0.9395949    
# 12.cop.start    -19.2619266 198.7132069  -0.0969 0.9227794    
# 13.cop.start    -17.1246828 210.1006818  -0.0815 0.9350387    
# 14.cop.start    -19.0459187 199.7807566  -0.0953 0.9240495    
# 15.cop.start    -19.0597550 199.7118588  -0.0954 0.9239683    
# 16.cop.start    -15.4105655 220.7992955  -0.0698 0.9443573    
# 17.cop.start    -17.8460741 206.0410090  -0.0866 0.9309782    
# 18.cop.start    -16.8947616 211.4458963  -0.0799 0.9363159    
# 19.cop.start    -19.1978533 199.0280814  -0.0965 0.9231568    
# 20.cop.start    -18.1758355 204.2620314  -0.0890 0.9290955    
# 21.cop.start    -15.5594410 219.8046557  -0.0708 0.9435668    
# 22.cop.start    -18.4146963 203.0018487  -0.0907 0.9277215    
# 23.cop.start    -17.5431931 207.7166170  -0.0845 0.9326928    
# 24.cop.start    -18.0886812 204.7277208  -0.0884 0.9295947    
# 25.cop.start    -18.5160008 202.4743873  -0.0914 0.9271361    
# 26.cop.start    -17.3278154 208.9333268  -0.0829 0.9339035    
# 27.cop.start    -18.8314734 200.8577925  -0.0938 0.9253036    
# 28.cop.start    -18.7176943 201.4363619  -0.0929 0.9259662    
# 29.cop.start      0.0000000 894.4271910   0.0000 1.0000000    
# 30.cop.start    -20.5433232 192.7140737  -0.1066 0.9151063    
# 1.cop.end         0.0000000 894.4271910   0.0000 1.0000000    
# 2.cop.end        -1.6515381   0.7071062  -2.3356 0.0195106 *  
# 3.cop.end        -1.3543802   0.1601281  -8.4581 < 2.2e-16 ***
# 4.cop.end        -1.7377527   0.3333333  -5.2133 1.856e-07 ***
# 5.cop.end         0.0000000 894.4271910   0.0000 1.0000000    
# 6.cop.end         0.0000000 894.4271910   0.0000 1.0000000    
# 7.cop.end         0.0000000 894.4271910   0.0000 1.0000000    
# 8.cop.end         0.0000000 894.4271910   0.0000 1.0000000    
# 9.cop.end         0.0000000 894.4271910   0.0000 1.0000000    
# 10.cop.end        0.0000000 894.4271910   0.0000 1.0000000    
# 11.cop.end        0.0000000 894.4271910   0.0000 1.0000000    
# 12.cop.end        0.0000000 894.4271910   0.0000 1.0000000    
# 13.cop.end        0.0000000 894.4271910   0.0000 1.0000000    
# 14.cop.end        0.0000000 894.4271910   0.0000 1.0000000    
# 15.cop.end        0.0000000 894.4271910   0.0000 1.0000000    
# 16.cop.end        0.0000000 894.4271910   0.0000 1.0000000    
# 17.cop.end        0.0000000 894.4271910   0.0000 1.0000000    
# 18.cop.end        0.0000000 894.4271910   0.0000 1.0000000    
# 19.cop.end        0.0000000 894.4271910   0.0000 1.0000000    
# 20.cop.end        0.0000000 894.4271910   0.0000 1.0000000    
# 21.cop.end        0.0000000 894.4271910   0.0000 1.0000000    
# 22.cop.end        0.0000000 894.4271910   0.0000 1.0000000    
# 23.cop.end        0.0000000 894.4271910   0.0000 1.0000000    
# 24.cop.end        0.0000000 894.4271910   0.0000 1.0000000    
# 25.cop.end        0.0000000 894.4271910   0.0000 1.0000000    
# 26.cop.end        0.0000000 894.4271910   0.0000 1.0000000    
# 27.cop.end        0.0000000 894.4271910   0.0000 1.0000000    
# 28.cop.end        0.0000000 894.4271910   0.0000 1.0000000    
# 29.cop.end        0.0000000 894.4271910   0.0000 1.0000000    
# 30.cop.end        0.0000000 894.4271910   0.0000 1.0000000    
# 1.depart         -8.7014746   0.2357022 -36.9172 < 2.2e-16 ***
# 2.depart         -8.6455924   0.2357022 -36.6802 < 2.2e-16 ***
# 3.depart         -8.6031331   0.2357022 -36.5000 < 2.2e-16 ***
# 4.depart         -8.2131081   0.2357022 -34.8453 < 2.2e-16 ***
# 5.depart         -8.7978526   0.2886750 -30.4767 < 2.2e-16 ***
# 6.depart         -8.1073118   0.7071048 -11.4655 < 2.2e-16 ***
# 7.depart         -8.4665832   0.2357022 -35.9207 < 2.2e-16 ***
# 8.depart         -7.6244610   0.4999993 -15.2489 < 2.2e-16 ***
# 9.depart         -8.2131177   0.5773492 -14.2256 < 2.2e-16 ***
# 10.depart        -8.3366519   0.3779642 -22.0567 < 2.2e-16 ***
# 11.depart        -6.1361062   0.5773494 -10.6281 < 2.2e-16 ***
# 12.depart        -8.4305661   0.5773491 -14.6022 < 2.2e-16 ***
# 13.depart        -6.3742619   0.9999954  -6.3743 1.838e-10 ***
# 14.depart        -8.4328617   0.7071047 -11.9259 < 2.2e-16 ***
# 15.depart        -8.4724875   0.9999941  -8.4725 < 2.2e-16 ***
# 16.depart        -4.5531868   0.9999965  -4.5532 5.284e-06 ***
# 17.depart        -7.3325942   0.9999948  -7.3326 2.256e-13 ***
# 18.depart        -6.1293192   0.9999955  -6.1293 8.824e-10 ***
# 19.depart        -8.8972145   0.9999938  -8.8973 < 2.2e-16 ***
# 20.depart        -7.6761506   0.9999946  -7.6762 1.643e-14 ***
# 21.depart        -4.7116752   0.9999964  -4.7117 2.457e-06 ***
# 22.depart        -7.7353839   0.9999945  -7.7354 1.021e-14 ***
# 23.depart        -7.0172213   0.9999950  -7.0173 2.263e-12 ***
# 24.depart        -7.3915096   0.9999948  -7.3915 1.452e-13 ***
# 25.depart        -7.8421735   0.9999945  -7.8422 4.441e-15 ***
# 26.depart        -8.0356182   0.9999944  -8.0357 8.882e-16 ***
# 27.depart        -8.6596852   0.9999940  -8.6597 < 2.2e-16 ***
# 28.depart        -8.1918068   0.9999943  -8.1919 2.220e-16 ***
# 29.depart        -7.0548164   0.9999950  -7.0549 1.728e-12 ***
# 30.depart        -5.7379132   0.1178511 -48.6878 < 2.2e-16 ***
# ---
# Signif. codes:  0 ‘***’ 0.001 ‘**’ 0.01 ‘*’ 0.05 ‘.’ 0.1 ‘ ’ 1
# Null deviance: 76628.21 on 4854 degrees of freedom
# Residual deviance: 29758.33 on 4611 degrees of freedom
# 	Chi-square: 46869.88 on 243 degrees of freedom, asymptotic p-value 0 
# AIC: 30244.33 AICC: 30270.05 BIC: 31820.81 
# Log posterior: -258117.5 
# Prior parameters: mu=0 sigma=1000 nu=4 

#The Global Rate in this model represents X.attack+X.Uattack rates (which were the same). Keep in mind that U.Uattack also encompasses "Ureceive"

##########################################
## BEST I CAN DO FOR ATTACK INTERCEPTS ##
##########################################


#Other non-attack related Intercept models


############################################


##### cop.start and cop.end #### 

##try a global cop.start and cop.end rate instead of fitting one for each male

#I have also added an ID.cop.start rate to differentiate between base rases of solicitation for the average male vs. a territory-holding male

sequences<-table(event.list$Date)

#create list of dummy intercept names

idno<-length(event.types[grep("cop.start",event.types)])

dummy.ints.attack<-paste(1:14,".attackFE", sep="")
dummy.ints.receive<-paste(1:13, ".receiveFE", sep="")


int.names<-c("Global.attackFE", "X.receiveFE", "X.UreceiveFE", "U.attackFE", "U.receiveFE", "U.UattackFE", dummy.ints.attack, dummy.ints.receive, "cop.startFE","cop.endFE", event.types[-c(grep("attack", event.types),grep("cop", event.types))])

int.list<-list()

for(i in 1:length(levels(event.list$Date))){

day.array<-array(0,dim=c(as.numeric(sequences[i]), length(event.types),length(int.names)))

#the third dimension is the stat slice.
dimnames(day.array)<-list(c(1:sequences[i]),as.character(event.types),as.character(int.names))

#add global attack effect, loops for "U" intercepts for attacks and receives, then one loop for typical intercepts

day.array[,grep("attack", event.types),1]<-1


for(h in 1:14){
day.array[,paste(h,".attack.",30, sep=""),2]<-1
}


for(h in 15:29){
day.array[,paste(h,".attack.",30, sep=""),3]<-1
}


for(h in 1:14){
	for(j in 15:29){	
day.array[,paste(j,".attack.", h, sep=""),4]<-1
}
}


for(h in 1:14){
	for(j in 15:29){
day.array[,paste(h,".attack.",j, sep=""),5]<-1
}
}

for(h in 15:29){
	for(j in 15:29){	
if(h!=j){
day.array[,paste(j,".attack.", h, sep=""),6]<-1
}
}
}

preloop<-6

#loop for dummy intercepts for attacks, loop for dummy intercepts for receives, add in cop.start and cop.end FEs, then one loop for typical intercepts

for(k in (preloop+1):(length(dummy.ints.attack)+preloop)){
	
	for(h in 1:14){
if(h!=(k-preloop)){
day.array[,paste((k-preloop),".attack.", h, sep=""),k]<-1
}
}
}

for(k in ((length(dummy.ints.attack)+preloop)+1):((length(dummy.ints.attack)+preloop)+length(dummy.ints.receive))){
	for(h in 1:14){
if(h!=(k-(length(dummy.ints.attack)+preloop))){
day.array[,paste(h,".attack.",(k-(length(dummy.ints.attack)+preloop)), sep=""),k]<-1
}
}
}

#cop.startFE
for(h in 1:idno){
day.array[,paste(h,".cop.start", sep=""),((length(dummy.ints.attack)+preloop)+length(dummy.ints.receive)+1)]<-1
}

#cop.endFE
for(h in 1:idno){
day.array[,paste(h,".cop.end", sep=""),((length(dummy.ints.attack)+preloop)+length(dummy.ints.receive)+2)]<-1
}

for(k in (((length(dummy.ints.attack)+preloop)+length(dummy.ints.receive)+3):length(int.names))){
	
	day.array[,int.names[k],k]<-1
	
}

#weird extra formatting to make it match the package example, seems like it becomes relevant *punintended!* if I want to fit day-specific effects

global<-list()

global[[1]]<-day.array

int.list[[i]]<-global

names(int.list[[i]])<-"global"

}

tic()
intfit.sup<-rem(evlist, int.list, supplist=supplist, timing="interval", estimator="BPM")
toc()

summary(intfit.sup)


# Egocentric Relational Event Model (Interval Likelihood)
# 
#                   Post.Mode     Post.SD  Z value  Pr(>|z|)    
# Global.attackFE  -9.5259220   0.2886747 -32.9988 < 2.2e-16 ***
# X.receiveFE       3.7712555   0.2921739  12.9076 < 2.2e-16 ***
# X.UreceiveFE      3.9025235   0.3370996  11.5768 < 2.2e-16 ***
# U.attackFE       -1.2734082   0.6454967  -1.9728 0.0485233 *  
# U.receiveFE       1.7877281   0.3145761   5.6830 1.324e-08 ***
# U.UattackFE       2.9176286   0.3248928   8.9803 < 2.2e-16 ***
# 1.attackFE        1.0330374   0.3944376   2.6190 0.0088184 ** 
# 2.attackFE        1.2763648   0.3921908   3.2544 0.0011361 ** 
# 3.attackFE        1.6061436   0.3891684   4.1271 3.673e-05 ***
# 4.attackFE        2.5265882   0.3860257   6.5451 5.944e-11 ***
# 5.attackFE        0.3110660   0.4195142   0.7415 0.4583958    
# 6.attackFE       -0.0207165   0.8047343  -0.0257 0.9794621    
# 7.attackFE        1.3366638   0.3932445   3.3991 0.0006762 ***
# 8.attackFE       -1.4929214   1.0698668  -1.3954 0.1628870    
# 9.attackFE      -12.5407387 243.0010039  -0.0516 0.9588412    
# 10.attackFE       1.5764042   0.4001704   3.9393 8.171e-05 ***
# 11.attackFE     -10.4618029 264.1957839  -0.0396 0.9684131    
# 12.attackFE      -0.0121458   0.5502168  -0.0221 0.9823884    
# 13.attackFE       0.9853197   1.0715358   0.9195 0.3578134    
# 14.attackFE       0.7284843   0.5224387   1.3944 0.1631993    
# 1.receiveFE      -0.7315755   0.2770237  -2.6408 0.0082700 ** 
# 2.receiveFE      -0.2483185   0.2681292  -0.9261 0.3543862    
# 3.receiveFE      -0.3750730   0.2725416  -1.3762 0.1687582    
# 4.receiveFE       1.0326554   0.2610273   3.9561 7.618e-05 ***
# 5.receiveFE      -0.8724350   0.2925982  -2.9817 0.0028667 ** 
# 6.receiveFE       0.7239207   0.3403191   2.1272 0.0334050 *  
# 7.receiveFE       0.2399268   0.2626681   0.9134 0.3610207    
# 8.receiveFE       0.0322648   0.3395839   0.0950 0.9243046    
# 9.receiveFE       0.6527401   0.3048317   2.1413 0.0322488 *  
# 10.receiveFE     -0.0083194   0.2918528  -0.0285 0.9772590    
# 11.receiveFE      1.6776185   0.4538049   3.6968 0.0002183 ***
# 12.receiveFE     -0.4512307   0.3601768  -1.2528 0.2102774    
# 13.receiveFE     -0.3517487   1.0308039  -0.3412 0.7329249    
# cop.startFE      -8.8434147   0.1178511 -75.0389 < 2.2e-16 ***
# cop.endFE        -1.4478174   0.1414214 -10.2376 < 2.2e-16 ***
# 1.initFO          6.7444301   0.1270001  53.1057 < 2.2e-16 ***
# 2.initFO          6.7001158   0.1048285  63.9150 < 2.2e-16 ***
# 3.initFO          6.7386788   0.1147079  58.7464 < 2.2e-16 ***
# 4.initFO          6.6846117   0.0753778  88.6814 < 2.2e-16 ***
# 5.initFO          6.7926857   0.1561738  43.4944 < 2.2e-16 ***
# 6.initFO          5.0619265   0.5773508   8.7675 < 2.2e-16 ***
# 7.initFO          6.7157484   0.0854358  78.6058 < 2.2e-16 ***
# 8.initFO          5.1159942   0.5000003  10.2320 < 2.2e-16 ***
# 9.initFO          3.3813905   1.0000015   3.3814 0.0007212 ***
# 10.initFO         6.6626326   0.1666667  39.9758 < 2.2e-16 ***
# 11.initFO        -6.7247399 321.8149019  -0.0209 0.9833284    
# 12.initFO         5.8091415   0.4472139  12.9896 < 2.2e-16 ***
# 13.initFO        -5.0627152 363.2563881  -0.0139 0.9888802    
# 14.initFO         5.4608346   0.5000003  10.9217 < 2.2e-16 ***
# 15.initFO         6.0968232   0.5000004  12.1936 < 2.2e-16 ***
# 16.initFO        -6.8411143 319.4179285  -0.0214 0.9829127    
# 17.initFO        -5.6466962 346.9319758  -0.0163 0.9870141    
# 18.initFO        -5.0627152 363.2563881  -0.0139 0.9888802    
# 19.initFO         4.4228431   1.0000021   4.4228 9.741e-06 ***
# 20.initFO         3.5065535   1.0000016   3.5065 0.0004540 ***
# 21.initFO        -5.0627152 363.2563881  -0.0139 0.9888802    
# 22.initFO         0.0000000 894.4271910   0.0000 1.0000000    
# 23.initFO        -5.0627152 363.2563881  -0.0139 0.9888802    
# 24.initFO        -7.5530467 305.8358503  -0.0247 0.9802971    
# 25.initFO         6.2146003   1.0000033   6.2146 5.146e-10 ***
# 26.initFO         6.9077510   0.7071081   9.7690 < 2.2e-16 ***
# 27.initFO         6.0604566   0.3333335  18.1814 < 2.2e-16 ***
# 28.initFO         6.5022861   0.7071080   9.1956 < 2.2e-16 ***
# 29.initFO         6.9077466   1.0000037   6.9077 4.925e-12 ***
# 30.initFO         3.3524069   0.2581989  12.9838 < 2.2e-16 ***
# 1.initCH          2.6172926   1.0000010   2.6173 0.0088631 ** 
# 2.initCH         -9.1855050 280.2587561  -0.0328 0.9738539    
# 3.initCH         -8.9884961 283.0089737  -0.0318 0.9746631    
# 4.initCH          1.5141258   1.0000003   1.5141 0.1299940    
# 5.initCH          3.0791100   1.0000013   3.0791 0.0020762 ** 
# 6.initCH          6.7359045   0.2500001  26.9436 < 2.2e-16 ***
# 7.initCH         -9.5410217 275.4923345  -0.0346 0.9723727    
# 8.initCH          6.5629143   0.2425357  27.0596 < 2.2e-16 ***
# 9.initCH          6.8779021   0.1740777  39.5105 < 2.2e-16 ***
# 10.initCH         4.4654068   0.5000003   8.9308 < 2.2e-16 ***
# 11.initCH         6.7536032   0.4082485  16.5429 < 2.2e-16 ***
# 12.initCH        -7.3922442 308.7518796  -0.0239 0.9808986    
# 13.initCH        -5.0627152 363.2563881  -0.0139 0.9888802    
# 14.initCH         6.2717656   0.3333335  18.8153 < 2.2e-16 ***
# 15.initCH         5.4036745   0.7071078   7.6419 2.132e-14 ***
# 16.initCH         6.9077542   0.3535536  19.5381 < 2.2e-16 ***
# 17.initCH         6.2146003   1.0000033   6.2146 5.146e-10 ***
# 18.initCH         6.9077466   1.0000037   6.9077 4.925e-12 ***
# 19.initCH         5.8091412   0.5000004  11.6183 < 2.2e-16 ***
# 20.initCH         6.5976000   0.2132008  30.9455 < 2.2e-16 ***
# 21.initCH         6.9077466   1.0000037   6.9077 4.925e-12 ***
# 22.initCH         0.0000000 894.4271910   0.0000 1.0000000    
# 23.initCH        -5.0627152 363.2563881  -0.0139 0.9888802    
# 24.initCH         6.8505964   0.2425357  28.2457 < 2.2e-16 ***
# 25.initCH        -5.6466962 346.9319758  -0.0163 0.9870141    
# 26.initCH        -5.6466962 346.9319758  -0.0163 0.9870141    
# 27.initCH         6.0604566   0.3333335  18.1814 < 2.2e-16 ***
# 28.initCH        -5.9926922 338.2402555  -0.0177 0.9858644    
# 29.initCH        -5.0627152 363.2563881  -0.0139 0.9888802    
# 30.initCH         6.8243737   0.0455016 149.9810 < 2.2e-16 ***
# 1.initSF          4.9198803   0.3162278  15.5580 < 2.2e-16 ***
# 2.initSF          5.2337785   0.2182179  23.9842 < 2.2e-16 ***
# 3.initSF          5.0470025   0.2672613  18.8842 < 2.2e-16 ***
# 4.initSF          5.2753277   0.1524986  34.5926 < 2.2e-16 ***
# 5.initSF          4.4654068   0.5000003   8.9308 < 2.2e-16 ***
# 6.initSF         -7.6008103 304.9854879  -0.0249 0.9801173    
# 7.initSF          5.1630631   0.1856954  27.8039 < 2.2e-16 ***
# 8.initSF          4.8283117   0.5773507   8.3629 < 2.2e-16 ***
# 9.initSF         -8.1170250 296.2257322  -0.0274 0.9781395    
# 10.initSF         4.8708723   0.4082485  11.9311 < 2.2e-16 ***
# 11.initSF         4.9618389   1.0000025   4.9618 6.983e-07 ***
# 12.initSF         6.5022894   0.3162279  20.5620 < 2.2e-16 ***
# 13.initSF         6.9077466   1.0000037   6.9077 4.925e-12 ***
# 14.initSF         5.4608346   0.5000003  10.9217 < 2.2e-16 ***
# 15.initSF         5.8091406   0.5773508  10.0617 < 2.2e-16 ***
# 16.initSF        -6.8411143 319.4179285  -0.0214 0.9829127    
# 17.initSF         6.2146003   1.0000033   6.2146 5.146e-10 ***
# 18.initSF        -5.0627152 363.2563881  -0.0139 0.9888802    
# 19.initSF         6.3687576   0.3779647  16.8501 < 2.2e-16 ***
# 20.initSF         5.4524671   0.3779646  14.4259 < 2.2e-16 ***
# 21.initSF        -5.0627152 363.2563881  -0.0139 0.9888802    
# 22.initSF         0.0000000 894.4271910   0.0000 1.0000000    
# 23.initSF         6.9077466   1.0000037   6.9077 4.925e-12 ***
# 24.initSF         4.0173785   1.0000019   4.0174 5.885e-05 ***
# 25.initSF         6.2146003   1.0000033   6.2146 5.146e-10 ***
# 26.initSF        -5.6466962 346.9319758  -0.0163 0.9870141    
# 27.initSF         4.9618431   0.5773507   8.5942 < 2.2e-16 ***
# 28.initSF         5.8091357   1.0000030   5.8091 6.280e-09 ***
# 29.initSF        -5.0627152 363.2563881  -0.0139 0.9888802    
# 30.initSF         3.9401937   0.1924501  20.4738 < 2.2e-16 ***
# 1.disengage      -4.2622836   0.0626224 -68.0632 < 2.2e-16 ***
# 2.disengage      -5.1487697   0.0887356 -58.0237 < 2.2e-16 ***
# 3.disengage      -4.7750140   0.0636285 -75.0452 < 2.2e-16 ***
# 4.disengage      -4.6474529   0.0548821 -84.6806 < 2.2e-16 ***
# 5.disengage      -4.5737155   0.1025978 -44.5791 < 2.2e-16 ***
# 6.disengage      -6.4777863   0.5773494 -11.2199 < 2.2e-16 ***
# 7.disengage      -5.6831776   0.0894427 -63.5399 < 2.2e-16 ***
# 8.disengage      -3.0256324   0.1889822 -16.0101 < 2.2e-16 ***
# 9.disengage      -2.9513586   0.9999975  -2.9514 0.0031637 ** 
# 10.disengage     -4.6109312   0.0880451 -52.3701 < 2.2e-16 ***
# 11.disengage     -1.2519039   0.9999986  -1.2519 0.2106042    
# 12.disengage     -5.0999545   0.3333332 -15.2999 < 2.2e-16 ***
# 13.disengage     -3.0694899   0.9999975  -3.0695 0.0021442 ** 
# 14.disengage     -3.9995581   0.2000000 -19.9978 < 2.2e-16 ***
# 15.disengage     -3.7115624   0.3779643  -9.8199 < 2.2e-16 ***
# 16.disengage      0.0000000 894.4271910   0.0000 1.0000000    
# 17.disengage     -5.5854796   0.7071053  -7.8991 2.887e-15 ***
# 18.disengage      0.0000000 894.4271910   0.0000 1.0000000    
# 19.disengage     -3.3061330   0.2500000 -13.2245 < 2.2e-16 ***
# 20.disengage     -2.4505089   0.3535533  -6.9311 4.176e-12 ***
# 21.disengage      0.0000000 894.4271910   0.0000 1.0000000    
# 22.disengage     -3.5143729   0.9999972  -3.5144 0.0004408 ***
# 23.disengage    -12.1897533 246.2837949  -0.0495 0.9605250    
# 24.disengage     -1.8919041   0.9999982  -1.8919 0.0585033 .  
# 25.disengage     -2.4340658   0.2000000 -12.1703 < 2.2e-16 ***
# 26.disengage     -3.8049157   0.4082481  -9.3201 < 2.2e-16 ***
# 27.disengage     -5.3682937   0.3162276 -16.9760 < 2.2e-16 ***
# 28.disengage     -5.0921384   0.3535532 -14.4027 < 2.2e-16 ***
# 29.disengage     -5.0977870   0.4082480 -12.4870 < 2.2e-16 ***
# 30.disengage     -4.8310314   0.1622214 -29.7805 < 2.2e-16 ***
# 1.depart         -8.7014746   0.2357022 -36.9172 < 2.2e-16 ***
# 2.depart         -8.6455924   0.2357022 -36.6802 < 2.2e-16 ***
# 3.depart         -8.6031331   0.2357022 -36.5000 < 2.2e-16 ***
# 4.depart         -8.2131081   0.2357022 -34.8453 < 2.2e-16 ***
# 5.depart         -8.7978526   0.2886750 -30.4767 < 2.2e-16 ***
# 6.depart         -8.1073118   0.7071048 -11.4655 < 2.2e-16 ***
# 7.depart         -8.4665832   0.2357022 -35.9207 < 2.2e-16 ***
# 8.depart         -7.6244610   0.4999993 -15.2489 < 2.2e-16 ***
# 9.depart         -8.2131177   0.5773492 -14.2256 < 2.2e-16 ***
# 10.depart        -8.3366519   0.3779642 -22.0567 < 2.2e-16 ***
# 11.depart        -6.1361062   0.5773494 -10.6281 < 2.2e-16 ***
# 12.depart        -8.4305661   0.5773491 -14.6022 < 2.2e-16 ***
# 13.depart        -6.3742619   0.9999954  -6.3743 1.838e-10 ***
# 14.depart        -8.4328617   0.7071047 -11.9259 < 2.2e-16 ***
# 15.depart        -8.4724875   0.9999941  -8.4725 < 2.2e-16 ***
# 16.depart        -4.5531868   0.9999965  -4.5532 5.284e-06 ***
# 17.depart        -7.3325942   0.9999948  -7.3326 2.256e-13 ***
# 18.depart        -6.1293192   0.9999955  -6.1293 8.824e-10 ***
# 19.depart        -8.8972145   0.9999938  -8.8973 < 2.2e-16 ***
# 20.depart        -7.6761506   0.9999946  -7.6762 1.643e-14 ***
# 21.depart        -4.7116752   0.9999964  -4.7117 2.457e-06 ***
# 22.depart        -7.7353839   0.9999945  -7.7354 1.021e-14 ***
# 23.depart        -7.0172213   0.9999950  -7.0173 2.263e-12 ***
# 24.depart        -7.3915096   0.9999948  -7.3915 1.452e-13 ***
# 25.depart        -7.8421735   0.9999945  -7.8422 4.441e-15 ***
# 26.depart        -8.0356182   0.9999944  -8.0357 8.882e-16 ***
# 27.depart        -8.6596852   0.9999940  -8.6597 < 2.2e-16 ***
# 28.depart        -8.1918068   0.9999943  -8.1919 2.220e-16 ***
# 29.depart        -7.0548164   0.9999950  -7.0549 1.728e-12 ***
# 30.depart        -5.7379132   0.1178511 -48.6878 < 2.2e-16 ***
# ---
# Signif. codes:  0 ‘***’ 0.001 ‘**’ 0.01 ‘*’ 0.05 ‘.’ 0.1 ‘ ’ 1
# Null deviance: 76628.21 on 4854 degrees of freedom
# Residual deviance: 29921.62 on 4669 degrees of freedom
# 	Chi-square: 46706.6 on 185 degrees of freedom, asymptotic p-value 0 
# AIC: 30291.62 AICC: 30306.36 BIC: 31491.81 
# Log posterior: -200142.3 
# Prior parameters: mu=0 sigma=1000 nu=4 

##yep this model performs the best so far! cool. 


########### initFO ###############


sequences<-table(event.list$Date)

#create list of dummy intercept names

idno<-length(event.types[grep("cop.start",event.types)])

dummy.ints.attack<-paste(1:14,".attackFE", sep="")
dummy.ints.receive<-paste(1:13, ".receiveFE", sep="")


int.names<-c("Global.attackFE", "X.receiveFE", "X.UreceiveFE", "U.attackFE", "U.receiveFE", "U.UattackFE", dummy.ints.attack, dummy.ints.receive, "cop.startFE", "cop.endFE", "initFO.FE", event.types[-c(grep("attack", event.types),grep("cop", event.types),grep("initFO", event.types))])

int.list<-list()

for(i in 1:length(levels(event.list$Date))){

day.array<-array(0,dim=c(as.numeric(sequences[i]), length(event.types),length(int.names)))

#the third dimension is the stat slice.
dimnames(day.array)<-list(c(1:sequences[i]),as.character(event.types),as.character(int.names))

#add global attack effect, loops for "U" intercepts for attacks and receives, then one loop for typical intercepts

day.array[,grep("attack", event.types),1]<-1


for(h in 1:14){
day.array[,paste(h,".attack.",30, sep=""),2]<-1
}


for(h in 15:29){
day.array[,paste(h,".attack.",30, sep=""),3]<-1
}


for(h in 1:14){
	for(j in 15:29){	
day.array[,paste(j,".attack.", h, sep=""),4]<-1
}
}


for(h in 1:14){
	for(j in 15:29){
day.array[,paste(h,".attack.",j, sep=""),5]<-1
}
}

for(h in 15:29){
	for(j in 15:29){	
if(h!=j){
day.array[,paste(j,".attack.", h, sep=""),6]<-1
}
}
}

preloop<-6

#loop for dummy intercepts for attacks, loop for dummy intercepts for receives, add in cop.start and cop.end FEs, then one loop for typical intercepts

for(k in (preloop+1):(length(dummy.ints.attack)+preloop)){
	
	for(h in 1:14){
if(h!=(k-preloop)){
day.array[,paste((k-preloop),".attack.", h, sep=""),k]<-1
}
}
}

for(k in ((length(dummy.ints.attack)+preloop)+1):((length(dummy.ints.attack)+preloop)+length(dummy.ints.receive))){
	for(h in 1:14){
if(h!=(k-(length(dummy.ints.attack)+preloop))){
day.array[,paste(h,".attack.",(k-(length(dummy.ints.attack)+preloop)), sep=""),k]<-1
}
}
}

#cop.startFE
for(h in 1:idno){
day.array[,paste(h,".cop.start", sep=""),((length(dummy.ints.attack)+preloop)+length(dummy.ints.receive)+1)]<-1
}


#cop.endFE
for(h in 1:idno){
day.array[,paste(h,".cop.end", sep=""),((length(dummy.ints.attack)+preloop)+length(dummy.ints.receive)+2)]<-1
}

#initFO.FE
for(h in 1:idno){
day.array[,paste(h,".initFO", sep=""),((length(dummy.ints.attack)+preloop)+length(dummy.ints.receive)+3)]<-1
}


for(k in (((length(dummy.ints.attack)+preloop)+length(dummy.ints.receive)+4):length(int.names))){
	
	day.array[,int.names[k],k]<-1
	
}

#weird extra formatting to make it match the package example, seems like it becomes relevant *punintended!* if I want to fit day-specific effects

global<-list()

global[[1]]<-day.array

int.list[[i]]<-global

names(int.list[[i]])<-"global"

}


tic()
intfit.sup<-rem(evlist, int.list, supplist=supplist, timing="interval", estimator="BPM")
toc()

summary(intfit.sup)

# Egocentric Relational Event Model (Interval Likelihood)
# 
#                   Post.Mode     Post.SD  Z value  Pr(>|z|)    
# Global.attackFE  -9.5259220   0.2886747 -32.9988 < 2.2e-16 ***
# X.receiveFE       3.7712555   0.2921739  12.9076 < 2.2e-16 ***
# X.UreceiveFE      3.9025235   0.3370996  11.5768 < 2.2e-16 ***
# U.attackFE       -1.2734082   0.6454967  -1.9728 0.0485233 *  
# U.receiveFE       1.7877281   0.3145761   5.6830 1.324e-08 ***
# U.UattackFE       2.9176286   0.3248928   8.9803 < 2.2e-16 ***
# 1.attackFE        1.0330374   0.3944376   2.6190 0.0088184 ** 
# 2.attackFE        1.2763648   0.3921908   3.2544 0.0011361 ** 
# 3.attackFE        1.6061436   0.3891684   4.1271 3.673e-05 ***
# 4.attackFE        2.5265882   0.3860257   6.5451 5.944e-11 ***
# 5.attackFE        0.3110660   0.4195142   0.7415 0.4583958    
# 6.attackFE       -0.0207165   0.8047343  -0.0257 0.9794621    
# 7.attackFE        1.3366638   0.3932445   3.3991 0.0006762 ***
# 8.attackFE       -1.4929214   1.0698668  -1.3954 0.1628870    
# 9.attackFE      -12.5411760 243.0502196  -0.0516 0.9588481    
# 10.attackFE       1.5764042   0.4001704   3.9393 8.171e-05 ***
# 11.attackFE     -10.4618029 264.1957859  -0.0396 0.9684131    
# 12.attackFE      -0.0121458   0.5502168  -0.0221 0.9823884    
# 13.attackFE       0.9853197   1.0715358   0.9195 0.3578134    
# 14.attackFE       0.7284843   0.5224387   1.3944 0.1631993    
# 1.receiveFE      -0.7315755   0.2770237  -2.6408 0.0082700 ** 
# 2.receiveFE      -0.2483185   0.2681292  -0.9261 0.3543862    
# 3.receiveFE      -0.3750730   0.2725416  -1.3762 0.1687582    
# 4.receiveFE       1.0326554   0.2610273   3.9561 7.618e-05 ***
# 5.receiveFE      -0.8724350   0.2925982  -2.9817 0.0028667 ** 
# 6.receiveFE       0.7239207   0.3403191   2.1272 0.0334050 *  
# 7.receiveFE       0.2399268   0.2626681   0.9134 0.3610207    
# 8.receiveFE       0.0322648   0.3395839   0.0950 0.9243046    
# 9.receiveFE       0.6527401   0.3048317   2.1413 0.0322488 *  
# 10.receiveFE     -0.0083194   0.2918528  -0.0285 0.9772590    
# 11.receiveFE      1.6776185   0.4538049   3.6968 0.0002183 ***
# 12.receiveFE     -0.4512307   0.3601768  -1.2528 0.2102774    
# 13.receiveFE     -0.3517487   1.0308039  -0.3412 0.7329249    
# cop.startFE      -8.8434147   0.1178511 -75.0389 < 2.2e-16 ***
# cop.endFE        -1.4478174   0.1414214 -10.2376 < 2.2e-16 ***
# initFO.FE         6.1008012   0.0385758 158.1508 < 2.2e-16 ***
# 1.initCH          2.6172926   1.0000010   2.6173 0.0088631 ** 
# 2.initCH         -9.1855050 280.2587561  -0.0328 0.9738539    
# 3.initCH         -8.9884961 283.0089737  -0.0318 0.9746631    
# 4.initCH          1.5141258   1.0000003   1.5141 0.1299940    
# 5.initCH          3.0791100   1.0000013   3.0791 0.0020762 ** 
# 6.initCH          6.7359045   0.2500001  26.9436 < 2.2e-16 ***
# 7.initCH         -9.5410217 275.4923345  -0.0346 0.9723727    
# 8.initCH          6.5629143   0.2425357  27.0596 < 2.2e-16 ***
# 9.initCH          6.8779021   0.1740777  39.5105 < 2.2e-16 ***
# 10.initCH         4.4654068   0.5000003   8.9308 < 2.2e-16 ***
# 11.initCH         6.7536032   0.4082485  16.5429 < 2.2e-16 ***
# 12.initCH        -7.3922442 308.7518796  -0.0239 0.9808986    
# 13.initCH        -5.0627152 363.2563881  -0.0139 0.9888802    
# 14.initCH         6.2717656   0.3333335  18.8153 < 2.2e-16 ***
# 15.initCH         5.4036745   0.7071078   7.6419 2.132e-14 ***
# 16.initCH         6.9077542   0.3535536  19.5381 < 2.2e-16 ***
# 17.initCH         6.2146003   1.0000033   6.2146 5.146e-10 ***
# 18.initCH         6.9077466   1.0000037   6.9077 4.925e-12 ***
# 19.initCH         5.8091412   0.5000004  11.6183 < 2.2e-16 ***
# 20.initCH         6.5976000   0.2132008  30.9455 < 2.2e-16 ***
# 21.initCH         6.9077466   1.0000037   6.9077 4.925e-12 ***
# 22.initCH         0.0000000 894.4271910   0.0000 1.0000000    
# 23.initCH        -5.0627152 363.2563881  -0.0139 0.9888802    
# 24.initCH         6.8505964   0.2425357  28.2457 < 2.2e-16 ***
# 25.initCH        -5.6466962 346.9319758  -0.0163 0.9870141    
# 26.initCH        -5.6466962 346.9319758  -0.0163 0.9870141    
# 27.initCH         6.0604566   0.3333335  18.1814 < 2.2e-16 ***
# 28.initCH        -5.9926922 338.2402555  -0.0177 0.9858644    
# 29.initCH        -5.0627152 363.2563881  -0.0139 0.9888802    
# 30.initCH         6.8243737   0.0455016 149.9810 < 2.2e-16 ***
# 1.initSF          4.9198803   0.3162278  15.5580 < 2.2e-16 ***
# 2.initSF          5.2337785   0.2182179  23.9842 < 2.2e-16 ***
# 3.initSF          5.0470025   0.2672613  18.8842 < 2.2e-16 ***
# 4.initSF          5.2753277   0.1524986  34.5926 < 2.2e-16 ***
# 5.initSF          4.4654068   0.5000003   8.9308 < 2.2e-16 ***
# 6.initSF         -7.6008103 304.9854879  -0.0249 0.9801173    
# 7.initSF          5.1630631   0.1856954  27.8039 < 2.2e-16 ***
# 8.initSF          4.8283117   0.5773507   8.3629 < 2.2e-16 ***
# 9.initSF         -8.1170250 296.2257322  -0.0274 0.9781395    
# 10.initSF         4.8708723   0.4082485  11.9311 < 2.2e-16 ***
# 11.initSF         4.9618389   1.0000025   4.9618 6.983e-07 ***
# 12.initSF         6.5022894   0.3162279  20.5620 < 2.2e-16 ***
# 13.initSF         6.9077466   1.0000037   6.9077 4.925e-12 ***
# 14.initSF         5.4608346   0.5000003  10.9217 < 2.2e-16 ***
# 15.initSF         5.8091406   0.5773508  10.0617 < 2.2e-16 ***
# 16.initSF        -6.8411143 319.4179285  -0.0214 0.9829127    
# 17.initSF         6.2146003   1.0000033   6.2146 5.146e-10 ***
# 18.initSF        -5.0627152 363.2563881  -0.0139 0.9888802    
# 19.initSF         6.3687576   0.3779647  16.8501 < 2.2e-16 ***
# 20.initSF         5.4524671   0.3779646  14.4259 < 2.2e-16 ***
# 21.initSF        -5.0627152 363.2563881  -0.0139 0.9888802    
# 22.initSF         0.0000000 894.4271910   0.0000 1.0000000    
# 23.initSF         6.9077466   1.0000037   6.9077 4.925e-12 ***
# 24.initSF         4.0173785   1.0000019   4.0174 5.885e-05 ***
# 25.initSF         6.2146003   1.0000033   6.2146 5.146e-10 ***
# 26.initSF        -5.6466962 346.9319758  -0.0163 0.9870141    
# 27.initSF         4.9618431   0.5773507   8.5942 < 2.2e-16 ***
# 28.initSF         5.8091357   1.0000030   5.8091 6.280e-09 ***
# 29.initSF        -5.0627152 363.2563881  -0.0139 0.9888802    
# 30.initSF         3.9401937   0.1924501  20.4738 < 2.2e-16 ***
# 1.disengage      -4.2622836   0.0626224 -68.0632 < 2.2e-16 ***
# 2.disengage      -5.1487697   0.0887356 -58.0237 < 2.2e-16 ***
# 3.disengage      -4.7750140   0.0636285 -75.0452 < 2.2e-16 ***
# 4.disengage      -4.6474529   0.0548821 -84.6806 < 2.2e-16 ***
# 5.disengage      -4.5737155   0.1025978 -44.5791 < 2.2e-16 ***
# 6.disengage      -6.4777863   0.5773494 -11.2199 < 2.2e-16 ***
# 7.disengage      -5.6831776   0.0894427 -63.5399 < 2.2e-16 ***
# 8.disengage      -3.0256324   0.1889822 -16.0101 < 2.2e-16 ***
# 9.disengage      -2.9513586   0.9999975  -2.9514 0.0031637 ** 
# 10.disengage     -4.6109312   0.0880451 -52.3701 < 2.2e-16 ***
# 11.disengage     -1.2519039   0.9999986  -1.2519 0.2106042    
# 12.disengage     -5.0999545   0.3333332 -15.2999 < 2.2e-16 ***
# 13.disengage     -3.0694899   0.9999975  -3.0695 0.0021442 ** 
# 14.disengage     -3.9995581   0.2000000 -19.9978 < 2.2e-16 ***
# 15.disengage     -3.7115624   0.3779643  -9.8199 < 2.2e-16 ***
# 16.disengage      0.0000000 894.4271910   0.0000 1.0000000    
# 17.disengage     -5.5854796   0.7071053  -7.8991 2.887e-15 ***
# 18.disengage      0.0000000 894.4271910   0.0000 1.0000000    
# 19.disengage     -3.3061330   0.2500000 -13.2245 < 2.2e-16 ***
# 20.disengage     -2.4505089   0.3535533  -6.9311 4.176e-12 ***
# 21.disengage      0.0000000 894.4271910   0.0000 1.0000000    
# 22.disengage     -3.5143729   0.9999972  -3.5144 0.0004408 ***
# 23.disengage    -12.1897538 246.2838524  -0.0495 0.9605250    
# 24.disengage     -1.8919041   0.9999982  -1.8919 0.0585033 .  
# 25.disengage     -2.4340658   0.2000000 -12.1703 < 2.2e-16 ***
# 26.disengage     -3.8049157   0.4082481  -9.3201 < 2.2e-16 ***
# 27.disengage     -5.3682937   0.3162276 -16.9760 < 2.2e-16 ***
# 28.disengage     -5.0921384   0.3535532 -14.4027 < 2.2e-16 ***
# 29.disengage     -5.0977870   0.4082480 -12.4870 < 2.2e-16 ***
# 30.disengage     -4.8310314   0.1622214 -29.7805 < 2.2e-16 ***
# 1.depart         -8.7014746   0.2357022 -36.9172 < 2.2e-16 ***
# 2.depart         -8.6455924   0.2357022 -36.6802 < 2.2e-16 ***
# 3.depart         -8.6031331   0.2357022 -36.5000 < 2.2e-16 ***
# 4.depart         -8.2131081   0.2357022 -34.8453 < 2.2e-16 ***
# 5.depart         -8.7978526   0.2886750 -30.4767 < 2.2e-16 ***
# 6.depart         -8.1073118   0.7071048 -11.4655 < 2.2e-16 ***
# 7.depart         -8.4665832   0.2357022 -35.9207 < 2.2e-16 ***
# 8.depart         -7.6244610   0.4999993 -15.2489 < 2.2e-16 ***
# 9.depart         -8.2131177   0.5773492 -14.2256 < 2.2e-16 ***
# 10.depart        -8.3366519   0.3779642 -22.0567 < 2.2e-16 ***
# 11.depart        -6.1361062   0.5773494 -10.6281 < 2.2e-16 ***
# 12.depart        -8.4305661   0.5773491 -14.6022 < 2.2e-16 ***
# 13.depart        -6.3742619   0.9999954  -6.3743 1.838e-10 ***
# 14.depart        -8.4328617   0.7071047 -11.9259 < 2.2e-16 ***
# 15.depart        -8.4724875   0.9999941  -8.4725 < 2.2e-16 ***
# 16.depart        -4.5531868   0.9999965  -4.5532 5.284e-06 ***
# 17.depart        -7.3325942   0.9999948  -7.3326 2.256e-13 ***
# 18.depart        -6.1293192   0.9999955  -6.1293 8.824e-10 ***
# 19.depart        -8.8972145   0.9999938  -8.8973 < 2.2e-16 ***
# 20.depart        -7.6761506   0.9999946  -7.6762 1.643e-14 ***
# 21.depart        -4.7116752   0.9999964  -4.7117 2.457e-06 ***
# 22.depart        -7.7353839   0.9999945  -7.7354 1.021e-14 ***
# 23.depart        -7.0172213   0.9999950  -7.0173 2.263e-12 ***
# 24.depart        -7.3915096   0.9999948  -7.3915 1.452e-13 ***
# 25.depart        -7.8421735   0.9999945  -7.8422 4.441e-15 ***
# 26.depart        -8.0356182   0.9999944  -8.0357 8.882e-16 ***
# 27.depart        -8.6596852   0.9999940  -8.6597 < 2.2e-16 ***
# 28.depart        -8.1918068   0.9999943  -8.1919 2.220e-16 ***
# 29.depart        -7.0548164   0.9999950  -7.0549 1.728e-12 ***
# 30.depart        -5.7379132   0.1178511 -48.6878 < 2.2e-16 ***
# ---
# Signif. codes:  0 ‘***’ 0.001 ‘**’ 0.01 ‘*’ 0.05 ‘.’ 0.1 ‘ ’ 1
# Null deviance: 76628.21 on 4854 degrees of freedom
# Residual deviance: 30565.98 on 4698 degrees of freedom
# 	Chi-square: 46062.24 on 156 degrees of freedom, asymptotic p-value 0 
# AIC: 30877.98 AICC: 30888.41 BIC: 31890.04 
# Log posterior: -171436 
# Prior parameters: mu=0 sigma=1000 nu=4 

#significant FE but much lower model performance. keep digging 


### Partition initFO rate for ID males

sequences<-table(event.list$Date)

#create list of dummy intercept names

idno<-length(event.types[grep("cop.start",event.types)])

dummy.ints.attack<-paste(1:14,".attackFE", sep="")
dummy.ints.receive<-paste(1:13, ".receiveFE", sep="")


int.names<-c("Global.attackFE", "X.receiveFE", "X.UreceiveFE", "U.attackFE", "U.receiveFE", "U.UattackFE", dummy.ints.attack, dummy.ints.receive, "cop.startFE", "cop.endFE", "initFO.FE","ID.initFO.FE", event.types[-c(grep("attack", event.types),grep("cop", event.types),grep("initFO", event.types))])

int.list<-list()

for(i in 1:length(levels(event.list$Date))){

day.array<-array(0,dim=c(as.numeric(sequences[i]), length(event.types),length(int.names)))

#the third dimension is the stat slice.
dimnames(day.array)<-list(c(1:sequences[i]),as.character(event.types),as.character(int.names))

#add global attack effect, loops for "U" intercepts for attacks and receives, then one loop for typical intercepts

day.array[,grep("attack", event.types),1]<-1


for(h in 1:14){
day.array[,paste(h,".attack.",30, sep=""),2]<-1
}


for(h in 15:29){
day.array[,paste(h,".attack.",30, sep=""),3]<-1
}


for(h in 1:14){
	for(j in 15:29){	
day.array[,paste(j,".attack.", h, sep=""),4]<-1
}
}


for(h in 1:14){
	for(j in 15:29){
day.array[,paste(h,".attack.",j, sep=""),5]<-1
}
}

for(h in 15:29){
	for(j in 15:29){	
if(h!=j){
day.array[,paste(j,".attack.", h, sep=""),6]<-1
}
}
}

preloop<-6

#loop for dummy intercepts for attacks, loop for dummy intercepts for receives, add in cop.start and cop.end FEs, then one loop for typical intercepts

for(k in (preloop+1):(length(dummy.ints.attack)+preloop)){
	
	for(h in 1:14){
if(h!=(k-preloop)){
day.array[,paste((k-preloop),".attack.", h, sep=""),k]<-1
}
}
}

for(k in ((length(dummy.ints.attack)+preloop)+1):((length(dummy.ints.attack)+preloop)+length(dummy.ints.receive))){
	for(h in 1:14){
if(h!=(k-(length(dummy.ints.attack)+preloop))){
day.array[,paste(h,".attack.",(k-(length(dummy.ints.attack)+preloop)), sep=""),k]<-1
}
}
}

#cop.startFE
for(h in 1:idno){
day.array[,paste(h,".cop.start", sep=""),((length(dummy.ints.attack)+preloop)+length(dummy.ints.receive)+1)]<-1
}


#cop.endFE
for(h in 1:idno){
day.array[,paste(h,".cop.end", sep=""),((length(dummy.ints.attack)+preloop)+length(dummy.ints.receive)+2)]<-1
}

#initFO.FE
for(h in 1:idno){
day.array[,paste(h,".initFO", sep=""),((length(dummy.ints.attack)+preloop)+length(dummy.ints.receive)+3)]<-1
}

#ID.initFO.FE
for(h in 1:14){
day.array[,paste(h,".initFO", sep=""),((length(dummy.ints.attack)+preloop)+length(dummy.ints.receive)+4)]<-1
}

for(k in (((length(dummy.ints.attack)+preloop)+length(dummy.ints.receive)+5):length(int.names))){
	
	day.array[,int.names[k],k]<-1
	
}

#weird extra formatting to make it match the package example, seems like it becomes relevant *punintended!* if I want to fit day-specific effects

global<-list()

global[[1]]<-day.array

int.list[[i]]<-global

names(int.list[[i]])<-"global"

}

tic()
intfit.sup<-rem(evlist, int.list, supplist=supplist, timing="interval", estimator="BPM")
toc()

summary(intfit.sup)

# Egocentric Relational Event Model (Interval Likelihood)
# 
#                   Post.Mode     Post.SD  Z value  Pr(>|z|)    
# Global.attackFE  -9.5259220   0.2886747 -32.9988 < 2.2e-16 ***
# X.receiveFE       3.7712555   0.2921739  12.9076 < 2.2e-16 ***
# X.UreceiveFE      3.9025235   0.3370996  11.5768 < 2.2e-16 ***
# U.attackFE       -1.2734082   0.6454967  -1.9728 0.0485233 *  
# U.receiveFE       1.7877281   0.3145761   5.6830 1.324e-08 ***
# U.UattackFE       2.9176286   0.3248928   8.9803 < 2.2e-16 ***
# 1.attackFE        1.0330374   0.3944376   2.6190 0.0088184 ** 
# 2.attackFE        1.2763648   0.3921908   3.2544 0.0011361 ** 
# 3.attackFE        1.6061436   0.3891684   4.1271 3.673e-05 ***
# 4.attackFE        2.5265882   0.3860257   6.5451 5.944e-11 ***
# 5.attackFE        0.3110660   0.4195142   0.7415 0.4583958    
# 6.attackFE       -0.0207165   0.8047343  -0.0257 0.9794621    
# 7.attackFE        1.3366638   0.3932445   3.3991 0.0006762 ***
# 8.attackFE       -1.4929214   1.0698668  -1.3954 0.1628870    
# 9.attackFE      -12.5411760 243.0502149  -0.0516 0.9588481    
# 10.attackFE       1.5764042   0.4001704   3.9393 8.171e-05 ***
# 11.attackFE     -10.4618029 264.1957859  -0.0396 0.9684131    
# 12.attackFE      -0.0121458   0.5502168  -0.0221 0.9823884    
# 13.attackFE       0.9853197   1.0715358   0.9195 0.3578134    
# 14.attackFE       0.7284843   0.5224387   1.3944 0.1631993    
# 1.receiveFE      -0.7315755   0.2770237  -2.6408 0.0082700 ** 
# 2.receiveFE      -0.2483185   0.2681292  -0.9261 0.3543862    
# 3.receiveFE      -0.3750730   0.2725416  -1.3762 0.1687582    
# 4.receiveFE       1.0326554   0.2610273   3.9561 7.618e-05 ***
# 5.receiveFE      -0.8724350   0.2925982  -2.9817 0.0028667 ** 
# 6.receiveFE       0.7239207   0.3403191   2.1272 0.0334050 *  
# 7.receiveFE       0.2399268   0.2626681   0.9134 0.3610207    
# 8.receiveFE       0.0322648   0.3395839   0.0950 0.9243046    
# 9.receiveFE       0.6527401   0.3048317   2.1413 0.0322488 *  
# 10.receiveFE     -0.0083194   0.2918528  -0.0285 0.9772590    
# 11.receiveFE      1.6776185   0.4538049   3.6968 0.0002183 ***
# 12.receiveFE     -0.4512307   0.3601768  -1.2528 0.2102774    
# 13.receiveFE     -0.3517487   1.0308039  -0.3412 0.7329249    
# cop.startFE      -8.8434147   0.1178511 -75.0389 < 2.2e-16 ***
# cop.endFE        -1.4478174   0.1414214 -10.2376 < 2.2e-16 ***
# initFO.FE         4.0360756   0.1666667  24.2165 < 2.2e-16 ***
# ID.initFO.FE      2.5583850   0.1713187  14.9335 < 2.2e-16 ***
# 1.initCH          2.6172926   1.0000010   2.6173 0.0088631 ** 
# 2.initCH         -9.1855050 280.2587561  -0.0328 0.9738539    
# 3.initCH         -8.9884961 283.0089737  -0.0318 0.9746631    
# 4.initCH          1.5141258   1.0000003   1.5141 0.1299940    
# 5.initCH          3.0791100   1.0000013   3.0791 0.0020762 ** 
# 6.initCH          6.7359045   0.2500001  26.9436 < 2.2e-16 ***
# 7.initCH         -9.5410217 275.4923345  -0.0346 0.9723727    
# 8.initCH          6.5629143   0.2425357  27.0596 < 2.2e-16 ***
# 9.initCH          6.8779021   0.1740777  39.5105 < 2.2e-16 ***
# 10.initCH         4.4654068   0.5000003   8.9308 < 2.2e-16 ***
# 11.initCH         6.7536032   0.4082485  16.5429 < 2.2e-16 ***
# 12.initCH        -7.3922442 308.7518796  -0.0239 0.9808986    
# 13.initCH        -5.0627152 363.2563881  -0.0139 0.9888802    
# 14.initCH         6.2717656   0.3333335  18.8153 < 2.2e-16 ***
# 15.initCH         5.4036745   0.7071078   7.6419 2.132e-14 ***
# 16.initCH         6.9077542   0.3535536  19.5381 < 2.2e-16 ***
# 17.initCH         6.2146003   1.0000033   6.2146 5.146e-10 ***
# 18.initCH         6.9077466   1.0000037   6.9077 4.925e-12 ***
# 19.initCH         5.8091412   0.5000004  11.6183 < 2.2e-16 ***
# 20.initCH         6.5976000   0.2132008  30.9455 < 2.2e-16 ***
# 21.initCH         6.9077466   1.0000037   6.9077 4.925e-12 ***
# 22.initCH         0.0000000 894.4271910   0.0000 1.0000000    
# 23.initCH        -5.0627152 363.2563881  -0.0139 0.9888802    
# 24.initCH         6.8505964   0.2425357  28.2457 < 2.2e-16 ***
# 25.initCH        -5.6466962 346.9319758  -0.0163 0.9870141    
# 26.initCH        -5.6466962 346.9319758  -0.0163 0.9870141    
# 27.initCH         6.0604566   0.3333335  18.1814 < 2.2e-16 ***
# 28.initCH        -5.9926922 338.2402555  -0.0177 0.9858644    
# 29.initCH        -5.0627152 363.2563881  -0.0139 0.9888802    
# 30.initCH         6.8243737   0.0455016 149.9810 < 2.2e-16 ***
# 1.initSF          4.9198803   0.3162278  15.5580 < 2.2e-16 ***
# 2.initSF          5.2337785   0.2182179  23.9842 < 2.2e-16 ***
# 3.initSF          5.0470025   0.2672613  18.8842 < 2.2e-16 ***
# 4.initSF          5.2753277   0.1524986  34.5926 < 2.2e-16 ***
# 5.initSF          4.4654068   0.5000003   8.9308 < 2.2e-16 ***
# 6.initSF         -7.6008103 304.9854879  -0.0249 0.9801173    
# 7.initSF          5.1630631   0.1856954  27.8039 < 2.2e-16 ***
# 8.initSF          4.8283117   0.5773507   8.3629 < 2.2e-16 ***
# 9.initSF         -8.1170250 296.2257322  -0.0274 0.9781395    
# 10.initSF         4.8708723   0.4082485  11.9311 < 2.2e-16 ***
# 11.initSF         4.9618389   1.0000025   4.9618 6.983e-07 ***
# 12.initSF         6.5022894   0.3162279  20.5620 < 2.2e-16 ***
# 13.initSF         6.9077466   1.0000037   6.9077 4.925e-12 ***
# 14.initSF         5.4608346   0.5000003  10.9217 < 2.2e-16 ***
# 15.initSF         5.8091406   0.5773508  10.0617 < 2.2e-16 ***
# 16.initSF        -6.8411143 319.4179285  -0.0214 0.9829127    
# 17.initSF         6.2146003   1.0000033   6.2146 5.146e-10 ***
# 18.initSF        -5.0627152 363.2563881  -0.0139 0.9888802    
# 19.initSF         6.3687576   0.3779647  16.8501 < 2.2e-16 ***
# 20.initSF         5.4524671   0.3779646  14.4259 < 2.2e-16 ***
# 21.initSF        -5.0627152 363.2563881  -0.0139 0.9888802    
# 22.initSF         0.0000000 894.4271910   0.0000 1.0000000    
# 23.initSF         6.9077466   1.0000037   6.9077 4.925e-12 ***
# 24.initSF         4.0173785   1.0000019   4.0174 5.885e-05 ***
# 25.initSF         6.2146003   1.0000033   6.2146 5.146e-10 ***
# 26.initSF        -5.6466962 346.9319758  -0.0163 0.9870141    
# 27.initSF         4.9618431   0.5773507   8.5942 < 2.2e-16 ***
# 28.initSF         5.8091357   1.0000030   5.8091 6.280e-09 ***
# 29.initSF        -5.0627152 363.2563881  -0.0139 0.9888802    
# 30.initSF         3.9401937   0.1924501  20.4738 < 2.2e-16 ***
# 1.disengage      -4.2622836   0.0626224 -68.0632 < 2.2e-16 ***
# 2.disengage      -5.1487697   0.0887356 -58.0237 < 2.2e-16 ***
# 3.disengage      -4.7750140   0.0636285 -75.0452 < 2.2e-16 ***
# 4.disengage      -4.6474529   0.0548821 -84.6806 < 2.2e-16 ***
# 5.disengage      -4.5737155   0.1025978 -44.5791 < 2.2e-16 ***
# 6.disengage      -6.4777863   0.5773494 -11.2199 < 2.2e-16 ***
# 7.disengage      -5.6831776   0.0894427 -63.5399 < 2.2e-16 ***
# 8.disengage      -3.0256324   0.1889822 -16.0101 < 2.2e-16 ***
# 9.disengage      -2.9513586   0.9999975  -2.9514 0.0031637 ** 
# 10.disengage     -4.6109312   0.0880451 -52.3701 < 2.2e-16 ***
# 11.disengage     -1.2519039   0.9999986  -1.2519 0.2106042    
# 12.disengage     -5.0999545   0.3333332 -15.2999 < 2.2e-16 ***
# 13.disengage     -3.0694899   0.9999975  -3.0695 0.0021442 ** 
# 14.disengage     -3.9995581   0.2000000 -19.9978 < 2.2e-16 ***
# 15.disengage     -3.7115624   0.3779643  -9.8199 < 2.2e-16 ***
# 16.disengage      0.0000000 894.4271910   0.0000 1.0000000    
# 17.disengage     -5.5854796   0.7071053  -7.8991 2.887e-15 ***
# 18.disengage      0.0000000 894.4271910   0.0000 1.0000000    
# 19.disengage     -3.3061330   0.2500000 -13.2245 < 2.2e-16 ***
# 20.disengage     -2.4505089   0.3535533  -6.9311 4.176e-12 ***
# 21.disengage      0.0000000 894.4271910   0.0000 1.0000000    
# 22.disengage     -3.5143729   0.9999972  -3.5144 0.0004408 ***
# 23.disengage    -12.1897538 246.2838524  -0.0495 0.9605250    
# 24.disengage     -1.8919041   0.9999982  -1.8919 0.0585033 .  
# 25.disengage     -2.4340658   0.2000000 -12.1703 < 2.2e-16 ***
# 26.disengage     -3.8049157   0.4082481  -9.3201 < 2.2e-16 ***
# 27.disengage     -5.3682937   0.3162276 -16.9760 < 2.2e-16 ***
# 28.disengage     -5.0921384   0.3535532 -14.4027 < 2.2e-16 ***
# 29.disengage     -5.0977870   0.4082480 -12.4870 < 2.2e-16 ***
# 30.disengage     -4.8310314   0.1622214 -29.7805 < 2.2e-16 ***
# 1.depart         -8.7014746   0.2357022 -36.9172 < 2.2e-16 ***
# 2.depart         -8.6455924   0.2357022 -36.6802 < 2.2e-16 ***
# 3.depart         -8.6031331   0.2357022 -36.5000 < 2.2e-16 ***
# 4.depart         -8.2131081   0.2357022 -34.8453 < 2.2e-16 ***
# 5.depart         -8.7978526   0.2886750 -30.4767 < 2.2e-16 ***
# 6.depart         -8.1073118   0.7071048 -11.4655 < 2.2e-16 ***
# 7.depart         -8.4665832   0.2357022 -35.9207 < 2.2e-16 ***
# 8.depart         -7.6244610   0.4999993 -15.2489 < 2.2e-16 ***
# 9.depart         -8.2131177   0.5773492 -14.2256 < 2.2e-16 ***
# 10.depart        -8.3366519   0.3779642 -22.0567 < 2.2e-16 ***
# 11.depart        -6.1361062   0.5773494 -10.6281 < 2.2e-16 ***
# 12.depart        -8.4305661   0.5773491 -14.6022 < 2.2e-16 ***
# 13.depart        -6.3742619   0.9999954  -6.3743 1.838e-10 ***
# 14.depart        -8.4328617   0.7071047 -11.9259 < 2.2e-16 ***
# 15.depart        -8.4724875   0.9999941  -8.4725 < 2.2e-16 ***
# 16.depart        -4.5531868   0.9999965  -4.5532 5.284e-06 ***
# 17.depart        -7.3325942   0.9999948  -7.3326 2.256e-13 ***
# 18.depart        -6.1293192   0.9999955  -6.1293 8.824e-10 ***
# 19.depart        -8.8972145   0.9999938  -8.8973 < 2.2e-16 ***
# 20.depart        -7.6761506   0.9999946  -7.6762 1.643e-14 ***
# 21.depart        -4.7116752   0.9999964  -4.7117 2.457e-06 ***
# 22.depart        -7.7353839   0.9999945  -7.7354 1.021e-14 ***
# 23.depart        -7.0172213   0.9999950  -7.0173 2.263e-12 ***
# 24.depart        -7.3915096   0.9999948  -7.3915 1.452e-13 ***
# 25.depart        -7.8421735   0.9999945  -7.8422 4.441e-15 ***
# 26.depart        -8.0356182   0.9999944  -8.0357 8.882e-16 ***
# 27.depart        -8.6596852   0.9999940  -8.6597 < 2.2e-16 ***
# 28.depart        -8.1918068   0.9999943  -8.1919 2.220e-16 ***
# 29.depart        -7.0548164   0.9999950  -7.0549 1.728e-12 ***
# 30.depart        -5.7379132   0.1178511 -48.6878 < 2.2e-16 ***
# ---
# Signif. codes:  0 ‘***’ 0.001 ‘**’ 0.01 ‘*’ 0.05 ‘.’ 0.1 ‘ ’ 1
# Null deviance: 76628.21 on 4854 degrees of freedom
# Residual deviance: 30086.7 on 4697 degrees of freedom
# 	Chi-square: 46541.51 on 157 degrees of freedom, asymptotic p-value 0 
# AIC: 30400.7 AICC: 30411.27 BIC: 31419.25 
# Log posterior: -172197.3 
# Prior parameters: mu=0 sigma=1000 nu=4 

#Great! what about pulling out the U's?


## initFO: parition out IDs AND U's


sequences<-table(event.list$Date)

#create list of dummy intercept names

idno<-length(event.types[grep("cop.start",event.types)])

dummy.ints.attack<-paste(1:14,".attackFE", sep="")
dummy.ints.receive<-paste(1:13, ".receiveFE", sep="")


int.names<-c("Global.attackFE", "X.receiveFE", "X.UreceiveFE", "U.attackFE", "U.receiveFE", "U.UattackFE", dummy.ints.attack, dummy.ints.receive, "cop.startFE", "cop.endFE", "initFO.FE","ID.initFO.FE", "U.initFO.FE",event.types[-c(grep("attack", event.types),grep("cop", event.types),grep("initFO", event.types))])

int.list<-list()

for(i in 1:length(levels(event.list$Date))){

day.array<-array(0,dim=c(as.numeric(sequences[i]), length(event.types),length(int.names)))

#the third dimension is the stat slice.
dimnames(day.array)<-list(c(1:sequences[i]),as.character(event.types),as.character(int.names))

#add global attack effect, loops for "U" intercepts for attacks and receives, then one loop for typical intercepts

day.array[,grep("attack", event.types),1]<-1


for(h in 1:14){
day.array[,paste(h,".attack.",30, sep=""),2]<-1
}


for(h in 15:29){
day.array[,paste(h,".attack.",30, sep=""),3]<-1
}


for(h in 1:14){
	for(j in 15:29){	
day.array[,paste(j,".attack.", h, sep=""),4]<-1
}
}


for(h in 1:14){
	for(j in 15:29){
day.array[,paste(h,".attack.",j, sep=""),5]<-1
}
}

for(h in 15:29){
	for(j in 15:29){	
if(h!=j){
day.array[,paste(j,".attack.", h, sep=""),6]<-1
}
}
}

preloop<-6

#loop for dummy intercepts for attacks, loop for dummy intercepts for receives, add in cop.start and cop.end FEs, then one loop for typical intercepts

for(k in (preloop+1):(length(dummy.ints.attack)+preloop)){
	
	for(h in 1:14){
if(h!=(k-preloop)){
day.array[,paste((k-preloop),".attack.", h, sep=""),k]<-1
}
}
}

for(k in ((length(dummy.ints.attack)+preloop)+1):((length(dummy.ints.attack)+preloop)+length(dummy.ints.receive))){
	for(h in 1:14){
if(h!=(k-(length(dummy.ints.attack)+preloop))){
day.array[,paste(h,".attack.",(k-(length(dummy.ints.attack)+preloop)), sep=""),k]<-1
}
}
}

#cop.startFE
for(h in 1:idno){
day.array[,paste(h,".cop.start", sep=""),((length(dummy.ints.attack)+preloop)+length(dummy.ints.receive)+1)]<-1
}

#cop.endFE
for(h in 1:idno){
day.array[,paste(h,".cop.end", sep=""),((length(dummy.ints.attack)+preloop)+length(dummy.ints.receive)+2)]<-1
}

#initFO.FE
for(h in 1:idno){
day.array[,paste(h,".initFO", sep=""),((length(dummy.ints.attack)+preloop)+length(dummy.ints.receive)+3)]<-1
}

#ID.initFO.FE
for(h in 1:14){
day.array[,paste(h,".initFO", sep=""),((length(dummy.ints.attack)+preloop)+length(dummy.ints.receive)+4)]<-1
}

#U.initFO.FE
for(h in 15:29){
day.array[,paste(h,".initFO", sep=""),((length(dummy.ints.attack)+preloop)+length(dummy.ints.receive)+5)]<-1
}


for(k in (((length(dummy.ints.attack)+preloop)+length(dummy.ints.receive)+6):length(int.names))){
	
	day.array[,int.names[k],k]<-1
	
}

#weird extra formatting to make it match the package example, seems like it becomes relevant *punintended!* if I want to fit day-specific effects

global<-list()

global[[1]]<-day.array

int.list[[i]]<-global

names(int.list[[i]])<-"global"

}

tic()
intfit.sup<-rem(evlist, int.list, supplist=supplist, timing="interval", estimator="BPM")
toc()

summary(intfit.sup)

# Egocentric Relational Event Model (Interval Likelihood)
# 
#                   Post.Mode     Post.SD  Z value  Pr(>|z|)    
# Global.attackFE  -9.5259220   0.2886747 -32.9988 < 2.2e-16 ***
# X.receiveFE       3.7712555   0.2921739  12.9076 < 2.2e-16 ***
# X.UreceiveFE      3.9025235   0.3370996  11.5768 < 2.2e-16 ***
# U.attackFE       -1.2734082   0.6454967  -1.9728 0.0485233 *  
# U.receiveFE       1.7877281   0.3145761   5.6830 1.324e-08 ***
# U.UattackFE       2.9176286   0.3248928   8.9803 < 2.2e-16 ***
# 1.attackFE        1.0330374   0.3944376   2.6190 0.0088184 ** 
# 2.attackFE        1.2763648   0.3921908   3.2544 0.0011361 ** 
# 3.attackFE        1.6061436   0.3891684   4.1271 3.673e-05 ***
# 4.attackFE        2.5265882   0.3860257   6.5451 5.944e-11 ***
# 5.attackFE        0.3110660   0.4195142   0.7415 0.4583958    
# 6.attackFE       -0.0207165   0.8047343  -0.0257 0.9794621    
# 7.attackFE        1.3366638   0.3932445   3.3991 0.0006762 ***
# 8.attackFE       -1.4929214   1.0698668  -1.3954 0.1628870    
# 9.attackFE      -12.5411756 243.0501754  -0.0516 0.9588481    
# 10.attackFE       1.5764042   0.4001704   3.9393 8.171e-05 ***
# 11.attackFE     -10.4618029 264.1957859  -0.0396 0.9684131    
# 12.attackFE      -0.0121458   0.5502168  -0.0221 0.9823884    
# 13.attackFE       0.9853197   1.0715358   0.9195 0.3578134    
# 14.attackFE       0.7284843   0.5224387   1.3944 0.1631993    
# 1.receiveFE      -0.7315755   0.2770237  -2.6408 0.0082700 ** 
# 2.receiveFE      -0.2483185   0.2681292  -0.9261 0.3543862    
# 3.receiveFE      -0.3750730   0.2725416  -1.3762 0.1687582    
# 4.receiveFE       1.0326554   0.2610273   3.9561 7.618e-05 ***
# 5.receiveFE      -0.8724350   0.2925982  -2.9817 0.0028667 ** 
# 6.receiveFE       0.7239207   0.3403191   2.1272 0.0334050 *  
# 7.receiveFE       0.2399268   0.2626681   0.9134 0.3610207    
# 8.receiveFE       0.0322648   0.3395839   0.0950 0.9243046    
# 9.receiveFE       0.6527401   0.3048317   2.1413 0.0322488 *  
# 10.receiveFE     -0.0083194   0.2918528  -0.0285 0.9772590    
# 11.receiveFE      1.6776185   0.4538049   3.6968 0.0002183 ***
# 12.receiveFE     -0.4512307   0.3601768  -1.2528 0.2102774    
# 13.receiveFE     -0.3517487   1.0308039  -0.3412 0.7329249    
# cop.startFE      -8.8434147   0.1178511 -75.0389 < 2.2e-16 ***
# cop.endFE        -1.4478174   0.1414214 -10.2376 < 2.2e-16 ***
# initFO.FE         3.3524074   0.2581988  12.9838 < 2.2e-16 ***
# ID.initFO.FE      3.2420533   0.2612259  12.4109 < 2.2e-16 ***
# U.initFO.FE       1.8903400   0.3380617   5.5917 2.249e-08 ***
# 1.initCH          2.6172926   1.0000010   2.6173 0.0088631 ** 
# 2.initCH         -9.1855050 280.2587561  -0.0328 0.9738539    
# 3.initCH         -8.9884961 283.0089737  -0.0318 0.9746631    
# 4.initCH          1.5141258   1.0000003   1.5141 0.1299940    
# 5.initCH          3.0791100   1.0000013   3.0791 0.0020762 ** 
# 6.initCH          6.7359045   0.2500001  26.9436 < 2.2e-16 ***
# 7.initCH         -9.5410217 275.4923345  -0.0346 0.9723727    
# 8.initCH          6.5629143   0.2425357  27.0596 < 2.2e-16 ***
# 9.initCH          6.8779021   0.1740777  39.5105 < 2.2e-16 ***
# 10.initCH         4.4654068   0.5000003   8.9308 < 2.2e-16 ***
# 11.initCH         6.7536032   0.4082485  16.5429 < 2.2e-16 ***
# 12.initCH        -7.3922442 308.7518796  -0.0239 0.9808986    
# 13.initCH        -5.0627152 363.2563881  -0.0139 0.9888802    
# 14.initCH         6.2717656   0.3333335  18.8153 < 2.2e-16 ***
# 15.initCH         5.4036745   0.7071078   7.6419 2.132e-14 ***
# 16.initCH         6.9077542   0.3535536  19.5381 < 2.2e-16 ***
# 17.initCH         6.2146003   1.0000033   6.2146 5.146e-10 ***
# 18.initCH         6.9077466   1.0000037   6.9077 4.925e-12 ***
# 19.initCH         5.8091412   0.5000004  11.6183 < 2.2e-16 ***
# 20.initCH         6.5976000   0.2132008  30.9455 < 2.2e-16 ***
# 21.initCH         6.9077466   1.0000037   6.9077 4.925e-12 ***
# 22.initCH         0.0000000 894.4271910   0.0000 1.0000000    
# 23.initCH        -5.0627152 363.2563881  -0.0139 0.9888802    
# 24.initCH         6.8505964   0.2425357  28.2457 < 2.2e-16 ***
# 25.initCH        -5.6466962 346.9319758  -0.0163 0.9870141    
# 26.initCH        -5.6466962 346.9319758  -0.0163 0.9870141    
# 27.initCH         6.0604566   0.3333335  18.1814 < 2.2e-16 ***
# 28.initCH        -5.9926922 338.2402555  -0.0177 0.9858644    
# 29.initCH        -5.0627152 363.2563881  -0.0139 0.9888802    
# 30.initCH         6.8243737   0.0455016 149.9810 < 2.2e-16 ***
# 1.initSF          4.9198803   0.3162278  15.5580 < 2.2e-16 ***
# 2.initSF          5.2337785   0.2182179  23.9842 < 2.2e-16 ***
# 3.initSF          5.0470025   0.2672613  18.8842 < 2.2e-16 ***
# 4.initSF          5.2753277   0.1524986  34.5926 < 2.2e-16 ***
# 5.initSF          4.4654068   0.5000003   8.9308 < 2.2e-16 ***
# 6.initSF         -7.6008103 304.9854879  -0.0249 0.9801173    
# 7.initSF          5.1630631   0.1856954  27.8039 < 2.2e-16 ***
# 8.initSF          4.8283117   0.5773507   8.3629 < 2.2e-16 ***
# 9.initSF         -8.1170250 296.2257322  -0.0274 0.9781395    
# 10.initSF         4.8708723   0.4082485  11.9311 < 2.2e-16 ***
# 11.initSF         4.9618389   1.0000025   4.9618 6.983e-07 ***
# 12.initSF         6.5022894   0.3162279  20.5620 < 2.2e-16 ***
# 13.initSF         6.9077466   1.0000037   6.9077 4.925e-12 ***
# 14.initSF         5.4608346   0.5000003  10.9217 < 2.2e-16 ***
# 15.initSF         5.8091406   0.5773508  10.0617 < 2.2e-16 ***
# 16.initSF        -6.8411143 319.4179285  -0.0214 0.9829127    
# 17.initSF         6.2146003   1.0000033   6.2146 5.146e-10 ***
# 18.initSF        -5.0627152 363.2563881  -0.0139 0.9888802    
# 19.initSF         6.3687576   0.3779647  16.8501 < 2.2e-16 ***
# 20.initSF         5.4524671   0.3779646  14.4259 < 2.2e-16 ***
# 21.initSF        -5.0627152 363.2563881  -0.0139 0.9888802    
# 22.initSF         0.0000000 894.4271910   0.0000 1.0000000    
# 23.initSF         6.9077466   1.0000037   6.9077 4.925e-12 ***
# 24.initSF         4.0173785   1.0000019   4.0174 5.885e-05 ***
# 25.initSF         6.2146003   1.0000033   6.2146 5.146e-10 ***
# 26.initSF        -5.6466962 346.9319758  -0.0163 0.9870141    
# 27.initSF         4.9618431   0.5773507   8.5942 < 2.2e-16 ***
# 28.initSF         5.8091357   1.0000030   5.8091 6.280e-09 ***
# 29.initSF        -5.0627152 363.2563881  -0.0139 0.9888802    
# 30.initSF         3.9401937   0.1924501  20.4738 < 2.2e-16 ***
# 1.disengage      -4.2622836   0.0626224 -68.0632 < 2.2e-16 ***
# 2.disengage      -5.1487697   0.0887356 -58.0237 < 2.2e-16 ***
# 3.disengage      -4.7750140   0.0636285 -75.0452 < 2.2e-16 ***
# 4.disengage      -4.6474529   0.0548821 -84.6806 < 2.2e-16 ***
# 5.disengage      -4.5737155   0.1025978 -44.5791 < 2.2e-16 ***
# 6.disengage      -6.4777863   0.5773494 -11.2199 < 2.2e-16 ***
# 7.disengage      -5.6831776   0.0894427 -63.5399 < 2.2e-16 ***
# 8.disengage      -3.0256324   0.1889822 -16.0101 < 2.2e-16 ***
# 9.disengage      -2.9513586   0.9999975  -2.9514 0.0031637 ** 
# 10.disengage     -4.6109312   0.0880451 -52.3701 < 2.2e-16 ***
# 11.disengage     -1.2519039   0.9999986  -1.2519 0.2106042    
# 12.disengage     -5.0999545   0.3333332 -15.2999 < 2.2e-16 ***
# 13.disengage     -3.0694899   0.9999975  -3.0695 0.0021442 ** 
# 14.disengage     -3.9995581   0.2000000 -19.9978 < 2.2e-16 ***
# 15.disengage     -3.7115624   0.3779643  -9.8199 < 2.2e-16 ***
# 16.disengage      0.0000000 894.4271910   0.0000 1.0000000    
# 17.disengage     -5.5854796   0.7071053  -7.8991 2.887e-15 ***
# 18.disengage      0.0000000 894.4271910   0.0000 1.0000000    
# 19.disengage     -3.3061330   0.2500000 -13.2245 < 2.2e-16 ***
# 20.disengage     -2.4505089   0.3535533  -6.9311 4.176e-12 ***
# 21.disengage      0.0000000 894.4271910   0.0000 1.0000000    
# 22.disengage     -3.5143729   0.9999972  -3.5144 0.0004408 ***
# 23.disengage    -12.1897538 246.2838524  -0.0495 0.9605250    
# 24.disengage     -1.8919041   0.9999982  -1.8919 0.0585033 .  
# 25.disengage     -2.4340658   0.2000000 -12.1703 < 2.2e-16 ***
# 26.disengage     -3.8049157   0.4082481  -9.3201 < 2.2e-16 ***
# 27.disengage     -5.3682937   0.3162276 -16.9760 < 2.2e-16 ***
# 28.disengage     -5.0921384   0.3535532 -14.4027 < 2.2e-16 ***
# 29.disengage     -5.0977870   0.4082480 -12.4870 < 2.2e-16 ***
# 30.disengage     -4.8310314   0.1622214 -29.7805 < 2.2e-16 ***
# 1.depart         -8.7014746   0.2357022 -36.9172 < 2.2e-16 ***
# 2.depart         -8.6455924   0.2357022 -36.6802 < 2.2e-16 ***
# 3.depart         -8.6031331   0.2357022 -36.5000 < 2.2e-16 ***
# 4.depart         -8.2131081   0.2357022 -34.8453 < 2.2e-16 ***
# 5.depart         -8.7978526   0.2886750 -30.4767 < 2.2e-16 ***
# 6.depart         -8.1073118   0.7071048 -11.4655 < 2.2e-16 ***
# 7.depart         -8.4665832   0.2357022 -35.9207 < 2.2e-16 ***
# 8.depart         -7.6244610   0.4999993 -15.2489 < 2.2e-16 ***
# 9.depart         -8.2131177   0.5773492 -14.2256 < 2.2e-16 ***
# 10.depart        -8.3366519   0.3779642 -22.0567 < 2.2e-16 ***
# 11.depart        -6.1361062   0.5773494 -10.6281 < 2.2e-16 ***
# 12.depart        -8.4305661   0.5773491 -14.6022 < 2.2e-16 ***
# 13.depart        -6.3742619   0.9999954  -6.3743 1.838e-10 ***
# 14.depart        -8.4328617   0.7071047 -11.9259 < 2.2e-16 ***
# 15.depart        -8.4724875   0.9999941  -8.4725 < 2.2e-16 ***
# 16.depart        -4.5531868   0.9999965  -4.5532 5.284e-06 ***
# 17.depart        -7.3325942   0.9999948  -7.3326 2.256e-13 ***
# 18.depart        -6.1293192   0.9999955  -6.1293 8.824e-10 ***
# 19.depart        -8.8972145   0.9999938  -8.8973 < 2.2e-16 ***
# 20.depart        -7.6761506   0.9999946  -7.6762 1.643e-14 ***
# 21.depart        -4.7116752   0.9999964  -4.7117 2.457e-06 ***
# 22.depart        -7.7353839   0.9999945  -7.7354 1.021e-14 ***
# 23.depart        -7.0172213   0.9999950  -7.0173 2.263e-12 ***
# 24.depart        -7.3915096   0.9999948  -7.3915 1.452e-13 ***
# 25.depart        -7.8421735   0.9999945  -7.8422 4.441e-15 ***
# 26.depart        -8.0356182   0.9999944  -8.0357 8.882e-16 ***
# 27.depart        -8.6596852   0.9999940  -8.6597 < 2.2e-16 ***
# 28.depart        -8.1918068   0.9999943  -8.1919 2.220e-16 ***
# 29.depart        -7.0548164   0.9999950  -7.0549 1.728e-12 ***
# 30.depart        -5.7379132   0.1178511 -48.6878 < 2.2e-16 ***
# ---
# Signif. codes:  0 ‘***’ 0.001 ‘**’ 0.01 ‘*’ 0.05 ‘.’ 0.1 ‘ ’ 1
# Null deviance: 76628.21 on 4854 degrees of freedom
# Residual deviance: 30056.53 on 4696 degrees of freedom
# 	Chi-square: 46571.68 on 158 degrees of freedom, asymptotic p-value 0 
# AIC: 30372.53 AICC: 30383.23 BIC: 31397.57 
# Log posterior: -173183.2 
# Prior parameters: mu=0 sigma=1000 nu=4 

#best of the FO intercept combinations so far!


#####


### for completeness, check only partitioning out X males and lumping ID and U males together


sequences<-table(event.list$Date)

#create list of dummy intercept names

idno<-length(event.types[grep("cop.start",event.types)])

dummy.ints.attack<-paste(1:14,".attackFE", sep="")
dummy.ints.receive<-paste(1:13, ".receiveFE", sep="")


int.names<-c("Global.attackFE", "X.receiveFE", "X.UreceiveFE", "U.attackFE", "U.receiveFE", "U.UattackFE", dummy.ints.attack, dummy.ints.receive, "cop.startFE", "cop.endFE", "initFO.FE","X.initFO.FE", event.types[-c(grep("attack", event.types),grep("cop", event.types),grep("initFO", event.types))])

int.list<-list()

for(i in 1:length(levels(event.list$Date))){

day.array<-array(0,dim=c(as.numeric(sequences[i]), length(event.types),length(int.names)))

#the third dimension is the stat slice.
dimnames(day.array)<-list(c(1:sequences[i]),as.character(event.types),as.character(int.names))

#add global attack effect, loops for "U" intercepts for attacks and receives, then one loop for typical intercepts

day.array[,grep("attack", event.types),1]<-1


for(h in 1:14){
day.array[,paste(h,".attack.",30, sep=""),2]<-1
}


for(h in 15:29){
day.array[,paste(h,".attack.",30, sep=""),3]<-1
}


for(h in 1:14){
	for(j in 15:29){	
day.array[,paste(j,".attack.", h, sep=""),4]<-1
}
}


for(h in 1:14){
	for(j in 15:29){
day.array[,paste(h,".attack.",j, sep=""),5]<-1
}
}

for(h in 15:29){
	for(j in 15:29){	
if(h!=j){
day.array[,paste(j,".attack.", h, sep=""),6]<-1
}
}
}

preloop<-6

#loop for dummy intercepts for attacks, loop for dummy intercepts for receives, add in cop.start and cop.end FEs, then one loop for typical intercepts

for(k in (preloop+1):(length(dummy.ints.attack)+preloop)){
	
	for(h in 1:14){
if(h!=(k-preloop)){
day.array[,paste((k-preloop),".attack.", h, sep=""),k]<-1
}
}
}

for(k in ((length(dummy.ints.attack)+preloop)+1):((length(dummy.ints.attack)+preloop)+length(dummy.ints.receive))){
	for(h in 1:14){
if(h!=(k-(length(dummy.ints.attack)+preloop))){
day.array[,paste(h,".attack.",(k-(length(dummy.ints.attack)+preloop)), sep=""),k]<-1
}
}
}

#cop.startFE
for(h in 1:idno){
day.array[,paste(h,".cop.start", sep=""),((length(dummy.ints.attack)+preloop)+length(dummy.ints.receive)+1)]<-1
}

#cop.endFE
for(h in 1:idno){
day.array[,paste(h,".cop.end", sep=""),((length(dummy.ints.attack)+preloop)+length(dummy.ints.receive)+2)]<-1
}

#initFO.FE
for(h in 1:idno){
day.array[,paste(h,".initFO", sep=""),((length(dummy.ints.attack)+preloop)+length(dummy.ints.receive)+3)]<-1
}

#X.initFO.FE
day.array[,paste(30,".initFO", sep=""),((length(dummy.ints.attack)+preloop)+length(dummy.ints.receive)+4)]<-1


for(k in (((length(dummy.ints.attack)+preloop)+length(dummy.ints.receive)+5):length(int.names))){
	
	day.array[,int.names[k],k]<-1
	
}

#weird extra formatting to make it match the package example, seems like it becomes relevant *punintended!* if I want to fit day-specific effects

global<-list()

global[[1]]<-day.array

int.list[[i]]<-global

names(int.list[[i]])<-"global"

}

tic()
intfit.sup<-rem(evlist, int.list, supplist=supplist, timing="interval", estimator="BPM")
toc()

summary(intfit.sup)


# Egocentric Relational Event Model (Interval Likelihood)
# 
#                   Post.Mode     Post.SD  Z value  Pr(>|z|)    
# Global.attackFE  -9.5259220   0.2886747 -32.9988 < 2.2e-16 ***
# X.receiveFE       3.7712555   0.2921739  12.9076 < 2.2e-16 ***
# X.UreceiveFE      3.9025235   0.3370996  11.5768 < 2.2e-16 ***
# U.attackFE       -1.2734082   0.6454967  -1.9728 0.0485233 *  
# U.receiveFE       1.7877281   0.3145761   5.6830 1.324e-08 ***
# U.UattackFE       2.9176286   0.3248928   8.9803 < 2.2e-16 ***
# 1.attackFE        1.0330374   0.3944376   2.6190 0.0088184 ** 
# 2.attackFE        1.2763648   0.3921908   3.2544 0.0011361 ** 
# 3.attackFE        1.6061436   0.3891684   4.1271 3.673e-05 ***
# 4.attackFE        2.5265882   0.3860257   6.5451 5.944e-11 ***
# 5.attackFE        0.3110660   0.4195142   0.7415 0.4583958    
# 6.attackFE       -0.0207165   0.8047343  -0.0257 0.9794621    
# 7.attackFE        1.3366638   0.3932445   3.3991 0.0006762 ***
# 8.attackFE       -1.4929214   1.0698668  -1.3954 0.1628870    
# 9.attackFE      -12.5411749 243.0500918  -0.0516 0.9588481    
# 10.attackFE       1.5764042   0.4001704   3.9393 8.171e-05 ***
# 11.attackFE     -10.4618029 264.1957859  -0.0396 0.9684131    
# 12.attackFE      -0.0121458   0.5502168  -0.0221 0.9823884    
# 13.attackFE       0.9853197   1.0715358   0.9195 0.3578134    
# 14.attackFE       0.7284843   0.5224387   1.3944 0.1631993    
# 1.receiveFE      -0.7315755   0.2770237  -2.6408 0.0082700 ** 
# 2.receiveFE      -0.2483185   0.2681292  -0.9261 0.3543862    
# 3.receiveFE      -0.3750730   0.2725416  -1.3762 0.1687582    
# 4.receiveFE       1.0326554   0.2610273   3.9561 7.618e-05 ***
# 5.receiveFE      -0.8724350   0.2925982  -2.9817 0.0028667 ** 
# 6.receiveFE       0.7239207   0.3403191   2.1272 0.0334050 *  
# 7.receiveFE       0.2399268   0.2626681   0.9134 0.3610207    
# 8.receiveFE       0.0322648   0.3395839   0.0950 0.9243046    
# 9.receiveFE       0.6527401   0.3048317   2.1413 0.0322488 *  
# 10.receiveFE     -0.0083194   0.2918528  -0.0285 0.9772590    
# 11.receiveFE      1.6776185   0.4538049   3.6968 0.0002183 ***
# 12.receiveFE     -0.4512307   0.3601768  -1.2528 0.2102774    
# 13.receiveFE     -0.3517487   1.0308039  -0.3412 0.7329249    
# cop.startFE      -8.8434147   0.1178511 -75.0389 < 2.2e-16 ***
# cop.endFE        -1.4478174   0.1414214 -10.2376 < 2.2e-16 ***
# initFO.FE         6.5068668   0.0390137 166.7841 < 2.2e-16 ***
# X.initFO.FE      -3.1544593   0.2611297 -12.0800 < 2.2e-16 ***
# 1.initCH          2.6172926   1.0000010   2.6173 0.0088631 ** 
# 2.initCH         -9.1855050 280.2587561  -0.0328 0.9738539    
# 3.initCH         -8.9884961 283.0089737  -0.0318 0.9746631    
# 4.initCH          1.5141258   1.0000003   1.5141 0.1299940    
# 5.initCH          3.0791100   1.0000013   3.0791 0.0020762 ** 
# 6.initCH          6.7359045   0.2500001  26.9436 < 2.2e-16 ***
# 7.initCH         -9.5410217 275.4923345  -0.0346 0.9723727    
# 8.initCH          6.5629143   0.2425357  27.0596 < 2.2e-16 ***
# 9.initCH          6.8779021   0.1740777  39.5105 < 2.2e-16 ***
# 10.initCH         4.4654068   0.5000003   8.9308 < 2.2e-16 ***
# 11.initCH         6.7536032   0.4082485  16.5429 < 2.2e-16 ***
# 12.initCH        -7.3922442 308.7518796  -0.0239 0.9808986    
# 13.initCH        -5.0627152 363.2563881  -0.0139 0.9888802    
# 14.initCH         6.2717656   0.3333335  18.8153 < 2.2e-16 ***
# 15.initCH         5.4036745   0.7071078   7.6419 2.132e-14 ***
# 16.initCH         6.9077542   0.3535536  19.5381 < 2.2e-16 ***
# 17.initCH         6.2146003   1.0000033   6.2146 5.146e-10 ***
# 18.initCH         6.9077466   1.0000037   6.9077 4.925e-12 ***
# 19.initCH         5.8091412   0.5000004  11.6183 < 2.2e-16 ***
# 20.initCH         6.5976000   0.2132008  30.9455 < 2.2e-16 ***
# 21.initCH         6.9077466   1.0000037   6.9077 4.925e-12 ***
# 22.initCH         0.0000000 894.4271910   0.0000 1.0000000    
# 23.initCH        -5.0627152 363.2563881  -0.0139 0.9888802    
# 24.initCH         6.8505964   0.2425357  28.2457 < 2.2e-16 ***
# 25.initCH        -5.6466962 346.9319758  -0.0163 0.9870141    
# 26.initCH        -5.6466962 346.9319758  -0.0163 0.9870141    
# 27.initCH         6.0604566   0.3333335  18.1814 < 2.2e-16 ***
# 28.initCH        -5.9926922 338.2402555  -0.0177 0.9858644    
# 29.initCH        -5.0627152 363.2563881  -0.0139 0.9888802    
# 30.initCH         6.8243737   0.0455016 149.9810 < 2.2e-16 ***
# 1.initSF          4.9198803   0.3162278  15.5580 < 2.2e-16 ***
# 2.initSF          5.2337785   0.2182179  23.9842 < 2.2e-16 ***
# 3.initSF          5.0470025   0.2672613  18.8842 < 2.2e-16 ***
# 4.initSF          5.2753277   0.1524986  34.5926 < 2.2e-16 ***
# 5.initSF          4.4654068   0.5000003   8.9308 < 2.2e-16 ***
# 6.initSF         -7.6008103 304.9854879  -0.0249 0.9801173    
# 7.initSF          5.1630631   0.1856954  27.8039 < 2.2e-16 ***
# 8.initSF          4.8283117   0.5773507   8.3629 < 2.2e-16 ***
# 9.initSF         -8.1170250 296.2257322  -0.0274 0.9781395    
# 10.initSF         4.8708723   0.4082485  11.9311 < 2.2e-16 ***
# 11.initSF         4.9618389   1.0000025   4.9618 6.983e-07 ***
# 12.initSF         6.5022894   0.3162279  20.5620 < 2.2e-16 ***
# 13.initSF         6.9077466   1.0000037   6.9077 4.925e-12 ***
# 14.initSF         5.4608346   0.5000003  10.9217 < 2.2e-16 ***
# 15.initSF         5.8091406   0.5773508  10.0617 < 2.2e-16 ***
# 16.initSF        -6.8411143 319.4179285  -0.0214 0.9829127    
# 17.initSF         6.2146003   1.0000033   6.2146 5.146e-10 ***
# 18.initSF        -5.0627152 363.2563881  -0.0139 0.9888802    
# 19.initSF         6.3687576   0.3779647  16.8501 < 2.2e-16 ***
# 20.initSF         5.4524671   0.3779646  14.4259 < 2.2e-16 ***
# 21.initSF        -5.0627152 363.2563881  -0.0139 0.9888802    
# 22.initSF         0.0000000 894.4271910   0.0000 1.0000000    
# 23.initSF         6.9077466   1.0000037   6.9077 4.925e-12 ***
# 24.initSF         4.0173785   1.0000019   4.0174 5.885e-05 ***
# 25.initSF         6.2146003   1.0000033   6.2146 5.146e-10 ***
# 26.initSF        -5.6466962 346.9319758  -0.0163 0.9870141    
# 27.initSF         4.9618431   0.5773507   8.5942 < 2.2e-16 ***
# 28.initSF         5.8091357   1.0000030   5.8091 6.280e-09 ***
# 29.initSF        -5.0627152 363.2563881  -0.0139 0.9888802    
# 30.initSF         3.9401937   0.1924501  20.4738 < 2.2e-16 ***
# 1.disengage      -4.2622836   0.0626224 -68.0632 < 2.2e-16 ***
# 2.disengage      -5.1487697   0.0887356 -58.0237 < 2.2e-16 ***
# 3.disengage      -4.7750140   0.0636285 -75.0452 < 2.2e-16 ***
# 4.disengage      -4.6474529   0.0548821 -84.6806 < 2.2e-16 ***
# 5.disengage      -4.5737155   0.1025978 -44.5791 < 2.2e-16 ***
# 6.disengage      -6.4777863   0.5773494 -11.2199 < 2.2e-16 ***
# 7.disengage      -5.6831776   0.0894427 -63.5399 < 2.2e-16 ***
# 8.disengage      -3.0256324   0.1889822 -16.0101 < 2.2e-16 ***
# 9.disengage      -2.9513586   0.9999975  -2.9514 0.0031637 ** 
# 10.disengage     -4.6109312   0.0880451 -52.3701 < 2.2e-16 ***
# 11.disengage     -1.2519039   0.9999986  -1.2519 0.2106042    
# 12.disengage     -5.0999545   0.3333332 -15.2999 < 2.2e-16 ***
# 13.disengage     -3.0694899   0.9999975  -3.0695 0.0021442 ** 
# 14.disengage     -3.9995581   0.2000000 -19.9978 < 2.2e-16 ***
# 15.disengage     -3.7115624   0.3779643  -9.8199 < 2.2e-16 ***
# 16.disengage      0.0000000 894.4271910   0.0000 1.0000000    
# 17.disengage     -5.5854796   0.7071053  -7.8991 2.887e-15 ***
# 18.disengage      0.0000000 894.4271910   0.0000 1.0000000    
# 19.disengage     -3.3061330   0.2500000 -13.2245 < 2.2e-16 ***
# 20.disengage     -2.4505089   0.3535533  -6.9311 4.176e-12 ***
# 21.disengage      0.0000000 894.4271910   0.0000 1.0000000    
# 22.disengage     -3.5143729   0.9999972  -3.5144 0.0004408 ***
# 23.disengage    -12.1897538 246.2838524  -0.0495 0.9605250    
# 24.disengage     -1.8919041   0.9999982  -1.8919 0.0585033 .  
# 25.disengage     -2.4340658   0.2000000 -12.1703 < 2.2e-16 ***
# 26.disengage     -3.8049157   0.4082481  -9.3201 < 2.2e-16 ***
# 27.disengage     -5.3682937   0.3162276 -16.9760 < 2.2e-16 ***
# 28.disengage     -5.0921384   0.3535532 -14.4027 < 2.2e-16 ***
# 29.disengage     -5.0977870   0.4082480 -12.4870 < 2.2e-16 ***
# 30.disengage     -4.8310314   0.1622214 -29.7805 < 2.2e-16 ***
# 1.depart         -8.7014746   0.2357022 -36.9172 < 2.2e-16 ***
# 2.depart         -8.6455924   0.2357022 -36.6802 < 2.2e-16 ***
# 3.depart         -8.6031331   0.2357022 -36.5000 < 2.2e-16 ***
# 4.depart         -8.2131081   0.2357022 -34.8453 < 2.2e-16 ***
# 5.depart         -8.7978526   0.2886750 -30.4767 < 2.2e-16 ***
# 6.depart         -8.1073118   0.7071048 -11.4655 < 2.2e-16 ***
# 7.depart         -8.4665832   0.2357022 -35.9207 < 2.2e-16 ***
# 8.depart         -7.6244610   0.4999993 -15.2489 < 2.2e-16 ***
# 9.depart         -8.2131177   0.5773492 -14.2256 < 2.2e-16 ***
# 10.depart        -8.3366519   0.3779642 -22.0567 < 2.2e-16 ***
# 11.depart        -6.1361062   0.5773494 -10.6281 < 2.2e-16 ***
# 12.depart        -8.4305661   0.5773491 -14.6022 < 2.2e-16 ***
# 13.depart        -6.3742619   0.9999954  -6.3743 1.838e-10 ***
# 14.depart        -8.4328617   0.7071047 -11.9259 < 2.2e-16 ***
# 15.depart        -8.4724875   0.9999941  -8.4725 < 2.2e-16 ***
# 16.depart        -4.5531868   0.9999965  -4.5532 5.284e-06 ***
# 17.depart        -7.3325942   0.9999948  -7.3326 2.256e-13 ***
# 18.depart        -6.1293192   0.9999955  -6.1293 8.824e-10 ***
# 19.depart        -8.8972145   0.9999938  -8.8973 < 2.2e-16 ***
# 20.depart        -7.6761506   0.9999946  -7.6762 1.643e-14 ***
# 21.depart        -4.7116752   0.9999964  -4.7117 2.457e-06 ***
# 22.depart        -7.7353839   0.9999945  -7.7354 1.021e-14 ***
# 23.depart        -7.0172213   0.9999950  -7.0173 2.263e-12 ***
# 24.depart        -7.3915096   0.9999948  -7.3915 1.452e-13 ***
# 25.depart        -7.8421735   0.9999945  -7.8422 4.441e-15 ***
# 26.depart        -8.0356182   0.9999944  -8.0357 8.882e-16 ***
# 27.depart        -8.6596852   0.9999940  -8.6597 < 2.2e-16 ***
# 28.depart        -8.1918068   0.9999943  -8.1919 2.220e-16 ***
# 29.depart        -7.0548164   0.9999950  -7.0549 1.728e-12 ***
# 30.depart        -5.7379132   0.1178511 -48.6878 < 2.2e-16 ***
# ---
# Signif. codes:  0 ‘***’ 0.001 ‘**’ 0.01 ‘*’ 0.05 ‘.’ 0.1 ‘ ’ 1
# Null deviance: 76628.21 on 4854 degrees of freedom
# Residual deviance: 30114.86 on 4697 degrees of freedom
# 	Chi-square: 46513.36 on 157 degrees of freedom, asymptotic p-value 0 
# AIC: 30428.86 AICC: 30439.42 BIC: 31447.4 
# Log posterior: -172211.4 
# Prior parameters: mu=0 sigma=1000 nu=4


#ok so in this case partitions for BOTH IDs and Us performs best


########### initSF ###############

sequences<-table(event.list$Date)

#create list of dummy intercept names

idno<-length(event.types[grep("cop.start",event.types)])

dummy.ints.attack<-paste(1:14,".attackFE", sep="")
dummy.ints.receive<-paste(1:13, ".receiveFE", sep="")


int.names<-c("Global.attackFE", "X.receiveFE", "X.UreceiveFE", "U.attackFE", "U.receiveFE", "U.UattackFE", dummy.ints.attack, dummy.ints.receive, "cop.startFE", "cop.endFE", "initFO.FE","ID.initFO.FE", "U.initFO.FE","initSF.FE",event.types[-c(grep("attack", event.types),grep("cop", event.types),grep("initFO", event.types),grep("initSF", event.types))])

int.list<-list()

for(i in 1:length(levels(event.list$Date))){

day.array<-array(0,dim=c(as.numeric(sequences[i]), length(event.types),length(int.names)))

#the third dimension is the stat slice.
dimnames(day.array)<-list(c(1:sequences[i]),as.character(event.types),as.character(int.names))

#add global attack effect, loops for "U" intercepts for attacks and receives, then one loop for typical intercepts

day.array[,grep("attack", event.types),1]<-1


for(h in 1:14){
day.array[,paste(h,".attack.",30, sep=""),2]<-1
}


for(h in 15:29){
day.array[,paste(h,".attack.",30, sep=""),3]<-1
}


for(h in 1:14){
	for(j in 15:29){	
day.array[,paste(j,".attack.", h, sep=""),4]<-1
}
}


for(h in 1:14){
	for(j in 15:29){
day.array[,paste(h,".attack.",j, sep=""),5]<-1
}
}

for(h in 15:29){
	for(j in 15:29){	
if(h!=j){
day.array[,paste(j,".attack.", h, sep=""),6]<-1
}
}
}

preloop<-6

#loop for dummy intercepts for attacks, loop for dummy intercepts for receives, add in cop.start and cop.end FEs, then one loop for typical intercepts

for(k in (preloop+1):(length(dummy.ints.attack)+preloop)){
	
	for(h in 1:14){
if(h!=(k-preloop)){
day.array[,paste((k-preloop),".attack.", h, sep=""),k]<-1
}
}
}

for(k in ((length(dummy.ints.attack)+preloop)+1):((length(dummy.ints.attack)+preloop)+length(dummy.ints.receive))){
	for(h in 1:14){
if(h!=(k-(length(dummy.ints.attack)+preloop))){
day.array[,paste(h,".attack.",(k-(length(dummy.ints.attack)+preloop)), sep=""),k]<-1
}
}
}

#cop.startFE
for(h in 1:idno){
day.array[,paste(h,".cop.start", sep=""),((length(dummy.ints.attack)+preloop)+length(dummy.ints.receive)+1)]<-1
}


#cop.endFE
for(h in 1:idno){
day.array[,paste(h,".cop.end", sep=""),((length(dummy.ints.attack)+preloop)+length(dummy.ints.receive)+2)]<-1
}

#initFO.FE
for(h in 1:idno){
day.array[,paste(h,".initFO", sep=""),((length(dummy.ints.attack)+preloop)+length(dummy.ints.receive)+3)]<-1
}

#ID.initFO.FE
for(h in 1:14){
day.array[,paste(h,".initFO", sep=""),((length(dummy.ints.attack)+preloop)+length(dummy.ints.receive)+4)]<-1
}

#U.initFO.FE
for(h in 15:29){
day.array[,paste(h,".initFO", sep=""),((length(dummy.ints.attack)+preloop)+length(dummy.ints.receive)+5)]<-1
}

#initSF.FE
for(h in 1:idno){
day.array[,paste(h,".initSF", sep=""),((length(dummy.ints.attack)+preloop)+length(dummy.ints.receive)+6)]<-1
}


#everything else
for(k in (((length(dummy.ints.attack)+preloop)+length(dummy.ints.receive)+7):length(int.names))){
	
	day.array[,int.names[k],k]<-1
	
}

#weird extra formatting to make it match the package example, seems like it becomes relevant *punintended!* if I want to fit day-specific effects

global<-list()

global[[1]]<-day.array

int.list[[i]]<-global

names(int.list[[i]])<-"global"

}

tic()
intfit.sup<-rem(evlist, int.list, supplist=supplist, timing="interval", estimator="BPM")
toc()

summary(intfit.sup)

# Egocentric Relational Event Model (Interval Likelihood)
# 
#                   Post.Mode     Post.SD  Z value  Pr(>|z|)    
# Global.attackFE  -9.5259220   0.2886747 -32.9988 < 2.2e-16 ***
# X.receiveFE       3.7712555   0.2921739  12.9076 < 2.2e-16 ***
# X.UreceiveFE      3.9025235   0.3370996  11.5768 < 2.2e-16 ***
# U.attackFE       -1.2734082   0.6454967  -1.9728 0.0485233 *  
# U.receiveFE       1.7877281   0.3145761   5.6830 1.324e-08 ***
# U.UattackFE       2.9176286   0.3248928   8.9803 < 2.2e-16 ***
# 1.attackFE        1.0330374   0.3944376   2.6190 0.0088184 ** 
# 2.attackFE        1.2763648   0.3921908   3.2544 0.0011361 ** 
# 3.attackFE        1.6061436   0.3891684   4.1271 3.673e-05 ***
# 4.attackFE        2.5265882   0.3860257   6.5451 5.944e-11 ***
# 5.attackFE        0.3110660   0.4195142   0.7415 0.4583958    
# 6.attackFE       -0.0207165   0.8047343  -0.0257 0.9794621    
# 7.attackFE        1.3366638   0.3932445   3.3991 0.0006762 ***
# 8.attackFE       -1.4929214   1.0698668  -1.3954 0.1628870    
# 9.attackFE      -12.5412379 243.0571914  -0.0516 0.9588491    
# 10.attackFE       1.5764042   0.4001704   3.9393 8.171e-05 ***
# 11.attackFE     -10.4618029 264.1957859  -0.0396 0.9684131    
# 12.attackFE      -0.0121458   0.5502168  -0.0221 0.9823884    
# 13.attackFE       0.9853197   1.0715358   0.9195 0.3578134    
# 14.attackFE       0.7284843   0.5224387   1.3944 0.1631993    
# 1.receiveFE      -0.7315755   0.2770237  -2.6408 0.0082700 ** 
# 2.receiveFE      -0.2483185   0.2681292  -0.9261 0.3543862    
# 3.receiveFE      -0.3750730   0.2725416  -1.3762 0.1687582    
# 4.receiveFE       1.0326554   0.2610273   3.9561 7.618e-05 ***
# 5.receiveFE      -0.8724350   0.2925982  -2.9817 0.0028667 ** 
# 6.receiveFE       0.7239207   0.3403191   2.1272 0.0334050 *  
# 7.receiveFE       0.2399268   0.2626681   0.9134 0.3610207    
# 8.receiveFE       0.0322648   0.3395839   0.0950 0.9243046    
# 9.receiveFE       0.6527401   0.3048317   2.1413 0.0322488 *  
# 10.receiveFE     -0.0083194   0.2918528  -0.0285 0.9772590    
# 11.receiveFE      1.6776185   0.4538049   3.6968 0.0002183 ***
# 12.receiveFE     -0.4512307   0.3601768  -1.2528 0.2102774    
# 13.receiveFE     -0.3517487   1.0308039  -0.3412 0.7329249    
# cop.startFE      -8.8434147   0.1178511 -75.0389 < 2.2e-16 ***
# cop.endFE        -1.4478174   0.1414214 -10.2376 < 2.2e-16 ***
# initFO.FE         3.3524074   0.2581988  12.9838 < 2.2e-16 ***
# ID.initFO.FE      3.2420533   0.2612259  12.4109 < 2.2e-16 ***
# U.initFO.FE       1.8903400   0.3380617   5.5917 2.249e-08 ***
# initSF.FE         4.8788099   0.0710669  68.6509 < 2.2e-16 ***
# 1.initCH          2.6172926   1.0000010   2.6173 0.0088631 ** 
# 2.initCH         -9.1855050 280.2587561  -0.0328 0.9738539    
# 3.initCH         -8.9884961 283.0089737  -0.0318 0.9746631    
# 4.initCH          1.5141258   1.0000003   1.5141 0.1299940    
# 5.initCH          3.0791100   1.0000013   3.0791 0.0020762 ** 
# 6.initCH          6.7359045   0.2500001  26.9436 < 2.2e-16 ***
# 7.initCH         -9.5410217 275.4923345  -0.0346 0.9723727    
# 8.initCH          6.5629143   0.2425357  27.0596 < 2.2e-16 ***
# 9.initCH          6.8779021   0.1740777  39.5105 < 2.2e-16 ***
# 10.initCH         4.4654068   0.5000003   8.9308 < 2.2e-16 ***
# 11.initCH         6.7536032   0.4082485  16.5429 < 2.2e-16 ***
# 12.initCH        -7.3922442 308.7518796  -0.0239 0.9808986    
# 13.initCH        -5.0627152 363.2563881  -0.0139 0.9888802    
# 14.initCH         6.2717656   0.3333335  18.8153 < 2.2e-16 ***
# 15.initCH         5.4036745   0.7071078   7.6419 2.132e-14 ***
# 16.initCH         6.9077542   0.3535536  19.5381 < 2.2e-16 ***
# 17.initCH         6.2146003   1.0000033   6.2146 5.146e-10 ***
# 18.initCH         6.9077466   1.0000037   6.9077 4.925e-12 ***
# 19.initCH         5.8091412   0.5000004  11.6183 < 2.2e-16 ***
# 20.initCH         6.5976000   0.2132008  30.9455 < 2.2e-16 ***
# 21.initCH         6.9077466   1.0000037   6.9077 4.925e-12 ***
# 22.initCH         0.0000000 894.4271910   0.0000 1.0000000    
# 23.initCH        -5.0627152 363.2563881  -0.0139 0.9888802    
# 24.initCH         6.8505964   0.2425357  28.2457 < 2.2e-16 ***
# 25.initCH        -5.6466962 346.9319758  -0.0163 0.9870141    
# 26.initCH        -5.6466962 346.9319758  -0.0163 0.9870141    
# 27.initCH         6.0604566   0.3333335  18.1814 < 2.2e-16 ***
# 28.initCH        -5.9926922 338.2402555  -0.0177 0.9858644    
# 29.initCH        -5.0627152 363.2563881  -0.0139 0.9888802    
# 30.initCH         6.8243737   0.0455016 149.9810 < 2.2e-16 ***
# 1.disengage      -4.2622836   0.0626224 -68.0632 < 2.2e-16 ***
# 2.disengage      -5.1487697   0.0887356 -58.0237 < 2.2e-16 ***
# 3.disengage      -4.7750140   0.0636285 -75.0452 < 2.2e-16 ***
# 4.disengage      -4.6474529   0.0548821 -84.6806 < 2.2e-16 ***
# 5.disengage      -4.5737155   0.1025978 -44.5791 < 2.2e-16 ***
# 6.disengage      -6.4777863   0.5773494 -11.2199 < 2.2e-16 ***
# 7.disengage      -5.6831776   0.0894427 -63.5399 < 2.2e-16 ***
# 8.disengage      -3.0256324   0.1889822 -16.0101 < 2.2e-16 ***
# 9.disengage      -2.9513586   0.9999975  -2.9514 0.0031637 ** 
# 10.disengage     -4.6109312   0.0880451 -52.3701 < 2.2e-16 ***
# 11.disengage     -1.2519039   0.9999986  -1.2519 0.2106042    
# 12.disengage     -5.0999545   0.3333332 -15.2999 < 2.2e-16 ***
# 13.disengage     -3.0694899   0.9999975  -3.0695 0.0021442 ** 
# 14.disengage     -3.9995581   0.2000000 -19.9978 < 2.2e-16 ***
# 15.disengage     -3.7115624   0.3779643  -9.8199 < 2.2e-16 ***
# 16.disengage      0.0000000 894.4271910   0.0000 1.0000000    
# 17.disengage     -5.5854796   0.7071053  -7.8991 2.887e-15 ***
# 18.disengage      0.0000000 894.4271910   0.0000 1.0000000    
# 19.disengage     -3.3061330   0.2500000 -13.2245 < 2.2e-16 ***
# 20.disengage     -2.4505089   0.3535533  -6.9311 4.176e-12 ***
# 21.disengage      0.0000000 894.4271910   0.0000 1.0000000    
# 22.disengage     -3.5143729   0.9999972  -3.5144 0.0004408 ***
# 23.disengage    -12.1897538 246.2838524  -0.0495 0.9605250    
# 24.disengage     -1.8919041   0.9999982  -1.8919 0.0585033 .  
# 25.disengage     -2.4340658   0.2000000 -12.1703 < 2.2e-16 ***
# 26.disengage     -3.8049157   0.4082481  -9.3201 < 2.2e-16 ***
# 27.disengage     -5.3682937   0.3162276 -16.9760 < 2.2e-16 ***
# 28.disengage     -5.0921384   0.3535532 -14.4027 < 2.2e-16 ***
# 29.disengage     -5.0977870   0.4082480 -12.4870 < 2.2e-16 ***
# 30.disengage     -4.8310314   0.1622214 -29.7805 < 2.2e-16 ***
# 1.depart         -8.7014746   0.2357022 -36.9172 < 2.2e-16 ***
# 2.depart         -8.6455924   0.2357022 -36.6802 < 2.2e-16 ***
# 3.depart         -8.6031331   0.2357022 -36.5000 < 2.2e-16 ***
# 4.depart         -8.2131081   0.2357022 -34.8453 < 2.2e-16 ***
# 5.depart         -8.7978526   0.2886750 -30.4767 < 2.2e-16 ***
# 6.depart         -8.1073118   0.7071048 -11.4655 < 2.2e-16 ***
# 7.depart         -8.4665832   0.2357022 -35.9207 < 2.2e-16 ***
# 8.depart         -7.6244610   0.4999993 -15.2489 < 2.2e-16 ***
# 9.depart         -8.2131177   0.5773492 -14.2256 < 2.2e-16 ***
# 10.depart        -8.3366519   0.3779642 -22.0567 < 2.2e-16 ***
# 11.depart        -6.1361062   0.5773494 -10.6281 < 2.2e-16 ***
# 12.depart        -8.4305661   0.5773491 -14.6022 < 2.2e-16 ***
# 13.depart        -6.3742619   0.9999954  -6.3743 1.838e-10 ***
# 14.depart        -8.4328617   0.7071047 -11.9259 < 2.2e-16 ***
# 15.depart        -8.4724875   0.9999941  -8.4725 < 2.2e-16 ***
# 16.depart        -4.5531868   0.9999965  -4.5532 5.284e-06 ***
# 17.depart        -7.3325942   0.9999948  -7.3326 2.256e-13 ***
# 18.depart        -6.1293192   0.9999955  -6.1293 8.824e-10 ***
# 19.depart        -8.8972145   0.9999938  -8.8973 < 2.2e-16 ***
# 20.depart        -7.6761506   0.9999946  -7.6762 1.643e-14 ***
# 21.depart        -4.7116752   0.9999964  -4.7117 2.457e-06 ***
# 22.depart        -7.7353839   0.9999945  -7.7354 1.021e-14 ***
# 23.depart        -7.0172213   0.9999950  -7.0173 2.263e-12 ***
# 24.depart        -7.3915096   0.9999948  -7.3915 1.452e-13 ***
# 25.depart        -7.8421735   0.9999945  -7.8422 4.441e-15 ***
# 26.depart        -8.0356182   0.9999944  -8.0357 8.882e-16 ***
# 27.depart        -8.6596852   0.9999940  -8.6597 < 2.2e-16 ***
# 28.depart        -8.1918068   0.9999943  -8.1919 2.220e-16 ***
# 29.depart        -7.0548164   0.9999950  -7.0549 1.728e-12 ***
# 30.depart        -5.7379132   0.1178511 -48.6878 < 2.2e-16 ***
# ---
# Signif. codes:  0 ‘***’ 0.001 ‘**’ 0.01 ‘*’ 0.05 ‘.’ 0.1 ‘ ’ 1
# Null deviance: 76628.21 on 4854 degrees of freedom
# Residual deviance: 30159.03 on 4725 degrees of freedom
# 	Chi-square: 46469.19 on 129 degrees of freedom, asymptotic p-value 0 
# AIC: 30417.03 AICC: 30424.13 BIC: 31253.92 
# Log posterior: -144206 
# Prior parameters: mu=0 sigma=1000 nu=4 

#great!


### Partition out ID.initSF


sequences<-table(event.list$Date)

#create list of dummy intercept names

idno<-length(event.types[grep("cop.start",event.types)])

dummy.ints.attack<-paste(1:14,".attackFE", sep="")
dummy.ints.receive<-paste(1:13, ".receiveFE", sep="")


int.names<-c("Global.attackFE", "X.receiveFE", "X.UreceiveFE", "U.attackFE", "U.receiveFE", "U.UattackFE", dummy.ints.attack, dummy.ints.receive, "cop.startFE", "cop.endFE", "initFO.FE","ID.initFO.FE", "U.initFO.FE","initSF.FE","ID.initSF.FE",event.types[-c(grep("attack", event.types),grep("cop", event.types),grep("initFO", event.types),grep("initSF", event.types))])

int.list<-list()

for(i in 1:length(levels(event.list$Date))){

day.array<-array(0,dim=c(as.numeric(sequences[i]), length(event.types),length(int.names)))

#the third dimension is the stat slice.
dimnames(day.array)<-list(c(1:sequences[i]),as.character(event.types),as.character(int.names))

#add global attack effect, loops for "U" intercepts for attacks and receives, then one loop for typical intercepts

day.array[,grep("attack", event.types),1]<-1


for(h in 1:14){
day.array[,paste(h,".attack.",30, sep=""),2]<-1
}


for(h in 15:29){
day.array[,paste(h,".attack.",30, sep=""),3]<-1
}


for(h in 1:14){
	for(j in 15:29){	
day.array[,paste(j,".attack.", h, sep=""),4]<-1
}
}


for(h in 1:14){
	for(j in 15:29){
day.array[,paste(h,".attack.",j, sep=""),5]<-1
}
}

for(h in 15:29){
	for(j in 15:29){	
if(h!=j){
day.array[,paste(j,".attack.", h, sep=""),6]<-1
}
}
}

preloop<-6

#loop for dummy intercepts for attacks, loop for dummy intercepts for receives, add in cop.start and cop.end FEs, then one loop for typical intercepts

for(k in (preloop+1):(length(dummy.ints.attack)+preloop)){
	
	for(h in 1:14){
if(h!=(k-preloop)){
day.array[,paste((k-preloop),".attack.", h, sep=""),k]<-1
}
}
}

for(k in ((length(dummy.ints.attack)+preloop)+1):((length(dummy.ints.attack)+preloop)+length(dummy.ints.receive))){
	for(h in 1:14){
if(h!=(k-(length(dummy.ints.attack)+preloop))){
day.array[,paste(h,".attack.",(k-(length(dummy.ints.attack)+preloop)), sep=""),k]<-1
}
}
}

#cop.startFE
for(h in 1:idno){
day.array[,paste(h,".cop.start", sep=""),((length(dummy.ints.attack)+preloop)+length(dummy.ints.receive)+1)]<-1
}


#cop.endFE
for(h in 1:idno){
day.array[,paste(h,".cop.end", sep=""),((length(dummy.ints.attack)+preloop)+length(dummy.ints.receive)+2)]<-1
}

#initFO.FE
for(h in 1:idno){
day.array[,paste(h,".initFO", sep=""),((length(dummy.ints.attack)+preloop)+length(dummy.ints.receive)+3)]<-1
}

#ID.initFO.FE
for(h in 1:14){
day.array[,paste(h,".initFO", sep=""),((length(dummy.ints.attack)+preloop)+length(dummy.ints.receive)+4)]<-1
}

#U.initFO.FE
for(h in 15:29){
day.array[,paste(h,".initFO", sep=""),((length(dummy.ints.attack)+preloop)+length(dummy.ints.receive)+5)]<-1
}

#initSF.FE
for(h in 1:idno){
day.array[,paste(h,".initSF", sep=""),((length(dummy.ints.attack)+preloop)+length(dummy.ints.receive)+6)]<-1
}

#ID.initSF.FE
for(h in 1:14){
day.array[,paste(h,".initSF", sep=""),((length(dummy.ints.attack)+preloop)+length(dummy.ints.receive)+7)]<-1
}

#everything else
for(k in (((length(dummy.ints.attack)+preloop)+length(dummy.ints.receive)+8):length(int.names))){
	
	day.array[,int.names[k],k]<-1
	
}

#weird extra formatting to make it match the package example, seems like it becomes relevant *punintended!* if I want to fit day-specific effects

global<-list()

global[[1]]<-day.array

int.list[[i]]<-global

names(int.list[[i]])<-"global"

}


tic()
intfit.sup<-rem(evlist, int.list, supplist=supplist, timing="interval", estimator="BPM")
toc()

summary(intfit.sup)


# Egocentric Relational Event Model (Interval Likelihood)
# 
#                   Post.Mode     Post.SD  Z value  Pr(>|z|)    
# Global.attackFE  -9.5259220   0.2886747 -32.9988 < 2.2e-16 ***
# X.receiveFE       3.7712555   0.2921739  12.9076 < 2.2e-16 ***
# X.UreceiveFE      3.9025235   0.3370996  11.5768 < 2.2e-16 ***
# U.attackFE       -1.2734082   0.6454967  -1.9728 0.0485233 *  
# U.receiveFE       1.7877281   0.3145761   5.6830 1.324e-08 ***
# U.UattackFE       2.9176286   0.3248928   8.9803 < 2.2e-16 ***
# 1.attackFE        1.0330374   0.3944376   2.6190 0.0088184 ** 
# 2.attackFE        1.2763648   0.3921908   3.2544 0.0011361 ** 
# 3.attackFE        1.6061436   0.3891684   4.1271 3.673e-05 ***
# 4.attackFE        2.5265882   0.3860257   6.5451 5.944e-11 ***
# 5.attackFE        0.3110660   0.4195142   0.7415 0.4583958    
# 6.attackFE       -0.0207165   0.8047343  -0.0257 0.9794621    
# 7.attackFE        1.3366638   0.3932445   3.3991 0.0006762 ***
# 8.attackFE       -1.4929214   1.0698668  -1.3954 0.1628870    
# 9.attackFE      -12.5412382 243.0572191  -0.0516 0.9588491    
# 10.attackFE       1.5764042   0.4001704   3.9393 8.171e-05 ***
# 11.attackFE     -10.4618029 264.1957859  -0.0396 0.9684131    
# 12.attackFE      -0.0121458   0.5502168  -0.0221 0.9823884    
# 13.attackFE       0.9853197   1.0715358   0.9195 0.3578134    
# 14.attackFE       0.7284843   0.5224387   1.3944 0.1631993    
# 1.receiveFE      -0.7315755   0.2770237  -2.6408 0.0082700 ** 
# 2.receiveFE      -0.2483185   0.2681292  -0.9261 0.3543862    
# 3.receiveFE      -0.3750730   0.2725416  -1.3762 0.1687582    
# 4.receiveFE       1.0326554   0.2610273   3.9561 7.618e-05 ***
# 5.receiveFE      -0.8724350   0.2925982  -2.9817 0.0028667 ** 
# 6.receiveFE       0.7239207   0.3403191   2.1272 0.0334050 *  
# 7.receiveFE       0.2399268   0.2626681   0.9134 0.3610207    
# 8.receiveFE       0.0322648   0.3395839   0.0950 0.9243046    
# 9.receiveFE       0.6527401   0.3048317   2.1413 0.0322488 *  
# 10.receiveFE     -0.0083194   0.2918528  -0.0285 0.9772590    
# 11.receiveFE      1.6776185   0.4538049   3.6968 0.0002183 ***
# 12.receiveFE     -0.4512307   0.3601768  -1.2528 0.2102774    
# 13.receiveFE     -0.3517487   1.0308039  -0.3412 0.7329249    
# cop.startFE      -8.8434147   0.1178511 -75.0389 < 2.2e-16 ***
# cop.endFE        -1.4478174   0.1414214 -10.2376 < 2.2e-16 ***
# initFO.FE         3.3524074   0.2581988  12.9838 < 2.2e-16 ***
# ID.initFO.FE      3.2420533   0.2612259  12.4109 < 2.2e-16 ***
# U.initFO.FE       1.8903400   0.3380617   5.5917 2.249e-08 ***
# initSF.FE         4.4038003   0.1386751  31.7563 < 2.2e-16 ***
# ID.initSF.FE      0.7190683   0.1614933   4.4526 8.483e-06 ***
# 1.initCH          2.6172926   1.0000010   2.6173 0.0088631 ** 
# 2.initCH         -9.1855050 280.2587561  -0.0328 0.9738539    
# 3.initCH         -8.9884961 283.0089737  -0.0318 0.9746631    
# 4.initCH          1.5141258   1.0000003   1.5141 0.1299940    
# 5.initCH          3.0791100   1.0000013   3.0791 0.0020762 ** 
# 6.initCH          6.7359045   0.2500001  26.9436 < 2.2e-16 ***
# 7.initCH         -9.5410217 275.4923345  -0.0346 0.9723727    
# 8.initCH          6.5629143   0.2425357  27.0596 < 2.2e-16 ***
# 9.initCH          6.8779021   0.1740777  39.5105 < 2.2e-16 ***
# 10.initCH         4.4654068   0.5000003   8.9308 < 2.2e-16 ***
# 11.initCH         6.7536032   0.4082485  16.5429 < 2.2e-16 ***
# 12.initCH        -7.3922442 308.7518796  -0.0239 0.9808986    
# 13.initCH        -5.0627152 363.2563881  -0.0139 0.9888802    
# 14.initCH         6.2717656   0.3333335  18.8153 < 2.2e-16 ***
# 15.initCH         5.4036745   0.7071078   7.6419 2.132e-14 ***
# 16.initCH         6.9077542   0.3535536  19.5381 < 2.2e-16 ***
# 17.initCH         6.2146003   1.0000033   6.2146 5.146e-10 ***
# 18.initCH         6.9077466   1.0000037   6.9077 4.925e-12 ***
# 19.initCH         5.8091412   0.5000004  11.6183 < 2.2e-16 ***
# 20.initCH         6.5976000   0.2132008  30.9455 < 2.2e-16 ***
# 21.initCH         6.9077466   1.0000037   6.9077 4.925e-12 ***
# 22.initCH         0.0000000 894.4271910   0.0000 1.0000000    
# 23.initCH        -5.0627152 363.2563881  -0.0139 0.9888802    
# 24.initCH         6.8505964   0.2425357  28.2457 < 2.2e-16 ***
# 25.initCH        -5.6466962 346.9319758  -0.0163 0.9870141    
# 26.initCH        -5.6466962 346.9319758  -0.0163 0.9870141    
# 27.initCH         6.0604566   0.3333335  18.1814 < 2.2e-16 ***
# 28.initCH        -5.9926922 338.2402555  -0.0177 0.9858644    
# 29.initCH        -5.0627152 363.2563881  -0.0139 0.9888802    
# 30.initCH         6.8243737   0.0455016 149.9810 < 2.2e-16 ***
# 1.disengage      -4.2622836   0.0626224 -68.0632 < 2.2e-16 ***
# 2.disengage      -5.1487697   0.0887356 -58.0237 < 2.2e-16 ***
# 3.disengage      -4.7750140   0.0636285 -75.0452 < 2.2e-16 ***
# 4.disengage      -4.6474529   0.0548821 -84.6806 < 2.2e-16 ***
# 5.disengage      -4.5737155   0.1025978 -44.5791 < 2.2e-16 ***
# 6.disengage      -6.4777863   0.5773494 -11.2199 < 2.2e-16 ***
# 7.disengage      -5.6831776   0.0894427 -63.5399 < 2.2e-16 ***
# 8.disengage      -3.0256324   0.1889822 -16.0101 < 2.2e-16 ***
# 9.disengage      -2.9513586   0.9999975  -2.9514 0.0031637 ** 
# 10.disengage     -4.6109312   0.0880451 -52.3701 < 2.2e-16 ***
# 11.disengage     -1.2519039   0.9999986  -1.2519 0.2106042    
# 12.disengage     -5.0999545   0.3333332 -15.2999 < 2.2e-16 ***
# 13.disengage     -3.0694899   0.9999975  -3.0695 0.0021442 ** 
# 14.disengage     -3.9995581   0.2000000 -19.9978 < 2.2e-16 ***
# 15.disengage     -3.7115624   0.3779643  -9.8199 < 2.2e-16 ***
# 16.disengage      0.0000000 894.4271910   0.0000 1.0000000    
# 17.disengage     -5.5854796   0.7071053  -7.8991 2.887e-15 ***
# 18.disengage      0.0000000 894.4271910   0.0000 1.0000000    
# 19.disengage     -3.3061330   0.2500000 -13.2245 < 2.2e-16 ***
# 20.disengage     -2.4505089   0.3535533  -6.9311 4.176e-12 ***
# 21.disengage      0.0000000 894.4271910   0.0000 1.0000000    
# 22.disengage     -3.5143729   0.9999972  -3.5144 0.0004408 ***
# 23.disengage    -12.1897538 246.2838524  -0.0495 0.9605250    
# 24.disengage     -1.8919041   0.9999982  -1.8919 0.0585033 .  
# 25.disengage     -2.4340658   0.2000000 -12.1703 < 2.2e-16 ***
# 26.disengage     -3.8049157   0.4082481  -9.3201 < 2.2e-16 ***
# 27.disengage     -5.3682937   0.3162276 -16.9760 < 2.2e-16 ***
# 28.disengage     -5.0921384   0.3535532 -14.4027 < 2.2e-16 ***
# 29.disengage     -5.0977870   0.4082480 -12.4870 < 2.2e-16 ***
# 30.disengage     -4.8310314   0.1622214 -29.7805 < 2.2e-16 ***
# 1.depart         -8.7014746   0.2357022 -36.9172 < 2.2e-16 ***
# 2.depart         -8.6455924   0.2357022 -36.6802 < 2.2e-16 ***
# 3.depart         -8.6031331   0.2357022 -36.5000 < 2.2e-16 ***
# 4.depart         -8.2131081   0.2357022 -34.8453 < 2.2e-16 ***
# 5.depart         -8.7978526   0.2886750 -30.4767 < 2.2e-16 ***
# 6.depart         -8.1073118   0.7071048 -11.4655 < 2.2e-16 ***
# 7.depart         -8.4665832   0.2357022 -35.9207 < 2.2e-16 ***
# 8.depart         -7.6244610   0.4999993 -15.2489 < 2.2e-16 ***
# 9.depart         -8.2131177   0.5773492 -14.2256 < 2.2e-16 ***
# 10.depart        -8.3366519   0.3779642 -22.0567 < 2.2e-16 ***
# 11.depart        -6.1361062   0.5773494 -10.6281 < 2.2e-16 ***
# 12.depart        -8.4305661   0.5773491 -14.6022 < 2.2e-16 ***
# 13.depart        -6.3742619   0.9999954  -6.3743 1.838e-10 ***
# 14.depart        -8.4328617   0.7071047 -11.9259 < 2.2e-16 ***
# 15.depart        -8.4724875   0.9999941  -8.4725 < 2.2e-16 ***
# 16.depart        -4.5531868   0.9999965  -4.5532 5.284e-06 ***
# 17.depart        -7.3325942   0.9999948  -7.3326 2.256e-13 ***
# 18.depart        -6.1293192   0.9999955  -6.1293 8.824e-10 ***
# 19.depart        -8.8972145   0.9999938  -8.8973 < 2.2e-16 ***
# 20.depart        -7.6761506   0.9999946  -7.6762 1.643e-14 ***
# 21.depart        -4.7116752   0.9999964  -4.7117 2.457e-06 ***
# 22.depart        -7.7353839   0.9999945  -7.7354 1.021e-14 ***
# 23.depart        -7.0172213   0.9999950  -7.0173 2.263e-12 ***
# 24.depart        -7.3915096   0.9999948  -7.3915 1.452e-13 ***
# 25.depart        -7.8421735   0.9999945  -7.8422 4.441e-15 ***
# 26.depart        -8.0356182   0.9999944  -8.0357 8.882e-16 ***
# 27.depart        -8.6596852   0.9999940  -8.6597 < 2.2e-16 ***
# 28.depart        -8.1918068   0.9999943  -8.1919 2.220e-16 ***
# 29.depart        -7.0548164   0.9999950  -7.0549 1.728e-12 ***
# 30.depart        -5.7379132   0.1178511 -48.6878 < 2.2e-16 ***
# ---
# Signif. codes:  0 ‘***’ 0.001 ‘**’ 0.01 ‘*’ 0.05 ‘.’ 0.1 ‘ ’ 1
# Null deviance: 76628.21 on 4854 degrees of freedom
# Residual deviance: 30137.16 on 4724 degrees of freedom
# 	Chi-square: 46491.05 on 130 degrees of freedom, asymptotic p-value 0 
# AIC: 30397.16 AICC: 30404.37 BIC: 31240.55 
# Log posterior: -145196.1 
# Prior parameters: mu=0 sigma=1000 nu=4 

#even better!


#### Partition out IDs AND U's for initSF


sequences<-table(event.list$Date)

#create list of dummy intercept names

idno<-length(event.types[grep("cop.start",event.types)])

dummy.ints.attack<-paste(1:14,".attackFE", sep="")
dummy.ints.receive<-paste(1:13, ".receiveFE", sep="")


int.names<-c("Global.attackFE", "X.receiveFE", "X.UreceiveFE", "U.attackFE", "U.receiveFE", "U.UattackFE", dummy.ints.attack, dummy.ints.receive, "cop.startFE", "cop.endFE", "initFO.FE","ID.initFO.FE", "U.initFO.FE","initSF.FE","ID.initSF.FE","U.initSF.FE",event.types[-c(grep("attack", event.types),grep("cop", event.types),grep("initFO", event.types),grep("initSF", event.types))])

int.list<-list()

for(i in 1:length(levels(event.list$Date))){

day.array<-array(0,dim=c(as.numeric(sequences[i]), length(event.types),length(int.names)))

#the third dimension is the stat slice.
dimnames(day.array)<-list(c(1:sequences[i]),as.character(event.types),as.character(int.names))

#add global attack effect, loops for "U" intercepts for attacks and receives, then one loop for typical intercepts

day.array[,grep("attack", event.types),1]<-1


for(h in 1:14){
day.array[,paste(h,".attack.",30, sep=""),2]<-1
}


for(h in 15:29){
day.array[,paste(h,".attack.",30, sep=""),3]<-1
}


for(h in 1:14){
	for(j in 15:29){	
day.array[,paste(j,".attack.", h, sep=""),4]<-1
}
}


for(h in 1:14){
	for(j in 15:29){
day.array[,paste(h,".attack.",j, sep=""),5]<-1
}
}

for(h in 15:29){
	for(j in 15:29){	
if(h!=j){
day.array[,paste(j,".attack.", h, sep=""),6]<-1
}
}
}

preloop<-6

#loop for dummy intercepts for attacks, loop for dummy intercepts for receives, add in cop.start and cop.end FEs, then one loop for typical intercepts

for(k in (preloop+1):(length(dummy.ints.attack)+preloop)){
	
	for(h in 1:14){
if(h!=(k-preloop)){
day.array[,paste((k-preloop),".attack.", h, sep=""),k]<-1
}
}
}

for(k in ((length(dummy.ints.attack)+preloop)+1):((length(dummy.ints.attack)+preloop)+length(dummy.ints.receive))){
	for(h in 1:14){
if(h!=(k-(length(dummy.ints.attack)+preloop))){
day.array[,paste(h,".attack.",(k-(length(dummy.ints.attack)+preloop)), sep=""),k]<-1
}
}
}

#cop.startFE
for(h in 1:idno){
day.array[,paste(h,".cop.start", sep=""),((length(dummy.ints.attack)+preloop)+length(dummy.ints.receive)+1)]<-1
}

#cop.endFE
for(h in 1:idno){
day.array[,paste(h,".cop.end", sep=""),((length(dummy.ints.attack)+preloop)+length(dummy.ints.receive)+2)]<-1
}

#initFO.FE
for(h in 1:idno){
day.array[,paste(h,".initFO", sep=""),((length(dummy.ints.attack)+preloop)+length(dummy.ints.receive)+3)]<-1
}

#ID.initFO.FE
for(h in 1:14){
day.array[,paste(h,".initFO", sep=""),((length(dummy.ints.attack)+preloop)+length(dummy.ints.receive)+4)]<-1
}

#U.initFO.FE
for(h in 15:29){
day.array[,paste(h,".initFO", sep=""),((length(dummy.ints.attack)+preloop)+length(dummy.ints.receive)+5)]<-1
}

#initSF.FE
for(h in 1:idno){
day.array[,paste(h,".initSF", sep=""),((length(dummy.ints.attack)+preloop)+length(dummy.ints.receive)+6)]<-1
}

#ID.initSF.FE
for(h in 1:14){
day.array[,paste(h,".initSF", sep=""),((length(dummy.ints.attack)+preloop)+length(dummy.ints.receive)+7)]<-1
}

#U.initSF.FE
for(h in 15:29){
day.array[,paste(h,".initSF", sep=""),((length(dummy.ints.attack)+preloop)+length(dummy.ints.receive)+8)]<-1
}

#everything else
for(k in (((length(dummy.ints.attack)+preloop)+length(dummy.ints.receive)+9):length(int.names))){
	
	day.array[,int.names[k],k]<-1
	
}

#weird extra formatting to make it match the package example, seems like it becomes relevant *punintended!* if I want to fit day-specific effects

global<-list()

global[[1]]<-day.array

int.list[[i]]<-global

names(int.list[[i]])<-"global"

}


tic()
intfit.sup<-rem(evlist, int.list, supplist=supplist, timing="interval", estimator="BPM")
toc()

summary(intfit.sup)

# Egocentric Relational Event Model (Interval Likelihood)
# 
#                   Post.Mode     Post.SD  Z value  Pr(>|z|)    
# Global.attackFE  -9.5259220   0.2886747 -32.9988 < 2.2e-16 ***
# X.receiveFE       3.7712555   0.2921739  12.9076 < 2.2e-16 ***
# X.UreceiveFE      3.9025235   0.3370996  11.5768 < 2.2e-16 ***
# U.attackFE       -1.2734082   0.6454967  -1.9728 0.0485233 *  
# U.receiveFE       1.7877281   0.3145761   5.6830 1.324e-08 ***
# U.UattackFE       2.9176286   0.3248928   8.9803 < 2.2e-16 ***
# 1.attackFE        1.0330374   0.3944376   2.6190 0.0088184 ** 
# 2.attackFE        1.2763648   0.3921908   3.2544 0.0011361 ** 
# 3.attackFE        1.6061436   0.3891684   4.1271 3.673e-05 ***
# 4.attackFE        2.5265882   0.3860257   6.5451 5.944e-11 ***
# 5.attackFE        0.3110660   0.4195142   0.7415 0.4583958    
# 6.attackFE       -0.0207165   0.8047343  -0.0257 0.9794621    
# 7.attackFE        1.3366638   0.3932445   3.3991 0.0006762 ***
# 8.attackFE       -1.4929214   1.0698668  -1.3954 0.1628870    
# 9.attackFE      -12.5412380 243.0572031  -0.0516 0.9588491    
# 10.attackFE       1.5764042   0.4001704   3.9393 8.171e-05 ***
# 11.attackFE     -10.4618029 264.1957859  -0.0396 0.9684131    
# 12.attackFE      -0.0121458   0.5502168  -0.0221 0.9823884    
# 13.attackFE       0.9853197   1.0715358   0.9195 0.3578134    
# 14.attackFE       0.7284843   0.5224387   1.3944 0.1631993    
# 1.receiveFE      -0.7315755   0.2770237  -2.6408 0.0082700 ** 
# 2.receiveFE      -0.2483185   0.2681292  -0.9261 0.3543862    
# 3.receiveFE      -0.3750730   0.2725416  -1.3762 0.1687582    
# 4.receiveFE       1.0326554   0.2610273   3.9561 7.618e-05 ***
# 5.receiveFE      -0.8724350   0.2925982  -2.9817 0.0028667 ** 
# 6.receiveFE       0.7239207   0.3403191   2.1272 0.0334050 *  
# 7.receiveFE       0.2399268   0.2626681   0.9134 0.3610207    
# 8.receiveFE       0.0322648   0.3395839   0.0950 0.9243046    
# 9.receiveFE       0.6527401   0.3048317   2.1413 0.0322488 *  
# 10.receiveFE     -0.0083194   0.2918528  -0.0285 0.9772590    
# 11.receiveFE      1.6776185   0.4538049   3.6968 0.0002183 ***
# 12.receiveFE     -0.4512307   0.3601768  -1.2528 0.2102774    
# 13.receiveFE     -0.3517487   1.0308039  -0.3412 0.7329249    
# cop.startFE      -8.8434147   0.1178511 -75.0389 < 2.2e-16 ***
# cop.endFE        -1.4478174   0.1414214 -10.2376 < 2.2e-16 ***
# initFO.FE         3.3524074   0.2581988  12.9838 < 2.2e-16 ***
# ID.initFO.FE      3.2420533   0.2612259  12.4109 < 2.2e-16 ***
# U.initFO.FE       1.8903400   0.3380617   5.5917 2.249e-08 ***
# initSF.FE         3.9401938   0.1924501  20.4738 < 2.2e-16 ***
# ID.initSF.FE      1.1826749   0.2094907   5.6455 1.647e-08 ***
# U.initSF.FE       1.4769070   0.2775555   5.3211 1.031e-07 ***
# 1.initCH          2.6172926   1.0000010   2.6173 0.0088631 ** 
# 2.initCH         -9.1855050 280.2587561  -0.0328 0.9738539    
# 3.initCH         -8.9884961 283.0089737  -0.0318 0.9746631    
# 4.initCH          1.5141258   1.0000003   1.5141 0.1299940    
# 5.initCH          3.0791100   1.0000013   3.0791 0.0020762 ** 
# 6.initCH          6.7359045   0.2500001  26.9436 < 2.2e-16 ***
# 7.initCH         -9.5410217 275.4923345  -0.0346 0.9723727    
# 8.initCH          6.5629143   0.2425357  27.0596 < 2.2e-16 ***
# 9.initCH          6.8779021   0.1740777  39.5105 < 2.2e-16 ***
# 10.initCH         4.4654068   0.5000003   8.9308 < 2.2e-16 ***
# 11.initCH         6.7536032   0.4082485  16.5429 < 2.2e-16 ***
# 12.initCH        -7.3922442 308.7518796  -0.0239 0.9808986    
# 13.initCH        -5.0627152 363.2563881  -0.0139 0.9888802    
# 14.initCH         6.2717656   0.3333335  18.8153 < 2.2e-16 ***
# 15.initCH         5.4036745   0.7071078   7.6419 2.132e-14 ***
# 16.initCH         6.9077542   0.3535536  19.5381 < 2.2e-16 ***
# 17.initCH         6.2146003   1.0000033   6.2146 5.146e-10 ***
# 18.initCH         6.9077466   1.0000037   6.9077 4.925e-12 ***
# 19.initCH         5.8091412   0.5000004  11.6183 < 2.2e-16 ***
# 20.initCH         6.5976000   0.2132008  30.9455 < 2.2e-16 ***
# 21.initCH         6.9077466   1.0000037   6.9077 4.925e-12 ***
# 22.initCH         0.0000000 894.4271910   0.0000 1.0000000    
# 23.initCH        -5.0627152 363.2563881  -0.0139 0.9888802    
# 24.initCH         6.8505964   0.2425357  28.2457 < 2.2e-16 ***
# 25.initCH        -5.6466962 346.9319758  -0.0163 0.9870141    
# 26.initCH        -5.6466962 346.9319758  -0.0163 0.9870141    
# 27.initCH         6.0604566   0.3333335  18.1814 < 2.2e-16 ***
# 28.initCH        -5.9926922 338.2402555  -0.0177 0.9858644    
# 29.initCH        -5.0627152 363.2563881  -0.0139 0.9888802    
# 30.initCH         6.8243737   0.0455016 149.9810 < 2.2e-16 ***
# 1.disengage      -4.2622836   0.0626224 -68.0632 < 2.2e-16 ***
# 2.disengage      -5.1487697   0.0887356 -58.0237 < 2.2e-16 ***
# 3.disengage      -4.7750140   0.0636285 -75.0452 < 2.2e-16 ***
# 4.disengage      -4.6474529   0.0548821 -84.6806 < 2.2e-16 ***
# 5.disengage      -4.5737155   0.1025978 -44.5791 < 2.2e-16 ***
# 6.disengage      -6.4777863   0.5773494 -11.2199 < 2.2e-16 ***
# 7.disengage      -5.6831776   0.0894427 -63.5399 < 2.2e-16 ***
# 8.disengage      -3.0256324   0.1889822 -16.0101 < 2.2e-16 ***
# 9.disengage      -2.9513586   0.9999975  -2.9514 0.0031637 ** 
# 10.disengage     -4.6109312   0.0880451 -52.3701 < 2.2e-16 ***
# 11.disengage     -1.2519039   0.9999986  -1.2519 0.2106042    
# 12.disengage     -5.0999545   0.3333332 -15.2999 < 2.2e-16 ***
# 13.disengage     -3.0694899   0.9999975  -3.0695 0.0021442 ** 
# 14.disengage     -3.9995581   0.2000000 -19.9978 < 2.2e-16 ***
# 15.disengage     -3.7115624   0.3779643  -9.8199 < 2.2e-16 ***
# 16.disengage      0.0000000 894.4271910   0.0000 1.0000000    
# 17.disengage     -5.5854796   0.7071053  -7.8991 2.887e-15 ***
# 18.disengage      0.0000000 894.4271910   0.0000 1.0000000    
# 19.disengage     -3.3061330   0.2500000 -13.2245 < 2.2e-16 ***
# 20.disengage     -2.4505089   0.3535533  -6.9311 4.176e-12 ***
# 21.disengage      0.0000000 894.4271910   0.0000 1.0000000    
# 22.disengage     -3.5143729   0.9999972  -3.5144 0.0004408 ***
# 23.disengage    -12.1897538 246.2838524  -0.0495 0.9605250    
# 24.disengage     -1.8919041   0.9999982  -1.8919 0.0585033 .  
# 25.disengage     -2.4340658   0.2000000 -12.1703 < 2.2e-16 ***
# 26.disengage     -3.8049157   0.4082481  -9.3201 < 2.2e-16 ***
# 27.disengage     -5.3682937   0.3162276 -16.9760 < 2.2e-16 ***
# 28.disengage     -5.0921384   0.3535532 -14.4027 < 2.2e-16 ***
# 29.disengage     -5.0977870   0.4082480 -12.4870 < 2.2e-16 ***
# 30.disengage     -4.8310314   0.1622214 -29.7805 < 2.2e-16 ***
# 1.depart         -8.7014746   0.2357022 -36.9172 < 2.2e-16 ***
# 2.depart         -8.6455924   0.2357022 -36.6802 < 2.2e-16 ***
# 3.depart         -8.6031331   0.2357022 -36.5000 < 2.2e-16 ***
# 4.depart         -8.2131081   0.2357022 -34.8453 < 2.2e-16 ***
# 5.depart         -8.7978526   0.2886750 -30.4767 < 2.2e-16 ***
# 6.depart         -8.1073118   0.7071048 -11.4655 < 2.2e-16 ***
# 7.depart         -8.4665832   0.2357022 -35.9207 < 2.2e-16 ***
# 8.depart         -7.6244610   0.4999993 -15.2489 < 2.2e-16 ***
# 9.depart         -8.2131177   0.5773492 -14.2256 < 2.2e-16 ***
# 10.depart        -8.3366519   0.3779642 -22.0567 < 2.2e-16 ***
# 11.depart        -6.1361062   0.5773494 -10.6281 < 2.2e-16 ***
# 12.depart        -8.4305661   0.5773491 -14.6022 < 2.2e-16 ***
# 13.depart        -6.3742619   0.9999954  -6.3743 1.838e-10 ***
# 14.depart        -8.4328617   0.7071047 -11.9259 < 2.2e-16 ***
# 15.depart        -8.4724875   0.9999941  -8.4725 < 2.2e-16 ***
# 16.depart        -4.5531868   0.9999965  -4.5532 5.284e-06 ***
# 17.depart        -7.3325942   0.9999948  -7.3326 2.256e-13 ***
# 18.depart        -6.1293192   0.9999955  -6.1293 8.824e-10 ***
# 19.depart        -8.8972145   0.9999938  -8.8973 < 2.2e-16 ***
# 20.depart        -7.6761506   0.9999946  -7.6762 1.643e-14 ***
# 21.depart        -4.7116752   0.9999964  -4.7117 2.457e-06 ***
# 22.depart        -7.7353839   0.9999945  -7.7354 1.021e-14 ***
# 23.depart        -7.0172213   0.9999950  -7.0173 2.263e-12 ***
# 24.depart        -7.3915096   0.9999948  -7.3915 1.452e-13 ***
# 25.depart        -7.8421735   0.9999945  -7.8422 4.441e-15 ***
# 26.depart        -8.0356182   0.9999944  -8.0357 8.882e-16 ***
# 27.depart        -8.6596852   0.9999940  -8.6597 < 2.2e-16 ***
# 28.depart        -8.1918068   0.9999943  -8.1919 2.220e-16 ***
# 29.depart        -7.0548164   0.9999950  -7.0549 1.728e-12 ***
# 30.depart        -5.7379132   0.1178511 -48.6878 < 2.2e-16 ***
# ---
# Signif. codes:  0 ‘***’ 0.001 ‘**’ 0.01 ‘*’ 0.05 ‘.’ 0.1 ‘ ’ 1
# Null deviance: 76628.21 on 4854 degrees of freedom
# Residual deviance: 30111.53 on 4723 degrees of freedom
# 	Chi-square: 46516.68 on 131 degrees of freedom, asymptotic p-value 0 
# AIC: 30373.53 AICC: 30380.86 BIC: 31223.4 
# Log posterior: -146184.3 
# Prior parameters: mu=0 sigma=1000 nu=4 


#another new best! one more


#What about just taking out ?? males and lumping the rest?


sequences<-table(event.list$Date)

#create list of dummy intercept names

idno<-length(event.types[grep("cop.start",event.types)])

dummy.ints.attack<-paste(1:14,".attackFE", sep="")
dummy.ints.receive<-paste(1:13, ".receiveFE", sep="")


int.names<-c("Global.attackFE", "X.receiveFE", "X.UreceiveFE", "U.attackFE", "U.receiveFE", "U.UattackFE", dummy.ints.attack, dummy.ints.receive, "cop.startFE", "cop.endFE", "initFO.FE","ID.initFO.FE", "U.initFO.FE","initSF.FE","X.initSF.FE", event.types[-c(grep("attack", event.types),grep("cop", event.types),grep("initFO", event.types),grep("initSF", event.types))])

int.list<-list()

for(i in 1:length(levels(event.list$Date))){

day.array<-array(0,dim=c(as.numeric(sequences[i]), length(event.types),length(int.names)))

#the third dimension is the stat slice.
dimnames(day.array)<-list(c(1:sequences[i]),as.character(event.types),as.character(int.names))

#add global attack effect, loops for "U" intercepts for attacks and receives, then one loop for typical intercepts

day.array[,grep("attack", event.types),1]<-1


for(h in 1:14){
day.array[,paste(h,".attack.",30, sep=""),2]<-1
}


for(h in 15:29){
day.array[,paste(h,".attack.",30, sep=""),3]<-1
}


for(h in 1:14){
	for(j in 15:29){	
day.array[,paste(j,".attack.", h, sep=""),4]<-1
}
}


for(h in 1:14){
	for(j in 15:29){
day.array[,paste(h,".attack.",j, sep=""),5]<-1
}
}

for(h in 15:29){
	for(j in 15:29){	
if(h!=j){
day.array[,paste(j,".attack.", h, sep=""),6]<-1
}
}
}

preloop<-6

#loop for dummy intercepts for attacks, loop for dummy intercepts for receives, add in cop.start and cop.end FEs, then one loop for typical intercepts

for(k in (preloop+1):(length(dummy.ints.attack)+preloop)){
	
	for(h in 1:14){
if(h!=(k-preloop)){
day.array[,paste((k-preloop),".attack.", h, sep=""),k]<-1
}
}
}

for(k in ((length(dummy.ints.attack)+preloop)+1):((length(dummy.ints.attack)+preloop)+length(dummy.ints.receive))){
	for(h in 1:14){
if(h!=(k-(length(dummy.ints.attack)+preloop))){
day.array[,paste(h,".attack.",(k-(length(dummy.ints.attack)+preloop)), sep=""),k]<-1
}
}
}

#cop.startFE
for(h in 1:idno){
day.array[,paste(h,".cop.start", sep=""),((length(dummy.ints.attack)+preloop)+length(dummy.ints.receive)+1)]<-1
}


#cop.endFE
for(h in 1:idno){
day.array[,paste(h,".cop.end", sep=""),((length(dummy.ints.attack)+preloop)+length(dummy.ints.receive)+2)]<-1
}

#initFO.FE
for(h in 1:idno){
day.array[,paste(h,".initFO", sep=""),((length(dummy.ints.attack)+preloop)+length(dummy.ints.receive)+3)]<-1
}

#ID.initFO.FE
for(h in 1:14){
day.array[,paste(h,".initFO", sep=""),((length(dummy.ints.attack)+preloop)+length(dummy.ints.receive)+4)]<-1
}

#U.initFO.FE
for(h in 15:29){
day.array[,paste(h,".initFO", sep=""),((length(dummy.ints.attack)+preloop)+length(dummy.ints.receive)+5)]<-1
}

#initSF.FE
for(h in 1:idno){
day.array[,paste(h,".initSF", sep=""),((length(dummy.ints.attack)+preloop)+length(dummy.ints.receive)+6)]<-1
}

#X.initSF.FE

day.array[,paste(30,".initSF", sep=""),((length(dummy.ints.attack)+preloop)+length(dummy.ints.receive)+7)]<-1


#everything else
for(k in (((length(dummy.ints.attack)+preloop)+length(dummy.ints.receive)+8):length(int.names))){
	
	day.array[,int.names[k],k]<-1
	
}

#weird extra formatting to make it match the package example, seems like it becomes relevant *punintended!* if I want to fit day-specific effects

global<-list()

global[[1]]<-day.array

int.list[[i]]<-global

names(int.list[[i]])<-"global"

}


tic()
intfit.sup<-rem(evlist, int.list, supplist=supplist, timing="interval", estimator="BPM")
toc()

summary(intfit.sup)

# Egocentric Relational Event Model (Interval Likelihood)
# 
#                   Post.Mode     Post.SD  Z value  Pr(>|z|)    
# Global.attackFE  -9.5259220   0.2886747 -32.9988 < 2.2e-16 ***
# X.receiveFE       3.7712555   0.2921739  12.9076 < 2.2e-16 ***
# X.UreceiveFE      3.9025235   0.3370996  11.5768 < 2.2e-16 ***
# U.attackFE       -1.2734082   0.6454967  -1.9728 0.0485233 *  
# U.receiveFE       1.7877281   0.3145761   5.6830 1.324e-08 ***
# U.UattackFE       2.9176286   0.3248928   8.9803 < 2.2e-16 ***
# 1.attackFE        1.0330374   0.3944376   2.6190 0.0088184 ** 
# 2.attackFE        1.2763648   0.3921908   3.2544 0.0011361 ** 
# 3.attackFE        1.6061436   0.3891684   4.1271 3.673e-05 ***
# 4.attackFE        2.5265882   0.3860257   6.5451 5.944e-11 ***
# 5.attackFE        0.3110660   0.4195142   0.7415 0.4583958    
# 6.attackFE       -0.0207165   0.8047343  -0.0257 0.9794621    
# 7.attackFE        1.3366638   0.3932445   3.3991 0.0006762 ***
# 8.attackFE       -1.4929214   1.0698668  -1.3954 0.1628870    
# 9.attackFE      -12.5412370 243.0570882  -0.0516 0.9588491    
# 10.attackFE       1.5764042   0.4001704   3.9393 8.171e-05 ***
# 11.attackFE     -10.4618029 264.1957859  -0.0396 0.9684131    
# 12.attackFE      -0.0121458   0.5502168  -0.0221 0.9823884    
# 13.attackFE       0.9853197   1.0715358   0.9195 0.3578134    
# 14.attackFE       0.7284843   0.5224387   1.3944 0.1631993    
# 1.receiveFE      -0.7315755   0.2770237  -2.6408 0.0082700 ** 
# 2.receiveFE      -0.2483185   0.2681292  -0.9261 0.3543862    
# 3.receiveFE      -0.3750730   0.2725416  -1.3762 0.1687582    
# 4.receiveFE       1.0326554   0.2610273   3.9561 7.618e-05 ***
# 5.receiveFE      -0.8724350   0.2925982  -2.9817 0.0028667 ** 
# 6.receiveFE       0.7239207   0.3403191   2.1272 0.0334050 *  
# 7.receiveFE       0.2399268   0.2626681   0.9134 0.3610207    
# 8.receiveFE       0.0322648   0.3395839   0.0950 0.9243046    
# 9.receiveFE       0.6527401   0.3048317   2.1413 0.0322488 *  
# 10.receiveFE     -0.0083194   0.2918528  -0.0285 0.9772590    
# 11.receiveFE      1.6776185   0.4538049   3.6968 0.0002183 ***
# 12.receiveFE     -0.4512307   0.3601768  -1.2528 0.2102774    
# 13.receiveFE     -0.3517487   1.0308039  -0.3412 0.7329249    
# cop.startFE      -8.8434147   0.1178511 -75.0389 < 2.2e-16 ***
# cop.endFE        -1.4478174   0.1414214 -10.2376 < 2.2e-16 ***
# initFO.FE         3.3524074   0.2581988  12.9838 < 2.2e-16 ***
# ID.initFO.FE      3.2420533   0.2612259  12.4109 < 2.2e-16 ***
# U.initFO.FE       1.8903400   0.3380617   5.5917 2.249e-08 ***
# initSF.FE         5.1608463   0.0764719  67.4868 < 2.2e-16 ***
# X.initSF.FE      -1.2206524   0.2070869  -5.8944 3.761e-09 ***
# 1.initCH          2.6172926   1.0000010   2.6173 0.0088631 ** 
# 2.initCH         -9.1855050 280.2587561  -0.0328 0.9738539    
# 3.initCH         -8.9884961 283.0089737  -0.0318 0.9746631    
# 4.initCH          1.5141258   1.0000003   1.5141 0.1299940    
# 5.initCH          3.0791100   1.0000013   3.0791 0.0020762 ** 
# 6.initCH          6.7359045   0.2500001  26.9436 < 2.2e-16 ***
# 7.initCH         -9.5410217 275.4923345  -0.0346 0.9723727    
# 8.initCH          6.5629143   0.2425357  27.0596 < 2.2e-16 ***
# 9.initCH          6.8779021   0.1740777  39.5105 < 2.2e-16 ***
# 10.initCH         4.4654068   0.5000003   8.9308 < 2.2e-16 ***
# 11.initCH         6.7536032   0.4082485  16.5429 < 2.2e-16 ***
# 12.initCH        -7.3922442 308.7518796  -0.0239 0.9808986    
# 13.initCH        -5.0627152 363.2563881  -0.0139 0.9888802    
# 14.initCH         6.2717656   0.3333335  18.8153 < 2.2e-16 ***
# 15.initCH         5.4036745   0.7071078   7.6419 2.132e-14 ***
# 16.initCH         6.9077542   0.3535536  19.5381 < 2.2e-16 ***
# 17.initCH         6.2146003   1.0000033   6.2146 5.146e-10 ***
# 18.initCH         6.9077466   1.0000037   6.9077 4.925e-12 ***
# 19.initCH         5.8091412   0.5000004  11.6183 < 2.2e-16 ***
# 20.initCH         6.5976000   0.2132008  30.9455 < 2.2e-16 ***
# 21.initCH         6.9077466   1.0000037   6.9077 4.925e-12 ***
# 22.initCH         0.0000000 894.4271910   0.0000 1.0000000    
# 23.initCH        -5.0627152 363.2563881  -0.0139 0.9888802    
# 24.initCH         6.8505964   0.2425357  28.2457 < 2.2e-16 ***
# 25.initCH        -5.6466962 346.9319758  -0.0163 0.9870141    
# 26.initCH        -5.6466962 346.9319758  -0.0163 0.9870141    
# 27.initCH         6.0604566   0.3333335  18.1814 < 2.2e-16 ***
# 28.initCH        -5.9926922 338.2402555  -0.0177 0.9858644    
# 29.initCH        -5.0627152 363.2563881  -0.0139 0.9888802    
# 30.initCH         6.8243737   0.0455016 149.9810 < 2.2e-16 ***
# 1.disengage      -4.2622836   0.0626224 -68.0632 < 2.2e-16 ***
# 2.disengage      -5.1487697   0.0887356 -58.0237 < 2.2e-16 ***
# 3.disengage      -4.7750140   0.0636285 -75.0452 < 2.2e-16 ***
# 4.disengage      -4.6474529   0.0548821 -84.6806 < 2.2e-16 ***
# 5.disengage      -4.5737155   0.1025978 -44.5791 < 2.2e-16 ***
# 6.disengage      -6.4777863   0.5773494 -11.2199 < 2.2e-16 ***
# 7.disengage      -5.6831776   0.0894427 -63.5399 < 2.2e-16 ***
# 8.disengage      -3.0256324   0.1889822 -16.0101 < 2.2e-16 ***
# 9.disengage      -2.9513586   0.9999975  -2.9514 0.0031637 ** 
# 10.disengage     -4.6109312   0.0880451 -52.3701 < 2.2e-16 ***
# 11.disengage     -1.2519039   0.9999986  -1.2519 0.2106042    
# 12.disengage     -5.0999545   0.3333332 -15.2999 < 2.2e-16 ***
# 13.disengage     -3.0694899   0.9999975  -3.0695 0.0021442 ** 
# 14.disengage     -3.9995581   0.2000000 -19.9978 < 2.2e-16 ***
# 15.disengage     -3.7115624   0.3779643  -9.8199 < 2.2e-16 ***
# 16.disengage      0.0000000 894.4271910   0.0000 1.0000000    
# 17.disengage     -5.5854796   0.7071053  -7.8991 2.887e-15 ***
# 18.disengage      0.0000000 894.4271910   0.0000 1.0000000    
# 19.disengage     -3.3061330   0.2500000 -13.2245 < 2.2e-16 ***
# 20.disengage     -2.4505089   0.3535533  -6.9311 4.176e-12 ***
# 21.disengage      0.0000000 894.4271910   0.0000 1.0000000    
# 22.disengage     -3.5143729   0.9999972  -3.5144 0.0004408 ***
# 23.disengage    -12.1897538 246.2838524  -0.0495 0.9605250    
# 24.disengage     -1.8919041   0.9999982  -1.8919 0.0585033 .  
# 25.disengage     -2.4340658   0.2000000 -12.1703 < 2.2e-16 ***
# 26.disengage     -3.8049157   0.4082481  -9.3201 < 2.2e-16 ***
# 27.disengage     -5.3682937   0.3162276 -16.9760 < 2.2e-16 ***
# 28.disengage     -5.0921384   0.3535532 -14.4027 < 2.2e-16 ***
# 29.disengage     -5.0977870   0.4082480 -12.4870 < 2.2e-16 ***
# 30.disengage     -4.8310314   0.1622214 -29.7805 < 2.2e-16 ***
# 1.depart         -8.7014746   0.2357022 -36.9172 < 2.2e-16 ***
# 2.depart         -8.6455924   0.2357022 -36.6802 < 2.2e-16 ***
# 3.depart         -8.6031331   0.2357022 -36.5000 < 2.2e-16 ***
# 4.depart         -8.2131081   0.2357022 -34.8453 < 2.2e-16 ***
# 5.depart         -8.7978526   0.2886750 -30.4767 < 2.2e-16 ***
# 6.depart         -8.1073118   0.7071048 -11.4655 < 2.2e-16 ***
# 7.depart         -8.4665832   0.2357022 -35.9207 < 2.2e-16 ***
# 8.depart         -7.6244610   0.4999993 -15.2489 < 2.2e-16 ***
# 9.depart         -8.2131177   0.5773492 -14.2256 < 2.2e-16 ***
# 10.depart        -8.3366519   0.3779642 -22.0567 < 2.2e-16 ***
# 11.depart        -6.1361062   0.5773494 -10.6281 < 2.2e-16 ***
# 12.depart        -8.4305661   0.5773491 -14.6022 < 2.2e-16 ***
# 13.depart        -6.3742619   0.9999954  -6.3743 1.838e-10 ***
# 14.depart        -8.4328617   0.7071047 -11.9259 < 2.2e-16 ***
# 15.depart        -8.4724875   0.9999941  -8.4725 < 2.2e-16 ***
# 16.depart        -4.5531868   0.9999965  -4.5532 5.284e-06 ***
# 17.depart        -7.3325942   0.9999948  -7.3326 2.256e-13 ***
# 18.depart        -6.1293192   0.9999955  -6.1293 8.824e-10 ***
# 19.depart        -8.8972145   0.9999938  -8.8973 < 2.2e-16 ***
# 20.depart        -7.6761506   0.9999946  -7.6762 1.643e-14 ***
# 21.depart        -4.7116752   0.9999964  -4.7117 2.457e-06 ***
# 22.depart        -7.7353839   0.9999945  -7.7354 1.021e-14 ***
# 23.depart        -7.0172213   0.9999950  -7.0173 2.263e-12 ***
# 24.depart        -7.3915096   0.9999948  -7.3915 1.452e-13 ***
# 25.depart        -7.8421735   0.9999945  -7.8422 4.441e-15 ***
# 26.depart        -8.0356182   0.9999944  -8.0357 8.882e-16 ***
# 27.depart        -8.6596852   0.9999940  -8.6597 < 2.2e-16 ***
# 28.depart        -8.1918068   0.9999943  -8.1919 2.220e-16 ***
# 29.depart        -7.0548164   0.9999950  -7.0549 1.728e-12 ***
# 30.depart        -5.7379132   0.1178511 -48.6878 < 2.2e-16 ***
# ---
# Signif. codes:  0 ‘***’ 0.001 ‘**’ 0.01 ‘*’ 0.05 ‘.’ 0.1 ‘ ’ 1
# Null deviance: 76628.21 on 4854 degrees of freedom
# Residual deviance: 30113.26 on 4724 degrees of freedom
# 	Chi-square: 46514.96 on 130 degrees of freedom, asymptotic p-value 0 
# AIC: 30373.26 AICC: 30380.47 BIC: 31216.64 
# Log posterior: -145184.1 
# Prior parameters: mu=0 sigma=1000 nu=4 

#yes- lumping U and ID performs better in this case- so the "global" rate here actually represents them, whereas now we have a single partitioned rate for "X" males 

########### initCH ############### 


####
#I have a feeling that initCH will need to be fit to individuals (at least ID'd indivs)- I think individuals really do vary in their propensity to get chased once attacked. 

##see if this is bourne out in model selection


### Global initCH rate 


sequences<-table(event.list$Date)

#create list of dummy intercept names

idno<-length(event.types[grep("cop.start",event.types)])

dummy.ints.attack<-paste(1:14,".attackFE", sep="")
dummy.ints.receive<-paste(1:13, ".receiveFE", sep="")


int.names<-c("Global.attackFE", "X.receiveFE", "X.UreceiveFE", "U.attackFE", "U.receiveFE", "U.UattackFE", dummy.ints.attack, dummy.ints.receive, "cop.startFE", "cop.endFE", "initFO.FE","ID.initFO.FE", "U.initFO.FE","initSF.FE","X.initSF.FE", "initCH.FE", event.types[-c(grep("attack", event.types),grep("cop", event.types),grep("initFO", event.types),grep("initSF", event.types),grep("initCH", event.types))])

int.list<-list()

for(i in 1:length(levels(event.list$Date))){

day.array<-array(0,dim=c(as.numeric(sequences[i]), length(event.types),length(int.names)))

#the third dimension is the stat slice.
dimnames(day.array)<-list(c(1:sequences[i]),as.character(event.types),as.character(int.names))

#add global attack effect, loops for "U" intercepts for attacks and receives, then one loop for typical intercepts

day.array[,grep("attack", event.types),1]<-1


for(h in 1:14){
day.array[,paste(h,".attack.",30, sep=""),2]<-1
}


for(h in 15:29){
day.array[,paste(h,".attack.",30, sep=""),3]<-1
}


for(h in 1:14){
	for(j in 15:29){	
day.array[,paste(j,".attack.", h, sep=""),4]<-1
}
}


for(h in 1:14){
	for(j in 15:29){
day.array[,paste(h,".attack.",j, sep=""),5]<-1
}
}

for(h in 15:29){
	for(j in 15:29){	
if(h!=j){
day.array[,paste(j,".attack.", h, sep=""),6]<-1
}
}
}

preloop<-6

#loop for dummy intercepts for attacks, loop for dummy intercepts for receives, add in cop.start and cop.end FEs, then one loop for typical intercepts

for(k in (preloop+1):(length(dummy.ints.attack)+preloop)){
	
	for(h in 1:14){
if(h!=(k-preloop)){
day.array[,paste((k-preloop),".attack.", h, sep=""),k]<-1
}
}
}

for(k in ((length(dummy.ints.attack)+preloop)+1):((length(dummy.ints.attack)+preloop)+length(dummy.ints.receive))){
	for(h in 1:14){
if(h!=(k-(length(dummy.ints.attack)+preloop))){
day.array[,paste(h,".attack.",(k-(length(dummy.ints.attack)+preloop)), sep=""),k]<-1
}
}
}

#cop.startFE
for(h in 1:idno){
day.array[,paste(h,".cop.start", sep=""),((length(dummy.ints.attack)+preloop)+length(dummy.ints.receive)+1)]<-1
}


#cop.endFE
for(h in 1:idno){
day.array[,paste(h,".cop.end", sep=""),((length(dummy.ints.attack)+preloop)+length(dummy.ints.receive)+2)]<-1
}

#initFO.FE
for(h in 1:idno){
day.array[,paste(h,".initFO", sep=""),((length(dummy.ints.attack)+preloop)+length(dummy.ints.receive)+3)]<-1
}

#ID.initFO.FE
for(h in 1:14){
day.array[,paste(h,".initFO", sep=""),((length(dummy.ints.attack)+preloop)+length(dummy.ints.receive)+4)]<-1
}

#U.initFO.FE
for(h in 15:29){
day.array[,paste(h,".initFO", sep=""),((length(dummy.ints.attack)+preloop)+length(dummy.ints.receive)+5)]<-1
}

#initSF.FE
for(h in 1:idno){
day.array[,paste(h,".initSF", sep=""),((length(dummy.ints.attack)+preloop)+length(dummy.ints.receive)+6)]<-1
}

#X.initSF.FE

day.array[,paste(30,".initSF", sep=""),((length(dummy.ints.attack)+preloop)+length(dummy.ints.receive)+7)]<-1

#initCH.FE
for(h in 1:idno){
day.array[,paste(h,".initCH", sep=""),((length(dummy.ints.attack)+preloop)+length(dummy.ints.receive)+8)]<-1
}

#everything else
for(k in (((length(dummy.ints.attack)+preloop)+length(dummy.ints.receive)+9):length(int.names))){
	
	day.array[,int.names[k],k]<-1
	
}

#weird extra formatting to make it match the package example, seems like it becomes relevant *punintended!* if I want to fit day-specific effects

global<-list()

global[[1]]<-day.array

int.list[[i]]<-global

names(int.list[[i]])<-"global"

}

tic()
intfit.sup<-rem(evlist, int.list, supplist=supplist, timing="interval", estimator="BPM")
toc()

summary(intfit.sup)

# Egocentric Relational Event Model (Interval Likelihood)
# 
#                   Post.Mode     Post.SD  Z value  Pr(>|z|)    
# Global.attackFE  -9.5259220   0.2886747 -32.9988 < 2.2e-16 ***
# X.receiveFE       3.7712555   0.2921739  12.9076 < 2.2e-16 ***
# X.UreceiveFE      3.9025235   0.3370996  11.5768 < 2.2e-16 ***
# U.attackFE       -1.2734082   0.6454967  -1.9728 0.0485233 *  
# U.receiveFE       1.7877281   0.3145761   5.6830 1.324e-08 ***
# U.UattackFE       2.9176286   0.3248928   8.9803 < 2.2e-16 ***
# 1.attackFE        1.0330374   0.3944376   2.6190 0.0088184 ** 
# 2.attackFE        1.2763648   0.3921908   3.2544 0.0011361 ** 
# 3.attackFE        1.6061436   0.3891684   4.1271 3.673e-05 ***
# 4.attackFE        2.5265882   0.3860257   6.5451 5.944e-11 ***
# 5.attackFE        0.3110660   0.4195142   0.7415 0.4583958    
# 6.attackFE       -0.0207165   0.8047343  -0.0257 0.9794621    
# 7.attackFE        1.3366638   0.3932445   3.3991 0.0006762 ***
# 8.attackFE       -1.4929214   1.0698668  -1.3954 0.1628870    
# 9.attackFE      -12.5412493 243.0584661  -0.0516 0.9588493    
# 10.attackFE       1.5764042   0.4001704   3.9393 8.171e-05 ***
# 11.attackFE     -10.4618029 264.1957859  -0.0396 0.9684131    
# 12.attackFE      -0.0121458   0.5502168  -0.0221 0.9823884    
# 13.attackFE       0.9853197   1.0715358   0.9195 0.3578134    
# 14.attackFE       0.7284843   0.5224387   1.3944 0.1631993    
# 1.receiveFE      -0.7315755   0.2770237  -2.6408 0.0082700 ** 
# 2.receiveFE      -0.2483185   0.2681292  -0.9261 0.3543862    
# 3.receiveFE      -0.3750730   0.2725416  -1.3762 0.1687582    
# 4.receiveFE       1.0326554   0.2610273   3.9561 7.618e-05 ***
# 5.receiveFE      -0.8724350   0.2925982  -2.9817 0.0028667 ** 
# 6.receiveFE       0.7239207   0.3403191   2.1272 0.0334050 *  
# 7.receiveFE       0.2399268   0.2626681   0.9134 0.3610207    
# 8.receiveFE       0.0322648   0.3395839   0.0950 0.9243046    
# 9.receiveFE       0.6527401   0.3048317   2.1413 0.0322488 *  
# 10.receiveFE     -0.0083194   0.2918528  -0.0285 0.9772590    
# 11.receiveFE      1.6776185   0.4538049   3.6968 0.0002183 ***
# 12.receiveFE     -0.4512307   0.3601768  -1.2528 0.2102774    
# 13.receiveFE     -0.3517487   1.0308039  -0.3412 0.7329249    
# cop.startFE      -8.8434147   0.1178511 -75.0389 < 2.2e-16 ***
# cop.endFE        -1.4478174   0.1414214 -10.2376 < 2.2e-16 ***
# initFO.FE         3.3524074   0.2581988  12.9838 < 2.2e-16 ***
# ID.initFO.FE      3.2420533   0.2612259  12.4109 < 2.2e-16 ***
# U.initFO.FE       1.8903400   0.3380617   5.5917 2.249e-08 ***
# initSF.FE         5.1608463   0.0764719  67.4868 < 2.2e-16 ***
# X.initSF.FE      -1.2206524   0.2070869  -5.8944 3.761e-09 ***
# initCH.FE         6.0457414   0.0396526 152.4678 < 2.2e-16 ***
# 1.disengage      -4.2622836   0.0626224 -68.0632 < 2.2e-16 ***
# 2.disengage      -5.1487697   0.0887356 -58.0237 < 2.2e-16 ***
# 3.disengage      -4.7750140   0.0636285 -75.0452 < 2.2e-16 ***
# 4.disengage      -4.6474529   0.0548821 -84.6806 < 2.2e-16 ***
# 5.disengage      -4.5737155   0.1025978 -44.5791 < 2.2e-16 ***
# 6.disengage      -6.4777863   0.5773494 -11.2199 < 2.2e-16 ***
# 7.disengage      -5.6831776   0.0894427 -63.5399 < 2.2e-16 ***
# 8.disengage      -3.0256324   0.1889822 -16.0101 < 2.2e-16 ***
# 9.disengage      -2.9513586   0.9999975  -2.9514 0.0031637 ** 
# 10.disengage     -4.6109312   0.0880451 -52.3701 < 2.2e-16 ***
# 11.disengage     -1.2519039   0.9999986  -1.2519 0.2106042    
# 12.disengage     -5.0999545   0.3333332 -15.2999 < 2.2e-16 ***
# 13.disengage     -3.0694899   0.9999975  -3.0695 0.0021442 ** 
# 14.disengage     -3.9995581   0.2000000 -19.9978 < 2.2e-16 ***
# 15.disengage     -3.7115624   0.3779643  -9.8199 < 2.2e-16 ***
# 16.disengage      0.0000000 894.4271910   0.0000 1.0000000    
# 17.disengage     -5.5854796   0.7071053  -7.8991 2.887e-15 ***
# 18.disengage      0.0000000 894.4271910   0.0000 1.0000000    
# 19.disengage     -3.3061330   0.2500000 -13.2245 < 2.2e-16 ***
# 20.disengage     -2.4505089   0.3535533  -6.9311 4.176e-12 ***
# 21.disengage      0.0000000 894.4271910   0.0000 1.0000000    
# 22.disengage     -3.5143729   0.9999972  -3.5144 0.0004408 ***
# 23.disengage    -12.1897538 246.2838524  -0.0495 0.9605250    
# 24.disengage     -1.8919041   0.9999982  -1.8919 0.0585033 .  
# 25.disengage     -2.4340658   0.2000000 -12.1703 < 2.2e-16 ***
# 26.disengage     -3.8049157   0.4082481  -9.3201 < 2.2e-16 ***
# 27.disengage     -5.3682937   0.3162276 -16.9760 < 2.2e-16 ***
# 28.disengage     -5.0921384   0.3535532 -14.4027 < 2.2e-16 ***
# 29.disengage     -5.0977870   0.4082480 -12.4870 < 2.2e-16 ***
# 30.disengage     -4.8310314   0.1622214 -29.7805 < 2.2e-16 ***
# 1.depart         -8.7014746   0.2357022 -36.9172 < 2.2e-16 ***
# 2.depart         -8.6455924   0.2357022 -36.6802 < 2.2e-16 ***
# 3.depart         -8.6031331   0.2357022 -36.5000 < 2.2e-16 ***
# 4.depart         -8.2131081   0.2357022 -34.8453 < 2.2e-16 ***
# 5.depart         -8.7978526   0.2886750 -30.4767 < 2.2e-16 ***
# 6.depart         -8.1073118   0.7071048 -11.4655 < 2.2e-16 ***
# 7.depart         -8.4665832   0.2357022 -35.9207 < 2.2e-16 ***
# 8.depart         -7.6244610   0.4999993 -15.2489 < 2.2e-16 ***
# 9.depart         -8.2131177   0.5773492 -14.2256 < 2.2e-16 ***
# 10.depart        -8.3366519   0.3779642 -22.0567 < 2.2e-16 ***
# 11.depart        -6.1361062   0.5773494 -10.6281 < 2.2e-16 ***
# 12.depart        -8.4305661   0.5773491 -14.6022 < 2.2e-16 ***
# 13.depart        -6.3742619   0.9999954  -6.3743 1.838e-10 ***
# 14.depart        -8.4328617   0.7071047 -11.9259 < 2.2e-16 ***
# 15.depart        -8.4724875   0.9999941  -8.4725 < 2.2e-16 ***
# 16.depart        -4.5531868   0.9999965  -4.5532 5.284e-06 ***
# 17.depart        -7.3325942   0.9999948  -7.3326 2.256e-13 ***
# 18.depart        -6.1293192   0.9999955  -6.1293 8.824e-10 ***
# 19.depart        -8.8972145   0.9999938  -8.8973 < 2.2e-16 ***
# 20.depart        -7.6761506   0.9999946  -7.6762 1.643e-14 ***
# 21.depart        -4.7116752   0.9999964  -4.7117 2.457e-06 ***
# 22.depart        -7.7353839   0.9999945  -7.7354 1.021e-14 ***
# 23.depart        -7.0172213   0.9999950  -7.0173 2.263e-12 ***
# 24.depart        -7.3915096   0.9999948  -7.3915 1.452e-13 ***
# 25.depart        -7.8421735   0.9999945  -7.8422 4.441e-15 ***
# 26.depart        -8.0356182   0.9999944  -8.0357 8.882e-16 ***
# 27.depart        -8.6596852   0.9999940  -8.6597 < 2.2e-16 ***
# 28.depart        -8.1918068   0.9999943  -8.1919 2.220e-16 ***
# 29.depart        -7.0548164   0.9999950  -7.0549 1.728e-12 ***
# 30.depart        -5.7379132   0.1178511 -48.6878 < 2.2e-16 ***
# ---
# Signif. codes:  0 ‘***’ 0.001 ‘**’ 0.01 ‘*’ 0.05 ‘.’ 0.1 ‘ ’ 1
# Null deviance: 76628.21 on 4854 degrees of freedom
# Residual deviance: 31003.1 on 4753 degrees of freedom
# 	Chi-square: 45625.11 on 101 degrees of freedom, asymptotic p-value 0 
# AIC: 31205.1 AICC: 31209.44 BIC: 31860.34 
# Log posterior: -116600.6 
# Prior parameters: mu=0 sigma=1000 nu=4 


#yep, this model is much worse


#### Partition out ID males 


sequences<-table(event.list$Date)

#create list of dummy intercept names

idno<-length(event.types[grep("cop.start",event.types)])

dummy.ints.attack<-paste(1:14,".attackFE", sep="")
dummy.ints.receive<-paste(1:13, ".receiveFE", sep="")


int.names<-c("Global.attackFE", "X.receiveFE", "X.UreceiveFE", "U.attackFE", "U.receiveFE", "U.UattackFE", dummy.ints.attack, dummy.ints.receive, "cop.startFE", "cop.endFE", "initFO.FE","ID.initFO.FE", "U.initFO.FE","initSF.FE","X.initSF.FE", "initCH.FE", "ID.initCH.FE", event.types[-c(grep("attack", event.types),grep("cop", event.types),grep("initFO", event.types),grep("initSF", event.types),grep("initCH", event.types))])

int.list<-list()

for(i in 1:length(levels(event.list$Date))){

day.array<-array(0,dim=c(as.numeric(sequences[i]), length(event.types),length(int.names)))

#the third dimension is the stat slice.
dimnames(day.array)<-list(c(1:sequences[i]),as.character(event.types),as.character(int.names))

#add global attack effect, loops for "U" intercepts for attacks and receives, then one loop for typical intercepts

day.array[,grep("attack", event.types),1]<-1


for(h in 1:14){
day.array[,paste(h,".attack.",30, sep=""),2]<-1
}


for(h in 15:29){
day.array[,paste(h,".attack.",30, sep=""),3]<-1
}


for(h in 1:14){
	for(j in 15:29){	
day.array[,paste(j,".attack.", h, sep=""),4]<-1
}
}


for(h in 1:14){
	for(j in 15:29){
day.array[,paste(h,".attack.",j, sep=""),5]<-1
}
}

for(h in 15:29){
	for(j in 15:29){	
if(h!=j){
day.array[,paste(j,".attack.", h, sep=""),6]<-1
}
}
}

preloop<-6

#loop for dummy intercepts for attacks, loop for dummy intercepts for receives, add in cop.start and cop.end FEs, then one loop for typical intercepts

for(k in (preloop+1):(length(dummy.ints.attack)+preloop)){
	
	for(h in 1:14){
if(h!=(k-preloop)){
day.array[,paste((k-preloop),".attack.", h, sep=""),k]<-1
}
}
}

for(k in ((length(dummy.ints.attack)+preloop)+1):((length(dummy.ints.attack)+preloop)+length(dummy.ints.receive))){
	for(h in 1:14){
if(h!=(k-(length(dummy.ints.attack)+preloop))){
day.array[,paste(h,".attack.",(k-(length(dummy.ints.attack)+preloop)), sep=""),k]<-1
}
}
}

#cop.startFE
for(h in 1:idno){
day.array[,paste(h,".cop.start", sep=""),((length(dummy.ints.attack)+preloop)+length(dummy.ints.receive)+1)]<-1
}


#cop.endFE
for(h in 1:idno){
day.array[,paste(h,".cop.end", sep=""),((length(dummy.ints.attack)+preloop)+length(dummy.ints.receive)+2)]<-1
}

#initFO.FE
for(h in 1:idno){
day.array[,paste(h,".initFO", sep=""),((length(dummy.ints.attack)+preloop)+length(dummy.ints.receive)+3)]<-1
}

#ID.initFO.FE
for(h in 1:14){
day.array[,paste(h,".initFO", sep=""),((length(dummy.ints.attack)+preloop)+length(dummy.ints.receive)+4)]<-1
}

#U.initFO.FE
for(h in 15:29){
day.array[,paste(h,".initFO", sep=""),((length(dummy.ints.attack)+preloop)+length(dummy.ints.receive)+5)]<-1
}

#initSF.FE
for(h in 1:idno){
day.array[,paste(h,".initSF", sep=""),((length(dummy.ints.attack)+preloop)+length(dummy.ints.receive)+6)]<-1
}

#X.initSF.FE

day.array[,paste(30,".initSF", sep=""),((length(dummy.ints.attack)+preloop)+length(dummy.ints.receive)+7)]<-1

#initCH.FE
for(h in 1:idno){
day.array[,paste(h,".initCH", sep=""),((length(dummy.ints.attack)+preloop)+length(dummy.ints.receive)+8)]<-1
}

#ID.initCH.FE
for(h in 1:14){
day.array[,paste(h,".initCH", sep=""),((length(dummy.ints.attack)+preloop)+length(dummy.ints.receive)+9)]<-1
}


#everything else
for(k in (((length(dummy.ints.attack)+preloop)+length(dummy.ints.receive)+10):length(int.names))){
	
	day.array[,int.names[k],k]<-1
	
}

#weird extra formatting to make it match the package example, seems like it becomes relevant *punintended!* if I want to fit day-specific effects

global<-list()

global[[1]]<-day.array

int.list[[i]]<-global

names(int.list[[i]])<-"global"

}


tic()
intfit.sup<-rem(evlist, int.list, supplist=supplist, timing="interval", estimator="BPM")
toc()

summary(intfit.sup)


# Egocentric Relational Event Model (Interval Likelihood)
# 
#                   Post.Mode     Post.SD  Z value  Pr(>|z|)    
# Global.attackFE  -9.5259220   0.2886747 -32.9988 < 2.2e-16 ***
# X.receiveFE       3.7712555   0.2921739  12.9076 < 2.2e-16 ***
# X.UreceiveFE      3.9025235   0.3370996  11.5768 < 2.2e-16 ***
# U.attackFE       -1.2734082   0.6454967  -1.9728 0.0485233 *  
# U.receiveFE       1.7877281   0.3145761   5.6830 1.324e-08 ***
# U.UattackFE       2.9176286   0.3248928   8.9803 < 2.2e-16 ***
# 1.attackFE        1.0330374   0.3944376   2.6190 0.0088184 ** 
# 2.attackFE        1.2763648   0.3921908   3.2544 0.0011361 ** 
# 3.attackFE        1.6061436   0.3891684   4.1271 3.673e-05 ***
# 4.attackFE        2.5265882   0.3860257   6.5451 5.944e-11 ***
# 5.attackFE        0.3110660   0.4195142   0.7415 0.4583958    
# 6.attackFE       -0.0207165   0.8047343  -0.0257 0.9794621    
# 7.attackFE        1.3366638   0.3932445   3.3991 0.0006762 ***
# 8.attackFE       -1.4929214   1.0698668  -1.3954 0.1628870    
# 9.attackFE      -12.5412493 243.0584695  -0.0516 0.9588493    
# 10.attackFE       1.5764042   0.4001704   3.9393 8.171e-05 ***
# 11.attackFE     -10.4618029 264.1957859  -0.0396 0.9684131    
# 12.attackFE      -0.0121458   0.5502168  -0.0221 0.9823884    
# 13.attackFE       0.9853197   1.0715358   0.9195 0.3578134    
# 14.attackFE       0.7284843   0.5224387   1.3944 0.1631993    
# 1.receiveFE      -0.7315755   0.2770237  -2.6408 0.0082700 ** 
# 2.receiveFE      -0.2483185   0.2681292  -0.9261 0.3543862    
# 3.receiveFE      -0.3750730   0.2725416  -1.3762 0.1687582    
# 4.receiveFE       1.0326554   0.2610273   3.9561 7.618e-05 ***
# 5.receiveFE      -0.8724350   0.2925982  -2.9817 0.0028667 ** 
# 6.receiveFE       0.7239207   0.3403191   2.1272 0.0334050 *  
# 7.receiveFE       0.2399268   0.2626681   0.9134 0.3610207    
# 8.receiveFE       0.0322648   0.3395839   0.0950 0.9243046    
# 9.receiveFE       0.6527401   0.3048317   2.1413 0.0322488 *  
# 10.receiveFE     -0.0083194   0.2918528  -0.0285 0.9772590    
# 11.receiveFE      1.6776185   0.4538049   3.6968 0.0002183 ***
# 12.receiveFE     -0.4512307   0.3601768  -1.2528 0.2102774    
# 13.receiveFE     -0.3517487   1.0308039  -0.3412 0.7329249    
# cop.startFE      -8.8434147   0.1178511 -75.0389 < 2.2e-16 ***
# cop.endFE        -1.4478174   0.1414214 -10.2376 < 2.2e-16 ***
# initFO.FE         3.3524074   0.2581988  12.9838 < 2.2e-16 ***
# ID.initFO.FE      3.2420533   0.2612259  12.4109 < 2.2e-16 ***
# U.initFO.FE       1.8903400   0.3380617   5.5917 2.249e-08 ***
# initSF.FE         5.1608463   0.0764719  67.4868 < 2.2e-16 ***
# X.initSF.FE      -1.2206524   0.2070869  -5.8944 3.761e-09 ***
# initCH.FE         6.7588320   0.0427179 158.2202 < 2.2e-16 ***
# ID.initCH.FE     -2.1422331   0.1148410 -18.6539 < 2.2e-16 ***
# 1.disengage      -4.2622836   0.0626224 -68.0632 < 2.2e-16 ***
# 2.disengage      -5.1487697   0.0887356 -58.0237 < 2.2e-16 ***
# 3.disengage      -4.7750140   0.0636285 -75.0452 < 2.2e-16 ***
# 4.disengage      -4.6474529   0.0548821 -84.6806 < 2.2e-16 ***
# 5.disengage      -4.5737155   0.1025978 -44.5791 < 2.2e-16 ***
# 6.disengage      -6.4777863   0.5773494 -11.2199 < 2.2e-16 ***
# 7.disengage      -5.6831776   0.0894427 -63.5399 < 2.2e-16 ***
# 8.disengage      -3.0256324   0.1889822 -16.0101 < 2.2e-16 ***
# 9.disengage      -2.9513586   0.9999975  -2.9514 0.0031637 ** 
# 10.disengage     -4.6109312   0.0880451 -52.3701 < 2.2e-16 ***
# 11.disengage     -1.2519039   0.9999986  -1.2519 0.2106042    
# 12.disengage     -5.0999545   0.3333332 -15.2999 < 2.2e-16 ***
# 13.disengage     -3.0694899   0.9999975  -3.0695 0.0021442 ** 
# 14.disengage     -3.9995581   0.2000000 -19.9978 < 2.2e-16 ***
# 15.disengage     -3.7115624   0.3779643  -9.8199 < 2.2e-16 ***
# 16.disengage      0.0000000 894.4271910   0.0000 1.0000000    
# 17.disengage     -5.5854796   0.7071053  -7.8991 2.887e-15 ***
# 18.disengage      0.0000000 894.4271910   0.0000 1.0000000    
# 19.disengage     -3.3061330   0.2500000 -13.2245 < 2.2e-16 ***
# 20.disengage     -2.4505089   0.3535533  -6.9311 4.176e-12 ***
# 21.disengage      0.0000000 894.4271910   0.0000 1.0000000    
# 22.disengage     -3.5143729   0.9999972  -3.5144 0.0004408 ***
# 23.disengage    -12.1897538 246.2838524  -0.0495 0.9605250    
# 24.disengage     -1.8919041   0.9999982  -1.8919 0.0585033 .  
# 25.disengage     -2.4340658   0.2000000 -12.1703 < 2.2e-16 ***
# 26.disengage     -3.8049157   0.4082481  -9.3201 < 2.2e-16 ***
# 27.disengage     -5.3682937   0.3162276 -16.9760 < 2.2e-16 ***
# 28.disengage     -5.0921384   0.3535532 -14.4027 < 2.2e-16 ***
# 29.disengage     -5.0977870   0.4082480 -12.4870 < 2.2e-16 ***
# 30.disengage     -4.8310314   0.1622214 -29.7805 < 2.2e-16 ***
# 1.depart         -8.7014746   0.2357022 -36.9172 < 2.2e-16 ***
# 2.depart         -8.6455924   0.2357022 -36.6802 < 2.2e-16 ***
# 3.depart         -8.6031331   0.2357022 -36.5000 < 2.2e-16 ***
# 4.depart         -8.2131081   0.2357022 -34.8453 < 2.2e-16 ***
# 5.depart         -8.7978526   0.2886750 -30.4767 < 2.2e-16 ***
# 6.depart         -8.1073118   0.7071048 -11.4655 < 2.2e-16 ***
# 7.depart         -8.4665832   0.2357022 -35.9207 < 2.2e-16 ***
# 8.depart         -7.6244610   0.4999993 -15.2489 < 2.2e-16 ***
# 9.depart         -8.2131177   0.5773492 -14.2256 < 2.2e-16 ***
# 10.depart        -8.3366519   0.3779642 -22.0567 < 2.2e-16 ***
# 11.depart        -6.1361062   0.5773494 -10.6281 < 2.2e-16 ***
# 12.depart        -8.4305661   0.5773491 -14.6022 < 2.2e-16 ***
# 13.depart        -6.3742619   0.9999954  -6.3743 1.838e-10 ***
# 14.depart        -8.4328617   0.7071047 -11.9259 < 2.2e-16 ***
# 15.depart        -8.4724875   0.9999941  -8.4725 < 2.2e-16 ***
# 16.depart        -4.5531868   0.9999965  -4.5532 5.284e-06 ***
# 17.depart        -7.3325942   0.9999948  -7.3326 2.256e-13 ***
# 18.depart        -6.1293192   0.9999955  -6.1293 8.824e-10 ***
# 19.depart        -8.8972145   0.9999938  -8.8973 < 2.2e-16 ***
# 20.depart        -7.6761506   0.9999946  -7.6762 1.643e-14 ***
# 21.depart        -4.7116752   0.9999964  -4.7117 2.457e-06 ***
# 22.depart        -7.7353839   0.9999945  -7.7354 1.021e-14 ***
# 23.depart        -7.0172213   0.9999950  -7.0173 2.263e-12 ***
# 24.depart        -7.3915096   0.9999948  -7.3915 1.452e-13 ***
# 25.depart        -7.8421735   0.9999945  -7.8422 4.441e-15 ***
# 26.depart        -8.0356182   0.9999944  -8.0357 8.882e-16 ***
# 27.depart        -8.6596852   0.9999940  -8.6597 < 2.2e-16 ***
# 28.depart        -8.1918068   0.9999943  -8.1919 2.220e-16 ***
# 29.depart        -7.0548164   0.9999950  -7.0549 1.728e-12 ***
# 30.depart        -5.7379132   0.1178511 -48.6878 < 2.2e-16 ***
# ---
# Signif. codes:  0 ‘***’ 0.001 ‘**’ 0.01 ‘*’ 0.05 ‘.’ 0.1 ‘ ’ 1
# Null deviance: 76628.21 on 4854 degrees of freedom
# Residual deviance: 30473.08 on 4752 degrees of freedom
# 	Chi-square: 46155.13 on 102 degrees of freedom, asymptotic p-value 0 
# AIC: 30677.08 AICC: 30681.51 BIC: 31338.81 
# Log posterior: -117336.6 
# Prior parameters: mu=0 sigma=1000 nu=4 


#better- obviously Id males initiate chases way less than U or ?? males 


## Partition out ID AND U males


sequences<-table(event.list$Date)

#create list of dummy intercept names

idno<-length(event.types[grep("cop.start",event.types)])

dummy.ints.attack<-paste(1:14,".attackFE", sep="")
dummy.ints.receive<-paste(1:13, ".receiveFE", sep="")


int.names<-c("Global.attackFE", "X.receiveFE", "X.UreceiveFE", "U.attackFE", "U.receiveFE", "U.UattackFE", dummy.ints.attack, dummy.ints.receive, "cop.startFE", "cop.endFE", "initFO.FE","ID.initFO.FE", "U.initFO.FE","initSF.FE","X.initSF.FE", "initCH.FE", "ID.initCH.FE", "U.initCH.FE", event.types[-c(grep("attack", event.types),grep("cop", event.types),grep("initFO", event.types),grep("initSF", event.types),grep("initCH", event.types))])

int.list<-list()

for(i in 1:length(levels(event.list$Date))){

day.array<-array(0,dim=c(as.numeric(sequences[i]), length(event.types),length(int.names)))

#the third dimension is the stat slice.
dimnames(day.array)<-list(c(1:sequences[i]),as.character(event.types),as.character(int.names))

#add global attack effect, loops for "U" intercepts for attacks and receives, then one loop for typical intercepts

day.array[,grep("attack", event.types),1]<-1


for(h in 1:14){
day.array[,paste(h,".attack.",30, sep=""),2]<-1
}


for(h in 15:29){
day.array[,paste(h,".attack.",30, sep=""),3]<-1
}


for(h in 1:14){
	for(j in 15:29){	
day.array[,paste(j,".attack.", h, sep=""),4]<-1
}
}


for(h in 1:14){
	for(j in 15:29){
day.array[,paste(h,".attack.",j, sep=""),5]<-1
}
}

for(h in 15:29){
	for(j in 15:29){	
if(h!=j){
day.array[,paste(j,".attack.", h, sep=""),6]<-1
}
}
}

preloop<-6

#loop for dummy intercepts for attacks, loop for dummy intercepts for receives, add in cop.start and cop.end FEs, then one loop for typical intercepts

for(k in (preloop+1):(length(dummy.ints.attack)+preloop)){
	
	for(h in 1:14){
if(h!=(k-preloop)){
day.array[,paste((k-preloop),".attack.", h, sep=""),k]<-1
}
}
}

for(k in ((length(dummy.ints.attack)+preloop)+1):((length(dummy.ints.attack)+preloop)+length(dummy.ints.receive))){
	for(h in 1:14){
if(h!=(k-(length(dummy.ints.attack)+preloop))){
day.array[,paste(h,".attack.",(k-(length(dummy.ints.attack)+preloop)), sep=""),k]<-1
}
}
}

#cop.startFE
for(h in 1:idno){
day.array[,paste(h,".cop.start", sep=""),((length(dummy.ints.attack)+preloop)+length(dummy.ints.receive)+1)]<-1
}


#cop.endFE
for(h in 1:idno){
day.array[,paste(h,".cop.end", sep=""),((length(dummy.ints.attack)+preloop)+length(dummy.ints.receive)+2)]<-1
}

#initFO.FE
for(h in 1:idno){
day.array[,paste(h,".initFO", sep=""),((length(dummy.ints.attack)+preloop)+length(dummy.ints.receive)+3)]<-1
}

#ID.initFO.FE
for(h in 1:14){
day.array[,paste(h,".initFO", sep=""),((length(dummy.ints.attack)+preloop)+length(dummy.ints.receive)+4)]<-1
}

#U.initFO.FE
for(h in 15:29){
day.array[,paste(h,".initFO", sep=""),((length(dummy.ints.attack)+preloop)+length(dummy.ints.receive)+5)]<-1
}

#initSF.FE
for(h in 1:idno){
day.array[,paste(h,".initSF", sep=""),((length(dummy.ints.attack)+preloop)+length(dummy.ints.receive)+6)]<-1
}

#X.initSF.FE

day.array[,paste(30,".initSF", sep=""),((length(dummy.ints.attack)+preloop)+length(dummy.ints.receive)+7)]<-1

#initCH.FE
for(h in 1:idno){
day.array[,paste(h,".initCH", sep=""),((length(dummy.ints.attack)+preloop)+length(dummy.ints.receive)+8)]<-1
}

#ID.initCH.FE
for(h in 1:14){
day.array[,paste(h,".initCH", sep=""),((length(dummy.ints.attack)+preloop)+length(dummy.ints.receive)+9)]<-1
}

#U.initCH.FE
for(h in 15:29){
day.array[,paste(h,".initCH", sep=""),((length(dummy.ints.attack)+preloop)+length(dummy.ints.receive)+10)]<-1
}

#everything else
for(k in (((length(dummy.ints.attack)+preloop)+length(dummy.ints.receive)+11):length(int.names))){
	
	day.array[,int.names[k],k]<-1
	
}

#weird extra formatting to make it match the package example, seems like it becomes relevant *punintended!* if I want to fit day-specific effects

global<-list()

global[[1]]<-day.array

int.list[[i]]<-global

names(int.list[[i]])<-"global"

}

tic()
intfit.sup<-rem(evlist, int.list, supplist=supplist, timing="interval", estimator="BPM")
toc()

summary(intfit.sup) 


# Egocentric Relational Event Model (Interval Likelihood)
# 
#                   Post.Mode     Post.SD  Z value  Pr(>|z|)    
# Global.attackFE  -9.5259220   0.2886747 -32.9988 < 2.2e-16 ***
# X.receiveFE       3.7712555   0.2921739  12.9076 < 2.2e-16 ***
# X.UreceiveFE      3.9025235   0.3370996  11.5768 < 2.2e-16 ***
# U.attackFE       -1.2734082   0.6454967  -1.9728 0.0485233 *  
# U.receiveFE       1.7877281   0.3145761   5.6830 1.324e-08 ***
# U.UattackFE       2.9176286   0.3248928   8.9803 < 2.2e-16 ***
# 1.attackFE        1.0330374   0.3944376   2.6190 0.0088184 ** 
# 2.attackFE        1.2763648   0.3921908   3.2544 0.0011361 ** 
# 3.attackFE        1.6061436   0.3891684   4.1271 3.673e-05 ***
# 4.attackFE        2.5265882   0.3860257   6.5451 5.944e-11 ***
# 5.attackFE        0.3110660   0.4195142   0.7415 0.4583958    
# 6.attackFE       -0.0207165   0.8047343  -0.0257 0.9794621    
# 7.attackFE        1.3366638   0.3932445   3.3991 0.0006762 ***
# 8.attackFE       -1.4929214   1.0698668  -1.3954 0.1628870    
# 9.attackFE      -12.5412493 243.0584731  -0.0516 0.9588493    
# 10.attackFE       1.5764042   0.4001704   3.9393 8.171e-05 ***
# 11.attackFE     -10.4618029 264.1957859  -0.0396 0.9684131    
# 12.attackFE      -0.0121458   0.5502168  -0.0221 0.9823884    
# 13.attackFE       0.9853197   1.0715358   0.9195 0.3578134    
# 14.attackFE       0.7284843   0.5224387   1.3944 0.1631993    
# 1.receiveFE      -0.7315755   0.2770237  -2.6408 0.0082700 ** 
# 2.receiveFE      -0.2483185   0.2681292  -0.9261 0.3543862    
# 3.receiveFE      -0.3750730   0.2725416  -1.3762 0.1687582    
# 4.receiveFE       1.0326554   0.2610273   3.9561 7.618e-05 ***
# 5.receiveFE      -0.8724350   0.2925982  -2.9817 0.0028667 ** 
# 6.receiveFE       0.7239207   0.3403191   2.1272 0.0334050 *  
# 7.receiveFE       0.2399268   0.2626681   0.9134 0.3610207    
# 8.receiveFE       0.0322648   0.3395839   0.0950 0.9243046    
# 9.receiveFE       0.6527401   0.3048317   2.1413 0.0322488 *  
# 10.receiveFE     -0.0083194   0.2918528  -0.0285 0.9772590    
# 11.receiveFE      1.6776185   0.4538049   3.6968 0.0002183 ***
# 12.receiveFE     -0.4512307   0.3601768  -1.2528 0.2102774    
# 13.receiveFE     -0.3517487   1.0308039  -0.3412 0.7329249    
# cop.startFE      -8.8434147   0.1178511 -75.0389 < 2.2e-16 ***
# cop.endFE        -1.4478174   0.1414214 -10.2376 < 2.2e-16 ***
# initFO.FE         3.3524074   0.2581988  12.9838 < 2.2e-16 ***
# ID.initFO.FE      3.2420533   0.2612259  12.4109 < 2.2e-16 ***
# U.initFO.FE       1.8903400   0.3380617   5.5917 2.249e-08 ***
# initSF.FE         5.1608463   0.0764719  67.4868 < 2.2e-16 ***
# X.initSF.FE      -1.2206524   0.2070869  -5.8944 3.761e-09 ***
# initCH.FE         6.8243736   0.0455016 149.9810 < 2.2e-16 ***
# ID.initCH.FE     -2.2077747   0.1159053 -19.0481 < 2.2e-16 ***
# U.initCH.FE      -0.4517613   0.1321174  -3.4194 0.0006276 ***
# 1.disengage      -4.2622836   0.0626224 -68.0632 < 2.2e-16 ***
# 2.disengage      -5.1487697   0.0887356 -58.0237 < 2.2e-16 ***
# 3.disengage      -4.7750140   0.0636285 -75.0452 < 2.2e-16 ***
# 4.disengage      -4.6474529   0.0548821 -84.6806 < 2.2e-16 ***
# 5.disengage      -4.5737155   0.1025978 -44.5791 < 2.2e-16 ***
# 6.disengage      -6.4777863   0.5773494 -11.2199 < 2.2e-16 ***
# 7.disengage      -5.6831776   0.0894427 -63.5399 < 2.2e-16 ***
# 8.disengage      -3.0256324   0.1889822 -16.0101 < 2.2e-16 ***
# 9.disengage      -2.9513586   0.9999975  -2.9514 0.0031637 ** 
# 10.disengage     -4.6109312   0.0880451 -52.3701 < 2.2e-16 ***
# 11.disengage     -1.2519039   0.9999986  -1.2519 0.2106042    
# 12.disengage     -5.0999545   0.3333332 -15.2999 < 2.2e-16 ***
# 13.disengage     -3.0694899   0.9999975  -3.0695 0.0021442 ** 
# 14.disengage     -3.9995581   0.2000000 -19.9978 < 2.2e-16 ***
# 15.disengage     -3.7115624   0.3779643  -9.8199 < 2.2e-16 ***
# 16.disengage      0.0000000 894.4271910   0.0000 1.0000000    
# 17.disengage     -5.5854796   0.7071053  -7.8991 2.887e-15 ***
# 18.disengage      0.0000000 894.4271910   0.0000 1.0000000    
# 19.disengage     -3.3061330   0.2500000 -13.2245 < 2.2e-16 ***
# 20.disengage     -2.4505089   0.3535533  -6.9311 4.176e-12 ***
# 21.disengage      0.0000000 894.4271910   0.0000 1.0000000    
# 22.disengage     -3.5143729   0.9999972  -3.5144 0.0004408 ***
# 23.disengage    -12.1897538 246.2838524  -0.0495 0.9605250    
# 24.disengage     -1.8919041   0.9999982  -1.8919 0.0585033 .  
# 25.disengage     -2.4340658   0.2000000 -12.1703 < 2.2e-16 ***
# 26.disengage     -3.8049157   0.4082481  -9.3201 < 2.2e-16 ***
# 27.disengage     -5.3682937   0.3162276 -16.9760 < 2.2e-16 ***
# 28.disengage     -5.0921384   0.3535532 -14.4027 < 2.2e-16 ***
# 29.disengage     -5.0977870   0.4082480 -12.4870 < 2.2e-16 ***
# 30.disengage     -4.8310314   0.1622214 -29.7805 < 2.2e-16 ***
# 1.depart         -8.7014746   0.2357022 -36.9172 < 2.2e-16 ***
# 2.depart         -8.6455924   0.2357022 -36.6802 < 2.2e-16 ***
# 3.depart         -8.6031331   0.2357022 -36.5000 < 2.2e-16 ***
# 4.depart         -8.2131081   0.2357022 -34.8453 < 2.2e-16 ***
# 5.depart         -8.7978526   0.2886750 -30.4767 < 2.2e-16 ***
# 6.depart         -8.1073118   0.7071048 -11.4655 < 2.2e-16 ***
# 7.depart         -8.4665832   0.2357022 -35.9207 < 2.2e-16 ***
# 8.depart         -7.6244610   0.4999993 -15.2489 < 2.2e-16 ***
# 9.depart         -8.2131177   0.5773492 -14.2256 < 2.2e-16 ***
# 10.depart        -8.3366519   0.3779642 -22.0567 < 2.2e-16 ***
# 11.depart        -6.1361062   0.5773494 -10.6281 < 2.2e-16 ***
# 12.depart        -8.4305661   0.5773491 -14.6022 < 2.2e-16 ***
# 13.depart        -6.3742619   0.9999954  -6.3743 1.838e-10 ***
# 14.depart        -8.4328617   0.7071047 -11.9259 < 2.2e-16 ***
# 15.depart        -8.4724875   0.9999941  -8.4725 < 2.2e-16 ***
# 16.depart        -4.5531868   0.9999965  -4.5532 5.284e-06 ***
# 17.depart        -7.3325942   0.9999948  -7.3326 2.256e-13 ***
# 18.depart        -6.1293192   0.9999955  -6.1293 8.824e-10 ***
# 19.depart        -8.8972145   0.9999938  -8.8973 < 2.2e-16 ***
# 20.depart        -7.6761506   0.9999946  -7.6762 1.643e-14 ***
# 21.depart        -4.7116752   0.9999964  -4.7117 2.457e-06 ***
# 22.depart        -7.7353839   0.9999945  -7.7354 1.021e-14 ***
# 23.depart        -7.0172213   0.9999950  -7.0173 2.263e-12 ***
# 24.depart        -7.3915096   0.9999948  -7.3915 1.452e-13 ***
# 25.depart        -7.8421735   0.9999945  -7.8422 4.441e-15 ***
# 26.depart        -8.0356182   0.9999944  -8.0357 8.882e-16 ***
# 27.depart        -8.6596852   0.9999940  -8.6597 < 2.2e-16 ***
# 28.depart        -8.1918068   0.9999943  -8.1919 2.220e-16 ***
# 29.depart        -7.0548164   0.9999950  -7.0549 1.728e-12 ***
# 30.depart        -5.7379132   0.1178511 -48.6878 < 2.2e-16 ***
# ---
# Signif. codes:  0 ‘***’ 0.001 ‘**’ 0.01 ‘*’ 0.05 ‘.’ 0.1 ‘ ’ 1
# Null deviance: 76628.21 on 4854 degrees of freedom
# Residual deviance: 30459.98 on 4751 degrees of freedom
# 	Chi-square: 46168.23 on 103 degrees of freedom, asymptotic p-value 0 
# AIC: 30665.98 AICC: 30670.49 BIC: 31334.2 
# Log posterior: -118331 
# Prior parameters: mu=0 sigma=1000 nu=4 

#close, but not quite as good as fitting individual FEs for initCH. I think its so close though- the U and ?? categories are probably pretty real. I'll try fitting just individual effects for initCH for the ID males like I had in the previous model formulations


### Individual initCH Effects for IDmales, but still partition out U and a global rate

sequences<-table(event.list$Date)

#create list of dummy intercept names

idno<-length(event.types[grep("cop.start",event.types)])

dummy.ints.attack<-paste(1:14,".attackFE", sep="")
dummy.ints.receive<-paste(1:13, ".receiveFE", sep="")

IDintsCH<-paste(1:14,"initCH",sep=".")

int.names<-c("Global.attackFE", "X.receiveFE", "X.UreceiveFE", "U.attackFE", "U.receiveFE", "U.UattackFE", dummy.ints.attack, dummy.ints.receive, "cop.startFE", "cop.endFE", "initFO.FE","ID.initFO.FE", "U.initFO.FE","initSF.FE","X.initSF.FE", "initCH.FE",  "U.initCH.FE", IDintsCH, event.types[-c(grep("attack", event.types),grep("cop", event.types),grep("initFO", event.types),grep("initSF", event.types), grep("initCH", event.types))])

int.list<-list()

for(i in 1:length(levels(event.list$Date))){

day.array<-array(0,dim=c(as.numeric(sequences[i]), length(event.types),length(int.names)))

#the third dimension is the stat slice.
dimnames(day.array)<-list(c(1:sequences[i]),as.character(event.types),as.character(int.names))

#add global attack effect, loops for "U" intercepts for attacks and receives, then one loop for typical intercepts

day.array[,grep("attack", event.types),1]<-1


for(h in 1:14){
day.array[,paste(h,".attack.",30, sep=""),2]<-1
}


for(h in 15:29){
day.array[,paste(h,".attack.",30, sep=""),3]<-1
}


for(h in 1:14){
	for(j in 15:29){	
day.array[,paste(j,".attack.", h, sep=""),4]<-1
}
}


for(h in 1:14){
	for(j in 15:29){
day.array[,paste(h,".attack.",j, sep=""),5]<-1
}
}

for(h in 15:29){
	for(j in 15:29){	
if(h!=j){
day.array[,paste(j,".attack.", h, sep=""),6]<-1
}
}
}

preloop<-6

#loop for dummy intercepts for attacks, loop for dummy intercepts for receives, add in cop.start and cop.end FEs, then one loop for typical intercepts

for(k in (preloop+1):(length(dummy.ints.attack)+preloop)){
	
	for(h in 1:14){
if(h!=(k-preloop)){
day.array[,paste((k-preloop),".attack.", h, sep=""),k]<-1
}
}
}

for(k in ((length(dummy.ints.attack)+preloop)+1):((length(dummy.ints.attack)+preloop)+length(dummy.ints.receive))){
	for(h in 1:14){
if(h!=(k-(length(dummy.ints.attack)+preloop))){
day.array[,paste(h,".attack.",(k-(length(dummy.ints.attack)+preloop)), sep=""),k]<-1
}
}
}

#cop.startFE
for(h in 1:idno){
day.array[,paste(h,".cop.start", sep=""),((length(dummy.ints.attack)+preloop)+length(dummy.ints.receive)+1)]<-1
}


#cop.endFE
for(h in 1:idno){
day.array[,paste(h,".cop.end", sep=""),((length(dummy.ints.attack)+preloop)+length(dummy.ints.receive)+2)]<-1
}

#initFO.FE
for(h in 1:idno){
day.array[,paste(h,".initFO", sep=""),((length(dummy.ints.attack)+preloop)+length(dummy.ints.receive)+3)]<-1
}

#ID.initFO.FE
for(h in 1:14){
day.array[,paste(h,".initFO", sep=""),((length(dummy.ints.attack)+preloop)+length(dummy.ints.receive)+4)]<-1
}

#U.initFO.FE
for(h in 15:29){
day.array[,paste(h,".initFO", sep=""),((length(dummy.ints.attack)+preloop)+length(dummy.ints.receive)+5)]<-1
}

#initSF.FE
for(h in 1:idno){
day.array[,paste(h,".initSF", sep=""),((length(dummy.ints.attack)+preloop)+length(dummy.ints.receive)+6)]<-1
}

#X.initSF.FE
day.array[,paste(30,".initSF", sep=""),((length(dummy.ints.attack)+preloop)+length(dummy.ints.receive)+7)]<-1


#initCH.FE
for(h in 1:idno){
day.array[,paste(h,".initCH", sep=""),((length(dummy.ints.attack)+preloop)+length(dummy.ints.receive)+8)]<-1
}


#U.initCH.FE
for(h in 15:29){
day.array[,paste(h,".initCH", sep=""),((length(dummy.ints.attack)+preloop)+length(dummy.ints.receive)+9)]<-1
}

#everything else
for(k in (((length(dummy.ints.attack)+preloop)+length(dummy.ints.receive)+10):length(int.names))){
	
	day.array[,int.names[k],k]<-1
	
}

#weird extra formatting to make it match the package example, seems like it becomes relevant *punintended!* if I want to fit day-specific effects

global<-list()

global[[1]]<-day.array

int.list[[i]]<-global

names(int.list[[i]])<-"global"

}


tic()
intfit.sup<-rem(evlist, int.list, supplist=supplist, timing="interval", estimator="BPM")
toc()

summary(intfit.sup)

# Egocentric Relational Event Model (Interval Likelihood)
# 
#                   Post.Mode     Post.SD  Z value  Pr(>|z|)    
# Global.attackFE  -9.5259220   0.2886747 -32.9988 < 2.2e-16 ***
# X.receiveFE       3.7712555   0.2921739  12.9076 < 2.2e-16 ***
# X.UreceiveFE      3.9025235   0.3370996  11.5768 < 2.2e-16 ***
# U.attackFE       -1.2734082   0.6454967  -1.9728 0.0485233 *  
# U.receiveFE       1.7877281   0.3145761   5.6830 1.324e-08 ***
# U.UattackFE       2.9176286   0.3248928   8.9803 < 2.2e-16 ***
# 1.attackFE        1.0330374   0.3944376   2.6190 0.0088184 ** 
# 2.attackFE        1.2763648   0.3921908   3.2544 0.0011361 ** 
# 3.attackFE        1.6061436   0.3891684   4.1271 3.673e-05 ***
# 4.attackFE        2.5265882   0.3860257   6.5451 5.944e-11 ***
# 5.attackFE        0.3110660   0.4195142   0.7415 0.4583958    
# 6.attackFE       -0.0207165   0.8047343  -0.0257 0.9794621    
# 7.attackFE        1.3366638   0.3932445   3.3991 0.0006762 ***
# 8.attackFE       -1.4929214   1.0698668  -1.3954 0.1628870    
# 9.attackFE      -12.5413190 243.0663149  -0.0516 0.9588504    
# 10.attackFE       1.5764042   0.4001704   3.9393 8.171e-05 ***
# 11.attackFE     -10.4618029 264.1957859  -0.0396 0.9684131    
# 12.attackFE      -0.0121458   0.5502168  -0.0221 0.9823884    
# 13.attackFE       0.9853197   1.0715358   0.9195 0.3578134    
# 14.attackFE       0.7284843   0.5224387   1.3944 0.1631993    
# 1.receiveFE      -0.7315755   0.2770237  -2.6408 0.0082700 ** 
# 2.receiveFE      -0.2483185   0.2681292  -0.9261 0.3543862    
# 3.receiveFE      -0.3750730   0.2725416  -1.3762 0.1687582    
# 4.receiveFE       1.0326554   0.2610273   3.9561 7.618e-05 ***
# 5.receiveFE      -0.8724350   0.2925982  -2.9817 0.0028667 ** 
# 6.receiveFE       0.7239207   0.3403191   2.1272 0.0334050 *  
# 7.receiveFE       0.2399268   0.2626681   0.9134 0.3610207    
# 8.receiveFE       0.0322648   0.3395839   0.0950 0.9243046    
# 9.receiveFE       0.6527401   0.3048317   2.1413 0.0322488 *  
# 10.receiveFE     -0.0083194   0.2918528  -0.0285 0.9772590    
# 11.receiveFE      1.6776185   0.4538049   3.6968 0.0002183 ***
# 12.receiveFE     -0.4512307   0.3601768  -1.2528 0.2102774    
# 13.receiveFE     -0.3517487   1.0308039  -0.3412 0.7329249    
# cop.startFE      -8.8434147   0.1178511 -75.0389 < 2.2e-16 ***
# cop.endFE        -1.4478174   0.1414214 -10.2376 < 2.2e-16 ***
# initFO.FE         3.3524074   0.2581988  12.9838 < 2.2e-16 ***
# ID.initFO.FE      3.2420533   0.2612259  12.4109 < 2.2e-16 ***
# U.initFO.FE       1.8903400   0.3380617   5.5917 2.249e-08 ***
# initSF.FE         5.1608463   0.0764719  67.4868 < 2.2e-16 ***
# X.initSF.FE      -1.2206524   0.2070869  -5.8944 3.761e-09 ***
# initCH.FE         6.8243734   0.0455016 149.9810 < 2.2e-16 ***
# U.initCH.FE      -0.4517611   0.1321174  -3.4194 0.0006276 ***
# 1.initCH         -4.2070723   1.0010314  -4.2027 2.637e-05 ***
# 2.initCH        -15.4875101 220.2835462  -0.0703 0.9439492    
# 3.initCH        -15.2821666 221.6680693  -0.0689 0.9450361    
# 4.initCH         -5.3102391   1.0010307  -5.3048 1.128e-07 ***
# 5.initCH         -3.7452549   1.0010317  -3.7414 0.0001830 ***
# 6.initCH         -0.0884684   0.2541070  -0.3482 0.7277245    
# 7.initCH        -15.8573990 217.8536648  -0.0728 0.9419738    
# 8.initCH         -0.2614586   0.2467669  -1.0595 0.2893554    
# 9.initCH          0.0535289   0.1799262   0.2975 0.7660812    
# 10.initCH        -2.3589644   0.5020659  -4.6985 2.621e-06 ***
# 11.initCH        -0.0707688   0.4107761  -0.1723 0.8632168    
# 12.initCH       -13.6065317 234.0361119  -0.0581 0.9536382    
# 13.initCH       -11.1018997 257.1141752  -0.0432 0.9655589    
# 14.initCH        -0.5526068   0.3364246  -1.6426 0.1004683    
# 1.disengage      -4.2622836   0.0626224 -68.0632 < 2.2e-16 ***
# 2.disengage      -5.1487697   0.0887356 -58.0237 < 2.2e-16 ***
# 3.disengage      -4.7750140   0.0636285 -75.0452 < 2.2e-16 ***
# 4.disengage      -4.6474529   0.0548821 -84.6806 < 2.2e-16 ***
# 5.disengage      -4.5737155   0.1025978 -44.5791 < 2.2e-16 ***
# 6.disengage      -6.4777863   0.5773494 -11.2199 < 2.2e-16 ***
# 7.disengage      -5.6831776   0.0894427 -63.5399 < 2.2e-16 ***
# 8.disengage      -3.0256324   0.1889822 -16.0101 < 2.2e-16 ***
# 9.disengage      -2.9513586   0.9999975  -2.9514 0.0031637 ** 
# 10.disengage     -4.6109312   0.0880451 -52.3701 < 2.2e-16 ***
# 11.disengage     -1.2519039   0.9999986  -1.2519 0.2106042    
# 12.disengage     -5.0999545   0.3333332 -15.2999 < 2.2e-16 ***
# 13.disengage     -3.0694899   0.9999975  -3.0695 0.0021442 ** 
# 14.disengage     -3.9995581   0.2000000 -19.9978 < 2.2e-16 ***
# 15.disengage     -3.7115624   0.3779643  -9.8199 < 2.2e-16 ***
# 16.disengage      0.0000000 894.4271910   0.0000 1.0000000    
# 17.disengage     -5.5854796   0.7071053  -7.8991 2.887e-15 ***
# 18.disengage      0.0000000 894.4271910   0.0000 1.0000000    
# 19.disengage     -3.3061330   0.2500000 -13.2245 < 2.2e-16 ***
# 20.disengage     -2.4505089   0.3535533  -6.9311 4.176e-12 ***
# 21.disengage      0.0000000 894.4271910   0.0000 1.0000000    
# 22.disengage     -3.5143729   0.9999972  -3.5144 0.0004408 ***
# 23.disengage    -12.1897538 246.2838524  -0.0495 0.9605250    
# 24.disengage     -1.8919041   0.9999982  -1.8919 0.0585033 .  
# 25.disengage     -2.4340658   0.2000000 -12.1703 < 2.2e-16 ***
# 26.disengage     -3.8049157   0.4082481  -9.3201 < 2.2e-16 ***
# 27.disengage     -5.3682937   0.3162276 -16.9760 < 2.2e-16 ***
# 28.disengage     -5.0921384   0.3535532 -14.4027 < 2.2e-16 ***
# 29.disengage     -5.0977870   0.4082480 -12.4870 < 2.2e-16 ***
# 30.disengage     -4.8310314   0.1622214 -29.7805 < 2.2e-16 ***
# 1.depart         -8.7014746   0.2357022 -36.9172 < 2.2e-16 ***
# 2.depart         -8.6455924   0.2357022 -36.6802 < 2.2e-16 ***
# 3.depart         -8.6031331   0.2357022 -36.5000 < 2.2e-16 ***
# 4.depart         -8.2131081   0.2357022 -34.8453 < 2.2e-16 ***
# 5.depart         -8.7978526   0.2886750 -30.4767 < 2.2e-16 ***
# 6.depart         -8.1073118   0.7071048 -11.4655 < 2.2e-16 ***
# 7.depart         -8.4665832   0.2357022 -35.9207 < 2.2e-16 ***
# 8.depart         -7.6244610   0.4999993 -15.2489 < 2.2e-16 ***
# 9.depart         -8.2131177   0.5773492 -14.2256 < 2.2e-16 ***
# 10.depart        -8.3366519   0.3779642 -22.0567 < 2.2e-16 ***
# 11.depart        -6.1361062   0.5773494 -10.6281 < 2.2e-16 ***
# 12.depart        -8.4305661   0.5773491 -14.6022 < 2.2e-16 ***
# 13.depart        -6.3742619   0.9999954  -6.3743 1.838e-10 ***
# 14.depart        -8.4328617   0.7071047 -11.9259 < 2.2e-16 ***
# 15.depart        -8.4724875   0.9999941  -8.4725 < 2.2e-16 ***
# 16.depart        -4.5531868   0.9999965  -4.5532 5.284e-06 ***
# 17.depart        -7.3325942   0.9999948  -7.3326 2.256e-13 ***
# 18.depart        -6.1293192   0.9999955  -6.1293 8.824e-10 ***
# 19.depart        -8.8972145   0.9999938  -8.8973 < 2.2e-16 ***
# 20.depart        -7.6761506   0.9999946  -7.6762 1.643e-14 ***
# 21.depart        -4.7116752   0.9999964  -4.7117 2.457e-06 ***
# 22.depart        -7.7353839   0.9999945  -7.7354 1.021e-14 ***
# 23.depart        -7.0172213   0.9999950  -7.0173 2.263e-12 ***
# 24.depart        -7.3915096   0.9999948  -7.3915 1.452e-13 ***
# 25.depart        -7.8421735   0.9999945  -7.8422 4.441e-15 ***
# 26.depart        -8.0356182   0.9999944  -8.0357 8.882e-16 ***
# 27.depart        -8.6596852   0.9999940  -8.6597 < 2.2e-16 ***
# 28.depart        -8.1918068   0.9999943  -8.1919 2.220e-16 ***
# 29.depart        -7.0548164   0.9999950  -7.0549 1.728e-12 ***
# 30.depart        -5.7379132   0.1178511 -48.6878 < 2.2e-16 ***
# ---
# Signif. codes:  0 ‘***’ 0.001 ‘**’ 0.01 ‘*’ 0.05 ‘.’ 0.1 ‘ ’ 1
# Null deviance: 76628.21 on 4854 degrees of freedom
# Residual deviance: 30135.79 on 4738 degrees of freedom
# 	Chi-square: 46492.42 on 116 degrees of freedom, asymptotic p-value 0 
# AIC: 30367.79 AICC: 30373.52 BIC: 31120.35 
# Log posterior: -131181.7 
# Prior parameters: mu=0 sigma=1000 nu=4 

#best among fixed chase effects models so far! 


### ID FE and individual effects for U's 

sequences<-table(event.list$Date)

#create list of dummy intercept names

idno<-length(event.types[grep("cop.start",event.types)])

dummy.ints.attack<-paste(1:14,".attackFE", sep="")
dummy.ints.receive<-paste(1:13, ".receiveFE", sep="")

UintsCH<-paste(15:29,"initCH",sep=".")

int.names<-c("Global.attackFE", "X.receiveFE", "X.UreceiveFE", "U.attackFE", "U.receiveFE", "U.UattackFE", dummy.ints.attack, dummy.ints.receive, "cop.startFE", "cop.endFE", "initFO.FE","ID.initFO.FE", "U.initFO.FE","initSF.FE","X.initSF.FE", "initCH.FE", "ID.initCH.FE", UintsCH, event.types[-c(grep("attack", event.types),grep("cop", event.types),grep("initFO", event.types),grep("initSF", event.types), grep("initCH", event.types))])

int.list<-list()

for(i in 1:length(levels(event.list$Date))){

day.array<-array(0,dim=c(as.numeric(sequences[i]), length(event.types),length(int.names)))

#the third dimension is the stat slice.
dimnames(day.array)<-list(c(1:sequences[i]),as.character(event.types),as.character(int.names))

#add global attack effect, loops for "U" intercepts for attacks and receives, then one loop for typical intercepts

day.array[,grep("attack", event.types),1]<-1


for(h in 1:14){
day.array[,paste(h,".attack.",30, sep=""),2]<-1
}


for(h in 15:29){
day.array[,paste(h,".attack.",30, sep=""),3]<-1
}


for(h in 1:14){
	for(j in 15:29){	
day.array[,paste(j,".attack.", h, sep=""),4]<-1
}
}


for(h in 1:14){
	for(j in 15:29){
day.array[,paste(h,".attack.",j, sep=""),5]<-1
}
}

for(h in 15:29){
	for(j in 15:29){	
if(h!=j){
day.array[,paste(j,".attack.", h, sep=""),6]<-1
}
}
}

preloop<-6

#loop for dummy intercepts for attacks, loop for dummy intercepts for receives, add in cop.start and cop.end FEs, then one loop for typical intercepts

for(k in (preloop+1):(length(dummy.ints.attack)+preloop)){
	
	for(h in 1:14){
if(h!=(k-preloop)){
day.array[,paste((k-preloop),".attack.", h, sep=""),k]<-1
}
}
}

for(k in ((length(dummy.ints.attack)+preloop)+1):((length(dummy.ints.attack)+preloop)+length(dummy.ints.receive))){
	for(h in 1:14){
if(h!=(k-(length(dummy.ints.attack)+preloop))){
day.array[,paste(h,".attack.",(k-(length(dummy.ints.attack)+preloop)), sep=""),k]<-1
}
}
}

#cop.startFE
for(h in 1:idno){
day.array[,paste(h,".cop.start", sep=""),((length(dummy.ints.attack)+preloop)+length(dummy.ints.receive)+1)]<-1
}


#cop.endFE
for(h in 1:idno){
day.array[,paste(h,".cop.end", sep=""),((length(dummy.ints.attack)+preloop)+length(dummy.ints.receive)+2)]<-1
}

#initFO.FE
for(h in 1:idno){
day.array[,paste(h,".initFO", sep=""),((length(dummy.ints.attack)+preloop)+length(dummy.ints.receive)+3)]<-1
}

#ID.initFO.FE
for(h in 1:14){
day.array[,paste(h,".initFO", sep=""),((length(dummy.ints.attack)+preloop)+length(dummy.ints.receive)+4)]<-1
}

#U.initFO.FE
for(h in 15:29){
day.array[,paste(h,".initFO", sep=""),((length(dummy.ints.attack)+preloop)+length(dummy.ints.receive)+5)]<-1
}

#initSF.FE
for(h in 1:idno){
day.array[,paste(h,".initSF", sep=""),((length(dummy.ints.attack)+preloop)+length(dummy.ints.receive)+6)]<-1
}

#X.initSF.FE
day.array[,paste(30,".initSF", sep=""),((length(dummy.ints.attack)+preloop)+length(dummy.ints.receive)+7)]<-1

#initCH.FE
for(h in 1:idno){
day.array[,paste(h,".initCH", sep=""),((length(dummy.ints.attack)+preloop)+length(dummy.ints.receive)+8)]<-1
}

#ID.initCH.FE
for(h in 1:14){
day.array[,paste(h,".initCH", sep=""),((length(dummy.ints.attack)+preloop)+length(dummy.ints.receive)+9)]<-1
}

# #U.initCH.FE
# for(h in 15:29){
# day.array[,paste(h,".initCH", sep=""),((length(dummy.ints.attack)+preloop)+length(dummy.ints.receive)+11)]<-1
# }

#everything else
for(k in (((length(dummy.ints.attack)+preloop)+length(dummy.ints.receive)+10):length(int.names))){
	
	day.array[,int.names[k],k]<-1
	
}

#weird extra formatting to make it match the package example, seems like it becomes relevant *punintended!* if I want to fit day-specific effects

global<-list()

global[[1]]<-day.array
```
