## Supplementary figures and images for "Fighting isn’t sexy in lekking Greater Sage-grouse (*Centrocercus urophasianus*)"

### attendfight_combined.pdf

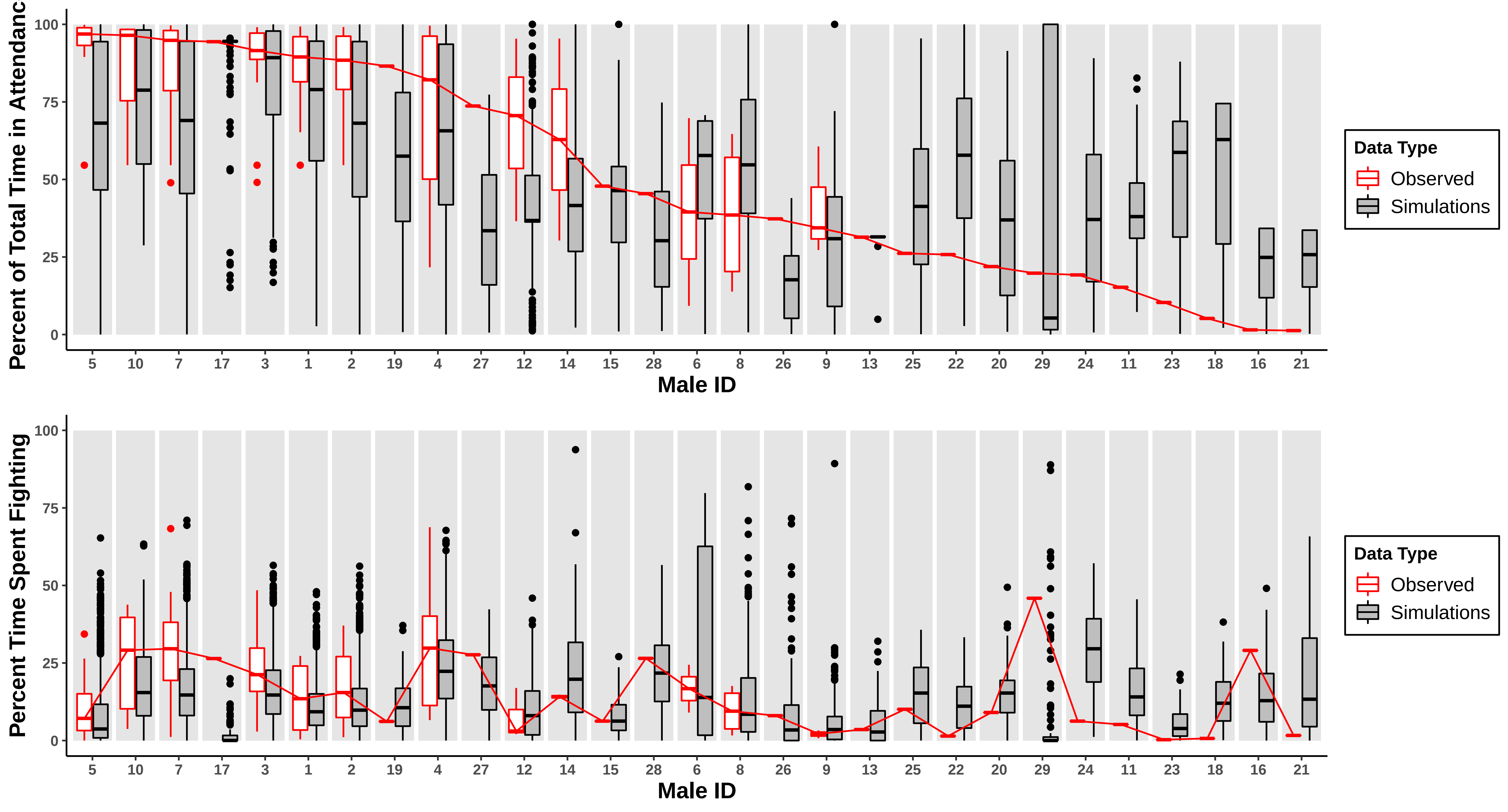

### attendfight_combined.pdf

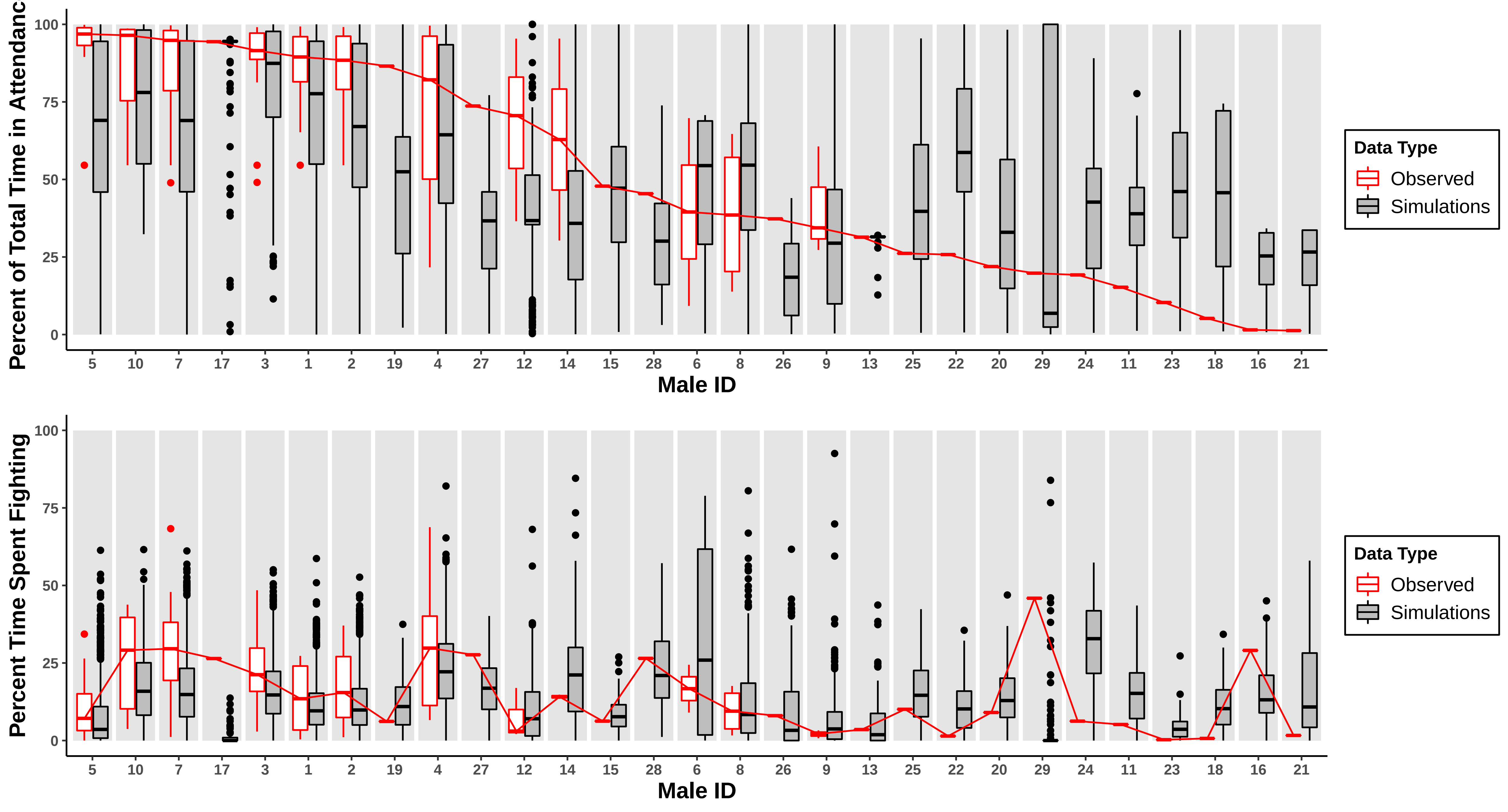

### copint_combined.pdf

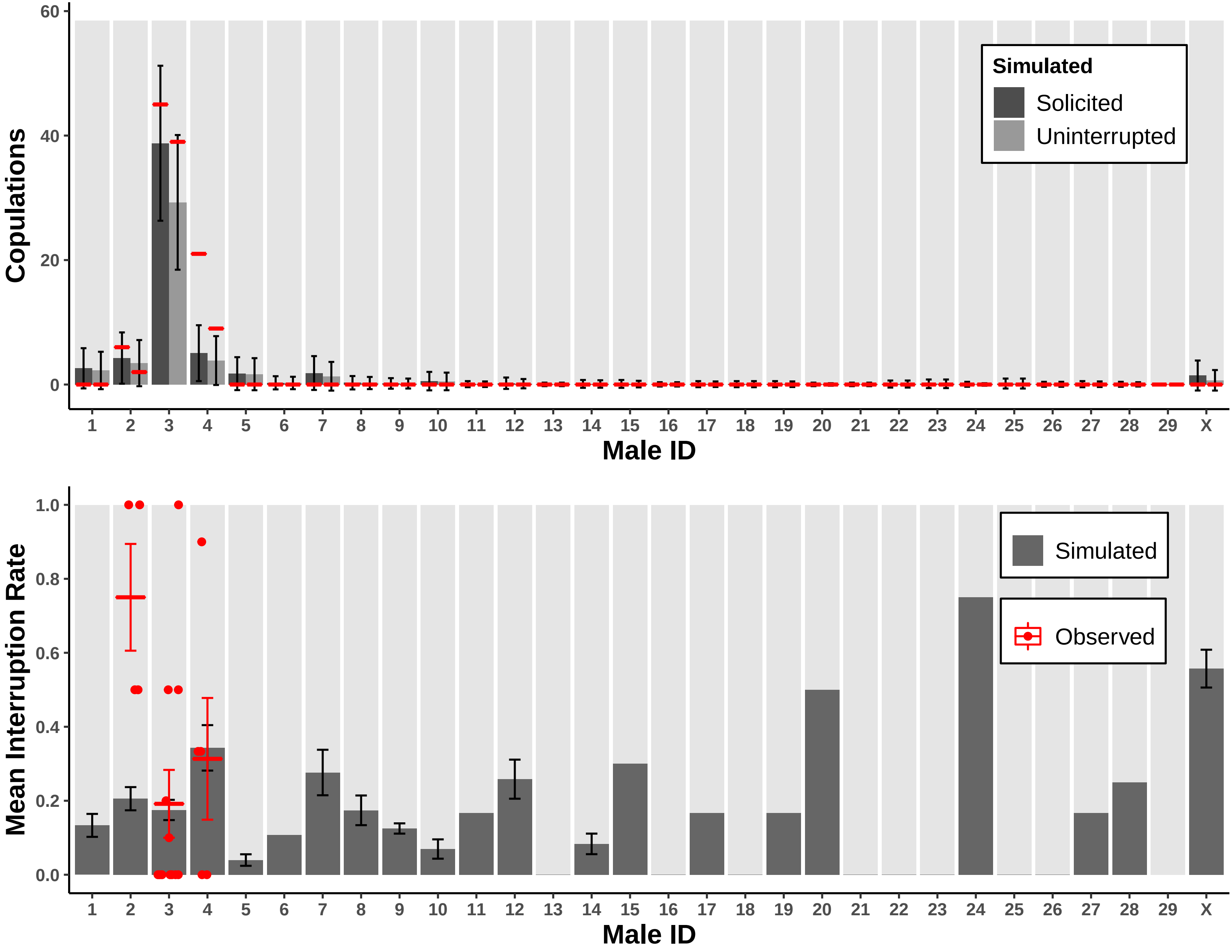

### copint_combined.pdf

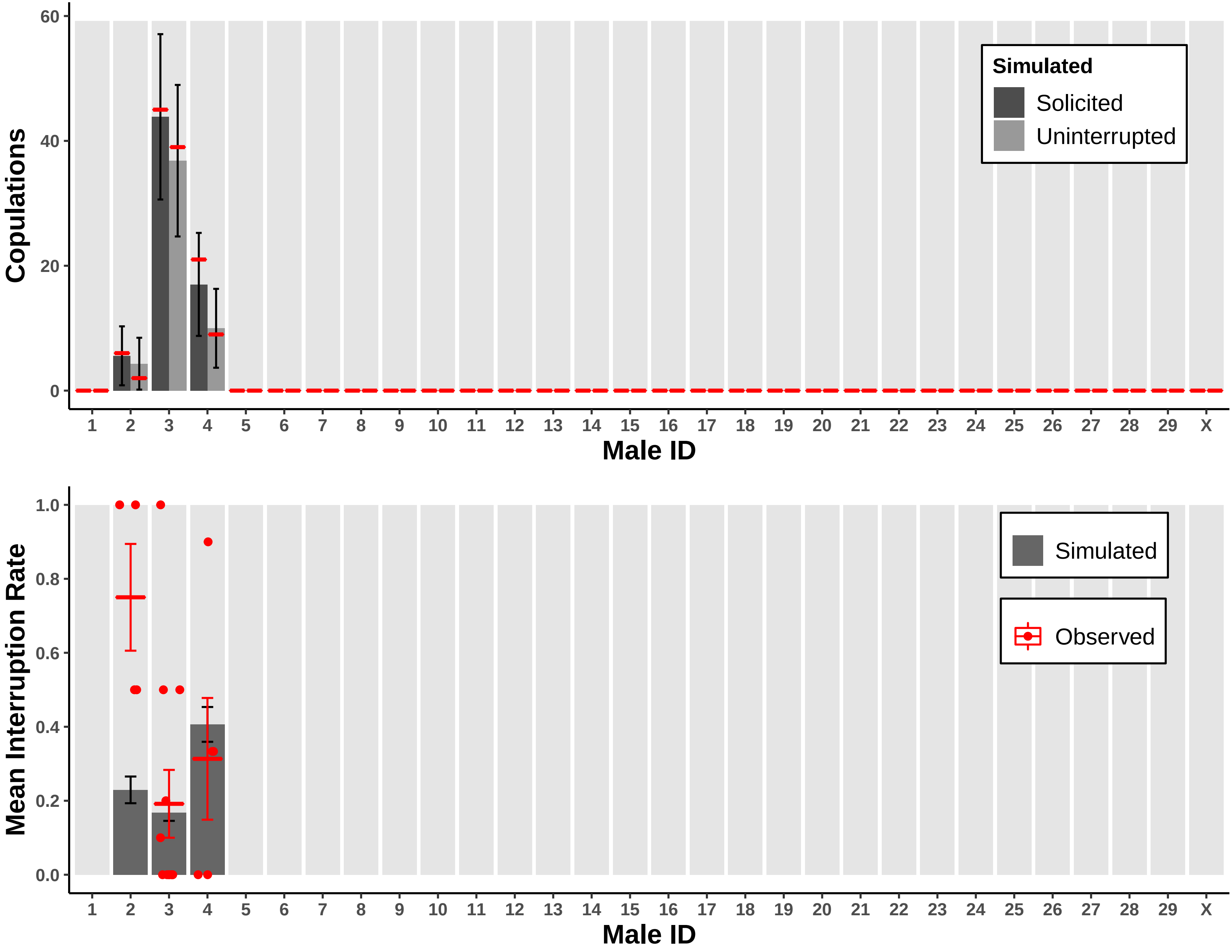

### copint_small.pdf

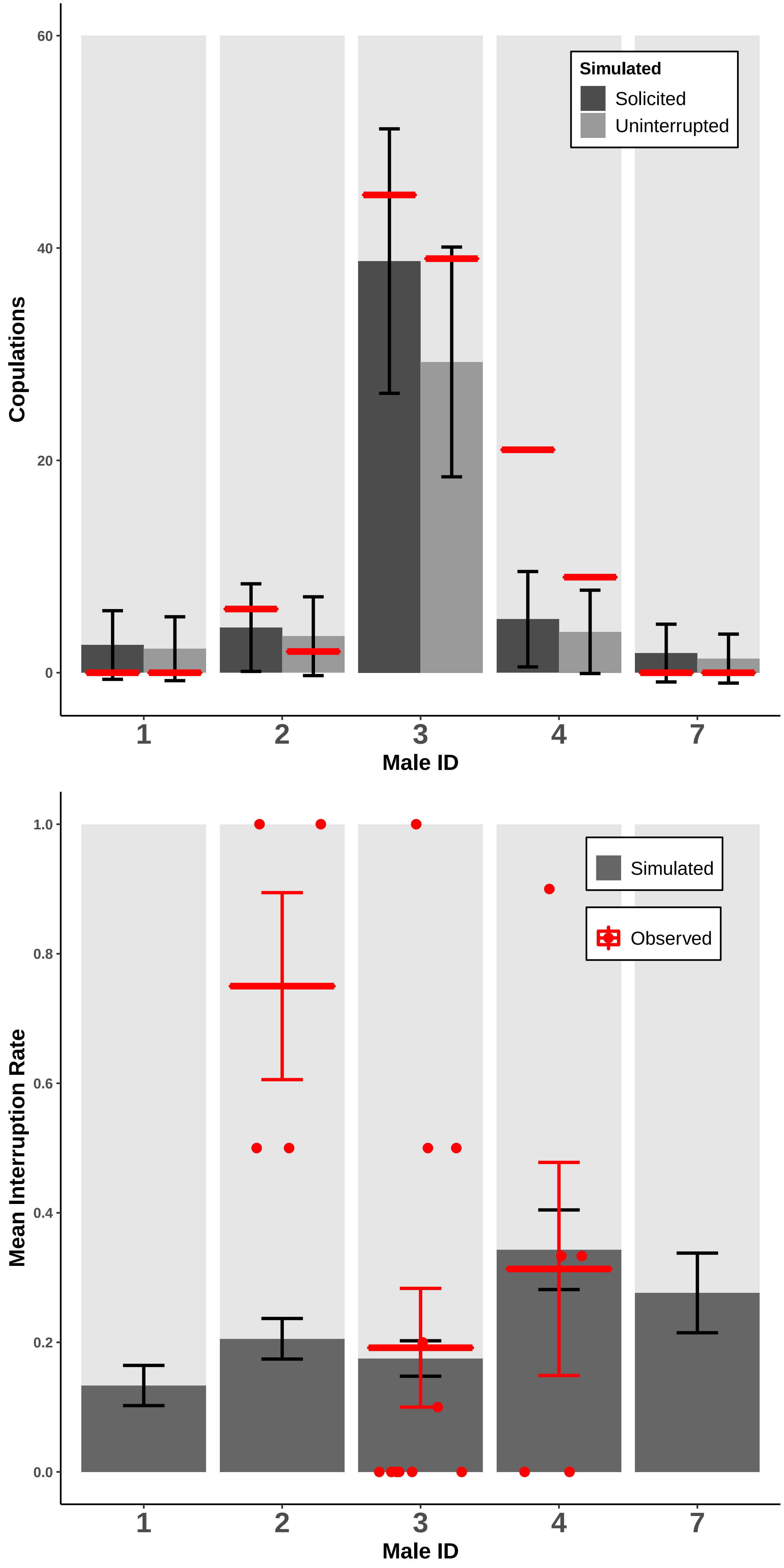

### copint_small.pdf

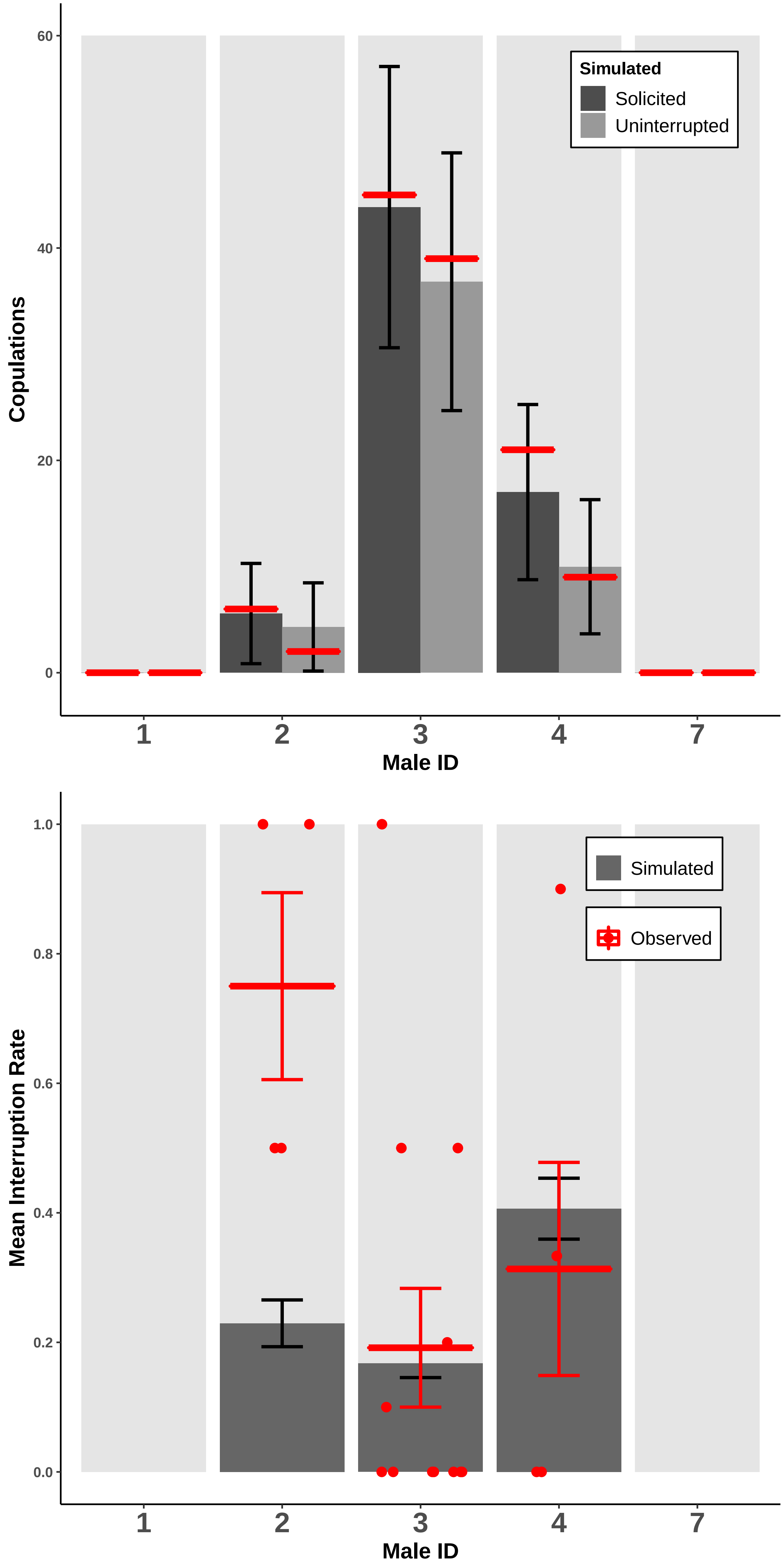

### fight_count_fig.pdf

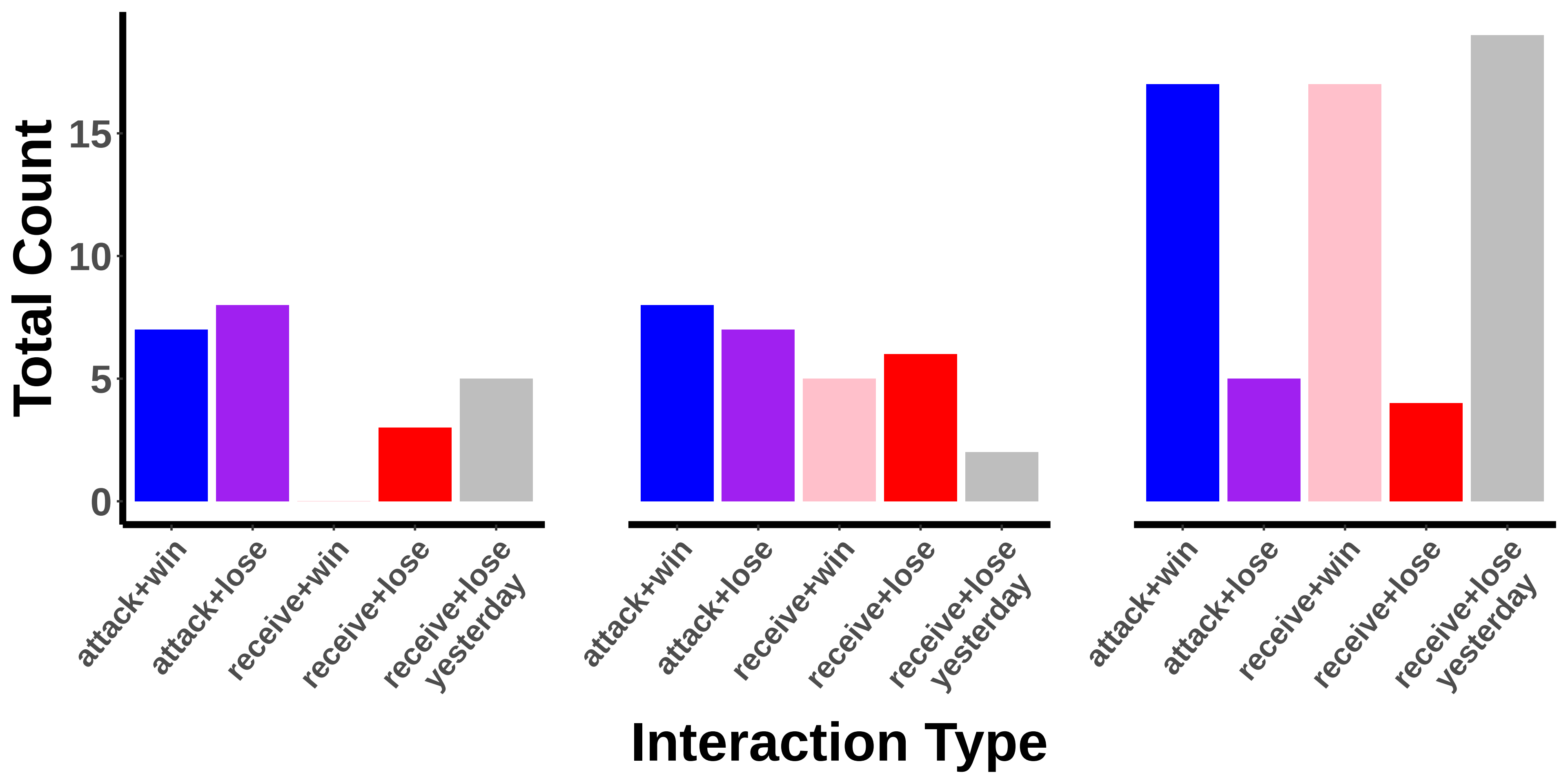

### fight_effect_fig.pdf

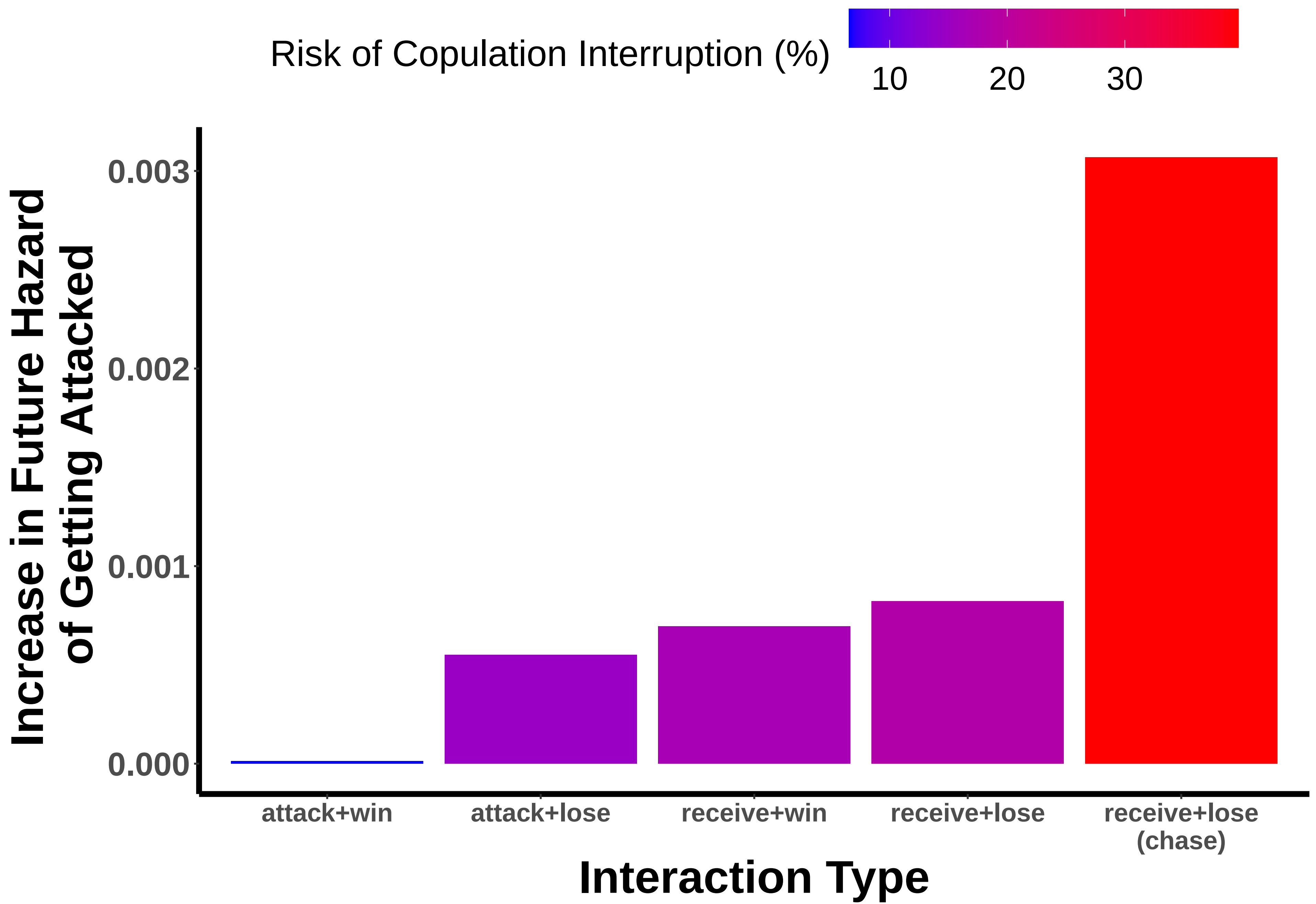

### matebase.pdf

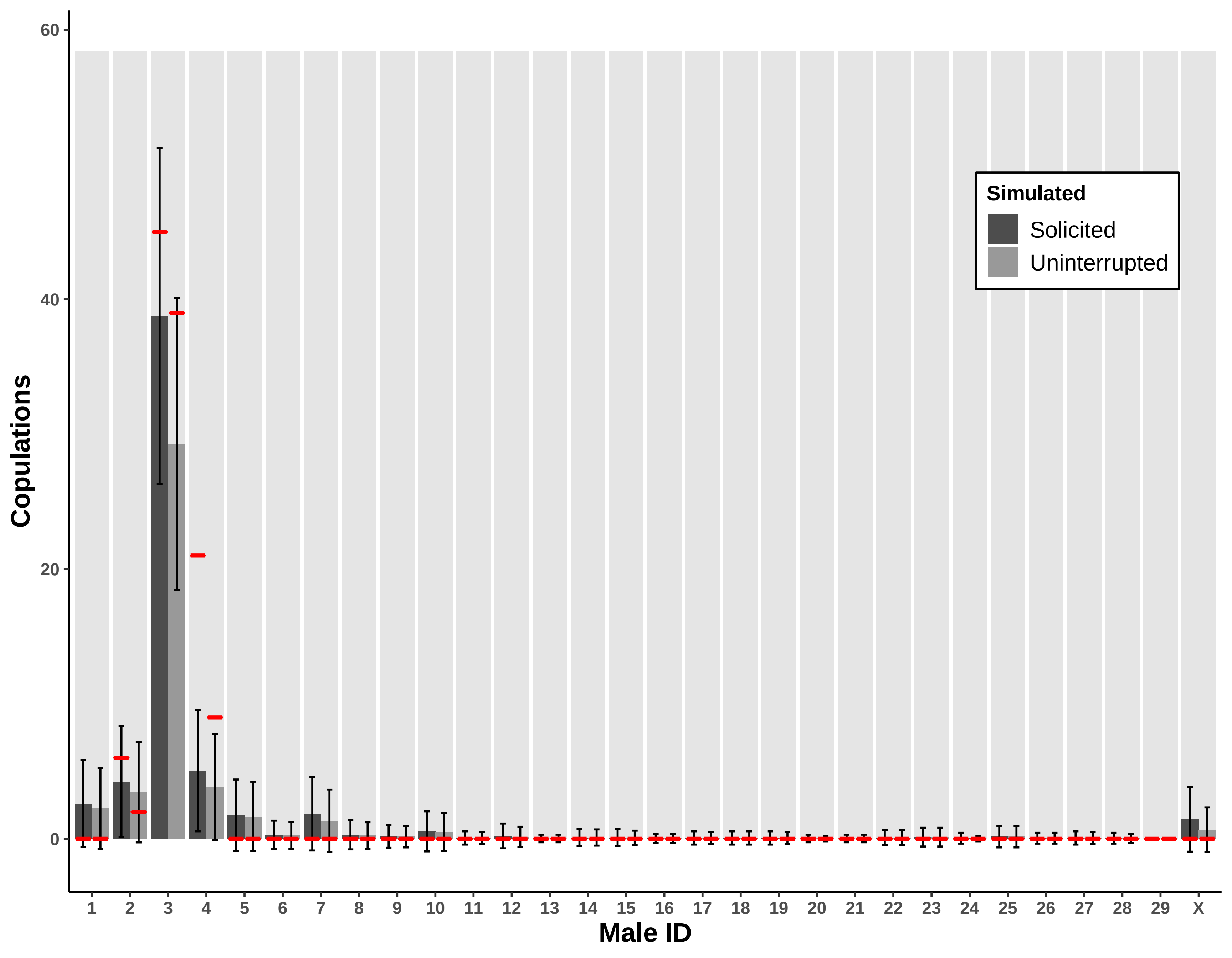

### matebase.pdf

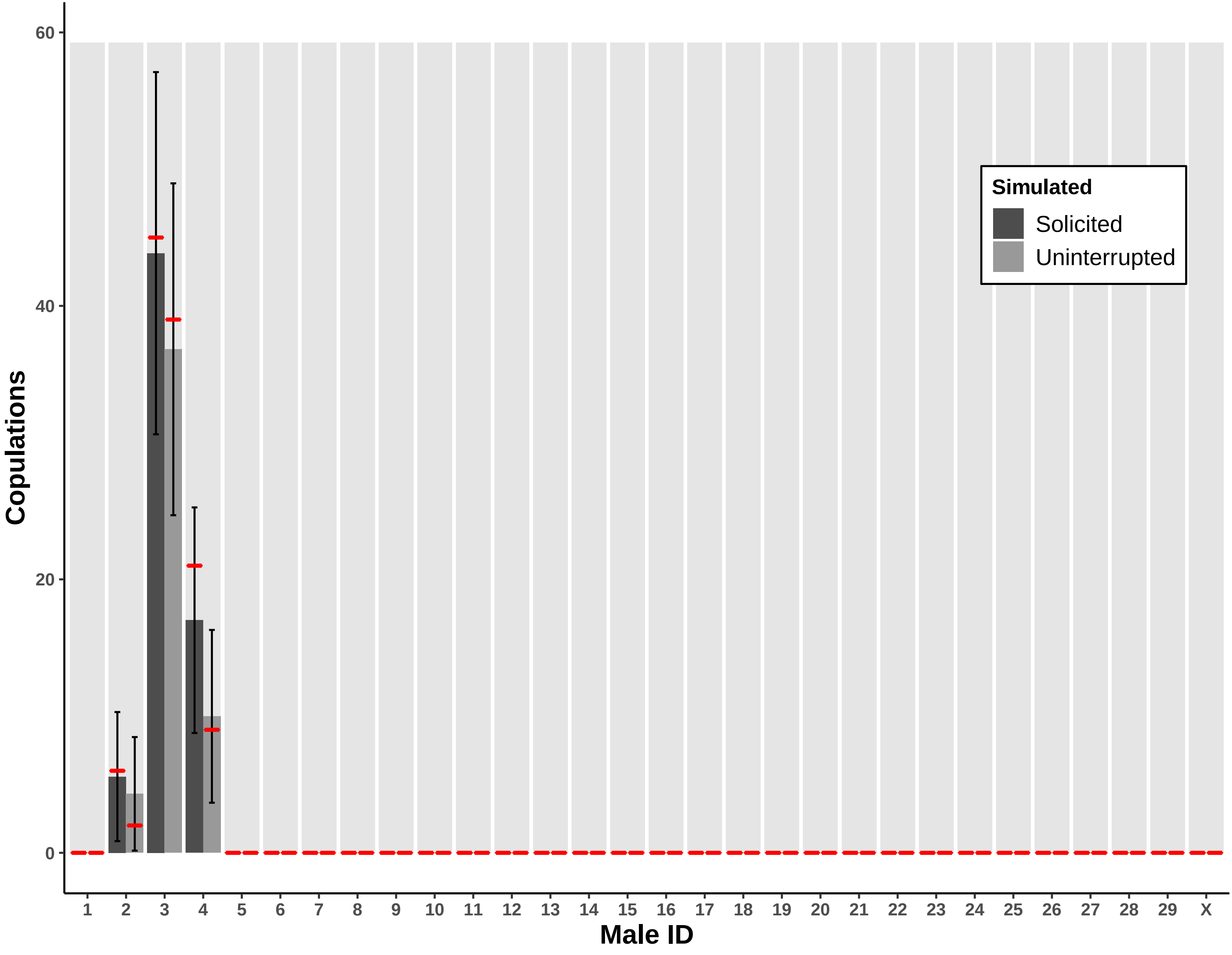

### matesmall.pdf

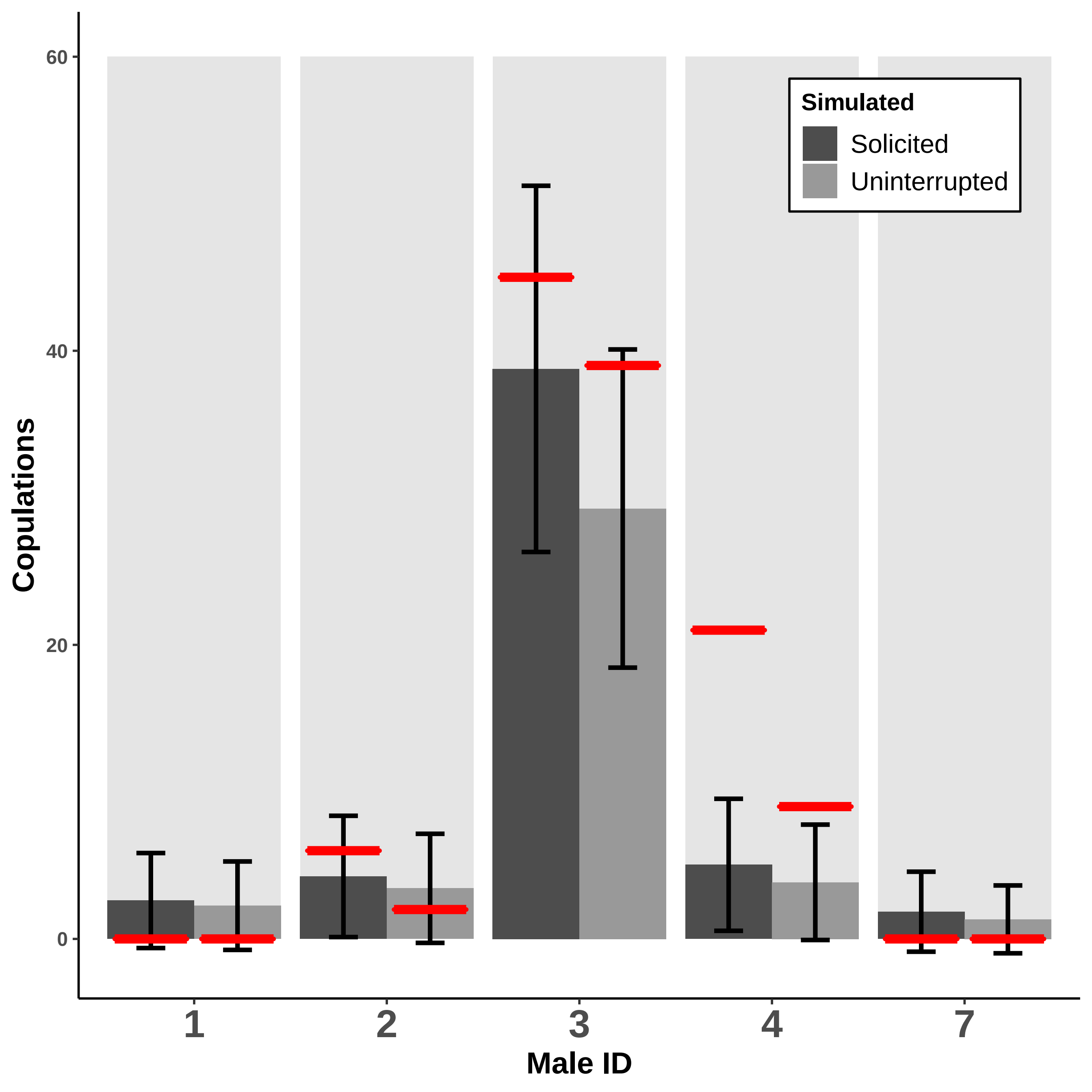

### matesmall.pdf

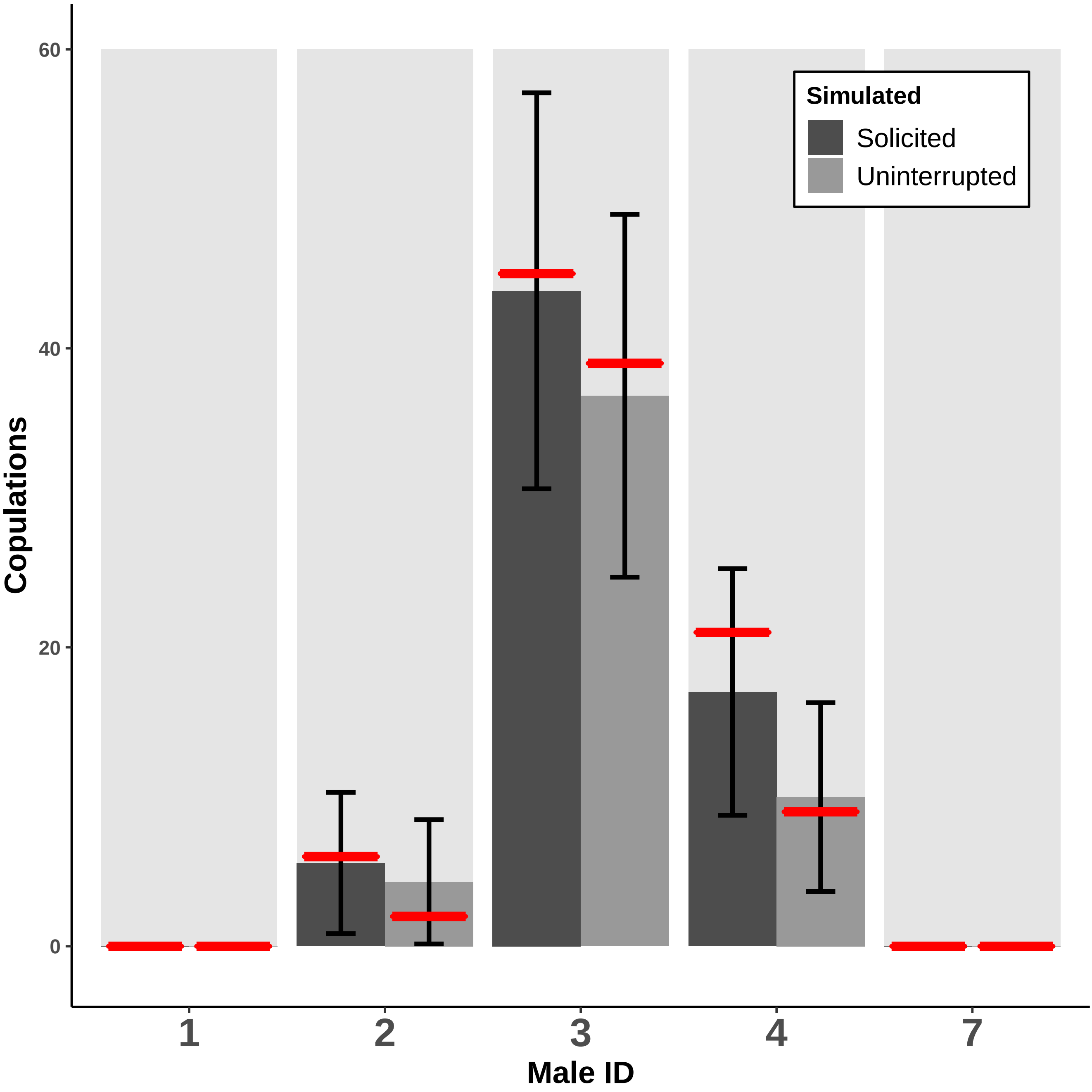

### matingplot_new_statlevs_newlabs.pdf

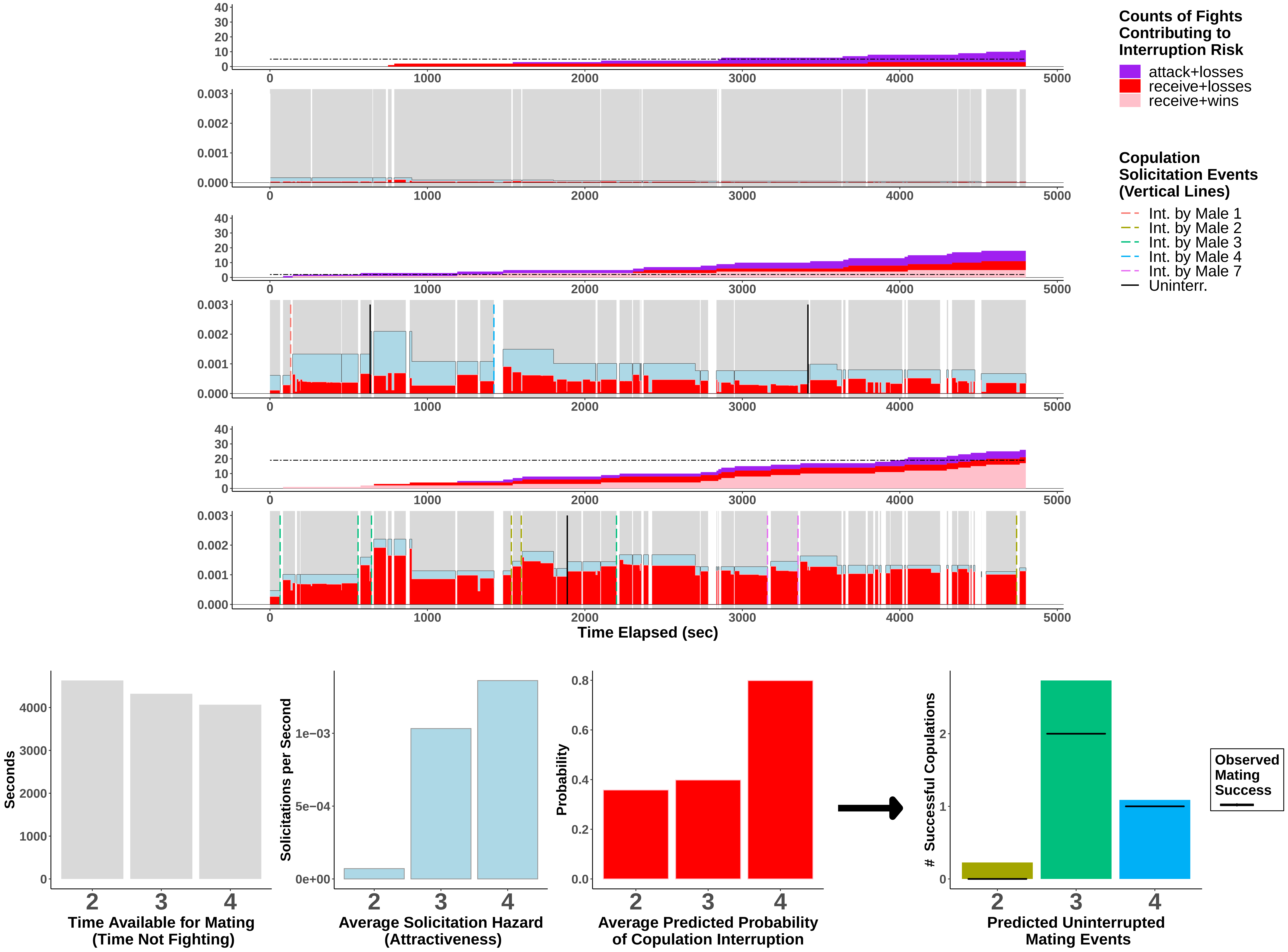

### overallcount_plot.pdf

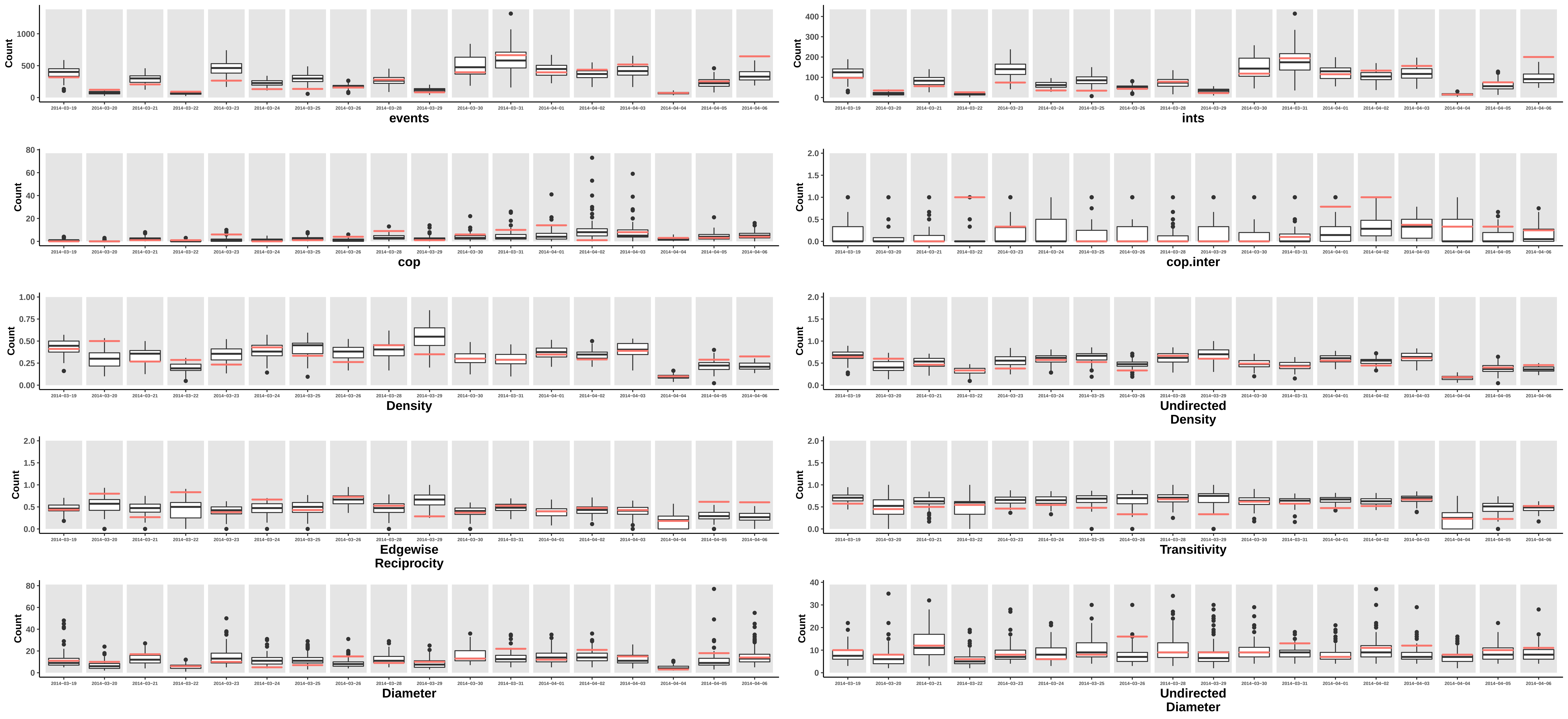

### overallcount_plot.pdf

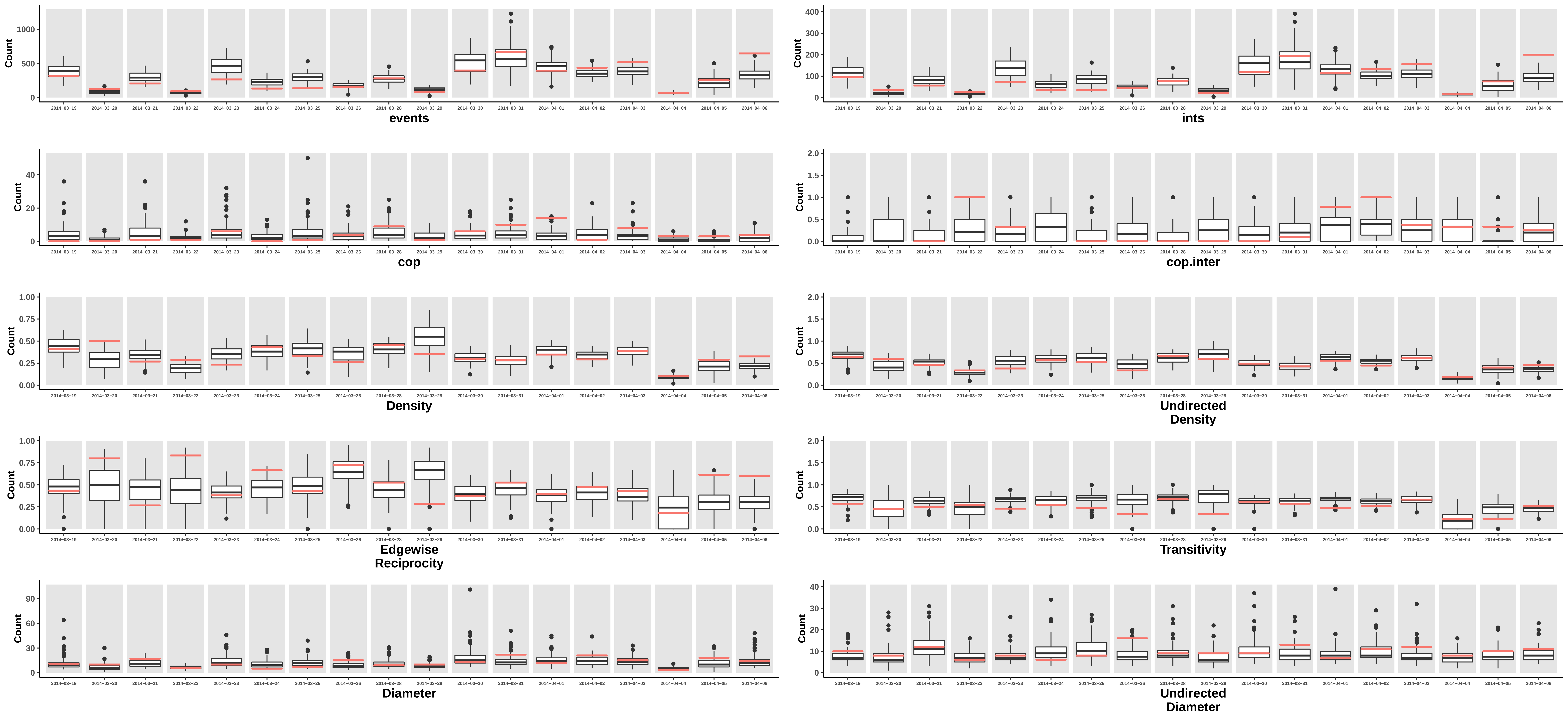

### overallfightplot.pdf

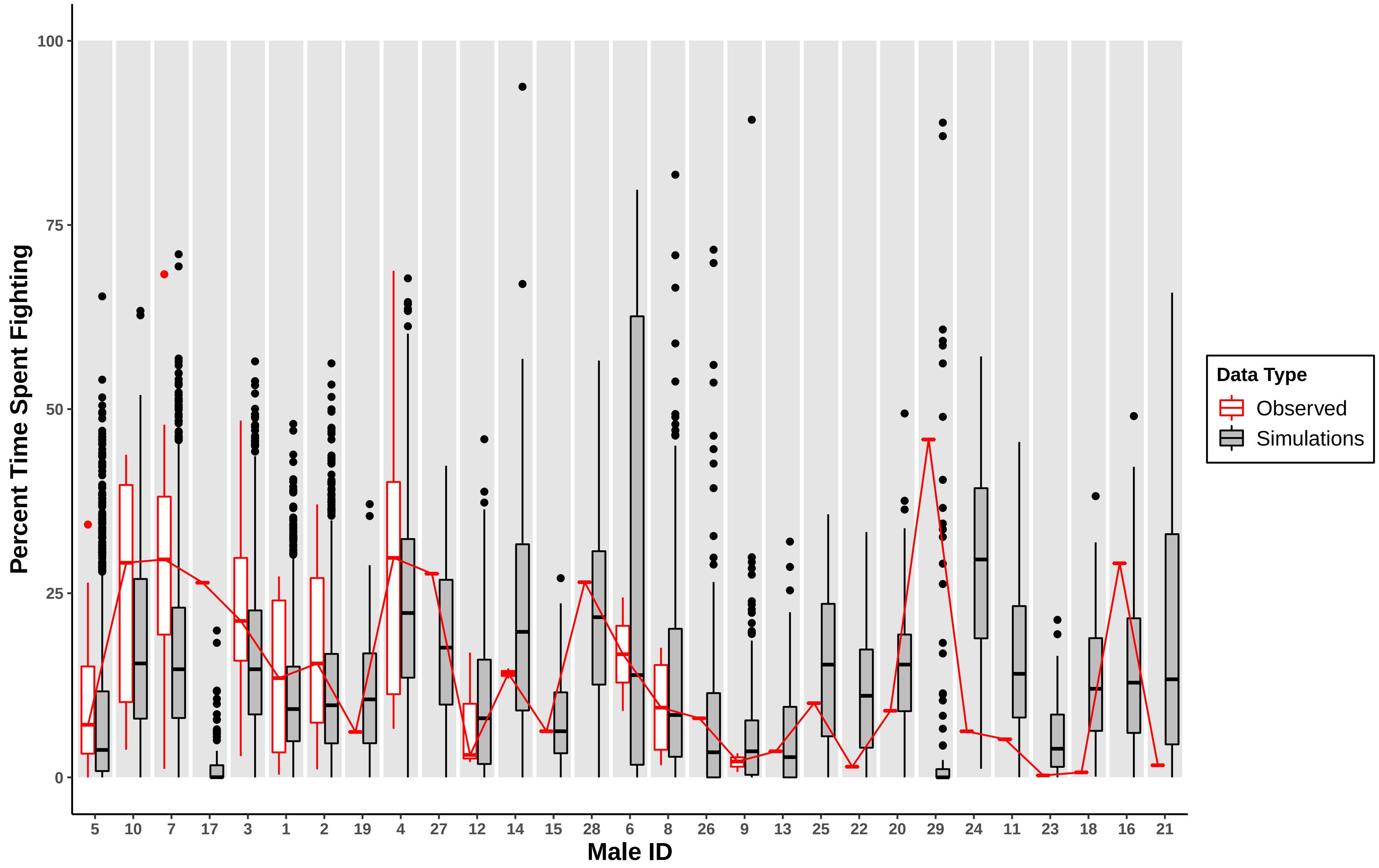

### overallfightplot.pdf

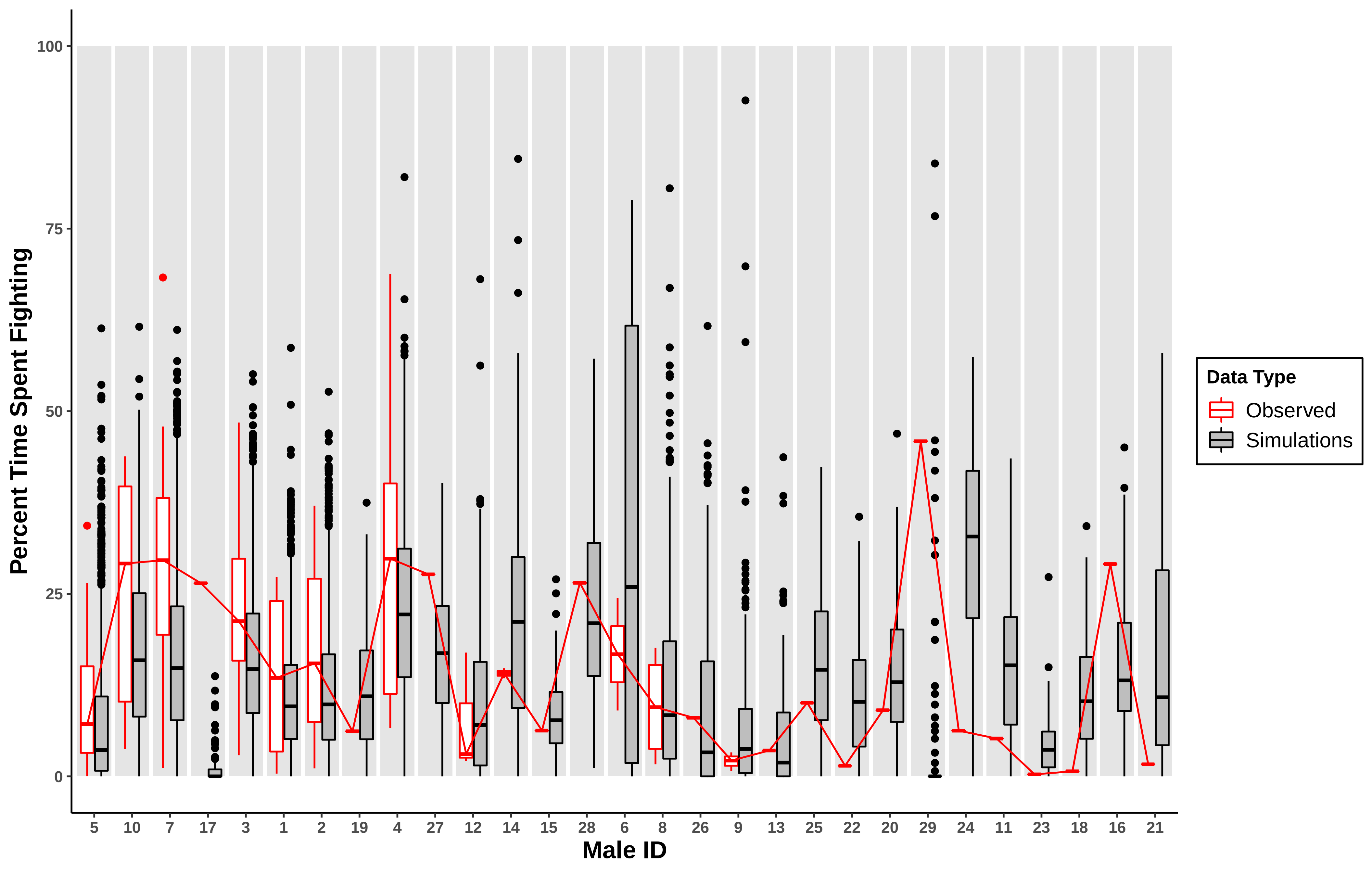

### overallnet_plot.pdf

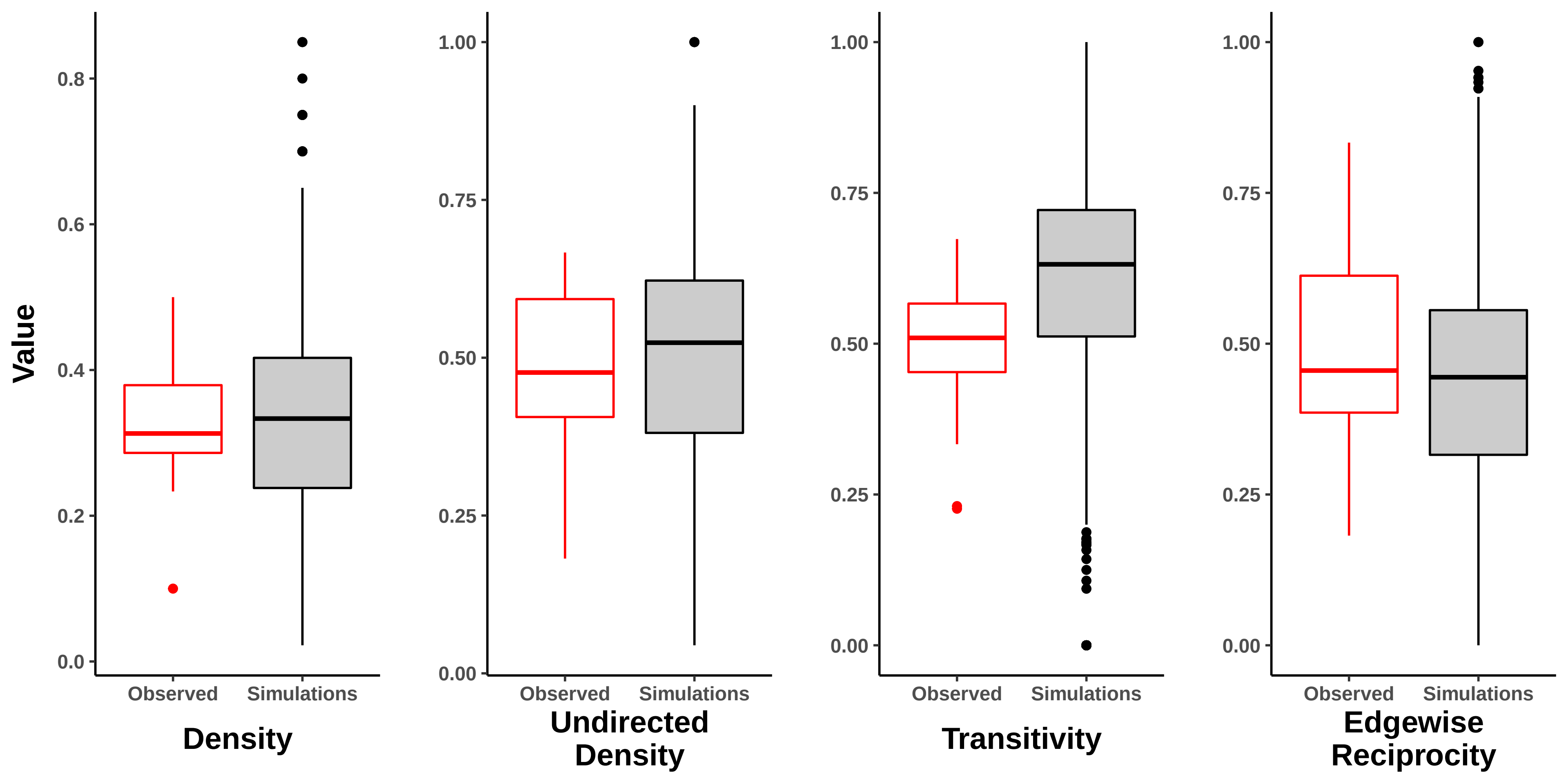

### overallnet_plot.pdf

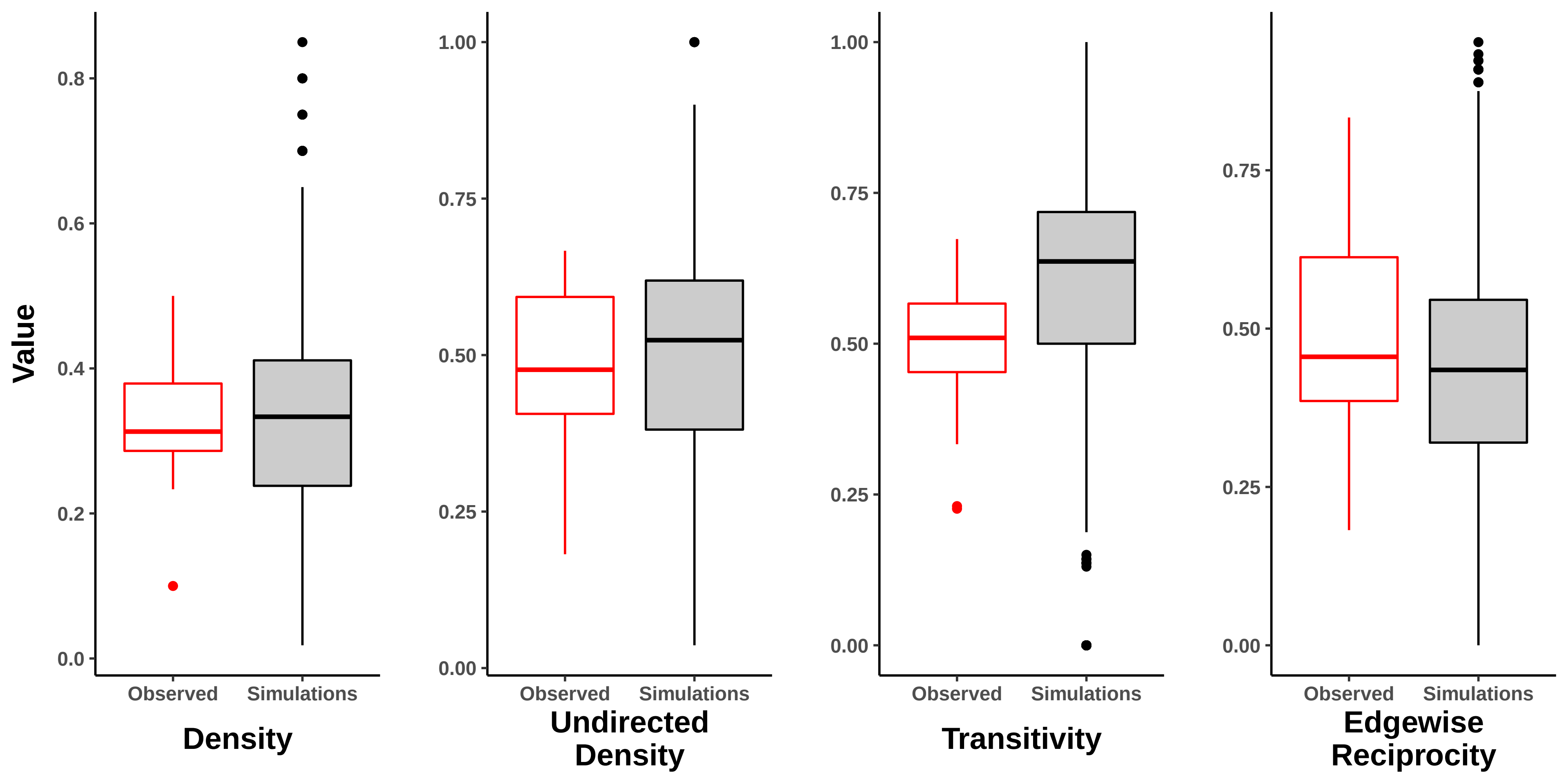

### supp_corplot_fight_by_mating.pdf

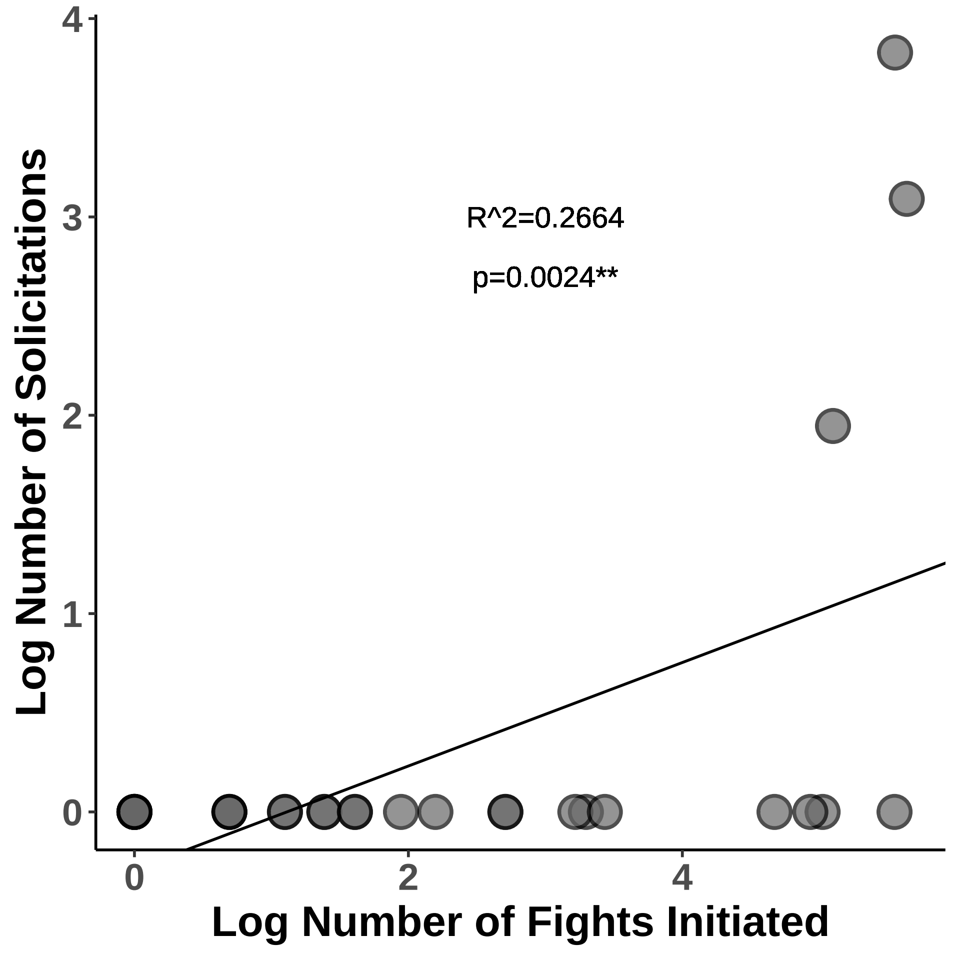

### time_fighting_fig.pdf

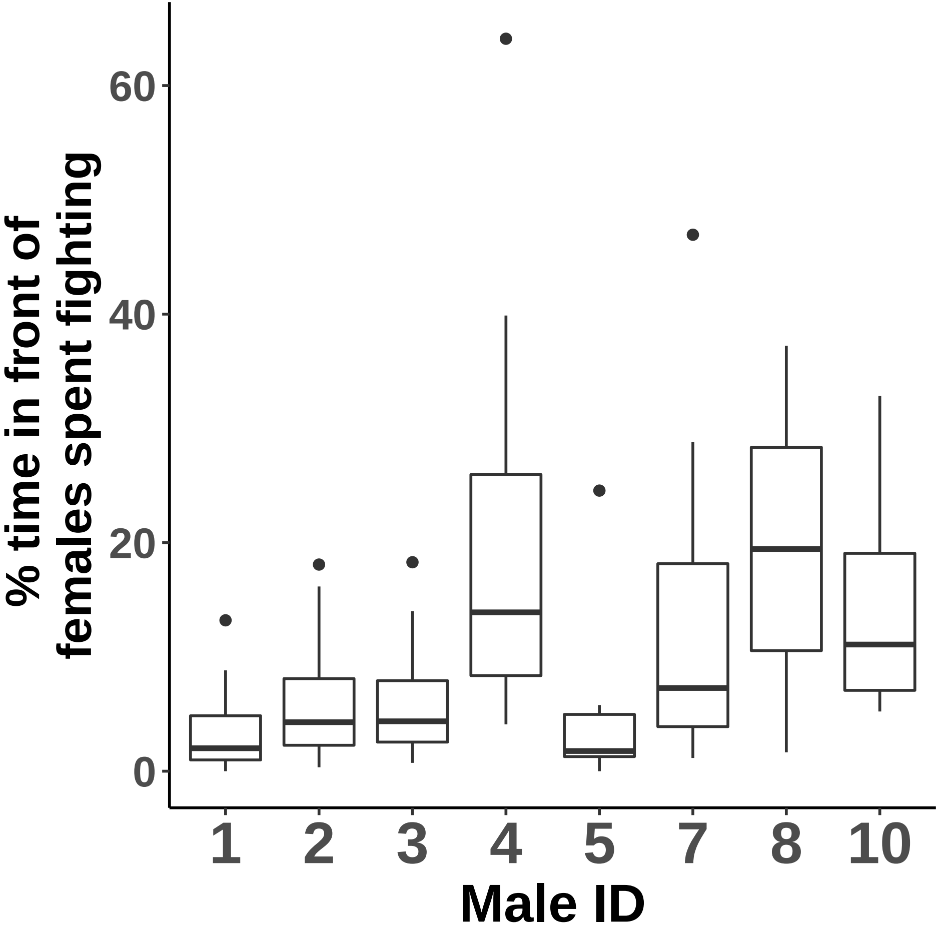

### totalcops.plot.pdf

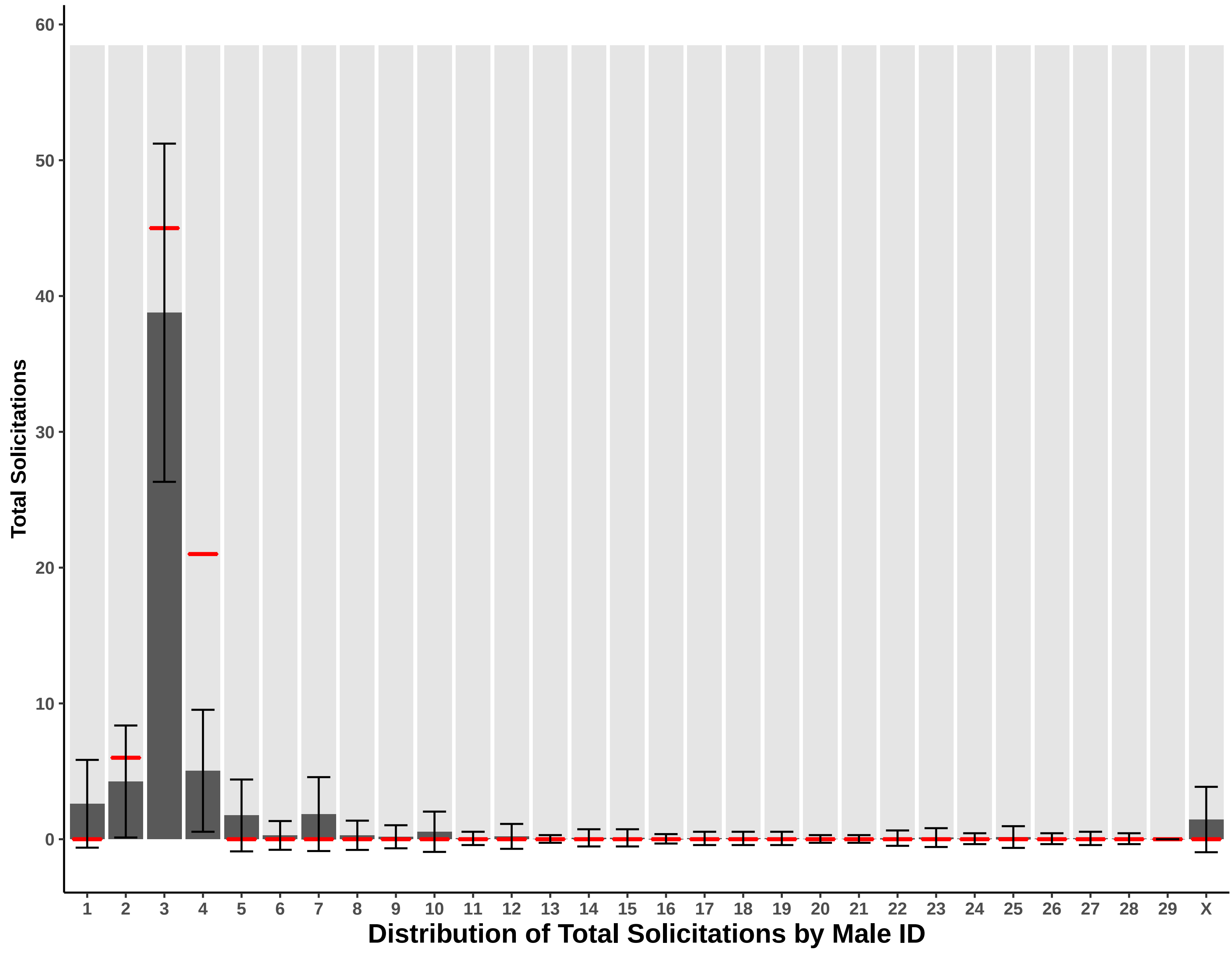

### totalcops.plot.pdf

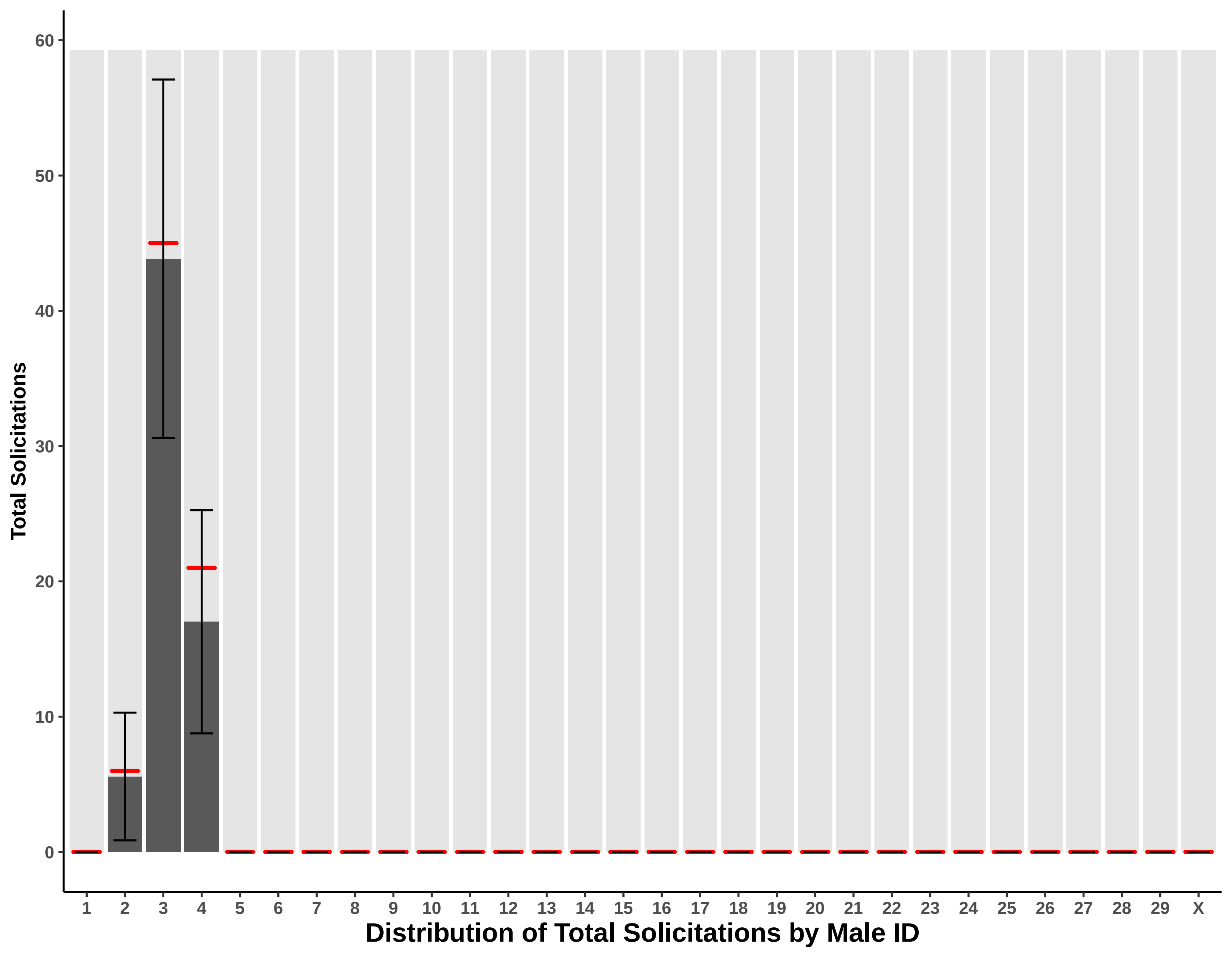

### totalends.plot.pdf

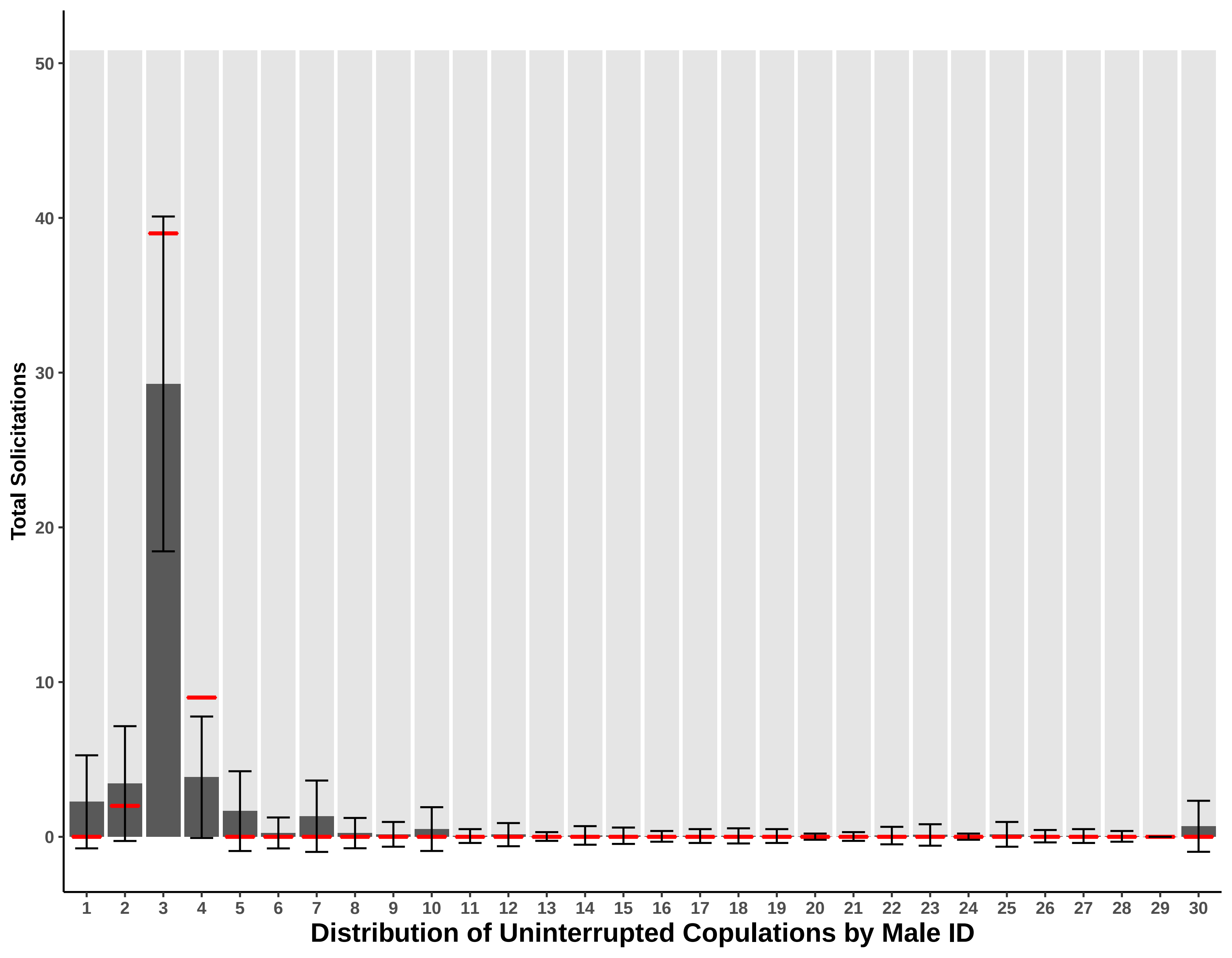

### totalends.plot.pdf

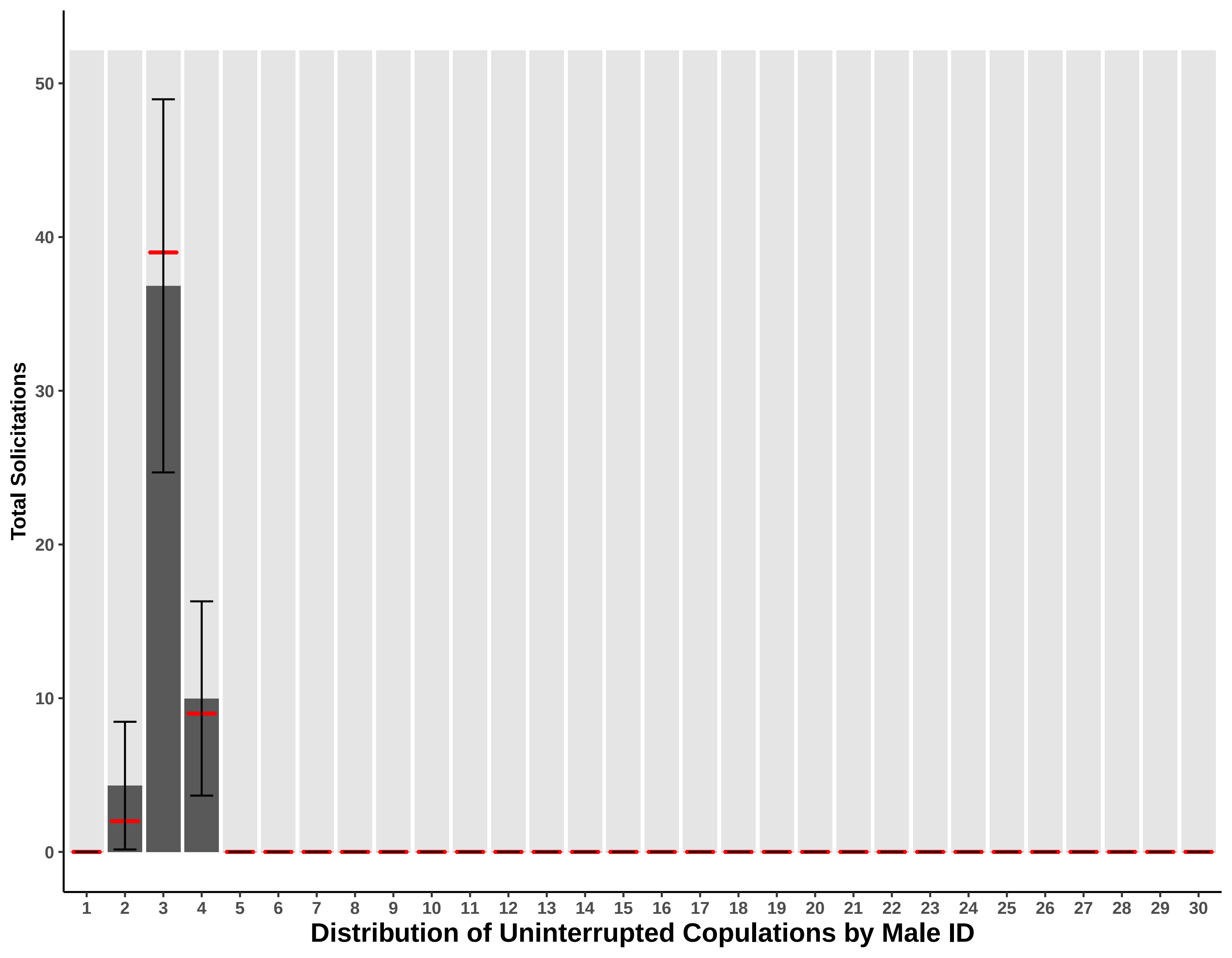

### totalints.plot.pdf

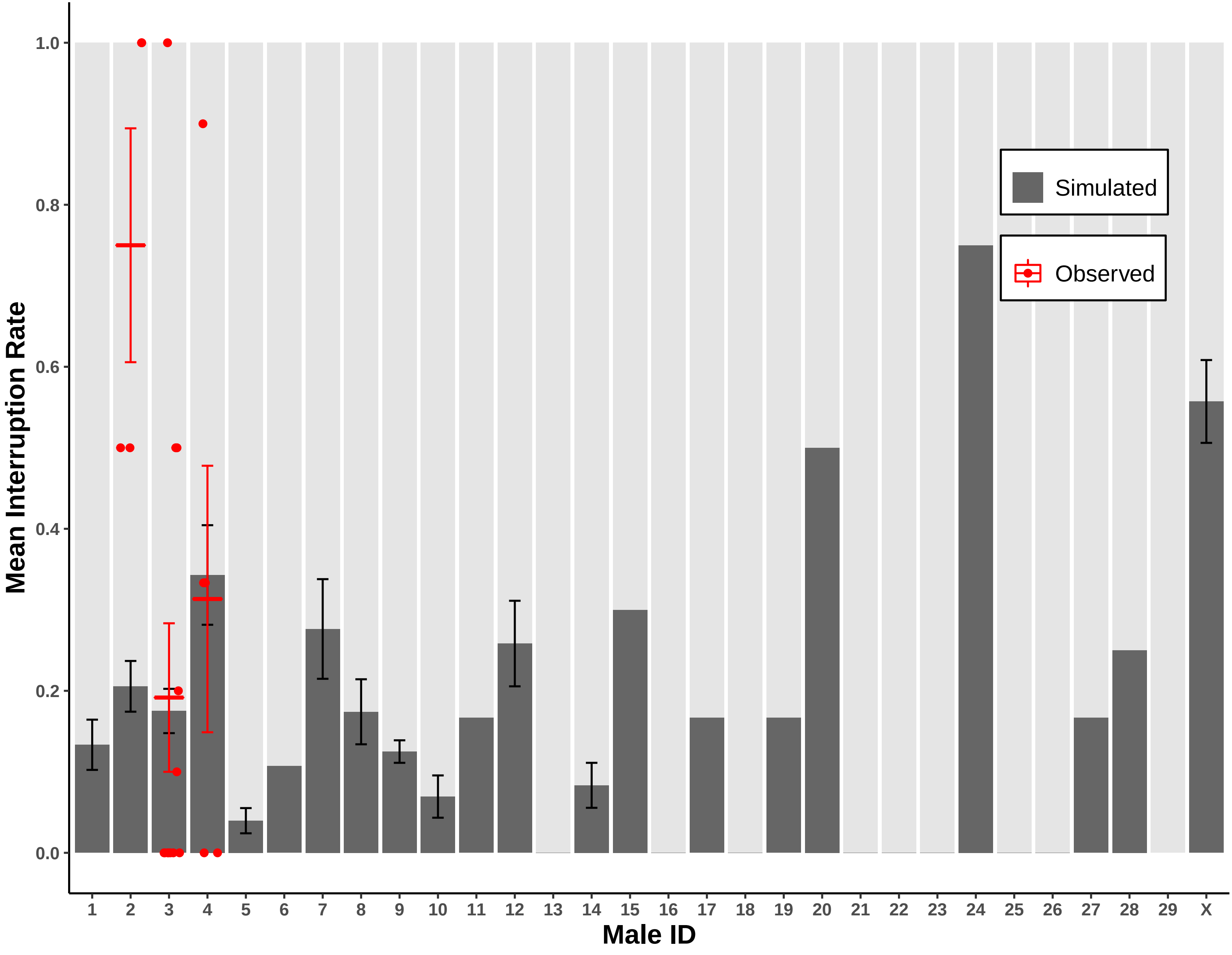

### totalints.plot.pdf

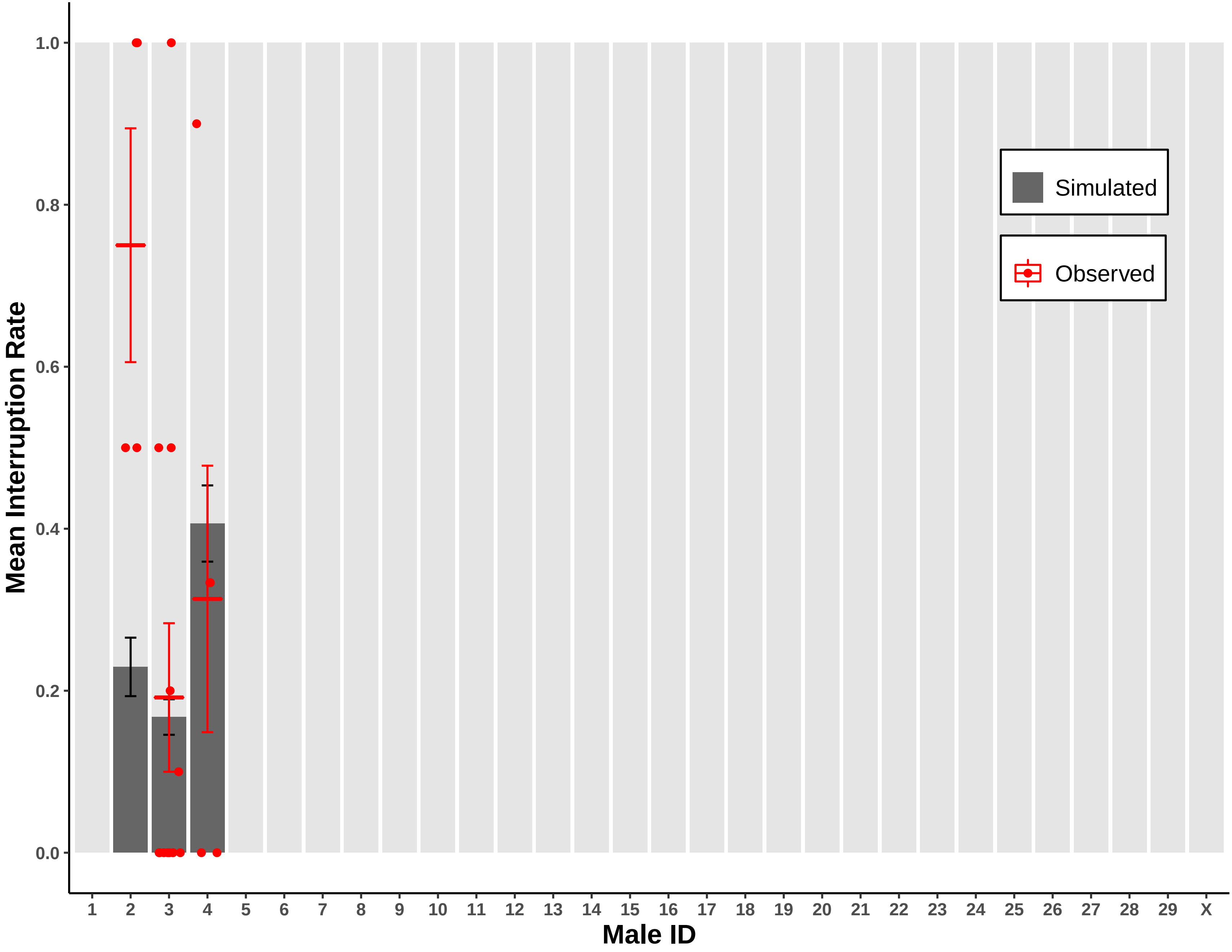

### totalints.small.pdf

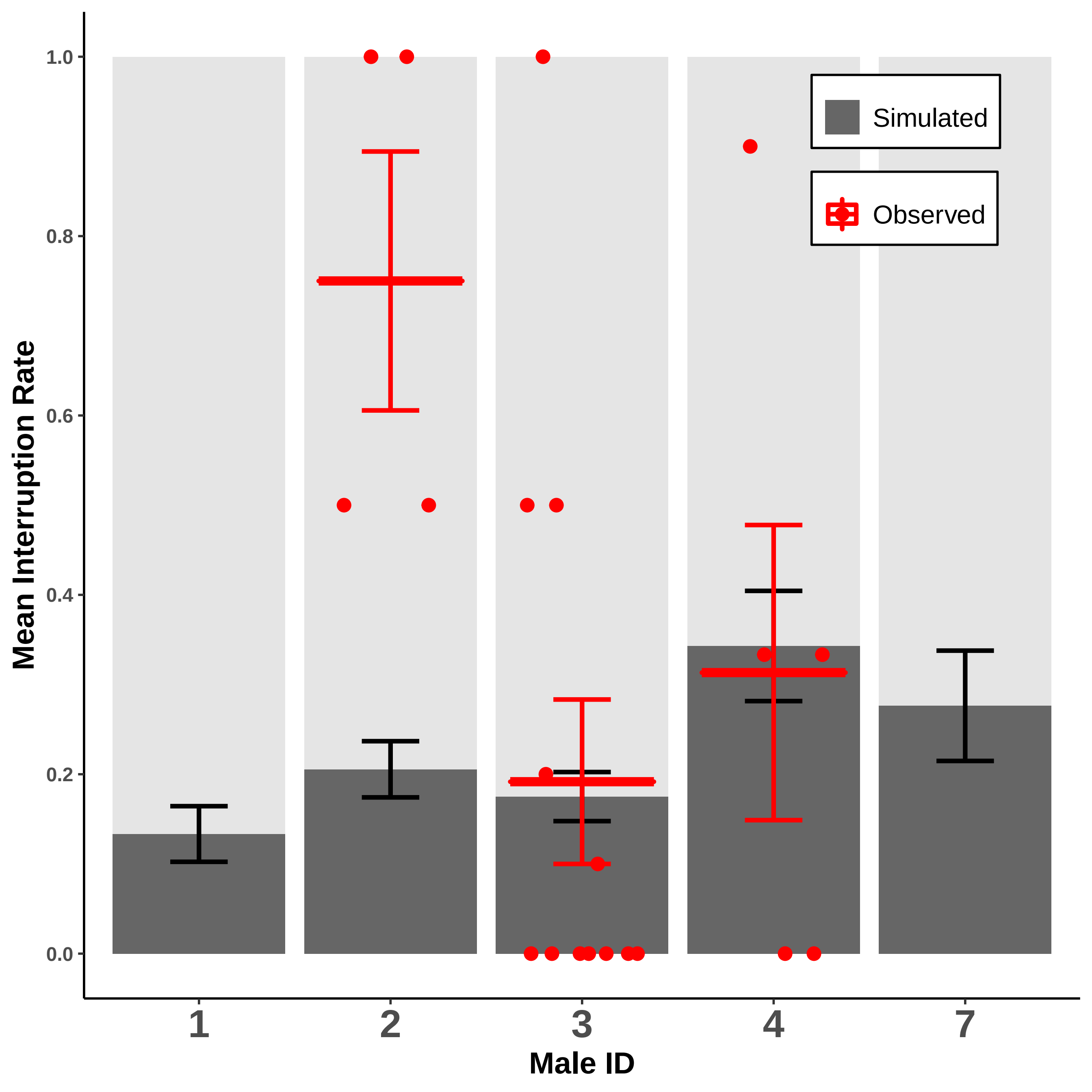

### totalints.small.pdf

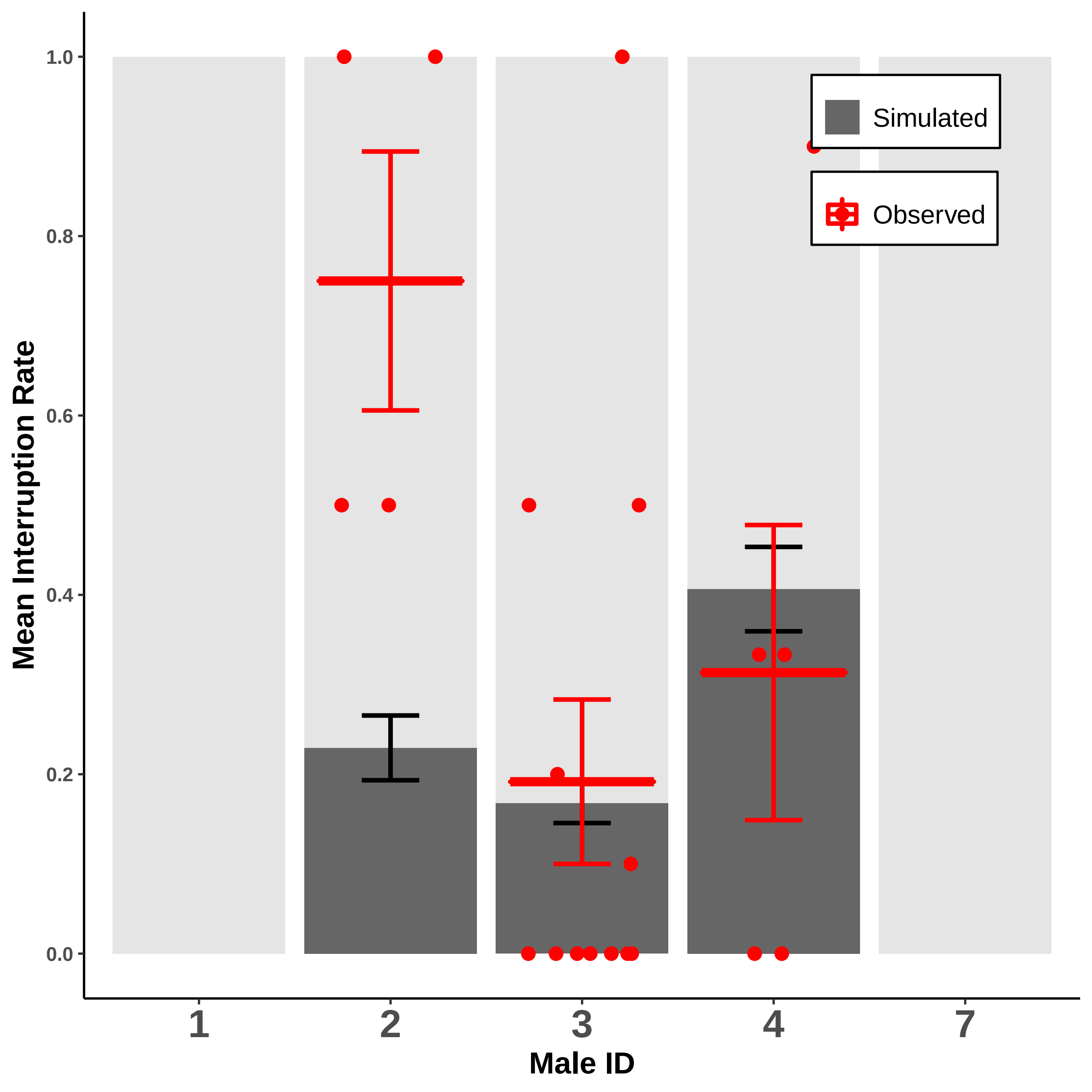
